# Pantetheinamides that inhibit the growth of intracellular *Salmonella* Typhimurium

**DOI:** 10.1101/2025.10.28.685111

**Authors:** Jacob Pierscianowski, Dustin Duncan, Chunling Blue Lan, Rohit Kholiya, Suzana Diaconescu-Grabari, Farhan R. Chowdhury, Ahmed B. Abdelaal, Aracelli Chang, Iain Roe, Shay Heans Moreyra, Sanjay Campbell, Alexandre Pierret, Kamal Adhikari, Srobona Ghosh, Angelos Pistofidis, Marcel A. Behr, T. Martin Schmeing, Brandon Findlay, Andréanne Lupien, Karine Auclair

## Abstract

The metabolic adaptability of intracellular pathogens, such as the Gram-negative *Salmonella enterica* serovar Typhimurium (STm), enables their survival in nutrient-restricted host environments, while also presenting an opportunity for selective antimicrobial targeting. Herein, we report the synthesis and screening of a small pantetheinamide library aimed at inhibiting the proliferation of STm within macrophages. Two lead compounds exhibited selective activity against intracellular STm and reduced bacterial burden in a murine colitis model. Mechanistic studies suggest their primary mode of action to be the inhibition of coenzyme A (CoA) biosynthesis via the PanD–PanZ regulatory complex, which is absent in host cells. Importantly, resistance pressure to our compound was shown to be significantly stronger in nutrient-limited media compared to nutrient-rich media, supporting the targeting of nutrient stress as a strategy to delay resistance. This work highlights the value of inhibiting pathogen-specific metabolic vulnerabilities in combination with host defence mechanisms to develop novel antimicrobials.

## Introduction

On the quest for innovative antibiotics to help fight the antimicrobial resistance crisis, an opportunity arises to design new treatments that can surpass the quality of care achieved by traditional antibiotics. For example, antibiotic-associated dysbiosis is the disturbance to our bodies that can result from a broad spectrum antibiotic damaging our microbiota and occasionally leading to chronic health issues (*1*). A healthy gut microbiota plays a critical role in preventing colonization by enteric pathogens such as *Salmonella enterica* serovar Typhimurium (STm), *Escherichia coli,* and *Clostridium difficile*, and supports recovery during infection (*2*). Accordingly, antimicrobial strategies that selectively target pathogenic bacteria while sparing beneficial commensals offer significant therapeutic advantages, including reduced side effects, lower rates of reinfection, and diminished pressure for resistance development.

Our approach to achieve this selectivity is to target intracellular (facultative or obligate) bacteria, including STm, *Mycobacterium tuberculosis* (Mtb), and *Legionella pneumophila*, which can adopt a pathogenic lifestyle within host cells, particularly macrophages (*3*). These immune cells phagocytose bacteria and treat them to a range of bactericidal conditions, including acidification, nutrient restriction, and the production of antimicrobial metabolites. One such endogenous antibacterial is itaconate: a Michael acceptor-bearing dicarboxylate with multiple cellular targets (**Fig. 1a**) (*4*). However, some pathogens evade the harmful effects of itaconate through specialized metabolic adaptations and detoxification systems. Inhibiting these could re-sensitize bacteria to our innate immunity and provide a basis for pathogen-specific antimicrobial strategies. In turn, this would reduce the resistance pressure to a narrow range of specific conditions.

**Fig 1:**
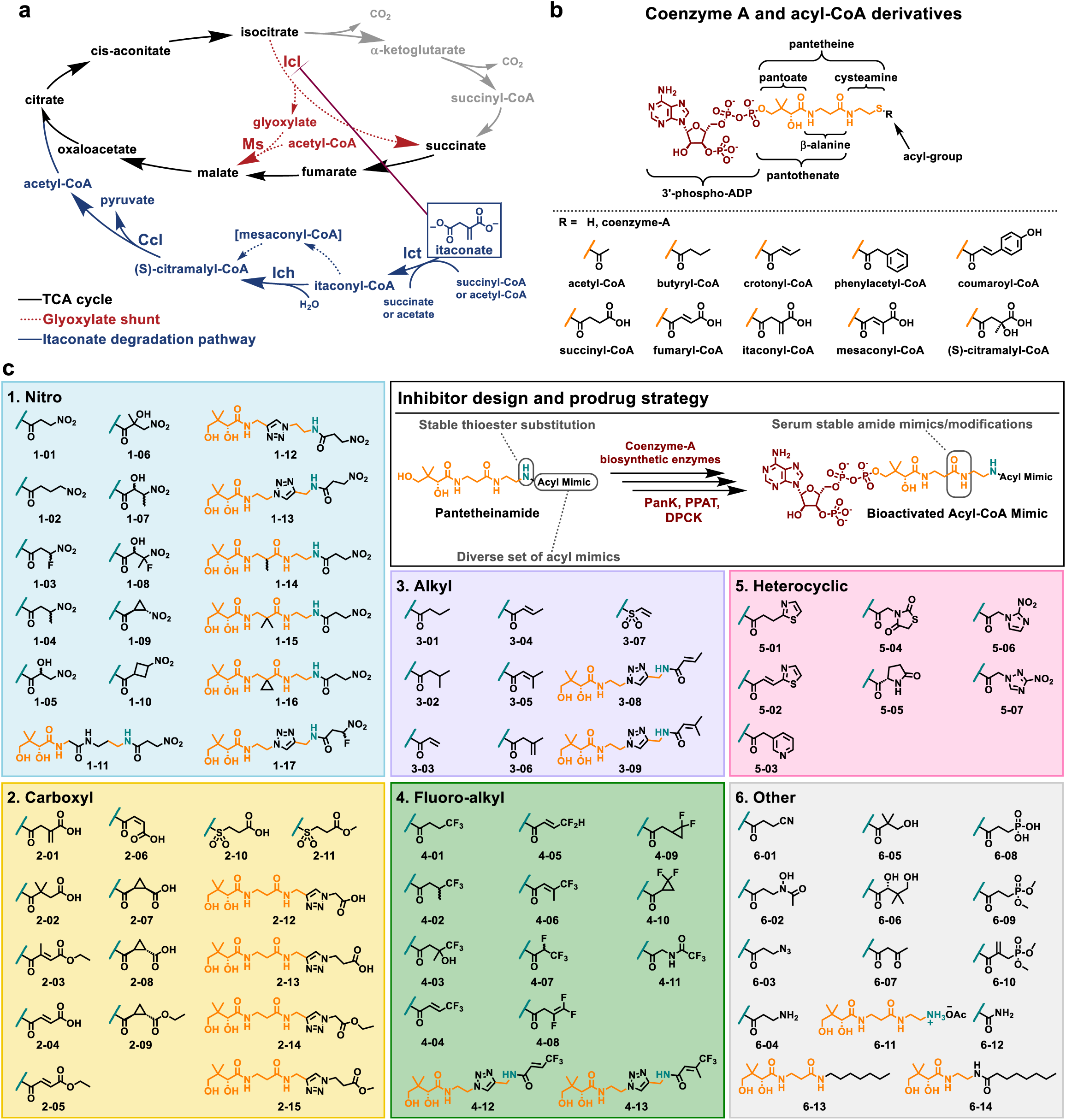
General structures, pathways and novel pantetheinamides library. **a,** The TCA cycle, glyoxylate shunt and itaconate degradation pathway all use CoA-acting enzymes and are important for STm intracellular survival. Itaconate is a phagocyte-derived antimicrobial molecule that inhibits bacterial Icl. It can be detoxified and degraded by STm into AcCoA and pyruvate by three enzymes: Ict, Ich, and Ccl. Ict utilizes succinyl- and acetyl-CoA as CoA donors, while Ich isomerizes itaconyl-CoA into the intermediate, mesaconyl-CoA, prior to hydration to *S*-citramalyl-CoA. **b,** The structure of CoA and its *S*-acyl derivatives contain three primary moieties (from left to the right): i) the 3’-phospho-ADP group (in red), ii) the pantetheine backbone (in orange), and iii) the acyl group (R) that results in the diversity of CoA molecules. Some of the most common cellular CoA species are shown, which served to inspire the mimicry strategy used in inhibitor design. The pantetheine backbone can be further subdivided into different components that reflect the many steps involved in CoA biosynthesis. **c,** The general inhibitor design is an acylated derivative of amino-pantetheine which is expected to be bioactivated by bacterial CoA biosynthetic enzymes to produce the full acyl-CoA mimic. The compound library is classified into six categories by the nature of the acyl mimic of the thioester-substituting amide bond in the general structure.

The Gram-negative bacterium STm, recognized as a high-priority pathogen by the WHO (*5*), causes significant global morbidity and mortality. It has a particularly flexible metabolism that allows it to persist in nutrient-limited intracellular environments (*6*, *7*). For example, in the absence of glucose, STm is dependent on the β-oxidation of fatty acids to acetyl-coenzyme A (AcCoA) and uses the glyoxylate shunt of the tricarboxylic acid cycle (TCA) to preserve carbon (**Fig. 1a**) (*8*). Macrophage-derived itaconate, which is present at up to 8 mM in activated murine macrophages (*9*, *10*), targets this conditionally essential pathway through inhibition of the first enzyme in the glyoxylate shunt, isocitrate lyase (Icl) (*11*, *12*). To combat this, STm employs a three-enzyme itaconate degradation pathway that detoxifies itaconate, converting it into AcCoA and pyruvate through the action of itaconate CoA transferase (Ict), itaconyl-CoA hydratase (Ich), and citramalyl-CoA lyase (Ccl). Notably, the TCA cycle, glyoxylate shunt, β-oxidation, and itaconate degradation all rely on CoA, a biologically ubiquitous cofactor derived from pantothenate (vitamin B_5_, **Fig. 1b**). CoA is essential for approximately 9% of bacterial enzymes and serves as a carrier of acyl groups to be used directly by enzymes, or indirectly as the precursor of the phosphopantetheinyl group of acyl-carrier proteins (ACP) (*13*). This centrality makes the CoA biosynthetic pathway a compelling target for antimicrobial intervention.

Pantothenamides are amide analogues of pantothenate that were first reported in the 1970s and have since been actively studied for their antimicrobial properties (*14*, *15*). These compounds are metabolically activated by three enzymes in the CoA biosynthetic pathway—pantothenate kinase (PanK), phosphopantetheine adenylyltransferase (PPAT), and dephospho-CoA kinase (DPCK)— into CoA antimetabolites that interfere with essential metabolic processes (*16*, *17*). Their modes of action (MoA) are multifaceted and vary depending on the organism in question, with the general consensus being that pantothenamides disrupt the homeostasis of CoA precursors and/or derivatives (*14*, *15, 18–21*). A structurally related class of molecules are the pantetheinamides (**Fig. 1c**), which are mimics of acyl-pantetheine with the thioester replaced by a more structurally stable amide. The antimicrobial activity of pantetheinamides has been underexplored, with their primary use in literature as tools for studying ACPs, highlighting that they too can be metabolically activated to CoA derivatives (*22–25*). Nonetheless, the structural similarity of bioactivated pantetheinamides to natural CoA intermediates suggests that they may possess a comparable or complementary MoA to that of pantothenamides.

In this study, we synthesized a library of novel pantetheinamides and evaluated their antimicrobial activity against STm, with emphasis on their ability to exploit the stress experienced during intracellular infection. To this effect, a multi-stage screen identified multiple hits with intramacrophage activity and low host-cell cytotoxicity. Remarkably, two compounds exhibited activity in a murine model of colitis. The mechanistic investigations presented herein suggest that the dominant MoA of these compounds is the inhibition of pantothenate biosynthesis via the PanD-PanZ regulatory complex. Resistance generation was only achieved in nutrient poor media and not in rich media, supporting our hypothesis that selective targeting of pathogenic populations may slow the development of resistance.

## Results

### Design of the pantetheinamide library

In this work, a small library of pantetheinamides (**Fig. 1c**) was designed by mimicking the wide breadth of natural acyl-CoAs, and relying on the promiscuity of bacterial CoA biosynthetic enzymes to ensure selectivity for bacterial targets and restrict potential cytotoxicity towards mammalian cells (*26*). In total, 70 novel compounds were synthesized. In addition to these, three compounds from one of our previous studies (*25*), **1-01**, **1-02**, and **3-03**, were included in this library, as well as two known pantothenamides, **6-13** (also known as N7-Pan) (*27*) and **6-14** (the reverse-amide analogue of N7-Pan) (*28*). Given that pantothenamides have not been tested in this context against STm, they were included to allow for comparison between the two classes. The library was classified into six groups based on the dominant functionality of the acyl mimic (relative to acyl CoA derivatives). Additionally, pantothenamides are known to have poor serum stability due to the presence of pantetheinases in serum that hydrolyze pantothenamides at the amide linking the β-alanine and cysteamine moieties (*29*, *30*). For this reason, some compounds were synthesized that include modifications to the pantetheine backbone that are known to improve serum stability (*28*, *31*, *32*).

### Screening of the pantetheinamide library

The goal is to identify molecules that selectively inhibit the growth of intracellular STm, without antibacterial activity in other contexts. To accomplish this, the compounds were screened for their ability to reduce STm growth at a high concentration (500 μM) in various media that reflect nutrient-related facets of either extracellular or intracellular environments (**Fig. 2a**). Mueller-Hinton broth (MHB)—the medium most widely employed for antibacterial testing—was used to represent nutrient-rich extracellular environments such as the gut, while aspects of the nutrient-limited intramacrophage environment were mimicked by using M9-minimal media containing either acetate (M9A) or itaconate (M9I) as the sole carbon source. Growth in M9A requires the glyoxylate shunt, simulating the use of fatty acids (*33*), whereas growth in M9I, as shown in our previous study, requires an active itaconate-degradation pathway and induces excess oxidative stress that is reflective of the intramacrophage environment (*34*). Of the 75 compounds tested, 31 inhibited growth (by >50%) in M9I, six in M9A, and one in MHB. Compounds from all structural classifications were found to be active in M9I. In particular, the pantothenamides **6-13** and **6-14** were active in M9A as well as M9I, with **6-13** being the only MHB active compounds. The other M9A active compounds were **1-03** and **1-17** from the “nitro” category and **4-02** and **4-08** from the “fluoro-alkyl” category.

**Fig. 2:**
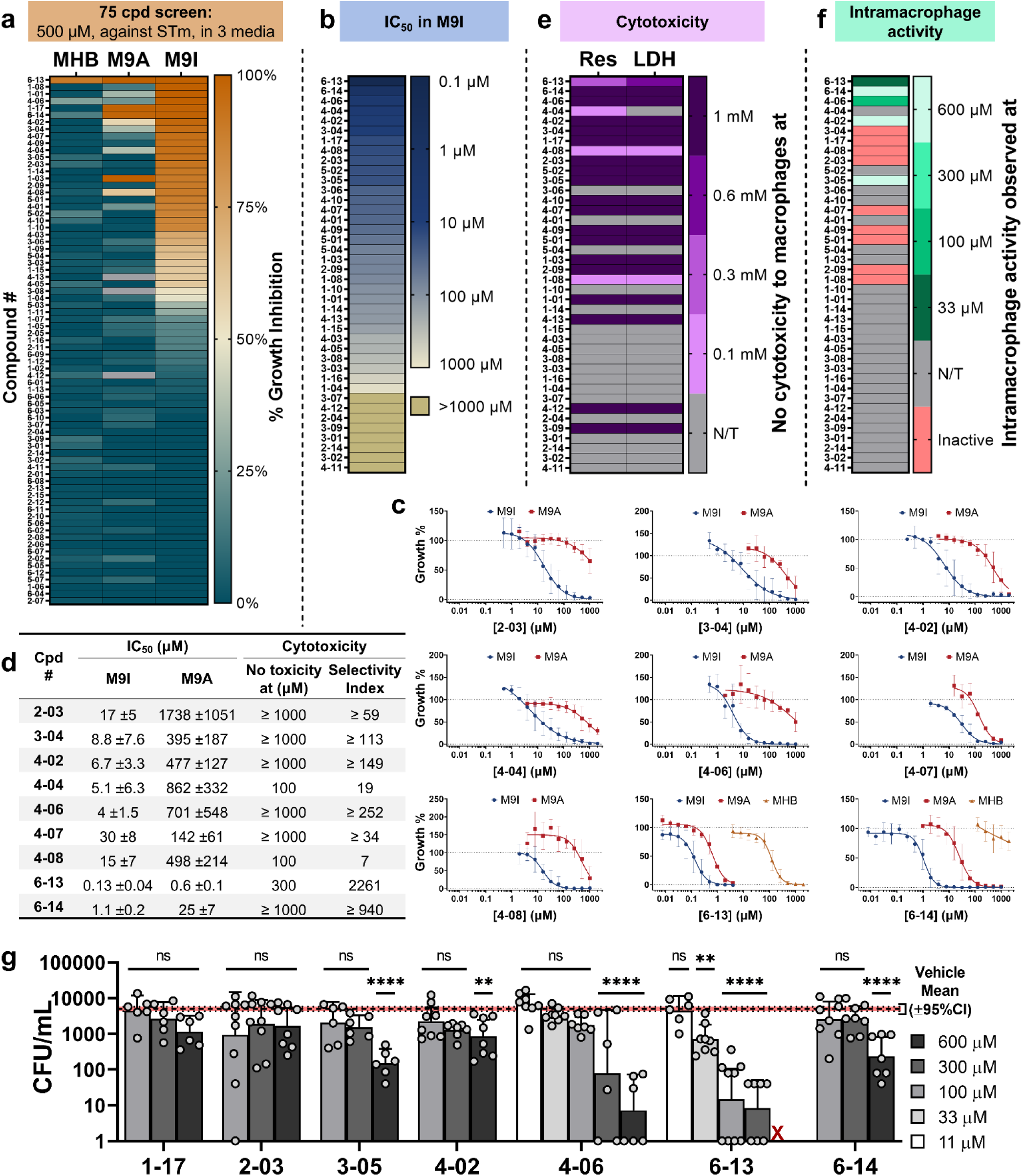
Screening the pantetheinamides library. **a,** Percentage growth inhibition of STm by each of the inhibitors in the library, tested at 500 µM in MHB, M9A, or M9I as the sole carbon source. Inhibition % values are summarized in **Supplementary Data 1**. **b,** The IC_50_ in M9I of a subset of the inhibitor library. The full dataset is shown in **Fig. S1** and summarized in **Supplementary Data 2**. **c,** Dose-response curves in M9I, M9A, and MHB for selected inhibitors, fit to a four-parameter logarithmic curve. **d,** Tabulated IC_50_ (± 95% CI), cytotoxicity and selectivity index (given as the cytoxocity/IC_50_ in M9I) for the same inhibitors as in **c**. In cases where no cytotoxicity was seen at the highest concentration tested (1 mM), a minimum index value is shown. **e,** Cytotoxicity of the compounds tested against RAW 264.7 murine macrophages after 24 h. Results are shown as the lowest concentration where no toxicity was seen in the two methods used: RES = resazurin, as a measurement of macrophage viability or LDH = lactate dehydrogenase enzyme activity released from dead cells. The lower concentration was used for selectivity index calculations when the two methods did not agree. **f,** Summary of the lowest concentrations where compounds significantly (p < 0.05) inhibited the proliferation of STm inside RAW 264.7 macrophages after 22 h. The full dataset is shown in **Fig. S4a**. **g,** Sample of results summarized in **f** showing the CFU of surviving STm inside the compound treated macrophages after 22 h. Data bars show the geometric mean of CFU/mL with error bars depicting the 95% CI. The vehicle control (± 95% CI) is shown as the red line. All experiments were independently repeated a minimum of 3 times, where n ≥ 4 for each condition. Significant results were determined using a one-way ANOVA with Dunnett’s multiple comparisons against the vehicle. ✱, p < 0.05; ✱✱, p < 0.01; ✱✱✱, p < 0.001; ✱✱✱✱, p < 0.0001; ns, non-significant, p > 0.05. “X” denotes that a concentration was omitted from the experiment due to toxicity.

The full dose-response curve was collected for all active molecules in M9I, in addition to some inactive molecules (**Fig. 2b**). Many compounds exhibited broad dose-response curves (**Fig. 2c** and **S1**), consistent with bacteriostatic antibiotics that target bacterial metabolism, as is proposed for pantetheinamides (*35*). Consequently, the activity was reported as an IC_50_ value, rather than traditional MICs. A sub-micromolar IC_50_ in M9I was observed for the pantothenamide **6-13** (130 nM), and 6 compounds exhibited an IC_50_ between 1 and 10 μM (**Fig. 2d**; **6-14**: 1.1 μM, **4-06**: 4.0 μM, **4-04**: 5.1 μM, **4-02**: 6.7 μM, and **3-04**: 8.8 μM). Using this data, important structure-activity relationships (SARs) can be established (**Fig. S2**). The fluoro-alkyl category of acyl mimics had the greatest number of molecules with an IC_50_ below 100 μM (8 of 13) and contained the three most active pantetheinamides. All other categories had 2-4 molecules with sub-100 μM IC_50_ values. The general trend in activities for the substituents β to the acyl amide saw CF_3_ (*e.g.* **4-01**) > thiazole (*e.g.* **5-01**) > NO_2_ (*e.g.* **1-01**) > CH_3_ (*e.g.* **3-01**). Introducing α,β-unsaturation (*e.g.* **4-04**), cyclopropyl (*e.g.* **4-09**), or cyclobutyl (*e.g.* **1-10**) groups into the acyl mimic increased the activity when compared to their saturated or linear counterparts, suggesting that the rigidity imparted by these structures may be beneficial. Within the carboxyl category, only esters exhibited activity, which could be the result of the charged acid not being able to pass through cell membranes to reach the target. Replacement of the thioester with either a sulfonamide or triazole, rather than an amide, was detrimental to activity. Compounds with mimics/modifications to the pantetheinase-labile amide exhibited decreased potency when compared to their unmodified counterpart, with the exception of **1-17**. The addition of larger groups resulted in a greater loss of potency: replacement of the amide with a triazole had a greater impact than reversing the amide or adding a small α-substituent. Furthermore, increasing the conformational rigidity of the α-substituent correlated with higher IC_50_ values (*e.g*, **1-14**, methyl < **1-15**, dimethyl < **1-16**, cyclopropyl). It appears that the reduced flexibility in the pantetheine backbone restricts the ability of these molecules to bind to their target in the optimal conformation. Overall, smaller, strategically located modifications to the pantetheine backbone are more conducive to better antibacterial activity.

Next, IC_50_ values for select compounds were also measured against STm in M9A and MHB (**Fig. 2c-d**). In all cases, the IC_50_ increased from M9I, to M9A, and to MHB growth media. However, the extent of the IC_50_ shift between M9I and M9A was found to vary considerably between compounds, possibly due to dissimilarities in cellular targets. Moreover, we validated that the greater sensitivity of STm to compounds in M9I was not the result of known mutations acquired during growth in M9I (*34*), by comparing the activities of **2-03**, **4-06**, **6-13**, and **6-14** in M9A for STm strains not adapted to M9I, subcultured twice in M9I, or subcultured eight times in M9I (**Fig. S3**). Further elaboration of the screen involved measuring the cytotoxicity of molecules on RAW 264.7 murine macrophages (**Fig. 2e**). All 22 compounds screened were non-toxic at 100 μM with 18 of them being non-toxic up to 1 mM. The selectivity index (SI) was also calculated (**Fig. 2d**) to further refine compound prioritization.

The final stage of the screening assessed the ability of selected compounds to inhibit the proliferation of intramacrophage STm (**Fig. 2f-g** and **Fig. S4a**). Five molecules significantly reduced the survival of STm inside macrophages after a 22 h period. Consistent with the M9I IC_50_ data, **6-13** was the most potent, exhibiting activity at concentrations as low as 33 μM. The intramacrophage screen involved a pre-incubation of both bacteria and macrophages with compound prior to infection, however, separate experiments using **4-06** demonstrated that neither pre-incubation step had any influence on compound activity (**Fig. S4b**). A time course experiment measuring STm survival at 2, 4, 14, and 22 h in response to treatment with **4-06** or **6-13** revealed that the compounds do not disrupt the ability of STm to infect the macrophages (even with pre-incubation), but rather, reduce their survival over time, reaching a minimum at 14 h (**Fig. S4c-d**). Overall, three novel pantetheinamides and two pantothenamides were identified that selectively inhibit the survival of intramacrophage STm, with **4-06** and **6-13** standing out as the molecules with the greatest activity and highest selectivity index from each structural class.

### The activity of pantetheinamides in M9I is not the result of itaconate degradation inhibition

We hypothesized that a plausible explanation for the greater activity in M9I compared to M9A could be due to the inhibition of itaconate degradation. To test this, two sets of experiments were employed. First, a checkerboard assay in M9A was performed to see if there was any pharmacological interaction between itaconate and the pantetheinamides (**Fig. S5**). If the pantetheinamides inhibit itaconate degradation, then the susceptibility of STm to itaconate would increase at higher pantetheinamide concentrations. None of the five compounds tested exhibited such an effect, for both wt STm or Δ*ich* STm, which cannot degrade itaconate (*34*). These results were confirmed by testing **4-06** and **6-13** (bioactivated and non-bioactivated) for their ability to block the activity of purified Ict, Ich, and Ccl enzymes (**Fig. S6**). No inhibition was detected. Together, these results suggest that itaconate degradation inhibition is not the MoA.

### The dominant mechanism of pantetheinamides is via disruption of PanD-PanZ regulation

The next step in exploring the MoA of the pantetheinamides was to explore whether they share a common mechanism with pantothenamides (*14*, *21*, *36*). For example, the pantothenamide *N*-pentyl pantothenamide (N5-Pan) was reported to inhibit pantothenate biosynthesis in *E. coli* (**Fig 3a**) (*18*, *37*). The first committed step of this pathway is the decarboxylation of L-aspartate to form β-alanine by the enzyme aspartate-1-decarboxylase (PanD), which is produced as a zymogen (pro-PanD). Autocatalysis of pro-PanD into its active form is regulated by PanZ (*37–41*). When PanZ binds CoA or AcCoA, it changes conformation, allowing it to bind pro-PanD and triggering its autocatalysis. However, PanD remains inactive as long as it is part of the PanD-PanZ-CoA complex, thus creating a negative feedback loop at high CoA concentrations. When CoA levels decrease, CoA and PanZ dissociate from the now active PanD, which produces β-alanine for the biosynthesis of pantothenate. The bioactivated N5-Pan antimetabolite (ethyldethiacoenzyme A) was shown to bind to PanZ with the same affinity as AcCoA, suggesting that PanZ may be its target (*37*). Furthermore, N5-Pan toxicity to *E. coli* grown on M9-glucose could be rescued by the addition of 0.5 mM β-alanine. Taking inspiration from this rescue experiment, the activity of both pantetheinamide **4-06** and pantothenamide **6-13** were tested against STm in M9I in a checkerboard assay with either β-alanine, L-aspartate, D-pantolactone (known to be metabolized to D-pantoate) (*42*), D-pantothenate, or L-alanine (**Fig. 3b**). Importantly, the inclusion of L-alanine in this dataset allows us to eliminate any effect that would arise from the addition of an easily metabolized new carbon and nitrogen source on the activity of the compounds. The results clearly assert that the addition of β-alanine or D-pantothenate can reduce the activity of both inhibitors with effects seen at concentrations as low as 100 nM of supplement. The larger effect on the activity of **4-06** compared to **6-13** suggests that the difference in IC_50_ (in M9I) between **4-06** and **6-13** could be predominantly due to other MoAs of **6-13**, while PanD-PanZ inhibition could be the main MoA of **4-06**. The lack of rescue by D-pantolactone emphasizes that it is the β-alanine moiety of D-pantothenate that allows for its rescue phenotype. Further supporting the PanD-PanZ inhibition mechanism is the small effect seen for L-aspartate.

**Fig. 3:**
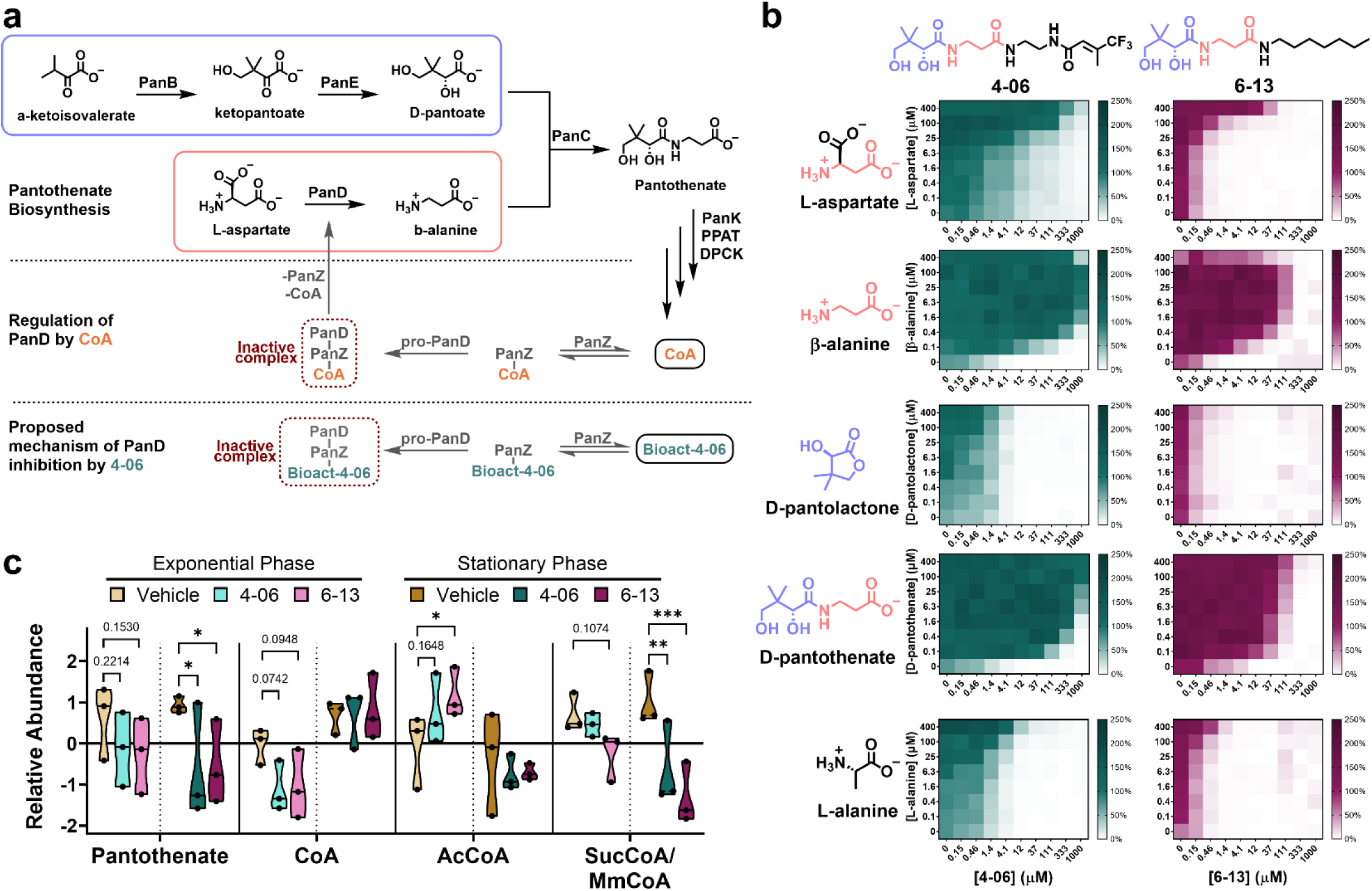
Inhibition of pantothenate biosynthesis is a major mechanism of action for 4-06 and 6-13. **a**, The pantothenate and CoA biosynthetic pathways in STm. The autocatalysis of pro-PanD is regulated by PanZ in response to intracellular CoA levels, where PanD remains inactive when complexed with PanZ and CoA. The proposed mechanism is that the PanK/PPAT/DPCK bioactived derivatives of **4-06** and **6-13** bind PanZ similar to CoA and inhibit PanD activity. **b**, Checkerboard assay showing the shift in the STm dose-response curve (in M9I) of **4-06** and **6-13** with the supplementation of metabolites from pantothenate biosynthesis (L-Asp, β-Ala, D-pantolactone, D-pantothenate) or a control metabolite (L-Ala). Colour gradient depicts the growth % compared to the vehicle control, n ≥ 3 for all conditions. **c**, Targeted metabolomics (LC/MS/MS) showing the relative abundance of the intracellular metabolites pantothenate, CoA, AcCoA, and SucCoA/MmCoA (isomers with identical elution and fragmentation pattern) in response to 10x the IC_50_ (in M9I) of **4-06** and **6-13** when grown starting at an OD_600_ = 0.1. Samples were collected at the same point along their respective growth curves during the mid-exponential phase (0.3 × growth plateau OD) or the early-stationary phase (0.9 × growth plateau OD) of growth. Significant results were determined using a two-way ANOVA against the vehicle within each of growth phase. ✱, p < 0.05; ✱✱, p < 0.01; ✱✱✱, p < 0.001; Non-significant (*p* > 0.05) comparisons are either not shown or shown as the exact *p*-value.

To corroborate this finding, metabolomics was used to examine the intracellular concentration of pantothenate and CoA in response to treatment (**Fig. 3c**). STm was grown in the presence of either **4-06**, **6-13** or vehicle and metabolic profiles at an exponential phase (EP) and at stationary phase (SP) of growth were evaluated. At both phases, lower levels of pantothenate were observed in response to **4-06** and **6-13** compared to the vehicle, with the effect being significant at SP. CoA levels were also reduced, but only at EP. In the case of AcCoA, an increase was seen at EP, which could be related to the fact that AcCoA is a direct product of itaconate catabolism and could have accumulated while other metabolic processes slowed. At SP, when metabolic rates have slowed, a negligible decrease in AcCoA is observed. Furthermore, succinyl-CoA (SucCoA) and its isomer methylmalonyl-CoA (MmCoA) levels decreased in both phases of growth, with significance observed at SP. The overall trends point towards both compounds decreasing the general pantothenate and CoA pools, with the more active compound, **6-13**, consistently exhibited a larger effect. While this is consistent with the compounds targeting PanZ and affecting its regulation of PanD, a more direct binding assay will be necessary to confirm this in the future.

### STm resistance to pantetheinamide 4-06

To validate our hypothesis that pantetheinamides may present a reduced potential for resistance in extracellular settings, resistance generation to **4-06** was attempted in both rich and poor media. This was accomplished using the soft agar gradient evolution (SAGE) method, which has been shown to better identify clinically relevant resistance mechanisms than other laboratory resistance generation approaches (*43*, *44*). SAGE experiments involve growing motile bacteria across soft agar plates with an increasing gradient of the antibiotic of interest (**Fig. S7**). Resistance generation was tested on both M9I agar and Mueller-Hinton agar (MHA) matrices using **4-06** at a maximum concentration of 5x the MIC measured in M9I. Background mutations observed in the M9I-adapted parent strain are consistent with those found in our previous findings (**Tables S1 and S2 and Supplementary Text 1**) (*34*).

Substantial resistance to **4-06** was achieved in M9I, with an MIC shift from 32 µM (pre-SAGE) to ≥ 512 µM (**Table 1**). Conversely, in MHA, a two-fold increase in the MIC of **4-06** was observed in only half of the replicates, supporting our hypothesis that resistance generation in rich media is slower for pantetheinamides. Five mutations were identified at high frequencies (>50%) and across all evolution experiments in M9I with **4-06**, suggesting that they contribute to resistance (**Table 2**). The first mutation is a one nucleotide (+C) insertion into the promoter region of *panB* (which also controls expression of *panC*), between the -10 and -35 elements, which was observed at 100% frequency in all four evolved populations. This exact mutation was previously shown to increase the expression of the *panBC* operon and increase the cellular concentrations of pantothenate and CoA, thus directly supporting the proposed MoA of **4-06** (*45*). Given that pantoate did not reverse the activity of **4-06** or **6-13**, it is likely the increased *panC* transcription that is beneficial. The second mutation is a single nucleotide polymorphism (SNP) in the putative transporter *yfdC*, resulting in amino acid 61 being mutated from alanine to threonine. This was seen at a frequency between 78-100% in all replicates. YfdC transporters belong to their own poorly characterized subfamily within the formate/nitrite transporter family (*46*). However, computational analysis of the *E. coli* YfdC (87% identity with STm YfdC) have suggested that this subfamily transports neutral or possibly cationic substrates, rather than anionic (*47*). As such, it is plausible that neutral analogs of pantothenate/pantetheine are natural substrates of YfdC, and mutation either prevents their import or increases their efflux, thus warranting further exploration into its natural function. The three other mutations identified were a nonsense and frameshift deletion in the genes encoding the MarT and IscR transcription factors, respectively, as well as a SNP in the ArcB sensor kinase of the aerobic response two-component system. The exact effect of these could not be surmised from the literature describing these transcription factor’s regulons (**Supplementary Text 2**).

**Table 1:** MIC values before and after SAGE resistance generation to compound 4-06.

| SAGE Exp. | MIC (μM) |  |  |  |
| --- | --- | --- | --- | --- |
|  | Rep<sup>a</sup> 1 | Rep<sup>a</sup> 2 | Rep<sup>a</sup> 3 | Rep<sup>a</sup> 4 |
| **Parent Strain** | 32 (range: 8 – 64<sup>b</sup>) |  |  |  |
| **M9I + 4-06** | >512 | 512 | 512 | >512 |
| **MHA + 4-06** | 64 | 64 | 32 | 32 |

<sup>a</sup> Each replicate (rep) refers to independent evolutions under each condition.
<sup>b</sup> Range of MICs found across 5 repeated measurements of the parent strain MIC. The median and mode MIC is 32 μM.

**Table 2:**
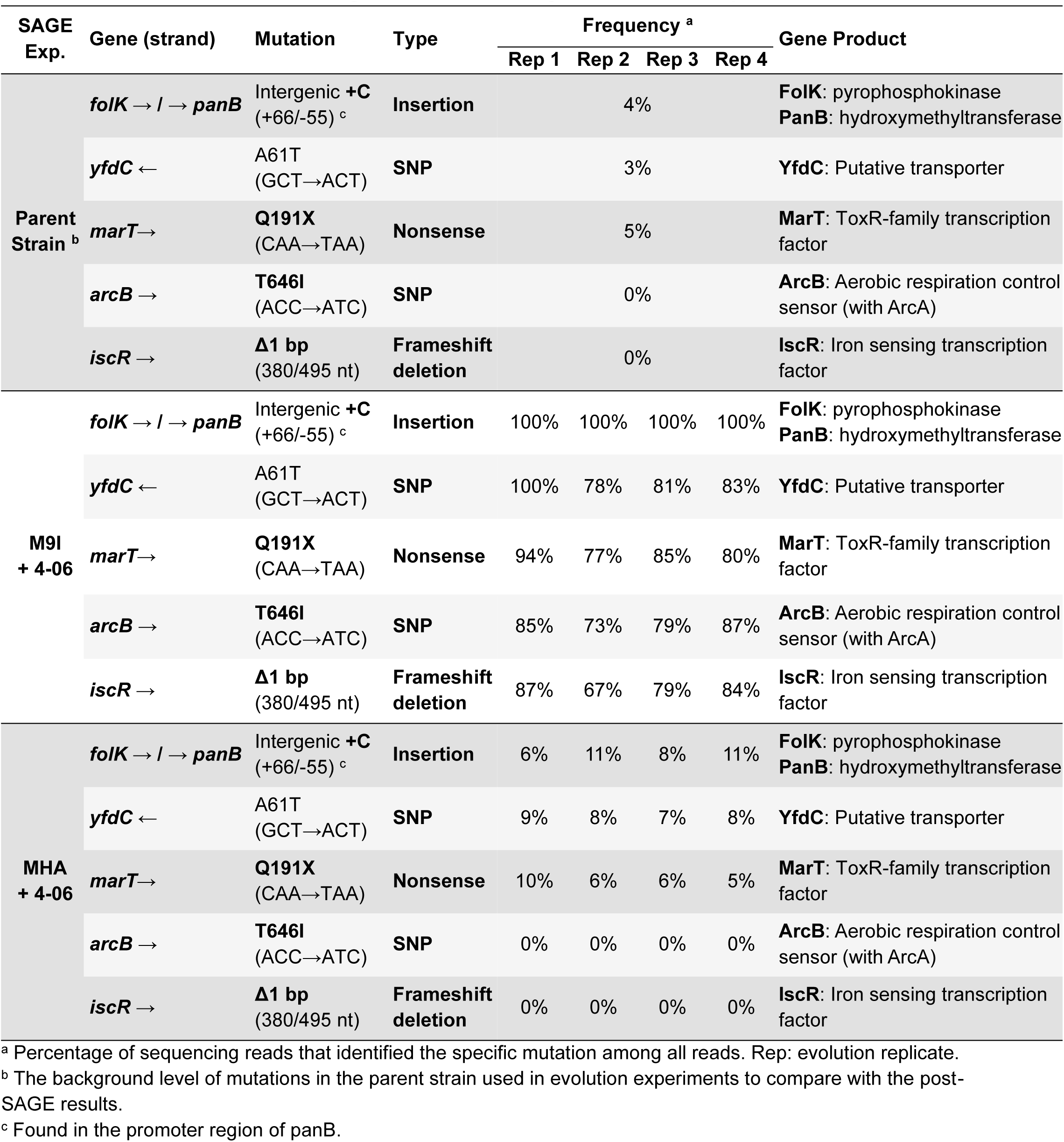
Mutations identified during SAGE resistance generation to compound 4-06.

Control evolutions without **4-06** were also done to identify mutations that may increase fitness during SAGE and determine if they impact the activity of **4-06**. Surprisingly, bacteria in two of the four M9I control (without **4-06)** evolutions were found to have acquired resistance to **4-06**, while none of the MHA control evolutions became resistant (**Table S3**). Genomic analysis identified that the *panB* mutation was the only one to be enriched in both resistant M9I control evolutions (**Table S4**). This highlights that the mutation in the *panBC* promoter region is sufficient for resistance (**Supplementary Text 3**) and further supports the proposed mechanism of action of pantetheinamides.

### In vivo efficacy of 4-06 and 6-13 in a murine model of colitis

To evaluate whether the selective intramacrophage activity of **4-06** and **6-13** could translate to *in vivo* efficacy, both compounds were tested in an STm colitis murine model (**Fig. 4a**) (*48*). In this model, streptomycin-treated Balb/c mice are infected with STm by oral gavage (p.o). The initial enteric infection progresses into a systemic infection over a four-day period, at which time the mice are humanely euthanized and the bacterial burden of different organs is quantified. Treatment was started 24 h post-infection (h.p.i), and both compounds, which are stable at pHs ranging from 3.8 to 8.0 for at least 24 h (**Fig. S8**), were administered separately at 100 and 200 mg/kg p.o. twice daily. Administration of **4-06** resulted in a significant reduction in bacterial burden in the large intestine (1.4 log) at 200 mg/kg (**Fig. 4b**). In the case of **6-13,** significant reductions were seen in the spleen (2.0 log) at 100 mg/kg, as well as in the small intestine (2.2 log) at 200 mg/kg (**Fig. 4c**). Minor reductions could also be seen in the cecum (1.4 log, *p* = 0.065) and large intestine (1.2 log, *p* = 0.076) at 200 mg/kg. It appears that the intramacrophage activity of **4-06** and **6-13** similarly translated to *in vivo* efficacy. Given that these compounds selectively target intraphagocyte nutrient stress, it follows that the most pronounced burden reductions are expected in the tissues with larger proportions of lymphoid tissue, specifically, the spleen and the Peyer’s patches of the small intestines (*49*). One concern that may have limited the therapeutic potential of both compounds is their metabolism by serum pantetheinases. These are a known liability for pantothenamides like **6-13** (*14*), and thus is likely to be a factor limiting the bioavailability of **4-06** as well. Nonetheless, this is the first instance of pantetheinamides or pantothenamides displaying activity against STm *in vivo*.

**Fig. 4:**
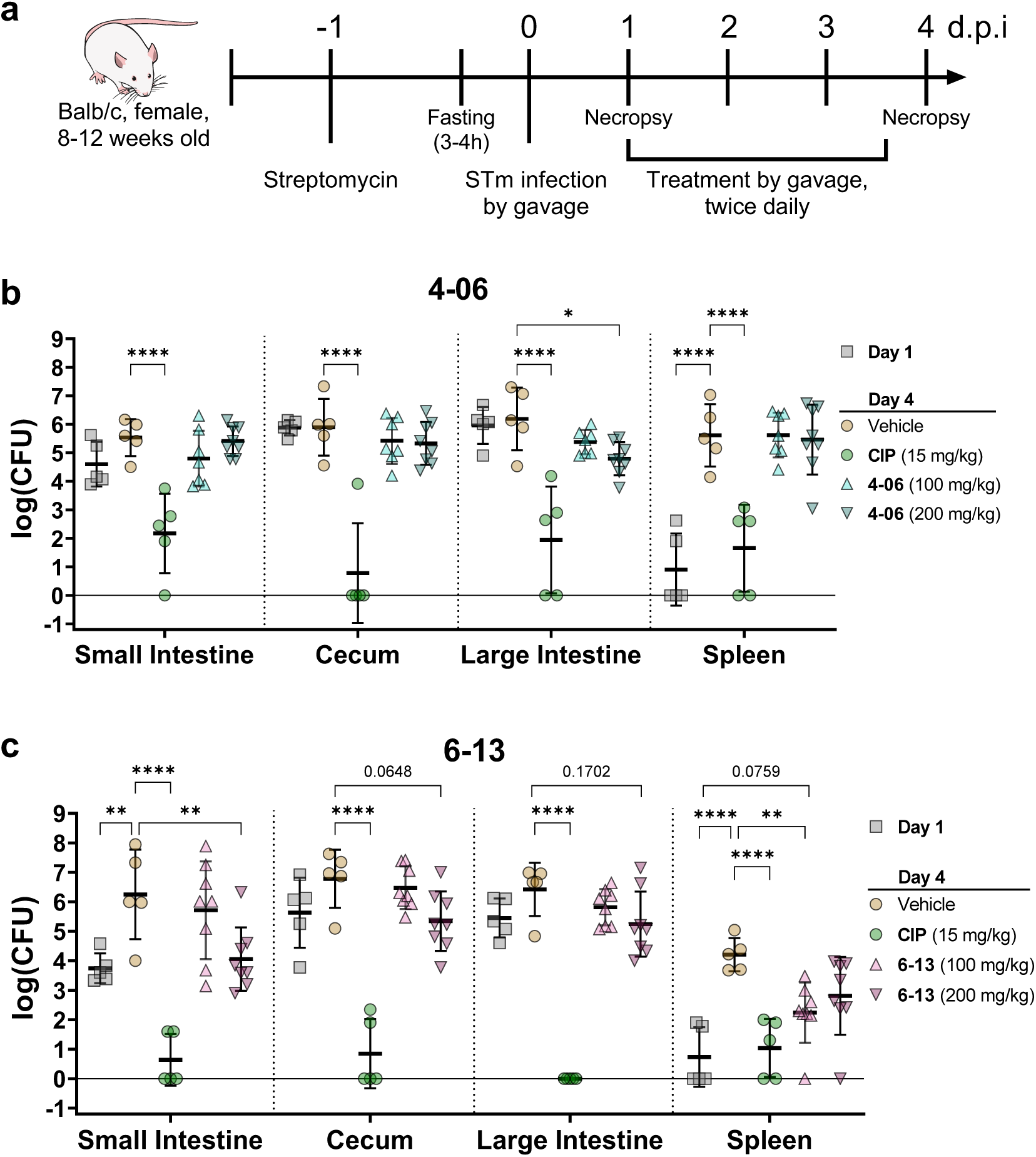
*In vivo* efficacy of 4-06 and 6-13 in an STm murine colitis model. **a,** Streptomycin-treated fasted Balb/c mice were infected with 1 × 10^9^ CFU of STm by gavage (*48*). Treatments with 100 and 200 mg/kg of either **b, 4-06** or **c, 6-13**, alongside their appropriate vehicle control and a ciprofloxacin (CIP) treatment at 15 mg/kg, were initiated at day 1 and administered twice daily for 3 days. Necropsy was done at day 1 (when treatment was initiated) and at day 4 after treatment. STm burdens in the small intestine, cecum, large intestine and spleen were measured by CFU counting. Data points depict the log(CFU) count per organ with bars showing the mean ± SD. n ≥ 5 mice per condition. Significant results were determined using a two-way ANOVA with Dunnett’s multiple comparisons against the vehicle. ✱, p < 0.05; ✱✱, p < 0.01; ✱✱✱, p < 0.001; ✱✱✱✱, p < 0.0001; Non-significant (*p* > 0.05) comparisons are either not shown or shown as the exact *p*-value.

## Discussion

Intracellular bacteria have a metabolic plasticity that is integral for growth and survival in diverse host cell environments (*50*). However, this feature is also a liability that can be exploited in the development of novel antibiotics. In nutrient-limited environments that are common at sites of infection, many metabolic pathways become essential for survival and are thus attractive targets for drug discovery (*51*). These deeply interconnected pathways, such as the biosynthesis of CoA or other vitamin-derived cofactors, present an opportunity for the development of pathogen-specific antibiotics, given that commensal populations exist in less stringent metabolic environments (*50*). Furthermore, designing drugs that synergize with the antimicrobial defences employed by our immune cells (*e.g.* itaconate) could result in effective and selective treatments with a slower development of resistance than broad-spectrum monotherapies.

CoA biosynthesis, in particular, has been linked to purine and thiamine biosynthesis; two essential pathways during STm pathogenesis (*51*, *53*). Previously, it was shown that STm *panB*/*E* mutants (that have reduced intracellular levels of CoA) show thiamine auxotrophy when flux through purine biosynthesis is low (*54*, *55*). Addition of pantothenate alleviated this thiamine requirement, linking the two pathways through purine biosynthesis. In addition to its other targets, itaconate has been shown to inhibit *de novo* purine biosynthesis in STm (*56*). As such, inhibition of pantothenate biosynthesis by pantetheinamides could synergize with the effects of itaconate through these connections.

In this study, a library of pantetheinamides and pantothenamides (known to be converted into CoA antimetabolites in bacteria) was synthesized as a means of potentially targeting the wide array of cellular functions that rely on CoA. A multi-step screen assessed their ability to inhibit the growth of STm experiencing various levels of nutrient stress, cytotoxicity to macrophages and capacity to reduce intramacrophage STm survival. Ultimately two compounds, **4-06** and **6-13**, were identified which reduced infection burden in a mouse colitis model. Through nutrient supplementation, metabolomics and resistance experiments, an apparent primary MoA for both molecules was identified: the inhibition of the PanD-PanZ complex that regulates CoA levels in STm and other enteric γ-proteobacteria (*37*, *57*). A recent study validated this target, by showing that STm *panD* knockout mutants have impaired intramacrophage replication and a reduced ability to colonize systemic infections in mice (*52*). Given that PanZ is not ubiquitous in bacteria, and not found in eukaryotes, it presents a selective target for the creation of narrow-spectrum antibiotics useful for the treatment of the many high-priority pathogens from this family that are known to be multi-drug resistant, including *E. coli*, *S.* Typhi, *Y. pestis*, *Shigella spp*. and *Klebsiella pneumoniae* (*5*, *57*).

Going beyond the pathogen-specificity of this approach, we also showed that there are limited environments where selective pressures exist for resistance to these compounds to develop. This theory is supported by the observation that resistance to **4-06** could only be generated under nutrient-limited conditions, while the same high concentration of **4-06** in rich media saw near negligible changes in susceptibility. Overall, the selective targeting of bacteria in pathogen-associated environments presents a promising strategy for the discovery of more effective therapies, reduced generation of antimicrobial resistance, and ultimately, better patient outcomes.

## Methods

### Chemicals, media and bacteria strains

All chemicals, antibiotics, BD Difco™ Nutrient broth/agar (NB/NA), and Mueller-Hinton broth/agar II (MHB/MHA) were purchased from Millipore-Sigma (Canada), Thermo Fisher Scientific (Canada) or Chem-Impex (USA) unless otherwise stated. AcCoA and CoA were purchased from CoALA Biosciences (USA). M9 media was prepared from the following components: M9 salts (48 mM Na_2_HPO_4_, 22 mM KH_2_PO_4_, 8.6 mM NaCl, 18.7 mM NH_4_Cl), 2 mM MgSO_4_, 0.1 mM CaCl_2_, trace metals (30 μM FeCl_3_, 25 μM Na_2_EDTA, 8 μM ZnCl_2_, 3 μM H_3_BO_3_, 3 μM MnCl_2_, 2 μM (NH_4_)_6_Mo_7_O_24_, 2 μM CuCl_2_, 2 μM CoCl_2_), and either 0.4% (w/v) sodium acetate for M9A, or 10 mM itaconic acid for M9I. The media was adjusted to pH 7.2 before filter sterilizing. *Salmonella* Typhimurium ATCC 14028, *E. coli* BL21 (DE3) and RAW 264.7 (TIB-71) macrophages were purchased from Cedarlane (Canada). The Δ*ich* STm mutant is from BEI Resources (www.beiresources.org) and was originally constructed by Porwollik *et al.* (2014) (*58*).

### General bacterial growth procedures

Bacteria were routinely grown by first streaking from a glycerol stock on nutrient agar plates (with 50 μg/mL kanamycin for the STm Δ*ich*) and grown overnight at 37⁰C. Generally, 3-5 colonies were picked and transferred to 5 mL of NB or MHB. Cultures were grown overnight at 37⁰C and 250 rpm. Stationary phase cultures were subcultured at a minimum 1/100 dilution into the appropriate fresh media and grown to a mid-exponential phase (OD_600_ ≈ 0.3-0.7) before use in experiments. Bacteria grown in minimal media were subcultured additional times (2 subcultures minimum in M9A and 3 subcultures minimum in M9I, unless otherwise stated) before use in experiments to allow for the bacteria to sufficiently adapt their metabolism to the media; improving the reproducibility of the experiments. ODs were monitored using either a Cary 60 UV-Vis spectrophotometer (Agilent Technologies) or a SpectraMax i3x microplate reader (Molecular Devices).

### Bacterial growth inhibition and checkerboard assays

Antimicrobial susceptibility was assessed using the traditional CLSI minimum inhibitory concentration (MIC) microplate dilution methods with a few modifications (*59*). Briefly, each compound (dissolved in MQ H_2_O for all pantetheinamides, or DMSO for pantothenamides) was serially diluted in 96-well plates to the desired concentrations. In the case of checkerboard assays, secondary compounds were separately diluted and added to the plate. Mid-exponential phase media-adapted bacteria were added at 5 × 10^5^ CFU/mL and in the case of M9I or M9A cultures, the plates were sealed with a gas permeable moisture barrier seal (FroggaBio, Canada) to prevent excessive evaporation, before incubation at 37⁰C and 350 rpm, using a tabletop orbital plate shaker (OHAUS, USA). Growth was quantified based on absorbance at 600 nm (OD_600_) using a microplate reader. The plate was monitored until there was visible growth in the untreated control, but without allowing for overgrowth: ∼14-16h (OD_600_ > 0.3) in MHB, ∼36 h (OD_600_ > 0.2) in M9A, and ∼72-120 h (OD_600_ > 0.15) in M9I. The IC_50_ was determined using a four-parameter logarithmic curve on Prism v9.0. All experiments were done at minimum in a biological triplicate.

### Macrophage infection assay

Raw 264.7 mouse macrophages were cultured in Dulbecco’s Modified Eagle Medium (DMEM) with 10% fetal bovine serum (FBS) and grown at 37⁰C in a 5% CO_2_ atmosphere. A cell scraper was used to lift cells off the flask when passaging (rather than trypsin). Macrophages were seeded in 96-well plates at 1 x 10^4^ cells per well (100 μL; measured by hemocytometer) and allowed to adhere for approximately 16 h before the experiment. On the day of the experiment, STm strains were subcultured 1/100 in NB and allowed to grow to mid-exponential phase. The culture was diluted in NB to an OD_600_ = 0.1 and diluted 1/10 into DMEM + 10% FBS to opsonize the bacteria. At this stage bacteria or macrophages or neither were pre-incubated with compound or vehicle (DMSO) at the desired concentrations (maintained throughout the experiment) and allowed to incubate statically for 3 h at 37⁰C and 5% CO_2_. Bacteria were then further diluted to achieve a multiplicity of infection (MOI) of 10 bacteria cells per macrophage cell in each well (in 100 μL). The plates were centrifuged at 500 × *g* for 2 min to sediment the STm cells and increase contact with the macrophages, before incubating at 37⁰C and 5% CO_2_. This was designated time zero. After 30 min of infection, the media was removed and replaced with fresh DMEM (10% FBS + compound) containing 100 μg/mL gentamicin to kill the extracellular bacteria. The plates were incubated for 30 min at 37⁰C in 5% CO_2_, before removing and replacing the media once again with DMEM (10% FBS + compound) with 10 μg/mL gentamicin. The infected cells were incubated for 22 h, unless otherwise stated. Surviving intracellular STm were quantified by CFU counting. Macrophages were first washed with phosphate-buffered saline (PBS; 2 × 200 µL per well), lysed with 0.1% Triton X-100 in PBS, scraped using a pipette tip and mixed by pipetting. The lysates were diluted as needed in PBS and plated on NA. To minimize random error, the experiment was performed in two separate and identical plates, the lysates of which were combined before plating. Within each plate, each condition was tested in technical duplicate, and the experiment was repeated independently a minimum of three times (n ≥ 6). The data was log normalized for analysis, outliers were removed using the ROUT method on Graphpad Prism and statistical differences were calculated using a one-way ANOVA with multiple comparisons against the vehicle-treated control.

### Cloning, expression, and purification of LpIct, LpIch, and LpCcl

The *Legionella pneumonia* genes encoding Ict (WP_129813656.1), Ich (WP_129813652.1), and Ccl (WP_129813654.1) were codon optimized for *E. coli,* synthesized by Biobasic (www.biobasic.com), and subcloned into the pBacP expression vector with an N-terminal TEV cleavable His_8_-tag (*60*). The expression plasmid was transformed into *E. coli* BL21 (DE3). A single colony was picked and cultured in 100 mL LB medium supplemented with 50 µg/mL kanamycin (KAN). Subsequently, 10 mL of an overnight culture was added to 1 L of LB (50 µg/mL KAN) and grown at 37°C until an OD_600_ of 0.6. The cultures were then cooled on ice for 30-45 min. Induction of protein expression was initiated by the addition of 0.5 mM (LpIct) or 0.1 mM (LpIch and LpCcl) isopropyl β-D-1-thiogalactopyranoside (IPTG) followed by incubation for 16 h at 20° C (LpIct) or 16° C (LpIch and LpCcl). After induction, the cells were harvested by centrifugation (JLA8.1000 rotor) at 4000 rpm for 25 min and stored at -80 °C until needed.

A typical protein purification protocol was used. Briefly, cell pellets were homogenized in IMAC binding buffer (50 mM Tris-HCl, 300 mM NaCl, pH 7.5, 2 mM β-mercaptoethanol (BME)), and 1 mM phenylmethylsulfonyl fluoride (PMSF), supplemented with a few crystals of DNase I (Bioshop) and one complete EDTA-free protease inhibitor tablet (Roche). Cells were lysed by sonication (5 min total pulse at 50% amplitude, 10 s on, 20 s off) followed by centrifugation at 18, 000 × *g* for 35 min to collect the supernatant. The supernatant was loaded on a pre-equilibrated (with binding buffer) 1 mL HisTrap HP (Cytiva) or 5 mL HisTrap IMAC FF column (GE Healthcare) connected to an AKTA purifier. Bound protein was washed with at least 20 mL of binding buffer containing 10 mM imidazole before elution with 30 mL of elution buffer A (50 mM HEPES, 200 mM NaCl, 250 mM imidazole, 2 mM BME, pH 7.4) for the 1 mL HisTrap HP, or on a 100 mL gradient to 100% elution buffer B (50 mM Tris-HCl, 300 mM NaCl, 300 mM imidazole, 2 mM BME, pH 7.5) and the fractions containing the desired protein were identified by SDS-PAGE. For LpIch and LpCcl, the affinity tag was removed by adding TEV protease (1 mg of TEV to 40 mg of protein) followed by overnight dialysis at room temperature against the buffer (50 mM Tris-HCl pH 7.5, 150 NaCl, 2 mM βME). The dialyzed sample was reapplied to the HisTrap column (pre-equilibrated with binding buffer) and the flow-through was collected. Eluted His-tagged LpIct or His-tag-cleaved LpIch and LpCcl were concentrated using an Amicon centrifugation concentrator (10 kDa, EMD Millipore) and further purified by size exclusion chromatography (SEC, HiLoad 16/600 Superdex 200 column, GE Healthcare) equilibrated with buffer (50 mM Tris-HCl pH 7.5, 150 NaCl, 0.3mM TCEP). The highest purity fractions were pooled, concentrated, flash-frozen and stored at -80 °C.

### HPLC purification of CoA thioesters

Coenzyme-A derivatives (RCoA) were routinely analyzed by HPLC (Agilent 1100 series, Agilent Technologies) with or without coupling to a mass spectrometry detector (MS, Agilent G6120B single quadrupole, ESI, Agilent Technologies). Separations were performed using an Agilent ZORBAX Sb-Aq column (250 mm × 4.6 mm; particle size 5 µm), with mobile phase A, consisting of Milli-Q H_2_O supplemented with 0.1% (v/v) HPLC-grade trifluoroacetic acid (TFA, Sigma-Aldrich) or 0.1% (v/v) LC/MS-grade formic acid (FA, Fisher Chemical) and mobile phase B, which consisted of pure HPLC-grade acetonitrile. A flow rate of 1 mL/min was used beginning at 99:1 (A:B) with a linear gradient to 70:30 over 18 min; a 1 min hold; a gradient to 1:99 from 19-24 min; followed by a hold for 2 min before returning to initial conditions. The retention times for the RCoA species, detected at 260 nm and confirmed by LR-ESI-MS (negative mode), were: 9.5 min for free CoA ([M-H]^-^ = 766.1), 11.7 min for (*S*)-citramalyl-CoA (CmCoA, [M-H]^-^ = 896.2), 11.8 min for acetyl-CoA (AcCoA, [M-H]^-^ = 808.1), and 13.7 min for itaconyl-CoA (ItaCoA, [M-H]^-^ = 878.1). Quantification of the RCoA species was achieved using an AcCoA calibration curve.

(*S*)-Citramalyl-CoA was synthesized enzymatically using Ict and Ich before purification by HPLC. Generally, the reaction contained 100 mM sodium phosphate (pH 7.2), 100 mM sodium itaconate (ItaNa_2_), 5 mM AcCoA, 1 µM Ict and 1 µM Ich. The reaction mixture was incubated at 37°C for 3 h, ensuring that all AcCoA had reacted before quenching with 1 M HCl at a ratio (v/v) of 1:5 (HCl:reaction mixture) and put on ice for ≥5 min. The mixture was centrifuged at 21 000 × *g* (4°C) for 5 min to remove precipitated protein, filtered (0.22 µm, nylon, Chromatographic Specialties), and separated by HPLC using the method above, collecting the CmCoA fractions, which were rapidly lyophilized.

### Enzymatic bioactivation of pantetheinamides and pantothenamides

Pantetheinamides and pantothenamides were successfully bioactived *in vitro* into the full CoA derivatives through the use of PanK, PPAT, and DPCK enzymes from *E. coli*, purified as described previously (*17*, *25*). The reaction buffer contained 25 mM Tris-HCl, 5 mM KCl, 2.5 mM MgCl_2_, 2.5 mM dithiothreitol (DTT), pH 7.2. The reactions contained 2.5 mM of compound (**4-06**, **6-13**, or pantetheine), 8 mM ATP (3.2 equivalents), and 1 µM each of PanK, PPAT, and DPCK. A no compound control instead contained 0.5 mM ATP and 5 mM ADP to match the expected product found in a 100% complete reaction. The reaction was left at rt for 24 h. Reaction products were separated from the protein by filtering through a Corning Spin-X UF concentrator (0.5 mL 10 kDa cutoff, Corning, USA) at 12 000 × *g* and rt for 15 min and collecting the protein-free supernatant. Twice, MQ H_2_O was added to the remaining fraction, which was centrifuged again to maximize recovery of the bioactivated product. The collected supernatants were lyophilized to concentrate the crude mixture. Samples were redissolved in MilliQ H_2_O, and analyzed by HPLC-MS to quantify the concentration the dephospho-CoA (dCoA) species (PPAT product) and full CoA species (DPCK product), by UV at 260 nm, based on an AcCoA standard curve. Products were separated using the method described for CoA thioester purification. Retention times at 260 nm were: 17.1 min for **4-06-**dCoA ([M-H]^-2^ = 402.1), 18.2 min for **4-06**-CoA ([M-H]^-2^ = 442.0), 22.0 min for **6-13**-dCoA ([M-H]^-1^ = 724.2), and 21.2 min for **6-13-**CoA ([M-H]^-1^ = 804.2). Crude reaction mixtures were used directly in enzymatic reactions.

### HPLC enzyme inhibition assays

The inhibition of LpIct, LpIch, and LpCcl by **4-06**, **6-13**, pantetheine or their bioactivated products were measured by HPLC. All reactions were carried out in 100 mM sodium phosphate (pH 7.2), and in the case of Ccl, with 5 mM MgCl_2_. Variable amounts of inhibitor were added at the stated concentrations. All reactions were performed at room temperature (rt, 25°C) for 15, 30, or 60 min and reactions were quenched as previously described. Ict reactions contained 200 nM LpIct, 100 mM ItaNa_2_ and 0.8 mM AcCoA. Reaction progress was monitored at 260 nm for the consumption of substrate (AcCoA) and formation of product (ItaCoA). Due to the instability of the Ich substrate, ItaCoA, purification was not feasible. As such, ItaCoA was synthesized enzymatically, and the crude product mixture was used directly to assay Ich. Briefly, ItaCoA was synthesized using 5 µM Ict, 100 mM ItaNa_2_, and 5 mM AcCoA, allowing the reaction progress at rt for 15 min and then separating the LpIct from the crude mixture by filtering through a Spin-X® protein concentrator (0.5 mL 10 kDa cutoff) at 12 000 × *g* and 4°C, 15 min. The flow-through was diluted to obtain a reaction containing ∼0.5 mM ItaCoA (exact concentration confirmed by HPLC each time) and 750 nM LpIch. Reactions were quenched as previously described, and analyzed by HPLC at 260 nm for the production of CmCoA. Ccl was assayed using 0.1 mM of purified CmCoA and 200 nM LpCcl. Reactions were quenched with HCl as described, and then phenylhydrazine was added to a final concentration of 10 mM to react with the Ccl product, pyruvate. Formation of the pyruvate phenylhydrazone proceeded at 37°C over 30 min, prior to HPLC analysis. Ccl reaction progress was measured by quantifying pyruvate phenylhydrazone at 324 nm. All results were repeated in triplicate for pure inhibitors or duplicate for bioactivated inhibitors. Data is presented as the percent inhibition compared to the vehicle control and were graphed using Graphpad Prism v9.0 (Graphpad Software, USA).

### Differential scanning fluorimetry of LpIch

Differential scanning fluorimetry (DSF) was performed on an Applied Biosystems StepOnePlus qPCR (Thermo Fisher). Samples (25 µL) were prepared in buffer (25 mM HEPES, 150 mM NaCl, 15 % PEG, pH 6.0) in 96-well (Applied Biosystem MicroAmp Potical) plates with 2 µM of protein, 1x SYPRO Orange dye (ThermoFisher Scientific), and CmCoA or crude bioactived **6-13** at 1 mM, where noted. The temperature program ramped from 5°C to 95°C at a 0.4 % ramp rate while monitoring fluorescence at ex = 480 nm and em = 568 nm. Data was analyzed using Protein ThermalShift™ software (ThermoFisher Scientific).

### STm growth and extraction for metabolomics

M9I-adapted STm (as described above) was washed once with M9I before resuspending to an OD_600_ of 0.1 in 50 mL of media in a 250 mL Erlenmeyer flask before incubating at 37⁰C, 250 rpm. Compounds **4-06** or **6-13** were added at 37 µM or 1.3 µM, respectively (10× their IC_50_ in M9I). Samples were taken at two phases of growth based on the OD_600_ plateau at stationary phase. Mid-exponential phase (EP) samples were collected at 0.3 × the plateau OD_600_ and early-stationary phase (SP) samples were collected at 0.9 × the plateau OD_600_. Sample across all conditions were normalized by the total amount of biomass collected. At each endpoint, enough culture was collected to obtain 13 mL of OD_600_ = 0.25, which was pelleted by centrifugation in a 15 mL tube at 14 000 × *g* and 4⁰C for 10 min. All but 1.5 mL of the supernatant was removed, then the cells were resuspended and transferred to a 2 mL microfuge tube to be pelleted again. The cells were washed 1× with ice-cold 150 mM ammonium formate, pH 7.4, before centrifugation at 21 000 × *g* and 4⁰C for 5 min. The supernatant was removed and the cells were frozen at -80 ⁰C for 1-2 days before extraction. A “no-cell” media control was prepared by following the same procedure with spent M9I media. This was done to discern between intracellular metabolites and those from residual media. Samples were kept on ice throughout the extraction process. Cell pellets were lysed and extracted by first resuspending them in 380 μL of 50% MeOH/H_2_O (v/v), before adding 220 µL of MeCN and 6 ceramic beads (1.4 mm) per sample. Cells were lysed and homogenized by bead-beating for 2 min at 30 Hz using a QIAGEN TissueLyser II (Germany). Next, 600 μL of ice-cold DCM and 400 μL of ice-cold water were added to each sample, and the mixture was vortexed for 1 min before allowing the metabolites to partition on ice for 10 min. Lastly, the mixtures were centrifuged at 1500 × *g* and 1⁰C for 10 min. A 700 μL aliquot of the aqueous layer was collected and dried by vacuum centrifugation (Labconco, USA) at -4°C before being stored at -80 ⁰C until analysis.

### Data collection for targeted metabolomics

Two separation and data collection methods were used for the targeted metabolomics of short chain CoA metabolites and a general ion pairing method. Chromatographic separation and MS/MS detection for both methods were completed using an Agilent 1290 Infinity ultra-performance quaternary pump liquid chromatography system coupled with a 6470 Triple Quadrupole (QQQ)–LC–MS/MS (Agilent Technologies, USA). The mass spectrometer was equipped with a Jet StreamTM electrospray ionization source, and samples were analyzed in negative mode. Samples were prepared by re-suspending the dry extracts in 50 μL of chilled H_2_O, clarified by centrifugation at 1°C, and transferred to glass inserts before injection. Sample injection volumes for analyses were 5 μL. They were first injected for the CoA collection method and then the ion pairing method. Multiple reaction monitoring parameters (qualifier/quantifier ions and retention times) were optimized and validated using authentic metabolite standards. Metabolite standards (Sigma-Aldrich) were run at multiple stages along the data acquisition workflow, which were used as quality control (QC) samples in the analysis.

For the CoA method, the source-gas temperature and flow were set at 300°C and 5 L/min, respectively, the nebulizer pressure was set at 45 psi, and the capillary voltage was set at 3,500 V. Separation was achieved using a XBridge C18 column 3.5 μm, 2.1 × 150 mm (Waters) with the gradient starting at 100% mobile phase A (10 mM ammonium acetate in water), followed by a 15 min gradient to 35% B (MeCN) at a flow rate of 0.2 mL/min. Next, an increase to 100% B over 0.1 min, a 4.9 min hold time at 100% mobile phase B and a subsequent re-equilibration time (6 min) at 100% A before the next injection.

For the general ion pairing method used a source-gas temperature and flow set at 150°C and 13 L min^−1^, respectively, the nebulizer pressure was set at 45 psi, and the capillary voltage was set at 2,000 V. Separation of metabolites was achieved by using a Zorbax Extend C18 column 1.8 μm, 2.1 × 150 mm with guard column 1.8 μm, 2.1 × 5 mm (Agilent Technologies). The gradient started at 100% mobile phase C (97% water, 3% methanol, 10 mM tributylamine, 15 mM acetic acid, 5 µM medronic acid) at a flow rate of 0.25 mL min^−1^ for 2.5 min, followed by a 5 min gradient to 20% mobile phase D (methanol, 10 mM tributylamine, 15 mM acetic acid, 5 µM medronic acid), a 5.5 min gradient to 45% D, a 7 min gradient to 99% D, and finally, a 4 min hold at 100% mobile phase D. The column was restored by washing with 99% mobile phase E (90% aqueous MeCN) for 3 min at 0.25 mL min^−1^, followed by increasing the flow rate to 0.8 mL min^−1^ over 0.5 min and holding for 3.85 min, before returning to 0.6 mL min^−1^ over 0.15 min. The column was then re-equilibrated at 100% C over 0.75 min, during which the flow rate was decreased to 0.4 mL min^−1^, and held for 7.65 min. One min before the next injection, the flow rate was brought back to 0.25 mL min^−1^. The column temperature was maintained at 35°C.

### Metabolomics analysis

Peak area data was carefully reviewed to ensure the accuracy and robustness of the analysis. Before analysis, ions with weak or no signal were removed, while injection blanks and the “no-cell” media controls were used to identify and remove metabolite signals that could not be resolved from the background. Metaboanalyst 6.0 (*61*) was used to perform all metabolomics analyses unless otherwise stated. The data (**Supplementary Data 3)** was filtered for reliability and to remove non-informative variables by excluding metabolites outside of the 5% interquantile range across all samples, and mean intensities within 5% of the limit of detection. The data was log_10_ transformed, auto scaled, and normalized to ensure a Gaussian distribution before analysis. Violin plots were made using Graphpad prism using the normalized abundances computed by Metaboanalyst.

### Soft agar gradient evolution (SAGE) of STm resistance to compound 4-06

SAGE evolutions were performed using M9I-adapted STm that had been subcultured in M9I 5 times (referred to as the “parent strain”) prior to storage at -80°C as 50% glycerol stocks. When starting cultures, frozen stocks were pelleted, washed 3× with M9I, and then resuspended in M9I. Minimum inhibitor concentrations (MICs) for the parent strain and all evolved strains were measured using the procedures previously described, but evaluating the MIC as the lowest concentration without visible growth. Four SAGE plate evolutions were performed: MHA alone, MHA with **4-06**, M9I alone, or M9I with **4-06**. MHA SAGE plates were prepared using established procedures (*43*). M9I SAGE plates were prepared by mixing a 1.5× concentrated stock of M9I media at a 2:1 ratio with an autoclaved mixture of 0.45% agar and 0.3% xanthan gum in ddH_2_O to form the 1× M9I SAGE medium (*62*). Evolutions with **4-06** contained a max concentration of 160 µM (5× MIC of the parent strain, 32 µM) pre-mixed with the appropriate SAGE medium. Experiments were initiated by inoculation in a line on the side of the plate where the concentration of antibiotic was lowest using 50 µL of an overnight culture. The wells were then covered with 2.5 mL of mineral oil to prevent desiccation and incubated at 37 °C. Cultures were allowed to grow throughout the plate (1 day for MHA alone and MHA with **4-06**, 6 days for M9I alone, and 8 days for M9I with **4-06**) and mutants were harvested by pipetting 20 μL of gel from within the last 1.5 cm of the well. The soft agar extract was cultured in 5 mL of the appropriate media and incubated overnight at 37 °C. Evolved strains were stored at -80 °C as 25% glycerol stocks. The DNA of the evolved strains was extracted using a BioBasic (Canada) EZ-10 genomic DNA miniprep kit. Whole genome sequencing was performed at SeqCenter (Pittsburgh, PA, USA) using the Illumina NovaSeq X Plus. The sequencing depth was on average 350×. The Breseq v0.37.1 pipeline (*63*) was used for variant calling with the polymorphism flag. The genomes were deposited on the NCBI database, Submission ID: SUB15249609 and BioProject: ID PRJNA1249526. The reference sequence for *Salmonella enterica* serovar Typhimurium ATCC 14028 used for variant calling is available at: genomes.atcc.org/genomes/814482244337467e?tab=overview-tab.

### *In vivo* treatment of a murine mouse model

To assess the *in vivo* activity of compounds **4-06** and **6-13** in a colitis model of STm in mice, a model previously described by Barther *et al* was used (*48*). Briefly, 10-12-week-old female Balb/c mice (JAX) were pre-treated by gavage with 0.1 mL streptomycin at a concentration of 200 mg/mL. Twenty hours following the streptomycin treatment, mice were fasted for 3-4 h before intragastric inoculation with STm. Fasting normalizes differences in colonization resulting from acute variation in feeding behaviour between mice for this model (*64*). A streptomycin-resistant clone was obtained by first passaging the parental ATCC 14028 strain in mice to allow adaptation to the intestinal environment. Gut tissues were collected, and one bacterial clone recovered from the gut was plated on Mueller-Hinton agar containing 200 μg/mL streptomycin to isolate resistant mutants, as previously described (*65*). The ATCC14028 streptomycin-resistant strain used for *in vivo* testing harbours the mutation C271T (p. Pro91Thr) in *rpsL* and has an MIC ≥ 250µg/mL to streptomycin. The genome was deposited on the NCBI SRA database: Submission ID: SUB15686508 and BioProject ID: PRJNA1338244. A bacterial suspension of STm ATCC 14028, resistant to streptomycin, was prepared at a concentration of 1 × 10^9^ CFU/mL in sterile PBS, and 0.2 mL of the bacterial suspension was administered by gavage. Treatment began 1 day after infection (Day 1). Mice were either treated with the vehicle controls (ddH_2_O or 20 % hydroxypropyl-β-cyclodextrin pH 7.0), ciprofloxacin (CIP, 15 mg/kg) or the tested compounds (100 and 200 mg/kg) twice daily by gavage (0.2 mL) for 3 days. Compound **4-06** was prepared in saline (0.9% NaCl in ddH_2_O), compound **6-13** was dissolved in 20% hydroxypropyl-β-cyclodextrin (pH 7.0), and CIP was prepared in either saline or pure ddH_2_O, for the respective treatments. The drug solutions were sonicated at room temperature for 15 min, aliquoted for daily use, and stored at –20 °C. Each day, a fresh aliquot was thawed and kept at 4°C during use. The day after the final treatment (Day 4), mice were humanely euthanized by CO_2_ exposure and serial dilutions of the gut (small intestine, cecum and large intestine, washed 3× in 5 mL PBS) and spleen homogenates were plated on MacConkey agar plates containing streptomycin at 100 µg/mL. This study was conducted following the animal ethics regulations at the Research Institute of the McGill University Health Center’s and performed under Animal Use Protocol 10184 (Lupien).

## Acknowledgements

We would like to thank Prof. Eric Brown at McMaster University for gifting us the Δ*ich Salmonella* Typhimurium mutant strain. Metabolites were analysed at the Rosalind and Morris Goodman Cancer Institute’s Metabolomics Innovation Resource, McGill University. This facility is supported by the Terry Fox Foundation, Quebec Breast Cancer Foundation, The Dr. R. John Fraser and Mrs. Clara M. Fraser Memorial Trust, and McGill University.

## Funding

This research was funded by the Canadian Institute of Health Research (CIHR grant PJT-166175) to K.A.* and M.A.B., the FRQ-funded Réseau Québecois de Recherche sur les Médicaments (RQRM grant 78855) to K.A.* and B.L.F. K.A.* and T.M.S are members of the Centre de recherche en biologie structurale, funded by FRQS Research Centre Grant #288558. NSERC, FRQS, and FRQNT are also thanked for scholarships to J.P. (CGSM and PGSD), F.R.C. (B2X Graduate Studentship), and C.B.L. (B2X Graduate Studentship), respectively. The funders had no role in study design, data collection and analysis, decision to publish, or preparation of the manuscript.

## Author Contributions

**Conceptualization**: JP, KA^1,10^*

**Data curation**: JP, FRC

**Formal analysis**: JP, FRC,

**Funding acquisition**: MAB, AL, TMS, BLF, KA^1,10^*

**Investigation**: JP, SDG, AC, SHM, FRC, DD, CBL, RK, ABA, SC, AP^1^, IR, AL, KA^7^, SG, AP^7^

**Methodology**: JP, FRC, BLF, AL, KA^1,10^*

**Project administration**: KA^1,10^*

**Supervision**: MAB, AL, TMS, BLF, KA^1,10^*

**Validation**: JP, KA^1,10^*

**Visualization**: JP

**Writing –original draft: JP**

**Writing –review & editing**: JP, SDG, SHM, FRC, DD, CBL, SC, IR, MAB, AL, TMS, BLF, KA^1,10^*

## Competing Interests

The authors declare that they have no competing interests.

## Data Availability Statement

The genomic data from the STm resistance experiments are available at the NCBI Sequence Read Archive under BioProject PRJNA1249526 and BioProject PRJNA1338244. All other data needed to evaluate the conclusions in the paper are present in the paper and/or the Supplementary Materials.

## Notes

### Competing Interest Statement

The authors have declared no competing interest.

