## Supporting Information #1 for "Pantetheinamides that inhibit the growth of intracellular *Salmonella* Typhimurium"

### Table of Contents

**Supplementary Text 1** Regarding Tables S1 and S2

**Supplementary Text 2** Regarding Tables 1 and 2

**Supplementary Text 3** Regarding Tables 1, 2, S3, and S4

**Fig. S1** All $IC_{50}$ curves for the compounds tested against STm in M9I, where the final IC50 values are shown in the Fig 3C heatmap.

**Fig. S2** Structure-activity relationships (SARs) for the inhibitor library.

**Fig. S3** Mutations associated with the M9I-adaptation do not result in increased susceptibility to pantetheinamides or pantothenamides.

**Fig. S4** Treatment of RAW 264.7 macrophages infected with STm.

**Fig. S5** Itaconate and pantetheinamides do not exhibit interactive effects that would indicate itaconate degradation inhibition as a possible MoA.

**Fig. S6** The activity of pantetheinamides and pantothenamides are not the result of direct itaconate degradation inhibition.

**Fig. S7** Resistance generation to compound 4-06 by SAGE.

**Fig. S8** Stability of compounds 4-06 and 6-13 in buffer at various pH values.

**Table S1** Substantive mutations (>50%) in M9I-adapted parent STm strain used for SAGE experiments.

**Table S2** Genes encompassed in the [rpoS]–[mutS] Δ15 401 bp mutation.

**Table S3** MIC shift after control SAGE experiments without the presence of 4-06.

**Table S4** Mutations identified in the control SAGE experiments without the presence of 4-06.

**References**

**Supplementary Text 1: Regarding Tables S1 and S2**

M9I-adapted STm was used as the parent strain for all SAGE experiments. The genomic mutations associated with M9I-adaptation were identified (**Tables S1 and S2**). The most substantial mutation observed was a 15 401 base pair deletion between the genes for the stationary phase alternative sigma factor, *rpoS,* and the mismatch DNA repair protein, *mutS*, along with the 14 genes in between (**Tables S2**). Our previous work exploring STm adaptation to M9I identified that mutations resulting in a non-functional RpoS were acquired to completion within three subcultures in M9I.[1] These results were consistent with the literature understanding that mutations in *rpoS* are often selected for by enteropathogenic bacteria when grown in nutrient limiting conditions.[2–4] Mutations in *mutS* have also been associated with these conditions, where the bacteria increase their propensity for mutations that confers a higher ability produce *de novo* mutations and adapt in nutrient limiting environments.[3] While the complete deletion of the *rpoS*-*mutS* region observed in the parent strain for the SAGE experiments is a dead end mutation, the fact that this is only in 97% of the population indicates that a fraction of the population maintains this genomic region.

**Supplementary Text 2: Regarding Tables 1 and 2**

The third mutation identified that is associated with the resistant phenotype is a nonsense mutation in the marT transcription factor at Gln 181. MarT is a ToxR-family transcription factor that has been shown to positively regulate gut colonization and biofilm formation, and negatively regulate motility.[5–7] As such, it could be that increased motility is the fitness benefit of a non-functional MarT, allowing for improved movement across the SAGE plates, rather than a direct effect on the activity of **4-06**. The fourth mutation is a SNP in the *arcB* gene, encoding the sensor kinase in the ArcAB aerobic response two-component system that controls a large regulon involved in both metabolism and virulence.[8,9] This SNP is found in the linking region between the central receiver domain and the phosphotransfer domain of the catalytic region, however, the exact effect of this mutation could not be surmised from previous studies. The fifth mutation is a frameshift deletion of one base pair in the *iscR* gene, which encodes an iron-sensing transcriptional regulator. Interestingly, we previously identified that mutations in the IscR binding site upstream of the gene for the dicarboxylate transporter DctA results in reduced IscR repression that was beneficial for growth in M9I.[1] This was also noted for other DctA substrates.[10,11] It is therefore plausible that the benefit of IscR deletion is derived indirectly through halting IscR repression of DctA, resulting in improved itaconate uptake and catabolism, which may be protective against the effects of **4-06**.

**Supplementary Text 3: Regarding Tables 1, 2, S3, and S4**

In the wt STm strain, none of the resistance associated mutations (in *panB, yfdC*, *marT*, *arcB*, and *iscR*) were identified by sequencing, indicating that they are either *de novo* mutations or found in very low frequencies within the population. In the M9I-adapted parent strain used for the SAGE experiments, the *panB, yfdC*, and *marT* mutations were found at 4%, 3% and 5%, respectively, while the *arcB* and *iscR* mutations were not found. This highlights that the *panB, yfdC*, and *marT* mutations are beneficial at low frequencies for growth under nutrient limiting conditions in M9I liquid cultures. In the case of *panB*, this mutations has been shown to not only increase intracellular pantothenate levels, but also extracellular pantothenate levels.[12] As such, it is possible that the 4% frequency of this mutation in liquid M9I cultures is sufficient to supply the pantothenate for the whole population. However, during growth on agar, the ability of bacteria to share nutrients is diminished. Following this, the enrichment of the *panB* mutation across all of the control evolutions in M9I (without **4-06**) to ≥ 20% could be due to a generalized increased need for pantothenate under these conditions. Importantly, the enrichment difference between replicates 1 and 2 at approximately 20% to replicates 3 and 4 with 43% and 63%, respectively, is where resistance to **4-06** is acquired. In replicate 3, both *yfdC* and *marT* are also enriched to approximately 50% each, however, this is not found in replicate 4, indicating that they are not necessary for resistance.

In the case of the *arcB* and *iscR* mutations, these are not found in the parent strain, the MHA with **4-06** evolutions, nor the control evolution strains (M9I and MHA). While this suggests that they are not necessary for resistance to **4-06**, given the fact that they appear in all four evolutions in M9I cultures with **4-06** at high frequencies and from a starting point below the sequencing threshold (or as *de novo* mutations), reinforces that they do contribute to a resistance phenotype. The full extent of this resistant phenotype, and of those for the other identified resistance-associated mutations will need to be examined using single-gene mutants in future studies.

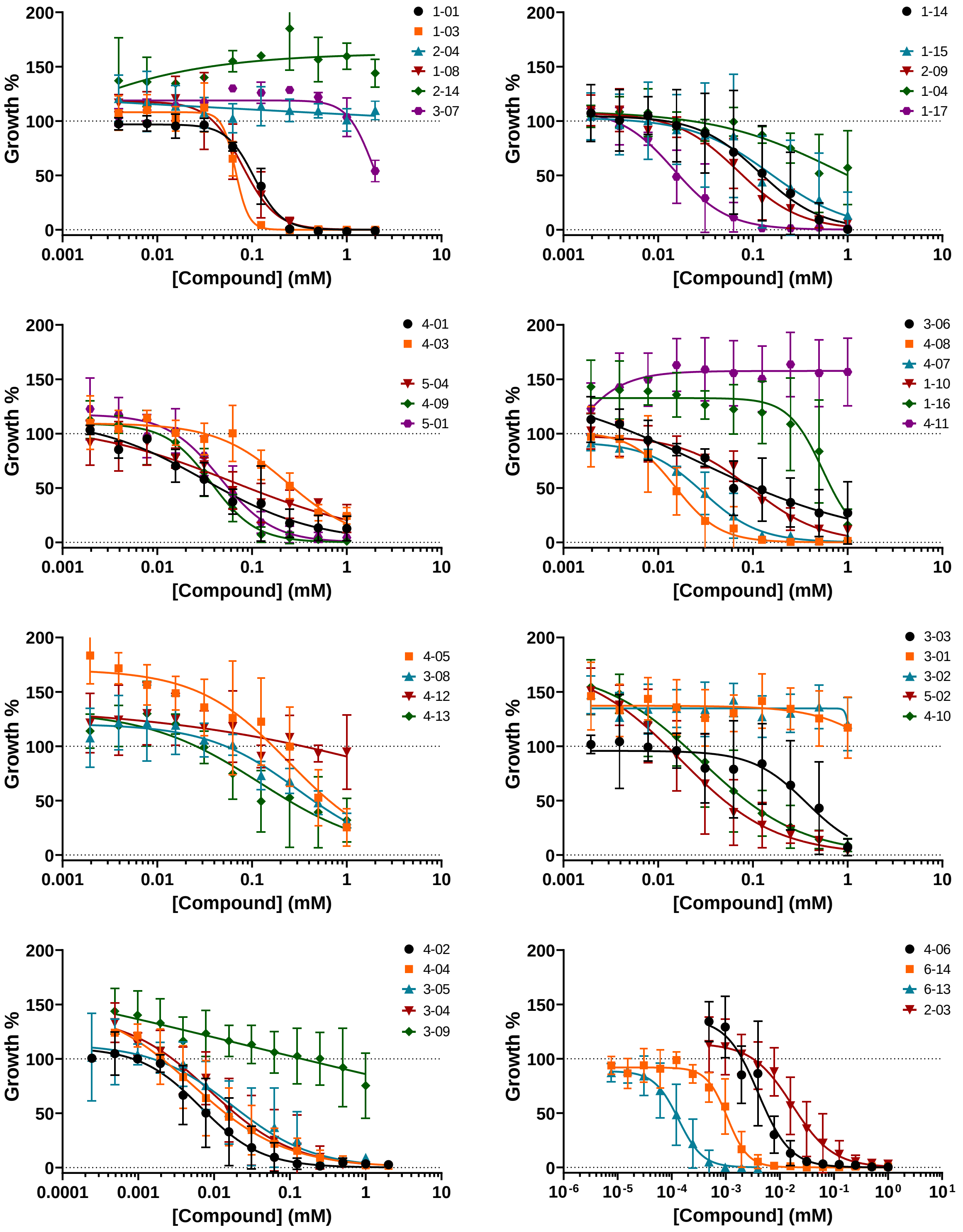


**Fig. S1: All $IC_{50}$ curves for the compounds tested against STm in M9I.** The final $IC_{50}$ values are shown in the Fig 3C heatmap and Supplementary Data 2.

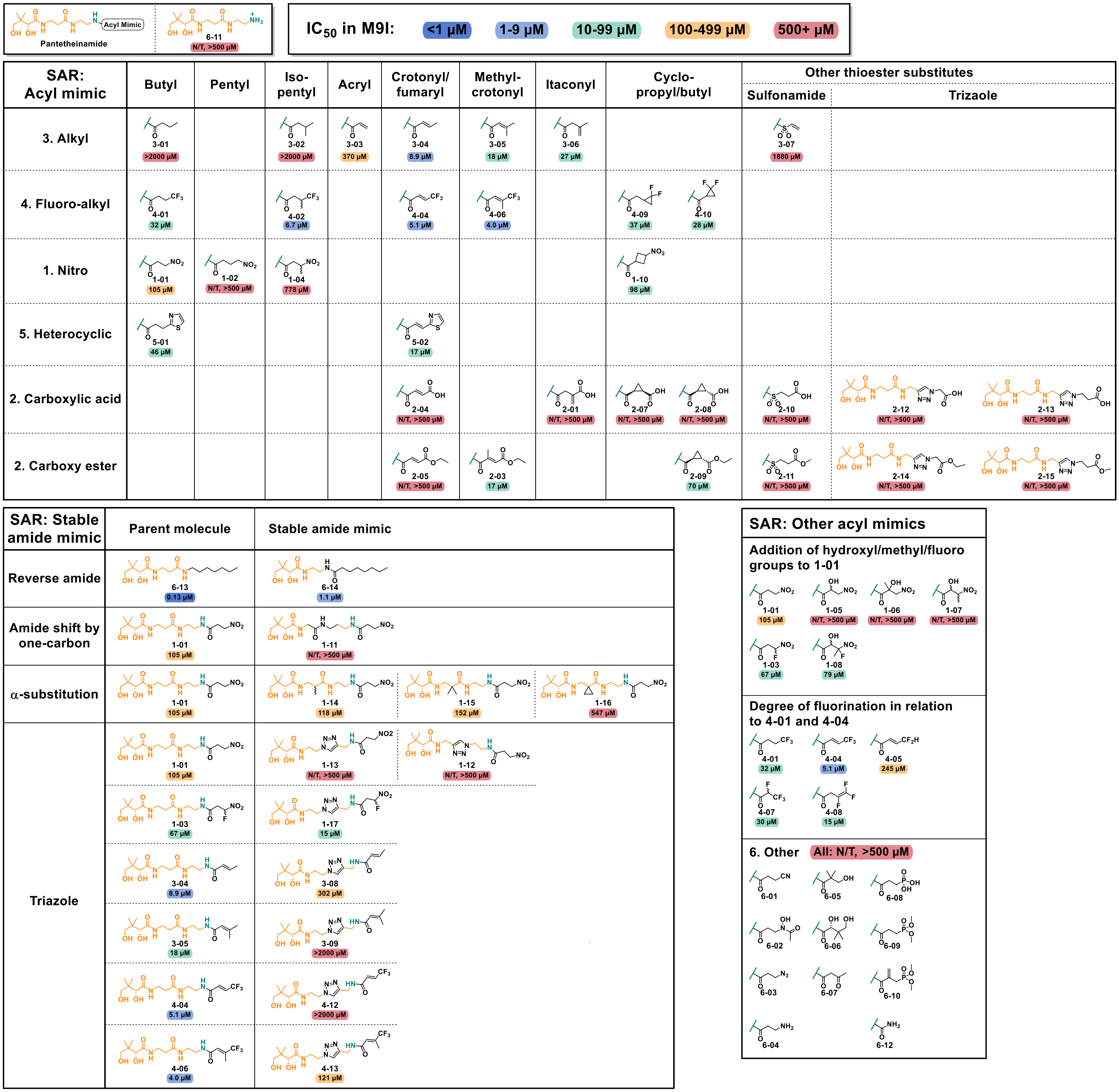


**Fig. S2: Structure-activity relationships (SARs) for the inhibitor library**. The SARs are based on the $IC_{50}$ and % inhibition of each compound against STm grown in M9I. The $IC_{50}$ values are color-coded into 5 ranges to highlight changes in activity due to structural changes, as listed in the legend. In cases where molecules did not show activity in the 500 µM screen in M9I and the $IC_{50}$ was not tested (N/T), then the $IC_{50}$ values are listed as > 500 µM.

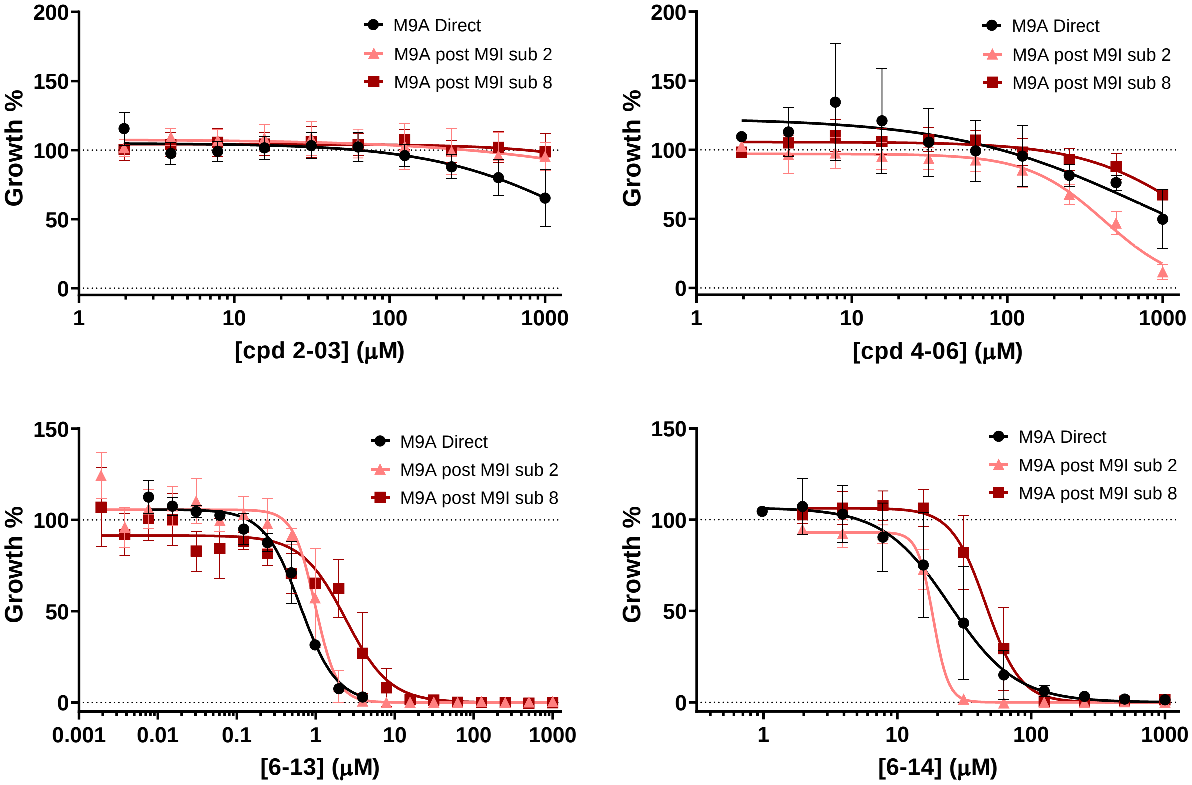


**Fig. S3**: **Mutations associated with the M9I-adaptation do not result in increased susceptibility to pantetheinamides or pantothenamides.** Compound dose-response curves tested in M9A using STm that was either not adapted to M9I (M9A direct), or subcultured twice (M9A post M9I sub 2) or 8 times (M9A post M9I sub 8) in M9I prior to assessing the dose-response in M9I.

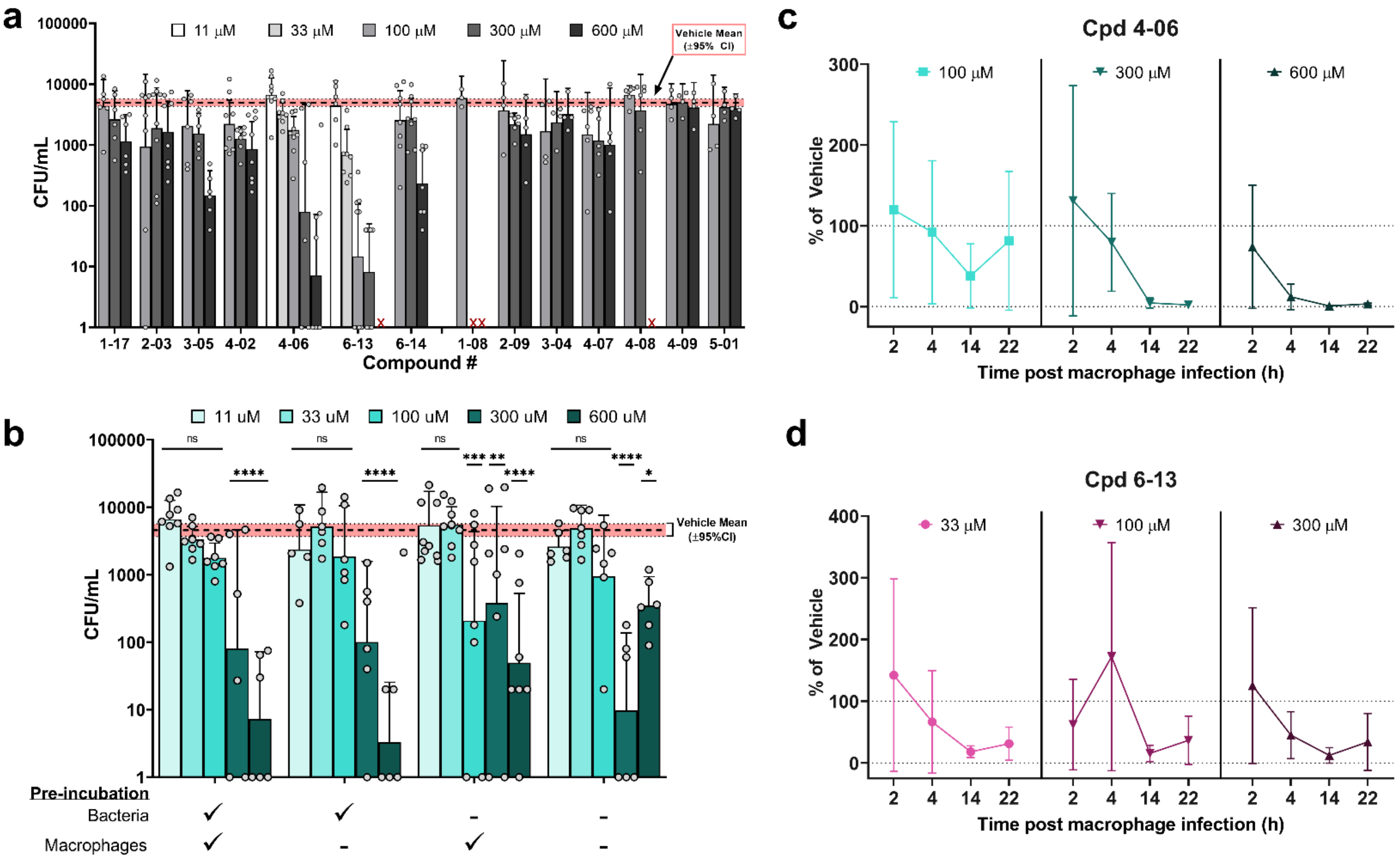


**Fig. S4**: **Treatment of RAW 264.7 macrophages infected with STm**. **a,** The compounds were tested by gentamicin protection assay, where the surviving STm inside the macrophages are quantified by CFU counting after 22 h. Both the bacteria and macrophages were incubated with compound for 3 h prior to infection. Compounds shown in the main text are to the left of **1-08**. **b,** Testing the effect of pre-incubation with or without bacteria and/or macrophages on compound activity using **4-06**. **c,** a time course of the intramacrophage STm population in response to **4-06** and **d, 6-13**, are shown at three different concentrations over the course of 22 h. Data bars (**a, b**) show the mean CFU/mL, data points (**c, d**) show the mean of each CFU count as % of the vehicle, and error bars depict the 95% CI. The vehicle mean (± 95% CI) is shown as the red line (**a, b**). All experiments were independently repeated a minimum of 3 times, with n ≥ 4 for each condition. Significant results were determined using a one-way ANOVA with Dunnett's multiple comparisons against the vehicle. ∗, $p < 0.05$; ∗∗, $p < 0.01$; ∗∗∗, $p < 0.001$; ∗∗∗∗, $p < 0.0001$; ns, non-significant, $p > 0.05$. The red "X" denotes concentrations that were not tested due to cytotoxicity.

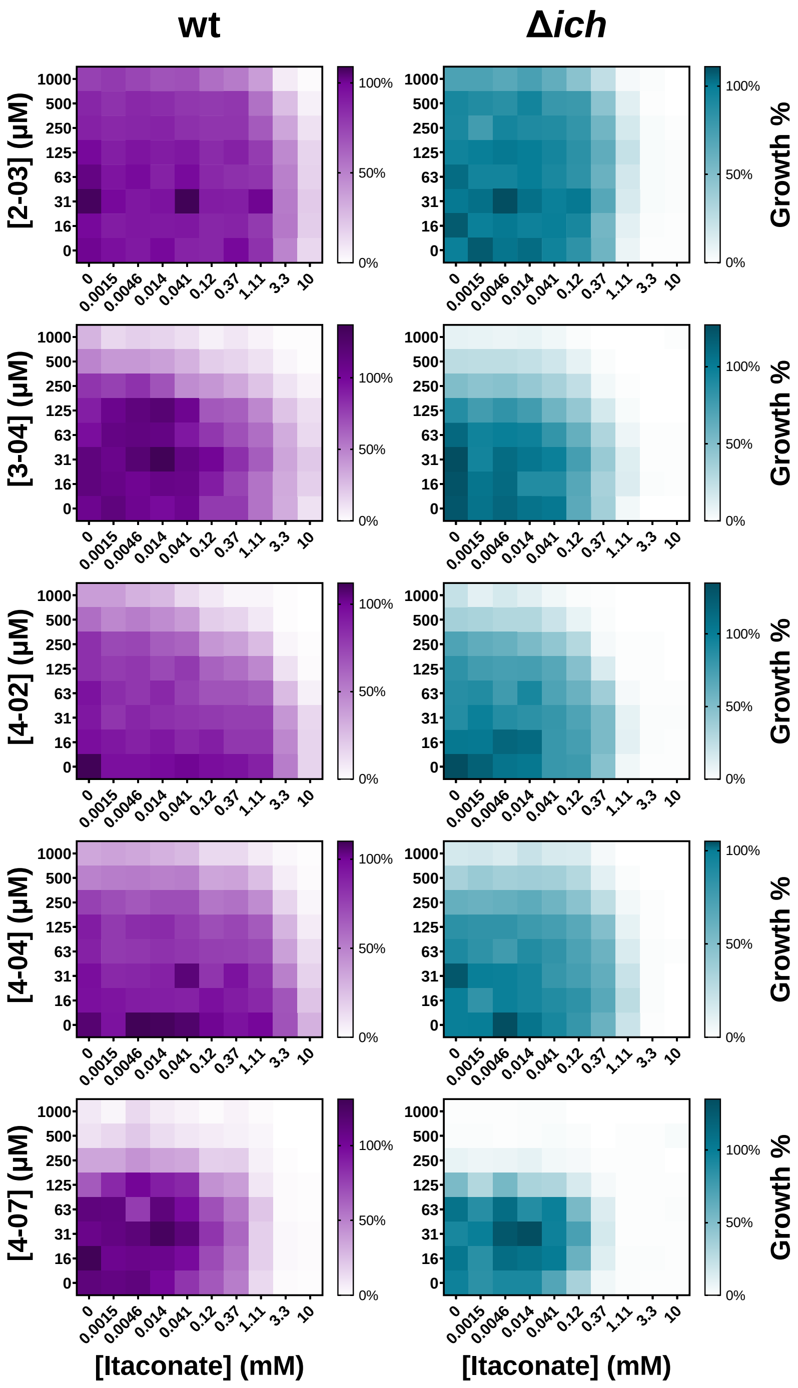


**Fig. S5: Itaconate and pantetheinamides do not exhibit interactive effects that would indicate itaconate degradation inhibition as a possible MoA**. Checkerboard assays with various compounds in M9A tested against both STm wt and Δ*ich* mutant. Synergy between both itaconate and inhibitor in STm wt would suggest that itaconate-degradation inhibition is at least one of the mechanisms of action. No synergy suggests the opposite. If synergy is seen in the wt, but lost in the IRO mutant (Δ*ich*), this would support itaconate-degradation inhibition as the mechanism, while the opposite would be true if synergy remains (indicating that another mechanism is dominant). Across all cases, little to no synergy is observed between the compounds and itaconate in the wt strain. Some additive effects exist at high concentrations of compound, but the trend appears the same in the IRO mutants as well. This supports that a mechanism other than inhibition of itaconate-degradation is the dominant mechanism of action for these compounds. Furthermore, the observation that some of these molecules have activity in M9A (in some cases needing to go to 1 mM or higher to begin to see the effect) again supports the theory that an alternative mechanism of action, rather than itaconate-degradation inhibition, is the dominant mechanism of action.

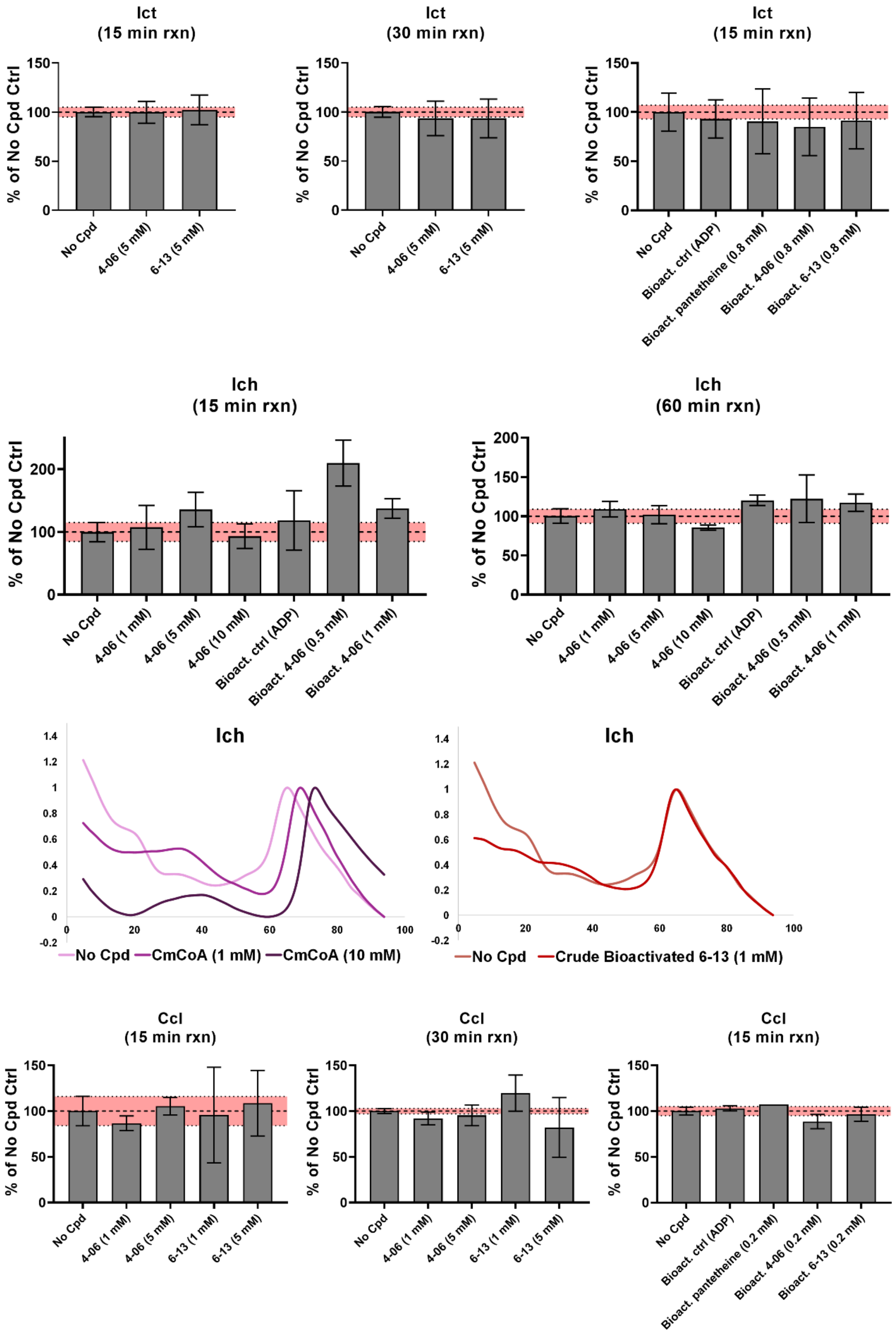


**Fig. S6**: **The activity of pantetheinamides and pantothenamides are not the result of direct itaconate degradation inhibition.** The *Legionella pneumophila* homologous enzymes of STm Ict (46% ID/65% similarity), Ich (57% ID/74% similarity), and Ccl (39% ID/59% similarity) were used due to poor expression of the STm enzymes. Inhibition was tested by measuring product formation for each enzyme by HPLC (bar graphs) using molecules **4-06** (all 3 enzymes) or **6-13** (Ict and Ccl) and their bioactivated CoA ("Bioact.") derivatives at the concentrations noted in brackets. Data bars show the % activity of the enzyme for each compound compared to that of the no compound (No Cpd) control set to 100%. Error bars depict the SD. The mean of the No Cpd control (± SD) is shown as the red band. $n \geq 3$ for each condition. All compound showed non-significant inhibition ($p > 0.05$) as determined by a one-way ANOVA with Dunnett's multiple comparisons against the No Cpd. Bioactivated **6-13** was tested against Ich by DSF and compared with the product, CmCoA. No change in the melting temperature ($T_m$) of the enzyme was observed with bioactived **6-13**. XY charts depict the fluorescence intensity (y-axis) over the temperature (x-axis) from 5 – 95°C.

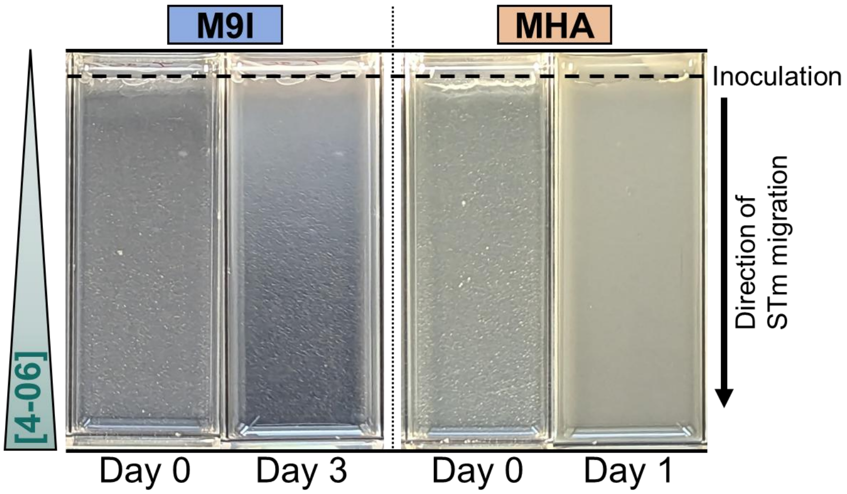


**Fig. S7: Resistance generation to compound 4-06 by SAGE**. SAGE plate set-up where the concentration of **4-06** increases from the inoculation site to the opposite end where there is a maximal concentration of 5x the MIC (32 µM x 5 = 160 µM). Evolutions were performed on M9I agar and Mueller-Hinton agar (MHA) both with or without **4-06**. n = 4 evolutions. The images depict an example of **4-06**-containing plates on day 0 when the plate is inoculated and in the case of MHA, the final day when samples were collected for sequencing and MIC testing. In the case of M9I, this depicts partial growth after only 3 days, whereas the full experiment and generation of resistant strains took 8 days. STm growth can be seen as the opaque gradient in the Day 3 and Day 1 plates.

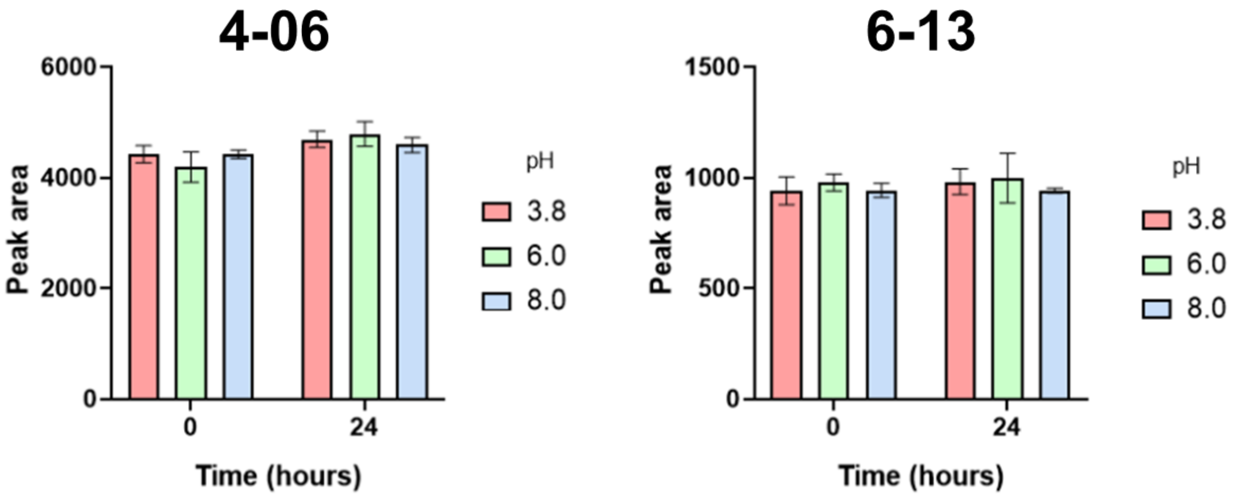


**Fig. S8: Stability of compounds 4-06 and 6-13 in buffer at various pH values.** Compounds were dissolved at 5 mM in buffers of three different pH and the concentration of the compounds was measured by HPLC after 0 h and 24 h at 37°C. Buffers at pH 3.8 and 6.0 were a mix of citrate (100 mM) and phosphate (200 mM), while the buffer at pH 8.0 was phosphate (200 mM) alone.

**Table S1: Substantive mutations (> 50%) in M9I-adapted parent STm strain used for SAGE experiments.**

| Gene (strand) | Mutation | Type | Frequency | Gene Product |
|---|---|---|---|---|
| **[rpoS]–[mutS]** | **Δ15 401 bp** (236/993 nt) | **Deletion** | 97.2% | **RpoS**: RNA polymerase sigma factor **MutS**: DNA mismatch repair protein |
| **barA←** | **C615F** (TGC→TTC) | **SNP** | 100% | hybrid sensory histidine kinase BarA (part two-component regulatory system with UvrY) |
| **nadB←** | **I530F** (ATC→TTC) | **SNP** | 100% | L-aspartate oxidase (quinolinate synthetase B). |
| **flhB→** | **L94L** (CTA→TTA) | **SNP** | 100% | flagellar biosynthetic protein FlhB |
| **gbpR_2←** | **+T** coding (5/924 nt) | **Frameshift insertion** | 100% | putative LysR family transcriptional regulator |
| **STM1620→** | **D196N** (GAT→AAT) | **SNP** | 100% | Putative oxidase, lactate-2-monooxygenase (PALNJAND_02238) |
| **phoP→** | **R139H** (CGC→CAC) | **SNP** | 100% | transcriptional regulatory protein PhoP, regul ator of virulence determinants in two-component system with PhoQ |
| **sfmH_1←** | **I157F** (ATT→TTT) | **SNP** | 100% | minor fimbrial subunit |
| **crl←** | **+A** coding (96/402nt) | **Frameshift insertion** | 100% | curlin genes transcriptional activator; interact s with sigma factor RpoS |

**Table S2: Genes encompassed in the [rpoS]–[mutS] Δ15 401 bp mutation.**

| Gene (strand) | Gene Product |
|---|---|
| **rpoS** | RNA polymerase sigma factor RpoS |
| **bsdD** | Involved in the non-oxidative decarboxylation and detoxification of phenolic derivatives |
| **edcC** | Involved in the non-oxidative decarboxylation and detoxification of phenolic derivatives |
| **ecdB** | catalyzes the synthesis of the prenylated FMN cofactor (prenyl FMN) for phenolic acid decarboxylase C |
| **hosA** | Transcriptional regulator HosA, may be involved in swimming motility and cellular aggregation |
| **glcR_1** | Transcription regulator, possible repressor of the lactose catabolism operon. |
| **ltnD** | Catalyzes oxidation of L-threonate to 2-oxo-tetronate. |
| **otnK** | Catalyzes the ATP-dependent phosphorylation of 3-oxo-tetronate to 3-oxo-tetronate 4-phosphate. |
| **otnC** | 3-oxo-tetronate 4-phosphate decarboxylase OtnC |
| **otnI** | 2-oxo-tetronate isomerase |
| **denD** | D-erythronate dehydrogenase |
| **gntT** | Part of the gluconate utilization system Gnt-I; high-affinity intake of gluconate. |
| **dmlR** | Transcriptional regulator required for the aerobic growth on D-malate as the sole carbon source. |
| **STM2911** | Hypothetical protein |
| **mdtD** | putative multidrug efflux pump |
| **mutS** | DNA mismatch repair protein |

**Table S3: MIC shift after control SAGE experiments without the presence of 4-06.**

| SAGE Exp. | MIC (µM) | | | |
|---|---|---|---|---|
| | **Rep[a] 1** | **Rep[a] 2** | **Rep[a] 3** | **Rep[a] 4** |
| **Parent Strain** | 32 (range: 8 – 64 [b]) | | | |
| **M9I (No 4-06)** | 8 | 32 | 512 | >512 |
| **MHA (No 4-06)** | 32 | 32 | 32 | 32 |

[a] Each replicate (rep) refers to independent evolutions under each condition.

[b] Range of MICs found across 5 repeated measurements of the parent strain MIC. The median and mode MIC is 32 µM.

**Table S4: Mutations identified in the control SAGE experiments without the presence of 4-06.**

| SAGE Exp. | Gene (strand) | Mutation | Type | Frequency [a] | | | | Gene Product |
|---|---|---|---|---|---|---|---|---|
| | | | | Rep 1 | Rep 2 | Rep 3 | Rep 4 | |
| Parent Strain [b] | ***folK* → / → *panB*** | Intergenic **+C** (+66/-55) [c] | **Insertion** | 4% | | | | **FolK**: pyrophosphokinase **PanB**: hydroxymethyltransferase |
| | ***yfdC* ←** | **A61T** (GCT→ACT) | **SNP** | 3% | | | | **YfdC**: Putative transporter |
| | ***marT*→** | **Q191X** (CAA→TAA) | **Nonsense** | 5% | | | | **MarT**: ToxR-family transcription factor |
| | ***arcB* →** | **T646I** (ACC→ATC) | **SNP** | 0% | | | | **ArcB**: Aerobic respiration control sensor (with ArcA) |
| | ***iscR* →** | **Δ1 bp** (380/495 nt) | **Frameshift deletion** | 0% | | | | **IscR**: Iron sensing transcription factor |
| M9I (No 4-06) | ***folK* → / → *panB*** | Intergenic **+C** (+66/-55) [c] | **Insertion** | 20% | 21% | 43% | 63% | **FolK**: pyrophosphokinase **PanB**: hydroxymethyltransferase |
| | ***yfdC* ←** | **A61T** (GCT→ACT) | **SNP** | 4% | 6% | 48% | 3% | **YfdC**: Putative transporter |
| | ***marT*→** | **Q191X** (CAA→TAA) | **Nonsense** | 5% | 5% | 51% | 4% | **MarT**: ToxR-family transcription factor |
| | ***arcB* →** | **T646I** (ACC→ATC) | **SNP** | 0% | 0% | 0% | 0% | **ArcB**: Aerobic respiration control sensor (with ArcA) |
| | ***iscR* →** | **Δ1 bp** (380/495 nt) | **Frameshift deletion** | 0% | 0% | 0% | 0% | **IscR**: Iron sensing transcription factor |
| MHA (No 4-06) | ***folK* → / → *panB*** | Intergenic **+C** (+66/-55) [c] | **Insertion** | 5% | 3% | 3% | 3% | **FolK**: pyrophosphokinase **PanB**: hydroxymethyltransferase |
| | ***yfdC* ←** | **A61T** (GCT→ACT) | **SNP** | 1% | 3% | 1% | 3% | **YfdC**: Putative transporter |
| | ***marT*→** | **Q191X** (CAA→TAA) | **Nonsense** | 2% | 2% | 1% | 4% | **MarT**: ToxR-family transcription factor |
| | ***arcB* →** | **T646I** (ACC→ATC) | **SNP** | 0% | 0% | 0% | 0% | **ArcB**: Aerobic respiration control sensor (with ArcA) |
| | ***iscR* →** | **Δ1 bp** (380/495 nt) | **Frameshift deletion** | 0% | 0% | 0% | 0% | **IscR**: Iron sensing transcription factor |

[a] Percentage of sequencing reads that identified the specific mutation among all reads. Rep: evolution replicate.

[b] The background level of mutations in the parent strain used in evolution experiments to compare with the post-SAGE results.

[c] Found in the promoter region of *panB*.
