## Supporting Information #2 for "Pantetheinamides that inhibit the growth of intracellular *Salmonella* Typhimurium"

#### Table of final compounds

| Cpd # in paper | Cpd # in methods |
|---|---|
| 1-01 | 6a (RK) |
| 1-02 | DD-06-143 |
| 1-03 | DD-07-066 |
| 1-04 | 6i (RK) |
| 1-05 | DD-04-021 |
| 1-06 | DD-03-001 |
| 1-07 | DD-03-085 |
| 1-08 | DD-04-011 |
| 1-09 | 6h (RK) |
| 1-10 | BCL-5-67 |
| 1-11 | DD-06-070 |
| 1-12 | DD-04-109 |
| 1-13 | DD-07-076 |
| 1-14 | 22a (RK) |
| 1-15 | 22b (RK) |
| 1-16 | 22c (RK) |
| 1-17 | DD-07-102 |
| 2-01 | DD-07-139 |
| 2-02 | 7m (RK) |
| 2-03 | BCL-7-31 |
| 2-04 | DD-06-147 |
| 2-05 | DD-04-007 |
| 2-06 | DD-07-129 |
| 2-07 | 12 (RK) |
| 2-08 | 9k (RK) |
| 2-09 | 6k (RK) |
| 2-10 | 15 (RK) |
| 2-11 | 16 (RK) |
| 2-12 | DD-07-141 |
| 2-13 | DD-08-001 |
| 2-14 | DD-07-081 |
| 2-15 | DD-07-015 |
| 3-01 | SCC-3-103 |
| 3-02 | SCC-3-102 |
| 3-03 | BCL-5-9 |
| 3-04 | BCL-5-41 |
| 3-05 | BCL-7-11 |
| 3-06 | BCL-4-95 |

| Cpd # in paper | Cpd # in methods |
|---|---|
| 3-07 | BCL-4-154 |
| 3-08 | ABA-1-18 |
| 3-09 | BCL-7-75 |
| 4-01 | 6b (RK) |
| 4-02 | 6c (RK) |
| 4-03 | 6d (RK) |
| 4-04 | BCL-4-148 |
| 4-05 | BCL-7-77 |
| 4-06 | BCL-7-45 |
| 4-07 | BCL-5-54 |
| 4-08 | BCL-4-119 |
| 4-09 | BCL-4-52 |
| 4-10 | BCL-7-76 |
| 4-11 | SCC-3-64 |
| 4-12 | ABA-1-19 |
| 4-13 | ABA-1-26 |
| 5-01 | BCL-4-83 |
| 5-02 | BCL-7-58 |
| 5-03 | BCL-1-144 |
| 5-04 | BCL-4-11 |
| 5-05 | BCL-5-29 |
| 5-06 | BCL-4-66 |
| 5-07 | BCL-4-138 |
| 6-01 | SCC-2-99 |
| 6-02 | BCL-2-56 |
| 6-03 | 6l (RK) |
| 6-04 | 10 (RK) |
| 6-05 | 6e (RK) |
| 6-06 | DD-07-134 |
| 6-07 | DD-07-148 |
| 6-08 | 7n (RK) |
| 6-09 | 6f (RK) |
| 6-10 | 6g (RK) |
| 6-11 | DD-07-115 |
| 6-12 | DD-06-126 |
| 6-13 | ABA-1-34 |
| 6-14 | BCL-7-49 |

#### General information

Reaction mixtures that require heating were heated in an oil bath with digital temperature control. Room temperature in this study is between 22 and 23°C. Analytical thin-layer chromatography (TLC) was carried out using 200 μm aluminum-backed silica gel plates with the F-254 indicator, purchased from Silicycle (TLA-R10011B-323). Visualization of the compounds on TLCs was accomplished under UV light (254 nm) and/or with a potassium permanganate stain. Product purification was achieved by flash column chromatography on a Biotage Isolera One using 40-63 μm silica gel purchased from Silicycle (R12030B) or RediSep Rf Gold Silica flash chromatography columns from Teledyne ISCO. All flash column chromatography purifications were performed using an isocratic or step solvent gradient. Triethylamine and diisopropylethylamine were distilled from KOH and stored in the presence of KOH pellets. Unless otherwise noted, reagents were purchased from commercial sources (Sigma-Aldrich, Thermo Scientific, Oakwood, Combi-Blocks, Ambeed), and used as received. Dry solvents were obtained from an Innovative Tech Pure Solve MD-7 Solvent purification system. MilliQ-quality water (MQ) was used whenever water is mentioned. Proton nuclear magnetic resonance ($^{1}$H-NMR) and proton-decoupled carbon nuclear magnetic resonance ($^{13}$C-NMR) were recorded on a Bruker AVIIIHD 500 MHz NMR spectrometer at 298 K. Chemical shifts ($\delta$) for $^{1}$H-NMR are reported in parts per million (ppm) and referenced to residual protons in the solvent; $CDCl_3$ ($\delta$ = 7.26 ppm), $CD_3OD$ ($\delta$ = 3.31 ppm), DMSO-$d_6$ ($\delta$ = 2.50 ppm), $D_2O$ ($\delta$ = 4.79 ppm). Chemical shifts ($\delta$) for $^{13}$C-NMR are reported in ppm and referenced to the carbon signal of the solvent; $CDCl_3$ ($\delta$ = 77.16 ppm), $CD_3OD$ ($\delta$ = 49.00 ppm), DMSO-$d_6$ ($\delta$ = 39.52 ppm). NMR splitting patterns are reported as singlet (s), broad singlet (bs), doublet (d), triplet (t), quartet (q), quintet (p), or sextet (h). High-resolution mass spectra (HR-MS) were obtained by electrospray ionization (ESI) or atmospheric pressure chemical ionization (APCI) on a Bruker Maxis atmospheric pressure ionization (API) Quadruple Time-of-Flight (QTOF) mass spectrometer.

#### General Protocols

**General procedure A:** synthesis of 19a-c

Compounds **19a-c** were synthesized in two steps from **17a-c** following a modified literature protocol.[1] One of **17a-c** and diethyl amine were dissolved in MeOH. The mixture was warmed to 65°C and stirred for 5 min. D-Pantolactone was added to the reaction mixture before stirring at 65°C for 48-72 h. The solvent was evaporated *in vacuo*. The residue was redissolved in water (10 mL) and loaded on an Amberlite IR-120 ($H^+$) ion exchange column (2 × 5 cm). The product was eluted with water until neutrality was obtained. The excess water was evaporated *in vacuo* to obtain crude intermediates **18a-c**, which were directly used in the next step without any purification.

One of **18a-c**, was next dissolved in dry DMF, and CSA was added, followed by *p*-methoxybenzaldehyde dimethyl acetal. The mixture was stirred at rt overnight. The next day, the reaction mixture was diluted in EA and washed with $H_2O$. The organic layer was then dried over $Na_2SO_4$, filtered and concentrated *in vacuo*. The solid obtained was washed with chilled $Et_2O$ to afford the desired compounds **19a**, **19b** or **19c**.

**General procedure B**: peptide coupling to synthesize **5a-n** and **21a-c**

One of the acid intermediates, **4a-n**, or **4q**, was dissolved in dry THF. NHS was added, followed by DCC. The reaction mixture was stirred at rt for 4 h, at which point one of the amine intermediates **3** or **20a-c** pre-dissolved in dry THF was added, followed by the addition of $Et_3N$. The mixture was stirred at rt overnight. The dicyclohexyl urea precipitate was removed by filtration and washed with EA. The organic layers were combined, and the solvents were removed *in vacuo*. The compound was purified by silica gel column chromatography.

**General procedure C**: PMP deprotection to generate **6a-n, 9k, 12, 15, 16**, and **22a-c**

One of the compounds **5a-n**, **8k**, **11**, **13**, **14**, or **21a-c** was dissolved in acetic acid and water (9:1 v/v, 15 mg/mL) and the solution was stirred at rt. After completion of the reaction as monitored by TLC (ca. 4 h), the solvents were removed *in vacuo*. The product was purified by silica gel column chromatography.

**General procedure D**: benzyl deprotection to generate **7m** and **7n**

One of the compounds **6m** or **6n** was stirred with Pd/C (10% w/w) in MeOH, and $H_2$ gas was purged into the reaction mixture at rt for 2 h. The reaction mixture was filtered through a celite pad, the solvent was evaporated *in vacuo*, and the residue was purified by silica gel column chromatography if mentioned.

**General procedure E**: EDCI/DMAP amide coupling between target acid and Boc-ethylenediamine

To a 25 mL round bottom flask were added the target acid (1.1 equiv.), Boc-ethylenediamine (1.0 equiv.), DMAP (0.1 equiv.), and DCM (5 mL/1 mmol amine). The mixture was cooled to 0°C and EDCI (1.1 equiv.) was added in small portions. The reaction mixture was stirred at rt overnight. The mixture was diluted in DCM (20 mL), then washed with sat. $NaHCO_3$ (2 × 20 mL), 1 M $KH_2PO_4$ (2 × 20 mL), and brine (1 × 20 mL). The organic layer was dried over anhydrous $Na_2SO_4$, filtered, and the solvent was evaporated under vacuum. Purification by flash chromatography afforded the product.

**General procedure F**: Boc deprotection

To a 25 mL round bottom flask were added the Boc-protected compound and DCM (1.5 mL/1 mmol compound). TFA (1.5 mL/1 mmol compound) was added and the mixture was stirred at rt for 2 h, after which the solvent was evaporated under vacuum. The residue was redissolved in DCM and the solvent was evaporated under vacuum. The residue was neutralized with excess amount of DIPEA, and used without further purification.

**General procedure G**: EDCI/DMAP amide coupling between PMP-pantothenate and target amine

To a 25 mL round bottom flask were added PMP-pantothenate (1.0 equiv.), target amine from Boc deprotection (1.0 equiv.), DMAP (0.1 equiv.), and DCM (5 mL/1 mmol amine). The mixture was cooled to 0°C and EDCI (1.1 equiv.) was added in small portions. The reaction mixture was stirred at rt overnight. The mixture was diluted in DCM (20 mL), then washed with sat. $NaHCO_3$ (2 × 20 mL), 1 M $KH_2PO_4$ (2 × 20 mL), and brine (1 × 20 mL). The organic layer was dried over anhydrous

$Na_2SO_4$, filtered, and the solvent was evaporated under vacuum. Purification by flash chromatography afforded the product.

**General procedure H**: acidic PMP deprotection

To a 25 mL round bottom flask were added the PMP-protected compound (1.0 equiv.) and 80% HOAc (5 mL/1 mmol compound). The reaction mixture was stirred at rt for 5 h, after which the solvent was evaporated under vacuum. The residue was dissolved in water (10 mL), and the aqueous layer was washed with DCM (2 × 10 mL) and EA (1 × 10 mL). The water of the aqueous layer was then evaporated under vacuum. The residue was dissolved in methanol (15 mL), and anhydrous $Na_2SO_4$ as well as solid $NaHCO_3$ were added. The mixture was allowed to stand for 5 min and then filtered. The solvent was evaporated under vacuum. Purification by flash chromatography, followed by lyophilization, afforded the product.

**General procedure I**: Reductive hydrogenation

To a 25 mL round bottom flask were added the starting material (1.0 equiv.), 10% Pd/C (100 mg/1 mmol compound), and MeOH (5 mL/1 mmol compound). $H_2$ was bubbled into the solution for 5 min, then a $H_2$ balloon was attached to maintain the $H_2$ atmosphere. The reaction mixture was stirred at rt overnight. Next, the mixture was filtered through celite under vacuum. The solvent was evaporated under vacuum to afford the product.

**General procedure J**: EDCI/HOBt amide coupling between target acid and amine (Boc-ethylenediamine or PMP-$NH_2$)

To a 25 mL round bottom flask were added the target acid (1.1 equiv.), HOBt hydrate (80%, 1.1 equiv.), DIPEA (2.5 equiv.), and DCM (5 mL/1 mmol amine). The mixture was cooled to 0°C and EDCI (1.1 equiv.) was added in small portions. The reaction mixture was stirred at 0°C for 30 min, then amine (1.0 equiv.) was added. The mixture was stirred at rt overnight. Next, it was diluted in DCM (20 mL), then washed with sat. $NaHCO_3$ (2 × 20 mL), 1 M $KH_2PO_4$ (2 × 20 mL), and brine (1 × 20 mL). The organic layer was dried over anhydrous $Na_2SO_4$, filtered, and the solvent was evaporated under vacuum. Purification by flash chromatography afforded the product.

**General procedure K**: EDCI/HOBt amide coupling between PMP-pantothenate and target amine

To a 25 mL round bottom flask were added the PMP-pantothenate (1.1 equiv.), HOBt hydrate (80%, 1.1 equiv.), DIPEA (2.5 equiv.), and DCM (5 mL/1 mmol amine). EDCI (1.1 equiv.) was added in small portions and the mixture was stirred at 0ºC for 30 min. The amine from Boc deprotection (1.0 equiv.) was then added. The reaction mixture was stirred at rt overnight. Next, it was diluted in DCM (20 mL), then washed with sat. $NaHCO_3$ (2 × 20 mL), 1 M $KH_2PO_4$ (2 × 20 mL), and brine (1 × 20 mL). The organic layer was dried over anhydrous $Na_2SO_4$, filtered, and the solvent was evaporated under vacuum. Purification by flash chromatography afforded the product.

**Procedure L**: Amide coupling between carboxylic acid and amine

In a 25 mL round-bottom flask, the carboxylic acid (1.1 equiv.) was added, followed by HOBt hydrate (80 % w/w, 1.1 equiv.), anhydrous DCM (5 mL/mmol amine) while stirring at 0°C. This mixture was stirred for another 5 min. Afterwards, DIPEA (2.5 equiv.) and EDCI (1.1 equiv.) were added, and the reaction mixture was stirred at 0°C for 1 h. Finally, the amine (1.0 equiv.) was added before stirring at rt overnight. The mixture was then diluted in DCM (20 mL), washed with sat. aq. $NaHCO_3$ (3 × 20 mL) and 1 M aq. $KH_2PO_4$ (2 × 20 mL), and dried over anhydrous $Na_2SO_4$. The solvent was evaporated under vacuum and purification by flash chromatography afforded the final product.

**Procedure M**: Amide coupling between acid chloride and amine

To a 25 mL round-bottom flask were added the amine (1.0 equiv.) and anhydrous DCM (4 mL/mmol amine). The mixture was stirred at 0°C for 5 min. Afterwards, TEA (1.5 equiv.) was added, followed by the acid chloride (1.2 equiv.) dropwise, at 0°C. The reaction mixture was left to stir overnight at rt. The mixture was then diluted in DCM 20 mL), washed with sat. aq. $NaHCO_3$ (2 × 20 mL), and dried over anhydrous $Na_2SO_4$. The solvent was evaporated under vacuum and purification by flash chromatography yielded the final product.

**Procedure N**: Boc deprotection with AcCl and MeOH

In a 25 mL round-bottom flask, the Boc-protected compound (1.0 equiv.) was added, followed by EA (2 mL/mmol compound) and MeOH (4 M). The flask was then cooled in an ice bath, and AcCl (4 M) was added dropwise to the reaction mixture, before stirring for 2-24 h at rt. The resulting white precipitate was then collected by filtration under vacuum and washed with EA.

The filter cake was next added to sat. aq. $Na_2CO_3$ (30 mL), and the product was extracted in DCM (2 × 30 mL). Following this, the pH of the aqueous layer was adjusted to 14 using NaOH pellets, and the product was extracted once more in DCM (1 × 30 mL). The organic layers were combined and dried over anhydrous $Na_2SO_4$, and the solvent was evaporated under vacuum to yield the product, which was used in the next step without further purification.

**Procedure O**: Lactone aminolysis using TBD

To a 50 mL round-bottom flask were added the amine (1.0 equiv.), D-pantolactone (2.0 equiv.), TBD (0.1 equiv.), and toluene (1 mL/mmol amine). The reaction mixture was stirred at rt for 24 h. The solvent was removed under vacuum, and purification by flash chromatography yielded the desired product.

**Procedure P**: Copper-catalyzed azide-alkyne cycloaddition (CuAAC)

To a 25 mL round-bottom flask were added the alkyne (1.0 equiv.), azide (1.0 equiv.), sodium ascorbate (1.0 equiv.), $CuSO_4$ (0.1 equiv.), and a 1:1 mixture of EtOH/$H_2O$ (10 mL/mmol alkyne). The reaction mixture was allowed to stir at rt for 1 h. Following this, the reaction was quenched with aq. EDTA (0.1 equiv. dissolved in sat. $Na_2CO_3$ solution to a 30 mM concentration). The solvent was evaporated under vacuum and the remaining residue was dissolved in MeOH (15 mL). The resulting precipitate was removed by gravity filtration and the solvent of the filtrate was removed under vacuum. Purification by flash chromatography yielded the desired product.

**Procedure Q**: Amide coupling between *N*-Boc-ethylenediamine and carboxylic acid

To a round-bottom flask were added *N*-Boc-ethylenediamine (1.0 equiv.), acid (1.1 equiv.), DMAP (11 mol%), and anhydrous DCM. The mixture was cooled to 0°C, and EDCI (1.1 equiv.) was added in portions. The reaction mixture was stirred at rt overnight. The mixture was then diluted in DCM (20 mL), and washed with sat. aq. $NaHCO_3$ (2 × 25 mL), 1 M aq. $KH_2PO_4$ (2 × 25 mL), and brine (1 × 25 mL). The organic layer was collected, dried over anhydrous $Na_2SO_4$, and concentrated under reduced pressure. Purification by flash chromatography afforded the desired product.

**<u>Procedure R</u>**: <u>Amide coupling between Boc-protected amine and **PMP-pantothenate**</u>

Before amide coupling, the Boc-protected amine was first deprotected using TFA with the following protocol: To a round-bottom flask, the Boc-protected amine (1.0 equiv.) was added, followed by anhydrous DCM and TFA (amount varies with each substrate). The reaction was stirred at rt for 2 h. Afterwards, DIPEA was added dropwise to adjust the pH to 7. The mixture was then concentrated under reduced pressure, and the remaining product was used without further purification.

For the amide coupling step: To a round-bottom flask containing the crude deprotected amine were added **PMP-pantothenate** (1.0 equiv.), DMAP (0.1 equiv.), and anhydrous DCM (30 mL). The reaction mixture was cooled to 0°C, and EDCI (1.0 equiv.) was added in portions. The reaction was then stirred at rt overnight. Afterwards, the reaction was diluted with DCM (20 mL), and the mixture was washed with $NaHCO_3$ (2 × 25 mL), 1 M $KH_2PO_4$ (2 × 25 mL), and brine (1 × 25 mL). The organic layer was collected, dried over anhydrous $Na_2SO_4$, and concentrated under reduced pressure. Purification by flash chromatography afforded the desired product.

**<u>Procedure S</u>**: <u>PMP deprotection</u>

The PMP-protected compound (1.0 equiv.) was dissolved in a 9:1 mixture of AcOH/$H_2O$. The reaction mixture was stirred at rt for 3 h. Afterwards, the mixture was diluted in $H_2O$ (10 mL). The aqueous layer was washed with DCM (3 × 10 mL) and EA (2 × 10 mL). The aqueous layer was then evaporated under reduced pressure, affording the final product.

#### Purity trace elution methods:

<u>In total there are 17 methods included:</u>

**<u>Elution methods 1-2</u>** use a SYNERGI 4 μm Hydro-RP 80 Å Phenomenex, 4.60 × 260 mm column at a flow rate of 0.5 mL/min, with detection at 214 nm and water as mobile phase A and acetonitrile as phase B.
**Elution Method 1**: 0-3 min: 1% B in A, 3-5 min: 1% to 10% B, 5-10 min: 10% to 15% B, 10-13 min: 15% to 30% B, 13-26 min: 30% B, 26-30 min: 30% to 1% B.
**Elution Method 2**: 0-8 min: 1% B in A, 8-15 min: 1% to 50% B, 15-25 min: 50% B, 25-30 min: 50% B to 100% B, 30-38 min: 100% B, 38-40 min: 100% B to 1% B.

**<u>Elution methods 3-4</u>** use a Zorbax NH2 5 μm, 4.60 × 150 mm column at a flow rate of 0.5 mL/min, with detection at 256 nm and water as mobile phase A and acetonitrile as phase B.
**Elution Method 3**: 0-3 min: 99% B in A, 3-5 min: 99% to 90% B, 5-10 min: 90% to 85% B, 10-13 min: 85% to 70% B, 13-26 min: 70% B, 26-30 min: 70% to 99% B.
**Elution Method 4**: 0-8 min: 99% B in A, 8-15 min: 99% to 60% B, 15-25 min: 60% B, 25-30 min: 60% B to 0% B, 30-38 min: 0% B, 38-40 min: 0% B to 99% B.

**<u>Elution methods 5-6</u>** use a Luna 5 μm C18(2) 100Å Phenomenex, 4.60 × 260 mm column at a flow rate of 0.5 mL/min, with detection at 214 nm and water as mobile phase A and acetonitrile as phase B.
**Method 5**: 0-8 min: 1% B in A, 8-15 min: 1% to 50% B, 15-25 min: 50% B, 25-30 min: 50% B to 100% B, 30-38 min: 100% B, 38-40 min: 100% B to 1% B.
**Method 6**: 0-2 min: 1% B in A, 2-10 min: 1% to 100% B, 10-20 min: 100% B, 20-24 min: 100% to 1% B, 24-25 min: 1% B.

**<u>Elution methods 7-9</u>** use a Luna 5 μm C18(2) 100 Å, LC Column 250 x 4.6 mm at a flow rate of 0.5 mL/min with detection at 214 nm.
**Method 7**: 0-2 min: 1% B in A, 2-10 min: 1% to 100% B, 10-20 min: 100% B, 20-24 min: 100% to 1% B, 25-26 min: 1% B.

**Method 8**: 0-2 min: 1% B in A, 2-10 min: 1% to 50% B, 10-20 min: 50% B, 20-24 min: 50% to 100% B, 25-35 min: 100% B, 35-40 min: 100% to 1% B, 40-42 min: 1% B.

**Method 9**: 0-2 min: 1% B in A, 2-5 min: 1% to 15% B, 5-10 min: 15% to 30% B, 10-13 min: 30% to 50% B, 13-18 min: 50% to 80% B, 18-24 min: 80% to 100% B, 24-28 min: 100% B, 28-30 min: 100% to 1% B.

**Elution methods 10-11** use a Zorbax NH2 5 um, 4.60 × 150 mm column at a flow rate of 0.5 mL/min, with detection at 256 nm.

**Method 10**: 0-3 min: 99% B in A, 3-5 min: 99% to 90% B, 5-10 min: 90% to 85% B, 10-13 min: 85% to 70% B, 13-26 min: 70% B, 26-30 min: 70% to 99% B.

**Method 11**: 0-1 min: 99% B in A, 1-8 min: 99% to 60% B, 8-10 min: 60% to 0% B, 10-15 min: 0% B, 15-25 min: 0% to 99% B, 25-30 min: 99% B.

**Elution methods 12-15** use a Phenomenex Luna® 5 µm, 200 × 4.6 mm, 100 Å, C18 column at a flow rate of 1 mL/min, with detection at 214 nm and water as mobile phase A and acetonitrile as phase B. All changes in solvent ratio during elution were linear.

**Elution method 12**: 0-3 min: 1% B in A, 3-5 min: 1% to 10% B, 5-10 min: 10% to 15% B, 10-13 min: 15% to 30% B, 13-26 min: 30% B, 26-30 min: 30% to 99% B, 30-32 min: 99% B, 32-35 min: 99% B to 1% B, 35-38 min: 1% B.

**Elution method 13**: 0-8 min: 1% B in A, 8-15 min: 1% to 50% B, 15-25 min: 50% B, 25-30 min: 50% to 99% B, 30-38 min: 99% B, 38-40 min: 99% to 1% B, 40-43 min: 1% B.

**Elution method 14**: 0-5 min: 1% B in A, 5-15 min: 1% B in A to 50% B in A, 15-20 min: 50% B in A, 20-25 min: 50% B in A to 99% B in A, 25-30 min: 99% B in A, 30-32 min: 99% B in A to 1% B in A, 32-35 min: 1% B in A.

**Elution method 15**: 0-10 min: 1% B in A to 99% B in A, 10-20 min: 99% B in A, 20-22 min: 99% B in A to 1% B in A, 22-25 min: 1% B in A.

#### Synthesis of intermediates and final compounds

*Synthesis Route for Compound 1-02*

##### Synthesis and characterization of DD-05-140

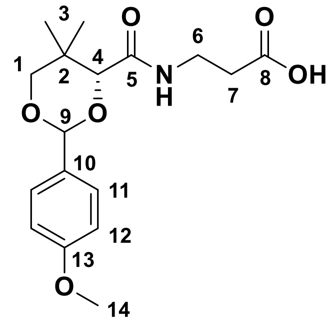

Synthesis of the known compound **DD-05-140** followed previously reported literature.[2] Yield: 2.66 g (47%). $^1$H NMR (300 MHz, $CDCl_3$): δ 7.41 (m, 2H, H-11), 7.03 (t, *J* = 6.4 Hz, 1H, NH), 6.91 (m, 2H, H-12), 5.46 (s, 1H, H-9), 4.09 (s, 1H, H-4), 3.81 (s, 3H, H-14), 3.70 (m, 2H, H-1), 3.54 (m, 2H, H-6), 2.61 (t, *J* = 6.2 Hz, 2H, H-7), 1.10 (s, 3H, H), 1.10 (s, 3H, H).

##### Synthesis and characterization of DD-05-006

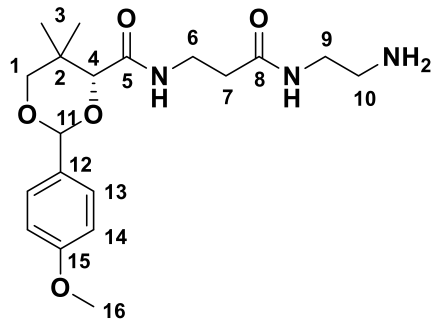

Synthesis of the known compound **DD-05-006** followed previously reported literature.[2] Yield: 1.18 g (53%). $^1$H NMR (300 MHz, $CDCl_3$): δ 7.51 (t, *J* = 5.6 Hz, 1H, NH), 7.43 (m, 2H, H-13), 7.06 (t, *J* = 6.3 Hz , 1H, NH-1 or NH-2), 6.91 (m, 2H, H-14), 5.45 (s, 1H, H-11), 4.06 (s, 1H, H-4), 3.81 (s, 3H, H-16), 3.68 (m, 2H, H-1), 3.52 (m, 2H, H-6 or H-9), 3.47 (m, 2H, H-6 or H9), 2.86 (m, 2H, H-10), 2.40 (t, *J* = 6.4 Hz, 2H, H-7), 1.09 (s, 3H, H-3), 1.06 (s, 3H, H-3).

##### Synthesis and characterization of DD-06-123

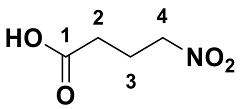

Synthesis of the known compound **DD-06-123** followed the previously reported procedure with modifications.[3] The reactants methyl 4-nitrobutanoate (0.165 g, 1.21 mmol) and lithium hydroxide monohydrate (0.099 g, 2.36 mmol) were dissolved in 1:1 THF:water (10 mL). This reaction mixture was stirred at room temperature for 16 h. The reaction mixture was washed dichloromethane (2 × 5 mL). The pH was reduced to less than 2 using 2 N HCl. The product was extracted in ethyl acetate (3 × 5 mL) and the combined organic layers were dried over $Na_2SO_4$, filtered, and concentrated *in vacuo* to afford the product as a yellow oil. Yield: 0.101 g (68%). $R_f$ = 0.40 (MeOH/DCM = 1:9). $^1$H NMR (500 MHz, $CDCl_3$): 4.48 (t, *J* = 6.8 Hz, 2H, H-4), 2.54 (t, *J* = 7.0 Hz, 2H, H-2), 2.31 (p, *J* = 6.9 Hz, 2H, H-3). $^{13}$C NMR ($CDCl_3$, 126 MHz): 178.0, 74.2, 30.3, 22.1.

**Synthesis and characterization of DD-06-124**

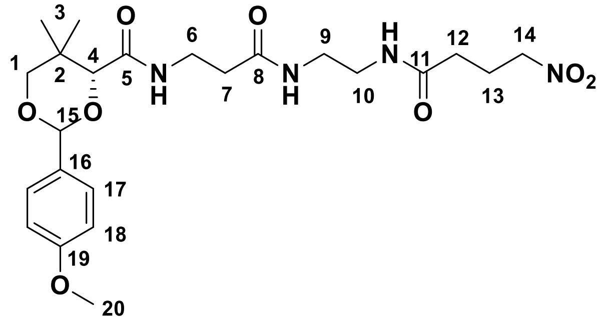

Synthesis of the known compound **DD-06-124** followed previously reported literature.[2] Yield: 0.044 g (46%). $R_f$: 0.30 (MeOH/DCM = 1:9). $^1$H NMR (500 MHz, $CDCl_3$): δ 7.42 (m, 2H, H-17), 7.05 (t, *J* = 6.3 Hz, 1H, NH), 6.91 (m, 2H, H-18), 6.83 (t, *J* = 5.2 Hz, 1H, NH), 6.76 (m, 1H, NH), 5.46 (s, 1H, H-15), 4.42 (m, 2H, H-14), 4.07 (s, 1H, H-4), 3.81 (s, 3H, H-20), 3.69 (m, 2H, H-1), 3.52 (q, *J* = 6.4 Hz, 2H, H-6), 3.40 – 3.24 (m, 4H, H-9 and H-10), 2.41 (t, *J* = 6.3 Hz, 2H, H-7), 2.27 – 2.23 (m, 4H, H-12 and H-13), 1.19 (s, 3H, H-3), 1.06 (s, 3H, H-3). $^{13}$C NMR (126 MHz, $CDCl_3$): δ 172.2 (C-8), 171.8 (C-11), 169.9 (C-5), 160.4 (C-19), 130.2 (C-16), 127.7 (C-17), 113.9 (C-18), 101.5 (C-15), 83.9 (C-4), 78.5 (C-1), 74.9 (C-14), 55.5 (C-20), 40.1 (C-10), 39.6 (C-9), 36.4 (C-7), 35.2 (C-6), 33.2 (C-2),

32.2 (C-12), 23.0 (C-13), 22.0 (C-3), 19.2 (C-3). HRMS (ESI$^+$) *m/z* $[C_{23}H_{34}N_4NaO_8]^+$ calcd. 517.2269, found 517.2254.

**Synthesis and characterization of DD-06-143 (1-02)**

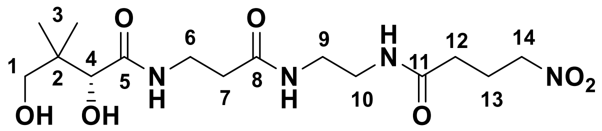

Synthesis of the known compound **DD-06-143** followed previously reported literature.[2] Yield: 0.0194 g (74%). $R_f$ = 0.10 (10% MeOH/DCM). HPLC purity 96%, RT = 12.35 min with method 6. $^1$H NMR (500 MHz, MeOD): δ 4.51 (t, *J* = 6.6 Hz, 2H, H-14), 3.89 (s, 1H, H-4), 3.53 – 3.37 (m, 4H, H-1, H-6), 3.28 – 3.27 (m, 4H, H-9, H-10), 2.42 (t, *J* = 6.6, 2H, H-7), 2.32 (m, 2H, H-12), 2.26 (m, 2H, H-13), 0.92 (s, 6H, H-3). $^{13}$C NMR (126 MHz, MeOD): δ 176.1 (C-5), 174.4 (C-8), 174.2 (C-11), 77.4 (C-4), 75.6 (C-14), 70.3 (C-1), 40.4 (C-2), 40.0 (C-9 or C-10), 40.0 (C-9 or C-10), 36.6 (C-7), 36.4 (C-6), 33.2 (C-12), 24.3 (C-13), 21.3 (C-3), 20.9 (C-3). HRMS (ESI$^+$) *m/z* $[C_{15}H_{28}N_4NaO_7]^+$ calcd. 399.1850, found 399.1838.

*Synthesis Route for Compound 1-03*

**Synthesis and characterization of DD-05-131**

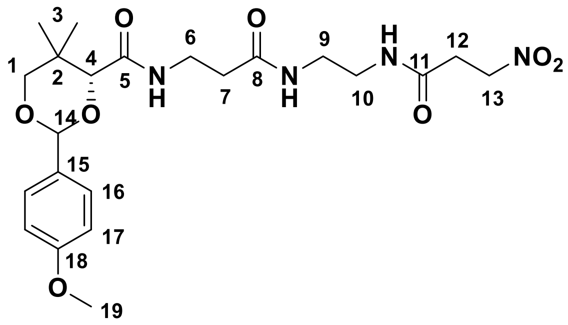

Synthesis of the known compound **DD-05-131** followed previously reported literature.[2] Yield: 0.154 g (42%). $^1$H NMR (300 MHz, $CDCl_3$): δ 7.43 (m, 2H, H-16), 7.13 – 7.03 (m, 2H, NH), 6.96 – 6.86 (m, 3H, H-17 and NH), 5.45 (s, 1H, H-14), 4.63 (m, 2H, H-13), 4.07 (s, 1H, H-4), 3.81 (s, 3H, H-19), 3.69 (m, 2H, H-1), 3.51 (m, H-6), 3.42 – 3.25 (m, 4H, H-

9 and H-10), 2.74 (m, 2H, H-12), 2.40 (t, $J$ = 6.4 Hz, 2H, H-7), 1.09 (s, 3H, H-3), 1.06 (s, 6H, H-3).

**Synthesis and characterization of compound DD-06-111**

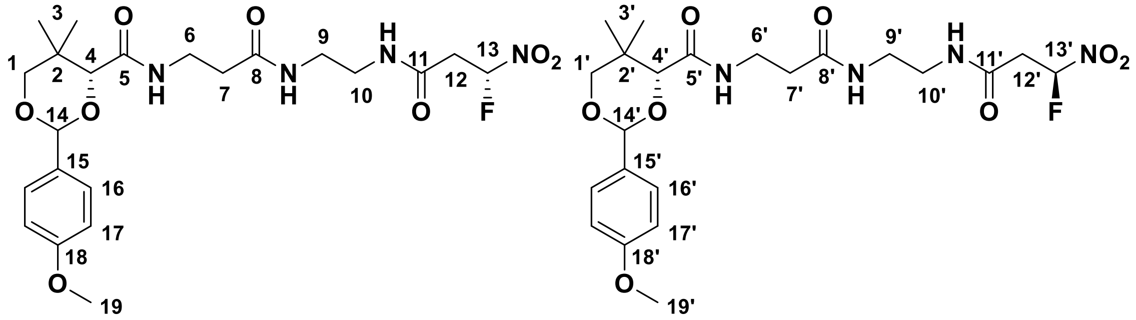

The reactant compound **DD-05-131** (0.101 g, 0.210 mmol) was dissolved into aqueous acetonitrile (1:1 MeCN:$H_2O$) and to this solution was added lithium hydroxide monohydrate (0.010 g, 0.24). The reaction mixture was stirred for 5 min at room temperature. To this solution was added Selectfluor® (0.082 g, 0.23 mol) and the reaction was stirred at room temperature for 5 min. The reaction was quenched by adding to a separatory funnel containing DCM, brine, and distilled water (10 mL, 6 mL, and 2 mL, respectively). The organic layer was collected. The product was further extracted from the aqueous layer into DCM (2 × 10 mL), and the combined organic layers were dried over $Na_2SO_4$, then filtered and concentrated *in vacuo*. Purification was achieved with a gradient of 3 – 7% MeOH/DCM to give the product as a white powder. Yield: 0.097 g (92%). $R_f$: 0.35 (MeOH/DCM = 1:9). $^1$H NMR (500 MHz, $CDCl_3$): δ 7.42 (m, 4H, H-16, H-16'), 7.20 (t, $J$ = 4.9 Hz, 2H, NH and NH'), 7.02 (t, $J$ = 6.5 Hz, 2H, NH and NH'), 6.92 (m, 4H, H-17, H-17'), 6.69 (m, 2H, NH and NH'), 6.17 (ddd, $J$ = 49.2, 6.6, 4.4 Hz; 2H, H-13 and H-13'), 5.46 (s, 2H, H-14 and H-14'), 4.07 (s, 2H, H-4 and H-4'), 3.82 (s, 6H, H-19 and H-19'), 3.69 (m, 4H, H-1 and H-1'), 3.53 (m, 4H, H-6), 3.44 – 3.27 (m, 8H, H-9 and H-10), 3.08 (ddd, $J$ = 24.4, 15.6, 4.4 Hz, 1H, H-12 or H-12'), 2.92 (ddd, $J$ = 16.9, 15.6, 6.7 Hz, 1H, H-12 or H-12'), 2.41 (m, 4H, H-7 and H-7'), 1.10 (s, 3H, H-3, H-3a'), 1.10 (s, 3H, H-3, H-3'), 1.07 (s, 6H, H-3, H-3'). $^{13}$C NMR (126 MHz, $CDCl_3$): δ 172.5 (C-8 and C-8'), 170.1 (C-5 and C-5'), 165.6 (d, $J$ = 4.1 Hz, C-11 and C-11'), 160.5 (C-18 and C-18'), 130.2 (C-15 and C-15'), 127.7 (C-16 and C-16'), 113.9 (C-17 and C-17'), 108.4 (d, $J$ = 239.0 Hz, C-13 or C-13'), 108.4 (d, $J$ = 239.0 Hz, C-13 or C-13'), 101.5 (C-14 and C-14'), 83.9

(C-4 or C-4'), 83.9 (C-4 or C-4'), 78.5 (C-1 and C-1'), 55.5 (C-19 and C-19'), 40.5 (C-9, C-9', C-10, or C-10'), 40.4 (C-9, C-9', C-10, or C-10'), 39.8 (d, $J$ = 21.5 Hz, C-12 and C-12'), 39.3 (C-9, C-9', C-10, or C-10'), 39.3 (C-9, C-9', C-10, or C-10'), 36.7 (C-7 and C-7'), 35.2 (C-6 or C-6'), 35.2 (C-6 or C-6'), 33.2 (C-2 and C-2'), 22.0 (C-3 and C-3'), 19.2 (C-3 and C-3'). $^{19}$F NMR (471 MHz, $CDCl_3$): δ -146.7 (ddd, 49.3, 24.3, 17.3 Hz; 1F, F-13 and F-13'). HRMS (ESI$^+$) *m/z* $[C_{22}H_{31}FN_4NaO_8]^+$ calcd. 521.2018, found 521.2019.

**Synthesis and characterization of compound DD-07-066 (1-03)**

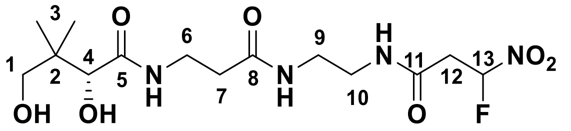

The reactant compound **DD-06-111** (0.0604 g, 0.121 mmol) was dissolved into a solution of 9:1 AcOH:$H_2O$ (6 mL) and the mixture was stirred at room temperature for 3 h. The reaction mixture was concentrated *in vacuo*. The residue was redissolved into distilled water (10 mL) and washed with DCM (5 × 10 mL). The aqueous layer was collected and concentrated *in vacuo* to afford the product as a white solid. Yield: 0.0433 g (94%). $R_f$: 0.13 (MeOH/DCM = 1:9). HPLC purity 94%, RT = 14.37 min with method 6. $^1$H NMR (500 MHz, MeOD): δ 6.35 (ddd, $J$ = 48.6, 6.5, 4.1 Hz; 1H, H-13), 3.89 (s, 1H, H-4), 3.54 – 3.38 (m, 4H, H-1 and H-6), 3.33 – 3.27 (m, 4H, H-9 and H-10), 3.22 (ddd, 23.6, 15.9, 4.2 Hz; 1H, H-12a), 3.11 (ddd, $J$ = 19.9, 15.8, 6.5 Hz; 1H, H-12b), 2.42 (t, $J$ = 6.6, 2H, H-7), 0.92 (s, 6H, H-3). $^{13}$C NMR (126 MHz, MeOD): δ 176.1 (C-5), 174.2 (C-8), 168.2 (d, $J$ = 3.8 Hz, C-11), 109.7 (d, $J$ = 236.7 Hz, C-13), 77.3 (C-4), 70.3 (C-1), 40.4 (C-9 or C-10), 40.2 (d, $J$ = 21.5 Hz, C-12), 40.1 (C-2), 39.9 (C-9 or C-10), 36.6 (C-7), 36.4 (C-6), 21.3 (C-3), 20.9 (C-3). $^{19}$F NMR (MeOD, 471 MHz): δ -148.9 (m, 1F, F-13). HRMS (ESI$^+$) *m/z* $[C_{14}H_{25}FN_4NaO_7]^+$ calcd. 383.1360, found 383.1360.

*Synthesis Route for Compound 1-05*

**Synthesis and characterization of compound DD-02-088**

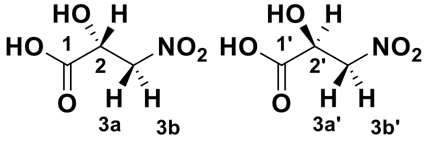

Synthesis of the known compound **DD-02-088** followed the previously reported procedure with modifications.[4,5] The reactants ethyl glyoxylate (in 50% toluene w/w, (5.1 g, 25 mmol) and lithium hydroxide monohydrate (1.08 g, 25.7 mmol) were dissolved in distilled water (100 mL) and stirred at room temperature for 5 min. The reactant nitromethane (1.4 mL, 1.6 g, 26 mmol) was added, then the mixture was stirred at room temperature for 16 h. The reaction mixture was washed with DCM (2 × 25 mL), and then acidified to pH 1 with 2 N HCl before extracting with the product in EtOAc (4 × 50 mL). The combined organic layers were dried over $Na_2SO_4$ and filtered. The filtrate was concentrated *in vacuo* to give the product as a red oil. Yield: 1.08 g (32%). $R_f$: 0.05 (MeOH/DCM = 1:9). $^1$H NMR (500 MHz, $d_6$-DMSO): δ 4.85 (dd, *J* = 13.4, 3.9 Hz, 2H, H-3a and H-3b'), 4.72 (dd, *J* = 13.4, 7.1 Hz, 2H, H-3b and H-3a'), 4.59 (dd, *J* = 7.1, 3.9 Hz, 2H, H-2 and H-2'). $^{13}$C NMR (126 MHz, $d_6$-DMSO): δ 172.0 (C-1 and C-1'), 78.3 (C-3 and C-3'), 67.5 (C-2 and C-2').

**Synthesis and characterization of compound DD-03-038**

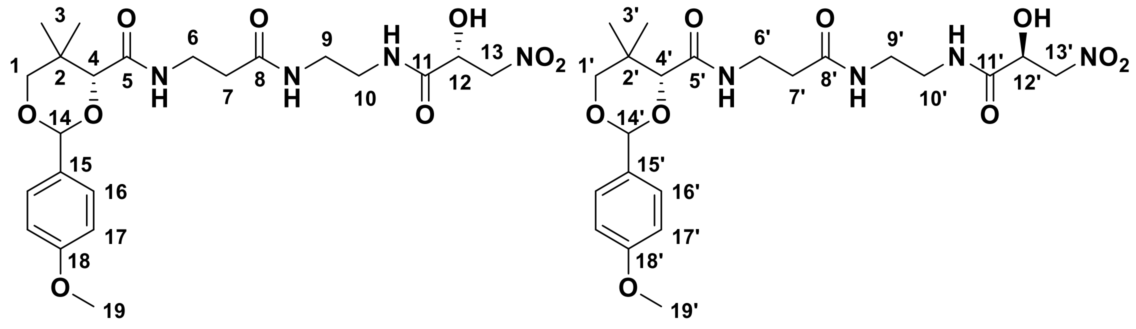

The reactants compound **DD-02-088** (0.073 g, 0.54 mmol), *N*-hydroxysuccinimide (0.066 g, 0.57 mmol), and *N,N'*-dicyclohexylcarbodiimide (0.123 g, 0.596 mmol) were dissolved in THF (5 mL) and the mixture was stirred at room temperature for 4.5 h. Compound **DD-05-006** (0.087 g, 0.23 mmol) dissolved in THF (5 mL) was added next, and the mixture was stirred at room temperature for 22.5 h. The reaction mixture was filtered and concentrated *in vacuo*. The residue was redissolved into DCM (5 mL) and the solution was washed with distilled water (5 mL) and brine (5 mL), before drying the organic layer over $Na_2SO_4$, filtering it, and concentrating it *in vacuo*. Purification was achieved with a gradient of 3 – 6% MeOH/DCM to give the product as a yellow solid. Yield: 0.045 g (37%). $R_f$: 0.57 (MeOH/DCM = 1:9). $^1$H NMR (500 MHz, $CDCl_3$): δ 7.63 (m, 2H, NH and NH'), 7.41 (m, 4H, H-16 and H-16'), 7.10 (m, 2H, NH and NH'), 7.01 (m, 2H, NH and NH'), 6.89

(m, 4H, H-17 and H-17'), 5.44 (s, 2H, H-14 and H-14'), 4.76 (m, 2H, H-12 and H-12'), 4.70 – 4.61 (m, 4H, H-13 and H-13'), 4.06 (s, 1H, H-4 or H-4'), 4.06 (s, 1H, H-4 or H-4'), 3.79 (s, 6H, H-19 and H-19'), 3.65 (m, 4H, H-1 and H-1'), 3.47 (m, 4H, H-6 and H-6'), 3.39 – 3.27 (m, 8H, H-9, H-9', H-10, and H-10'), 2.38 (t, $J$ = 6.6 Hz, 4H, H-7 and H-7'), 1.07 (s, 6H, H-3 and H-3'), 1.02 (s, 6H, H-3 and H-3'). $^{13}$C NMR (126 MHz, $CDCl_3$): δ 172.4 (C-8 or C-8'), 172.40 (C-8 or C-8'), 171.0 (C-11 or C-11'), 171.0 (C-11 or C-11'), 170.3 (C-5 or C-5'), 170.2 (C-5 or C-5'), 160.4 (C-18 and C-18'), 130.2 (C-15 and C-15'), 127.7 (C-16 or C-16'), 127.7 (C-16 or C-16'), 113.8 (C-17 and C-17'), 101.5 (C-14 or C-14'), 101.5 (C-14 or C-14'), 83.9 (C-4 and C-4'), 78.4 (C-1 and C-1'), 78.2 (C-13 and C-13'), 69.3 (C-12 or C-12'), 69.3 (C-12 or C-12'), 55.5 (C-19 and C-19'), 39.5 (C-9 and C-10, or C-9' and C-10'), 39.3 (C-9, C-9', C-10, or C-10'), 39.3 (C-9, C-9', C-10, or C-10'), 36.2 (C-7 and C-7'), 35.4 (C-6 or C-6'), 35.3 (C-6 or C-6'), 33.2 (C-2 and C-2'), 21.9 (C-3 and C-3'), 19.2 (C-3 and C-3'), 19.22 (C-3 and C-3'). HRMS (ESI$^+$) *m/z* $[C_{22}H_{32}N_4NaO_9]^+$ calcd. 519.2061, found 519.2062.

**Synthesis and characterization of compound DD-04-021 (1-05)**

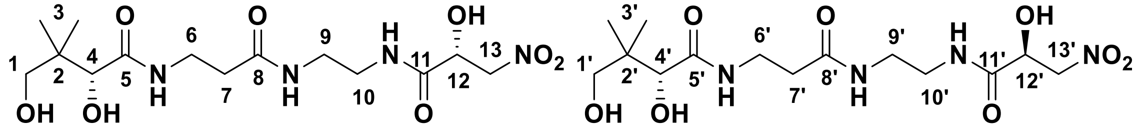

The reactant compound **DD-03-038** (0.107 g, 0.215 mmol) was dissolved into a solution of 9:1 AcOH:$H_2O$ (11 mL) and the mixture was stirred at room temperature for 3.5 h. The reaction mixture was next diluted into distilled water (10 mL) and washed with toluene (3 × 20 mL). The aqueous layer was collected and concentrated *in vacuo* to afford the product as a pale, red oil. Yield: 0.083 g (99%). $R_f$: 0.14 (MeOH/DCM = 1:9). HPLC purity 96%, RT = 12.23 min with method 6. $^1$H NMR (500 MHz, MeOD): δ 4.81 (m, 2H, H-13 and H-13'), 4.70 – 4.63 (m, 4H, H-12, H-12', H-13, and H-13'), 3.89 (s, 2H, H-4 and H-4'), 3.50 – 3.38 (m, 8H, H-1, H-1', H-6, and H-6'), 3.33 – 3.29 (m, 8H, H-9, H-9', H-10 and H-10'), 2.42 (t, $J$ = 6.7 Hz, 4H, H-7 and H-7'), 0.92 (s, 12H, H-3 and H-3'). $^{13}$C NMR (126 MHz, MeOD): δ 176.1 (C-5 and C-5'), 174.3 (C-8 and C-8'), 173.2 (C-11 and C-11'), 79.5 (C-13 and C-13'), 77.4 (C-4 and C-4'), 70.4 (C-12 and C-12'), 70.3 (C-1 and C-1'), 40.4 (C-2 and C-2'), 40.1 (C-9 and C-9' or C-10 and C-10'), 39.9 (C-9 and C'9' or C-10 and C-

10’), 36.6 (C-7 and C-7’), 36.4 (C-6 and C-6’), 21.3 (C-3 and C-3’), 20.9 (C-3 and C-3’). HRMS (ESI$^+$) *m/z* [$C_{14}H_{26}N_4NaO_8$]$^+$ calcd. 401.1643, found 401.1625.

*Synthesis Route for Compound 1-06*

**Synthesis and characterization of compound DD-02-134**

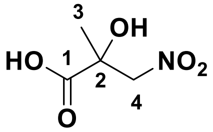

The reactants pyruvic acid (710 μL, 0.887 g, 10.1 mmol), nitromethane (550 μL, 0.620 g, 10.2 mmol), and lithium hydroxide monohydrate (0.857 g, 20.4 mmol) were dissolved in distilled water (30 mL) and the mixture was stirred at room temperature for 10.5 h. The reaction mixture was acidified to pH 1 and the product extracted in EtOAc (3 × 30 mL). The combined organic layers were dried over $Na_2SO_4$ and filtered. The filtrate was concentrated *in vacuo* to give the product as a red oil. Yield: 0.397 g (26%). $R_f$: 0.00 (MeOH/DCM = 1:9). $^1$H NMR (300 MHz, $CDCl_3$): δ 4.85 (d, $J$ = 13.7 Hz, 1H, H-4), 4.53, (d, $J$ = 13.7 Hz, 1H, H-4), 1.45 (s, 3H, H-3). $^{13}$C NMR (76 MHz, $CDCl_3$): δ 175.6 (C-1), 81.2 (C-4), 72.3 (C-2), 23.9 (C-3). HRMS (ESI$^+$) *m/z* [$C_4H_6N_1Na_2O_5$]$^+$ calcd. 194.0036, found 194.0031.

**Synthesis and characterization of compound DD-02-143**

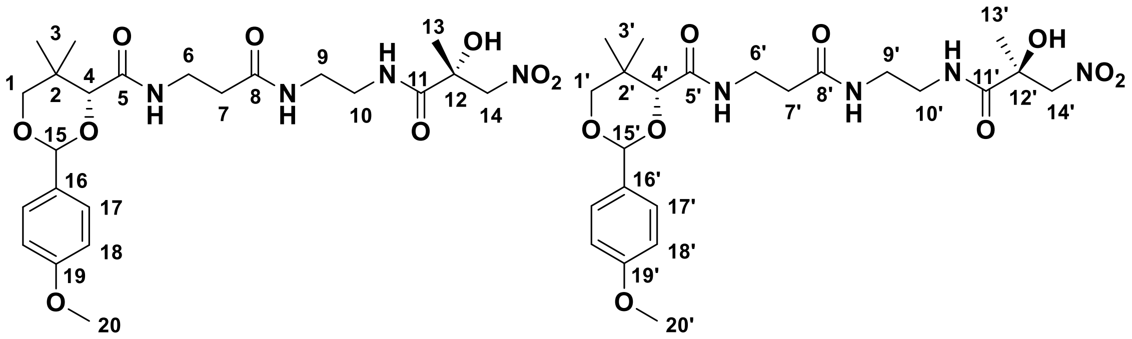

The reactants compound **DD-02-134** (0.106 g, 0.711 mmol), *N*-hydroxysuccinimide (0.085 g, 0.74 mmol), and *N,N’*-dicyclohexylcarbodiimide (0.144 g, 0.698 mmol) were dissolved in THF (10 mL) and the mixture was stirred at room temperature for 4.5 h. The reactant compound **DD-05-006** (0.101 g, 0.266 mmol) was dissolved in THF (5 mL) and added. The mixture was stirred at room temperature for 26 h. The reaction mixture was concentrated *in vacuo*, redissolved into DCM (5 mL) and the solution washed with distilled water (5 mL) and brine (5 mL). The combined organic layers were dried over $Na_2SO_4$, filtered, and concentrated *in vacuo*. Purification was achieved with a gradient of 2 – 6% MeOH/DCM to give the product as a yellow powder. Yield: 0.065 g (48%). $R_f$: 0.51 (MeOH/DCM = 1:9). HPLC purity 96%, RT = 4.26 min with method 3. $^1$H NMR (500 MHz, $CDCl_3$): δ 7.47 – 7.41 (m, 5H, H-17, H-17', and NH or NH'), 7.03 (t, *J* = 6.4 Hz, 2H, NH and NH'), 7.36 (t, *J* = 5.6 Hz, 1H, NH or NH'), 6.92 (m, 4H, H-18 and H-18'), 6.67 (m, 2H, NH and NH'), 5.46 (s, 2H, H-15 and H-15'), 4.96 (d, *J* = 13.1 Hz, 1H, H-14 or H-14'), 4.96 (d, *J* = 13.1 Hz, 1H, H-14 or H-14'), 4.44 (d, *J* = 13.1 Hz, 1H, H-14 or H-14'), 4.42 (d, *J* = 13.1 Hz, 1H, H-14 or H-14'), 4.08 (s, 1H, H-4 or H-4'), 4.08 (s, 1H, H-4 or H-4'), 3.81 (s, 6H, H-20 and H-20'), 3.67 (m, 4H, H-1 and H-1'), 3.55 – 3.25 (m, 12H, H-6, H-6', H-9, H-9', H-10, and H-10'), 2.41 (t, *J* = 6.4 Hz, 2H, H-7 or H-7'), 2.41 (t, *J* = 6.4 Hz, 2H, H-7 or H-7'), 1.44 (s, 6H, H-13 and H-13'), 1.10 (s, 6H, H-3 and H-3'), 1.06 (s, 6H, H-3 and H-3'). $^{13}$C NMR (126 MHz, $CDCl_3$): δ 173.7 (C-11 or C-11'), 173.7 (C-11 or C-11''), 172.2 (C-8 or C-8'), 172.1 (C-8 or C-8'), 170.1 (C-5 or C-5'), 170.1 (C-5 or C-5'), 160.4 (C-19 and C-19'), 130.2 (C-16 and C-16'), 127.7 (C-17 and C-17'), 113.9 (C-18 and C-18'), 101.5 (C-15 and C-15'), 84.0 (C-4 or C-4'), 83.9 (C-4 or C-4'), 81.5 (C-14 or C-14'), 81.5 (C-14 or C-14'), 78.6 (C-1 or C-1'), 78.5 (C-1 or C-1'), 74.2 (C-12 and C-12'), 55.5 (C-20 and C-20'), 39.8 (C-9', C-9', C-10, or C-10'), 39.8 (C-9, C-9', C-10, or C-10'), 39.4 (C-9, C-9', C-10, or C-10'), 39.4 (C-9, C-9', C-10, or C-10'), 36.5 (C-7 or C-7'), 36.4 (C-7 or C-7'), 35.2 (C-6 and C-6'), 33.2 (C-2 and C-2'), 24.3 (C-13 and C-13'), 22.0 (C-3, C-3'), 22.0 (C-3, C-3'), 19.3 (C-3, C-3a'). HRMS (ESI$^+$) *m/z* $[C_{23}H_{34}N_4NaO_9]^+$ calcd. 533.2218, found 533.2201.

**Synthesis and characterization of compound DD-03-001 (1-06)**

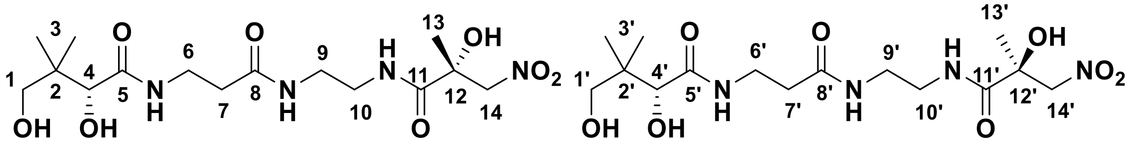

The reactant compound **DD-02-143** (0.037 g, 0.072 mmol) was dissolved into a solution of 9:1 AcOH:$H_2O$ (4 mL) and the mixture was stirred at room temperature for 4 h. The reaction mixture was diluted into distilled water (2 mL) and washed with toluene (3 × 5

mL). The aqueous layer was collected and lyophilized to afford the product as a yellow oil. Yield: 0.024 g (85%). $R_f$: 0.19 (MeOH/DCM = 1:9). HPLC purity 91%, RT = 7.12 min with method 1 and HPLC purity 93%, RT = 12.26 min with method 6. $^1$H NMR (500 MHz, MeOD): δ 4.96 (d, *J* = 12.5 Hz, 2H, H-14 and H-14'), 4.51 (d, *J* = 12.5 Hz, 2H, H-14 and H-14'), 3.89 (s, 2H, H-4 and H-4'), 3.50 – 3.38 (m, 12H, H-1, H-1', H-6, H-6', H-9, and H-9' or H-10 and H-10'), 3.32 (m, 4H, H-9 and H-9' or H-10 and H-10'), 2.43 (t, *J* = 6.7, 4H, H-7 and H-7'), 1.42 (s, 6H, H-13 and H-13'), 0.92 (s, 12H, H-3a and H-3'). $^{13}$C NMR (126 MHz, MeOD): δ 176.2 (C-11 and C-11'), 176.1 (C-5 and C-5'), 174.2 (C-8 and C-8'), 83.2 (C-14 and C-14'), 77.3 (C-4 and C-4'), 74.7 (C-12 and C-12'), 70.3 (C-1 and C-1'), 40.4 (C-2 and C-2'), 40.1 (C-9 and C-9' or C-10 and C-10'), 39.9 (C-9 and C-9' or C-10 and C-10'), 36.5 (C-7 and C-7'), 36.4 (C-6 and C-6'), 24.7 (C-13 and C-13'), 21.3 (C-3a and C-3'), 20.90 (C-3 and C-3'). HRMS (ESI$^+$) *m/z* $[C_{15}H_{28}N_4NaO_8]^+$ calcd. 415.1799, found 415.1798.

*Synthesis Route for Compound 1-07*

**Synthesis and characterization of compound DD-02-105**

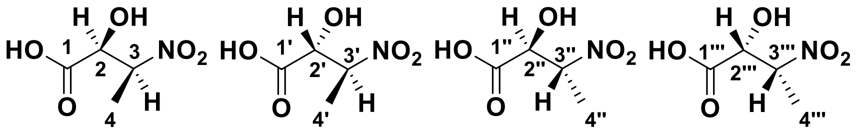

The reactants ethyl glyoxylate (50% toluene w/w; 2.04 g, 10.0 mmol) and lithium hydroxide monohydrate (0.948 g, 22.6 mmol) were dissolved in distilled water (50 mL) and the mixture was stirred at room temperature for 5 min. The reactant nitroethane (1.5 mL, 1.6 g, 21 mmol) was added, then the reaction was stirred at room temperature for 24 h. The reaction mixture was washed with DCM (3 × 10 mL), and then the aqueous layer was acidified to pH 2 with 2 N HCl before extraction of the product in EtOAc (3 × 50 mL). The combined organic layers were dried over $Na_2SO_4$ and filtered. The filtrate was concentrated *in vacuo* to give the product as a yellow oil. Yield: 0.668 g (45%). $R_f$: 0.00 (MeOH/DCM = 1:9). $^1$H NMR (500 MHz, $CDCl_3$): δ 4.97 (m, 4H, H-3, H-3', H-3'', and H-3'''), 4.88 (d, *J* = 3.3 Hz, 2H, H-2 and H-2''' or H-2' and H-2''), 4.39 (d, *J* = 4.5 Hz, 2H, H-2 and H-2''' or H-2' and H-2''), 1.60 (d, *J* = 6.8 Hz, 6H, H-4 and H-4''' or H-4' and H-4''), 1.49 (d, *J* = 6.8 Hz, 6H, H-4 and H-4''' or H-4' and H-4''). $^{13}$C NMR (126 MHz, $CDCl_3$): δ

174.0 (C-1 and C-1''' or C-1' and C-1''), 173.6 (C-1 and C-1''' or C-1' and C-1''), 85.2 (C-3 and C-3''' or C-3' and C-3''), 84.7 (C-3 and C-3''' or C-3' and C-3''), 73.3 (C-2 and C-2''' or C-2' and C-2''), 72.8 (C-2 and C-2''' or C-2' and C-2''), 15.6 (C-4 and C-4''' or C-4' and C-4''), 12.2 (C-4 and C-4''' or C-4' and C-4''). HRMS (ESI$^+$) *m/z* $[C_4H_6N_1Na_2O_5]^+$ 194.0036, found 194.0030.

**Synthesis and characterization of compound DD-03-081**

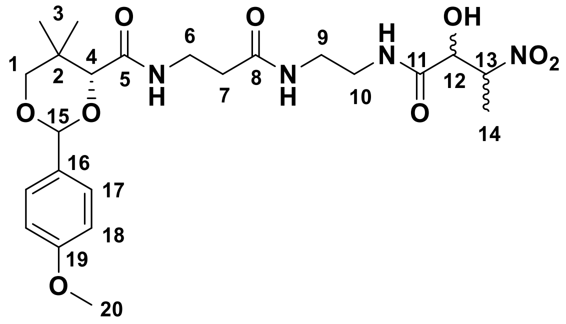

The reactants compound **DD-02-105** (0.120 g, 0.805 mmol), *N*-hydroxysuccinimide (0.108 g, 0.938 mmol), and *N,N'*-dicyclohexylcarbodiimide (0.209 g, 1.01 mmol) were dissolved in THF (5 mL) and stirred at room temperature for 4 h. The reactant compound **DD-05-006** (0.103 g, 0.271 mmol) was dissolved in THF (5 mL) and added. The mixture was stirred at room temperature for 18.5 h. The reaction mixture was filtered and concentrated *in vacuo*. Purification was achieved with a gradient of 2 – 6% MeOH/DCM to give the product as a pale yellow, stringy solid. Yield: 0.066 g (all diastereomers combined) (48%). $R_f$: 0.53 (*R,R* and *S,S* or *R,S* and *S,R*), 0.58 (*R,R* and *S,S* or *R,S* and *S,R*) (MeOH/DCM = 1:9). HPLC purity 96%, RT = 3.93 min, 4.27 min with method 3. Diastereomeric pair 1: *R,R* and *S,S* or *R,S* and *S,R*: $^1$H NMR (500 MHz, $CDCl_3$): δ 7.58 (m, 1H, NH), 7.40 (m, 2H, H-17), 7.08 (m, 1H, NH), 6.96 (m, 2H, H-18), 6.89 (m, 1H, NH), 5.44 (s, 1H, H-15), 4.92 (qd, *J* = 6.9, 2.7 Hz, 1H, H-13), 4.84 (d, *J* = 2.7 Hz, 0.5H, H-12), 4.83 (d, *J* = 2.8 Hz, 0.5H, H-12), 4.06 (s, 0.5H, H-4), 4.06 (s, 0.5H, H-4), 3.79 (s, 3H, H-20), 3.66 (m, 2H, H-1), 3.47 (q, *J* = 6.5 Hz, 2H, H-6), 3.42 – 3.23 (m, 4H, H-9 and H-10), 2.38 (t, *J* = 6.6 Hz, 2H, H-7), 1.41 (d, *J* = 6.8 Hz, 3H, H-14), 1.07 (s, 3H, H-3), 1.03 (s, 3H, H-3). $^{13}$C NMR (126 MHz, $CDCl_3$): δ 172.3 (C-5), 172.22 (C-5), 170.9 (C-8), 170.8 (C-8), 170.2 (C-11), 160.4 (C-19), 130.2 (C-16), 127.7 (C-17), 127.7 (C-17), 113.8 (C-18), 101.5 (C-15), 101.4 (C-15), 83.9 (C-4 and C-13), 78.4 (C-1), 72.6 (C-12), 72.6 (C-12), 55.4 (C-20), 39.5 (C-9 or C-10), 39.4 (C-9 and C-10), 39.4 (C-9 or C-10), 36.2 (C-7), 36.2 (C-7),

35.3 (C-6), 35.3 (C-6), 33.2 (C-2), 33.2 (C-2), 21.9 (C-3), 19.2 (C-3), 19.2 (C-3), 11.5 (C-14), 11.5 (C-14). HRMS (ESI$^+$) *m/z* $[C_{23}H_{34}N_4NaO_9]^+$ calcd. 533.2218, found 533.2218. Diastereomeric pair 2: *R,R* and *S,S* or *R,S* and *S,R*: $^1$H NMR (500 MHz, $CDCl_3$): δ 7.52 (q, *J* = 6.1 Hz, 1H, NH), 7.42 (m, 2H, H-17), 7.05 (m, 1H, NH), 6.90 (m, 2H, H-18), 6.89 (q, *J* = 5.8 Hz, 1H, NH), 5.45 (s, 1H, H-15), 5.01 (m, 1H, H-13), 4.26 (d, *J* = 3.8 Hz, 1H, H-12), 4.24 (d, *J* = 3.8 Hz, 1H, H-12), 4.06 (s, 0.5H, H-4), 4.06 (s, 0.5H, H-4), 3.80 (s, 3H, H-20), 3.67 (m, 2H, H-1), 3.52 – 3.28 (m, 6H, H-6, H-9, and H-10), 2.38 (t, *J* = 6.4 Hz, 1H, H-7,), 2.37 (t, *J* = 6.6 Hz, 1H, H-7), 1.60 (d, *J* = 7.0 Hz, 1.5H, H-14), 1.60 (d, *J* = 7.0 Hz, 1.5H, H-14), 1.08 (s, 3H, H-3), 1.04 (s, 1.5H, H-3), 1.03 (s, 1.5H, H-3). $^{13}$C NMR (126 MHz, $CDCl_3$): δ 172.2 (C-5), 172.2 (C-5), 171.6 (C-8), 171.6 (C-8), 170.2 (C-11), 170.1 (C-11), 160.4 (C-19'), 130.2 (C-16), 127.7 (C-17), 127.7 (C-17), 113.9 (C-18), 101.5 (C-15), 83.9 (C-4), 83.9 (C-4), 83.8 (C-13), 83.8 (C-13), 78.5 (C-1), 73.5 (C-12), 73.4 (C-12), 55.5 (C-20), 39.4 (C-9 or C-10), 39.4 (C-9 and C-10), 39.3 (C-9 or C-10), 36.3 (C-7), 36.3 (C-7), 35.3 (C-6), 35.3 (C-6), 33.2 (C-2), 21.9 (C-3), 19.2 (C-3), 19.2 (C-3), 15.6 (C-14), 15.6 (C-14). HRMS (ESI$^+$) *m/z* $[C_{23}H_{34}N_4NaO_9]^+$ calcd. 533.2218, found 533.2208.

**Synthesis and characterization of compound DD-03-085 (1-07)**

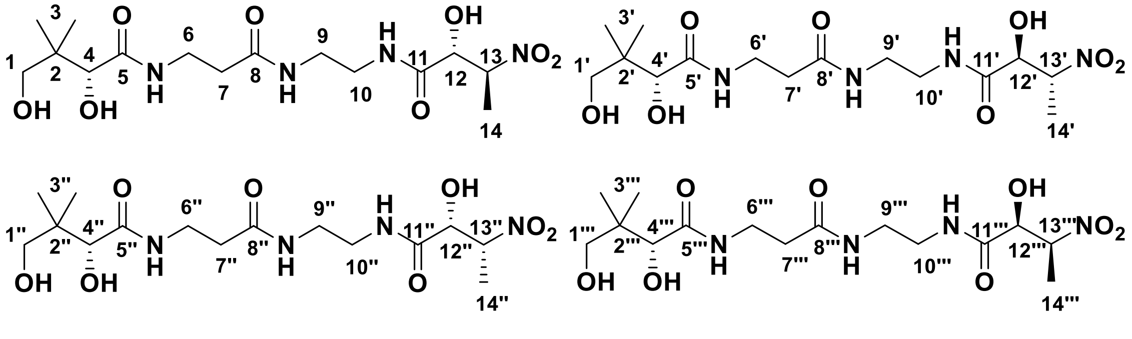

The reactant compound **DD-03-081** (0.029 g, 0.057 mmol) was dissolved into a solution of 9:1 AcOH:$H_2O$ (3 mL) and the mixture was stirred at room temperature for 4 h. The reaction mixture was diluted into distilled water (3 mL) and washed with toluene (3 × 5 mL). The aqueous layer was collected and concentrated *in vacuo* to afford the product as a pale, yellow oil. Yield: 0.019 g (85%). $R_f$: 0.20 (MeOH/DCM = 1:9). HPLC purity 96%, RT = 12.34 min with method 6 for diastereomeric pair 1, HPLC purity 93%, RT = 12.37 min with method 6 for diastereomeric pair 2. Diastereomeric pair *R,R* and *S,S* or *R,S* and *S,R*: $^1$H NMR (500 MHz, MeOD): δ 4.99 (m, 2H, H-13 and H-13' or H-13'' and H-13'''),

4.28 (d, $J$ = 4.1 Hz, 2H, H-12 and H-12' or H-12" and H-12'''), 3.89 (s, 2H, H-4 and H-4' or H-4" and H-4'''), 3.52 – 3.36 (m, 8H, H-1, H-1', H-6, and H-6' or H-1", H-1''', H-6", and H-6'''), 3.31 (m, 8H, H-9, H-9', H-10, and H-10' or H-9", H-9''', H-10", and H-10'''), 2.43 (t, $J$ = 6.6, 4H, H-7 and H-7' or H-7" and H-7'''), 1.58 (d, $J$ = 6.8 Hz, 6H, H-14 and H-14' or H-14" and H-14'''), 0.92 (s, 12H, H-3, H-3', H-3", and H-3'''). $^{13}$C NMR (126 MHz, MeOD): δ 176.1 (C-5 and C-5' or C-5" and C-5'''), 174.3 (C-8 and C-8' or C-8" and C-8'''), 173.7 (C-11 and C-11' or C-11" and C-11'''), 85.2 (C-13, C-13', C-13", or C-13'''), 85.1 (C-13, C-13', C-13", or C-13'''), 77.3 (C-4 and C-4' or C-4" and C-4'''), 74.3 (C-12 and C-12' or C-12" and C-12'''), 70.3 (C-1 and C-1' or C-1" and C-1'''), 40.3 (C-2 and C-2' or C-2" and C-2'''), 39.9 (C-9, C-9', C-9", C-9''', C-10, C-10', C-10", or C-10'''), 39.9 (C-9, C-9', C-9", C-9''', C-10, C-10', C-10", or C-10'''), 39.9 (C-9, C-9', C-9", C-9''', C-10, C-10', C-10", or C-10'''), 39.9 (C-9, C-9', C-9", C-9''', C-10, C-10', C-10", or C-10'''), 36.6 (C-7 and C-7' or C-7" and C-7'''), 36.4 (C-6 and C-6' or C-6" and C-6'''), 21.3 (C-3, C-3', C-3", and C-3'''), 15.70 (C-14, C-14', C-14", or C-14'''), 15.69 (C-14, C-14', C-14", or C-14'''). HRMS (ESI$^+$) *m/z* $[C_{15}H_{28}N_4NaO_8]^+$ calcd. 415.1799, found 415.1789.

Diastereomeric pair *R,R* and *S,S* or *R,S* and *S,R*: $^1$H NMR (500 MHz, MeOD): δ 4.95 (qd, $J$ = 6.8, 2.9 Hz, 2H, H-13 and H-13' or H-13" and H-13'''), 4.82 (d, $J$ = 2.9 Hz, 2H, H-12 and H-12' or H-12" and H-12'''), 3.90 (s, 2H, H-4 and H-4' or H-4" and H-4'''), 3.51 – 3.38 (m, 8H, H-1, H-1', H-6, and H-6' or H-1", H-1''', H-6", and H-6'''), 3.34 – 3.32 (m, 8H, H-9, H-9', H-10, and H-10' or H-9", H-9''', H-10" and H-10'''), 2.42 (t, $J$ = 6.6, 4H, H-7 and H-7' or H-7" and H-7'''), 1.42 (d, $J$ = 6.7 Hz, 6H, H-14 and H-14' or H-14" and H-14'''), 0.92 (s, 12H, H-3, H-3', H-3", and H-3'''). $^{13}$C NMR (126 MHz, MeOD) δ 176.1 (C-5 and C-5' or C-5" and C-5'''), 174.2 (C-8 and C-8' or C-8" and C-8'''), 173.2 (C-11 and C-11' or C-11" and C-11'''), 85.1 (C-13 and C-13' or C-13" and C-13'''), 77.3 (C-4 and C-4' or C-4" and C-4'''), 73.8 (C-12 and C-12' or C-12" and C-12'''), 70.3 (C-1 and C-1' or C-1" and C-1'''), 40.3 (C-2 and C-2' or C-2" and C-2'''), 39.9 (C-9 and C-9' or C-9" and C-9''' or C-10 and C-10' or C-10" and C-10'''), 39.9 (C-9 and C-9' or C-9" and C-9''' or C-10 and C-10' or C-10" and C-10'''), 36.6 (C-7 and C-7' or C-7" and C-7'''), 36.4 (C-6 and C-6' or C-6" and C-6'''), 21.3 (C-3 and C-3' or C-3" and C-3'''), 20.9 (C-3 and C-3' or C-3" and C-3'''), 11.6 (C-14 and C-14' or C-14" and C-14'''). HRMS (ESI$^+$) *m/z* $[C_{15}H_{28}N_4NaO_8]^+$ calcd. 415.1799, found 415.1791.

*Synthesis Route for Compound 1-08*

**Synthesis and characterization of compound DD-03-098**

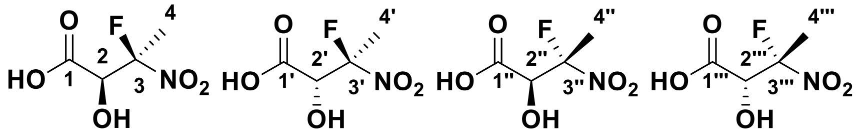

The reactants compound **DD-02-105** (0.444 g, 2.97 mmol) and lithium hydroxide monohydrate (0.253 g, 6.03 mmol) were dissolved into distilled water (40 mL) and the mixture was stirred at room temperature for 5 min. To this reaction mixture was added Selectfluor® (1.07 g, 3.02 mmol) before heating to 50°C for 2 h. The reaction mixture was acidified to pH 1, and the product was extracted into EtOAc (3 × 25 mL). The combined organic layers were dried over $Na_2SO_4$. The filtrate was concentrated *in vacuo* to give the product as a yellow oil. Yield: 0.285 g (57%). $R_f$: 0.75 (MeOH/DCM = 1:9). $^1$H NMR (300 MHz, $CDCl_3$): δ 4.83 (d, *J* = 22.5 Hz, 2H, H-2 and H-2'''), 4.80 (d, *J* = 19.7 Hz, 2H, H-2' and H-2''), 2.04 (d, *J* = 19.6 Hz, 6H, H-4 and H-4'''), 2.01 (d, *J* = 20.4 Hz, 6H, H-4' and H-4''). $^{13}$C NMR (76 MHz, $CDCl_3$): δ 170.6 (C-1, C-1', C-1'', and C-1'''), 118.5 (d, *J* = 244.5 Hz, C-3 and C-3''' or C-3' and C-3''), 118.1 (d, *J* = 245.1 Hz, C-3 and C-3''' or C-3' and C-3''), 83.3 (d, *J* = 23.1 Hz, C-2, C-2', C-2'', or C-2'''), 73.0 (d, *J* = 23.4 Hz, C-2, C-2', C-2'', or C-2'''), 72.9 (d, *J* = 24.5 Hz, C-2, C-2', C-2'', or C-2'''), 21.4 (d, *J* = 22.8 Hz, C-4 and C-4''' or C-4' and C-4''), 21.3 (d, *J* = 22.5 Hz, C-4 and C-4' or C-4'' and C-4'''). $^{19}$F NMR (283 MHz, $CDCl_3$): δ -127.2 (m, F-3' and F-3''), -132.0 (m, F-3 and F-3'''). HRMS (ESI$^+$) *m/z* $[C_4H_6FNNaO_5]^+$ calcd. 190.0122, found 190.0116.

**Synthesis and characterization of compound DD-03-100**

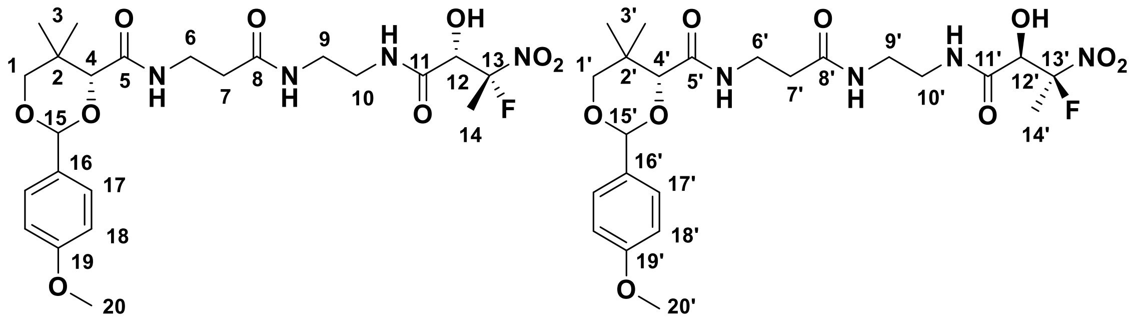

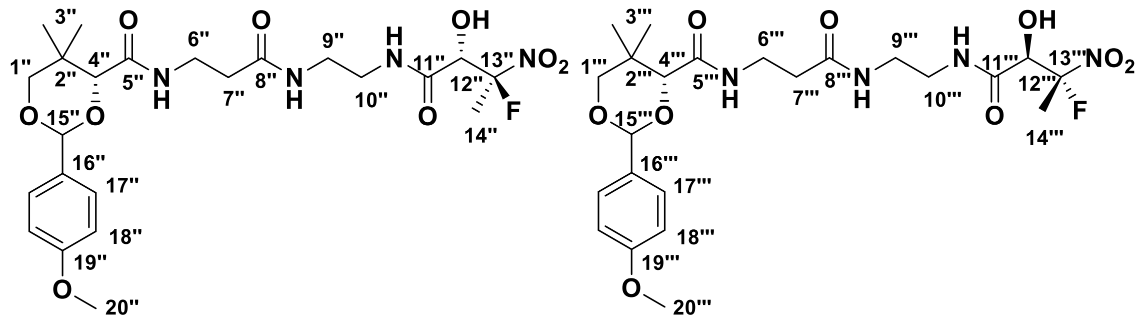

The reactants compound **DD-03-098** (0.245 g, 1.47 mmol), *N*-hydroxysuccinimide (0.192 g, 1.67 mmol), and *N,N'*-dicyclohexylcarbodiimide (0.335 g, 1.62 mmol) were dissolved in THF (5 mL) and the mixture was stirred at room temperature for 3 h. The reactant compound **DD-05-006** (0.161 g, 0.424 mmol) was dissolved in THF (5 mL) and added. The mixture was stirred at room temperature for 27.5 h. The reaction mixture was filtered and concentrated *in vacuo*. Purification was achieved with a gradient of 2 – 5% MeOH/DCM to give the product as a white solid. Yield: 0.160 g (71%), 0.044 g (20%) *S,R* and *R,S* mixture, 0.023 g (10%) *S,S* and *R,R* mixture, 0.093 g (41%). Mixture of diastereomers: HPLC purity 99%, RT = 3.79 min with method 3, and HPLC purity 91%, RT = 14.28 min with method 6.

Diastereomeric pair: *S,R* and *R,S* (**4.52**, **4.52'**). $R_f$: 0.53 (MeOH/DCM = 1:9). $^1$H NMR (500 MHz, MeOD): δ 7.52 (t, *J* = 5.8 Hz, 1H, NH or NH'), 7.46 (m, 1H, NH, or NH'), 7.42 (m, 4H, H-17 and H-17'), 7.04 (m, 2H, NH and NH'), 6.91 (m, 4H, H-18, and H-18'), 6.78 (m, 2H, NH and NH'), 5.47 (s, 1H, H-15 or H-15'), 5.46 (s, 1H, H-15 or H-15'), 4.70 (d, *J* = 22.8 Hz, 1H, H-12 or H-12'), 4.68 (d, *J* = 23.0 Hz, 1H, H-12 or H-12'), 4.10 (s, 1H, H-4 or H-4'), 4.08 (s, 1H, H-4 or H-4'), 3.81 (s, 6H, H-20 and H-20'), 3.68 (m, 4H, H-1 and H-1'), 3.57 – 3.21 (m, 12H, H-6, H-6', H-9, H-9', H-10, and H-10'), 2.38 (m, 4H, H-7 and H-7'), 2.00 (d, *J* = 19.8 Hz, 6H, H-14 and H-14'), 1.08 (s, 6H, H-3 and H-3'), 1.03 (s, 3H, H-3 and H-3'), 1.03 (s, 3H, H-3 and H-3'). $^{13}$C NMR (126 MHz, MeOD) δ 172.2 (C-5 or C-5'), 172.2 (C-5 or C-5'), 170.6 (C-8 or C-8'), 170.5 (C-8 or C-8'), 168.4 (C-11 or C-11'), 168.3 (C-11 or C-11'), 160.5 (C-19 and C-19'), 130.2 (C-16 and C-16'), 127.8 (C-17 or C-17'), 127.7 (C-17 or C-17'), 119.7 (d, *J* = 244.0 Hz, C-13 or C-13'), 119.6 (d, *J* = 244.0 Hz, C-13 or C-13'), 113.2 (C-18a and C-18'), 101.6 (C-15 and C-15'), 83.9 (C-4 or C-4'), 83.8 (C-4 or C-4'), 78.5 (C-1 or C-1'), 78.5 (C-1 or C-1'), 73.9 (d, *J* = 22.7 Hz, C-12 or C-12'), 73.8 (d, *J* = 22.8, C-12 or C-12'), 55.5 (C-20 and C-20'), 39.5 (C-9 and C-9' or C-10 and

C-10’), 39.4 (C-9, C-9’, C-10, or C-10’), 39.2 (C-9, C-9’, C-10, or C-10’), 36.6 (C-7 or C-7’), 36.5 (C-7 or C-7’), 35.3 (C-6 and C-6’), 33.2 (C-2 and C-2’), 21.9 (C-3 and C-3’), 21.9 (C-3 and C-3’), 21.4 (d, *J* = 22.3 Hz, C-14 and C-14’), 19.3 (C-3 and C-3’), 19.2 (C-3 and C-3’). $^{19}$F NMR (471 MHz, MeOD): δ -132.0 (m, F-13 and F-13’). HRMS (ESI$^+$) *m/z* $[C_{23}H_{33}FN_4NaO_9]^+$ calcd. 551.2124, found 551.2120.

Diastereomeric pair: *S,S* and *R,R* (**4.52”, 4.52’”**). $R_f$: 0.51 (MeOH/DCM = 1:9). $^1$H NMR (500 MHz, MeOD): δ 7.51 (m, 2H, NH” and NH’”), 7.42 (m, 4H, H-17” and H-17’”), 7.06 (m, 2H, NH” and NH’”), 6.90 (m, 4H, H-18” and H-18’”), 6.85 (m, 2H, NH” and NH’”), 5.45 (s, 2H, H-15” and H-15’”), 4.61 (d, *J* = 16.5 Hz, 1H, H-12” or H-12’”), 4.60 (d, *J* = 16.8 Hz, 1H, H-12” or H-12’”), 4.07 (s, 1H, H-4” or H-4’”), 4.06 (s, 1H, H-4” or H-4’”), 3.80 (s, 6H, H-20” and H-20’”), 3.67 (m, 4H, H-1” and H-1’”), 3.48 (m, 4H, H-6” and H-6’”), 3.38 – 3.29 (m, 8H, H-9”, H-9’”, H-10”, and H-10’”), 2.38 (t, *J* = 6.4 Hz, 4H, H-7” and H-7’”), 1.94 (d, *J* = 20.8 Hz, 3H, H-14” or H-14’”), 1.93 (d, *J* = 20.8 Hz, 3H, H-14” or H-14’”), 1.08 (s, 6H, H-3” and H-3’”), 1.04 (s, 6H, H-3” and H-3’”). $^{13}$C NMR (126 MHz, MeOD): δ 172.4 (C-5” or C-5’”), 172.3 (C-5” or C-5’”), 170.2 (C-8” or C-8’”), 170.2 (C-8” or C-8’”), 168.4 (C-11” or C-11’”), 168.4 (C-11” or C-11’”), 160.4 (C-19” and C-19’”), 130.2 (C-16” and C-16’”), 127.7 (C-17” or C-17’”), 127.7 (C-17 or C-17’”), 119.0 (d, *J* = 242.5 Hz, C-13” or C-13’”), 118.9 (d, *J* = 242.5 Hz, C-13” or C-13’”), 113.9 (C-18” and C-18’”), 101.5 (C-15” and C-15’”), 83.9 (C-4” or C-4’”), 83.8 (C-4” or C-4’”), 78.5 (C-1” and C-1’”), 73.6 (d, *J* = 22.4 Hz, C-12” and C-12’”), 55.5 (C-20” and C-20’”), 39.4 (C-9” and C-9’” or C-10” and C-10’”), 39.3 (C-9”, C-9’”, C-10”, or C-10’”), 39.3 (C-9”, C-9’”, C-10”, or C-10’”), 36.2 (C-7” and C-7’”), 35.3 (C-6” or C-6’”), 35.3 (C-6” or C-6’”), 33.2 (C-2” and C-2’”), 21.9 (C-3” and C-3’”), 21.3 (d, *J* = 22.4 Hz, C-14” or C-14’”), 21.2 (d, *J* = 22.4 Hz, C-14” or C-14’”), 19.2 (C-3a” and C-3’”), 19.22 (C-3” and C-3’”). $^{19}$F NMR (471 MHz, MeOD): δ -124.5 (m, F-13” or F-13’”), -124.8 (m, F-13” or F-13’”).

**Synthesis and characterization of compound DD-04-011 (1-08)**

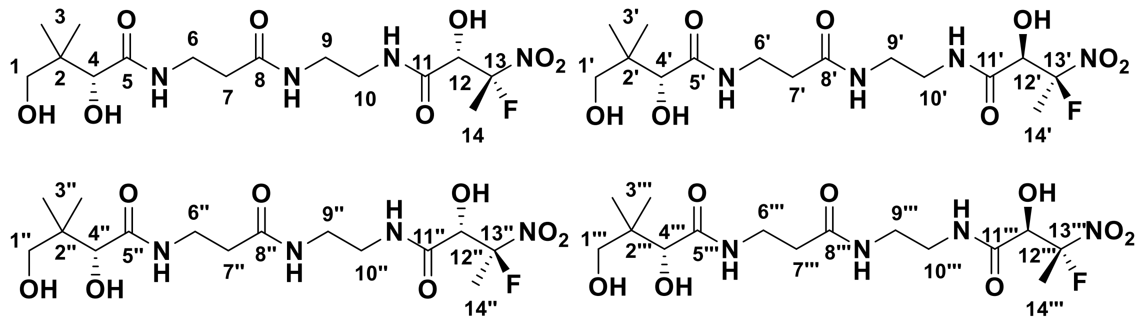

The reactant compound **DD-03-100** (0.091 g, 0.17 mmol) was dissolved into a solution of 9:1 AcOH:$H_2O$ (9 mL) and the mixture was stirred at room temperature for 4 h. The reaction mixture was diluted into distilled water (9 mL) and washed with toluene (3 × 20 mL). The aqueous layer was collected and concentrated *in vacuo* to afford a pale yellow oil as two pairs of diastereomers (2:1 of *S,R* and *R,S* : *S,S* and *R,R* (*i.e.* **4.30** and **4.30'** to **4.30"** and **4.30'''**). Yield: 0.024 g (34%). $R_f$: 0.06 (MeOH/DCM = 1:9). HPLC purity 95%, RT = 7.13 min with method 1, and HPLC purity 94%, RT = 12.44 min with method 6. $^1$H NMR (500 MHz, MeOD): δ 4.70 (d, *J* = 22.8 Hz, 2H, H-12 and H-12'), 4.66 (d, *J* = 17.4 Hz, 2H, H-12" and H-12'''), 3.89 (s, 4H, H-4, H-4', H-4", and H-4'''), 3.51 – 3.32 (m, 32H, H-1, H-1', H-1", H-1''', H-6, H-6', H-6", H-6''', H-9, H-9', H-9", H-9''', H-10, H-10', H-10", and H-10'''), 2.44 – 2.40 (m, 8H, H-7, H-7', H-7", and H-7'''), 1.97 (d, *J* = 19.8 Hz, 6H, H-14 and H-14'), 1.97 (d, *J* = 14.1 Hz, 6H, H-14" and H-14'''), 0.92 (s, 24H, H-3, H-3', H3", H-3'''). $^{13}$C NMR (126 MHz, MeOD): δ 176.1 (C-5, C-5', C-5", and C-5'''), 174.3 (C-8, C-8', C-8", and C-8'''), 170.8 (C-11, C-11', C-11", and C-11'''), 121.1 (d, *J* = 242.6 Hz, C-13, C-13', C-13", and C-13'''), 77.4 (C-4, C-4', C-4", and C-4'''), 74.6 (d, *J* = 22.6, C-12 and C-12'), 74.5 (d, *J* = 24.2, C-12" and C-12'''), 70.3 (C-1, C-1', C-1", and C-1'''), 40.4 (C-2, C-2', C-2", and C-2'''), 39.9 (C-9, C-9', C-9", C-9''', C-10, C-10', C-10", and C-10'''), 36.6 (C-7, C-7', C-7", and C-7'''), 36.4 (C-6, C-6', C-6", and C-6'''), 21.7 (d, *J* = 23.1 Hz, C-14 and C-14' or C-14" and C-14'''), 21.6 (d, *J* = 23.3 Hz, C-14 and C-14' or C-14" and C-14'''), 21.3 (C-3a, C-3a', C-3a", and C-3a''' or C-3b, C-3b', C-3b", and C-3b'''), 20.9 (C-3, C-3', C-3", and C-3'''), $^{19}$F NMR (471 MHz, MeOD) δ -126.5 (m, F-13" and F-13'''), -133.0 (m, F-13 and F-13'). HRMS (ESI$^+$) *m/z* $[C_{15}H_{27}FN_4NaO_8]^+$ calcd. 433.1705, found 433.1708.

*Synthesis Route for Compound 1-11*

**Synthesis and characterization of compound DD-05-090**

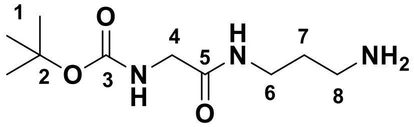

The reactants (*tert*-butoxycarbonyl)glycine (0.876 g, 5.00 mmol), *N*-hydroxysuccinimide (0.637 g, 5.53 mmol), and *N,N'*-dicyclohexylcarbodiimide (1.394 g, 6.756 mmol) were dissolved in THF (50 mL) and the mixture was stirred at room temperature for 4 h. A solution of 1,3-diaminopropane (1.869 g, 2.1 mL, 25 mmol) in THF (50 mL) was next added and the mixture was stirred at room temperature for 21 h. The reaction mixture was filtered and concentrated *in vacuo*. Purification was achieved with a gradient of 5:94:1 to 15:84:1 MeOH:DCM:$NEt_3$ then to 15:83:2 MeOH:DCM:$NH_4OH_{(aq)}$ (28%) to give the product as a colourless oil. Yield: 1.007 g (87%). $R_f$: 0.05 (MeOH/DCM/$NEt_3$ = 10:89:1). $^1$H NMR (500 MHz, $CDCl_3$): δ 7.22 (bs, 1H, NH), 5.49 (t, *J* = 5.8 Hz, 1H, NH), 3.73 (m, 2H, H-4), 3.34 (q, *J* = 6.2 Hz, 2H, H-6), 2.76 (t, *J* = 6.4 Hz, 2H, H-8), 2.05 (bs, 3H, NH), 1.62 (p, *J* = 6.5, 2H, H-7), 1.41 (s, 9H, H-1) .$^{13}$C NMR (126 MHz, $CDCl_3$) δ 169.8 (C-5), 156.3 (C-3), 80.1 (C-2), 44.4 (C-4), 39.9 (C-8), 37.8 (C-6), 32.0 (C-7), 28.4 (C-1). HRMS (ESI$^+$) *m/z* $[C_{10}H_{21}N_3NaO_3]^+$ calcd. 254.1475, found 254.1473.

**Synthesis and characterization of compound DD-05-095**

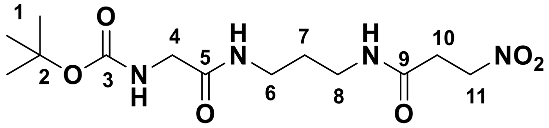

The reactants 3-nitropropionic acid (0.242 g, 2.03 mmol), *N*-hydroxysuccinimide (0.260 g, 2.26 mmol), and *N,N'*-dicyclohexylcarbodiimide (0.462 g, 2.24 mmol) were dissolved in THF (10 mL) and the mixture was stirred at room temperature for 4 h. A solution of compound **DD-05-090** (0.464 g, 2.01 mmol) in THF (10 mL) was next added and the mixture was stirred at room temperature for 15 h. The reaction mixture was filtered and concentrated *in vacuo*. The residue was redissolved into DCM (10 mL) and washed with distilled water (2 × 10 mL). The aqueous layer was further extracted with DCM (2 × 10 mL). The combined organic layers were dried over $Na_2SO_4$ and filtered. The filtrate was concentrated *in vacuo*. Purification was achieved with a gradient of 0 – 5% MeOH/DCM to give the product as a white solid. Yield: 0.154 g (23%). $R_f$: 0.38 (MeOH/DCM = 1:9). $^1$H NMR (500 MHz, $CDCl_3$): δ 7.18 (t, *J* = 6.1 Hz, 1H, NH), 7.04 (s, 1H, NH), 5.59 (t, *J* = 5.9 Hz, 1H, NH), 4.67 (m, 2H, H-11), 3.72 (m, 2H, H-4), 3.28 – 3.21 (m, 4H, H-6 and H-8), 2.81 (m, 2H, H-10), 1.62 (p, *J* = 6.3, 2H, H-7), 1.41 (s, 9H, H-1) .$^{13}$C NMR (126 MHz, $CDCl_3$): δ 170.7 (C-9), 169.2 (C-5), 156.4 (C-3), 80.4 (C-2), 70.4 (C-11), 44.3 (C-4), 36.5 (C-6 or C-8), 36.4 (C-6 or C-8), 32.5 (C-10), 29.2 (C-7), 28.3 (C-1). HRMS (ESI$^+$) *m/z* $[C_{13}H_{24}N_4NaO_6]^+$ calcd. 355.1588, found 355.1571.

**Synthesis and characterization of compound DD-05-101**

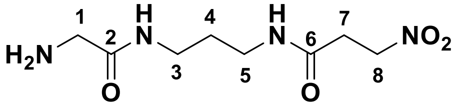

The reactant compound **DD-05-095** (0.027 g, 0.081 mmol) was dissolved into a solution of 1:1 DCM:TFA (3 mL). The reaction mixture was stirred at room temperature for 2.5 h. The solvent was removed *in vacuo* and the product did not require any purification. The product was a yellow oil. Yield: 0.030 g (99%). $R_f$: 0.05 (MeOH/DCM = 1:9). $^1$H NMR (500 MHz, MeOD): δ 4.71 (m, 2H, H-8), 3.66 (s, 2H, H-1), 3.72 (t, 2H, *J* = 6.8 Hz, H-3), 3.23 (t, 2H, *J* = 6.9 Hz, H-5), 2.85 (m, 2H, H-7), 1.71 (p, 2H, *J*: 6.8 Hz, H-4).$^{13}$C NMR (126 MHz, MeOD): δ 171.7 (C-6), 167.3 (C-2), 71.3 (C-8), 41.5 (C-1), 37.8 (C-3), 37.7 (C-5), 33.1 (C-7), 30.1 (C-4). HRMS (ESI$^+$) *m/z* $[C_8H_{16}N_4NaO_4]^+$ calcd. 255.1064, found 255.1059.

**Synthesis and characterization of compound DD-06-070 (1-11)**

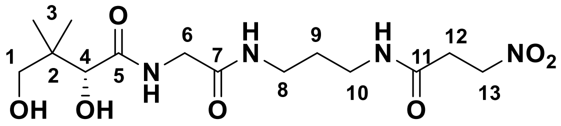

The reactants compound **DD-05-101** (0.350 g, 1.51 mmol) and D-pantolactone (0.980 g, 7.53 mmol) were dissolved in EtOH (3 mL) and the mixture was stirred at 115$^o$C in a sealed container for 14 h. The reaction mixture was concentrated *in vacuo*. Purification was achieved with a gradient of 5 – 12% MeOH/DCM to give the product as yellow oil. Yield: 0.061 g (11%). $R_f$: 0.07 (MeOH/DCM = 1:9). HPLC purity 96%, RT = 12.34 min with method 6. $^1$H NMR (500 MHz, MeOD): δ 4.72 (m, 2H, H-13), 3.91 (s, 1H, H-4), 3.86 (m,

2H, H-6), 3.45 (m, 2H, H-1), 3.25 – 3.21 (m, 4H, H-8 and H-10), 2.86 (m, 2H, H-12), 1.68 (p, $J$ = 6.7 Hz, 2H, H-9), 0.99 (s, 3H, H-3), 0.95 (s, 3H, H-3). $^{13}$C NMR (126 MHz, MeOD): δ 176.9 (C-5), 171.8 (C-7 and C-11), 77.6 (C-1), 71.3 (C-13), 61.5 (C-4), 43.1 (C-6), 40.6 (C-2), 37.7 (C-8 or C-10), 37.5 (C-8 or C-10), 33.1 (C-12), 30.1 (C-9), 22.1 (C-3), 21.3 (C-3). HRMS (ESI$^+$) *m/z* $[C_{14}H_{26}N_4NaO_7]^+$ calcd. 385.1694, found 385.1693.

*Synthesis Route for Compound 1-12*

**Synthesis and characterization of compound DD-04-104**

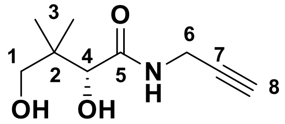

The known compound **DD-04-104** was prepared following a previously reported procedure.[6] Yield: 0.054 g (30%). $R_f$: 0.35 (MeOH/DCM = 1:9). $^1$H NMR (500 MHz, $CDCl_3$): δ 7.21 (t, $J$ = 5.7 Hz, 1H, NH), 4.05 – 4.03 (m, 3H, H-4 and H-6), 3.49 (s, 2H, H-1), 2.24 (t, $J$ = 2.5 Hz, 1H, H-8), 0.98 (s, 3H, H-3), 0.91 (s, 3H, H-3).$^{13}$C NMR (126 MHz, $CDCl_3$): δ 173.1 (C-5), 79.3 (C-7), 77.5 (C-4), 71.7 (C-8), 71.3 (C-1), 39.5 (C-2), 28.8 (C-6), 21.1 (C-3), 20.5 (C-3).

**Synthesis and characterization of compound DD-07-111**

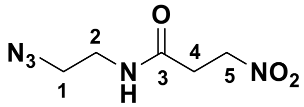

The reactants 2-bromoethylamine hydrobromide (0.629 g, 3.07 mmol) and sodium azide (0.401 g, 6.17 mmol) were dissolved into water (15 mL) before heating to reflux for 22 h. The product was extracted into DCM (3 × 15 mL) and the combined organic layers were concentrated *in vacuo* to a volume of ca. 10 mL, before dilution into THF (10 mL) and concentration *in vacuo* to remove the remaining DCM. This 2-azidoethylamine product was used without further purification. Separately, the reactants 3-nitropropionic acid (0.130 g, 1.09 mmol), *N*-hydroxysuccinimide (0.148 g, 1.29 mmol), and *N,N'*-dicyclohexylcarbodiimide (0.263 g, 1.27 mmol) were dissolved in THF (10 mL) and the mixture was stirred at room temperature for 4 h. The solution of 2-azidoethylamine was next added and the mixture was stirred at room temperature for 18 h. The reaction mixture was concentrated *in vacuo*. Purification was achieved with 100% DCM to give the product as white solid. Yield: 0.066 g (11%). $R_f$: 0.70 (MeOH/DCM = 1:9). $^1$H NMR (500 MHz, $CDCl_3$): δ 4.75 (t, *J* = 6.3 Hz, 2H, H-5), 3.35 (t, *J* = 6.3 Hz, 2H, H-4), 2.86 (s, 4H, H-1 and H-2). $^{13}$C NMR (126 MHz, $CDCl_3$): δ 168.6 (C-3), 68.7 (C-5), 28.5 (C-4), 25.7 (C-1 and C-2). HRMS (ESI$^+$) *m/z* $[C_5H_9N_5NaO_3]^+$ calcd. 210.0598, found 210.0592.

**Synthesis and characterization of compound DD-04-109 (1-12)**

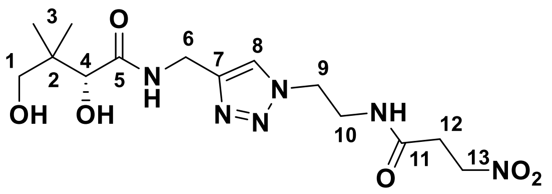

The reactants compound **DD-04-104** (0.038 g, 0.21 mmol), compound **DD-07-111** (0.064 g, 0.34 mmol), and sodium ascorbate (0.019 g, 0.19 mmol) were dissolved into a solution of 1:1 *i*PrOH:$H_2O$ (2 mL). Copper sulfate pentahydrate (0.008 g, 0.05 mmol) was then added and the reaction mixture was stirred at room temperature for 40 min. The reaction mixture was diluted into distilled water (5 mL) and washed with DCM (2 × 5 mL). The aqueous layer was concentrated *in vacuo*. Purification was achieved with a gradient of 3 – 10% MeOH/DCM to give the product as an off-white foam. Yield: 0.050 g (66%). $R_f$: 0.16 (MeOH/DCM = 1:9). HPLC purity 98%, RT = 4.80 min with method 5, and HPLC purity 98%, RT = 12.33 min with method 6. $^1$H NMR (500 MHz, MeOD): δ 7.82 (s, 1H, H-8), 4.69 (m, 2H, H-13), 4.52 – 4.44 (m, 4H, H-6 and H-9), 3.94 (s, 1H, H-4), 3.65 (m, 2H, H-10), 3.43 (m, 2H, H-1), 2.81 (m, 2H, H-12), 0.92 (s, 6H, H-3). $^{13}$C NMR (126 MHz, MeOD): δ 176.1 (C-5), 172.0 (C-11), 146.2 (C-7), 124.7 (C-8), 77.4 (C-4), 71.1 (C-13), 70.2 (C-1), 50.4 (C-9), 40.6 (C-2), 40.5 (C-10), 35.1 (C-6), 32.9 (C-12), 21.4 (C-3), 20.9 (C-3). HRMS (ESI$^+$) *m/z* $[C_{14}H_{24}N_6NaO_6]^+$ calcd. 395.1650, found 395.1642.

*Synthesis Route for Compound 1-13*

**Synthesis and characterization of compound DD-07-051**

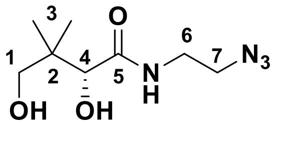

The known compound **DD-07-051** was prepared following a previously reported procedure with modifications.[7] The reactant sodium azide (0.391 g, 6.01 mmol) was dissolved into water (10 mL) before addition of 2-bromoethylamine hydrobromide (compound **5.41**; 0.415 g, 2.02 mmol) and the mixture was heated to reflux for 24 h. The reaction mixture was then transferred to a separatory funnel, and the pH was increased to >12 using 5 M NaOH. The intermediate 2-azidoethylamine was extracted into ethyl acetate (5 mL). The organic layer was dried over $Na_2SO_4$ and filtered. To this solution were added D-pantolactone (0.392 g, 3.01 mmol) and triethylamine (300 µL, 2.15 mmol) before heating to 115°C in a sealed vessel for 3 days. The reaction mixture was concentrated *in vacuo*, and then purification was achieved with a gradient of 0 – 4% MeOH/DCM to afford the product as a colourless oil. Yield: 0.049 g (11%). $R_f$: 0.40 (MeOH/DCM = 1:9). $^1$H NMR (500 MHz, $CDCl_3$): δ 7.13 (s, 1H, NH), 4.06 (s, 1H, H-4), 3.56 – 3.41 (m, 6H, H-1 and H-6 and H-7), 1.03 (s, 3H, H-3), 0.93 (s, 3H, H-3).

**Synthesis and characterization of compound DD-07-014**

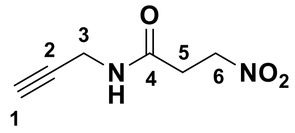

The reactants 3-nitropropionic acid (0.724 g, 6.08 mmol), *N*-hydroxysuccinimide (0.838 g, 7.28 mmol), and *N*,*N*'-dicyclohexylcarbodiimide (1.496 g, 7.25 mmol) were dissolved into THF (18 mL) and the mixture was stirred at room temperature for 4 h. To this solution was added propargylamine (compound **5.40**; 0.390 µL, 6.08 mmol) dissolved in THF (12 mL), and the reaction mixture was stirred at room temperature for 20.5 h. The reaction mixture was concentrated *in vacuo*, and the residue was redissolved into DCM (10 mL), before washing with water (2 × 10 mL), followed by extraction of the product from the aqueous layer with DCM (5 × 10 mL). The combined organic layers were dried over $Na_2SO_4$, filtered, then concentrated *in vacuo*. The residue was redissolved in chilled MeCN (10 mL) and cooled at -20°C. The solution was filtered, and the filtrate was concentrated *in vacuo*. Purification was achieved with 100% DCM to afford the product as a fluffy white powder. Yield: 0.390 g (41%). $R_f$: 0.55 (MeOH/DCM = 1:9). HPLC purity 95%, RT = 12.95 min with method 6. $^1$H NMR (500 MHz, $CDCl_3$): δ 6.16 (bs, 1H, NH),

4.71 (m, 2H, H-6), 4.07 (dd, *J* = 5.3, 2.6 Hz; 2H, H-3), 2.86 (m, 2H, H-5), 2.26 (t, *J* = 2.6 Hz, 1H, H-1). $^{13}$C NMR (126 MHz, $CDCl_3$): δ 168.2 (C-4), 79.1 (C-2), 72.1 (C-1), 70.1 (C-6), 32.4 (C-5), 29.6 (C-3). HRMS (ESI$^+$) *m/z* $[C_6H_8N_2NaO_3]^+$ calcd. 179.0427, found 179.0421.

**Synthesis and characterization of compound DD-07-076 (1-13)**

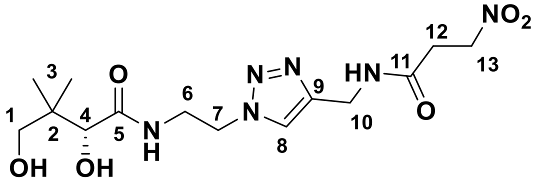

The reactants compound **DD-07-051** (0.057 g, 0.26 mmol), compound **DD-07-014** (0.034 g, 0.22 mmol), and sodium ascorbate (0.050 g, 0.25 mmol) were dissolved in aqueous isopropanol (1:1, 5 mL). To this solution was added copper sulfate (0.008 g, 0.05 mmol), and the mixture was stirred at room temperature for 1 h. The reaction was quenched using 3 drops of a sat. aq. ethylenediamine tetraacetic acid (EDTA) solution at pH 12. The product was extracted into chloroform (5 mL), followed by chloroform:isopropanol (2:3, 5 mL). The combined organic layers were dried over $Na_2SO_4$, then filtered and concentrated *in vacuo*. Purification was achieved with a gradient of 5 – 12% MeOH/DCM to give the product as a white solid. Yield: 0.066 g (81%). $R_f$: 0.14 (MeOH/DCM = 1:9). HPLC purity 98%, RT = 7.14 min with method 1. $^1$H NMR (500 MHZ, MeOD): δ 7.85 (s, 1H, H-8), 4.73 (m, 2H, H-13), 4.52 (t, *J* = 6.0 Hz, 2H, H-7), 4.44 (s, 2H, H-10), 3.86 (s, 1H, H-4), 3.70 (m, 2H, H-6), 3.37 (m, 2H, H-1), 2.89 (m, 2H, H-12), 0.86 (s, 3H, H-3), 0.86 (s, 3H, H-3). $^{13}$C NMR (126 MHz, MeOD): δ 176.6 (C-5), 171.6 (C-11), 146.4 (C-9), 124.7 (C-8), 77.3 (C-4), 71.2 (C-13), 70.2 (C-1), 50.5 (C-7), 40.3 (C-2), 39.9 (C-6), 35.7 (C-10), 33.0 (C-12), 21.3 (C3), 20.8 (C-3). HRMS (ESI$^+$) *m/z* $[C_{14}H_{24}N_6NaO_6]^+$ calcd. 395.1650, found 395.1642.

*Synthesis Route for Compound 1-17*

**Synthesis and characterization of compound DD-07-088**

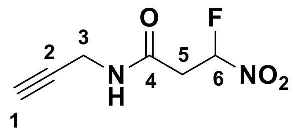

The reactants compound **DD-07-014** (0.076 g, 0.49 mmol) and lithium hydroxide monohydrate (0.022 g, 0.52 mmol) were dissolved into aqueous acetonitrile (1:1 MeCN:$H_2O$) and stirred at room temperature for 5 min. To this solution was added Selectfluor® (0.180 g, 0.508 mmol) and the mixture was stirred at room temperature for 5 min. The reaction was quenched by addition into DCM, brine, and water (10 mL, 3 mL, 2 mL) in a separatory funnel. The organic phase was collected, and the product was further extracted into DCM (2 × 10 mL). The organic layers were combined and dried over $Na_2SO_4$, filtered, then concentrated *in vacuo*. Purification was achieved with 100% DCM to afford the product as a white solid. Yield: 0.063 g (75%). $R_f$: 0.62 (MeOH/DCM = 1:9). HPLC purity 95%, RT = 13.55 min with method 6. $^1$H NMR (500 MHz, $CDCl_3$): δ 6.34 (ddd, *J* = 48.7, 6.6, 4.1 Hz; 1H, H-6), 3.98 (d, *J* = 2.6 Hz, 2H, H-3), 3.22 (ddd, *J* = 23.7, 15.9, 4.0 Hz; 1H, H-5), 3.10 (ddd, *J* = 19.8, 15.9, 6.6 Hz, 1H, H-5), 2.61 (t, *J* = 2.6 Hz, 1H, H-1). $^{13}$C NMR (126 MHz, $CDCl_3$): δ 167.4 (d, *J* = 3.7 Hz, C-4), 109.6 (d, *J* = 236.6 Hz, C-6), 80.0 (C-2), 72.6 (C-1), 39.9 (d, *J* = 21.5 Hz, C-5), 29.6 (C-3). $^{19}$F NMR (471 MHz, $CDCl_3$): δ -149.0 (ddd, 48.8, 23.7, 19.9 Hz, F-6). HRMS (ESI$^+$) *m/z* $[C_6H_7FN_2NaO_3]^+$ calcd. 197.0333, found 197.0332.

**Synthesis and characterization of compound DD-07-102 (1-17)**

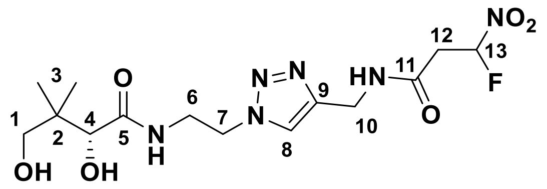

The reactants compound **DD-07-051** (0.049 g, 0.23 mmol), compound **DD-07-088** (0.037 g, 0.21 mmol), and sodium ascorbate (0.045 g, 0.23 mmol) were dissolved in aqueous isopropanol (1:1, 5 mL). To this solution was added copper sulfate (0.009 g, 0.06 mmol), and the mixture was stirred at room temperature for 1 h. The reaction was quenched using 3 drops of sat. aq. EDTA solution at pH 12. The mixture was concentrated *in vacuo*. Purification was achieved with a gradient of 5 – 10% MeOH/DCM to give the product as a white solid. Yield: 0.066 g (80%). $R_f$: 0.16 (MeOH/DCM = 1:9). HPLC purity 97%, RT = 7.20 min with method 1, and HPLC purity 95%, RT = 7.10 min with method 2. $^1$H NMR (500 MHz, MeOD): δ 7.88 (s, 1H, H-8), 6.36 (ddd, *J* = 48.7, 6.5, 4.1 Hz, 1H, H-13), 4.53 (t, *J* = 6.0 Hz, 2H, H-7), 4.45 (s, 2H, H-10), 3.87 (s, 1H, H-4), 3.71 (m, 2H, H-6), 3.34 (m, 2H, H-1), 3.18 (m, 2H, H-12), 0.86 (s, 3H, H-3), 0.86 (s, 3H, H-3). $^{13}$C NMR (126 MHz, MeOD): δ 176.5 (C-5), 167.8 (d, *J* = 3.7 Hz, C-11), 145.8 (C-9), 124.7 (C-8), 109.6 (d, *J* = 236.7 Hz, C-13), 77.3 (C-4), 70.2 (C-1), 50.5 (C-7), 40.3 (C-2), 40.0 (d, *J* = 21.5, C-12), 39.9 (C-6), 35.7 (C-10), 21.3 (C3), 20.8 (C-3). $^{19}$F NMR (471 MHz, MeOD): δ -148.8 (ddd, *J* = 48.7 23.9, 20.1 Hz, F-13). HRMS (ESI$^+$) *m/z* $[C_{14}H_{23}FN_6NaO_6]^+$ calcd. 413.1555, found 413.1540.

*Synthesis Route for Compound 2-01*

**Synthesis and characterization of compound DD-03-117**

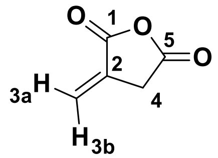

Synthesis of the known compound **DD-03-117** followed previously reported literature.[8] Yield: 1.07 g (90%). $^1$H NMR (300 MHz, $CDCl_3$): δ 6.48 (m, 1H, H-3a), 5.90 (m, 1H, H-3b), 3.60 (m, 2H, H-4). $^{13}$C NMR (75 MHz, $CDCl_3$): δ 168.21 (C-1), 164.80 (C-5), 130.37 (C-2), 126.70 (C-3), 33.57 (C-4).

**Synthesis and characterization of compound DD-07-131**

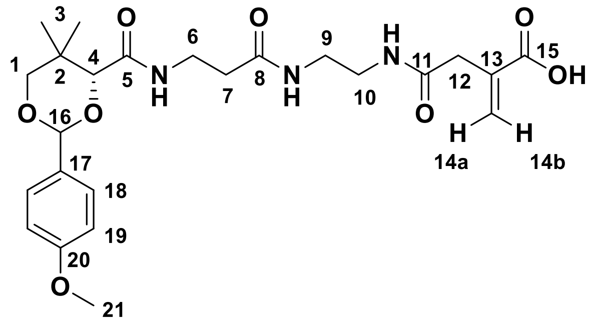

The reactant compound **DD-05-006** (0.053 g, 0.14 mmol) was dissolved in DCM (1 mL) at 0°C. Compound **DD-03-117** (0.018 g, 0.18 mmol) was next added and the mixture was stirred at 0°C for 25 min. The reaction mixture was concentrated *in vacuo*. Purification was achieved with a gradient of 3 – 12% MeOH/DCM to give the product as a white powder. Yield: 0.026 g (38%). $R_f$: 0.07 (MeOH/DCM = 1:9). $^1$H NMR (500 MHz, $CDCl_3$): δ 7.41 (m, 2H, H-18), 7.14 (t, *J* = 6.2 Hz, 1H, NH), 6.90 (m, 2H, H-19), 6.32 (bs, 1H, H-14b), 5.75 (bs, 1H, H-14a), 5.44 (s, 1H, H-16), 4.07 (s, 1H, H-4), 3.80 (s, 3H, H-21), 3.66 (m, 2H, H-1), 3.50 (m, 2H, H-6), 3.37 – 3.26 (m, 2H, H-9 and H-10), 3.16 (s, 2H, H-12), 2.39 (m, 2H, H-7), 1.08 (s, 3H, H-3), 1.05 (s, 3H, H-3). $^{13}$C NMR (126 MHz, $CDCl_3$): δ 172.0 (C-8), 171.8 (C-11), 170.4 (C-5), 169.9 (C-15), 160.4 (C-20), 135.0 (C-13), 130.2 (C-17), 129.8 (C-14) 127.7 (C-18), 113.9 (C-19), 101.5 (C-16), 83.8 (C-4), 78.5 (C-1), 55.5 (C-21), 40.5 (C-12), 39.4 (C-9 or C-10), 39.2 (C-9 or C-10), 36.0 (C-7), 35.7 (C-6), 33.2 (C-2), 22.0 (C3), 19.2 (C-3). HRMS (ESI$^+$) *m/z* $[C_{24}H_{33}N_3NaO_8]^+$ calcd. 514.2160, found 514.2140.

**Synthesis and characterization of compound DD-07-139 (2-01)**

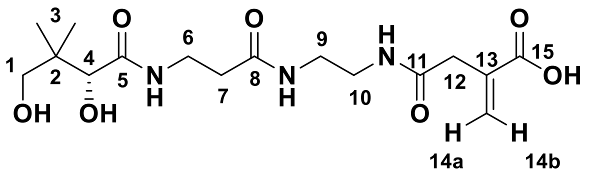

The reactant compound **DD-07-131** (0.019 g, 0.039 mmol) was dissolved into a solution of 9:1 AcOH:$H_2O$ (2 mL), and stirred at room temperature for 4.5 h. The reaction mixture was concentrated *in vacuo* and redissolved in distilled water (5 mL). The solution was then washed with DCM (5 × 5 mL). The aqueous layer was collected and concentrated *in vacuo* to afford the product as a colourless oil. Yield: 0.011 g (76%). $R_f$: 0.03 (MeOH/DCM = 1:9). $^1$H NMR (500 MHz, MeOD): δ 6.29 (bs, 1H, H-14b), 5.75 (bs, 1H, H-14a), 3.92 (s, 1H, H-4), 3.55 – 3.38 (m, 4H, H-1 and H-6), 3.38 – 3.26 (m, 4H, H-9 and H-10), 3.23 (s, 2H, H-12), 2.44 (t, 2H, *J* = 6.6 Hz, H-7), 0.94 (s, 3H, H-3), 0.94 (s, 3H, H-3). $^{13}$C NMR (126 MHz, MeOD): δ 176.1 (C-5), 174.1 (C-8), 173.7 (C-11), 170.5 (C-15), 137.6 (C-13), 128.6 (C-14), 77.3 (C-4), 70.3 (C-1), 40.6 (C-12), 40.6 (C-2), 40.1 (C-9 or C-10), 40.0 (C-9 or C-10), 36.6 (C-7), 36.4 (C-6), 21.3 (C-3), 21.0 (C-3). HRMS (ESI$^+$) *m/z* $[C_{16}H_{26}N_3Na_2O_7]^+$ calcd. 418.1561, found 418.1561.

*Synthesis Route for Compound 2-04*

**Synthesis and characterization of compound DD-03-141**

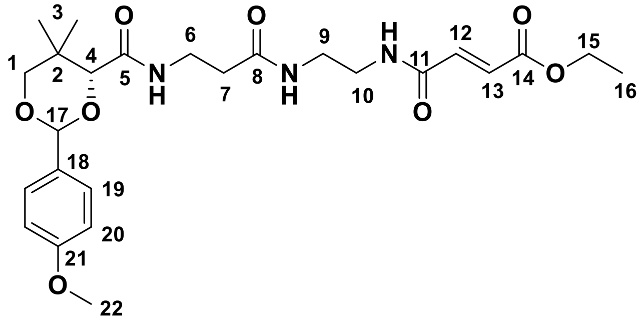

The reactants monoethyl fumarate (0.078 g, 0.54 mmol), *N*-hydroxysuccinimide (0.066 g, 0.57 mmol), and *N,N'*-dicyclohexylcarbodiimide (0.115 g, 0.557 mmol) were dissolved in THF (10 mL) and stirred at room temperature for 3.5 h. Separately, compound **DD-05-006** (0.190 g, 0.501 mmol) was dissolved in THF (10 mL) before addition to the initial mixture and stirring at room temperature for 19.5 h. The reaction mixture was filtered and concentrated *in vacuo*. Purification was achieved with a gradient of 2 – 5% MeOH/DCM to give the product as colourless oil. Yield: 0.082 g (32%) $R_f$: 0.50 (MeOH/DCM = 1:9). HPLC purity 97%, RT = 3.89 min with method 3, and HPLC purity 96%, RT = 5.73 min with method 4. $^1$H NMR (500 MHz, $CDCl_3$): δ 7.44 (m, 2H, H-19), 7.18 (t, *J* = 5.2 Hz, 1H, NH), 7.02 (t, *J* = 6.4 Hz, 1H, NH), 6.95 – 6.91 (m, 3H, H-20 and H-12 or H-13), 6.82 – 6.78 (m, 2H, H-12 or H-13 and NH), 5.49 (s, 1H, H-17), 4.25 (q, *J* = 7.2 Hz, 2H, H-15), 4.11 (s, 1H, H-4), 3.84 (s, 3H, H-22), 3.72 (m, 2H, H-1), 3.56 (m, 2H, H-6), 3.47 – 3.36 (m, 4H, H-9 and H-10), 2.45 (t, *J* = 6.2 Hz, 2H, H-7), 1.33 (t, *J* = 7.1 Hz, 3H, H-16), 1.12 (s, 3H, H-3), 1.10 (s, 3H, H-3). $^{13}$C NMR (126 MHz, $CDCl_3$): δ 172.2 (C-8), 170.1 (C-5), 165.7 (C-14), 164.5 (C-11), 160.5 (C-21), 136.4 (C-12 or C-13), 130.5 (C-12 or C-13), 130.2 (C-18), 127.7 (C-19), 113.9 (C-20), 101.5 (C-17), 84.0 (C-4), 78.5 (C-1), 61.3 (C-15), 55.5 (C-22), 40.4 (C-10), 39.7 (C-9), 36.7 (C-7), 35.2 (C-6), 33.3 (C-2), 22.0 (C-3), 19.3 (C-3), 14.3 (C-16). HRMS (ESI$^+$) *m/z* $[C_{25}H_{35}N_3NaO_8]^+$ calcd. 528.2316, found 528.2301.

**Synthesis and characterization of compound DD-06-147 (2-04)**

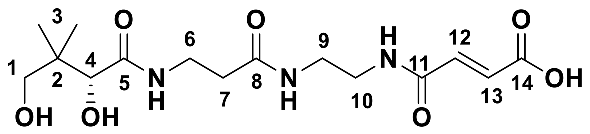

The reactant compound **DD-03-141** (0.0466 g, 0.0922 mmol) was dissolved into a solution of 1:1 THF:$H_2O$ (3 mL). To this solution was added lithium hydroxide monohydrate (0.005 g, 0.1 mmol) and the reaction mixture was stirred at room temperature for 19 h. The mixture was diluted into 3 mL distilled water and to it was added 5 drops of AcOH.The product was extracted in EtOAc (3 × 5 mL) and the combined the organic layers were dried over $Na_2SO_4$. The filtrate was concentrated *in vacuo*, redissolved in 9:1 AcOH:$H_2O$ (4.5 mL) and stirred at room temperature for 2.75 h. The reaction mixture was concentrated *in vacuo*, then the residue was redissolved into distilled water (5 mL) and washed with DCM (5 × 5 mL). The aqueous layer was concentrated *in vacuo* to afford a colourless oil. Yield: 0.0268 g (81%). $R_f$: 0.02 (MeOH/DCM = 1:9). $^1$H NMR (500 MHz, MeOD): δ 6.91 (d, *J* = 15.5 Hz, 1H, H-12 or H-13), 6.71 (d, *J* = 15.5 Hz, 1H, H-12 or H-13), 3.89 (s, 1H, H-4), 3.51 – 3.37 (m, 6H, H-1 and H-6 and H-9 or H-10) 3.35 (m, 2H, H-9 or H-10), 2.42 (t, *J* = 6.6, 2H, H-7), 0.92 (s, 6H, H-3). $^{13}$C NMR (126 MHz, MeOD): δ 176.1 (C-5), 174.2 (C-8), 169.3 (C-14), 167.0 (C-11), 136.6 (C-12 or C-13), 132.9 (C-12 or C-13), 77.3 (C-4), 70.3 (C-1), 40.6 (C-2), 40.2 (C-9 or C-10), 39.9 (C-9 or C-10), 36.6 (C-7), 36.4 (C-6), 21.4 (C-3), 20.9 (C-3). HRMS (ESI$^+$) *m/z* [$C_{15}H_{25}N_3Na_2O_7$]$^+$ calcd. 404.1404, found 404.1399.

*Synthesis Route for Compound 2-05*

**Synthesis and characterization of compound DD-04-007 (2-05)**

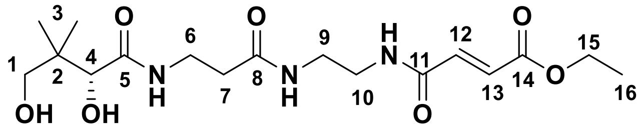

The reactant compound **DD-03-141** (0.081 g, 0.16 mmol) was dissolved into a solution of 9:1 AcOH:$H_2O$ (8 mL) and stirred at room temperature for 23 h. The reaction mixture was diluted into distilled water (8 mL) and washed with toluene (3 × 20 mL), then the product was extracted into EtOAc (3 × 20 mL) and the combined organic layers were dried over $Na_2SO_4$. The filtrate was concentrated *in vacuo* to afford the product as an off-white solid.

Yield: 0.046 g (72%). $R_f$: 0.10 (MeOH/DCM = 1:9). HPLC purity 95%, RT = 4.96 min with method 5, and HPLC purity 93%, RT = 12.49 min with method 6. $^1$H NMR (500 MHz, MeOD): δ 7.00 (d, $J$ = 15.5 Hz, 1H, H-12 or H-13), 6.71 (d, $J$ = 15.5 Hz, 1H, H-12 or H-13), 4.24 (q, $J$ = 7.1 Hz, H-15), 3.89 (s, 1H, H-4), 3.51 – 3.32 (m, 8H, H-1 and H-6 and H-9 and H-10), 2.42 (t, $J$ = 6.6, 2H, H-7), 1.30 (t, $J$ = 7.1 Hz, 3H, H-16), 0.92 (s, 6H, H-3). $^{13}$C NMR (126 MHz, MeOD): δ 176.0 (C-5), 174.2 (C-8), 166.8 (C-14), 166.4 (C-11), 137.6 (C-12 or C-13), 130.9 (C-12 or C-13), 77.6 (C-4), 70.3 (C-1), 62.2 (C-15), 40.3 (C-2), 40.3 (C-9 or C-10), 39.8 (C-9 or C-10), 36.6 (C-7), 36.4 (C-6), 21.4 (C-3), 20.9 (C-3), 14.4 (C-16). HRMS (ESI$^+$) $m/z$ $[C_{17}H_{29}N_3NaO_7]^+$ calcd. 410.1898, found 410.1912.

*Synthesis Route for Compound 2-06*

**Synthesis and characterization of compound DD-03-140**

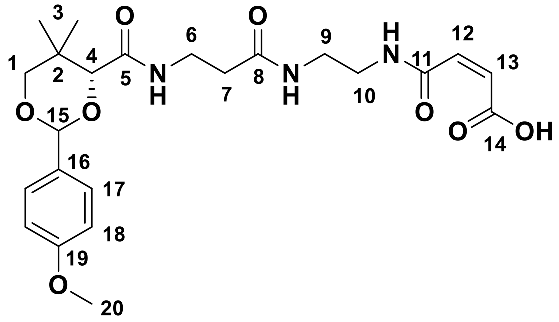

The reactants compound **DD-05-006** (0.152 g, 0.400 mmol) and maleic anhydride (0.045 g, 0.46 mmol) were dissolved in DCM (5 mL) and stirred at room temperature for 0.5 h. The reaction mixture was concentrated *in vacuo.* Purification was achieved with 5% MeOH/DCM to give the product as a white powder. Yield: 0.137 g (72%) $R_f$: 0.15 (MeOH/DCM = 1:9). $^1$H NMR (500 MHz, $CDCl_3$): δ 8.53 (t, $J$ = 5.1 Hz, 1H, NH), 7.43 (m, 2H, H-17), 7.08 (t, $J$ = 5.3 Hz, 1H, NH), 7.04 (t, $J$ = 6.5 Hz, 1H, NH), 6.92 (m, 2H, H-18), 6.34 (d, $J$ = 12.8 Hz, 1H, H-13), 6.24 (d, $J$ = 12.8 Hz, 1H, H-12), 5.46 (s, 1H, H-15), 4.08 (s, 1H, H-4), 3.81 (s, 3H, H-20), 3.68 (m, 2H, H-1), 3.56 (m 2H, H-6), 3.47 – 3.40 (m, 4H, H-9 and H-10), 2.43 (m, 2H, H-7), 1.11 (s, 3H, H-3), 1.05 (s, 3H, H-3). $^{13}$C NMR (126 MHz, $CDCl_3$): δ 172.8 (C-8), 170.4 (C-5), 166.5 (C-11), 165.8 (C-14), 160.5 (C-19), 135.5 (C-13), 132.1 (C-12), 130.1 (C-16), 127.7 (C-17), 113.9 (C-18), 101.6 (C-15), 83.9 (C-4), 78.5

(C-1), 55.5 (C-20), 41.0 (C-9), 38.6 (C-10), 36.9 (C-7), 35.6 (C-6), 33.3 (C-2), 22.0 (C-3), 19.2 (C-3). HRMS (ESI$^+$) *m/z* [$C_{23}H_{31}N_3NaO_8$]$^+$ calcd. 500.2003, found 500.1988.

**Synthesis and characterization of compound DD-07-129 (2-06)**

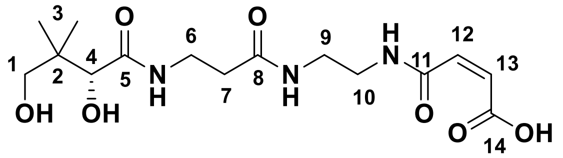

The reactant compound **DD-03-140** (0.0459 g, 0.0961 mmol) was dissolved into a solution of 9:1 AcOH:$H_2O$ (4.5 mL) and stirred at room temperature for 2.5 h. The reaction mixture was concentrated *in vacuo*. The residue was dissolved into distilled water (5 mL) and washed with DCM (4 × 5 mL). The aqueous layer was collected and concentrated *in vacuo* to afford a yellow solid. Yield: 0.0335 g (97%). $R_f$: 0.03 (MeOH/DCM = 1:9). $^1$H NMR (500 MHz, MeOD): δ 6.45 (d, $J$ = 12.5 Hz, 1H, H-12 or H-13), 6.25 (d, $J$ = 12.5 Hz, 1H, H-12 or H-13), 3.89 (s, 1H, H-4), 3.54 – 3.34 (m, 8H, H-1, H-6, H-9 and H-10), 2.43 (t, $J$ = 6.6, 2H, H-7), 0.92 (s, 6H, H-3). $^{13}$C NMR (126 MHz, MeOD): δ 176.1 (C-5), 174.2 (C-8), 168.4 (C-11 and C-14), 134.4 (C-12 or C-13), 132.9 (C-12 or C-13), 77.3 (C-4), 70.3 (C-1), 40.4 (C-2), 40.3 (C-9 or C-10), 39.5 (C-9 or C-10), 36.6 (C-7), 36.4 (C-6), 21.4 (C-3), 20.9 (C-3). HRMS (ESI$^+$) *m/z* [$C_{15}H_{25}N_3NaO_7$]$^+$ calcd. 382.1585, found 382.1571.

*Synthesis Route for Compound 2-12*

**Synthesis and characterization of compound DD-06-119**

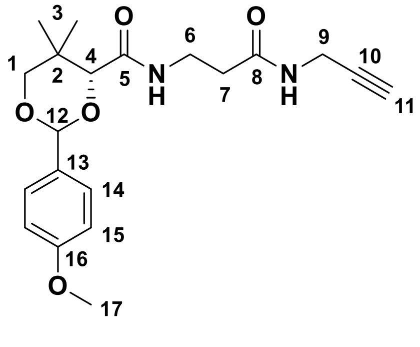

The reactants compound **DD-05-140** (0.338 g, 1.00 mmol), *N*-hydroxysuccinimide (0.133 g, 1.16 mmol), and *N,N'*-dicyclohexylcarbodiimide (0.241 g, 1.17 mmol) were dissolved in THF (10 mL) and stirred at room temperature for 4 h. To this solution was added propargylamine (75 μL, 0.065 g, 1.19 mmol) dissolved in THF (5 mL) and the mixture was stirred at room temperature for 19 h. The reaction mixture was filtered, concentrated *in vacuo*, redissolved into DCM (5 mL), and washed with distilled water (2 × 5 mL), then dried over $Na_2SO_4$ and filtered. The filtrate was concentrated *in vacuo*. Purification was achieved with a gradient of 0 – 2% MeOH/DCM to give the product as a pale yellow solid. Yield: 0.205 g (55%). $R_f$ = 0.42 (MeOH/DCM = 1:9). HPLC purity 96%, RT = 3.63 min with method 3, and HPLC purity 96%, RT = 5.57 min with method 4. $^1$H NMR (500 MHz, $CDCl_3$): δ 7.43 (m, 2H, H-14), 7.01 (t, *J* = 6.3 Hz, 1H, NH), 6.92 (m, 2H, H-15), 6.23 (t, *J* = 5.7 Hz, 1H, NH), 5.46 (s, 1H, H-12), 4.08 (s, 1H, H-4), 4.01 (m, 2H, H-9) 3.82 (s, 3H, H-17), 3.68 (m, 2H, H-1), 3.59 (m, 1H, H-6), 3.50 (m, 1H, H-6), 2.45 (t, *J* = 6.3 Hz, 2H, H-7), 2.19 (t, *J* = 2.5 Hz, 1H, H-11), 1.10 (s, 3H, H-3), 1.10 (s, 3H, H-3). $^{13}$C NMR (126 MHz, $CDCl_3$): δ 170.6 (C-8), 169.8 (C-5), 160.4 (C-16), 130.3 (C-13), 127.7 (C-14), 113.9 (C-15), 101.5 (C-12), 84.0 (C-4), 79.5 (C-10), 78.6 (C-1), 71.7 (C-11), 55.5 (C-17), 36.1 (C-7), 34.9 (C-6), 33.2 (C-2), 29.3 (C-9), 22.0 (C-3), 19.3 (C-3). HRMS (ESI$^+$) *m/z* $[C_{20}H_{26}N_2NaO_5]^+$ calcd. 397.1734, found 397.1718.

**Synthesis and characterization of compound DD-07-037**

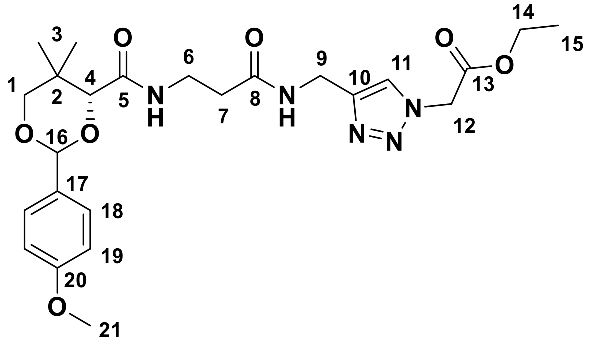

The reactants sodium azide (0.202 g, 3.11 mmol) and ethyl 2-bromoacetate (0.174 g, 1.04 mmol) were dissolved into aqueous acetone (1:1, 5 mL) and heated to reflux for 20 h. The reaction mixture was transferred to a separatory funnel, and the product ethyl 2-azidoacetate was extracted into diethyl ether (3 × 5 mL). The combined organic layers were dried over $Na_2SO_4$. Isopropanol (3 mL) was added into the organic layer and the ether was removed *in vacuo*. The reactant compound **DD-06-119** (0.033 g, 0.066 mmol) was dissolved into the solution of ethyl 2-azidoacetate in isopropanol, and to it was added a solution of copper sulfate (0.025 g, 0.016 mmol) in water (3 mL). Following brief mixing, sodium ascorbate (0.076 g, 0.38 mmol) was added. The mixture was stirred at room temperature for 21 h. The reaction was quenched with 5 drops of a sat. aq. EDTA solution (pH 12) before being transferred to a separatory funnel. The product was extracted in DCM (3 × 5 mL) and the combined organic layers were dried over $Na_2SO_4$, filtered, and concentrated *in vacuo*. Purification was achieved with a gradient of 0 – 3% MeOH/DCM to give the product as a colourless oil. Yield: 0.079 g (53%). $R_f$: 0.38 (MeOH/DCM = 1:9). HPLC purity 98%, RT = 3.84 min with method 3, and HPLC purity 100%, RT = 5.66 min with method 4. $^1$H NMR (500 MHz, $CDCl_3$): δ 7.59 (s, 1H, H-11), 7.43 (m, 2H, H-18), 7.09 (t, $J$ = 6.3 Hz, 1H, NH), 6.92 (m, 2H, H-19), 6.74 (t, $J$ = 5.8 Hz, 1H, NH-), 5.45 (s, 1H, H-16), 5.01 (m, 2H, H-12), 4.47 (m, 2H, H-9), 4.24 (q, $J$ = 7.1 Hz, 2H, H-14), 4.05 (s, 1H, H-4), 3.81 (s, 3H, H-21), 3.67 (m, 2H, H-1), 3.53 (m, 2H, H-6), 2.43 (t, $J$ = 6.3 Hz, 2H, H-7), 1.28 (t, $J$ = 7.1 Hz, 3H, H-15), 1.05 (s, 3H, H-3), 1.01 (s, 3H, H-3). $^{13}$C NMR (126 MHz, $CDCl_3$): 171.2 (C-8), 169.5 (C-5), 166.3 (C-13), 160.3 (C-20), 145.10 (C-10), 130.40 (C-17), 127.6 (C-18), 123.9 (C-11), 113.9 (C-19), 101.3 (C-16), 83.9 (C-4), 78.6 (C-1), 62.6 (C-14), 55.5 (C-21), 50.9 (C-12), 35.8 (C-7), 35.0 (C-9), 34.9 (C-6), 33.2 (C-2), 21.9 (C-3), 19.1 (C-3), 14.2 (C-15). HRMS (ESI$^+$) *m/z* $[C_{24}H_{33}N_5NaO_7]^+$ calcd. 526.2272, found 526.2260.

**Synthesis and characterization of compound DD-07-141 (2-12)**

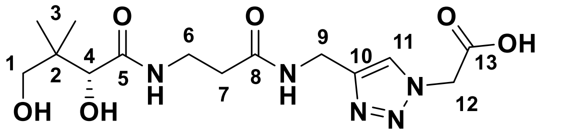

The reactant compound **DD-07-037** (0.085 g, 0.17 mmol) was dissolved into THF (2 mL) and to it was added a solution of lithium hydroxide monohydrate (0.008 g, 0.2 mmol) dissolved into distilled water (2mL). The mixture was stirred at room temperature for 25 h. The reaction mixture was then concentrated *in vacuo* and the residue was redissolved into distilled water (10 mL), before acidification with AcOH (5 drops). The product was extracted into DCM (3 × 10 mL) and the combined organic layers were dried over $Na_2SO_4$, filtered, then concentrated *in vacuo*. The residue was redissolved into a solution of 9:1 AcOH:$H_2O$ (3 mL), and stirred at room temperature for 3 h. The reaction mixture was concentrated *in vacuo* and redissolved in distilled water (5 mL). The solution was then washed with DCM (5 × 5 mL). The aqueous layer was collected and concentrated *in vacuo* to afford the product as a colourless oil. Yield: 0.017 g (28%). $R_f$: 0.00 (MeOH/DCM = 1:9). $^1$H NMR (500 MHz, MeOD): δ 7.91 (s, 1H, H-11), 5.21 (s, 2H, H-12), 4.46 (s, 2H, H-9), 3.88 (s, 1H, H-4), 3.50 (m, 2H, H-6), 3.41 (m, 2H, H-1), 2.46 (t, *J* = 6.6, 2H, H-7), 0.90 (s, 6H, H-3). $^{13}$C NMR (126 MHz, MeOD): δ 176.1 (C-5), 175.2 (C-8), 173.7 (C-13), 146.2 (C-10), 125.9 (C-11), 77.3 (C-4), 70.3 (C-1), 49.9 (C-9), 40.3 (C-2), 36.4 (C-7), 36.3 (C-6), 35.6 (C-12), 21.3 (C-3), 20.9 (C-3). HRMS (ESI$^+$) *m/z* $[C_{14}H_{22}N_5Na_2O_6]^+$ calcd. 402.1360, found 402.1342.

*Synthesis Route for Compound 2-13*

**Synthesis and characterization of compound DD-02-026**

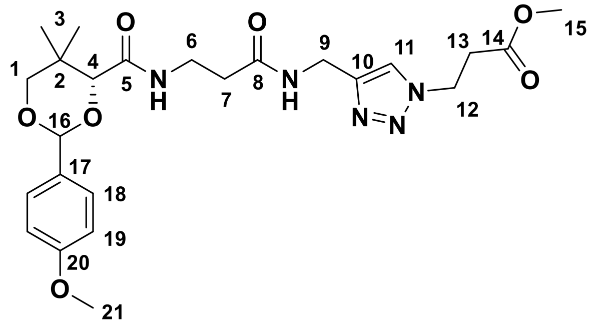

The reactants compound **DD-06-119** (0.064 g, 0.17 mmol) and methyl 3-azidopropanoate (0.033 g, 0.26 mmol) were dissolved into isopropanol (1.5 mL). To this solution was added a solution of copper sulfate pentahydrate (0.005 g, 0.03 mmol) in distilled water (1.5 mL). To this mixture was added sodium ascorbate (0.035 g, 0.18 mmol) and the reaction was stirred for 14.5 h at room temperature. The reaction was quenched by adding two drops of sat. aq. EDTA solution (pH 12). The product was extracted into DCM (2 × 5 mL) and the combined organic layers were washed with distilled water (10 mL). The organic layer was dried over $Na_2SO_4$, filtered, and concentrated *in vacuo*. Purification was achieved with a gradient of 1 – 3% MeOH/DCM to give the product as an off-white solid. Yield: 0.054 g (63%). $R_f$: 0.39 (MeOH/DCM = 1:9). HPLC purity 93%, RT = 13.94 min with method 6. $^1$H NMR (500 MHz, $CDCl_3$): δ 7.54 (s, 1H, H-11), 7.41 (m, 2H, H-18), 7.08 (t, *J* = 6.2 Hz, 1H, NH), 6.97 (t, *J* = 5.5 Hz, 1H, NH), 6.90 (m, 2H, H-19), 5.44 (s, 1H, H-15),

4.51 (t, *J* = 6.6 Hz, 2H, H-12), 4.42 (d, *J* = 5.6 Hz, 2H, H-9), 4.04 (s, 1H, H-4), 3.79 (s, 3H, H-21), 3.68 – 3.61 (m, 5H, H-1 and H-15), 3.51 (m, 2H, H-6), 2.89 (t, *J* = 6.6 Hz, 2H, H-13), 2.42 (t, *J* = 6.3 Hz, 2H, H-7), 1.04 (s, 3H, H-3), 1.01 (s, 3H, H-3). $^{13}$C NMR (126 MHz, $CDCl_3$): δ 171.2 (C-8), 170.9 (C-14), 169.4 (C-5), 160.3 (C-20), 144.6 (C-10), 130.3 (C-17), 127.6 (C-18), 123.1 (C-11), 113.8 (C-19), 101.2 (C-15), 83.9 (C-4), 78.5 (C-1), 55.4 (C-21), 52.2 (C-13), 45.6 (C-12), 35.7 (C-7), 34.9 (C-6), 34.8 (C-9), 34.3 (C-13), 33.1 (C-2), 21.9 (C-3a), 19.1 (C-3). HRMS (ESI$^+$) *m/z* $[C_{24}H_{33}N_5NaO_7]^+$ calcd. 526.2272, found 526.2278.

**Synthesis and characterization of compound DD-08-001 (2-13)**

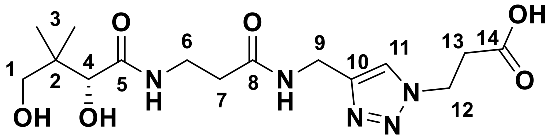

The reactant compound **DD-02-026** (0.085 g, 0.17 mmol) was dissolved into THF (2 mL) and to it was added a solution of lithium hydroxide monohydrate (0.009 g, 0.2 mmol) dissolved into distilled water (2mL). The mixture was stirred at room temperature for 19.5 h. The saponified intermediate was extracted into acidic EtOAc (2 × 10 mL, with 5 drops of AcOH). The combined organic layers were concentrated *in vacuo* and the residue was redissolved into a solution of 9:1 AcOH:$H_2O$ (8 mL), before stirring at room temperature for 4.5 h. The reaction mixture was concentrated *in vacuo* and redissolved in distilled water (10 mL). The solution was then washed with DCM (5 × 10 mL). The aqueous layer was collected and concentrated *in vacuo* to afford the product as a colourless oil. Yield: 0.043 g (63%). $R_f$: 0.00 (MeOH/DCM = 1:9). $^1$H NMR (500 MHz, MeOD): δ 7.88 (s, 1H, H-11), 4.62 (t, *J* = 6.7 Hz, 2H, H-12), 4.43 (m, 2H, H-9), 3.89 (s, 1H, H-4), 3.50 (t, *J* = 6.6 Hz, 2H, H-6), 3.41 (m, 2H, H-1), 2.84 (t, *J* = 6.7 Hz, 2H, H-13), 2.45 (t, *J* = 6.6, 2H, H-7), 0.90 (s, 3H, H-3), 0.90 (s, 3H, H-3). $^{13}$C NMR (126 MHz, MeOD): δ 176.1 (C-5), 175.8 (C-14), 173.7 (C-8), 146.0 (C-10), 124.6 (C-11), 77.2 (C-4), 70.3 (C-1), 48.0 (C-12), 40.4 (C-2), 37.1 (C-13), 36.3 (C-7), 36.3 (C-6), 35.6 (C-9), 21.3 (C-3), 20.9 (C-3). HRMS (ESI$^+$) *m/z* $[C_{15}H_{24}N_5Na_2O_6]^+$ calcd. 416.1516, found 416.1510.

*Synthesis Route for Compound 2-14*

**Synthesis and characterization of compound DD-07-081 (2-14)**

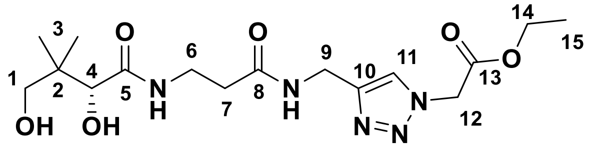

The reactant compound **DD-07-037** (0.064 g, 0.13 mmol) was dissolved into a solution of 9:1 AcOH:$H_2O$ (6 mL) and the mixture was stirred at room temperature for 4 h. The reaction mixture was concentrated *in vacuo*, and the residue was redissolved in distilled water (10 mL) before washing with DCM (5 × 10 mL). The aqueous layer was collected and concentrated *in vacuo* to afford the product as a white solid. Yield: 0.038 g (78%). $R_f$: 0.13 (MeOH/DCM = 1:9). HPLC purity 98%, RT = 4.72 min with method 5, and HPLC purity 97%, RT = 12.46 min with method 6. $^1$H NMR (500 MHz, MeOD): δ 7.92 (s, 1H, H-11), 5.30 (s, 2H, H-12), 4.46 (s, 2H, H-9), 4.25 (q, *J* = 7.1 Hz, 2H, H-14), 3.88 (s, 1H, H-4), 3.54 – 3.35 (m, 4H, H-1 and H-6), 2.46 (t, *J* = 6.6, 2H, H-7), 1.29 (t, *J* = 7.1 Hz, 3H, H-15), 0.90 (s, 6H, H-3). $^{13}$C NMR (126 MHz, MeOD): δ 176.1 (C-5), 173.8 (C-8), 168.4 (C-13), 146.3 (C-10), 125.8 (C-11), 77.3 (C-4), 70.3 (C-1), 63.2 (C-14), 51.7 (C-12), 40.4 (C-2), 36.4 (C-7), 36.3 (C-6), 35.6 (C-9), 21.3 (C-3), 20.9 (C-3), 14.4 (C-15). HRMS (ESI$^+$) *m/z* $[C_{16}H_{27}N_5NaO_6]^+$ calcd. 408.1854, found 408.1847.

*Synthesis Route for Compound 2-15*

**Synthesis and characterization of compound DD-07-015 (2-15)**

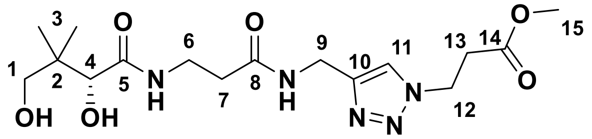

The reactant compound **DD-02-026** (0.057 g, 0.11 mmol) was dissolved into a solution of 9:1 AcOH:$H_2O$ (6 mL) and the mixture was stirred at room temperature for 3.5 h. The reaction mixture was concentrated *in vacuo*, then redissolved in distilled water (10 mL) and washed with DCM (5 × 10 mL). The aqueous layer was collected and concentrated *in vacuo* to afford the product as a colourless oil. Yield: 0.036 g (81%). $R_f$: 0.22 (MeOH/DCM = 1:9). HPLC purity 97%, RT = 6.88 min with method 2. $^1$H NMR (500 MHz,

MeOD): δ 7.88 (s, 1H, H-11), 4.65 (t, $J$ = 6.6 Hz, 2H, H-12), 4.41 (s, 2H, H-9), 3.88 (s, 1H, H-4), 3.67 (s, 3H, H-15), 3.57 – 3.34 (m, 4H, H-1 and H-6), 2.99 (t, $J$ = 6.6 Hz, 2H, H-13), 2.45 (t, $J$ = 6.6, 2H, H-7), 0.90 (s, 6H, H-3). $^{13}$C NMR (126 MHz, MeOD): δ 176.1 (C-5), 173.7 (C-8), 172.6 (C-14), 146.1 (C-10), 124.8 (C-11), 77.3 (C-4), 70.3 (C-1), 52.5 (C-15), 46.9 (C-12), 40.4 (C-2), 36.4 (C-7), 36.3 (C-6), 35.6 (C-9), 35.1 (C-13), 21.3 (C-3), 20.9 (C-3). HRMS (ESI$^+$) *m/z* $[C_{16}H_{27}N_5NaO_6]^+$ calcd. 408.1854, found 408.1854.

*Synthesis Route for Compound 6-06*

**Synthesis and characterization of compound DD-07-125**

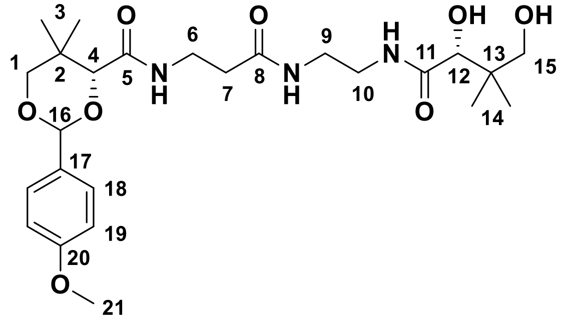

The reactants compound **DD-05-006** (0.077 g, 0.20 mmol), D-pantolactone (0.077 g, 0.059 mmol), and triethylamine (50 µL, 0.036 g, 0.36 mmol) were dissolved in ethanol (1.5 mL) and heated to 115°C for 45.5 h. The reaction mixture was concentrated *in vacuo*. Purification was achieved with a gradient of 3 – 7% MeOH/DCM to afford the product as a white powder. Yield: 0.064 g (63%). $R_f$: 0.38 (MeOH/DCM = 1:9). HPLC purity 100%, RT = 4.28 min with method 3, and HPLC purity 97%, RT = 5.64 min with method 4. $^1$H NMR (500 MHz, $CDCl_3$): δ 7.43 (m, 2H, H-18), 7.21 (bs, 1H, NH), 6.99 (t, $J$ = 6.4 Hz, 1H, NH), 6.92 (m, 2H, H-19), 6.77 (bs, 1H, NH), 5.46 (s, 1H, H-16), 4.08 (s, 1H, H-4), 3.97 (s, 1H, H-12), 3.82 (s, 3H, H-21), 3.68 (m, 2H, H-1), 3.52 (m, 2H, H-6), 3.48 (m, 2H, H-15), 3.38 (m, 4H, H-9 and H-10), 2.40 (t, $J$ = 6.4, 2H, H-7), 1.10 (s, 3H, H-3), 1.07 (s, 3H, H-3), 1.02 (s, 3H, H-14), 0.93 (s, 3H, H-14). $^{13}$C NMR (126 MHz, $CDCl_3$): δ 174.6 (C-11), 171.8 (C-8), 170.2 (C-5), 160.5 (C-20), 130.2 (C-17), 127.7 (C-18), 113.9 (C-19), 101.6 (C-16), 83.9 (C-4), 78.6 (C-1), 78.2 (C-12), 70.9 (C-15), 55.5 (C-21), 40.0 (C-9 or C-10), 39.4 (C-13), 39.3 (C-9 or C-10), 36.7 (C-7), 35.4 (C-6), 33.3 (C-2), 22.0 (C-3), 21.9 (C-

14), 20.9 (C-14), 19.3 (C-3). HRMS (ESI+) *m/z* $[C_{25}H_{39}N_3NaO_8]^+$ calcd. 532.2629, found 532.2631.

**Synthesis and characterization of compound DD-07-134 (6-06)**

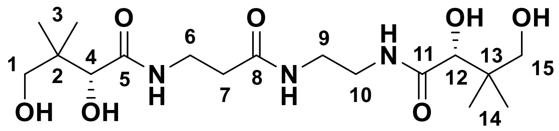

The reactant compound **DD-07-125** (0.053 g, 0.10 mmol) was dissolved into a solution of 9:1 AcOH:$H_2O$ (5 mL) and stirred at room temperature for 5.5 h. The reaction mixture was concentrated *in vacuo*. The residue was redissolved into $H_2O$ (10 mL) and washed with DCM (5 × 10 mL). The aqueous layer was collected and concentrated *in vacuo* to afford the product as a colourless oil. Yield: 0.035 g (86%). $R_f$: 0.02 (MeOH/DCM = 1:9). HPLC purity 93%, RT = 7.14 min with method 1. $^1$H NMR (MeOD, 500 MHz): δ 3.90 (s, 1H, H-4 or H-12), 3.89 (s, 1H, H-4 or H-12), 3.54 – 3.36 (m, 6H, H-1, H-6, H-15), 3.35 – 3.26 (m, 4H, H-9, H-10), 2.41 (t, *J* = 6.7, 2H, H-7), 0.93 (s, 3H, H-3 or H-14), 0.92 (s, 3H, H-3 or H-14), 0.92 (s, 6H, H-3 and H-14). $^{13}$C NMR (126 MHz, MeOD): δ 176.5 (C-5 or C-11), 176.1 (C-5 or C-11), 174.2 (C-8), 77.5 (C-4 or C-12), 77.3 (C-4 or C-12), 70.3 (C-1 or C-15), 70.3 (C-1 or C-15), 40.4 (C-2 and C-13), 40.1 (C-9 or C-10), 39.6 (C-9 or C-10), 36.6 (C-7), 36.4 (C-6), 21.5 (C-3 or C-14), 21.3 (C-3 or C-14), 21.0 (C-3 or C-14), 20.9 (C-3 or C-14). HRMS (ESI+) *m/z* $[C_{17}H_{33}N_3NaO_7]^+$ calcd. 414.2211, found 414.2217.

*Synthesis Route for Compound 6-07*

**Synthesis and characterization of compound DD-05-078**

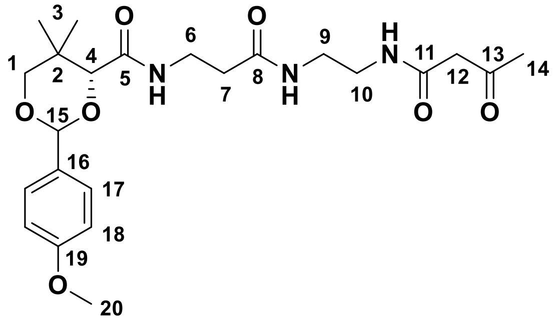

The reactant compound **DD-05-006** (0.100 g, 0.264 mmol) was dissolved in DCM (5 mL). Diketene (0.081 g, 0.963 mmol) was added and the mixture was stirred at room temperature for 4 h. The reaction mixture was concentrated *in vacuo*. Purification was achieved with a gradient of 0 – 5% MeOH/DCM to give a pure product as a red solid. Yield: 0.089 g (73%). $R_f$: 0.33 (MeOH/DCM = 1:9). $^1$H NMR (500 MHz, $CDCl_3$): δ 7.42 (m, 2H, H-17), 7.28 (bs, 1H, NH), 7.04 (t, $J$ = 6.4 Hz, NH), 6.92 (m, 2H, H-18), 6.58 (bs, 1H, NH), 5.46 (s, 1H, H-15), 4.08 (s, 1H, H-4), 3.82 (s, 3H, H-20), 3.69 (m, 2H, H-1), 3.54 (m, 2H, H-6), 3.44 – 3.33 (m, 6H, H-9 and H-10 and H-12), 2.42 (t, $J$ = 6.2 Hz, 2H, H-7), 2.23 (s, 3H, H-14), 1.10 (s, 3H, H-3), 1.08 (s, 3H, H-3). $^{13}$C NMR (126 MHz, $CDCl_3$): δ 204.4 (C-13), 171.8 (C-8), 169.8 (C-5), 166.8 (C-11), 160.4 (C-19), 130.3 (C-16), 127.7 (C-17), 113.9 (C-18), 101.5 (C-15), 84.0 (C-4), 78.6 (C-1), 55.5 (C-20), 50.1 (C-12), 39.8 (C-9 or C-10), 39.7 (C-9 or C-10), 36.5 (C-7), 35.2 (C-6), 33.2 (C-2), 31.1 (C-14), 22.0 (C3), 19.3 (C-3). HRMS (ESI$^+$) *m/z* $[C_{23}H_{33}N_3NaO_7]^+$ calcd. 486.2211, found 486.2219.

**Synthesis and characterization of compound DD-07-148 (6-07)**

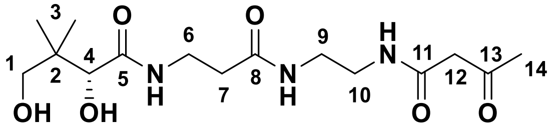

The reactant compound **DD-05-078** (0.089 g, 0.192 mmol) was dissolved into a solution of 9:1 AcOH:$H_2O$ (9 mL) and stirred at room temperature for 5.5 h. The reaction mixture was concentrated *in vacuo* then the residue was redissolved into distilled water (5 mL) and washed with DCM (4 × 5 mL). The aqueous layer was collected and concentrated *in vacuo*. Purification was achieved with a gradient of 3 – 10% MeOH/DCM to give the product as a pale, yellow oil. Yield: 0.040 g (60%). $R_f$: 0.09 (MeOH/DCM = 1:9). HPLC purity 92%, RT = 7.25 min with method 2, and HPLC purity 98%, RT = 4.50 min with method 5. $^1$H NMR (500 MHz, $d_6$-DMSO): δ 8.08 (t, $J$ = 5.5 Hz, 1H, NH), 7.90 (t, $J$ = 5.5 Hz, 1H, NH), 7.70 (t, $J$ = 5.9 Hz, 1H, NH), 5.36 (d, $J$ = 5.6 Hz, 1H, OH), 4.46 (t, $J$ = 5.6 Hz, 1H, OH), 3.69 (d, $J$ = 5.5 Hz, H-4), 3.38 – 3.14 (m, 7H, H-1 and H-6 and H-14), 3.12 – 3.05 (m, 4H, H-9 and H-10), 2.25 (m, 2H, H-7), 2.14 (s, 2H, H-12), 0.80 (s, 3H, H-3), 0.78 (s, 3H, H-3). $^{13}$C NMR (126 MHz, $d_6$-DMSO): δ 203.0 (C-13), 172.8 (C-5), 170.7 (C-8), 166.2 (C-11), 75.0 (C-4), 68.0 (C-1), 51.3 (C-12), 39.0 (C-2), 38.4 (C-10), 38.1 (C-9),

35.2 (C-7), 34.8 (C-6), 30.0 (C-14), 20.9 (C-3), 20.3 (C-3). HRMS (ESI+) *m/z* $[C_{15}H_{27}N_3NaO_6]^+$ calcd. 368.1792, found 368.1779.

*Synthesis Route for Compound 6-11*

**Synthesis and characterization of compound DD-07-115 (6-11)**

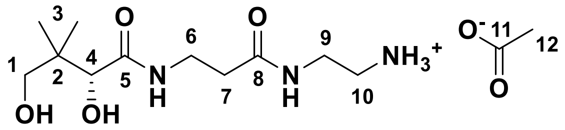

Compound **DD-05-006** (0.056 g, 0.15 mmol) was dissolved into a solution of 9:1 AcOH:$H_2O$ (5 mL) and stirred at room temperature for 24 h. The reaction mixture was concentrated *in vacuo*, then redissolved into water (10 mL) and washed with DCM (5 × 10 mL). The aqueous layer was concentrated *in vacuo* to afford the product as a yellow oil. Yield: 0.042 g (94%). $R_f$: 0.00 (MeOH/DCM = 1:9). $^1$H NMR (500 MHz, MeOD): δ 3.90 (s, 1H, H-4), 3.50 (t, *J* = 6.4 Hz, 2H, H-6) 3.49 – 3.36 (m, 4H, H-1 and H-9), 3.04 (t, *J* = 5.8 Hz, 2H, H-10), 2.44 (m, 2H, H-7), 1.92 (s, 3H, H-12), 0.92 (s, 3H, H-3), 0.92 (s, 3H, H-3). $^{13}$C NMR (126 MHz, MeOD): δ 179.2 (C-11), 176.3 (C-5), 175.3 (C-8), 77.3 (C-4), 70.3 (C-1), 40.9 (C-10), 40.4 (C-2), 38.3 (C-9), 36.9 (C-7), 36.5 (C-6), 23.5 (C-12), 21.4 (C-3), 20.9 (C-3). HRMS (ESI+) *m/z* $[C_{11}H_{23}N_3NaO_4]^+$ calcd. 284.1581, found 284.1575.

*Synthesis Route for Compound 6-12*

**Synthesis and characterization of compound DD-06-106**

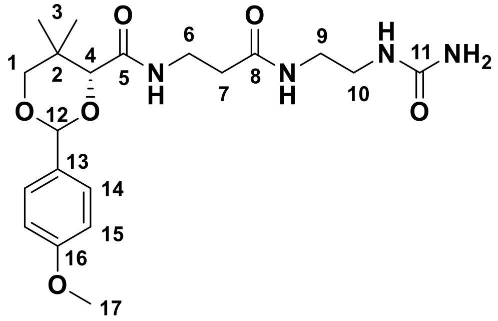

The reactant compound **DD-05-006** (0.097 g, 0.26 mmol) was dissolved into DCM (5 mL). To this solution was added ethyl isocyanatotrimethylsilane (50 µL, 0.042 g, 0.37 mmol) and the mixture was stirred at room temperature for 50 min. The reaction mixture was concentrated *in vacuo.* Purification was achieved with a gradient of 4 – 9% MeOH/DCM to give the product as a white powder. Yield: 0.080 g (74%). $R_f$: 0.14 (MeOH/DCM = 1:9).

HPLC purity 98%, RT = 6.14 min with method 1. $^{1}H$ NMR (500 MHz, $CDCl_3$): δ 7.43 (m, 2H, H-14), 7.11 (t, *J* = 5.3 Hz, 1H, NH), 7.06 (t, *J* = 6.3 Hz, 1H, NH), 6.91 (m, 2H, H-15), 5.75 (t, *J* = 5.6 Hz, 1H, NH), 5.45 (s, 1H, H-12), 4.91 (bs, 2H, $NH_2$), 4.07 (s, 1H, H-4), 3.80 (s, 3H, H-17), 3.68 (m, 2H, H-1), 3.51 (m, 2H, H-6), 3.27 (m, 2H, H-9), 3.21 (m, 2H, H-10), 2.39 (t, *J* = 6.5 Hz, 2H, H-7), 1.09 (s, 3H, H-3), 1.05 (s, 6H, H-3). $^{13}C$ NMR (126 MHz, $CDCl_3$): δ 172.0 (C-8), 169.8 (C-5), 160.4 (C-16), 160.0 (C-11), 130.2 (C-13), 127.7 (C-14), 113.9 (C-15), 101.5 (C-12), 84.0 (C-4), 78.5 (C-1), 55.5 (C-17), 40.5 (C-9), 44.0 (C-10), 36.2 (C-7), 35.4 (C-6), 33.2 (C-2), 22.0 (C-3), 19.3 (C-3). HRMS ($ESI^+$) *m/z* $[C_{20}H_{30}N_4NaO_6]^+$ calcd. 445.2058, found 445.2056.

**Synthesis and characterization of compound DD-06-126 (6-12)**

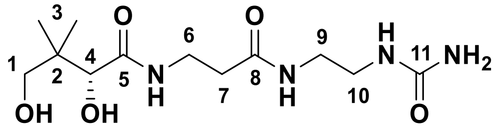

The reactant compound **DD-06-106** (0.047 g, 0.11 mmol) was dissolved into a solution of 9:1 AcOH:$H_2O$ (5 mL) and stirred at room temperature for 3 h. The reaction mixture was concentrated *in vacuo*. The residue was redissolved into distilled water (10 mL) and washed with DCM (5 × 10 mL). The aqueous layer was collected and concentrated *in vacuo* to afford the product as a colourless oil. Yield: 0.030 g (88%). $R_f$: 0.04 (MeOH/DCM = 1:9). HPLC purity 94%, RT = 14.37 min with method 6. $^{1}H$ NMR (500 MHz, MeOD): δ 3.92 (s, 1H, H-4), 3.52 – 3.40 (m, 4H, H-1 and H-6), 3.28 – 3.21 (m, 4H, H-9 and H-10), 2.44 (t, *J* = 6.6, 2H, H-7), 0.94 (s, 6H, H-3). $^{13}C$ NMR (126 MHz, MeOD): δ 176.1 (C-5), 174.2 (C-8), 162.3 (C-11), 77.3 (C-4), 70.3 (C-1), 40.9 (C-9 or C-10), 40.4 (C-2), 40.3 (C-9 or C-10), 36.6 (C-7), 36.4 (C-6), 21.4 (C-3), 20.9 (C-3). HRMS ($ESI^+$) *m/z* $[C_{12}H_{24}N_4NaO_5]^+$ calcd. 383.1360, found 383.1360.

Synthesis of **4f**

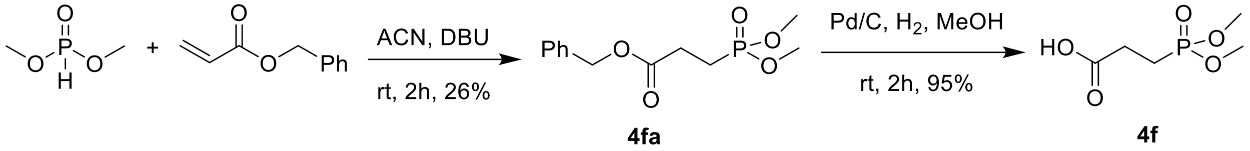

To a magnetically stirred solution of dimethyl phosphite (0.42 mL, 4.55 mmol) and benzyl acrylate (1 mL, 6.82 mmol) in acetonitrile (15 mL) in a 50 mL round bottom flask was added DBU (0.4 mL, 2.27 mmol) and the reaction mixture was stirred at rt for 2 h. Solvents were evaporated *in vacuo*. The residue was purified by column chromatography ($SiO_2$, Hex/EA = 40:60), to afford **4fa** (330 mg, 26%) as a clear oil. $R_f$ = 0.40 (EA); Purity 90%, RT = 14.96 min with method 8; $^1H$ NMR ($CDCl_3$, 500 MHz) $\delta$ 7.43 – 7.29 (m, 5H), 5.14 (s, 2H), 3.72 (d, $J$ = 10.8 Hz, 3H), 3.72 (d, $J$ = 10.8 Hz, 3H), 2.65 (ddd, $J$ = 12.5, 8.7, 7.1 Hz, 2H), 2.18 – 2.03 (m, 2H); $^{13}C$ NMR (126 MHz, $CDCl_3$) $\delta$ 171.8 (d, $J$ = 17.9 Hz), 135.6, 128.6, 128.4, 128.3, 66.8, 52.5 (d, $J$ = 6.6 Hz), 27.5 (d, $J$ = 3.8 Hz), 20.0 (d, $J$ = 144.7 Hz); HRMS (ESI$^+$) *m/z* $[C_{12}H_{17}O_5PNa]^+$ calcd: 295.0711; found: 295.0700.

Intermediate **4fa** (300 mg, 1.10 mmol) was dissolved in MeOH (12 mL) in a 50 mL round bottom flask, Pd/C (30 mg) was added to it, and hydrogen gas was purged in the reaction mixture for 2 h. The reaction mixture was filtered through a celite pad, and the filtrate was concentrated *in vacuo* to obtain the product **4f** (190 mg, 95%) as a clear oil. $R_f$ = 0.40 (DCM/MeOH = 90:10); Purity 90%, RT = 14.56 min with method 7; RT = 20.30 min with method 8; $^1H$ NMR ($CD_3OD$, 500 MHz) $\delta$ 3.76 (s, 3H), 3.74 (s, 3H), 2.60 – 2.50 (m, 2H), 2.17 – 2.06 (m, 2H); $^{13}C$ NMR ($CDCl_3$, 126 MHz) $\delta$ 174.8 (dd, $J$ = 17.4, 6.9 Hz), 52.8 (d, $J$ = 6.5 Hz), 27.1 (d, $J$ = 4.1 Hz), 20.3 (s), 19.2 (s); $^{31}P$ NMR ($CDCl_3$, 203 MHz) $\delta$ 33.49; HRMS (ESI$^+$) *m/z* $[C_5H_{10}O_5PNa_2]^+$ calcd: 227.0067; found: 227.0050.

Synthesis of **4g**

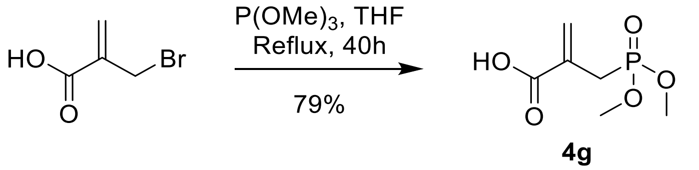

Compound **4g** was synthesized by a reported procedure.[9] In a 25 mL round bottom flask, 2-(bromomethyl)acrylic acid (500 mg, 3.03 mmol) was dissolved in THF (6 mL) and trimethyl phosphite (0.38 mL, 3.18 mmol) was added. The reaction mixture was refluxed for 40 h. The solvent was evaporated *in vacuo*, and the residue was purified by column chromatography ($SiO_2$, DCM/MeOH = 97:3 to 96:4) to afford the product (465 mg, 79%) as a clear oil. $R_f$ = 0.50 (DCM/MeOH = 90:10, visualized by $KMnO_4$ stain). $^1$H NMR ($CDCl_3$, 500 MHz) $\delta$ 6.47 (d, $J$ = 5.7 Hz, 1H), 5.90 (d, $J$ = 5.6 Hz, 1H), 3.77 (d, $J$ = 11.0 Hz, 6H), 2.99 (d, $J$ = 22.3 Hz, 2H); $^{31}$P NMR ($CDCl_3$, 203 MHz) $\delta$ 28.74.

Synthesis of **4h**

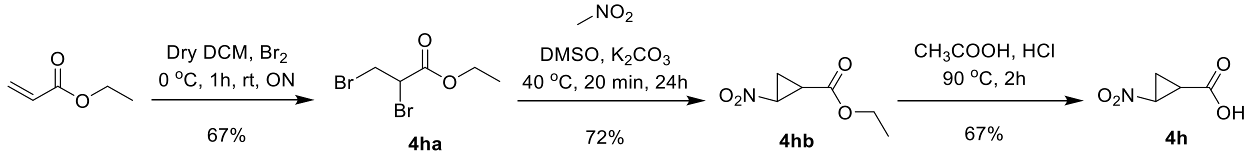

Compound **4ha** was synthesized and characterized following a literature protocol.[10] In a 50 mL round bottom flask, to a solution of ethyl acrylate (1 mL, 10 mmol) in dry DCM (6 mL), bromine (0.52 mL, 20 mmol) was added dropwise over 15 min at 0°C under an inert atmosphere. The reaction mixture was allowed to stir at 0°C for 1 h and then at rt overnight. The reaction mixture was quenched with a sat. aq. $Na_2S_2O_3$ solution (20 ml). The mixture was filtered, followed by filtrate extraction with DCM (3 × 20 mL). The combined organic layers were washed with water (2 × 25 mL) and brine (1 × 20 mL), dried over anhydrous $Na_2SO_4$ and concentrated *in vacuo* to yield compound **4ha** (1.98 g, 67%) as a clear oil, which was directly used in the next step. $^1$H NMR ($CDCl_3$, 300 MHz) $\delta$ 4.42 (dd, $J$ = 11.4, 4.4 Hz, 1H), 4.30 (q, $J$ = 7.1 Hz, 2H), 3.92 (m, 1H), 3.67 (m, 1H), 1.33 (t, $J$ = 7.1 Hz, 3H).

Compound **4hb** was synthesized and characterized following a modified literature protocol.[11–13] In a 25 mL round bottom flask, nitromethane (127 mg, 2.08 mmol) was dissolved in dry DMSO (2 mL), and anhydrous $K_2CO_3$ (796 mg, 5.77 mmol) was added. The mixture was stirred at 40°C for 20 min under an inert atmosphere. Compound **4ha** (500 mg, 1.92 mmol) pre-dissolved in dry DMSO (5 mL) was added dropwise to the mixture through a syringe over 6 h before stirring for 18 h at 40°C. The reaction mixture was then cooled to rt and poured into ice-cold water (15 mL).

The product was extracted in $Et_2O$ (3 × 20 mL). The combined organic layers were washed with water (2 × 20 mL) and brine (20 mL), dried over anhydrous $Na_2SO_4$ and concentrated *in vacuo* to yield the compound **4hb** (220 mg, 72%) as a yellowish oil. $R_f$ = 0.40 (Hex/EA 80:20); $^1$H NMR ($CDCl_3$, 300 MHz) $\delta$ 4.59 (ddd, $J$ = 7.5, 4.4, 2.7 Hz, 1H), 4.20 (qd, $J$ = 7.1, 0.7 Hz, 2H), 2.73 (ddd, $J$ = 10.1, 7.1, 2.7 Hz, 1H), 2.06 (ddd, $J$ = 10.2, 5.8, 4.4 Hz, 1H), 1.73 (td, $J$ = 7.3, 5.8 Hz, 1H), 1.29 (t, $J$ = 7.1 Hz, 3H).

Compound **4h** was synthesized and characterized following a literature protocol.[Error! Bookmark not defined.] In a 25 mL round bottom flask, a suspension of **4hb** (190 mg, 1.19 mmol) in a 1:1 mixture of AcOH and 12N aq. HCl (2 ml) was stirred at 90°C for 2 h. The solvent was evaporated *in vacuo*, and the product was purified by column chromatography ($SiO_2$, DCM/MeOH = 96:4) to afford compound **4h** (105 mg, 67%) as a white solid. $R_f$ = 0.30 (DCM/MeOH = 90:10, visualized by iodine stain); $^1$H NMR ($CDCl_3$, 300 MHz) $\delta$ 9.11 (bs, 1H), 4.65 (m, 1H), 2.76 (m, 1H), 2.15 (m, 1H), 1.79 (m, 1H).

Synthesis of **4i**

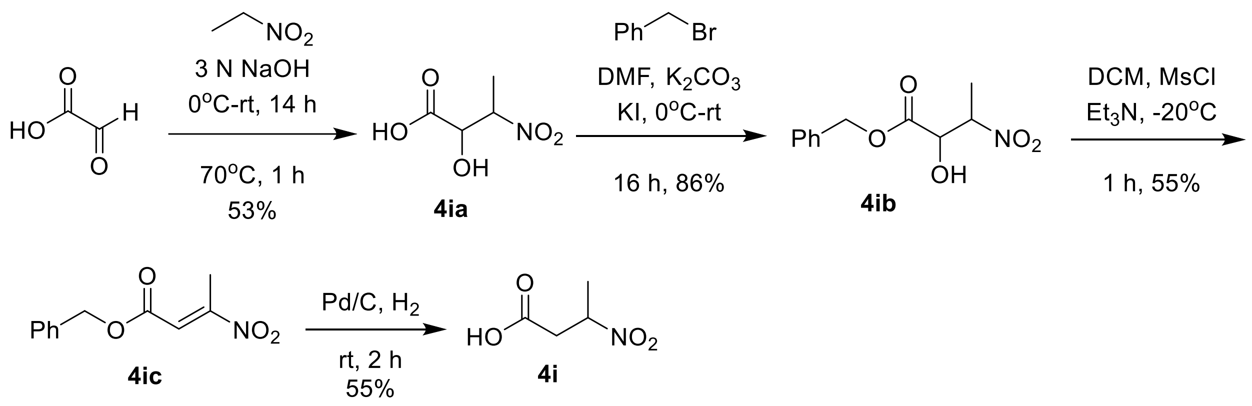

Compound **4ia** was synthesized by a reported procedure.[14] In a 50 mL round bottom flask, glyoxalic acid monohydrate (2 g, 21.73 mmol) was dissolved in 3 N aq. NaOH (913 mg, 22.81 mmol in 7.5 mL $H_2O$) and cooled to 0°C. Nitroethane (1.5 mL, 23.90 mmol) was next added to the mixture, before stirrinf at rt for 14 h, and then at 70°C for 1 h. The reaction mixture was cooled to 0°C and 6 N $H_2SO_4$ was added (pH 2), before extraction of the product in EA (8 × 50 mL). The combined organic layers were dried over anhydrous $Na_2SO_4$ and concentrated *in vacuo* to yield the crude product as a viscous yellow oil. To remove excess nitroethane, the crude product was dissolved in DCM (20 mL) before addition of hexane (30 mL). The liquid was

removed by decantation, and the product was dried *in vacuo* to yield **4ia** (1.7 g, 53%) as a yellow oil. $R_f$ = 0.20 (DCM/MeOH = 90:10). This crude product was used directly in the next step.
Compound **4ib** was synthesized using reported procedures.[2,15] In a 25 ml round bottom flask, to a solution of **4ia** (1 g, 6.70 mmol) in DMF (8 mL) at 0°C was added $K_2CO_3$ (926 mg, 6.7 mmol) in portions, before KI (56 mg, 0.33 mmol) and benzyl bromide (0.8 mL, 6.7 mmol). The resulting solution was stirred at rt for 14 h. The reaction mixture was next poured into an $NH_4Cl$ solution (1.5 M, 25 mL) and the product was extracted in EA (4 × 30 mL). The combined organic phase was washed with water (2 × 30 mL), dried over $Na_2SO_4$, and concentrated *in vacuo* to afford **4ib** (807 mg, 86%) as a brown oil, which was used as a crude mixture of diastereoisomers. $R_f$ = 0.70 (DCM/MeOH = 90:10, visualized by $KMnO_4$ stain).
Compound **4ic** was synthesized using a reported procedure.**Error! Bookmark not defined.** In a 50 mL round bottom flask, a solution of **4ib** (800 mg, 3.34 mmol) in dry DCM (10 mL) was added MsCl (0.78 mL, 10.03 mmol) and the reaction mixture was cooled to -20°C. $Et_3N$ (1.4 mL, 10.03 mmol) was added dropwise and the mixture was stirred at -20°C for 1 h. The reaction mixture was poured in water (20 mL) and the product was extracted in EA (3 × 30 mL). The combined organic phase was washed with sat. aq. $CuSO_4$ (30 mL), sat. aq. $NaHCO_3$ (30 mL), and brine (30 mL), successively. The organic layer was dried over $MgSO_4$ and concentrated *in vacuo,* and the residue was purified by column chromatography ($SiO_2$, Hex/EA = 97:3 to 96:4) to afford **4ic** (404 mg, 55%) as a yellow oil. $R_f$ = 0.70 (Hex/EA = 85:15); $^1H$ NMR ($CDCl_3$, 300 MHz) $\delta$ 7.43 – 7.36 (m, 5H), 7.11 (q, $J$ = 1.2 Hz, 1H), 5.23 (s, 2H), 2.60 (d, $J$ = 1.1 Hz, 3H); $^{13}C$ NMR ($CDCl_3$, 75 MHz) $\delta$ 164.1, 160.4, 134.9, 128.9, 128.6, 121.2, 67.6, 14.2.
Compound **4ic** (150 mg 0.68 mmol) was dissolved in MeOH (15 mL) in a 50 mL round bottom flask. Pd/C (30 mg) was added and hydrogen gas was purged in the mixture for 2 h. The reaction mixture was filtered through a celite pad and concentrated *in vacuo*. The residue was purified by column chromatography ($SiO_2$, DCM/MeOH = 94:6 to 92:8) to yield **4i** (50 mg, 55%) as an off-white solid. $R_f$ = 0.50 (DCM/MeOH = 90:10); Purity 96%, RT = 14.36 min with method 7; RT = 18.20 min with method 8; $^1H$ NMR ($CDCl_3$, 300 MHz) $\delta$ 10.12 (bs, 1H), 4.92 (dqd, $J$ = 8.9, 6.9, 4.7 Hz, 1H), 3.22 (dd, $J$ = 17.7, 8.9 Hz, 1H), 2.76 (dd, $J$ = 17.7, 4.6 Hz, 1H), 1.64 (d, $J$ = 6.9 Hz, 3H); $^{13}C$ NMR ($CDCl_3$, 126 MHz) $\delta$ 175.2, 78.2, 38.1, 19.3; HRMS (ESI$^-$) $m/z$ $[C_4H_6NO_4]^-$ calcd: 132.0302; found: 132.0301.

Synthesis of **4k**

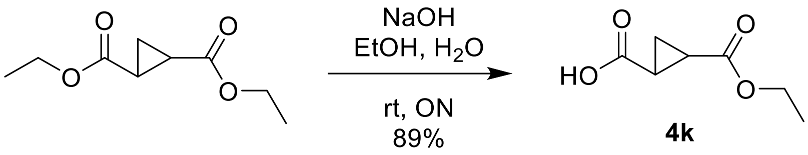

Compound **4k** was synthesized and characterized following a literature protocol.[16] In a 50 mL round bottom flask, diethyl *trans*-1,2-cyclopropanedicarboxylate (500 mg, 2.69 mmol) and NaOH (110 mg, 2.74 mmol) were dissolved in EtOH:$H_2O$ (12:1, 10 mL) and stirred at rt overnight. The mixture was concentrated *in vacuo* and the residue was dissolved in $H_2O$ (15 mL). The aqueous layer was washed with $Et_2O$ (2 × 15 mL) to remove unreacted starting material and non-polar impurities. The aqueous solution was then adjusted to pH 2 with 2 N HCl and the product was extracted in EA (2 × 15 mL). The combined organic extracts were dried over anhydrous $Na_2SO_4$ and concentrated *in vacuo* to yield **4k** (394 mg, 89%) as a clear oil. $^1$H NMR ($CDCl_3$, 300 MHz) $\delta$ 4.16 (q, $J$ = 7.2 Hz, 2H), 2.31 – 2.07 (m, 2H), 1.59 – 1.41 (m, 2H), 1.27 (t, $J$ = 7.1 Hz, 3H).

Synthesis of **4l**

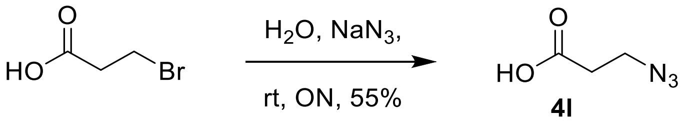

A modified reported procedure was used for the synthesis of compound **4l**.[17] 3-Bromopropionic acid (1 g, 6.54 mmol) was dissolved in $H_2O$ (10 mL) in a 50 mL round bottom flask, and $NaN_3$ (850 mg, 13.07) was added before stirring overnight at rt. The mixture was concentrated *in vacuo*, and the product was purified by column chromatography ($SiO_2$, DCM/$H_2O$ = 96:4) to afford **4l** (410 mg, 55%) as a clear oil. $R_f$ = 0.60 (DCM/MeOH = 90:10); $^1$H NMR ($CDCl_3$, 300 MHz) $\delta$ 3.59 (t, $J$ = 6.4 Hz, 2H), 2.63 (t, $J$ = 6.4 Hz, 2H).

Synthesis of **4m**

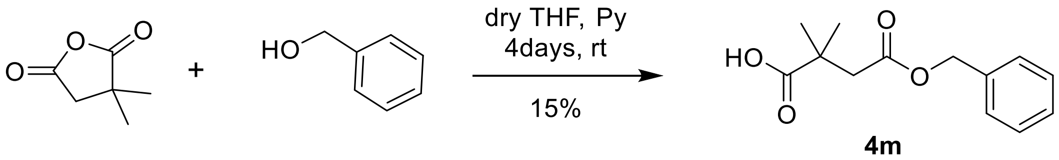

Compound **4m** was synthesized by a reported procedure.[18] 2,2-Dimethyl succinic anhydride (1.5 g, 11.71 mmol) and benzyl alcohol (1.34 mL, 12.88 mmol) were dissolved in dry THF (1.5 mL) in a 50 mL round bottom flask. Pyridine (0.5 mL, 6.21 mmol) was added and the mixture was stirred for 4 days at rt. The reaction mixture was diluted in sat. aq. $NaHCO_3$ (15 mL). The aqueous phase was separated and acidified with 2 N HCl to a pH around 4. The mixture of esters was extracted in diethyl ether (30 mL). The organic layer was dried over $Na_2SO_4$ and concentrated *in vacuo* to obtain the isomeric esters. The desired significant ester **4m** (441 mg, 15%) was crystallized from hexane at -20°C to produce a white solid. $R_f$ = 0.40 (Hex/EA = 70:30). $^1$H NMR ($CDCl_3$, 500 MHz) $\delta$ 7.37 – 7.30 (m, 5H), 5.12 (s, 2H), 2.67 (s, 2H), 1.30 (s, 6H); $^{13}$C NMR ($CDCl_3$, 126 MHz) $\delta$ 183.5, 171.0, 135.7, 128.6, 128.2, 66.4, 44.0, 40.6, 25.2.

Synthesis of **4n**

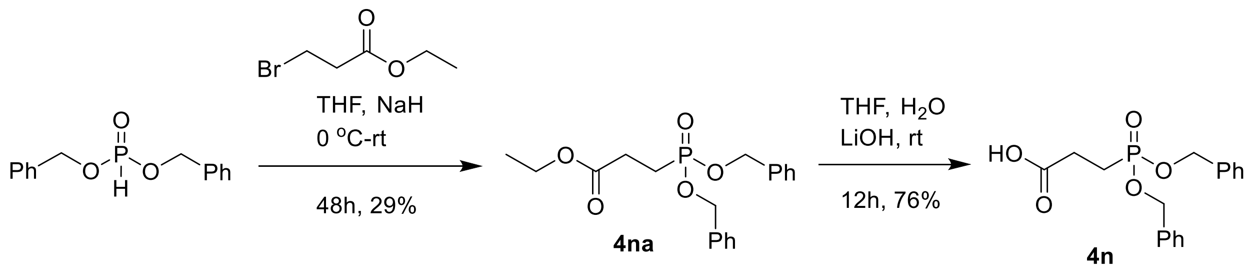

Compounds **4n** was synthesized by a reported procedure.[19] In a 50 mL round bottom flask, to a solution of dibenzylphosphite (0.92 mL, 4.14 mmol) at 0°C in THF (10 mL) was added NaH (166 mg, 4.14 mmol). After 15 min, ethyl-3-bromopropionate was added at rt, and the mixture was stirred at rt for 48 h. The reaction was quenched with sat. aq. $NH_4Cl$ (15 mL) and the product was extracted in diethyl ether (3 × 20 mL). The combined organic layers were dried over $Na_2SO_4$, filtered and evaporated to dryness, and the residue was purified by column chromatography ($SiO_2$, Hex/EA = 50:50 to 35:65) to afford **4na** (298 mg, 29%) as a yellow oil. $R_f$ = 0.50 (Hex/EA = 30:70); $^1$H NMR ($CDCl_3$, 300 MHz) $\delta$ 7.42 – 7.27 (m, 10H), 5.16 – 4.88 (m, 4H), 4.08 (q, $J$ = 7.1 Hz, 2H), 2.63 – 2.46 (m, 2H), 2.19 – 1.99 (m, 2H), 1.21 (t, $J$ = 7.1 Hz, 3H).

In a 25 mL round bottom flask, to a solution of **4na** (290 mg, 0.80 mmol) in THF:$H_2O$ (1:1, 5 mL) was added LiOH.$H_2O$ (51 mg, 1.2 mmol) and the reaction mixture was stirred at rt for 12 h. $H_2O$ (10 mL) was added and the product was washed with EA (2 × 20 mL) to remove unreacted starting material and non-polar impurities. The aqueous layer was acidified to pH 2 with 4 M HCl, and the product was extracted in EA (3 × 20 mL). The combined organic layers were dried over $Na_2SO_4$, filtered and evaporated to dryness to afford **4n** (203 mg, 76%) as a clear oil. $R_f$ = 0.10 (Hex/EA = 30:70); $^1$H NMR ($CDCl_3$, 500 MHz) $\delta$ 7.50 – 7.24 (m, 10H), 5.06-4.94 (m, 4H), 2.67 – 2.53 (m, 2H), 2.14-2.07 (m, 2H).

<u>Synthesis of **4p**</u>

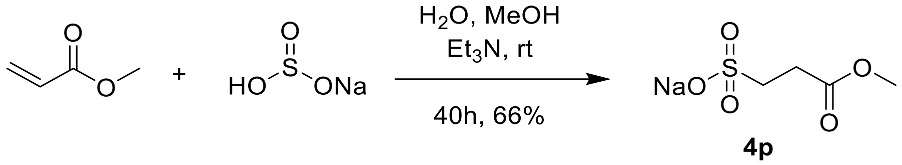

Compound **4p** was synthesized by a reported procedure.[20] In a 250 mL round bottom flask, methyl acrylate (616 mg, 7 mmol), MeOH (70 mL), $Et_3N$ (0.4 mL, 2.8 mmol) and aq. sodium bisulfite (874 mg, 8.4 mmol, in 17 mL $H_2O$) were added sequentially and the mixture was stirred vigorously at rt for 40 h. The reaction mixture was filtered through a celite pad, and the filtrate was concentrated *in vacuo*. The resultant solid was washed with EtOH (20 mL) to obtain **4p** (878 mg, 66%) as a white solid. $^1$H NMR ($D_2O$, 500 MHz) $\delta$ 3.74 (s, 3H), 3.23 (t, $J$ = 7.3 Hz, 2H), 2.83 (t, $J$ = 7.3 Hz, 2H).

<u>Synthesis of **2**</u>

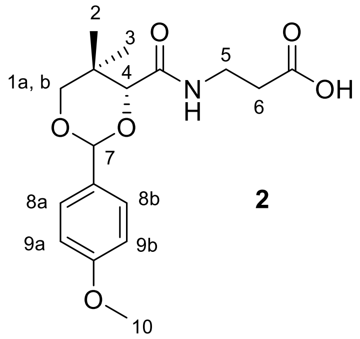

**2**

Known compound **2** was synthesized and characterized following a modified literature protocol.[21] D-Pantothenate hemicalcium salt (5 g, 20.98 mmol) was dissolved in the MQ water (30 mL) and loaded onto a Amberlite® IR-120 ($H^+$) column (2.5 × 10 cm) further eluted with

MQ water until pH of the eluent reached neutral. Water was evaporated *in vacuo* to obtain compound **1** (D-pantothenic acid) as a viscous yellowish oil which was dissolved in dry DMF (50 mL). CSA (244 mg, 1.05 mmol) was added, followed by *p*-methoxybenzaldehyde dimethyl acetal (3.93 mL, 23.08 mmol), and the mixture was stirred at rt overnight. The next day, the reaction mixture was diluted in EA (300 mL) and washed with water (3 × 150 mL). EA layer was then dried over $Na_2SO_4$, filtered and evaporated *in vacuo*. The solid obtained was washed with chilled $Et_2O$ (30 mL) to afford the desired compound **2** (4.74 g, 67%) as a white solid. $R_f$ = 0.30 (EA). $^1$H NMR ($CD_3OD$, 500 MHz) $\delta$ 7.64 (m, 1H, -NH), 7.45 (d, *J* = 8.8 Hz, 2H, H-8a and H-8b), 6.92 (d, *J* = 8.8 Hz, 2H, H-9a and H-9b), 5.53 (s, 1H, H-7), 4.15 (s, 1H, H-4), 3.80 (s, 3H, H-10), 3.73 (d, *J* = 11.3 Hz, 1H, H-1a or H-1b), 3.67 (d, *J* = 11.3 Hz, 1H, H-1a or H-1b), 3.47 (m, 2H, H-5), 2.52 (t, *J* = 6.6 Hz, 2H, H-6), 1.10 (s, 3H, H-2 or H-3), 1.04 (s, 3H, H-2 or H-3).

<u>Synthesis of **19a**</u>

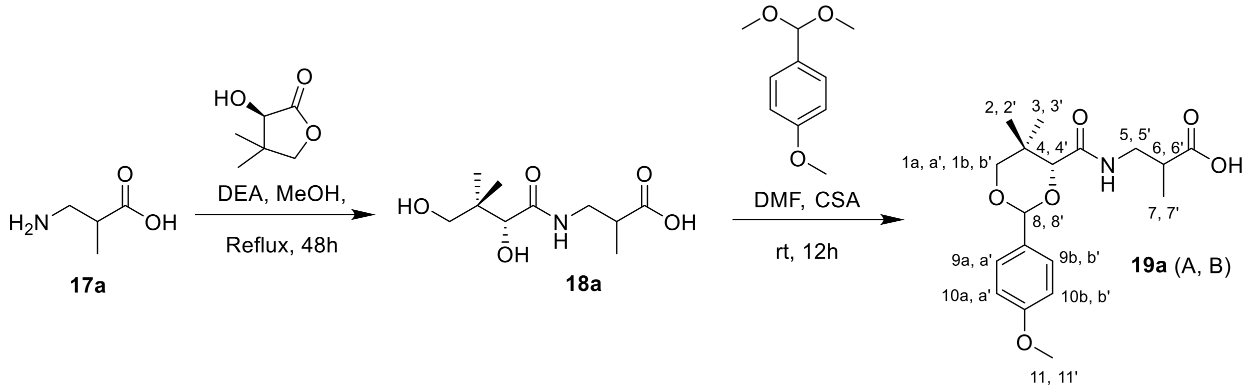

Compound **19a** was synthesized from **17a** following the general protocol A. Compound **17a** (500 mg, 4.13 mmol), diethyl amine (1.3 mL, 12.38 mmol), MeOH (8 mL) and D-pantolactone (537 mg, 4.13 mmol) were used. The crude intermediate **18a** (862 mg, quantitative) was obtained as a viscous, yellowish oil for this pair of diastereomers. $R_f$ = 0.50 (DCM/MeOH = 85:15, visualized by $KMnO_4$ stain).

**18a** (862 mg, 3.69 mmol) was dissolved in dry DMF (5 mL). CSA (43 mg, 0.18 mmol) was added, followed by *p*-methoxybenzaldehyde dimethyl acetal (672 mg, 3.69 mmol), and the mixture was stirred at rt overnight. The reaction mixture was diluted in EA (40 mL) and washed with water (3 × 30 mL). After solvent evaporation, the solid obtained was washed with chilled $Et_2O$ (8 mL) to afford **19a** (1.30 g, 67%) as a crystalline white solid for this pair of diastereomers. $R_f$ = 0.30 (EA); Purity >99%, RT = 15.30 min with method 7; RT = 23.78 min with method 8; $^1$H NMR ($CDCl_3$, 300 MHz) $\delta$ 7.47 – 7.34 (m, 4H, H-9a, H-9b, H-9a' and H-9b'), 7.03 (bs, 2H, -NH and –NH'), 6.96 – 6.84 (m, 4H, H-10a, H-10b, H-10a' and H-10b'), 5.46 (s, 1H, H-8 or H-8'), 5.44 (s, 1H, H-8 or H-8'), 4.10 (s, 1H, H-4 or H-4'), 4.09 (s, 1H, H-4 or H-4'), 3.86 – 3.75 (m, 6H, H-11 and H-11'), 3.71 (d, $J$ = 11.4 Hz, 2H, H-1a and H-1a'), 3.65 (d, $J$ = 11.5 Hz, 2H, H-1b and H-1b'), 3.56 (ddd, $J$ = 13.7, 7.1, 4.8 Hz, 1H, H-5a, H-5b, H-5a', or H-5b'), 3.48 – 3.39 (m, 2H, H-5a, H-5b, H-5a', or H-5b'), 3.30 (ddd, $J$ = 13.7, 8.2, 5.6 Hz, 1H, H-5a, H-5b, H-5a', or H-5b'), 2.86 – 2.64 (m, 2H, H-6 and H-6'), 1.24 (d, $J$ = 4.2 Hz, 3H, H-7 or H-7'), 1.21 (d, $J$ = 4.1 Hz, 3H, H-7 or H-7'), 1.15 – 1.04 (m, 12H, H-2, H-3, H-2', and H-3'); $^{13}$C NMR ($CDCl_3$, 126 MHz) $\delta$ 180.1, 180.1, 169.6, 169.6, 160.2, 160.2, 130.2, 127.4, 127.4, 113.8, 113.7, 101.2, 101.1, 83.7, 83.7, 78.5, 78.5, 55.3, 55.3, 40.9, 40.8, 39.5, 39.3, 33.1, 33.1, 21.8, 21.8, 19.2, 19.1, 14.8, 14.8; HRMS (ESI$^+$) $m/z$ $[C_{18}H_{25}NO_6Na]^+$ calcd: 374.1580; found: 374.1562.

Synthesis of **19b**

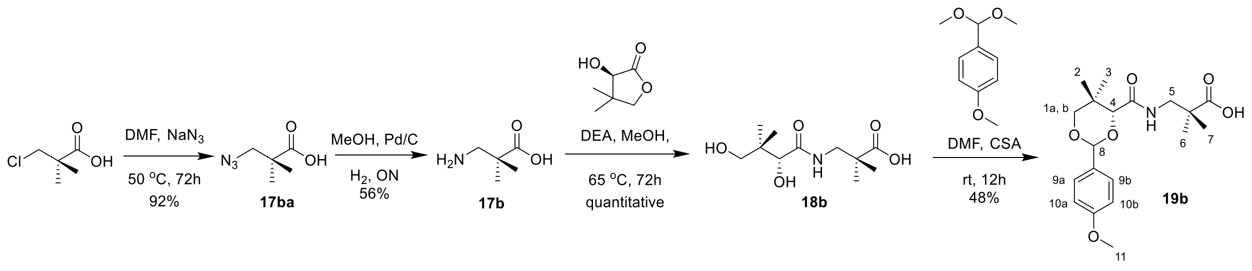

Compound **17ba** was synthesized by a reported procedure.[22] 2,2-Dimethyl-3-chloropropionic acid (1.0 g, 7.32 mmol) was dissolved in dry DMF (10 mL), and $NaN_3$ (2.38 g, 36.61 mmol) was added to it. The reaction mixture was stirred at 50°C under a nitrogen atmosphere for 72 h. The reaction mixture was cooled to rt, diluted in DCM, and filtered over a celite pad to remove unreacted $NaN_3$. The solvent was removed *in vacuo* to yield **17ba** (959 mg, 92%) as a dark yellow oil. $^1$H NMR ($CDCl_3$, 300 MHz) $\delta$ 3.43 (s, 2H), 1.25 (s, 6H).

Compound **17b** was synthesized by a reported procedure.[23] Compound **17ba** (940 mg, 6.57 mmol) was dissolved in MeOH (15 mL), Pd/C (94 mg) was added to it, and the reaction mixture was stirred under a hydrogen atmosphere overnight. The reaction mixture was filtered through a celite pad, and the filtrate was concentrated *in vacuo* to obtain **17b** (434 mg, 56%) as a white solid. $^1$H NMR ($CD_3OD$, 500 MHz) $\delta$ 2.88 (s, 2H), 1.19 (s, 6H).

Compound **19b** was synthesized following the general procedure A from **17b** over two steps. **17b** (420 mg, 3.59 mmol), and diethyl amine (1.2 mL, 10.76 mmol) were dissolved in MeOH (5 mL) and warmed. D-Pantolactone (467 mg, 3.59 mmol) was added to the reaction mixture and stirred at 65°C for 72 h. Crude intermediate **18b** (887 mg, quantitative) was obtained as a clear oil. $R_f$ = 0.50 (DCM/MeOH = 85:15, visualized by $KMnO_4$ stain).

Compound **18b** (887 mg, 3.57 mmol) was dissolved in dry DMF (10 mL). CSA (42 mg, 0.18 mmol) was added, followed by *p*-methoxybenzaldehyde dimethyl acetal (654 mg, 3.59 mmol), and the mixture was stirred at rt overnight. The reaction mixture was diluted in EA (40 mL) and washed with water (3 × 30 mL). After solvent evaporation, the solid obtained was washed with chilled $Et_2O$ (10 mL) and purified by column chromatography ($SiO_2$, Hex/EA = 50:50) to afford **19b** (626 mg, 48%) as a white solid. $R_f$ = 0.30 (Hex/EA = 50:50). Purity 99%, RT = 16.02 min with method 7; RT = 27.65 min with method 8; $^1$H NMR ($CDCl_3$, 500 MHz) $\delta$ 7.40 (m, 2H, H-9a, H-9b), 7.15 (t, $J$ = 6.5 Hz, 1H, -NH), 6.90 (m, 2H, H-10a, H-10b), 5.46 (s, 1H, H-8), 4.12 (s, 1H, H-4), 3.79 (s, 3H, H-11), 3.68 (m, 2H, H-1a, H-1b), 3.40 (ddd, $J$ = 53.5, 13.7, 6.5 Hz, 2H, H-5), 1.25 (s, 3H, H-6 or H-7), 1.23 (s, 3H, H-6 or H-7), 1.13 (s, 3H, H-2 or H-3), 1.10 (s, 3H, H-2 or H-3); $^{13}$C NMR ($CDCl_3$, 126 MHz) $\delta$ 182.7, 169.6, 160.1, 130.2, 127.3, 113.7, 101.1, 83.7, 78.5, 55.2, 45.8, 43.1, 33.1, 23.2, 23.2, 21.8, 19.3; HRMS (ESI$^+$) *m/z* $[C_{19}H_{27}NO_6Na]^+$ calcd: 388.1736; found: 388.1727.

Synthesis of **19c**

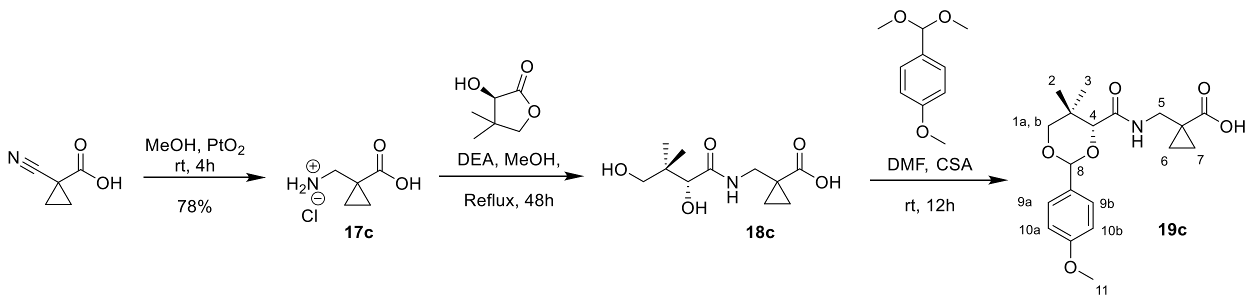

Compound **17c** was synthesized by a reported procedure.[24] 1-Cyanocyclopropane-1-carboxylic acid (600 mg, 5.40 mmol) was dissolved in MeOH (18 mL), before adding $PtO_2$ (120 mg) and conc. HCl (0.9 mL) dropwise. The reaction mixture was stirred at rt under an argon atmosphere for 4 h. The mixture was filtered through a celite pad, and the filtrate was concentrated *in vacuo*. The product was recrystallized from the residue using cold $CHCl_3$ to obtain **17c** (636 mg, 78%)

as a white solid. $^1$H NMR ($CD_3OD$, 500 MHz) $\delta$ 3.09 (s, 2H), 1.38 – 1.37 (m, 2H), 1.08 – 1.06 (m, 2H).

Compound **19c** was synthesized following the general procedure A from **17c** over two steps. Compound **17c** (580 mg, 3.83 mmol) and diethyl amine (1.2 mL, 11.48 mmol) were dissolved in MeOH and warmed, then D-pantolactone (498 mg, 3.83 mmol) was added and the mixture was stirred at 65°C for 72 h. **18c** (939 mg, quantitative) was obtained as a clear oil. $R_f$ = 0.60 (DCM/MeOH = 85:15, visualized by $KMnO_4$ stain).

Compound **18c** (939 mg, 3.83 mmol) was dissolved in dry DMF (4 mL). CSA (45 mg, 0.19 mmol) was added, followed by *p*-methoxybenzaldehyde dimethyl acetal (698 mg, 3.83 mmol), and the mixture was stirred at rt overnight. The reaction mixture was diluted in EA (40 mL) and washed with water (3 × 30 mL). After solvent evaporation, the solid obtained was washed with chilled $Et_2O$ (10 mL) and purified by column chromatography ($SiO_2$, Hex/EA = 50:50) to afford **19c** (357 mg, 26%) as a white solid. $R_f$ = 0.20 (Hex/EA = 50:50); Purity >99%, RT = 15.73 min with method 7; RT = 25.98 min with method 8; $^1$H NMR ($CDCl_3$, 500 MHz) $\delta$ 7.42 (m, 2H, H-9a, H-9b), 7.33 (t, *J* = 6.5 Hz, 1H, -NH), 6.88 (m, 2H, H-10a, H-10b), 5.47 (s, 1H, H-8), 4.11 (s, 1H, H-4), 3.79 (s, 3H, H-11), 3.69 (q, *J* = 11.4 Hz, 2H, H-1a, H-1b), 3.50 (m, 2H, H-5), 1.34-1.31 (m, 2H, H-6 or H-7), 1.11 (s, 3H, H-2 or H-3), 1.09 (s, 3H, H-2 or H-3), 1.08 – 1.06 (m, 2H, H-6 or H-7); $^{13}$C NMR ($CDCl_3$, 126 MHz) $\delta$ 180.9, 169.4, 160.1, 130.3, 127.3, 113.7, 101.0, 83.7, 76.8, 55.2, 41.3, 33.1, 24.6, 21.9, 19.1, 15.8, 15.6; HRMS (ESI$^+$) *m/z* $[C_{19}H_{25}NO_6Na]^+$ calcd: 386.1580; found: 386.1568.

<u>Synthesis of **3**</u>

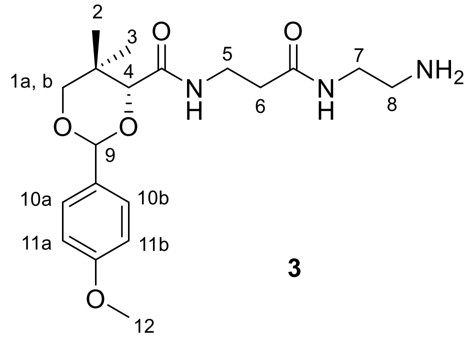

**3**

Compound **3** was synthesized by a reported protocol.[2] Compound **2** (3 g, 8.89 mmol), NHS (1.02 g, 8.89 mmol) and DCC (1.93 g, 9.34 mmol) were dissolved in dry THF (50 mL). The reaction mixture was stirred at rt for 4 h, at which point the precipitate was removed by filtration and was washed with EA (30 mL). The combined organic solvents were evaporated to afford the NHS ester, which was redissolved in dry THF (40 mL) and added slowly to a previously prepared solution of ethylene diamine (5.94 mL, 88.92 mmol) in dry THF (30 mL). The mixture was stirred at rt overnight. The dicyclohexyl urea product was removed by filtration and washed with EA (30 mL). The solvents and diethyl amine were removed under reduced pressure, and the residue was dissolved into DCM (20 mL) and filtered again. Solvents were removed *in vacuo*, and the residue was purified by column chromatography ($SiO_2$, DCM/MeOH = 90:10) to afford compound **3** as a sticky yellow solid (2.23 g, 66%). $R_f = 0.20$ (DCM/MeOH = 85:15); $^1$H NMR ($CDCl_3$, 500 MHz) $\delta$ 7.42 (d, $J$ = 8.7 Hz, 2H, H-10a, H-10b), 7.07 (m, 1H, -NH), 6.91 (d, $J$ = 8.7 Hz, 2H, H-11a and H-11b), 6.34 (bs, 1H, -NH), 5.45 (s, 1H, H-9), 4.07 (s, 1H, H-4), 3.81 (s, 3H, H-12), 3.70 (d, $J$ = 11.4 Hz, 1H, H-1a), 3.65 (d, $J$ = 11.4 Hz, 1H, H-1b), 3.60-3.49 (m, 2H, H-5 or H-7), 3.30-3.20 (m, 2H, H-5 or H-7), 2.75 (t, $J$ = 5.9 Hz, 2H, H-8), 2.42 (t, $J$ = 6.3 Hz, 2H, H-6), 1.10 (s, 3H, H-2 or H-3), 1.09 (s, 3H, H-2 or H-3).

<u>Synthesis of **20a**</u>

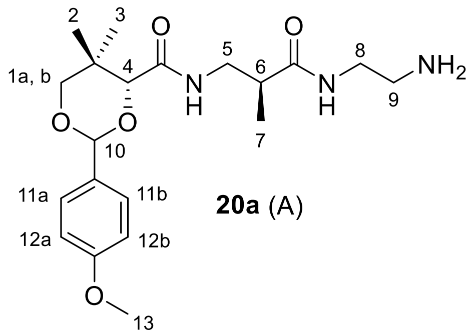

**20a** (A)

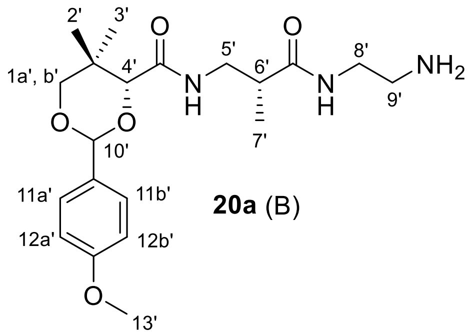

**20a** (B)

Compound **19a** (800 mg, 2.28 mmol) was dissolved in dry THF (15 mL). NHS (288 mg, 2.50 mmol) was added followed by DCC (517 mg, 2.50 mmol). The reaction mixture was stirred at rt for 4 h at which point the precipitate was filtered off and washed with EA (25 mL). The combined organic solvents were evaporated to afford the NHS ester which was used without purification. The NHS ester was redissolved in dry THF (10 mL) and added slowly to a previously prepared solution of ethylene diamine (1.52 mL, 22.76 mmol) in dry THF (5 mL). The mixture was stirred at rt overnight. The dicyclohexyl urea product was removed by filtration and washed with EA (25 mL). The organic solvents and diethyl amine were removed under reduced pressure, and the residue was dissolved in DCM (25 mL) and filtered again. After solvent evaporation, the residue was purified by column chromatography ($SiO_2$, DCM/MeOH = 90:10 to 88:12) to afford **20a** (748 mg, 83%) as a yellow oil for this pair of diastereomers. $R_f$ = 0.50 (MeOH/DCM/$Et_3N$ = 14:85:1); Purity 78%, RT = 16.04 min with method 10; RT = 11.21 min with method 11; $^1$H NMR ($CDCl_3$, 300 MHz) $\delta$ 7.50 – 7.36 (m, 4H, H-11a, H-11b and H-11a', H-11b'), 7.15 – 7.00 (m, 2H, -NH and –NH'), 6.92 (d, $J$ = 8.7 Hz, 4H, H-12a, H-12b, H-12a' and H-12b'), 6.36 – 6.18 (m, 2H, -NH and –NH'), 5.46 (s, 1H, H-10 or H-10'), 5.45 (s, 1H, H-10 or H-10'), 4.09 (s, 1H, H-4 or H-4'), 4.08 (s, 1H, H-4 or H-4'), 3.82 (s, 6H, H-13 and H-13'), 3.71 (d, $J$ = 11.4 Hz, 2H, H-1a, H-1b, or H-1a', H-1b'), 3.65 (d, $J$ = 11.5 Hz, 2H, H-1a, H-1b, or H-1a', H-1b'), 3.48 – 3.12 (m, 8H, H-5, H-8, H-5' and H-8'), 2.76 (t, $J$ = 5.9 Hz, 2H, H-9 or H-9'), 2.68 (t, $J$ = 5.9 Hz, 2H, H-9 or H-9'), 2.62 – 2.46 (m, 2H, H-6 and H-6'), 1.18 – 1.03 (m, 18H, H-2, H-3, H-7, H-2', H-3' and H-7'); $^{13}$C NMR ($CDCl_3$, 126 MHz) $\delta$ 175.0, 175.0, 169.6, 169.6, 160.3, 160.2, 130.2, 130.1, 127.5, 127.5, 113.7, 101.4, 101.3, 84.0, 83.8, 78.5, 78.5, 55.4, 42.1, 42.0, 41.3, 41.3, 41.1, 41.1, 41.0, 40.8, 33.1, 33.1, 21.9, 21.8, 19.2, 19.2, 15.7, 15.5; HRMS (ESI$^+$) *m/z* $[C_{20}H_{31}N_3O_5Na]^+$ calcd: 416.2161; found: 416.2147.

Synthesis of **20b**

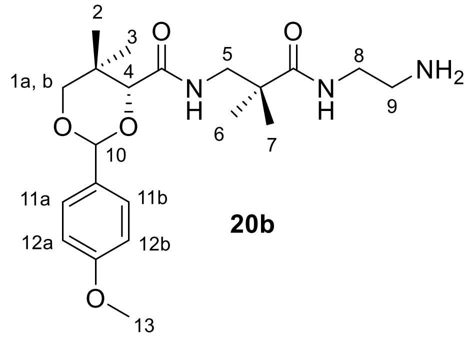

**20b**

Compound **19b** (586 mg, 1.60 mmol) was dissolved in dry THF (15 mL). NHS (185 mg, 1.60 mmol) was added followed by DCC (348 mg, 1.68 mmol). The reaction mixture was stirred at rt for 4 h, at which point the precipitate was filtered off and washed with EA (20 mL). The combined organic solvents were evaporated to afford the NHS ester which was used without purification. The NHS ester was redissolved in dry THF (10 mL) and added slowly to a previously prepared solution of ethylene diamine (1.1 mL, 16.04 mmol) in THF (5 mL). The mixture was stirred at rt overnight. The dicyclohexyl urea precipitate was removed by filtration and washed with EA (20 mL). The organic layers were combined, and the solvents and diethyl amine were removed under reduced pressure. The residue was dissolved into DCM (20 mL) and filtered again. After evaporation of DCM, the residue was purified by silica gel column chromatography with a gradient of DCM/MeOH/$Et_3N$ 89:10:1. Compound **20b** was isolated as a white solid (454 mg, 70%). $R_f$ = 0.50 (14:85:1 MeOH/DCM/$Et_3N$); $^1$H NMR ($CDCl_3$, 500 MHz) $\delta$ 7.44 (m, 2H, H-11a, H-11b), 7.15 (t, $J$ = 6.6 Hz, 1H, -NH), 6.92 (m, 2H, H-12a, H-12b), 6.45 (t, $J$ = 5.6 Hz, 1H, -NH), 5.46 (s, 1H, H-10), 4.10 (s, 1H, H-4), 3.82 (s, 3H, H-13), 3.76 – 3.62 (m, 2H, H-1a, H-1b), 3.41 (m, 2H, H-5), 3.21 (m, 2H, H-8), 2.69 (t, $J$ = 5.9 Hz, 2H, H-9), 1.20 (s, 3H, H-6 or H-7), 1.18 (s, 3H, H-6 or H-7), 1.12 (s, 3H, H-2 or H-3), 1.09 (s, 3H, H-2 or H-3); $^{13}$C NMR ($CDCl_3$, 126 MHz) $\delta$ 176.7, 169.6, 160.2, 130.3, 127.5, 113.7, 101.3, 84.0, 78.5, 55.4, 46.8, 43.3, 41.7, 41.1, 33.1, 24.0, 23.7, 21.9, 19.3; HRMS (ESI$^+$) $m/z$ $[C_{21}H_{34}N_3O_5]^+$ calcd: 408.2493; found: 408.2484.

Synthesis of **20c**

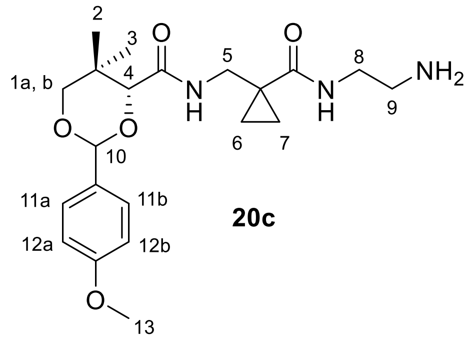

**20c**

Compound **19c** (330 mg, 0.91 mmol) was dissolved in dry THF (10 mL). NHS (105 mg, 0.91 mmol) was added followed by DCC (197 mg, 0.95 mmol). The reaction mixture was stirred at rt for 4 h, at which point the precipitate was filtered off and washed with EA (15 mL). The solvents were evaporated to afford the NHS ester which was used without purification. The NHS ester was redissolved in dry THF (10 mL) and added slowly to a previously prepared solution of ethylene diamine (0.61 mL, 9.08 mmol) in dry THF (5 mL). The mixture was stirred at rt overnight. The dicyclohexyl urea precipitate was removed by filtration and washed with EA (15 mL). The organic layers were combined, and the solvents and diethyl amine were removed under reduced pressure. The residue was dissolved in DCM (15 mL) and filtered again. After evaporation, the residue was purified by column chromatography ($SiO_2$, DCM/MeOH/$Et_3N$ =

89:10:1) to afford compound **20c** (244 mg, 66%) as a white solid. $R_f = 0.50$ (MeOH/DCM/$Et_3N$ = 14:85:1); $^1H$ NMR ($CDCl_3$, 500 MHz) $\delta$ 7.72 (t, $J$ = 5.4 Hz, 1H, -NH), 7.42 (m, 2H, H-11a, H-11b), 6.99 – 6.88 (m, 3H, H-12a, H-12b, -NH), 5.47 (s, 1H, H-10), 4.12 (s, 1H, H-4), 3.82 (s, 3H, H-13), 3.73 – 3.65 (m, 2H, H-1a, H-1b), 3.62-3.17 (m, 4H, H-5, H-8), 2.80 (t, $J$ = 6.0 Hz, 2H, H-9), 1.25 (m, 2H, H-6 or H-7), 1.09 (s, 6H, H-2, H-3), 0.67 (m, 2H, H-6 or H-7); $^{13}C$ NMR ($CDCl_3$, 126 MHz) $\delta$ 172.8, 170.4, 160.3, 130.0, 127.6, 113.8, 101.5, 83.8, 78.4, 55.3, 43.2, 42.6, 41.6, 33.2, 25.3, 21.9, 19.1, 14.8, 14.2; HRMS (ESI$^+$) *m/z* $[C_{21}H_{32}N_3O_5]^+$ calcd: 406.2336; found: 406.2332.

<u>Synthesis of **5a**</u>

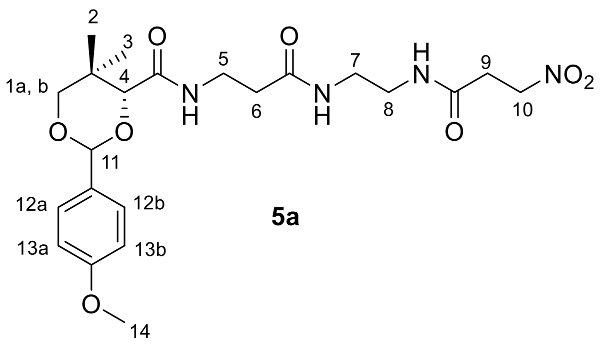

**5a**

Known compound **5a** was obtained following the general procedure B from compound **3**.**Error! Bookmark not defined.** To a solution of 3-nitropropanoic acid **4a** (1 g, 8.34 mmol), NHS (1.0 g, 8.76 mmol) and DCC (1.89 g, 9.17 mmol) in dry THF (30 mL) was added a solution of compound **3** (3.32 g, 8.76 mmol) in dry THF (20 mL) followed by $Et_3N$ (3.5 mL, 25.02 mmol) addition. Product purification by column chromatography ($SiO_2$, DCM/MeOH = 95:5) afforded **5a** (2.88 g, 72%) as an off-white solid. $R_f = 0.50$ (DCM/MeOH = 90:10); Purity >99%, RT = 14.66 min with method 7; RT = 14.05 min with method 8; $^1H$ NMR ($CDCl_3$, 500 MHz) $\delta$ 7.43 (m, 2H, H-12a, H-12b), 7.01 (t, $J$ = 6.2 Hz, 1H, -NH), 6.94 (m, 2H, H-13a, H-13b), 6.82 (s, 1H, -NH), 6.51 (s, 1H, -NH), 5.47 (s, 1H, H-11), 4.65 (m, 2H, H-10), 4.09 (s, 1H, H-4), 3.82 (s, 3H, H-14), 3.69 (m, 2H, H-1a, H-1b), 3.56 (m, 2H, H-5), 3.45 – 3.29 (m, 4H, H-7, H-8), 2.74 (m, 2H, H-9), 2.44 (m, 2H, H-6), 1.11 (s, 3H, H-2 or H-3), 1.08 (s, 3H, H-2 or H-3).

<u>Synthesis of **5b**</u>

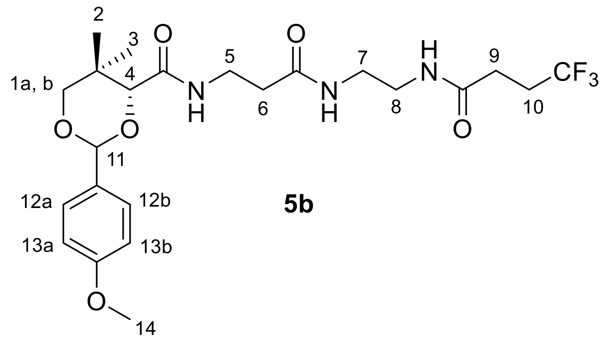

**5b**

Compound **5b** was obtained following the general procedure B from compound **3**. To a solution of 4,4,4-trifluorobutanoic acid **4b** (80 mg, 0.56 mmol), NHS (65 mg, 0.56 mmol) and DCC (122 mg, 0.59 mmol) in dry THF (8 mL) was added a solution of compound **3** (213 mg, 0.56 mmol) in dry THF (5 mL), followed by $Et_3N$ (0.20 mL, 1.40 mmol). Product purification by column chromatography ($SiO_2$, DCM/MeOH = 95:5) afforded **5b** (192 mg, 66%) as an off-white solid. $R_f$ = 0.50 (DCM/MeOH = 90:10); Purity 92%, RT = 14.58 min with method 7; RT = 21.66 min with method 8; $^1$H NMR ($CDCl_3$, 300 MHz) $\delta$ 7.43 (d, $J$ = 8.6 Hz, 2H, H-12a, H-12b), 7.02 (m, 1H, -NH), 6.92 (d, $J$ = 8.8 Hz, 2H, H-13a, H-13b), 6.76 – 6.56 (m, 2H, -NH), 5.46 (s, 1H, H-11), 4.08 (s, 1H, H-4), 3.82 (s, 3H, H-14), 3.72 (d, $J$ = 11.5 Hz, 1H, H-1a), 3.65 (d, $J$ = 11.5 Hz, 1H, H-1b), 3.53 (m, 2H, H-5), 3.46 – 3.23 (m, 4H, H-7, H-8), 2.58 – 2.29 (m, 6H, H-6, H-9, H-10), 1.10 (s, 3H, H-2 or H-3), 1.08 (s, 3H, H-2 or H-3); $^{13}$C NMR ($CD_3OD$, 75 MHz) $\delta$ 174.1, 173.0, 171.6, 161.7, 131.8, 128.8, 114.5, 102.7, 85.1, 79.3, 55.7, 40.1, 39.9, 36.5, 36.3, 34.0, 30.4 (q, $J$ = 29.6 Hz), 29.3 (q, $J$ = 3.1 Hz), 22.2, 19.7; $^{19}$F NMR ($CDCl_3$, 282 MHz) $\delta$ -66.71 (t, $J$ = 10.3 Hz); HRMS (ESI$^+$) $m/z$ $[C_{23}H_{33}F_3N_3O_6Na]^+$ calcd: 526.2141; found: 526.2130.

Synthesis of **5c**

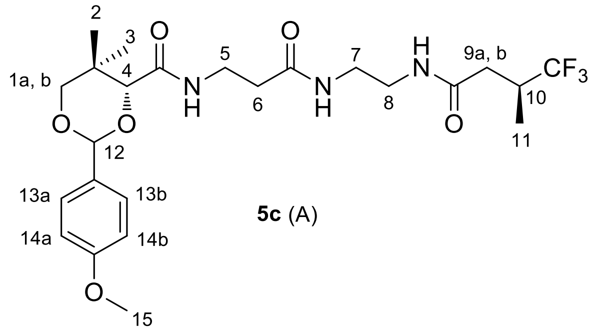

**5c** (A)

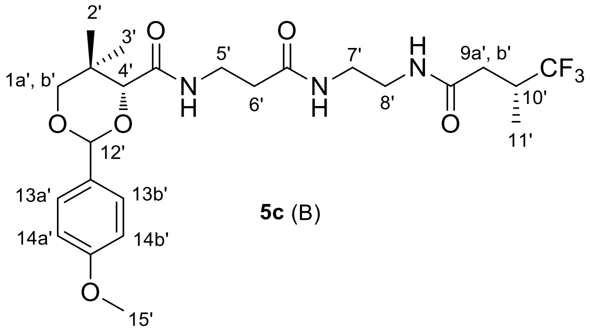

**5c** (B)

Compound **5c** was obtained following the general procedure B from compound **3.** To a solution of 4,4,4-trifluoro-3-methylbutanoic acid **4c** (80 mg, 0.51 mmol), NHS (59 mg, 0.51 mmol) and DCC (111 mg, 0.54 mmol) in dry THF (8 mL) was added a solution of compound **3** (194 mg, 0.51 mmol) in dry THF (5 mL), followed by $Et_3N$ (0.18 mL, 1.281 mmol). Product purification by column chromatography ($SiO_2$, DCM/MeOH = 95:5) afforded **5c** (208 mg, 76%) as a white solid for this pair of diastereomers. $R_f$ = 0.50 (DCM/MeOH = 90:10); Purity 93%, RT = 14.92 min with method 7; RT = 23.73 min with method 8; $^1$H NMR ($CDCl_3$, 300 MHz) $\delta$ 7.44 (m, 4H, H-13a, H-13b and H-13a', H-13b'), 7.06 – 6.94 (bs, 2H, -NH and –NH'), 6.91 (m, 4H, 14a, 14b and 14a', 14b'), 6.64 (bs, 4H, -NH, -NH and –NH', -NH'), 5.46 (s, 2H, H-12 and H-12'), 4.07 (s, 2H, H-4 and H-4'), 3.81 (s, 6H, H-15 and H-15'), 3.76 – 3.60 (m, 4H, H-1a, H-b and H-1a', H-b'), 3.60 – 3.45 (m, 4H, H-5 and H-5'), 3.47 – 3.18 (m, 8H, H-7, H-8, and H-7', H-8'), 2.92 – 2.72 (m, 2H, H-10 and H-10'), 2.57 – 2.45 (m, 2H, H-9a and H-9a'), 2.42 (t, $J$ = 6.1 Hz, 4H, H-6 and H-6'), 2.15 – 2.00 (m, 2H, H-9b and H-9b'), 1.12 – 1.08 (m, 18H, H-2, H-3, H-11, H-2', H-3', and H-11'). $^{13}$C NMR (126 MHz, $CDCl_3$) $\delta$ 172.0, 172.0, 170.5, 169.8, 169.7, 160.3, 130.1, 128.0 (q, $J$ = 278.8 Hz), 127.5, 113.7, 101.4, 83.8, 78.4, 77.3, 77.1, 76.8, 55.3, 40.0, 40.0, 39.6, 36.5, 36.4, 36.3, 36.2, 35.1 (q, $J$ = 27.5 Hz), 35.0, 33.1, 21.8, 19.1, 13.0, 13.0; $^{19}$F NMR ($CDCl_3$, 471 MHz) $\delta$ -73.50 – -73.58 (m); HRMS (ESI$^+$) $m/z$ $[C_{24}H_{34}F_3N_3O_6Na]^+$ calcd: 540.2297; found: 540.2284.

Synthesis of **5d**

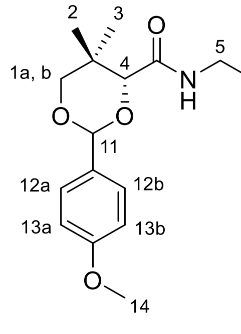

**5d** (A)

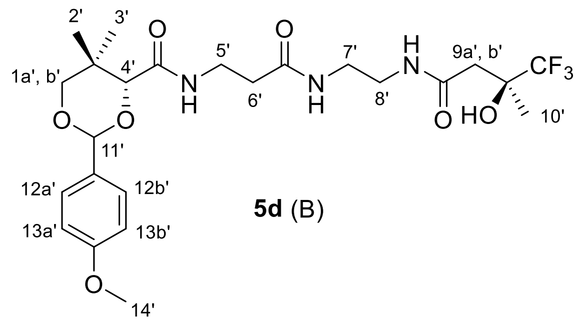

**5d** (B)

Compound **5d** was obtained following the general procedure B from compound **3**. To a solution of 4,4,4-trifluoro-3-hydroxy-3-methylbutanoic acid (**4d**, 60 mg, 0.35 mmol), NHS (40 mg, 0.35 mmol) and DCC (76 mg, 0.37 mmol) in dry THF (10 mL) was added a solution of compound **3** (132 mg, 0.35 mmol) in dry THF (5 mL), followed by $Et_3N$ (0.05 mL, 0.35 mmol). Product purification by column chromatography ($SiO_2$, DCM/MeOH = 96:5) afforded **5d** (116 mg, 62%) as a white solid for this pair of diastereomers. $R_f$ = 0.50 (DCM/MeOH = 90:10); Purity 89%, RT = 14.50 min with method 7; RT = 21.05 min with method 8; $^1$H NMR ($CDCl_3$, 300 MHz) *δ* 7.43 (m, 4H, H-12a, H-12b and H-12a', H-12b'), 7.14 – 6.96 (m, 4H, -NH, -NH and –NH', -NH'), 6.93 (m, 4H, H-13a, H-13b, H-13a', and H-13b'), 6.73 – 6.54 (m, 2H, -NH and –NH'), 5.46 (s, 2H, H-11 and H-11'), 4.09 (s, 2H, H-4 and H-4'), 3.82 (s, 6H, H-14 and H-14'), 3.77 – 3.60 (m, 4H, H-1a, H-1b, H-1a', and H-1b'), 3.63 – 3.18 (m, 14H, H-5, H-7, H-8, -OH, H-5', H-7', H-8', and -OH'), 2.58 – 2.34 (m, 8H, H-6, H-9a, H-9b, H-6', H-9a', and H-9b') 1.40 (s, 6H, H-10 and H-10'), 1.20 – 1.03 (m, 12H, H-2, H-3, H-2', and H-3'); $^{13}$C NMR ($CDCl_3$, 75 MHz) *δ* $^{13}$C NMR ($CDCl_3$, 75 MHz) *δ* 172.1, 171.1, 171.1, 169.9, 169.9, 160.2, 130.0, 127.5, 125.9 (q, *J* = 285.7 Hz), 113.7, 101.3, 101.3, 83.8, 83.7, 78.3, 72.4 (q, *J* = 28.8 Hz), 55.3, 55.3, 55.3, 55.3, 39.6, 39.6, 39.5, 39.2, 36.2, 36.1, 35.2, 35.1, 33.1, 33.0, 21.8, 21.5, 19.0; $^{19}$F NMR ($CDCl_3$, 282 MHz) *δ* -82.55, -82.63; HRMS (ESI$^+$) *m/z* $[C_{24}H_{34}F_3N_3O_7Na]^+$ calcd: 556.2247; found: 556.2228.

Synthesis of **5e**

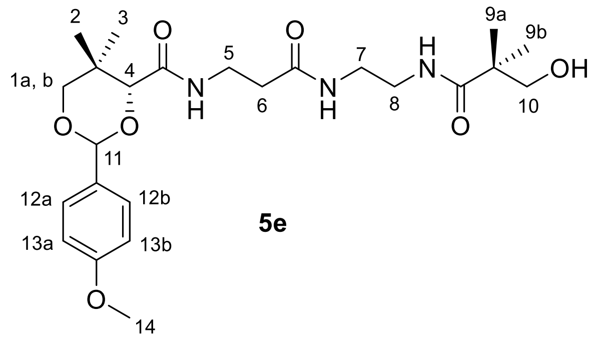

**5e**

Compound **5e** was obtained following the general procedure B from compound **3**. To a solution of hydroxypivalic acid (**4e**, 467 mg, 3.95 mmol), NHS (91 mg, 0.79 mmol) and DCC (163 mg, 0.79 mmol) in dry THF (20 mL) was added a solution of compound **3** (300 mg, 0.79 mmol) in dry THF (10 mL), followed by addition of $Et_3N$ (0.12 mL, 0.79 mmol). Product purification by column chromatography ($SiO_2$, DCM/MeOH = 94:6) afforded **5e** (155 mg, 41%) as a white solid. $R_f$ = 0.50 (DCM/MeOH = 90:10); Purity 98%, RT = 13.59 min with method 7; RT = 17.40 min with method 8; $^1$H NMR ($CDCl_3$, 500 MHz) *δ* 7.43 (m, 2H, H-12a, H-12b), 7.03 (t, *J* = 6.2 Hz, 1H, -NH), 6.91 (m, 2H, H-13a, H-13b), 6.81 (dd, *J* = 12.3, 6.4 Hz, 2H, -NH and -NH), 5.46 (s, 1H, H-11), 4.08 (s, 1H, H-4), 3.82 (s, 3H, H-14), 3.71 (d, *J* = 11.4 Hz, 1H, H-1a), 3.65 (d, *J* =

11.5 Hz, 1H, H-1b), 3.58 – 3.44 (m, 4H, H-5 and H-10), 3.43 – 3.28 (m, 4H, H-7, H-8), 2.41 (t, *J* = 6.5 Hz, 2H, H-6), 1.13 (s, 6H, H-9a, H-9b), 1.09 (s, 3H, H-2 or H-3), 1.08 (s, 3H, H-2 or H-3); $^{13}$C NMR ($CDCl_3$, 101 MHz) $\delta$ 178.8, 172.3, 169.8, 160.3, 130.2, 127.6, 113.8, 101.4, 83.9, 78.5, 69.9, 55.4, 43.3, 40.0, 39.4, 36.2, 35.3, 33.1, 22.5, 21.9, 19.2; HRMS (ESI$^+$) *m/z* $[C_{24}H_{37}N_3O_7Na]^+$ calcd: 502.2529; found: 502.2523.

Synthesis of **5f**

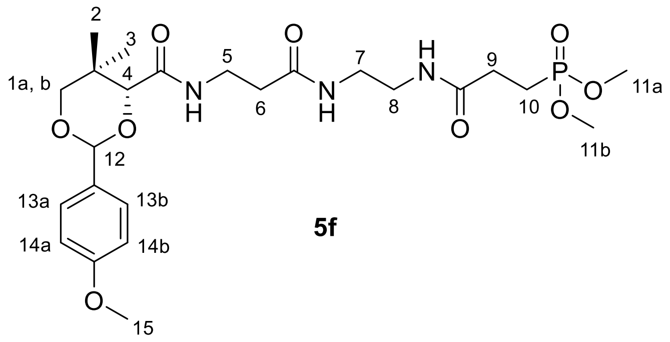

**5f**

Compound **5f** was obtained following the general procedure B from compound **3**. To a solution of **4f** (120 mg, 0.66 mmol), NHS (76 mg, 0.66 mmol) and DCC (143 mg, 0.69 mmol) in dry THF (12 mL) was added a solution of compound **3** (250 mg, 0.66 mmol) in dry THF (6 mL), followed by $Et_3N$ (0.1 mL, 0.66 mmol). Product purification by column chromatography ($SiO_2$, DCM/MeOH = 94:6) afforded **5f** (166 mg, 46%) as a white solid. $R_f$ = 0.50 (DCM/MeOH = 90:10); Purity >99%, RT = 13.03 min with method 7; RT = 16.69 min with method 8; $^1$H NMR ($CDCl_3$, 500 MHz) $\delta$ 7.43 (m, 2H, H-13a, H-13b), 7.13 (t, *J* = 5.3 Hz, 1H, -NH), 7.01 (t, *J* = 6.2 Hz, 1H, -NH), 6.91 (m, 2H, H-14a, H-14b), 6.73 (t, *J* = 5.3 Hz, 1H, -NH), 5.46 (s, 1H, H-12), 4.08 (s, 1H, H-4), 3.82 (s, 3H, H-15), 3.75 – 3.62 (m, 8H, H-1a, H-1b, H-11a and H-11b), 3.62 – 3.50 (m, 2H, H-7 or H-8), 3.40 – 3.33 (m, 2H, H-7 or H-8), 3.29 (m, 2H, H-5), 2.47 – 2.40 (m, 4H, H-6, H-9), 2.11 (m, 2H, H-10), 1.11 (s, 3H, H-2 or H-3), 1.08 (s, 3H, H-2 or H-3); $^{13}$C NMR ($CDCl_3$,126 MHz) $\delta$ 171.6, 171.5, 169.4, 160.1, 130.0, 127.4, 113.6, 101.2, 83.8, 78.3, 52.5 (dd, *J* = 6.6, 1.2 Hz), 55.2, 39.5, 39.1, 36.0, 35.1, 33.0, 28.8 (d, *J* = 4.0 Hz), 21.7, 19.9 (d, *J* = 142.8 Hz), 19.0; $^{31}$P NMR ($CDCl_3$, 203 MHz) $\delta$ 33.61; HRMS (ESI$^+$) *m/z* $[C_{24}H_{38}N_3O_9PNa]^+$ calcd: 566.2243; found: 566.2223.

Synthesis of **5g**

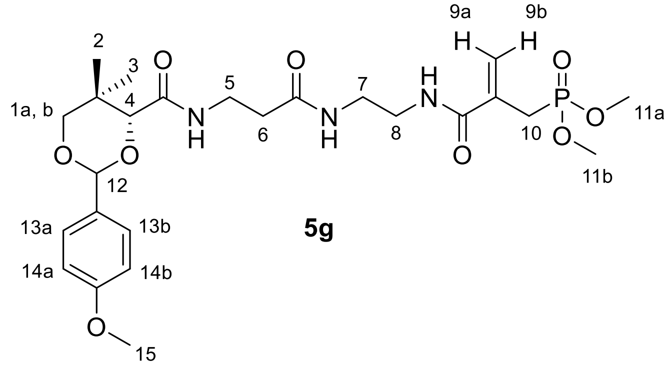

**5g**

Compound **5g** was obtained following the general procedure B from compound **3**. To a solution of **4g** (178 mg, 0.92 mmol), NHS (106 mg, 0.92 mmol) and DCC (284 mg, 1.38 mmol) in dry THF (8 mL) was added a solution of compound **3** (348 mg, 0.92 mmol) in dry THF (5 mL), followed by $Et_3N$ (0.32 mL, 2.3 mmol). Product purification by column chromatography ($SiO_2$, DCM/MeOH = 95:5) afforded **5g** (170 mg, 33%) as a white solid. $R_f$ = 0.50 (DCM/MeOH = 90:10); Purity >99%, RT = 13.30 min with method 7; RT = 17.17 min with method 8; $^1$H NMR ($CDCl_3$, 500 MHz) $\delta$ 7.43 (m, 2H, H-13a, H-13b), 7.18 (m, 2H, -NH and -NH), 7.03 (t, $J$ = 6.3 Hz, 1H, -NH), 6.92 (m, 2H, H-14a, H-14b), 5.91 (d, $J$ = 5.4 Hz, 1H, H-9a), 5.47 (d, $J$ = 5.8 Hz, 1H, H-9b), 5.46 (s, 1H, H-12), 4.07 (s, 1H, H-4), 3.82 (s, 3H, H-15), 3.75 (d, $J$ = 2.3 Hz, 3H, H-11a or H-11b), 3.72 (d, $J$ = 2.3 Hz, 3H, H-11a or H-11b), 3.72 – 3.63 (m, 2H, H-1a, H-1b), 3.57 (m, 2H, H-5), 3.45 – 3.32 (m, 4H, H-7, H-8), 2.95 (m, 2H, H-10), 2.41 (t, $J$ = 6.2 Hz, 2H, H-6), 1.12 (s, 3H, H-2 or H-3), 1.07 (s, 3H, H-2 or H-3); $^{13}$C NMR ($CDCl_3$,126 MHz) $\delta$ 171.8, 169.3, 167.8, 160.0, 135.1 (d, $J$ = 10.3 Hz), 130.0, 127.4, 122.7 (d, $J$ = 11.7 Hz), 113.6, 101.2, 83.7, 78.3, 55.2, 52.9 (dd, $J$ = 6.5, 3.0 Hz), 39.8, 38.7, 35.8, 35.0, 32.9, 30.5 – 27.4 (m), 21.7, 19.0; $^{31}$P NMR ($CDCl_3$, 203 MHz) $\delta$ 28.21; HRMS (ESI$^+$) *m/z* $[C_{25}H_{38}N_3O_9PNa]^+$ calcd: 578.2243; found: 578.2200.

Synthesis of **5h**

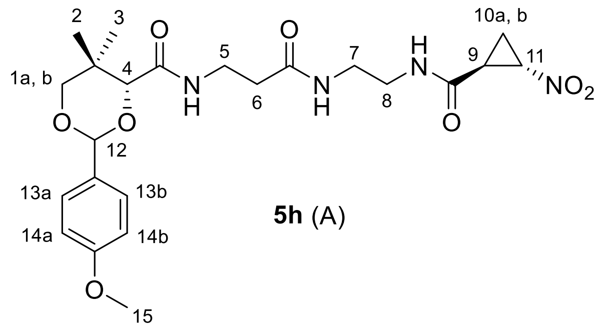

**5h** (A)

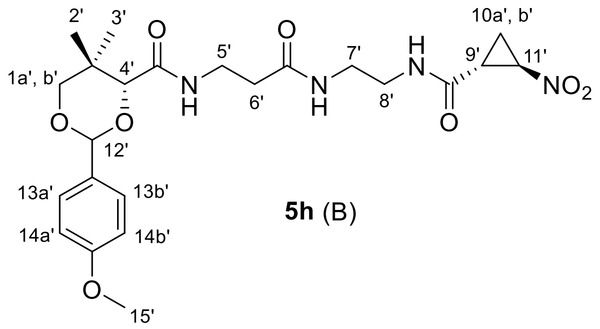

**5h** (B)

Compound **5h** was obtained following the general procedure B from compound **3.** To a solution of **4h** (105 mg, 0.80 mmol), NHS (92 mg, 0.80 mmol) and DCC (174 mg, 0.84 mmol) in dry THF (10 mL) was added a solution of compound **3** (304 mg, 0.80 mmol) in dry THF (5 mL), followed by $Et_3N$ (0.28 mL, 2.00 mmol). Product purification by column chromatography ($SiO_2$, DCM/MeOH = 94:6) afforded **5h** (160 mg, 41%) as a white solid for this pair of diastereomers. $R_f$ = 0.60 (DCM/MeOH = 90:10); Purity 97%, RT = 14.34 min with method 7; RT = 20.29 min with method 8; $^1$H NMR ($CDCl_3$,500 MHz) $\delta$ 7.46 – 7.40 (m, 4H, H-13a, H-13b, H-13a', and H-13b'), 7.25 (bs, 2H, -NH and –NH'), 7.03 – 6.96 (m, 2H, -NH and –NH'), 6.96 – 6.89 (m, 4H, H-14a, H-14b, H-14a' and H-14b'), 6.70 – 6.61 (m, 2H, -NH and –NH'), 5.52 – 5.42 (m, 2H, H-12 and H-12'), 4.61 – 4.48 (m, 2H, H-11 and H-11'), 4.12 – 4.03 (m, 2H, H-4 and H-4'), 3.82 (s, 6H, H-15 and H-15'), 3.75 – 3.63 (m, 4H, H-1a, H-1b, H-1a', H-1b'), 3.63 – 3.46 (m, 4H, H-5 and H-5'), 3.45 – 3.27 (m, 8H, H-7, H-8, H-7' and H-8'), 2.61 – 2.52 (m, 2H, H-9 and H-9'), 2.47 – 2.37 (m, 4H, H-6 and H-6'), 2.00 – 1.87 (m, 2H, H-10a and H-10a'), 1.74 – 1.68 (m, 2H, H-10b and H-10b'), 1.13 – 1.08 (m, 6H, H-2 or H-3, and H-2' or H-3'), 1.09 – 1.05 (s, 6H, H-2 or H-3, and H-2' or H-3'); $^{13}$C NMR ($CDCl_3$,75 MHz) $\delta$ 167.2, 167.2, 165.1, 165.0, 163.4, 155.5, 125.3, 122.8, 108.9, 96.5, 79.0, 73.5, 72.0, 54.4, 50.5, 35.0, 35.0, 34.6, 34.5, 31.3, 31.3, 30.4, 28.3, 28.2, 21.9, 17.0, 14.4, 14.3, 11.8; HRMS (ESI$^+$) *m/z* $[C_{23}H_{32}N_4O_8Na]^+$ calcd: 515.2118; found: 515.2128.

Synthesis of **5i**

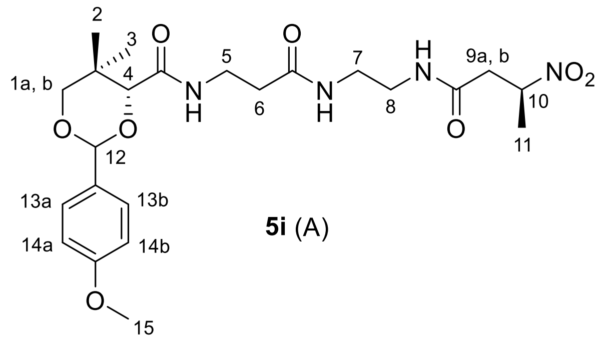

**5i** (A)

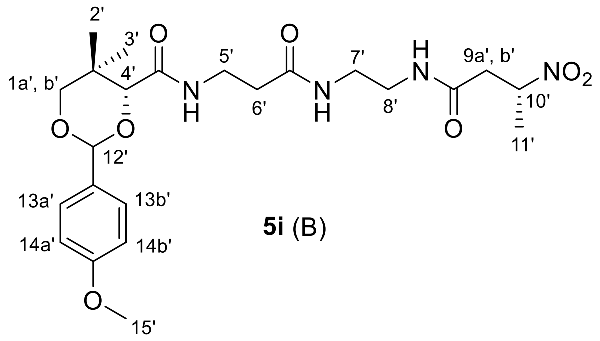

**5i** (B)

Compound **5i** was obtained following the general procedure B from compound **3**. To a solution of **4i** (95 mg, 0.71 mmol), NHS (82 mg, 0.71 mmol) and DCC (155 mg, 0.75 mmol) in dry THF (8 mL) was added a solution of compound **3** (271 mg, 0.71 mmol) in dry THF (4 mL), followed by $Et_3N$ (0.1 mL, 0.71 mmol). Product purification by column chromatography ($SiO_2$, DCM/MeOH = 95:5) afforded **5i** (224 mg, 63%) as an off-white solid for this pair of diastereomers. $R_f$ = 0.50 (DCM/MeOH = 90:10); Purity 92%, RT = 14.22 min with method 7; RT = 19.92 min with method 8; $^1$H NMR ($CDCl_3$, 500 MHz) $\delta$ 7.47 – 7.40 (m, 4H, H-13a, H-13b, H-13a’ and H-13b’), 7.01 (t, $J$ = 6.3 Hz, 2H, -NH and -NH’), 6.96 – 6.89 (m, 4H, H-14a, H-14b, H-14a’ and H-14b’), 6.79 (bs, 1H, -NH or -NH’), 6.73 (bs, 1H, -NH or -NH’), 6.54 – 6.52 (m, 2H, -NH, -NH’), 5.47 (s, 2H, H-12 and H-12’), 5.02 (dddd, $J$ = 9.2, 7.1, 4.6, 2.6 Hz, 2H, H-10 and H-10’), 4.09 (s, 1H, H-4 or H-4’), 4.09 (s, 1H, H-4 or H-4’), 3.82 (s, 6H, H-15 and H-15’), 3.71 (dd, $J$ = 11.4, 2.2 Hz, 2H, H-1a or H-1a’), 3.66 (d, $J$ = 11.5 Hz, 2H, H-1b or H-1b’), 3.63 – 3.47 (m, 4H, H-5 and H-5’), 3.46 – 3.21 (m, 8H, H-7, H-8, H-7’ and H-8’), 2.89 (m, 2H, H-9a and H-9b, or H-9a’ and H-9b’), 2.51 (m, 2H, H-9a and H-9b, or H-9a’ and H-9b’), 2.42 (td, $J$ = 6.3, 3.2 Hz, 4H, H-6 and H-6’), 1.58 (d, $J$ = 2.7 Hz, 3H, H-11 or H-11’), 1.57 (d, $J$ = 2.7 Hz, 3H, H-11 or H-11’), 1.11 (d, $J$ = 6.8 Hz, 6H, H-2, H-2’ or H-3, H-3’), 1.08 (d, $J$ = 2.7 Hz, 6H, H-2, H-2’ or H-3, H-3’); $^{13}$C NMR ($CDCl_3$, 101 MHz) $\delta$ 172.3, 172.3, 169.7, 169.7, 169.2, 160.3, 130.2, 127.6, 113.8, 101.4, 83.9, 83.9, 79.4, 79.3, 78.5, 55.4, 40.3, 40.2, 40.0, 39.9, 39.4, 36.2, 36.1, 35.2, 35.1, 33.2, 22.0, 19.4, 19.2; HRMS (ESI$^+$) $m/z$ $[C_{23}H_{34}N_4O_8Na]^+$ calcd: 517.2274; found: 517.2261.

Synthesis of **5j**

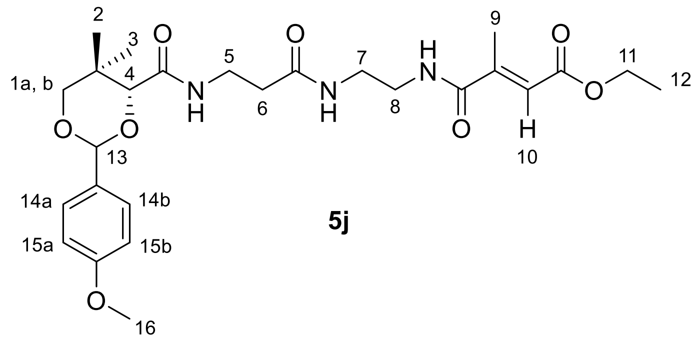

**5j**

Compound **5j** was obtained following the general procedure B from compound **3**. To a solution of **4j** (104 mg, 0.66 mmol), NHS (76 mg, 0.66 mmol) and DCC (143 mg, 0.69 mmol) in dry THF (8 mL) was added a solution of compound **3** (250 mg, 0.66 mmol) in dry THF (4 mL) followed by addition of $Et_3N$ (0.1 mL, 0.66 mmol). Product purification by column chromatography ($SiO_2$, DCM/MeOH = 96:4) afforded compound **5j** (194 mg, 57%) as a crystalline white solid. $R_f$ = 0.60 (DCM/MeOH = 90:10); Purity 98%, RT = 14.75 min with method 7; RT = 22.47 min with method 8; $^1$H NMR ($CDCl_3$, 500 MHz) $\delta$ 7.42 (m, 2H, H-14a and H-14b), 7.06 (s, 1H, -NH), 6.97 (t, $J$ = 6.4 Hz, 1H, -NH), 6.91 (m, 2H, H-15a and H-15b), 6.73 (s, 1H, -NH), 6.46 (q, $J$ = 1.5 Hz, 1H, H-10), 5.46 (s, 1H, H-13), 4.19 (q, $J$ = 7.1 Hz, 2H, H-11), 4.08 (s, 1H, H-4), 3.82 (s, 3H, H-16), 3.71 (d, $J$ = 11.5 Hz, 1H, H-1a), 3.65 (d, $J$ = 11.1 Hz, 1H, H-1b), 3.54 (m, 2H, H-5), 3.46 – 3.35 (m, 4H, H-7 and H-8), 2.45 (m, 2H, H-6), 2.29 (d, $J$ = 1.5 Hz, 3H, H-9), 1.29 (t, $J$ = 7.1 Hz, 3H, H-12), 1.09 (s, 3H, H-2 or H-3), 1.08 (s, 3H, H-2 or H-3); $^{13}$C NMR ($CDCl_3$, 126 MHz) $\delta$ 172.4, 169.9, 168.8, 166.2, 160.4, 147.7, 130.2, 127.6, 122.6, 113.8, 101.5, 83.9, 78.5, 77.4, 77.2, 76.9, 60.6, 55.4, 41.1, 39.5, 36.4, 35.1, 33.2, 21.9, 19.2, 14.6, 14.3; HRMS (ESI$^+$) *m/z* $[C_{26}H_{37}N_3O_8Na]^+$ calcd: 542.2478; found: 542.2458.

Synthesis of **5k**

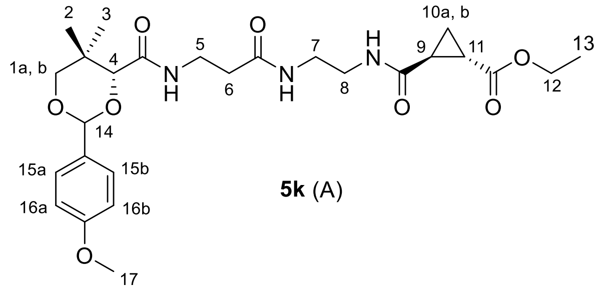

**5k** (A)

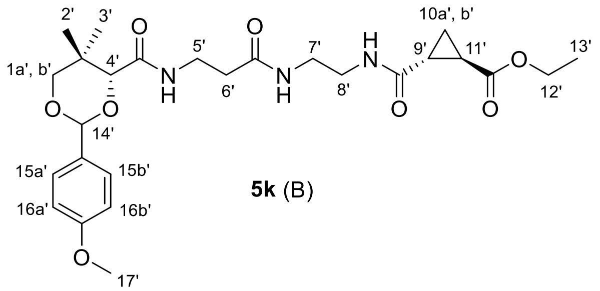

**5k** (B)

Compound **5k** was obtained following the general procedure B from compound **3**. To a solution of **4k** (200 mg, 1.26 mmol), NHS (146 mg, 1.26 mmol) and DCC (274 mg, 1.34 mmol) in dry THF (12 mL) was added a solution of compound **3** (504 mg, 1.33 mmol) in dry THF (6 mL), followed by $Et_3N$ (0.5 mL, 3.16 mmol). Product purification by column chromatography ($SiO_2$, DCM/MeOH = 96:4) afforded **5k** (386 mg, 59%) as a sticky white solid for this pair of diastereomers. $R_f$ = 0.60 (DCM/MeOH = 90:10); Purity 97%, RT = 14.45 min with method 7; RT = 20.71 min with method 8; $^1$H NMR ($CDCl_3$, 300 MHz) $\delta$ 7.43 (d, $J$ = 8.6 Hz, 4H, H-15a, H-15b, H-15a' and H-15b'), 7.07 – 6.96 (m, 2H, -NH and -NH'), 6.92 (d, $J$ = 8.7 Hz, 4H, H-16a, H-16b, H-16a' and H-16b'), 6.82 – 6.65 (m, 4H, -NH, -NH, –NH' and -NH'), 5.46 (s, 2H, H-14 and H-14'), 4.13 (q, $J$ = 7.2 Hz, 4H, H-12 and H-12'), 4.08 (s, 2H, H-4 and H-4'), 3.82 (s, 6H, H-17 and H-17'), 3.75 – 3.61 (m, 4H, H-1a, H-1b, H-1a' and H-1b'), 3.61 – 3.46 (m, 4H, H-5 and H-5'), 3.45 – 3.23 (m, 8H, H-7, H-8, H-7' and H-8'), 2.42 (t, $J$ = 6.2 Hz, 4H, H-6 and H-6'), 2.21 – 2.07 (m, 2H, H-9 or H-11, and H-9' or H-11'), 2.02 – 1.90 (m, 2H, H-9 or H-11, and H-9' or H-11'), 1.49 – 1.36 (m, 2H, H-10a and H-10a'), 1.35 – 1.29 (m, 2H, H-10b and H-10b'), 1.26 (t, $J$ = 7.1 Hz, 6H, H-13 and H-13'), 1.10 (s, 6H, H-2 or H-3, and H-2' or H-3'), 1.08 (s, 6H, H-2 or H-3, and H-2' or H-3'); $^{13}$C NMR ($CDCl_3$, 126 MHz) $\delta$ 172.7, 172.7, 171.8, 171.8, 171.2, 171.2, 169.8, 169.7, 160.3, 130.1, 127.6, 113.7, 101.4, 101.3, 83.8, 78.4, 61.0, 55.3, 40.0, 39.9, 39.8, 36.3, 36.2, 35.1, 33.1, 30.9, 24.0, 21.8, 21.8, 21.5, 19.1, 14.8, 14.2; HRMS (ESI$^+$) *m/z* $[C_{26}H_{37}O_8N_3Na]^+$ calcd: 542.2478; found: 542.2478.

Synthesis of **5l**

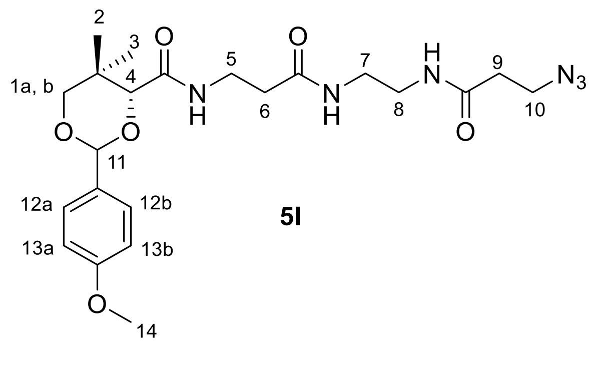

**5l**

Compound **5l** was obtained following the general procedure B from compound **3.** To a solution of **4l** (100 mg, 0.87 mmol), NHS (100 mg, 0.87 mmol) and DCC (188 mg, 0.91 mmol) in dry THF (8 mL) was added a solution of compound **3** (330 mg, 0.87 mmol) in dry THF (4 mL), followed by $Et_3N$ (0.12 mL, 0.87 mmol). Product purification by column chromatography ($SiO_2$, DCM/MeOH = 96:4) afforded **5l** (306 mg, 74%) as an off-white solid. $R_f$ = 0.50 (DCM/MeOH = 90:10); Purity 95%, RT = 14.17 min with method 7; RT = 18.95 min with method 8; $^1$H NMR ($CDCl_3$, 500 MHz) $\delta$ 7.44 (m, 2H, H-12a and H-12b), 7.01 (t, $J$ = 6.3 Hz, 1H, -NH), 6.93 (m, 2H, H-13a and H-13b), 6.59 (s, 2H, -NH and -NH), 5.47 (s, 1H, H-11), 4.09 (s, 1H, H-4), 3.82 (s, 3H, H-14), 3.71 (d, $J$ = 11.4 Hz, 1H, H-1a), 3.66 (d, $J$ = 11.5 Hz, 1H, H-1b), 3.58-3.53 (m, 4H, H-5 and H-10), 3.43 – 3.29 (m, 4H, H-7 and H-8), 2.43 (m, 2H, H-6), 2.38 (t, $J$ = 6.4 Hz, 2H, H-9), 1.10 (s, 3H, H-2 or H-3), 1.09 (s, 3H, H-2 or H-3); $^{13}$C NMR ($CDCl_3$, 126 MHz) $\delta$ 172.2, 170.9, 169.8, 160.4, 130.2, 127.7, 113.9, 101.5, 83.9, 78.5, 55.5, 47.4, 40.2, 39.7, 36.4, 35.8, 35.1, 33.2, 22.0, 19.2; HRMS (ESI$^+$) $m/z$ $[C_{22}H_{32}N_6O_6Na]^+$ calcd: 499.2281; found: 499.2284.

Synthesis of **5m**

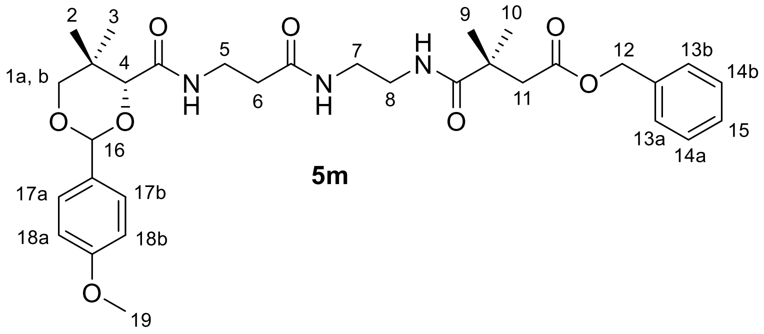

**5m**

Compound **5m** was obtained following the general procedure B from compound **3**. To a solution of **4m** (120 mg, 0.51 mmol), NHS (58 mg, 0.51 mmol) and DCC (110 mg, 0.53 mmol) in dry THF (8 mL) was added a solution of compound **3** (193 mg, 0.51 mmol) in dry THF (5 mL), followed by $Et_3N$ (0.18 mL, 1.27 mmol). Product purification by column chromatography ($SiO_2$, DCM/MeOH = 96:4) afforded **5m** (187 mg, 62%) as a clear oil. $R_f$ = 0.60 (DCM/MeOH = 90:10); Purity 85%, RT = 15.81 min with method 7; RT = 28.61 min with method 8; $^1$H NMR ($CDCl_3$, 500 MHz) $\delta$ 7.43 (d, $J$ = 8.7 Hz, 2H, H-17a, H-17b), 7.41 – 7.28 (m, 5H, H-13a, H-13b, H-14a, H-14b, H-15), 7.02 (m, 1H, -NH), 6.92 (d, $J$ = 8.7 Hz, 2H, H-18a, H-18b), 6.66 (bs, 1H, -NH), 6.50 (bs, 1H, -NH), 5.45 (s, 1H, H-16), 5.07 (s, 2H, H-12), 4.06 (s, 1H, H-4), 3.82 (s, 3H, H-19), 3.70 (d, $J$ = 11.4 Hz, 1H, H-1a), 3.64 (d, $J$ = 11.4 Hz, 1H, H-1b), 3.52 (m, 2H, H-5), 3.43 – 3.24 (m, 4H, H-7, H-8), 2.68 (s, 2H, H-11), 2.38 (dd, $J$ = 7.1, 5.3 Hz, 2H, H-6), 1.29 – 1.18 (m, 6H, H-9, H-10), 1.11 (s, 3H, H-2 or H-3), 1.07 (s, 3H, H-2 or H-3); $^{13}$C NMR ($CDCl_3$, 126 MHz)

$\delta$ 177.5, 172.1, 171.8, 169.4, 160.3, 135.7, 130.2, 128.6, 128.3, 128.1, 127.6, 113.7, 101.4, 83.9, 78.5, 66.4, 55.3, 44.7, 40.6, 40.0, 38.9, 36.2, 35.1, 33.1, 25.6, 25.6, 21.9, 19.1; HRMS (ESI$^+$) *m/z* $[C_{32}H_{43}N_3O_8Na]^+$ calcd: 620.2948; found: 620.2934.

Synthesis of **5n**

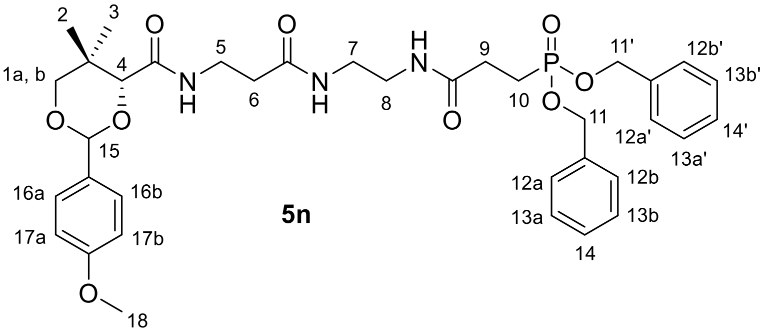

Compound **5n** was obtained following the general procedure B from compound **3**. To a solution of **4n** (200 mg, 0.60 mmol), NHS (69 mg, 0.60 mmol) and DCC (130 mg, 0.63 mmol) in dry THF (15 mL) was added a solution of compound **3** (227 mg, 0.60 mmol) in dry THF (6 mL) followed by $Et_3N$ (0.25 mL, 1.80 mmol). Product purification by column chromatography ($SiO_2$, DCM/MeOH = 95:5) afforded **5n** (290 mg, 70%) as a clear oil. $R_f$ = 0.50 (DCM/MeOH = 90:10); Purity 93%, RT = 15.32 min with method 7; RT = 27.91 min with method 8; $^1$H NMR ($CDCl_3$, 400 MHz) $\delta$ 7.43 (d, $J$ = 8.7 Hz, 2H, H-16a and H-16b), 7.37 – 7.31 (m, 10H, H-12a, 12a', 12b, 12b', 13a, 13a', 13b, 13b', 14 and 14'), 7.06 (t, $J$ = 5.0 Hz, 1H, -NH), 7.00 (t, $J$ = 6.4 Hz, 1H, -NH), 6.91 (d, $J$ = 8.7 Hz, 2H, H-17a and H-17b), 6.45 (t, $J$ = 5.8 Hz, 1H, -NH), 5.44 (s, 1H, H-15), 5.04 – 4.92 (m, 4H, H-11 and H-11'), 4.05 (s, 1H, H-4), 3.81 (s, 3H, H-18), 3.70 – 3.51 (m, 4H, H-1a, H-1b and H-5), 3.37-3.21 (m, 4H, H-7 and H-8), 2.49 – 2.27 (m, 4H, H-6 and H-9), 2.13 (dt, $J$ = 17.5, 7.3 Hz, 2H, H-10), 1.10 (s, 3H, H-2 or H-3), 1.06 (s, 3H, H-2 or H-3); $^{13}$C NMR ($CDCl_3$, 126 MHz) $\delta$ 171.6 (d, $J$ = 10.5 Hz), 171.6, 169.5, 160.3, 136.1 (d, $J$ = 5.8 Hz), 130.2, 128.7, 128.6, 128.0 (d, $J$ = 1.6 Hz), 127.6, 113.8, 101.4, 83.9, 78.5, 77.3, 77.1, 76.8, 67.5 (dd, $J$ = 6.4, 2.7 Hz), 55.4, 39.8, 39.1, 36.3, 35.1, 33.1, 29.2 (d, $J$ = 4.1 Hz), 21.9, 21.7 (d, $J$ = 142.6 Hz), 19.2; $^{31}$P NMR ($CDCl_3$, 203 MHz) $\delta$ 32.00; HRMS (ESI$^+$) *m/z* $[C_{36}H_{46}N_3O_9PNa]^+$ calcd: 718.2869; found: 718.2860.

Synthesis of **6m**

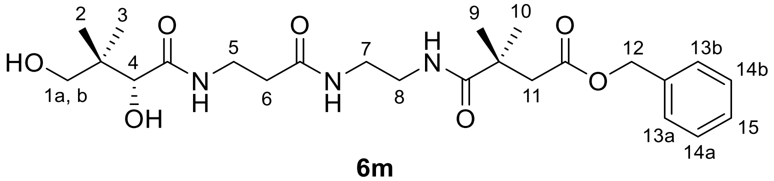

**6m**

Compound **6m** was obtained from compound **5m** using the general procedure C. Compound **5m** (255 mg, 0.43 mmol) was dissolved in a mixture of AcOH/$H_2O$ (9:1, 8 mL) and the solution was stirred at rt for 4 h. Purification by column chromatography ($SiO_2$, DCM/MeOH = 90:10) afforded **6m** (161 mg, 79%) as a clear oil. $R_f$ = 0.60 (DCM/MeOH = 85:15); Purity 89%, RT = 13.30 min with method 7; RT = 17.10 min with method 8; $^1$H NMR ($CD_3OD$, 500 MHz) $\delta$ 7.37 – 7.29 (m, 5H, H-13a, H-13b, H-14a, H-14b and H-15), 5.10 (s, 2H, H-12), 3.89 (s, 1H, H-4), 3.49 – 3.35 (m, 4H, H-1a, H-1b and H-5), 3.29 – 3.22 (m, 4H, H-7 and H-8), 2.66 (s, 2H, H-11), 2.40 (t, $J$ = 6.6 Hz, 2H, H-6), 1.24 (s, 6H, H-9 and H-10), 0.91 (s, 6H, H-2 and H-3); $^{13}$C NMR ($CD_3OD$, 126 MHz) $\delta$ 179.9, 176.0, 174.2, 173.1, 137.5, 129.5, 129.2, 77.3, 70.4, 67.3, 45.4, 41.8, 40.5, 40.4, 39.9, 36.6, 36.4, 25.9, 25.9, 21.3, 20.9; HRMS (ESI$^+$) *m/z* $[C_{24}H_{37}N_3O_7Na]^+$ calcd: 502.2529; found: 502.2520.

Synthesis of **6n**

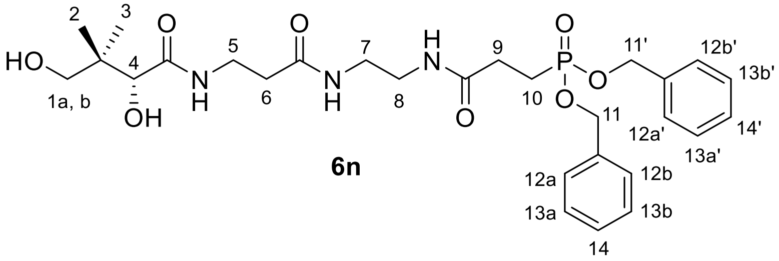

**6n**

Compound **6n** was obtained from compound **5n** using the general procedure C. Compound **5n** (104 mg, 0.15 mmol) was dissolved in a mixture of AcOH/$H_2O$ (9:1, 5 mL) and the solution was stirred at rt for 4 h. Purification by column chromatography ($SiO_2$, DCM/MeOH = 92:8) afforded **6n** (63 mg, 73%) as a gummy white solid. $R_f$ = 0.50 (DCM/MeOH = 90:10); Purity 95%, RT = 13.31 min with method 7; RT = 17.33 min with method 8; $^1$H NMR ($CDCl_3$, 400 MHz) $\delta$ 7.40 – 7.31 (m, 10H, H-12a, 12a', 12b, 12b', 13a, 13a', 13b, 13b', 14 and 14'), 7.06 (t, $J$ = 6.0 Hz, 1H, -NH), 6.51 (t, $J$ = 5.8 Hz, 1H, -NH), 5.13 – 4.86 (m, 5H, H-11, H-11' and -NH), 4.11 (bs, 1H, -OH), 4.00 (d, $J$ = 4.3 Hz, 1H, H-4), 3.73 – 3.09 (m, 8H, H-1a, H-1b, H-5, H-7 and H-8), 2.52 – 2.25 (m, 4H, H-6 and H-9), 2.12 (m, 2H, H-10), 1.01 (s, 3H, H-2 or H-3), 0.93 (s, 3H, H-2 or H-

3); $^{13}C$ NMR ($CDCl_3$, 126 MHz) $\delta$ 174.1, 172.4, 171.7 (d, $J$ = 11.2 Hz), 135.9 (d, $J$ = 2.9 Hz), 135.9 (d, $J$ = 3.1 Hz), 128.7, 128.7 (d, $J$ = 2.1 Hz), 128.0, 77.7, 70.9, 67.8 (d, $J$ = 6.7 Hz), 67.7 (d, $J$ = 6.6 Hz), 39.8, 39.3, 39.1, 35.9, 35.2, 29.0 (d, $J$ = 3.9 Hz), 22.01, 21.5, 20.9 , 20.6; $^{31}P$ NMR (203 MHz, $CD_3OD$) $\delta$ 32.70; HRMS ($ESI^+$) $m/z$ $[C_{28}H_{40}N_3O_8PNa]^+$ calcd: 600.2451; found: 600.2444.

Synthesis of **8k**

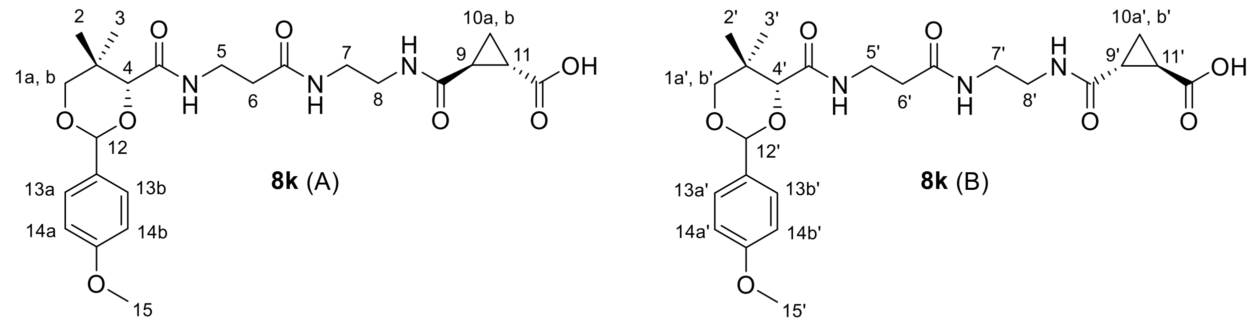

Compound **8k** was synthesized from **5k** (200 mg, 0.37 mmol), which was dissolved in THF: $H_2O$ (1:1, 3 mL). $LiOH.H_2O$ (24 mg, 0.56 mmol) was added to the solution before stirring at rt overnight. Excess THF was evaporated *in vacuo* below 40°C. The reaction mixture was cooled to 0°C and neutralized slowly with cold 0.5 N HCl (pH 5). The product was extracted in EA (3 × 25 mL), and the combined organic layers were dried over $Na_2SO_4$. The solvent was evaporated *in vacuo* and the product was purified from the residue by column chromatography ($SiO_2$, DCM/MeOH = 96:4) to afford **8k** (80 mg, 70%) as a white solid for this pair of diastereomers. $R_f$ = 0.40 (DCM/MeOH = 90:10); Purity >99%, RT = 16.92 min with method 7; RT = 30.74 min with method 8; $^1H$ NMR ($CDCl_3$, 500 MHz) $\delta$ 7.41 (d, $J$ = 8.5 Hz, 4H, H-13a, 13b, H-13a', H-13b'), 7.36 (bs, 2H, -NH and -NH'), 7.14 – 7.02 (m, 4H, -NH, -NH, -NH' and -NH'), 6.91 (m, 4H, H-14a, H-14b, H-14a' and H-14b'), 5.45 (s, 2H, H-12 and H-12'), 4.07 (s, 2H, H-4 and H-4'), 3.80 (s, 6H, H-15 and H-15'), 3.69 (d, $J$ = 11.4 Hz, 2H, H-1a and H-1b, or H-1a' and H-1b'), 3.64 (d, $J$ = 11.6 Hz, 2H, H-1a and H-1b, or H-1a' and H-1b'), 3.56 – 3.46 (m, 4H, H-5 and H-5'), 3.45 – 3.18 (m, 8H, H-7, H-8, H-7' and H-8'), 2.42 (t, $J$ = 6.5 Hz, 4H, H-6 and H-6'), 2.14 – 2.04 (m, 2H, H-11 and H-11'), 2.04 – 1.94 (m, 2H, H-9 and H-9'), 1.47 – 1.36 (m, 2H, H-10a and H-10a'), 1.35 – 1.25 (m, 2H, H-10b and H-10b'), 1.08 (s, 6H, H-2 or H-3, and H-2' or H-3'), 1.05 (s, 6H, H-2 or H-3, and H-2' or H-3'); $^{13}C$ NMR ($CDCl_3$, 126 MHz) $\delta$ 175.5, 172.2, 172.2,

171.6, 170.1, 160.3, 130.0, 127.6, 113.7, 101.3, 83.8, 78.3, 55.3, 39.6, 39.4, 39.4, 35.9, 35.9, 35.4, 33.1, 24.3, 21.8, 21.4, 19.1, 14.8; HRMS (ESI+) *m/z* $[C_{24}H_{33}N_3O_8Na_2]^+$ calcd: 537.2063; found: 537.1992.

Synthesis of **11**

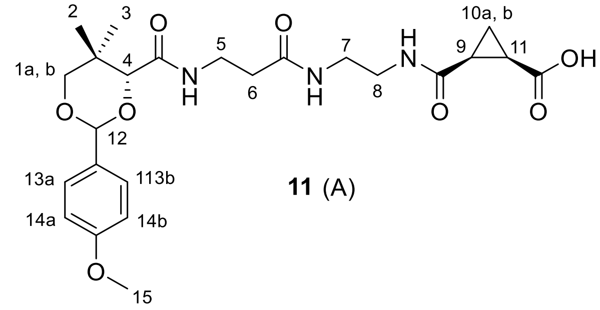

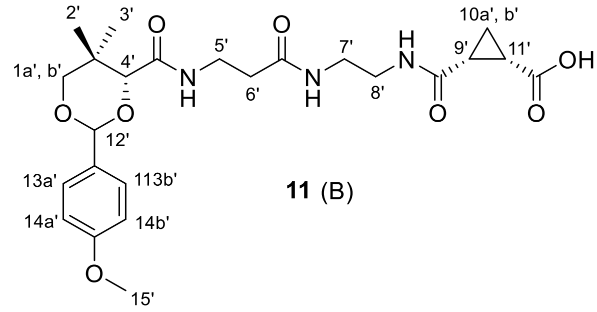

To a solution of compound **3** (150 mg, 0.40 mmol) in dry THF (3 mL) was added $Et_3N$ (0.06 mL, 0.40 mmol) before stirring at rt for 5 min. Then commercially available 3-oxabicyclo[3.1.0]hexane-2,4-dione (**4o**, 44 mg, 0.40 mmol) pre-dissolved in dry THF (2 mL) was added to the reaction mixture and the mixture was stirred at rt for 12 h. The solvent was evaporated *in vacuo*, and the product was purified by column chromatography ($SiO_2$, DCM/MeOH = 92:10) to afford compound **11** (80 mg, 41%) as a white solid for this pair of diastereomers. $R_f$ = 0.50 (DCM/MeOH = 85:15); Purity 99%, RT = 15.57 min with method 7; RT = 27.91 min with method 8; $^1$H NMR ($CDCl_3$, 300 MHz) $\delta$ 7.42 (d, $J$ = 8.2 Hz, 4H, H-13a, H-13b, H-13a'and H-13b'), 7.08 – 7.04 (m, 4H, -NH, -NH, –NH' and -NH'), 7.00 (bs, 2H, -NH and –NH'), 6.91 (d, $J$ = 8.4 Hz, 4H, H-14a, H-14b, H-14a', H-14b'), 5.48 – 5.46 (m, 2H, H-12 and H-12'), 4.09 (s, 2H, H-4 and H-4'), 3.82 (s, 6H, H-15 and H-15'), 3.73 – 3.63 (m, 4H, H-1a, H-1b, H-1a' and H-1b'), 3.55 – 3.18 (m, 12H, H-5, H-7, H-8, H-5', H-7' and H-8'), 2.58 – 2.31 (m, 4H, H-6 and H-6'), 2.12 – 1.98 (m, 4H, H-9, H-11, H-9' and H-11'), 1.63 – 1.55 (m, 2H, H-10a and H-10a'), 1.39 – 1.32 (m, 2H, 10b and 10b'), 1.09 (s, 6H, H-2 or H-3, and H-2' or H-3'), 1.06 (s, 6H, H-2 or H-3, and H-2' or H-3'); $^{13}$C NMR ($CDCl_3$, 75 MHz) $\delta$ 173.6, 173.6, 172.0, 171.5, 171.4, 170.3, 170.2, 160.2, 130.0, 127.5, 113.7, 101.3, 83.7, 83.6, 78.3, 55.3, 39.9, 38.9, 36.2, 36.1, 35.7, 35.6, 33.1, 24.3, 22.0, 21.8, 21.8, 19.1, 13.1, 13.1; HRMS (ESI+) *m/z* $[C_{24}H_{33}N_3O_8Na]^+$ calcd: 514.2165; found: 514.2157.

Synthesis of **13**

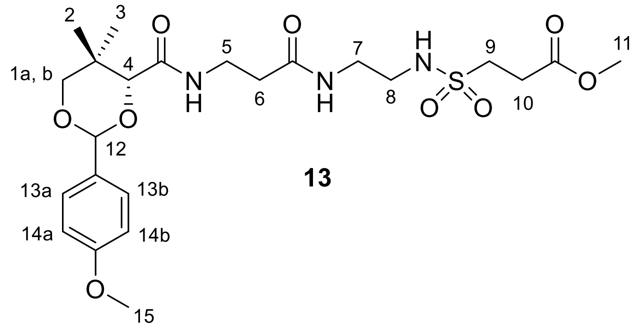

**13**

Compound **4p** (500 mg, 2.87 mmol) was dissolved in $POCl_3$ (2 mL) in a 10 mL round bottom flask under an argon atmosphere and refluxed for 4 h; then the reaction mixture was cooled to rt, diluted in DCM (5 mL), and eluted through a short silica pad which was washed with DCM (10 mL). The organic layers were combined and the solvent was evaporated *in vacuo* to afford the crude methyl 3-(chlorosulfonyl)propanoate (245 mg, 46%) as a light pink oil directly, which was used directly in the next step.

Compound **3** (500 mg, 1.32 mmol) was dissolved in dry DCM (10 mL), and $Et_3N$ (0.54 mL, 3.95 mmol) was added. The mixture was cooled to 0ºC. A solution of the crude methyl 3-(chlorosulfonyl)propanoate (245 mg, 1.32 mmol) in dry DCM (5 mL) was added dropwise to the reaction mixture before stirring at rt overnight. The reaction mixture was next concentrated *in vacuo* and the product was purified by column chromatography ($SiO_2$, DCM/MeOH = 97:3) to afford compound **13** (491 mg, 71%) as a sticky white solid. $R_f$ = 0.70 (DCM/MeOH = 90:10); Purity >99%, RT = 14.19 min with method 7; RT = 19.92 min with method 8; $^1$H NMR ($CDCl_3$, 400 MHz) $\delta$ 7.44 (m, 2H, H-13a and H-13b), 7.03 (t, $J$ = 6.4 Hz, 1H, -NH), 6.92 (m, 2H, H-14a and H-14b), 6.70 (bs, 1H, -NH), 5.60 (bs, 1H, -NH), 5.47 (s, 1H, H-12), 4.11 (s, 1H, H-4), 3.82 (s, 3H, H-15), 3.76 – 3.13 (m, 13H, H-1a, H-1b, H-5, H-7, H-8, H-9 and H-11), 2.79 (t, $J$ = 7.4 Hz, 2H, H-10), 2.44 (m, 2H, H-6), 1.10 (s, 3H, H-2 or H-3), 1.08 (s, 3H, H-2 or H-3); $^{13}$C NMR ($CDCl_3$, 126 MHz) $\delta$ 172.0, 171.2, 170.1, 160.3, 130.2, 127.6, 113.8, 101.4, 83.8, 78.4, 55.4, 52.4, 47.7, 42.9, 40.0, 36.5, 35.1, 33.1, 28.7, 21.8, 19.1; HRMS (ESI$^+$) *m/z* $[C_{23}H_{35}N_3O_9SNa]^+$ calcd: 552.1952; found: 552.1985.

Synthesis of **14**

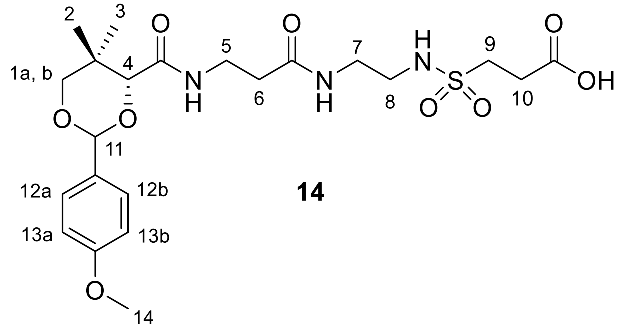

**14**

Compound **13** (320 mg, 0.60 mmol) was dissolved in THF: $H_2O$ (1:1, 12 mL) before adding LiOH.$H_2O$ (38 mg, 0.91 mmol) and stirring the mixture at rt overnight. Excess THF was evaporated *in vacuo* below 40ºC. The reaction mixture was next cooled to 0ºC and neutralized slowly with cold 0.5 N HCl (pH 5). The product was extracted in EA (25 mL × 3). The combined organic layers were dried over $Na_2SO_4$ and concentrated *in vacuo.* Purification by column chromatography ($SiO_2$, DCM/MeOH = 90:10) afforded the product as a white solid (272 mg, 87%). $R_f$ = 0.30 (DCM/MeOH = 90:10); Purity 97%, RT = 15.99 min with method 7; RT = 27.79 min with method 8; $^1$H NMR ($CDCl_3$, 400 MHz) $\delta$ 7.42 (d, $J$ = 8.7 Hz, 2H, H-12a and H-12b), 7.14 (t, $J$ = 6.2 Hz, 1H, -NH), 6.93-6.90 (m, 3H, H-13a, H-13b and -NH), 5.62 (bs, 1H, -NH), 5.46 (s, 1H, H-11), 4.11 (s, 1H, H-4), 3.81 (s, 3H, H-14), 3.71-3.63 (m, 2H, H-1a and H-1b), 3.52 (m, 2H, H-5), 3.42 – 3.28 (m, 4H, H-7 and H-8), 3.19 (m, 2H, H-9), 2.78 (t, $J$ = 6.0 Hz, 2H, H-10), 2.47 (m, 2H, H-6), 1.09 (s, 3H, H-2 or H-3), 1.05 (s, 3H, H-2 or H-3); $^{13}$C NMR ($CDCl_3$, 126 MHz) $\delta$ 173.4, 172.3, 170.6, 160.3, 130.1, 127.6, 113.8, 101.3, 83.7, 78.3, 55.4, 47.2, 42.6, 39.7, 35.9, 35.6, 33.1, 28.8, 21.8, 19.1; HRMS (ESI$^+$) *m/z* $[C_{22}H_{33}N_3O_9SNa]^+$ calcd: 538.1835; found: 538.1808.

Synthesis of **21a**

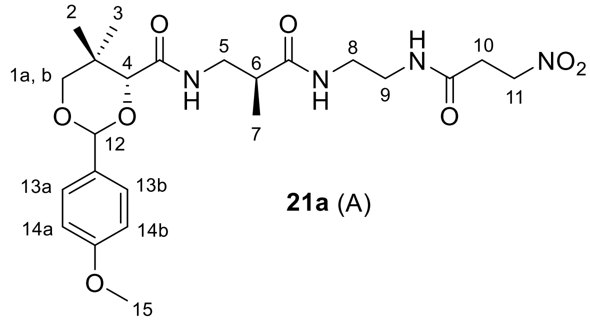

**21a** (A)

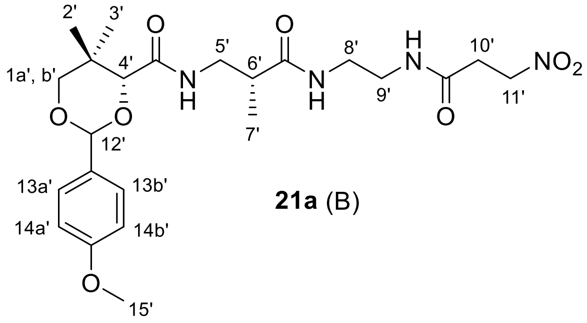

**21a** (B)

Compound **21a** was obtained following the general procedure B from compound **20a**. To a solution of **4a** (100 mg, 0.84 mmol), NHS (97 mg, 0.84 mmol) and DCC (182 mg, 0.88 mmol) in dry THF (15 mL) was added a solution of **20a** (330 mg, 0.88 mmol) in dry THF (10 mL), followed by $Et_3N$ (0.3 mL, 2.10 mmol). Purification by column chromatography ($SiO_2$, DCM/MeOH = 96:4) afforded **21a** (195 mg, 47%) as a yellow oil for this pair of diastereomers. $R_f$ = 0.60 (DCM/MeOH = 90:10); Purity 96%, RT = 14.15 min with method 7; RT = 19.67 min with method 8; $^1$H NMR ($CDCl_3$, 500 MHz) $\delta$ 7.46 – 7.39 (m, 4H, H-13a, H-13b, H-13a' and H-13b'), 7.00 – 6.89 (m, 7H, H-14a, H-14b and H-14a', H-14b', -NH, -NH' and -NH or -NH' ), 6.85 – 6.77 (m, 1H, -NH or –NH'), 6.46 – 6.38 (m, 2H, -NH, -NH'), 5.47 (s, 1H, H-12 or H-12'), 5.46 (s, 1H, H-12 or H-12'), 4.69 – 4.62 (m, 4H, H-11 and H-11'), 4.09 (s, 1H, H-4 or H-4'), 4.08 (s, 1H, H-4 or H-4'), 3.82 (s, 3H, H-15 or H-15'), 3.81 (s, 3H, H-15 or H-15'), 3.75 – 3.68 (m, 2H, H-1a and H-1a'), 3.66 (d, $J$ = 11.5 Hz, 2H, H-1b and H-1b'), 3.49 – 3.22 (m, 12H, H-5, H-8, H-9, H-5', H-8' and H-9'), 2.81 – 2.69 (m, 4H, H-10 and H-10'), 2.58 – 2.47 (m, 2H, H-6 and H-6'), 1.18 – 1.09 (m, 12H, H-7, H-7', H-2 or H-2' and H-3 or H-3'), 1.08 (s, 6H, H-2 or H-2' and H-3 or H-3'); $^{13}$C NMR ($CDCl_3$,126 MHz) $\delta$ 175.8, 169.8, 169.7, 169.2, 169.2, 169.2, 160.3, 130.2, 130.1, 127.6, 127.5, 113.8, 101.3, 101.3, 83.9, 83.9, 78.4, 70.3, 55.4, 41.9, 41.9, 41.0, 41.0, 40.7, 40.7, 40.1, 40.1, 40.0, 40.0, 39.4, 39.3, 33.1, 32.4, 32.4, 21.8, 19.2, 19.1, 15.6, 15.4; HRMS (ESI$^+$) $m/z$ $[C_{23}H_{34}N_4O_8Na]^+$ calcd: 517.2274; found: 517.2147.

Synthesis of **21b**

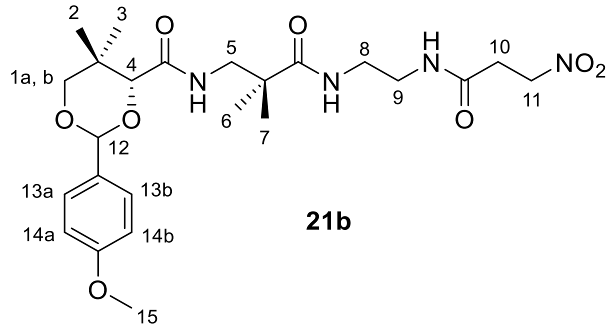

**21b**

Compound **21b** was obtained following the general procedure B from compound **20b**. To a solution of **4a** (123 mg, 1.03 mmol), NHS (119 mg, 1.03 mmol) and DCC (224 mg, 1.09 mmol) in dry THF (12 mL) was added a solution of **20b** (420 mg, 1.034 mmol) in dry THF (6 mL) followed by addition of $Et_3N$ (0.15 mL, 1.03 mmol). Purification by column chromatography

($SiO_2$, DCM/MeOH = 97:3 to 96:4) afforded **21b** (296 mg, 56%) as a white sticky solid. $R_f$ = 0.60 (DCM/MeOH = 90:10); Purity 99%, RT = 14.61 min with method 7; RT = 21.69 min with method 8; $^1$H NMR ($CDCl_3$, 500 MHz) $\delta$ 7.44 (m, 2H, H-13a and H-13b), 7.03 (t, $J$ = 6.8 Hz, 1H, -NH), 6.93 (m, 2H, H-14a and H-14b), 6.88 (bs, 1H, -NH), 6.67 (t, $J$ = 4.7 Hz, 1H, -NH), 5.48 (s, 1H, H-12), 4.67 (m, 2H, H-11), 4.10 (s, 1H, H-4), 3.82 (s, 3H, H-15), 3.71 (d, $J$ = 11.4 Hz, 1H, H-1a), 3.66 (d, $J$ = 11.4 Hz, 1H, H-1b), 3.45 – 3.24 (m, 6H, H-5, H-8 and H-9), 2.73 (m, 2H, H-10), 1.18 (s, 6H, H-6 and H-7), 1.12 (s, 3H, H-2 or H-3), 1.08 (s, 3H, H-2 or H-3); $^{13}$C NMR ($CDCl_3$, 126 MHz) $\delta$ 177.5, 169.8, 169.3, 160.3, 130.2, 127.5, 113.8, 101.2, 83.9, 78.4, 70.3, 55.4, 46.9, 43.1, 40.0, 39.8, 33.1, 32.5, 23.9, 23.7, 21.9, 19.2; HRMS (ESI$^+$) *m/z* $[C_{24}H_{36}N_4O_8Na]^+$ calcd: 531.2431; found: 531.2411.

Synthesis of **21c**

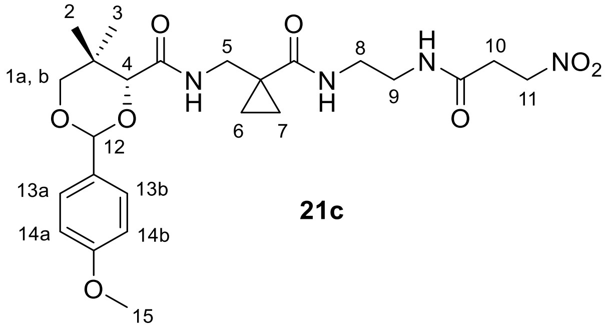

**21c**

Compound **21c** was obtained following the general procedure B from compound **20c**. To a solution of **4a** (62 mg, 0.52 mmol), NHS (60 mg, 0.52 mmol) and DCC (112 mg, 0.54 mmol) in dry THF (10 mL) was added a solution of compound **3** (210 mg, 0.52 mmol) in dry THF (4 mL), followed by addition of $Et_3N$ (0.10 mL, 0.52 mmol). Purification by column chromatography ($SiO_2$, DCM/MeOH = 97:3) afforded **21c** (173 mg, 66%) as a white solid. $R_f$ = 0.60 (DCM/MeOH = 90:10); Purity 91%, RT = 14.22 min with method 7; RT = 15.07 min with method 8; $^1$H NMR ($CDCl_3$, 500 MHz) $\delta$ 8.12 (bs, 1H, -NH), 7.43 (m, 2H, H-13a and H-13b), 7.03 (bs, 1H, -NH), 6.96 (t, $J$ = 6.7 Hz, 1H, -NH), 6.93 (m, 2H, H-14a and H-14b), 5.49 (s, 1H, H-12), 4.67 (m, 2H, H-11), 4.15 (s, 1H, H-4), 3.83 (s, 3H, H-15), 3.74 (d, $J$ = 11.5 Hz, 1H, H-1a or H-1b), 3.68 (d, $J$ = 11.5 Hz, 1H, H-1a or H-1b), 3.54 – 3.37 (m, 4H, H-8 and H-9), 3.33 (m, 2H, H-5), 2.79 (m, 2H, H-10), 1.27 (m, 2H, H-6 or H-7), 1.09 (s, 3H, H-2 or H-3), 1.09 (s, 3H, H-2 or H-3), 0.74 – 0.66 (m, 2H, H-6 or H-7); $^{13}$C NMR ($CDCl_3$, 126 MHz) $\delta$ 174.1, 171.0,

168.6, 160.5, 130.0, 127.7, 113.9, 101.6, 83.8, 78.4, 70.4, 55.5, 42.7, 41.3, 39.6, 33.3, 32.7, 25.6, 22.0, 19.2, 15.0, 14.8; HRMS (ESI$^+$) *m/z* $[C_{24}H_{34}N_4O_8Na]^+$ calcd: 529.2274; found: 529.2250.

Synthesis of **6a (RK-03-142) (1-01)**

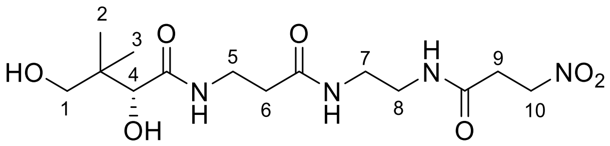

**6a**

Compound **6a** was obtained from compound **5a** using the general procedure C. Compound **5a** (1.4 g, 2.91 mmol) was dissolved in a mixture of AcOH/$H_2O$ (9:1, 25 mL) before stirring at rt for 4 h. Purification by column chromatography ($SiO_2$, DCM/MeOH = 90:10) afforded **6a** (718 mg, 68%) as a viscous, yellowish oil. $R_f$ = 0.30 (DCM/MeOH = 90:10); Purity >99%, RT = 14.66 min with method 7; RT = 14.05 min with method 8; $^1$H NMR ($CD_3OD$, 500 MHz) $\delta$ 4.72 (m, 2H, H-10), 4.59 (bs, 1H, -OH), 3.90 (s, 1H, H-4), 3.54 – 3.35 (m, 4H, H-1, H-5), 3.31 – 3.25 (m, 4H, H-7 and H-8), 2.88 (m, 2H, H-9), 2.42 (t, $J$ = 6.6 Hz, 2H, H-6), 0.93 (s, 6H, H-2 and H-3).

Synthesis of **6b (RK-01-031) (4-01)**

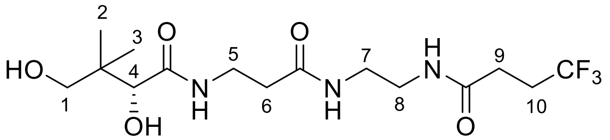

**6b**

Compound **6b** was obtained from compound **5b** using the general procedure C. Compound **5b** (100 mg, 0.2 mmol) was dissolved in a mixture of AcOH/$H_2O$ (9:1, 5 mL) and the solution was stirred at rt for 4 h. Purification by column chromatography ($SiO_2$, DCM/MeOH = 92:8) afforded **6b** (53 mg, 69%) as a sticky white solid. $R_f$ = 0.30 (DCM/MeOH = 85:15, visualized by $KMnO_4$ stain); Purity 98%, RT = 12.59 min with method 7; RT = 15.11 min with method 8; $^1$H NMR ($CD_3OD$, 300 MHz) $\delta$ 3.89 (s, 1H, H-4), 3.55 – 3.35 (m, 4H, H-7, H-8), 3.29 – 3.19 (m, 4H, H-1, H-5 ), 2.60 – 2.33 (m, 6H, H-6, H-9 and H-10), 0.92 (s, 6H, H-2, H-3); $^{13}$C NMR ($CD_3OD$, 75 MHz) $\delta$ 174.7, 172.7, 171.7, 128.9-125.2 (m), 75.9, 68.9, 38.9, 38.7, 38.5, 35.2, 35.0, 29.0 (q, $J$ = 29.5 Hz), 27.9 (q, $J$ = 3.1 Hz), 19.9, 19.5; $^{19}$F NMR ($CD_3OD$, 282 MHz) $\delta$ -68.45 (t, $J$ = 10.2 Hz); HRMS (ESI$^+$) *m/z* $[C_{15}H_{26}F_3N_3O_5Na]^+$ calcd: 408.1722; found: 408.1708.

Synthesis of **6c (RK-01-32) (4-02)**

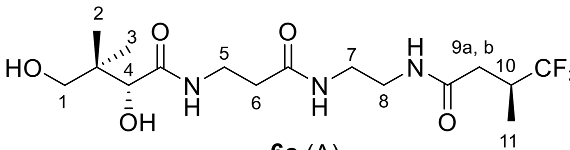

**6c** (A)

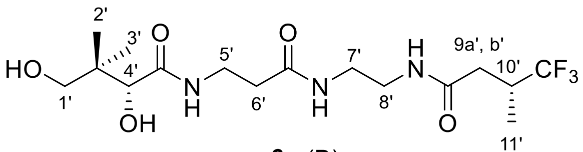

**6c** (B)

Compound **6c** was obtained from compound **5c** using the general procedure C. Compound **5c** (100 mg, 0.19 mmol) was dissolved in a mixture of AcOH/$H_2O$ (9:1 v/v, 5 mL) and the solution was stirred at rt for 4 h. Purification by column chromatography ($SiO_2$, DCM/MeOH = 92:8) afforded **6c** (31 mg, 40%) as a sticky white solid for this pair of diastereomers. $R_f$ = 0.50 (DCM/MeOH = 85:15, visualized by $KMnO_4$ stain); Purity 95%, RT = 12.61 min with method 7; RT = 15.60 min with method 8; $^1$H NMR ($CD_3OD$, 300 MHz) $\delta$ 3.86 (s, 2H, H-4 and H-4'), 3.54 – 3.33 (m, 8H, H-7, H-8, H-7' and H-8'), 3.27 – 3.15 (m, 8H, H-1, H-5, H-1' and H-5'), 2.88 – 2.63 (m, 2H, H-10 and H-10'), 2.62 – 2.47 (m, 2H, H-9a and H-9a'), 2.45 – 2.31 (m, 4H, H-6 and H-6'), 2.25 – 2.06 (m, 2H, H-9b and H-9b'), 1.21 – 1.02 (m, 6H, H-11 and H-11'), 0.89 (s, 12H, H-2, H-3, H-2' and H-3'); $^{13}$C NMR (75 MHz, $CD_3OD$) $\delta$ 174.6, 172.7, 171.3, 129.9, 126.2, 75.9, 68.9, 38.9, 38.6, 38.6, 35.6, 35.6, 35.5, 35.2, 35.2, 35.0, 34.9, 34.5, 19.9, 19.5, 11.7, 11.7; $^{19}$F NMR (282 MHz, $CD_3OD$) $\delta$ -75.25 (d, $J$ = 8.9 Hz); HRMS (ESI$^+$) *m/z* $[C_{16}H_{28}F_3N_3O_5Na]^+$ calcd: 422.1879; found: 422.1868.

Synthesis of **6d (RK-01-034) (4-03)**

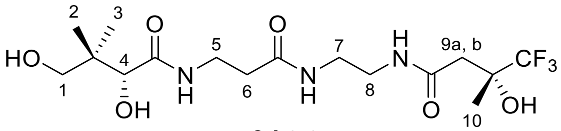

**6d** (A)

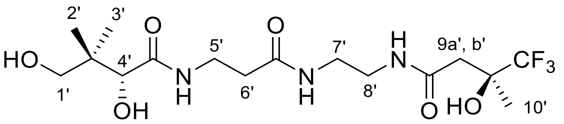

**6d** (B)

Compound **6d** was obtained from compound **5d** using the general procedure C. Compound **5d** (116 mg, 0.22 mmol) was dissolved in a mixture of AcOH/$H_2O$ (9:1, 6 mL) and the solution was stirred at rt for 4 h. Purification by column chromatography ($SiO_2$, DCM/MeOH = 90:10) afforded **6d** (52 mg, 57%) as a sticky white solid for this pair of diastereomers. $R_f$ = 0.30 (DCM/MeOH = 85:15, visualized by $KMnO_4$ stain); Purity 96%, RT = 12.54 min with method 7; RT = 14.89 min with method 8; $^1$H NMR ($CD_3OD$, 500 MHz) $\delta$ 3.91 (s, 2H, H-4 and H-4'), 3.53 – 3.33 (m, 12H, H-1, H-7, H-8, H-1', H-7' and H-8'), 3.31 – 3.27 (m, 4H, H-5 and H-5'), 2.64 – 2.61 (m, 2H, H-9a and H-9a'), 2.45 – 2.42 (m, 6H, H-H-9b, H-9b', H-6 and H-6'), 1.44 (s, 6H, H-10 and H-10'), 0.94 (s, 12H, H-2, H-3, H-2' and H-3'); $^{13}$C NMR ($CD_3OD$, 75 MHz) $\delta$ 174.7,

172.8, 170.7, 126.1 (q, $J$ = 285.0 Hz), 75.9, 72.0 (q, $J$ = 28.5 Hz), 68.9, 40.1, 39.0, 38.5, 38.5, 35.2, 35.0, 20.0, 19.7, 19.7, 19.5; $^{19}$F NMR ($CD_3OD$, 282 MHz) $\delta$ -84.61; HRMS (ESI$^+$) *m/z* $[C_{16}H_{29}F_3N_3O_6]^+$ calcd: 416.2003; found: 416.2002.

Synthesis of **6e (RK-04-016) (6-05)**

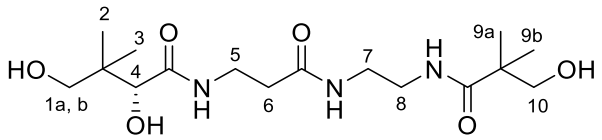

**6e**

Compound **6e** was obtained from compound **5e** using the general procedure C. Compound **5e** (90 mg, 0.23 mmol) was dissolved in a mixture of AcOH/$H_2O$ (9:1, 6 mL) and the solution was stirred at rt for 4 h. Purification by column chromatography ($SiO_2$, DCM/MeOH = 88:12) afforded **6d** (59 mg, 86%) as a sticky white solid. $R_f$ = 0.30 (DCM/MeOH = 85:15); Purity 97%, RT = 12.35 min with method 7; RT = 13.75 min with method 8; $^1$H NMR ($CD_3OD$, 500 MHz) $\delta$ 3.89 (s, 1H, H-4), 3.51 (s, 2H, H-10), 3.49 – 3.43 (m, 3H, H-1a and H-5), 3.39 (d, $J$ = 11.0 Hz, 1H, H-1b), 3.30 – 3.25 (m, 4H, H-7 and H-8), 2.41 (t, $J$ = 6.7 Hz, 2H, H-6), 1.13 (s, 6H, H-9a and H-9b), 0.92 (s, 6H, H-2 and H-3); $^{13}$C NMR (126 MHz, $CD_3OD$) $\delta$ 180.1, 176.1, 174.3, 77.3, 70.3, 70.3, 45.0, 40.4, 40.4, 40.0, 36.6, 36.4, 22.7, 21.3, 20.9; HRMS (ESI$^+$) *m/z* $[C_{16}H_{31}N_3O_6Na]^+$ calcd: 384.2111; found: 384.2103.

Synthesis of **6f (RK-03-020) (6-09)**

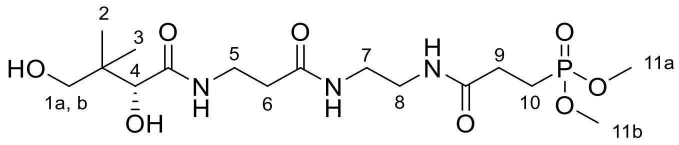

**6f**

Compound **6f** was obtained from compound **5f** using the general procedure C. Compound **5f** (140 mg, 0.26 mmol) was dissolved in a mixture of AcOH/$H_2O$ (9:1, 10 mL) and the solution was stirred at rt for 4 h. Purification by column chromatography ($SiO_2$, DCM/MeOH = 90:10) afforded **6f** (96 mg, 88%) as a white solid. $R_f$ = 0.30 (DCM/MeOH = 85:15, visualized by $KMnO_4$ stain); Purity >99%, RT = 12.36 min with method 7; RT = 13.83 min with method 8; $^1$H NMR ($CD_3OD$, 500 MHz) $\delta$ 3.91 (s, 1H, H-4), 3.78 (s, 3H, H-11a or H-11b), 3.76 (s, 3H, H-11a or H-11b), 3.54 – 3.46 (m, 3H, H-1a and H-5), 3.41 (d, $J$ = 10.9 Hz, 1H, H-1b), 3.30 – 3.25 (m,

4H, H-7, H-8), 2.50 – 2.42 (m, 4H, H-6, H-9), 2.15 (m, 2H, H-10), 0.94 (s, 6H, H-2 and H-3); $^{13}$C NMR ($CD_3OD$, 126 MHz) $\delta$ 174.6, 172.7, 172.5 (d, $J$ = 15.2 Hz), 76.0, 68.9, 52.0 (d, $J$ = 6.6 Hz), 39.0, 38.8, 38.6, 35.2, 35.0, 28.1 (d, $J$ = 4.1 Hz), 20.0, 19.5, 19.3 (d, $J$ = 143.7 Hz); $^{31}$P NMR ($CD_3OD$, 203 MHz) $\delta$ 34.60; HRMS (ESI$^+$) $m/z$ $[C_{16}H_{32}N_3O_8PNa]^+$ calcd: 448.1825; found: 448.1800.

Synthesis of **6g (RK-03-097) (6-10)**

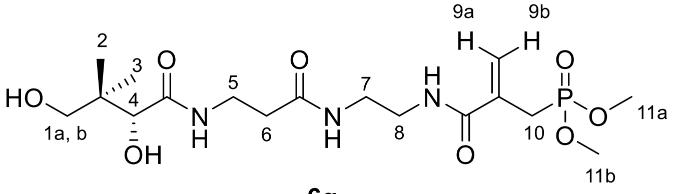

**6g**

Compound **6g** was obtained from compound **5g** using the general procedure C. Compound **5g** (110 mg, 0.20 mmol) was dissolved in a mixture of AcOH/$H_2O$ (9:1, 10 mL) and the solution was stirred at rt for 4 h. Purification by column chromatography ($SiO_2$, DCM/MeOH = 90:10) afforded **6g** (68 mg, 79%) as a clear oil. $R_f$ = 0.20 (DCM/MeOH = 85:15); Purity 98%, RT = 12.29 min with method 7; RT = 13.84 min with method 8; $^1$H NMR ($CD_3OD$, 500 MHz) $\delta$ 5.90 (d, $J$ = 5.4 Hz, 1H, H-9a), 5.64 (m, 1H, H-9b), 3.91 (s, 1H, H-4), 3.78 (s, 3H, H-11a or H-11b), 3.75 (s, 3H, H-11a or H-11b), 3.55 – 3.34 (m, 8H, H-1a, H-1b, H-5, H-7 and H-8), 3.07 (dd, $J$ = 21.9, 0.6 Hz, 2H, H-10), 2.45 (t, $J$ = 6.6 Hz, 2H, 2H, H-6), 0.94 (s, 6H, H-2 and H-3); $^{13}$C NMR ($CD_3OD$, 126 MHz) $\delta$ 176.0, 174.2, 170.0 (d, $J$ = 3.6 Hz), 136.7 (d, $J$ = 10.2 Hz), 123.4 (d, $J$ = 11.3 Hz), 77.3, 70.3, 53.7 (d, $J$ = 6.8 Hz), 40.5, 40.3, 39.9, 36.6, 36.4, 28.8 (d, $J$ = 139.2 Hz), 21.3, 20.9; $^{31}$P NMR ($CD_3OD$, 203 MHz) $\delta$ 29.39; HRMS (ESI$^+$) $m/z$ $[C_{17}H_{32}N_3O_8PNa]^+$ calcd: 460.1825; found: 460.1812.

Synthesis of **6h (RK-01-111) (1-09)**

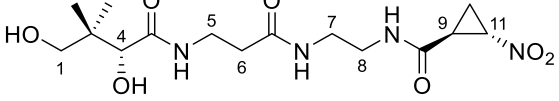

**6h** (A)

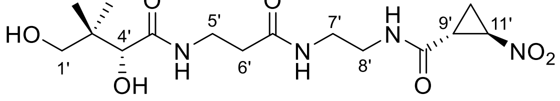

**6h** (B)

Compound **6h** was obtained from compound **5h** using the general procedure C. Compound **5h** (160 mg, 0.34 mmol) was dissolved in a mixture of AcOH/$H_2O$ (9:1, 8 mL) and the solution was stirred at rt for 4 h. Purification by column chromatography ($SiO_2$, DCM/MeOH = 92:8) afforded **6h** (114 mg, 93%) as a white solid for this pair of diastereomers. $R_f$ = 0.30 (DCM/MeOH = 85:15, visualized by $KMnO_4$ stain); Purity 96%, RT = 12.55 min with method 7; RT = 14.64 min with method 8; $^{1}$H NMR ($CDCl_3$, 300 MHz) $\delta$ 4.62 – 4.49 (m, 2H, H-11 and H-11'), 3.89 (s, 2H, H-4 and H-4'), 3.56 – 3.32 (m, 12H, H-1, H-5, H-7 or H-8, H-1', H-5' and H-7' or H-8'), 3.29 – 3.22 (m, 4H, H-7 or H-8 and, H-7' or H-8'), 2.79 – 2.65 (m, 2H, H-9 and H-9'), 2.47 – 2.34 (m, 4H, H-6 and H-6'), 2.03 – 1.90 (m, 2H, H-10a and H-10a'), 1.74 – 1.61 (m, 2H, H-10b and H-10b'), 0.92 (s, 12H, H-2, H-3, H-2' and H-3'); $^{13}$C NMR ($CD_3OD$, 126 MHz) $\delta$ 174.7, 172.8, 168.8, 76.0, 68.9, 58.6, 39.0, 38.5, 35.3, 35.0, 25.9, 20.0, 19.5, 15.5; HRMS (ESI$^+$) *m/z* $[C_{15}H_{26}N_4O_7Na]^+$ calcd: 397.1699; found: 397.1683.

<u>Synthesis of **6i (RK-04-015) (1-04)**</u>

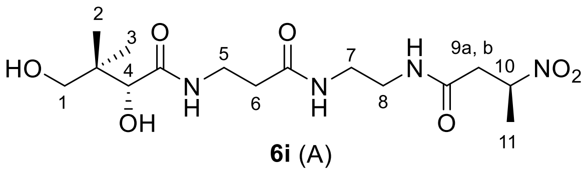

**6i** (A)

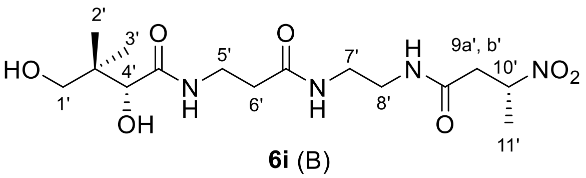

**6i** (B)

Compound **6i** was obtained from compound **5i** using the general procedure C. Compound **5i** (185 mg, 0.37 mmol) was dissolved in a mixture of AcOH/$H_2O$ (9:1, 10 mL) and the solution was stirred at rt for 4 h. Purification by column chromatography ($SiO_2$, DCM/MeOH = 90:10) afforded **6i** (131 mg, 93%) as a white solid for this pair of diastereomers. $R_f$ = 0.40 (DCM/MeOH = 85:15, visualized by $KMnO_4$ stain); Purity 99%, RT = 12.49 min with method 7; RT = 14.52 min with method 8; $^{1}$H NMR ($CD_3OD$, 500 MHz) $\delta$ 4.98 (dqd, $J$ = 8.9, 6.8, 4.8 Hz, 2H, H-10 and H-10'), 3.89 (s, 2H, H-4 and H-4'), 3.53 – 3.37 (m, 8H, H-1, H-5, H-1' and H-5'), 3.30 – 3.20 (m, 8H, H-7, H-8, H-7' and H-8'), 2.98 (dd, $J$ = 15.9, 8.9 Hz, 2H, H-9a and H-9a'), 2.66 (ddd, $J$ = 15.9, 4.9, 0.9 Hz, 2H, H-9b and H-9b'), 2.41 (t, $J$ = 6.6 Hz, 4H, H-6 and H-6'), 1.56 (d, $J$ = 6.9 Hz, 6H, H-11 and H-11'), 0.92 (s, 12H, H-2, H-3, H-2' and H-3'); $^{13}$C NMR ($CD_3OD$, 126 MHz) $\delta$ 176.0, 174.2, 171.5, 80.5, 77.3, 70.3, 40.8, 40.4, 40.0, 39.9, 36.6, 36.4, 21.4, 20.9, 19.7; HRMS (ESI$^+$) *m/z* $[C_{15}H_{28}N_4O_7Na]^+$ calcd: 399.1856; found: 399.1865.

<u>Synthesis of **6k (RK-04-003) (2-09)**</u>

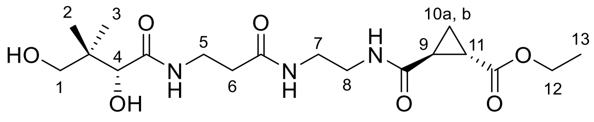

**6k** (A)

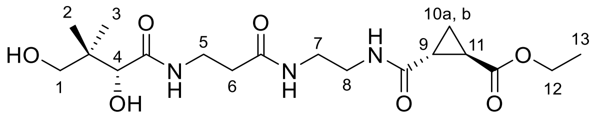

**6k** (B)

Compound **6k** was obtained from compound **5k** using the general procedure C. Compound **5k** (90 mg, 0.17 mmol) was dissolved in a mixture of AcOH/$H_2O$ (9:1, 6 mL) and the solution was stirred at rt for 4 h. Purification by column chromatography ($SiO_2$, DCM/MeOH = 90:10) afforded **6k** (58 mg, 83%) as a sticky white solid for this pair of diastereomers. $R_f$ = 0.50 (DCM/MeOH = 85:15, visualized by $KMnO_4$ stain); Purity 97%, RT = 12.52 min with method 7; RT = 15.04 min with method 8; $^{1}$H NMR ($CD_3OD$, 500 MHz) $\delta$ 4.15 (q, $J$ = 7.1 Hz, 4H, H-12 and H-12'), 3.91 (s, 2H, H-4 and H-4'), 3.54 – 3.40 (m, 8H, H-1, H-5, H-1' and H-5'), 3.35 – 3.27 (m, 8H, H-7, H-8, H-7' and H-8'), 2.43 (t, $J$ = 6.7 Hz, 4H, H-6 and H-6'), 2.12 (ddd, $J$ = 8.7, 5.8, 3.8 Hz, 2H, H-11 and H-11'), 2.04 (ddd, $J$ = 8.7, 5.7, 3.8 Hz, 2H, H-9 and H-9'), 1.42 – 1.29 (m, 4H, H-10a, H-10b, H-10a' and H-10b'), 1.27 (t, $J$ = 7.1 Hz, 6H, H-13 and H-13'), 0.94 (s, 12H, H-2, H-3, H-2' and H-3'); $^{13}$C NMR ($CD_3OD$, 126 MHz) $\delta$ 176.0, 174.1, 173.9, 173.1, 77.3, 70.3, 62.1, 40.3, 40.2, 40.1, 36.6, 36.4, 24.6, 22.2, 21.4, 20.9, 14.8, 14.5; HRMS (ESI$^+$) *m/z* $[C_{18}H_{31}N_3O_7Na]^+$ calcd: 424.2060; found: 424.2033.

<u>Synthesis of **6l (RK-03-057) (6-03)**</u>

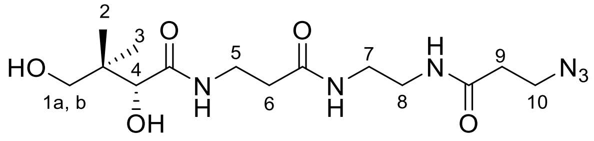

**6l**

Compound **6l** was obtained from compound **5l** using the general procedure C. Compound **5l** (130 mg, 0.27 mmol) was dissolved in a mixture of AcOH/$H_2O$ (9:1, 5 mL) and the solution was stirred at rt for 4 h. Purification by column chromatography ($SiO_2$, DCM/MeOH = 92:8) afforded **6l** (96 mg, 99%) as an off-white solid. $R_f$ = 0.50 (DCM/MeOH = 85:15, visualized by $KMnO_4$ stain); Purity 97%, RT = 15.47 min with method 9; RT = 10.79 min with method 11; $^{1}$H NMR ($CD_3OD$, 500 MHz) $\delta$ 3.92 (s, 1H, H-4), 3.59 (t, $J$ = 6.5 Hz, 2H, H-5), 3.56 – 3.39 (m, 4H, H-1a, H-1b and H-10), 3.36 – 3.27 (m, 4H, H-7 and H-8), 2.48 (t, $J$ = 6.5 Hz, 2H, H-6), 2.44 (t, $J$ = 6.7 Hz, 2H, H-9), 0.95 (s, 6H, H-2 and H-3); $^{13}$C NMR ($CD_3OD$, 126 MHz) $\delta$ 174.7, 172.8, 171.9,

75.9, 68.9, 39.0, 38.6, 38.6, 35.2, 35.0, 34.9, 20.0, 19.5; HRMS (ESI$^+$) *m/z* [$C_{14}H_{26}N_6O_5Na$]$^+$ calcd: 381.1862; found: 381.1872.

Synthesis of **7m (RK-04-090) (2-02)**

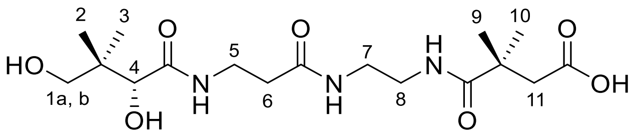

**7m**

Compound **7m** was obtained from compound **6m** using the general procedure D. Compound **6m** (150 mg, 0.31 mmol) was stirred with Pd/C (15 mg) in MeOH (20 mL) at rt and $H_2$ gas was purged in the mixture for 2 h. Purification by column chromatography ($SiO_2$, DCM/MeOH = 88:12) afforded **7m** (107 mg, 88%) as a white solid. $R_f$ = 0.20 (DCM/MeOH = 85:15, visualized by $KMnO_4$ stain); Purity 95%, RT = 12.66 min with method 7; RT = 15.50 min with method 8; $^1$H NMR ($CD_3OD$, 500 MHz) $\delta$ 3.89 (s, 1H, H-4), 3.52 – 3.47 (m, 3H, H-1a, H-5), 3.39 (d, *J* = 11.0 Hz, 1H, H-1b), 3.33 – 3.27 (m, 4H, H-7, H-8), 2.57 (s, 2H, H-11), 2.42 (t, *J* = 6.6 Hz, 2H, H-6), 1.25 (s, 6H, H-9, H-10), 0.92 (s, 6H, H-2, H-3); $^{13}$C NMR (126 MHz, $CD_3OD$) $\delta$ 180.4, 176.0, 175.7, 174.2, 77.3, 70.3, 45.8, 41.6, 40.3, 40.3, 39.9, 36.6, 36.4, 26.0, 26.0, 21.3, 20.9; HRMS (ESI$^+$) *m/z* [$C_{17}H_{31}N_3O_7Na$]$^+$ calcd: 412.2060; found: 412.2046.

Synthesis of **7n (RK-02-128) (6-08)**

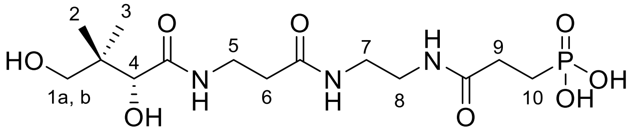

**7n**

Compound **7n** was obtained from compound **6n** using the general procedure D. Compound **6n** (62 mg, 0.11 mmol) was dissolved in MeOH (5 mL), before adding Pd/C (8 mg) and stirring at rt for 2 h while purging $H_2$ gas into the mixture. The reaction mixture was filtered through a celite pad, and the filtrate was concentrated *in vacuo* to obtain **7n** (42 mg, quantitative) as a white solid. $R_f$ = 0.10 (DCM/MeOH = 85:15, visualized by $KMnO_4$ stain); HPLC purity 89%, RT = 6.00 min with method 7; $^1$H NMR ($CD_3OD$, 500 MHz) $\delta$ 3.92 (s, 1H, H-4), 3.52 – 3.26 (m, 8H, H-1a, H-1b, H-5, H-7 and H-8), 2.49 (m, 2H, H-6), 2.44 (t, *J* = 6.6 Hz, 2H, H-9), 1.98 (m, 2H, H-10), 0.94 (s, 6H, H-2 and H-3). $^{13}$C NMR ($CD_3OD$, 126 MHz) $\delta$ 174.7, 173.9, 172.8, 75.9,

68.9, 39.0, 38.7, 38.6, 35.2, 35.0, 29.5, 23.3 (d, $J$ = 138.1 Hz), 20.0, 19.5; $^{31}$P NMR ($CD_3OD$, 203 MHz) $\delta$ 26.50; HRMS (ESI$^+$) *m/z* [$C_{14}H_{27}N_3O_8PNa_2$]$^+$ calcd: 442.1337; found: 442.1309.

Synthesis of **9k (RK-04-030) (2-08)**

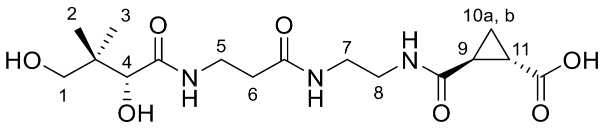

**9k** (A)

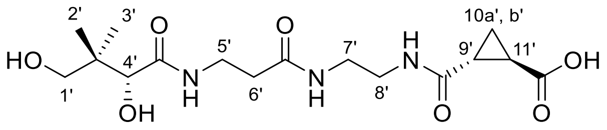

**9k** (B)

Compound **9k** was obtained from compound **8k** using the general procedure C. Compound **8k** (71 mg, 0.14 mmol) was dissolved in a mixture of AcOH/$H_2O$ (9:1, 6 mL) and the solution was stirred at rt for 4 h. Purification by column chromatography ($SiO_2$, DCM/MeOH = 90:10 to 88:12 v/v) afforded **6n** (31 mg, 40%) as a gummy white solid for this pair of diastereomers. $R_f$ = 0.20 (DCM/MeOH = 85:15, visualized by $KMnO_4$ stain); Purity 95%, RT = 16.94 min with method 7; RT = 17.29 min with method 8; $^1$H NMR ($CD_3OD$, 500 MHz) $\delta$ 3.89 (s, 2H, H-4 and H-4'), 3.53 – 3.33 (m, 12H, H-1, H-1', H-5, H-5', H-7 or H-8, and H-7' or H-8'), 3.30 – 3.23 (m, H-7 or H-8, and H-7' or H-8'), 2.49 – 2.36 (m, 4H, H-6 and H-6'), 2.17 – 2.07 (m, 2H, H-11 and H-11'), 2.02 – 1.94 (m, 2H, H-9 and H-9'), 1.37 – 1.31 (m, 2H, H-10a and H-10a'), 1.31 – 1.26 (m, 2H, H-10b and H-10b'), 0.92 (s, 12H, H-2, H-3, H-2' and H-3'); $^{13}$C NMR ($CD_3OD$, 126 MHz) $\delta$ 176.0, 175.8, 174.1, 173.3, 77.3, 70.3, 40.3, 40.2, 40.1, 36.6, 36.4, 24.6, 22.1, 21.4, 20.9, 14.8; HRMS (ESI$^+$) *m/z* [$C_{16}H_{27}N_3O_7Na$]$^+$ calcd: 396.1747; found: 396.1738.

Synthesis of **10 (RK-03-060) (6-04)**

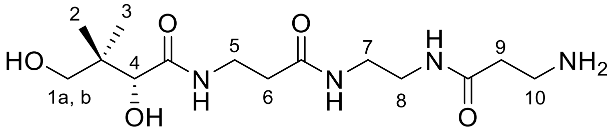

**10**

Compound **6l** (50 mg, 0.14 mmol) was dissolved in THF: $H_2O$ (9:1, 5 mL), and $PPh_3$ (73 mg, 0.28 mmol) was added before stirring the reaction mixture for 12 h. Solvents were evaporated *in vacuo*, water (10 mL) was added, and the reaction mixture was extracted with DCM (3 ×10 mL ) to remove excess $PPh_3$ and non-polar impurities. The aqueous layer was concentrated *in vacuo* to afford **10** (39 mg, 83%) as a sticky yellowish solid. $R_f$ = 0.10 (DCM/MeOH = 85:15); $^1$H NMR ($CD_3OD$, 500 MHz) $\delta$ 3.91 (s, 1H, H-4), 3.58 (t, $J$ = 6.5 Hz, 2H, H-10), 3.55 – 3.45 (m, 3H, H-

1a and H-5), 3.41 (d, $J$ = 10.9 Hz, 1H, H-1b), 3.35 – 3.26 (m, 4H, H-7 and H-8), 2.48 (t, $J$ = 6.5 Hz, 2H, H-6), 2.44 (t, $J$ = 6.6 Hz, 2H, H-9), 0.94 (s, 6H, H-2 and H-3). $^{13}$C NMR ($CD_3OD$, 126 MHz) $\delta$ 176.0, 174.2, 174.2, 77.3, 70.3, 40.3, 40.1, 39.9, 38.5, 37.8, 36.6, 36.4, 21.4, 20.9; HRMS (ESI$^+$) $m/z$ $[C_{14}H_{28}N_4O_5Na]^+$ calcd: 355.1957; found: 355.1935.

Synthesis of **12 (RK-04-020) (2-07)**

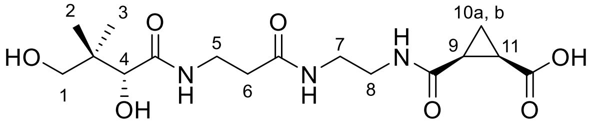

**12** (A)

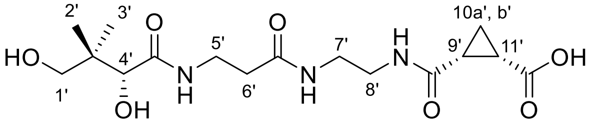

**12** (B)

Compound **12** was obtained from compound **11** using the general procedure C. Compound **11** (80 mg, 0.16 mmol) was dissolved in a mixture of AcOH/$H_2O$ (9:1, 4 mL) and the solution was stirred at rt for 4 h. Purification by column chromatography ($SiO_2$, DCM/MeOH = 88:12) afforded **12** (44 mg, 73%) as a white solid for this pair of diastereomers. $R_f$ = 0.30 (DCM/MeOH = 85:15, visualized by $KMnO_4$ stain); Purity 98%, RT = 13.63 min with method 7; RT = 16.69 min with method 8; $^1$H NMR ($CD_3OD$, 300 MHz) $\delta$ 3.88 (s, 2H, H-4 and H-4'), 3.53 – 3.34 (m, 8H, H-1, H-5, H-1' and H-5'), 3.30 – 3.20 (m, 8H, H-7, H-8, H-7' and H-8'), 2.47 – 2.34 (m, 4H, H-6 and H-6'), 2.16 – 1.92 (m, 4H, H-9, H-11, H-9' and H-11'), 1.59 – 1.41 (m, 2H, H-10a and H-10a'), 1.32 – 1.17 (m, 2H, H-10b and H-10b'), 0.91 (s, 12H, H-2, H-3, H-2' and H-3'); $^{13}$C NMR ($CD_3OD$, 75 MHz) $\delta$ 176.1, 175.3, 174.1, 172.5, 77.3, 70.3, 40.3, 40.1, 39.9, 36.6, 36.4, 25.2, 21.4, 20.9, 20.9, 12.4; HRMS (ESI$^+$) $m/z$ $[C_{16}H_{27}N_3O_7Na]^+$ calcd: 396.1747; found: 396.1727.

Synthesis of **15 (RK-04-104) (2-10)**

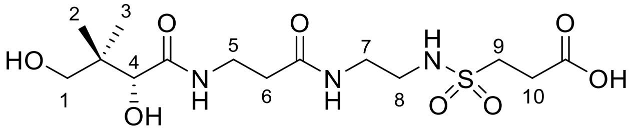

**15**

Compound **15** was obtained from compound **14** using the general procedure C. Compound **14** (272 mg, 0.53 mmol) was dissolved in a mixture of AcOH/$H_2O$ (9:1, 12 mL) and the solution was stirred at rt for 4 h. Purification by column chromatography ($SiO_2$, DCM/MeOH = 88:12) afforded **15** (78 mg, 37%) as a clear oil. $R_f$ = 0.20 (DCM/MeOH = 85:15, visualized by $KMnO_4$

stain); Purity 97%, RT = 14.66 min with method 7; RT = 18.08 min with method 8; $^{1}$H NMR ($CD_3OD$, 500 MHz) $\delta$ 3.91 (s, 1H, H-4), 3.57 – 3.28 (m, 8H, H-1, H-5, H-7 and H-8), 3.18 (m, 2H, H-9), 2.76 (m, 2H, H-10), 2.46 (t, $J$ = 6.6 Hz, 2H, H-6), 0.94 (s, 6H, H-2 and H-3); $^{13}$C NMR ($CD_3OD$, 126 MHz) $\delta$ 176.1, 174.2, 77.3, 70.3, 48.6, 43.1, 40.9, 40.4, 36.5, 36.4, 29.7, 21.3, 20.9; HRMS (ESI$^{+}$) $m/z$ $[C_{14}H_{27}N_3O_8SNa]^+$ calcd: 420.1417; found: 420.1395.

Synthesis of **16 (RK-04-103) (2-11)**

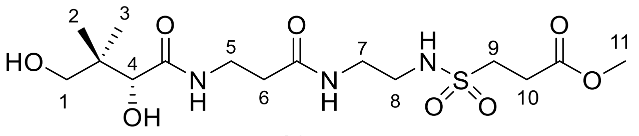

**16**

Compound **16** was obtained from compound **13** using the general procedure C. Compound **13** (168 mg, 0.32 mmol) was dissolved in a mixture of AcOH/$H_2O$ (9:1, 6 mL) and the solution was stirred at rt for 4 h. Purification by column chromatography ($SiO_2$, DCM/MeOH = 92:8) afforded **16** (94 mg, 72%) as a clear oil. $R_f$ = 0.40 (DCM/MeOH = 85:15, visualized by $KMnO_4$ stain); Purity >99%, RT = 12.38 min with method 7; RT = 14.38 min with method 8; $^{1}$H NMR ($CD_3OD$, 500 MHz) $\delta$ 3.90 (s, 1H, H-4), 3.72 (s, 3H, H-11), 3.59 – 3.23 (m, 8H, H-1, H-5, H-7, H-8), 3.16 (m, 2H, H-9), 2.79 (m, 2H, H-10), 2.44 (t, $J$ = 6.6 Hz, 2H, H-6), 0.93 (s, 6H, H-2 and H-3). $^{13}$C NMR ($CD_3OD$, 126 MHz) $\delta$ 176.0, 174.2, 172.8, 77.3, 70.3, 52.6, 48.3, 43.1, 40.9, 40.4, 36.6, 36.4, 29.5, 21.3, 20.9; HRMS (ESI$^{+}$) $m/z$ $[C_{15}H_{29}N_3O_8SNa]^+$ calcd: 434.1573; found: 434.1574.

Synthesis of **22a (RK-02-002) (1-14)**

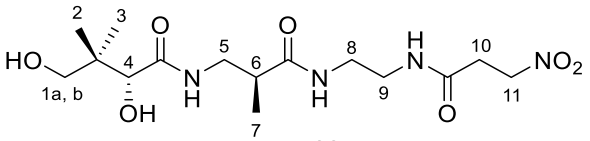

**22a** (A)

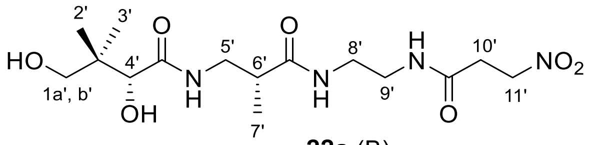

**22a** (B)

Compound **22a** was obtained from compound **21a** using the general procedure C. Compound **21a** (165 mg, 0.33 mmol) was dissolved in a mixture of AcOH/$H_2O$ (9:1, 10 mL) and the solution was stirred at rt for 4 h. Purification by column chromatography ($SiO_2$, DCM/MeOH = 90:10) afforded **22a** (77 mg, 61%) as a clear oil for this pair of diastereomers. $R_f$ = 0.30 (DCM/MeOH = 85:15, visualized by $KMnO_4$ stain); Purity 97%, RT = 12.43 min with method 7; RT = 14.26 min with method 8; $^{1}$H NMR ($CD_3OD$, 500 MHz) $\delta$ 4.72 – 4.70 (m, 4H, H-11 and H-

11’), 3.91 (s, 1H, H-4 or H-4’), 3.89 (s, 1H, H-4 or H-4’), 3.48 – 3.45 (m, 2H, H-1a and H-1a’), 3.42 – 3.22 (m, 14H, H-1b, H-1b’, H-5, H-8, H-9, H-5’, H-8’, H-9’), 2.88 – 2.85 (m, 4H, H-10 and H-10’), 2.62 – 2.54 (m, 2H, H-6 and H-6’), 1.13 (d, *J* = 1.7 Hz, 3H, H-7 or H-7’), 1.11 (d, *J* = 1.7 Hz, 3H, H-7 or H-7’), 0.92 (s, 12H, H-2, H-3, H-2’ and H-3’); $^{13}$C NMR ($CD_3OD$, 126 MHz) $\delta$ 177.7, 177.7, 176.1, 176.0, 171.8, 171.8, 77.3, 77.3, 71.2, 70.3, 70.2, 43.0, 43.0, 41.7, 41.6, 40.3, 40.3, 40.0, 40.0, 39.9, 33.0, 21.4, 21.3, 20.9, 15.9, 15.9; HRMS (ESI$^+$) *m/z* $[C_{15}H_{28}N_4O_7Na]^+$ calcd: 399.1856; found: 399.1849.

Synthesis of **22b (RK-03-015) (1-15)**

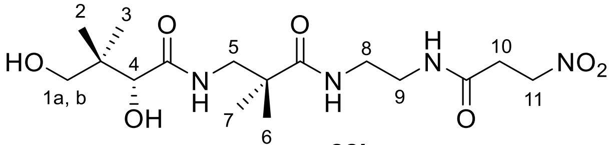

**22b**

Compound **22b** was obtained from compound **21b** using the general procedure C. Compound **21b** (190 mg, 0.37 mmol) was dissolved in a mixture of AcOH/$H_2O$ (9:1, 10 mL) and the solution was stirred at rt for 4 h. Purification by column chromatography ($SiO_2$, DCM/MeOH = 92:8) afforded **22b** (106 mg, 73%) as a white solid. $R_f$ = 0.50 (15:85 MeOH/DCM, visualized by $KMnO_4$ stain); Purity 97%, RT = 12.52 min with method 7; RT = 14.81 min with method 8; $^1$H NMR ($CD_3OD$, 500 MHz) $\delta$ 4.69 (m, 2H, H-11), 3.92 (s, 1H, H-4), 3.47 (d, *J* = 10.9 Hz, 1H, H-1a), 3.39 (d, *J* = 10.9 Hz, 1H, H-1b), 3.34 (m, 2H, H-5), 3.32 – 3.22 (m, 4H, H-8 and H-9), 2.87 (m, 2H, H-10), 1.18 (s, 3H, H-6 or H-7), 1.17 (s, 3H, H-6 or H-7), 0.93 (s, 6H, H-2 and H-3); $^{13}$C NMR ($CD_3OD$, 126 MHz) $\delta$ 179.53, 176.03, 172.04, 77.33, 71.20, 70.38, 48.04, 44.27, 40.70, 40.39, 39.93, 33.09, 23.99, 23.95, 21.30, 21.02; HRMS (ESI$^+$) *m/z* $[C_{16}H_{30}N_4O_7Na]^+$ calcd: 413.2012; found: 413.2004.

Synthesis of **22c (RK-03-016) (1-16)**

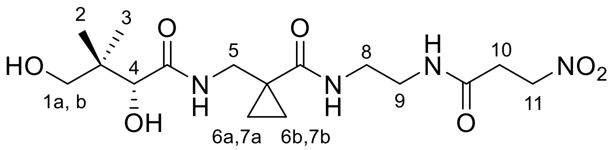

**22c**

Compound **22c** was obtained from compound **21c** using the general procedure C. Compound **21c** (145 mg, 0.29 mmol) was dissolved in a mixture of AcOH/$H_2O$ (9:1, 8 mL) and the solution was stirred at rt for 4 h. Purification by column chromatography ($SiO_2$, DCM/MeOH = 92:8) afforded **22c** (90 mg, 81%) as a white solid. $R_f$ = 0.50 (DCM/MeOH = 85:15, visualized by $KMnO_4$ stain); Purity >99%, RT = 12.54 min with method 7; RT = 14.73 min with method 8; $^1H$ NMR ($CD_3OD$, 500 MHz) $\delta$ 4.72 (m, 2H, H-11), 3.94 (s, 1H, H-4), 3.49-3.38 (m, 4H, H-1a and H-1b, H-5), 3.36 – 3.22 (m, 4H, H-8, H-9), 2.88 (m, 2H, H-10), 1.12 – 1.10 (m, 2H, H-6 or H-7), 0.92 (s, 6H, H-2 and H-3), 0.89 – 0.87 (m, 2H, H-6 or H-7); $^{13}C$ NMR ($CD_3OD$, 126 MHz) $\delta$ 176.6, 176.0, 172.0, 77.2, 71.2, 70.2, 43.4, 40.6, 40.4, 40.1, 33.2, 26.6, 21.4, 20.8, 14.6, 14.5; HRMS ($ESI^+$) *m/z* $[C_{16}H_{28}N_4O_7Na]^+$ calcd: 411.1856; found: 411.1845.

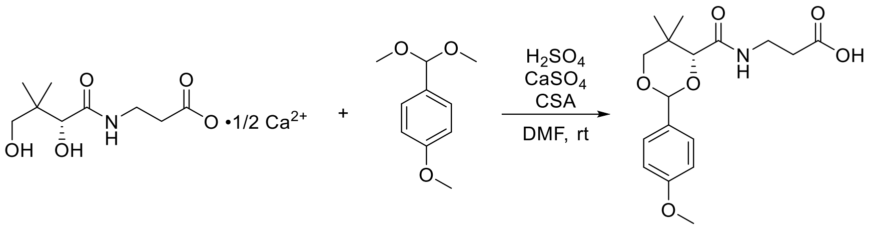

PMP-pantothenate

Synthesis of **PMP-pantothenate and Rac-PMP-pantothenate**

To a 100 mL round bottom flask were added racemic or D-pantothenic acid hemicalcium salt (4.77 g, 20 mmol, 1.0 equiv.), $CaSO_4$ powder (4 g) and anhydrous DMF (40 mL). Next, conc. $H_2SO_4$ (1.08 mL, 20 mmol, 1.0 equiv.) was added dropwise. The mixture was stirred at rt for 1 h. CSA (464.6 mg, 2 mmol, 0.1 equiv.) and *para*-anisaldehyde dimethyl acetal (5.10 mL, 30 mmol, 1.5 equiv.) were added and the reaction mixture was stirred at rt overnight. Upon reaction completion, the mixture was diluted in 100 mL water and filtered under vacuum. The filter cake was washed with EA, and the filtrate was washed with water (3 × 100 mL). The combined organic layers were dried over anhydrous $Na_2SO_4$, filtered, and the solvent was evaporated under vacuum to give **PMP-pantothenate** (2.96 g, 44%) as a white solid. To help crystalize the product to a white solid, Hex/diethyl ether = 1:1 can be added to the residue, and the resulting mixture filtered under vacuum. The filter cake is next washed with diethyl ether to give the product. **Rac-PMP-pantothenate** was prepared from the same procedure.[2]

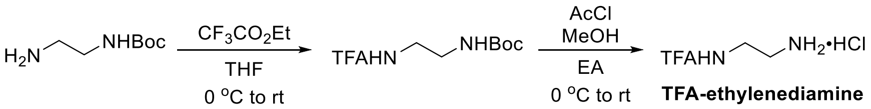

TFA-ethylenediamine hydrochloride

Synthesis of **TFA-ethylenediamine hydrochloride**

To a 100 mL round bottom flask were added Boc-ethylenediamine (6.24 mL, 40 mmol, 1.0 equiv.) and anhydrous THF (40 mL). The solution was cooled to 0°C and ethyl trifluoroacetate (4.80 mL, 40 mmol, 1.0 equiv.) was added. The reaction mixture was slowly returned to rt and stirred overnight. The solvent was then evaporated under vacuum, and the residue was dissolved in anhydrous EA (40 mL). MeOH (6.47 mL, 160 mmol, 4.0 equiv.) was added and the mixture was cooled to 0°C. Acetyl chloride (11.4 mL, 160 mmol, 4.0 equiv.) was added dropwise, and the reaction mixture was stirred at rt for 4 h. It was then filtered under vacuum and the filter cake was washed with EA to give **TFA-ethylenediamine hydrochloride** (6.66 g, 86% over two steps) as a white solid. $^1H$ NMR ($D_2O$, 500 MHz): $\delta$ 3.68 (t, $J$ = 6.0 Hz, 2H), 3.26 (t, $J$ = 6.0 Hz, 2H). Characterization matched previously reported data.[25]

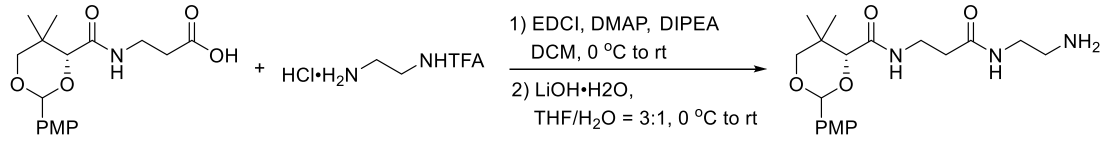

**PMP-NH2**

Synthesis of **PMP-NH2**

To a 50 mL round bottom flask were added TFA-ethylenediamine hydrochloride (481.4 mg, 2.5 mmol, 1.0 equiv.), PMP-pantothenate (927.7 mg, 2.75 mmol, 1.1 equiv.), DMAP (30.5 mg, 0.25 mmol), DIPEA (0.48 mL, 2.75 mmol, 1.1 equiv.), and DCM (10 mL). The mixture was cooled to 0°C and EDCI (527.2 mg, 2.75 mmol, 1.1 equiv.) was added in small portions. The reaction mixture was stirred at rt overnight. The mixture was diluted in DCM (20 mL), then washed with sat. $NaHCO_3$ (2 × 20 mL), 1 M $KH_2PO_4$ (2 × 20 mL), and brine (1 × 20 mL). The organic layer was dried over anhydrous $Na_2SO_4$, filtered, and evaporated under vacuum. Purification by flash chromatography (SiO2, DCM/EA = 1:1 then DCM/MeOH = 15:1) gave the intermediate (460 mg, 40%) as a white foam. To this crude intermediate (460 mg, 0.97 mmol, 1.0 equiv.) was added THF/$H_2O$ = 3:1 (16 mL). The mixture was cooled to 0°C and LiOH•$H_2O$ (83.9 mg, 2.0 mmol, 2.0 equiv.) was added. The mixture was stirred at rt for 3 h, after which water (40 mL) was added and the product was extracted in EA (3 × 25 mL). The combined organic layers were dried over anhydrous $Na_2SO_4$, filtered, and evaporated under vacuum to give **PMP-NH2** (220 mg, 60%) as a sticky light yellow oil. **Rac-PMP-NH2** was prepared from the same procedure.

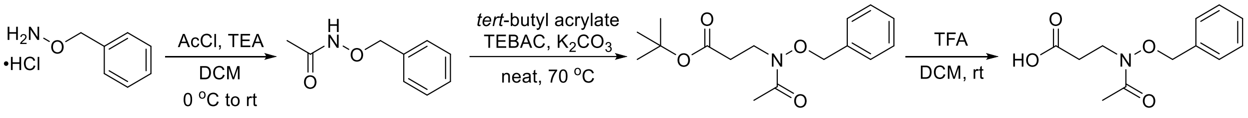

**BCL-1-105** **BCL-1-110** **BCL-1-115**

Synthesis of **BCL-1-115**

To a 100 mL round bottom flask were added *O*-benzylhydroxyamine hydrochloride (1.60 g, 10 mmol, 1.0 equiv.), DCM (20 mL), and TEA (3.10 mL, 22 mmol, 2.2 equiv.). The mixture was cooled to 0°C and acetyl chloride (0.86 mL, 12 mmol, 1.2 equiv.) was added dropwise. The reaction mixture was stirred at rt overnight. The reaction was then quenched with sat. aq. $NH_4Cl$ (20 mL), and diluted in water (20 mL). The product was extracted in DCM (3 × 20 mL). The combined organic layers were dried over anhydrous $Na_2SO_4$, filtered, and evaporated under vacuum.

Purification by flash chromatography ($SiO_2$, Hex/EA = 5:1 to 1:1) gave **BCL-1-105** (963 mg, 58%) as a colorless oil. $R_f$ = 0.15 (Hex/EA = 1:1); $^1$H NMR ($CDCl_3$, 500 MHz): $\delta$ (8.60, 8.36) (s, br, 1H), 7.37 (s, br, 5H), (4.88, 4.79 )(s, 2H), (2.01, 1.84) (s, 3H). Characterization matched previously reported data. [26]

Following a reported procedure,[27] to a 50 mL round bottom flask were added **BCL-1-105** (963 mg, 0.58 mmol, 1.0 equiv.), $K_2CO_3$ (483.7 mg, 3.5 mmol, 0.6 equiv.), TEBAC (79.7 mg, 0.35 mmol, 0.06 equiv.), and tert-butyl acrylate (0.85 mL), 5.8 mmol, 1.0 equiv.). The reaction was heated at 70°C for 4 h. The mixture was then cooled to rt, and water (15 mL) was added. The product was extracted in DCM (3 × 15 mL). The combined organic layers were dried over anhydrous $Na_2SO_4$, filtered, and evaporated under vacuum. Purification by flash chromatography ($SiO_2$, Hex/EA = 10:1 to 3:1) gave **BCL-1-110** (1.33 g, 78%) as a light yellow oil. $R_f$ = 0.15 (Hex/EA = 1:1); $^1$H NMR ($CDCl_3$, 500 MHz): $\delta$ 7.37 (bs, 5H), 4.82 (s, 2H), 3.90 (t, $J$ = 7.1 Hz, 2H), 2.53 (t, $J$ = 7.1 Hz, 2H), 2.05 (s, 3H), 1.41 (s, 9H). Characterization matched previously reported data.[28]

To a 50 mL round bottom flask were added **BCL-1-110** (1.33 g, 4.5 mmol, 1.0 equiv.), DCM (10 mL), and TFA (5 mL). The reaction mixture was stirred at rt for 3 h. The solvent was then evaporated under vacuum. The residue was dissolved in EA (30 mL), and the product was extracted in sat. $NaHCO_3$ (2 × 30 mL). The aqueous layer was washed with EA (1 × 20 mL), then acidified with 6 M HCl to *p*H = 4. The product was extracted in EA (3 × 20 mL). The combined organic layers were dried over anhydrous $Na_2SO_4$, filtered, and evaporated under vacuum to give **BCL-1-115** (641 mg, 60%) as a white solid. $^1$H NMR ($CDCl_3$, 500 MHz): $\delta$ 10.40 (bs 1H), 7.39-7.34 (m, 5H), 4.82 (s, 2H), 3.95 (t, $J$ = 7.1 Hz, 2H), 2.63 (t, $J$ = 7.0 Hz, 2H), 2.08 (s, 3H). Characterization matched previously reported data. [28]

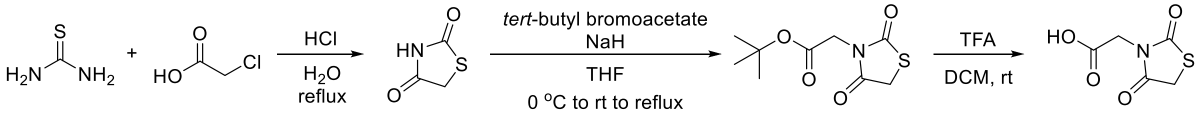

BCL-2-83 BCL-2-93 BCL-2-101

<u>Synthesis of **BCL-2-101**</u>

To a 100 mL round bottom flask were added thiourea (3.81 g, 50 mmol, 1.0 equiv.), chloroacetic acid (4.72 g, 50 mmol, 1.0 equiv.), and water (10 mL). The mixture was stirred at rt for 15 min. The mixture was then cooled to 0°C and conc. HCl (5 mL) was added. The reaction mixture was refluxed overnight. The mixture was then cooled to rt, and the precipitate was filtered under vacuum. The filter cake was washed with ice-cold water to give **BCL-2-83** (3.17 g, 54%) as a white solid. $^1$H NMR (DMSO-$d_6$, 500 MHz): $\delta$ 12.00 (bs, 1H), 4.13 (s, 2H). Characterization matched previously reported data.[29]

Following a reported procedure,[30] to a 100 mL round bottom flask were added **BCL-2-83** (1.17 g, 10 mmol, 1.0 equiv.) and anhydrous THF (25 mL). The mixture was cooled to 0°C and NaH (60%, 480 mg, 12 mmol, 1.2 equiv.) was added in one portion. The mixture was stirred at rt for 30 min. *tert*-Butyl bromoacetate (2.22 mL, 15 mmol, 1.5 equiv.) was added and the reaction mixture was refluxed for 4 h. The mixture was then cooled to rt, and quenched with sat. aq. $NH_4Cl$ (20 mL). The mixture was diluted in water (30 mL), and the product was extracted in EA (3 × 25 mL). The combined organic layers were dried over anhydrous $Na_2SO_4$, filtered, and evaporated under vacuum. Purification by flash chromatography ($SiO_2$, DCM) gave **BCL-2-93** (2.03 g, 88%) as a colorless oil, which solidifies upon standing. $R_f$ = 0.40 (DCM); $^1$H NMR ($CDCl_3$, 500 MHz): $\delta$ 4.24 (s, 2H), 4.01 (s, 2H), 1.46 (s, 9H). Characterization matched previously reported data.[30]

To a 50 mL round bottom flask were added **BCL-2-93** (2.03 g, 8.8 mmol, 1.0 equiv.), DCM (5 mL), and TFA (5 mL). The reaction mixture was stirred at rt for 2 h. The solvent was then evaporated under vacuum. The residue was dissolved in sat. $NaHCO_3$ until the solution was basic. The aqueous layer was washed with EA (3 × 20 mL), then acidified with 6 M HCl to *p*H = 2. The product was extracted in EA (3 × 25 mL). The combined organic layers were dried over anhydrous $Na_2SO_4$, filtered, and evaporated under vacuum to give **BCL-2-101** (1.54 g, quant.) as a yellow solid. $^1$H NMR (DMSO-$d_6$, 500 MHz): $\delta$ 4.32 (s, 2H), 4.20 (s, 2H). Characterization matched previously reported data. [31]

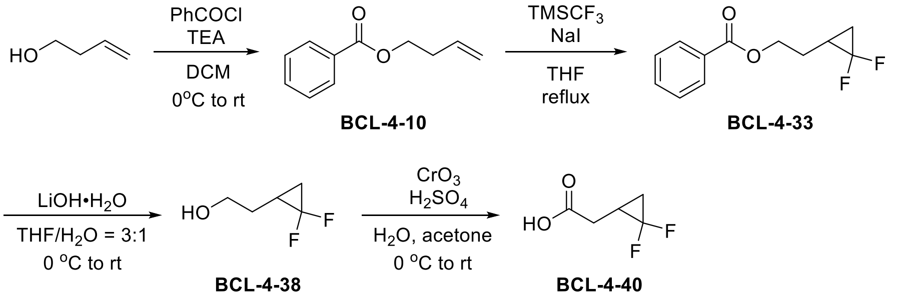

<u>Synthesis of **BCL-4-40**</u>

To a 100 mL round bottom flask were added distilled 3-buten-1-ol (2.58 mL, 30 mmol, 1.0 equiv.), TEA (5.02 mL, 36 mmol, 1.2 equiv.), and DCM (30 mL). The mixture was cooled to 0°C and benzoyl chloride (4.18 mL, 36 mmol, 1.2 equiv.) was added dropwise. The reaction mixture was stirred at rt overnight. Upon completion, the organic layer was washed with sat. $NaHCO_3$ (2 × 50 mL), 1 M $KH_2PO_4$ (2 × 50 mL), and brine (1 × 50 mL). The organic layer was dried over anhydrous $Na_2SO_4$, filtered, and evaporated under vacuum. Purification by vacuum distillation (14 mmHg, 118-120ºC) gave **BCL-4-10** (4.62 g, 87%) as a colorless oil. $^1$H NMR ($CDCl_3$, 500 MHz): $\delta$ 8.05 (m, 2H), 7.55 (m, 1H), 7.43 (m, 2H), 5.88 (ddt, $J$ = 17.1, 10.4, 6.8 Hz, 1H), 5.18 (m, 1H), 5.11 (m, 1H), 4.38 (t, $J$ = 6.7 Hz, 2H), 2.53 (m, 2H). Characterization matched previously reported data.[32]

Following a reported procedure with modification,[11] to a 100 mL round bottom flask were added **BCL-4-10** (1.76 g, 10 mmol, 1.0 equiv.), NaI (749.5 mg, 5 mmol, 0.5 equiv.), and anhydrous THF (20 mL). The atmosphere was changed to $N_2$ and a nitrogen balloon was attached to maintain the atmosphere. The mixture was heated to reflux and $TMSCF_3$ (3.70 mL, 25 mmol, 2.5 equiv.) was added dropwise over 30 min. The reaction mixture was refluxed overnight, after which TLC indicated that the reaction was not complete. An additional amount of NaI (749.5 mg, 5 mmol, 0.5 equiv.) and $TMSCF_3$ (3.70 mL, 25 mmol, 2.5 equiv.) was added and the reaction was refluxed for an additional 6 h. Additional amounts of NaI (749.5 mg, 5 mmol, 0.5 equiv.) and $TMSCF_3$ (3.70 mL, 25 mmol, 2.5 equiv.) were then added and the reaction mixture was refluxed overnight. The mixture was then cooled to rt, diluted in DCM (50 mL), and washed with water (2 × 50 mL), 10% $Na_2S_2O_3$ (2 × 50 mL), and brine (1 × 50 mL). The organic layer was dried over anhydrous $Na_2SO_4$, filtered, and evaporated under vacuum to give **BCL-4-33** (2.05 g, 91%) as a yellow oil, which was used directly without purification.

To a 250 mL round bottom flask were added **BCL-4-33** (2.05 g, 9.1 mmol) and THF/$H_2O$ = 3:1 (100 mL). The mixture was cooled to 0ºC and LiOH•$H_2O$ (1.15 g, 27.3 mmol, 3.0 equiv.) was added. The reaction mixture was stirred at rt overnight. The solvent was next evaporated under vacuum. The residue was dissolved in water (30 mL), and the product was extracted in EA (3 × 25 mL). The combined organic layers were washed with sat. $NaHCO_3$ (1 × 30 mL) and brine (1 × 30 mL), then dried over anhydrous $Na_2SO_4$, filtered, and evaporated under vacuum. Purification by flash chromatography ($SiO_2$, Hex/EA = 10:1 to 2:1) gave **BCL-4-38** (420 mg, 38%) as a yellow oil. $R_f$ = 0.45 (Hex/EA = 1:1); $^1$H NMR ($CDCl_3$, 500 MHz): $\delta$ 3.72-3.69 (m, 2H), 2.10 (bs, 1H), 1.85-1.51 (m, 3H), 1.40 (m, 1H), 0.94 (m, 1H); $^{19}$F NMR ($CDCl_3$, 471 MHz): $\delta$ -128.6 (d, $J$ = 155.4 Hz), -144.0 (d, $J$ = 155.4 Hz). Characterization matched previously reported data.[11]

Following a reported procedure,[33] to a 250 mL round bottom flask were added aq. $H_2SO_4$ (1.5 M, 30 mL). The solution was cooled to 0ºC and $CrO_3$ (1.80 g, 18 mmol, 5.3 mmol) was added. The mixture was stirred until homogenous to give a dark red solution as the Jones reagent. **BCL-4-38** (420 mg, 3.4 mmol) was dissolved in acetone (30 mL), then added to the mixture dropwise at 0°C. The reaction mixture was stirred at rt for 4 h. Upon completion, EtOH was added to destroy excess Jones reagent. The product was then extracted in diethyl ether (4 × 30 mL). The combined organic layers were washed with 2 M NaOH (4 × 30 mL). The combined aqueous layers were acidified with 6 M HCl to *p*H = 1. The product was extracted in diethyl ether (4 × 30 mL). The organic layers were dried over anhydrous $Na_2SO_4$, filtered, and evaporated under vacuum to give **BCL-4-40** (319 mg, 69%) as a colorless oil, which solidifies upon standing. $^1$H NMR ($CDCl_3$, 500 MHz): $\delta$ 10.74 (bs, 1H), 2.60 (ddt, $J$ = 17.4, 7.2, 1.5 Hz, 1H), 2.50 (dddd, $J$ = 17.4, 7.6, 2.7, 1.1 Hz, 1H), 1.84 (m, 1H), 1.55 (m, 1H), 1.08 (m, 1H); $^{19}$F NMR ($CDCl_3$, 471 MHz): $\delta$ -130.2 (d, $J$ = 160.1 Hz), -143.5 (d, $J$ = 160.1 Hz). Characterization matched previously reported data.[33]

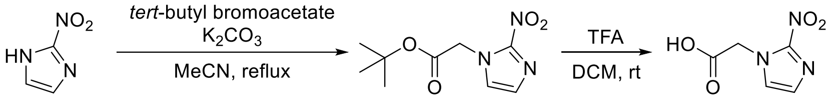

**BCL-4-46** **BCL-4-50**

<u>Synthesis of **BCL-4-50**</u>

To a 50 mL round bottom flask were added 2-nitroimidazole (565.4 mg, 5.0 mmol, 1.0 equiv.), $K_2CO_3$ (1.73 g, 12.5 mmol, 2.5 equiv.), *tert*-butyl bromoacetate (0.81 mL, 5.5 mmol, 1.1 equiv.), and MeCN (10 mL). The reaction mixture was refluxed for 3 h. The mixture was cooled to rt, and the solvent was evaporated under vacuum. Water (50 mL) was added to the residue and the product was extracted in EA (3 × 30 mL). The combined organic layers were washed with water (1 × 30 mL), sat. $NaHCO_3$ (1 × 30 mL), and brine (1 × 30 mL). The organic layer was dried over anhydrous $Na_2SO_4$, filtered, and evaporated under vacuum to give **BCL-4-46** (824 mg, 73%) as a yellow solid. This was directly used in next step without purification.

To a 50 mL round bottom flask were added **BCL-4-46** (824 mg, 3.62 mmol, 1.0 equiv.), DCM (2.5 mL), and TFA (2.5 mL). The reaction mixture was stirred at rt for 4 h before evaporating the solvent under vacuum. The residue was triturated in diethyl ether. The solid was collected by filtration under vacuum, and the filter cake was washed with diethyl ether and hexanes to give **BCL-4-50** (547 mg, 88%) as a light yellow solid. $^1$H NMR (DMSO-$d_6$, 500 MHz): $\delta$ 7.63 (d, $J$ = 1.1 Hz, 1H), 7.21 (d, $J$ = 1.1 Hz, 1H), 5.21 (s, 2H). Characterization matched previously reported data.[34]

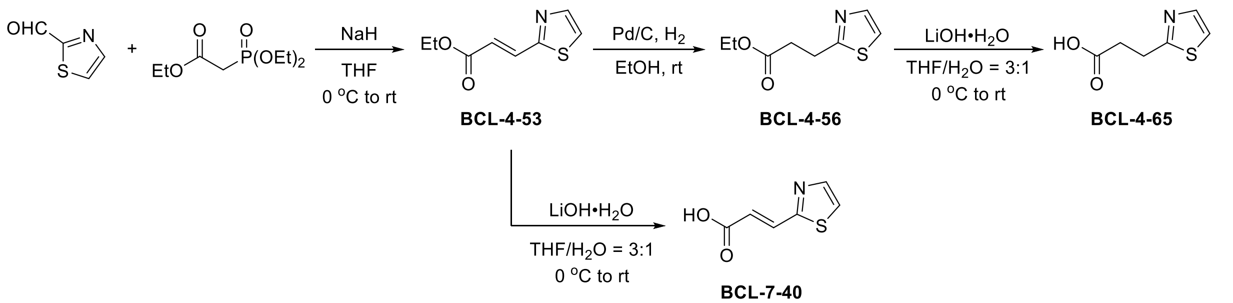

Synthesis of **BCL-4-65**

To a 100 mL round bottom flask was added NaH (60%, 424.0 mg, 10.6 mmol, 1.2 equiv.). The atmosphere was changed to $N_2$ and a $N_2$ balloon was attached to maintain the atmosphere. Anhydrous THF (5 mL) was added and the mixture was cooled to 0ºC. Triethyl phosphonoacetate (2.28 mL, 11.5 mmol, 1.3 equiv.) was dissolved in anhydrous THF (8 mL) and added dropwise to the mixture before stirring at 0ºC for 30 min. Thiazole 2-carbaldehyde (1.00 g, 8.84 mmol, 1.0 equiv.) was dissolved in anhydrous THF (8 mL) and added. The reaction mixture was stirred at rt overnight. Next, water (30 mL) was added and the product was extracted in EA (3 × 30 mL). The combined organic layers were washed with water (1 × 30 mL), sat. $NaHCO_3$ (1 × 20 mL), and brine (1 × 30 mL). The organic layer was dried over anhydrous $Na_2SO_4$, decolored with charcoal, filtered, and the solvent was evaporated under vacuum. Purification by flash chromatography ($SiO_2$, Hex/EA = 20:1 to 10:1) gave **BCL-4-53** (1.52 g, 94%) as a colorless oil. $R_f$ = 0.35 (Hex/EA = 5:1); $^1$H NMR ($CDCl_3$, 500 MHz): $\delta$ 7.87 (d, $J$ = 3.4 Hz, 1H), 7.75 (d, $J$ = 15.9 Hz, 1H), 7.39 (d, $J$ = 3.5 Hz, 1H), 6.67 (d, $J$ = 15.8 Hz, 1H), 4.23 (q, $J$ = 7.2 Hz, 2H), 1.29 (t, $J$ = 7.2 Hz, 3H). Characterization matched previously reported data.[35]

To a 100 mL round bottom flask were added **BCL-4-53** (1.52 g, 8.3 mmol, 1.0 equiv.), 10% Pd/C (150 mg), and EtOH (20 mL). $H_2$ was bubbled into the solution for 30 min, then a $H_2$ balloon was attached to the reaction flask. The reaction mixture was stirred at rt overnight. The mixture was then filtered over celite under vacuum, and the filter cake was washed with EtOH. The filtrate was evaporated under vacuum to give **BCL-4-56** (1.34 g, 87%) as a yellow oil. It was directly used in the next step without purification.

To a 250 mL round bottom flask were added **BCL-4-56** (1.34 g, 7.2 mmol, 1.0 equiv.) and THF/$H_2O$ = 3:1 (80 mL). The mixture was cooled to 0ºC and LiOH•$H_2O$ (759.4 mg, 18.1 mmol, 2.5 equiv.) was added. The reaction mixture was stirred at rt for 3 h. The solvent was then evaporated and the residue was redissolved in water (60 mL). The aqueous layer was washed with

EA (2 × 30 mL), then acidified with 6 M HCl to *p*H = 1. The product was extracted in EA (3 × 30 mL). The combined organic layers were dried over anhydrous $Na_2SO_4$, filtered, and evaporated under vacuum to give **BCL-4-65** (739 mg, 65%) as a white solid. $^1$H NMR (DMSO-$d_6$, 500 MHz): $\delta$ 12.26 (bs, 1H), 7.67 (d, $J$ = 3.3 Hz, 1H), 7.55 (d, $J$ = 3.3 Hz, 1H), 3.20 (t, $J$ = 7.1 Hz, 2H), 2.72 (t, $J$ = 7.1 Hz, 2H). Characterization matched previously reported data. [36]

To a 250 mL round bottom flask were added **BCL-4-53** (1.52 g, 8.5 mmol, 1.0 equiv.) and THF/$H_2O$ = 3:1 (90 mL). The mixture was cooled to 0ºC and LiOH•$H_2O$ (894.0 mg, 21.3 mmol, 2.5 equiv.) was added. The reaction mixture was stirred at rt for 3 h. The solvent was then evaporated and the residue was redissolved in water (60 mL). The aqueous layer was washed with EA (3 × 20 mL), then acidified with 6 M HCl to *p*H = 1. The product was extracted in EA (3 × 50 mL). The combined organic layers were dried over anhydrous $Na_2SO_4$, filtered, and evaporated under vacuum to give **BCL-7-40** (1.02 g, 77%) as a yellow solid. $^1$H NMR (DMSO-$d_6$, 500 MHz): $\delta$ 12.78 (bs, 1H), 7.97 (d, $J$ = 3.2 Hz, 1H), 7.88 (d, $J$ = 3.1 Hz, 1H), 7.67 (d, $J$ = 15.8 Hz, 1H), 6.66 (d, $J$ = 15.8 Hz, 1H). Characterization matched previously reported data.[37]

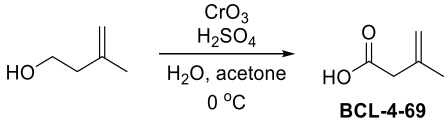

**BCL-4-69**

Synthesis of **BCL-4-69**

Following a reported procedure,[38] to a 50 mL round bottom flask was added water (4 mL). The solvent was cooled to 0ºC and $CrO_3$ (2.80 g, 28 mmol, 1.4 equiv.) was added. Conc. $H_2SO_4$ (2.7 mL, 50 mmol, 2.5 equiv.) was added dropwise followed by water (4 mL). The mixture was stirred until homogeneous to afford Jones reagent as a dark red solution. To a 250 mL round bottom flask were added distilled isoprenol (2.0 mL, 20 mmol, 1.0 equiv.) and acetone (100 mL). The mixture was cooled to 0ºC and the Jones reagent solution was added dropwise. The reaction mixture was stirred at 0ºC for 1 h. The solvent was then evaporated under vacuum, and the residue was redissolved in water (100 mL). The product was extracted in diethyl ether (4 × 50 mL). The combined organic layers were washed with 2 M NaOH (4 × 30 mL). The combined aqueous layers were acidified with 50% $H_2SO_4$ to pH = 1. The product was extracted in diethyl ether (4 × 50 mL). The combined organic layers were dried over anhydrous $Na_2SO_4$, filtered, and evaporated under vacuum. Purification by vacuum distillation (14 mmHg, 77-79ºC) gave **BCL-4-69** (664 mg, 33%)

as a colorless oil. $^1$H NMR ($CDCl_3$, 500 MHz): $\delta$ 4.94 (m, 1H), 4.88 (m, 1H), 3.07 (m, 2H), 1.83 (m, 3H). Characterization matched previously reported data.[38]

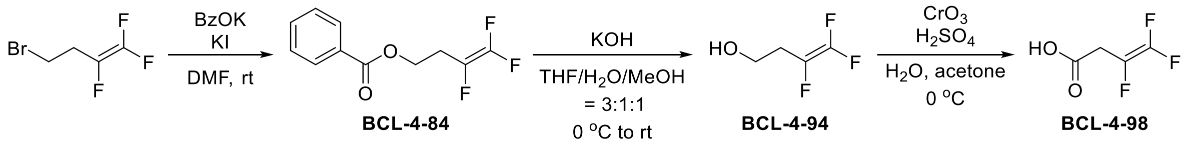

<u>Synthesis of **BCL-4-98**</u>

To a 50 mL round bottom flask were added 1,1,2-trifluoro-4-bromobut-1-ene (1.50 g, 7.93 mmol, 1.0 equiv.), potassium benzoate (1.53 g, 9.52 mmol, 1.2 equiv.), KI (132.8 mg, 0.8 mmol, 0.1 equiv.), and anhydrous DMF (15 mL). The reaction mixture was stirred at rt overnight. The mixture was then diluted in EA (70 mL), washed with water (2 × 50 mL), sat. $NaHCO_3$ (2 × 50 mL), and brine (1 × 40 mL). The organic layer was dried over anhydrous $Na_2SO_4$, filtered, and evaporated under vacuum to give **BCL-4-84** (630 mg, 35%) as a yellow oil. This crude product was directly used in the next step without purification. $^1$H NMR ($CDCl_3$, 500 MHz): $\delta$ 8.03 (m, 2H), 7.56 (m, 1H), 7.45 (m, 2H), 4.48 (t, $J$ = 6.2 Hz, 2H), 2.76 (m, 2H); $^{19}$F NMR ($CDCl_3$, 471 MHz): $\delta$ -103.0 (m), -123.7 (ddd, $J$ = 114.6, 85.0, 3.9 Hz), -175.0 (dd, $J$ = 114.7, 33.1 Hz). Characterization matched previously reported data.[39]

To a 250 mL round bottom flask were added **BCL-4-84** (1.72 g, 7.47 mmol, 1.0 equiv.) and THF/$H_2O$/MeOH =3:1 (90 mL). The mixture was cooled to 0°C and KOH (1.05 g, 18.68 mmol, 2.5 equiv.) was added. The reaction mixture was stirred at rt for 3 h. The solvent was then evaporated under vacuum. The residue was dissolved in water and the product was extracted in diethyl ether (4 × 25 mL). The combined organic layers were dried over anhydrous $Na_2SO_4$, filtered, and evaporated under vacuum to give **BCL-4-94** (374 mg, 40%) as a yellow oil. This volatile product was directly used in the next step without purification.

To a 50 mL round bottom flask was added water (2 mL). The solvent was cooled to 0°C and $CrO_3$ (414.4 mg, 4.15 mmol, 1.4 equiv.) was added. Conc. $H_2SO_4$ (0.40 mL, 7.4 mmol, 2.5 equiv.) was added dropwise, followed by water (1 mL). The mixture was stirred until homogeneous to afford Jones reagent as a dark red solution. To a 100 mL round bottom flask were added **BCL-4-94** (374 mg, 2.96 mmol, 1.0 equiv.) and acetone (15 mL). The mixture was cooled to 0°C and the Jones reagent solution was added dropwise. The reaction mixture was stirred at 0°C for 2 h. The solvent was then evaporated under vacuum, and the residue was redissolved in water (100 mL). The solution was extracted in diethyl ether (3 × 25 mL). The combined organic layers were washed with 1 M NaOH (3 × 25 mL). The combined aqueous layers were acidified with 50% $H_2SO_4$ to *p*H = 2. The product was extracted in diethyl ether (3 × 25 mL). The combined organic layers were dried over anhydrous $Na_2SO_4$, filtered, and evaporated under vacuum. This volatile product was immediately used in the next step without purification to avoid material loss.

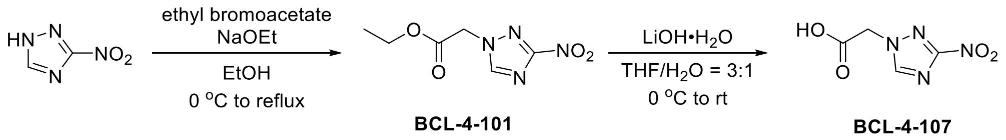

Synthesis of **BCL-4-107**

Following a reported procedure,[40] to a 100 mL round bottom flask was added EtOH (15 mL). Sodium metal (275.8 mg, 12 mmol, 1.2 equiv.) was added and the mixture was stirred at 0°C until the sodium had disappeared. To this freshly prepared NaOEt solution was added 3-nitro-1,2,4-triazole (1.14 g, 10 mmol, 1.0 equiv.) and the mixture was stirred at 0°C for 30 min. Ethyl bromoacetate (1.33 mL, 12 mmol, 1.2 equiv.) was added and the reaction mixture was refluxed overnight. Upon completion, 2 drops of water were added and the solvent was evaporated under vacuum. Purification by flash chromatography ($SiO_2$, DCM/EA = 20:1 to 10:1) gave **BCL-4-101** (1.80 g, 90%) as a yellow oil, which solidifies upon standing. $R_f$ = 0.34 (Hex/EA = 1:1); $^1$H NMR ($CDCl_3$, 500 MHz): $\delta$ 8.33 (s, 1H), 5.08 (s, 2H), 4.30 (q, $J$ = 7.2 Hz, 2H), 1.32 (t, $J$ = 7.2 Hz, 3H). Characterization matched previously reported data.[40]

To a 250 mL round bottom flask were added **BCL-4-101** (1.80 g, 9.0 mmol, 1.0 equiv.) and THF/$H_2O$ = 3:1 (100 mL). The mixture was cooled to 0°C and LiOH•$H_2O$ (944.1 mg, 22.5 mmol, 2.5 equiv.) was added. The reaction mixture was stirred at rt for 3 h. The solvent was then evaporated and the residue was redissolved in water (60 mL). This aqueous solution was washed with EA (2 × 30 mL), then acidified with 6 M HCl to *p*H = 1. The product was extracted in EA (4 × 25 mL). The combined organic layers were dried over anhydrous $Na_2SO_4$, filtered, and evaporated under vacuum to give **BCL-4-107** (1.38 g, 90%) as a white solid. $^1$H NMR (DMSO-$d_6$, 500 MHz): $\delta$ 8.86 (s, 1H), 5.29 (s, 2H). Characterization matched previously reported data.[40]

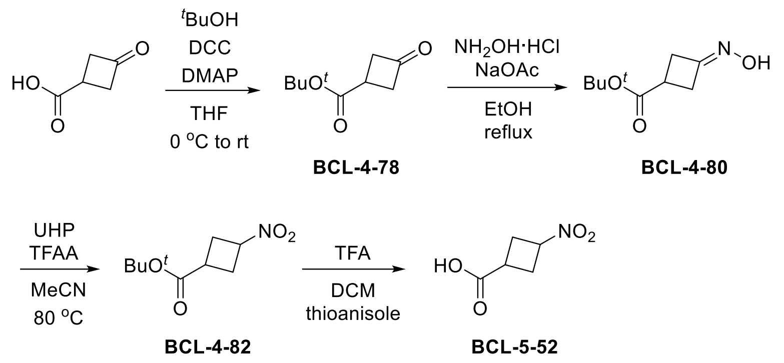

#### Synthesis of **BCL-5-52**

To a 100 mL round bottom flask were added 3-oxocyclobutanecarboxylic acid (2.62 g, 22.3 mmol, 1.0 equiv.), *tert*-butyl alcohol (2.96 mL, 31.2 mmol, 1.4 equiv.), DMAP (1.36 g, 11.2 mmol, 0.5 equiv.), and anhydrous THF (18 mL). The mixture was cooled to 0ºC. DCC (5.06 g, 24.5 mmol, 1.1 equiv.) was dissolved in anhydrous THF (7 mL) and added dropwise. The reaction mixture was stirred at rt overnight. Next, the precipitate was filtered over celite under vacuum, and the filtrate was concentrated under vacuum. The residue was dissolved in EA (75 mL) and washed with sat. $NaHCO_3$ (2 × 50 mL), 1 M $KH_2PO_4$ (2 × 50 mL), and brine (1 × 50 mL). The combined organic layers were dried over anhydrous $Na_2SO_4$, filtered, and evaporated under vacuum. The residue was redissolved in Hex/EA = 5:1 (100 mL) and passed through a short silica pad. The solvent was evaporated under vacuum to give **BCL-4-78** (3.49 g, 92%) as a light yellow oil that was directly used in next step without further purification.

To a 250 mL round bottom flask were added **BCL-4-78** (3.49 g, 20.5 mmol, 1.0 equiv.), sodium acetate (6.73 g, 82.0 mmol, 4.0 equiv.), hydroxylamine hydrochloride (2.83 g, 41.0 mmol, 2.0 equiv.), and EtOH (60 mL). The reaction mixture was refluxed for 4 h. The mixture was then cooled to rt, and approximately half of the solvent was evaporated under vacuum. The mixture was poured into ice water (100 mL), and the product was extracted in EA (3 × 50 mL). The combined organic layers were dried over anhydrous $Na_2SO_4$, filtered, and evaporated under vacuum. Impurities were removed from the residue by trituration with hexanes to give **BCL-4-80** (3.65 g, 88%) as a white solid. This was directly used in next step without purification.

Following a reported procedure,[41] to a 100 mL round bottom flask were added UHP (5.62 g, 59.7 mmol, 3.3 equiv.) and anhydrous MeCN (40 mL). The mixture was cooled to -10ºC. TFAA (8.3 mL, 59.7 mmol, 3.3 equiv.) was dissolved in anhydrous MeCN (20 mL) and added dropwise over 1 h. The mixture was stirred at -10ºC for 1 h to give the $H_2O_2$•TFAA solution. To a 250 mL round

bottom flask were added **BCL-4-80** (3.65 g, 18.1 mmol, 1.0 equiv.), $Na_2HPO_4$ (19.5 g, 137.6 mmol, 7.6 equiv.), and anhydrous MeCN (40 mL). The mixture was heated to 80°C, and the prepared $H_2O_2$•TFAA solution was added over 30 min. The reaction mixture was stirred at 80°C for another 30 min. The mixture was then allowed to cool to rt and filtered under vacuum. The filtrate was concentrated under vacuum and the residue was redissolved in EA (100 mL) before washing with water (1 × 50 mL) and brine (2 × 50 mL). Purification by flash chromatography (Hex/EA = 50:1 to 10:1) gave **BCL-4-82** (1.88 g, 52%) as a light yellow oil. *cis* isomer: $R_f$ = 0.25 (Hex/EA = 10:1); $^1$H NMR ($CDCl_3$, 500 MHz): $\delta$ 4.80 (m, 1H), 2.89-2.69 (m, 5H), 1.44 (s, 9H); *trans* isomer: $R_f$ = 0.40 (Hex/EA = 10:1); $^1$H NMR ($CDCl_3$, 500 MHz): $\delta$ 5.05 (m, 1H), 3.14 (m, 1H), 2.86 (m, 2H), 2.72 (m, 2H), 1.45 (s, 9H). Characterization matched previously reported data.[41]
To a 50 mL round bottom flask were added **BCL-4-82** (380.0 mg, 1.88 mmol, 1.0 equiv.), DCM (1.5 mL), thioanisole (1.5 mL), and TFA (1.5 mL). The reaction mixture was stirred at rt for 3 h. The solvent was then evaporated under vacuum, and the residue was redissolved in DCM (25 mL). The product was extracted in sat. $NaHCO_3$ (2 × 30 mL), and the combined aqueous layers were washed with DCM (2 × 25 mL) and EA (1 × 25 mL). The aqueous layer was acidified with 6 M HCl to *p*H = 2. The product was extracted in diethyl ether (3 × 25 mL). The combined organic layers were dried over anhydrous $Na_2SO_4$, filtered, and evaporated under vacuum. Impurities were removed from the residue by trituration with hexanes to give **BCL-5-52** (167 mg, 61%) as a white solid. $^1$H NMR ($CDCl_3$, 500 MHz): $\delta$ 11.52 (bs, 1H), 5.11 (m, 1H), 3.32 (m, 1H), 2.96 (m, 2H), 2.83 (m, 2H); $^{13}$C NMR ($CDCl_3$, 125 MHz): $\delta$ 180.86, 75.45, 32.73, 31.05; HRMS (ESI$^-$) *m/z* $[C_{10}H_{13}N_2O_8]^-$ calcd: 289.0677 found: 289.0673.

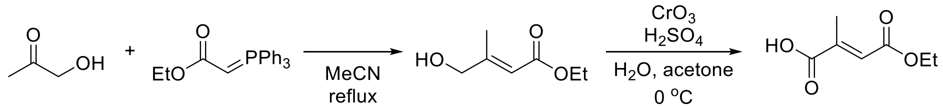

BCL-7-14 BCL-7-19

Synthesis of **BCL-7-19**

Following a reported procedure,[42] to a 100 mL round bottom flask were added hydroxyacetone (0.70 mL, 10 mmol, 1.0 equiv.), ethyl (triphenylphosphoranylidene)acetate (4.18 g, 12 mmol, 1.2 equiv.), and anhydrous MeCN (25 mL). The reaction mixture was refluxed overnight. The mixture was then cooled to rt, and the solvent was evaporated under vacuum. The residue was dissolved in diethyl ether (20 mL) and filtered under vacuum. The filtrate was concentrated under vacuum. Purification by flash chromatography (Hex/EA = 10:1 to 4:1) gave **BCL-7-14** (840 mg, 58%) as a yellow oil. $^1$H NMR ($CDCl_3$, 500 MHz): $\delta$ 5.94 (m, 1H), 4.13 (q, $J$ = 7.2 Hz, 2H), 4.08 (m, 2H), 2.82 (bs, 1H), 2.04 (s, 3H), 1.25 (t, $J$ = 7.1 Hz, 3H). Characterization matched previously reported data.[42]

To a 50 mL round bottom flask was added water (4 mL). The solvent was cooled to 0°C and $CrO_3$ (1.45 g, 14.5 mmol, 2.5 equiv.) was added. Conc. $H_2SO_4$ (1.40 mL, 26.1 mmol, 4.5 equiv.) was added dropwise, followed by water (2 mL). The mixture was stirred until homogeneous to afford Jones reagent as a dark red solution. To a 250 mL round bottom flask were added **BCL-7-14** (840.0 mg, 5.8 mmol, 1.0 equiv.) and acetone (60 mL). The mixture was cooled to 0°C and the Jones reagent solution was added dropwise. The reaction mixture was stirred at 0°C for 40 min. The solvent was then evaporated under vacuum, and the residue was redissolved in water (100 mL). Theproduct was extracted from this solution using diethyl ether (4 × 50 mL). The combined organic layers were washed with brine (4 × 50 mL), dried over anhydrous $Na_2SO_4$, filtered, and evaporated under vacuum to give **BCL-7-19** (856 mg, 93%) as a white solid. $^1$H NMR (DMSO-$d_6$, 500 MHz): $\delta$ 6.61 (q, $J$ = 1.6 Hz, 1H), 4.14 (q, $J$ = 7.1 Hz, 2H), 2.14 (d, $J$ = 1.7 Hz, 3H), 1.22 (t, $J$ = 7.1 Hz, 3H). Characterization matched previously reported data.[43]

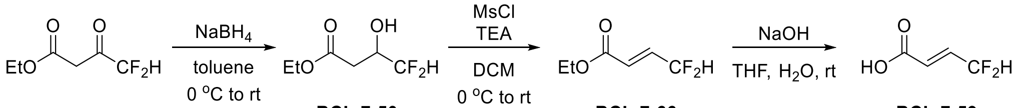

BCL-7-50 BCL-7-66 BCL-7-59

<u>Synthesis of **BCL-7-59**</u>

To a 250 mL round bottom flask were added ethyl 4,4-difluoroacetoacetate (2.61 mL, 20 mmol, 1.0 equiv.) and toluene (100 mL). The mixture was cooled to 0°C and $NaBH_4$ (794.4 mg, 21 mmol, 1.05 equiv.) was added in small portions. The reaction mixture was stirred at rt for 4 h. The reaction was then quenched with water (50 mL). The product was extracted in EA (2 × 50 mL). The combined organic layers were dried over anhydrous $Na_2SO_4$, filtered, and evaporated under vacuum to give **BCL-7-50** (2.67 g, 79%) as a colorless oil. This was used directly in the next step without purification.

To a 250 mL round bottom flask were added **BCL-7-50** (3.09 g, 18.4 mmol, 1.0 equiv.), TEA (10.3 mL, 73.6 mmol, 4.0 equiv.), and DCM (70 mL). The mixture was cooled to 0°C and MsCl (1.48 mL, 19.3 mmol, 1.05 equiv.) was added dropwise. The reaction mixture was stirred at rt overnight. The mixture was then washed with 1 M $KH_2PO_4$ (3 × 50 mL). The organic layer was dried over anhydrous $Na_2SO_4$, filtered, and evaporated under vacuum. Purification by vacuum distillation (48 mmHg, 43°C) gave **BCL-7-66** (2.12 g, 77%) as a volatile colorless oil. $^1$H NMR ($CDCl_3$, 500 MHz): $\delta$ 6.79 (dtd, $J$ = 15.9, 10.2, 4.1 Hz, 1H), 6.27 (dt, $J$ = 16.0, 3.1 Hz, 1H), 6.21 (tdd, $J$ = 54.8, 4.2, 1.1 Hz, 1H), 4.24 (q, $J$ = 7.1 Hz, 2H), 1.30 (t, $J$ = 7.1 Hz, 3H); $^{19}$F NMR ($CDCl_3$, 471 MHz): $\delta$ -116.0. Characterization matched previously reported data.[44]

To a 100 mL round bottom flask were added **BCL-7-66** (2.12 g, 14.1 mmol) and THF (27 mL). Next, 4 M NaOH (8.8 mL, 35.3 mmol, 2.5 equiv.) was added and the reaction mixture was stirred at rt for 3 h. The mixture was then diluted in MTBE (20 mL), and the aqueous layer was kept separately. The organic layer was washed with 1 M NaOH (2 × 10 mL), and the aqueous layers were combined. The aqueous layers were washed with MTBE (1 × 20 mL), and acidified with 6 M HCl to $p$H = 2. The product was extracted in MTBE (4 × 20 mL). The combined organic layers were dried over anhydrous $Na_2SO_4$, filtered, and evaporated under vacuum. The residue was put on ice to initiate crystallization. The crystals were collected by filtration and washed with hexanes to give **BCL-7-59** (1.27 g, 74%) as a white solid. $^1$H NMR ($CDCl_3$, 500 MHz): $\delta$ 10.57 (bs, 1H), 6.92 (dtd, $J$ = 15.9, 10.2, 4.0 Hz, 1H), 6.31 (dt, $J$ = 15.8, 2.3 Hz, 1H), 6.26 (tdd, $J$ = 54.7, 4.0, 0.8 Hz, 1H); $^{19}$F NMR ($CDCl_3$, 471 MHz): $\delta$ -116.8. Characterization matched previously reported data.[44]

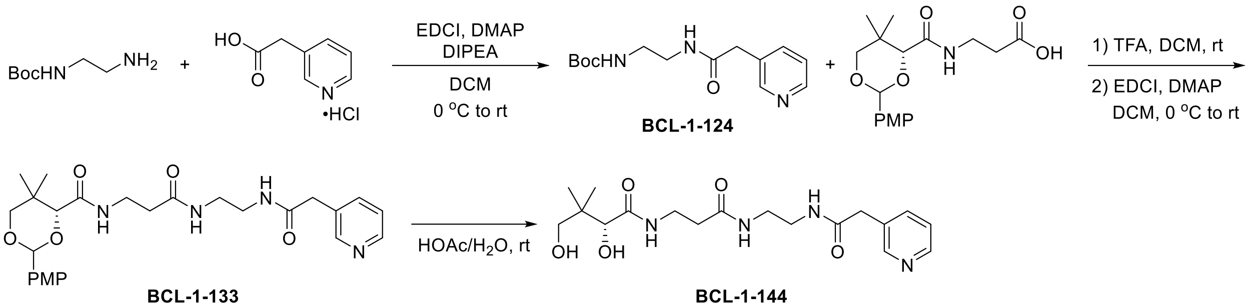

BCL-1-124

BCL-1-133

BCL-1-144

<u>Synthesis of **BCL-1-144 (5-03)**</u>

Following general procedure E with modifications, 3-pyridylacetic acid hydrochloride (286.4 mg, 1.65 mmol), Boc-ethylenediamine (0.24 mL, 1.5 mmol), DMAP (18.3 mg, 0.15 mmol), EDCI (316.3 mg, 1.65 mmol), and DCM (5 mL) were used. DIPEA (0.56 mL, 3.3 mmol, 2.2 equiv.) was also introduced with the acid at the beginning. Purification by flash chromatography ($SiO_2$, DCM/EA = 1:1, then DCM/MeOH = 15:1) gave **BCL-1-124** (200 mg, 48%) as an orange solid. $R_f$ = 0.55 (DCM/MeOH = 10:1); $^1H$ NMR ($CDCl_3$, 500 MHz): $\delta$ 8.47 (s, 2H), 7.64 (d, $J$ = 8.2 Hz, 1H), 7.23 (s, 1H), 6.77 (bs, 1H), 5.13 (bs, 1H), 3.49 (s, 2H), 3.31 (bs, 2H), 3.22 (bs, 2H), 1.40 (s, 9H); $^{13}C$ NMR ($CDCl_3$, 125 MHz): $\delta$ 170.6, 157.2, 150.3, 148.5, 137.0, 131.0, 123.7, 79.8, 41.3, 40.6, 40.2, 28.5; HRMS (ESI$^+$) *m/z* $[C_{14}H_{21}N_3O_3Na]^+$ calcd: 302.1475; found: 302.1467.

Following general procedure F, **BCL-1-124** (200.0 mg, 0.72 mmol), TFA (1 mL), and DCM (1 mL) were used. Next, general procedure G was followed with PMP-pantothenate (236.2 mg, 0.7 mmol), DMAP (8.6 mg, 0.07 mmol), EDCI (134.2 mg, 0.7 mmol), and DCM (4 mL). Purification by flash chromatography ($SiO_2$, DCM/MeOH = 15:1) gave **BCL-1-133** (79 mg, 22% over two steps) as a waxy light yellow solid. $R_f$ = 0.40 (DCM/MeOH = 10:1); $^1H$ NMR ($CDCl_3$, 500 MHz): $\delta$ 8.42 (bs, 2H), 7.60 (d, $J$ = 7.8 Hz, 1H), 7.40 (d, $J$ = 8.7 Hz, 2H), 7.19 (bs, 2H), 7.14 (bs, 1H), 7.09 (t, $J$ = 6.3 Hz, 1H), 6.87 (d, $J$ = 8.7 Hz, 2H), 5.42 (s, 1H), 4.04 (s, 1H), 3.76 (s, 3H), 3.67 (d, $J$ = 11.4 Hz, 1H), 3.61 (d, $J$ = 11.3 Hz, 1H), 3.48-3.40 (m, 4H), 3.27-3.23 (m, 4H), 2.32 (t, $J$ = 6.4 Hz, 2H), 1.06 (s, 3H), 1.03 (s, 3H); $^{13}C$ NMR ($CDCl_3$, 125 MHz): $\delta$ 172.2, 171.0, 169.7, 160.3, 150.1, 148.2, 137.1, 131.6 (bs), 130.2, 127.6, 123.2 (bs), 113.8, 101.4 83.9, 78.5, 55.4, 54.0, 42.2, 40.4, 40.3, 39.4, 36.0, 35.1, 33.1, 21.9, 19.2; HRMS (ESI$^+$) *m/z* $[C_{26}H_{34}N_4O_6Na]^+$ calcd: 521.2371; found: 521.2352.

Following general procedure H, **BCL-1-133** (61.0 mg, 0.12 mmol) and 80% HOAc (1 mL) were used. Purification by flash chromatography ($SiO_2$, DCM/MeOH = 10:1 to 5:1) gave **BCL-1-144** (33 mg, 72%) as a waxy light yellow solid. $R_f$ = 0.40 (DCM/MeOH = 5:1); Purity: >99%, RT =

12.72 min with method 12, RT = 14.89 min with method 13; $^1$H NMR ($CD_3OD$, 500 MHz): $\delta$ 8.50-8.46 (m, 2H), 7.81 (d, $J$ = 8.5 Hz, 1H), 7.42 (m, 1H), 3.90 (s, 1H), 3.59 (s, 2H), 3.53-3.43 (m, 3H), 3.42-3.33 (m, 2H), 3.33-3.26 (m, 3H), 2.39 (t, $J$ = 6.7 Hz, 2H), 0.93 (s, 6H); $^{13}$C NMR ($CD_3OD$, 125 MHz): $\delta$ 176.0, 174.1, 173.1, 150.6, 148.4, 139.1, 133.8, 125.3, 77.3, 70.3, 40.6, 40.3, 40.2, 40.0, 36.6, 36.4, 21.4, 20.9; HRMS (ESI$^+$) *m/z* $[C_{18}H_{28}N_4O_5Na]^+$ calcd: 403.1952; found: 403.1952.

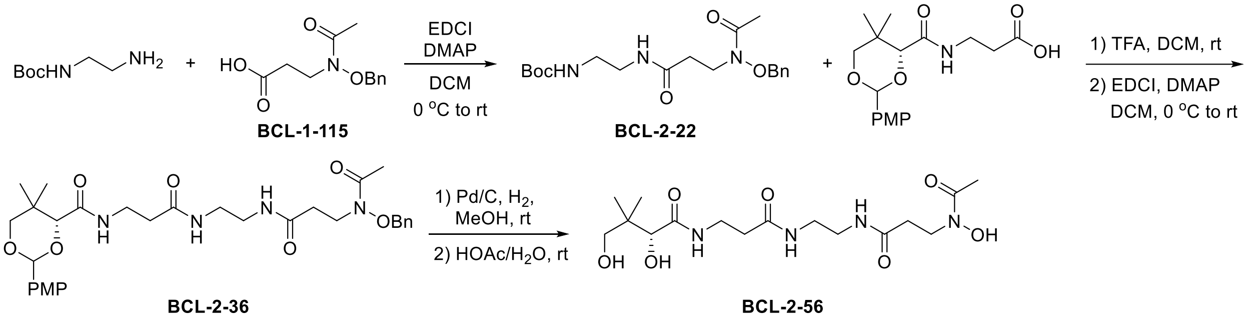

<u>Synthesis of **BCL-2-56 (6-02)**</u>

Following general procedure E, **BCL-1-115** (261.0 mg, 1.1 mmol), Boc-ethylenediamine (0.16 mL, 1.0 mmol), DMAP (12.2 mg, 0.1 mmol), EDCI (210.9 mg, 1.1 mmol), and DCM (5 mL) were used. Purification by flash chromatography ($SiO_2$, DCM/MeOH = 20:1 to 10:1) gave **BCL-2-22** (364 mg, 96%) as a white solid. $R_f$ = 0.62 (DCM/MeOH = 10:1); $^1$H NMR ($CDCl_3$, 500 MHz): $\delta$ 7.30-7.12 (m, 5H), 7.13 (t, $J$ = 5.5 Hz, 1H), 5.48 (t, $J$ = 5.9 Hz, 1H), 4.74 (s, 2H), 3.89 (t, $J$ = 6.6 Hz, 2H), 3.20 (m, 2H), 3.09 (m, 2H), 2.40 (t, $J$ = 6.7 Hz, 2H), 1.96 (s, 3H), 1.33 (s, 9H); $^{13}$C NMR ($CDCl_3$, 125 MHz): $\delta$ 172.6, 171.1, 156.3, 134.0, 129.3, 128.8, 128.5, 78.9, 76.0, 41.8, 39.9, 39.8, 33.9, 28.2, 20.3; HRMS (ESI$^+$) *m/z* $[C_{19}H_{29}N_3O_5Na]^+$ calcd: 402.1999; found: 402.1987.

Following general procedure F, **BCL-2-22** (364.0 mg, 0.96 mmol), TFA (1 mL), and DCM (1 mL) were used. Next, general procedure G was followed with PMP-pantothenate (323.9 mg, 0.96 mmol), DMAP (12.2 mg, 0.1 mmol), EDCI (184.0 mmol, 0.96 mmol), and DCM (5 mL). Purification by flash chromatography ($SiO_2$, DCM/MeOH = 20:1) gave **BCL-2-36** (366 mg, 65% over two steps) as a waxy yellow solid. $R_f$ = 0.59 (DCM/MeOH = 10:1); $^1$H NMR ($CDCl_3$, 500 MHz): $\delta$ 7.39 (d, $J$ = 8.7 Hz, 2H), 7.33 (s, 5H), 7.06 (t, $J$ = 6.3 Hz, 1H), 7.02 (bs, 1H), 6.99 (bs, 1H), 6.86 (d, $J$ = 8.7 Hz, 2H), 5.40 (s, 1H), 4.78 (s, 2H), 4.01 (s, 1H), 3.90 (s, 2H), 3.75 (s, 3H), 3.64 (d, $J$ = 11.4 Hz, 1H), 3.59 (d, $J$ = 11.4 Hz, 1H), 3.51-3.39 (m, 2H), 3.23-3.18 (m, 4H), 2.41 (t, $J$ = 6.7 Hz, 2H), 2.32 (t, $J$ = 6.4 Hz, 2H), 2.00 (s, 3H), 1.05 (s, 3H), 1.01 (s, 3H); $^{13}$C NMR ($CDCl_3$, 125 MHz): $\delta$ 171.7, 171.4, 169.5, 160.2, 134.1, 130.1, 129.4, 129.1, 128.7, 127.5, 113.7,

101.3, 83.8, 78.4, 76.3, 55.3, 50.4, 42.0, 39.4, 39.3, 36.0, 35.1, 34.1, 33.0, 21.8, 20.5, 19.1; HRMS (ESI$^+$) *m/z* $[C_{31}H_{42}N_4O_8Na]^+$ calcd: 621.2895; found: 621.2921.

Following general procedure I, **BCL-2-36** (366.0 mg, 0.65 mmol), Pd/C (35 mg), and MeOH (3 mL) was used. Next, general procedure H was followed using 80% HOAc (2 mL). No purification was needed to give **BCL-2-56** (114 mg, 45%) as a beige solid. Purity: 99%, RT = 10.73 min with method 12, RT = 14.42 min with method 13; $^1$H NMR ($CD_3OD$, 500 MHz): $\delta$ 3.91 (s, 1H), 3.86 (t, $J$ = 6.9 Hz, 2H), 3.47 (m, 2H), 3.44 (d, $J$ = 11.0 Hz, 1H), 3.37 (d, $J$ = 11.1 Hz, 1H), 3.29-3.18 (m, 4H), 2.50 (t, $J$ = 6.9 Hz, 2H), 2.42 (t, $J$ = 6.8 Hz, 2H), 2.08 (s, 3H), 0.91 (s, 3H), 0.90 (s, 3H); $^{13}$C NMR ($CD_3OD$, 125 MHz): $\delta$ 176.2, 174.1, 174.0, 173.7, 77.3, 70.2, 45.8, 40.3, 40.0, 36.52, 36.49, 34.6, 21.3, 20.3; HRMS (ESI$^+$) *m/z* $[C_{16}H_{30}N_4O_7Na]^+$ calcd: 413.2007; found: 413.1992.

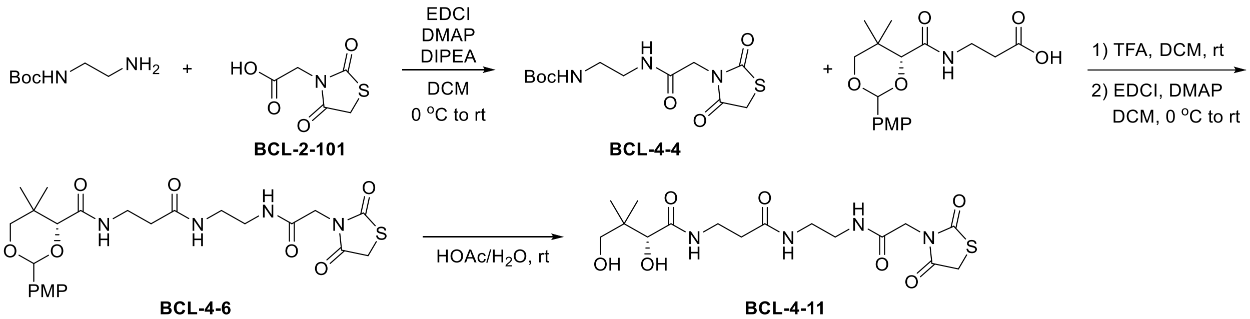

<u>Synthesis of **BCL-4-11 (5-04)**</u>

Following general procedure J with modification, **BCL-2-101** (289.0 mg, 1.65 mmol), Boc-ethylenediamine (0.24 mL, 1.5 mmol), HOBt (278.7 mg, 1.65 mmol), EDCI (316.3 mg, 1.65 mmol), DIPEA (0.29 mL, 1.5 equiv.) and DCM (8 mL) were used. Purification by flash chromatography ($SiO_2$, DCM/MeOH = 50:1) gave **BCL-4-4** (391 mg, 82%) as a white solid. $R_f$ = 0.58 (DCM/MeOH = 10:1); $^1$H NMR ($CDCl_3$, 500 MHz): $\delta$ 7.05 (bs, 1H), 5.08 (t, $J$ = 6.0 Hz, 1H), 4.24 (s, 2H), 4.05 (s, 2H), 3.34 (m, 2H), 3.24 (m, 2H), 1.43 (s, 9H); $^{13}$C NMR ($CDCl_3$, 125 MHz): $\delta$ 171.7, 171.2, 165.6, 157.6, 80.2, 43.9, 41.8, 40.1, 34.0, 28.5; HRMS (ESI$^+$) *m/z* $[C_{12}H_{19}N_3O_5SNa]^+$ calcd: 340.0938; found: 340.0932.

Following general procedure F, **BCL-4-4** (391.0 mg, 1.23 mmol), TFA (1.5 mL), and DCM (1.5 mL) were used. Next, general procedure K was followed with PMP-pantothenate (414.9 mg, 1.23 mmol), HOBt (207.7 mg, 1.23 mmol), EDCI (235.8 mg, 1.23 mmol), DIPEA (0.21 mL, 1.23 mmol), and DCM (5 mL). Purification by flash chromatography ($SiO_2$, DCM/MeOH = 50:1 to 20:1) gave **BCL-4-6** (415 mg, 63% over two steps) as a white foam. $R_f$ = 0.30 (DCM/MeOH = 10:1); $^1$H NMR ($CDCl_3$, 500 MHz): $\delta$ 7.41 (d, $J$ = 8.8 Hz, 2H), 7.33 (t, $J$ = 4.6 Hz, 1H), 7.06 (t, $J$

= 6.3 Hz, 1H), 6.89 (d, $J$ = 8.7 Hz, 2H), 6.86 (t, $J$ = 5.8 Hz, 1H), 5.44 (s, 1H), 4.13 (s, 2H), 4.05 (s, 1H), 3.96 (s, 2H), 3.79 (s, 3H), 3.68 (d, $J$ = 11.4 Hz, 1H), 3.63 (d, $J$ = 11.5 Hz, 1H), 3.50 (m, 2H), 3.34-3.23 (m, 4H), 2.39 (t, $J$ = 6.3 Hz, 2H), 1.08 (s, 3H), 1.04 (s, 3H); $^{13}$C NMR ($CDCl_3$, 125 MHz): $\delta$ 172.8, 171.8, 171.3, 169.9, 166.0, 160.3, 130.2, 127.7, 113.8, 101.4, 83.9, 78.5, 55.4, 43.8, 40.8, 39.0, 36.2, 35.2, 34.1, 33.2, 22.0, 19.2; HRMS (ESI$^+$) *m/z* $[C_{24}H_{32}N_4O_8SNa]^+$ calcd: 559.1833; found: 559.1828.

Following general procedure H, **BCL-4-6** (415.0 mg, 0.77 mmol) and 80% HOAc (3 mL) were used. Purification by flash chromatography (C1 $SiO_2$, DCM/MeOH = 20:1 to 15:1) gave **BCL-4-11** (79 mg, 25%) as a white solid. $R_f$ = 0.18 (DCM/MeOH = 10:1); Purity: 97%, RT = 12.62 min with method 12, RT = 15.04 min with method 13; $^1$H NMR ($D_2O$, 500 MHz): $\delta$ 4.35 (s, 2H), 4.24 (s, 1H), 3.99 (s, 1H), 3.57-3.44 (m, 3H), 3.42-3.29 (m, 5H), 2.49 (t, $J$ = 6.6 Hz, 2H), 0.92 (s, 3H), 0.89 (s, 3H); $^{13}$C NMR ($D_2O$, 125 MHz): $\delta$ 175.2, 175.1, 174.2, 174.1, 168.4, 75.8, 68.4, 43.3, 38.8, 38.59, 38.56, 35.5, 35.3, 34.3, 20.5, 19.1; HRMS (ESI$^+$) *m/z* $[C_{16}H_{26}N_4O_7SNa]^+$ calcd: 441.1414; found: 441.1423.

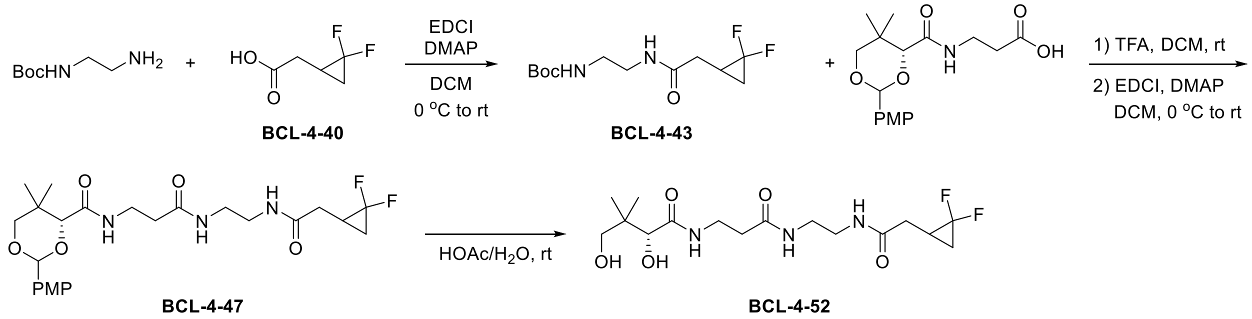

Synthesis of **BCL-4-52 (4-09)**

Following general procedure E, **BCL-4-40** (149.7 mg, 1.1 mmol), Boc-ethylenediamine (0.16 mL, 1.0 mmol), DMAP (12.2 mg, 0.1 mmol), EDCI (210.8 mg, 1.1 mmol), and DCM (5 mL) were used. Purification by flash chromatography ($SiO_2$, DCM/MeOH = 20:1) gave **BCL-4-43** (262 mg, 94%) as a white solid. $R_f$ = 0.55 (DCM/MeOH = 10:1); $^1$H NMR ($CDCl_3$, 500 MHz): $\delta$ 6.61 (bs, 1H), 5.06 (t, $J$ = 6.1 Hz, 1H), 3.35 (m, 2H), 3.26 (m, 2H), 2.41 (dd, $J$ = 16.0, 7.9 Hz, 1H), 2.27 (ddd, $J$ = 16.1, 8.0, 2.8 Hz, 1H), 1.85 (m, 1H), 1.52 (m, 1H), 1.41 (s, 9H), 1.04 (m, 1H); $^{19}$F NMR ($CDCl_3$, 471 MHz): $\delta$ -129.5 (d, $J$ = 160.1 Hz), -142.9 (d, $J$ = 160.1 Hz); $^{13}$C NMR ($CDCl_3$, 125 MHz): $\delta$ 171.1, 157.3, 113.6 (dd, $J$ = 283.7, 283.7 Hz), 79.9, 41.2, 40.2, 33.9 (d, $J$ = 5.0 Hz), 28.4, 18.8 (t, $J$ = 11.3 Hz), 16.3 (t, $J$ = 11.3 Hz); HRMS (ESI$^+$) *m/z* $[C_{12}H_{20}N_2O_3F_2Na]^+$ calcd: 301.1334; found: 301.1325.

Following general procedure F, **BCL-4-43** (262.0 mg, 0.94 mmol), TFA (1 mL), and DCM (1 mL) were used. Next, general procedure G was followed using PMP-pantothenate (317.2 mg, 0.94 mmol), DMAP (11.0 mg, 0.09 mmol), EDCI (180.2 mmol, 0.94 mmol), and DCM (5 mL). Purification by flash chromatography ($SiO_2$, DCM/MeOH = 50:1 to 20:1) gave **BCL-4-47** (341 mg, 73% over two steps) as a white solid. $R_f$ = 0.50 (DCM/MeOH = 10:1); $^1H$ NMR ($CDCl_3$, 500 MHz): $\delta$ 7.41 (d, $J$ = 8.7 Hz, 2H), 7.07-6.99 (m, 2H), 6.93-6.84 (m, 3H), 5.43 (s, 1H), 4.04 (s, 1H), 3.79 (s, 3H), 3.68 (d, $J$ = 11.4 Hz, 1H), 3.63 (d, $J$ = 11.5 Hz, 1H), 3.49 (m, 2H), 3.34-3.28 (m, 4H), 2.39 (t, $J$ = 6.4 Hz, 2H), 2.32 (dd, $J$ = 15.9, 7.6 Hz, 1H), 2.25 (ddd, $J$ = 15.9, 7.6, 2.5 Hz, 1H), 1.82 (m, 1H), 1.48 (m, 1H), 1.07 (s, 3H), 1.04 (s, 3H), 0.99 (m, 1H); $^{19}F$ NMR ($CDCl_3$, 471 MHz): $\delta$ (-129.35, -129.36) (d, $J$ = 155.4 Hz), (-142.88, -142.89) (d, $J$ = 155.4 Hz); $^{13}C$ NMR ($CDCl_3$ 12,5 MHz): $\delta$ 172.2, 171.4 (d, $J$ = 2.1 Hz), 169.7, 160.3, 130.2, 127.6, 113., 113.6 (dd, $J$ = 283.8, 283.8 Hz), 101., 83.9, 78.5, 55.4, 40.2, 39.6, 36.1, 35.1, 33.8 (d, $J$ = 3.9 Hz), 33.1, 21.9, 19.2, 18.7 (t, $J$ = 11.3 Hz), 16.2 (t, $J$ = 11.3 Hz); HRMS (ESI$^+$) *m/z* $[C_{24}H_{33}N_3O_6F_2Na]^+$ calcd: 520.2230; found: 520.2238.

Following general procedure H, **BCL-4-47** (341.0 mg, 0.68 mmol) and 80% HOAc (3 mL) were used. Purification by flash chromatography ($SiO_2$, DCM/MeOH = 20:1 to 10:1) gave **BCL-4-52** (132 mg, 51%) as a sticky white solid. $R_f$ = 0.20 (DCM/MeOH = 10:1); Purity: 98%, RT = 15.65 min with method 12, RT = 15.81 min with method 13; $^1H$ NMR ($D_2O$, 500 MHz): $\delta$ 3.98 (s, 1H), 3.54-3.50 (m, 3H), 3.51 (d, $J$ = 11.3 Hz, 1H), 3.37-3.28 (m, 4H), 2.49 (t, $J$ = 6.6 Hz, 2H), 2.47-2.37 (m, 2H), 1.89 (m, 1H), 1.61(m, 1H), 1.17 (m, 1H), 0.92 (s, 3H), 0.89 (s, 3H); $^{19}F$ NMR ($CDCl_3$, 471 MHz): $\delta$ -129.4 (d, $J$ = 155.4 Hz), (-143.09, -143.10) (d, $J$ = 155.4 Hz); $^{13}C$ NMR ($D_2O$, 125 MHz): $\delta$ 175.1, 174.4 (d, $J$ = 2.3 Hz), 174.1, 114.2 (dd, $J$ = 282.5, 282.5 Hz), 75.8 (×2), 68.3, 38.63, 38.60, 38.58, 35.4, 35.3, 33.1 (d, $J$ = 4.3 Hz), 20.5, 18.1 (t, $J$ = 11.3 Hz), 15.4 (t, $J$ = 11.3 Hz); HRMS (ESI$^+$) *m/z* $[C_{16}H_{27}N_3O_5F_2Na]^+$ calcd: 402.1811; found: 402.1812.

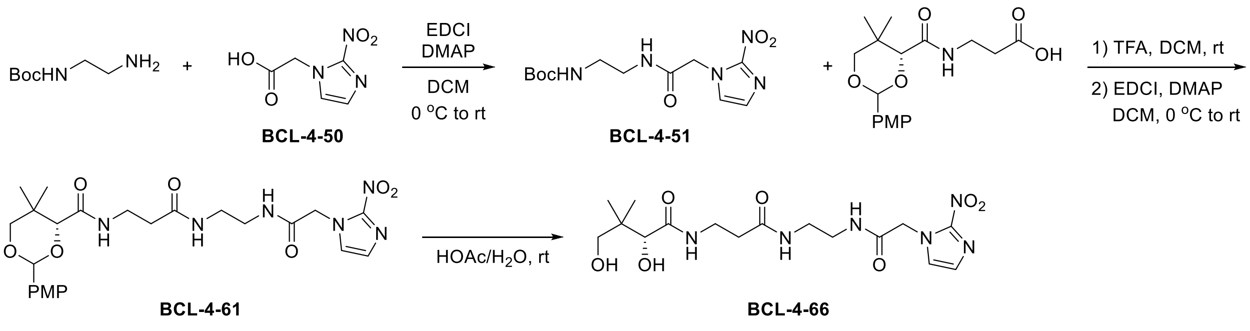

Synthesis of **BCL-4-66 (5-06)**

Following general procedure E, **BCL-4-50** (188.2 mg, 1.1 mmol), Boc-ethylenediamine (0.16 mL, 1.0 mmol), DMAP (12.2 mg, 0.1 mmol), EDCI (210.8 mg, 1.1 mmol), and DCM (5 mL) were used. Purification by flash chromatography ($SiO_2$, DCM/MeOH = 20:1) gave **BCL-4-51** (125 mg, 40%) as a white solid. $R_f$ = 0.46 (DCM/MeOH = 10:1); $^1H$ NMR ($CD_3OD$, 500 MHz): $\delta$ 7.45 (d, $J$ = 1.2 Hz, 1H), 7.19 (d, $J$ = 1.2 Hz, 1H), 5.16 (s, 2H), 3.31 (t, $J$ = 6.2 Hz, 2H), 3.18 (t, $J$ = 6.3 Hz, 2H), 1.45 (s, 9H); $^{13}C$ NMR ($CD_3OD$, 125 MHz): $\delta$ 168.3, 158.6, 146.4, 129.3, 128.4, 80.2, 52.8, 40.8, 40.7, 28.7; HRMS (ESI$^+$) $m/z$ $[C_{12}H_{19}N_5O_5Na]^+$ calcd: 336.1278; found: 336.1276.

Following general procedure F, **BCL-4-51** (265.0 mg, 0.85 mmol), TFA (1.5 mL), and DCM (1.5 mL) were used. Next, general procedure G was used with PMP-pantothenate (286.7 mg, 0.85 mmol), DMAP (11.0 mg, 0.09 mmol), EDCI (162.9 mmol, 0.85 mmol), and DCM (5 mL). Purification by flash chromatography ($SiO_2$, DCM/MeOH = 25:1 to 20:1) gave **BCL-4-61** (371 mg, 82% over two steps) as a yellow solid. $R_f$ = 0.39 (DCM/MeOH = 10:1); $^1H$ NMR ($CDCl_3$, 500 MHz): $\delta$ 7.61 (t, $J$ = 4.9 Hz, 1H), 7.40 (d, $J$ = 8.7 Hz, 2H), 7.10 (d, $J$ = 3.3 Hz, 1H), 7.09 (d, $J$ = 3.3 Hz, 1H), 7.07 (t, $J$ = 6.4 Hz, 1H), 6.92 (t, $J$ = 6.4 Hz, 1H), 6.86 (d, $J$ = 8.8 Hz, 2H), 5.43 (s, 1H), 4.90 (s, 2H), 4.04 (s, 1H), 3.77 (s, 3H), 3.68 (d, $J$ = 11.4 Hz, 1H), 3.62 (d, $J$ = 11.4 Hz, 1H), 3.49 (m, 2H), 3.30-3.27 (m, 4H), 2.36 (t, $J$ = 6.4 Hz, 2H), 1.08 (s, 3H), 1.03 (s, 3H); $^{13}C$ NMR ($CDCl_3$, 125 MHz): $\delta$ 172.6, 169.8, 166.0, 160.3, 145.0, 130.2, 128.3, 127.7, 127.6, 113.8, 101.4, 83.9, 78.5, 55.4, 52.2, 40.2, 39.2, 36.2, 35.2, 33.2, 22.0, 19.2; HRMS (ESI$^+$) $m/z$ $[C_{24}H_{32}N_6O_8Na]^+$ calcd: 555.2174; found: 555.2173.

Following general procedure H, **BCL-4-61** (371.0 mg, 0.70 mmol) and 80% HOAc (3 mL) were used. Purification by flash chromatography (C1 $SiO_2$, DCM/MeOH = 20:1 to 10:1) gave **BCL-4-66** (146 mg, 51%) as a white solid. $R_f$ = 0.30 (DCM/MeOH = 5:1); Purity: 97%, RT = 11.89 min with method 12, RT = 14.84 min with method 13; $^1H$ NMR ($D_2O$, 500 MHz): $\delta$ 7.47 (bs, 1H), 7.26 (bs, 1H), 5.21 (s, 2H), 3.97 (s, 1H), 3.53-3.46 (m, 3H), 3.43-3.27 (m, 5H), 2.49 (t, $J$ = 6.6 Hz, 2H), 0.92 (s, 3H), 0.88 (s, 3H); $^{13}C$ NMR ($D_2O$, 125 MHz): $\delta$ 175.1, 174.2, 168.5, 144.4, 128.9, 128.0, 75.8, 68.3, 51.9, 38.8, 38.62, 38.56, 35.4, 35.3, 20.4, 19.1; HRMS (ESI$^+$) $m/z$ $[C_{16}H_{26}N_6O_7Na]^+$ calcd: 437.1755; found: 437.1754.

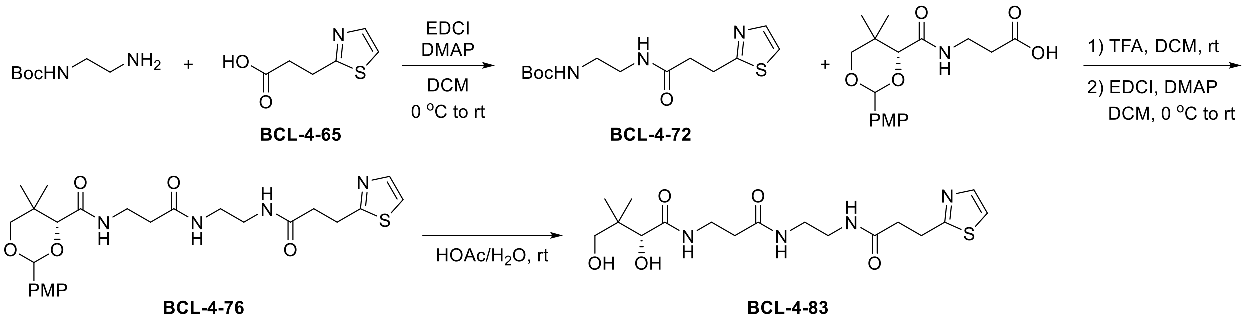

Synthesis of **BCL-4-83 (5-01)**

Following general procedure E, **BCL-4-65** (172.9 mg, 1.1 mmol), Boc-ethylenediamine (0.16 mL, 1.0 mmol), DMAP (12.2 mg, 0.1 mmol), EDCI (210.8 mg, 1.1 mmol), and DCM (5 mL) were used. Purification by flash chromatography ($SiO_2$, DCM/MeOH = 20:1) gave **BCL-4-72** (270 mg, 90%) as a light yellow solid. $R_f$ = 0.50 (DCM/MeOH = 10:1); $^1$H NMR ($CDCl_3$, 500 MHz): $\delta$ 7.64 (d, $J$ = 3.3 Hz, 1H), 7.17 (d, $J$ = 3.3 Hz, 1H), 6.76 (t, $J$ = 5.8 Hz, 1H), 5.23 (t, $J$ = 5.9 Hz, 1H), 3.32 (t, $J$ = 7.2 Hz, 2H), 3.31 (m, 2H), 3.20 (m, 2H), 2.66 (t, $J$ = 7.2 Hz, 2H), 1.39 (s, 9H); $^{13}$C NMR ($CDCl_3$, 125 MHz): $\delta$ 172.1, 169.4, 156.7, 142.3, 118.6, 79.6, 40.4, 40.3, 35.6, 28.7, 28.5; HRMS (ESI$^+$) *m/z* $[C_{13}H_{22}N_3O_3S]^+$ calcd: 300.1376; found: 300.1377.

Following general procedure F, **BCL-4-72** (270.0 mg, 0.9 mmol), TFA (1.5 mL), and DCM (1.5 mL) were used. Next, general procedure G was followed using PMP-pantothenate (303.6 mg, 0.9 mmol), DMAP (11.0 mg, 0.09 mmol), EDCI (172.5 mmol, 0.9 mmol), and DCM (5 mL). Purification by flash chromatography ($SiO_2$, DCM/MeOH = 20:1 to 15:1) gave **BCL-4-76** (433 mg, 93% over two steps) as a white solid. $R_f$ = 0.50 (DCM/MeOH = 10:1); $^1$H NMR ($CDCl_3$, 500 MHz): $\delta$ 7.61 (d, $J$ = 3.3 Hz, 1H), 7.40 (d, $J$ = 8.7 Hz, 2H), 7.17 (d, $J$ = 3.3 Hz, 1H), 7.04 (t, $J$ = 6.3 Hz, 1H), 6.95 (t, $J$ = 5.1 Hz, 1H), 6.91 (t, $J$ = 5.5 Hz, 1H), 6.88 (d, $J$ = 8.7 Hz, 2H), 5.43 (s, 1H), 4.04 (s, 1H), 3.78 (s, 3H), 3.67 (d, $J$ = 11.4 Hz, 1H), 3.62 (d, $J$ = 11.5 Hz, 1H), 3.48 (m, 2H), 3.30 (t, $J$ = 7.2 Hz, 2H), 3.28-3.21 (m, 4H), 2.64 (t, $J$ = 7.1 Hz, 2H), 2.35 (t, $J$ = 6.3 Hz, 2H), 1.07 (s, 3H), 1.04 (s, 3H); $^{13}$C NMR ($CDCl_3$, 125 MHz): $\delta$ 172.4, 171.8, 169.6, 169.4, 160.3, 142.1, 130.2, 127.6, 118.8, 113.8, 101.4, 83.9, 78.5, 55.4, 39.8, 39.6, 36.1, 35.5, 35.1, 33.1, 28.7, 21.9, 19.2; HRMS (ESI$^+$) *m/z* $[C_{25}H_{35}N_4O_6S]^+$ calcd: 519.2272; found: 519.2281.

Following general procedure H, **BCL-4-76** (433.0 mg, 0.83 mmol) and 80% HOAc (4 mL) were used. Purification by flash chromatography ($SiO_2$, DCM/MeOH = 20:1 to 10:1) gave **BCL-4-83** (205 mg, 61%) as a yellow semi-solid. $R_f$ = 0.16 (DCM/MeOH = 10:1); Purity: 98%, RT = 14.67 min with method 12, RT = 15.31 min with method 13; $^1$H NMR ($D_2O$, 500 MHz): $\delta$ 7.60 (d, $J$ =

3.4 Hz, 1H), 7.37 (d, $J$ = 3.3 Hz, 1H), 3.88 (s, 1H), 3.48-3.32 (m, 3H), 3.28-3.20 (m, 3H), 3.20-3.10 (m, 4H), 2.63 (t, $J$ = 7.1 Hz, 2H), 2.35 (t, $J$ = 6.6 Hz, 2H), 0.81 (s, 3H), 0.78 (s, 3H); $^{13}$C NMR ($D_2O$, 125 MHz): $\delta$ 175.1, 174.6, 174.1, 170.5, 141.4, 119.8, 75.7, 68.3, 38.7, 38.6, 38.4, 35.4, 35.30, 35.28, 28.0, 20.5, 19.1; HRMS (ESI$^+$) *m/z* $[C_{17}H_{29}N_4O_5S]^+$ calcd: 401.1453; found: 401.1856.

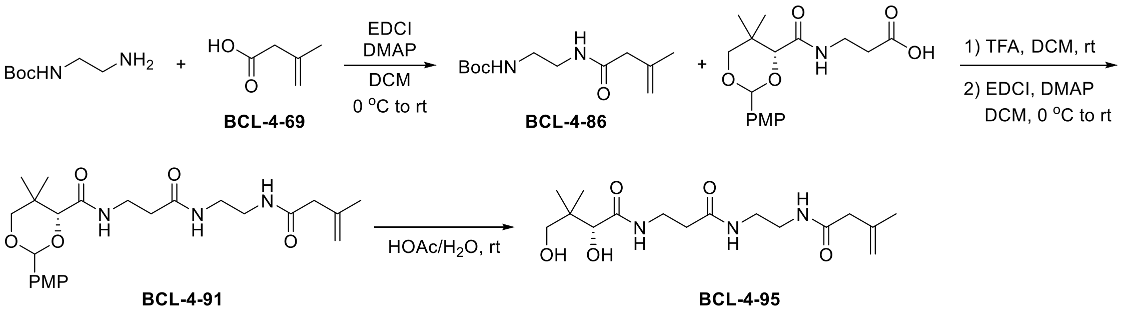

<u>Synthesis of **BCL-4-95 (3-06)**</u>

Following general procedure E, **BCL-4-69** (110.1 mg, 1.1 mmol), Boc-ethylenediamine (0.16 mL, 1.0 mmol), DMAP (12.2 mg, 0.1 mmol), EDCI (210.8 mg, 1.1 mmol), and DCM (5 mL) were used. Purification by flash chromatography ($SiO_2$, DCM/MeOH = 25:1) gave **BCL-4-86** (237 mg, 98%) as a white solid. $R_f$ = 0.60 (DCM/MeOH = 10:1); $^1$H NMR ($CDCl_3$, 500 MHz): $\delta$ 6.41 (bs, 1H), 5.02 (bs, 1H), 4.94 (m, 1H), 4.86 (m, 1H), 3.33 (m, 2H), 3.24 (m, 2H), 2.93 (d, $J$ = 1.1 Hz, 2H), 1.76 (t, $J$ = 1.2 Hz, 3H), 1.41 (s, 9H); $^{13}$C NMR ($CDCl_3$, 125 MHz): $\delta$ 171.3, 156.9, 140.1, 115.6, 79.7, 46.2, 40.7, 40.4, 28.5, 22.5; HRMS (ESI$^+$) *m/z* $[C_{12}H_{22}N_2O_3Na]^+$ calcd: 265.1523; found: 265.1517.

Following general procedure F, **BCL-4-86** (237.0 mg, 0.98 mmol), TFA (1.5 mL), and DCM (1.5 mL) were used. Next, general procedure G was followed using PMP-pantothenate (330.6 mg, 0.98 mmol), DMAP (12.2 mg, 0.1 mmol), EDCI (187.9 mmol, 0.98 mmol), and DCM (5 mL). Purification by flash chromatography ($SiO_2$, DCM/MeOH = 50:1 to 15:1) gave **BCL-4-91** (145 mg, 32% over two steps) as a white solid. $R_f$ = 0.23 (DCM/MeOH = 20:1); $^1$H NMR ($CDCl_3$, 500 MHz): $\delta$ 7.42 (d, $J$ = 8.8 Hz, 2H), 7.03 (m, 1H), 6.90 (d, $J$ = 8.7 Hz, 2H), 6.81 (t, $J$ = 4.8 Hz, 1H), 6.49 (m, 1H), 5.45 (s, 1H), 4.93 (m, 1H), 4.84 (m, 1H), 4.06 (s, 1H), 3.80 (s, 3H), 3.69 (d, $J$ = 11.4 Hz, 1H), 3.64 (d, $J$ = 11.4 Hz, 1H), 3.51 (m, 2H), 3.34-3.29 (m, 4H), 2.89 (d, $J$ = 1.1 Hz, 2H), 2.40 (t, $J$ = 6.3 Hz, 2H), 1.74 (s, 3H), 1.09 (s, 3H), 1.07 (s, 3H); $^{13}$C NMR ($CDCl_3$, 125 MHz): $\delta$ 172.0, 171.8, 169.7, 160.3, 140.0, 130.3, 127.6, 115.6, 113.8, 101.4, 84.0, 78.6, 55.4, 46.1, 39.98, 39.96,

36.1, 35.0, 33.2, 22.5, 22.0, 19.2; HRMS (ESI$^+$) *m/z* $[C_{24}H_{35}N_3O_6Na]^+$ calcd: 484.2418; found: 484.2409.

Following general procedure H, **BCL-4-91** (145.0 mg, 0.31 mmol) and 80% HOAc (2 mL) were used. Purification by flash chromatography ($SiO_2$, DCM/MeOH = 20:1 to 10:1) gave **BCL-4-95** (75 mg, 70%) as a yellow semi-solid. $R_f$ = 0.21 (DCM/MeOH = 10:1); Purity: 83%, RT = 14.85 min with method 12, RT = 15.46 min with method 13; $^1$H NMR ($D_2O$, 500 MHz): $\delta$ 4.98 (m, 1H), 4.88 (m, 1H), 3.98 (s, 1H), 3.53-3.46 (m, 3H), 3.39 (d, $J$ = 11.3 Hz, 1H), 3.33-3.30 (m, 4H), 2.99 (d, $J$ = 1.1 Hz, 2H), 2.48 (t, $J$ = 6.6 Hz, 2H), 1.74 (t, $J$ = 1.2 Hz, 3H), 0.92 (s, 3H), 0.88 (s, 3H); $^{13}$C NMR ($D_2O$, 125 MHz): $\delta$ 175.1, 174.6, 174.1, 139.8, 114.7, 75.8, 68.3, 44.9, 38.63, 38.63, 38.57, 35.4, 35.3, 21.3, 20.5, 19.1; HRMS (ESI$^+$) *m/z* $[C_{16}H_{29}N_3O_5Na]^+$ calcd: 366.1999; found: 366.1992.

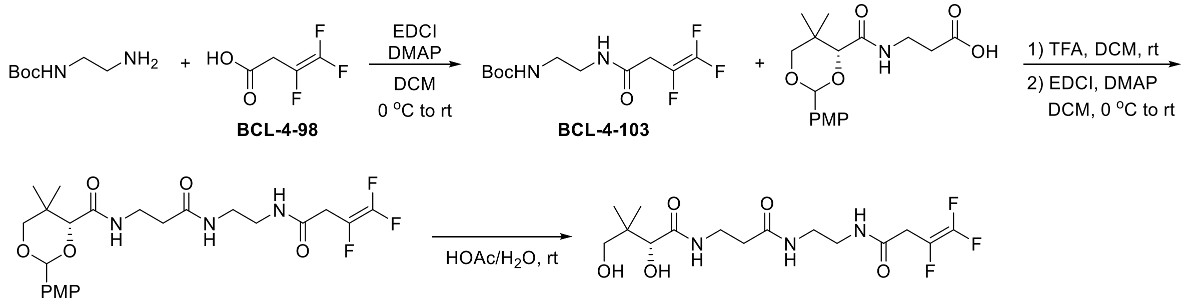

<u>Synthesis of **BCL-4-119 (4-08)**</u>

Following general procedure E, **BCL-4-98** (217.0 mg, 1.55 mmol), Boc-ethylenediamine (0.122 mL, 1.4 mmol), DMAP (19.5 mg, 0.16 mmol), EDCI (297.1 mg, 1.55 mmol), and DCM (7.5 mL) were used. Purification by flash chromatography ($SiO_2$, DCM/MeOH = 50:1 to 25:1) gave **BCL-4-103** (140 mg, 33%) as a white solid. $R_f$ = 0.60 (DCM/MeOH = 10:1); $^1$H NMR ($CDCl_3$, 500 MHz): $\delta$ 6.89 (bs, 1H), 4.97 (bs, 1H), 3.38 (m, 2H), 3.29 (m, 2H), 3.23 (dd, $J$ = 3.8, 2.5 Hz, 1H), 3.19 (dd, $J$ = 3.8, 2.6 Hz, 1H), 1.43 (s, 9H); $^{19}$F NMR ($CDCl_3$, 471 MHz): $\delta$ -101.7 (dd, $J$ = 79.5, 33.4 Hz), -121.1 (dd, $J$ = 115.3, 79.5 Hz), -124.9 (d, $J$ = 17.7 Hz); $^{13}$C NMR ($CDCl_3$, 125 MHz): $\delta$ 166.7, 157.6, 80.2, 42.1, 39.9, 34.1 (dd, $J$ = 23.3, 2.7 Hz), 28.34 HRMS (ESI$^+$) *m/z* $[C_{11}H_{17}N_2O_3F_3Na]^+$ calcd: 305.1084; found: 305.1084.

Following general procedure F, **BCL-4-103** (140.0 mg, 0.5 mmol), TFA (1 mL), and DCM (1 mL) were used. Next, general procedure G was followed using PMP-pantothenate (168.7 mg, 0.5 mmol), DMAP (6.1 mg, 0.05 mmol), EDCI (95.9 mmol, 0.5 mmol), and DCM (3 mL). Purification by flash chromatography ($SiO_2$, DCM/MeOH = 50:1 to 15:1) gave **BCL-4-112** (160 mg, 64% over two steps) as a beige solid. $R_f = 0.49$ (DCM/MeOH = 10:1); $^1$H NMR ($CDCl_3$, 500 MHz): $\delta$ 7.42 (d, $J = 8.7$ Hz, 2H), 7.11 (bs, 1H), 7.01 (t, $J = 6.3$ Hz, 1H), 6.91 (d, $J = 8.7$ Hz, 2H), 6.85 (bs, 1H), 5.45 (s, 1H), 4.06 (s, 1H), 3.80 (s, 3H), 3.70 (d, $J = 11.4$ Hz, 1H), 3.64 (d, $J = 11.5$ Hz, 1H), 3.46 (m, 2H), 3.35-3.30 (m, 4H), 3.17 (dd, $J = 3.8$, 2.5 Hz, 1H), 3.12 (dd, $J = 3.8$, 2.5 Hz, 1H), 2.40 (t, $J = 6.2$ Hz, 2H), 1.09 (s, 3H), 1.06 (s, 3H); $^{19}$F NMR ($CDCl_3$, 471 MHz): $\delta$ -101.8 (dd, $J = 79.7$, 33.1 Hz), -121.4 (dd, $J = 115.3$, 79.7 Hz), -124.9 (d, $J = 17.8$ Hz); $^{13}$C NMR ($CDCl_3$, 125 MHz): $\delta$ 172.5, 169.9, 167.0, 160.4, 130.2, 127.7, 113.8, 101.5, 83.9, 78.5, 55.4, 40.8, 39.3, 36.4, 35.1, 33.9 (d, $J = 23.3$ Hz), 33.2, 21.9, 19.2; HRMS (ESI$^+$) *m/z* $[C_{23}H_{30}N_3O_6F_3Na]^+$ calcd: 524.1979; found: 524.1976.

Following general procedure H, **BCL-4-112** (160.0 mg, 0.32 mmol) and 80% HOAc (2 mL) were used. Purification by flash chromatography ($SiO_2$, DCM/MeOH = 20:1 to 10:1) gave **BCL-4-119** (81 mg, 66%) as a white solid. $R_f = 0.10$ (DCM/MeOH = 10:1); Purity: 97%, RT = 15.53 min with method 12, RT = 15.78 min with method 13; $^1$H NMR ($D_2O$, 500 MHz): $\delta$ 3.98 (s, 1H), 3.55-3.44 (m, 3H), 3.47-3.29 (m, 7H), 2.48 (t, $J = 6.6$ Hz, 2H), 0.92 (s, 3H), 0.89 (s, 3H); $^{19}$F NMR ($CDCl_3$, 471 MHz): $\delta$ -102.5 (dd, $J = 81.8$, 32.5 Hz), -121.7 (dd, $J = 114.0$, 81.8 Hz), -125.6 (d, $J = 18.6$ Hz); $^{13}$C NMR ($D_2O$, 125 MHz): $\delta$ 175.1, 174.2, 170.1, 154.0 (ddd, $J = 283.8$, 45.0, 45.0 Hz), 124.4 (ddd, $J = 231.3$, 18.8, 18.8 Hz), 75.8, 68.3, 38.9, 38.6, 38.5, 35.4, 35.3, 33.0 (dd, $J = 22.8$, 2.7 Hz), 20.4, 19.0; HRMS (ESI$^+$) *m/z* $[C_{15}H_{24}N_3O_5F_3Na]^+$ calcd: 406.1560; found: 406.1560.

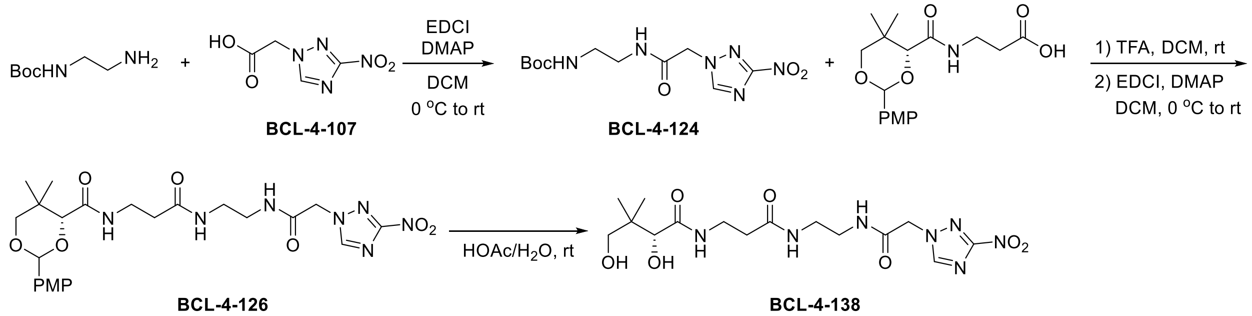

Synthesis of **BCL-4-138 (5-07)**

Following general procedure E, **BCL-4-107** (473.3 mg, 2.75 mmol), Boc-ethylenediamine (0.40 mL, 2.5 mmol), DMAP (30.5 mg, 0.25 mmol), EDCI (527.2 mg, 2.75 mmol), and DCM (10 mL) were used. Purification by flash chromatography ($SiO_2$, DCM/MeOH = 50:1 to 15:1) gave **BCL-4-124** (400 mg, 51%) as a white solid. $R_f = 0.45$ (DCM/MeOH = 10:1); $^1$H NMR ($CD_3OD$, 500 MHz): $\delta$ 8.60 (s, 1H), 5.05 (s, 2H), 3.27 (m, 2H), 3.14 (t, $J = 6.1$ Hz, 2H), 1.39 (s, 9H); $^{13}$C NMR

($CD_3OD$, 125 MHz): $\delta$ 167.3, 158.6, 148.8, 80.2, 53.8, 41.0, 40.6, 28.7; HRMS (ESI$^+$) *m/z* $[C_{11}H_{18}N_6O_5F_3Na]^+$ calcd: 337.1231; found: 337.1238.

Following general procedure F, **BCL-4-124** (400.0 mg, 1.76 mmol), TFA (2 mL), and DCM (2 mL) were used. Next, general procedure G was followed using PMP-pantothenate (593.8 mg, 1.76 mmol), DMAP (22.0 mg, 0.18 mmol), EDCI (337.4 mmol, 1.76 mmol), and DCM (15 mL). Purification by flash chromatography ($SiO_2$, DCM/MeOH = 50:1 to 15:1) gave **BCL-4-126** (554 mg, 59% over two steps) as a white solid. $R_f$ = 0.23 (DCM/MeOH = 20:1); $^1$H NMR ($CDCl_3$, 500 MHz): $\delta$ 8.32 (s, 1H), 7.70 (t, $J$ = 5.5 Hz, 1H), 7.40 (d, $J$ = 8.7 Hz, 2H), 7.06 (t, $J$ = 6.4 Hz, 1H), 6.91 (t, $J$ = 5.4 Hz, 1H), 6.87 (d, $J$ = 8.7 Hz, 2H), 5.45 (s, 1H), 4.90 (s, 2H), 4.07 (s, 1H), 3.78 (s, 3H), 3.69 (d, $J$ = 11.4 Hz, 1H), 3.64 (d, $J$ = 11.5 Hz, 1H), 3.48 (m, 2H), 3.33-3.28 (m, 4H), 2.37 (t, $J$ = 6.3 Hz, 2H), 1.07 (s, 3H), 1.03 (s, 3H); $^{13}$C NMR ($CDCl_3$, 125 MHz): $\delta$ 172.7, 170.1, 164.9, 162.7, 160.4, 147.0, 130.1, 127.7, 113.8, 101.4, 83.9, 78.4, 55.5, 53.3, 40.7, 39.0, 36.5, 35.4, 33.2, 22.0, 19.2; HRMS (ESI$^+$) *m/z* $[C_{23}H_{31}N_7O_8Na]^+$ calcd: 556.2126; found: 556.2151.

Following general procedure H, **BCL-4-126** (554.0 mg, 1.04 mmol) and 80% HOAc (6 mL) were used. Purification by flash chromatography ($SiO_2$, DCM/MeOH = 20:1 to 10:1) gave **BCL-4-138** (270 mg, 62%) as a white solid. $R_f$ = 0.05 (DCM/MeOH = 10:1); Purity: >99%, RT = 13.49 min with method 12, RT = 15.57 min with method 13; $^1$H NMR ($D_2O$, 500 MHz): $\delta$ 8.71 (s, 1H), 5.22 (s, 2H), 3.97 (s, 1H), 3.56-3.43 (m, 3H), 3.42-3.31 (m, 5H), 2.48 (t, $J$ = 6.6 Hz, 2H), 0.91 (s, 3H), 0.88 (s, 3H); $^{13}$C NMR ($D_2O$, 125 MHz): $\delta$ 175.1, 174.2, 167.1, 161.9, 148.1, 75.8, 68.3, 52.8, 39.0, 38.6, 38.5, 35.4, 35.3, 20.4, 19.1; HRMS (ESI$^+$) *m/z* $[C_{15}H_{25}N_7O_7Na]^+$ calcd: 438.1708; found: 438.1708.

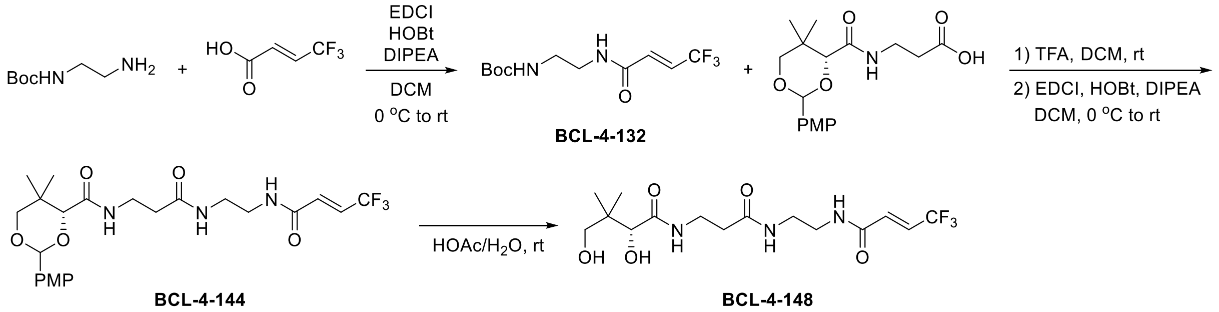

<u>Synthesis of **BCL-4-148 (4-04)**</u>

Following general procedure J, 4,4,4-trifluorocrotonic acid (154.1 mg, 1.1 mmol), Boc-ethylenediamine (0.16 mL, 1.0 mmol), HOBt (185.8 mg, 1.1 mmol), DIPEA (0.45 mL, 2.5 mmol), EDCI (210.9 mg, 1.1 mmol), and DCM (5 mL) were used. Purification by flash chromatography

(SiO$_2$, DCM/EA = 10:1) gave **BCL-4-132** (157 mg, 55%) as a white solid. $R_f$ = 0.58 (DCM/EA = 1:1); $^1$H NMR (CD$_3$OD, 500 MHz): $\delta$ 6.76-6.30 (m, 2H), 3.38 (t, $J$ = 6.1 Hz, 2H), 3.22 (t, $J$ = 6.1 Hz, 2H), 1.45 (s, 9H); $^{19}$F NMR (CD$_3$OD, 471 MHz): $\delta$ -66.6; $^{13}$C NMR (CD$_3$OD, 125 MHz): $\delta$ 165.2, 158.6, 132.9 (q, $J$ = 6.0 Hz), 128.2 (q, $J$ = 34.9 Hz), 124.2 (q, $J$ = 268.9 Hz), 80.2, 40.9, 40.6, 28.7; HRMS (ESI$^+$) $m/z$ [$C_{11}H_{17}N_2O_3F_3Na$]$^+$ calcd: 305.1089; found: 305.1064.

Following general procedure F, **BCL-4-132** (299.0 mg, 1.05 mmol), TFA (1.5 mL), and DCM (1.5 mL) were used. Next, general procedure K was followed using PMP-pantothenate (354.2 mg, 1.05 mmol), HOBt (177.3 mg, 1.05 mmol), DIPEA (0.46 mL, 2.6 mmol), EDCI (201.2 mg, 1.05 mmol), and DCM (5 mL). Purification by flash chromatography (SiO$_2$, DCM/MeOH = 50:1 to 15:1) gave **BCL-4-144** (428 mg, 81% over two steps) as a white solid. $R_f$ = 0.58 (DCM/MeOH = 20:1); $^1$H NMR (CD$_3$OD, 500 MHz): $\delta$ 7.68 (t, $J$ = 6.1 Hz, 1H), 7.48 (d, $J$ = 8.8 Hz, 2H), 6.94 (d, $J$ = 8.8 Hz, 1H), 6.81-6.67 (m, 2H), 5.54 (s, 1H), 4.16 (s, 1H), 3.81 (s, 3H), 3.74 (d, $J$ = 11.6 Hz, 1H), 3.69 (d, $J$ = 11.5 Hz, 1H), 3.50 (m, 2H), 3.37 (m, 2H), 3.31 (m, 2H), 2.42 (t, $J$ = 6.6 Hz, 2H), 1.13 (s, 3H), 1.06 (s, 3H); $^{19}$F NMR (CD$_3$OD, 471 MHz): $\delta$ -66.5 $^{13}$C NMR (CD$_3$OD, 125 MHz): $\delta$ 174.1, 171.6, 165.1, 161.7, 132.9 (q, $J$ = 6.0 Hz), 131.8, 128.8, 128.2 (q, $J$ = 34.8 Hz), 124.2 (q, $J$ = 268.9 Hz), 114.5, 102.7, 85.1, 79.3, 55.7, 40.3, 39.7, 36.4, 36.3, 34.0, 22.2, 19.7; HRMS (ESI$^+$) $m/z$ [$C_{23}H_{30}N_3O_6F_3Na$]$^+$ calcd: 524.1979; found: 524.1975.

Following general procedure H, **BCL-4-144** (428.0 mg, 0.85 mmol) and 80% HOAc (5 mL) were used. Purification by flash chromatography (SiO$_2$, DCM/MeOH = 20:1 to 10:1) gave **BCL-4-148** (165 mg, 51%) as a white solid. $R_f$ = 0.20 (DCM/MeOH = 10:1); Purity: 97%, RT = 16.91 min with method 12, RT = 16.57 min with method 13; $^1$H NMR (D$_2$O, 500 MHz): $\delta$ 6.78-6.69 (m, 2H), 3.97 (s, 1H), 3.53-3.24 (m, 8H), 2.48 (t, $J$ = 6.6 Hz, 2H), 0.91 (s, 3H), 0.88 (s, 3H); $^{19}$F NMR (D$_2$O, 471 MHz): $\delta$ -65.0; $^{13}$C NMR (D$_2$O, 125 MHz): $\delta$ 175.1, 174.2, 165.7, 130.9 (q, $J$ = 6.1 Hz), 127.7 (q, $J$ = 34.8 Hz), 122.5 (q, $J$ = 269.5 Hz), 75.8, 68.3, 39.0, 38.6, 38.5, 35.4, 35.3, 20.4, 19.0; HRMS (ESI$^+$) $m/z$ [$C_{15}H_{24}N_3O_5F_3Na$]$^+$ calcd: 406.1560; found: 406.1560.

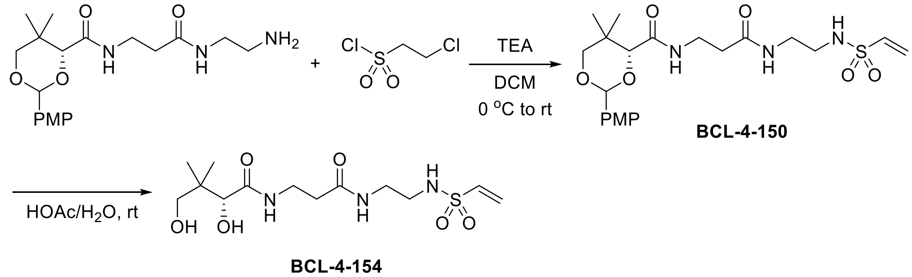

BCL-4-150

BCL-4-154

Synthesis of **BCL-4-154 (3-07)**

To a 25 mL round bottom flask were added PMP-$NH_2$ (189.7 mg, 0.5 mmol, 1.0 equiv.), TEA (0.28 mL, 2.0 mmol, 4.0 equiv.), and DCM (2 mL). The mixture was cooled to 0°C and a 3-chlorosulfonyl chloride solution (63 μL dissolved in 1 mL DCM, 0.6 mmol, 1.2 equiv.) was added dropwise. The reaction mixture was stirred at rt for 2 h. The mixture was then diluted in DCM (20 mL), washed with sat. $NaHCO_3$ (2 × 20 mL), 1 M $KH_2PO_4$ (2 × 20 mL), and brine (1 × 20 mL). The organic layer was dried over anhydrous $Na_2SO_4$, filtered, and the solvent was evaporated under vacuum. Purification by flash chromatography ($SiO_2$, DCM/MeOH = 50:1 to 15:1) gave **BCL-4-150** (205 mg, 87%) as a light yellow solid. $R_f$ = 0.49 (DCM/MeOH = 10:1); $^1$H NMR ($CDCl_3$, 500 MHz): $\delta$ 7.42 (d, $J$ = 8.7 Hz, 2H), 7.05 (t, $J$ = 6.4 Hz, 1H), 6.91 (d, $J$ = 8.7 Hz, 2H), 6.90 (m, 1H), 6.46 (dd, $J$ = 16.5, 9.9 Hz, 1H), 6.16 (d, $J$ = 16.6 Hz, 1H), 5.88 (m, 2H), 5.45 (s, 1H), 4.09 (s, 1H), 3.80 (s, 3H), 3.68 (d, $J$ = 11.4 Hz, 1H), 3.64 (d, $J$ = 11.5 Hz, 1H), 3.51 (m, 2H), 3.34 (m, 2H), 3.07 (m, 2H), 2.42 (m, 2H), 1.08 (s, 3H), 1.05 (s, 3H); $^{13}$C NMR ($CDCl_3$, 125 MHz): $\delta$ 172.0, 170.0, 160.4, 135.9, 130.2, 127.7, 126.6, 113.8, 101.4, 83.9, 78.5, 55.4, 42.6, 39.8, 36.3, 35.3, 33.2, 21.9, 19.6; HRMS (ESI$^+$) *m/z* $[C_{21}H_{31}N_3O_7SNa]^+$ calcd: 492.1775; found: 492.1760.

Following general procedure H, **BCL-4-150** (205.0 mg, 0.43 mmol) and 80% HOAc (3 mL) were used. Purification by flash chromatography ($SiO_2$, DCM/MeOH = 20:1 to 10:1) gave **BCL-4-154** (70 mg, 46%) as a sticky yellow oil. $R_f$ = 0.13 (DCM/MeOH = 10:1); Purity: 97%, RT = 12.99 min with method 12, RT = 15.12 min with method 13; $^1$H NMR ($D_2O$, 500 MHz): $\delta$ 6.70 (dd, $J$ = 16.5, 10.1 Hz, 1H), 6.26 (d, $J$ = 16.6 Hz, 1H), 6.13 (d, $J$ = 10.1 Hz, 1H), 3.99 (s, 1H), 3.59-3.46 (m, 3H), 3.40 (d, $J$ = 11.3 Hz, 1H), 3.33 (t, $J$ = 6.0 Hz, 2H), 3.13 (t, $J$ = 6.0 Hz, 2H), 2.52 (t, $J$ = 6.5 Hz, 2H), 0.93 (s, 3H), 0.89 (s, 3H); $^{13}$C NMR ($D_2O$, 125 MHz): $\delta$ 175.1, 174.2, 134.3, 128.2, 75.8, 68.4, 41.4, 39.3, 38.6, 35.4, 35.3, 20.5, 19.1; HRMS (ESI$^+$) *m/z* $[C_{13}H_{25}N_3O_6SNa]^+$ calcd: 374.1356; found: 374.1344.

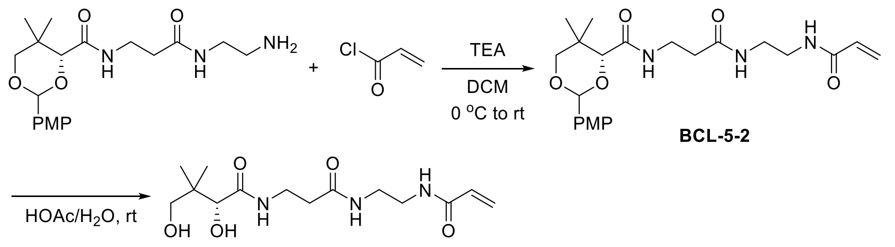

BCL-5-2

BCL-5-9

Synthesis of **BCL-5-9 (3-03)**

To a 25 mL round bottom flask were added PMP-$NH_2$ (189.7 mg, 0.5 mmol, 1.0 equiv.), TEA (105 μL, 0.75 mmol, 1.5 equiv.), and DCM (2 mL). The mixture was cooled to 0°C and an acryloyl chloride solution (49 μL dissolved in 1 mL DCM, 0.6 mmol, 1.2 equiv.) was added dropwise. The reaction mixture was stirred at rt overnight. The mixture was then diluted in DCM (20 mL), washed with sat. $NaHCO_3$ (2 × 20 mL), 1 M $KH_2PO_4$ (2 × 20 mL), and brine (1 × 20 mL). The organic layer was dried over anhydrous $Na_2SO_4$, filtered, and evaporated under vacuum. Purification by flash chromatography ($SiO_2$, DCM/MeOH = 50:1 to 15:1) gave **BCL-5-2** (136 mg, 63%) as a sticky white semi-solid. $R_f$ = 0.52 (DCM/MeOH = 10:1); $^1$H NMR ($CDCl_3$, 500 MHz): $\delta$ 7.42 (d, $J$ = 8.8 Hz, 2H), 7.04 (t, $J$ = 5.6 Hz, 1H), 6.98 (bs, 1H), 6.90 (d, $J$ = 8.7 Hz, 2H), 6.85 (bs, 1H), 6.22 (d, $J$ = 17.1 Hz, 1H), 6.07 (dd, $J$ = 17.0, 10.3 Hz, 1H), 5.59 (d, $J$ = 10.3 Hz, 1H), 5.44 (s, 1H), 4.06 (s, 1H), 3.80 (s, 3H), 3.69 (d, $J$ = 11.3 Hz, 1H), 3.64 (d, $J$ = 11.4 Hz, 1H), 3.51 (m, 2H), 3.41-3.29 (m, 4H), 2.40 (t, $J$ = 6.3 Hz, 2H), 1.08 (s, 3H), 1.06 (s, 3H); $^{13}$C NMR ($CDCl_3$, 125 MHz): $\delta$ 172.2, 169.7, 166.6, 160.4, 130.9, 130.2, 127.7, 126.6, 113.8, 101.4, 84.0, 78.5, 55.5, 40.2, 39.8, 36.2, 35.1, 33.2, 22.0, 19.2; HRMS (ESI$^+$) $m/z$ $[C_{22}H_{31}N_3O_6Na]^+$ calcd: 456.2105; found: 456.2109.

Following general procedure H, **BCL-5-2** (309.0 mg, 0.71 mmol) and 80% HOAc (4 mL) were used. Purification by flash chromatography ($SiO_2$, DCM/MeOH = 20:1 to 10:1) gave **BCL-5-9** (110 mg, 50%) as a sticky white solid. $R_f$ = 0.10 (DCM/MeOH = 10:1); Purity: 98%, RT = 11.20 min with method 12, RT = 14.65 min with method 13; $^1$H NMR ($D_2O$, 500 MHz): $\delta$ 6.26 (dd, $J$ = 17.1, 10.0 Hz, 1H), 6.18 (d, $J$ = 17.2 Hz, 1H), 5.77 (d, $J$ = 9.9 Hz, 1H), 3.97 (s, 1H), 3.56-3.43 (m, 3H), 3.43-3.31 (m, 5H), 2.48 (t, $J$ = 6.6 Hz, 2H), 0.91 (s, 3H), 0.87 (s, 3H); $^{13}$C NMR ($D_2O$, 125 MHz): $\delta$ 175.1, 174.2, 168.8, 129.9, 127.4, 75.7, 68.3, 38.7, 38.7, 38.6, 35.4, 35.3, 20.5, 19.0; HRMS (ESI$^+$) $m/z$ $[C_{14}H_{25}N_3O_5Na]^+$ calcd: 338.1686; found: 338.1681.

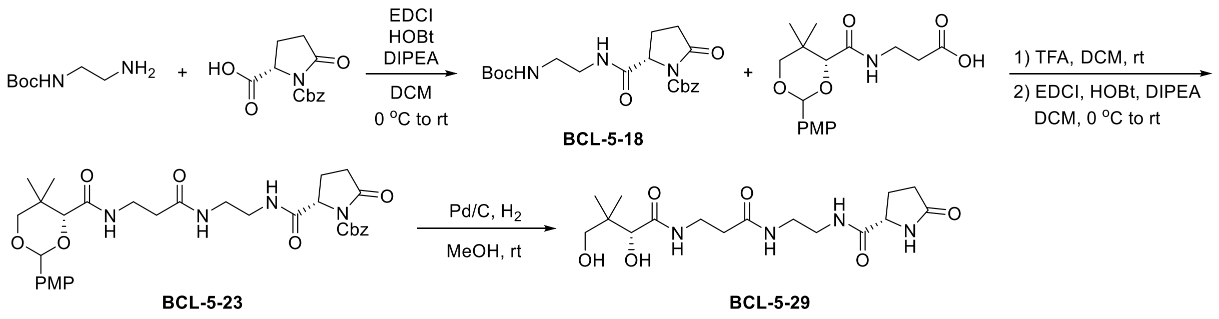

Synthesis of **BCL-5-29 (5-05)**

Following general procedure J, Z-Pyr-OH (289.5 mg, 1.1 mmol), Boc-ethylenediamine (0.16 mL, 1.0 mmol), HOBt (185.6 mg, 1.1 mmol), DIPEA (0.45 mL, 2.5 mmol), EDCI (210.9 mg, 1.1 mmol), and DCM (5 mL) were used. Purification by flash chromatography ($SiO_2$, DCM/MeOH = 50:1 to 15:1) gave **BCL-5-18** (367 mg, 91%) as a white solid. $R_f$ = 0.55 (DCM/MeOH = 10:1); $^1$H NMR ($CDCl_3$, 500 MHz): $\delta$ 7.47 (bs, 1H), 7.32-7.27 (m, 5H), 5.28 (bs, 1H), 5.20 (d, *J* = 12.3 Hz, 1H), 5.12 (d, *J* = 12.3 Hz, 1H), 4.56 (dd, *J* = 9.4, 3.0 Hz, 1H), 3.35 (m, 2H), 3.22 (m, 2H), 2.69 (dt, *J* = 17.6, 9.8 Hz, 1H), 2.41 (ddd, *J* = 17.6, 9.5, 3.5 Hz, 1H), 2.28 (dq, *J* = 13.3, 9.6 Hz, 1H), 2.12 (ddt, *J* = 13.2, 9.8, 3.4 Hz, 1H), 1.37 (s, 9H); $^{13}$C NMR ($CDCl_3$, 125 MHz): $\delta$ 175.2, 171.1, 156.6, 151.0, 134.8, 128.7, 128.6, 128.1, 79.3, 68.4, 60.7, 40.5, 40.3, 31.4, 28.5, 22.7; HRMS (ESI$^+$) *m/z* $[C_{20}H_{27}N_3O_6Na]^+$ calcd: 428.1792; found: 428.1795.

Following general procedure F, **BCL-5-18** (367.0 mg, 0.91 mmol), TFA (2 mL), and DCM (2 mL) were used. Next, general procedure K was used with PMP-pantothenate (307.2 mg, 0.91 mmol), HOBt (153.7 mg, 0.91 mmol), DIPEA (0.41 mL, 2.28 mmol), EDCI (174.4 mg, 0.91 mmol), and DCM (5 mL) Purification by flash chromatography ($SiO_2$, DCM/MeOH = 50:1 to 15:1) gave **BCL-5-23** (352 mg, 62% over two steps) as a white foam. $R_f$ = 0.40 (DCM/MeOH = 10:1); $^1$H NMR ($CDCl_3$, 500 MHz): $\delta$ 7.50 (bs, 1H), 7.41 (d, *J* = 8.8 Hz, 2H), 7.31-7.28 (m, 5H), 7.00 (t, *J* = 6.3 Hz, 1H), 6.89 (d, *J* = 8.7 Hz, 2H), 6.64 (bs, 1H), 5.42 (s, 1H), 5.21 (d, *J* = 12.4 Hz, 1H), 5.11 (d, *J* = 12.4 Hz, 1H), 4.53 (dd, *J* = 9.3, 3.1 Hz, 1H), 4.02 (s, 1H), 3.79 (s, 3H), 3.67 (d, *J* = 11.4 Hz, 1H), 3.61 (d, *J* = 11.4 Hz, 1H), 3.50 (m, 1H), 3.38 (m, 2H), 3.33-3.21 (m, 3H), 2.67 (dt, *J* = 17.5, 9.7 Hz, 1H), 2.42 (ddd, *J* = 17.6, 9.6, 3.7 Hz, 1H), 2.33-2.19 (m, 3H), 2.07 (ddt, *J* = 13.1, 9.7, 3.6 Hz, 1H), 1.07 (s, 3H), 1.02 (s, 3H); $^{13}$C NMR ($CDCl_3$, 125 MHz): $\delta$ 174.9, 171.7, 171.4, 169.7, 160.3, 151.4, 134.9, 130.2, 128.7, 128.6, 127.9, 127.6, 113.8, 101.4, 83.9, 78.5, 68.4, 60.5, 55.2, 39.9, 39.2, 36.2, 35.3, 33.1, 31.5, 22.5, 21.9, 19.2; HRMS (ESI$^+$) *m/z* $[C_{32}H_{40}N_4O_9Na]^+$ calcd: 647.2687; found: 647.2684.

Following general procedure I, **BCL-5-23** (352.0 mg, 0.56 mmol), Pd/C (30 mg), and MeOH (2 mL) were used. The residue was dissolved in water (10 mL) and washed with DCM (3 × 10 mL) and EA (3 × 10 mL). The water was removed under vacuum, and the residue was lyophilized to give **BCL-5-29** (141 mg, 67%) as a white solid. Purity: 95%, RT = 10.06 min with method 12, RT = 14.22 min with method 13; $^1$H NMR ($D_2O$, 500 MHz): $\delta$ 4.30 (dd, *J* = 9.0, 5.1 Hz, 1H), 3.98 (s, 1H), 3.58-3.43 (m, 3H), 3.43-3.26 (m, 5H), 2.59-2.32 (m, 5H), 2.05 (m, 1H), 0.92 (s, 3H), 0.89 (s,

3H); $^{13}$C NMR ($D_2O$, 125 MHz): $\delta$ 182.3, 175.2, 175.1, 174.2, 75.8, 68.3, 57.1, 38.8, 38.58, 38.55, 35.44, 35.35, 29.2, 25.2, 20.5, 19.0; HRMS (ESI$^+$) *m/z* $[C_{16}H_{28}N_4O_6Na]^+$ calcd: 395.1901; found: 395.1895.

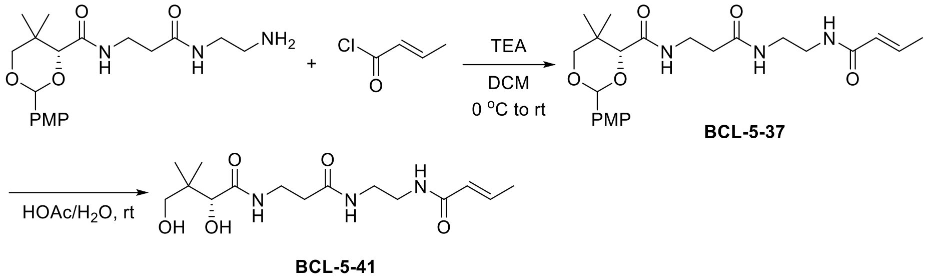

Synthesis of **BCL-5-41 (3-04)**

To a 25 mL round bottom flask were added PMP-$NH_2$ (189.7 mg, 0.5 mmol, 1.0 equiv.), TEA (105 μL, 0.75 mmol, 1.5 equiv.), and DCM (2 mL). The mixture was cooled to 0°C and a crotonyl chloride solution (70 μL dissolved in 1 mL DCM, 0.6 mmol, 1.2 equiv.) was added dropwise. The reaction mixture was stirred at rt overnight. The mixture was then diluted in DCM (20 mL), washed with sat. $NaHCO_3$ (2 × 20 mL), 1 M $KH_2PO_4$ (2 × 20 mL), and brine (1 × 20 mL). The organic layer was dried over anhydrous $Na_2SO_4$, filtered, and the solvent was evaporated under vacuum. Purification by flash chromatography ($SiO_2$, DCM/MeOH = 50:1 to 15:1) gave **BCL-5-37** (137 mg, 61%) as a sticky beige solid. $R_f$ = 0.44 (DCM/MeOH = 10:1); $^1$H NMR ($CDCl_3$, 500 MHz): $\delta$ 7.40 (d, $J$ = 8.7 Hz, 2H), 7.19 (t, $J$ = 6.2 Hz, 1H), 7.06 (t, $J$ = 6.2 Hz, 1H), 6.88 (d, $J$ = 8.8 Hz, 2H), 6.79 (t, $J$ = 5.0 Hz, 1H), 6.75 (dq, $J$ = 15.3, 6.9 Hz, 1H), 5.77 (dq, $J$ = 15.2, 1.7 Hz, 1H), 5.42 (s, 1H), 4.03 (s, 1H), 3.77 (s, 3H), 3.66 (d, $J$ = 11.4 Hz, 1H), 3.61 (d, $J$ = 11.4 Hz, 1H), 3.48 (m, 2H), 3.36-3.24 (m, 4H), 2.37 (t, $J$ = 6.4 Hz, 2H), 1.78 (dd, $J$ = 6.9, 1.7 Hz, 3H), 1.06 (s, 3H), 1.04 (s, 3H); $^{13}$C NMR ($CDCl_3$, 125 MHz): $\delta$ 172.1, 169.5, 167.0, 160.3, 139.9, 130.2, 127.6, 125.1, 113.7, 101.3, 83.9, 78.5, 55.4, 39.82, 39.76, 36.0, 35.1, 33.1, 21.9, 19.2, 17.7; HRMS (ESI$^+$) *m/z* $[C_{23}H_{33}N_3O_6Na]^+$ calcd: 470.2262; found: 470.2266.

Following general procedure H, **BCL-5-37** (137.0 mg, 0.31 mmol) and 80% HOAc (2 mL) were used. Purification by flash chromatography ($SiO_2$, DCM/MeOH = 20:1 to 10:1) gave **BCL-5-41** (54 mg, 53%) as a sticky white solid. $R_f$ = 0.05 (DCM/MeOH = 10:1); Purity: 98%, RT = 13.30 min with method 12, RT = 15.12 min with method 13; $^1$H NMR ($D_2O$, 500 MHz): $\delta$ 6.78 (m, 1H), 5.96 (m, 1H), 3.97 (s, 1H), 3.56-3.42 (m, 3H), 3.42-3.29 (m, 5H), 2.48 (t, $J$ = 6.5 Hz, 2H), 1.86 (m, 3H), 0.91 (s, 3H), 0.87 (s, 3H); $^{13}$C NMR ($D_2O$, 125 MHz): $\delta$ 175.1, 174.2, 169.2, 142.0, 123.7,

75.7, 68.3, 38.8, 38.6, 38.5, 35.4, 35.3, 20.5, 19.0, 17.2; HRMS (ESI$^+$) *m/z* $[C_{15}H_{27}N_3O_5Na]^+$ calcd: 352.1843; found: 352.1849.

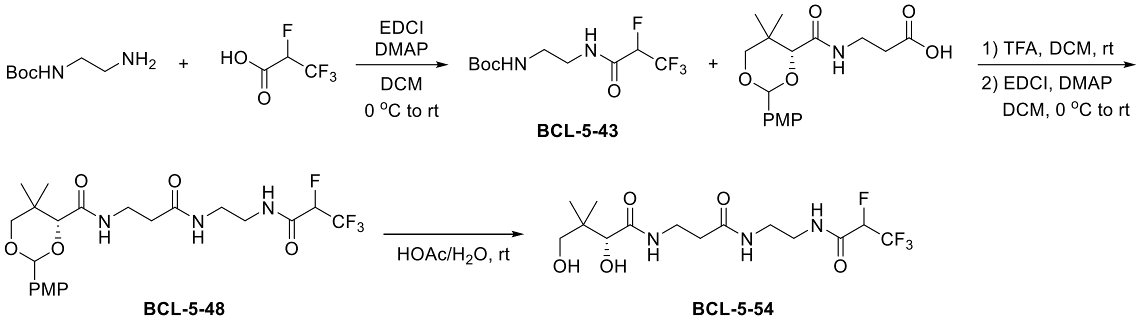

Synthesis of **BCL-5-54 (4-07)**

Following general procedure E, 2,3,3,3-tetrafluoropropanoic acid (160.6 mg, 1.1 mmol), Boc-ethylenediamine (0.16 mL, 1.0 mmol), DMAP (12.2 mg, 0.1 mmol), EDCI (210.9 mg, 1.1 mmol), and DCM (5 mL) were used. Purification by flash chromatography ($SiO_2$, Hex/EA = 4:1 to 2:1) gave **BCL-5-43** (213 mg, 74%) as a white solid. $R_f = 0.23$ (Hex/Ea = 2:1); $^1$H NMR ($CDCl_3$, 500 MHz): $\delta$ 7.49 (bs, 1H), 5.05 (dq, $J$ = 46.4, 6.4 Hz, 1H), 5.00 (bs, 1H), 3.44 (m, 2H), 3.31 (m, 2H), 1.42 (s, 9H); $^{19}$F NMR ($CDCl_3$, 471 MHz): $\delta$ -76.09, -76.11; $^{13}$C NMR ($CDCl_3$, 125 MHz): $\delta$ 162.0 (d, $J$ = 19.3 Hz), 157.3, 120.7 (qd, $J$ = 283.3, 27.5 Hz), 85.7 (dq, $J$ = 204.2, 33.8 Hz), 80.3, 41.3, 39.5, 28.3; HRMS (ESI$^+$) *m/z* $[C_{10}H_{16}N_2O_3F_4Na]^+$ calcd: 311.0986; found: 311.0985.

Following general procedure F, **BCL-5-43** (213.0 mg, 0.74 mmol), TFA (1.5 mL), and DCM (1.5 mL) were used. Next, general procedure G was followed using PMP-pantothenate (250.0 mg, 0.74 mmol), DMAP (8.6 mg, 0.07 mmol), EDCI (141.9 mmol, 0.74 mmol), and DCM (4 mL). Purification by flash chromatography ($SiO_2$, DCM/MeOH = 50:1 to 20:1) gave **BCL-5-48** (320 mg, 85% over two steps) as a white solid. $R_f = 0.63$ (DCM/MeOH = 10:1); $^1$H NMR ($CDCl_3$, 500 MHz): $\delta$ 7.71 (bs, 1H), 7.41 (d, $J$ = 8.7 Hz, 2H), 7.01 (t, $J$ = 6.5 Hz, 1H), 6.94 (bs, 1H), 6.90 (d, $J$ = 8.7 Hz, 2H), 5.44 (s, 1H), (5.01, 5.00) (dq, $J$ = 46.2, 6.6 Hz, 1H), 4.05 (s, 1H), 3.80 (s, 3H), 3.69 (d, $J$ = 11.4 Hz, 1H), 3.64 (d, $J$ = 11.4 Hz, 1H), 3.50 (m, 2H), 3.46-3.31 (m, 4H), 2.41 (m, 2H), 1.10-1.03 (m, 6H); $^{19}$F NMR ($CDCl_3$, 471 MHz): $\delta$ -75.98, -76.00; $^{13}$C NMR ($CDCl_3$, 125 MHz): $\delta$ (172.45, 172.42), 170.0, 162.0 (d, $J$ = 19.4 Hz), 160.4, (130.18, 130.16), (127.7, 127.6), 120.8 (qd, $J$ = 282.4, 25.5 Hz), 113.8, (101.50, 101.45), 85.6 (dq, $J$ = 203.7, 33.9 Hz), (83.92, 83.88), 78.5, 55.4, 40.3, 39.0, 36.3, 35.1, 33.2, 21.9, 19.2; HRMS (ESI$^+$) *m/z* $[C_{22}H_{29}N_3O_6F_4Na]^+$ calcd: 530.1885; found: 530.1888.

Following general procedure H, **BCL-5-48** (320.0 mg, 0.63 mmol) and 80% HOAc (4 mL) were used. Purification by flash chromatography ($SiO_2$, DCM/MeOH = 20:1 to 10:1) gave **BCL-5-54** (117 mg, 48%) as a white solid. $R_f$ = 0.15 (DCM/MeOH = 10:1); Purity: 98%, RT = 15.98 min with method 12, RT = 15.84 min with method 13; $^1$H NMR ($D_2O$, 500 MHz): $\delta$ 5.53 (dq, $J$ = 44.9, 6.6 Hz, 1H), 3.98 (s, 1H), 3.55-3.30 (m, 8H), 2.48 (t, $J$ = 6.6 Hz, 2H), 0.92 (s, 3H), 0.89 (s, 3H); $^{19}$F NMR ($D_2O$, 471 MHz): $\delta$ -76.32, -76.34; $^{13}$C NMR ($D_2O$, 125 MHz): $\delta$ 175.1, 174.2, 164.0 (d, $J$ = 20.2 Hz), 120.7 (qd, $J$ = 281.6, 25.8 Hz), 85.0 (dq, $J$ = 199.0, 33.7 Hz), 75.8, 68.3, 38.6, 38.6, 38.3, 35.4, 35.2, 20.4, 19.0; HRMS (ESI$^+$) *m/z* $[C_{14}H_{23}N_3O_5F_4Na]^+$ calcd: 412.1466; found: 412.1460.

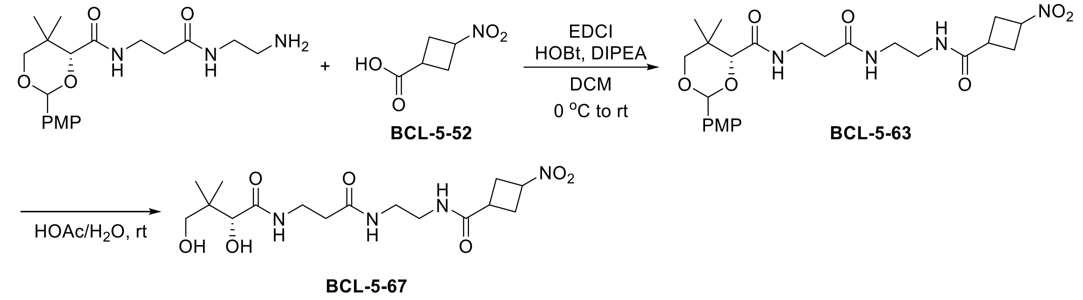

Synthesis of **BCL-5-67 (1-10)**

Following general procedure J, 3-nitrocyclobutane-1-carboxylic acid (120.4 mg, 0.83 mmol), PMP-$NH_2$ (284.6 mg, 0.75 mmol), HOBt (140.2 mg, 0.83 mmol), DIPEA (0.33 mL, 1.88 mmol), EDCI (159.1 mg, 0.83 mmol), and DCM (5 mL) were used. Purification by flash chromatography ($SiO_2$, DCM/MeOH = 50:1 to 15:1) gave **BCL-5-63** (310 mg, 82%) as a white solid. $R_f$ = 0.52/0.48 (DCM/MeOH = 10:1); $^1$H NMR ($CDCl_3$, 500 MHz): $\delta$ (7.41, 7.40) (d, $J$ = 8.9 Hz, 2H), 7.15-6.92 (m, 3H), 6.89 (d, $J$ = 8.7 Hz, 2H), 5.44 (s, 1H), (5.07. 4.72) (m, 1H), 4.05 (s, 1H), 3.79 (s, 3H), 3.68 (d, $J$ = 13.2 Hz, 1H), 3.64 (d, $J$ = 13.1 Hz, 1H), 3.50 (m, 2H), 3.32-3.29 (m, 4H), 3.14-2.54 (m, 5H), 2.43-2.36 (m, 2H), 1.34-0.80 (m, 6H); $^{13}$C NMR ($CDCl_3$, 125 MHz): $\delta$ (174.3, 172.8), (172.5, 172.2), (169.8, 169.7), (160.34, 160.32), (130.20, 130.15), 127.6, (113.79, 113.78), (101.4, 101.3), 83.9, 78.4, (76.1, 72.3), 55.4, (40.6, 40.4), (39.6, 39.4), (36.3, 36.2), (35.1, 34.2), 33.1, (31.6, 31.5), (31.34, 31.32), 31.2, 21.9, 19.2; HRMS (ESI$^+$) *m/z* $[C_{24}H_{34}N_4O_8Na]^+$ calcd: 529.2259; found: 529.2258.

Following general procedure H, **BCL-5-63** (310.0 mg, 0.61 mmol) and 80% HOAc (4 mL) were used. Purification by flash chromatography ($SiO_2$, DCM/MeOH = 20:1 to 10:1) gave **BCL-5-67** (156 mg, 66%) as a white solid. $R_f$ = 0.15/0.10 (DCM/MeOH = 10:1); Purity: 99%, RT = 14.63

min (66%) and 15.32 min (33%) with method 12, RT = 15.49 min (64%) and 15.69 min (35%) with method 13; $^1$H NMR ($D_2O$, 500 MHz): $\delta$ (5.20, 5.07) (m, 1H), 3.98 (s, 1H), 3.57-3.43 (m, 3H), 3.39 (d, $J$ = 11.3 Hz, 1H), 3.34-3.31 (m, 4H), 3.01-2.72 (m, 5H), 2.49 (m, 2H), 0.92 (s, 3H), 0.89 (s, 3H); $^{13}$C NMR ($D_2O$, 125 MHz): $\delta$ (176.9, 175.8), 175.1, 174.1, (76.2, 75.8), 72.5, 68.3, (38.77, 38.75), (38.64, 38.62), 38.6, 35.5, 35.3, 33.8, 31.2, (30.73, 30.68), 20.5, 19.1; HRMS (ESI$^+$) *m/z* $[C_{16}H_{28}N_4O_7Na]^+$ calcd: 411.1850; found: 411.1840.

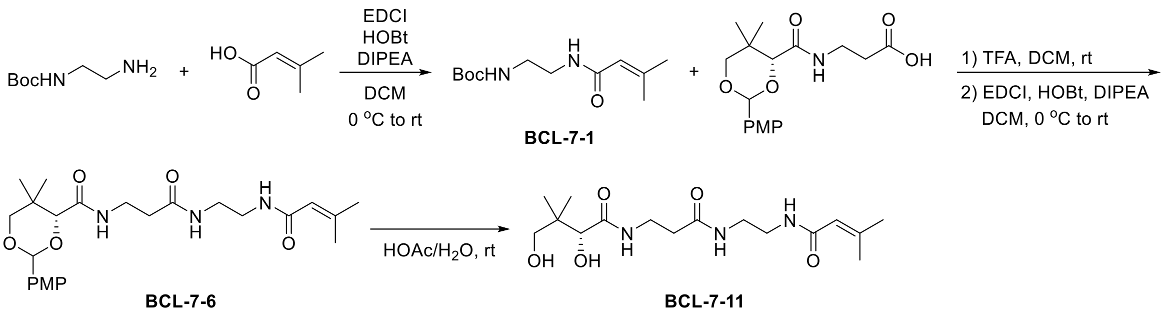

<u>Synthesis of **BCL-7-11 (3-05)**</u>

Following general procedure J, 3,3-dimethylacrylic acid (110.1 mg, 1.1 mmol), Boc-ethylenediamine (0.16 mL, 1.0 mmol), HOBt (185.8 mg, 1.1 mmol), DIPEA (0.45 mL, 2.5 mmol), EDCI (210.9 mg, 1.1 mmol), and DCM (5 mL) were used. Purification by flash chromatography ($SiO_2$, DCM/MeOH = 50:1 to 20:1) gave **BCL-7-1** (218 mg, 90%) as a white solid. $R_f$ = 0.60 (DCM/MeOH = 10:1); $^1$H NMR ($CDCl_3$, 500 MHz): $\delta$ 6.29 (bs, 1H), 5.54 (s, 1H), 5.19 (bs, 1H), 3.34 (m, 2H), 3.23 (m, 2H), 2.10 (s, 3H), 1.78 (s, 3H), 1.39 (s, 9H); $^{13}$C NMR ($CDCl_3$, 125 MHz): $\delta$ 167.8, 156.9, 150.9, 118.5, 79.5, 40.5, 40.1, 28.4, 27.2, 19.8; HRMS (ESI$^+$) *m/z* $[C_{12}H_{22}N_2O_3Na]^+$ calcd: 265.1523; found: 265.1514.

Following general procedure F, **BCL-7-1** (218.0 mg, 0.9 mmol), TFA (1 mL), and DCM (1 mL) were used. Next, general procedure K was followed using PMP-pantothenate (334.0 mg, 0.99 mmol), HOBt (189.7 mg, 0.99 mmol), DIPEA (0.40 mL, 2.25 mmol), EDCI (189.7 mg, 0.99 mmol), and DCM (5 mL). Purification by flash chromatography ($SiO_2$, DCM/MeOH = 50:1 to 20:1) gave **BCL-7-6** (363 mg, 87% over two steps) as a white foam. $R_f$ = 0.53 (DCM/MeOH = 10:1); $^1$H NMR ($CDCl_3$, 500 MHz): $\delta$ 7.40 (d, $J$ = 8.7 Hz, 2H), 7.12 (bs, 1H), 7.06 (t, $J$ = 6.3 Hz, 1H), 6.88 (d, $J$ = 8.7 Hz, 2H), 6.48 (bs, 1H), 5.52 (s, 1H), 5.42 (s, 1H), 4.03 (s, 1H), 3.77 (s, 3H), 3.66 (d, $J$ = 11.4 Hz, 1H), 3.61 (d, $J$ = 11.5 Hz, 1H), 3.47 (m, 2H), 3.31-3.25 (m, 4H), 2.37 (t, $J$ = 6.4 Hz, 2H), 2.09 (s, 3H), 1.77 (s, 3H), 1.06 (s, 3H), 1.04 (s, 3H); $^{13}$C NMR ($CDCl_3$, 125 MHz): $\delta$ 172.0, 169.5, 168.0, 160.2, 151.3, 130.2, 127.6, 118.3, 113.8, 101.3, 83.9, 78.5, 55.4, 40.1, 39.4,

35.9, 35.0, 33.1, 27.2, 21.9, 19.8, 19.2; HRMS (ESI$^+$) *m/z* $[C_{24}H_{35}N_3O_6Na]^+$ calcd: 484.2418; found: 484.2428.

Following general procedure H, **BCL-7-6** (363.0 mg, 0.78 mmol) and 80% HOAc (4 mL) were used. Purification by flash chromatography ($SiO_2$, DCM/MeOH = 20:1 to 10:1) gave **BCL-7-11** (207 mg, 77%) as a white solid. $R_f$ = 0.18 (DCM/MeOH = 10:1); Purity: >99%, RT = 8.48 min with method 12, RT = 13.45 min with method 13; $^1$H NMR ($D_2O$, 500 MHz): $\delta$ 5.70 (s, 1H), 3.97 (s, 1H), 3.54-3.44 (m, 3H), 3.40-3.29 (m, 5H), 2.49 (m, 2H), 2.01 (s, 3H), 1.85 (s, 3H), 0.91 (s, 3H), 0.87 (s, 3H); $^{13}$C NMR ($D_2O$, 125 MHz): $\delta$ 175.1, 174.1, 170.4, 152.1, 117.4, 75.8, 68.4, 38.9, 38.6, 38.2, 35.4, 35.3, 26.1, 20.5, 19.3, 19.1; HRMS (ESI$^+$) *m/z* $[C_{16}H_{29}N_3O_5Na]^+$ calcd: 366.1999; found: 366.1990.

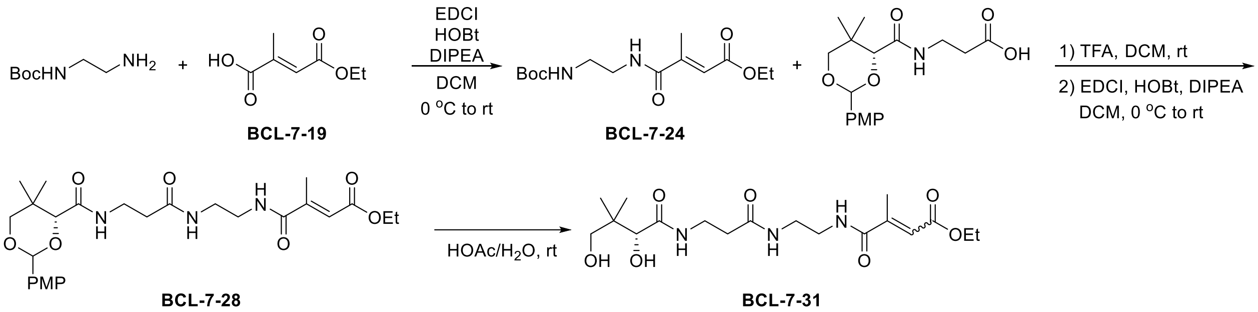

Synthesis of **BCL-7-31 (2-03)**

Following general procedure J, **BCL-7-19** (174.0 mg, 1.1 mmol), Boc-ethylenediamine (0.16 mL, 1.0 mmol), HOBt (185.8 mg, 1.1 mmol), DIPEA (0.45 mL, 2.5 mmol), EDCI (210.9 mg, 1.1 mmol), and DCM (5 mL) were used. Purification by flash chromatography ($SiO_2$, DCM/EA = 10:1 to 2:1) gave **BCL-7-24** (275 mg, 92%) as a white solid. $R_f$ = 0.37 (DCM/EA = 1:1); $^1$H NMR ($CDCl_3$, 500 MHz): $\delta$ 7.24 (bs, 1H), 6.40 (s, 1H), 5.31 (bs, 1H), 4.14 (q, $J$ = 7.1 Hz, 2H), 3.36 (m, 2H), 3.28 (m, 2H), 2.25 (s, 3H), 1.38 (s, 9H), 1.23 (t, $J$ = 7.1 Hz, 3H); $^{13}$C NMR ($CDCl_3$, 125 MHz): $\delta$ 168.7, 166.2, 157.5, 148.0, 122.3, 79.9, 60.4, 42.0, 39.7, 28.4, 14.5, 14.2; HRMS (ESI$^+$) *m/z* $[C_{14}H_{24}N_2O_5Na]^+$ calcd: 323.1577; found: 323.1572.

Following general procedure F, **BCL-7-24** (275.0 mg, 0.92 mmol), TFA (1.5 mL), and DCM (1.5 mL) were used. Next, general procedure K was followed using PMP-pantothenate (340.7 mg, 1.01 mmol), HOBt (170.6 mg, 1.01 mmol), DIPEA (0.41 mL, 2.3 mmol), EDCI (193.6 mg, 1.01 mmol), and DCM (5 mL). Purification by flash chromatography ($SiO_2$, DCM/MeOH = 50:1 to 20:1) gave **BCL-7-28** (424 mg, 89% over two steps) as a white solid. $R_f$ = 0.52 (DCM/MeOH = 10:1); $^1$H NMR ($CDCl_3$, 500 MHz): $\delta$ 7.38 (d, $J$ = 8.7 Hz, 2H), 7.34 (bs, 1H), 7.20 (bs, 1H), 7.02 (m, 1H),

6.86 (d, $J$ = 8.8 Hz, 2H), 6.40 (m, 1H), 5.41 (s, 1H), 4.14 (q, $J$ = 7.2 Hz, 2H), 4.02 (s, 1H), 3.76 (s, 3H), 3.66 (d, $J$ = 11.4 Hz, 1H), 3.60 (d, $J$ = 11.5 Hz, 1H), 3.48 (m, 2H), 3.37-3.31 (m, 4H), 2.38 (m, 2H), 2.24 (s, 3H), 1.24 (t, $J$ = 7.2 Hz, 3H), 1.05 (s, 3H), 1.02 (s, 3H); $^{13}$C NMR ($CDCl_3$, 125 MHz): $\delta$ 172.4, 169.7, 168.8, 166.1, 160.3, 147.8, 130.1, 127.6, 122.4, 113.7, 101.4, 83.8, 78.4, 60.5, 55.3, 40.8, 39.3, 36.1, 35.1, 33.1, 21.8, 19.1, 14.5, 14.20; HRMS (ESI$^+$) *m/z* $[C_{26}H_{37}N_3O_8Na]^+$ calcd: 542.2473; found: 542.2460.

Following general procedure H, **BCL-7-28** (424.0 mg, 0.82 mmol) and 80% HOAc (4 mL) were used. Purification by flash chromatography ($SiO_2$, DCM/MeOH = 20:1 to 10:1) gave **BCL-7-31** (185 mg, 56%) as a sticky white semi-solid. $R_f$ = 0.19 (DCM/MeOH = 10:1); Purity: >99%, RT = 8.69 min with method 12, RT = 13.33 min with method 13; $^1$H NMR ($D_2O$, 500 MHz): $\delta$ 6.33 (q, $J$ = 1.5 Hz, 1H), (4.25, 4.24) (q, $J$ = 7.2 Hz, 2H), 3.97 (s, 1H), 3.55-3.45 (m, 3H), 3.41-3.35 (m, 5H), 2.49 (t, $J$ = 6.6 Hz, 2H), 2.22 (d, $J$ = 1.6 Hz, 3H), (1.30, 1.29) (t, $J$ = 7.1, 3H), 0.91 (s, 3H), 0.88 (s, 3H); $^{13}$C NMR ($D_2O$, 125 MHz): $\delta$ (175.09, 175.07), (174.22, 174.18), (167.9, 167.8), 148.1, (122.12, 122.10), 75.8, 68.4, (61.81, 61.80), (39.20, 39.19), (38.58, 38.56), 38.5, 38.5, 35.4, 35.3, (20.45, 20.44), (19.07, 19.05), 14.4, (13.34, 13.32); HRMS (ESI$^+$) *m/z* $[C_{18}H_{31}N_3O_7Na]^+$ calcd: 424.2054; found: 424.2045.

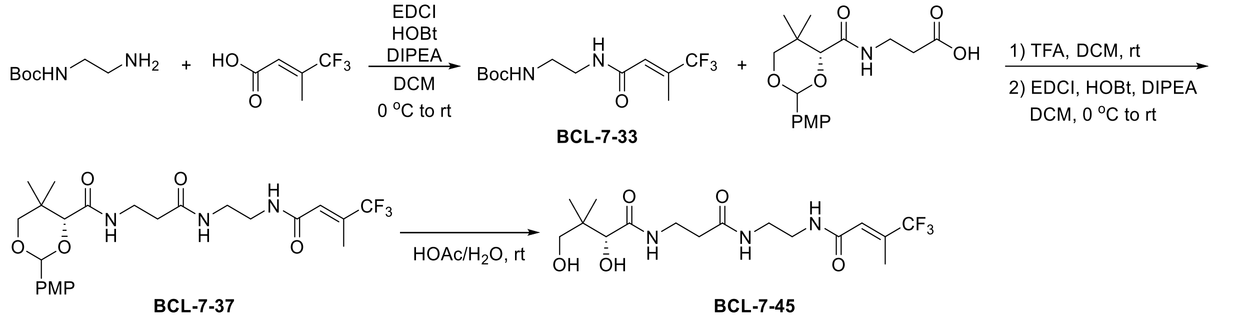

Synthesis of **BCL-7-45 (4-06)**

Following general procedure J, 3-(trifluoromethyl)crotonic acid (169.5 mg, 1.1 mmol), Boc-ethylenediamine (0.16 mL, 1.0 mmol), HOBt (185.8 mg, 1.1 mmol), DIPEA (0.45 mL, 2.5 mmol), EDCI (210.9 mg, 1.1 mmol), and DCM (5 mL) were used. Purification by flash chromatography ($SiO_2$, DCM/EA = 10:1 to 1:1) gave **BCL-7-33** (227 mg, 77%) as a white solid. $R_f$ = 0.62 (DCM/EA = 1:1); $^1$H NMR ($CD_3OD$, 500 MHz): $\delta$ 6.45 (m, 1H), 3.35 (t, $J$ = 6.1 Hz, 2H), 3.21 (t, $J$ = 6.1 Hz, 2H), 2.21 (d, $J$ = 1.5 Hz, 3H), 1.45 (s, 9H); $^{19}$F NMR ($CD_3OD$, 471 MHz): $\delta$ -72.4; $^{13}$C NMR ($CD_3OD$, 125 MHz): $\delta$ 167.0, 158.5, 137.9 (q, $J$ = 30.1 Hz), 125.6 (q, $J$ = 5.8 Hz), 125.0 (q,

$J$ = 273.1 Hz), 80.1, 40.7, 40.4, 28.7, 12.1 (m); HRMS (ESI$^+$) *m/z* $[C_{12}H_{19}N_2O_3F_3Na]^+$ calcd: 319.1240; found: 319.1243.

Following general procedure F, **BCL-7-33** (227.0 mg, 0.77 mmol), TFA (1.5 mL), and DCM (1.5 mL) were used. Next, general procedure K was followed using PMP-pantothenate (286.8 mg, 0.85 mmol), HOBt (162.9 mg, 0.85 mmol), DIPEA (0.35 mL, 1.93 mmol), EDCI (162.9 mg, 0.85 mmol), and DCM (4 mL). Purification by flash chromatography ($SiO_2$, DCM/MeOH = 50:1 to 10:1) gave **BCL-7-37** (344 mg, 87% over two steps) as a white solid. $R_f$ = 0.48 (DCM/MeOH = 10:1); $^1$H NMR ($CD_3OD$, 500 MHz): $\delta$ 7.48 (d, $J$ = 8.8 Hz, 2H), 6.95 (d, $J$ = 8.7 Hz, 2H), 6.45 (m, 1H), 5.54 (s, 1H), 4.16 (s, 1H), 3.82 (s, 3H), 3.73 (d, $J$ = 11.4 Hz, 1H), 3.70 (d, $J$ = 11.3 Hz, 1H), 3.51 (m, 2H), 3.36-3.28 (m, 4H), 2.43 (t, $J$ = 6.7 Hz, 2H), 2.22 (d, $J$ = 1.5 Hz, 3H), 1.14 (s, 3H), 1.07 (s, 3H); $^{19}$F NMR ($CD_3OD$, 471 MHz): $\delta$ -72.2; $^{13}$C NMR ($CD_3OD$, 125 MHz): $\delta$ 174.0, 171.5, 166.8, 161.6, 138.0 (q, $J$ = 29.8 Hz), 131.8, 128.8, 125.5 (q, $J$ = 5.7 Hz), 125.0 (q, $J$ = 273.2 Hz), 114.5, 102.6, 85.1, 79.2, 55.7, 39.8, 36.4, 36.2, 33.9, 22.2, 19.7, 12.1 (m); HRMS (ESI$^+$) *m/z* $[C_{24}H_{32}N_3O_6F_3Na]^+$ calcd: 538.2135; found: 538.2139.

Following general procedure H, **BCL-7-37** (344.0 mg, 0.82 mmol) and 80% HOAc (4 mL) were used. Purification by flash chromatography ($SiO_2$, DCM/MeOH = 20:1 to 10:1) gave **BCL-7-45** (71 mg, 27%) as a white solid. $R_f$ = 0.20 (DCM/MeOH = 10:1); Purity: >99%, RT = 8.71 min with method 12, RT = 13.37 min with method 13; $^1$H NMR ($D_2O$, 500 MHz): $\delta$ 6.49 (m, 1H), 3.97 (s, 1H), 3.55-3.46 (m, 3H), 3.41-3.35 (m, 5H), 2.49 (t, $J$ = 6.6 Hz, 2H), 2.08 (d, $J$ = 1.5 Hz, 3H), 0.91 (s, 3H), 0.88 (s, 3H); $^{19}$F NMR ($D_2O$, 471 MHz): $\delta$ -70.7; $^{13}$C NMR ($D_2O$, 125 MHz): $\delta$ 175.1, 174.2, 167.7, 136.0 (m), 124.2 (m), 123.3 (q, $J$ = 273.4 Hz), 75.8, 68.3, 38.6, 38.6, 38.5, 35.4, 35.3, 20.4, 19.0, 11.7 (m); HRMS (ESI$^+$) *m/z* $[C_{16}H_{26}N_3O_5F_3Na]^+$ calcd: 420.1717; found: 420.1719.

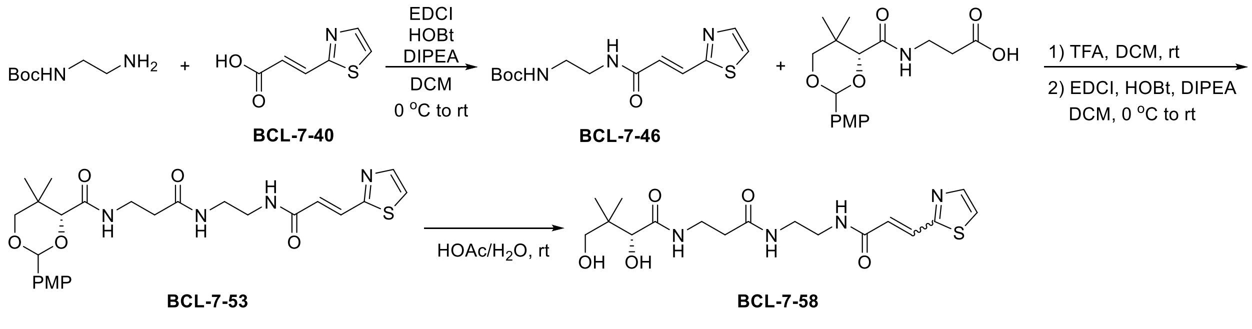

<u>Synthesis of **BCL-7-58 (5-02)**</u>

Following general procedure J, **BCL-7-40** (170.7 mg, 1.1 mmol), Boc-ethylenediamine (0.16 mL, 1.0 mmol), HOBt (185.8 mg, 1.1 mmol), DIPEA (0.45 mL, 2.5 mmol), EDCI (210.9 mg, 1.1

mmol), and DCM (5 mL) were used. Purification by flash chromatography ($SiO_2$, DCM/MeOH = 20:1 to 10:1) gave **BCL-7-46** (264 mg, 88%) as a yellow solid. $R_f$ = 0.48 (DCM/MeOH = 10:1); $^1$H NMR ($CDCl_3$, 500 MHz): $\delta$ 7.79 (m, 1H), 7.65 (dd, $J$ = 15.5, 2.1 Hz, 1H), 7.42 (bs, 1H), 7.32 (m, 1H), 6.79 (d, $J$ = 15.5 Hz, 1H), 5.53 (bs, 1H), 3.46 (m, 2H), 3.29 (m, 2H), 1.36 (s, 9H); $^{13}$C NMR ($CDCl_3$, 125 MHz): $\delta$ 165.6, 164.0, 157.0, 144.4, 132.0, 125.7, 120.8, 79.6, 40.8, 40.3, 28.4; HRMS (ESI$^+$) *m/z* $[C_{13}H_{19}N_3O_3SNa]^+$ calcd: 320.1039; found: 320.1037.

Following general procedure F, **BCL-7-46** (264.0 mg, 0.88 mmol), TFA (1.5 mL), and DCM (1.5 mL) were used. Next, general procedure K was used with PMP-pantothenate (327.2 mg, 0.97 mmol), HOBt (163.8 mg, 0.97 mmol), DIPEA (0.39 mL, 2.2 mmol), EDCI (185.9 mg, 0.97 mmol), and DCM (5 mL). Purification by flash chromatography ($SiO_2$, DCM/MeOH = 50:1 to 15:1) gave **BCL-7-53** (392 mg, 86% over two steps) as a white solid. $R_f$ = 0.48 (DCM/MeOH = 10:1); $^1$H NMR ($CDCl_3$, 500 MHz): $\delta$ 7.80 (m, 1H), 7.64 (dd, $J$ = 15.3, 1.7 Hz, 1H), 7.50 (bs, 1H), 7.38 (d, $J$ = 8.8 Hz, 2H), 7.33 (m, 1H), 7.24 (bs, 1H), 7.09 (t, $J$ = 6.4 Hz, 1H), 6.85 (d, $J$ = 8.9 Hz, 2H), 6.79 (d, $J$ = 15.4 Hz, 1H), 5.41 (s, 1H), 4.04 (s, 1H), 3.75 (s, 3H), 3.64 (d, $J$ = 11.4 Hz, 1H), 3.59 (d, $J$ = 11.4 Hz, 1H), 3.50 (m, 2H), 3.45-3.29 (m, 4H), 2.40 (t, $J$ = 6.6 Hz, 2H), 1.06 (s, 3H), 1.02 (s, 3H); $^{13}$C NMR ($CDCl_3$, 125 MHz): $\delta$ 172.1, 169.7, 165.7, 164.0, 160.2, 144.4, 132.0, 130.1, 127.6, 125.7, 120.9, 113.7, 101.3, 83.9, 78.4, 55.3, 39.9, 39.6, 36.1, 35.2, 33.1, 21.9, 19.2; HRMS (ESI$^+$) *m/z* $[C_{25}H_{32}N_4O_6SNa]^+$ calcd: 539.1935; found: 539.1939.

Following general procedure H, **BCL-7-53** (392.0 mg, 0.76 mmol) and 80% HOAc (4 mL) were used. Purification by flash chromatography ($SiO_2$, DCM/MeOH = 20:1 to 8:1) gave **BCL-7-58** (208 mg, 68%) as a white solid. $R_f$ = 0.12 (DCM/MeOH = 10:1); Purity: >99%, RT = 8.58 min with method 12, RT = 13.04 min with method 13; $^1$H NMR ($D_2O$, 500 MHz): $\delta$ (7.87, 7.84) (d, $J$ = 3.3 Hz, 1H), (7.67, 7.64) (d, $J$ = 3.3 Hz, 1H), (7.55, 7.49) (d, $J$ = 15.7 Hz, 1H), (6.79, 6.73) (d, $J$ = 15.6 Hz, 1H), (3.94, 3.93) (s, 1H), 3.51-3.31 (m, 8H), 2.48 (m, 2H), (0.87, 0.86) (s, 3H), (0.83, 0.82) (s, 3H); $^{13}$C NMR ($D_2O$, 125 MHz): $\delta$ (175.06, 175.03), (174.2, 174.1), (167.4, 167.2), (164.33, 164.25), (143.61, 143.59), (131.6, 131.5), (125.11, 125.06), 122.6, 75.7, 68.3, 38.9, 38.7, 38.5, 35.4, 35.3, 20.4, 19.0; HRMS (ESI$^+$) *m/z* $[C_{17}H_{26}N_4O_5SNa]^+$ calcd: 421.1516; found: 421.1514.

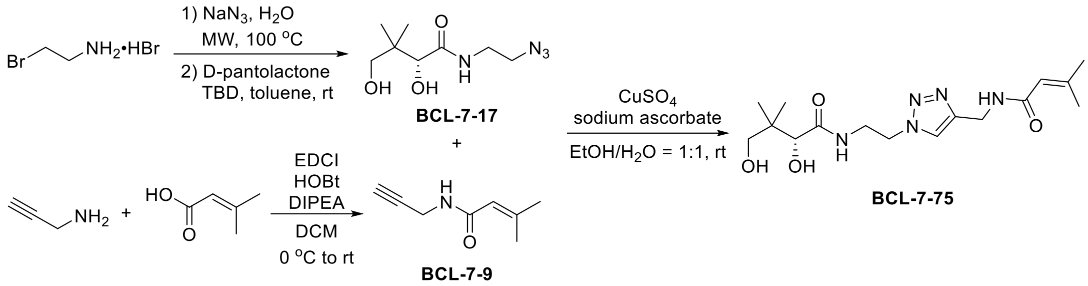

Synthesis of **BCL-7-75 (3-09)**

To a Biotage® 20 mL microwave vial were added 2-bromoethylamine hydrobromide (1.02 g, 5 mmol, 1.0 equiv.), $NaN_3$ (975.0 mg, 15 mmol, 3.0 equiv.), and water (10 mL). The reaction mixture was subjected to microwave irradiation at 100°C for 1 h. The mixture was cooled to rt and aq. NaOH (4 M) was added to adjust the *p*H = 14. The product was extracted in diethyl ether (3 × 25 mL). The combined organic layers were dried over anhydrous $Na_2SO_4$, filtered, and the solvent evaporated carefully under vacuum (cold water bath, low vacuum) to give the azide intermediate (400 mg, 93%). Following a reported procedure,[45] to a 25 mL round bottom flask were added the azide intermediate (400 mg, 4.65 mmol, 1.0 equiv.), D-pantolactone (1.21 g, 9.3 mmol, 2.0 equiv.), TBD (65.4 mg, 0.47 mmol, 0.1 equiv.), and toluene (5 mL). The reaction mixture was stirred at rt for 24 h. The solvent was evaporated under vacuum. Purification by flash chromatography ($SiO_2$, DCM/EA = 4:1, then DCM/MeOH = 15:1) gave **BCL-7-17** (379 mg, 38%) as a colorless oil. $R_f$ = 0.51 (DCM/MeOH = 10:1); $^1$H NMR ($CDCl_3$, 500 MHz): $\delta$ 7.37 (bs, 1H), 5.03 (m, 1H), 4.35 (bs, 1H), 3.98 (m, 1H), 3.46-3.36 (m, 6H), 0.90 (s, 3H), 0.87 (s, 3H); $^{13}$C NMR ($CDCl_3$, 125 MHz): $\delta$ 174.3, 76.8, 70.7, 50.7, 39.2, 38.3, 20.7, 20.5; HRMS (ESI$^+$) *m/z* $[C_8H_{16}N_4O_3Na]^+$ calcd: 239.1115; found: 239.1119.

Following general procedure J, 3,3-dimethylacrylic acid (275.3 mg, 2.75 mmol), distilled propargylamine (0.16 mL, 2.5 mmol), HOBt (464.5 mg, 2.75 mmol), DIPEA (1.11 mL, 6.25 mmol), EDCI (527.2 mg, 2.75 mmol), and DCM (10 mL) were used. Purification by flash chromatography ($SiO_2$, Hex/EA = 5:1 to 1:1) gave **BCL-7-9** (272 mg, 79%) as an off-white solid. $R_f$ = 0.34 (Hex/EA = 2:1); $^1$H NMR ($CDCl_3$, 500 MHz): $\delta$ 6.31 (bs, 1H), 5.57 (m, 1H), 4.00 (dd, *J* = 5.4, 2.6 Hz, 2H), 2.15 (t, *J* = 2.6 Hz, 1H), 2.09 (d, *J* = 1.5 Hz, 3H), 1.77 (d, *J* = 1.5 Hz, 3H); $^{13}$C NMR ($CDCl_3$, 125 MHz): $\delta$ 166.8, 151.9, 117.8, 80.1, 71.1, 28.7, 27.1, 19.8; HRMS (ESI$^+$) *m/z* $[C_8H_{11}NONa]^+$ calcd: 160.0733; found: 160.0733.

To a 25 mL round bottom flask were added **BCL-7-17** (136.7 mg, 0.63 mmol, 1.0 equiv.), sodium ascorbate (124.8 mg, 0.63 mmol, 1.0 equiv.), $CuSO_4$ (10.1 mg, 0.06 mmol), and EtOH/$H_2O$ = 1:1 (6 mL). **BCL-7-9** (86.7 mg, 0.63 mmol, 1.0 equiv.) was added and the reaction mixture was stirred at rt for 1 h. The reaction was quenched with aq. EDTA (17.5 mg, 0.06 mmol, 0.1 equiv., dissolved in 2 mL sat. $Na_2CO_3$). The solvent was evaporated under vacuum. The residue was dissolved in MeOH and filtered. The filtrate was dried over anhydrous $Na_2SO_4$, filtered, and the solvent was evaporated under vacuum. Purification by flash chromatography ($SiO_2$, DCM/MeOH = 20:1 to 10:1) gave **BCL-7-75** (187 mg, 84%) as a hygroscopic white solid. $R_f$ = 0.24 (DCM/MeOH = 10:1); Purity: >99%, RT = 8.56 min with method 12, RT = 13.56 min with method 13; $^1$H NMR ($D_2O$, 500 MHz): $\delta$ 7.91 (s, 1H), 5.72 (m, 1H), 4.58 (m, 2H), 4.46 (s, 2H), 3.89 (s, 1H), 3.82 (m, 1H), 3.71 (m, 1H), 3.41 (d, $J$ = 11.2 Hz, 1H), 3.27 (d, $J$ = 11.2 Hz, 1H), 2.02 (d, $J$ = 1.3 Hz, 3H), 1.85 (d, $J$ = 1.4 Hz, 3H), 0.73 (s, 3H), 0.70 (s, 3H); $^{13}$C NMR ($D_2O$, 125 MHz): $\delta$ 175.4, 169.8, 153.0, 145.1, 124.0, 117.1, 75.7, 68.3, 49.5, 38.5, 38.4, 33.9, 26.2, 20.3, 19.4, 18.8; HRMS (ESI$^+$) *m/z* $[C_{16}H_{27}N_5O_4Na]^+$ calcd: 376.1955; found: 376.1962.

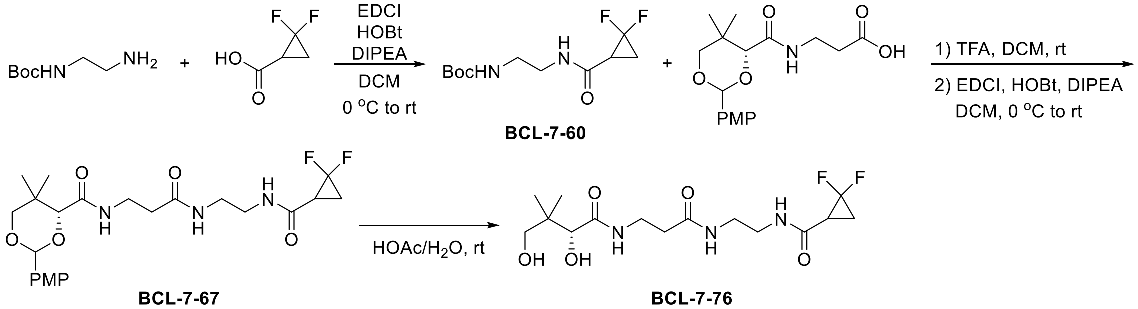

Synthesis of **BCL-7-76 (4-10)**

Following general procedure J, 2,2-difluorocyclopropanecarboxylic acid (134.3 mg, 1.1 mmol), Boc-ethylenediamine (0.16 mL, 1.0 mmol), HOBt (185.8 mg, 1.1 mmol), DIPEA (0.45 mL, 2.5 mmol), EDCI (210.9 mg, 1.1 mmol), and DCM (5 mL) were used. Purification by flash chromatography ($SiO_2$, DCM/MeOH = 50:1 to 10:1) gave **BCL-7-60** (141 mg, 53%) as a white solid. $R_f$ = 0.55 (DCM/MeOH = 10:1); $^1$H NMR ($CD_3OD$, 500 MHz): $\delta$ 3.32 (m, 2H), 3.20 (m, 2H), 2.50 (ddd, $J$ = 13.6, 10.9, 7.9 Hz, 1H), 2.01 (dtd, $J$ = 13.7, 7.8, 6.3 Hz, 1H), 1.75 (dddd, $J$ = 12.2, 10.8, 7.7, 4.7 Hz, 1H), 1.46 (s, 9H); $^{19}$F NMR ($CD_3OD$, 471 MHz): $\delta$ -127.5 (d, $J$ = 153.6 Hz), -143.3 (d, $J$ = 153.6 Hz); $^{13}$C NMR ($CD_3OD$, 125 MHz): $\delta$ 167.7, 158.6, 112.7 (dd, $J$ = 286.2, 281.9 Hz), 80.2, 40.83, 40.80, 28.7, 27.6 (t, $J$ = 10.8 Hz), 15.5 (dd, $J$ = 11.1, 10.0 Hz); HRMS (ESI$^+$) *m/z* $[C_{11}H_{18}N_2O_3F_2Na]^+$ calcd: 287.1178; found: 287.1174.

Following general procedure F, **BCL-7-60** (328.0 mg, 1.24 mmol), TFA (2 mL), and DCM (2 mL) were used. Next, general procedure K was followed using PMP-pantothenate (458.8 mg, 1.36 mmol), HOBt (229.7 mg, 1.36 mmol), DIPEA (0.55 mL, 3.1 mmol), EDCI (260.7 mg, 1.36 mmol), and DCM (6 mL). Purification by flash chromatography ($SiO_2$, DCM/MeOH = 50:1 to 20:1) gave **BCL-7-67** (553 mg, 92% over two steps) as a white solid. $R_f$ = 0.48 (DCM/MeOH = 10:1); $^1$H NMR ($CDCl_3$, 500 MHz): $\delta$ 7.40 (d, $J$ = 8.7 Hz, 2H), 7.37 (bs, 1H), 7.09-7.04 (m, 2H), 6.89 (d, $J$ = 8.7 Hz, 2H), 5.43 (s, 1H), 4.04 (s, 1H), 3.78 (s, 3H), 3.67 (d, $J$ = 11.5 Hz, 1H), 3.62 (d, $J$ = 11.6 Hz, 1H), 3.48 (m, 2H), 3.36-3.22 (m, 4H), 2.37 (t, $J$ = 6.2 Hz, 2H), 2.29 (m, 1H), 2.01 (m, 1H), 1.58 (m, 1H), 1.07 (s, 3H), 1.04 (s, 3H); $^{19}$F NMR ($CDCl_3$, 471 MHz): $\delta$ -125.65 (d, $J$ = 153.6), -125.67 (d, $J$ = 153.6), -141.36 (dd, $J$ = 153.5), -141.39 (dd, $J$ = 153.5); $^{13}$C NMR ($CDCl_3$, 125 MHz): $\delta$ (172.24, 172.19), (169.72, 169.69), 165.8, 160.3, (130.18, 130.17), 127.6, 113.8, 111.2 (dd, $J$ = 287.3, 283.1 Hz), 101.4, (83.89, 83.88), 78.4, 55.4, (40.13, 40.06), (39.51, 39.47), (36.1, 36.0), 35.1, (33.10, 33.09), 26.8 (t, $J$ = 10.7 Hz), 21.9, 19.1, 15.3 (t, $J$ = 10.3 Hz); HRMS (ESI$^+$) *m/z* $[C_{23}H_{31}N_3O_6F_2Na]^+$ calcd: 506.2073; found: 506.2091.

Following general procedure H, **BCL-7-67** (553.0 mg, 1.14 mmol) and 80% HOAc (5 mL) were used. Purification by flash chromatography ($SiO_2$, DCM/MeOH = 20:1 to 8:1) gave **BCL-7-76** (288 mg, 69%) as a white solid. $R_f$ = 0.10 (DCM/MeOH = 10:1); Purity: >99%, RT = 8.30 min with method 12, RT = 13.14 min with method 13; $^1$H NMR ($D_2O$, 500 MHz): $\delta$ 3.98 (s, 1H), 3.55-3.45 (m, 3H), 3.43-3.26 (m, 5H), 2.58 (ddd, $J$ = 13.1, 11.1, 7.9 Hz, 1H), 2.48 (t, $J$ = 6.6 Hz, 2H), 1.97 (dtd, $J$ = 13.6, 8.2, 5.8 Hz, 1H), 1.86 (m, 1H), 0.92 (s, 3H), 0.88 (s, 3H); $^{19}$F NMR ($D_2O$, 471 MHz): $\delta$ -126.6 (d, $J$ = 154.1 Hz), (-141.22, -141.23) (d, $J$ = 154.0 Hz); $^{13}$C NMR ($D_2O$, 125 MHz): $\delta$ 175.1, 174.2, 168.4, 111.4 (t, $J$ = 282.5 Hz), 75.8, 68.3, 38.9, 38.7, 38.6, 35.4, 35.3, 26.3 (t, $J$ = 10.8 Hz), 20.4, 19.0, 15.0 (t, $J$ = 10.3 Hz); HRMS (ESI$^+$) *m/z* $[C_{15}H_{25}N_3O_5F_2Na]^+$ calcd: 388.1654; found: 388.1649. HPLC method 12: RT = 8.30 min (>99%); method 13: 13.14 min (>99%).

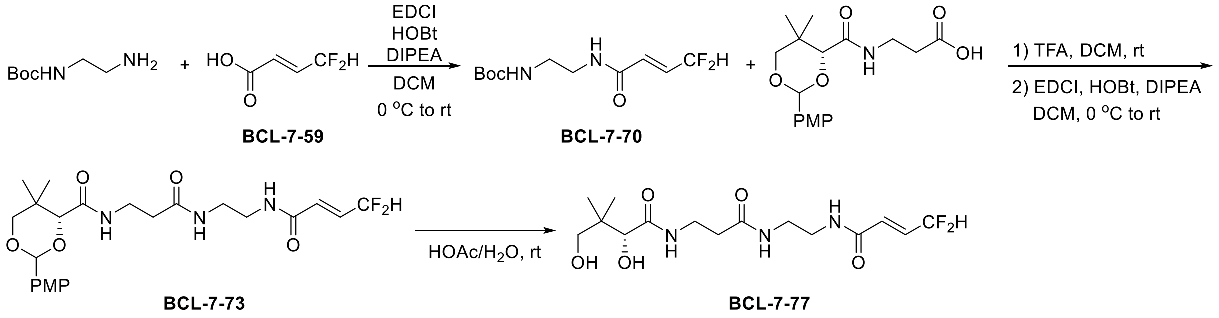

Synthesis of **BCL-7-77 (4-05)**

Following general procedure J, **BCL-7-59** (134.3 mg, 1.1 mmol), Boc-ethylenediamine (0.16 mL, 1.0 mmol), HOBt (185.8 mg, 1.1 mmol), DIPEA (0.45 mL, 2.5 mmol), EDCI (210.9 mg, 1.1 mmol), and DCM (5 mL) were used. Purification by flash chromatography ($SiO_2$, DCM/EA = 5:1 to 1:1) gave **BCL-7-70** (170 mg, 64%) as a white solid. $R_f$ = 0.32 (DCM/EA = 1:1); $^1$H NMR ($CD_3OD$, 500 MHz): $\delta$ 6.69 (dtd, $J$ = 15.8, 10.1, 4.4 Hz, 1H), 6.42 (dtd, $J$ = 15.7, 3.1, 0.9 Hz, 1H), 6.36 (td, $J$ = 55.2, 4.6 Hz, 1H), 3.34 (t, $J$ = 6.2 Hz, 2H), 3.19 (t, $J$ = 6.1 Hz, 2H), 1.43 (s, 9H); $^{19}$F NMR ($CD_3OD$, 471 MHz): $\delta$ -116.4; $^{13}$C NMR ($CD_3OD$, 125 MHz): $\delta$ 166.4, 158.6, 134.0 (t, $J$ = 24.2 Hz), 130.9 (t, $J$ = 10.5 Hz), 114.8 (t, $J$ = 234.3 Hz), 80.1, 40.74, 40.72, 28.7; HRMS (ESI$^+$) *m/z* $[C_{11}H_{18}N_2O_3F_2Na]^+$ calcd: 287.1178; found: 287.1176.

Following general procedure F, **BCL-7-70** (334.0 mg, 1.26 mmol), TFA (2 mL), and DCM (2 mL) were used. Next, general procedure K was followed using PMP-pantothenate (468.9 mg, 1.39 mmol), HOBt (234.8 mg, 1.39 mmol), DIPEA (0.56 mL, 3.15 mmol), EDCI (266.4 mg, 1.39 mmol), and DCM (7.5 mL). Purification by flash chromatography ($SiO_2$, DCM/MeOH = 50:1 to 15:1) gave **BCL-7-73** (586 mg, 96% over two steps) as a white solid. $R_f$ = 0.43 (DCM/MeOH = 10:1); $^1$H NMR (DMSO-$d_6$, 500 MHz): $\delta$ 8.44 (t, $J$ = 5.7 Hz, 1H), 7.98 (t, $J$ = 5.6 Hz, 1H), 7.49 (t, $J$ = 6.0 Hz, 1H), 7.44 (d, $J$ = 8.7 Hz, 2H), 6.93 (d, $J$ = 8.8 Hz, 2H), 6.76-6.50 (m, 2H), 6.45 (dt, $J$ = 15.6, 3.1 Hz, 1H), 5.52 (s, 1H), 4.08 (s, 1H), 3.76 (s, 3H), 3.65 (d, $J$ = 11.3 Hz, 1H), 3.61 (d, $J$ = 11.2 Hz, 1H), 3.36 (m, 1H), 3.28 (m, 1H), 3.18 (m, 2H), 3.13 (m, 2H), 2.28 (t, $J$ = 7.0 Hz, 2H), 1.01 (s, 3H), 0.95 (s, 3H); $^{19}$F NMR (DMSO-$d_6$, 471 MHz): $\delta$ -113.9; $^{13}$C NMR (DMSO-$d_6$, 125 MHz): $\delta$ 170.8, 168.2, 163.1, 159.6, 131.3 (t, $J$ = 23.6 Hz), 130.9 (t, $J$ = 10.5 Hz), 130.5, 127.7, 113.9 (t, $J$ = 233.2 Hz), 113.3, 100.4, 83.2, 77.4, 55.1, 38.6, 38.1, 35.1, 34.8, 32.5, 21.5, 19.0; HRMS (ESI$^+$) *m/z* $[C_{23}H_{31}N_3O_6F_2Na]^+$ calcd: 506.2073; found: 506.2071.

Following general procedure H, **BCL-7-73** (561.0 mg, 1.16 mmol) and 80% HOAc (5 mL) were used. Purification by flash chromatography ($SiO_2$, DCM/MeOH = 20:1 to 10:1) gave **BCL-7-77** (239 mg, 56%) as a white solid. $R_f$ = 0.08 (DCM/MeOH = 10:1); Purity: >99%, RT = 8.36 min with method 12, RT = 13.17 min with method 13; $^1$H NMR ($D_2O$, 500 MHz): $\delta$ 6.69 (dtd, $J$ = 15.9, 10.1, 4.4 Hz, 1H), 6.49 (dtd, $J$ = 15.6, 3.1, 0.9 Hz, 1H), 6.42 (tdd, $J$ = 55.3, 4.5, 1.0 Hz, 1H), 3.98 (s, 1H), 3.64-3.23 (m, 8H), 2.48 (t, $J$ = 6.6 Hz, 2H), 0.91 (s, 3H), 0.88 (s, 3H); $^{19}$F NMR ($D_2O$, 471 MHz): $\delta$ -115.2; $^{13}$C NMR ($D_2O$, 125 MHz): $\delta$ 175.1, 174.2, 166.7, 133.1 (t, $J$ = 24.1 Hz), 129.2 (t, $J$ = 10.8 Hz), 113.4 (t, $J$ = 234.1 Hz), 75.8, 68.3, 38.9, 38.6, 38.5, 35.4, 35.3, 20.5, 19.0; HRMS

(ESI$^+$) *m/z* $[C_{15}H_{25}N_3O_5F_2Na]^+$ calcd: 388.1654; found: 388.1653. HPLC method 12: RT = 8.36 min (>99%); method 13: 13.17 min (>99%).

Synthesis of **BCL-07-042**

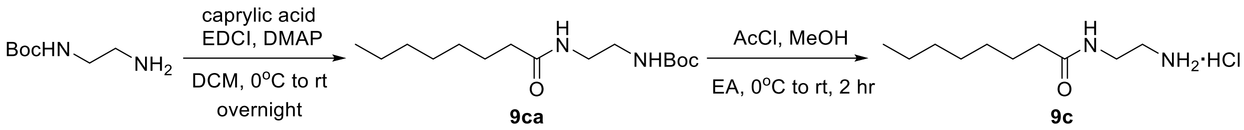

To a 100 mL round bottom flask were added *N*-Boc-ethylenediamine (1.60 g, 10 mmol, 1.0 equiv.), caprylic acid (1.74 mL, 11 mmol, 1.1 equiv.), DMAP (122.2 mg, 1 mmol, 0.1 equiv.), and anhydrous DCM (20 mL). The mixture was cooled to rt and EDCI (2.11 g, 11 mmol, 1.1 equiv.) was added in a few portions. The reaction mixture was warmed to rt and stirred overnight. The mixture was next diluted in DCM (40 mL), washed with sat. $NaHCO_3$ (2×50 mL), 1 M $KH_2PO_4$ (2×50 mL), and brine (1×50 mL). The organic layer was dried over anhydrous $Na_2SO_4$ and the solvent was evaporated under vacuum. The residue was dissolved in a mixture of hexanes and EA (Hex/EA = 100:1, 20 mL) and left in a freezer for 1 hour. The product was collected by filtration and washed with hexanes to afford **9ca** (2.62 g, 91%) as a white solid. This compound was previously reported.[13] It was used in the next step without further purification.
To a 100 mL round bottom flask were added **9ca** (2.62 g, 9.1 mmol, 1.0 equiv.), anhydrous EA (15 mL), and anhydrous MeOH (2.43 mL, 60 mmol). The mixture was cooled to 0°C and AcCl (4.28 mL, 60 mmol) was added dropwise. The mixture was stirred at rt for 2 hours. The product was collected by filtration and washed with EA to afford **9c** (2.09 g, quant.) as a very hygroscopic white solid. Mp: 111°C (decomp.); IR (neat, $cm^{-1}$): 3359 (w, br), 2922 (m), 2854 (m), 1665 (m), 1601 (m), 1521 (m); $^1H$ NMR (DMSO-$d_6$, 500 MHz): $\delta$ 8.74 (bs, 1H), 8.27 (bs, 3H), 3.24 (m, 2H), 2.76 (m, 2H), 2.00 (m, 2H), 1.38 (m, 2H), 1.14-1.11 (m, 8H), 0.74 (m, 3H); $^{13}C\{^1H\}$ NMR (DMSO-$d_6$, 125 MHz): $\delta$ 173.6, 39.0, 36.8, 35.8, 31.6, 29.1, 28.9, 25.5, 22.5, 14.3; HRMS (ESI-TOF) *m/z*: $[M - H]^-$ Cacld for $C_{10}H_{22}N_2OCl$ 221.1426; Found 221.1425.

**\Synthesis of BCL-7-49 (6-14)**

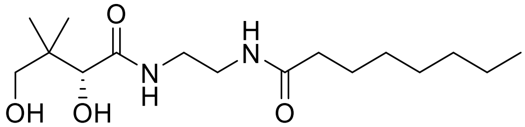

*N*-(2-Aminoethyl)octanamide (**BCL-07-042**, 93.1 mg, 0.5 mmol, 1.0 equiv.) and D-pantolactone (130.1 mg, 1.0 mmol, 2.0 equiv.) were reacted following the general procedure. Purification by flash chromatography ($SiO_2$, DCM/EA = 4:1, then DCM/MeOH = 15:1) gave the desired product **BCL-07-049** (148 mg, 94%) as a white solid. Alternatively, upon completion, the mixture was diluted in toluene (2 mL) and the product was collected by filtration. The filter cake was washed with toluene and hexanes to give the desired product **BCL-07-049** (107 mg, 68%) as a white solid. $R_f$ = 0.37 (DCM/MeOH = 10:1); $[\alpha]_D^{23}$ = +30.3 (c = 1.00, MeOH); Mp: 95-97°C; IR (neat, $cm^{-1}$): 3329 (m, br), 3287 (m, br), 2925 (w), 2854 (w), 1642 (s), 1558 (m), 1032 (m); $^1H$ NMR ($CDCl_3$, 500 MHz): $\delta$ 7.52 (t, $J$ = 5.7 Hz, 1H), 6.97 (t, $J$ = 5.3 Hz, 1H), 5.05 (d, $J$ = 5.0 Hz, 1H), 4.37 (t, $J$ = 5.4 Hz, 1H), 3.97 (d, $J$ = 4.9 Hz, 1H), 3.46 (dd, $J$ = 11.1, 5.4 Hz, 1H), 3.40 (dd, $J$ = 11.1, 5.5 Hz, 1H), 3.38-3.26 (m, 4H), 2.13 (t, $J$ = 7.7 Hz, 2H), 1.54 (m, 2H), 1.27-1.17 (m, 8H), 0.92 (s, 3H), 0.90 (s, 3H), 0.83 (t, $J$ = 6.9 Hz, 3H); $^{13}C\{^1H\}$ NMR ($CDCl_3$, 125 MHz): $\delta$ 174.9, 77.5, 70.8, 40.0, 39.3, 39.1, 36.7, 31.7, 29.3, 29.1, 25.8, 22.7, 21.1, 20.9, 14.1; HRMS (ESI-TOF) *m/z*: $[M + Na]^+$ Cacld for $C_{16}H_{32}N_2O_4Na$ 339.2254; Found 339.2259.

Synthesis of **ABA-1-1**

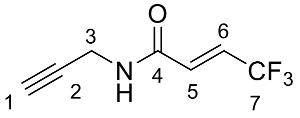

**ABA-1-1**

Following the general procedure L, 4,4,4-trifluorocrotonic acid (154 mg, 1.1 mmol), HOBt hydrate (80% w/w, 186 mg, 1.1 mmol), DCM (5 mL), DIPEA (0.44 mL, 2.5 mmol), EDCI (211 mg, 1.1 mmol), and propargylamine (64 μL, 1.0 mmol) were used. Purification by flash chromatography ($SiO_2$, DCM/EA = 100:1 to 50:1) yielded the product **ABA-1-1** (60 mg, 27%) as a white crystalline solid. $R_f$ = 0.38 (DCM/EA = 50:1); $^1$H NMR ($CDCl_3$, 500 MHz): $\delta$ 6.93 (bs, 1H, N-H), 6.78 (dq, $J$ = 15.4, 6.6 Hz, 1H, H-6), 6.60 (dq, $J$ = 15.4, 2.0 Hz, 1H, H-5), 4.15 (dd, $J$ = 5.3, 2.6 Hz, 2H, H-3), 2.27 (t, $J$ = 2.6 Hz, 1H, H-1); $^{19}$F NMR ($CDCl_3$, 471 MHz): $\delta$ -65.2; $^{13}$C NMR ($CDCl_3$, 125 MHz): $\delta$ 162.8, 130.3 (q, $J$ = 5.8 Hz), 129.2 (q, $J$ = 35.3 Hz), 122.5 (q, $J$ = 270.0 Hz), 78.5, 72.4, 29.8; HRMS (ESI$^+$) *m/z* $[C_7H_6NOF_3Na]^+$ calcd: 200.0294; found: 200.0297.

Synthesis of **ABA-1-3**

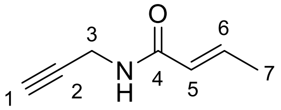

**ABA-1-3**

Following procedure M, propargylamine (160 μL, 2.5 mmol), DCM (10 mL), TEA (0.53 mL, 3.8 mmol), and crotonoyl chloride (0.29 mL, 3.0 mmol) were used. Purification by flash chromatography ($SiO_2$, Hex/EA = 2:1 to 1:1) yielded the product **ABA-1-3** (260 mg, 84%) as a white solid. $R_f$ = 0.62 (Hex/EA = 2:1); $^1$H NMR ($CDCl_3$, 500 MHz): $\delta$ 6.87 (dq, $J$ = 15.4, 6.9 Hz, 1H, H-6), 5.88 (bs, 1H, N-H), 5.81 (dd, $J$ = 15.3, 1.7 Hz, 1H, H-5), 4.10 (dd, $J$ = 5.3, 2.6 Hz, 2H, H-3), 2.22 (t, $J$ = 2.5 Hz, 1H, H-1), 1.85 (dd, $J$ = 6.9, 1.7 Hz, 3H, H-7); $^{13}$C NMR ($CDCl_3$, 125 MHz): $\delta$ 165.7, 141.0, 124.4, 79.7, 71.6, 29.3, 17.9; HRMS (ESI$^+$) *m/z* $[C_7H_{10}NO]^+$ calcd: 124.0757; found: 124.0757. The characterization matched that reported in the literature. [46]

Synthesis of **ABA-1-23**

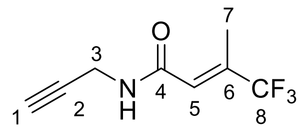

**ABA-1-23**

Following the general procedure L, 3-(trifluoromethyl)crotonic (170 mg, 1.1 mmol), HOBt hydrate (80% w/w, 186 mg, 1.1 mmol), DCM (5 mL), DIPEA (0.44 mL, 2.5 mmol), EDCI (211 mg, 1.1 mmol), and propargylamine (64 µL, 1.0 mmol) were used. Purification by flash chromatography ($SiO_2$, Hex/EA = 10:1 to 5:1) yielded the product **ABA-1-23** (67 mg, 35%) as an oil that solidifies upon standing. $R_f$ = 0.18 (Hex/EA = 5:1); $^1$H NMR ($CDCl_3$, 500 MHz): $\delta$ 6.27 (m, 1H, H-5), 6.23 (bs, 1H, N-H), 4.11 (dd, $J$ = 5.3, 2.6 Hz, 2H, H-3), 2.25 (t, $J$ = 2.6 Hz, 1H, H-1), 2.23 (d, $J$ = 1.5 Hz, 3H, H-7); $^{19}$F NMR ($CDCl_3$, 471 MHz): $\delta$ -71.2; $^{13}$C NMR ($CDCl_3$, 125 MHz): $\delta$ 164.2, 139.4 (q, $J$ = 30.2 Hz), 123.4 (q, $J$ = 274.0 Hz), 122.9 (q, $J$ = 5.6 Hz), 78.9, 72.2, 29.3, 12.2; HRMS (ESI$^+$) *m/z* $[C_8H_8NOF_3Na]^+$ calcd: 214.0450; found: 214.0446.

Synthesis of **ABA-1-9**

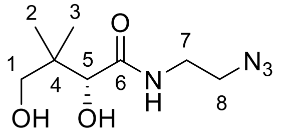

**ABA-1-9**

To a Biotage® 20 mL MW vial were added 2-bromoethylamine hydrobromide (3.10 g, 15 mmol, 1.0 equiv.), $NaN_3$ (2.40 g, 37.5 mmol, 2.5 equiv.), and water (15 mL). The reaction was allowed to proceed under MW radiation with stirring at 100°C for 1 h. Upon cooling to rt, the crude product was diluted in water (50 mL), and the pH was increased to 14 using NaOH pellets. The product was extracted in diethyl ether (4 × 30 mL) and the combined organic layers were washed with water (3 × 25 mL). The organic layer was then dried over anhydrous $Na_2SO_4$ and the solvent was evaporated under vacuum (carefully due to product volatility) to yield 2-azidoethan-1-amine as a yellow oil (1.30 g), which was used in the next step without purification. Following the general procedure O, the amine (1.30 g), D-pantolactone (3.90 g, 30 mmol), TBD (210 mg, 1.5 mmol), and toluene (15 mL) were used. Purification by flash chromatography ($SiO_2$, DCM/EA = 4:1 then DCM/MeOH = 20:1) afforded **ABA-1-9** (780 mg, 24% over two steps) as an oil with a yellow tinge. $R_f$ = 0.38 (DCM/MeOH = 10:1); $^1$H NMR ($CDCl_3$, 500 MHz): $\delta$ 7.25 (bs, 1H, NH), 4.04

(bs, 1H, OH), 4.02 (s, 2H, H-5, OH), 3.48 (s, 2H, H-1), 3.47 – 3.41 (m, 4H, H-7, H-8), 0.98 (s, 3H, H-3), 0.91 (s, 3H, H-2); $^{13}$C NMR ($CDCl_3$, 125 MHz): $\delta$ 173.9, 77.5, 71.3, 50.9, 39.2, 38.5, 21.1, 20.4. The characterization matched that reported in the literature. [7]

Synthesis of **ABA-1-32**

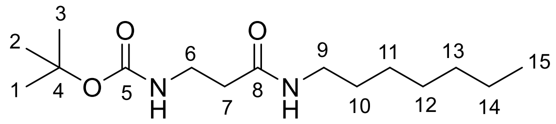

**ABA-1-32**

To a 100 mL round-bottom flask, *n*-heptylamine (1.48 mL, 10 mmol, 1.0 equiv.), Boc-β-alanine (2.08 g, 11 mmol, 1.1 equiv.), DMAP (122 mg, 1.0 mmol, 0.1 equiv.), and DCM (25 mL) were added. The mixture was left to stir for 1 h at 0°C. EDCI (2.11 g, 11 mmol, 1.1 equiv.) was then added and the mixture was left to stir for overnight at rt. The mixture was diluted with DCM (40 mL) then washed with sat. $Na_2CO_3$ (2 × 20 mL) and 1 M $KH_2PO_4$ (2 × 20 mL). The organic layer was then dried over anhydrous $Na_2SO_4$, and the solvent was evaporated under vacuum to yield a yellow oil that solidified upon standing. The solid residue was dissolved in a mixture of hexanes and EA (100:1, 20 mL) and placed in a freezer for 2 h. The product was then collected by filtration and washed with cold hexanes to yield **ABA-1-32** (2.32 g, 81%) as a white solid. $R_f$ = 0.38 (DCM/EA = 1:1); $^1$H NMR ($CDCl_3$, 500 MHz): $\delta$ 5.90 (bs, 1H, N-H), 5.18 (bs, 1H, N-H), 3.39 (t, $J$ = 6.1 Hz, 2H, H-6), 3.27 – 3.18 (m, 2H, H-9), 2.39 (t, $J$ = 6.0 Hz, 2H, H-7), 1.52 – 1.45 (m, 2H, H-10), 1.42 (s, 9H, H-1, H-2, H-3), 1.33 – 1.21 (m, 8H, H-11, H-12, H-13, H-14), 0.87 (t, $J$ = 6.7 Hz, 3H, H-15); $^{13}$C NMR ($CDCl_3$, 125 MHz): $\delta$ 171.4, 156.3, 79.5, 39.8, 36.9, 36.5, 31.8, 29.7, 29.1, 28.5, 27.0, 22.7, 14.2; HRMS ($ESI^+$) *m/z* $[C_{15}H_{30}N_2O_3Na]^+$ calcd: 309.2149; found: 309.2150.

Synthesis of **ABA-1-18 (3-08)**

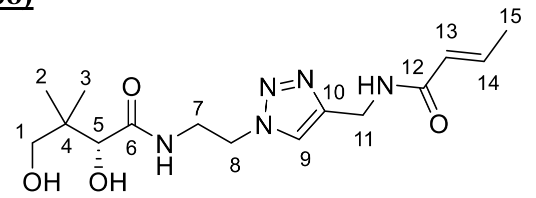

**ABA-1-18**

Following the general procedure P, **ABA-1-3** (62 mg, 0.5 mmol), **ABA-1-9** (108 mg, 0.5 mmol), sodium ascorbate (99 mg, 0.5 mmol), $CuSO_4$ (8 mg, 0.05 mmol), and EtOH/$H_2O$ = 1:1 (5 mL) were used. Purification by flash chromatography ($SiO_2$, DCM/MeOH = 20:1 to 10:1) afforded the desired product **ABA-1-18** (65 mg, 38%) as a fluffy white solid. $R_f$ = 0.25 (DCM/MeOH = 10:1); HPLC purity 99%, RT = 14.64 min with method 12, RT = 15.48 min with method 13; $^1$H NMR ($D_2O$, 500 MHz): $\delta$ 7.92 (s, 1H, H-9), 6.80 (m, 1H, H-14), 5.98 (dq, $J$ = 15.4, 1.7 Hz, 1H, H-13), 4.57 (t, $J$ = 5.0 Hz, 2H, H-8), 4.49 (s, 2H, H-11), 3.88 (s, 1H, H-5), 3.81 (m, 1H, H-7a), 3.71 (m, 1H, H-7b), 3.40 (d, $J$ = 11.2 Hz, 1H, H-1a), 3.26 (d, $J$ = 11.3 Hz, 1H, H-1b), 1.85 (dd, $J$ = 6.9, 1.8 Hz, 3H, H-15), 0.73 (s, 3H, H-2), 0.70 (s, 3H, H-3); $^{13}$C NMR ($D_2O$, 125 MHz): $\delta$ 175.4, 168.7, 144.8, 142.4, 124.0, 123.5, 75.6, 68.3, 49.5, 38.5, 38.4, 34.2, 20.3, 18.8, 17.2; HRMS (ESI$^+$) *m/z* $[C_{15}H_{26}N_5O_4]^+$ calcd: 340.1979; found: 340.1971.

Synthesis of **ABA-1-19 (4-12)**

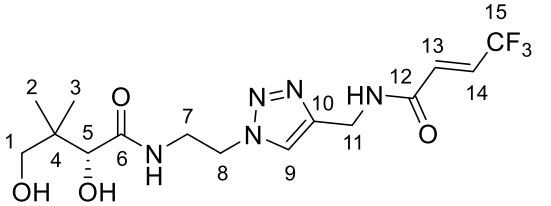

**ABA-1-19**

Following the general procedure P, **ABA-1-1** (51 mg, 0.3 mmol), **ABA-1-9** (63 mg, 0.3 mmol), sodium ascorbate (58 mg, 0.3 mmol), $CuSO_4$ (5 mg, 0.03 mmol), and EtOH/$H_2O$ = 1:1 (3 mL) were used. Purification by flash chromatography ($SiO_2$, DCM/MeOH = 20:1 to 10:1) yielded the desired product **ABA-1-19** (70 mg, 61%) as a fluffy white solid. $R_f$ = 0.25 (DCM/MeOH = 10:1); HPLC purity 99%, RT = 17.67 min with method 12, RT = 16.98 min with method 13; $^1$H NMR ($D_2O$, 500 MHz): $\delta$ 7.97 (s, 1H, H-9), 6.83 – 6.70 (m, 2H, H-13, H-14), 4.61 – 4.54 (m, 4H, H-8, H-11), 3.89 (s, 1H, H-5), 3.82 (m, 1H, H-7a), 3.71 (m, 1H, H-7b), 3.41 (d, $J$ = 11.2 Hz, 1H, H-1a), 3.26 (d, $J$ = 11.2 Hz, 1H, H-1b), 0.73 (s, 3H, H-2), 0.70 (s, 3H, H-3); $^{19}$F NMR ($D_2O$, 471 MHz):

$\delta$ -65.0; $^{13}$C NMR ($D_2O$, 125 MHz): $\delta$ 175.4, 165.2, 144.2, 130.7 (q, $J$ = 6.2 Hz), 128.0 (q, $J$ = 34.9 Hz), 124.2, 122.4 (q, $J$ = 269.3 Hz), 75.6, 68.3, 49.5, 38.5, 38.4, 34.5, 20.3, 18.8; HRMS (ESI$^+$) $m/z$ $[C_{15}H_{22}N_5O_4F_3Na]^+$ calcd: 416.1516; found: 416.1502.

Synthesis of **ABA-1-26 (4-13)**

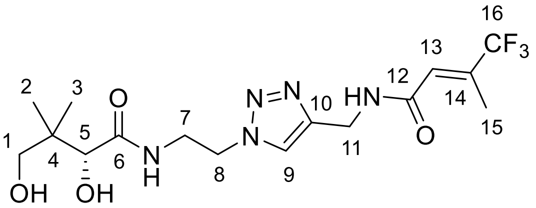

**ABA-1-26**

Following the general procedure P, **ABA-1-23** (51 mg, 0.3 mmol), **ABA-1-9** (63 mg, 0.3 mmol), sodium ascorbate (58 mg, 0.3 mmol), $CuSO_4$ (5 mg, 0.03 mmol), and EtOH/$H_2O$ = 1:1 (3 mL) were used. Purification by flash chromatography ($SiO_2$, DCM/MeOH = 20:1 to 10:1) yielded the desired product **ABA-1-26** (70 mg, 66%) as an off-white solid. $R_f$ = 0.31 (DCM/MeOH = 10:1); HPLC purity 95%, RT = 19.19 min with method 12, RT = 17.49 min with method 13; $^1$H NMR ($D_2O$, 500 MHz): $\delta$ 7.97 (s, 1H, H-9), 6.49 (m, 1H, H-13), 4.59 (t, $J$ = 5.6 Hz, 2H, H-8), 4.54 (s, 2H, H-11), 3.90 (s, 1H, H-5), 3.84 (m, 1H, H-7a), 3.72 (m, 1H, H-7b), 3.41 (d, $J$ = 11.2 Hz, 1H, H-1a), 3.27 (d, $J$ = 11.2 Hz, 1H, H-1b), 2.09 (d, $J$ = 1.5 Hz, 3H, H-15), 0.74 (s, 3H, H-2), 0.71 (s, 3H, H-3); $^{19}$F NMR ($D_2O$, 471 MHz): $\delta$ -70.8; $^{13}$C NMR ($D_2O$, 125 MHz): $\delta$ 175.4, 167.1, 144.4, 136.9 (d, $J$ = 30.2 Hz), 123.3 (q, $J$ = 273.5 Hz), 75.6, 68.3, 49.5, 38.5, 38.4, 34.2, 20.3, 18.8, 11.6; HRMS (ESI$^+$) $m/z$ $[C_{16}H_{24}N_5O_4F_3Na]^+$ calcd: 430.1673; found: 430.1685.

Synthesis of **ABA-1-34 (6-13)**

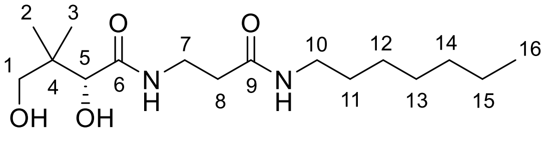

**ABA-1-34**

Following the general procedure N with modification, **ABA-1-32** (2.32 g, 8.1 mmol), EA (16 mL), MeOH (2.60 mL), and AcCl (4.57 mL) were used in a 100 mL round-bottom flask to yield **ABA-1-33** as a colorless oil (726 mg), which was utilized in the next step without further purification. Following the general procedure O, **ABA-1-33** (726 mg), D-pantolactone (1.02 g, 7.8 mmol), TBD

(54 mg, 0.4 mmol), and toluene (4 mL) were used. Purification by flash chromatography ($SiO_2$, DCM/EA = 4:1 then DCM/MeOH = 20:1) yielded the desired product **ABA-1-34** (656 mg, 26% over two steps) as a white solid. $R_f$ = 0.53 (DCM/MeOH = 10:1); HPLC purity 95%, RT = 30.67 min with method 12, RT = 19.24 min with method 13; $^1$H NMR ($CDCl_3$, 500 MHz): $\delta$ 7.51 (bs, 1H, NH), 6.38 (bs, 1H, NH), 4.59 (bs, 1H, OH), 3.98 (s, 1H, H-5), 3.63 – 3.33 (m, 4H, H-1, H-7), 3.22 – 3.12 (m, 2H, H-10), 2.42 (t, $J$ = 6.1 Hz, 2H, H-8), 1.53 – 1.40 (m, 2H, H-11), 1.36 – 1.16 (m, 8H, H-12, H-13, H-14, H-15), 0.97 (s, 3H, H-2), 0.91 (s, 3H, H-3), 0.86 (t, $J$ = 6.8 Hz, 3H, H-16); $^{13}$C NMR ($CDCl_3$, 125 MHz): $\delta$ 174.1, 171.6, 70.9, 39.9, 39.4, 35.9, 35.4, 31.8, 29.6, 29.1, 27.0, 22.7, 21.4, 20.6, 14.2; HRMS (ESI$^+$) $m/z$ $[C_{16}H_{32}N_2O_4Na]^+$ calcd: 339.2254; found: 339.2259. The characterization matched that reported in the literature.[47]

Synthesis of **SCC-3-98**

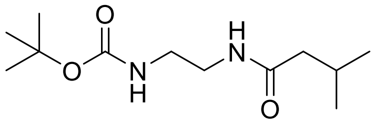

**SCC-3-98**

Following the general procedure Q, *N*-Boc-ethylenediamine (1.39 g, 8.70 mmol), 3-methylbutanoic acid (978 mg, 6.78 mmol), DMAP (117 mg, 0.96 mmol), anhydrous DCM (40 mL), and EDCI (1.83 g, 9.57 mmol) were used. Purification by flash chromatography ($SiO_2$, DCM/MeOH = 25:1 to 10:1) yielded **SCC-3-98** (1.74 g, 82%) as a white solid. $R_f$ = 0.51 (DCM/MeOH = 20:1). $^1$H NMR ($CDCl_3$, 500 MHz): $\delta$ 6.29 (bs, 1H), 5.06 (bs, 1H), 3.38 (m, 2H), 3.28 (q, $J$ = 5.9 Hz, 2H), 2.11 (m, 1H), 2.05 (d, $J$ = 6.5 Hz, 2H), 1.44 (s, 9H), 0.95 (d, $J$ = 6.4 Hz, 6H); $^{13}$C NMR ($CDCl_3$, 125 MHz): $\delta$ 173.4, 157.0, 79.6, 46.1, 40.6, 40.4, 28.4, 26.1, 22.5; HRMS (ESI$^+$) *m/z* $[C_{12}H_{24}N_2O_3Na]^+$ calcd: 267.1679 ; found: 267.1685.

Synthesis of **SCC-3-93**

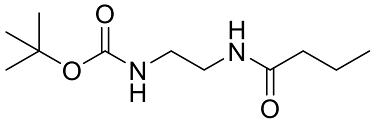

**SCC-3-93**

Following the general procedure Q, *N*-Boc-ethylenediamine (826 mg, 5.16 mmol), butyric acid (500 mg, 5.67 mmol), DMAP (70 mg, 0.57 mmol), anhydrous DCM (30 mL), and EDCI (1.09 g, 5.67 mmol) were used. Purification by flash chromatography ($SiO_2$, DCM/MeOH = 25:1 to 10:1) afforded **SCC-3-93** (840 mg, 70%) as a white solid. $R_f$ = 0.51 (DCM/MeOH = 20:1); $^1$H NMR ($CDCl_3$, 500 MHz): $\delta$ 6.29 (bs, 1H), 5.04 (bs, 1H), 3.39 (m, 2H), 3.18 (m, 2H), 2.17 (t, $J$ = 7.5 Hz, 2H), 1.66 (h, 2H), 1.45 (s, 9H), 0.95 (t, $J$ = 7.4 Hz, 3H); $^{13}$C NMR ($CDCl_3$, 125 MHz): $\delta$ 173.8, 157.0, 79.7, 40.7, 40.3, 38.6, 28.4, 19.1, 13.8; HRMS (ESI$^+$) *m/z* $[C_{11}H_{22}N_2O_3Na]^+$ calcd: 253.1523; found: 253.1518.

Synthesis of **SCC-2-96**

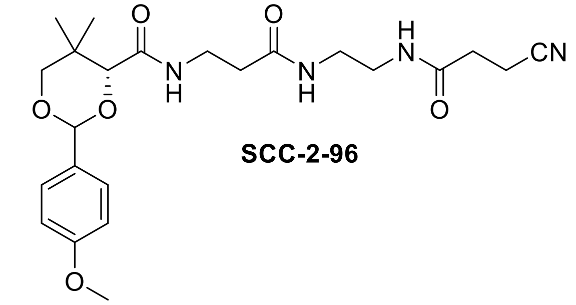

**SCC-2-96**

To **PMP-NH2** (475 mg, 1.25 mmol, 1.1 equiv.) was added anhydrous THF (40 mL), followed by 3-cyanopropanoic acid (130 mg, 1.32 mmol, 1.0 equiv.). A solution of DMAP (16 mg, 0.13 mmol, 0.1 equiv.) in anhydrous THF (2 mL), and a solution of DCC (271 mg, 1.32 mmol, 1.0 equiv.) in anhydrous THF (5 mL) were then added to the reaction mixture. The mixture was stirred at rt overnight. The mixture was then filtered, and the filter cake was washed with THF (10 mL). Upon collecting the filtrate, the solvent was evaporated under reduced pressure. The product was purified from the residue by flash chromatography ($SiO_2$, DCM/MeOH = 25:1 to 10:1), affording **SCC-2-96** (290 mg, 50%) as a white solid. $R_f$ = 0.65 (DCM/MeOH = 10:1); $^1$H NMR ($CDCl_3$, 500 MHz): $\delta$ 7.45 (m, 2H), 7.06 (t, $J$ = 6.3 Hz, 1H), 7.01 (m, 1H), 6.93 (m, 2H), 6.80 (m, 1H), 5.49 (s, 1H), 4.10 (s, 1H), 3.84 (s, 3H), 3.76-3.63 (m, 2H), 3.60-3.51 (m, 2H), 3.43-3.29 (m, 4H), 2.65 (t, $J$ = 7.0 Hz, 2H), 2.49 (t, $J$ = 7.5 Hz, 2H), 2.43 (m, 2H), 1.12 (s, 3H) 1.09 (s, 3H); $^{13}$C NMR ($CDCl_3$, 125 MHz): $\delta$ 172.2, 169.9, 169.7, 160.3, 130.1, 127.6, 119.3, 113.8, 101.4, 83.8, 78.4, 55.4, 40.2, 39.4, 36.4, 35.1, 33.1, 31.6, 21.9, 19.1, 13.5; HRMS (ESI$^+$) $m/z$ $[C_{23}H_{32}N_4O_6Na]^+$ calcd: 483.2214 found: 483.2198.

Synthesis of **SCC-3-43**

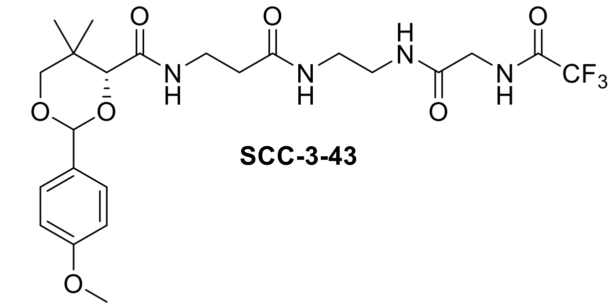

**SCC-3-43**

Following the general procedure Q, *N*-Boc-ethylenediamine (624 mg, 3.90 mmol), (2,2,2-trifluoroacetyl)glycine (734 mg, 4.29 mmol), DMAP (48 mg, 0.39 mmol), anhydrous DCM (30 mL), and EDCI (822 mg, 4.29 mmol) were used. The product, **SCC-3-39**, was used in the next step without further purification. Following procedure R with modification, **SCC-3-39** (470 mg of crude), anhydrous DCM (30 mL), and TFA (1.50 mL, 19.7 mmol) were used in the deprotection step. Afterwards, **PMP-pantothenate** (439 mg, 1.31 mmol), DMAP (17 mg, 0.14 mmol), EDCI (251 mg, 1.31 mmol), and anhydrous DCM (30 mL) were used for the amide coupling step of general procedure R. Purification by flash chromatography ($SiO_2$, DCM/MeOH = 25:1 to 10:1) afforded **SCC-3-34** (321 mg, 46% over two steps) as a white solid. $R_f$ = 0.55 (DCM/MeOH = 10:1); $^1$H NMR ($CDCl_3$, 500 MHz): $\delta$ 8.04 (t, $J$ = 5.4 Hz, 1H), 7.45 (t, $J$ = 5.0 Hz, 1H), 7.41 (d, $J$ = 8.8 Hz, 2H), 7.04 (t, $J$ = 6.4 Hz, 1H), 6.99 (m, 1H), 6.90 (d, $J$ = 8.8 Hz, 2H), 5.44 (s, 1H), 4.05 (s, 1H), 3.91 (d, $J$ = 5.3 Hz, 2H), 3.79 (s, 3H), 3.72 – 3.60 (m, 2H), 3.49 (m, 2H), 3.36 – 3.26 (m, 4H), 2.37 (t, $J$ = 6.5 Hz, 2H), 1.07 (s, 3H), 1.03 (s, 3H); $^{19}$F NMR ($CDCl_3$, 471 MHz): $\delta$ -75.6; $^{13}$C NMR ($CDCl_3$, 125 MHz): $\delta$ 172.3, 170.0, 167.9, 160.3, 157.7 (q, $J$ = 37.5 Hz), 130.1, 127.6, 116.9 (q, $J$ = 299.7 Hz), 113.8, 101.4, 83.8, 78.4, 55.3, 42.7, 40.0, 39.0, 36.3, 35.2, 33.1, 21.8, 19.1; HRMS (ESI$^+$) *m/z* $[C_{23}H_{31}N_4O_7F_3Na]^+$ calcd: 555.2037; found: 555.2027.

Synthesis of **SCC-3-95**

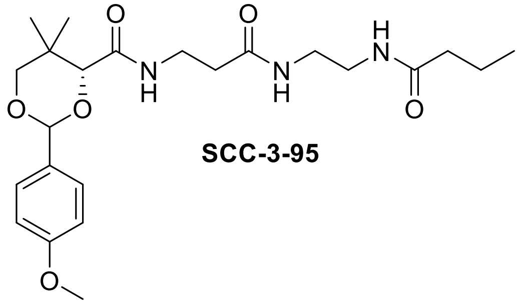

**SCC-3-95**

Following procedure R with modification, **SCC-3-93** (335 mg, 1.45 mmol), anhydrous DCM (30 mL), and TFA (1.50 mL, 19.7 mmol) were used in the deprotection step. Afterwards, **PMP-pantothenate** (521 mg, 1.54 mmol), DMAP (19 mg, 0.15 mmol), EDCI (296 mg, 1.54 mmol), and anhydrous DCM (30 mL) were used for the amide coupling step of procedure R. Purification by flash chromatography ($SiO_2$, DCM/MeOH = 25:1 to 10:1) afforded **SCC-3-95** (420 mg, 70% over two steps) as a white solid. $R_f$ = 0.60 (DCM/MeOH = 10:1); $^1$H NMR ($CDCl_3$, 500 MHz): $\delta$ 7.40 (d, $J$ = 8.7 Hz 2H), 7.12 – 7.02 (m, 2H), 6.88 (d, $J$ = 8.7 Hz, 2H), 6.61 (s, 1H), 5.43 (s, 1H), 4.04 (s, 1H), 3.78 (s, 3H), 3.72 – 3.59 (m, 2H), 3.57 – 3.46 (m, 2H), 3.40 – 3.17 (m, 4H), 2.38 (t, $J$ = 6.4 Hz, 2H), 2.15 – 2.02 (m, 2H), 1.58 (h, $J$ = 7.4 Hz, 2H), 1.07 (s, 3H) 1.05 (s, 3H), 0.90 (t, $J$ = 7.4 Hz, 3H); $^{13}$C NMR ($CDCl_3$, 125 MHz): $\delta$ 174.2, 172.0, 169.6, 160.2, 130.2, 127.5, 113.7, 101.3, 83.9, 78.4, 55.3, 39.8, 38.5, 36.0, 35.0, 33.1, 21.9, 19.1, 13.7; HRMS (ESI$^+$) *m/z* $[C_{23}H_{35}N_3O_6Na]^+$ calcd: 472.2418; found: 472.2410.

Synthesis of **SCC-3-99**

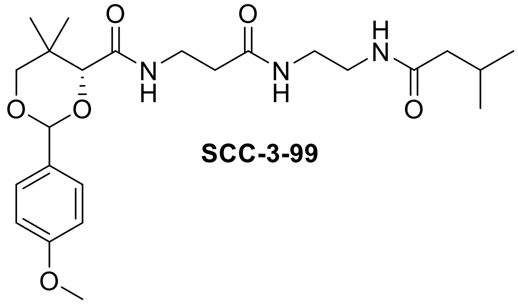

**SCC-3-99**

Following procedure R, **SCC-3-98** (361 mg, 1.48 mmol), anhydrous DCM (30 mL), and TFA (1.50 mL, 19.7 mmol) were used in the deprotection step. Afterwards, **PMP-pantothenate** (500 mg, 1.48 mmol), DMAP (18 mg, 0.15 mmol), EDCI (284 mg, 1.48 mmol), and anhydrous DCM (30 mL) were used for the amide coupling step of procedure R. Purification by flash chromatography ($SiO_2$, DCM/MeOH = 25:1 to 10:1) afforded **SCC-3-99** (370 mg, 54% over two steps) as a white solid. $R_f$ = 0.50 (DCM/MeOH = 10:1); $^1$H NMR ($CDCl_3$, 500 MHz): $\delta$ 7.45 (d, $J$ = 8.4 Hz, 2H), 7.06 (t, $J$ = 6.2 Hz, 1H), 6.93 (d, $J$ = 8.5 Hz, 2H), 6.87 (s, 1H), 6.40 (s, 1H), 5.48 (s, 1H), 4.09 (s, 1H), 3.83 (s, 3H), 3.72 (d, $J$ = 11.5 Hz, 1H), 3.67 (d, $J$ = 11.4 Hz, 1H), 3.61 – 3.47 (m, 2H), 3.43 – 3.24 (m, 4H), 2.43 (t, $J$ = 6.3 Hz, 2H), 2.13 – 2.04 (m, 1H), 2.02 (d, $J$ = 6.8 Hz, 2H), 1.12 (s, 3H), 1.10 (s, 3H), 0.93 (d, $J$ = 6.4, 1.5 Hz, 6H); $^{13}$C NMR ($CDCl_3$, 125 MHz): $\delta$ 173.7, 171.9, 169.6, 160.3, 130.2, 127.6, 113.8, 101.3, 83.9, 78.5, 55.4, 45.9, 40.1, 39.8, 36.1, 34.9, 33.1, 26.1, 22.4, 21.9, 19.1; HRMS (ESI$^+$) $m/z$ $[C_{24}H_{37}N_3O_6Na]^+$ calcd: 486.2575; found: 486.2576.

Synthesis of **SCC-3-39 (Crude)**

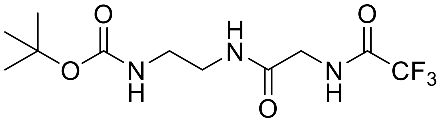

**SCC-3-39**

Following the general procedure Q, *N*-Boc-ethylenediamine (624 mg, 3.90 mmol), (2,2,2-trifluoroacetyl)glycine (734 mg, 4.29 mmol), DMAP (48 mg, 0.39 mmol), anhydrous DCM (30 mL), and EDCI (822 mg, 4.29 mmol) were used. Purification by flash chromatography ($SiO_2$, DCM/MeOH = 25:1 to 10:1) afforded **SCC-3-39** (281 mg, 23%) as a pale-yellow solid. $R_f$ = 0.40 (DCM/MeOH = 20:1); $^1$H NMR ($CDCl_3$, 500 MHz): $\delta$ 7.59 (s, 1H), 7.05 (s, 1H), 5.03 (s, 1H), 4.02 (s, 2H), 3.43 (s, 2H), 3.30 (s, 2H), 1.45 (s, 9H); HRMS (ESI$^+$) $m/z$ $[C_{11}H_{18}N_3O_4F_3Na]^+$ calcd: 336.1142; found: 336.1138.

Synthesis of **SCC-2-99 (6-01)**

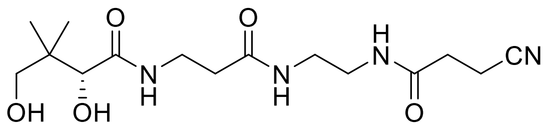

**SCC-2-99**

Following procedure S, **SCC-2-96** (89.3 mg, 0.19 mmol) and a 9:1 mixture of AcOH/$H_2O$ (10 mL) were used. **SCC-2-99** (53.1 mg, 80%) was obtained as a white solid. $R_f$ = 0.19 (DCM/MeOH = 10:1); HPLC purity 95%, RT = 12.44 min with method 14, RT = 6.18 min with method 15; $^1$H NMR ($CD_3OD$, 500 MHz): $\delta$ 3.92 (s, 1H), 3.64 – 3.38 (m, 4H), 3.40 – 3.24 (m, 4H), 2.78 – 2.67 (m, 2H), 2.64 – 2.56 (m, 2H), 2.45 (t, $J$ = 6.7 Hz, 2H), 0.95 (s, 6H); $^{13}$C NMR ($CD_3OD$, 125 MHz): $\delta$ 176.0, 174.2, 172.4, 120.6, 77.3, 70.3, 40.3, 40.1, 40.0, 36.6, 36.4, 32.1, 21.3, 20.9, 13.7; HRMS (ESI$^+$) *m/z* $[C_{15}H_{26}N_4O_5Na]^+$ calcd: 365.1795; found: 365.1787.

Synthesis of **SCC-3-103 (3-01)**

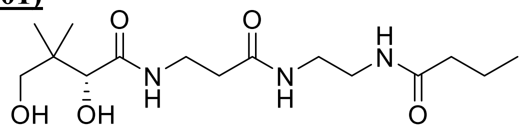

**SCC-3-103**

Following procedure S, **SCC-3-95** (160 mg, 0.36 mmol) and a 9:1 mixture of AcOH/$H_2O$ (5 mL) were used. **SCC-3-103** (100 mg, 85%) was obtained as a white solid. HPLC purity >99%, RT = 13.23 min with method 14, RT = 5.20 min with method 15; $^1$H NMR ($CD_3OD$, 500 MHz): $\delta$ 3.89 (s, 1H), 3.58 – 3.34 (m, 4H), 3.31 – 3.23 (m, 4H), 2.42 (t, $J$ = 6.7 Hz, 2H), 2.21 – 2.12 (m, 2H), 1.63 (h, $J$ = 7.4 Hz, 2H), 0.94 (t, $J$ = 7.5 Hz, 3H), 0.92 (s, 6H); $^{13}$C NMR ($CD_3OD$, 125 MHz): $\delta$ 176.4, 176.0, 174.1, 77.3, 70.3, 40.3, 40.1, 39.9, 39.0, 36.6, 36.4, 21.3, 20.9, 20.3, 14.0; HRMS (ESI$^+$) *m/z* $[C_{15}H_{29}N_3O_5Na]^+$ calcd: 354.1999; found: 354.1994.

Synthesis of **SCC-3-102 (3-02)**

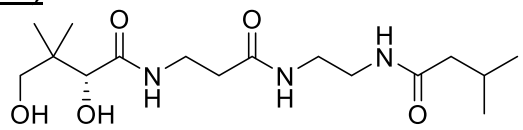

**SCC-3-102**

Following procedure S, **SCC-3-99** (128 mg, 0.28 mmol) and a 9:1 mixture of AcOH/$H_2O$ (10 mL) were used. **SCC-3-102** (82 mg, 86%) was obtained as a white solid. $R_f$ = 0.29 (DCM/MeOH = 10:1); HPLC purity 98%, RT = 14.03 min with method 14, RT = 5.17 min with method 15; $^1$H NMR ($CD_3OD$, 500 MHz); $\delta$ 3.89 (s, 1H), 3.59 – 3.35 (m, 4H), 3.34 – 3.20 (m, 4H), 2.42 (t, $J$ =

6.7 Hz, 2H), 2.18 – 1.95 (m, 3H), 0.94 (d, $J$ = 6.5 Hz, 6H), 0.92 (s, 6H); $^{13}$C NMR ($CD_3OD$, 125 MHz): $\delta$ 174.7, 174.5, 172.7, 76.0, 69.0, 45.0, 39.0, 38.8, 38.4, 35.2, 35.0, 25.9, 21.4, 20.0, 19.5; HRMS (ESI$^+$) $m/z$ $[C_{16}H_{31}N_3O_5Na]^+$ calcd: 368.2161; found: 368.2156.

Synthesis of **SCC-3-64 (4-11)**

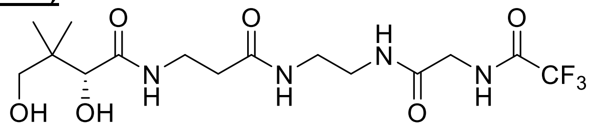

**SCC-3-64**

Following procedure S, **SCC-3-43** (127 mg, 0.24 mmol) and a 9:1 mixture of AcOH/$H_2O$ (10 mL) were used. **SCC-3-64** (80 mg, 81%) was obtained as a white solid. $R_f$ = 0.25 (DCM/MeOH = 10:1); HPLC purity 94%, RT = 13.60 min with method 14, RT = 6.82 min with method 15; $^1$H NMR ($CD_3OD$, 500 MHz): $\delta$ 3.96 (s, 2H), 3.92 (s, 1H), 3.58 – 3.45 (m, 3H), 3.45 – 3.26 (m, 5H), 2.42 (t, $J$ = 6.6 Hz, 2H), 0.94 (s, 6H); $^{13}$C NMR ($CD_3OD$, 125 MHz): $\delta$ 174.7, 172.9, 168.9, 158.2 (q, $J$ = 37.2 Hz), 116.0 (q, $J$ = 286.4 Hz), 76.0, 69.0, 42.0 (d, $J$ = 4.6 Hz), 39.0, 38.9, 38.5, 35.3, 35.0, 20.0, 19.5; HRMS (APCI$^+$) $m/z$ $[C_{15}H_{26}N_4O_6F_3]^+$ calcd: 415.1799; found: 415.1801.

### NMR spectra

#### [1]H spectrum of **DD-05-140**

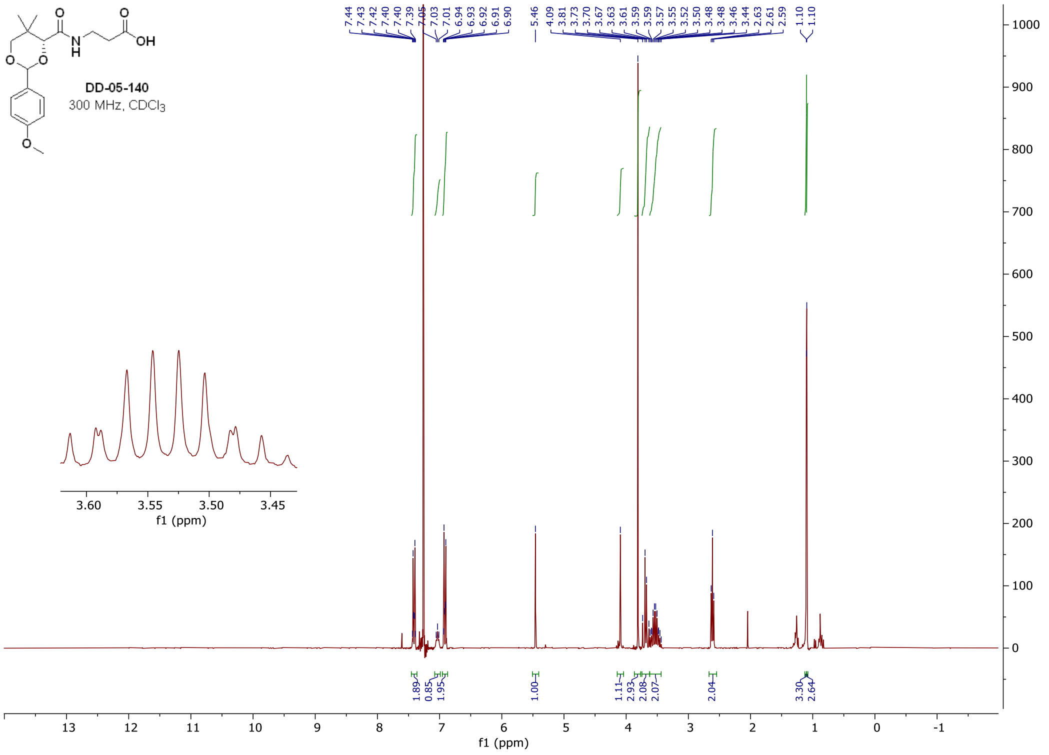

#### [1]H spectrum of **DD-05-006**

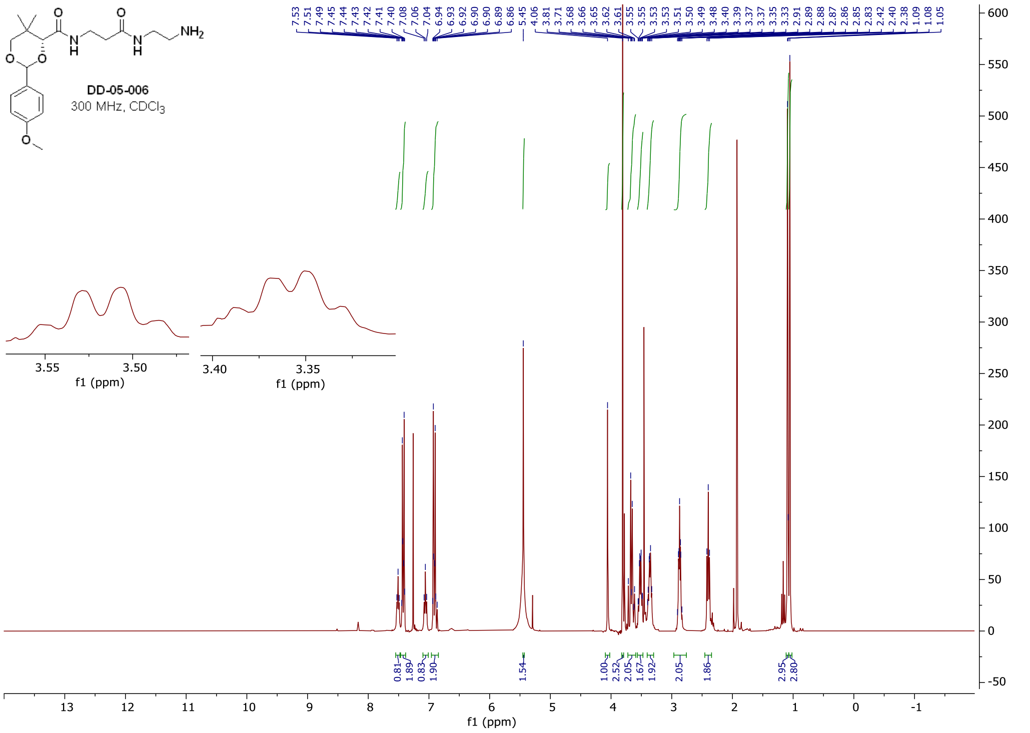

### $^{1}H$ and $^{13}C$ spectra of **DD-06-123**

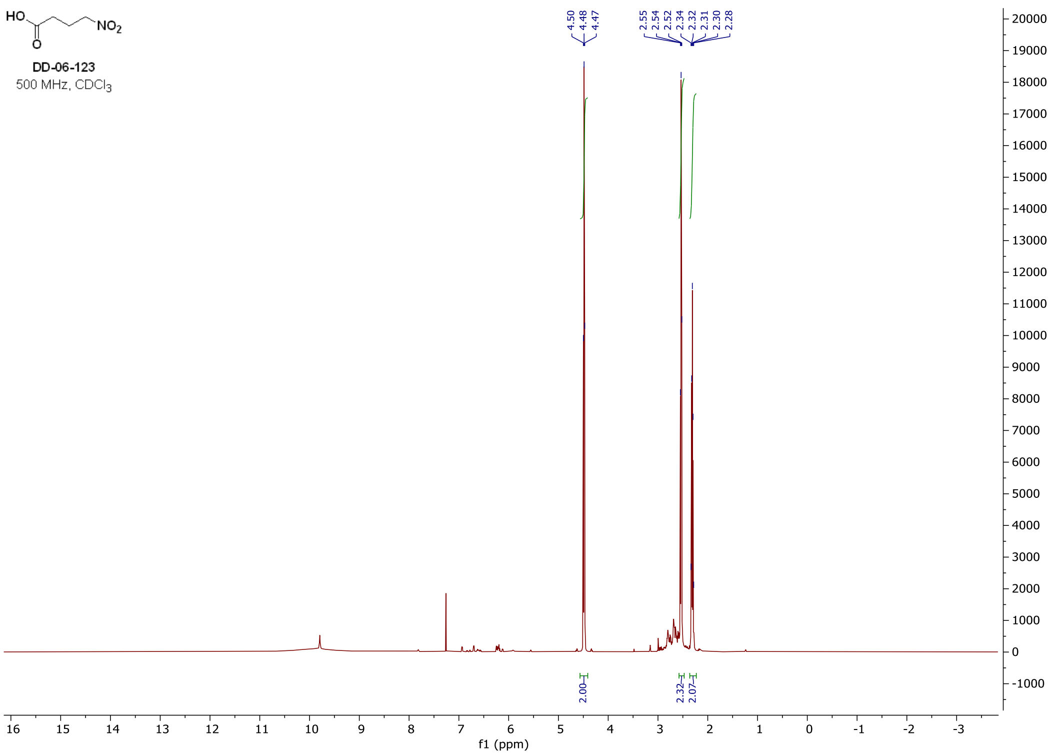

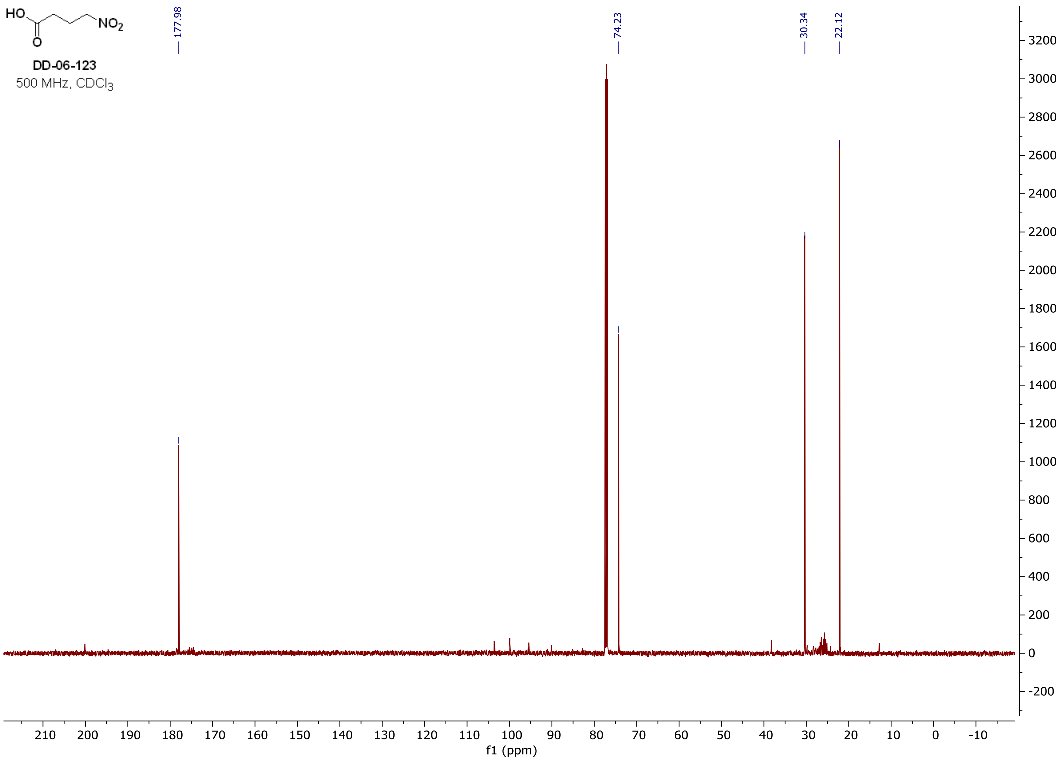

### [1]H and [13]C spectra of **DD-06-124**

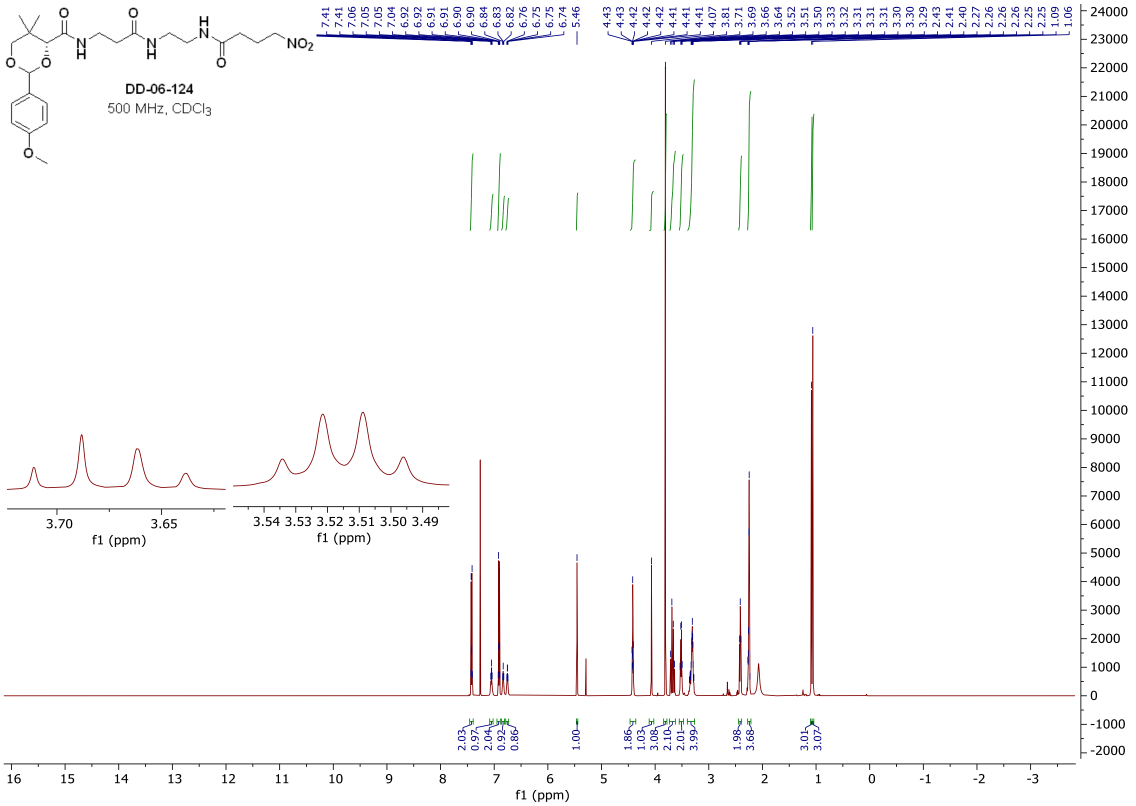

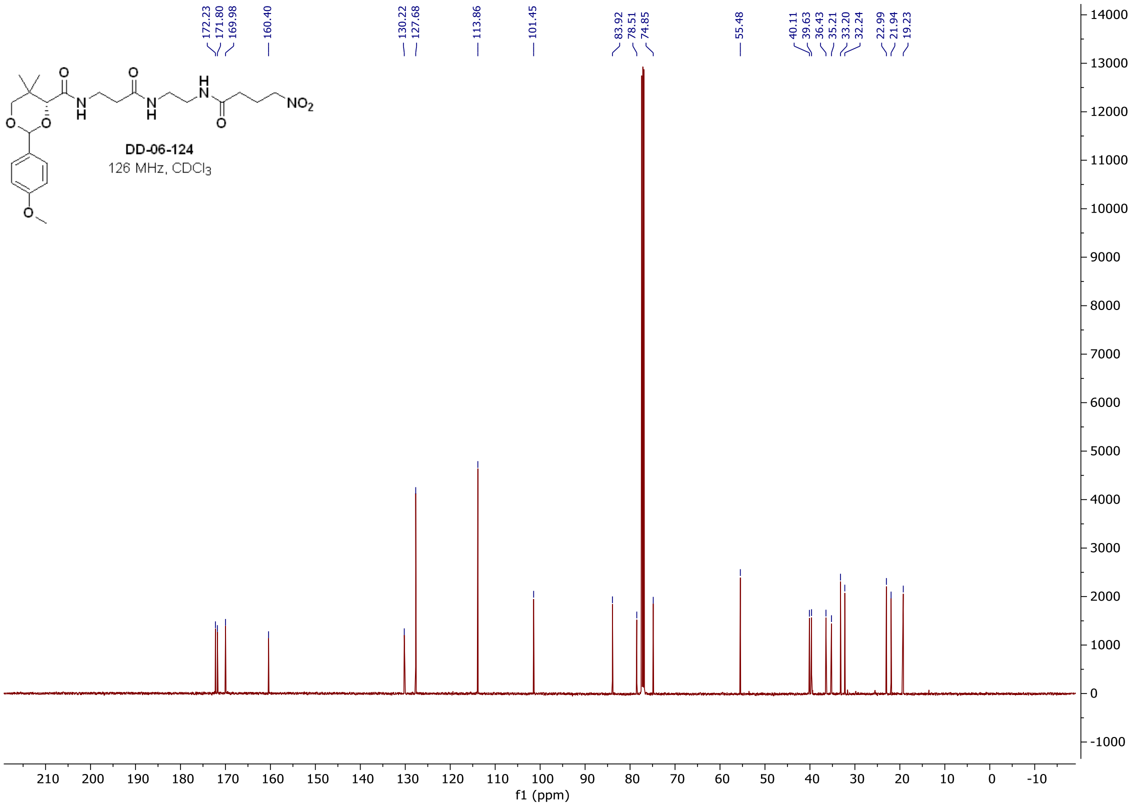

$^{1}H$ and $^{13}C$ spectra of **DD-06-143**

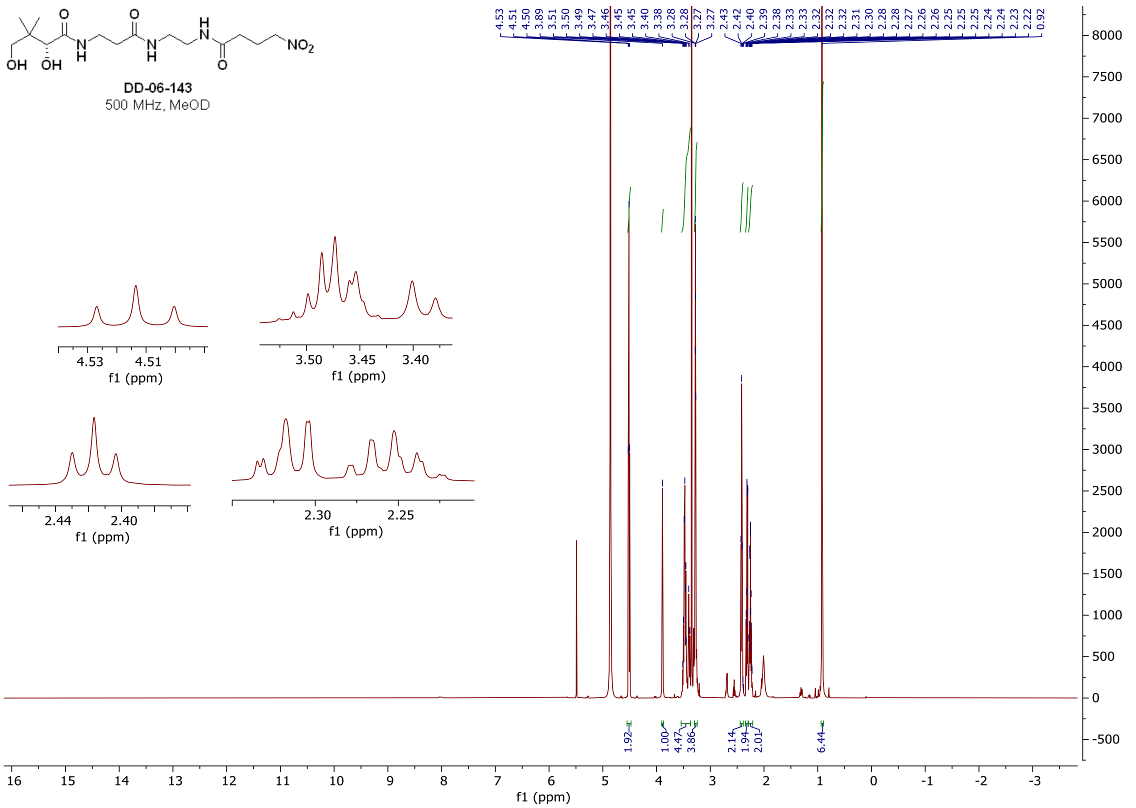

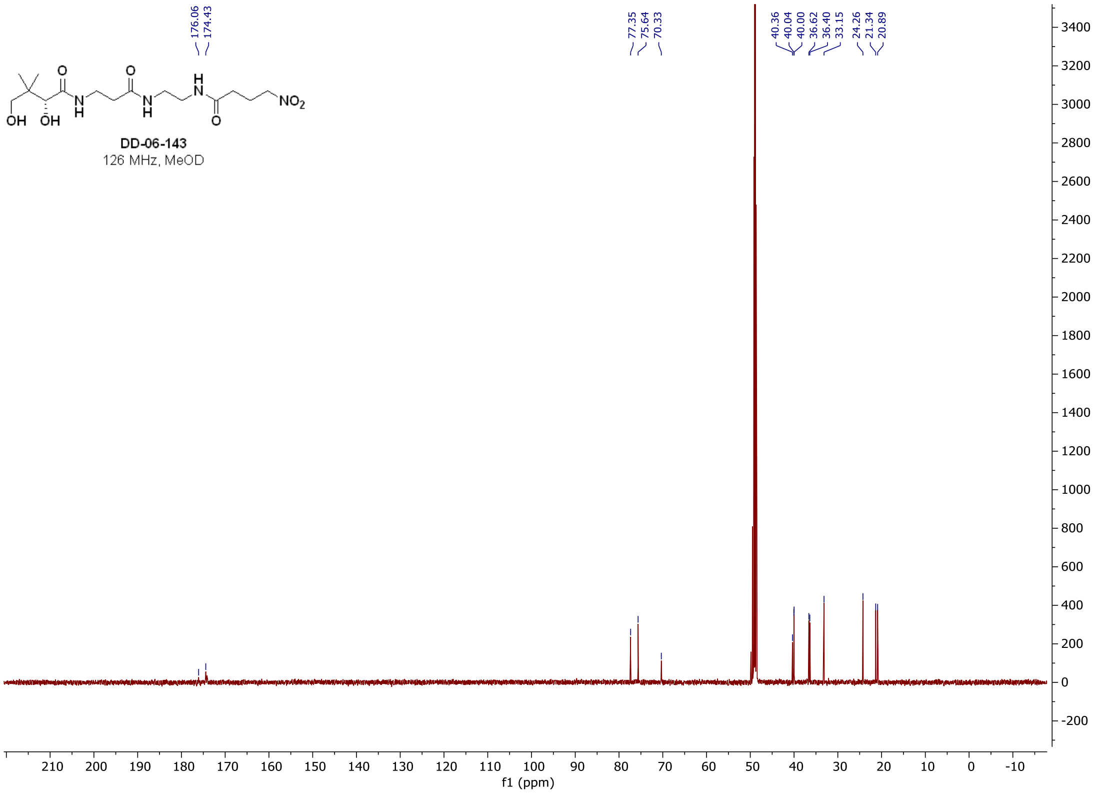

#### $^{1}$H spectrum of **DD-05-131**

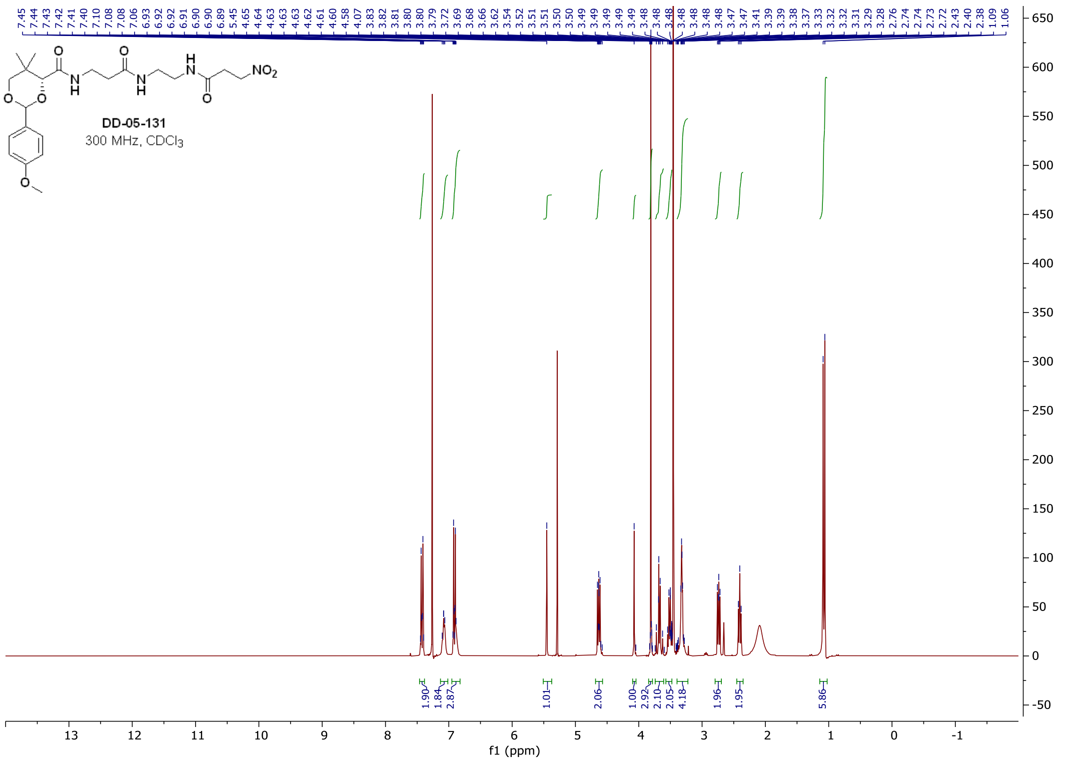

#### $^{1}$H, $^{13}$C, and $^{19}$F spectra of **DD-06-111**

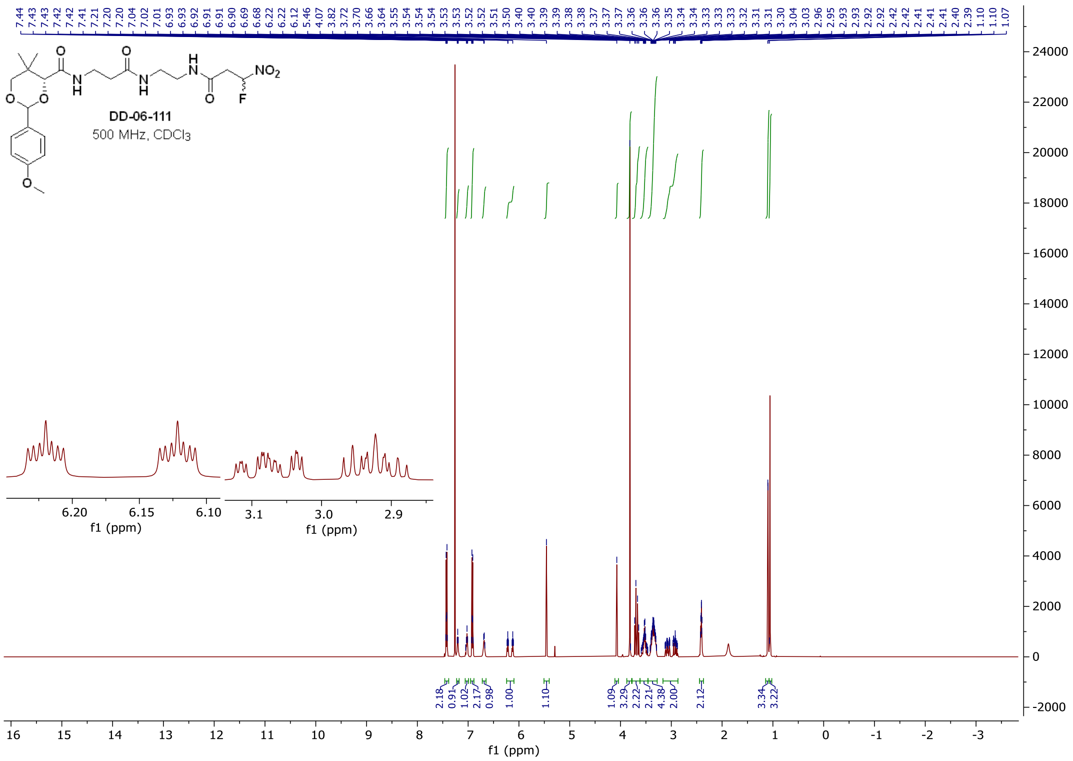

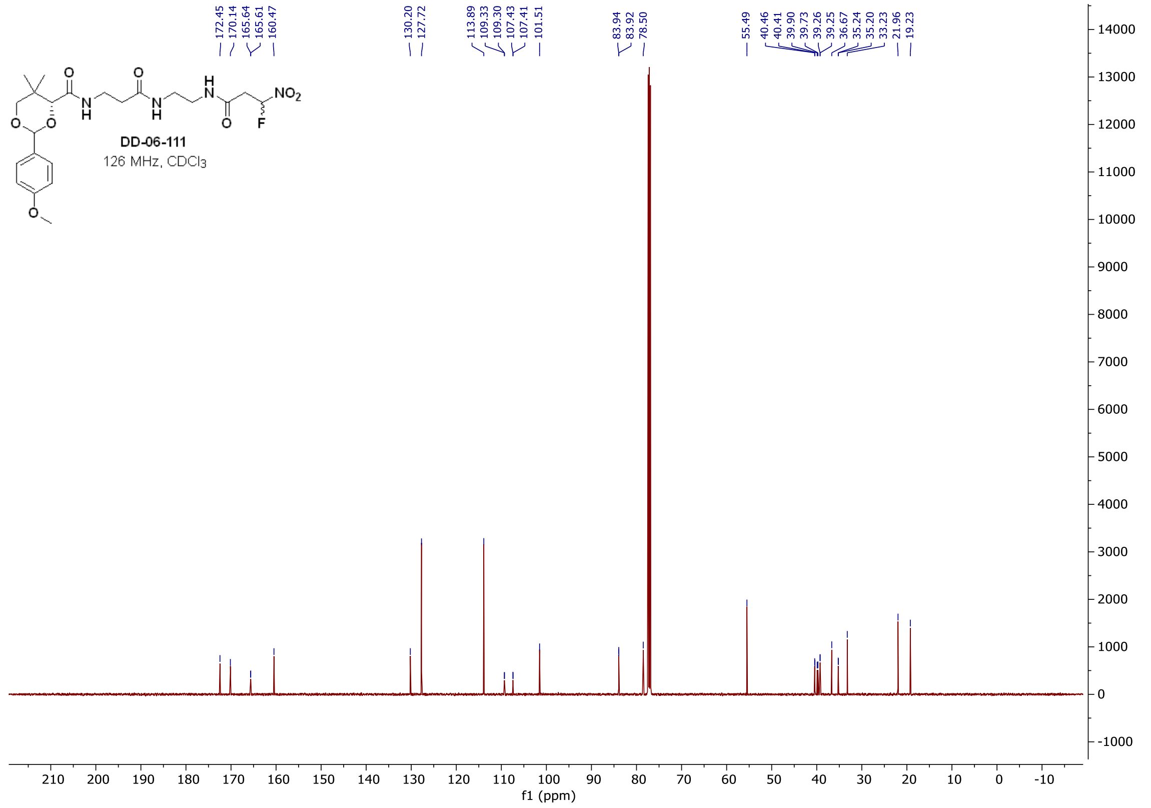
DD-06-111
126 MHz, CDCl3
172.45
170.14
165.64
165.61
160.47
130.20
127.72
113.89
109.33
109.30
107.43
107.41
101.51
83.94
83.92
78.50
55.49
40.46
40.41
39.90
39.73
39.26
39.25
36.67
35.24
35.20
33.23
21.96
19.23
f1 (ppm)

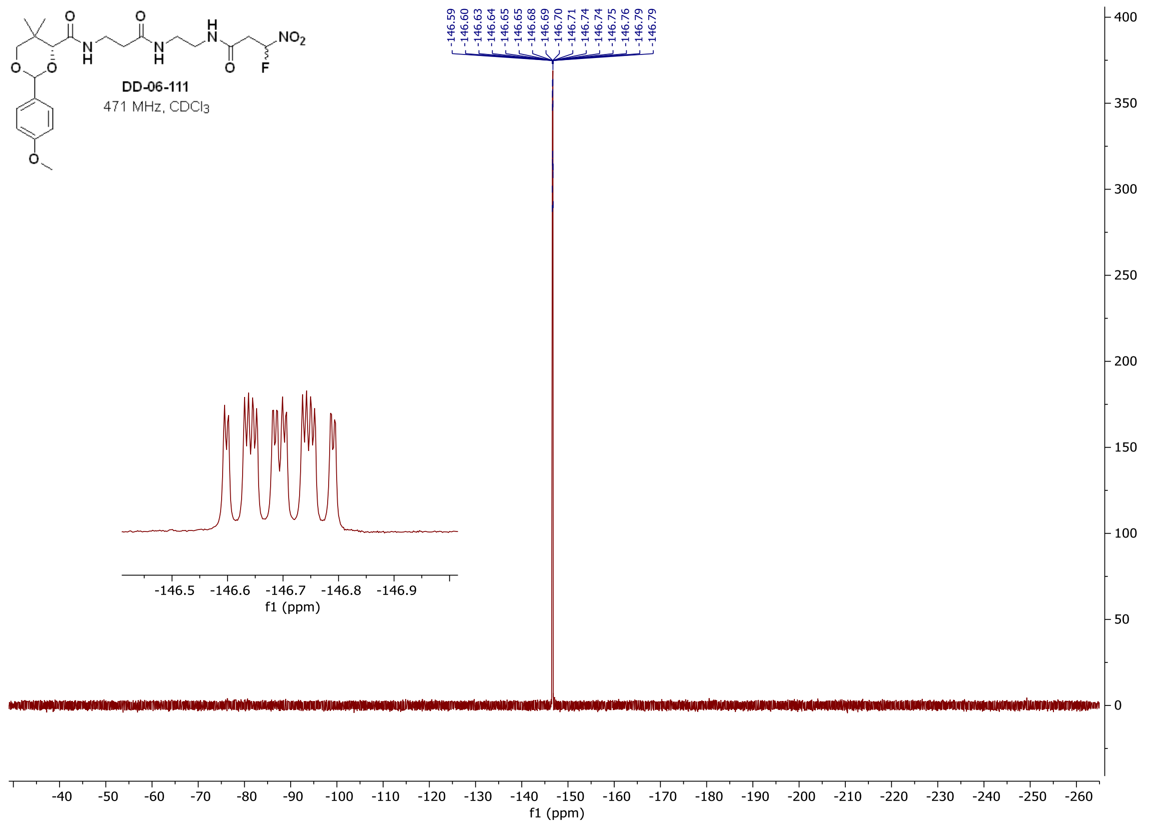
DD-06-111
471 MHz, CDCl3
-146.59
-146.60
-146.63
-146.64
-146.65
-146.65
-146.68
-146.69
-146.70
-146.71
-146.74
-146.75
-146.76
-146.79
-146.79
f1 (ppm)

$^{1}H$, $^{13}C$, and $^{19}F$ spectra of **DD-07-066**

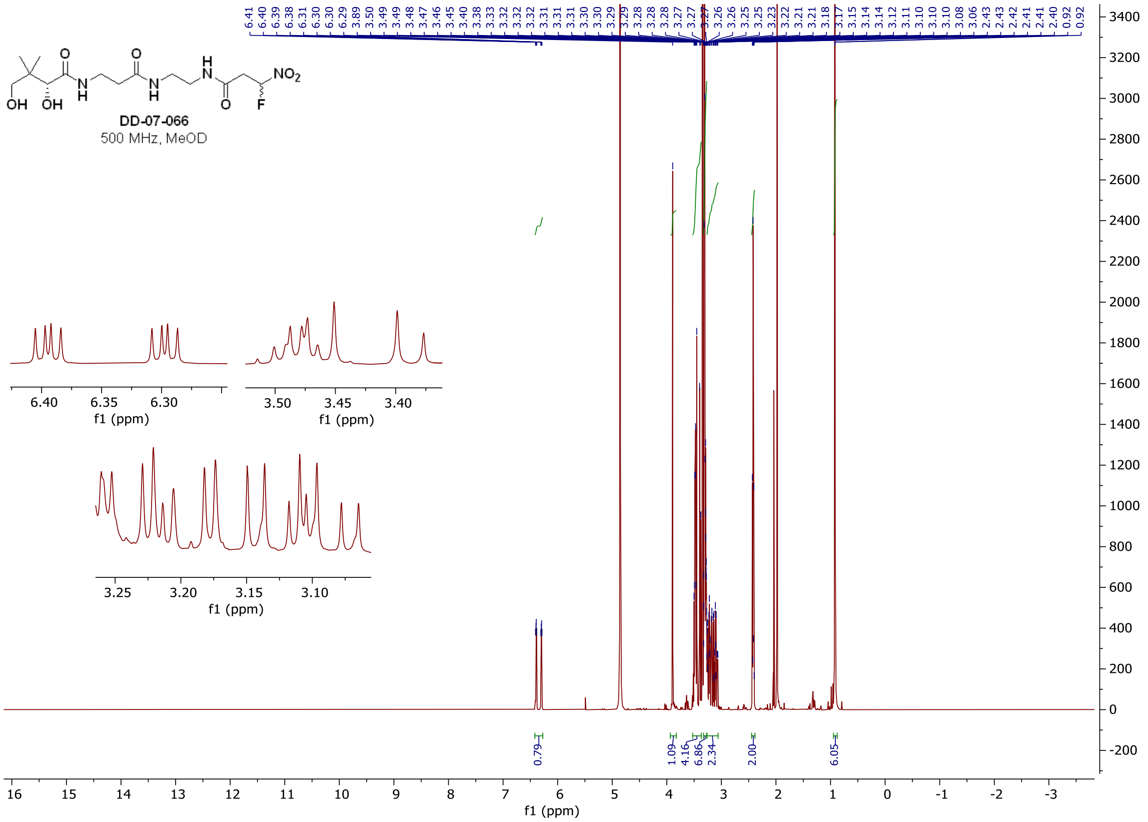

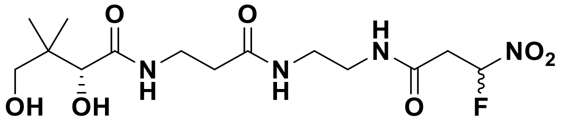

**DD-07-066**
471 MHz, MeOD

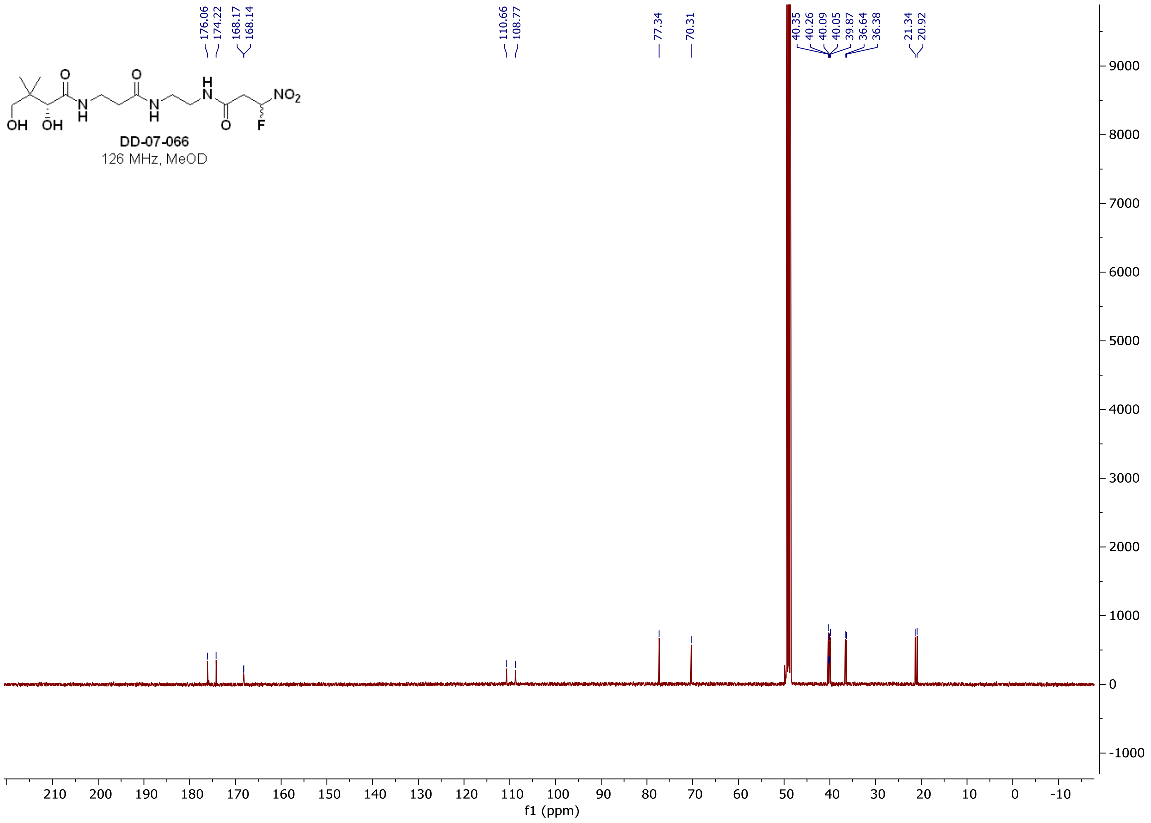
DD-07-066
126 MHz, MeOD
176.06
174.22
168.17
168.14
110.66
108.77
77.34
70.31
40.35
40.26
40.09
40.05
39.87
36.64
36.38
21.34
20.92
f1 (ppm)

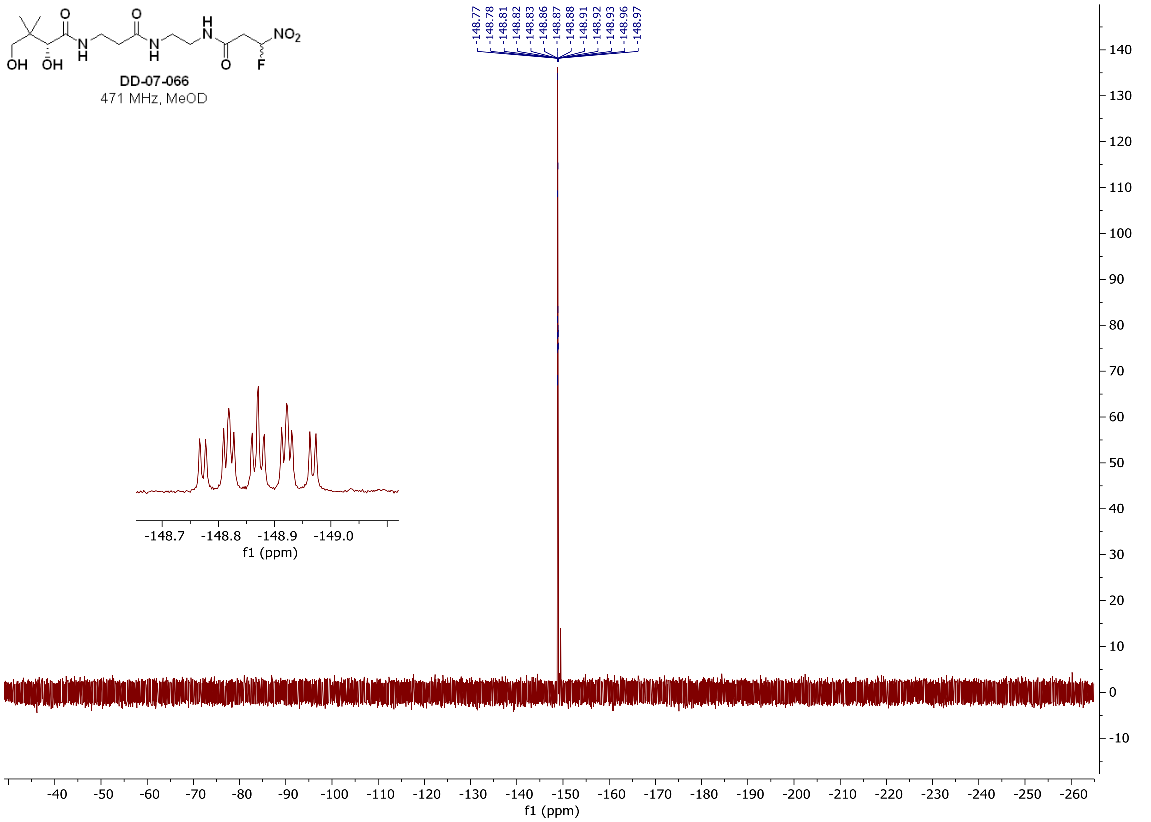
DD-07-066
471 MHz, MeOD
-148.77
-148.78
-148.81
-148.82
-148.83
-148.86
-148.87
-148.88
-148.91
-148.92
-148.93
-148.96
-148.97
f1 (ppm)

#### $^{1}H$ and $^{13}C$ spectra of **DD-02-088**

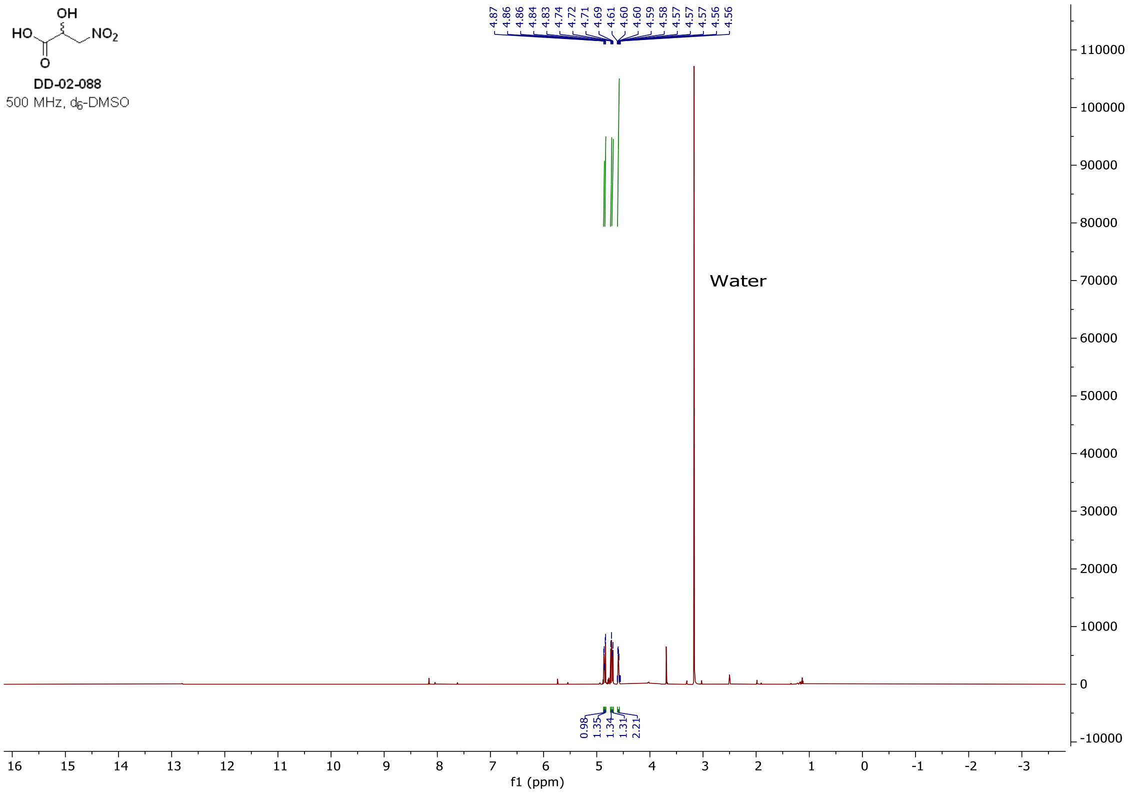

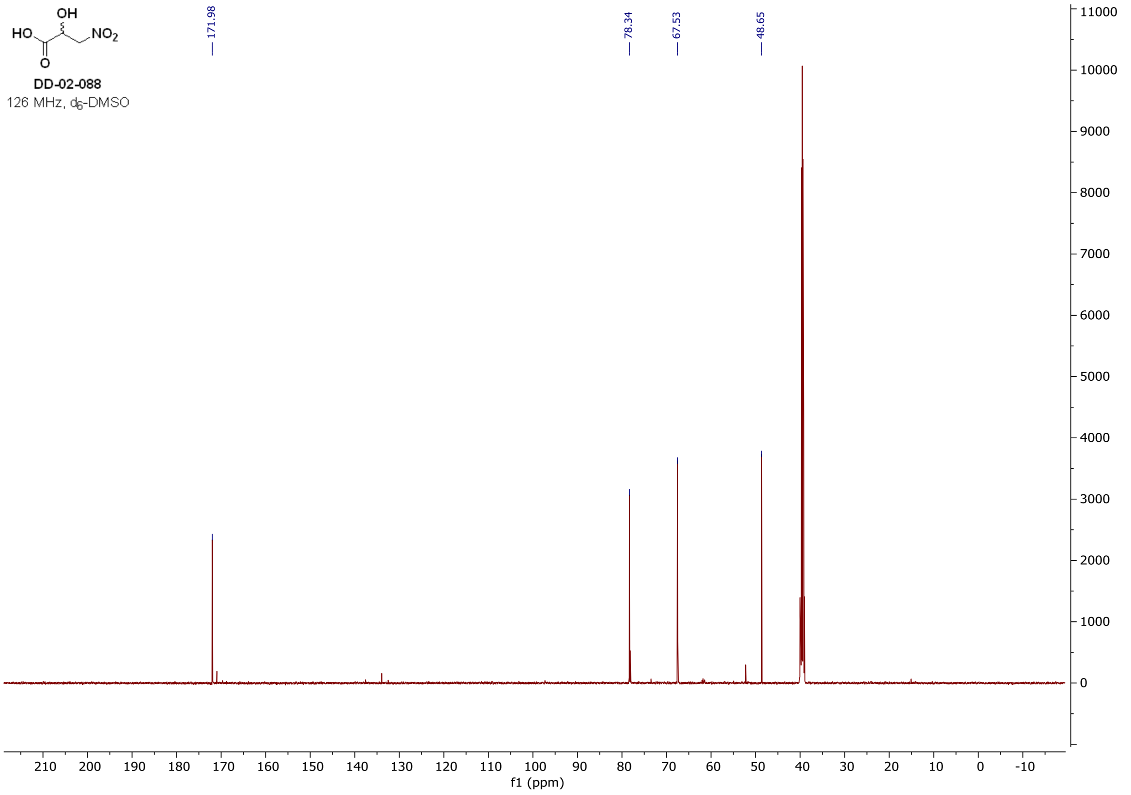

#### $^{1}H$ and $^{13}C$ spectra of **DD-03-038**

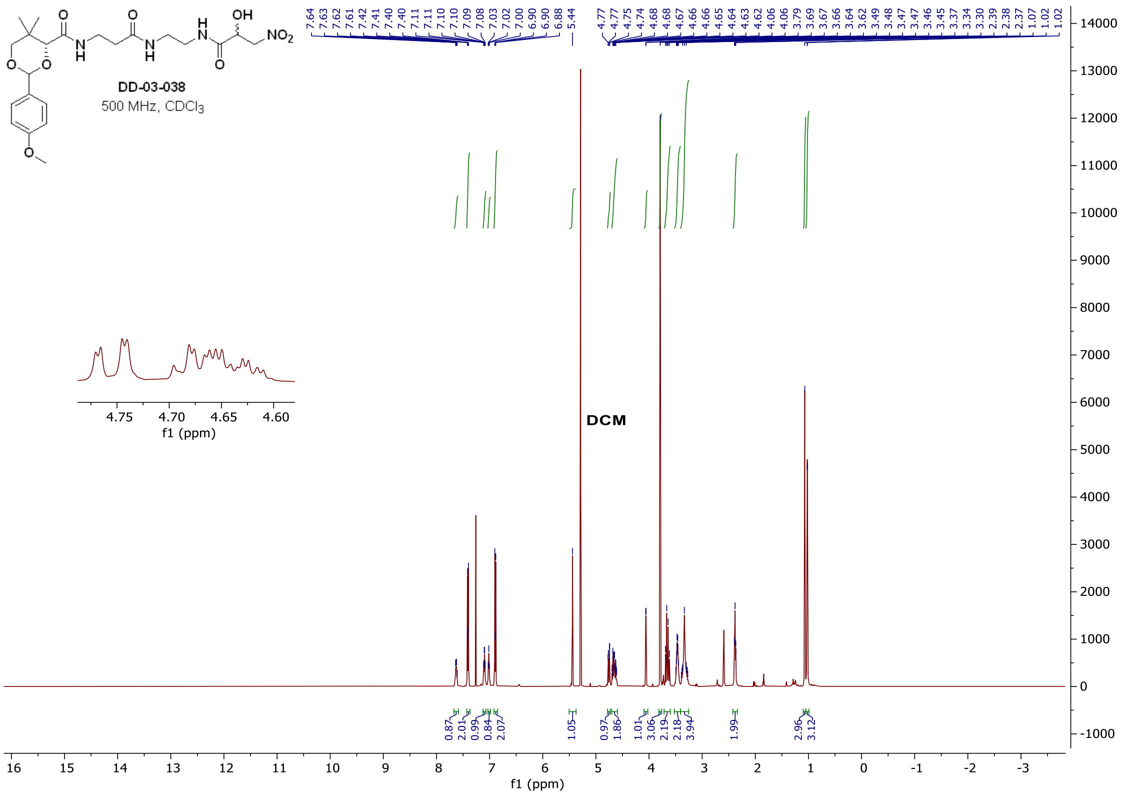

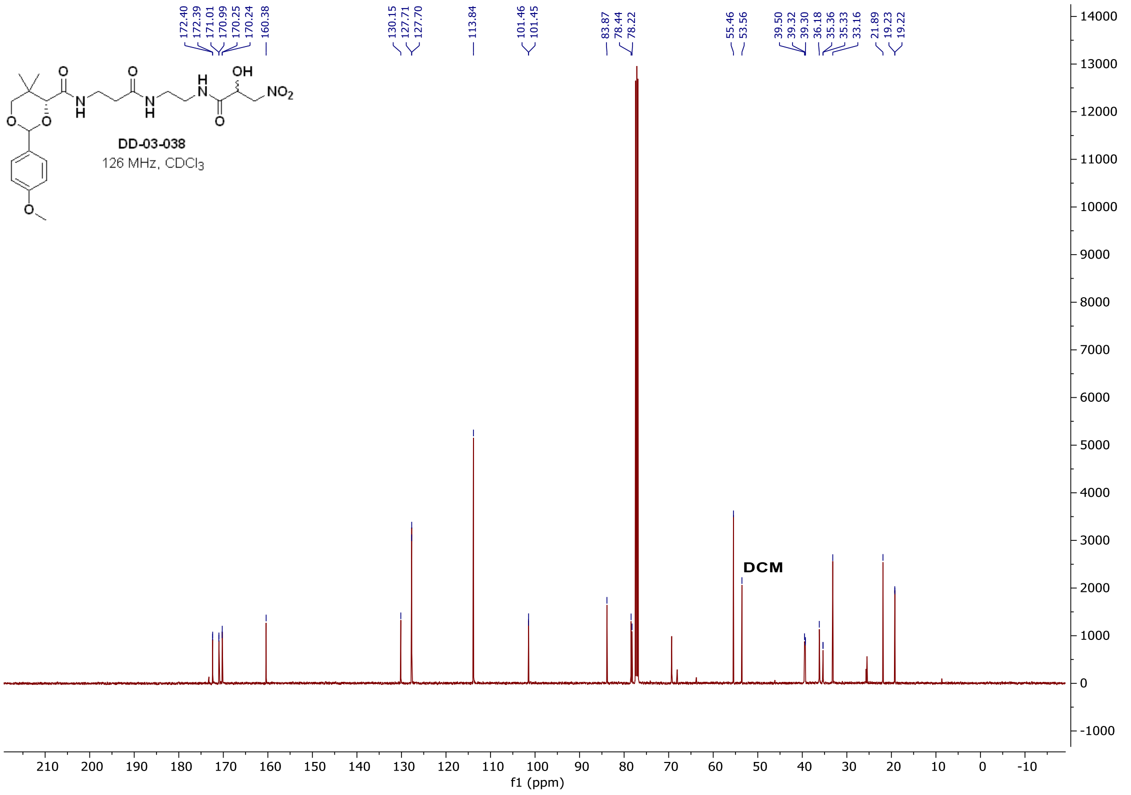

[1]H and [13]C spectra of **DD-04-021**

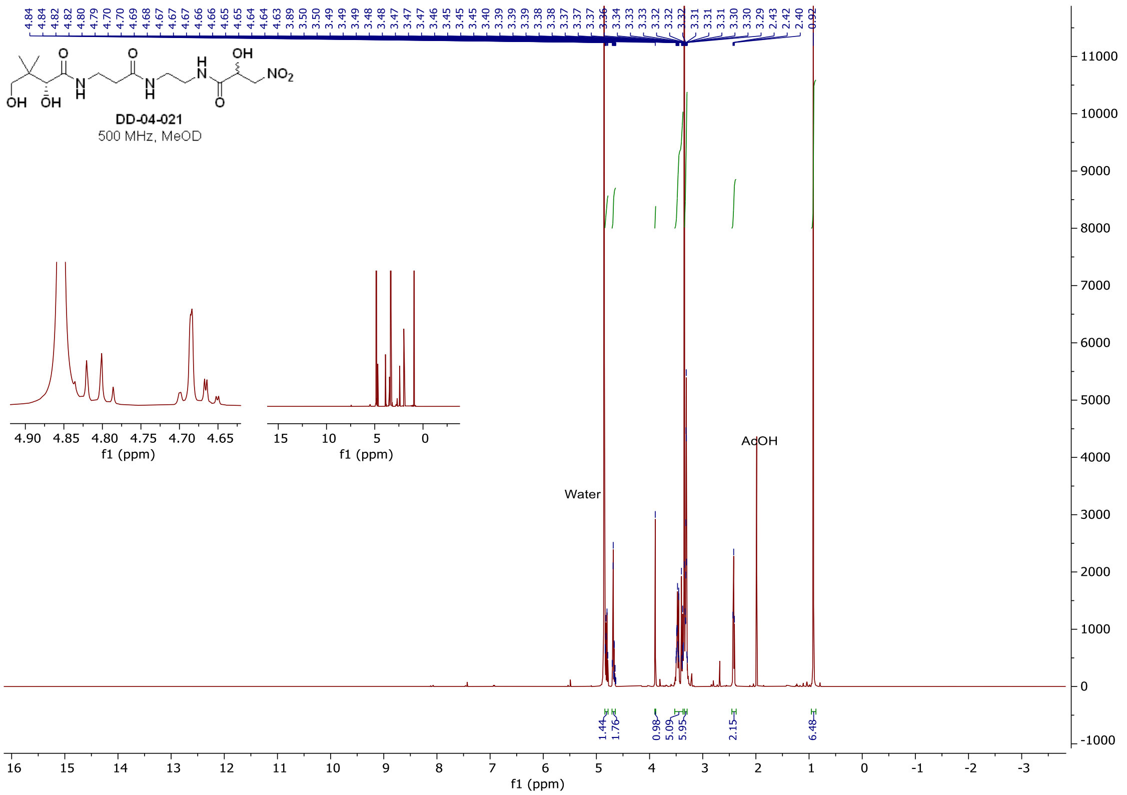

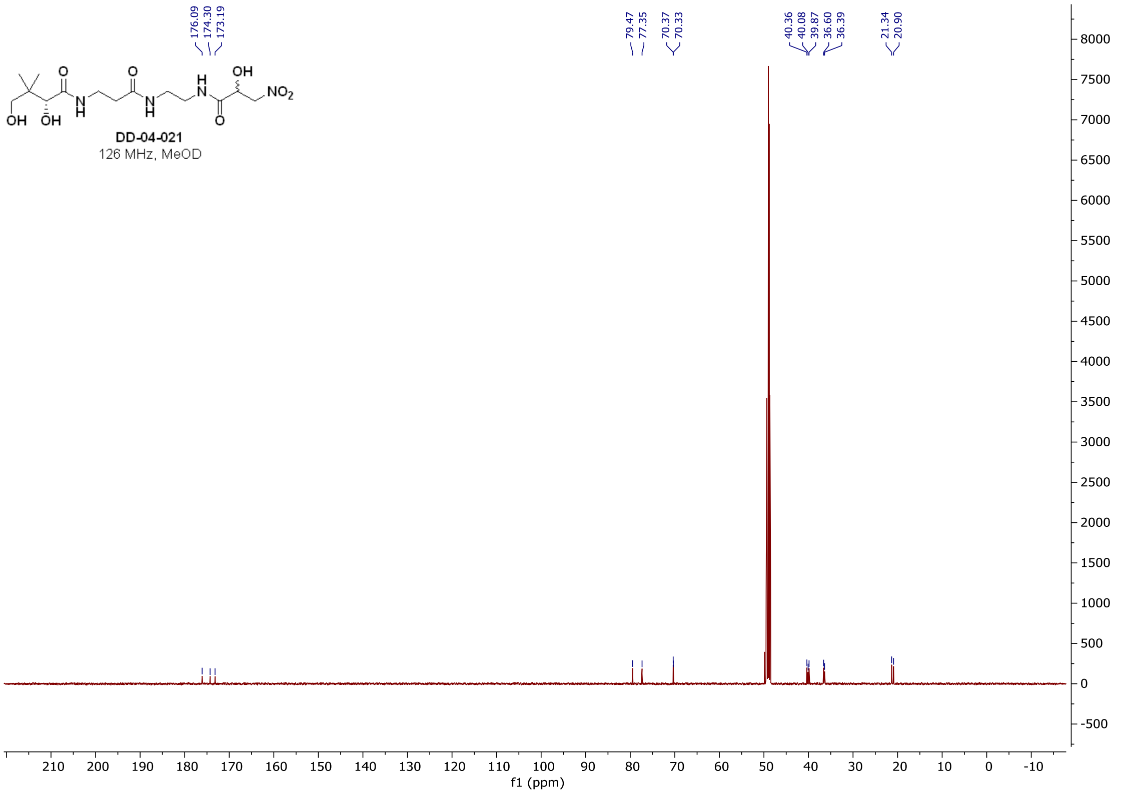

<sup></sup>

### [1]H and [13]C spectra of **DD-02-134**

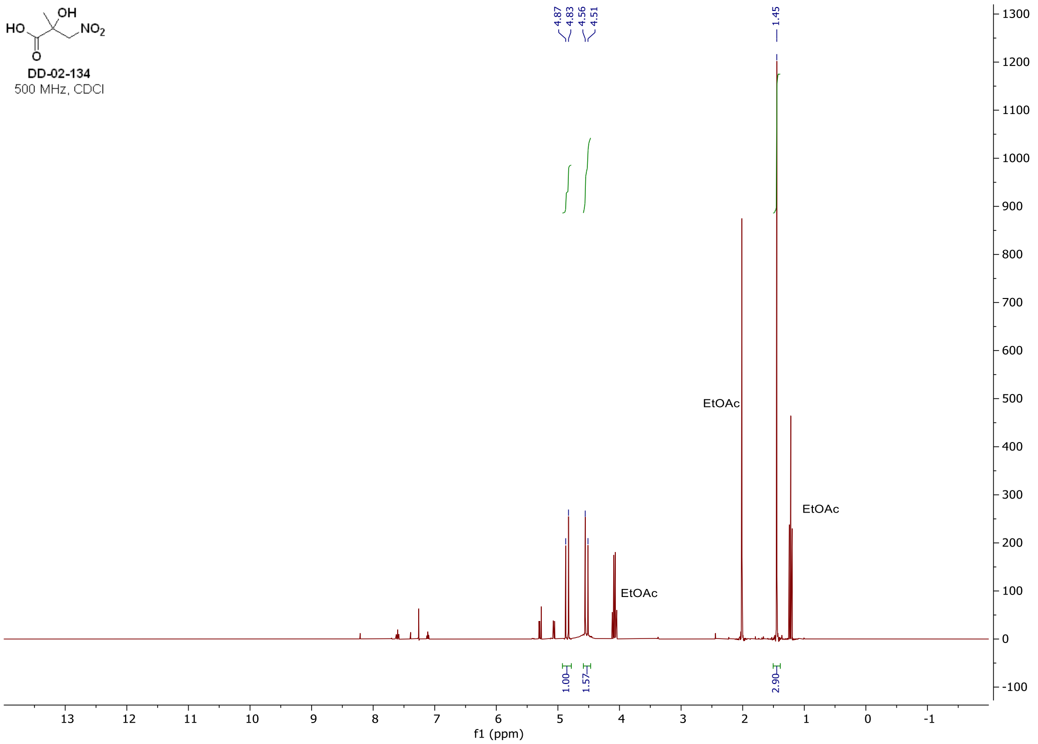

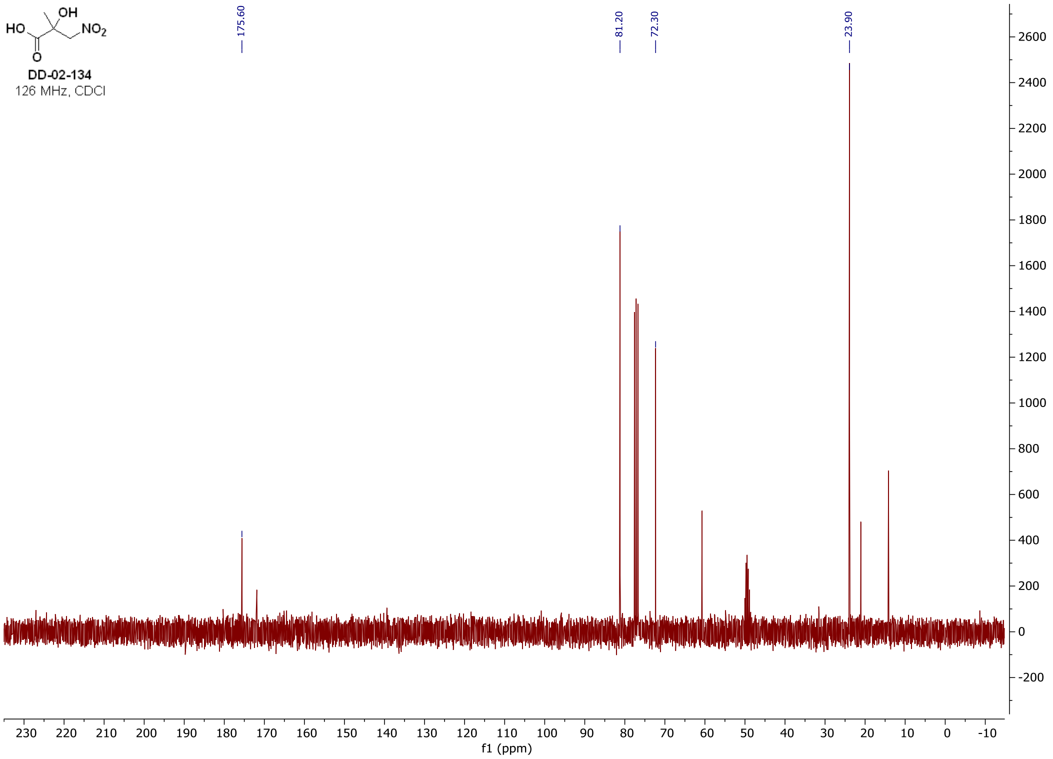

#### $^{1}H$ and $^{13}C$ spectra of **DD-02-143**

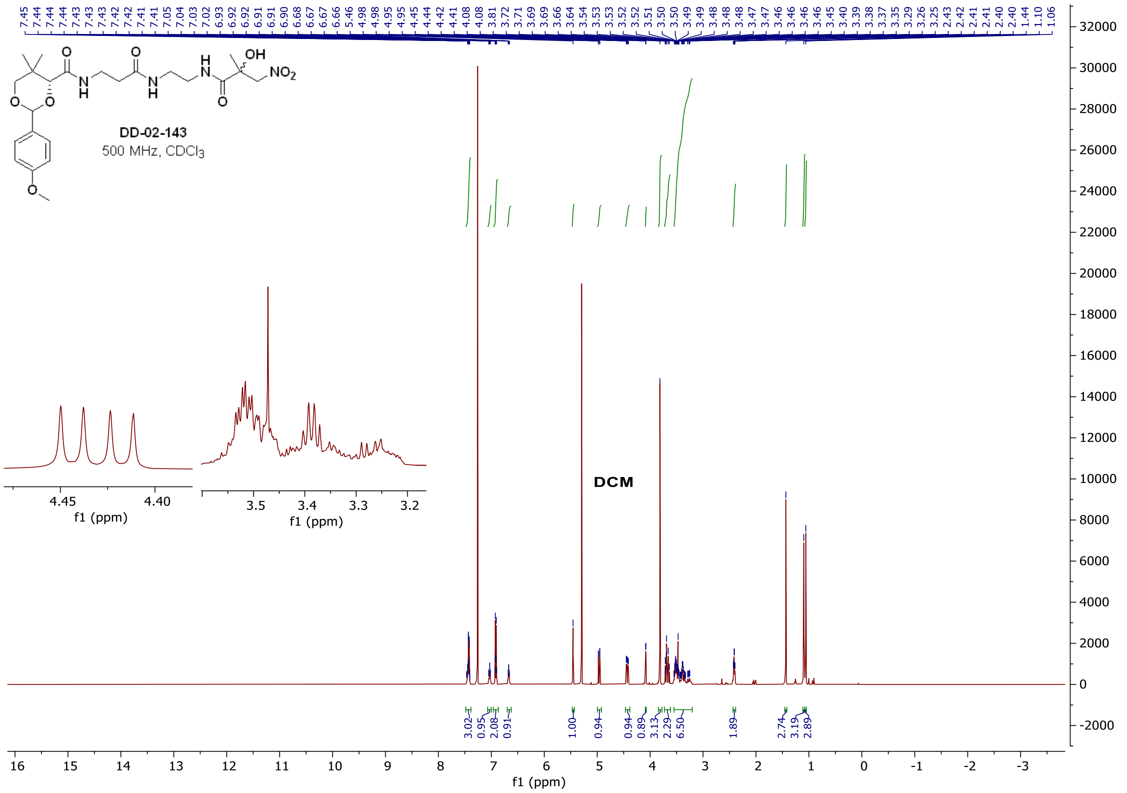

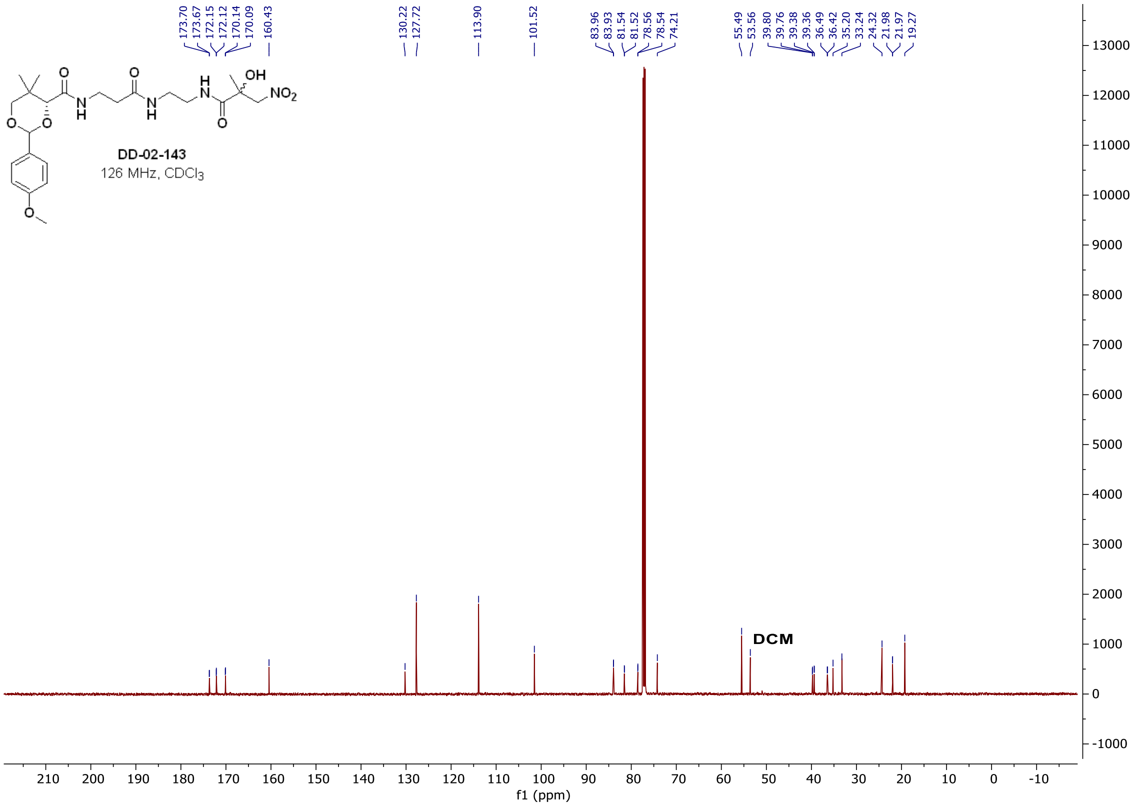

$^{1}H$ and $^{13}C$ spectra of **DD-03-001**

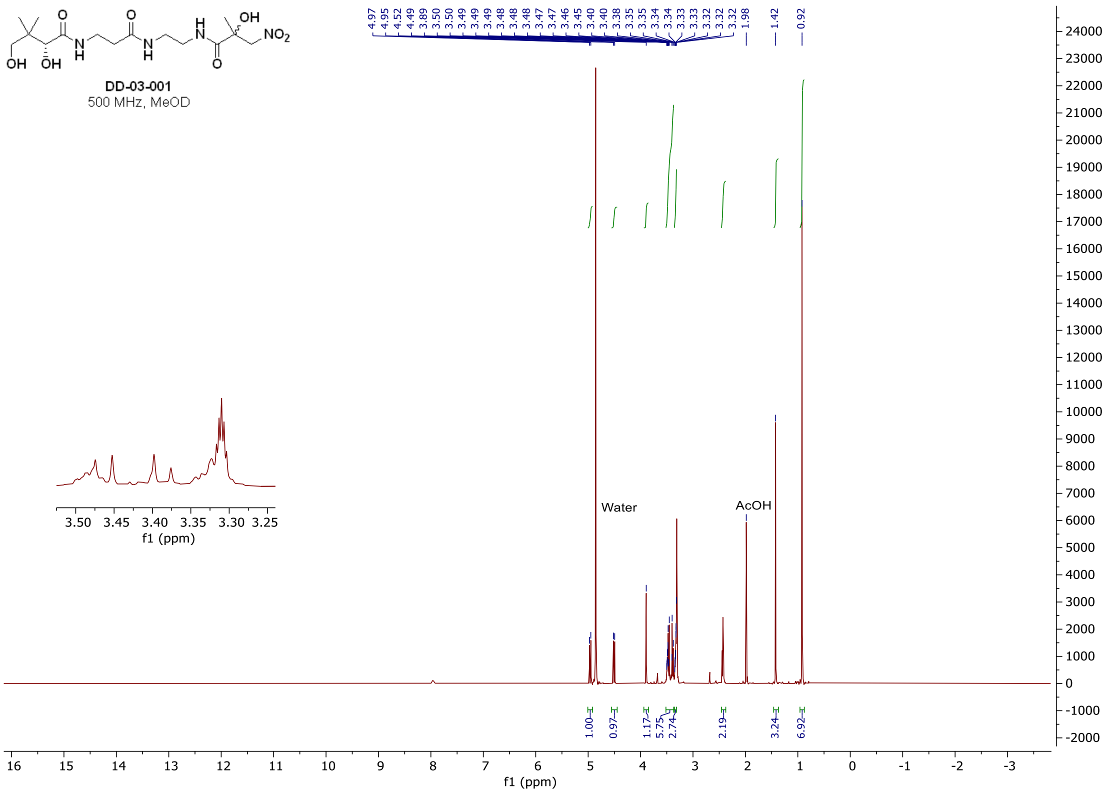

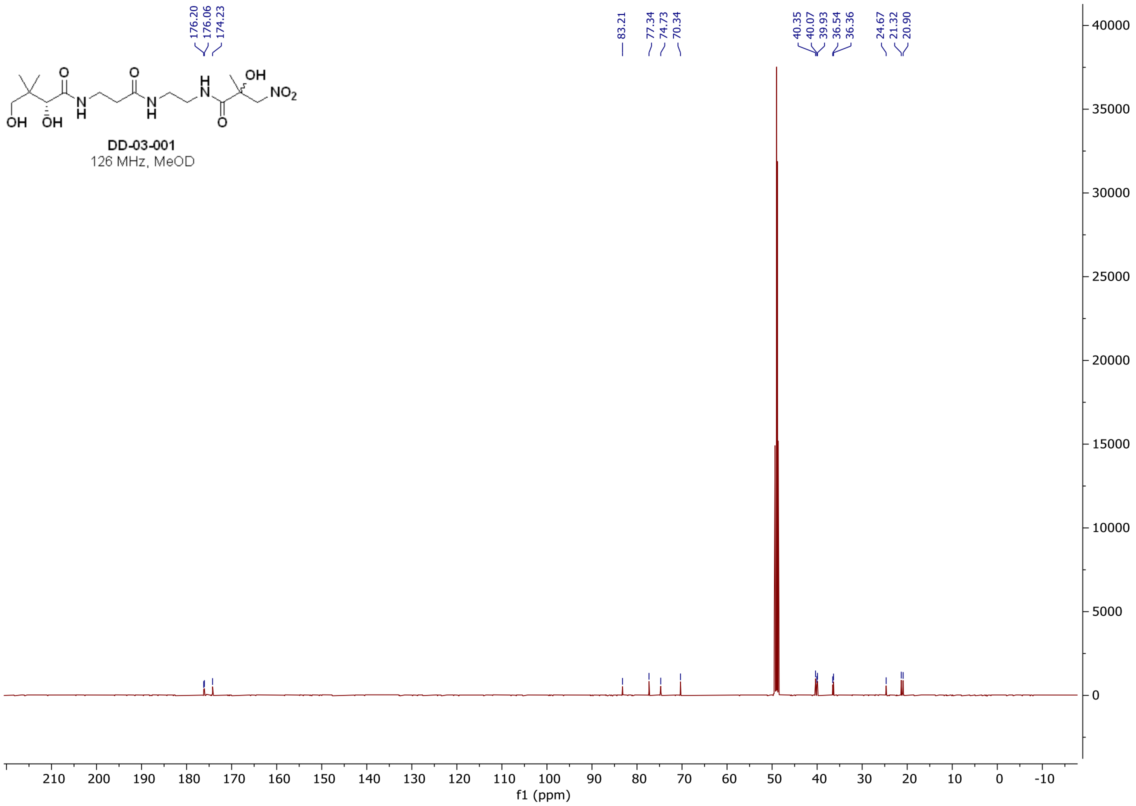

$^{1}H$ and $^{13}C$ spectra of **DD-02-105**

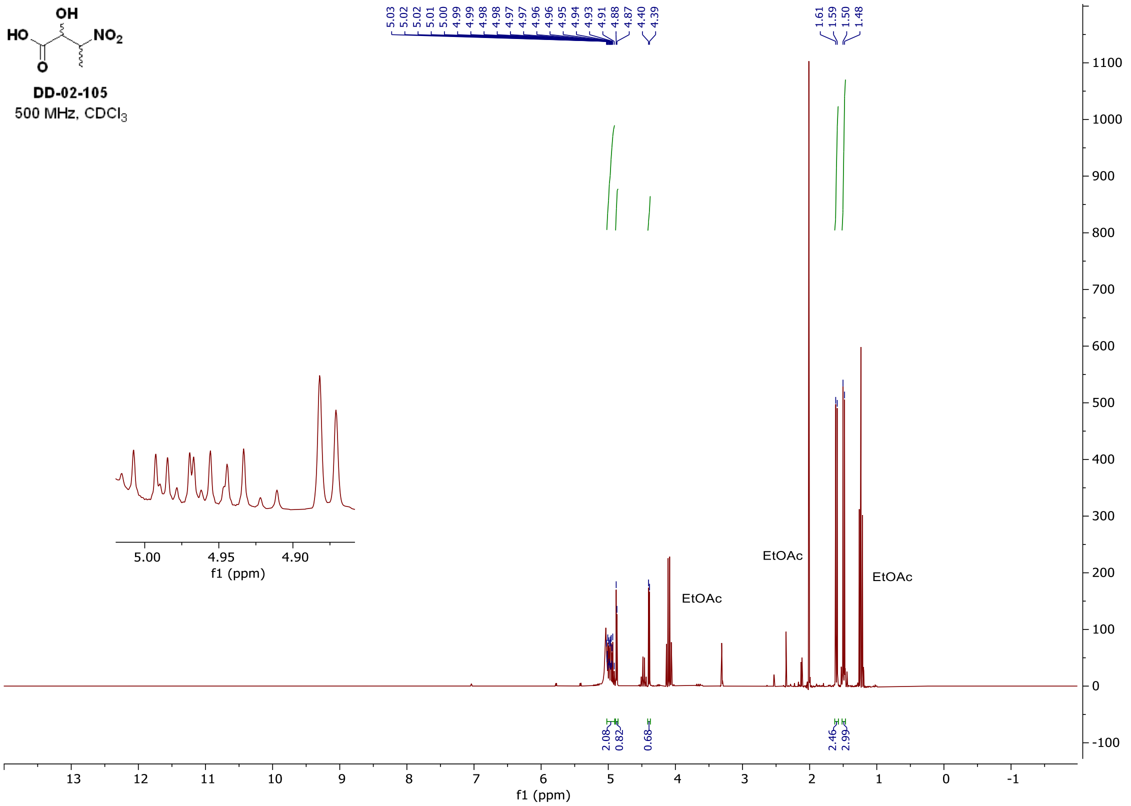

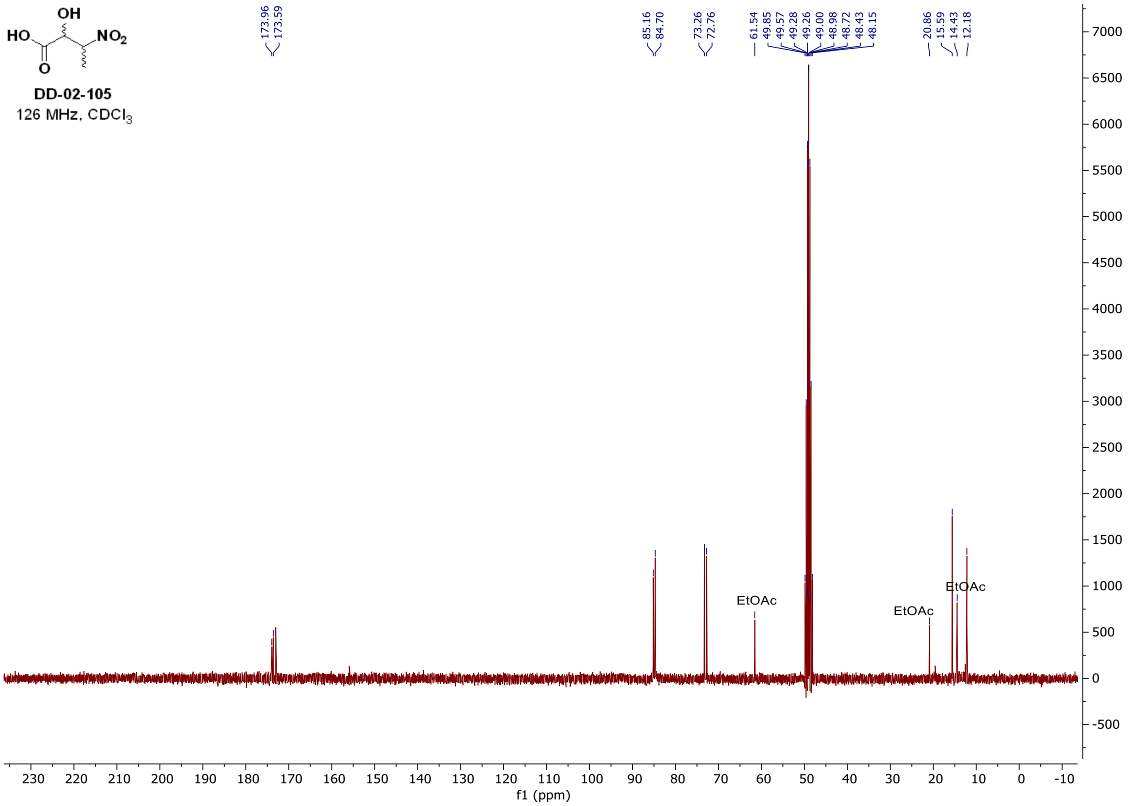

$^{1}H$ and $^{13}C$ **spectra of DD-03-081 (*R*,*R* and *S*,*S* or *R*,*S* and *S*,*R* diasteromeric pairs)**

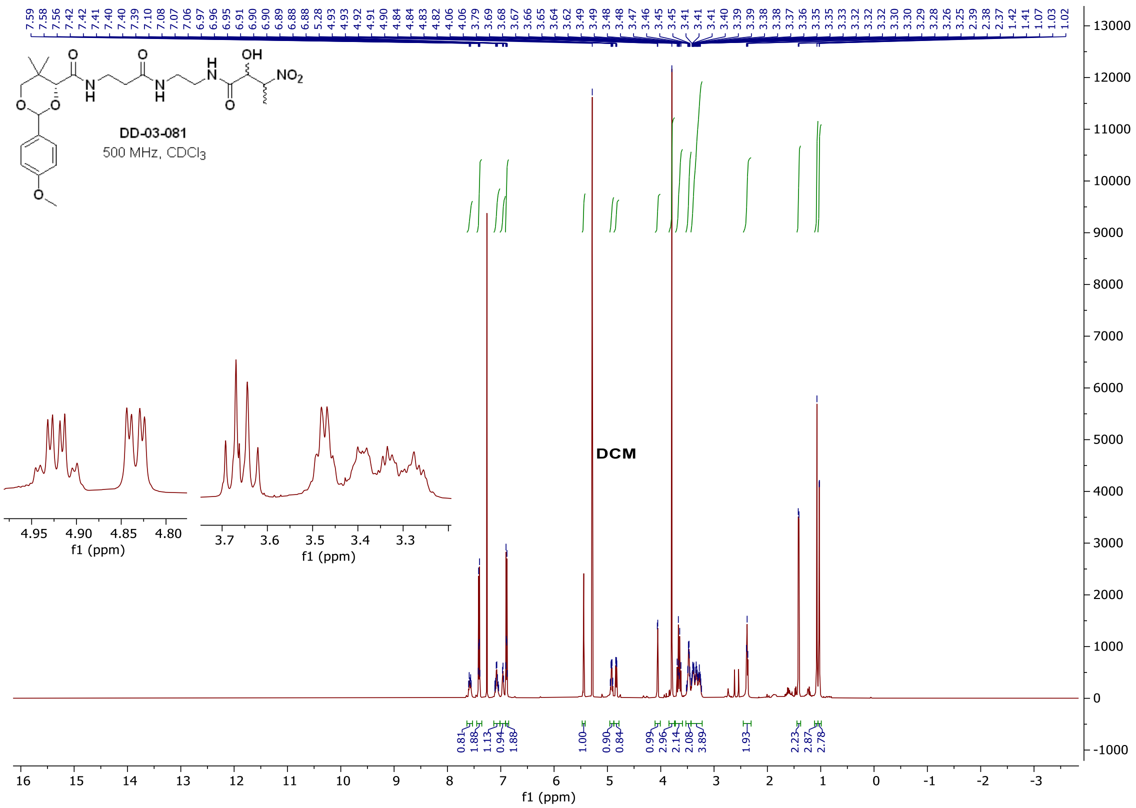

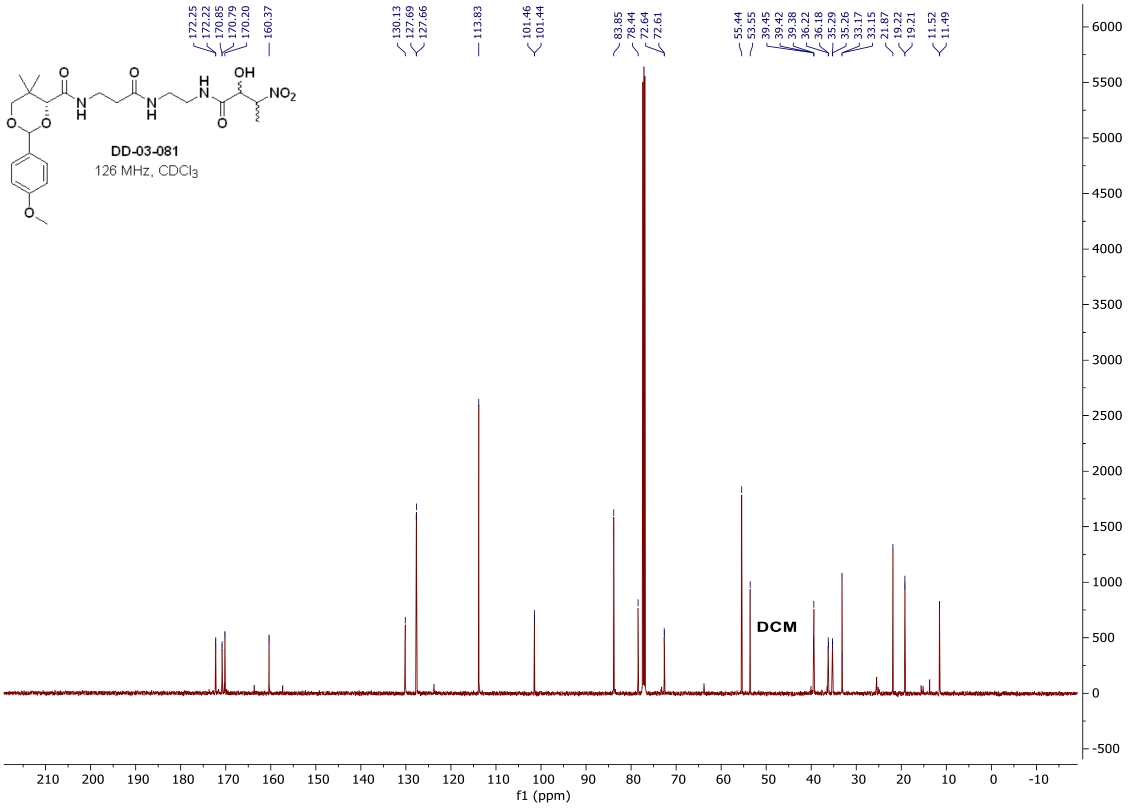

#### $^{1}H$ and $^{13}C$ **spectra of DD-03-081 (*R*,*R* and *S*,*S* or *R*,*S* and *S*,*R* diasteromeric pairs)**

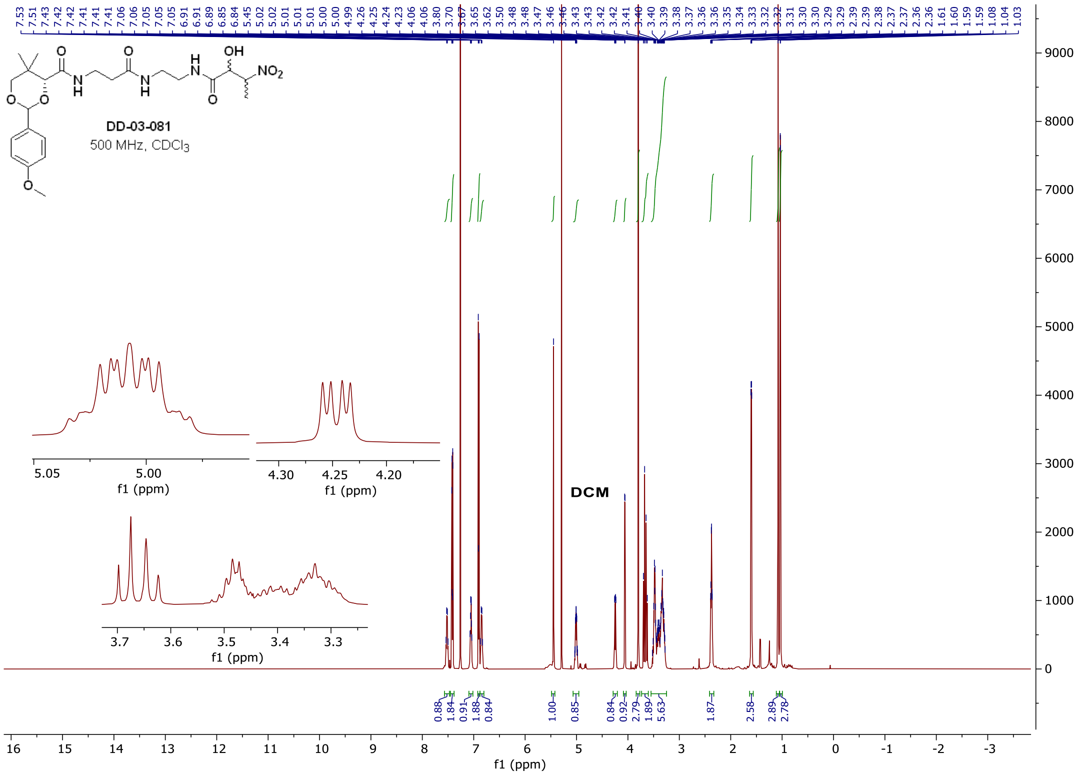

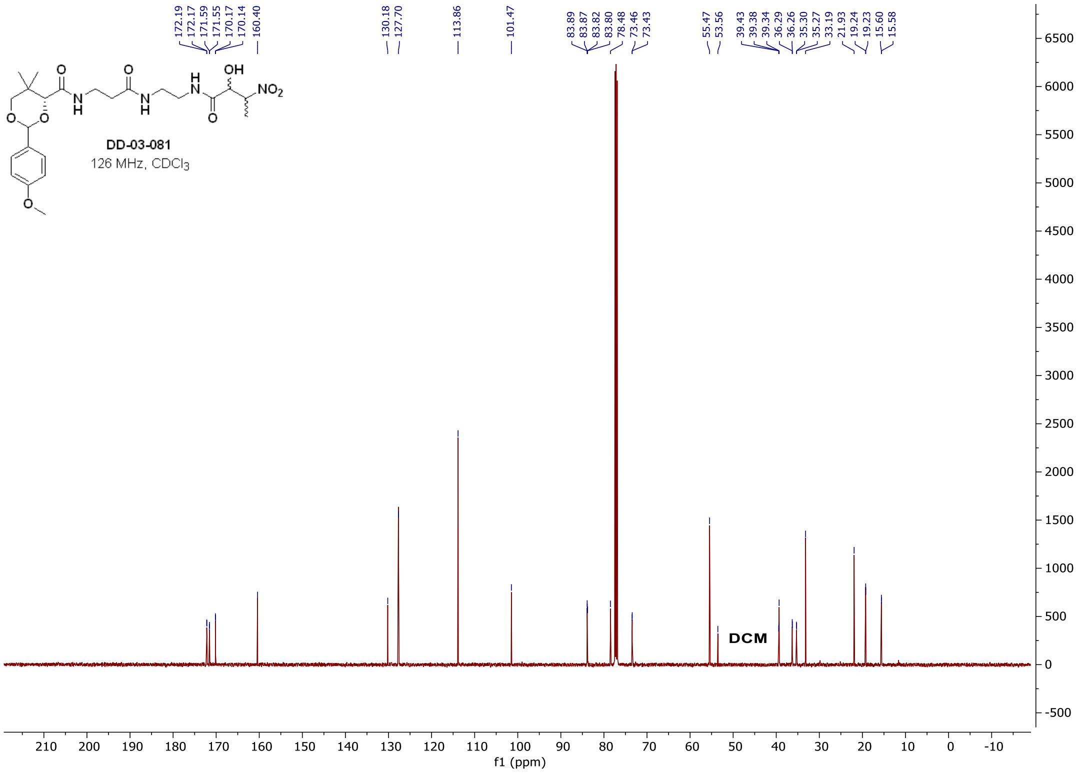

#### $^{1}H$ and $^{13}C$ **spectra of DD-03-085 (*R*,*R* and *S*,*S* or *R*,*S* and *S*,*R* diasteromeric pairs)**

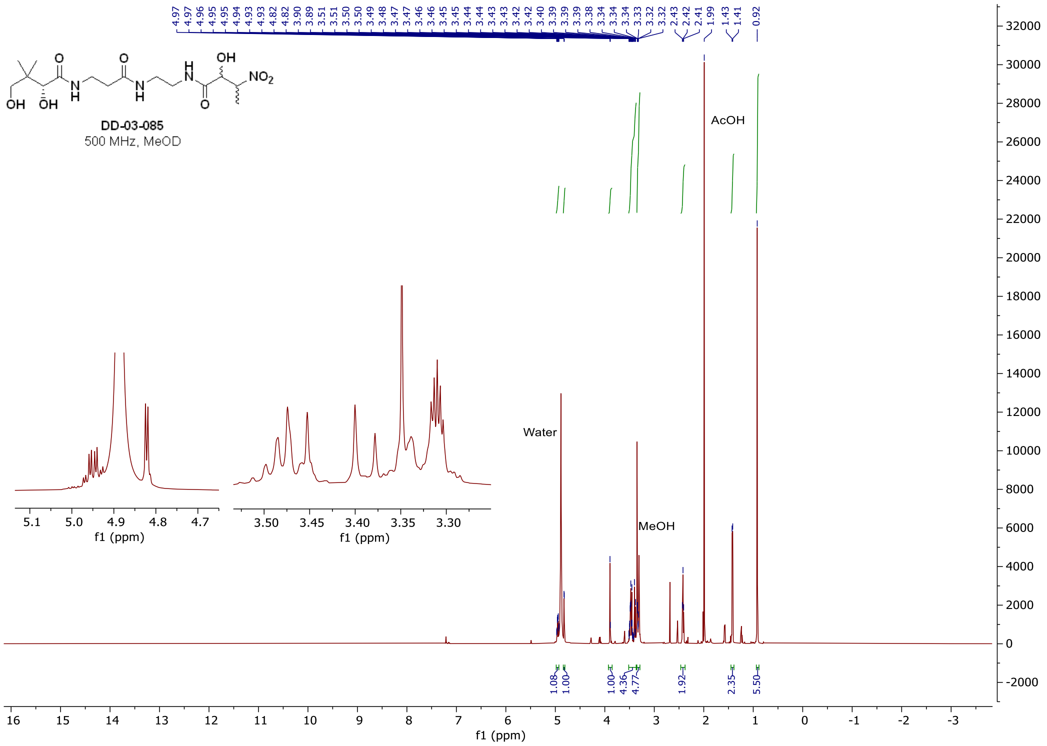

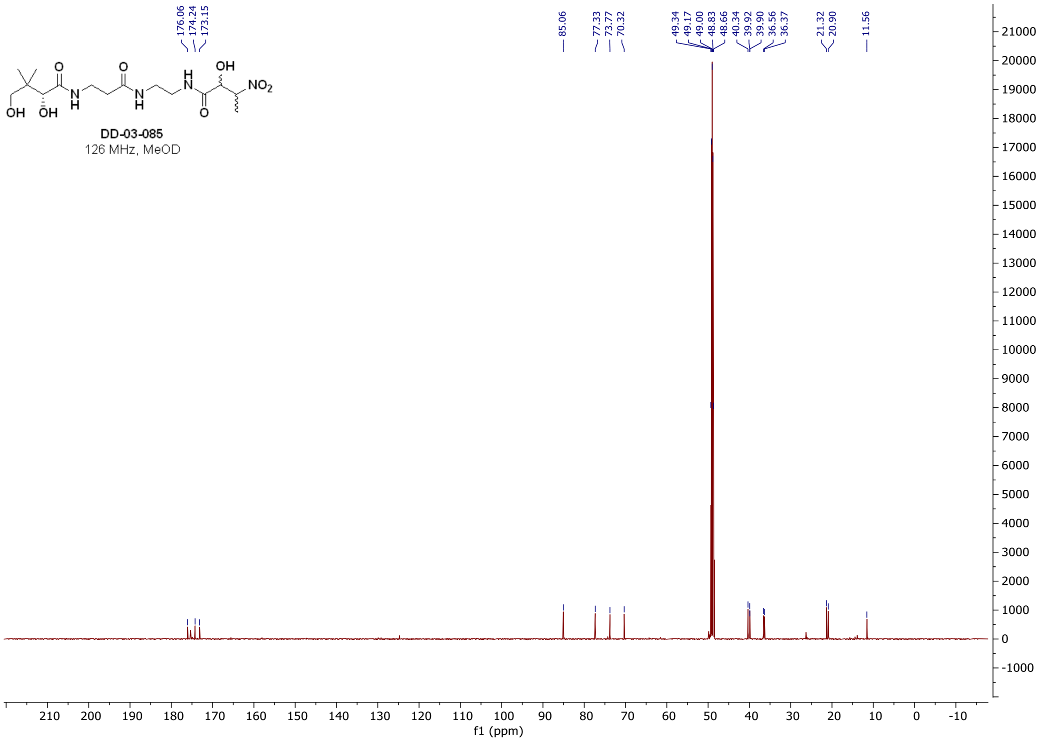

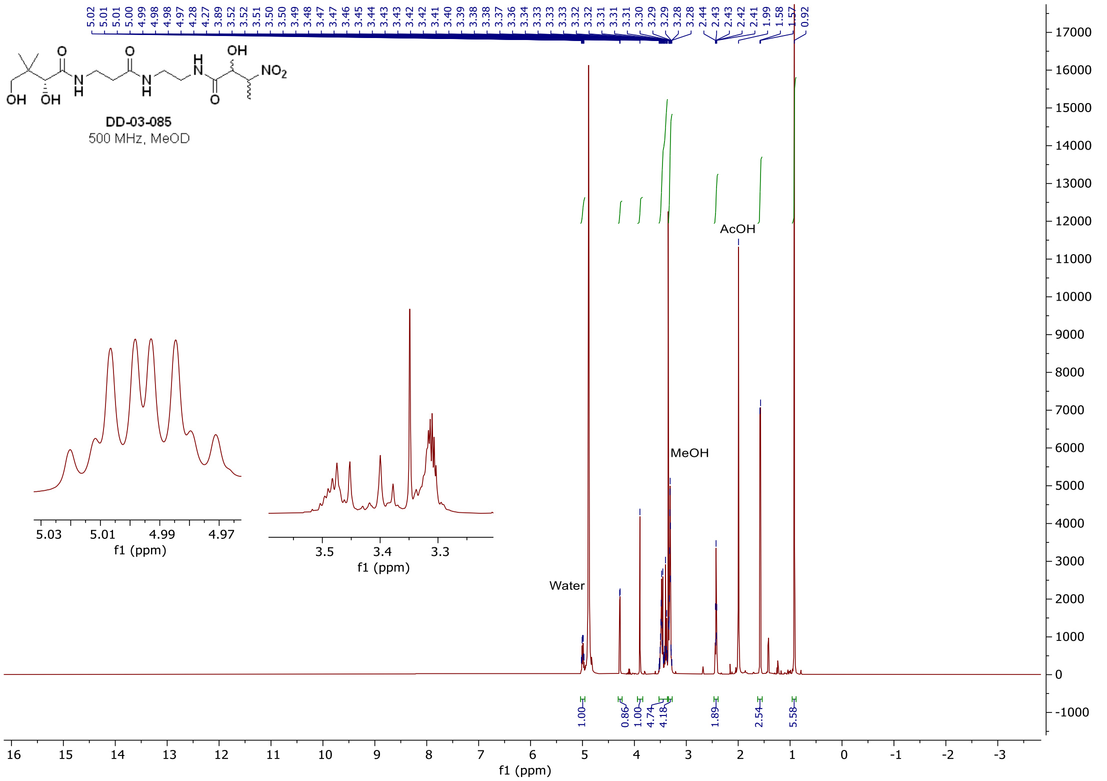
5.02 5.01 5.01 5.00 4.99 4.98 4.98 4.97 4.28 4.27 3.89 3.52 3.52 3.51 3.50 3.50 3.49 3.48 3.47 3.47 3.46 3.45 3.44 3.43 3.43 3.42 3.42 3.41 3.40 3.39 3.38 3.38 3.37 3.36 3.34 3.33 3.33 3.33 3.32 3.32 3.31 3.31 3.30 3.29 3.29 3.28 3.28 2.44 2.43 2.43 2.42 2.41 1.99 1.58 1.57 0.92
DD-03-085
500 MHz, MeOD
AcOH
MeOH
Water
1.00 0.86 1.00 4.74 4.18 1.89 2.54 5.58
f1 (ppm)

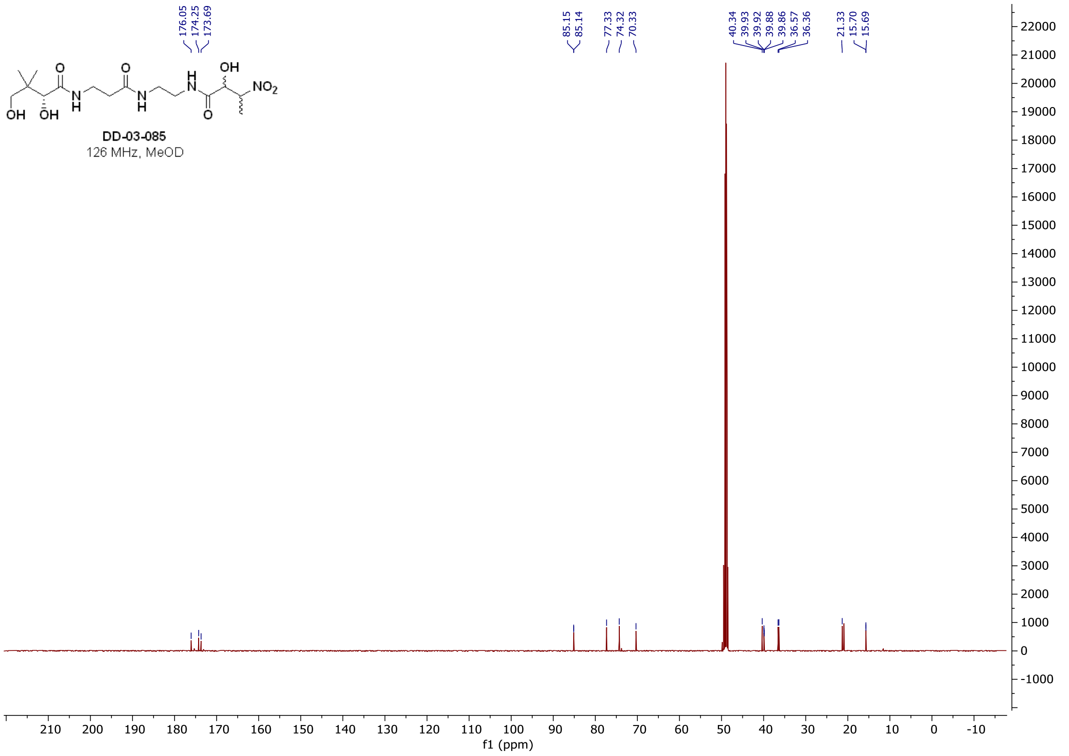
176.05 174.25 173.69
85.15 85.14 77.33 74.32 70.33
40.34 39.93 39.92 39.88 39.86 36.57 36.36
21.33 15.70 15.69
DD-03-085
126 MHz, MeOD
f1 (ppm)

#### $^{1}$H, $^{13}$C, and $^{19}$F **spectra of DD-03-098**

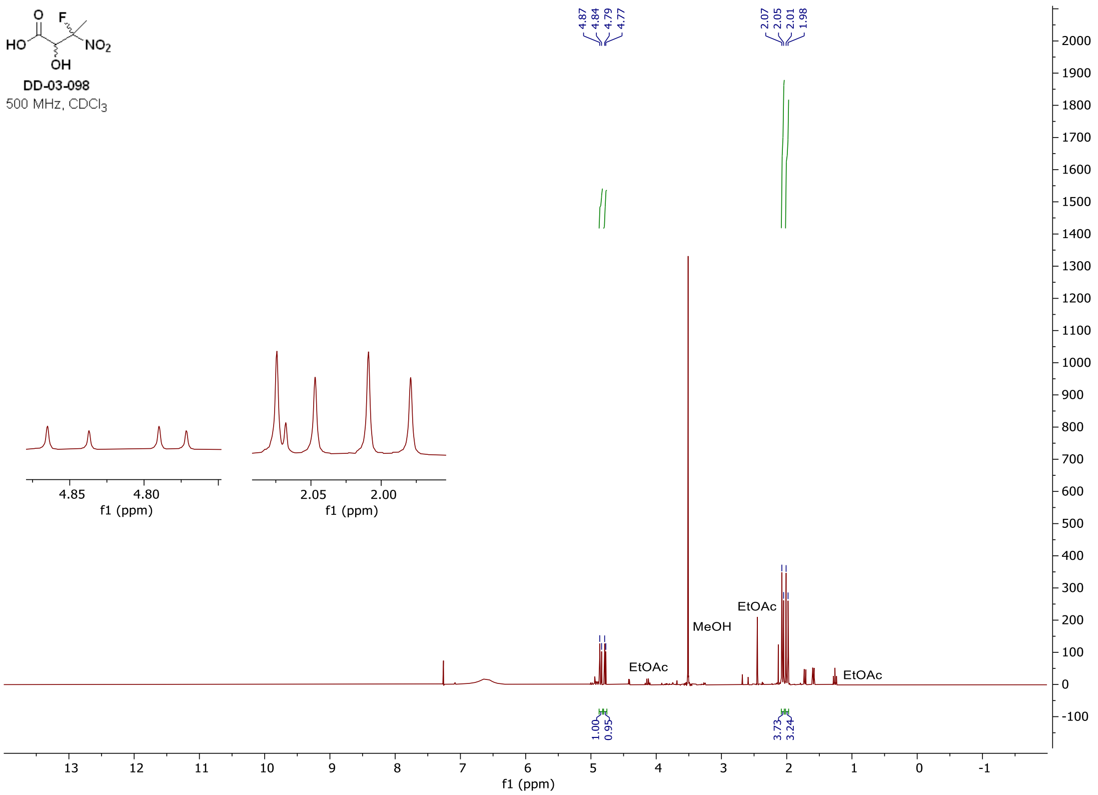

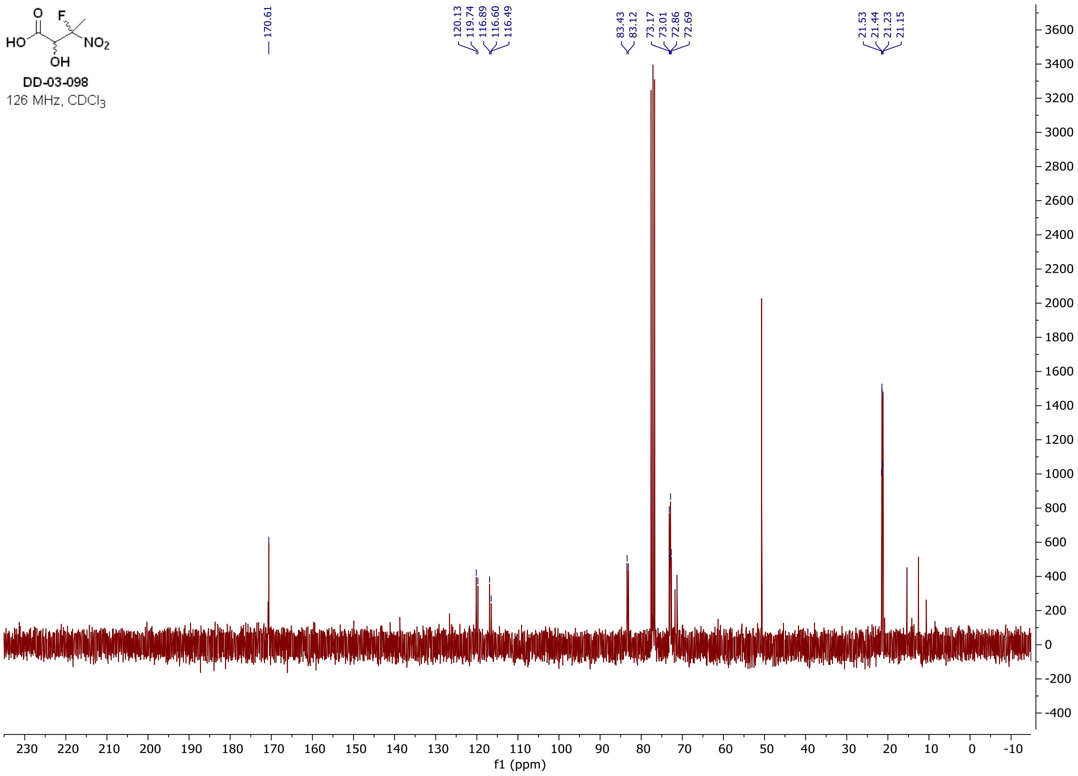

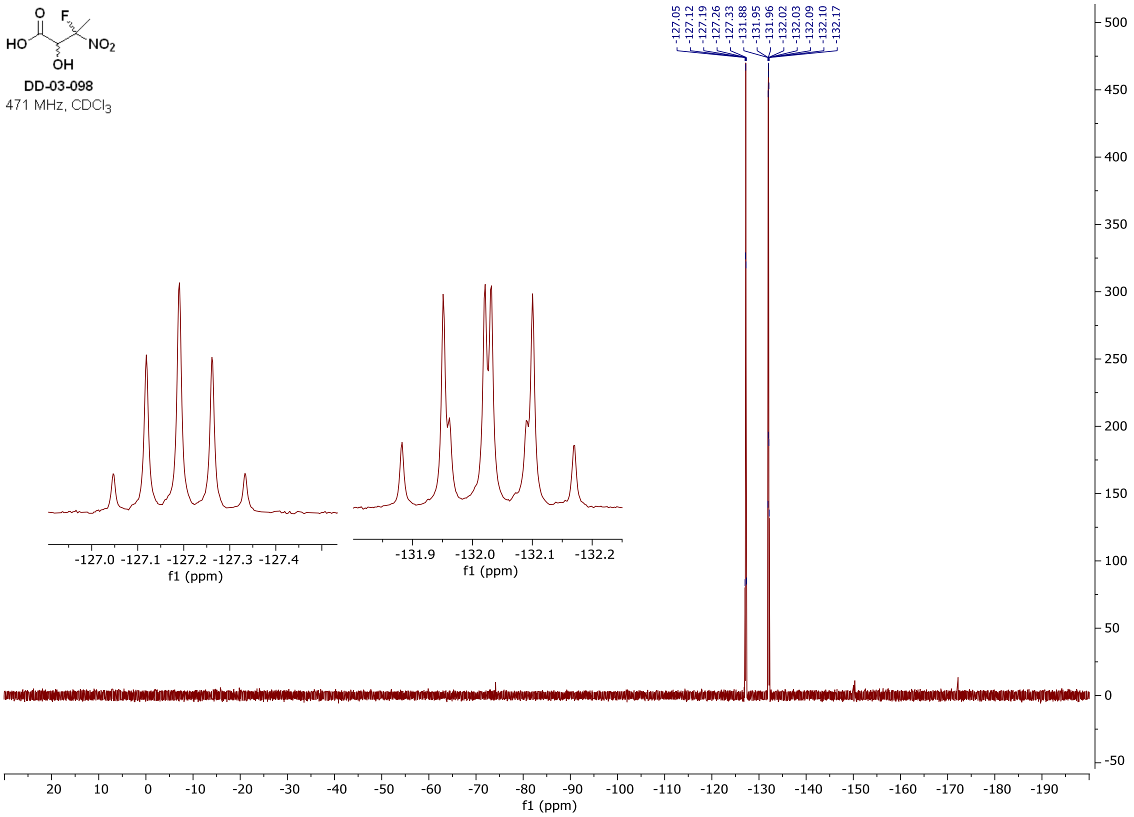

#### [1]H, [13]C, and [19]F **spectra of DD-03-100 (*S*,*R* and *R*,*S* diastereomeric pair)**

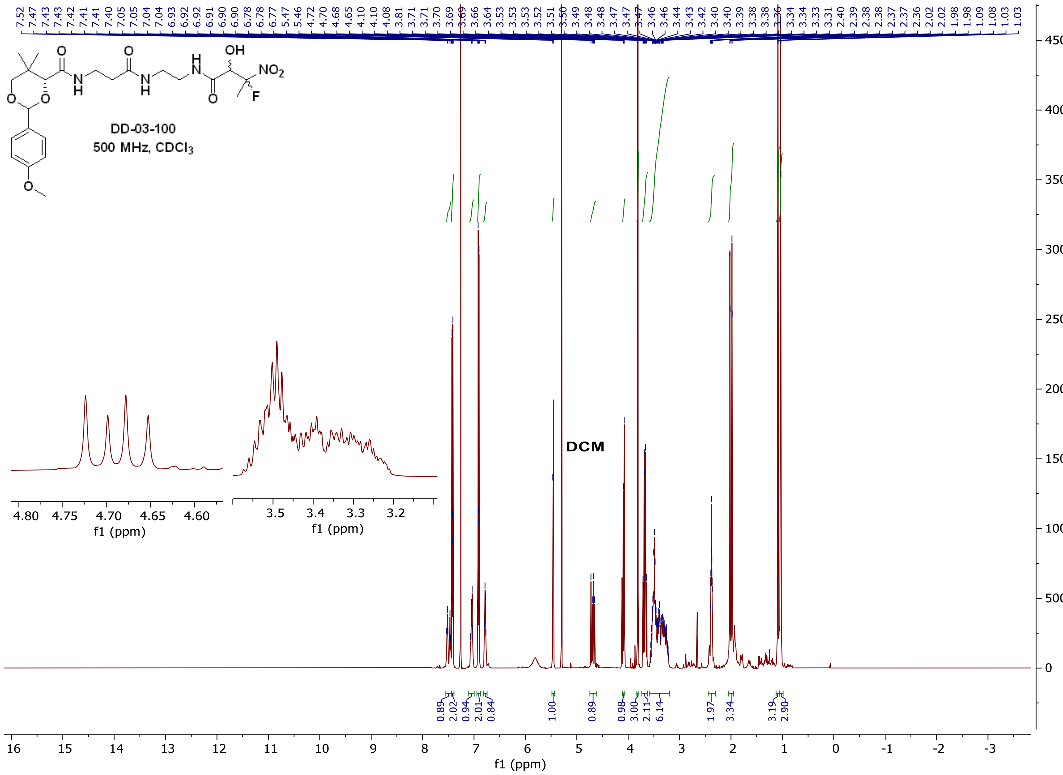

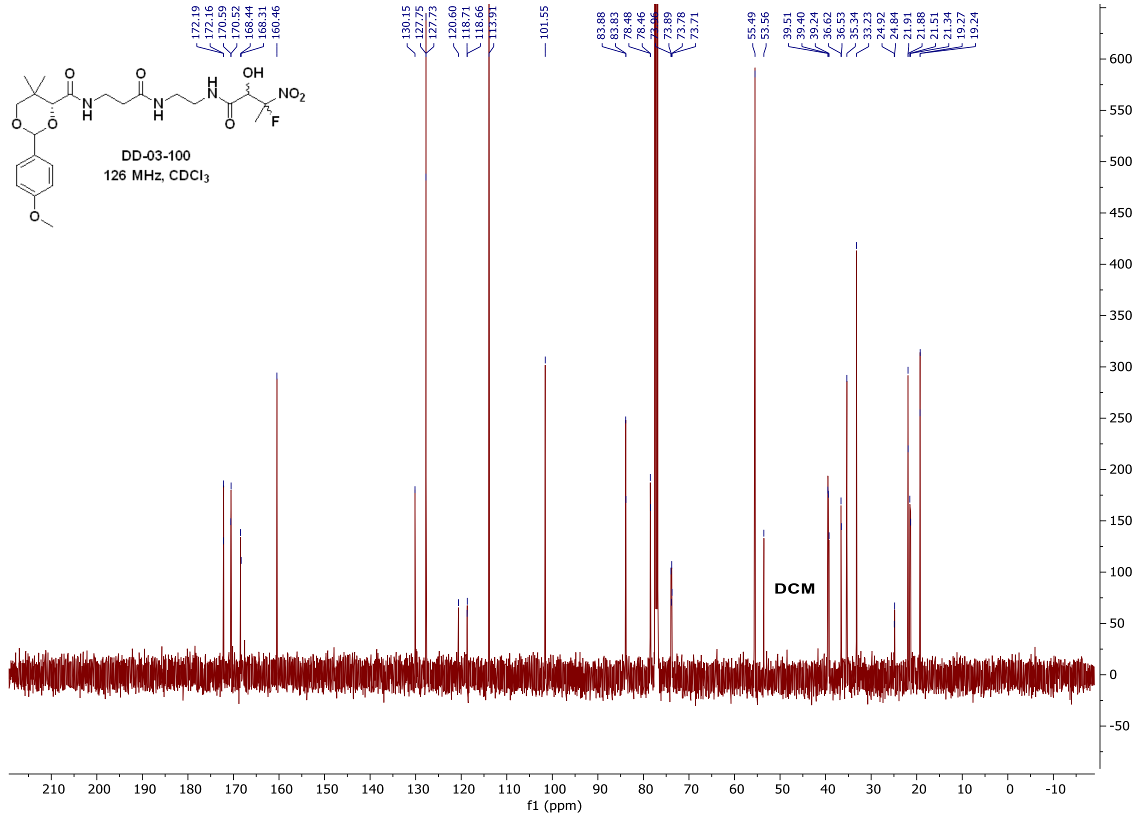
DD-03-100
126 MHz, CDCl3
DCM
f1 (ppm)

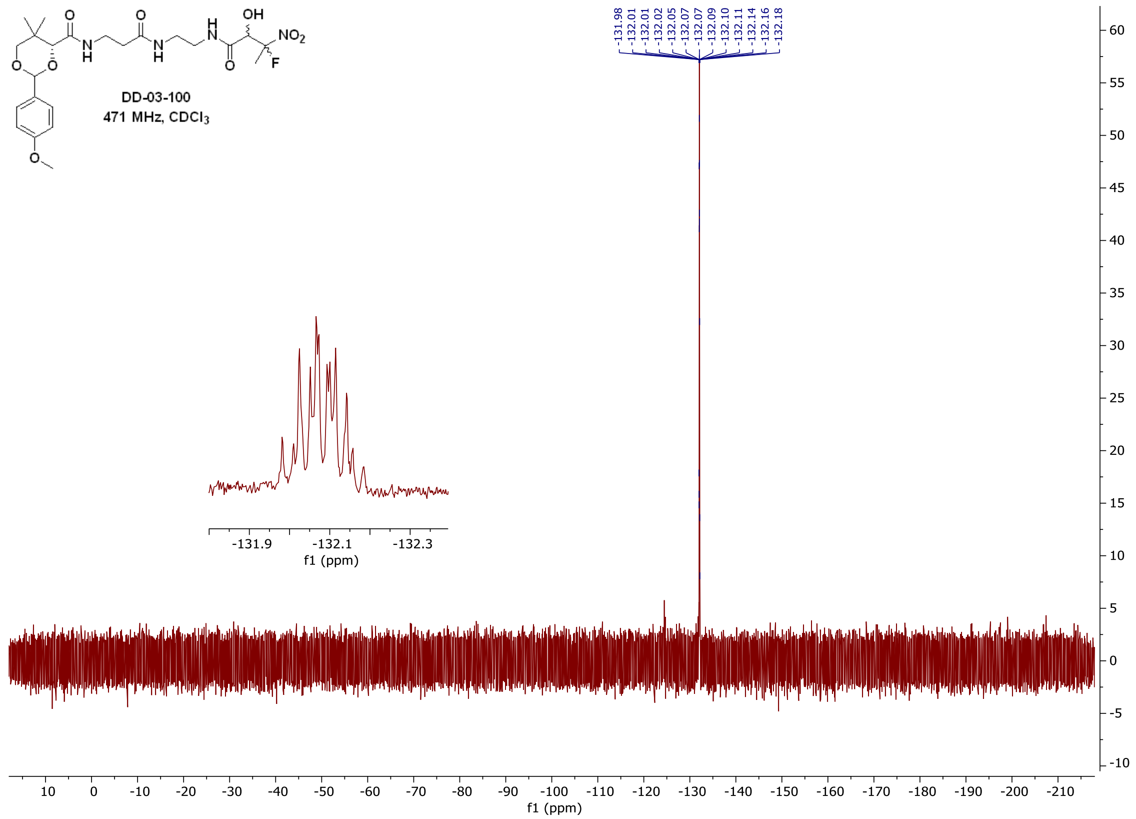
DD-03-100
471 MHz, CDCl3
f1 (ppm)

$^{1}$H, $^{13}$C, and $^{19}$F **spectra of DD-03-100 (*S,S* and *R,R* diastereomeric pair)**

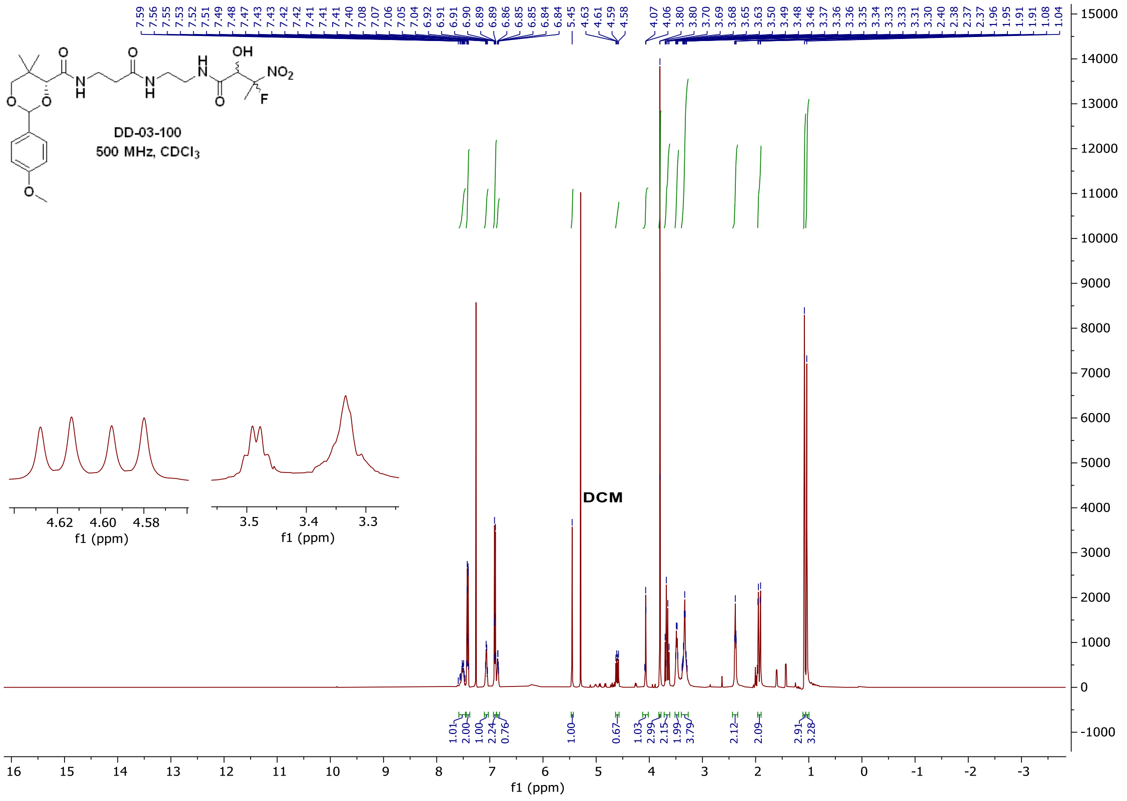

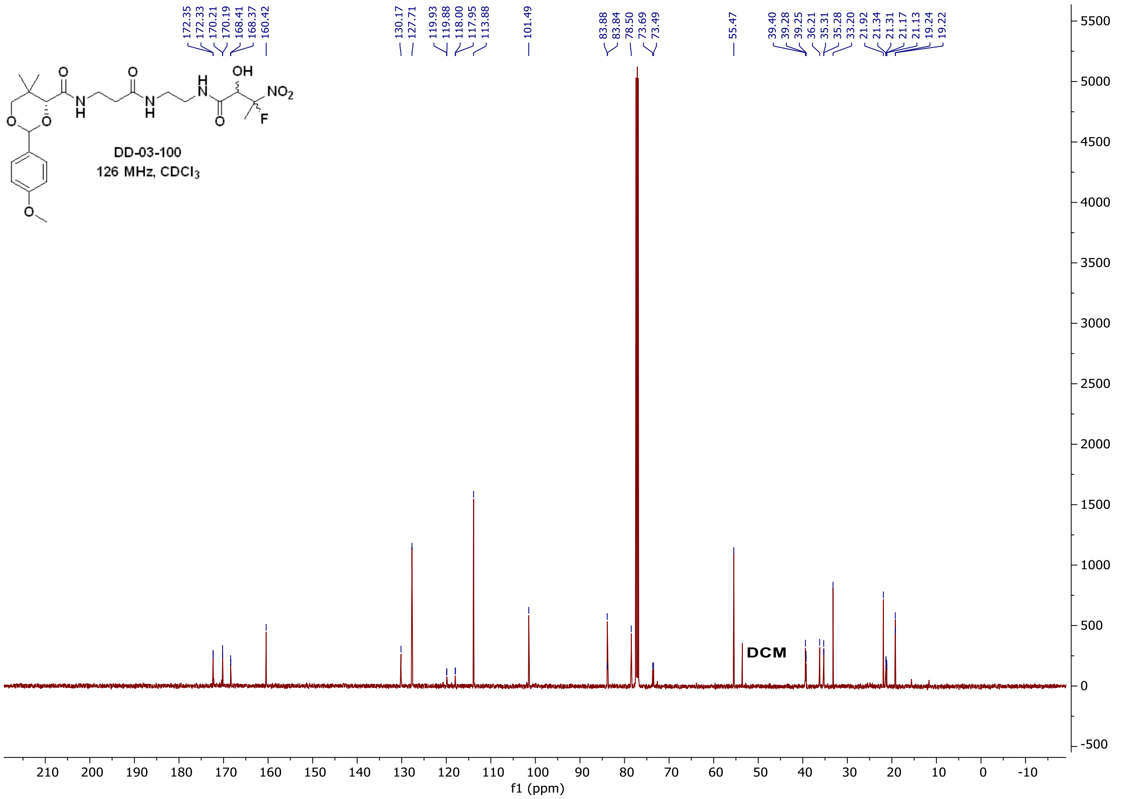

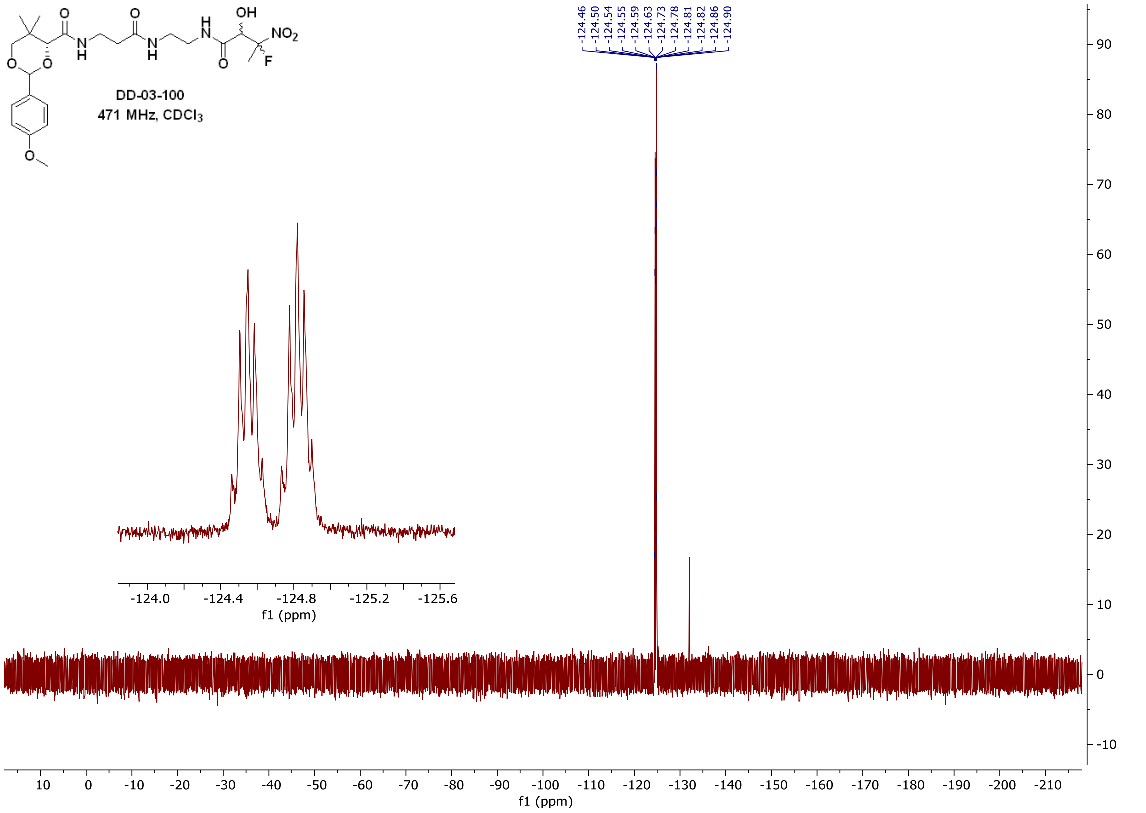

### $^{1}H$, $^{13}C$, and $^{19}F$ **spectra of DD-04-011**

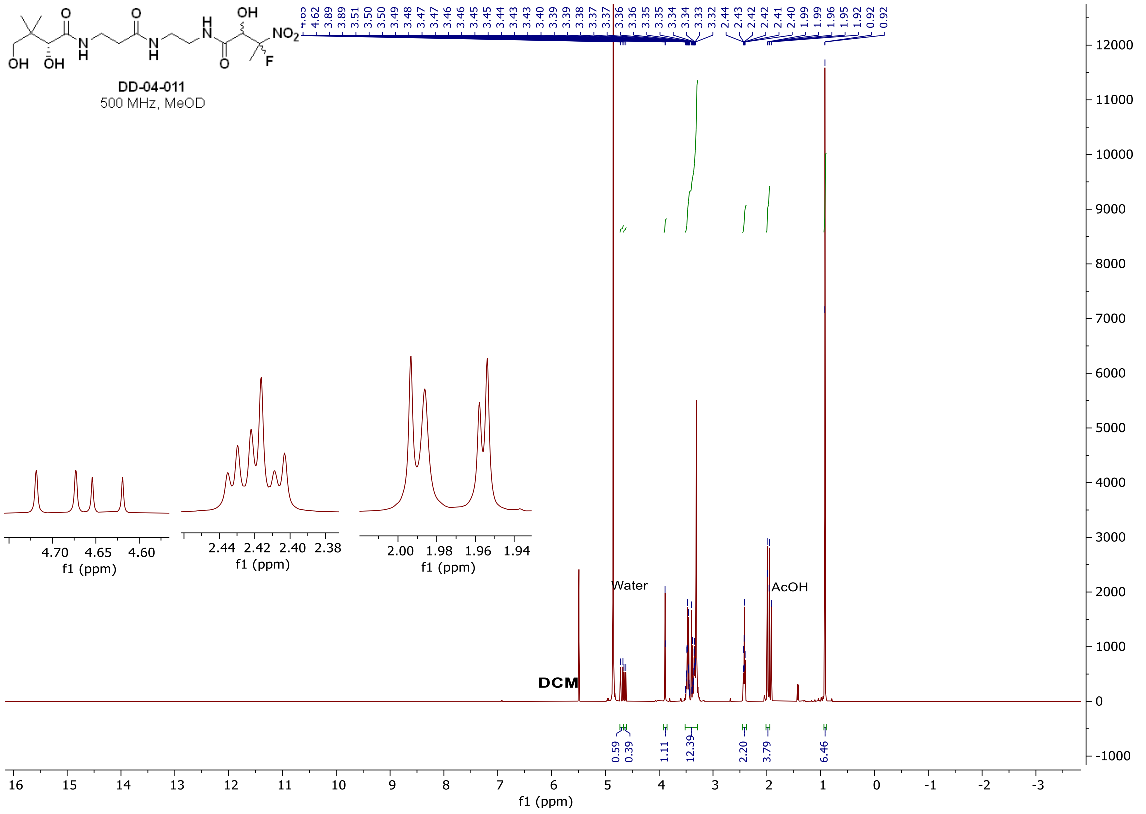

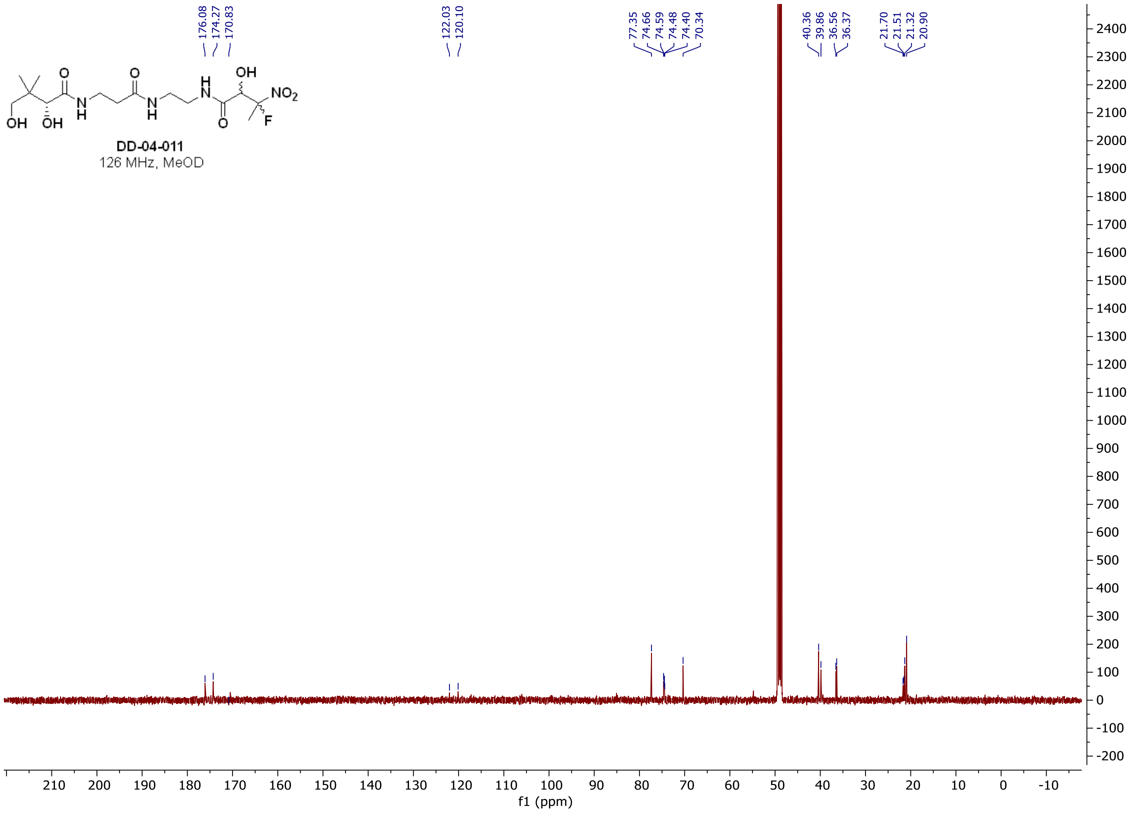
DD-04-011
126 MHz, MeOD
176.08
174.27
170.83
122.03
120.10
77.35
74.66
74.59
74.48
74.40
70.34
40.36
39.86
36.56
36.37
21.70
21.51
21.32
20.90
f1 (ppm)

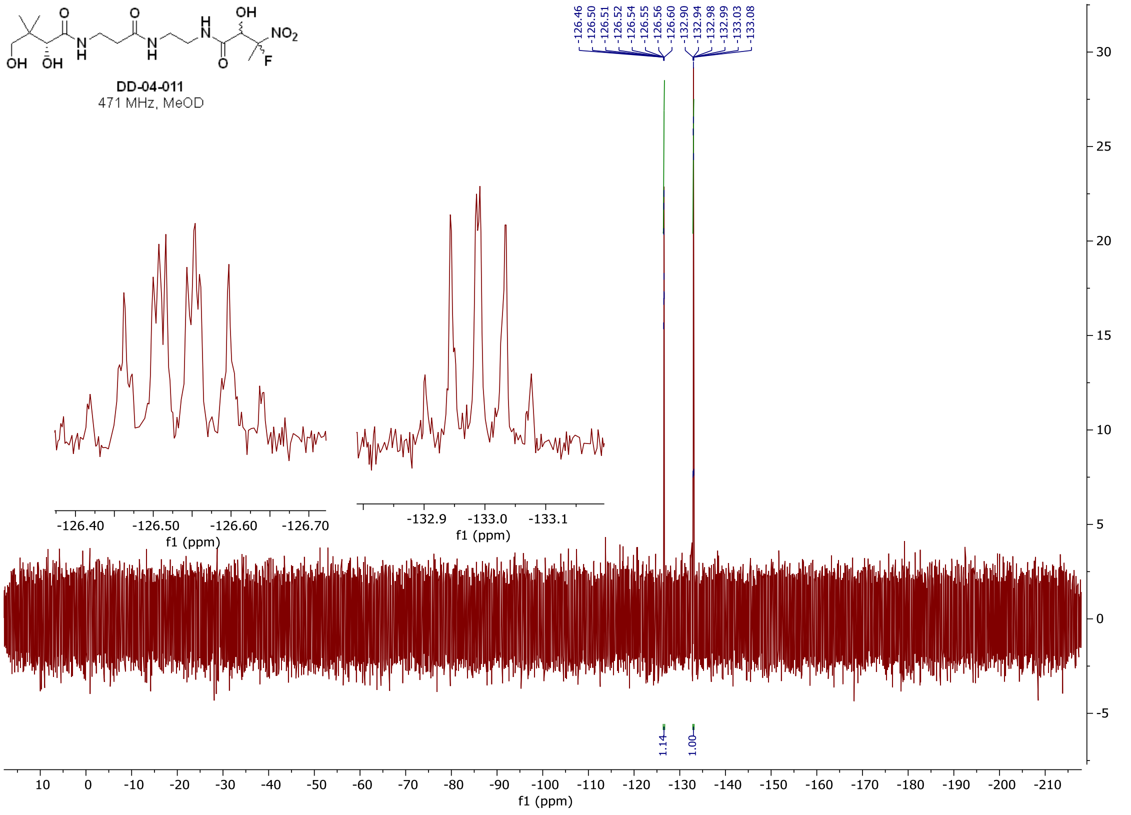
DD-04-011
471 MHz, MeOD
-126.46
-126.50
-126.51
-126.52
-126.54
-126.55
-126.56
-126.60
-132.90
-132.94
-132.98
-132.99
-133.03
-133.08
1.14
1.00
f1 (ppm)

#### $^{1}H$ and $^{13}C$ **spectra of DD-05-090**

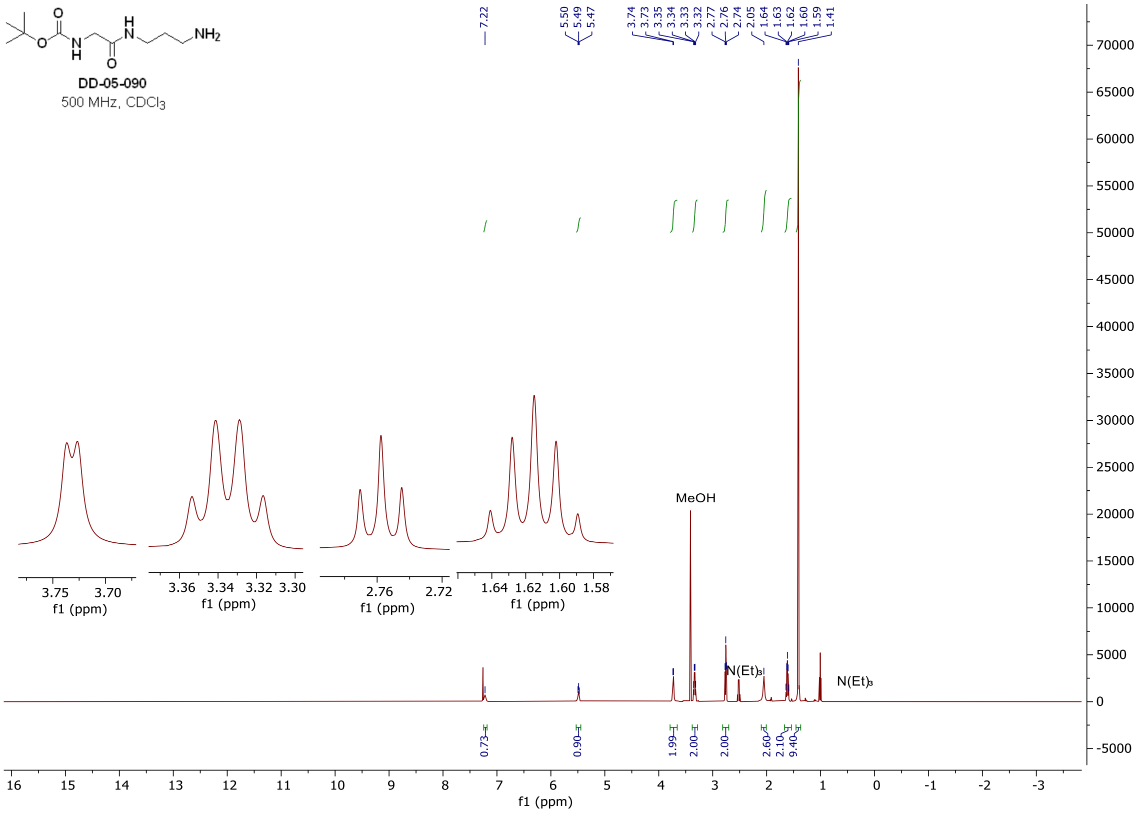

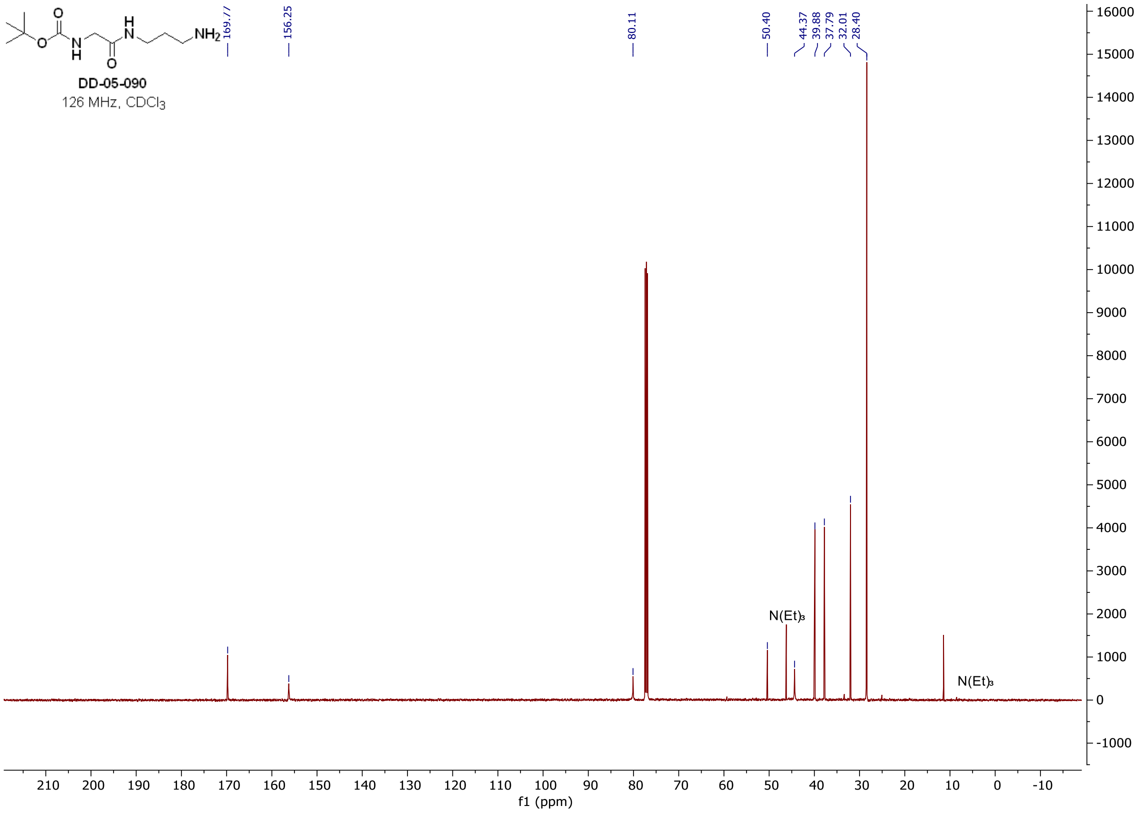

### $^{1}H$ and $^{13}C$ spectra of DD-05-095

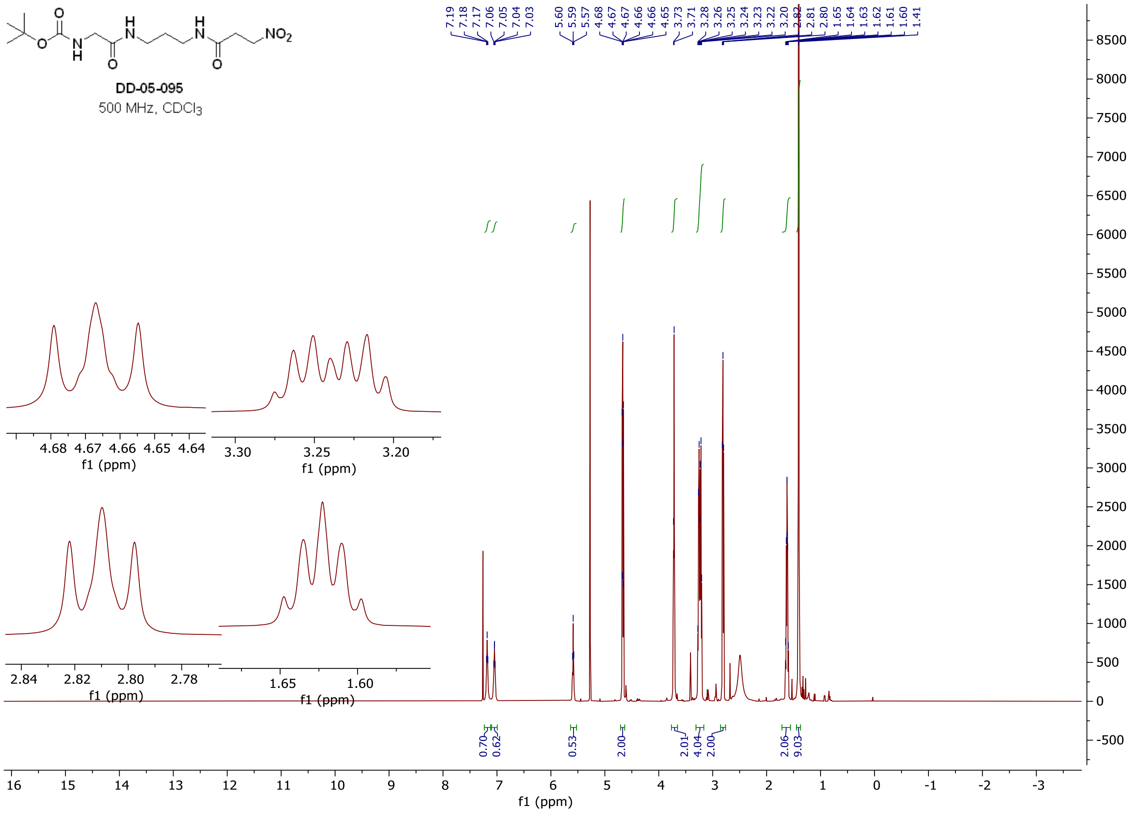

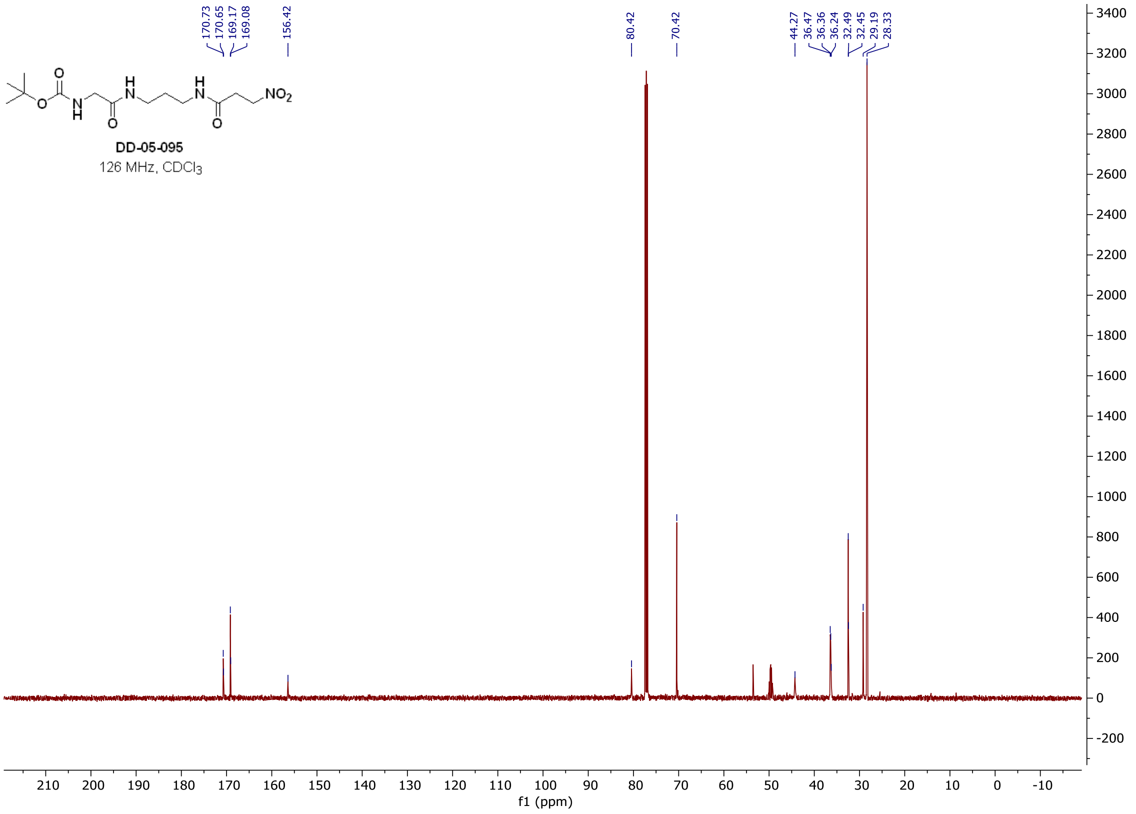

#### [1]H and [13]C **spectra of DD-05-101**

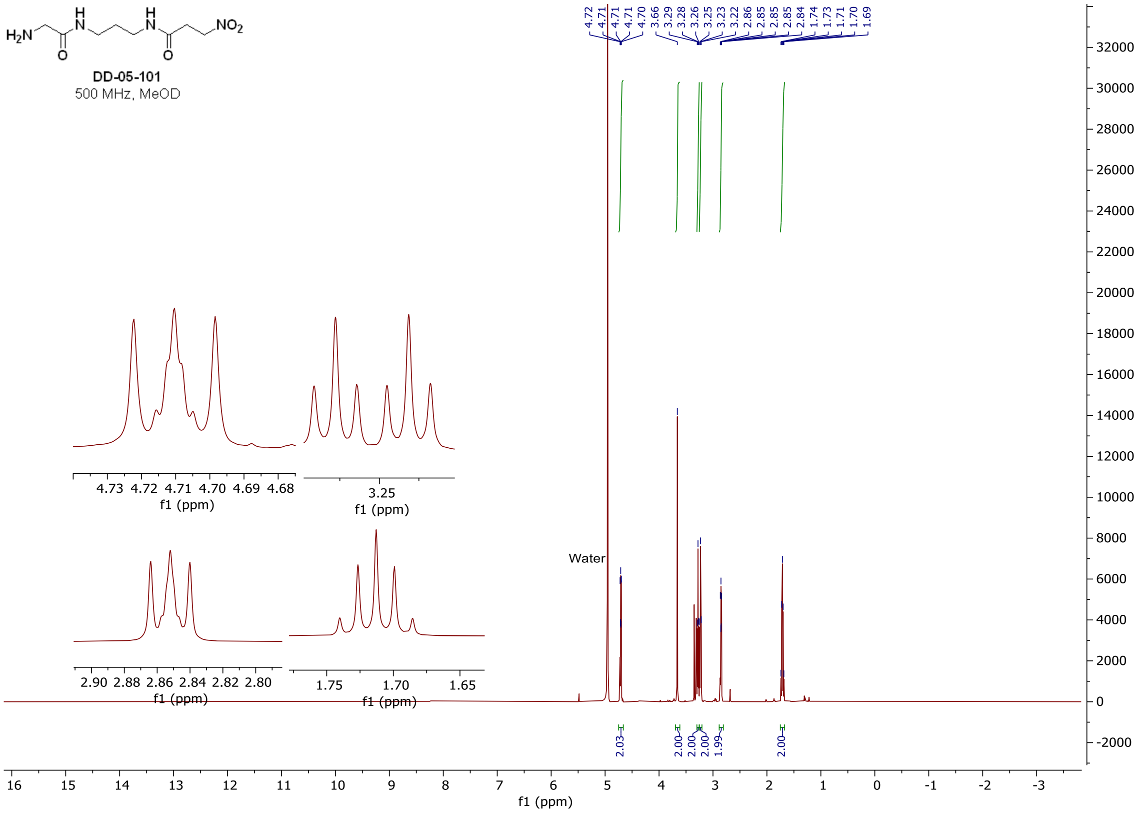

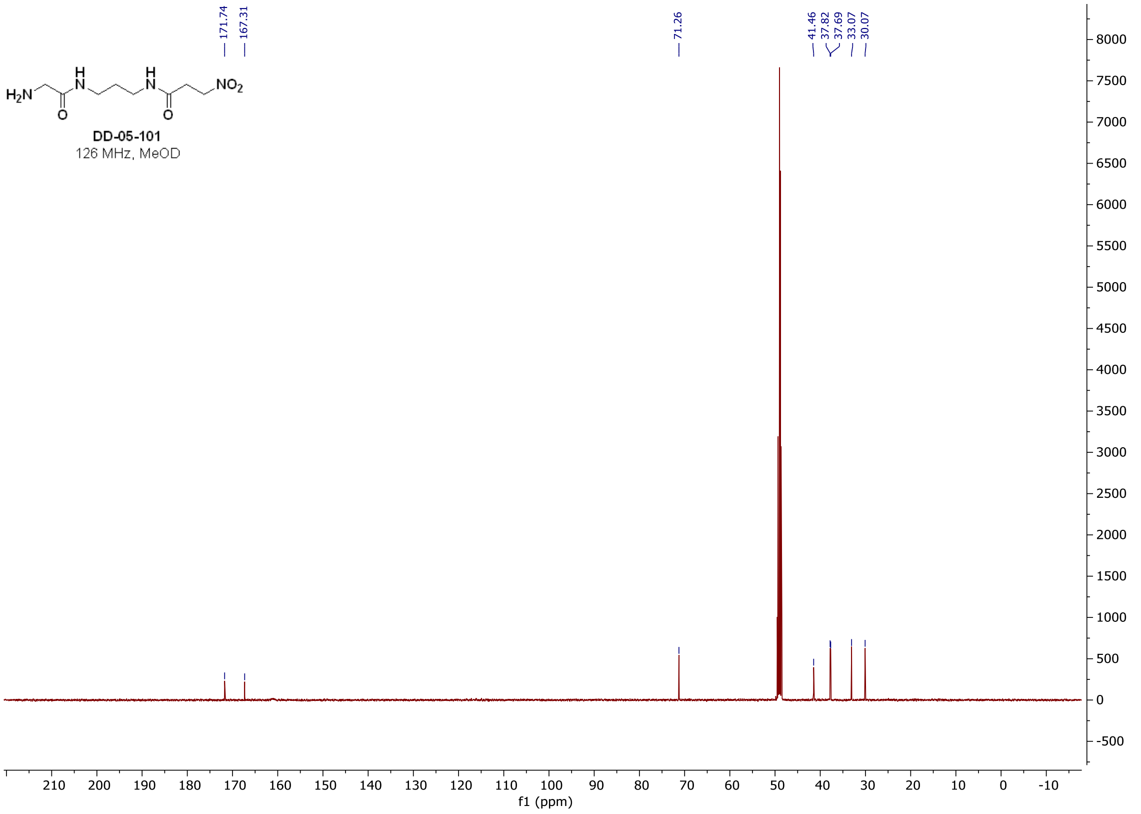

#### $^{1}H$ and $^{13}C$ spectra of DD-06-070

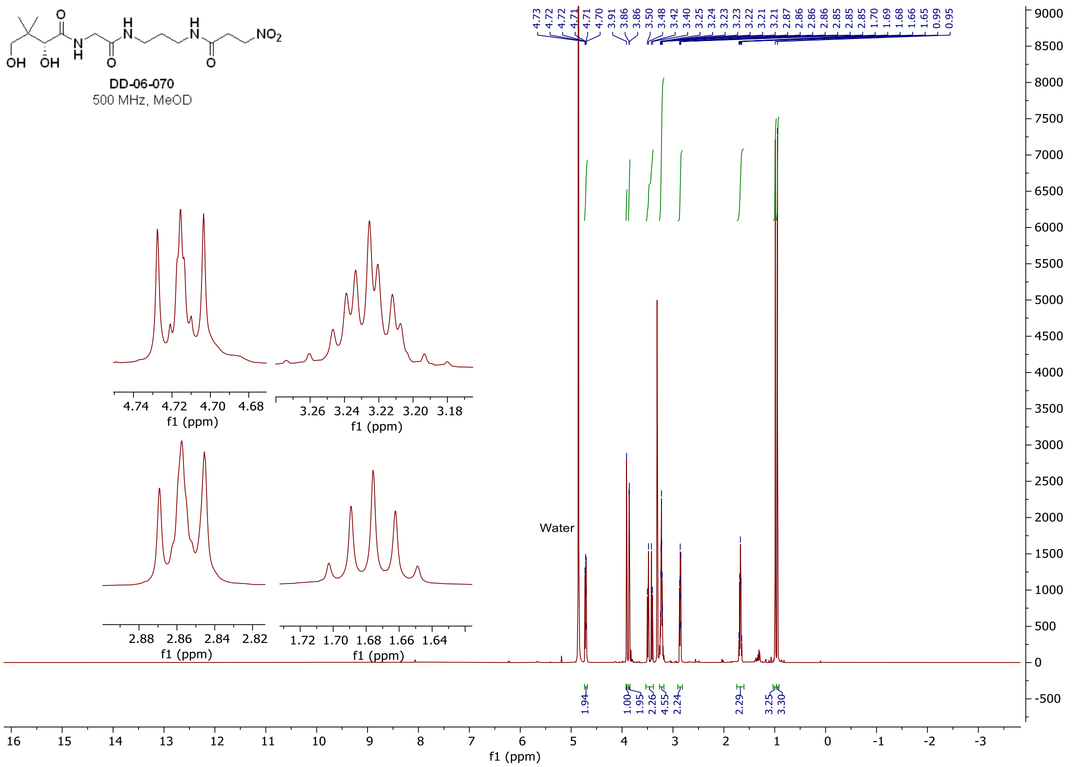

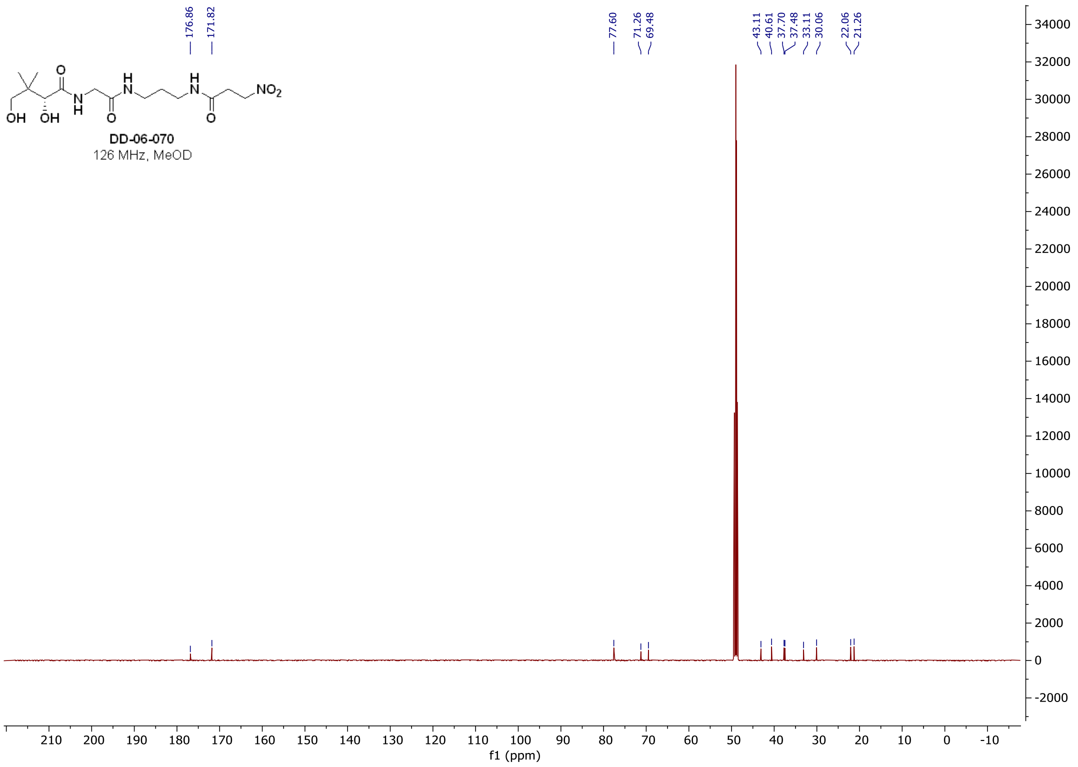

$^{1}$H and $^{13}$C **spectra of DD-04-104**

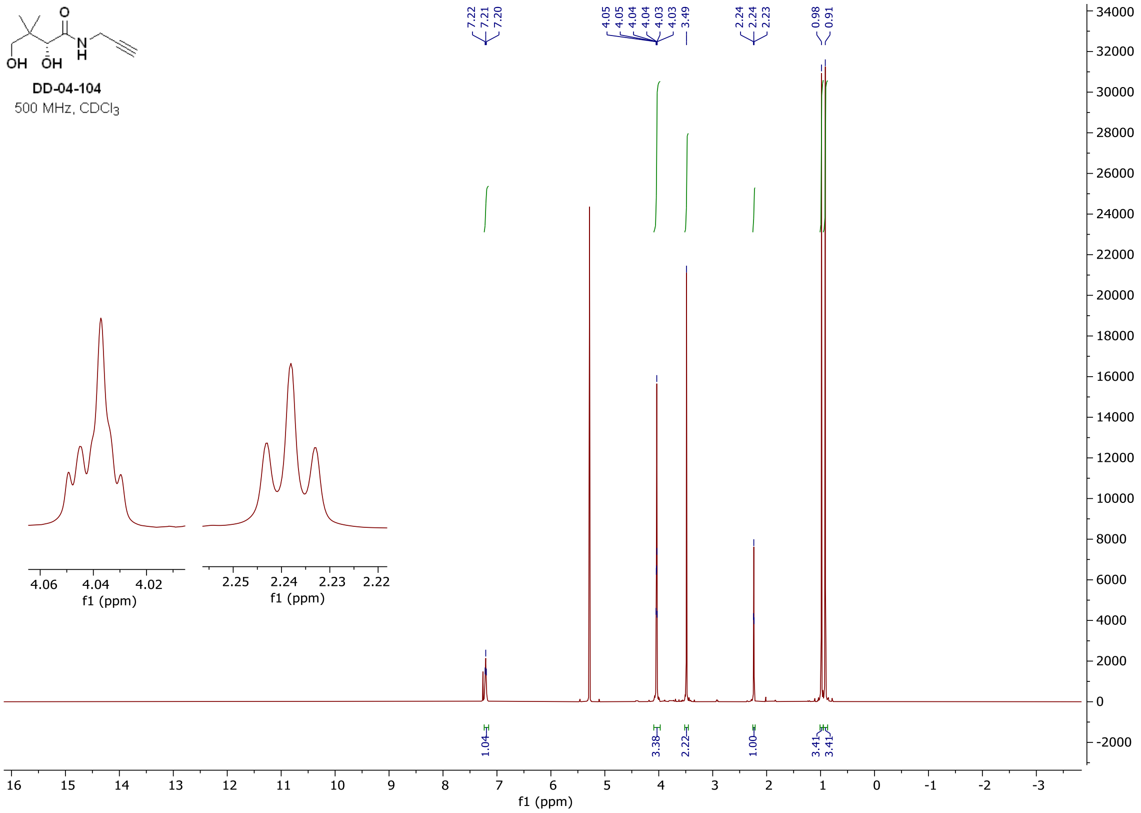

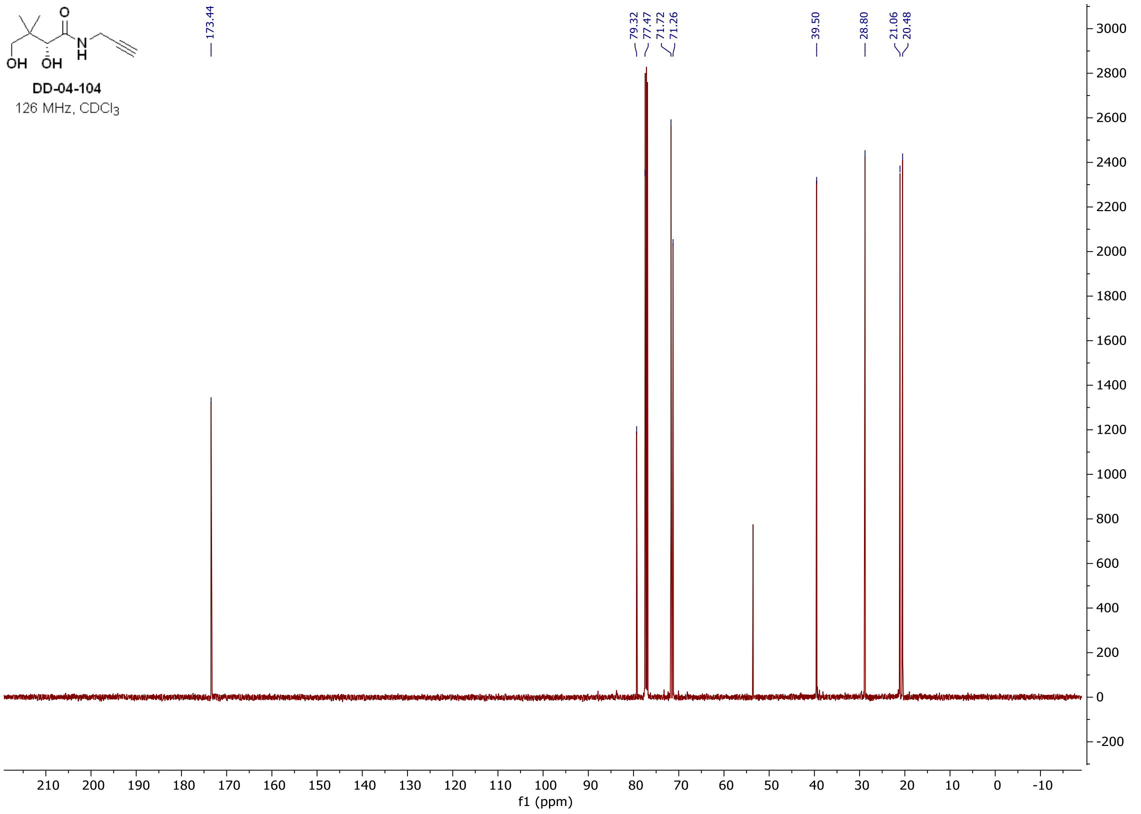

### [1]H and [13]C **spectra of DD-07-111**

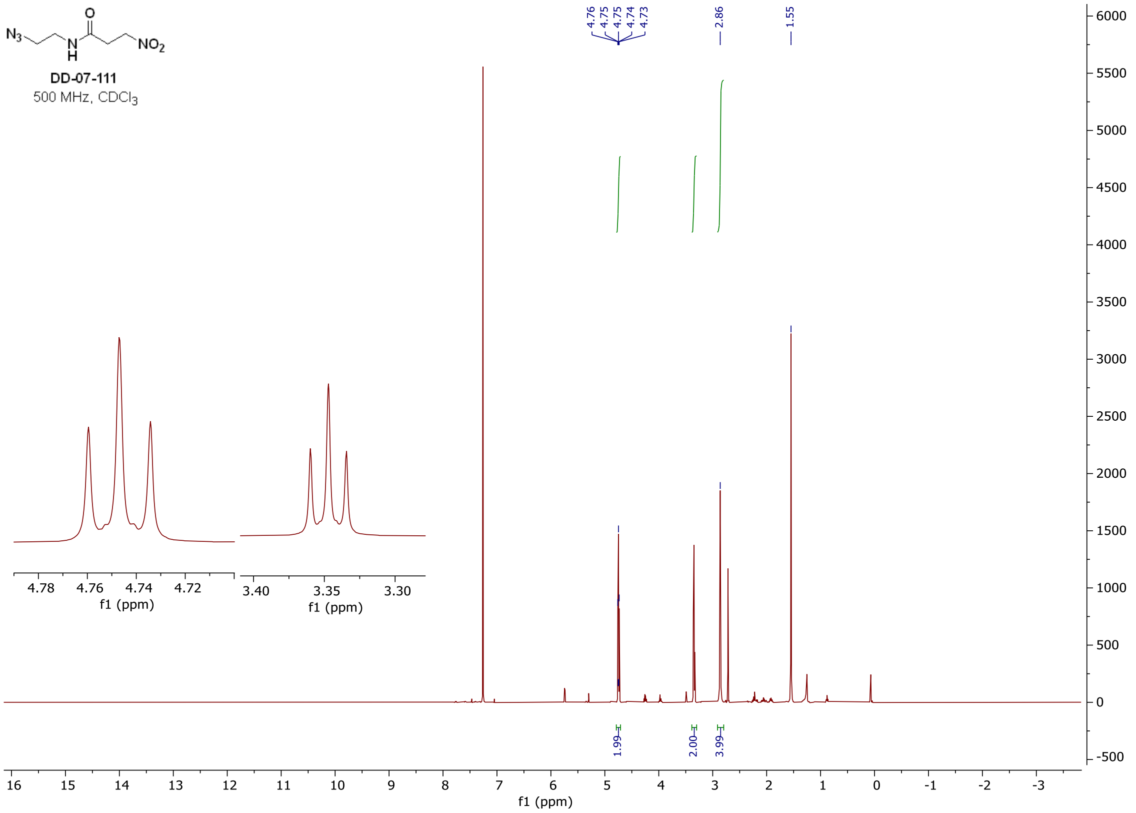

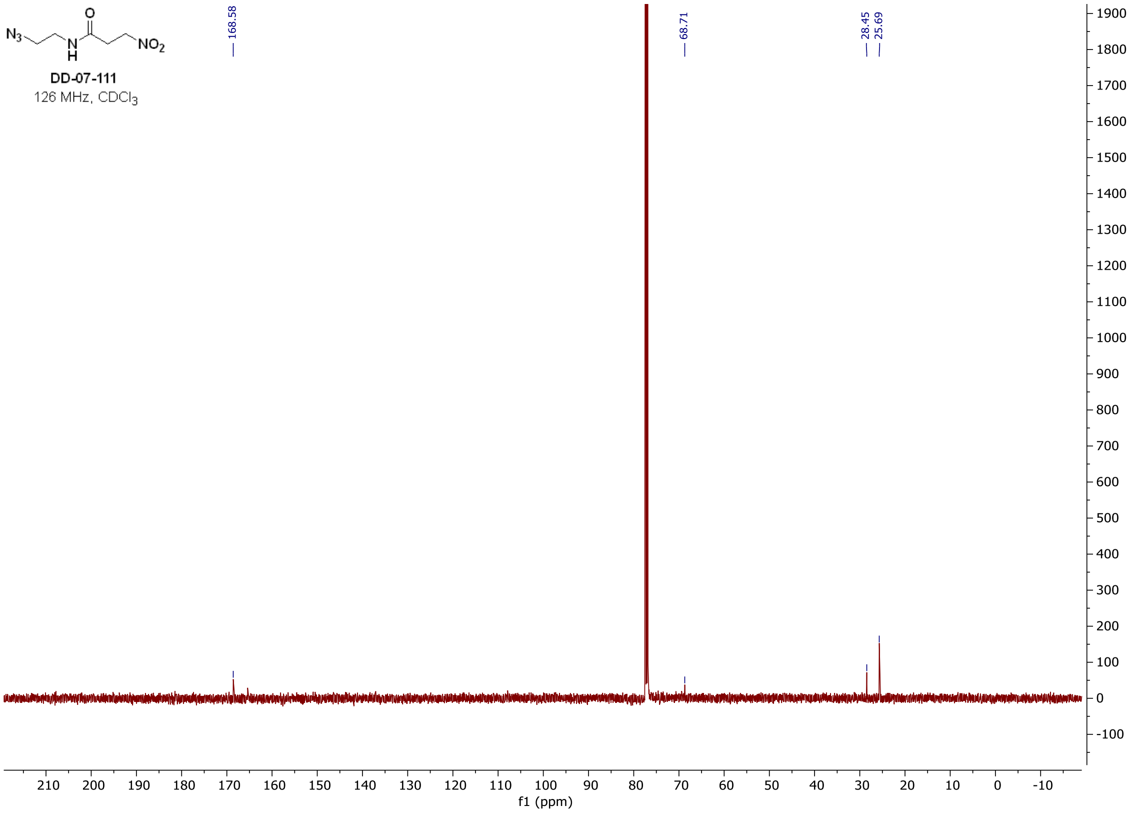

$^{1}H$ and $^{13}C$ **spectra of DD-04-109**

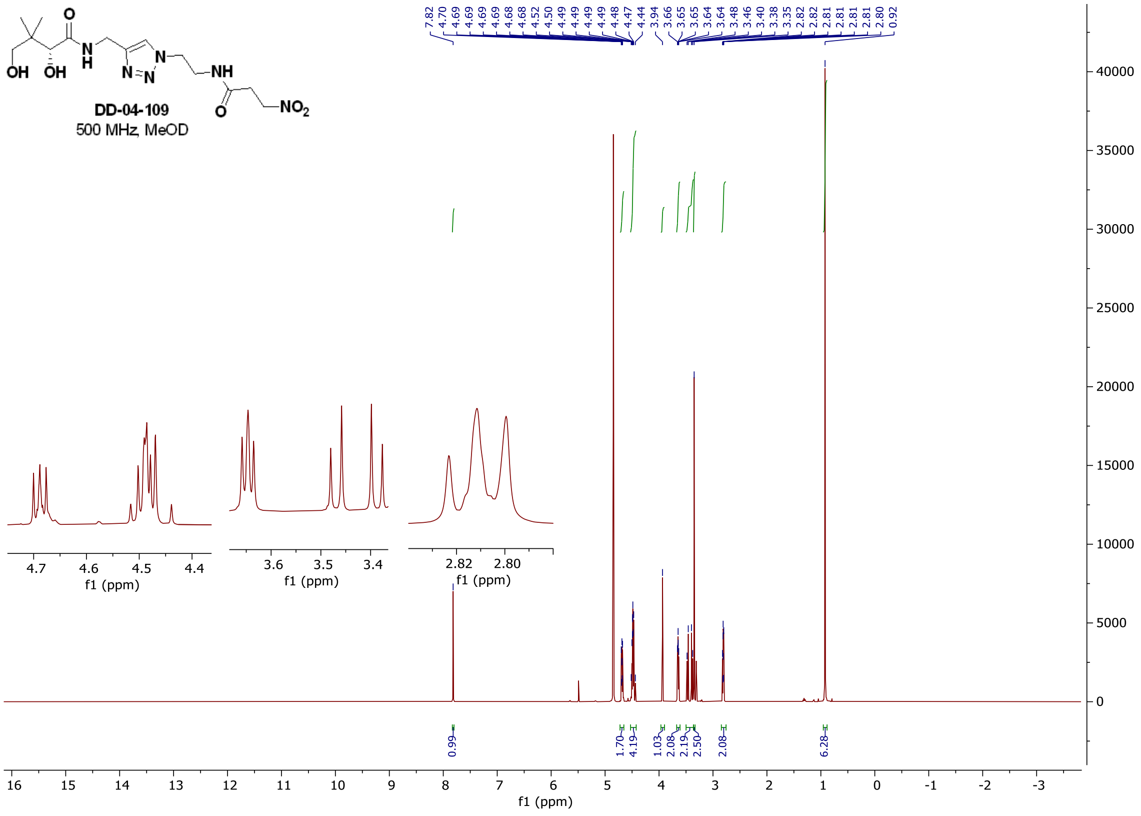

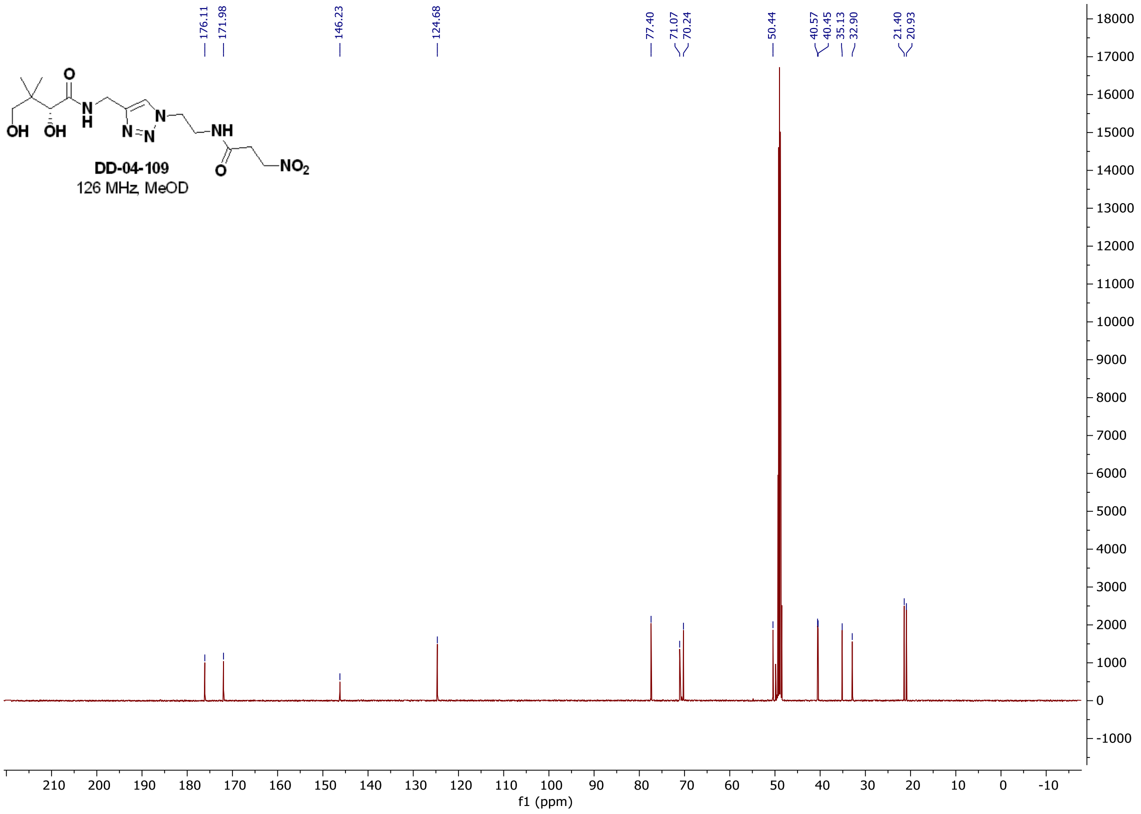

#### $^{1}$H **spectrum of DD-04-109**

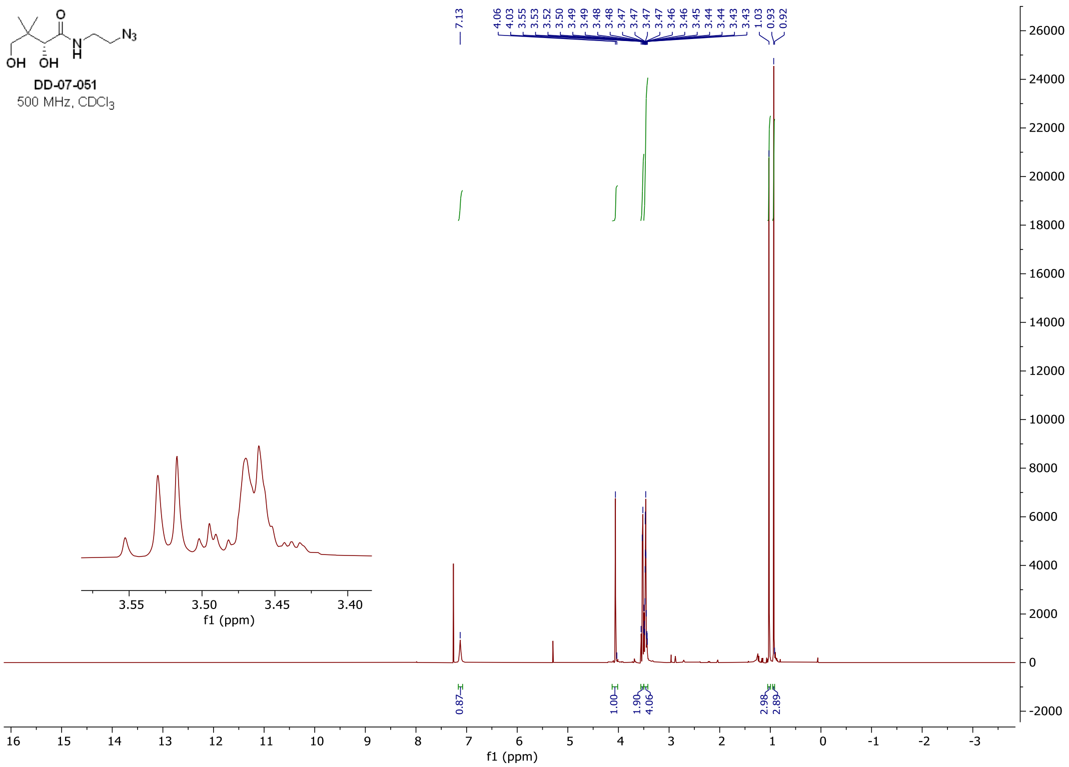

#### $^{1}$H and $^{13}$C **spectra of DD-07-014**

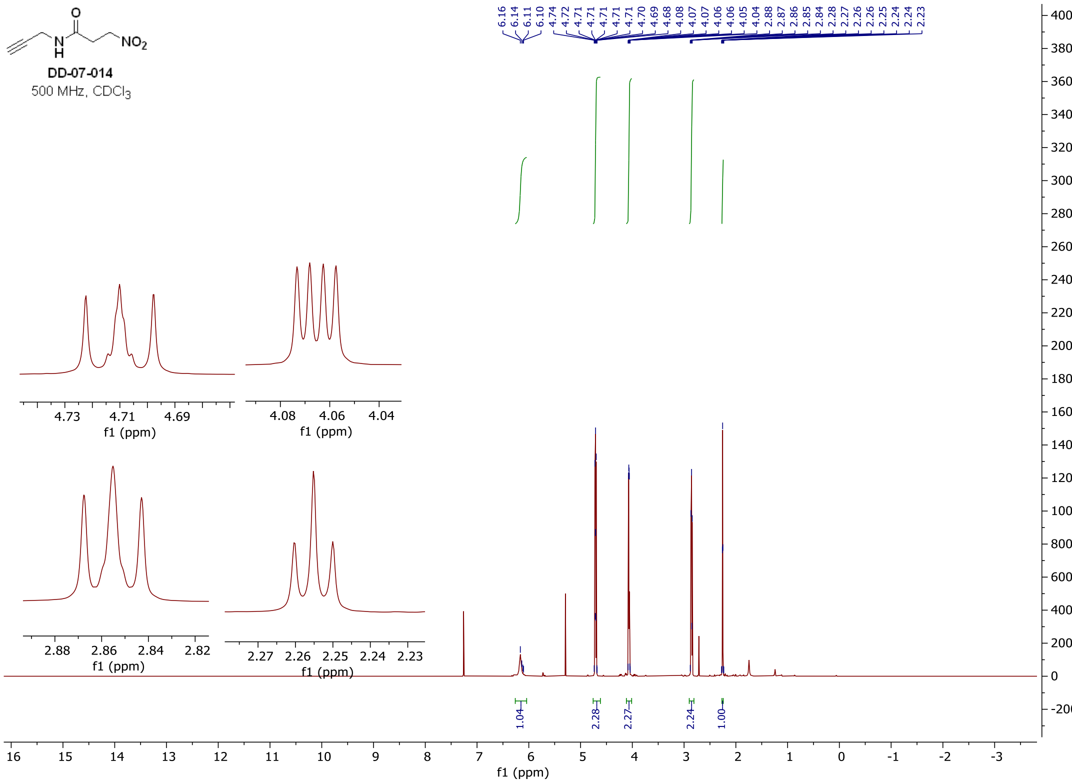

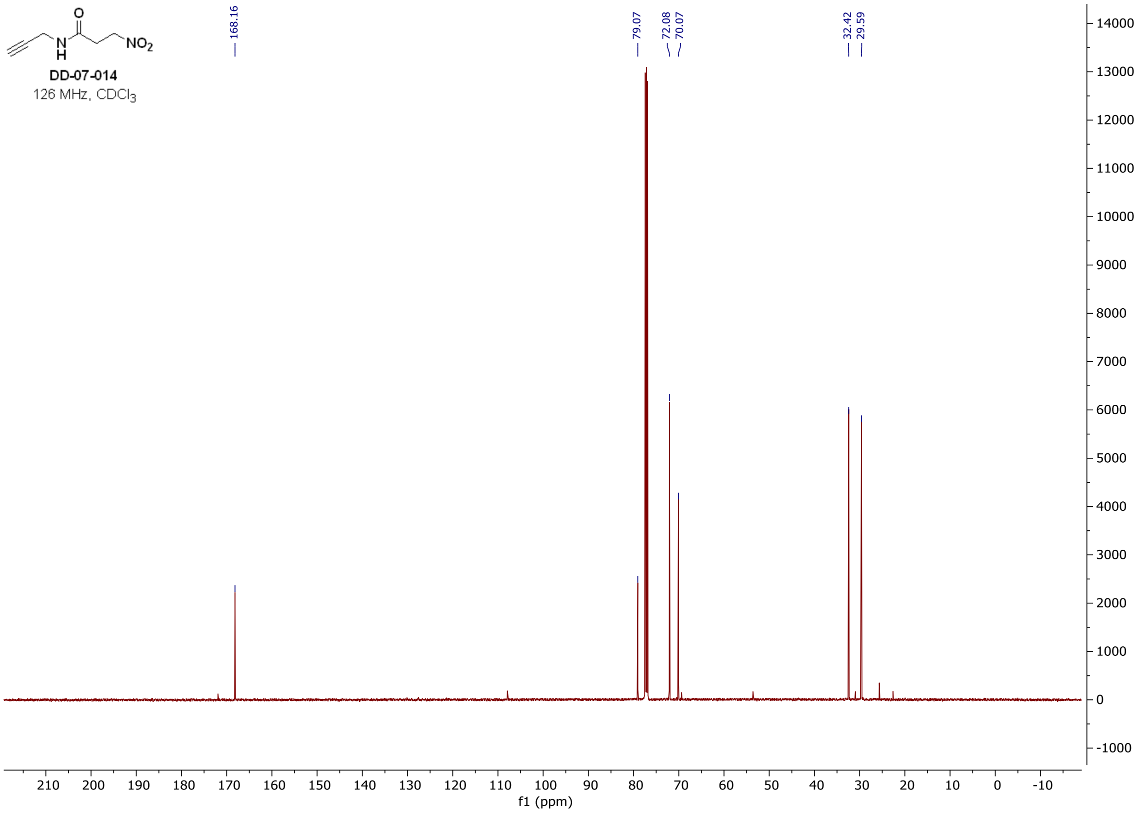

### [1]H and [13]C **spectra of DD-07-076**

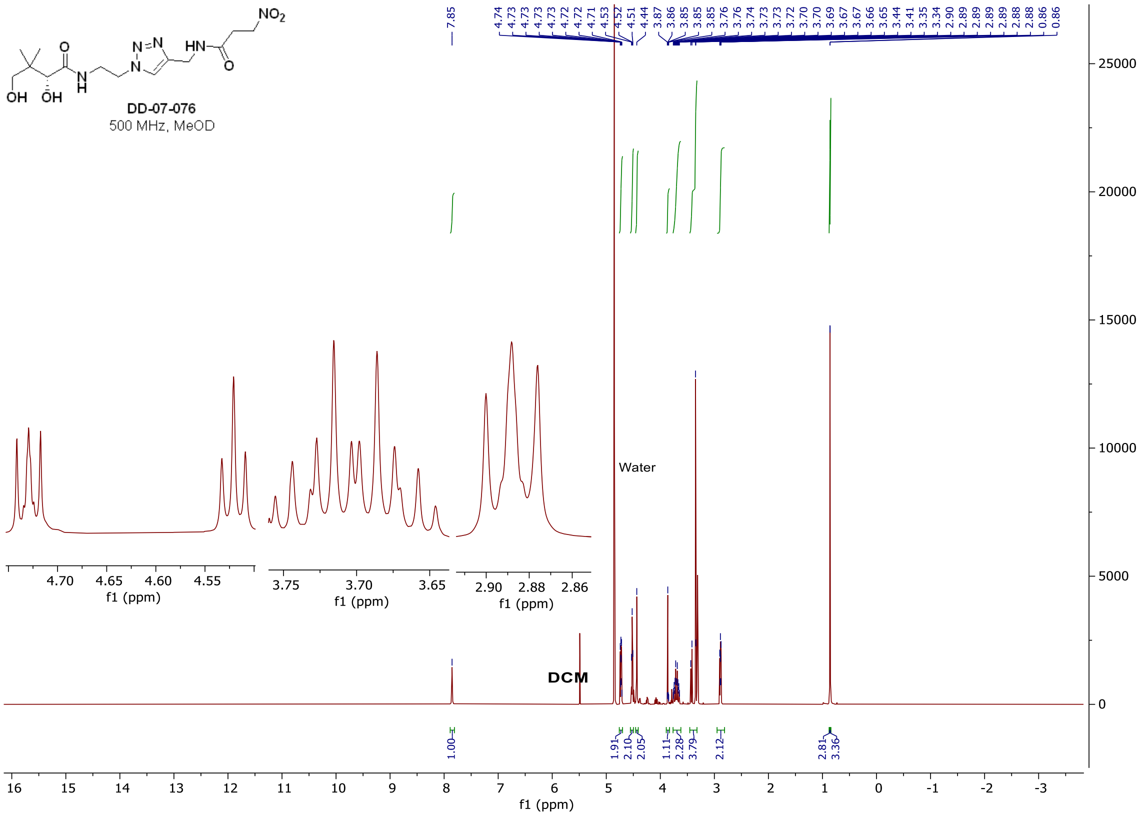

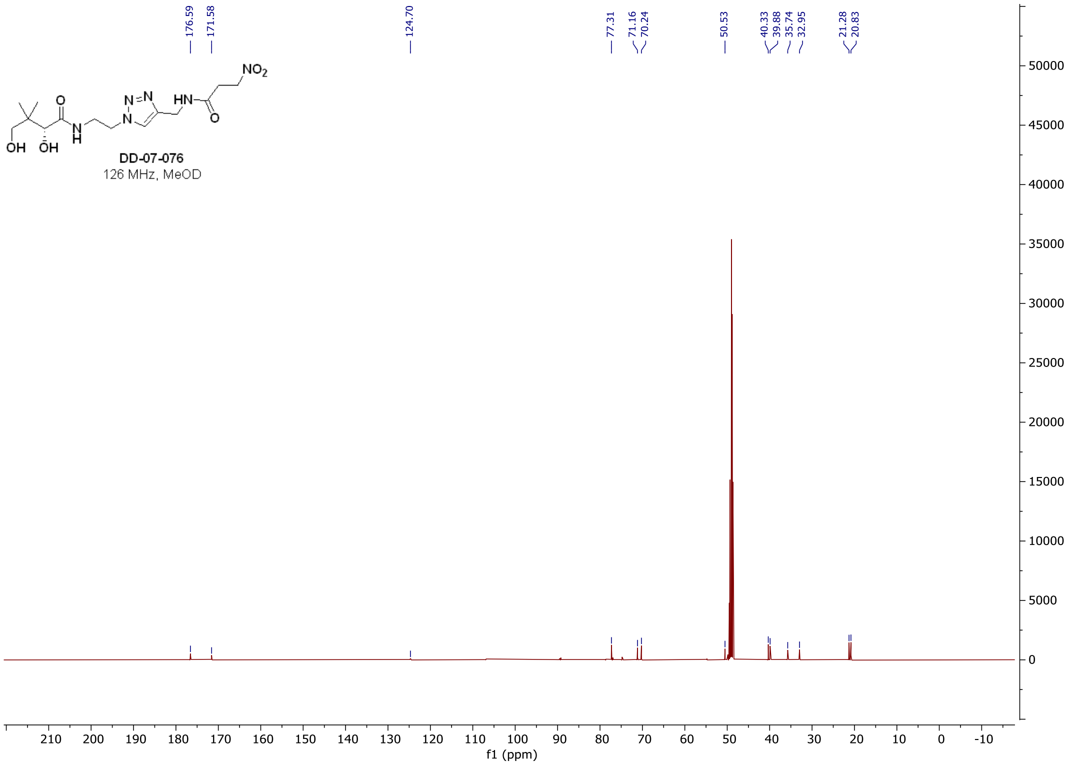

#### $^{1}$H, $^{13}$C, and $^{19}$F **spectra of DD-07-088**

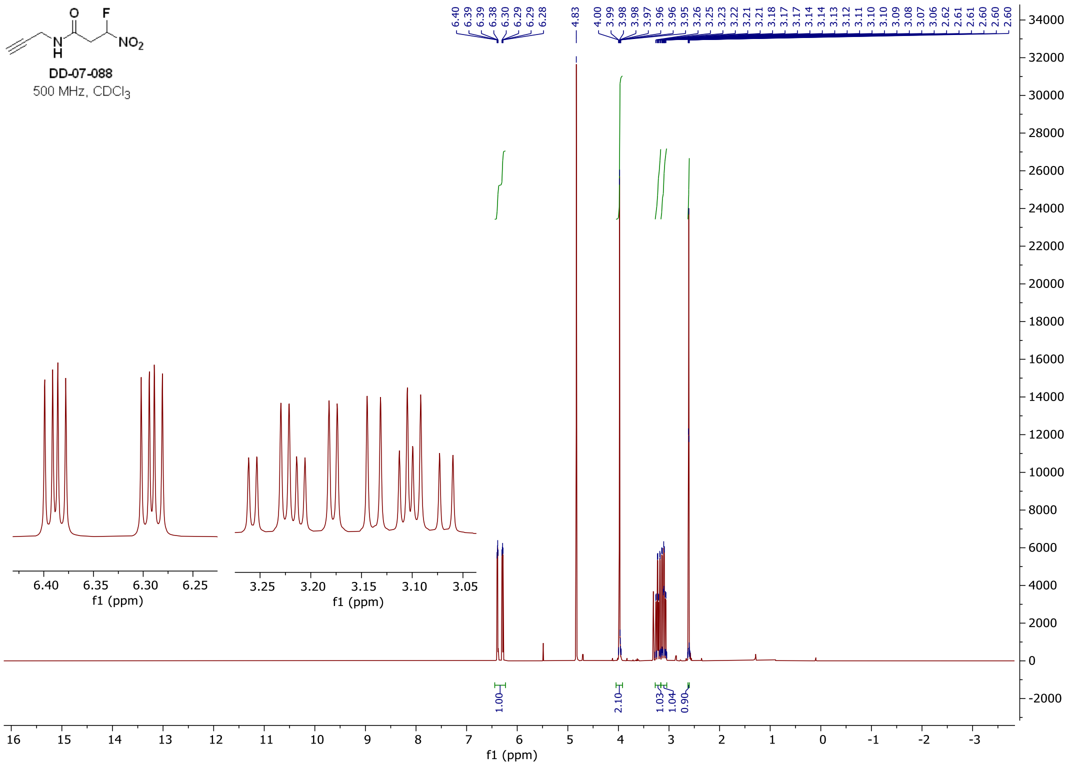

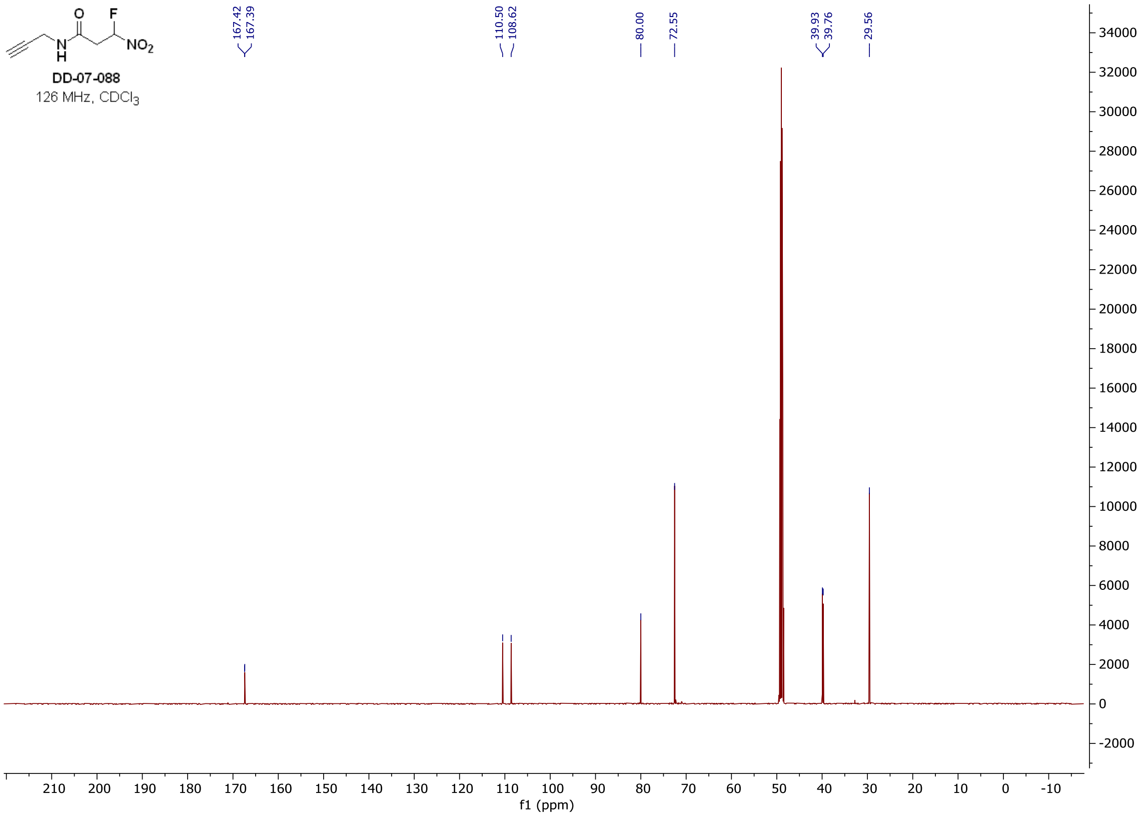
DD-07-088
126 MHz, CDCl3
167.42
167.39
110.50
108.62
80.00
72.55
39.93
39.76
29.56
f1 (ppm)

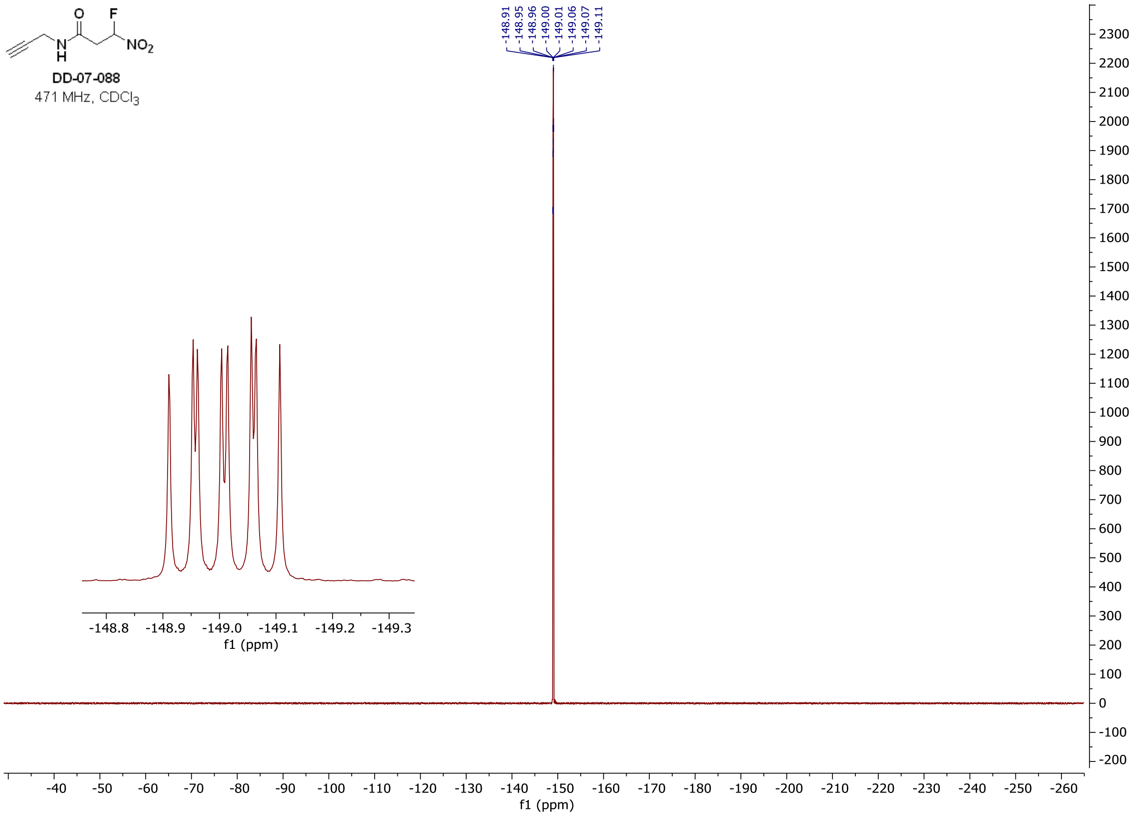
DD-07-088
471 MHz, CDCl3
-148.91
-148.95
-148.96
-149.00
-149.01
-149.06
-149.07
-149.11
f1 (ppm)

#### $^{1}H$, $^{13}C$, and $^{19}F$ **spectra of DD-07-102**

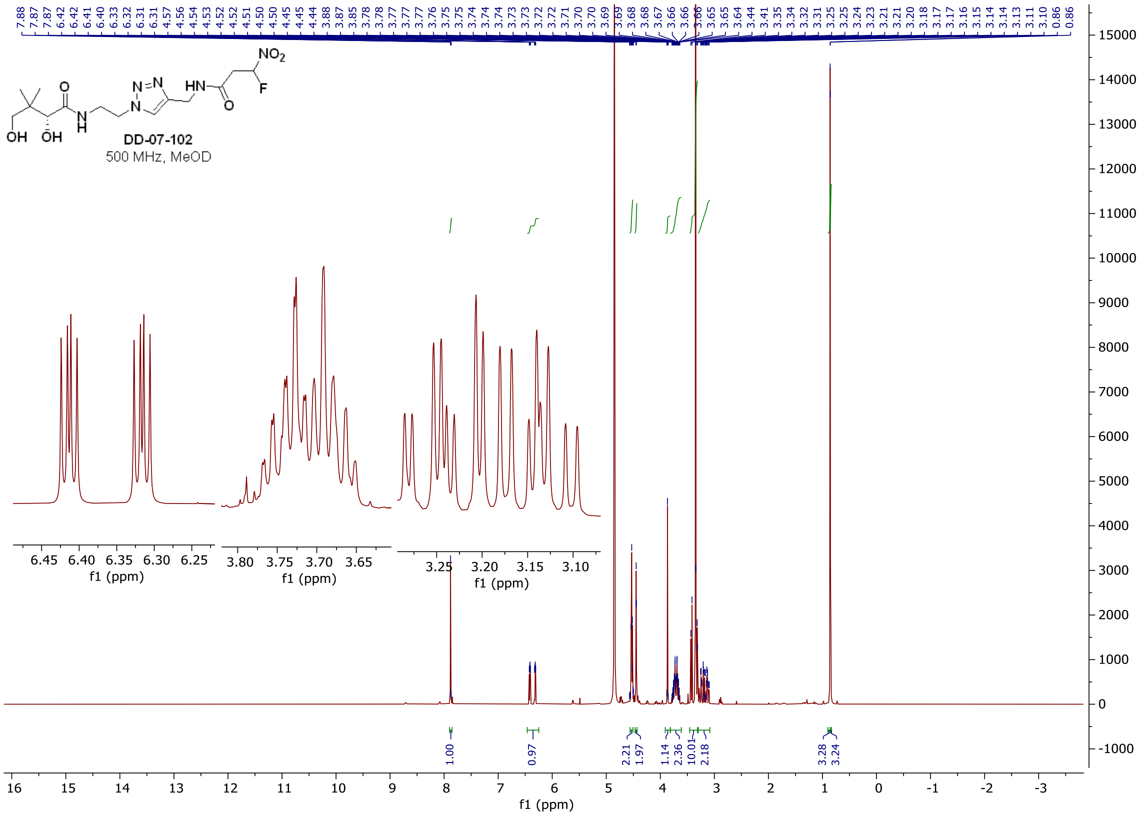

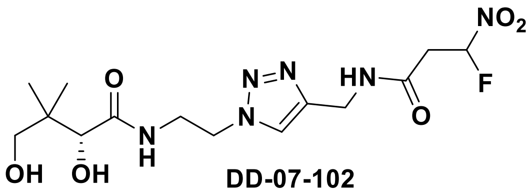

**DD-07-102**

471 MHz, MeOD

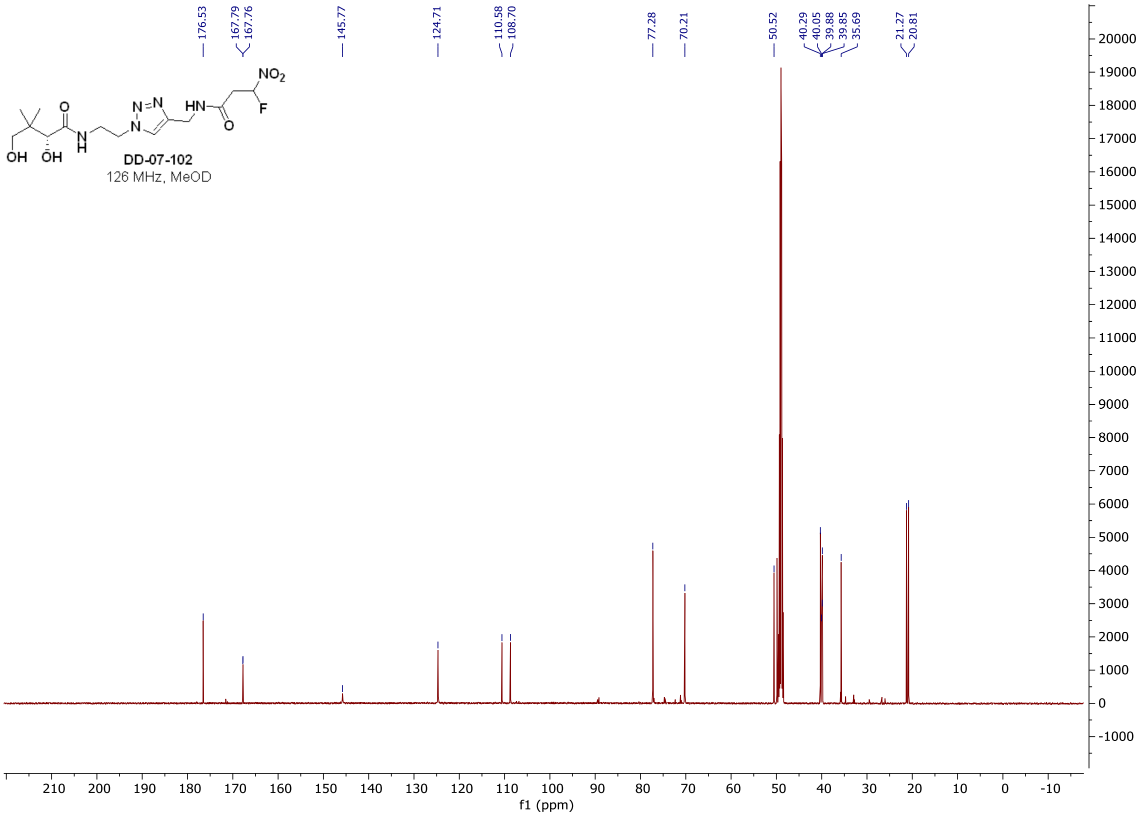
DD-07-102
126 MHz, MeOD

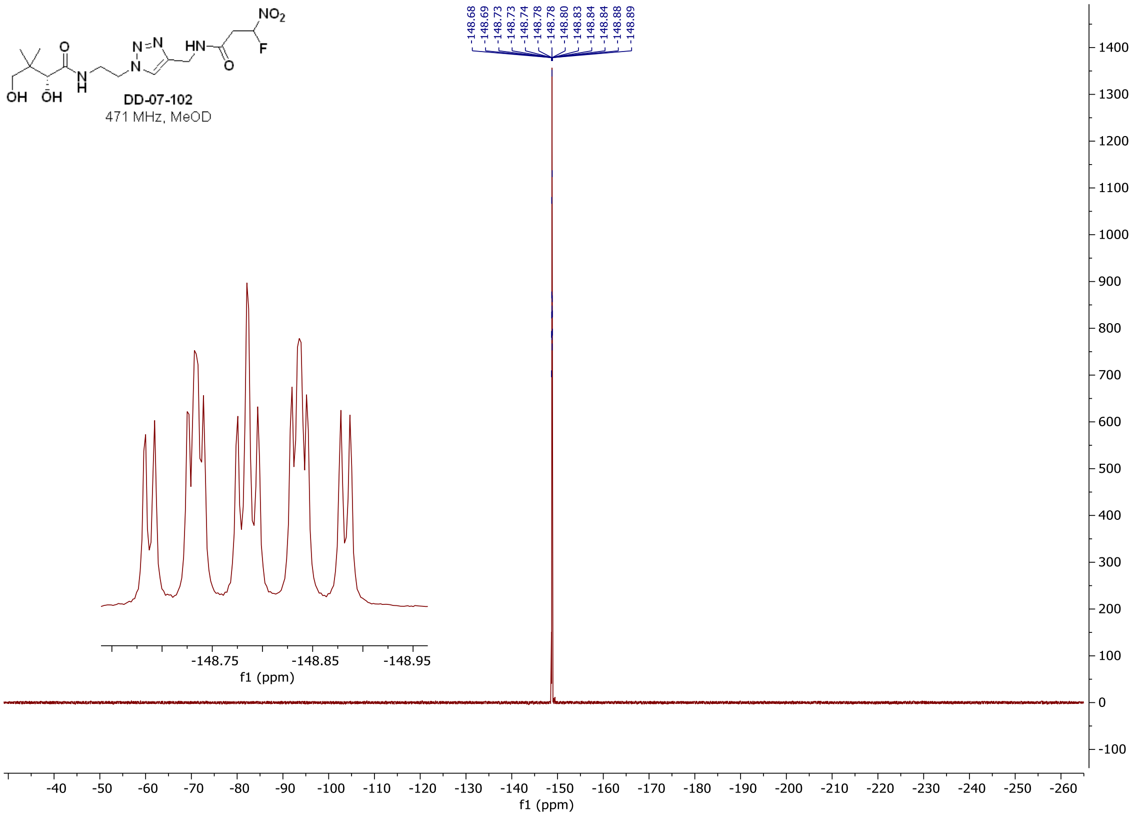
DD-07-102
471 MHz, MeOD

[1]H and [13]C **spectra of DD-03-117**

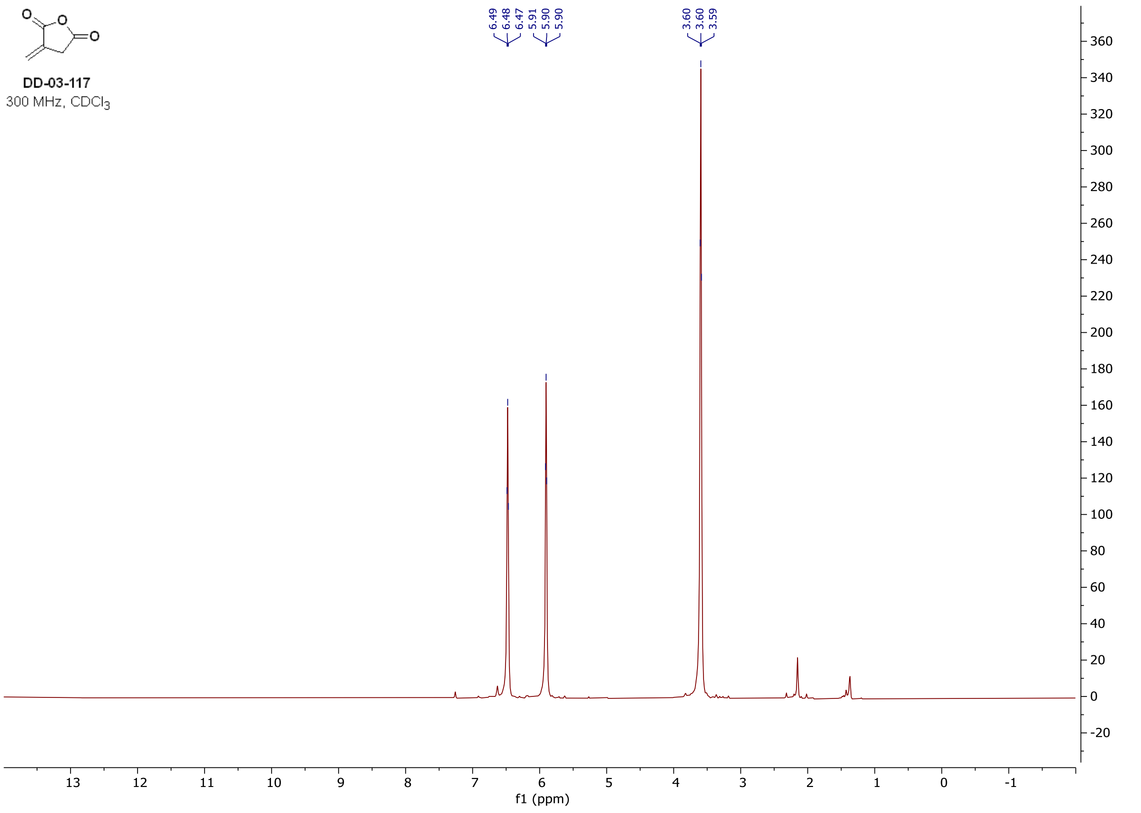

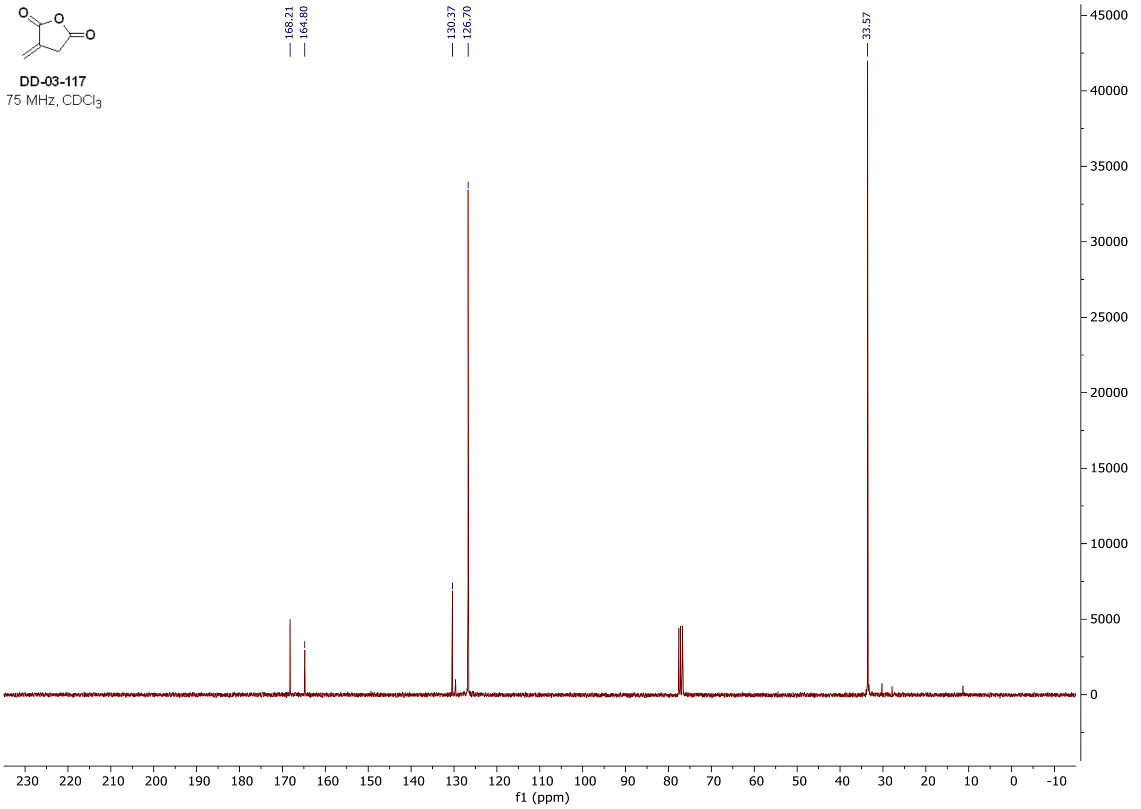

$^1$H and $^{13}$C **spectra of DD-07-131**

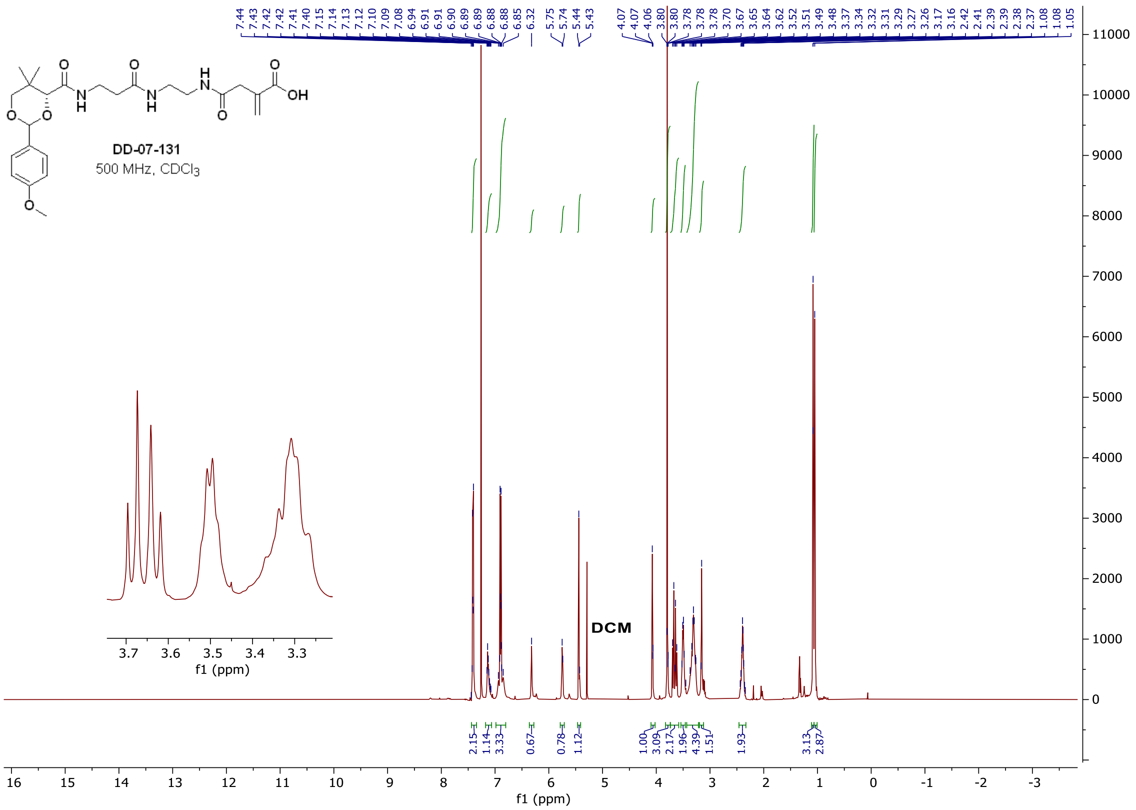

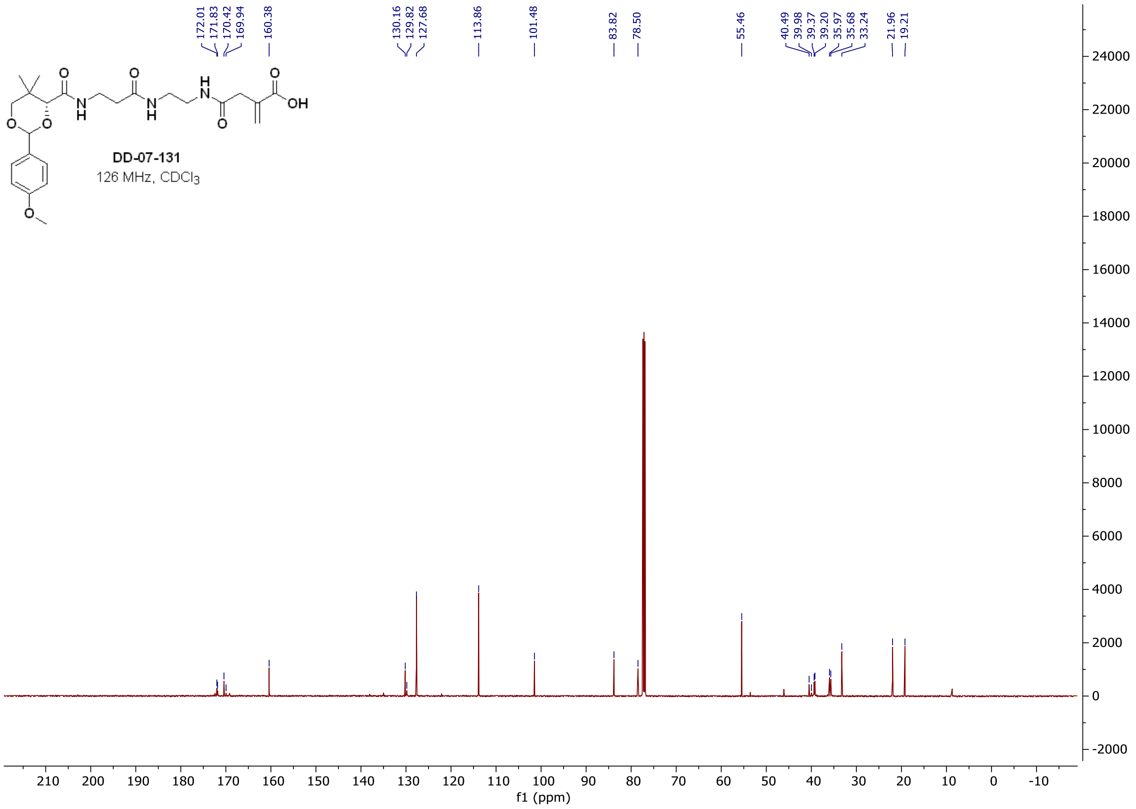

$^{1}H$ and $^{13}C$ **spectra of DD-07-139**

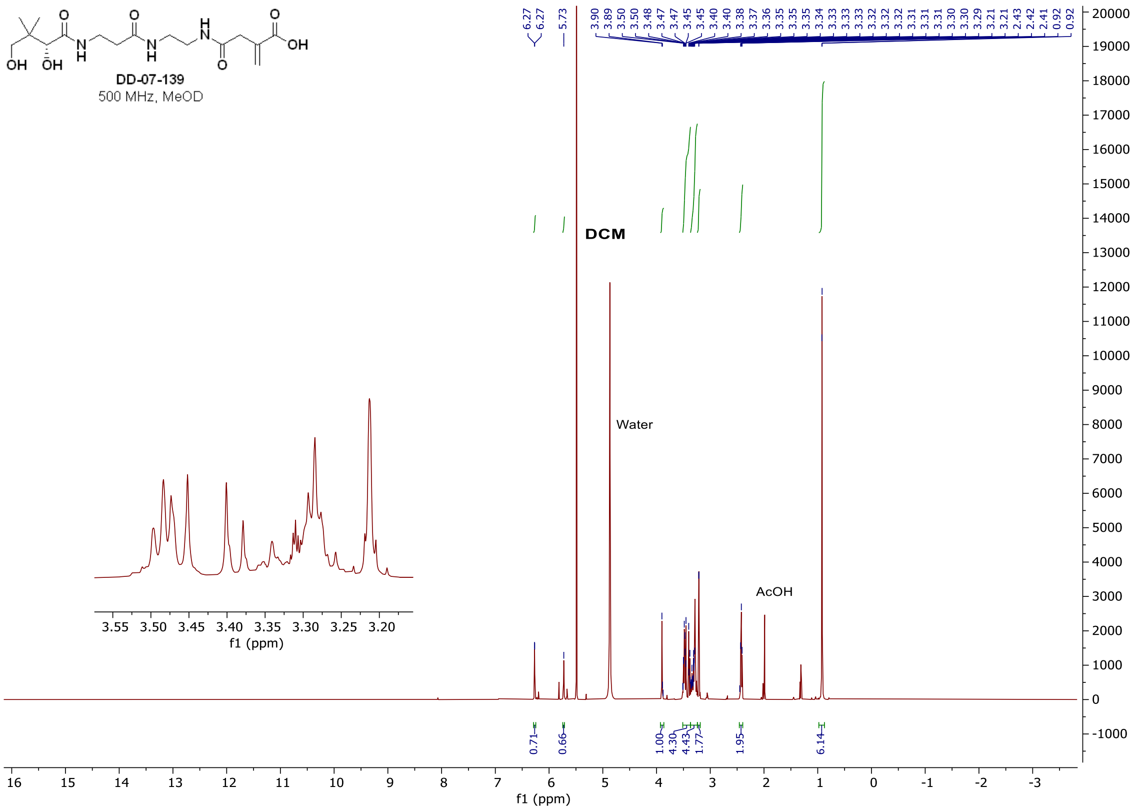

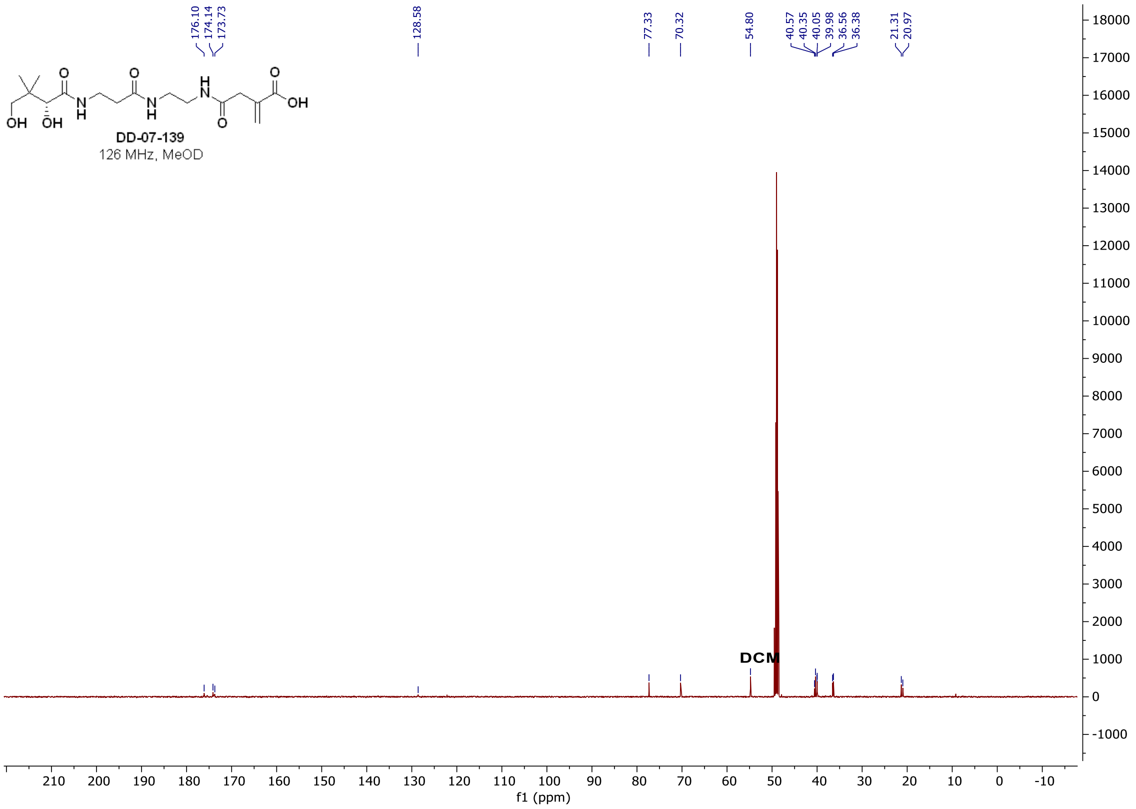

$^{1}$H and $^{13}$C **spectra of DD-03-141**

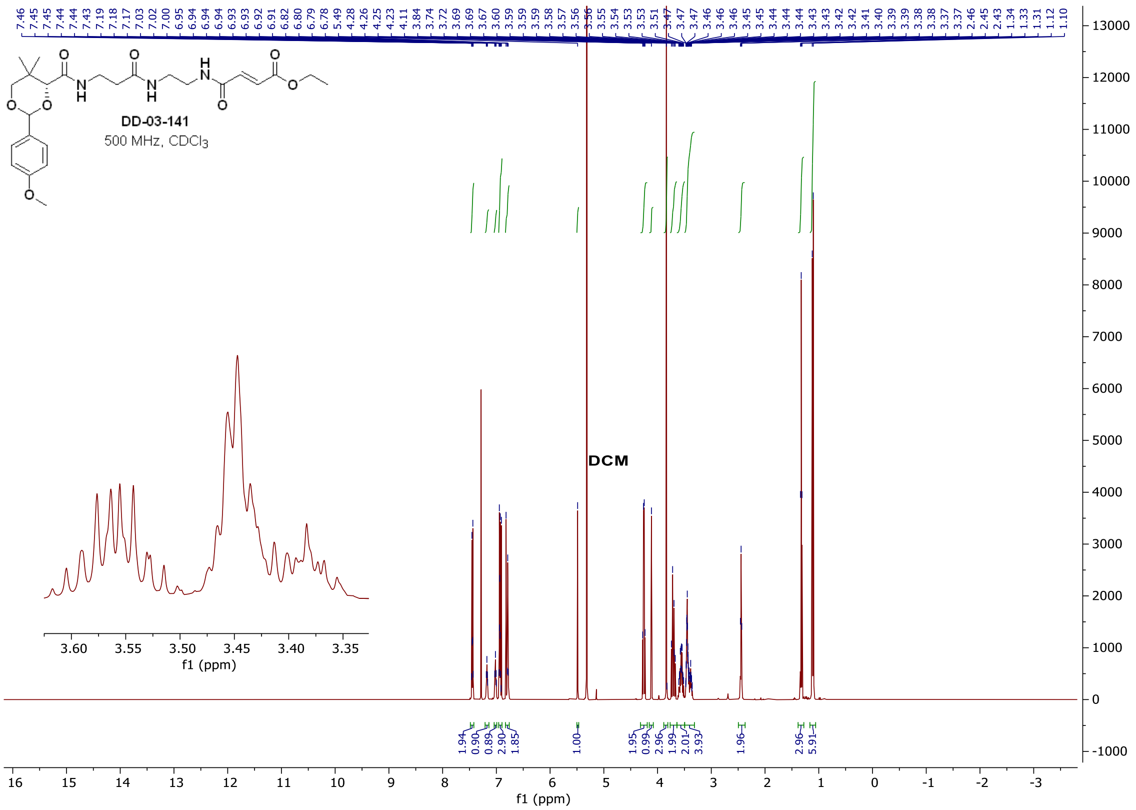

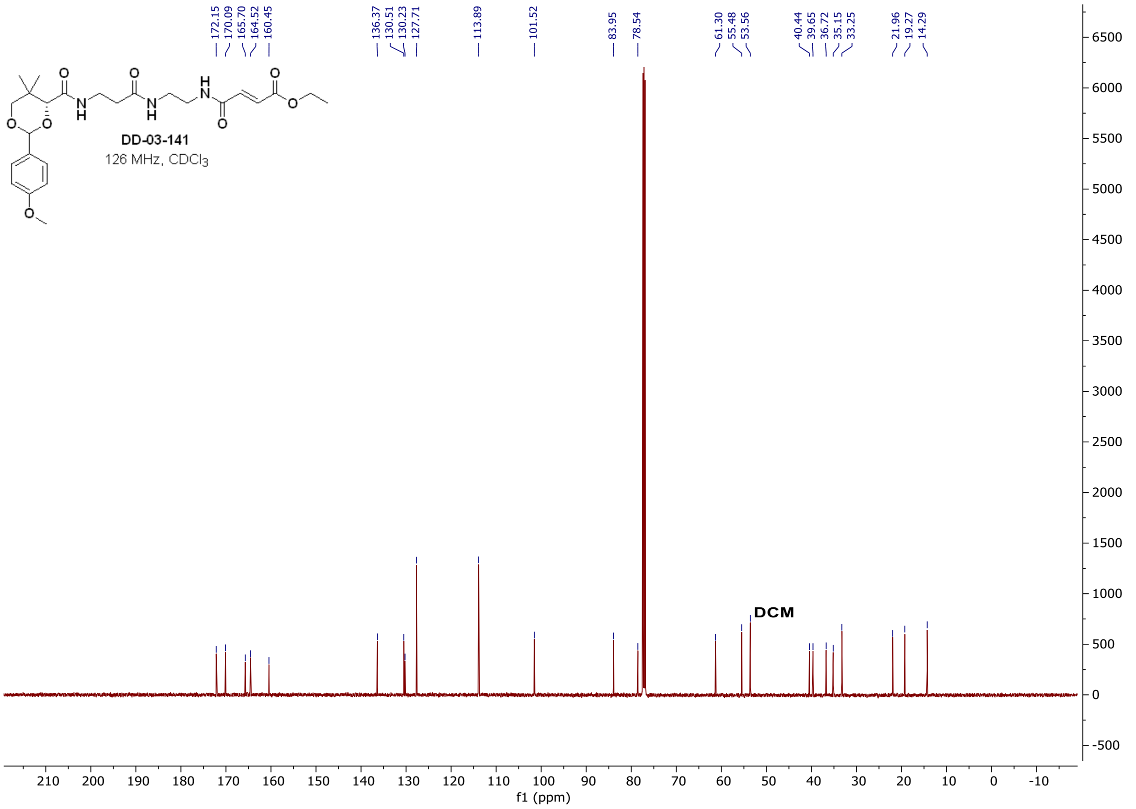

$^{1}H$ and $^{13}C$ **spectra of DD-06-147**

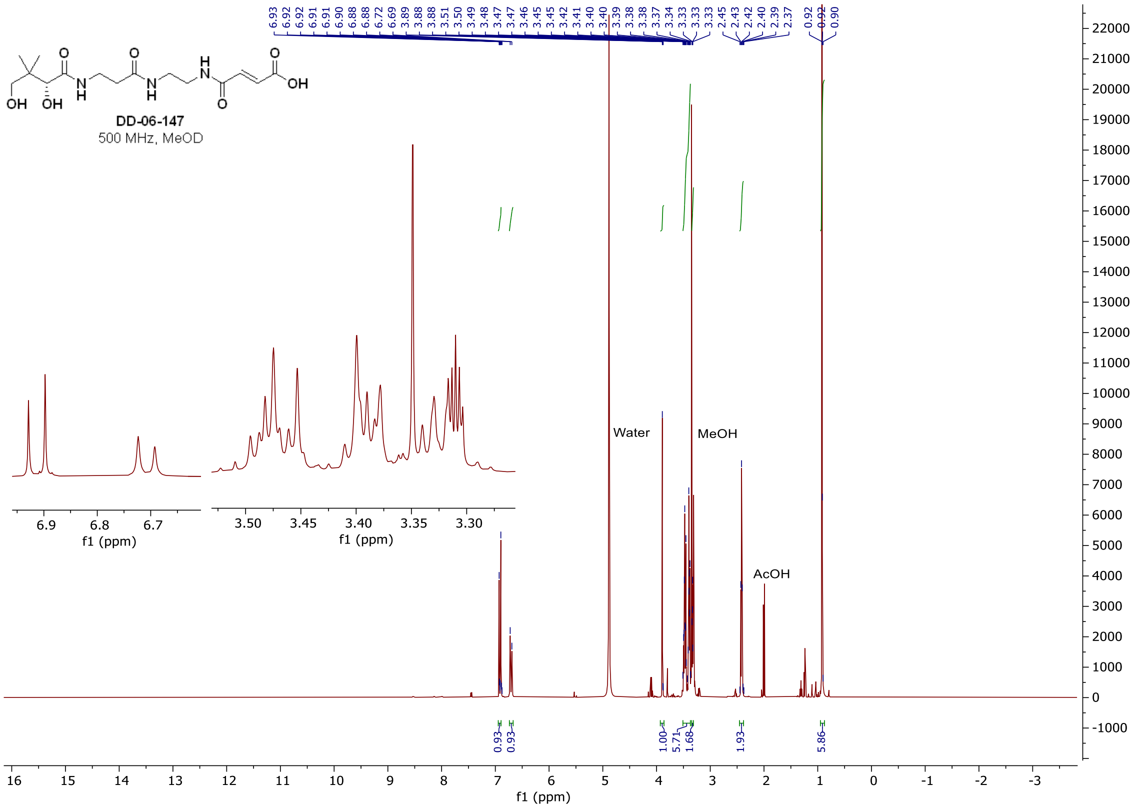

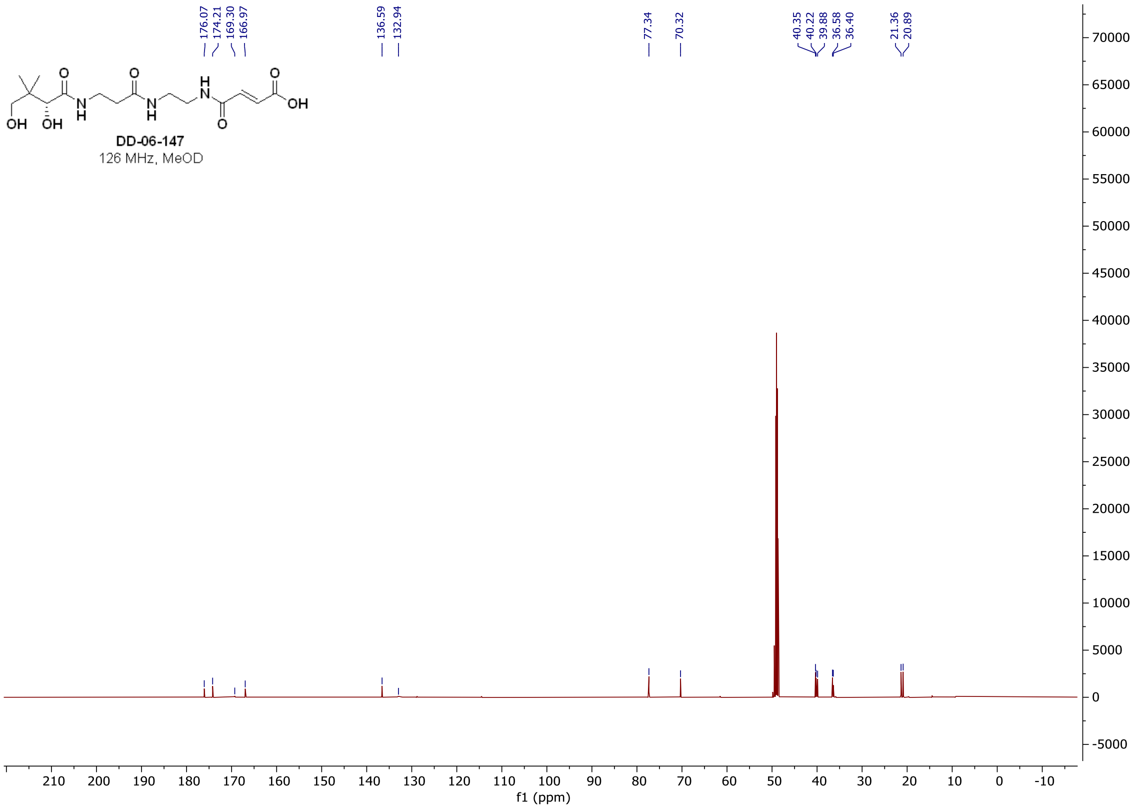

### $^{1}$H and $^{13}$C **spectra of DD-04-007**

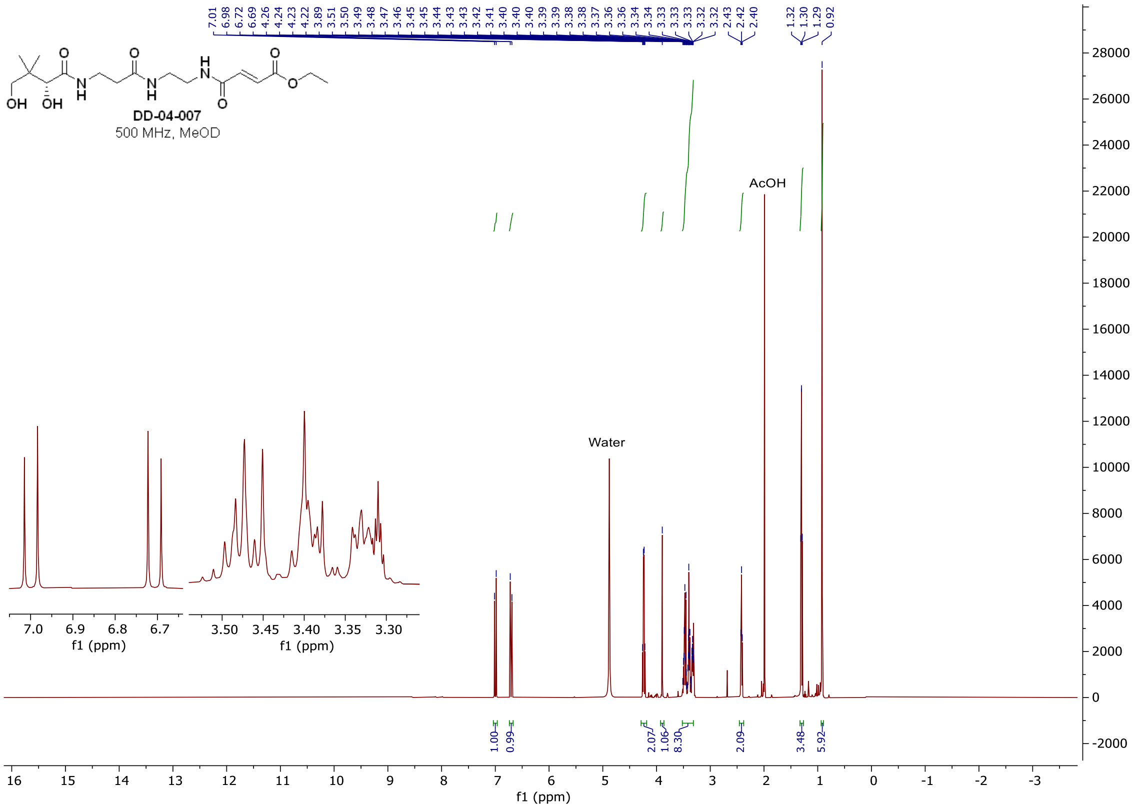

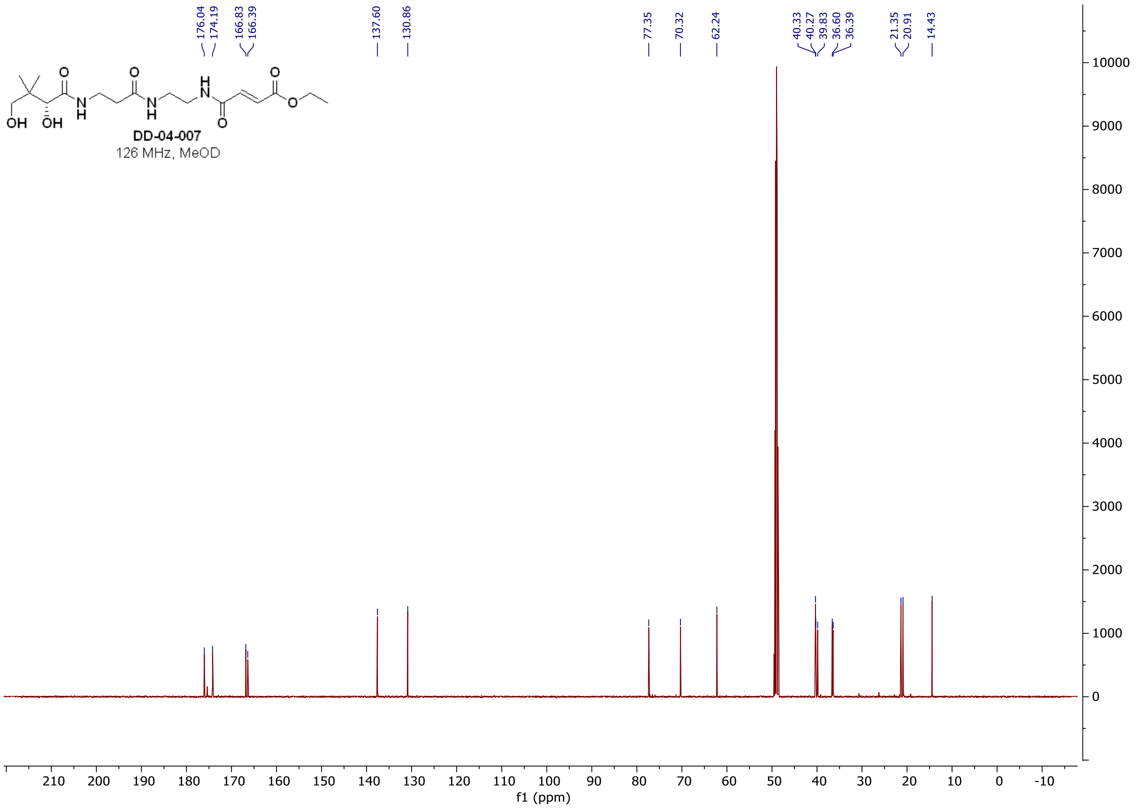

#### [1]H and [13]C spectra of DD-03-140

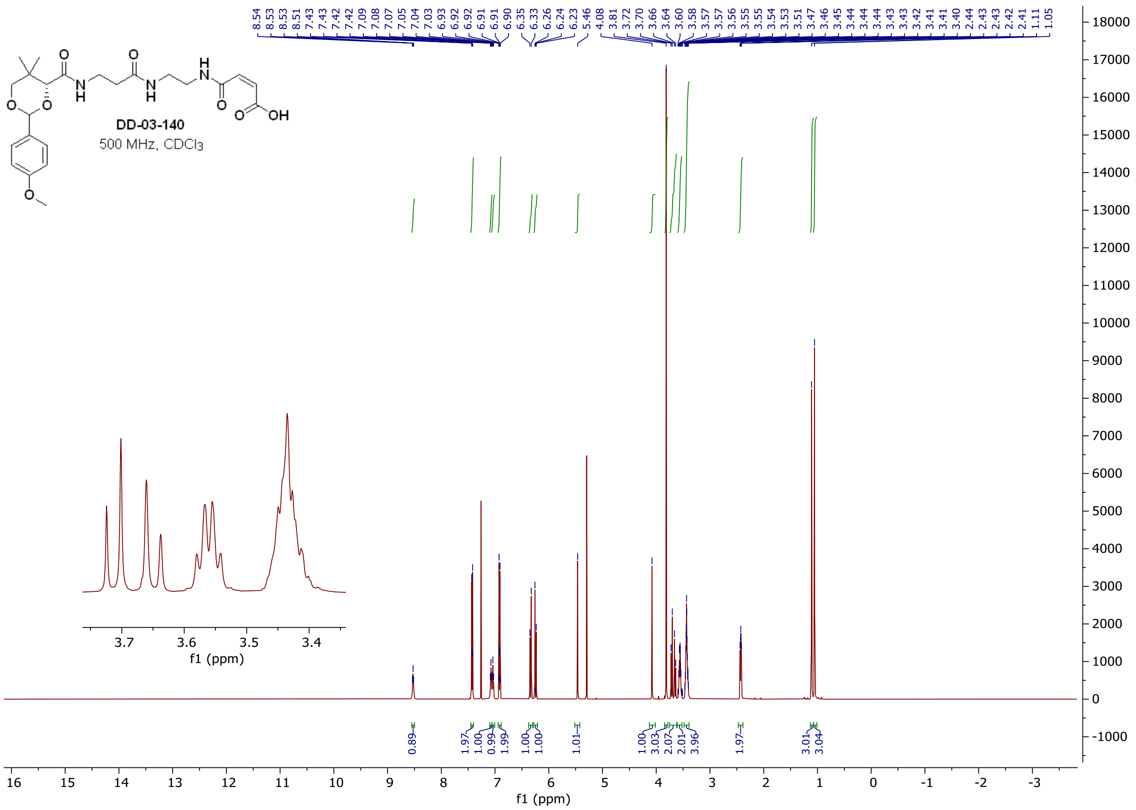

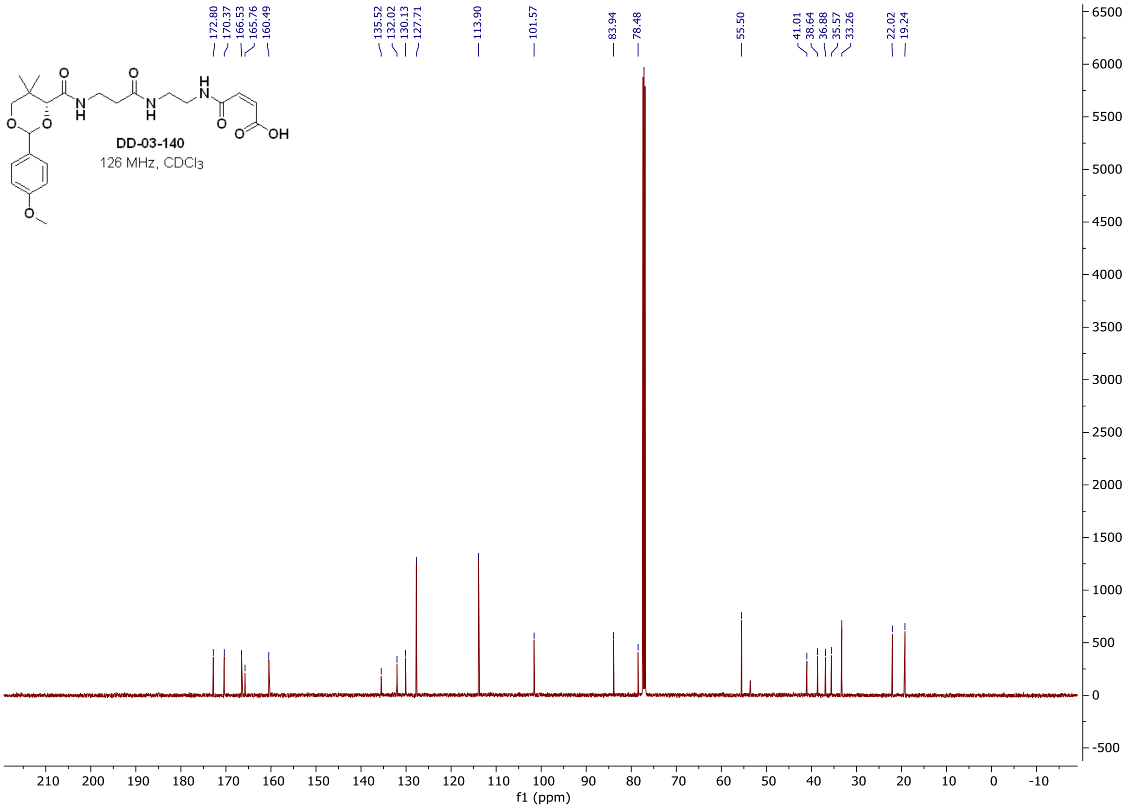

#### $^{1}$H and $^{13}$C **spectra of DD-07-129**

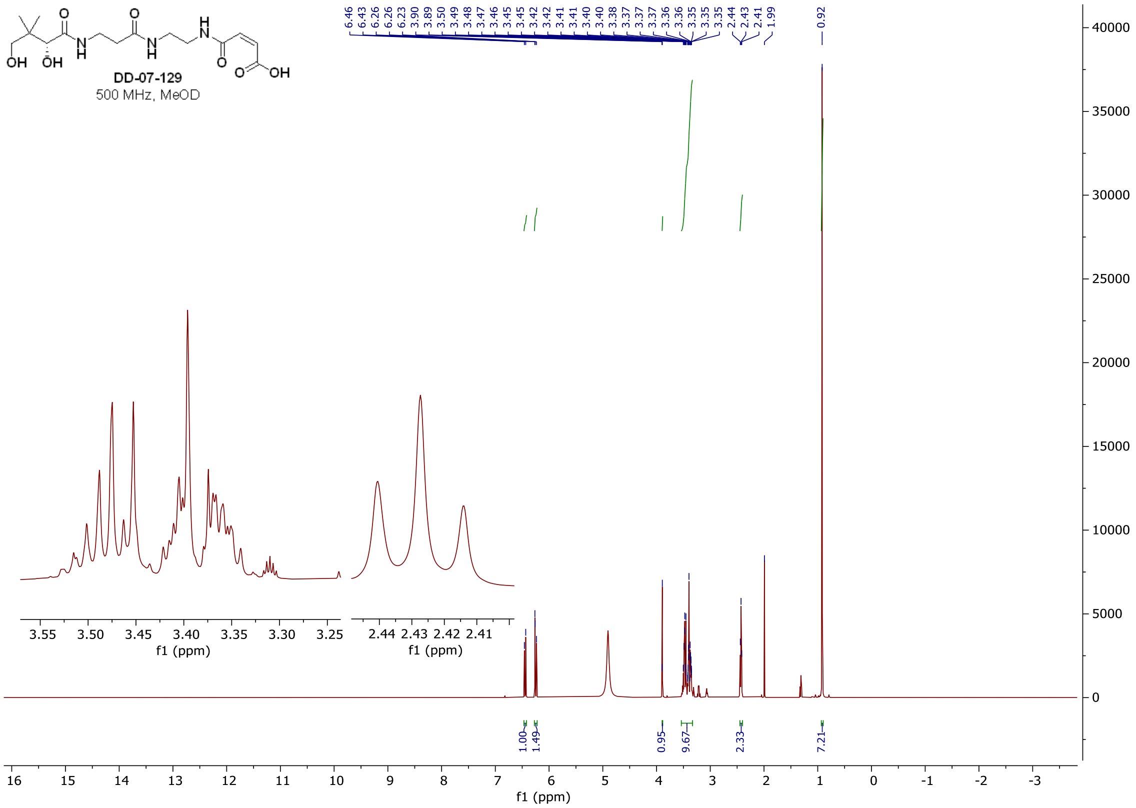

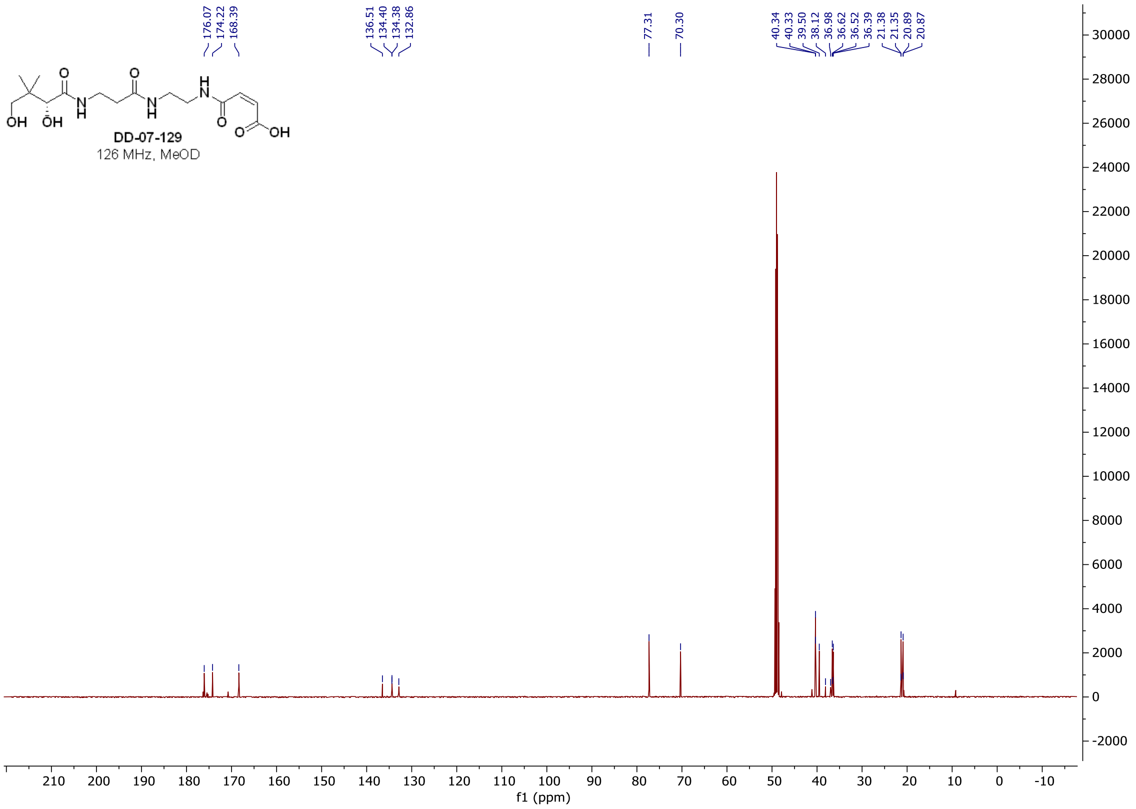

### [1]H and [13]C **spectra of DD-06-119**

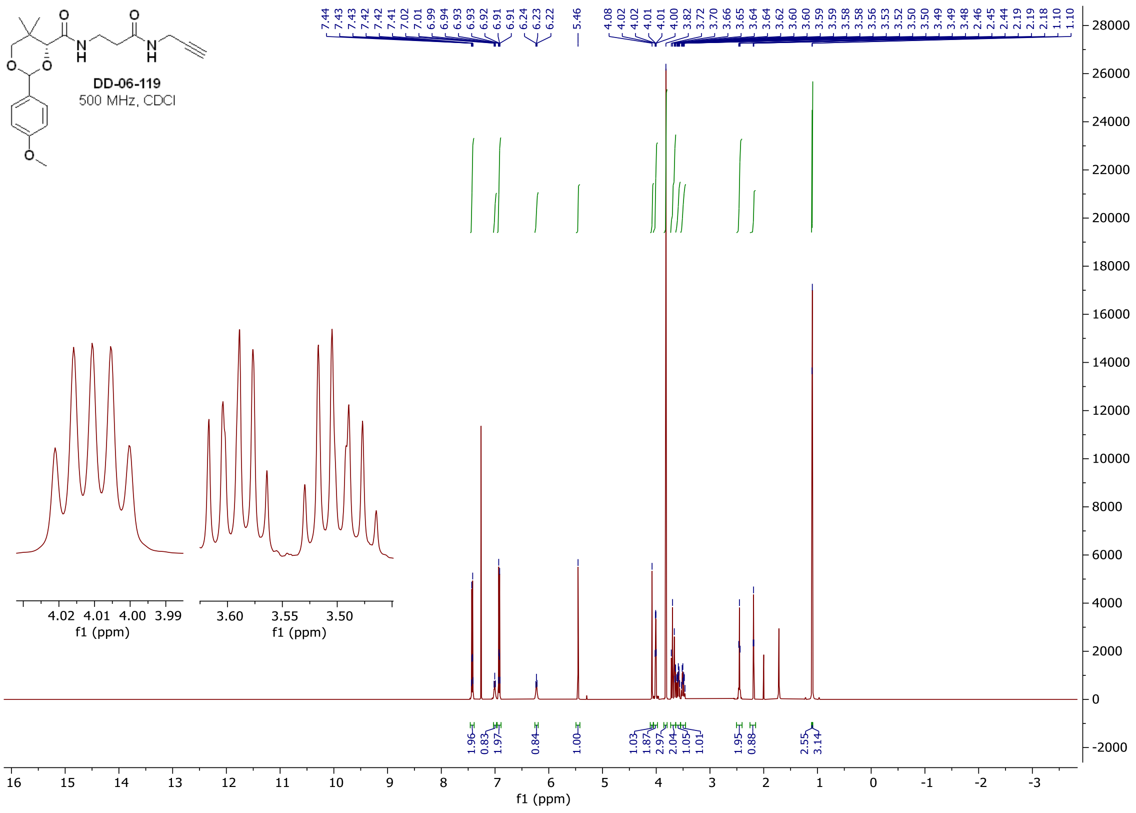

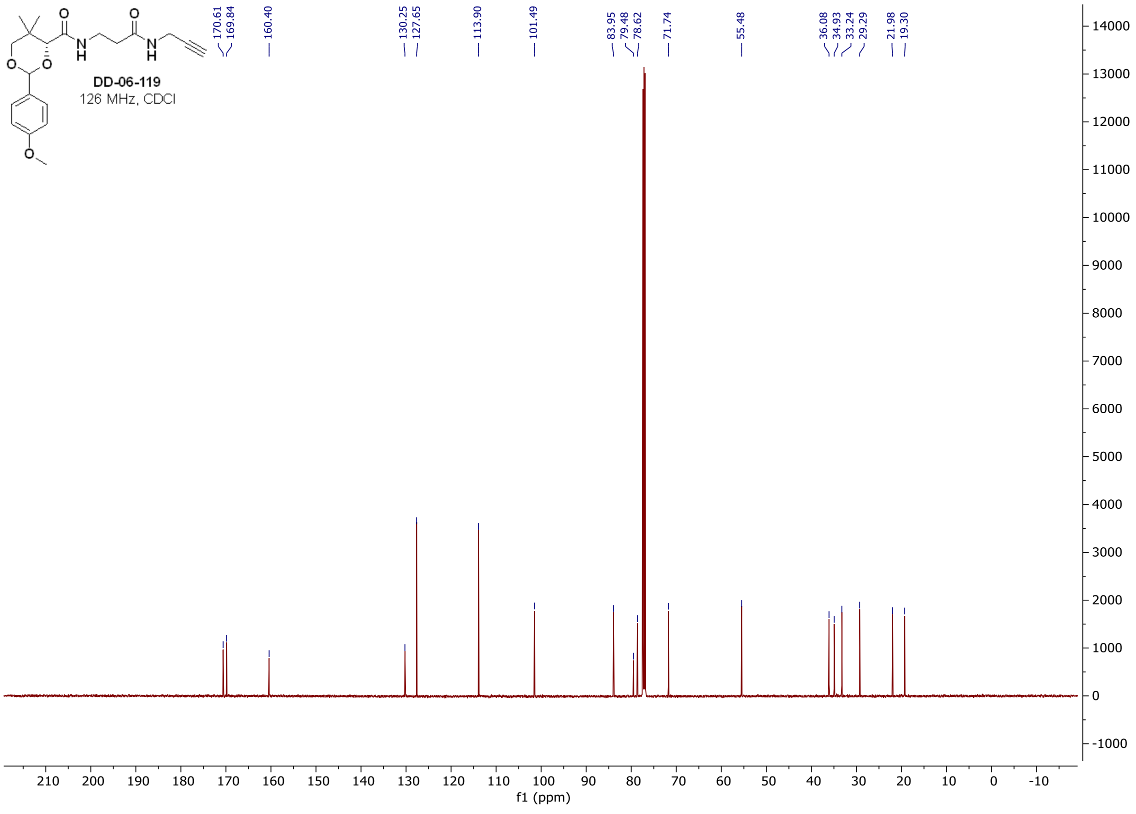

<sup></sup>

$^{1}$H and $^{13}$C **spectra of DD-07-037**

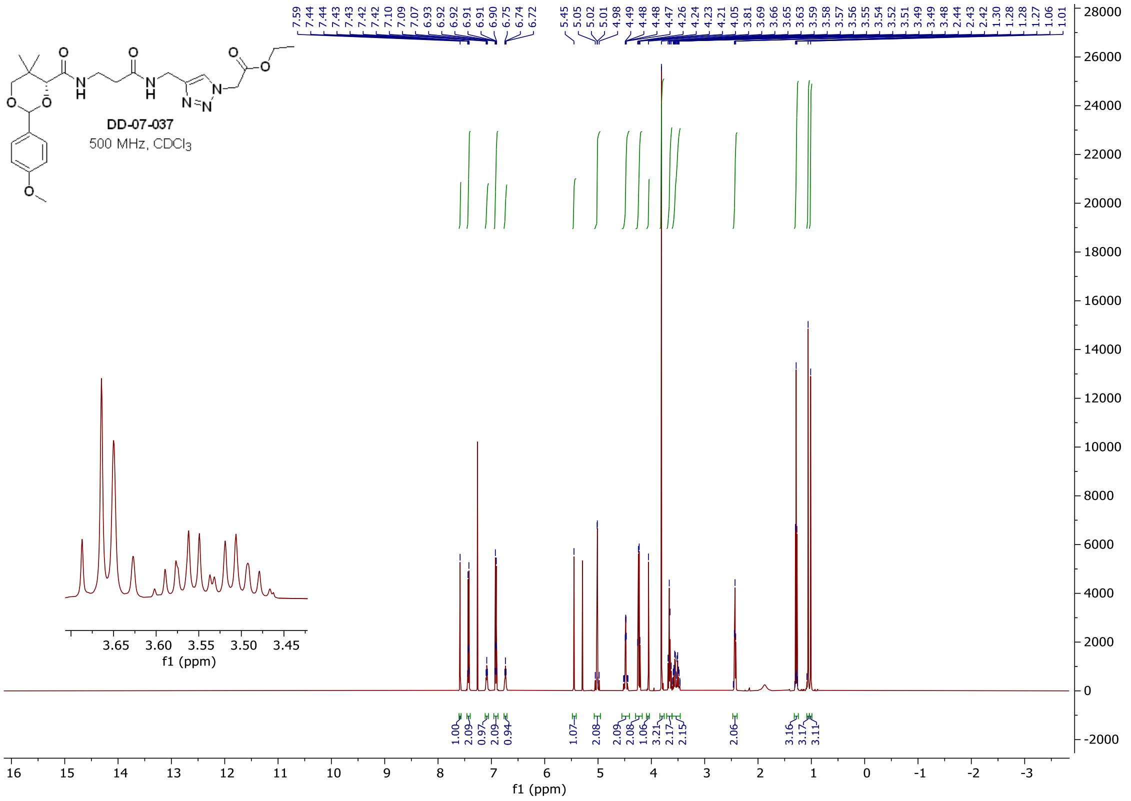

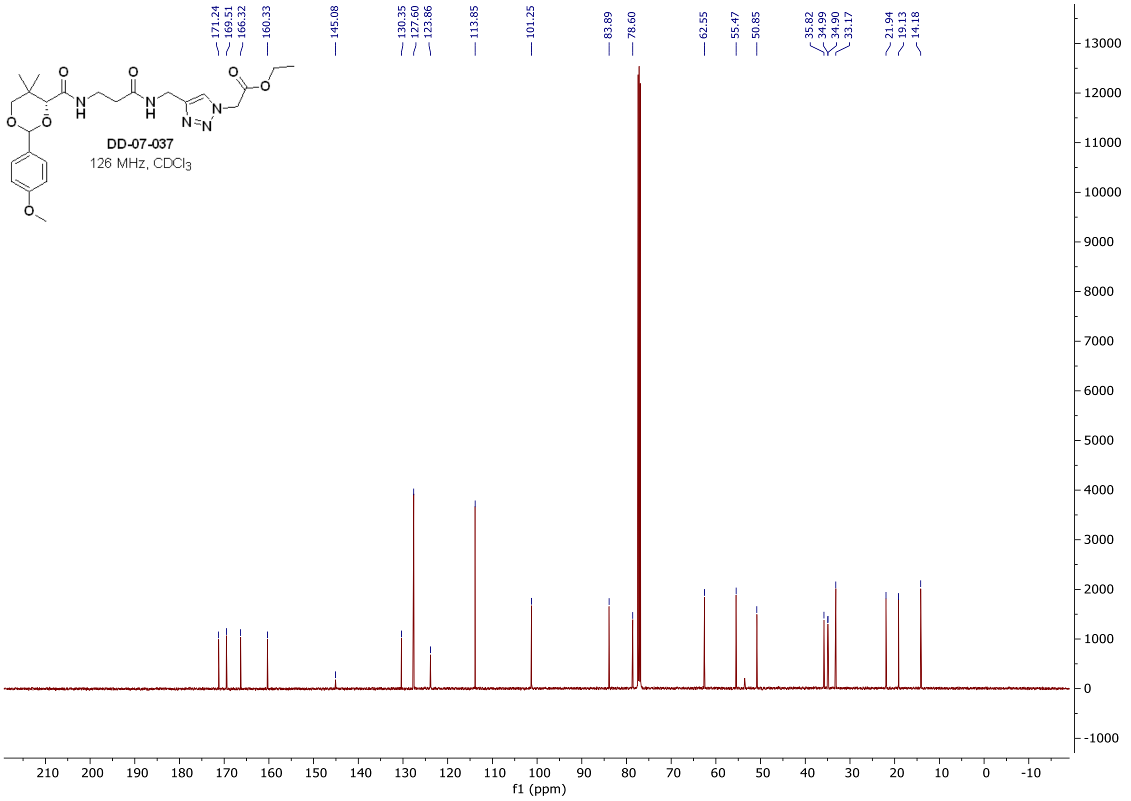

#### $^{1}$H and $^{13}$C spectra of DD-07-141

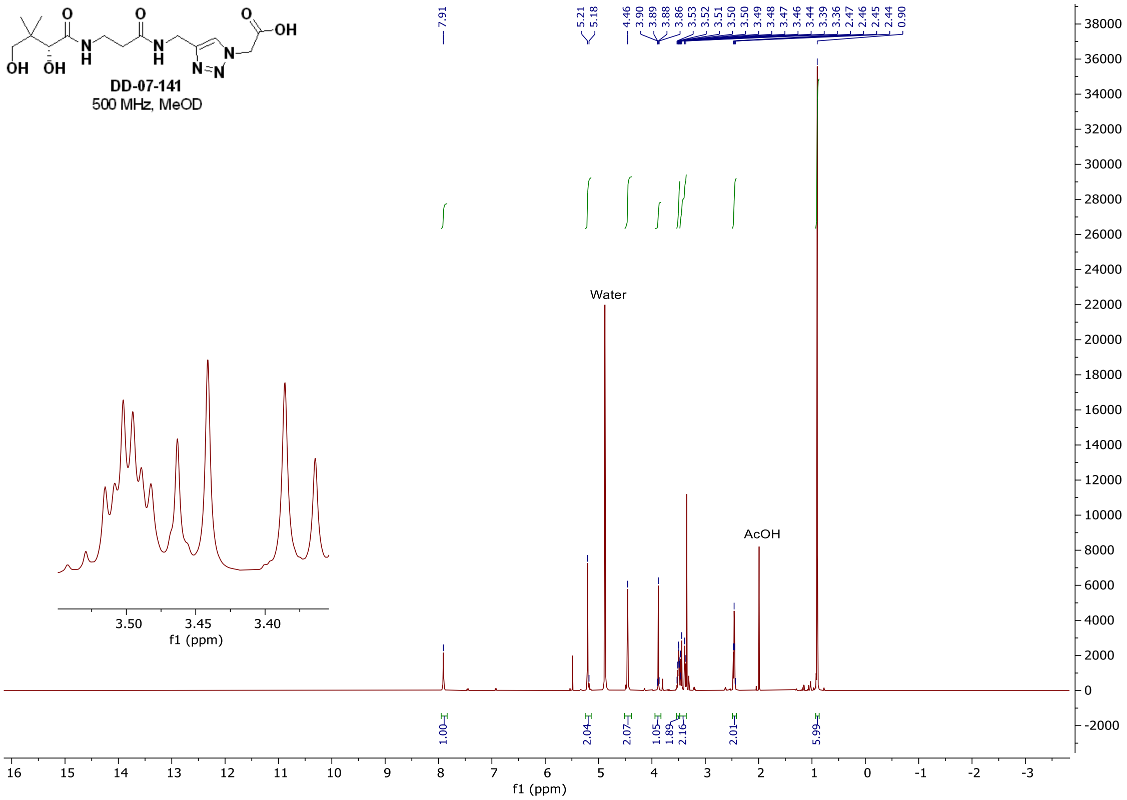

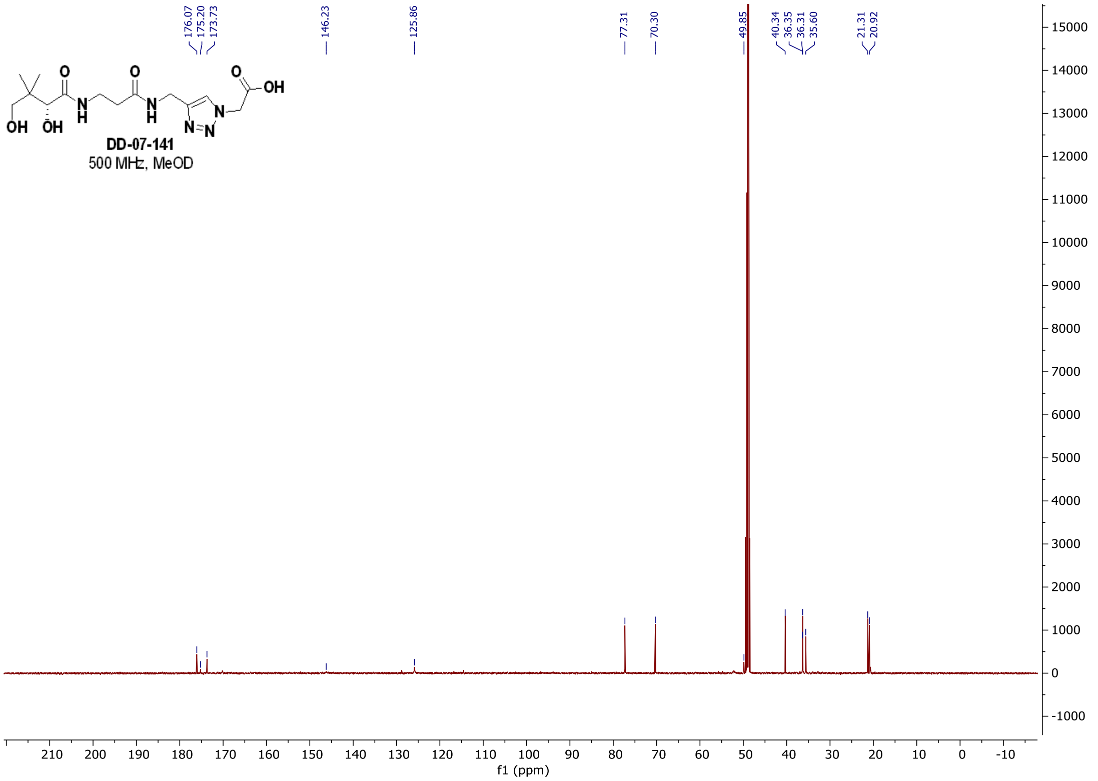

#### $^{1}$H and $^{13}$C **spectra of DD-02-026**

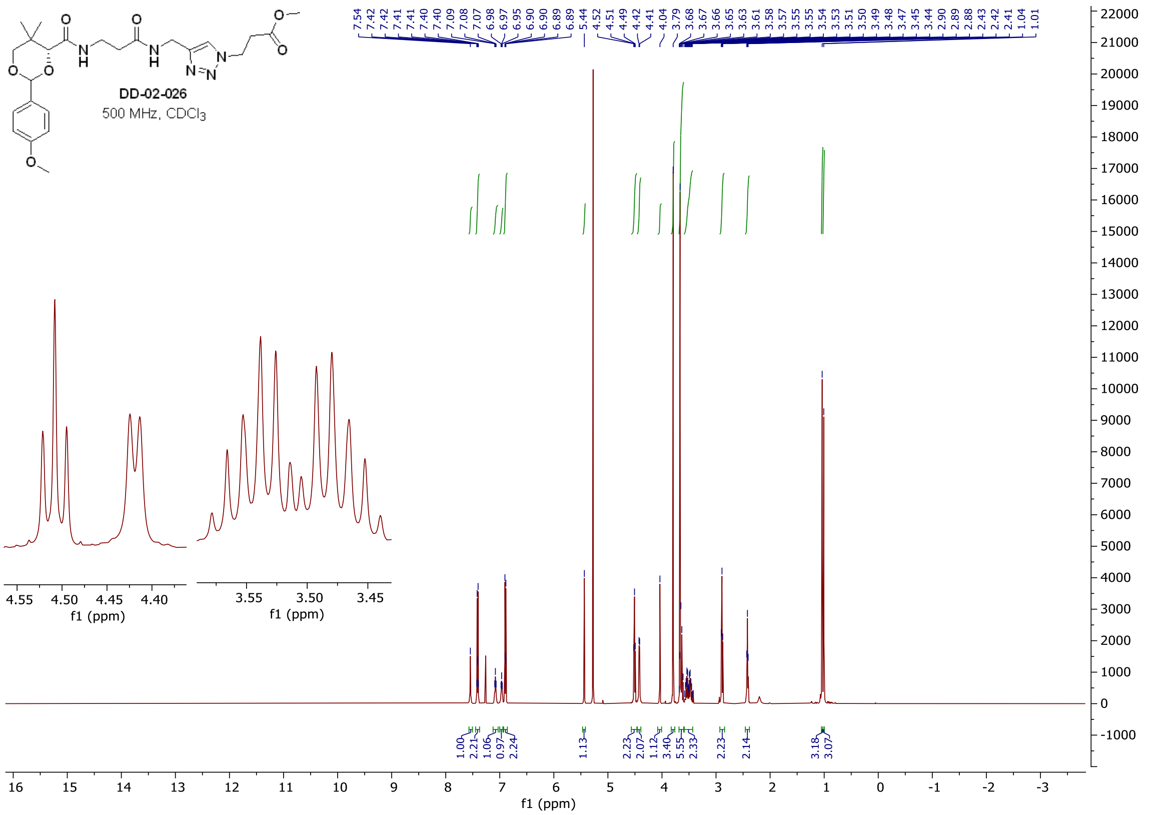

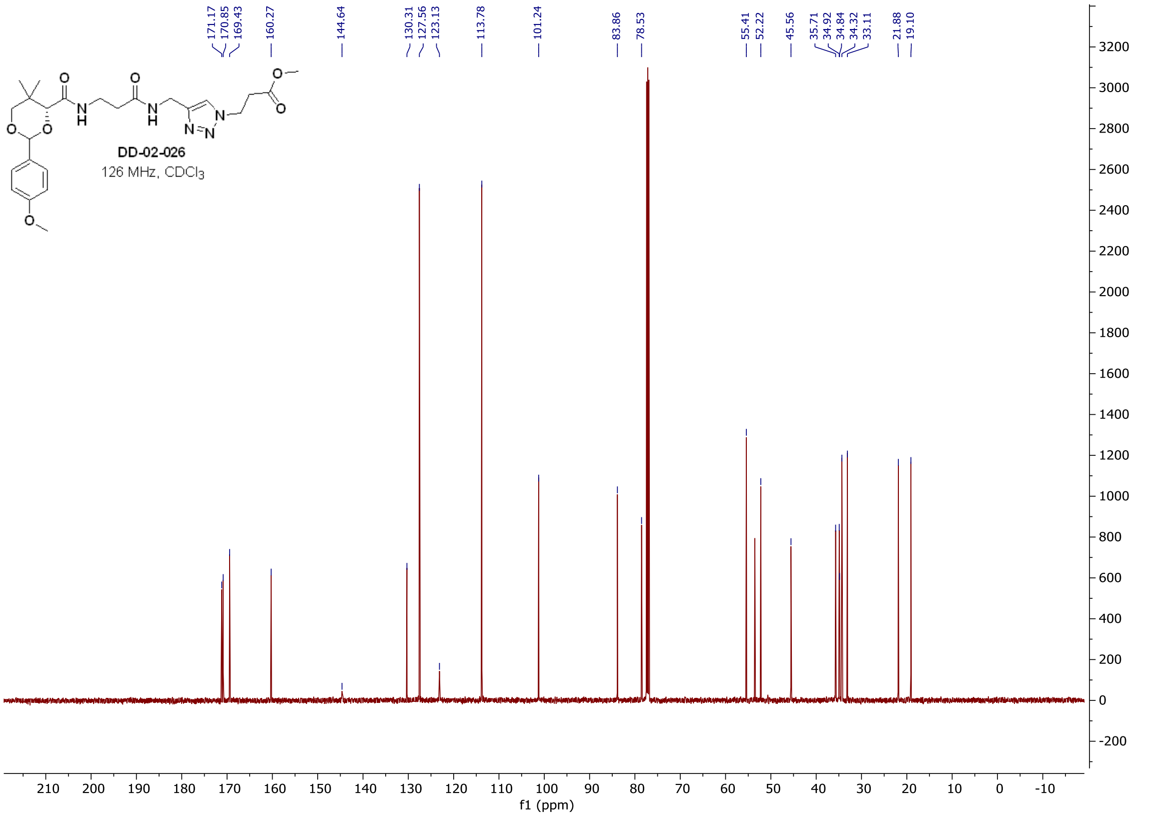

#### $^{1}H$ and $^{13}C$ **spectra of DD-08-001**

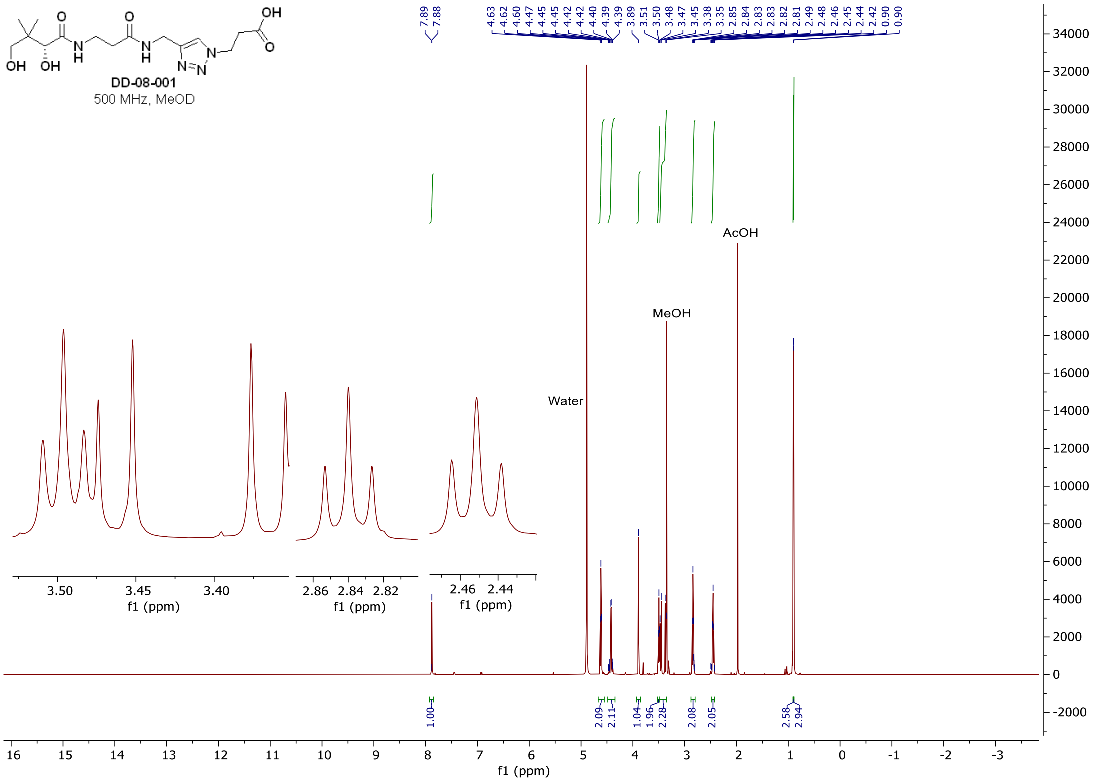

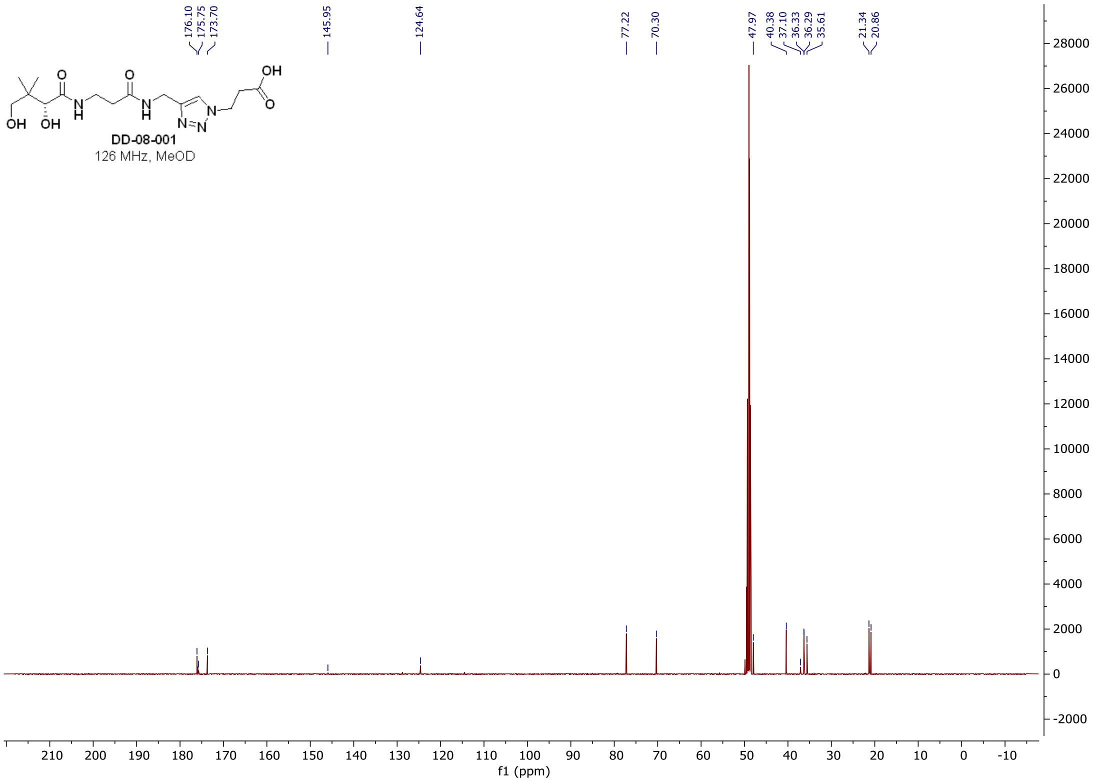

#### $^{1}H$ and $^{13}C$ **spectra of DD-07-081**

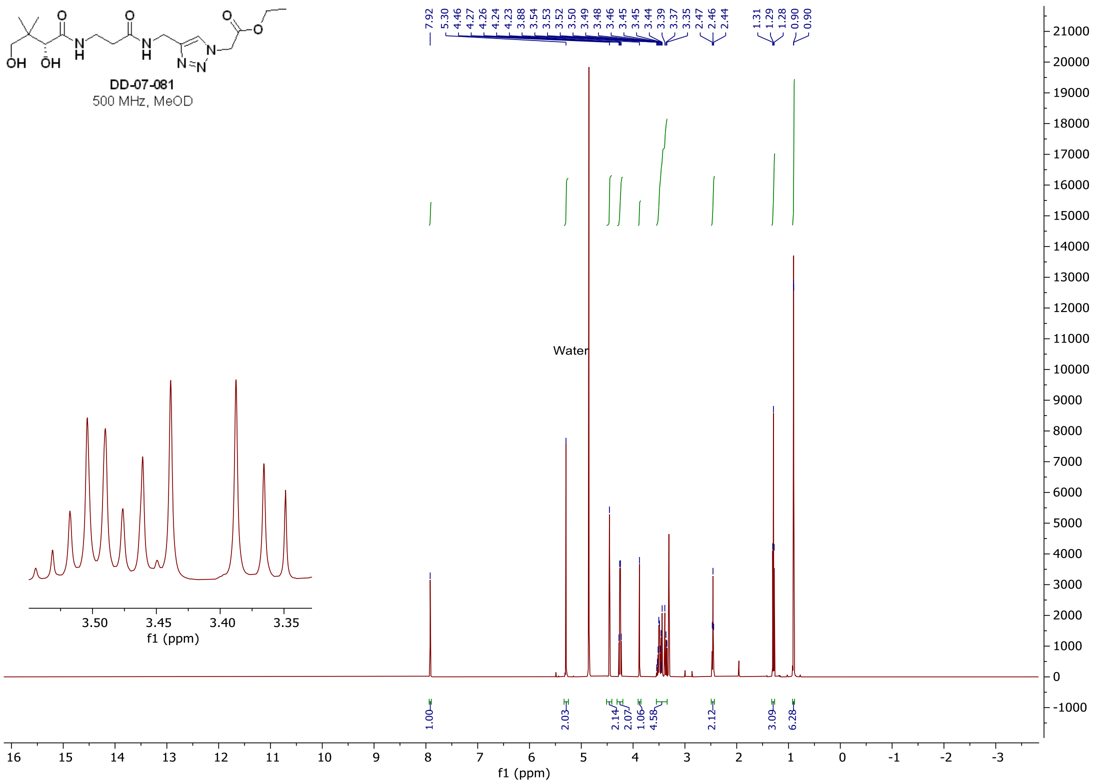

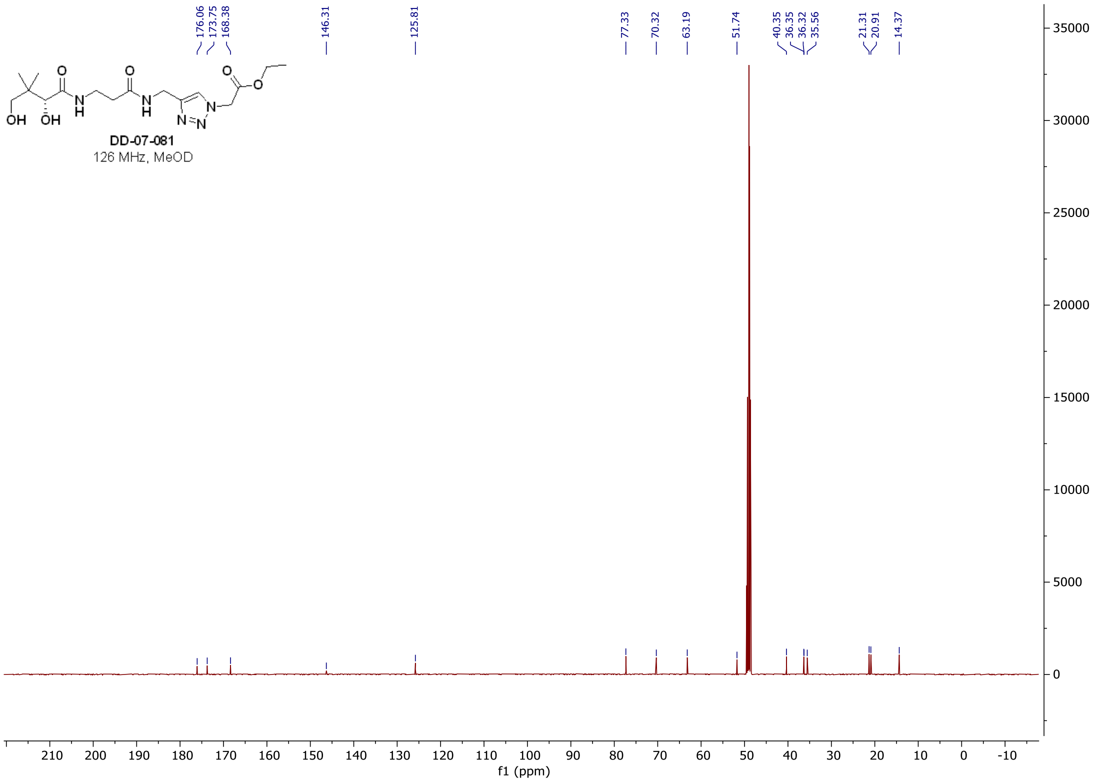

#### [1]H and [13]C **spectra of DD-07-015**

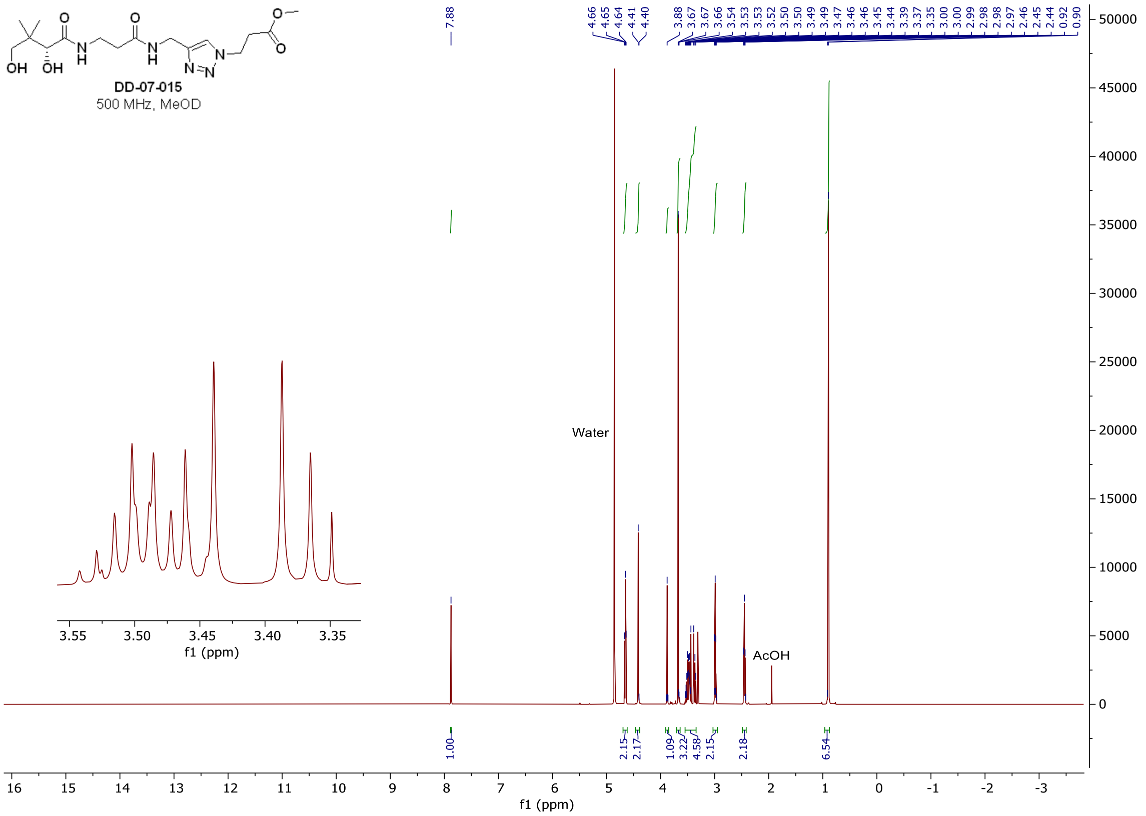

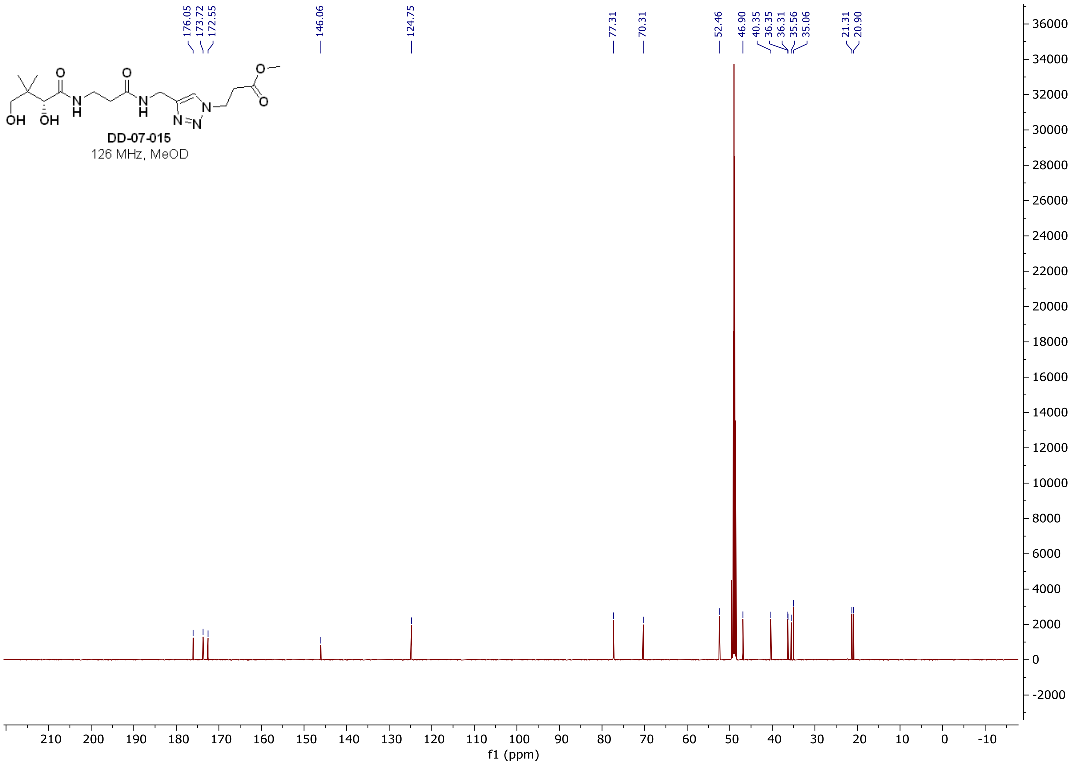

#### $^1H$ and $^{13}C$ spectra of DD-07-125

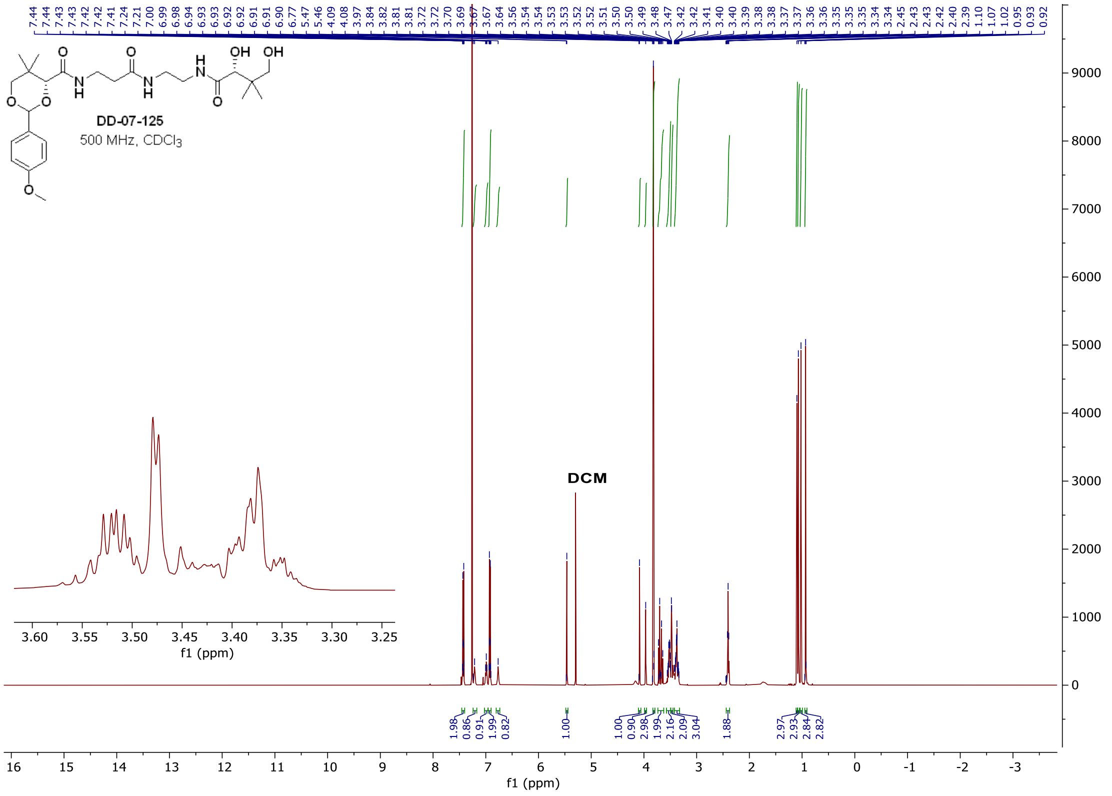

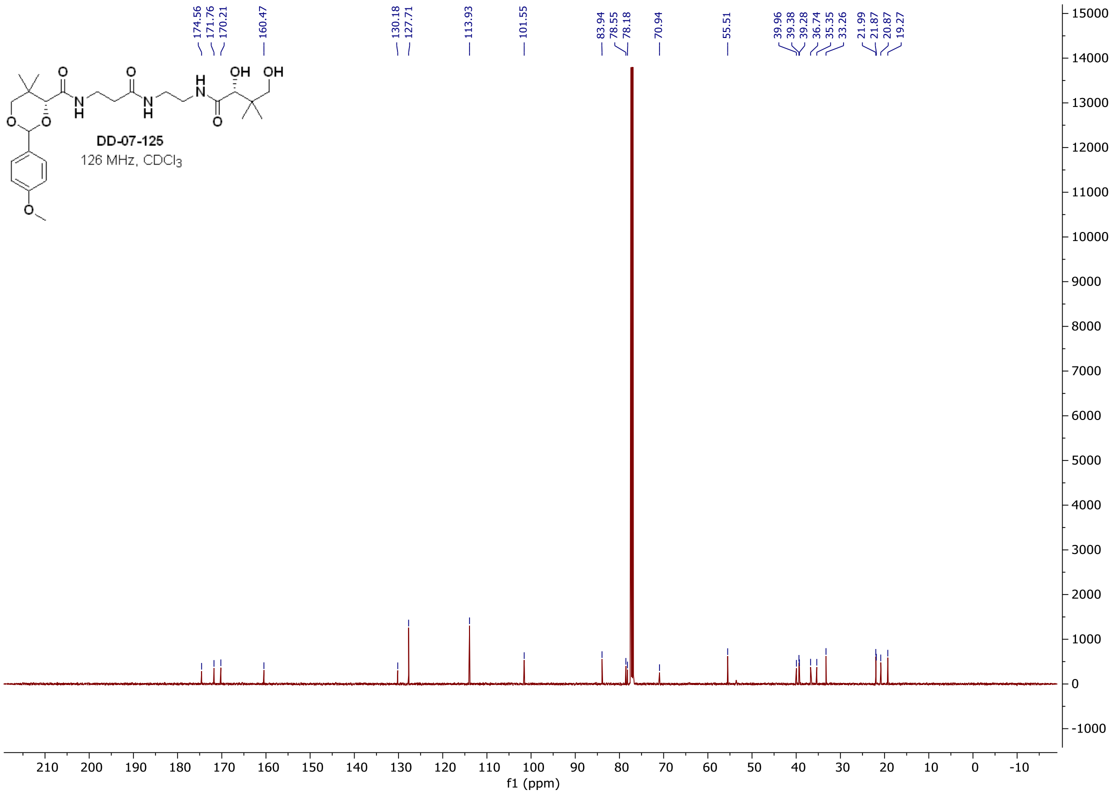

#### $^{1}H$ and $^{13}C$ spectra of DD-07-134

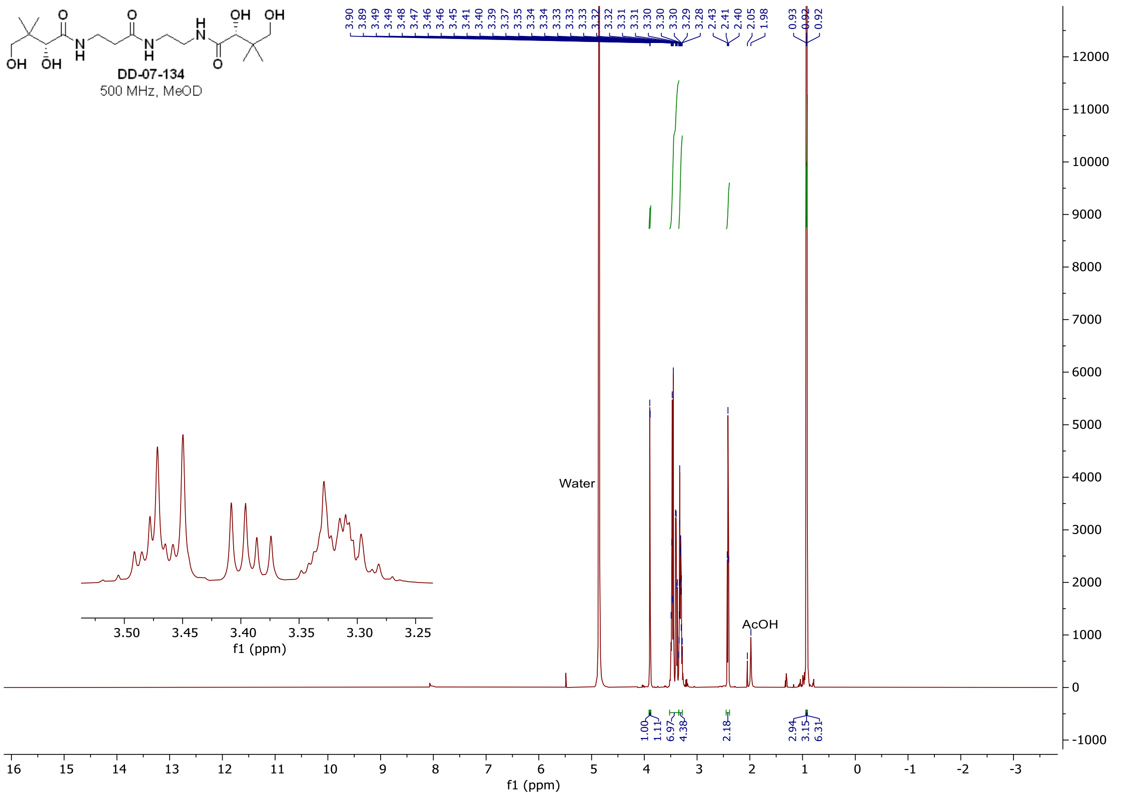

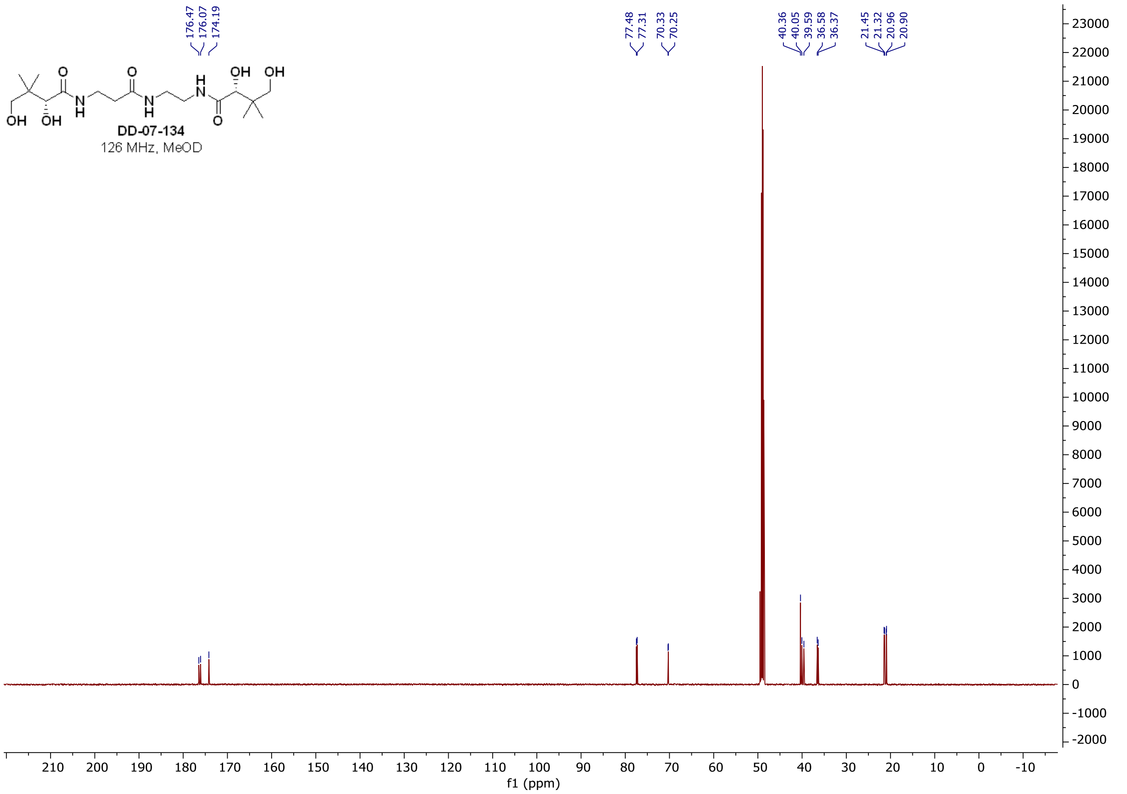

#### $^{1}H$ and $^{13}C$ **spectra of DD-05-078**

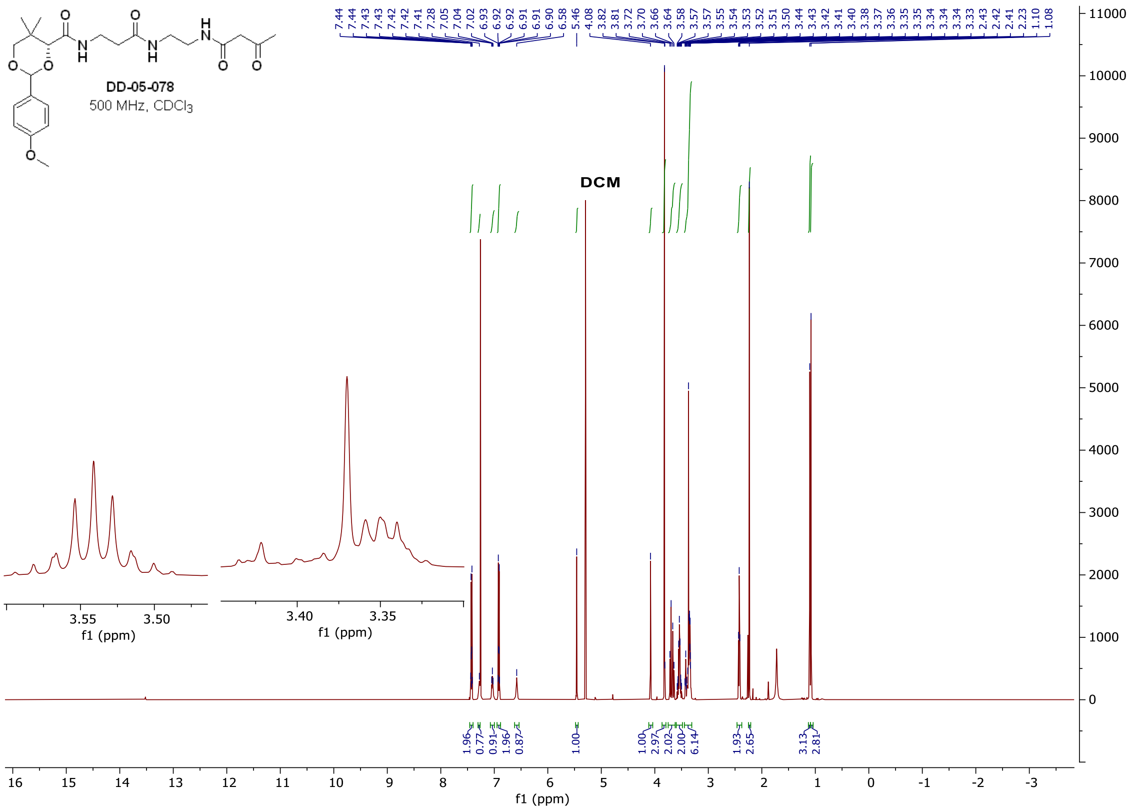

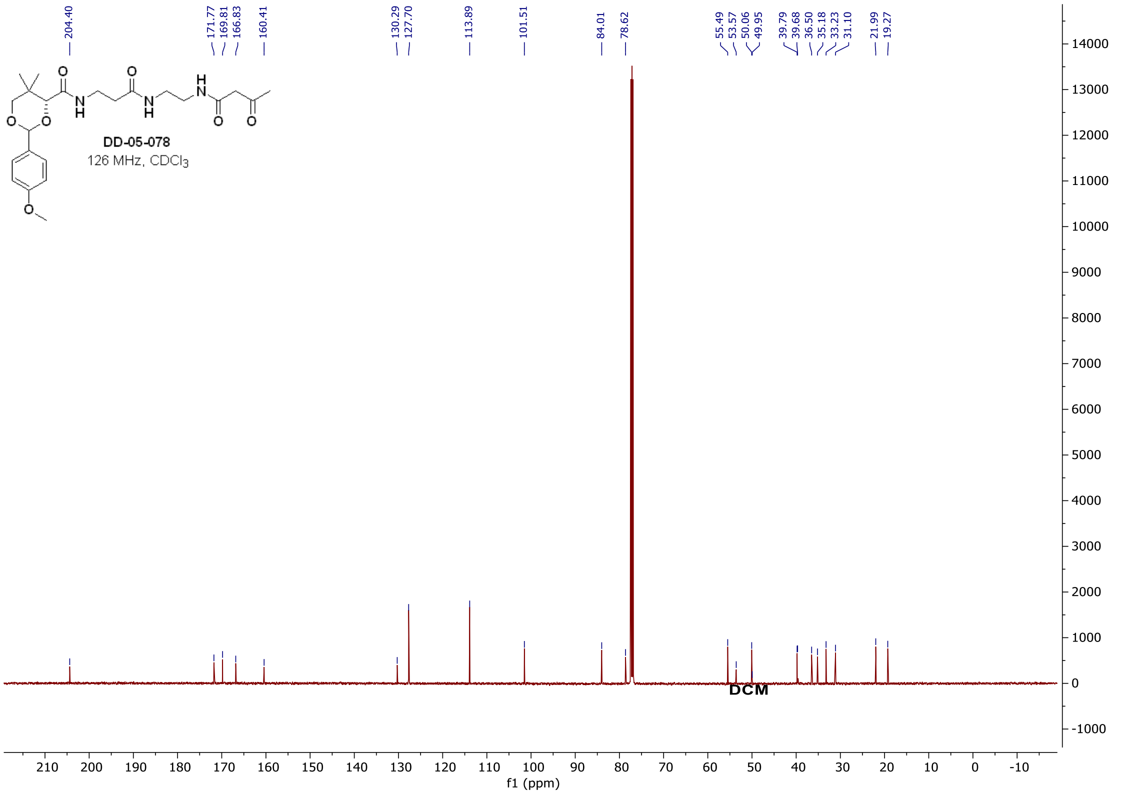

$^{1}$H and $^{13}$C **spectra of DD-07-148**

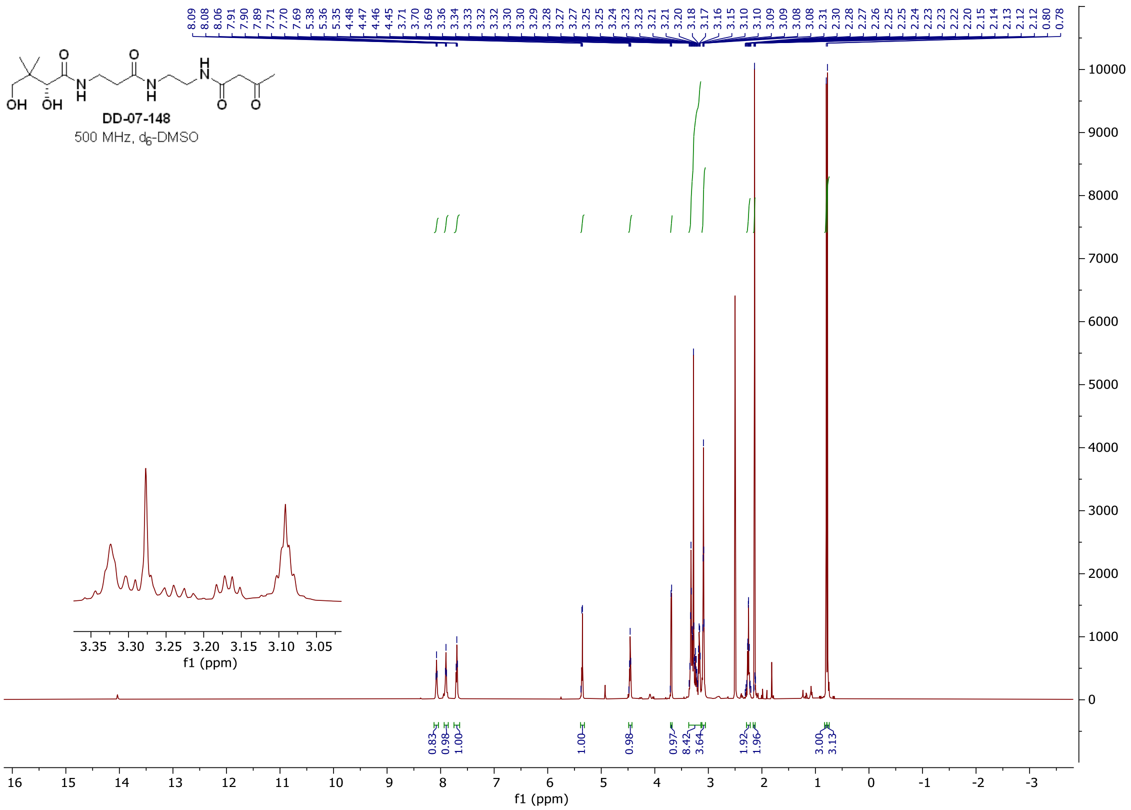

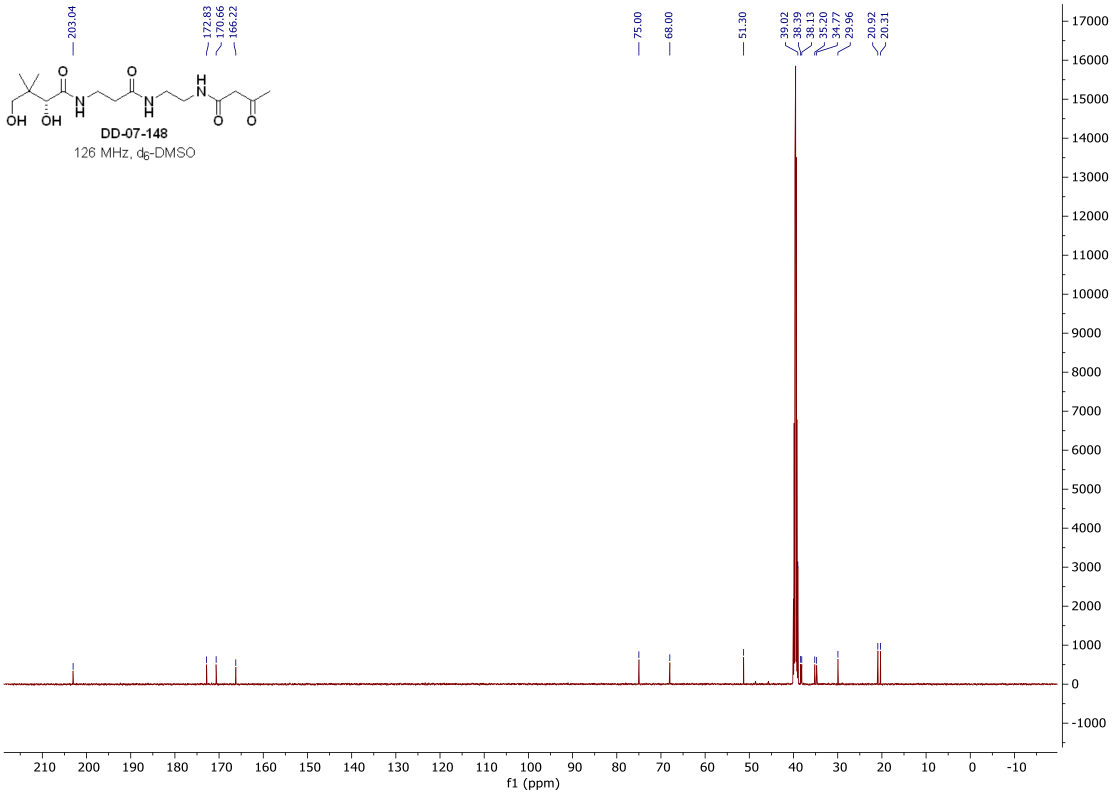

### $^{1}$H and $^{13}$C **spectra of DD-07-115**

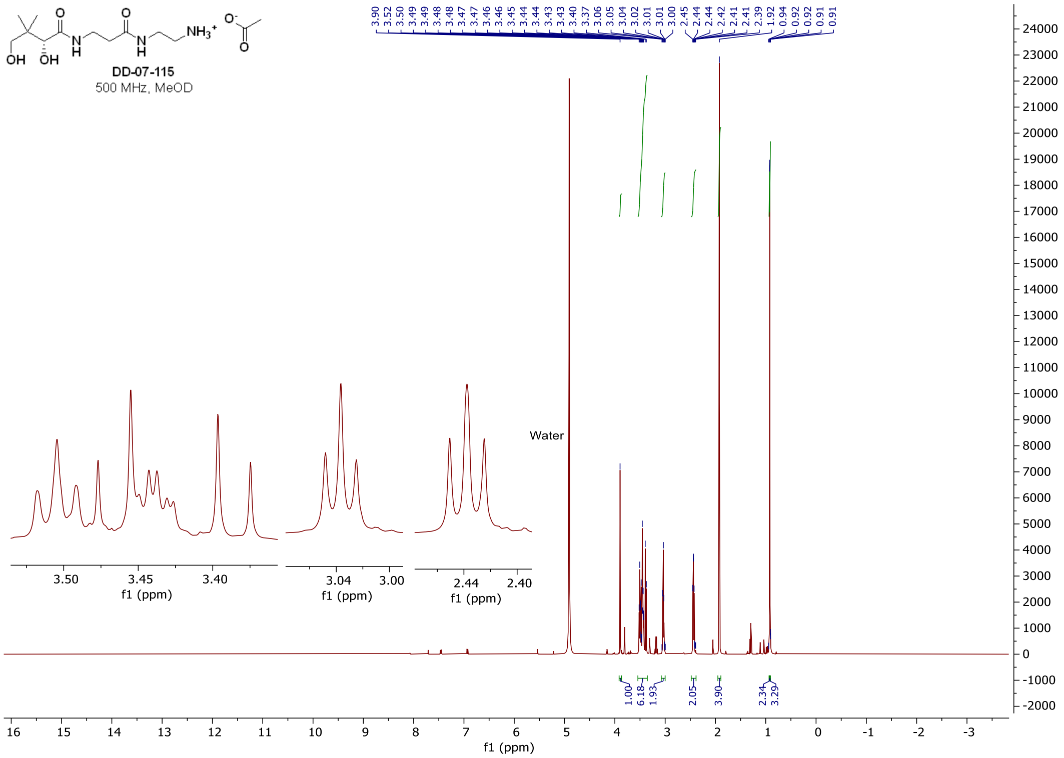

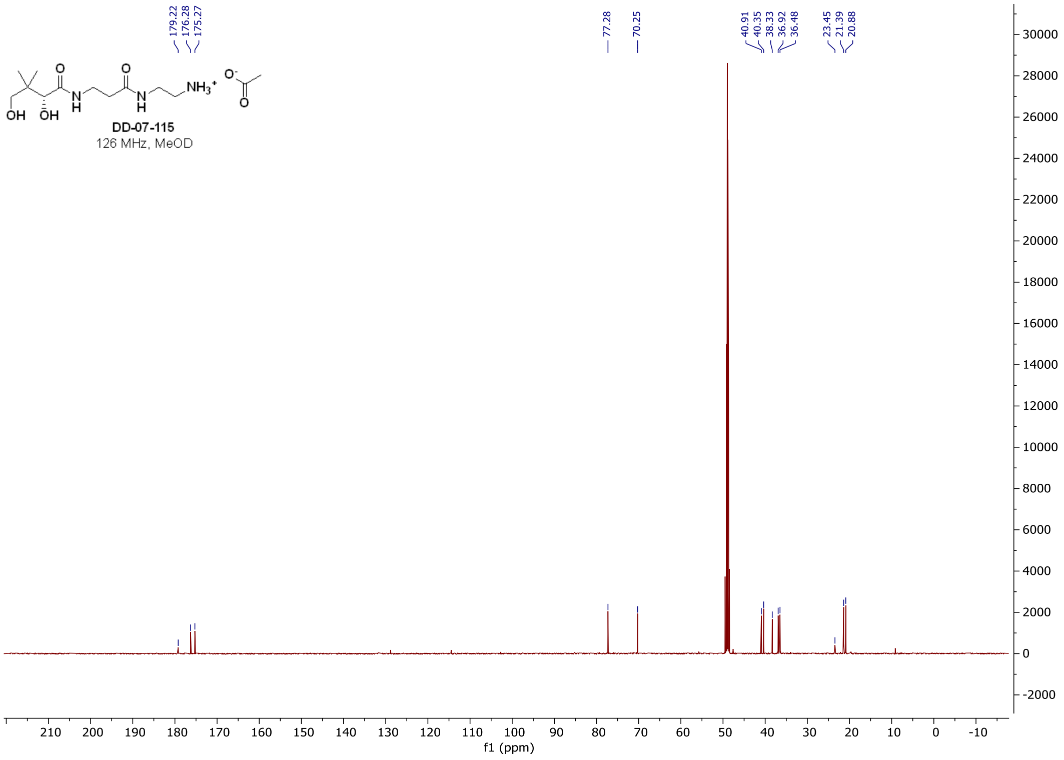

#### $^1$H and $^{13}$C **spectra of DD-06-106**

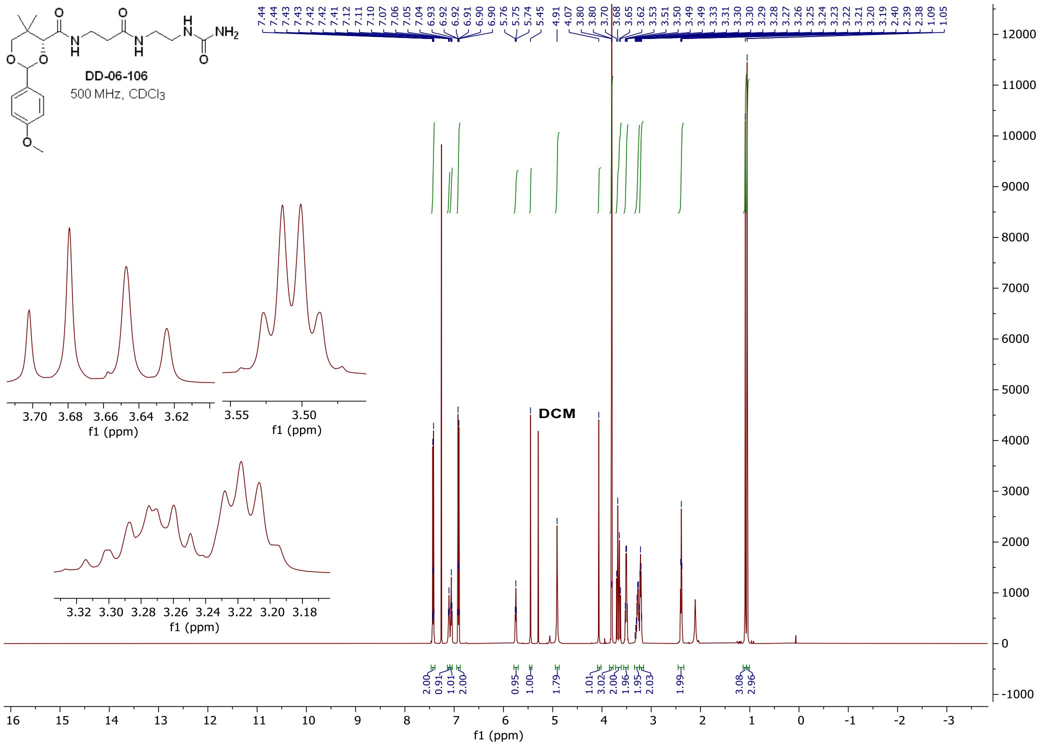

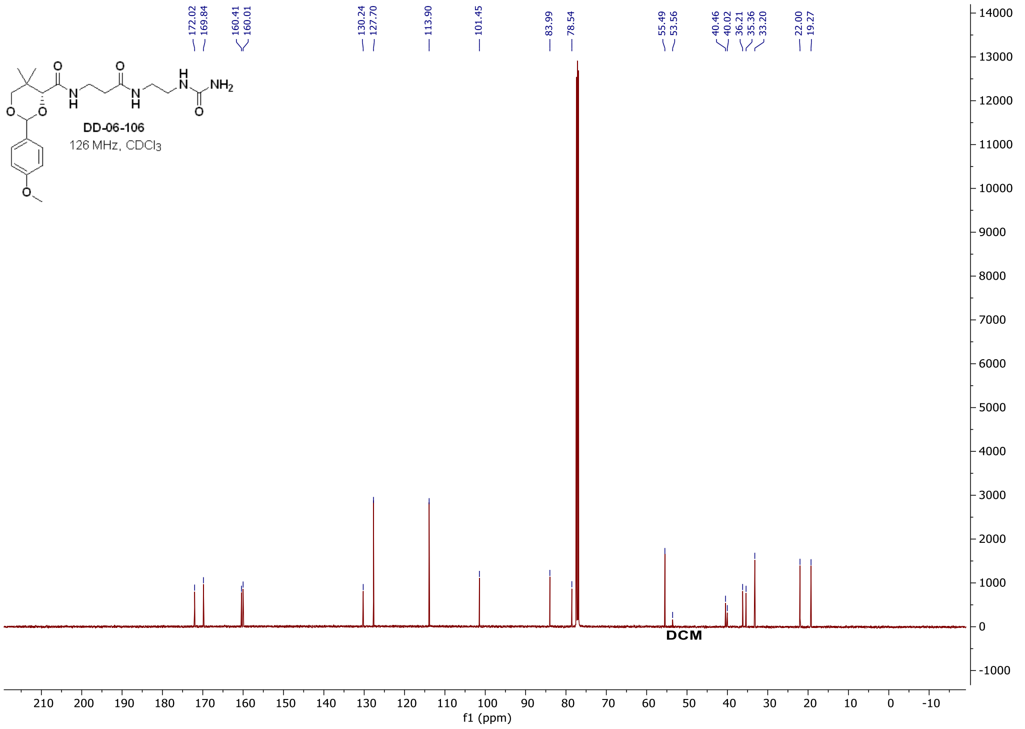

$^{1}H$ and $^{13}C$ **spectra of DD-06-126**

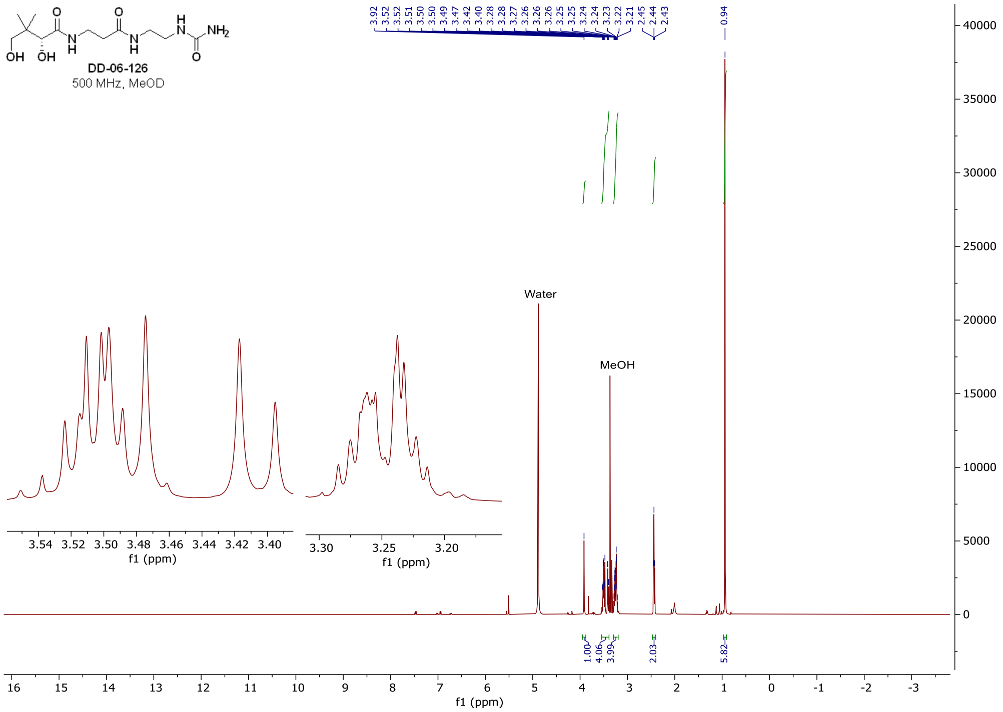

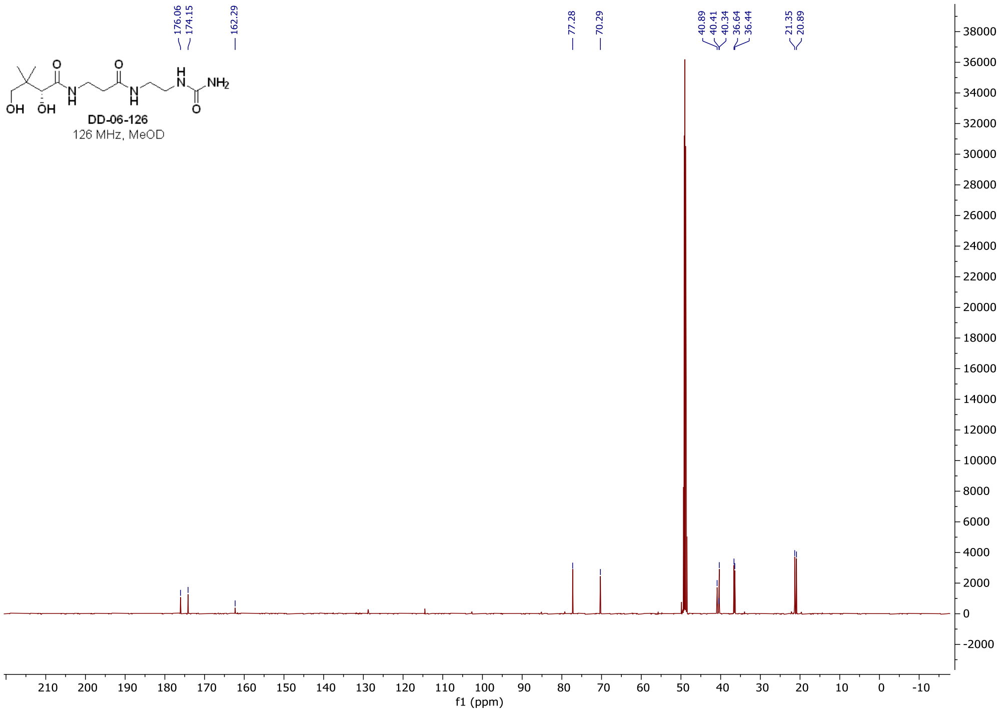

#### [1]H NMR spectrum of **4fa**

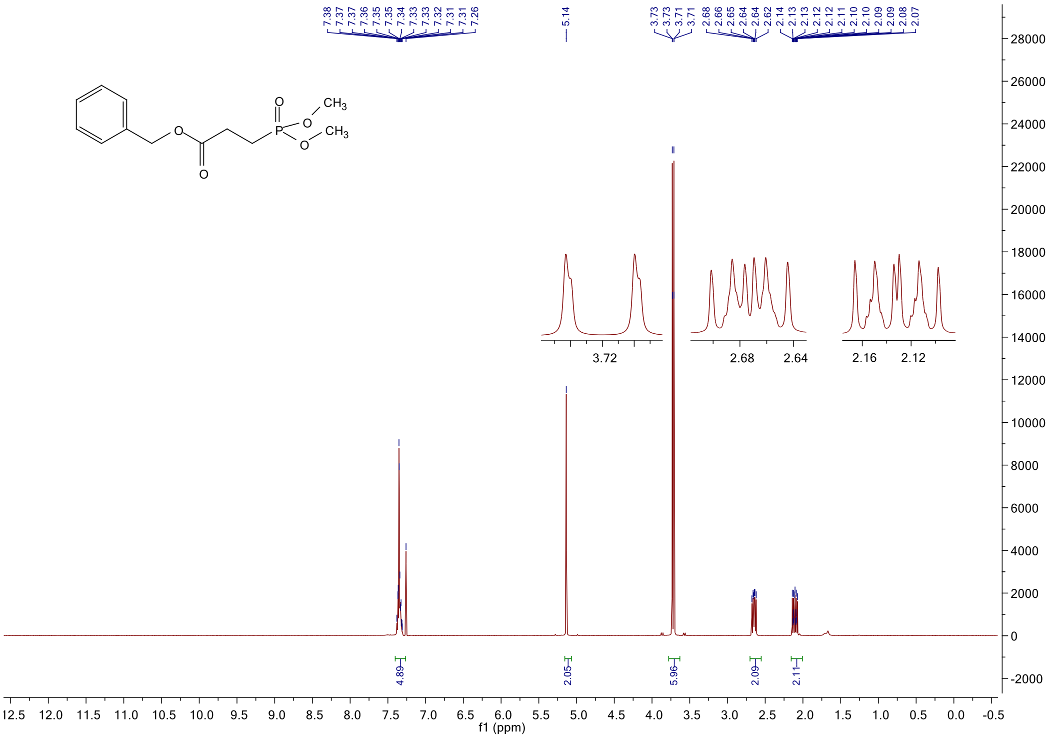

#### [13]C NMR spectrum of **4fa**

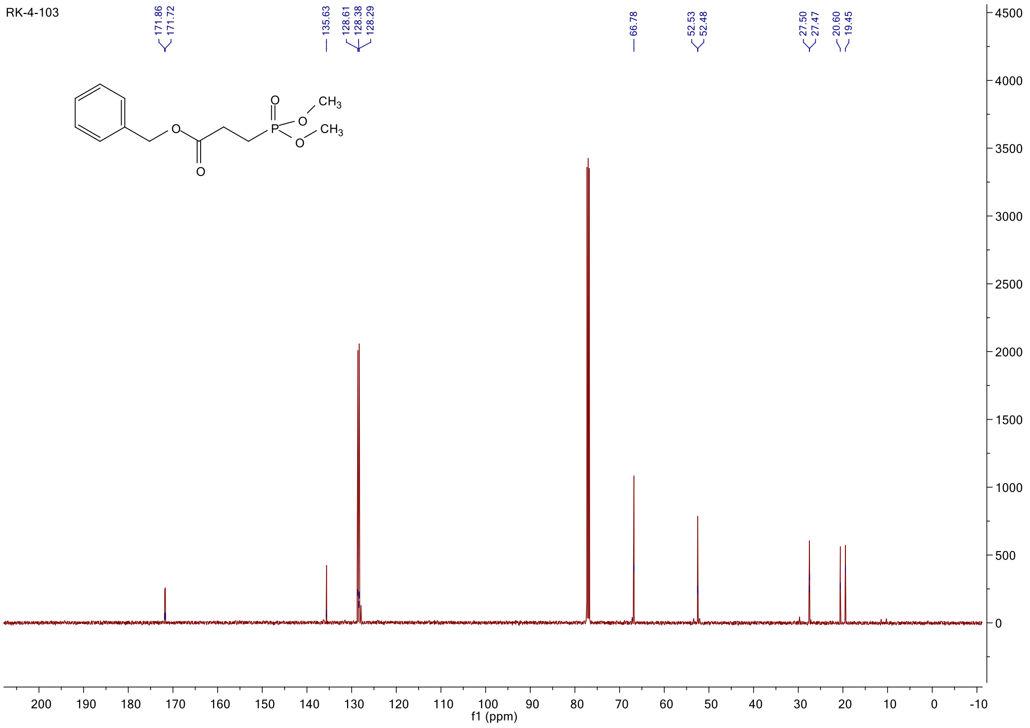

$^{1}$H NMR spectrum of **4f**

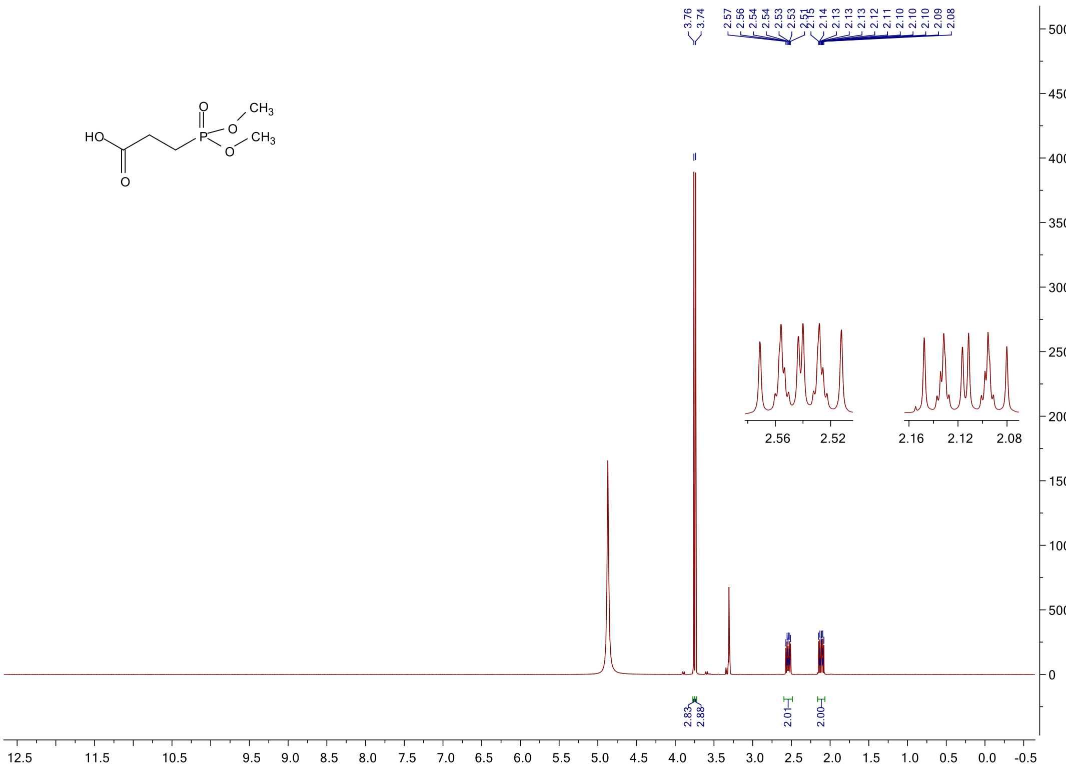

$^{13}$C NMR spectrum of **4f**

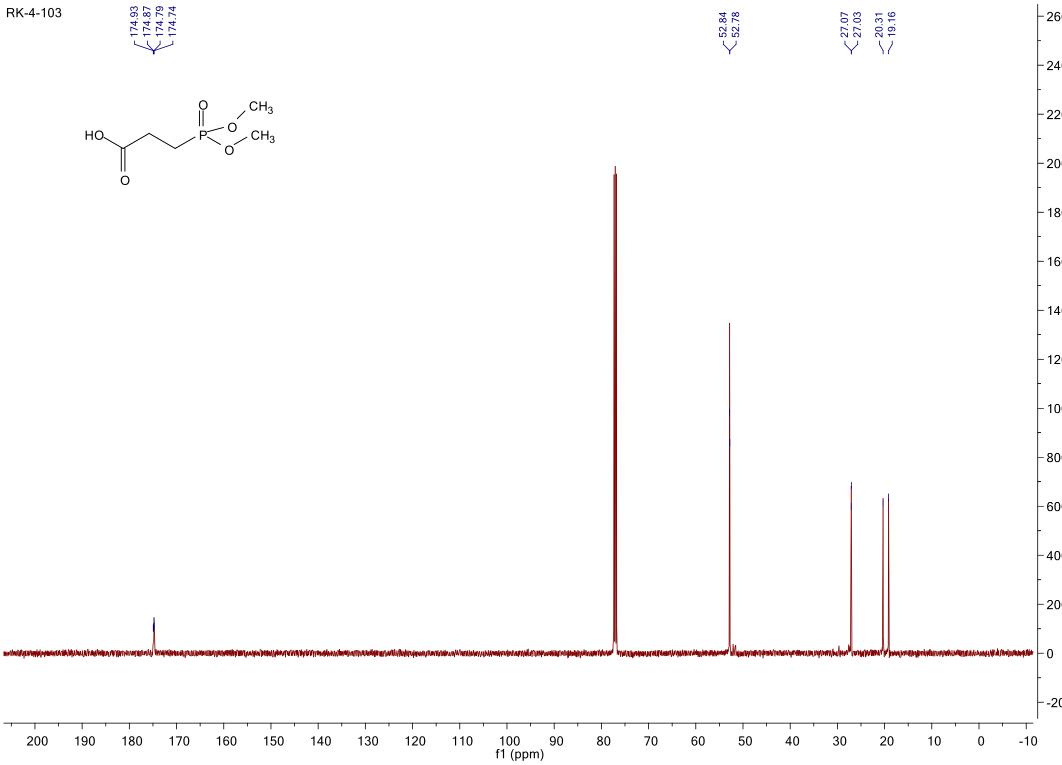

[31]P NMR spectrum of **4f**

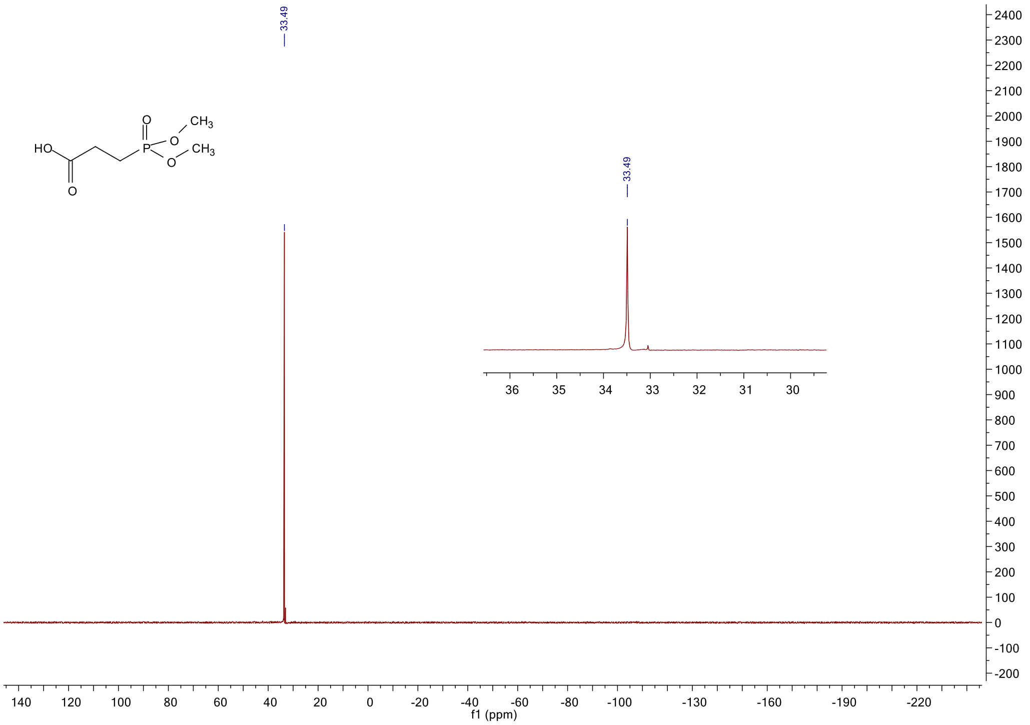

#### $^{1}$H NMR spectrum of **4g**

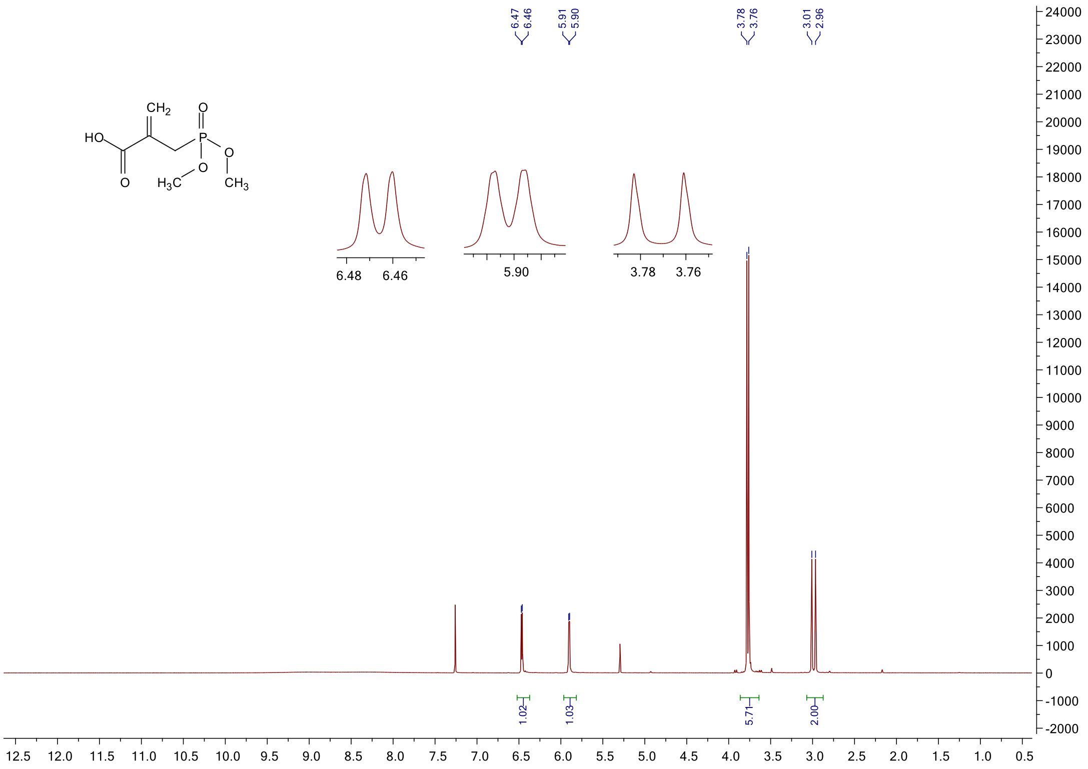

#### $^{31}$P NMR spectrum of **4g**

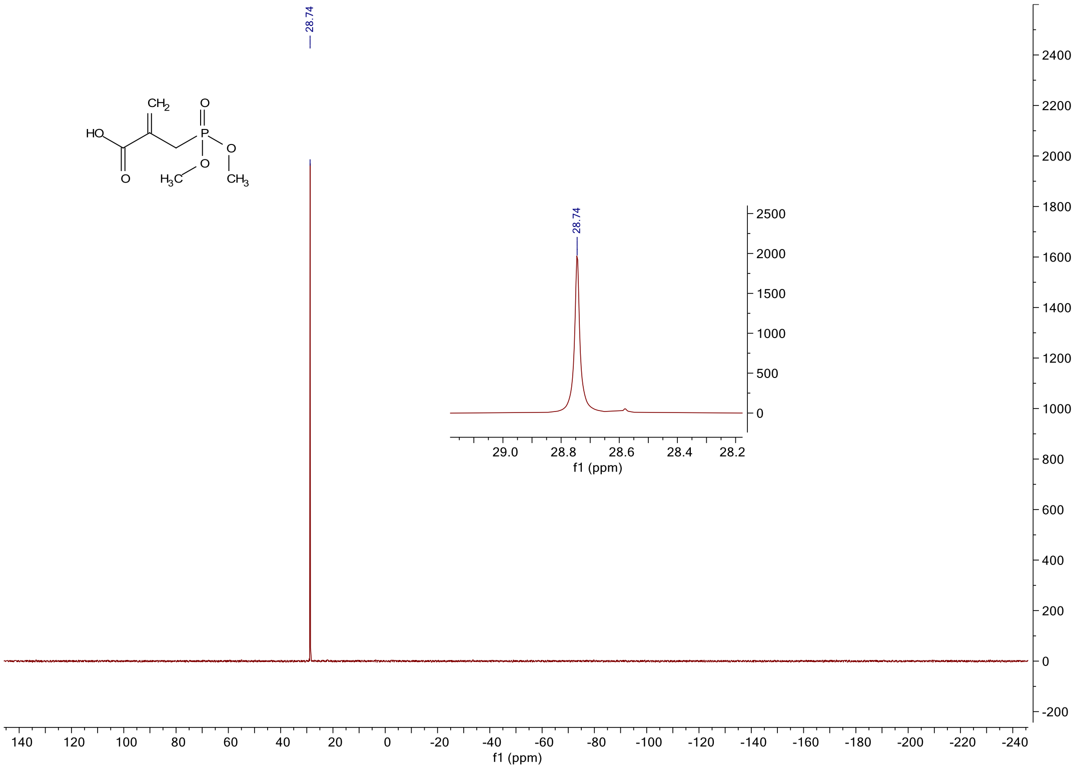

#### $^1$H NMR spectrum of **4ha**

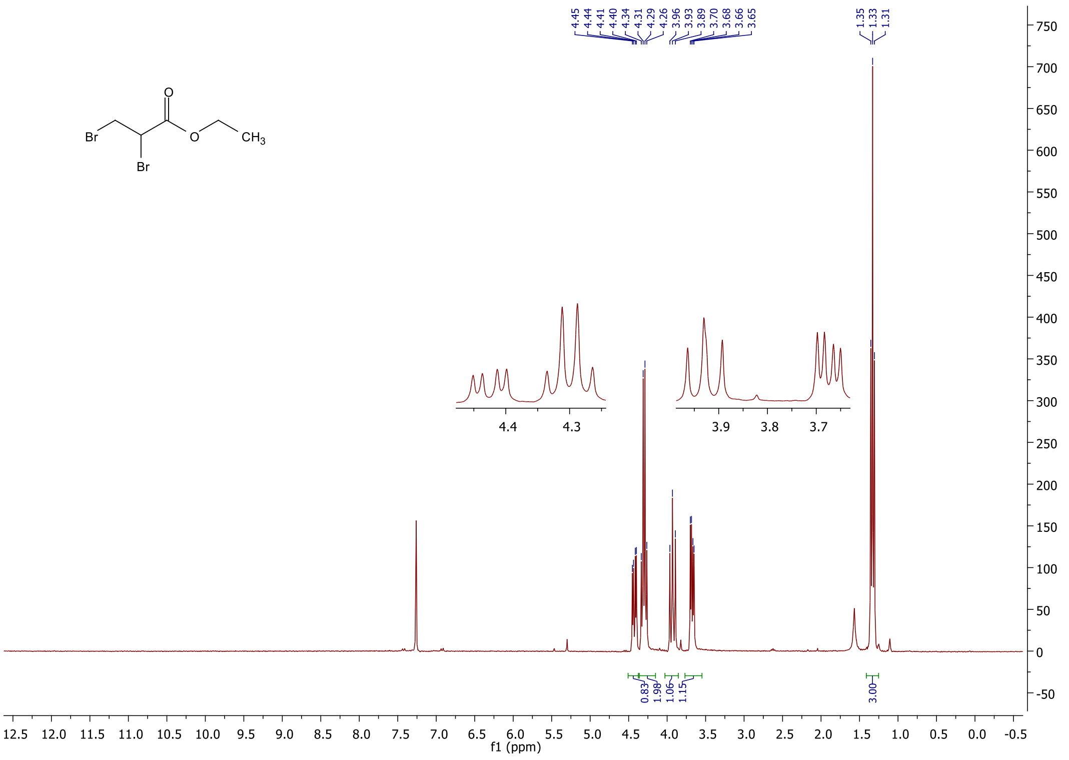

#### $^1$H NMR spectrum of **4hb**

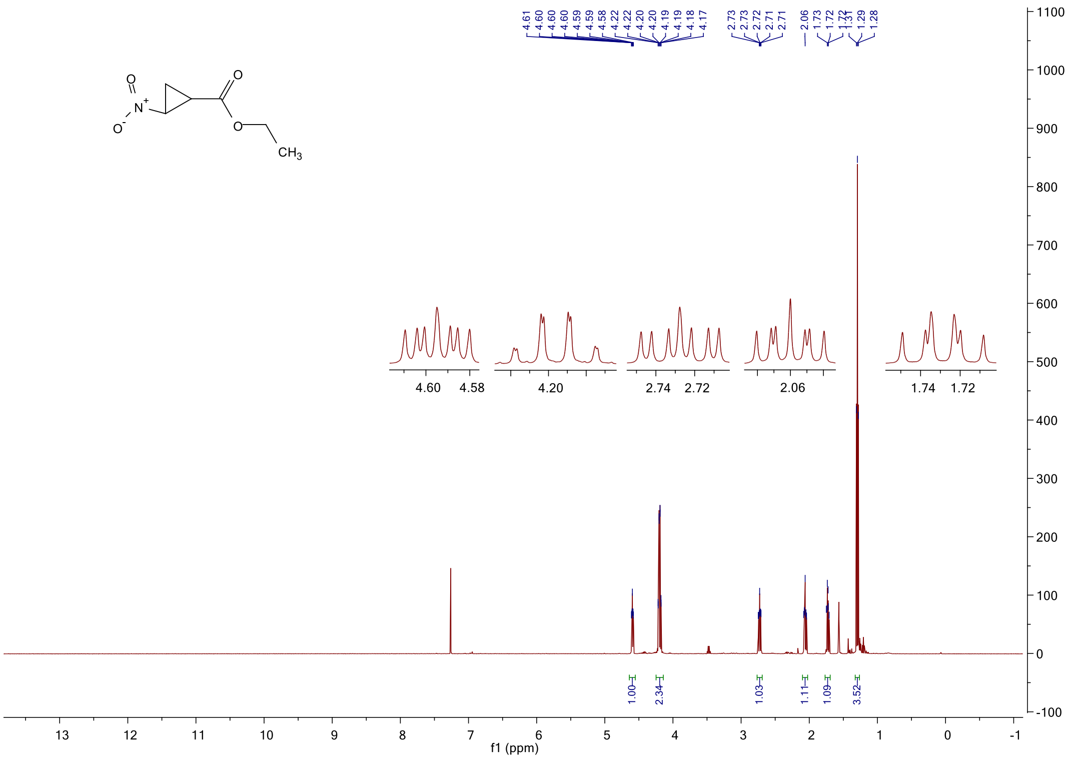

#### [1]H NMR spectrum of **4h**

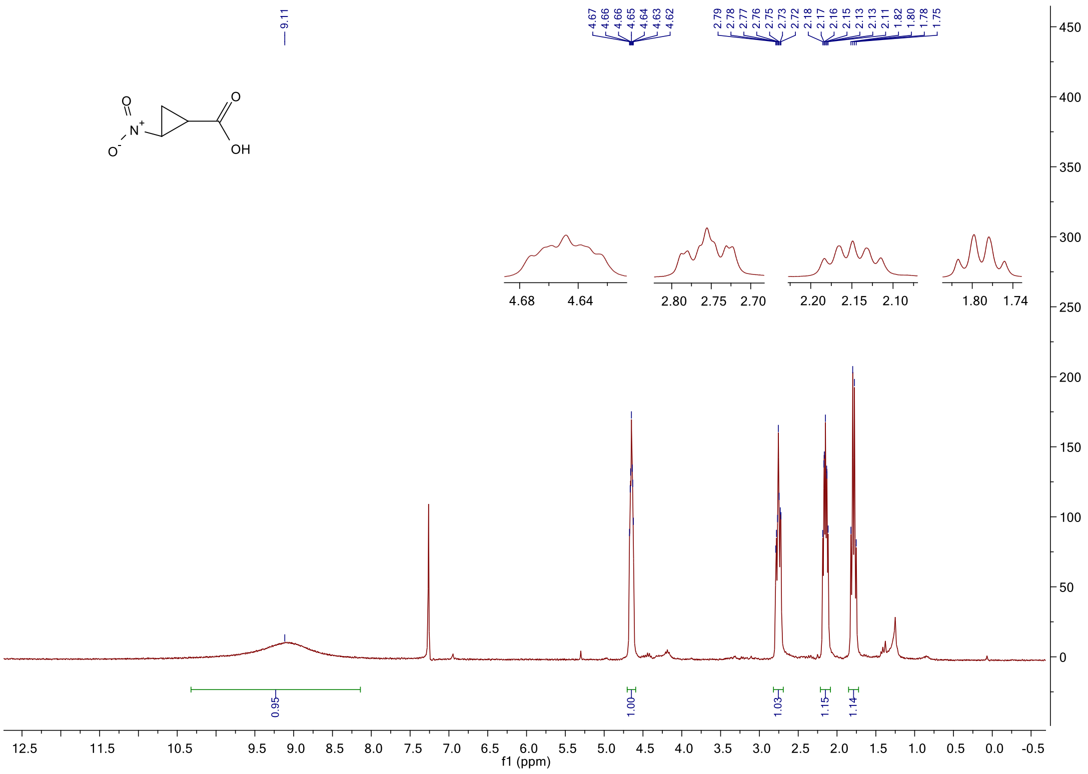

#### $^1$H NMR spectrum of **4ia**

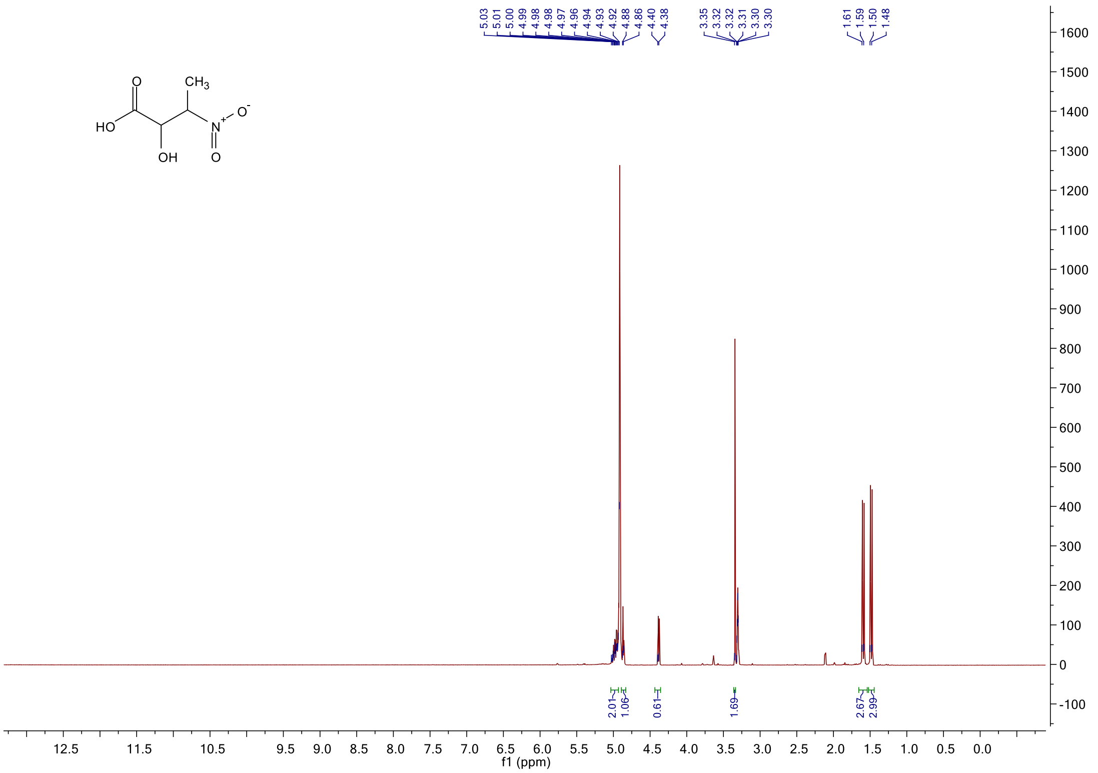

#### $^1$H NMR spectrum of **4ib**

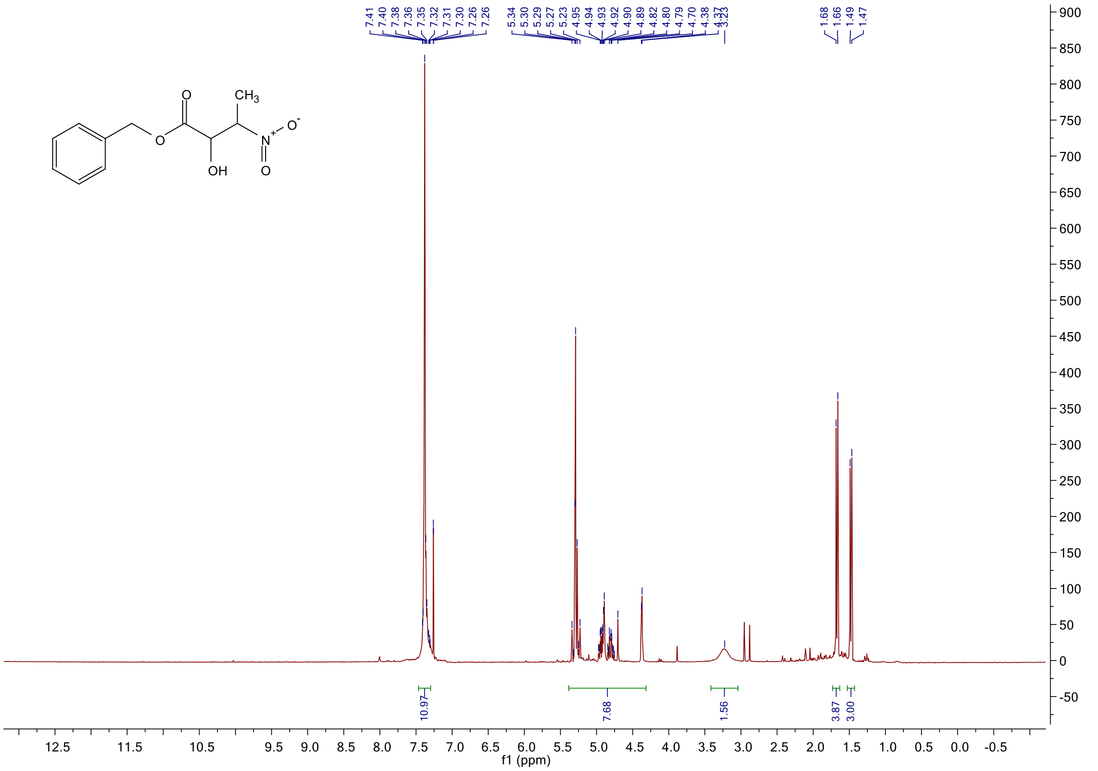

#### $^{1}$H NMR spectrum of **4ic**

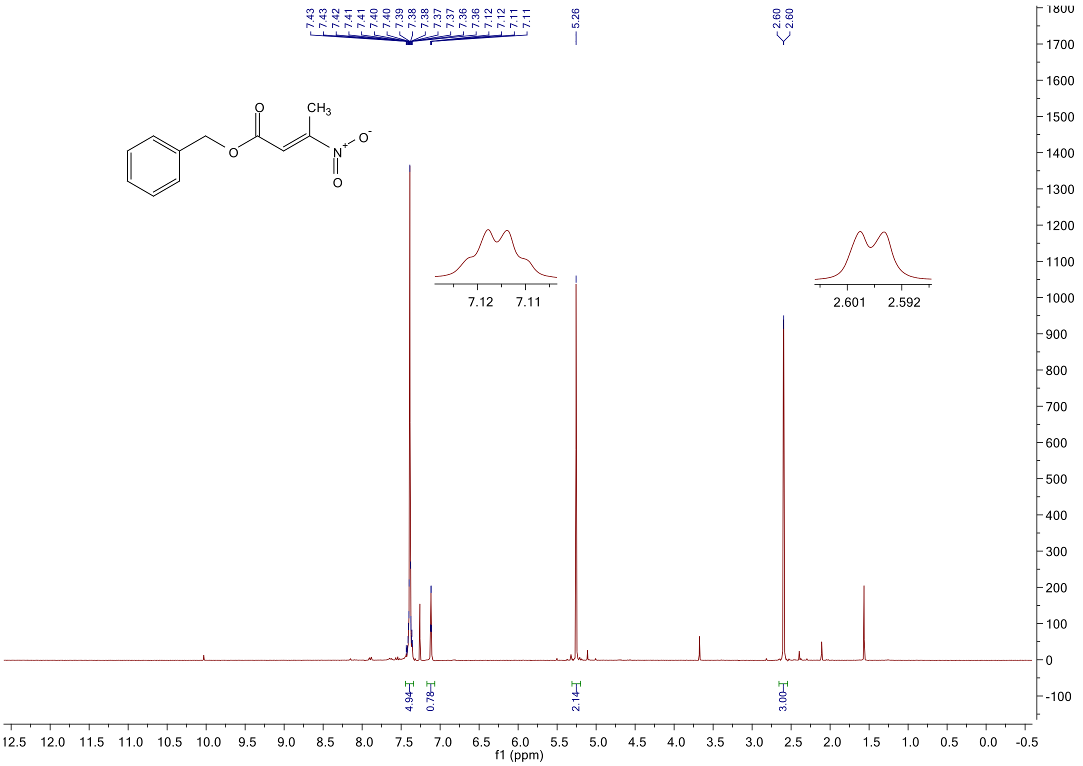

#### $^{13}$C NMR spectrum of **4ic**

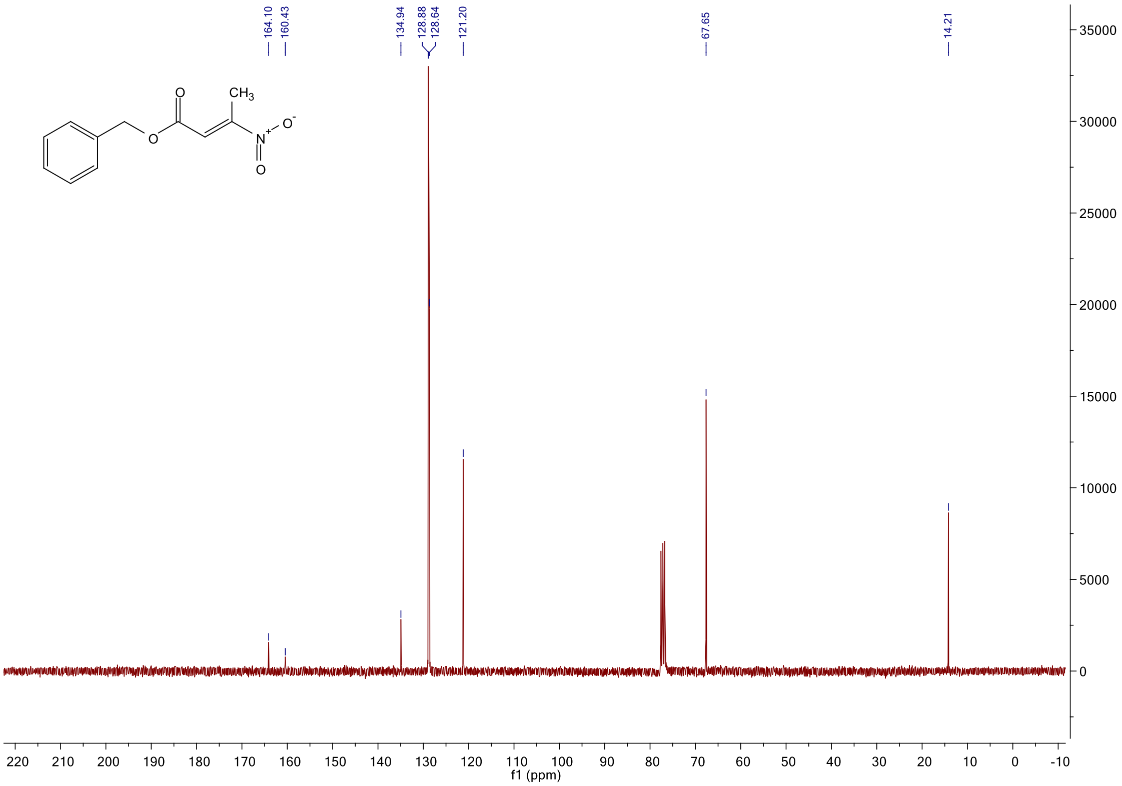

#### $^1$H NMR spectrum of **4i**

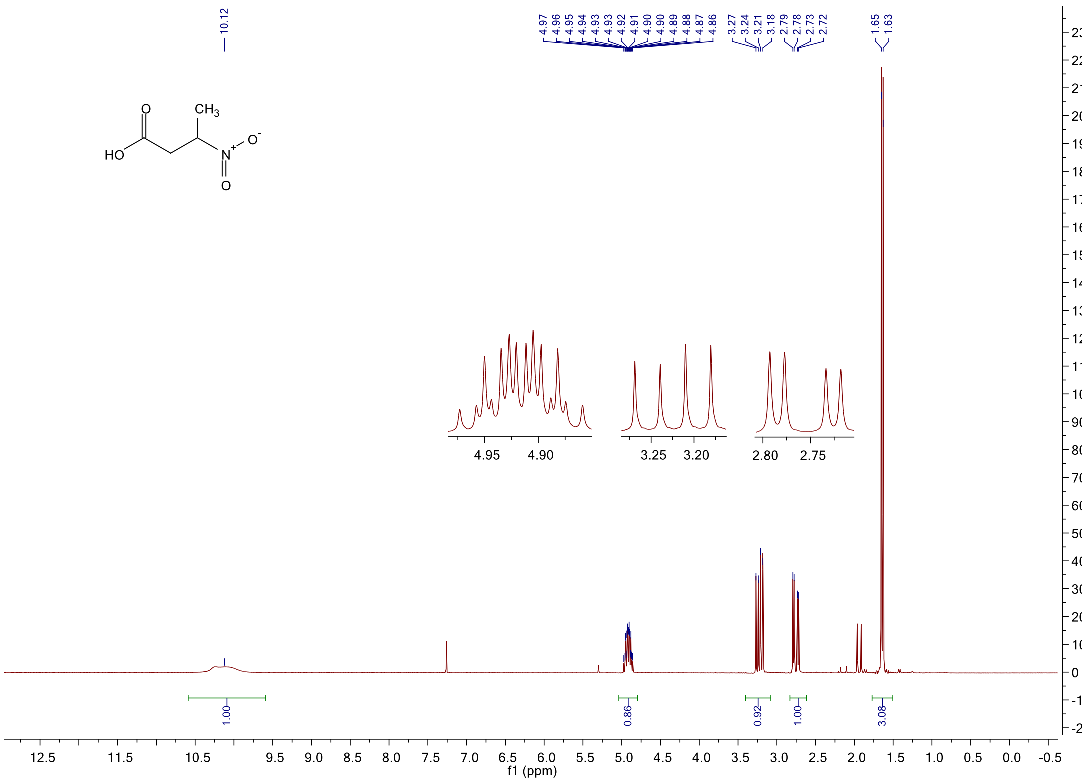

#### $^{13}$C NMR spectrum of **4i**

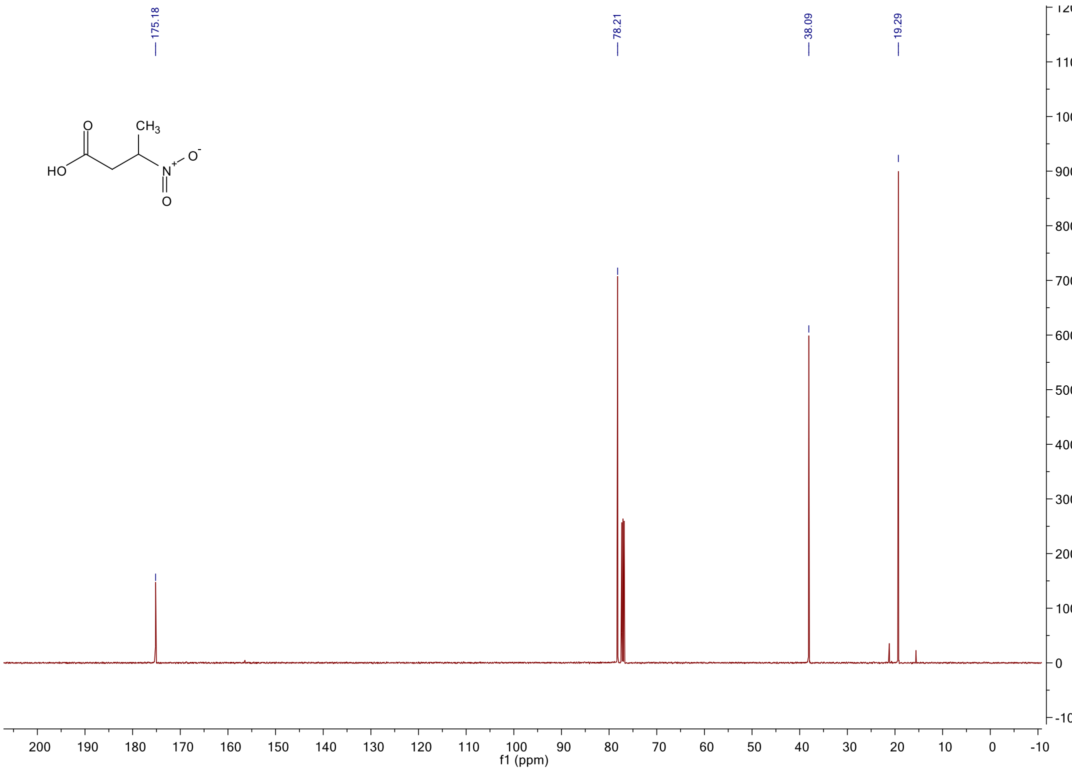

<sup></sup>

$^{1}$H NMR spectrum of **4k**

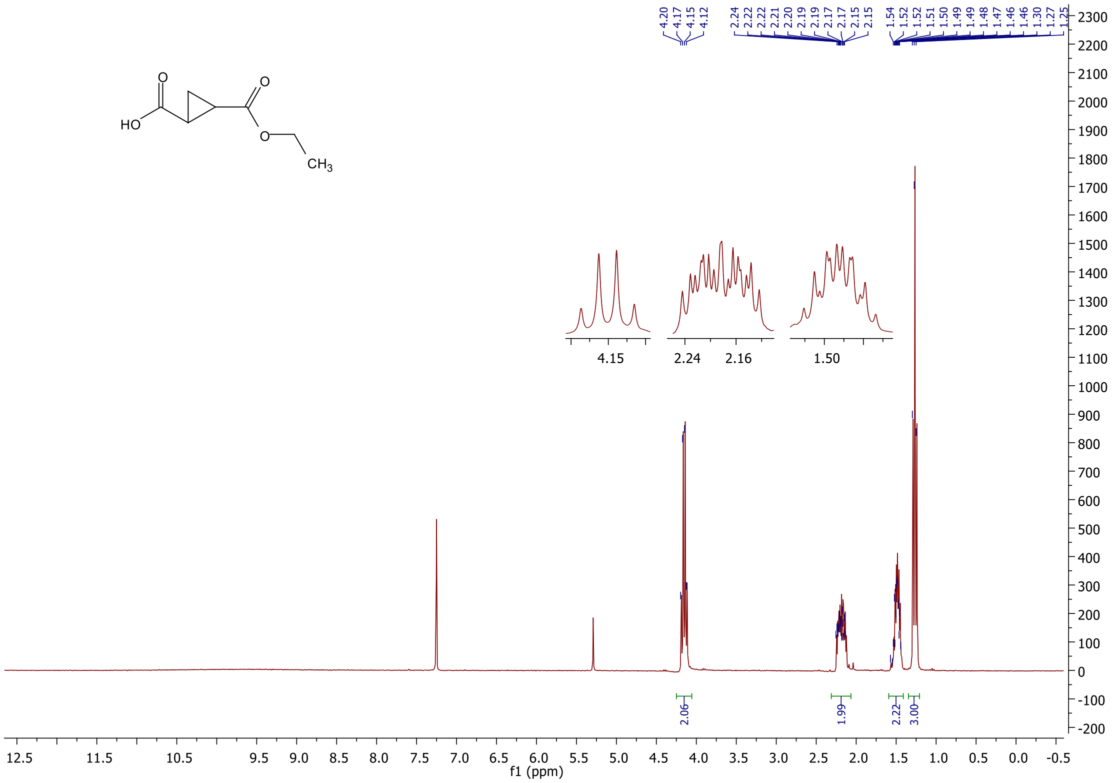

[1]H NMR spectrum of **4l**

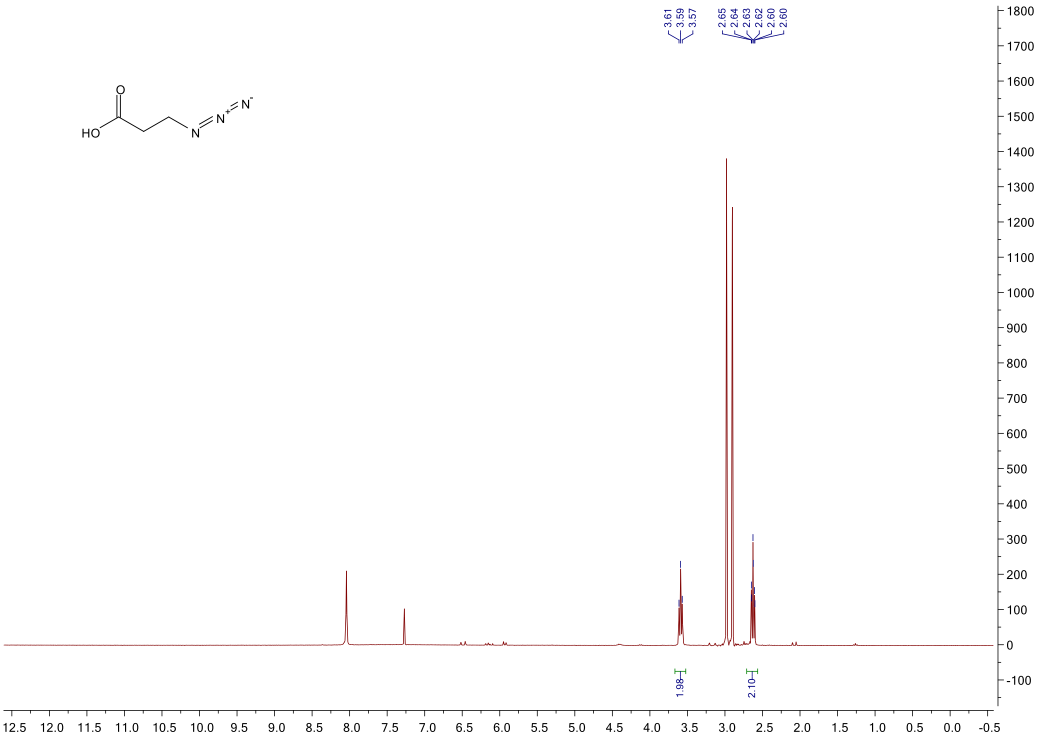

$^{1}$H NMR spectrum of **4m**

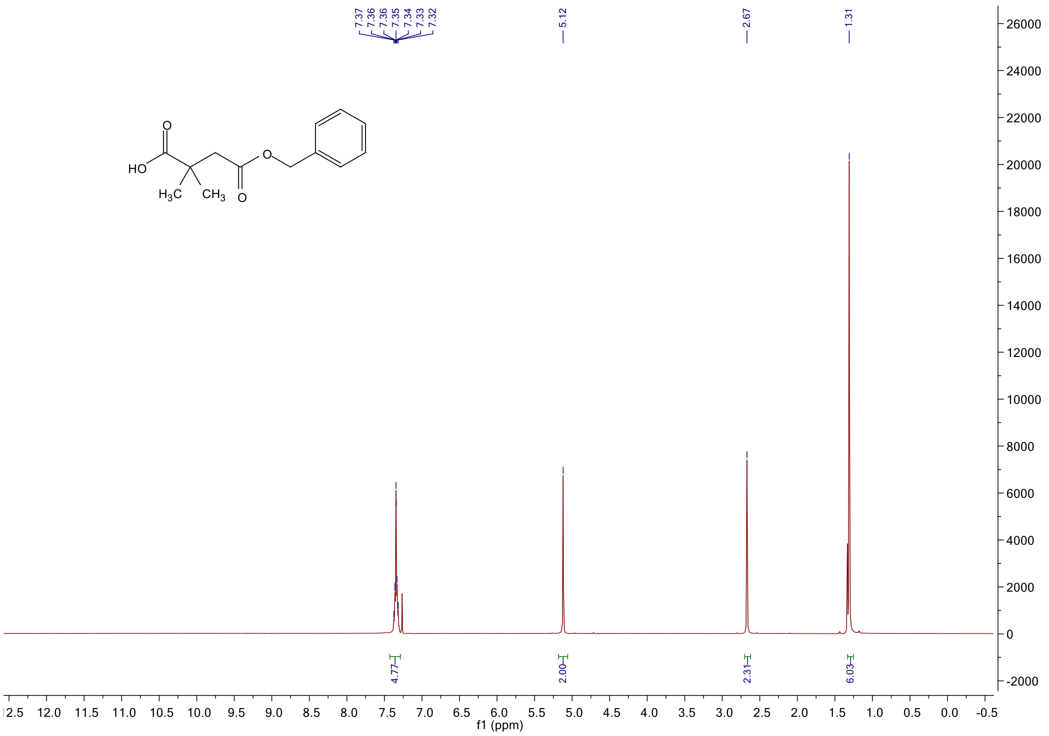

$^{13}$C NMR spectrum of **4m**

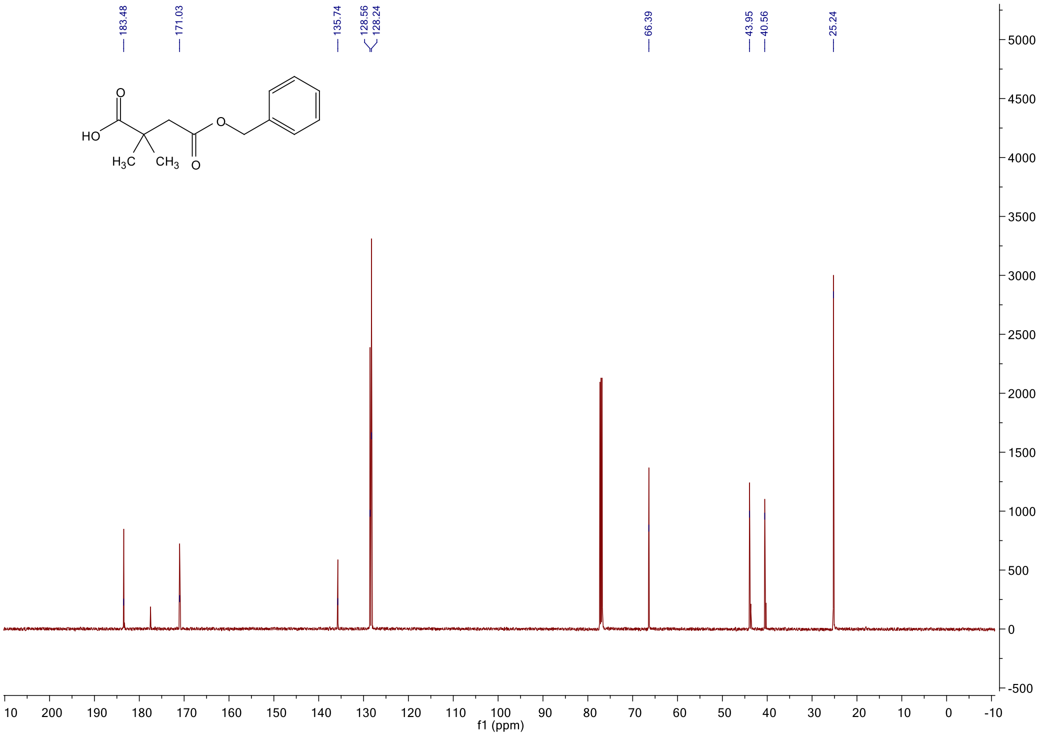

[1]H NMR spectrum of **4na**

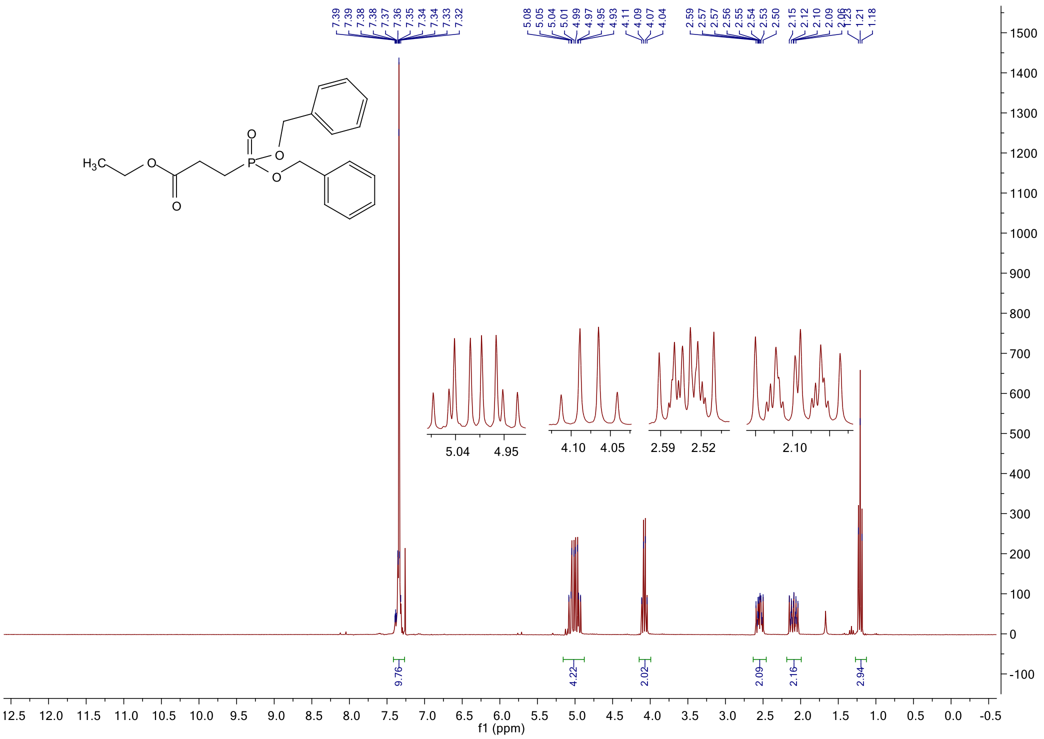

[1]H NMR spectrum of **4n**

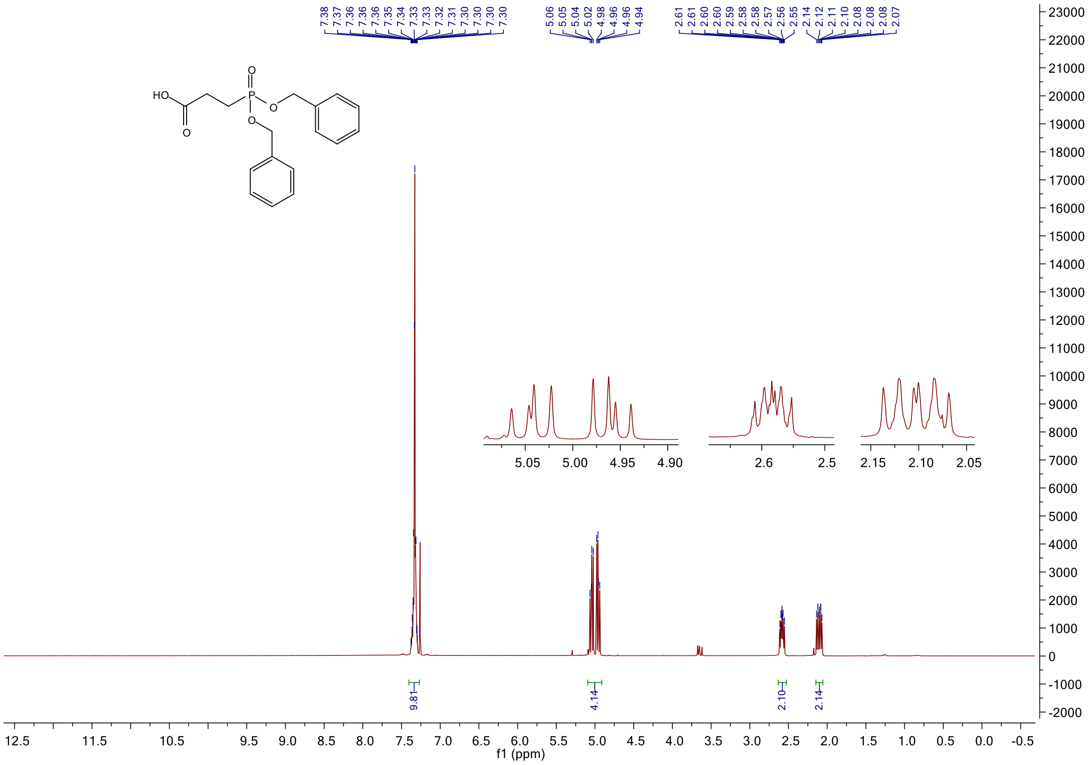

#### [1]H NMR spectrum of **4p**

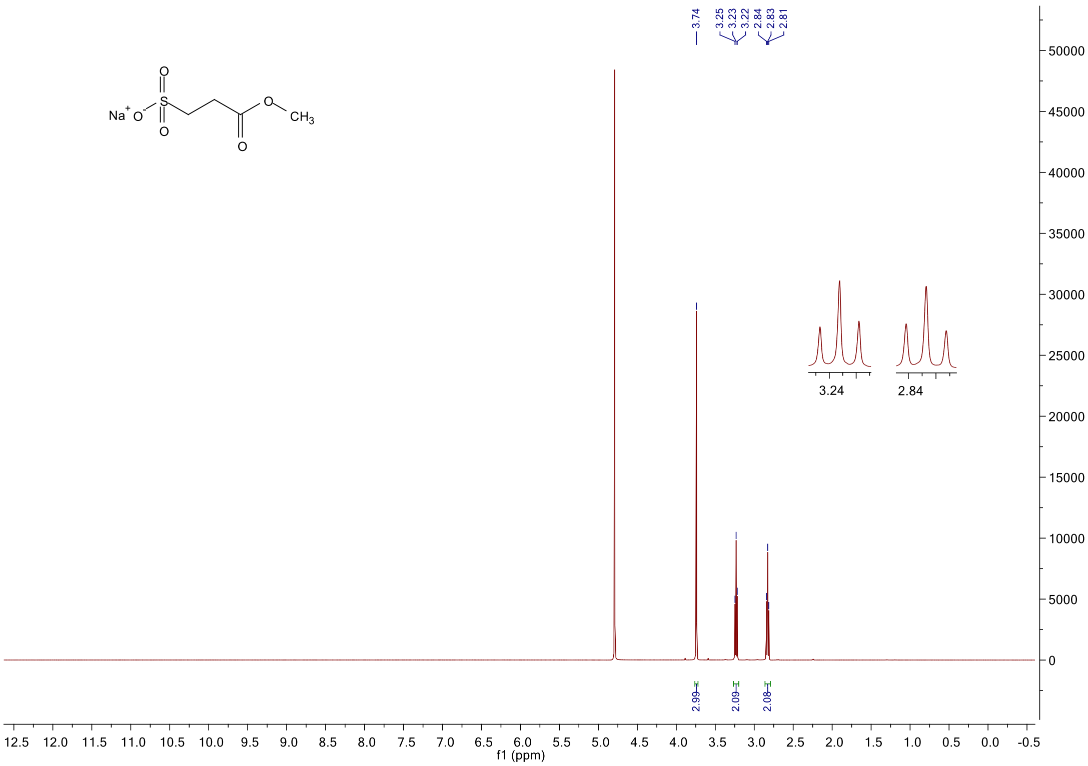

#### [1]H NMR spectrum of **2**

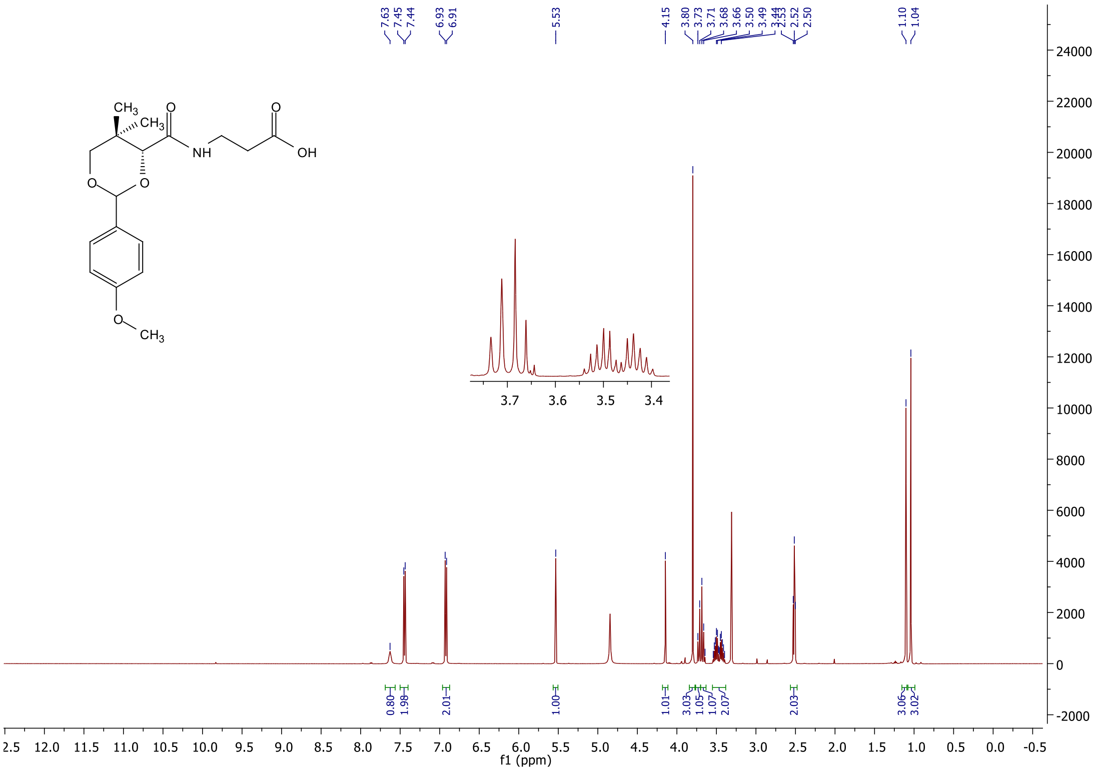

#### [1]H NMR spectrum of **19a**

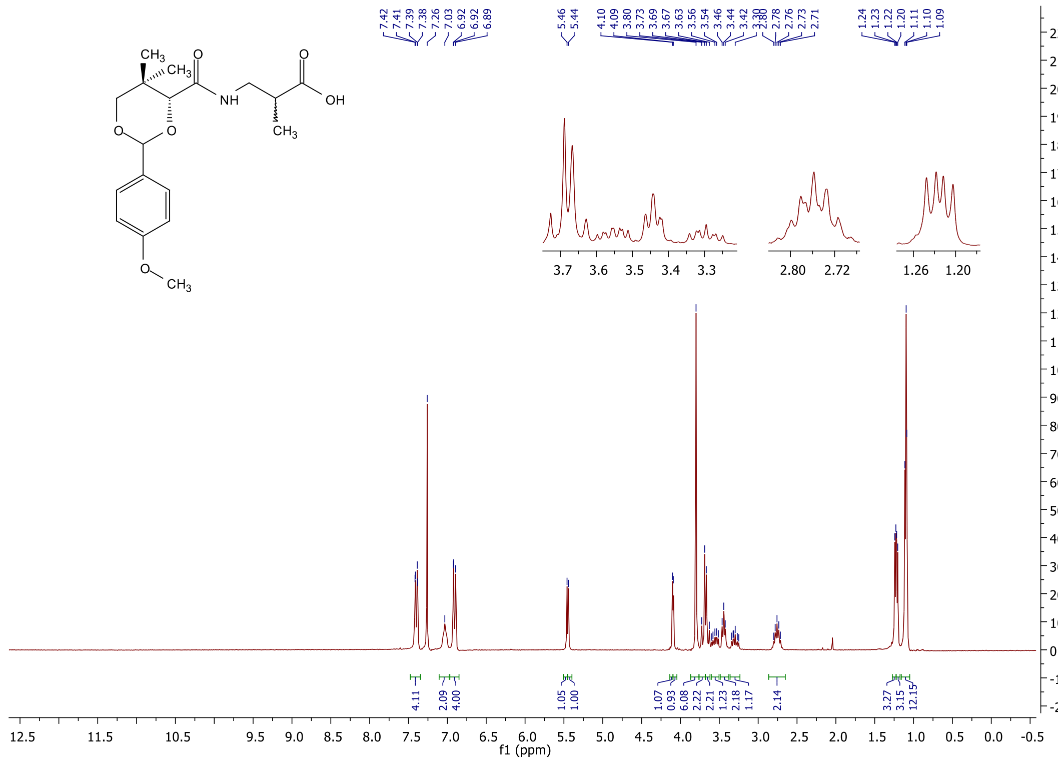

#### [13]C NMR spectrum of **19a**

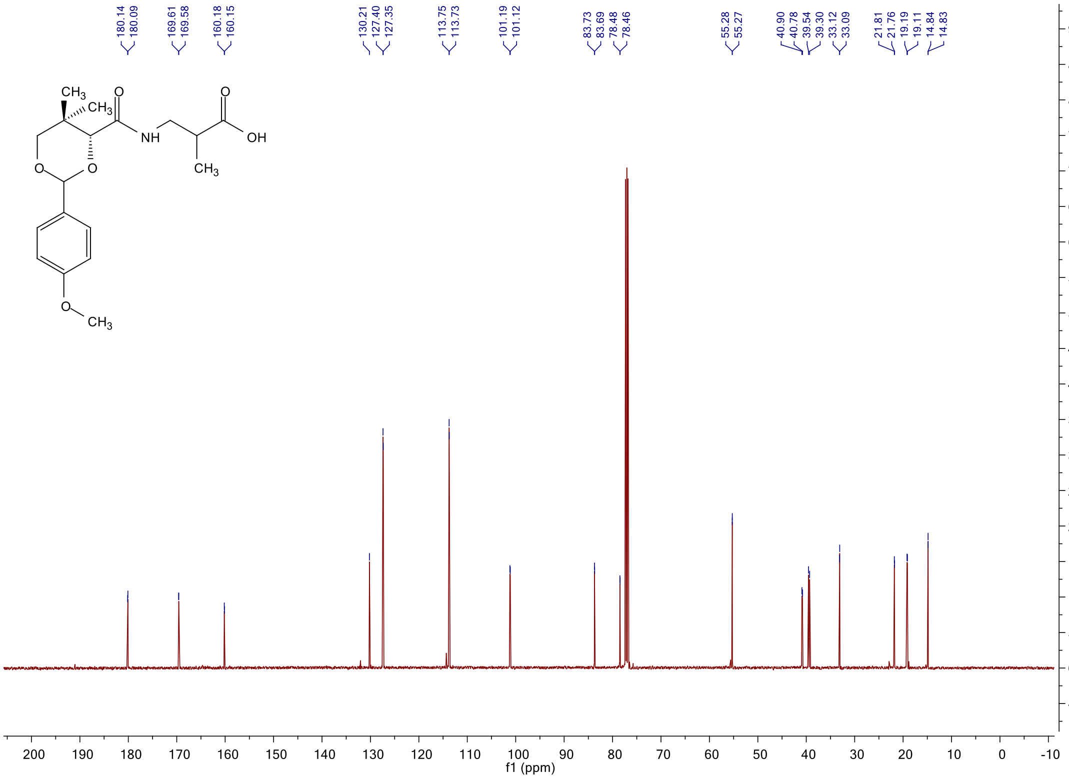

$^{1}$H NMR spectrum of **17ba**

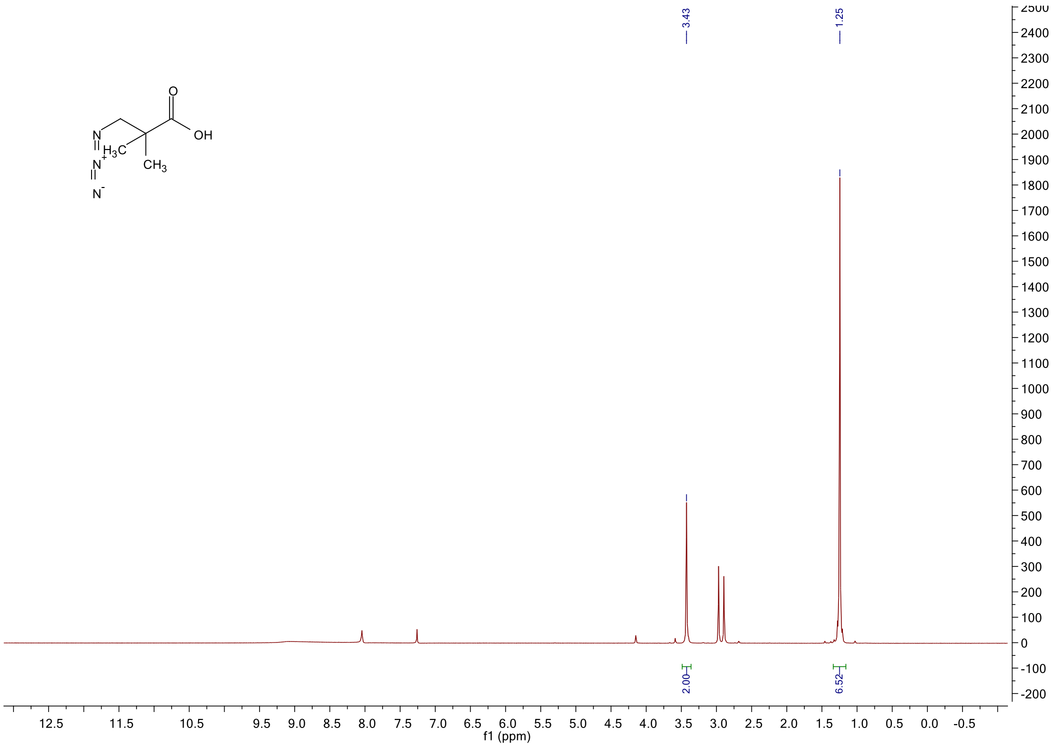

$^{1}$H NMR spectrum of **17b**

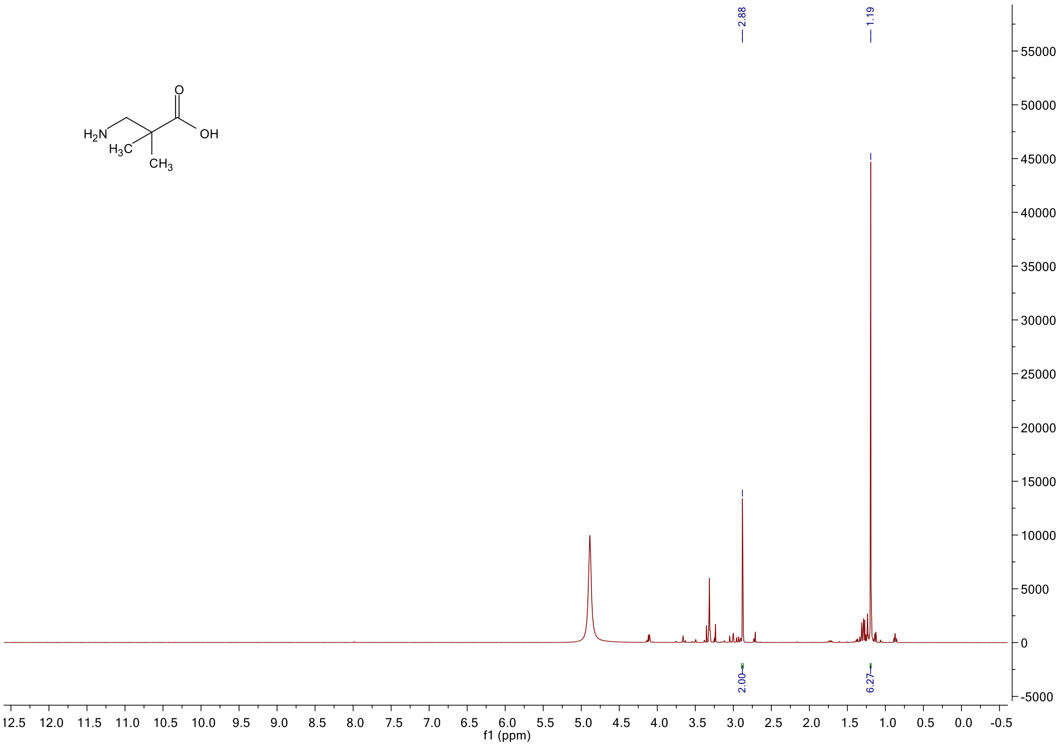

#### [1]H NMR spectrum of **19b**

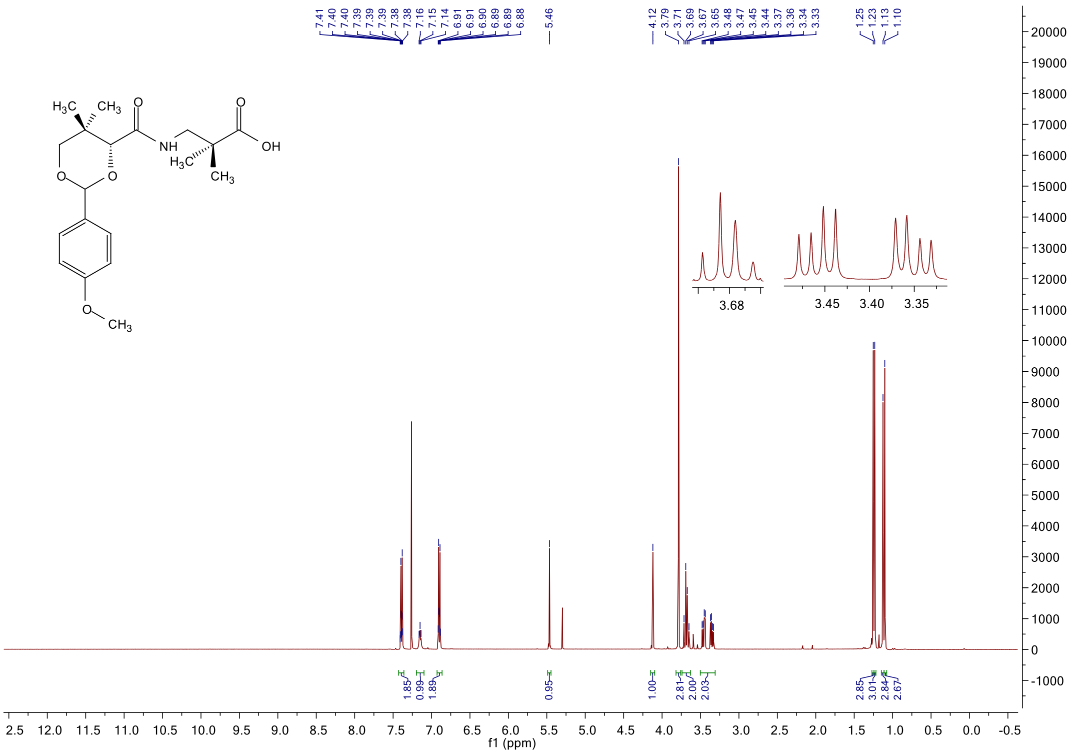

#### [13]C NMR spectrum of **19b**

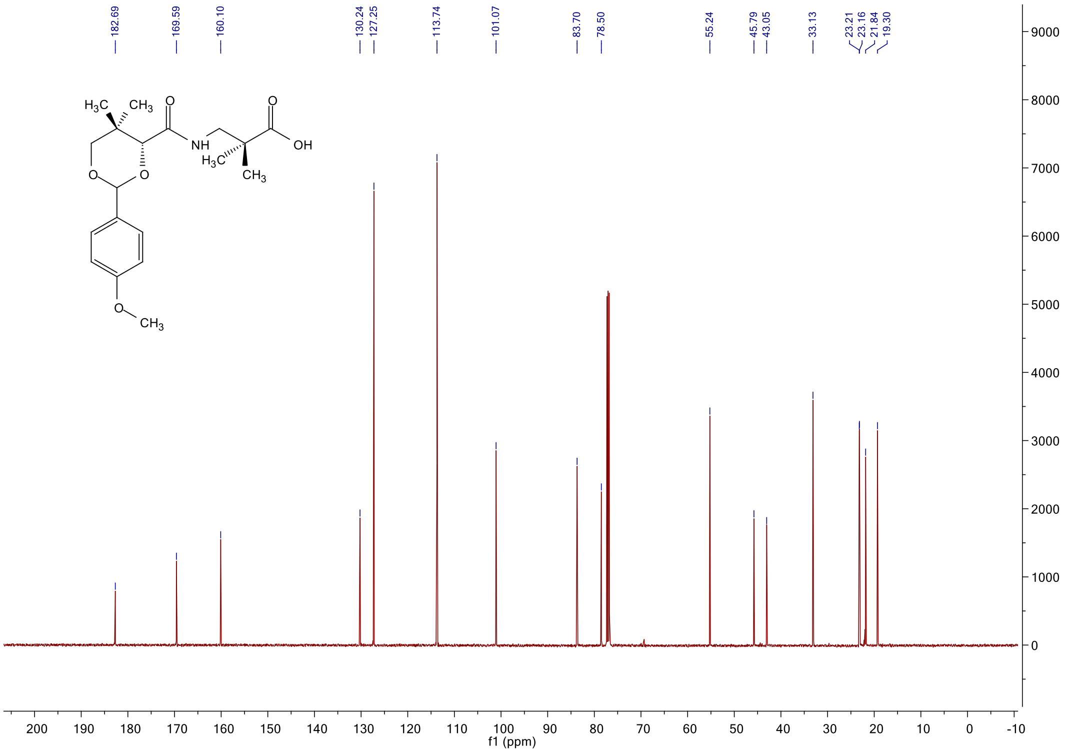

#### $^1$H NMR spectrum of **17c**

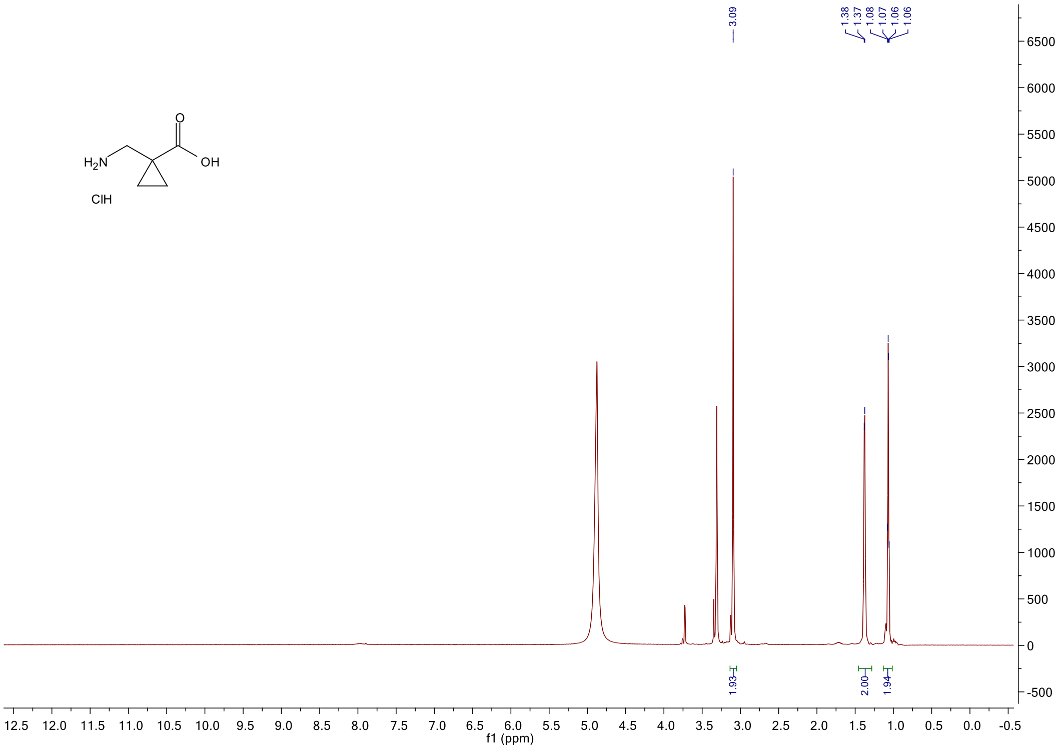

#### $^1$H NMR spectrum of **19c**

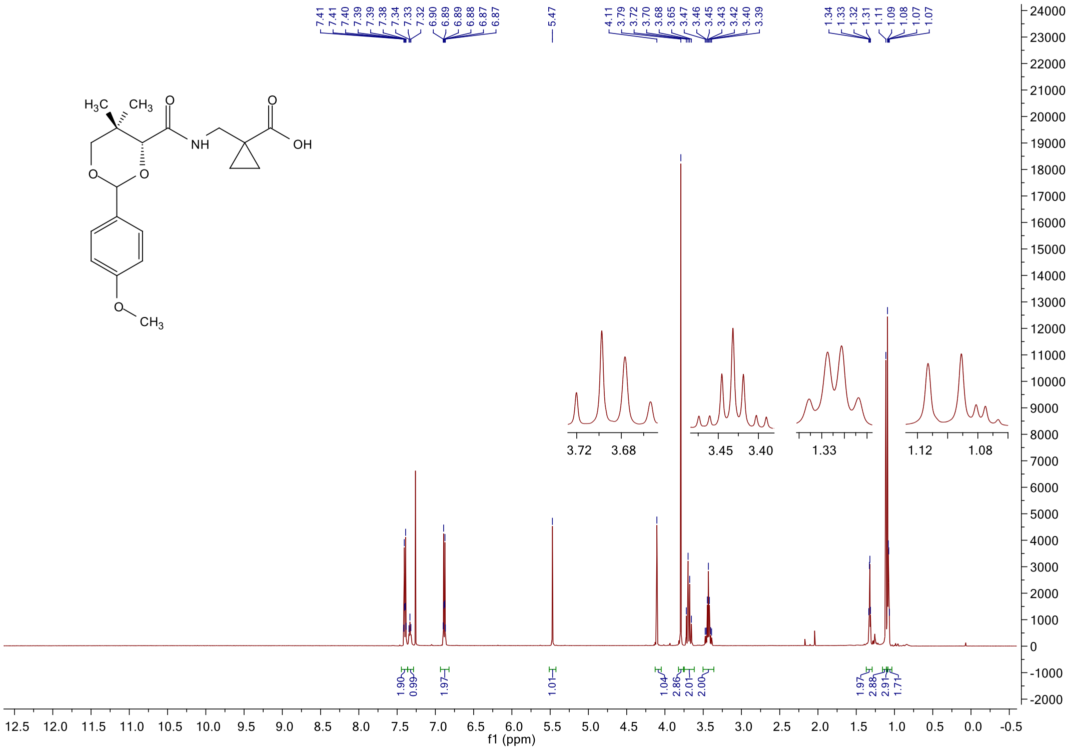

#### $^{13}$C NMR spectrum of **19c**

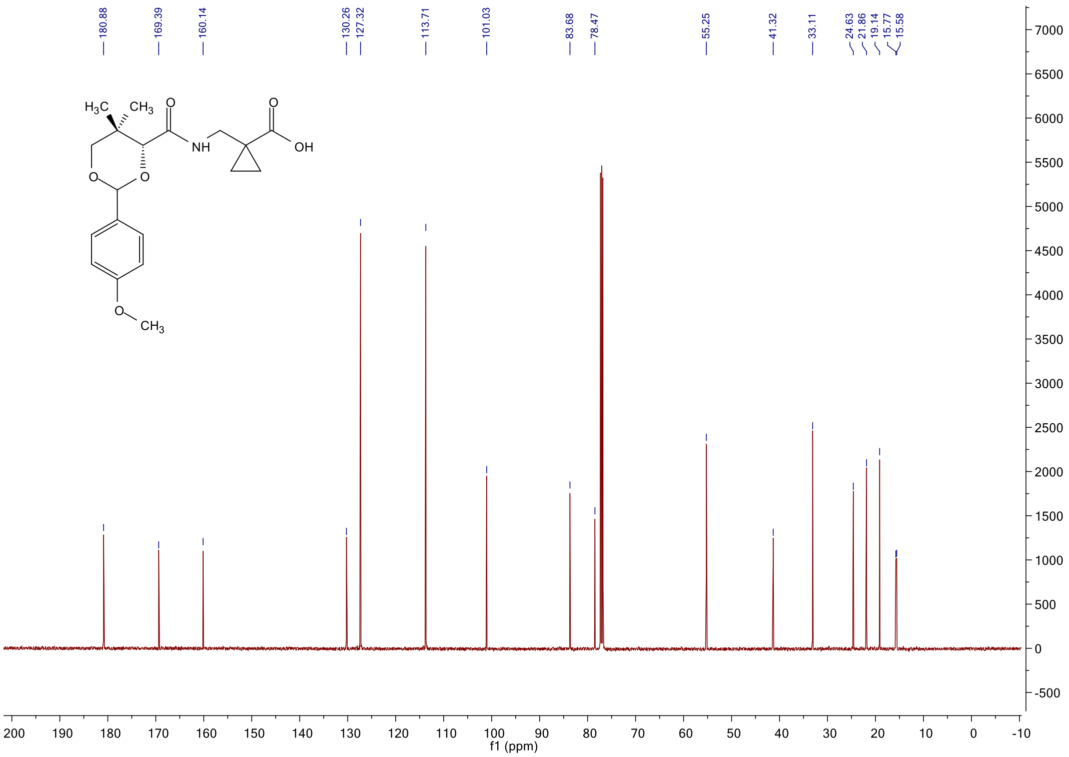

#### $^{1}$H NMR spectrum of **3**

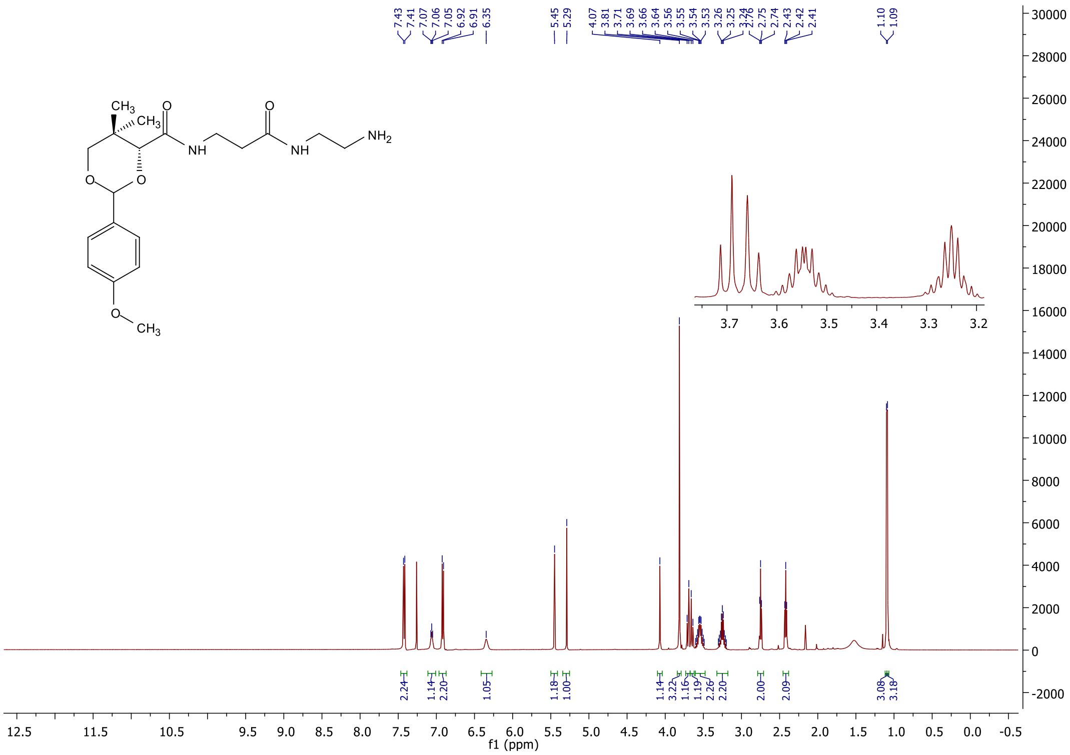

#### $^{1}$H NMR spectrum of **20a**

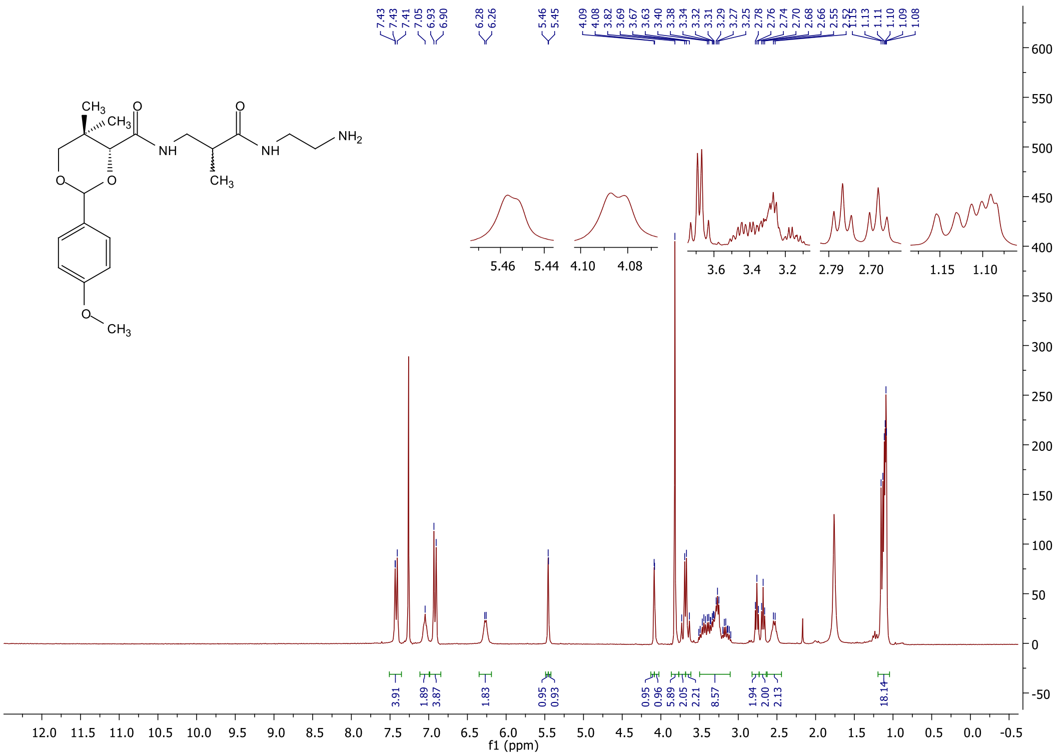

#### $^{13}$C NMR spectrum of **20a**

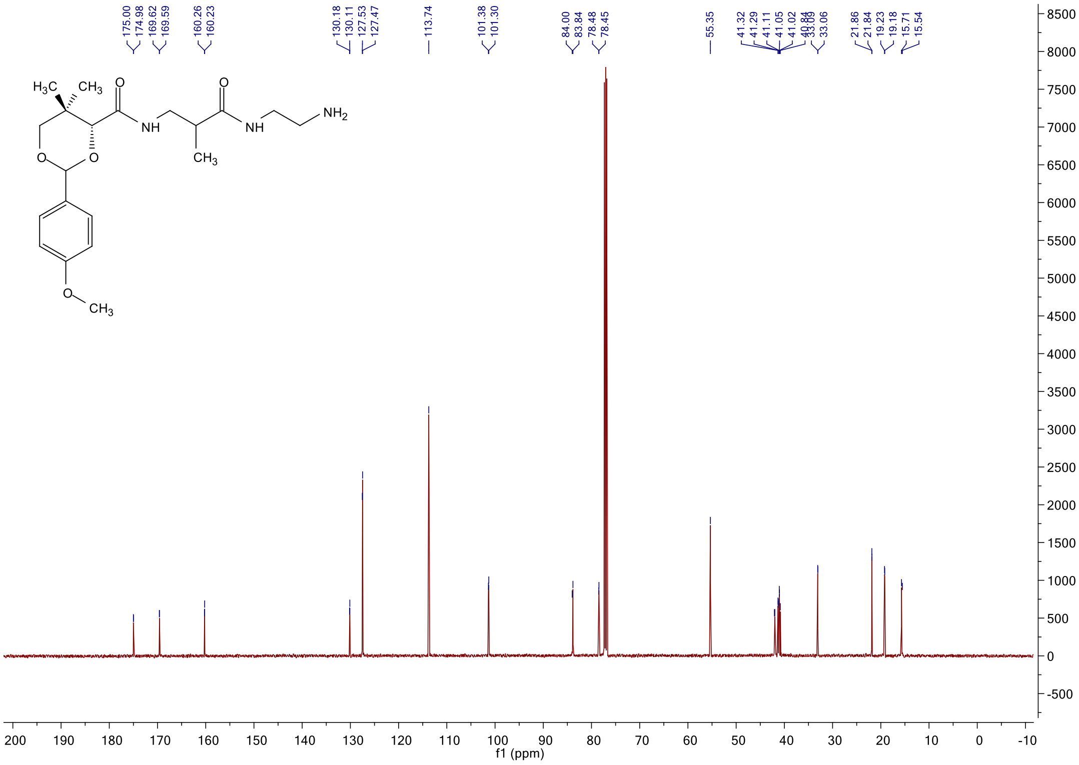

#### $^{1}$H NMR spectrum of **20b**

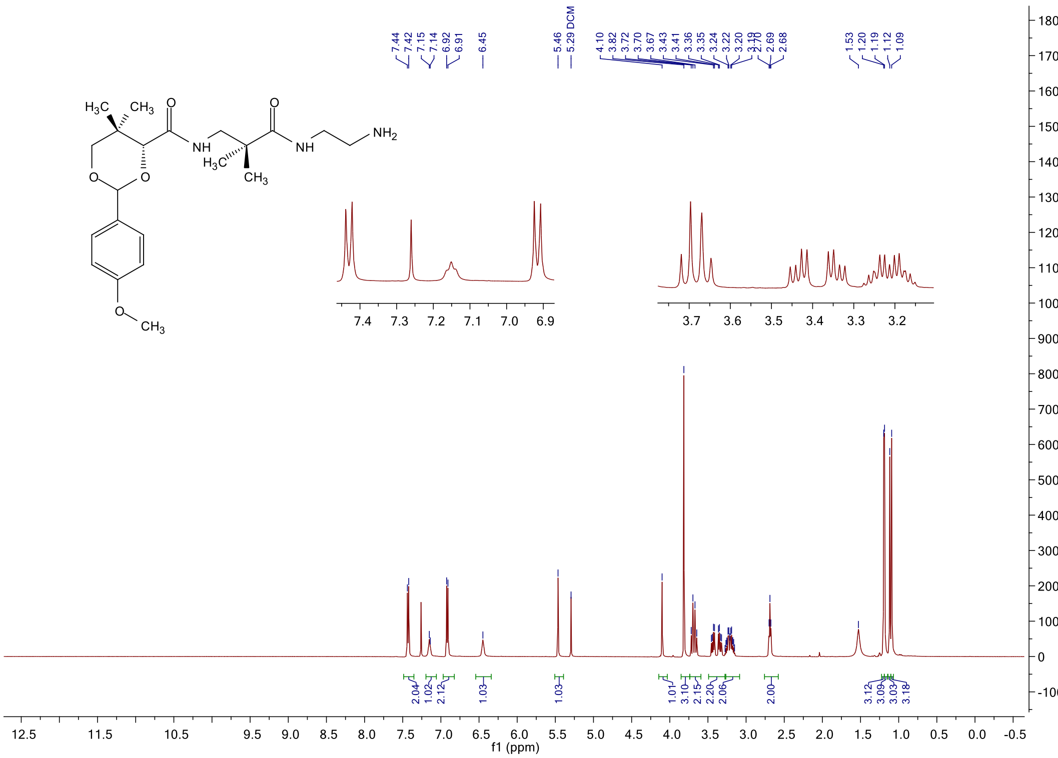

#### $^{13}$C NMR spectrum of **20b**

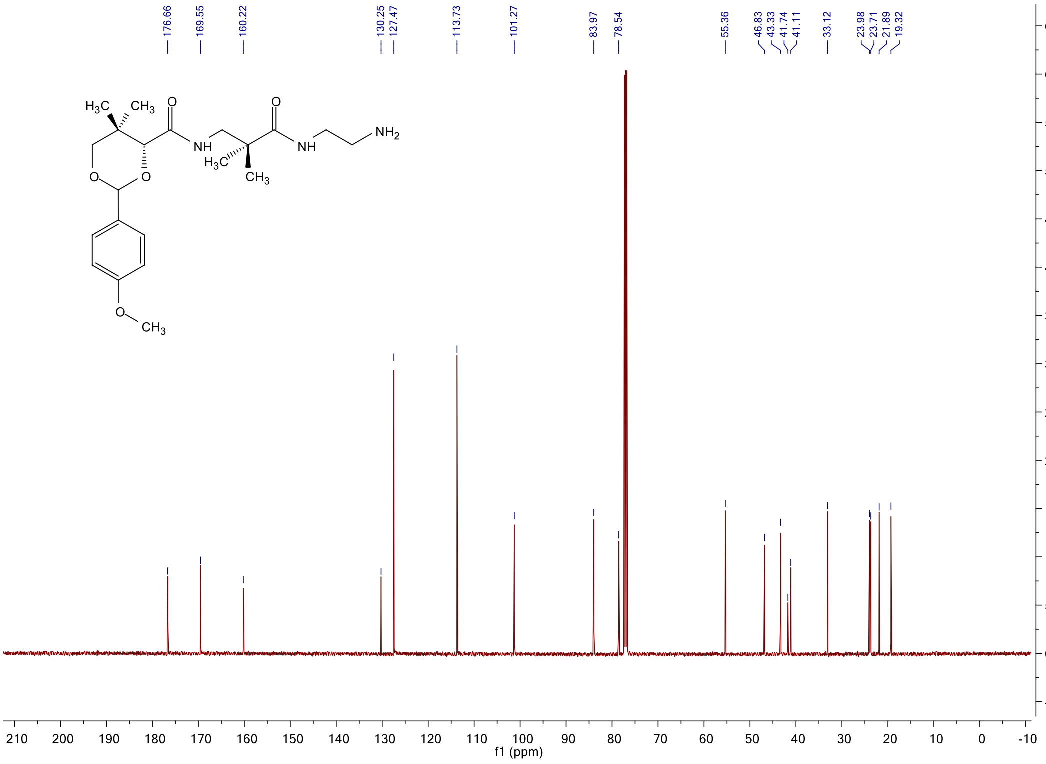

#### [1]H NMR spectrum of **20c**

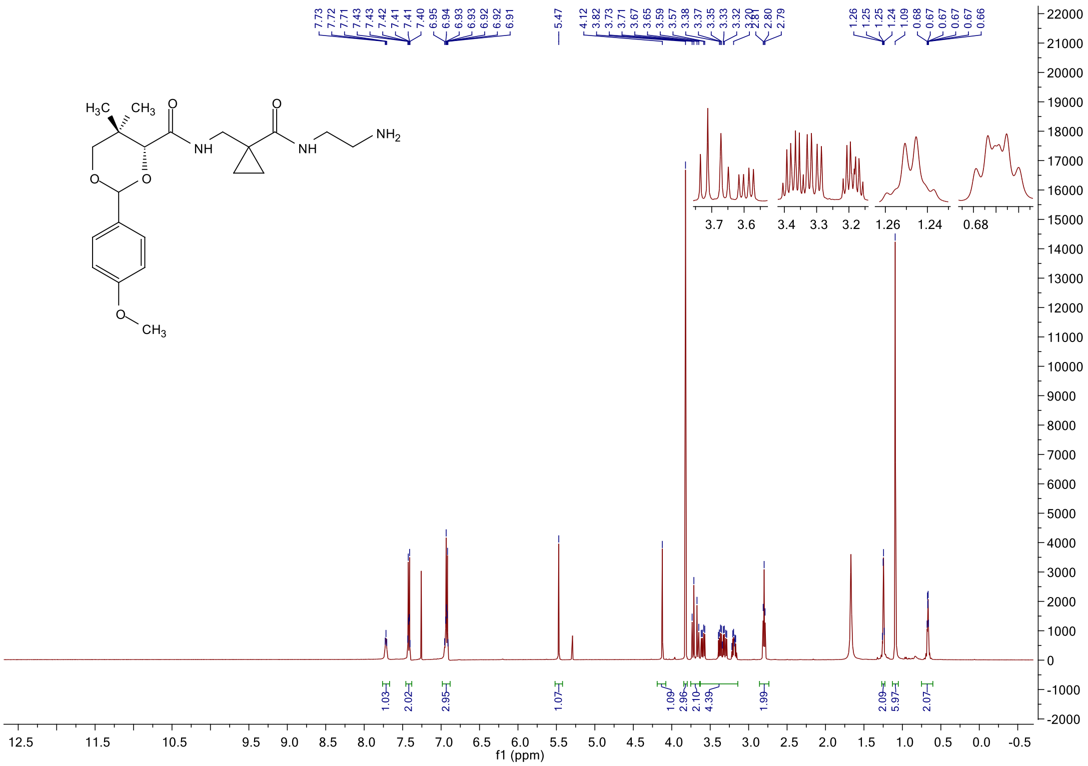

#### [13]C NMR spectrum of **20c**

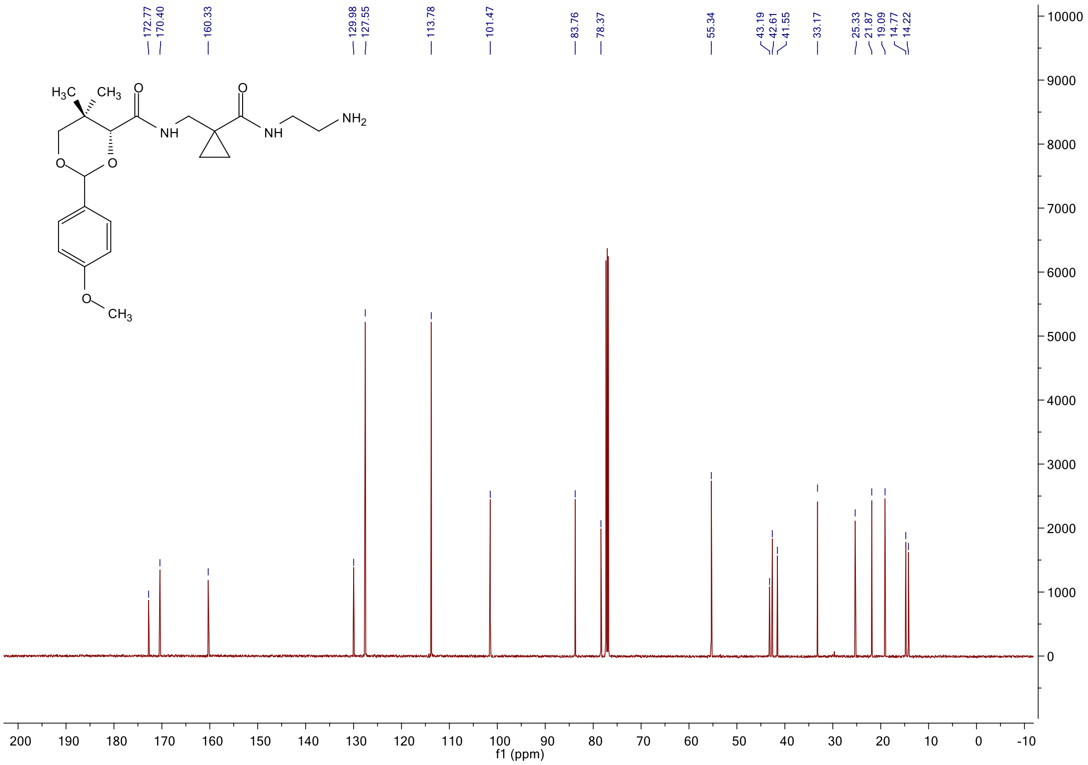

#### $^{1}$H NMR spectrum of **5a**

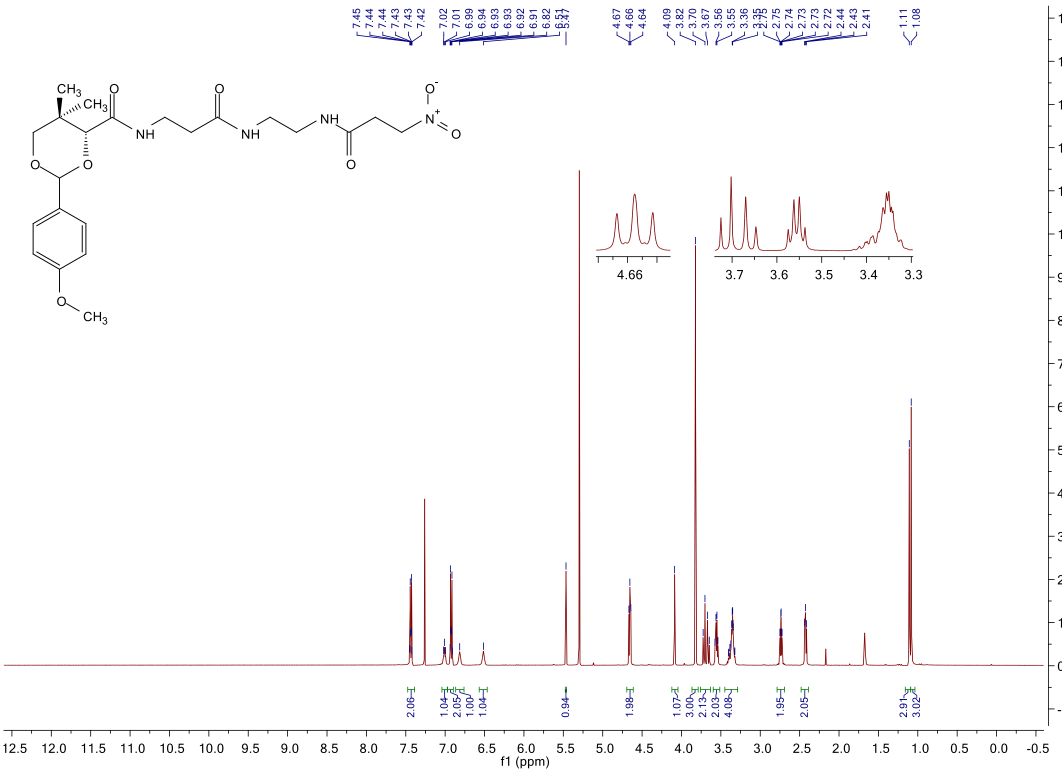

#### $^{1}$H NMR spectrum of **5b**

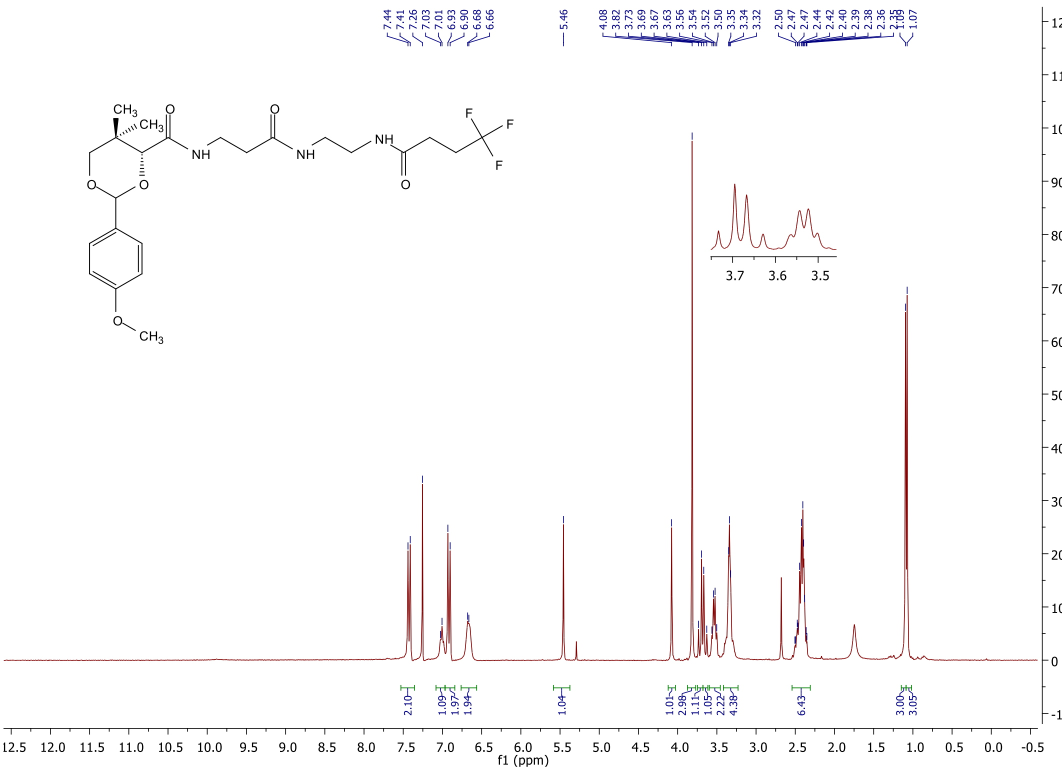

#### $^{13}$C NMR spectrum of **5b**

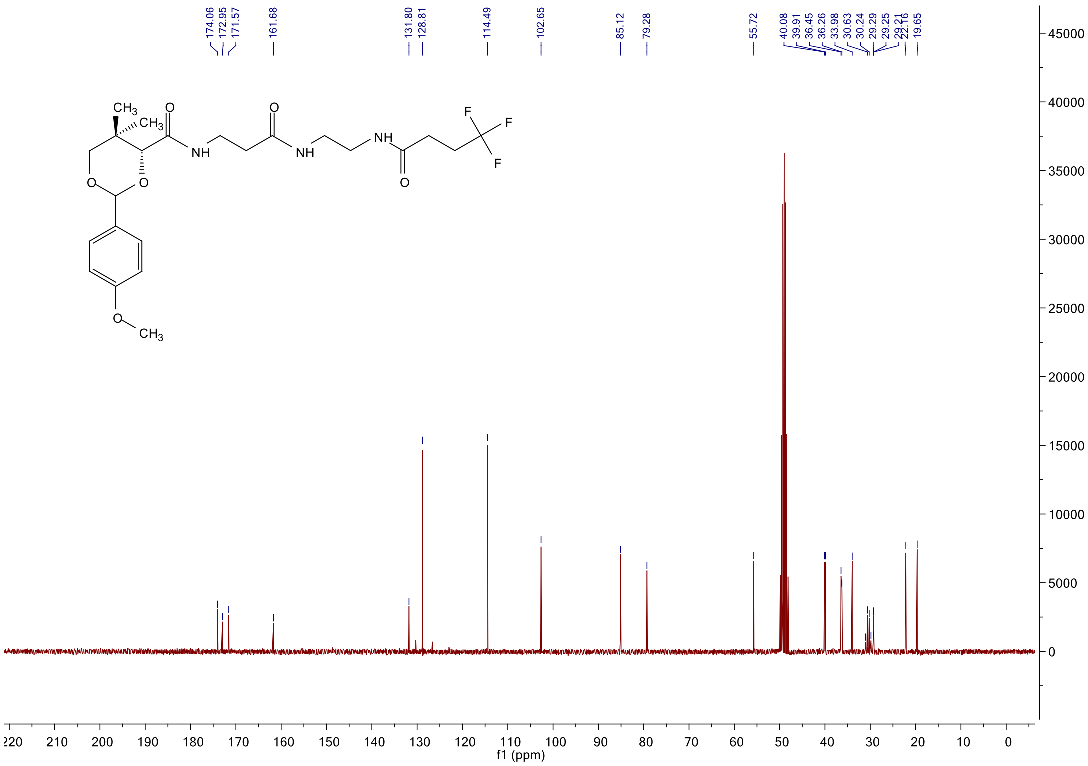

#### $^{19}$F NMR spectrum of **5b**

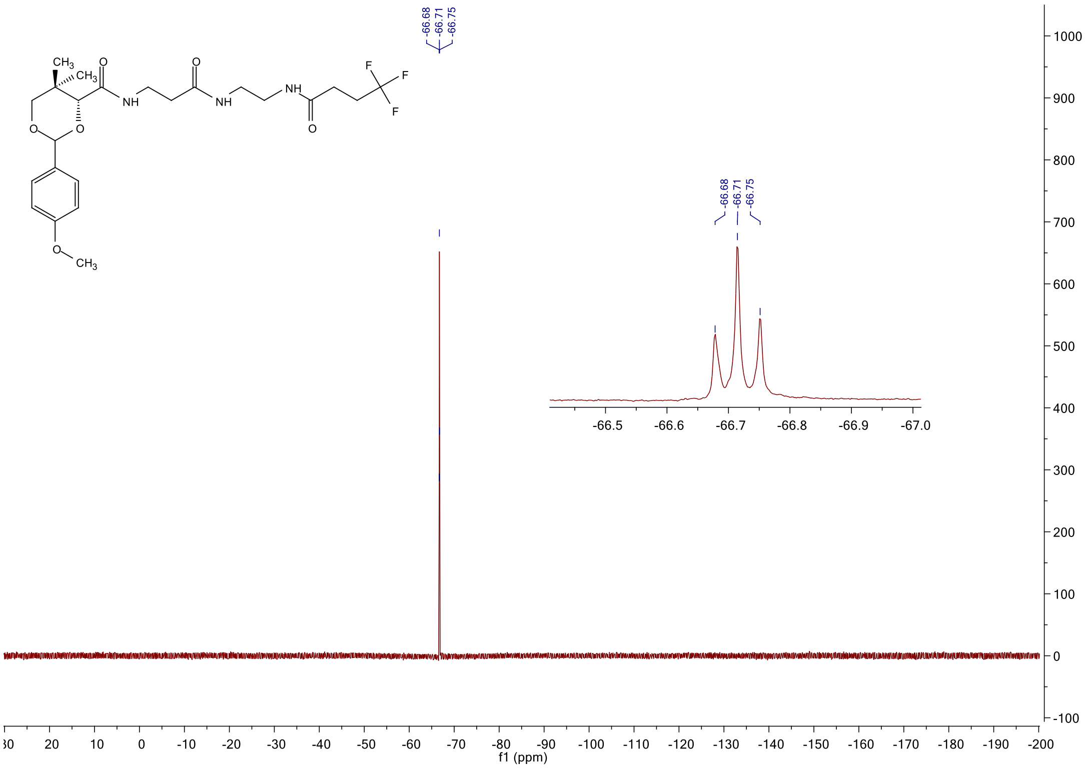

<sup></sup>

#### $^{1}$H NMR spectrum of **5c**

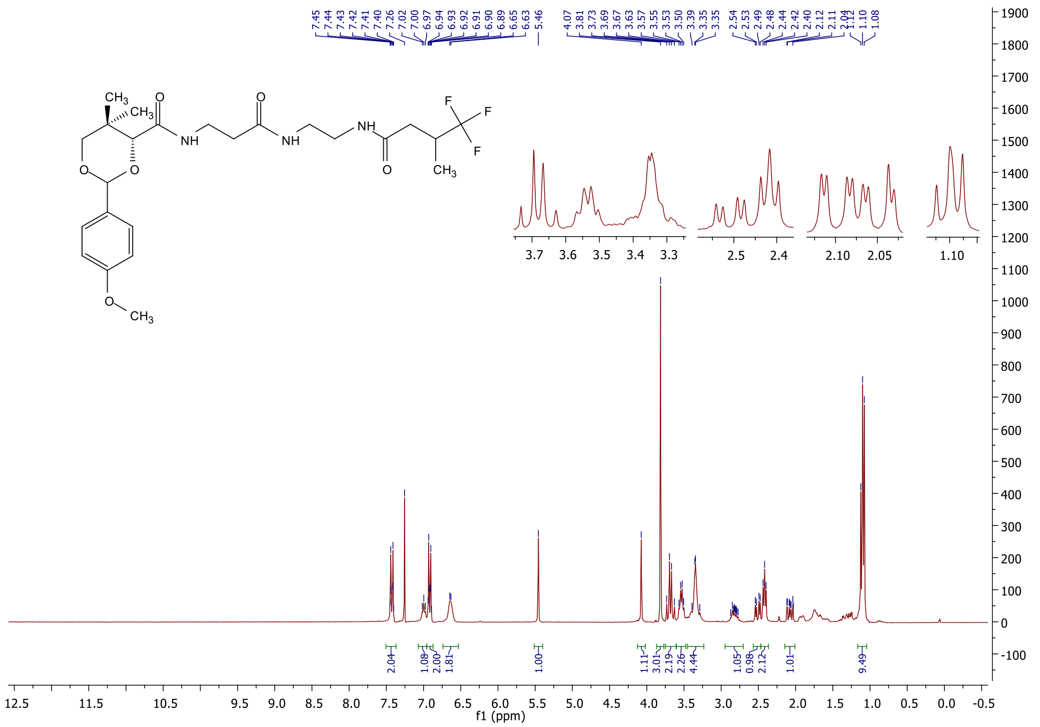

#### $^{13}$C NMR spectrum of **5c**

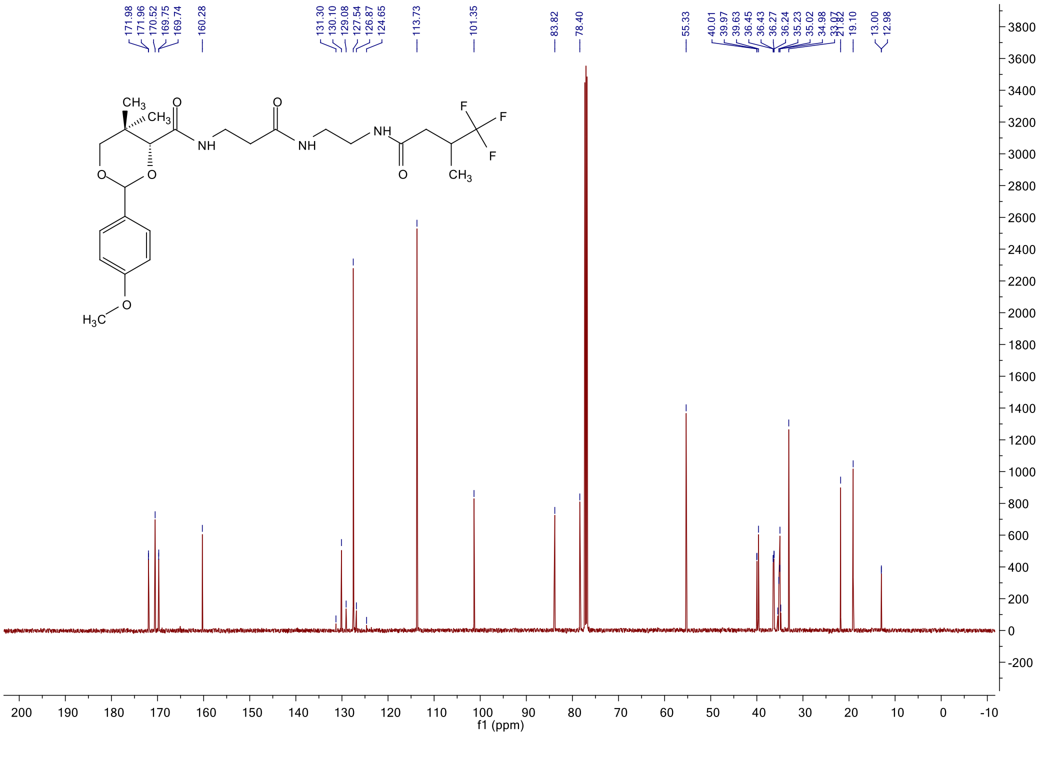

$^{19}$F NMR spectrum of **5c**

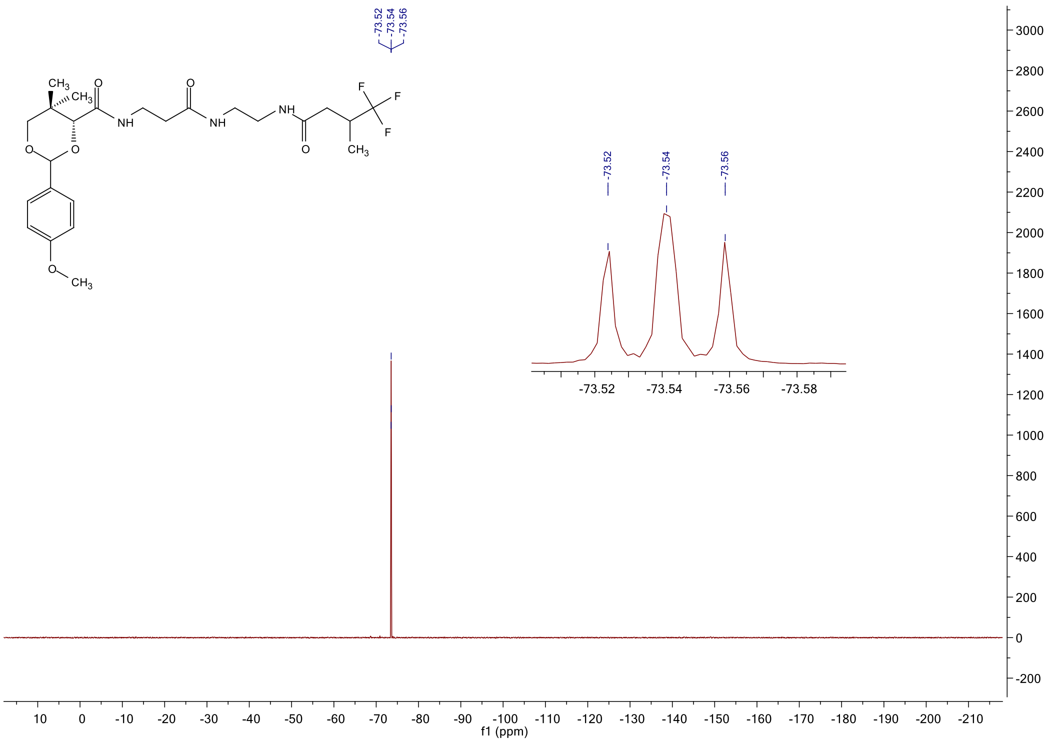

$^{1}$H NMR spectrum of **5d**

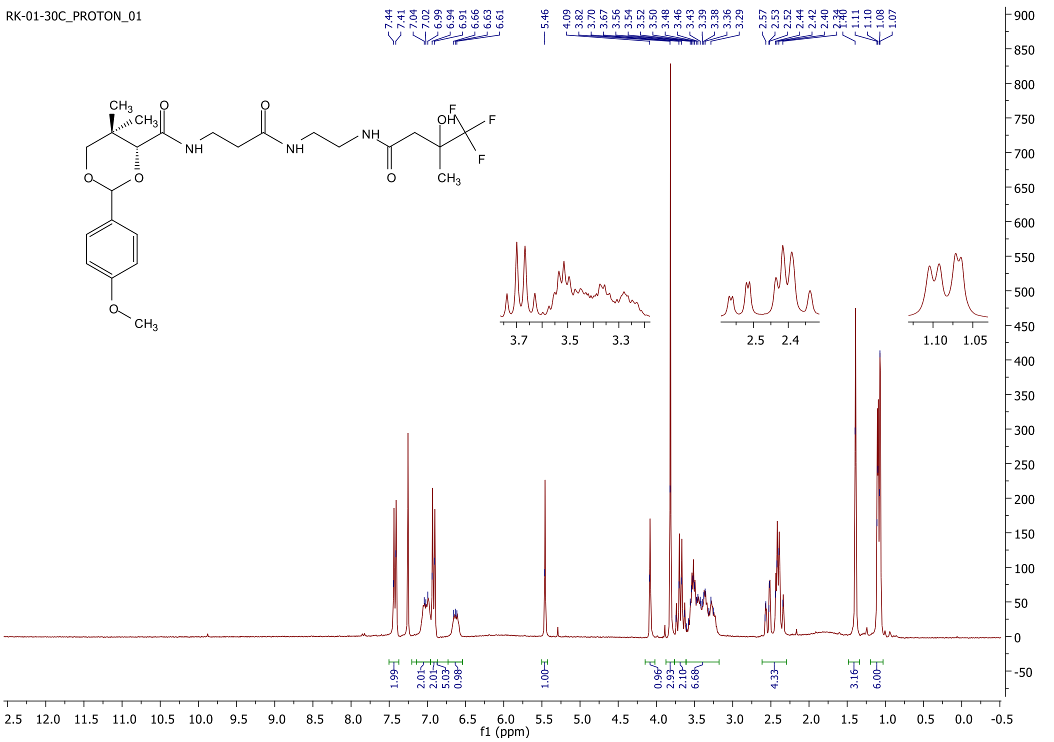

#### $^{13}$C NMR spectrum of **5d**

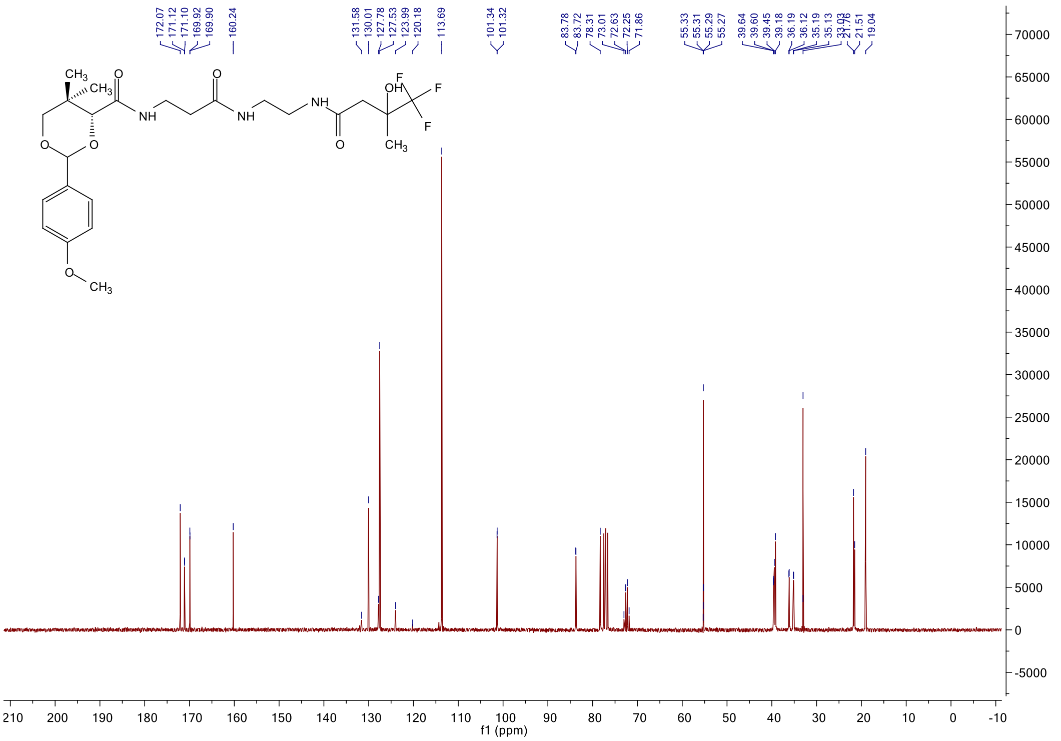

#### $^{19}$F NMR spectrum of **5d**

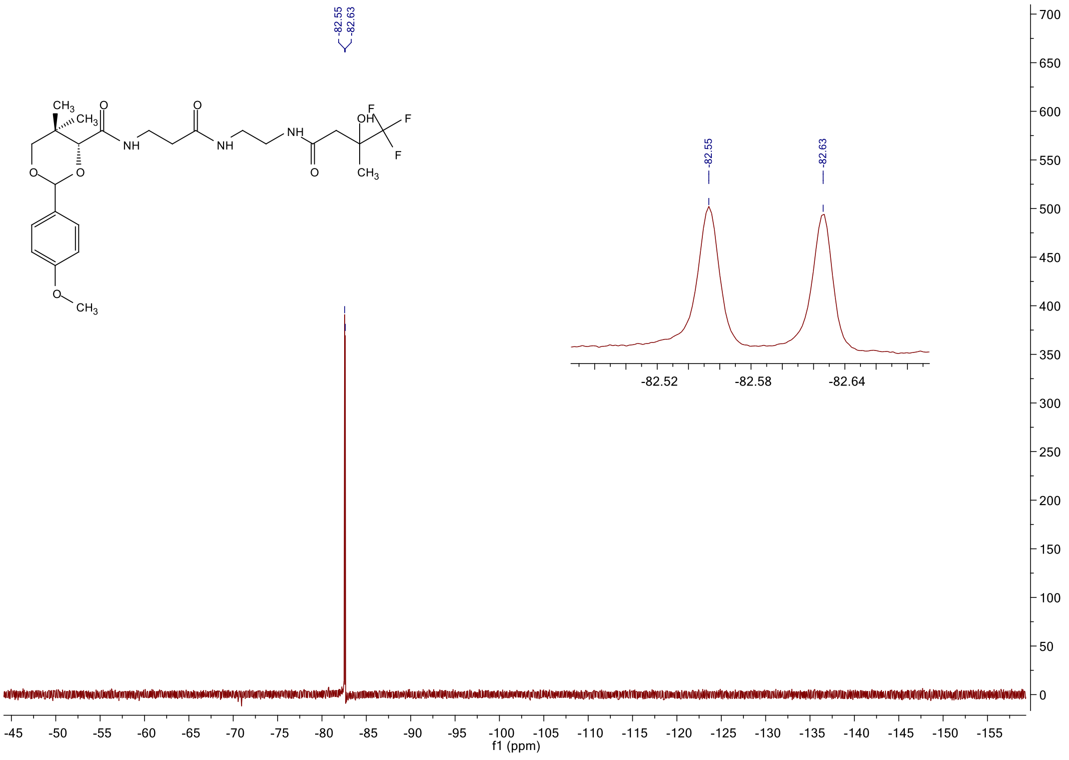

$^{1}$H NMR spectrum of **5e**

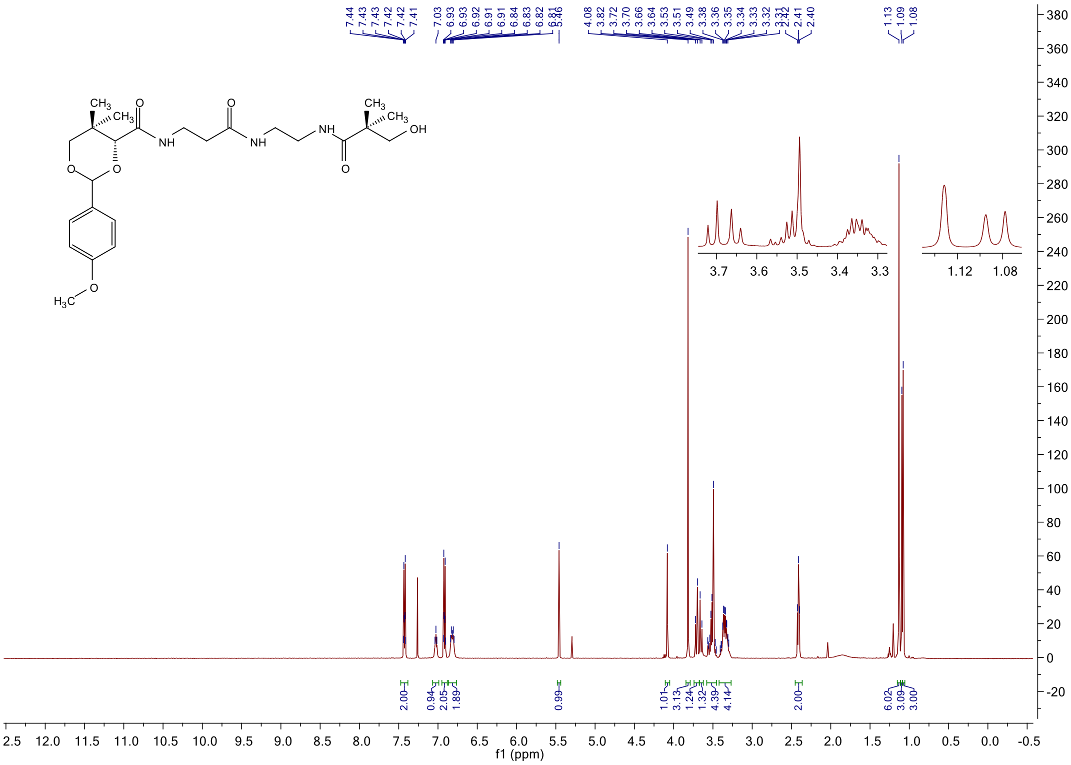

$^{13}$C NMR spectrum of **5e**

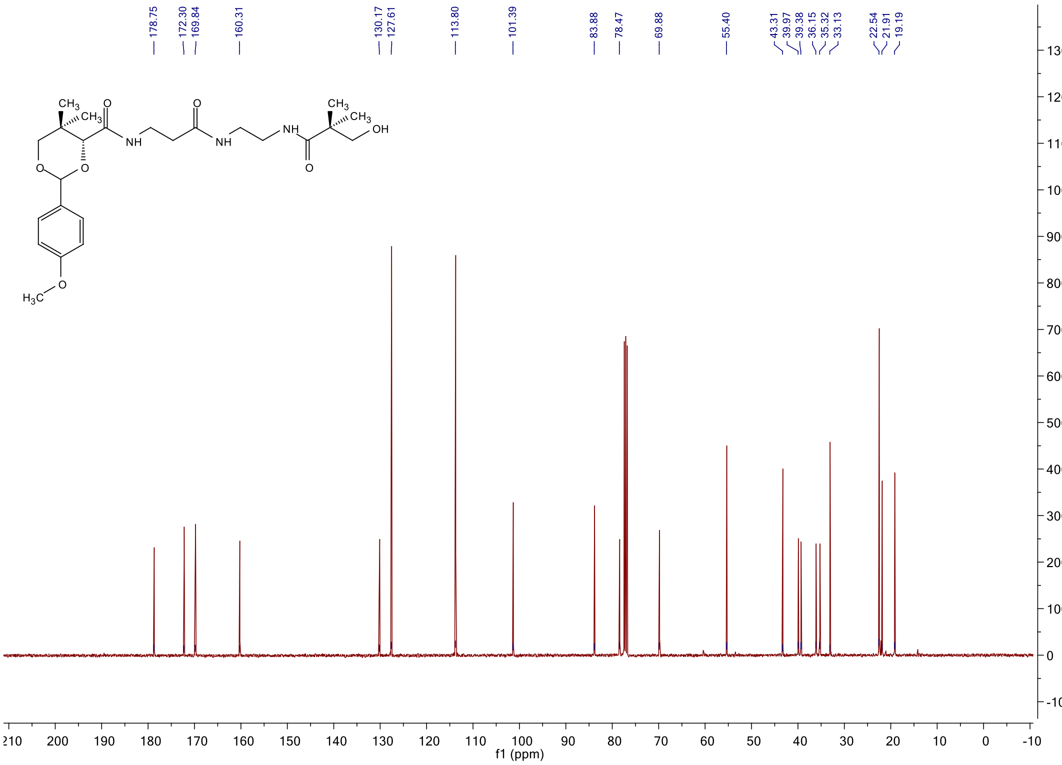

#### $^{1}$H NMR spectrum of **5f**

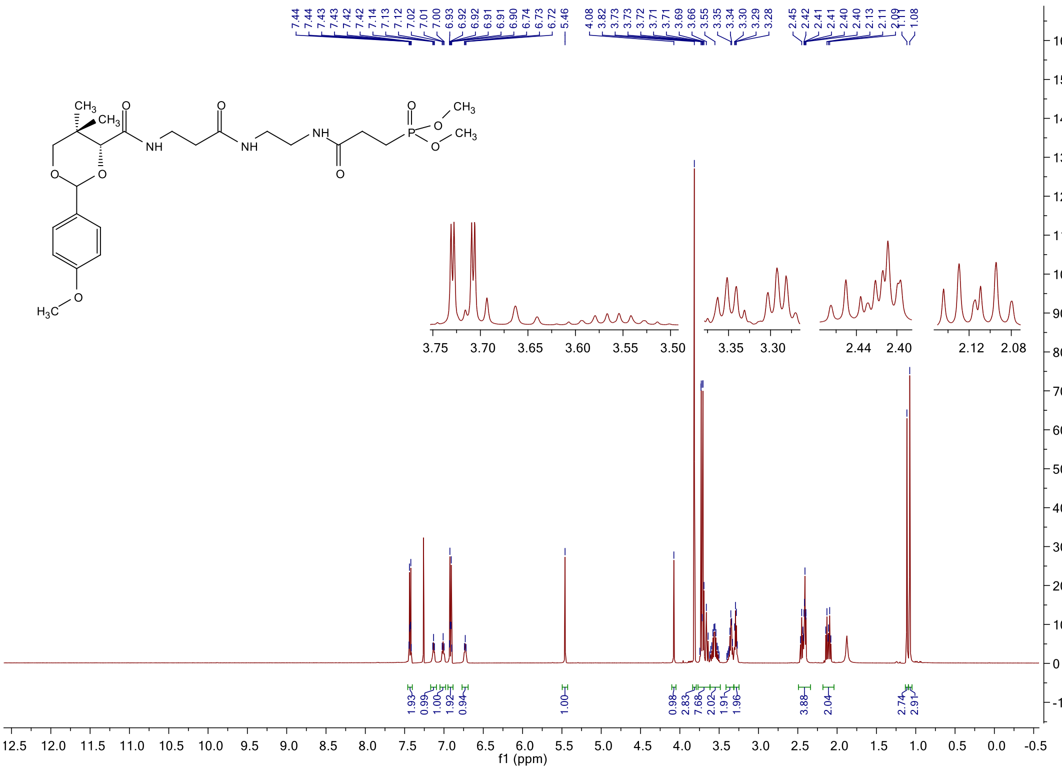

#### $^{13}$C NMR spectrum of **5f**

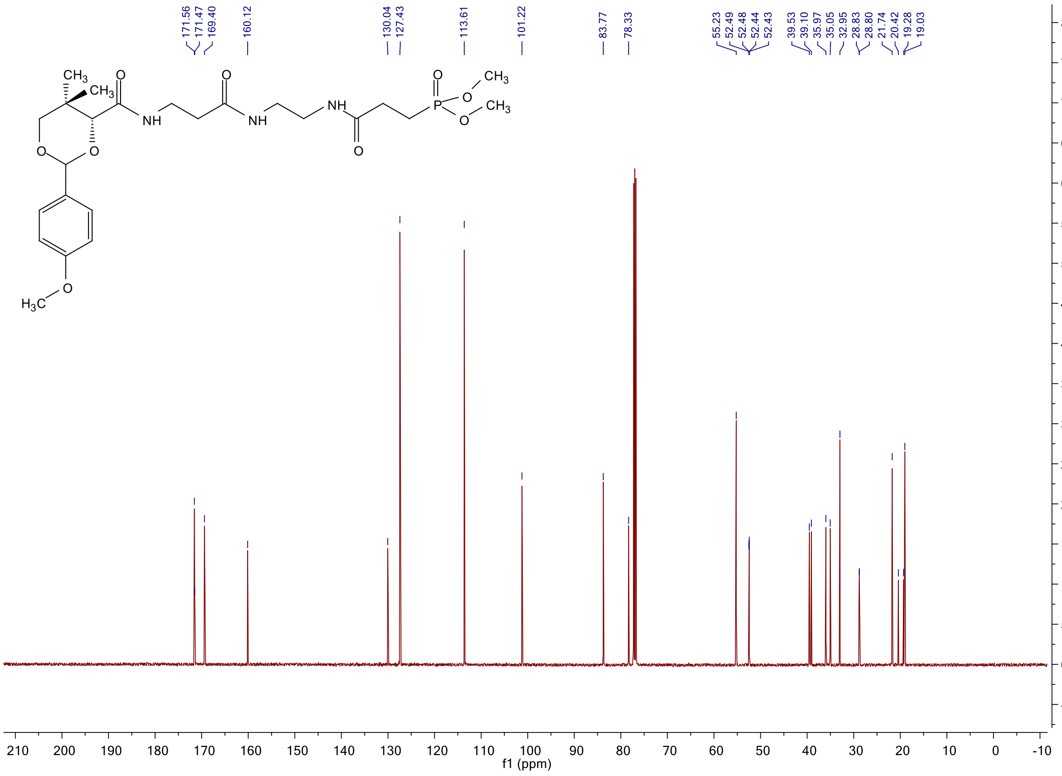

$^{31}$P NMR spectrum of **5f**

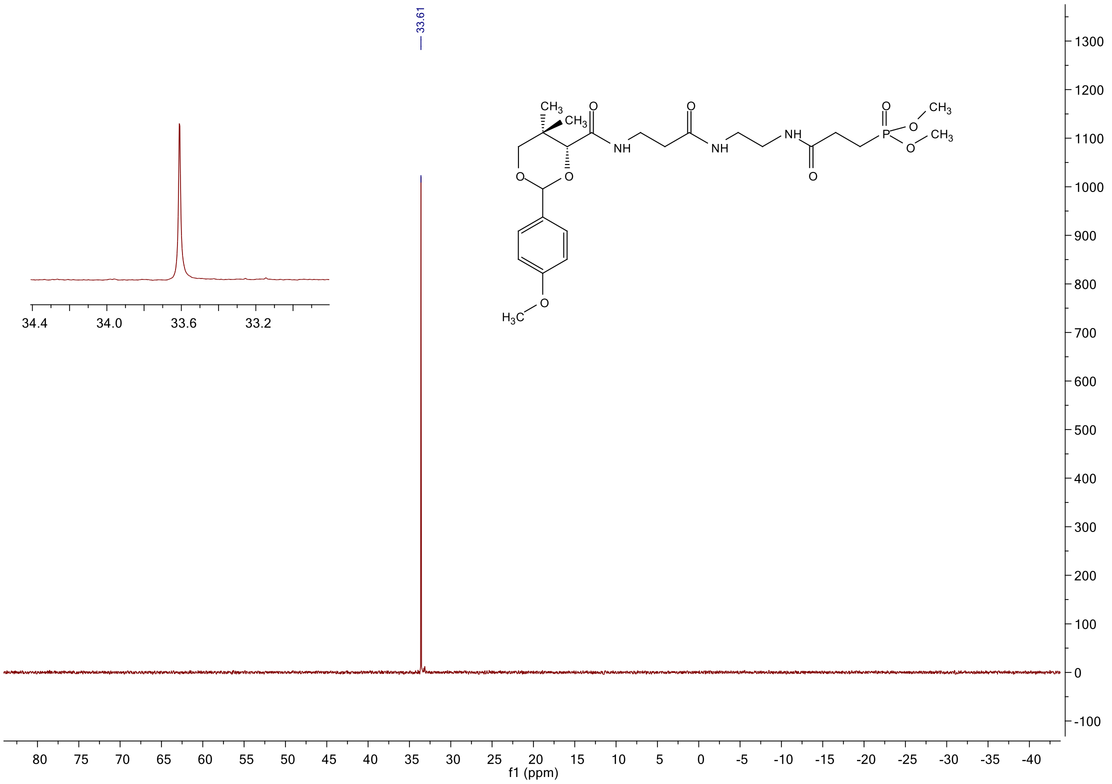

#### [1]H NMR spectrum of **5g**

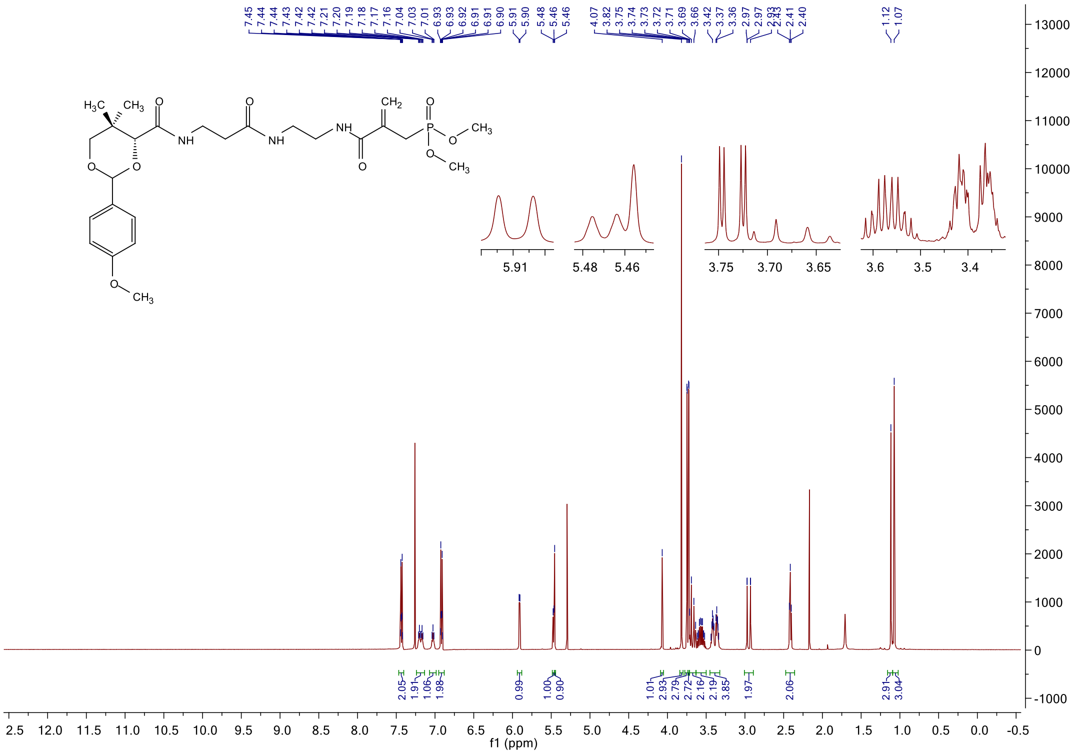

#### [13]C NMR spectrum of **5g**

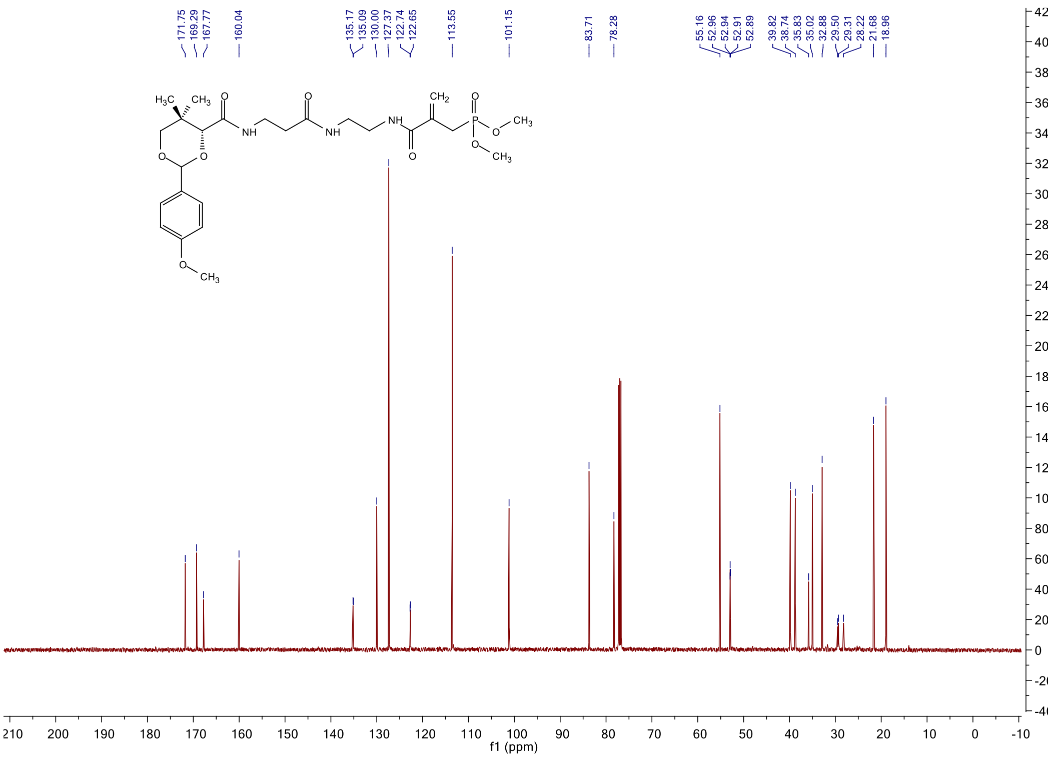

$^{31}$P NMR spectrum of **5g**

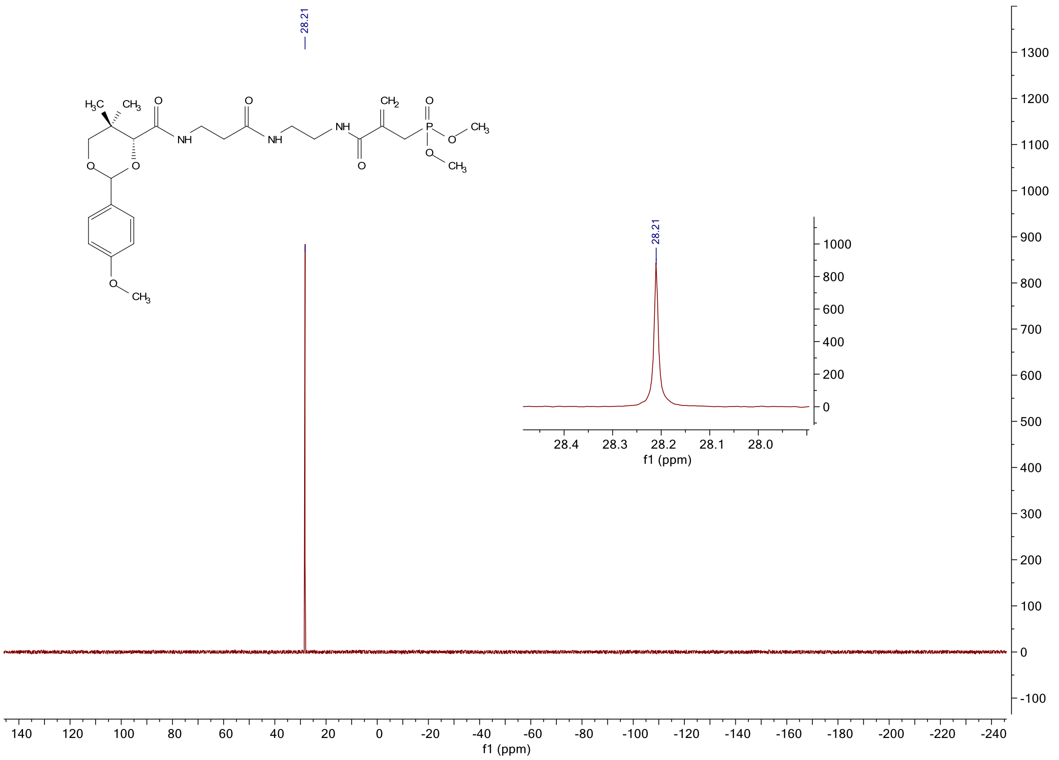

#### $^{1}$H NMR spectrum of **5h**

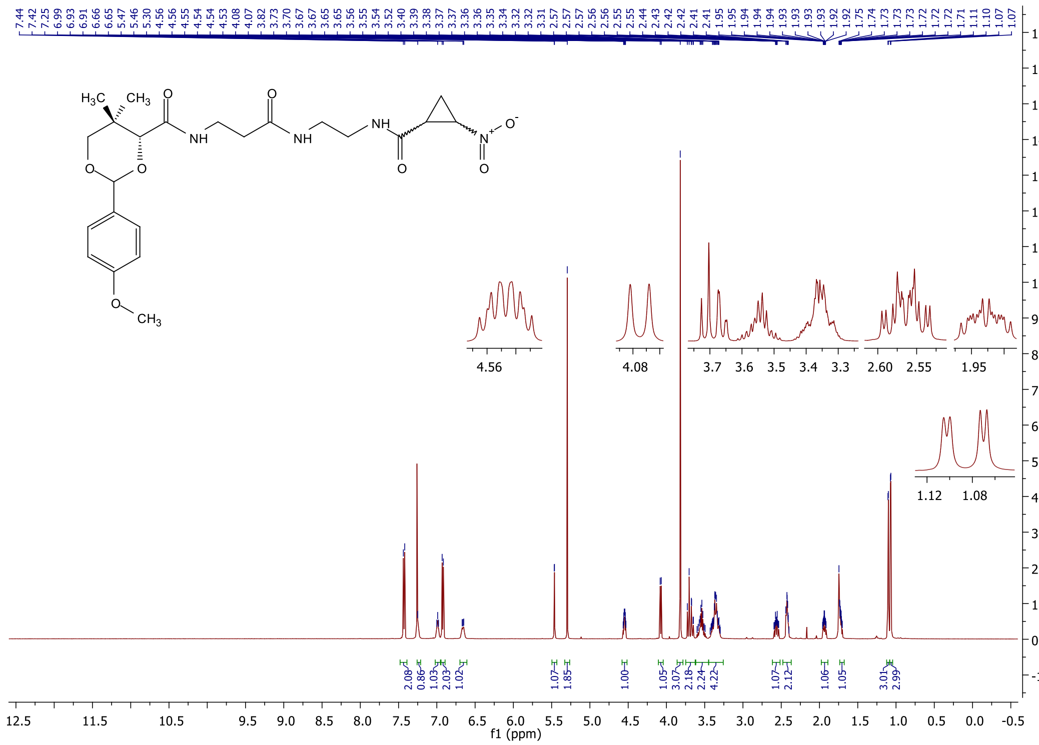

#### $^{13}$C NMR spectrum of **5h**

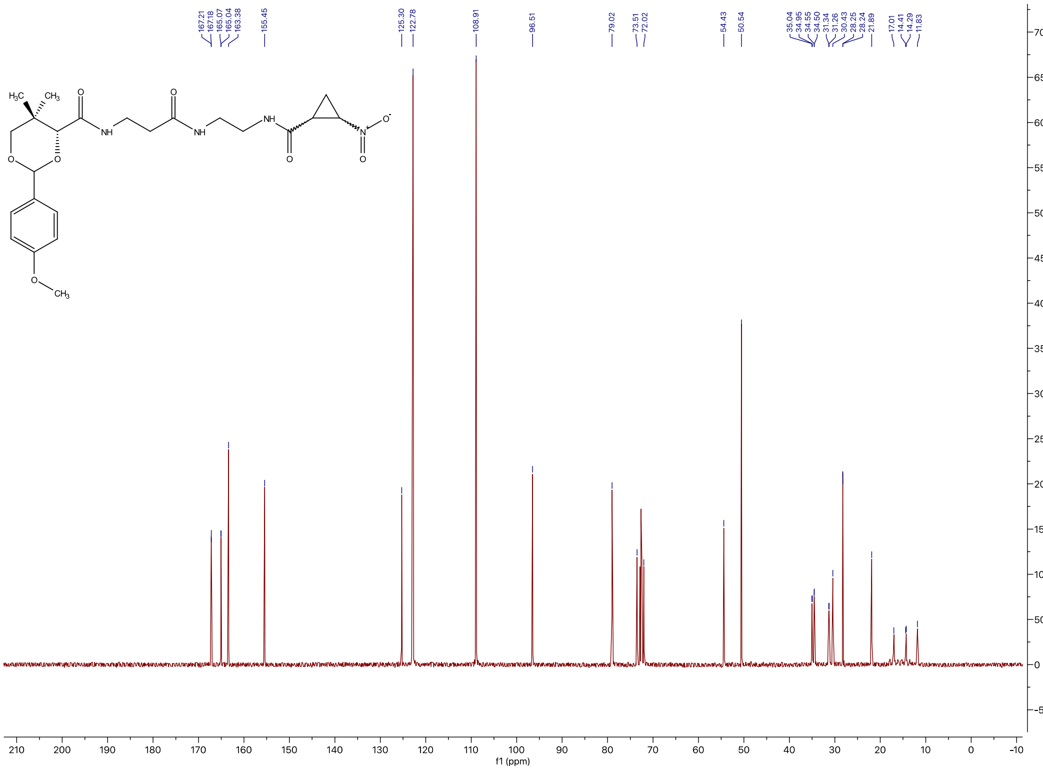

#### [1]H NMR spectrum of **5i**

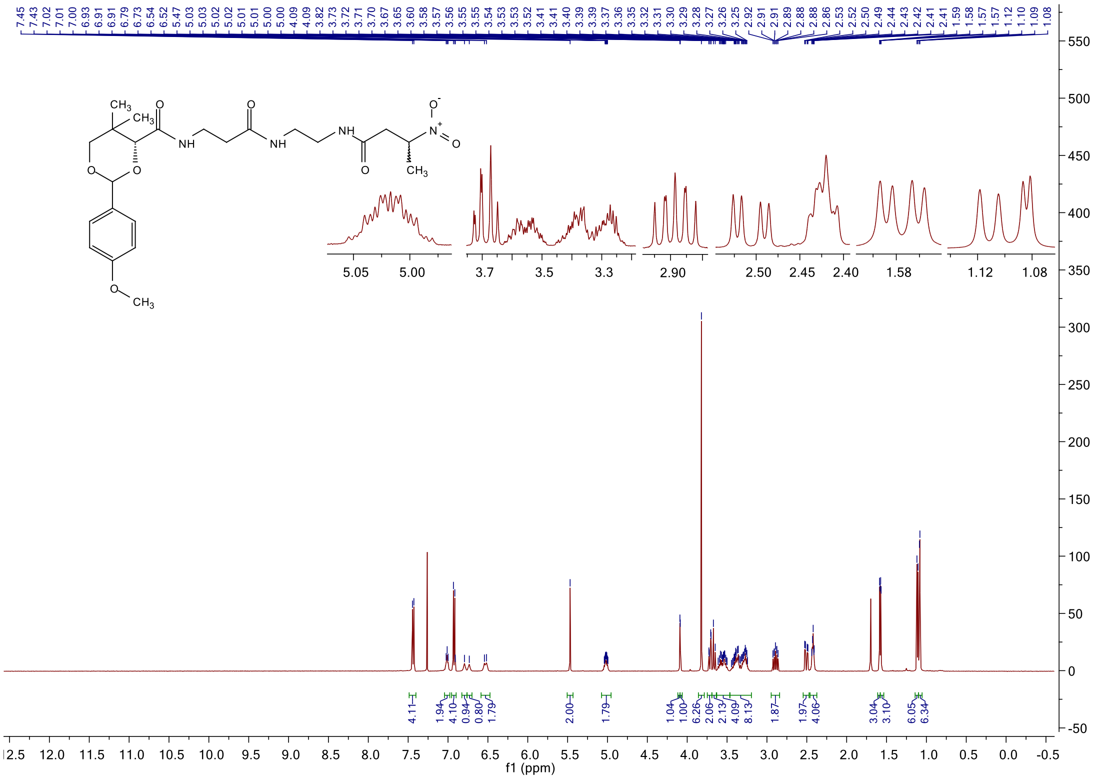

#### [13]C NMR spectrum of **5i**

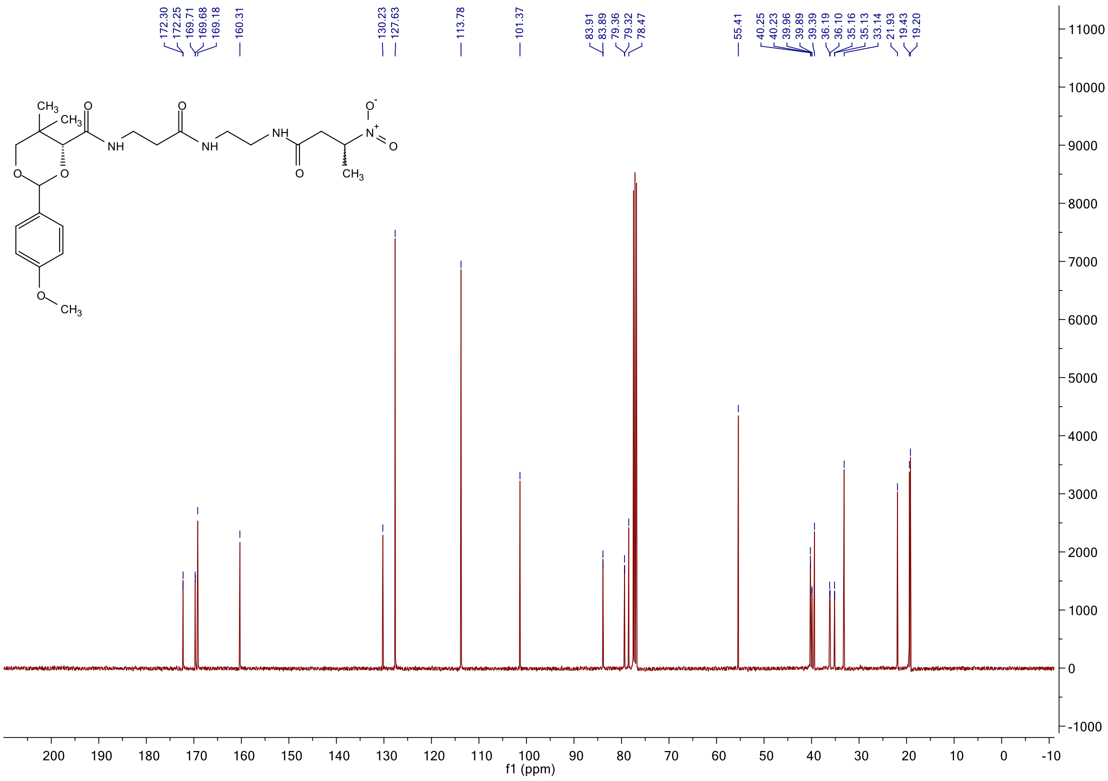

#### $^{1}$H NMR spectrum of **5k**

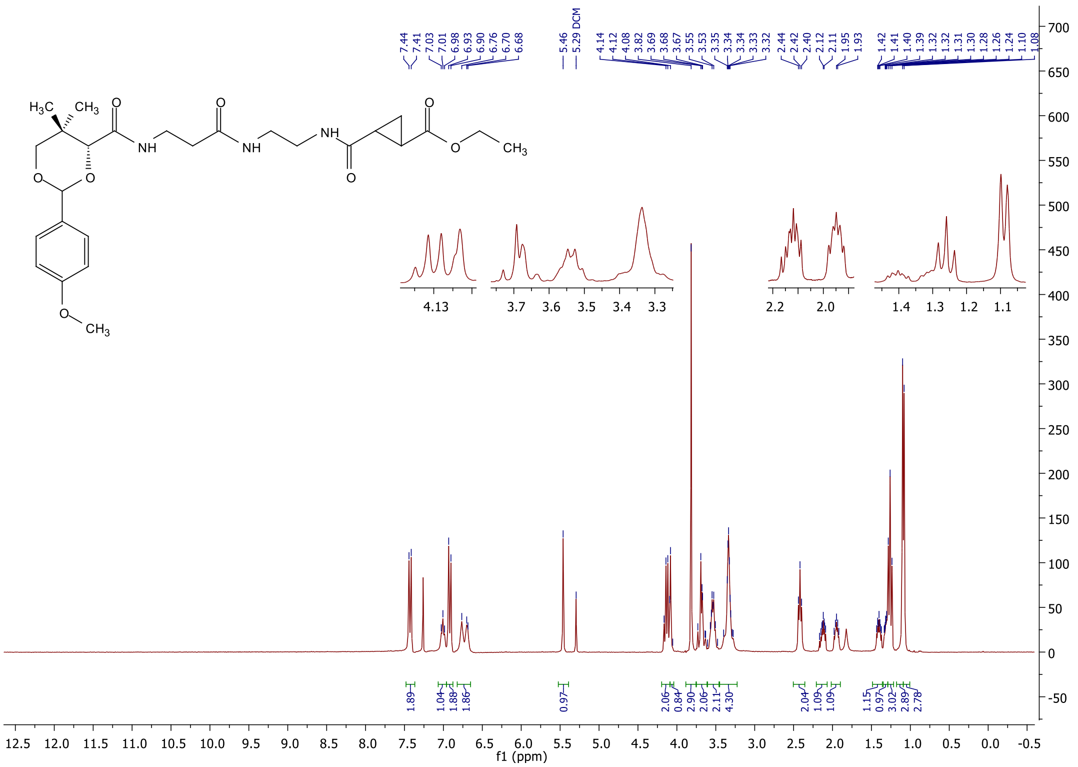

#### $^{13}$C NMR spectrum of **5k**

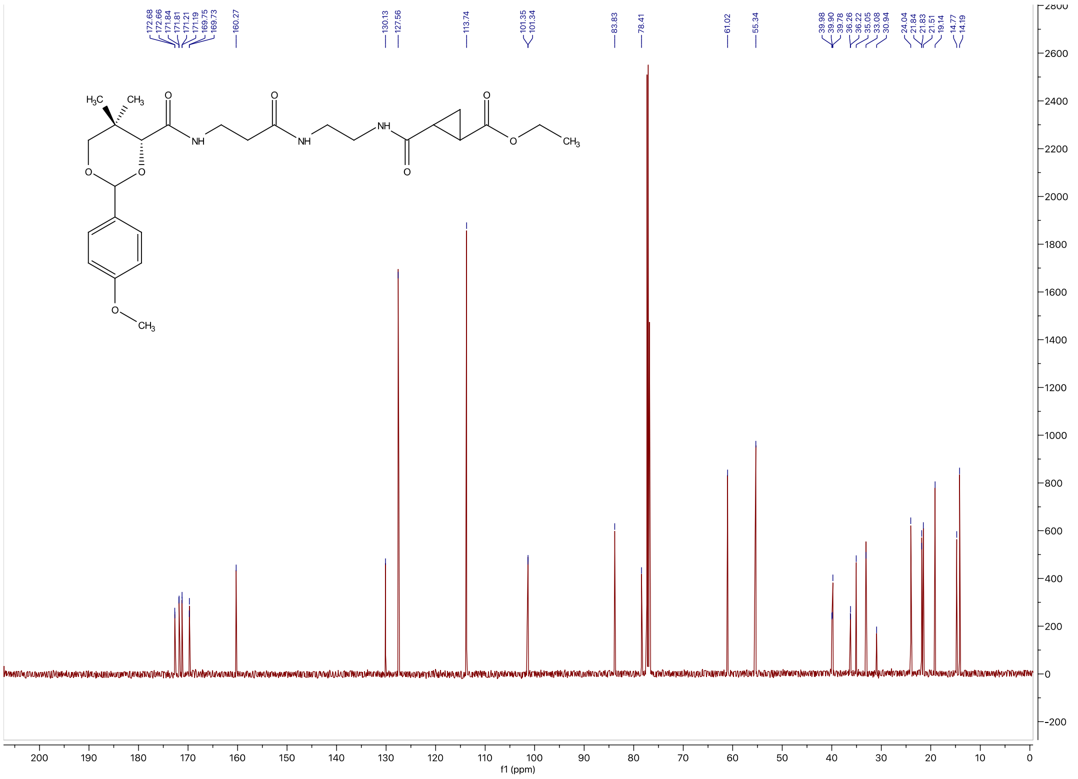

[1]H NMR spectrum of **5l**

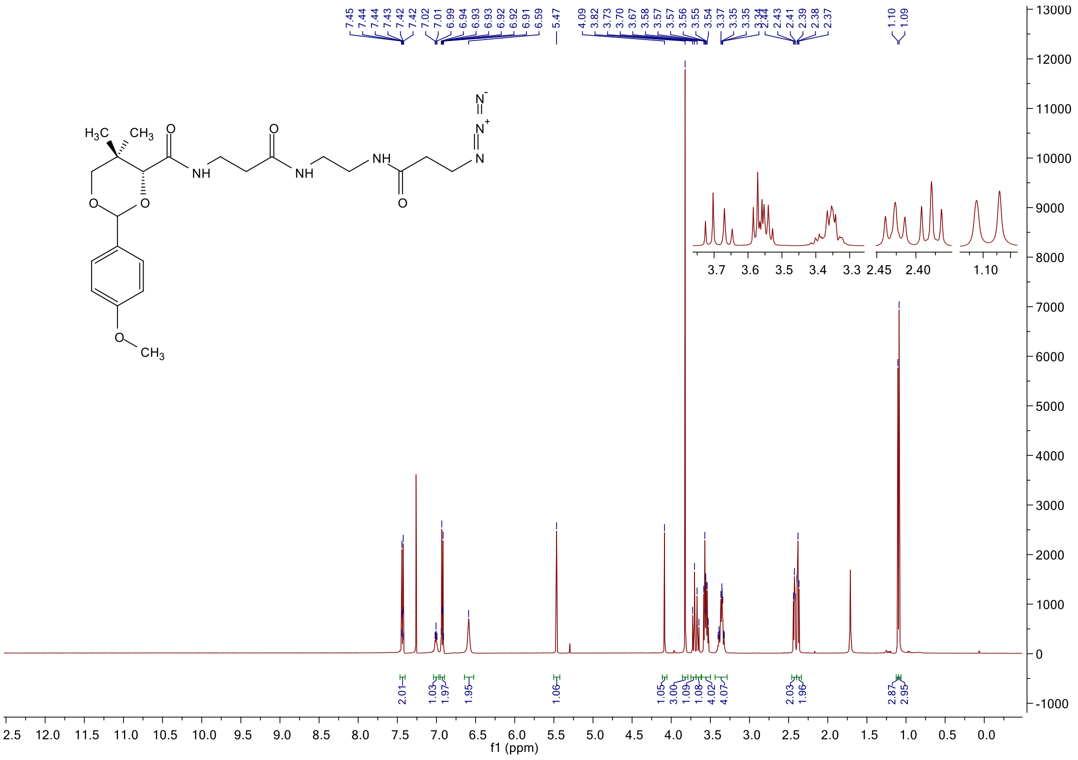

[13]C NMR spectrum of **5l**

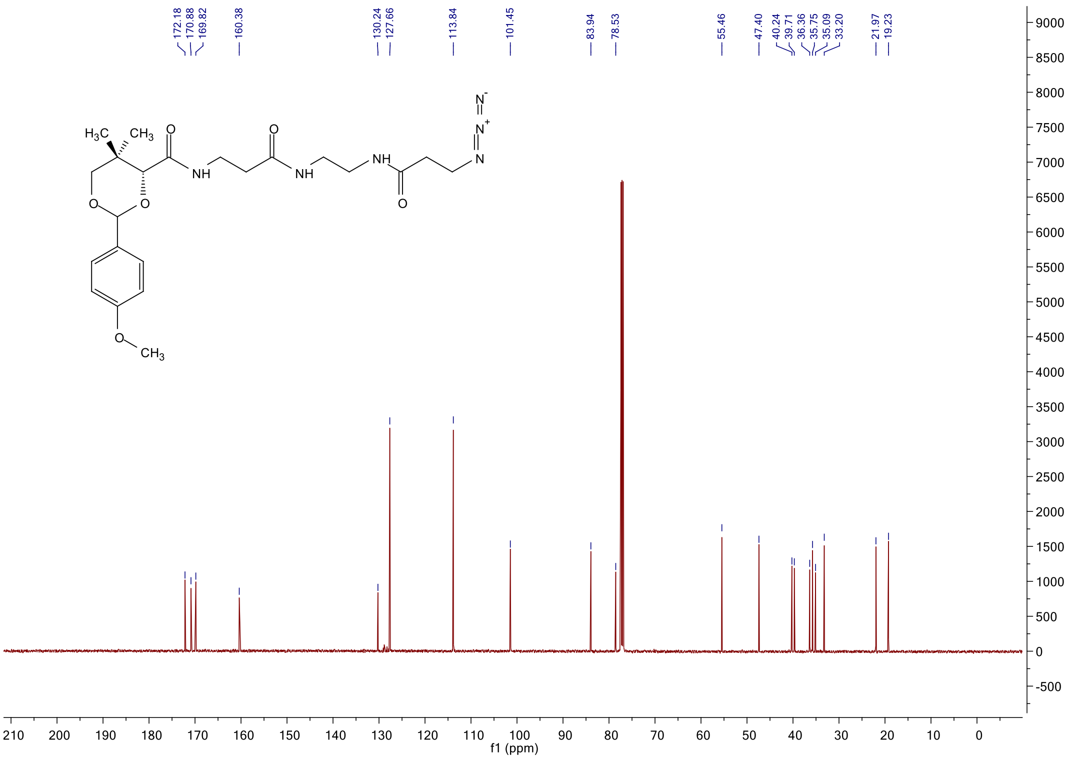

#### [1]H NMR spectrum of **5m**

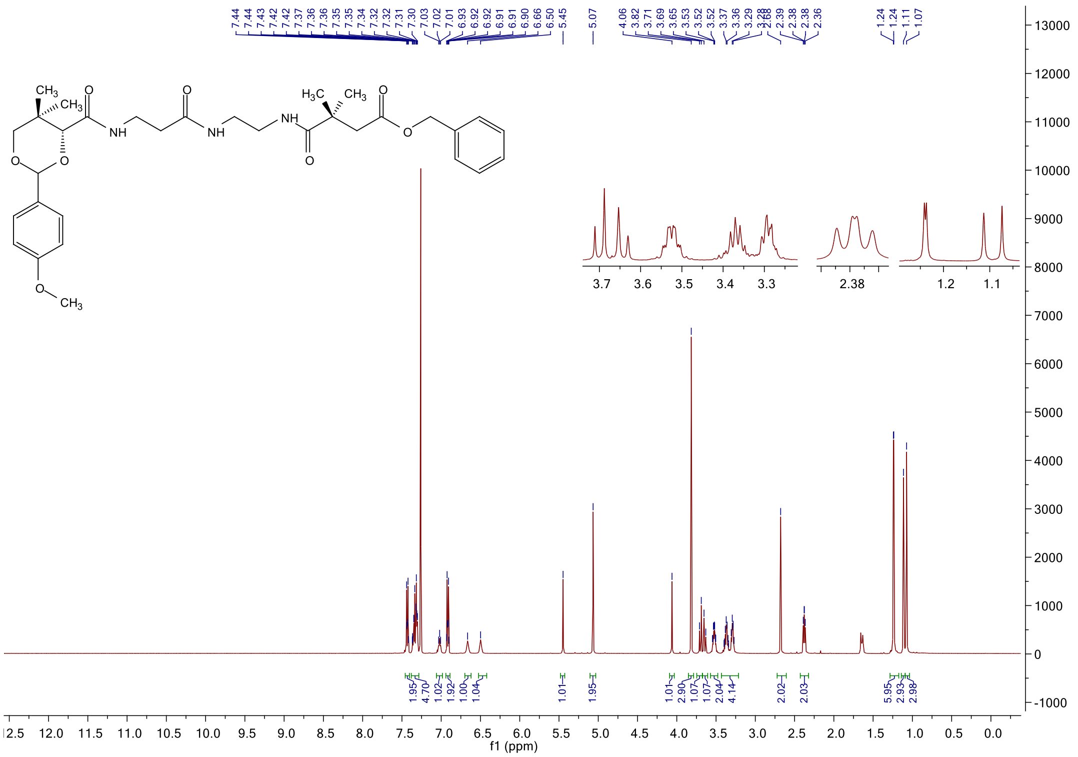

#### [13]C NMR spectrum of **5m**

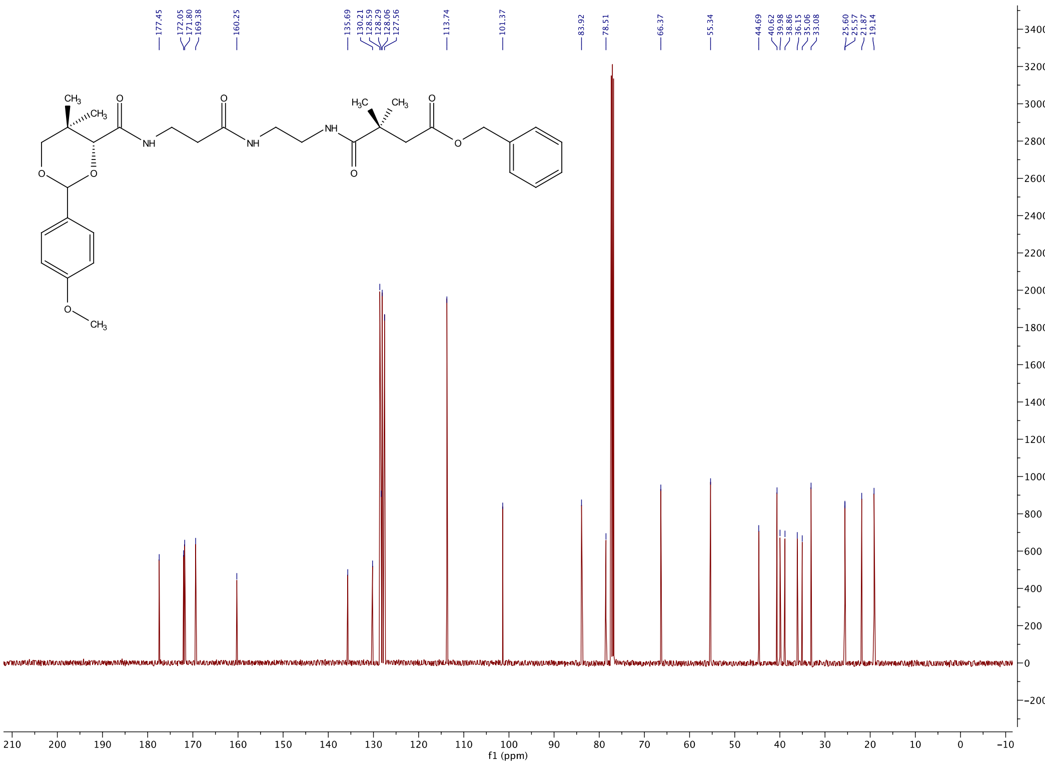

[1]H NMR spectrum of **5n**

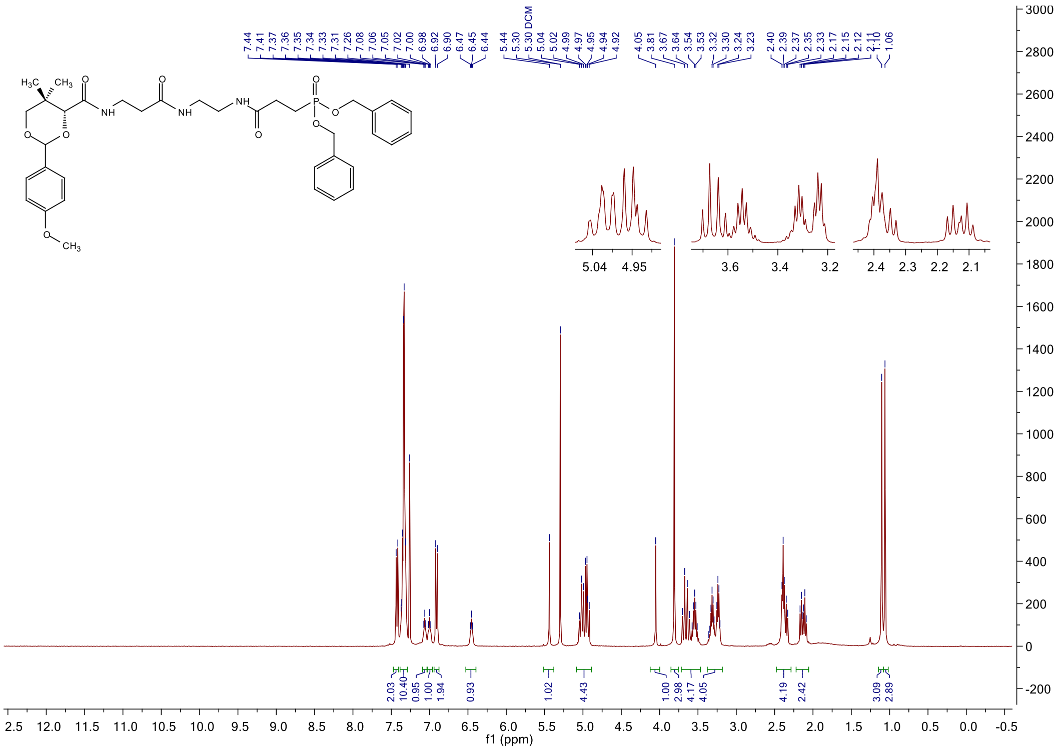

[13]C NMR spectrum of **5n**

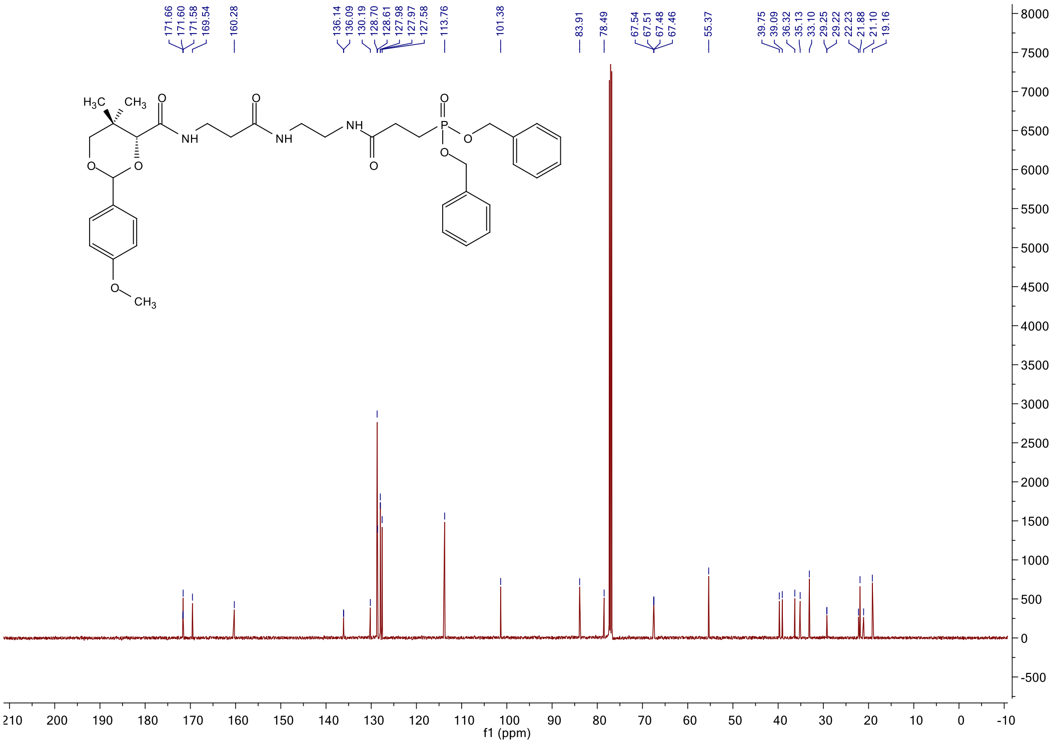

[31]P NMR spectrum of **5n**

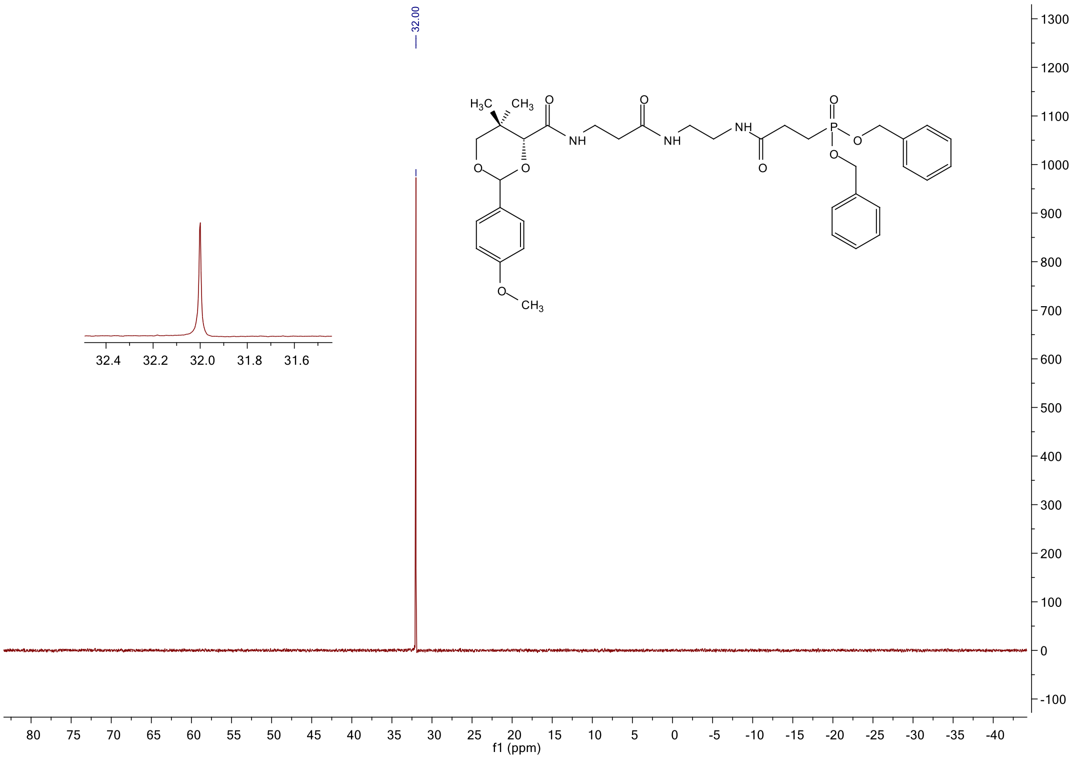

#### $^{1}$H NMR spectrum of **8k**

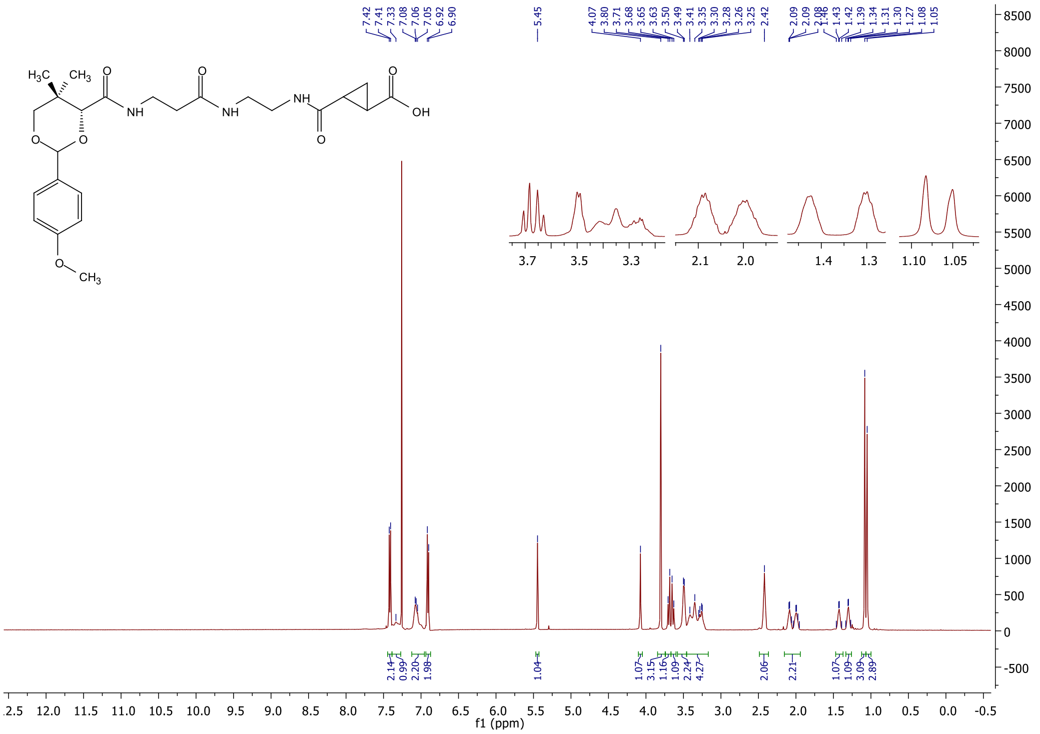

#### $^{13}$C NMR spectrum of **8k**

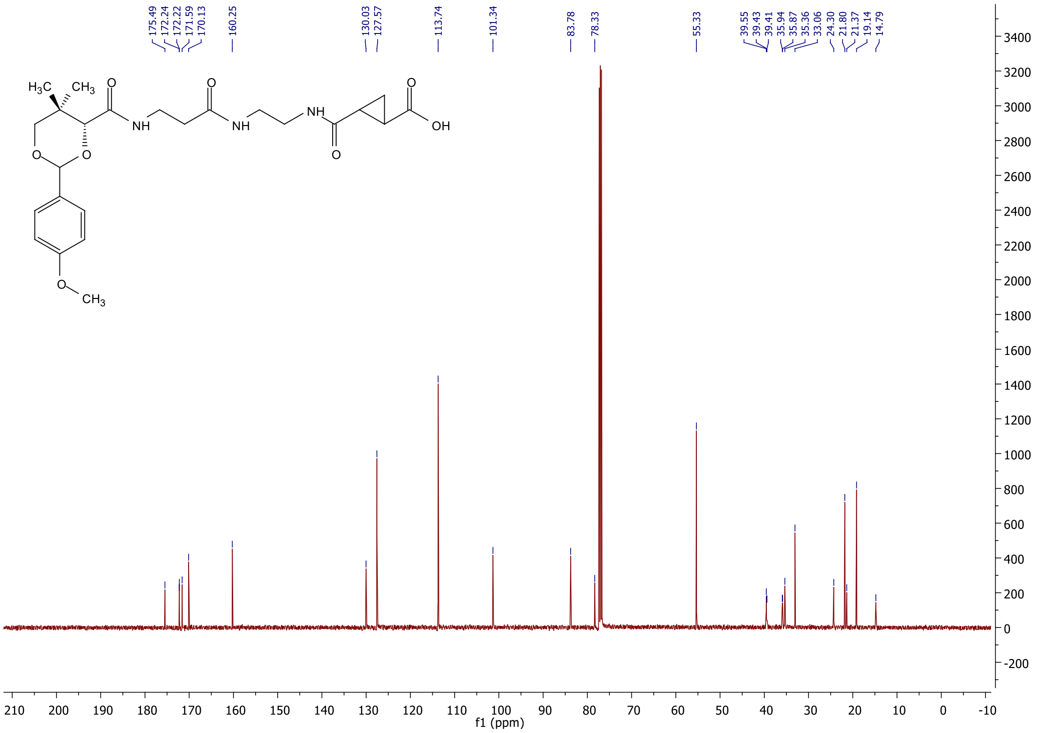

#### $^1$H NMR spectrum of **11**

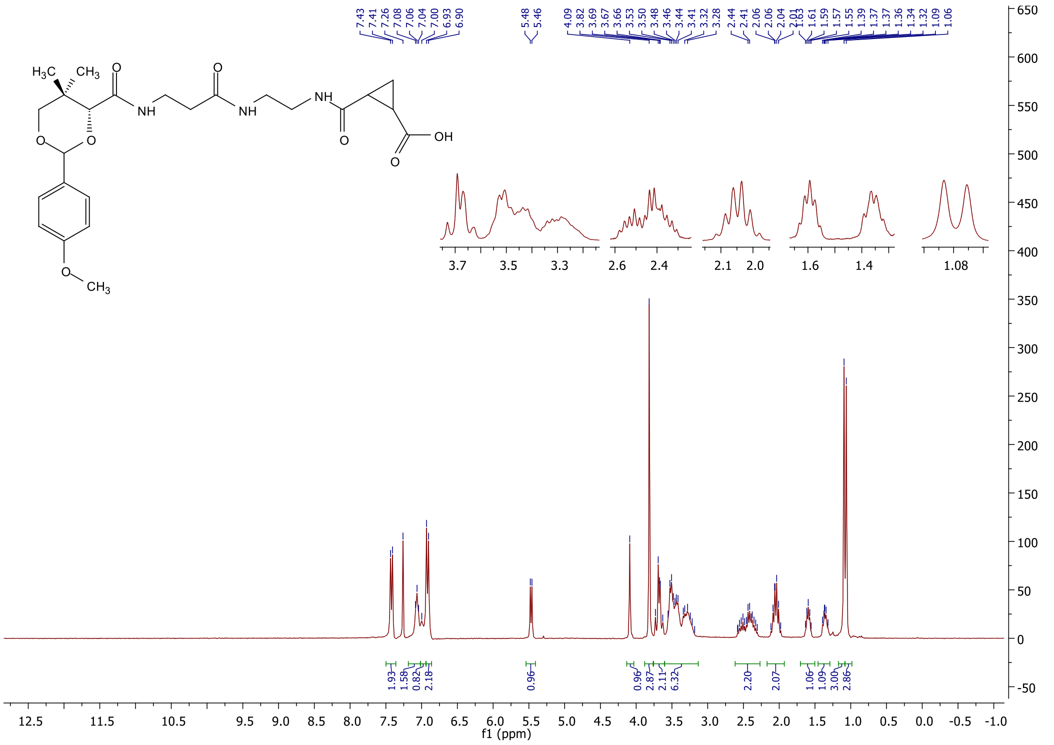

#### $^{13}$C NMR spectrum of **11**

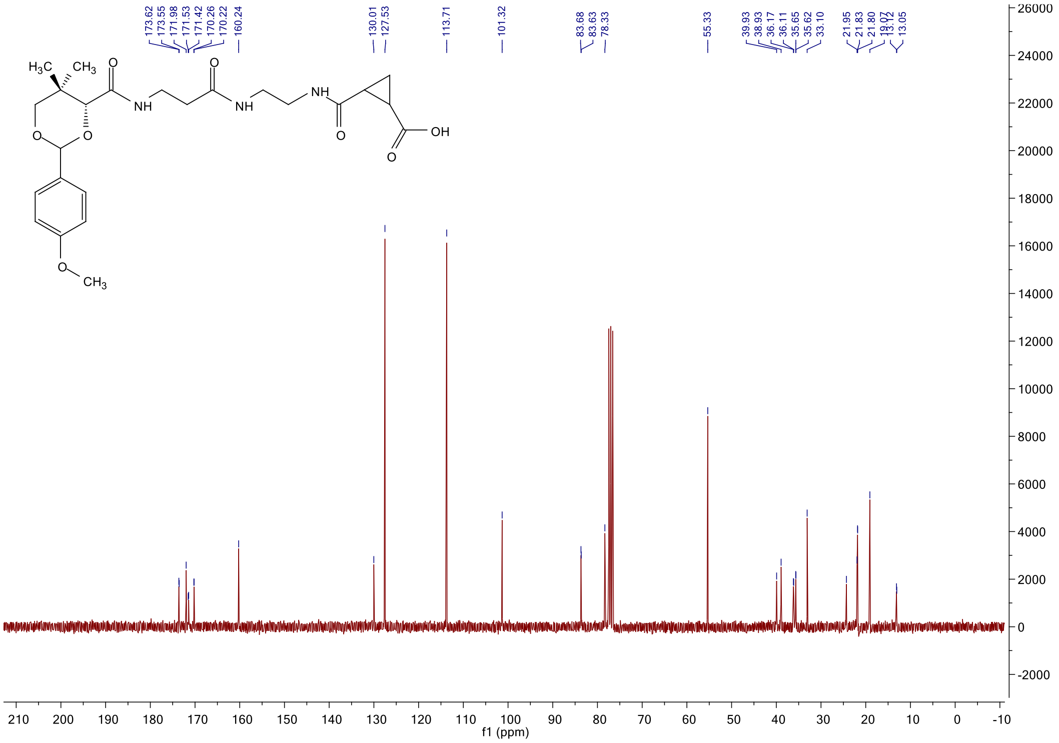

[1]H NMR spectrum of **13**

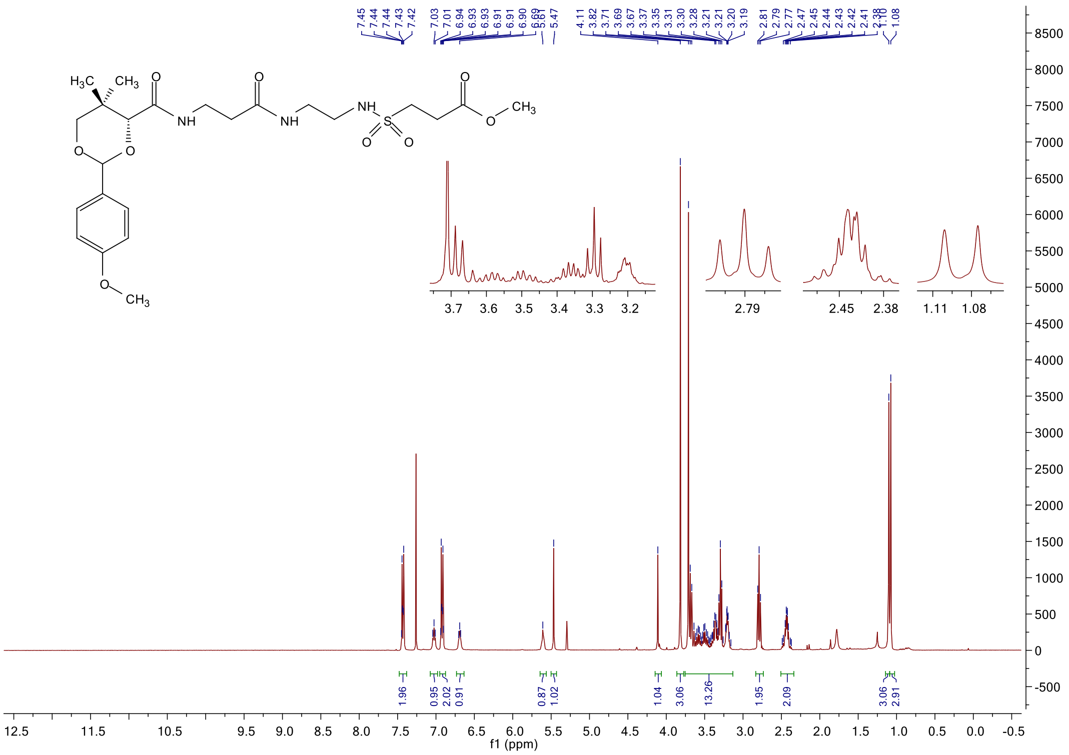

[13]C NMR spectrum of **13**

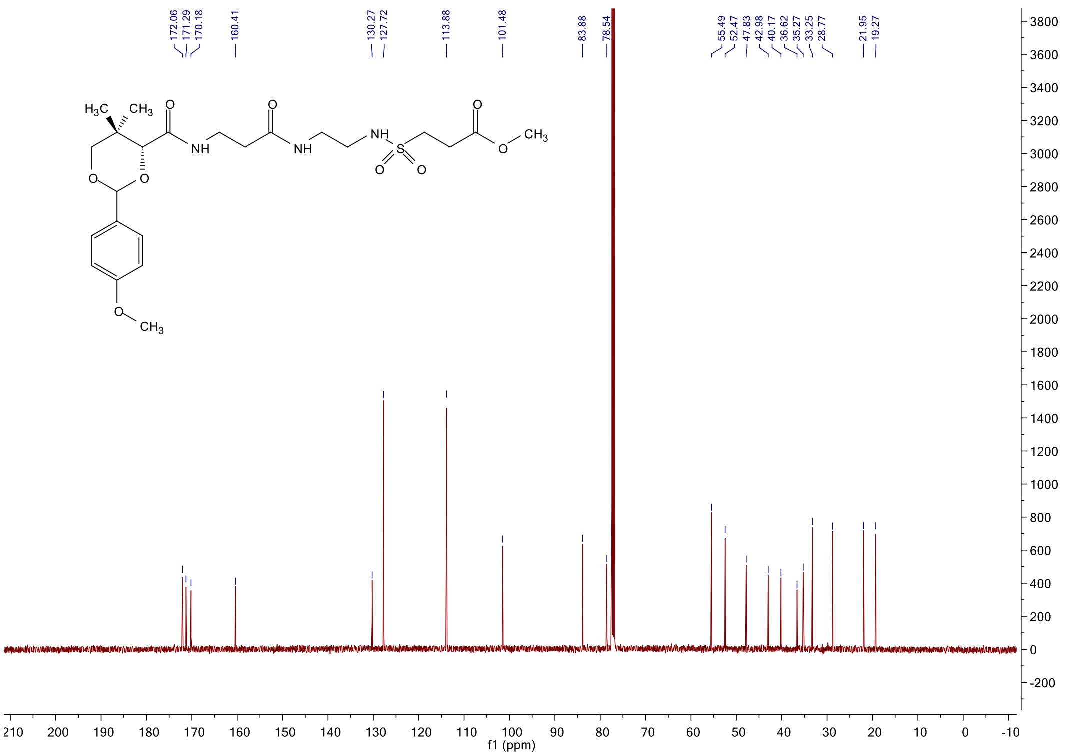

<sup></sup>

#### $^{1}$H NMR spectrum of **14**

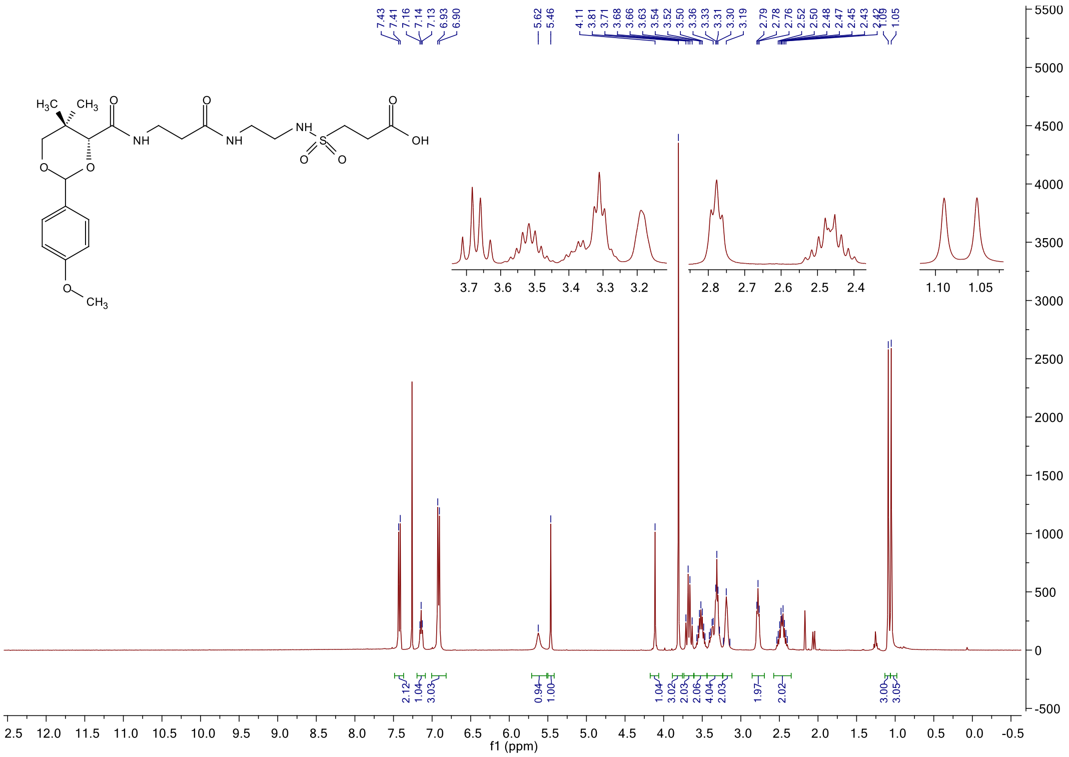

#### $^{13}$C NMR spectrum of **14**

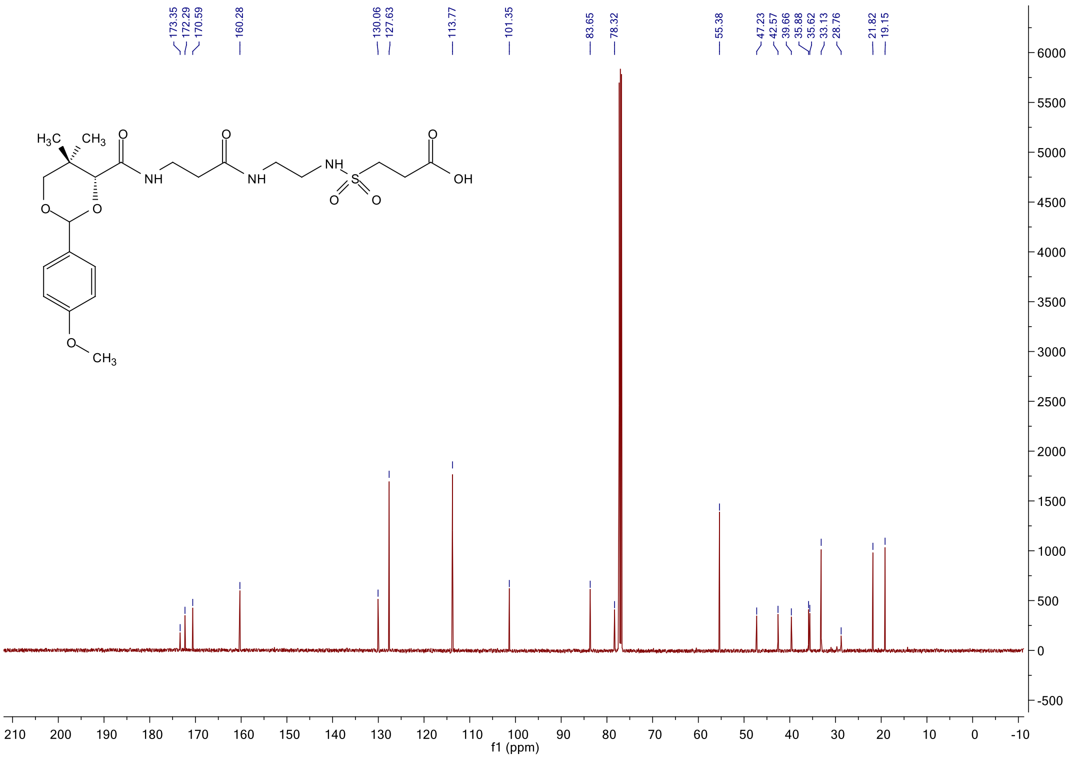

### $^1$H NMR spectrum of **21a**

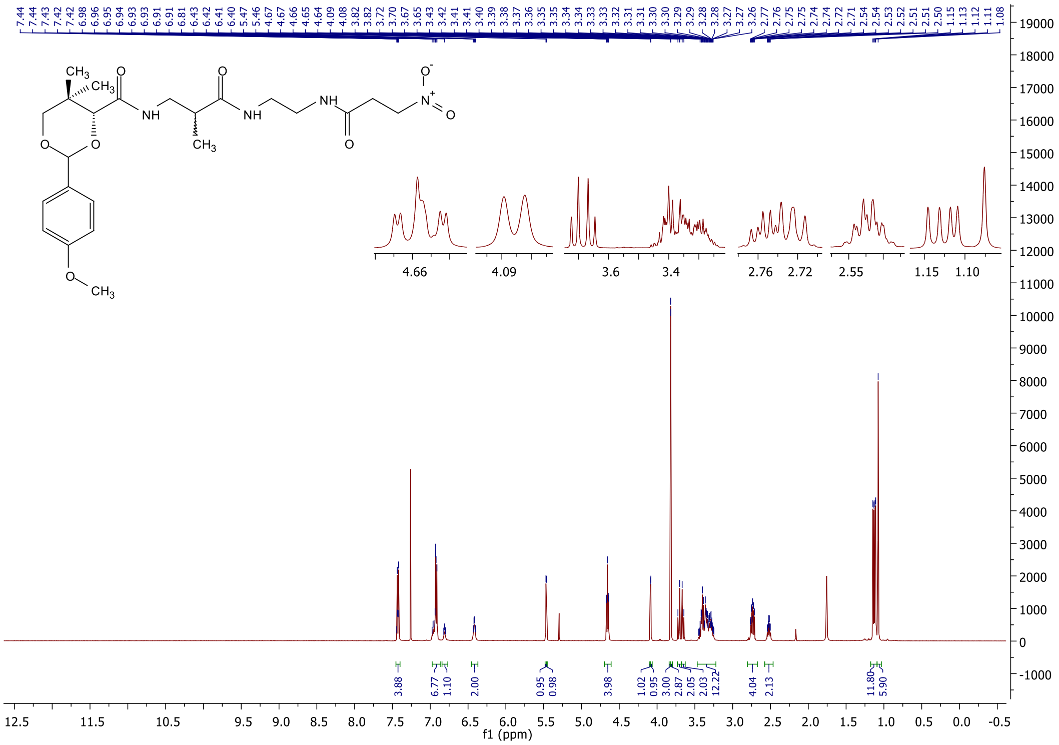

### $^{13}$C NMR spectrum of **21a**

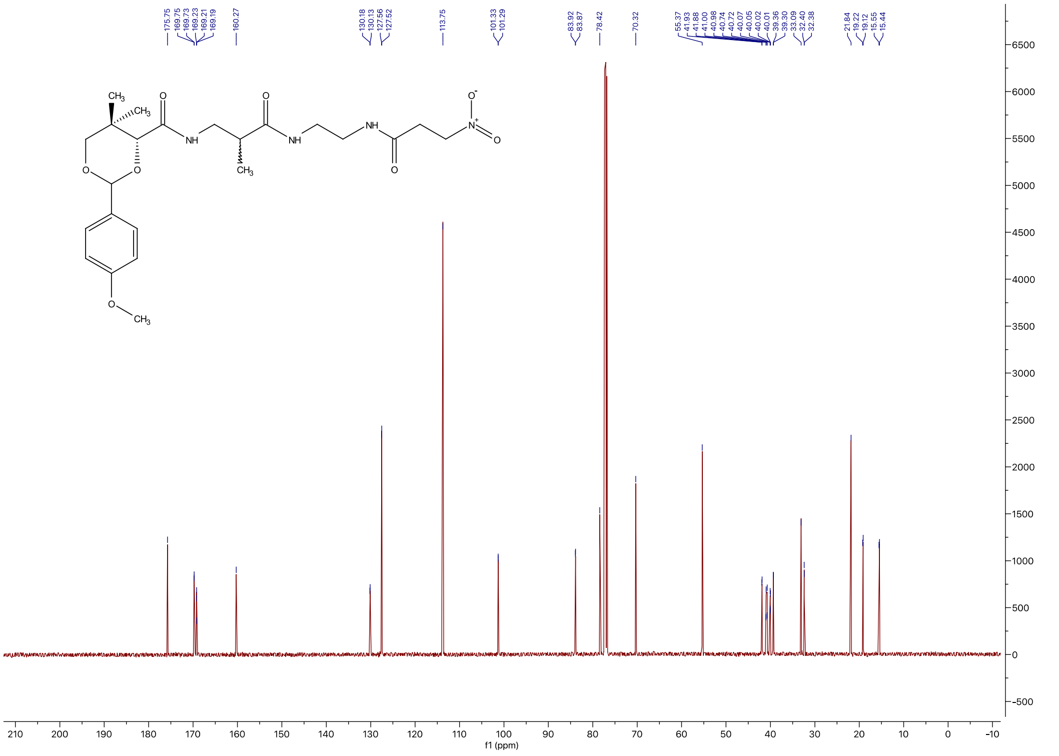

#### $^{1}$H NMR spectrum of **21b**

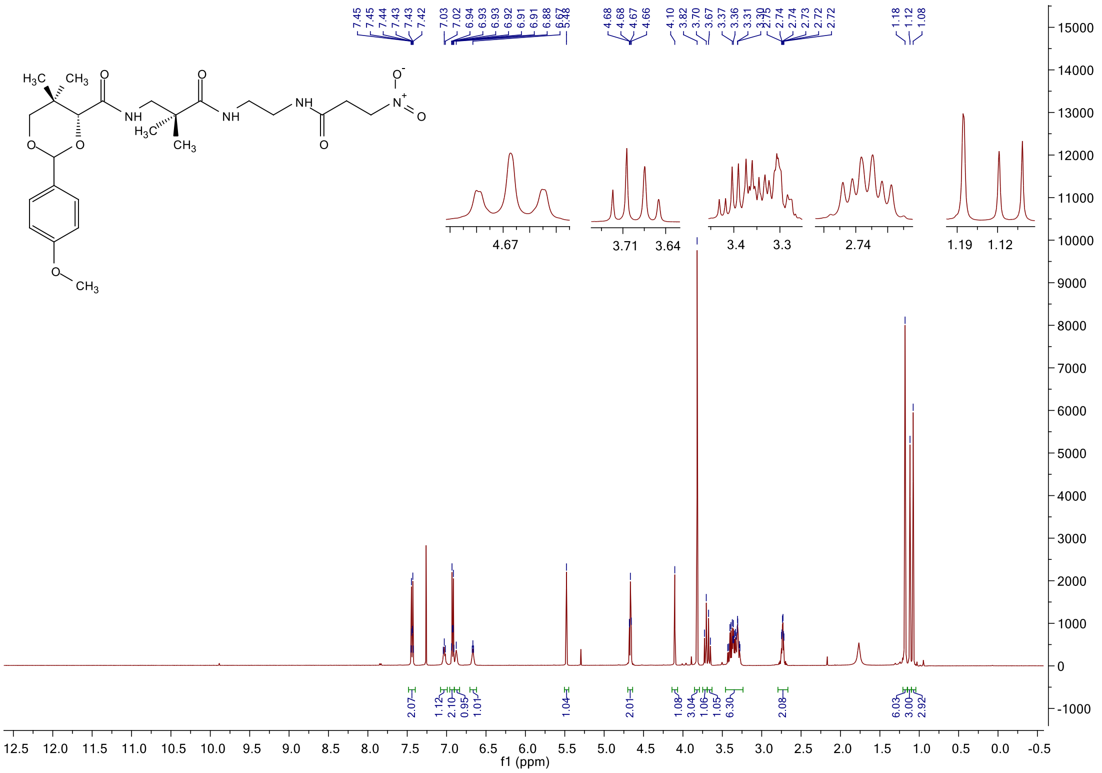

#### $^{13}$C NMR spectrum of **21b**

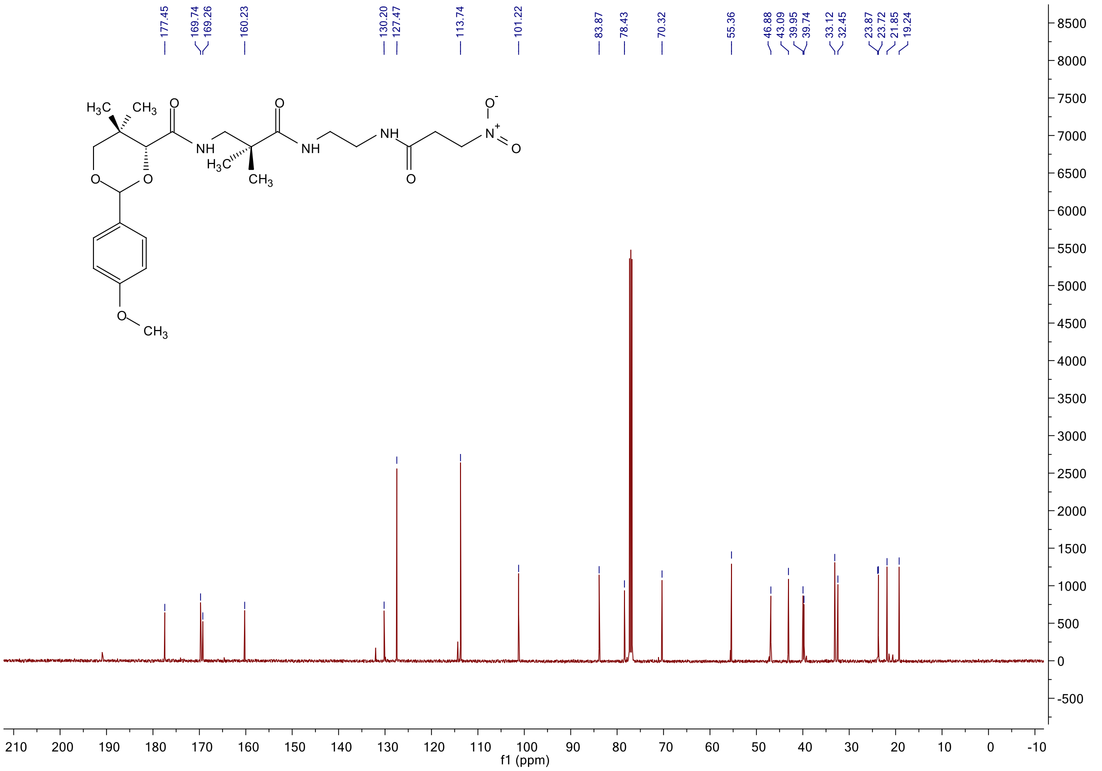

#### $^{1}$H NMR spectrum of **21c**

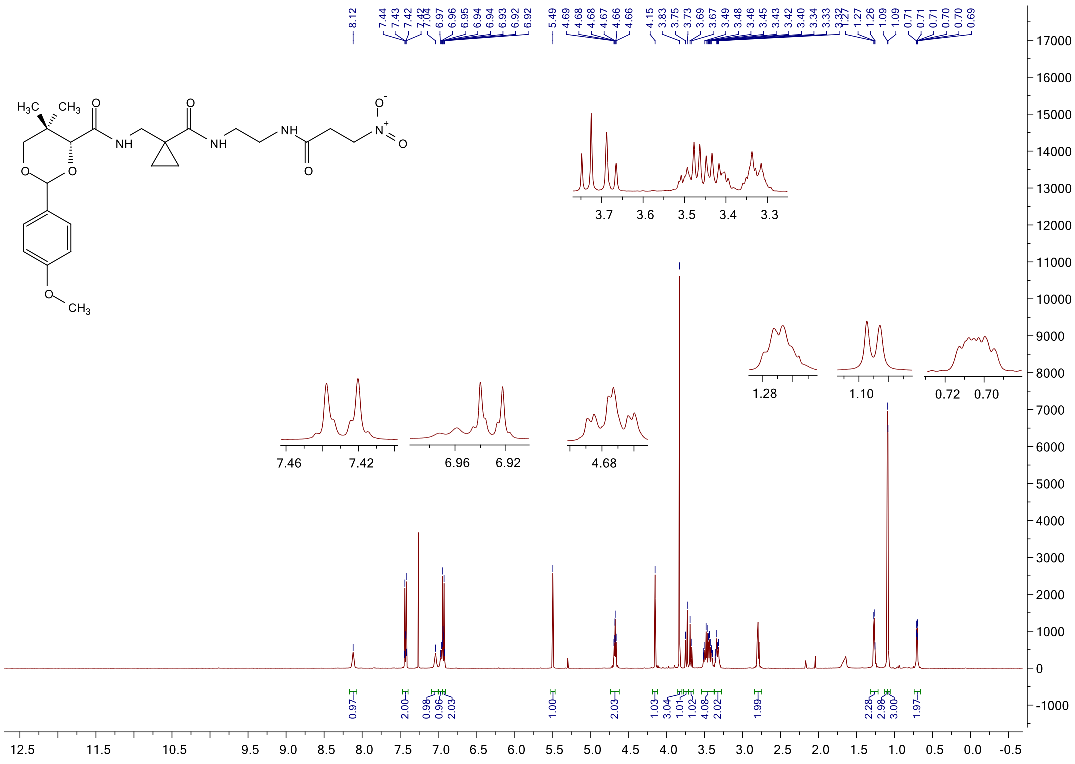

#### $^{13}$C NMR spectrum of **21c**

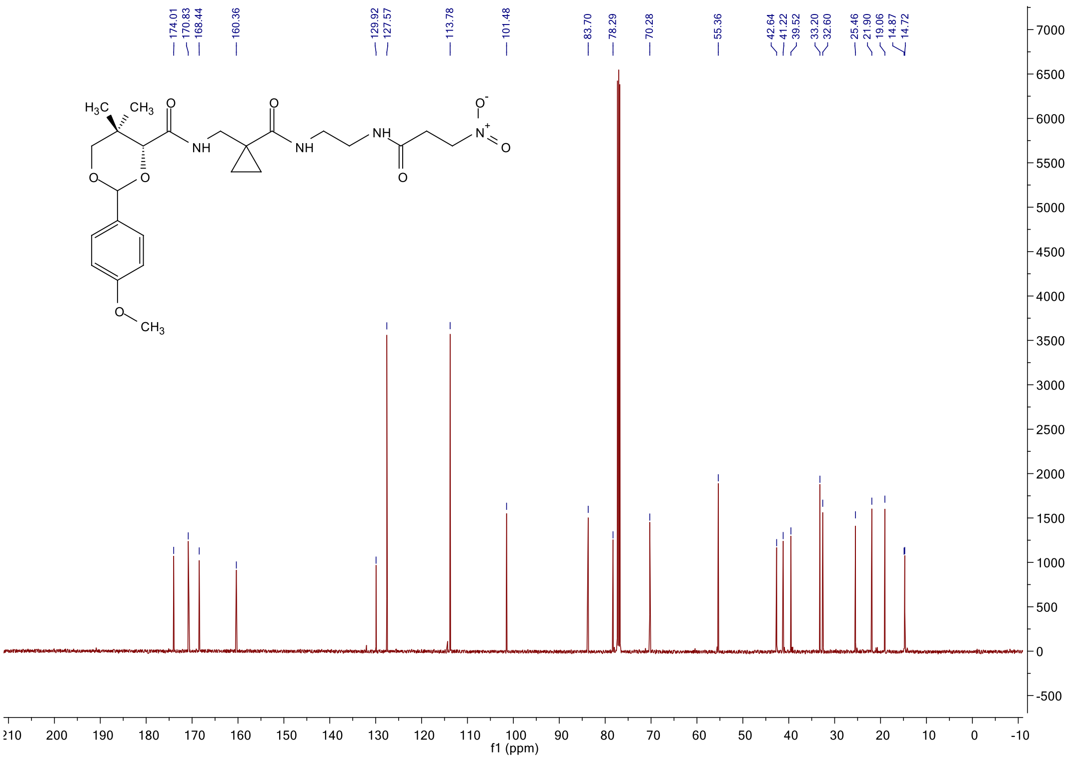

#### [1]H NMR spectrum of **6a**

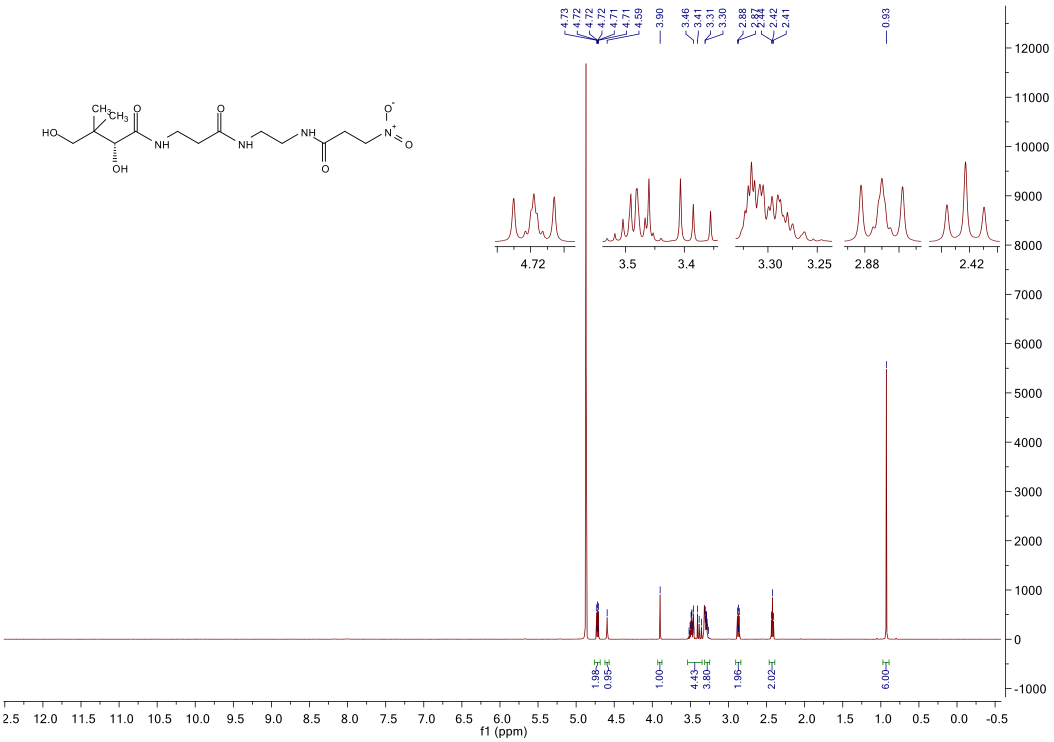

#### [1]H NMR spectrum of **6b**

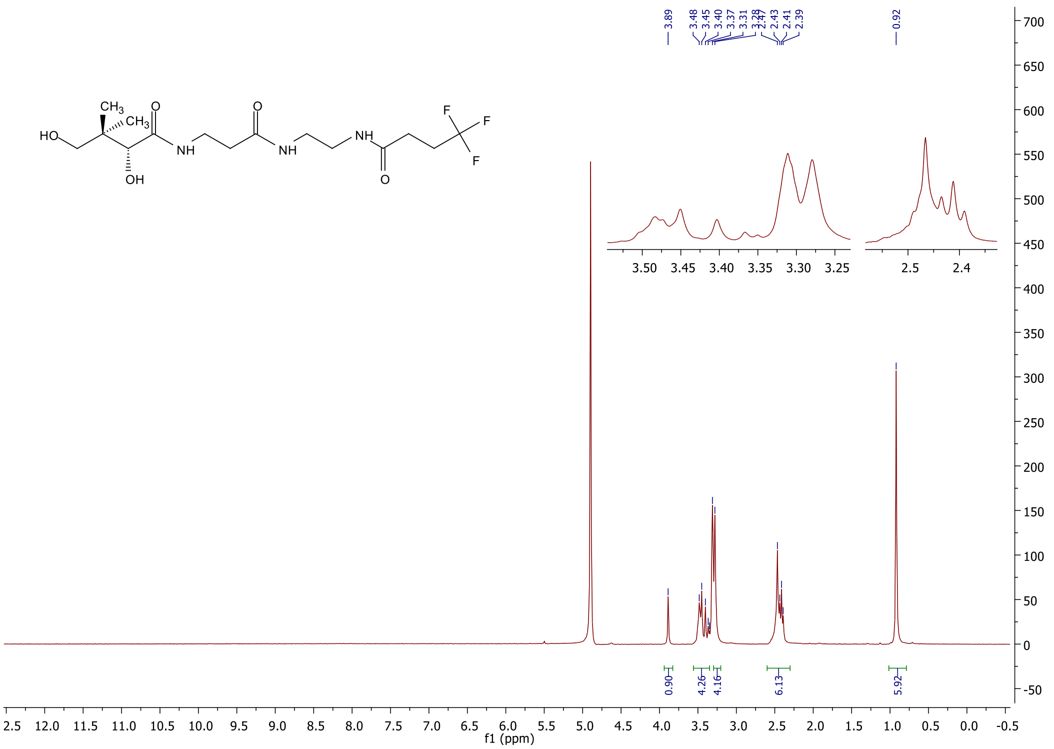

$^{13}$C NMR spectrum of **6b**

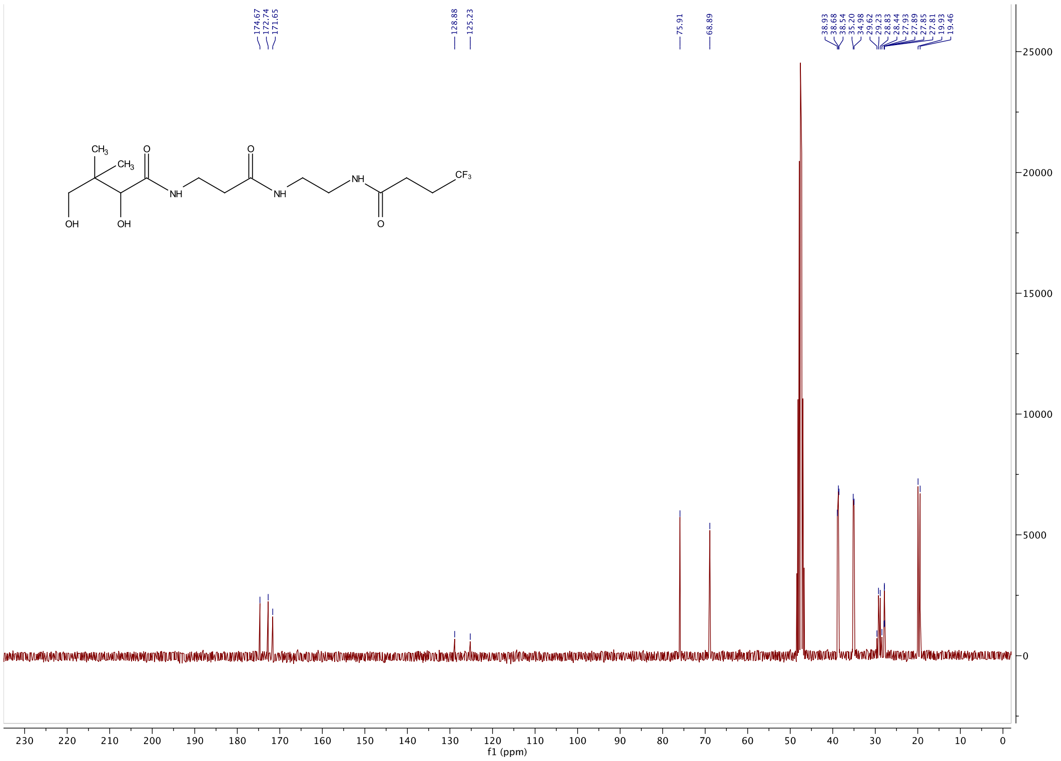

$^{19}$F NMR spectrum of **6b**

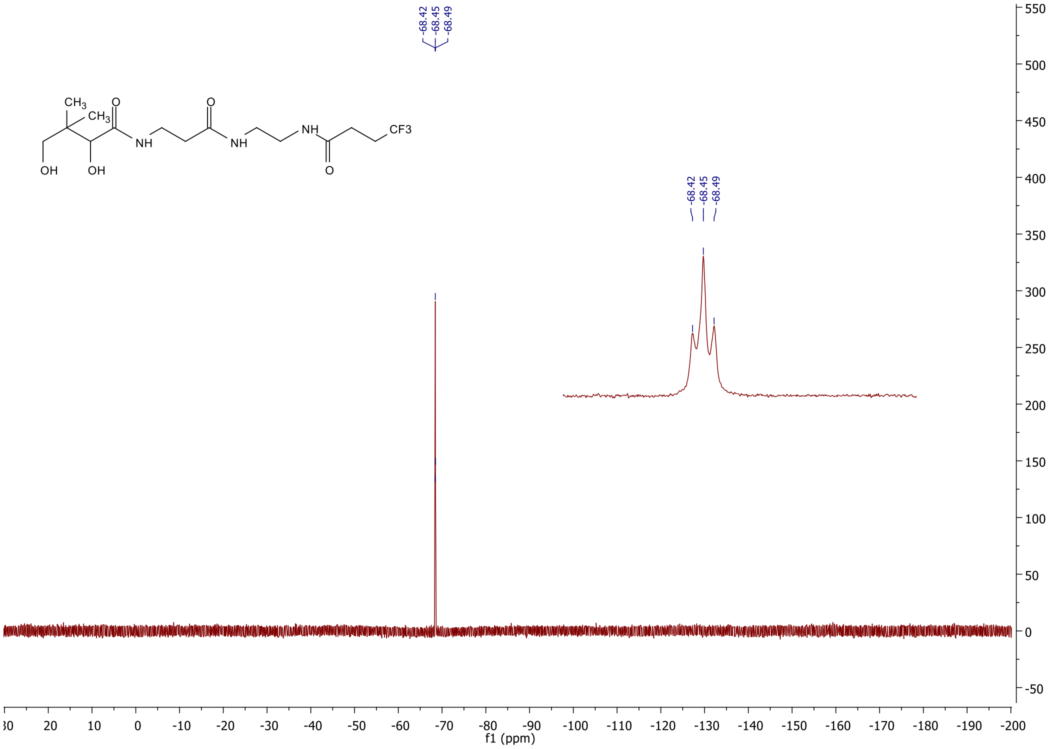

#### [1]H NMR spectrum of **6c**

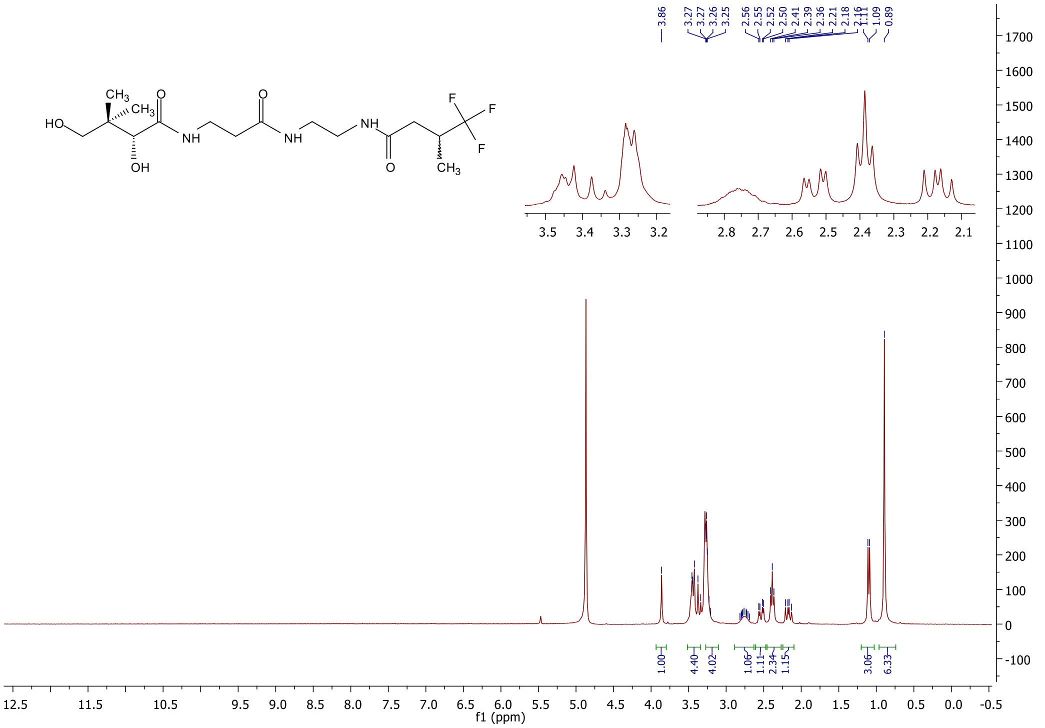

#### [13]C NMR spectrum of **6c**

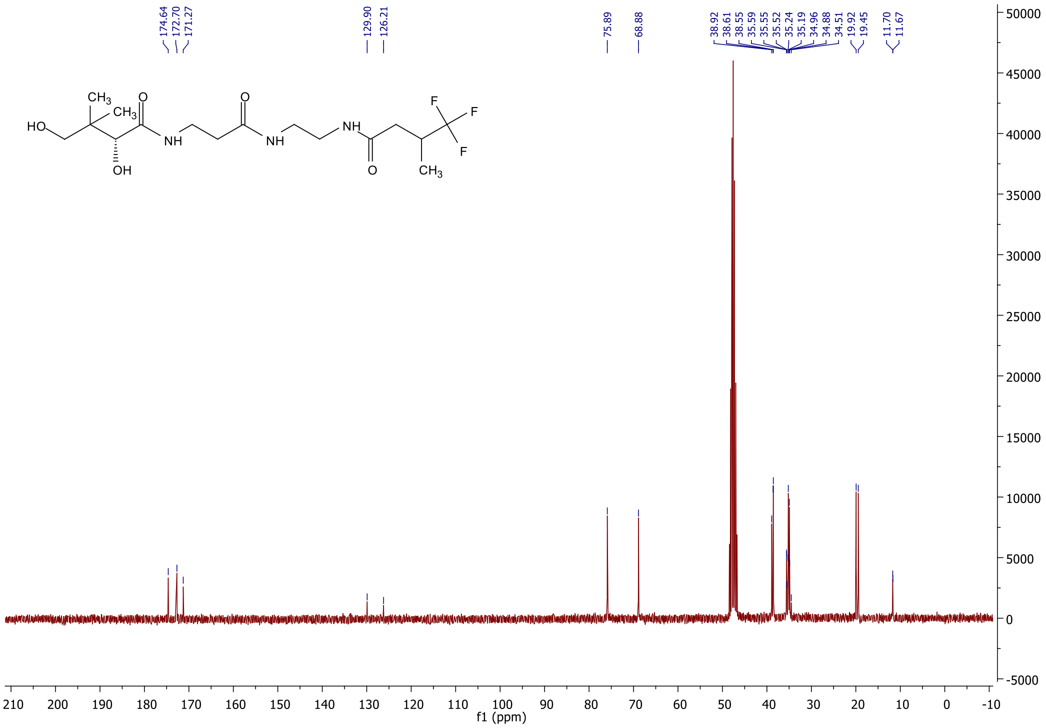

#### [19]F NMR spectrum of **6c**

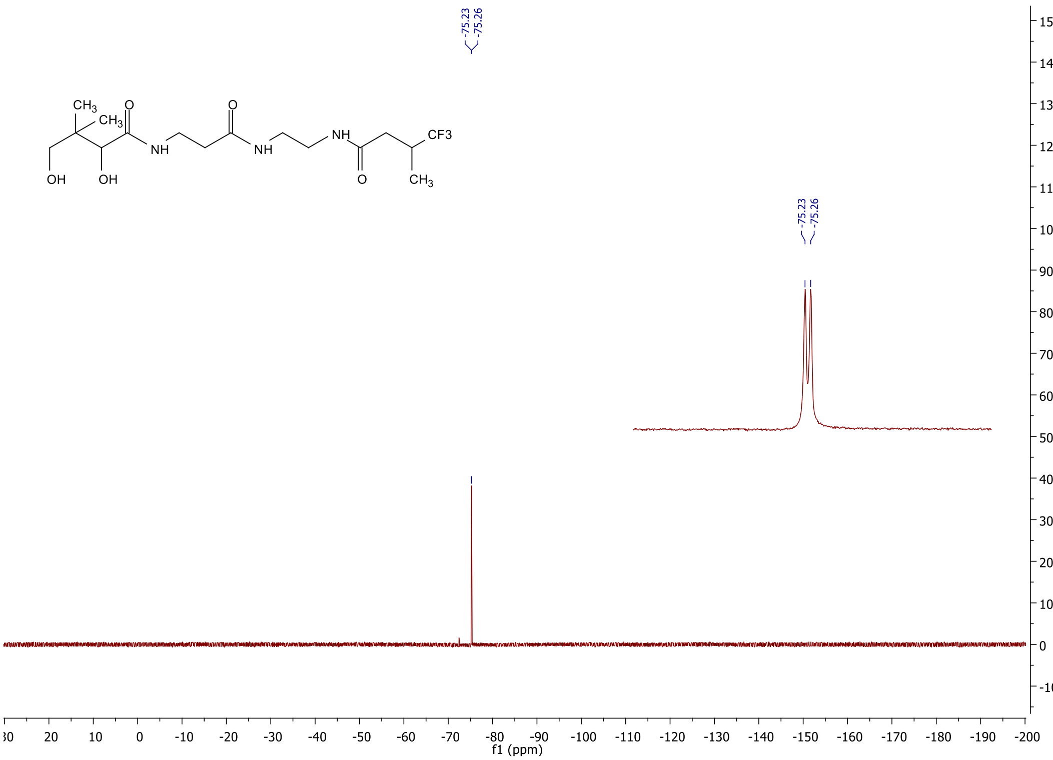

#### [1]H NMR spectrum of **6d**

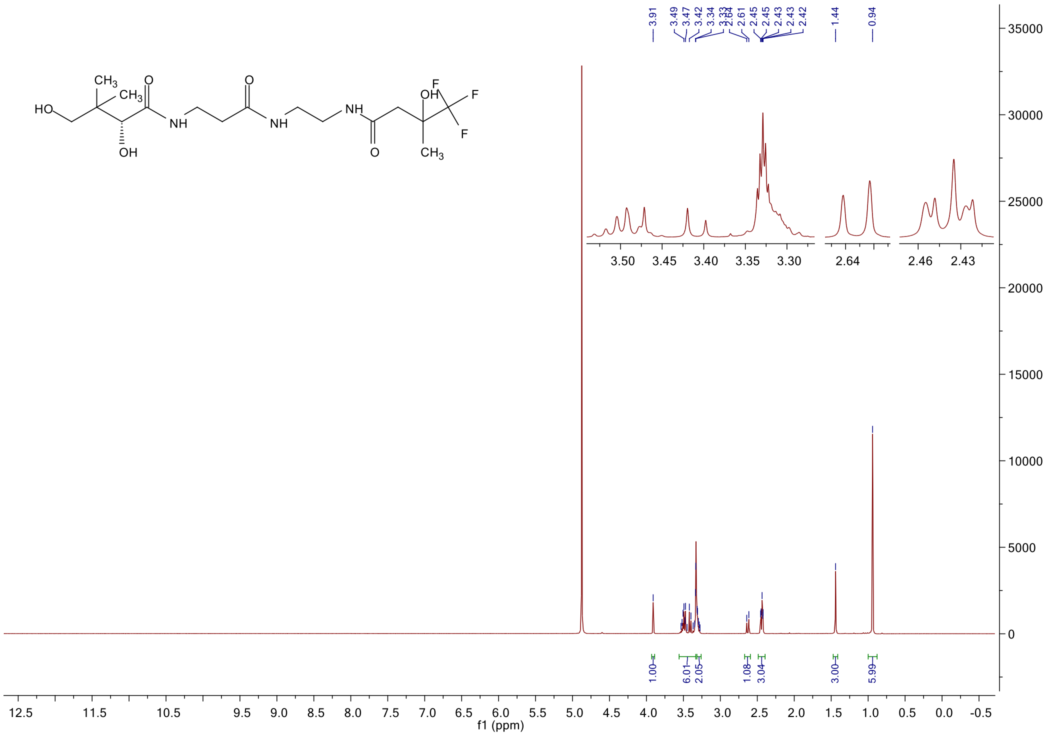

#### $^{13}C$ NMR spectrum of **6d**

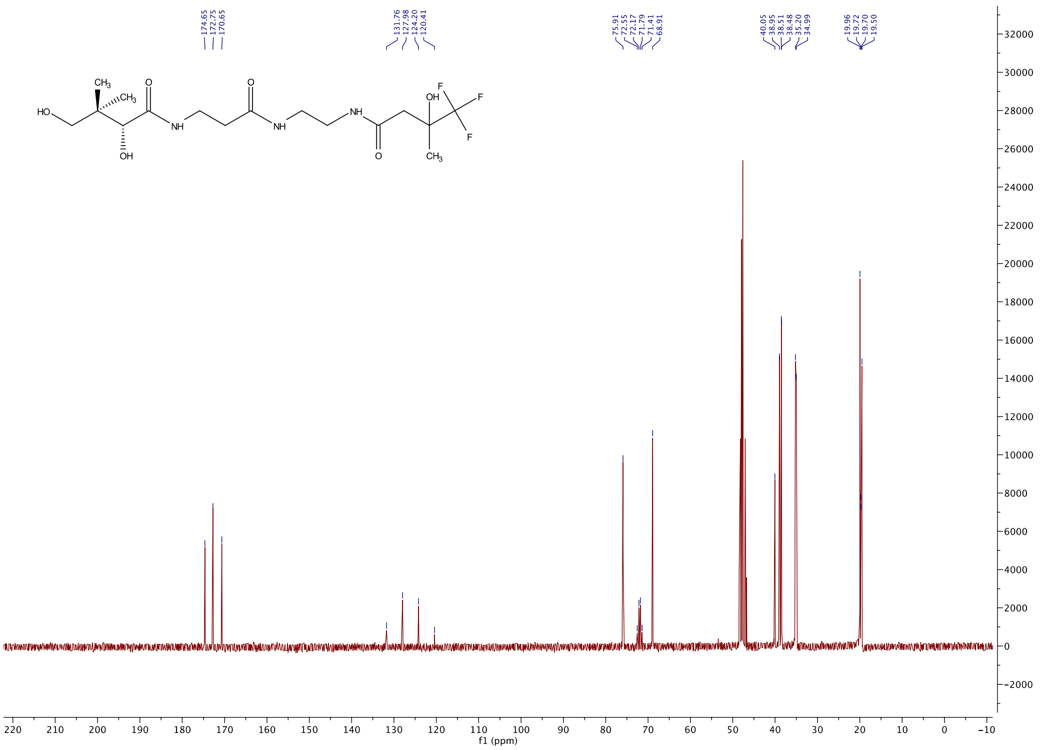

#### $^{19}F$ NMR spectrum of **6d**

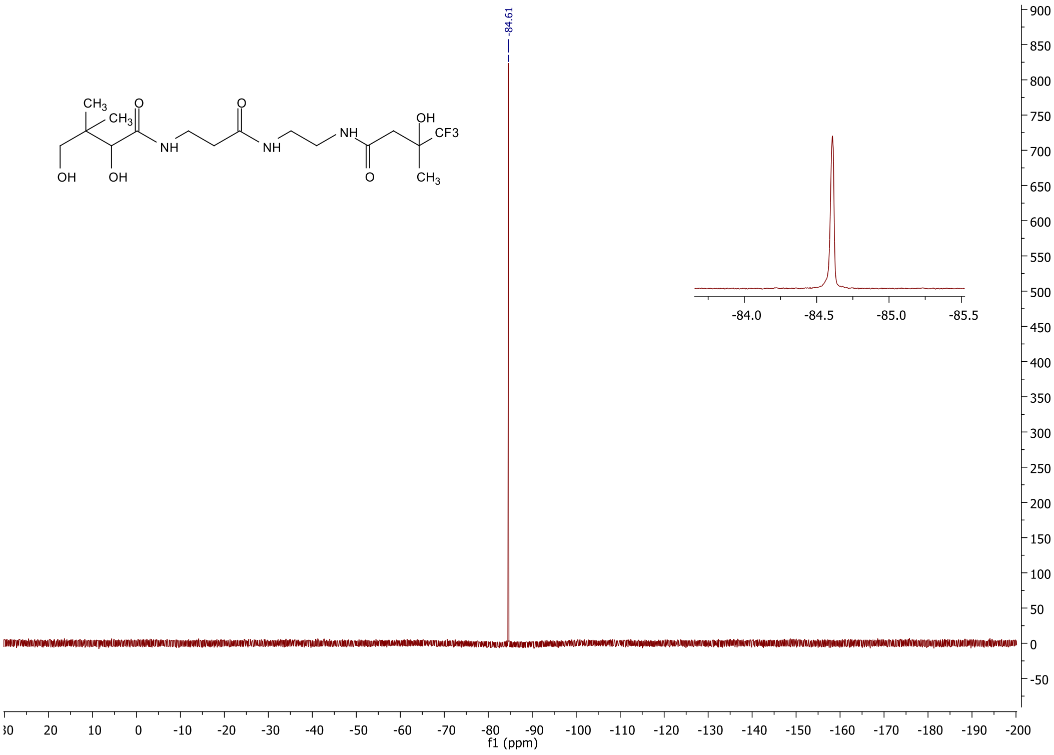

#### $^{1}$H NMR spectrum of **6e**

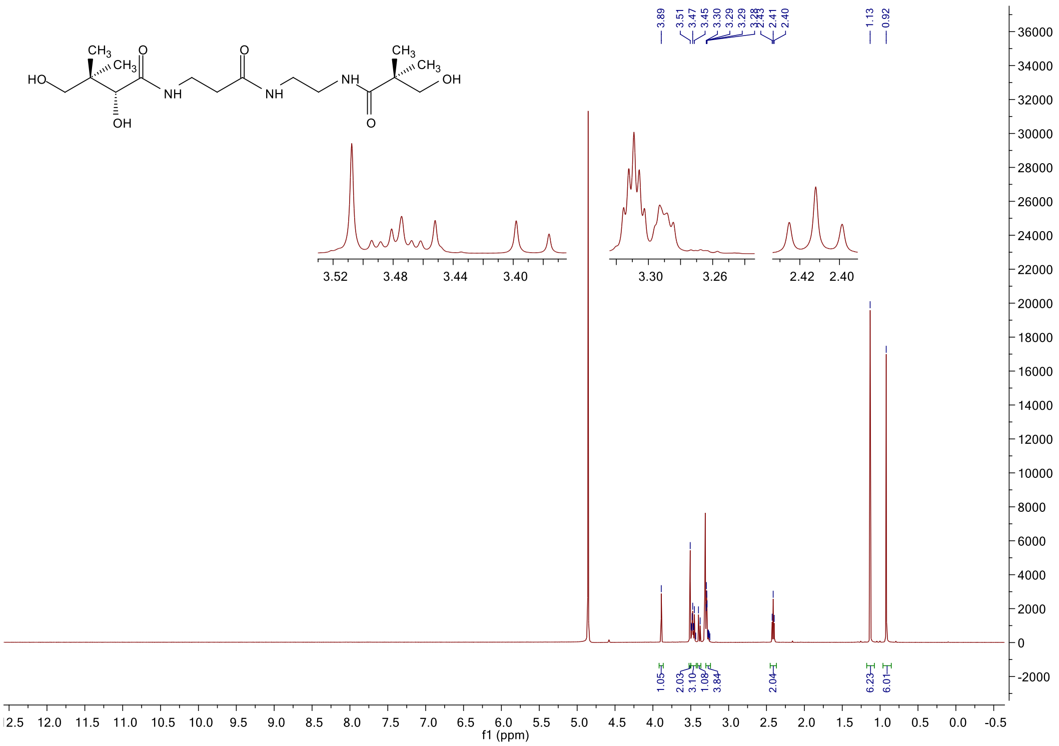

#### $^{13}$C NMR spectrum of **6e**

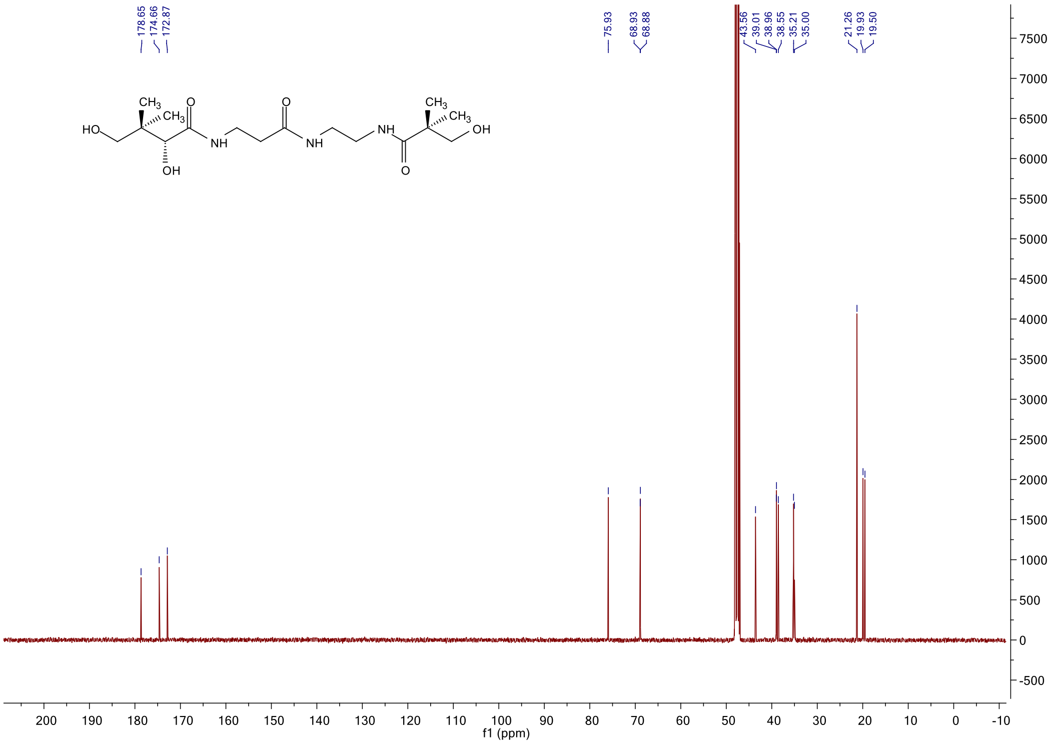

#### $^{1}$H NMR spectrum of **6f**

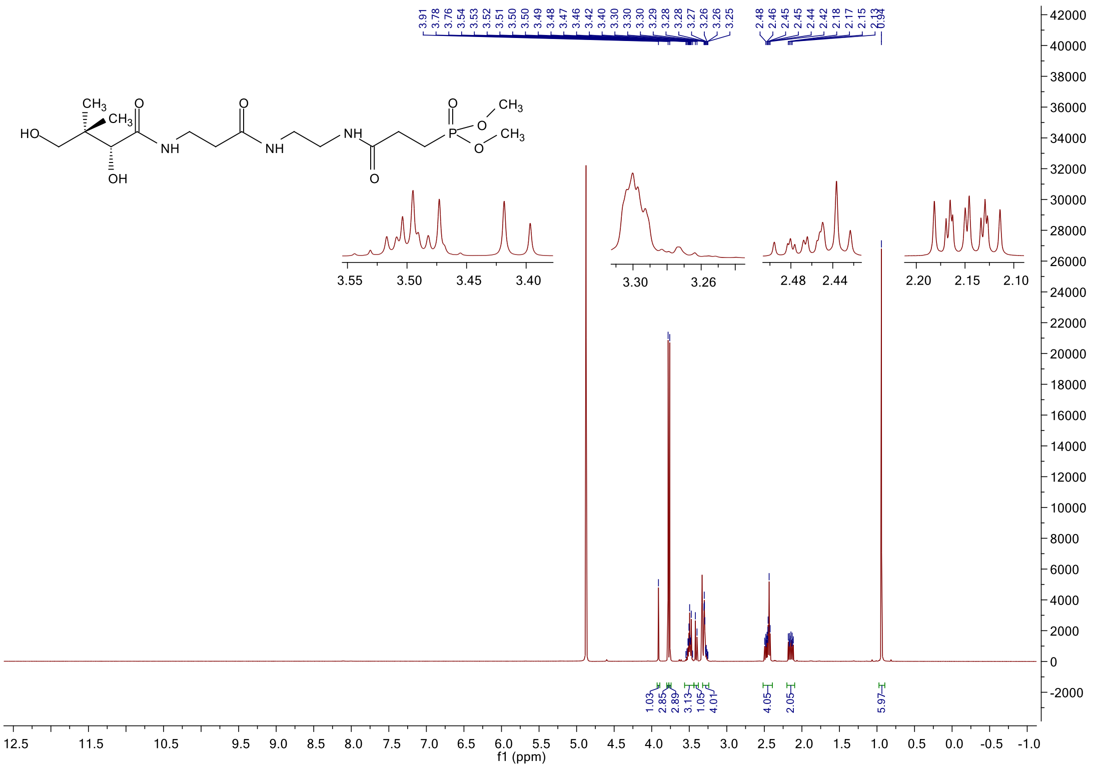

#### $^{13}$C NMR spectrum of **6f**

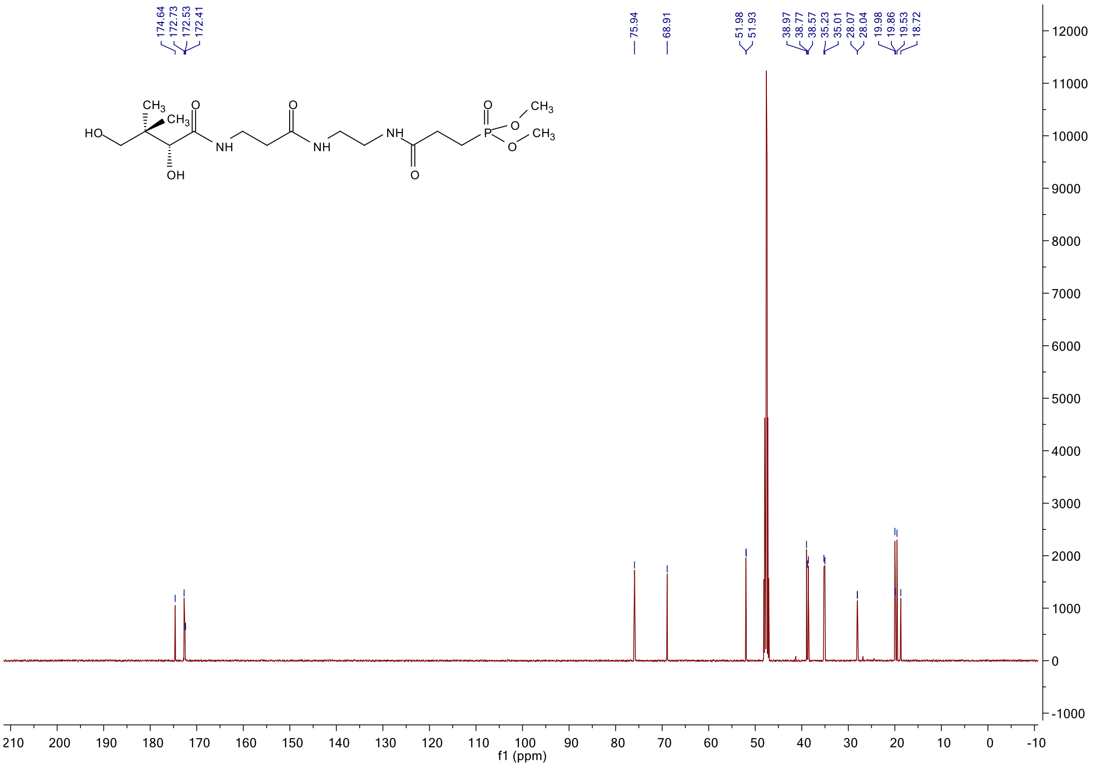

#### $^{31}$P NMR spectrum of **6f**

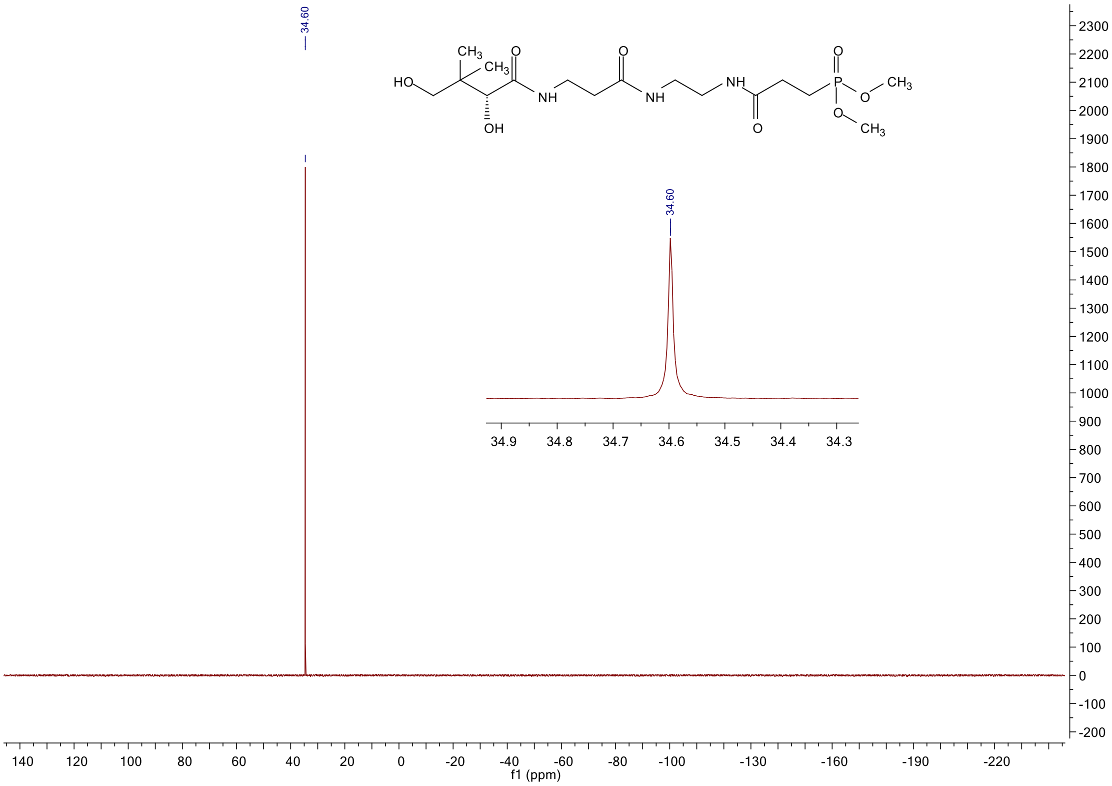

#### $^{1}$H NMR spectrum of **6g**

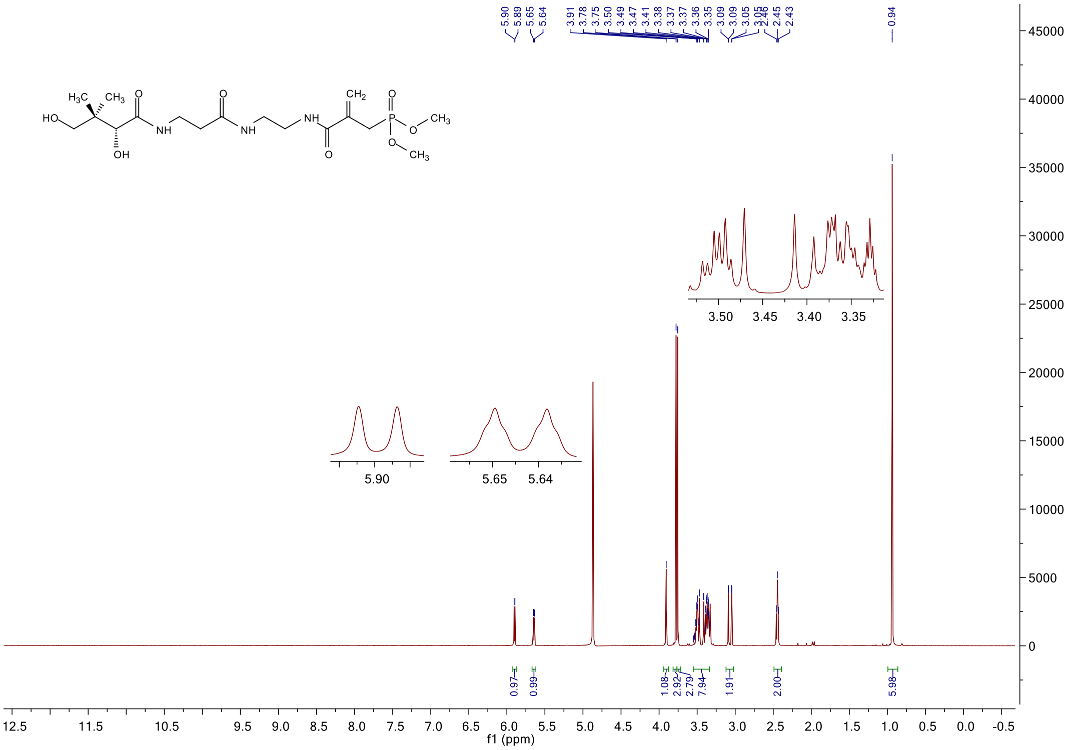

#### $^{13}$C NMR spectrum of **6g**

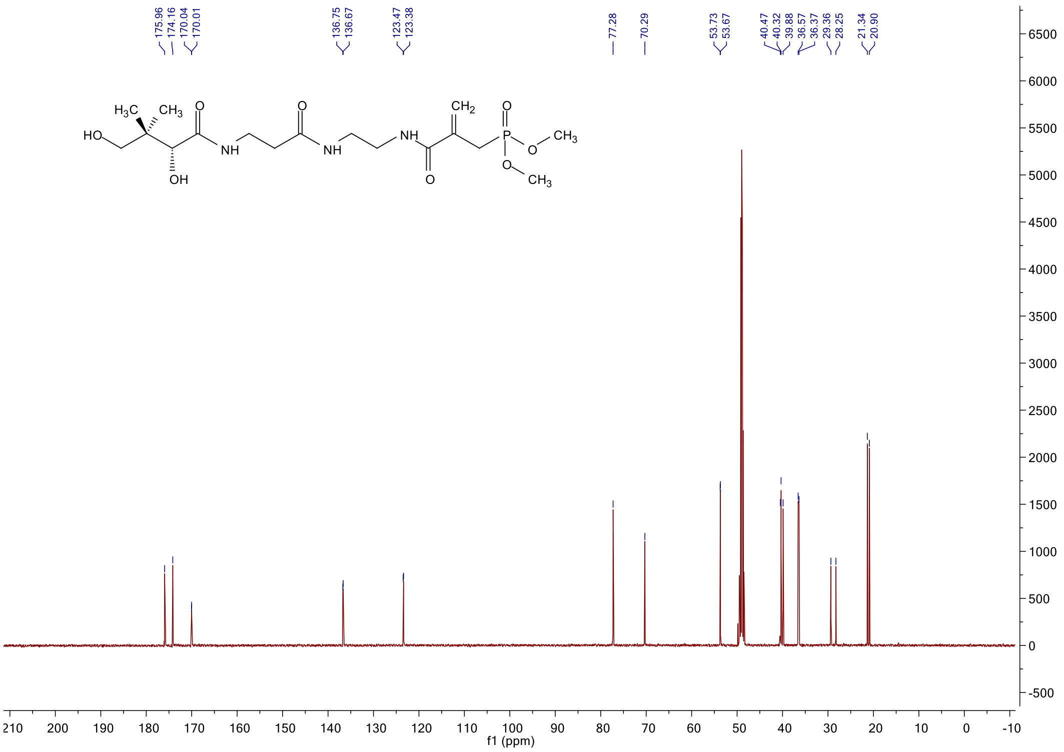

#### $^{31}$P NMR spectrum of **6g**

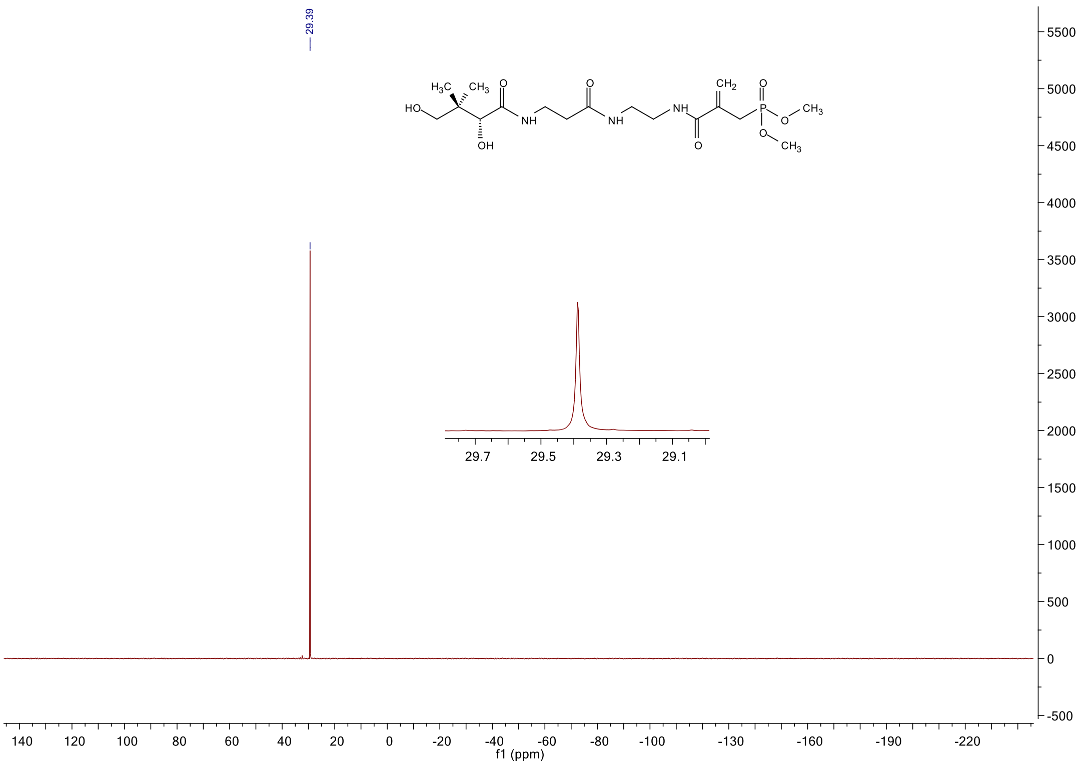

$^{1}$H NMR spectrum of **6h**

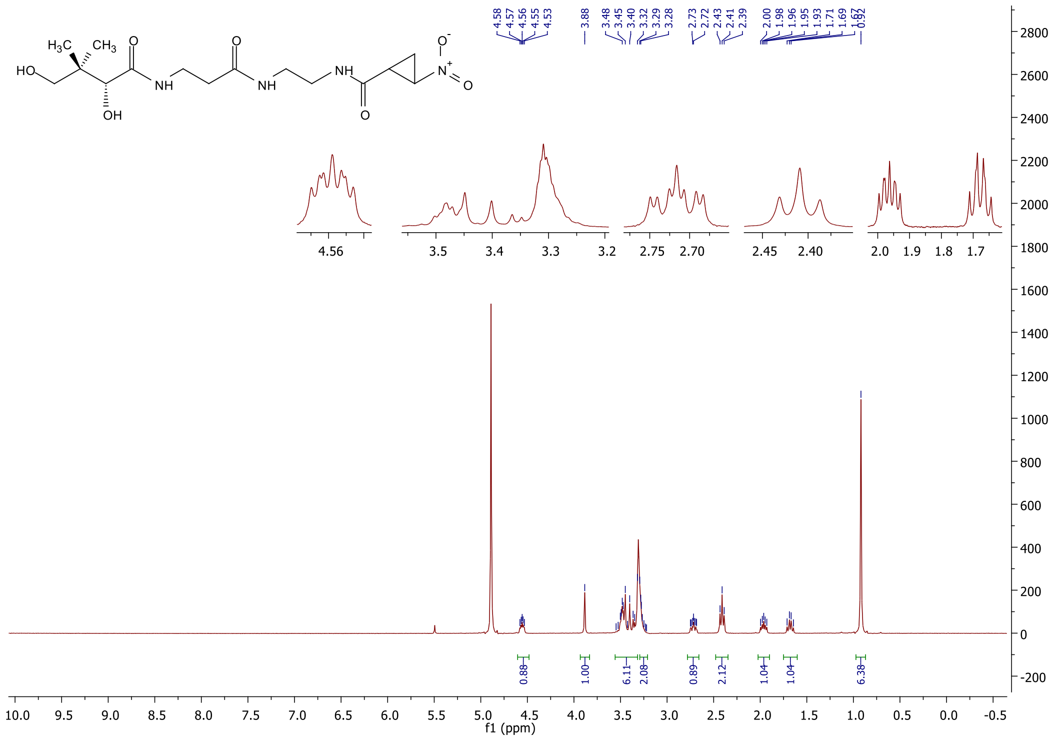

$^{13}$C NMR spectrum of **6h**

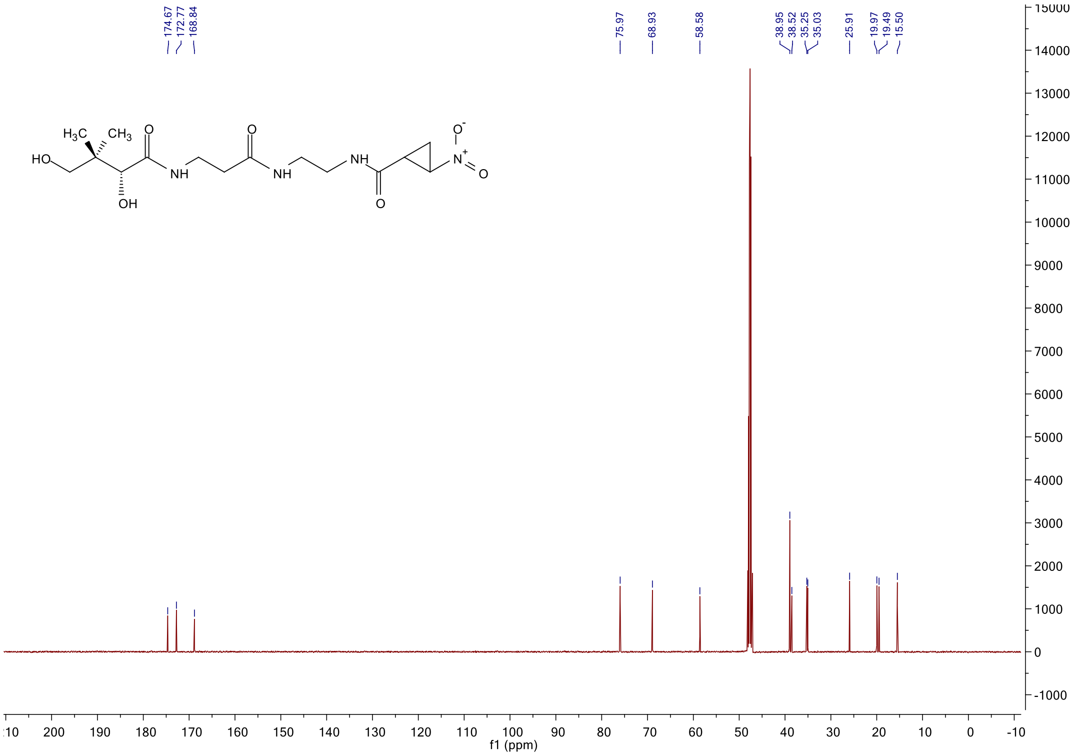

#### [1]H NMR spectrum of **6i**

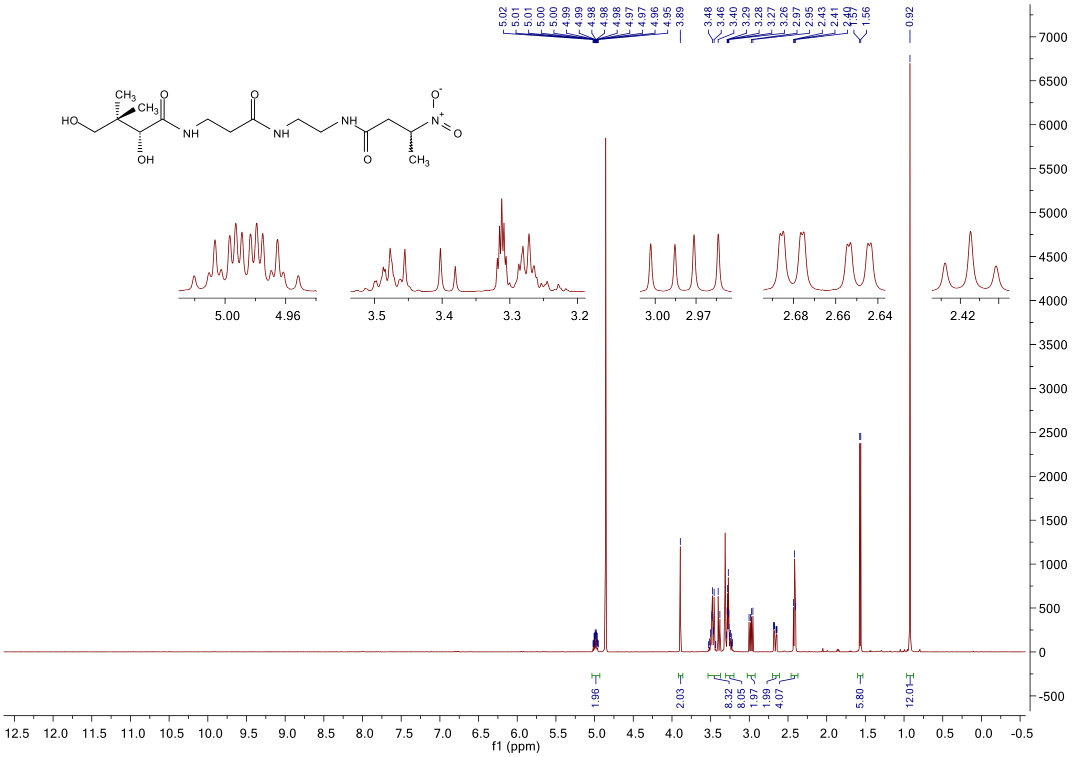

#### [13]C NMR spectrum of **6i**

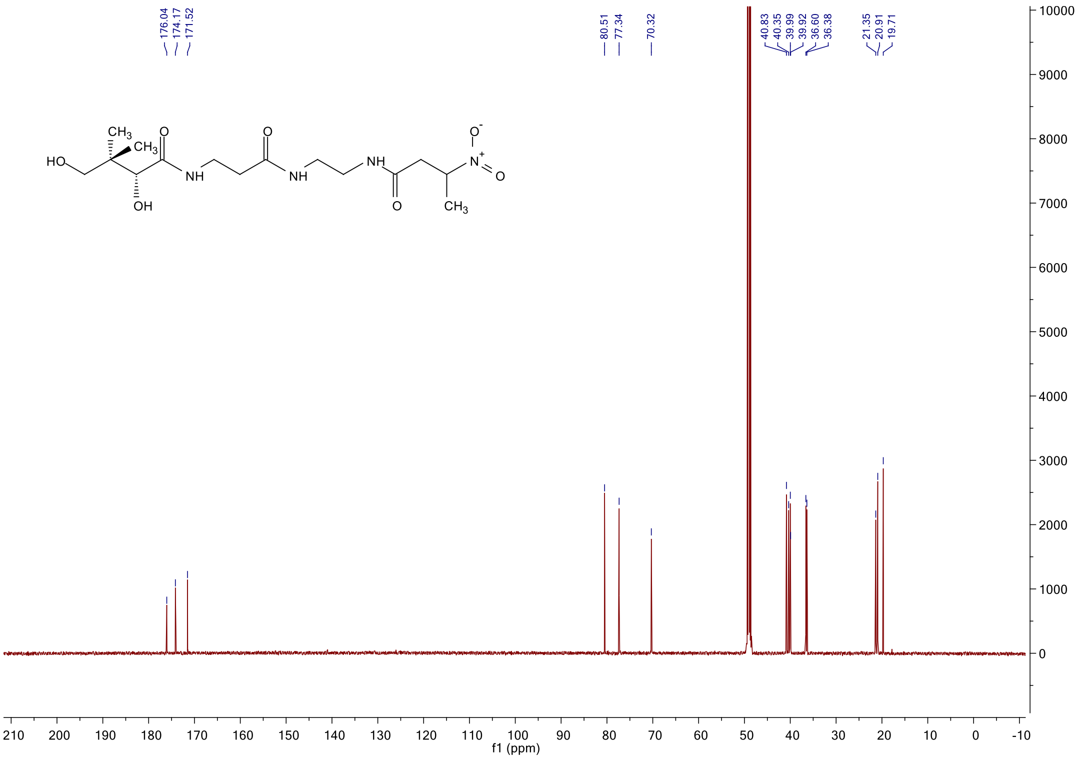

#### $^{1}$H NMR spectrum of **6k**

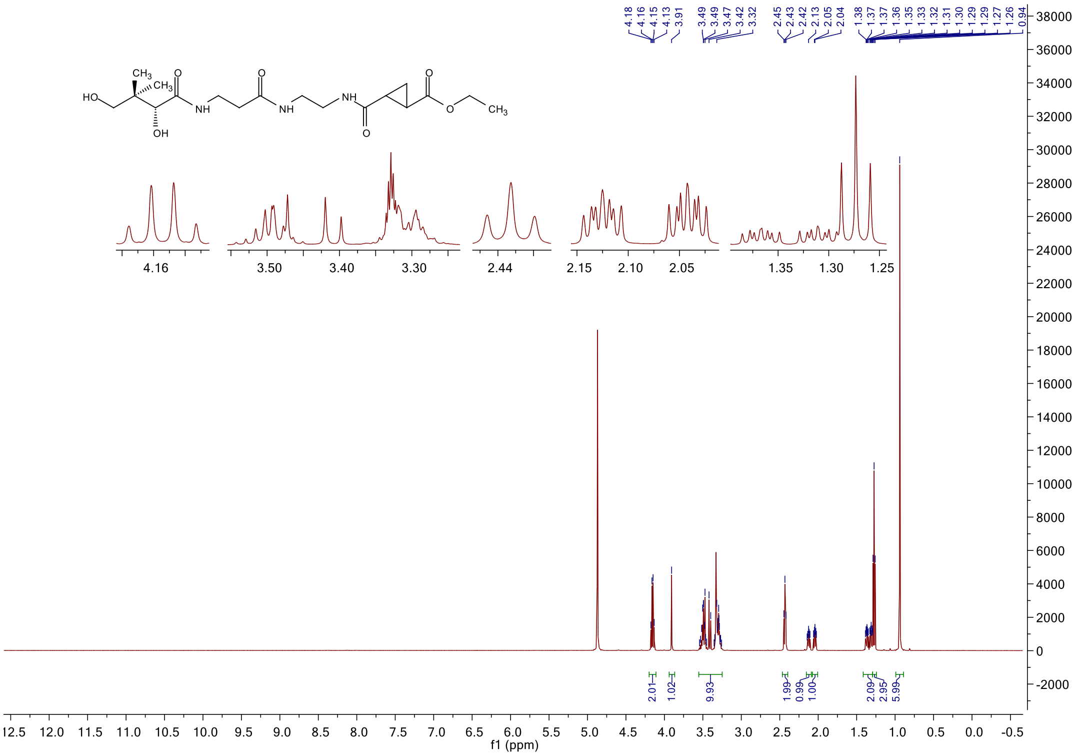

#### $^{13}$C NMR spectrum of **6k**

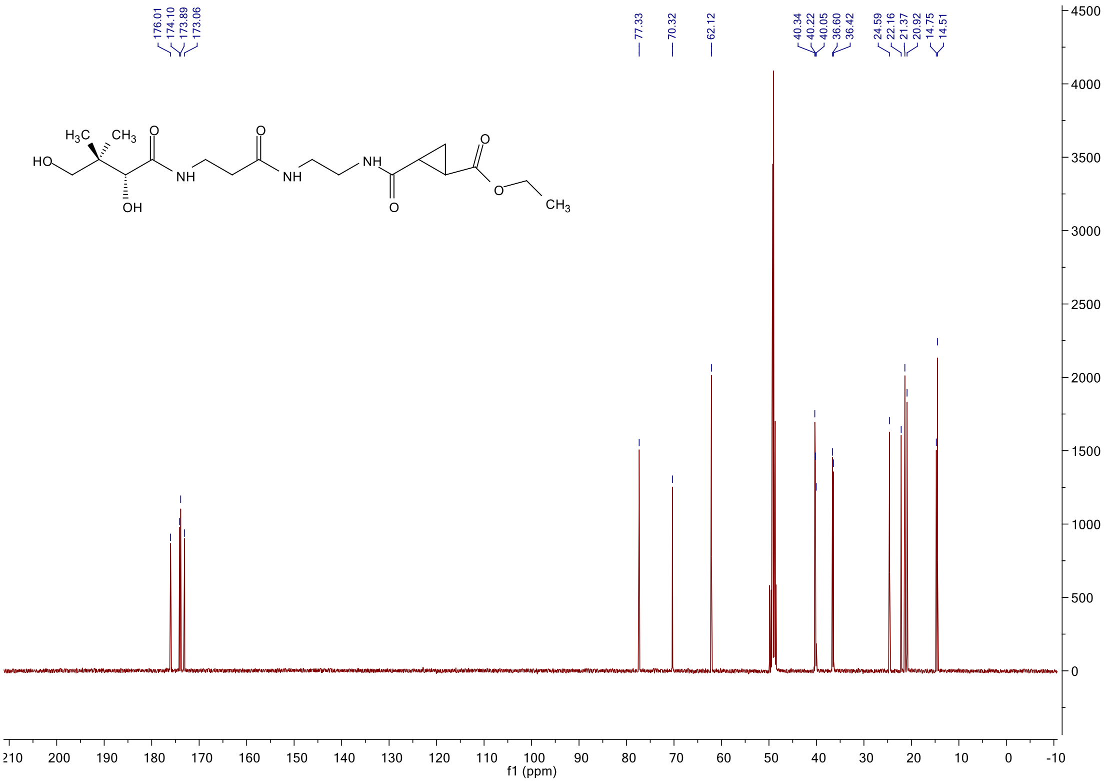

#### [1]H NMR spectrum of **6l**

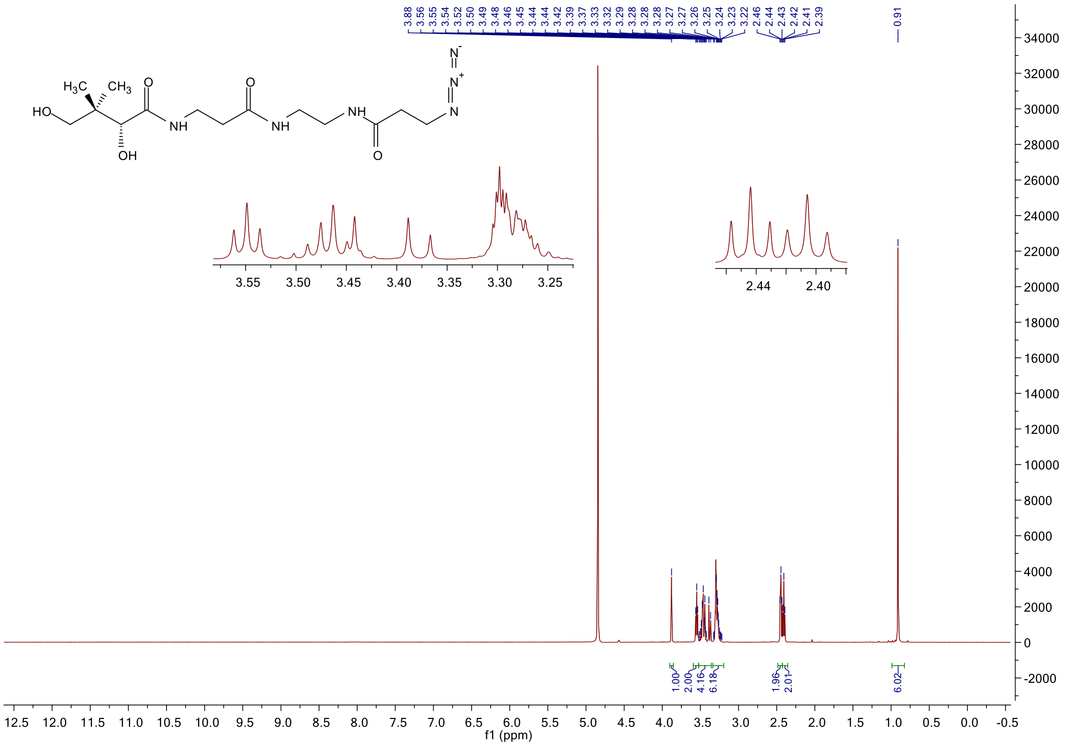

#### [13]C NMR spectrum of **6l**

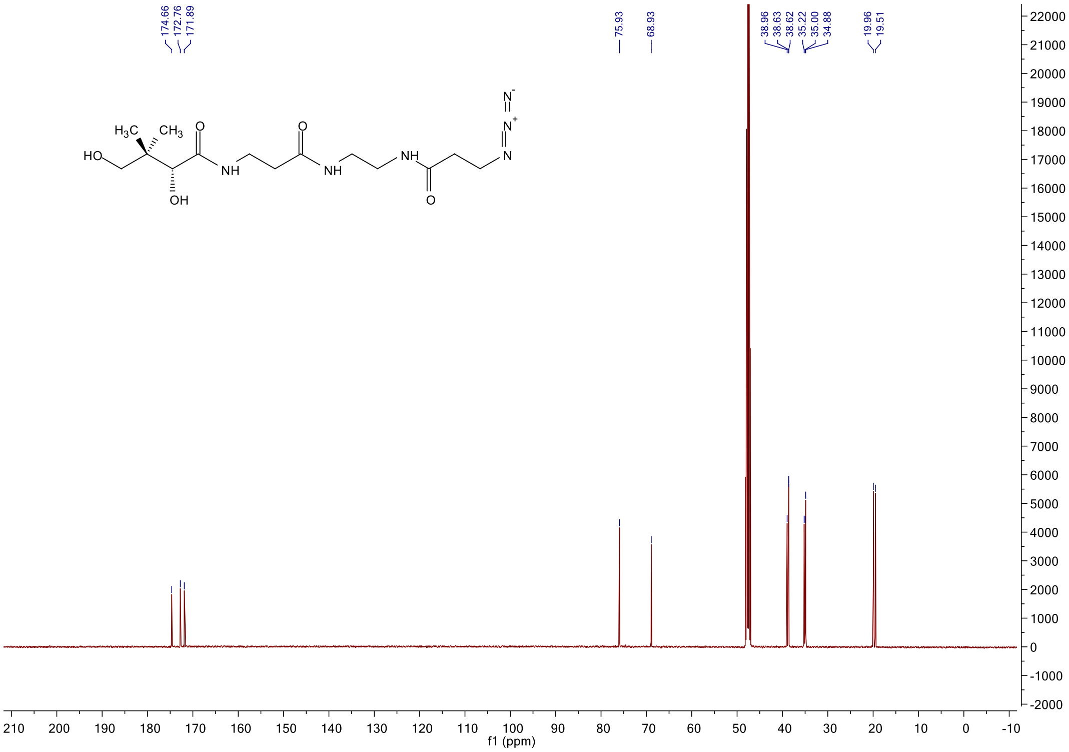

<sup></sup>

[1]H NMR spectrum of **6m**

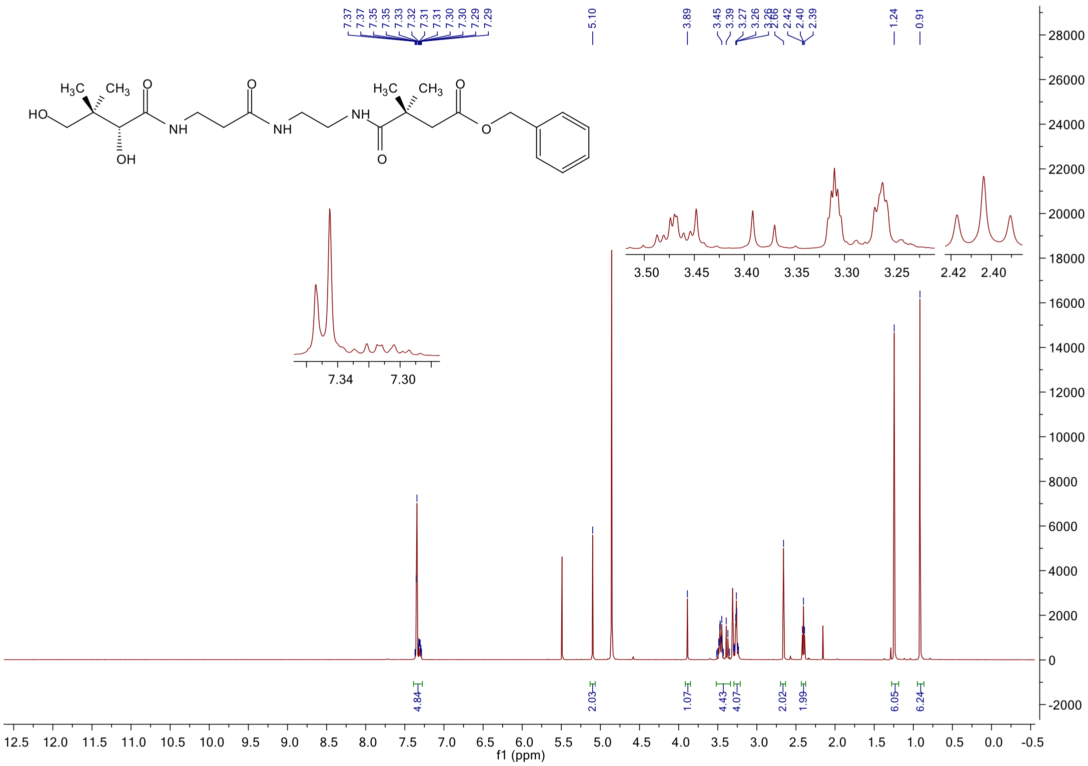

[13]C NMR spectrum of **6m**

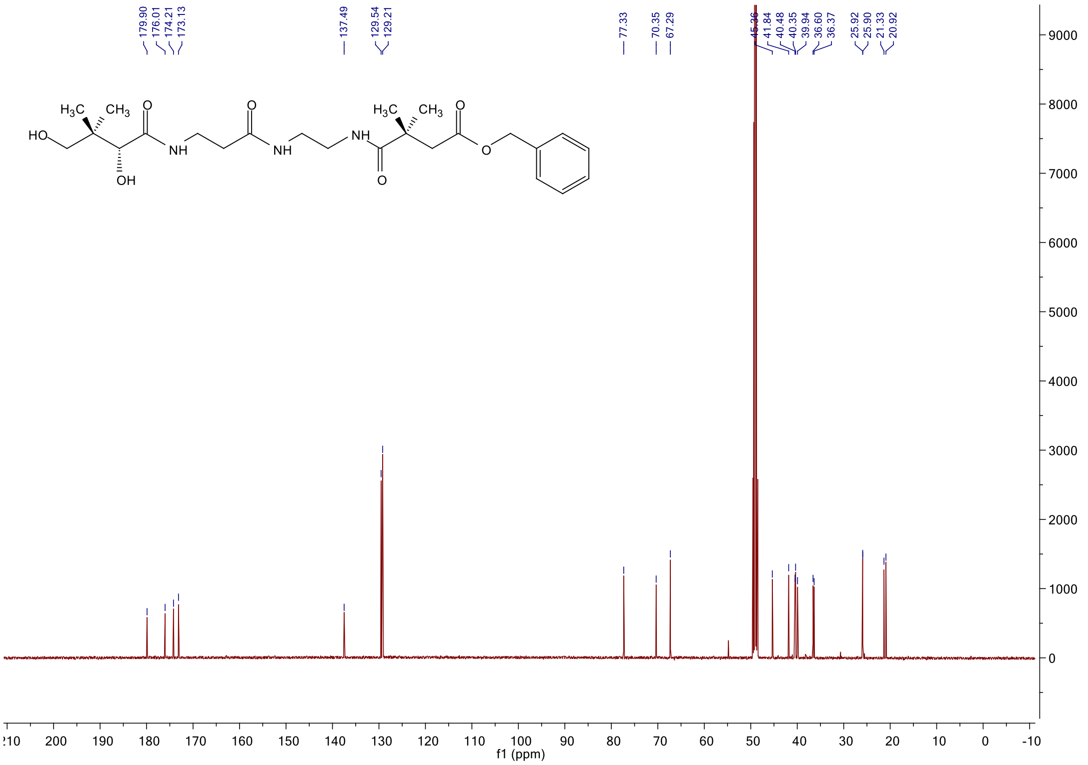

$^{1}$H NMR spectrum of **6n**

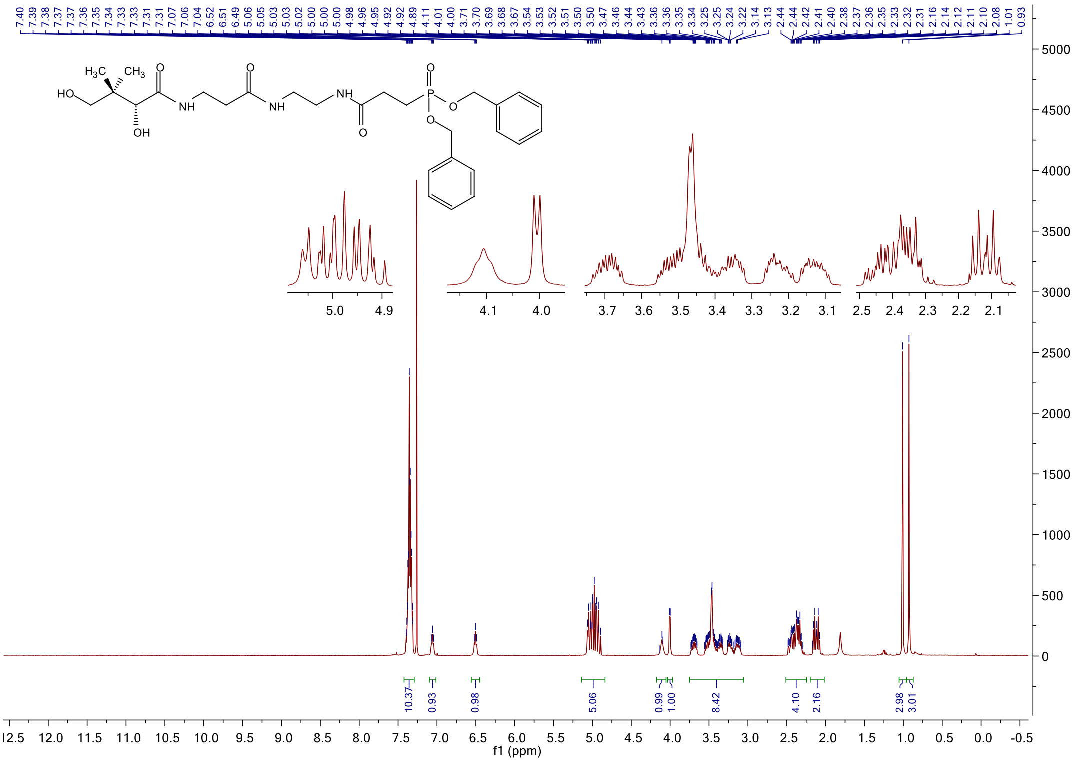

$^{13}$C NMR spectrum of **6n**

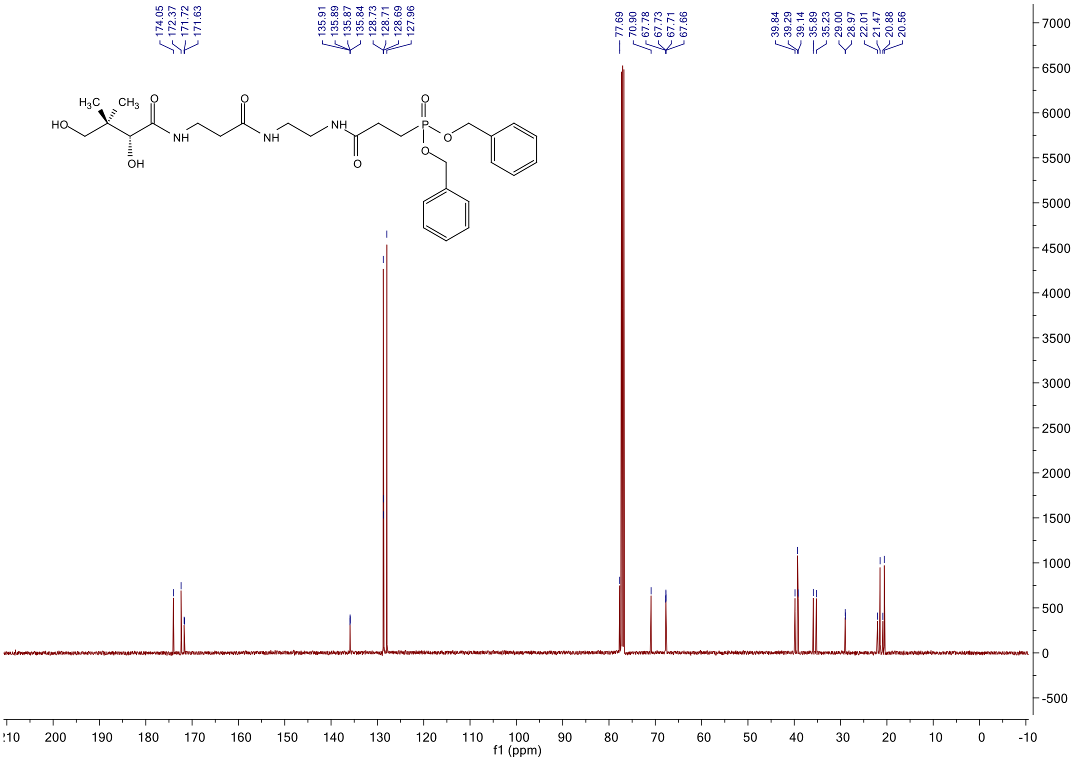

#### $^{31}$P NMR spectrum of **6n**

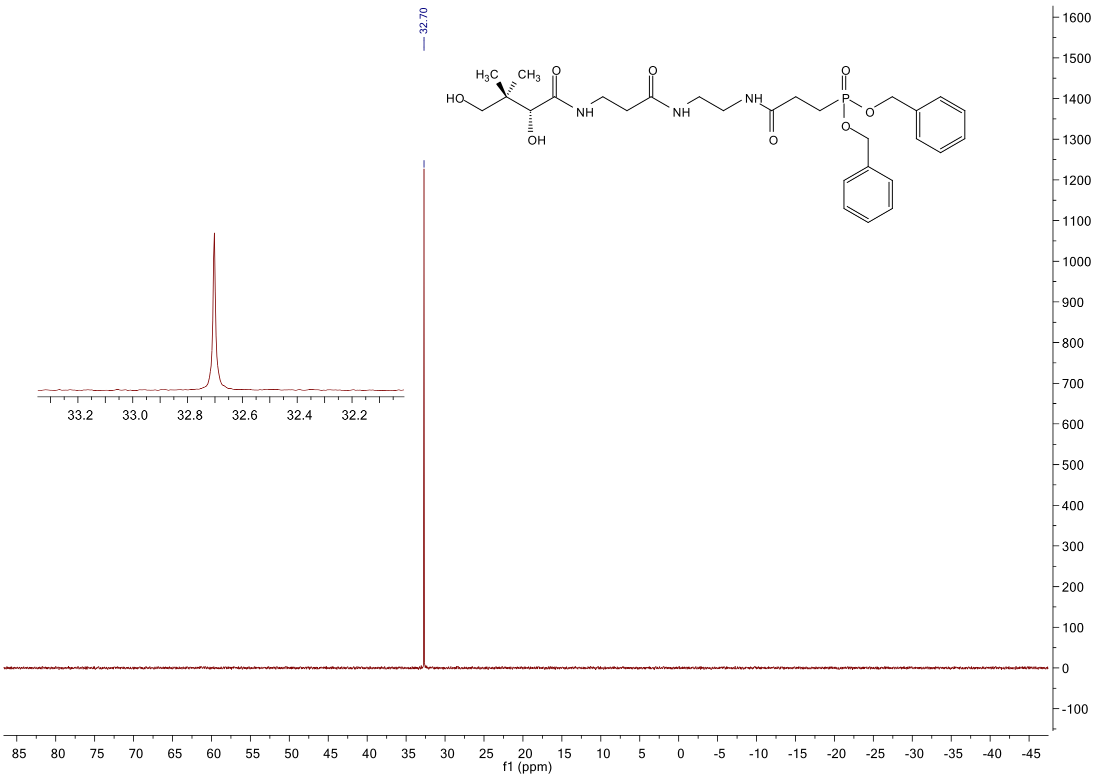

#### $^{1}$H NMR spectrum of **7m**

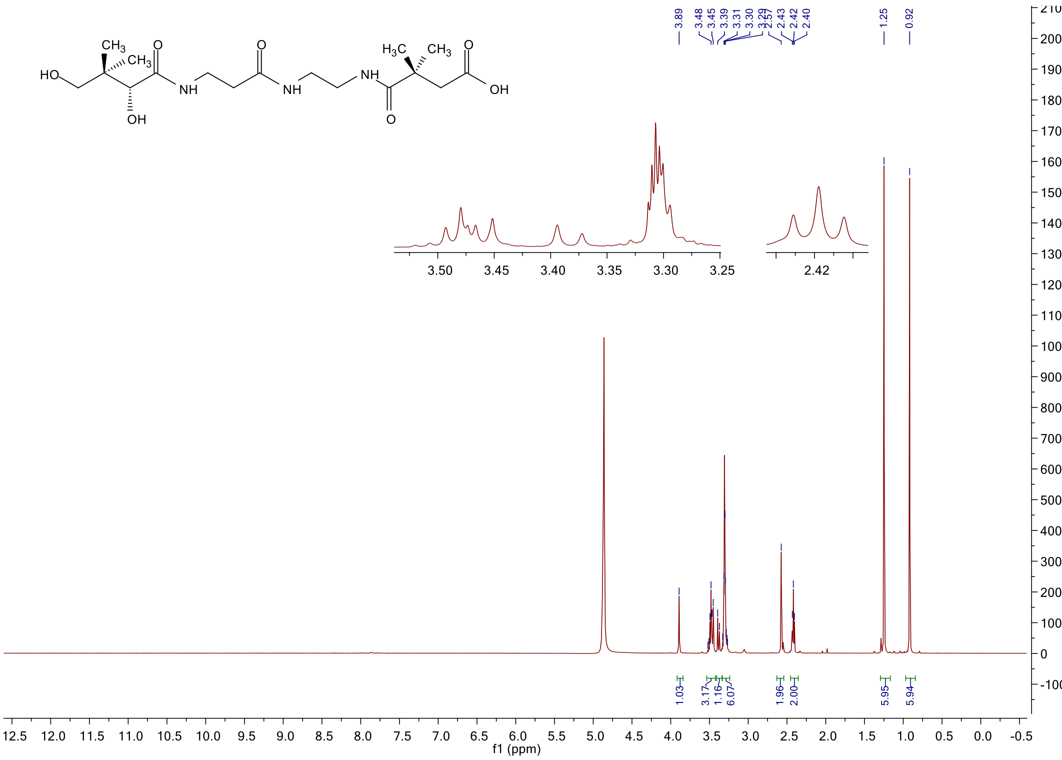

#### [13]C NMR spectrum of **7m**

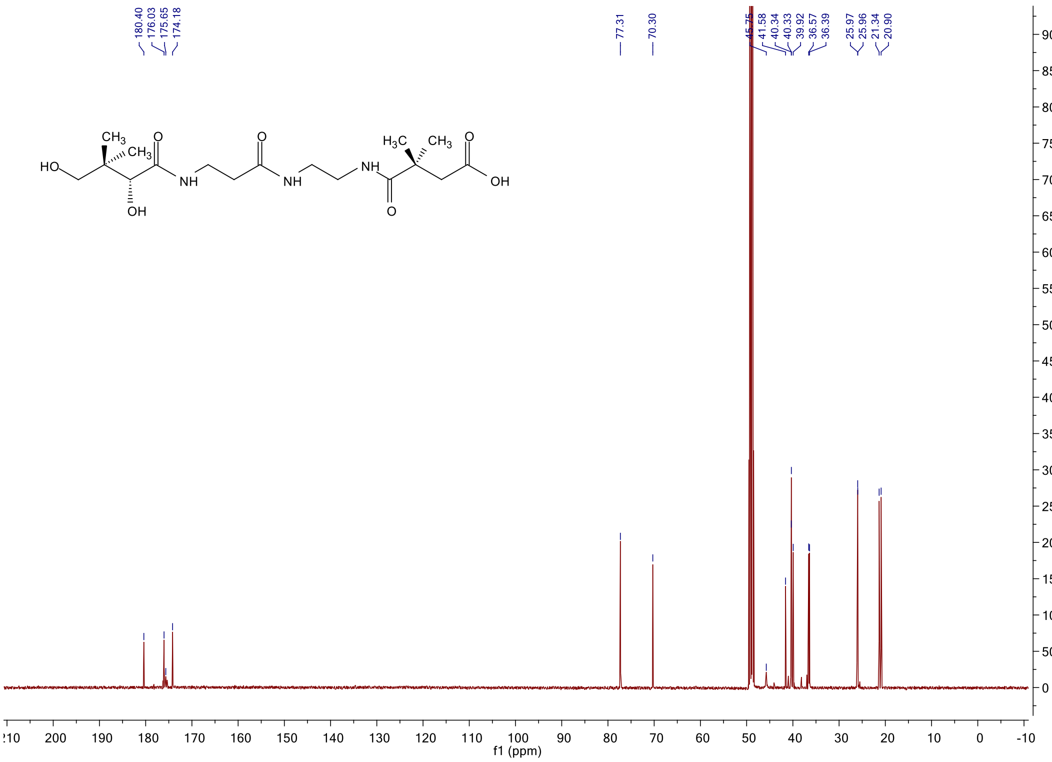

#### [1]H NMR spectrum of **7n**

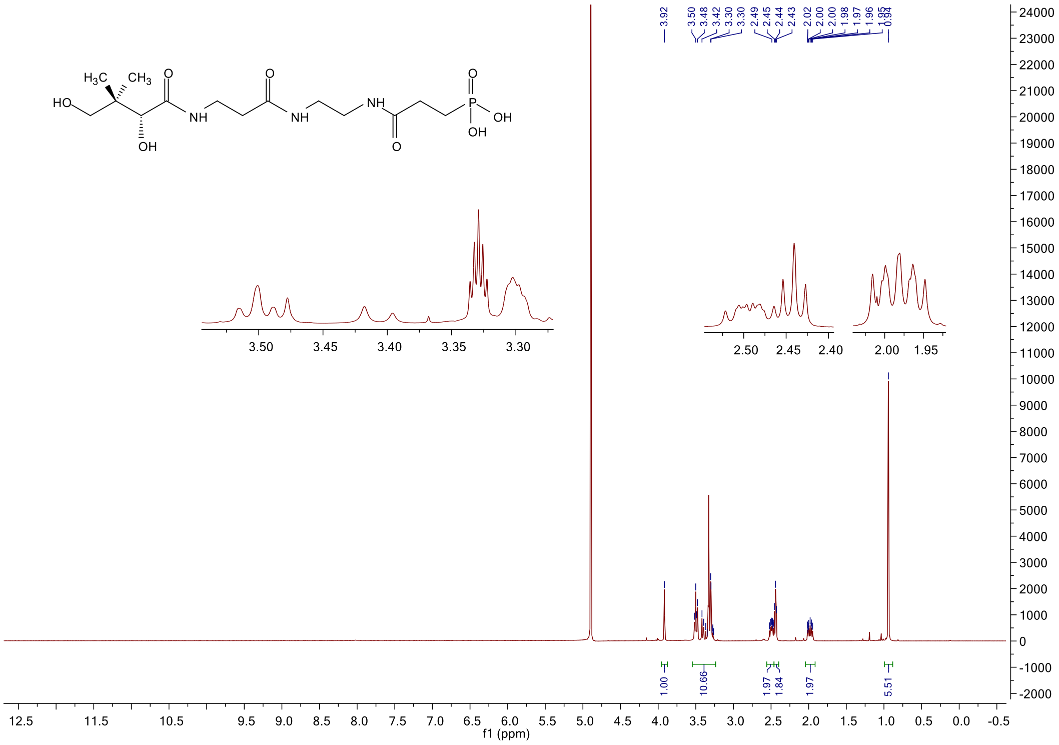

$^{13}$C NMR spectrum of **7n**

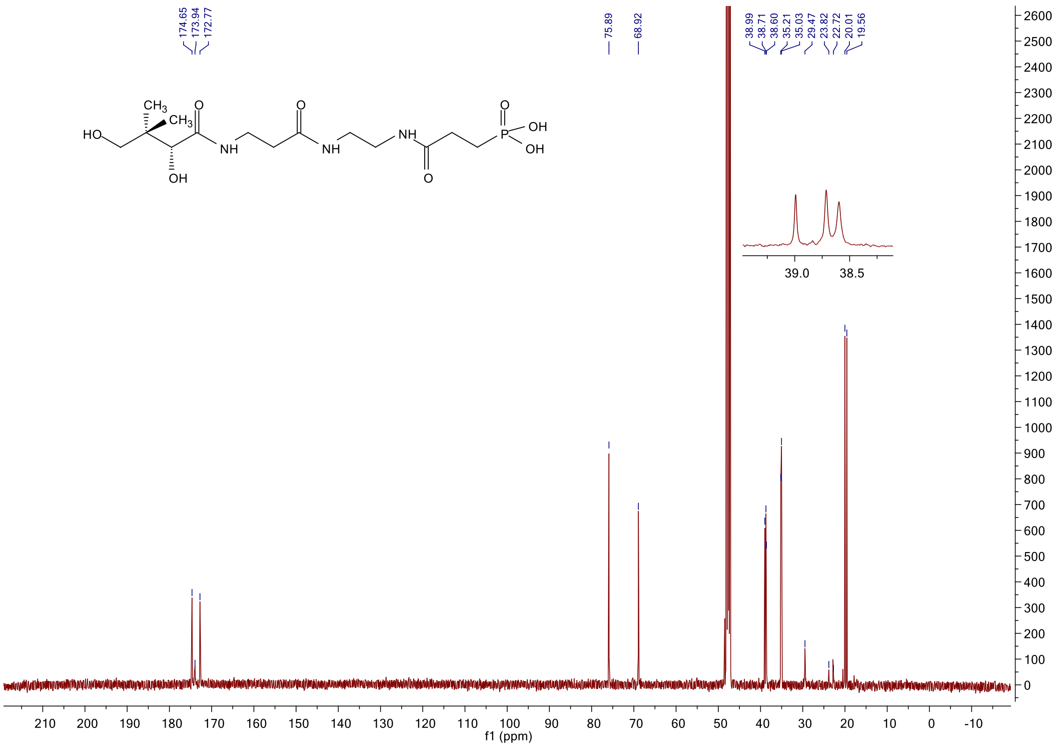

$^{31}$P NMR spectrum of **7n**

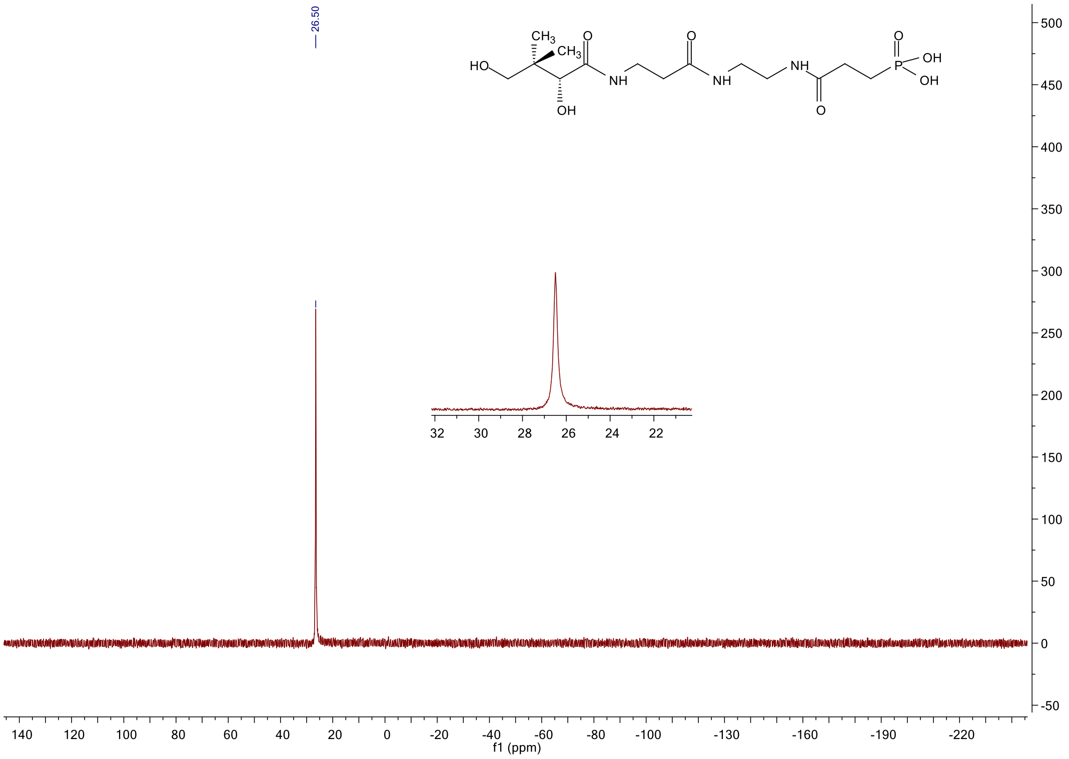

#### [1]H NMR spectrum of **9k**

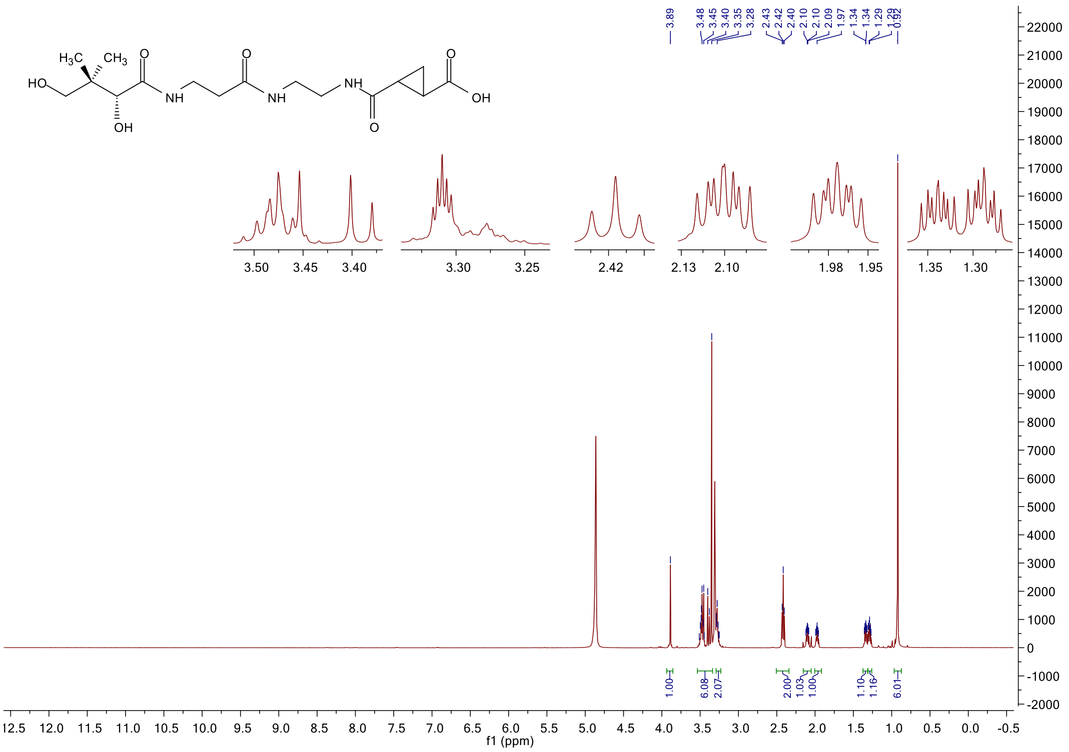

#### [13]C NMR spectrum of **9k**

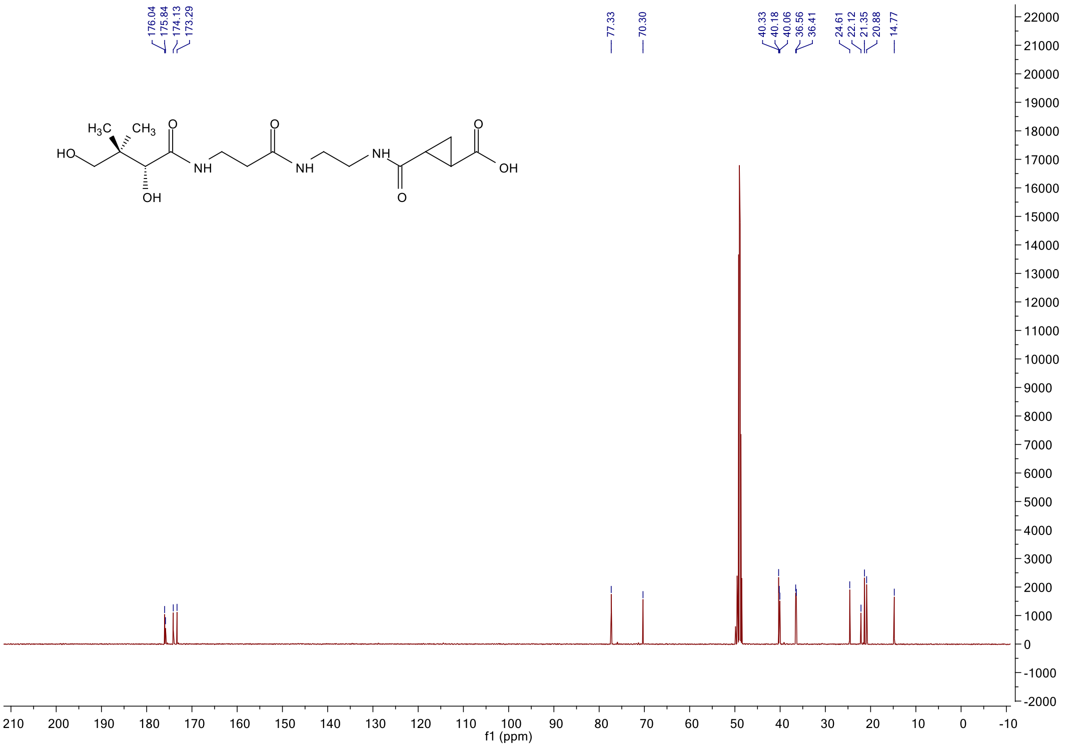

#### $^{1}$H NMR spectrum of **10**

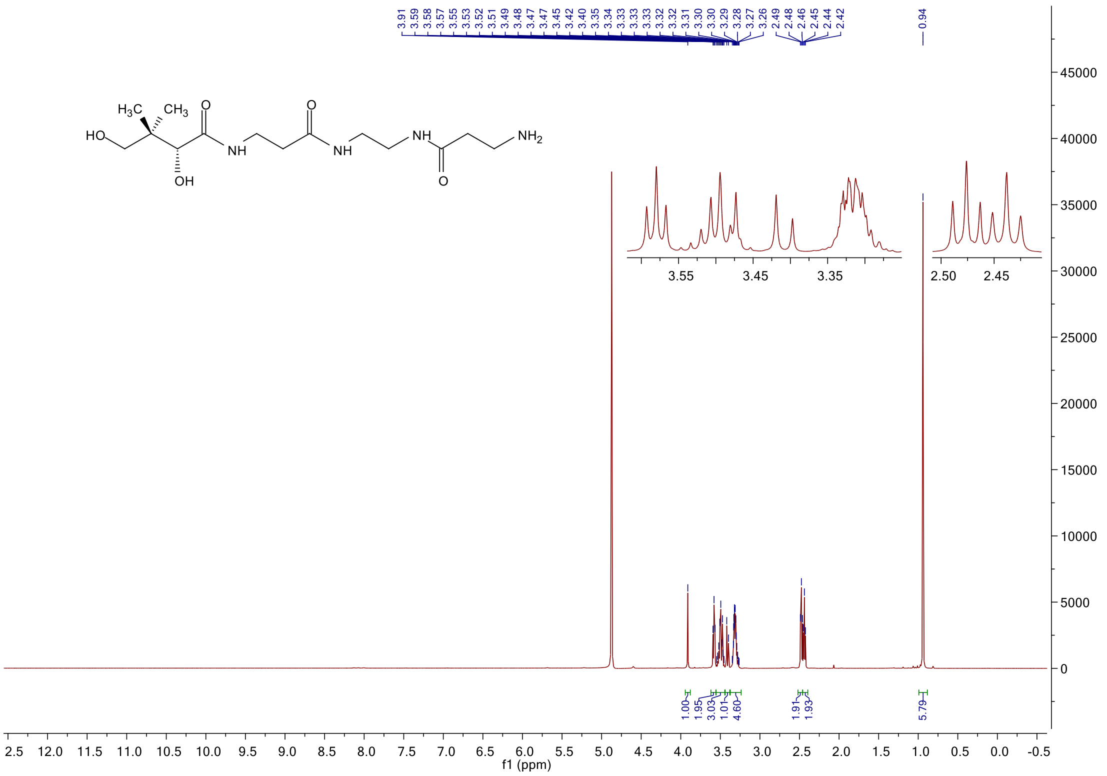

#### $^{13}$C NMR spectrum of **10**

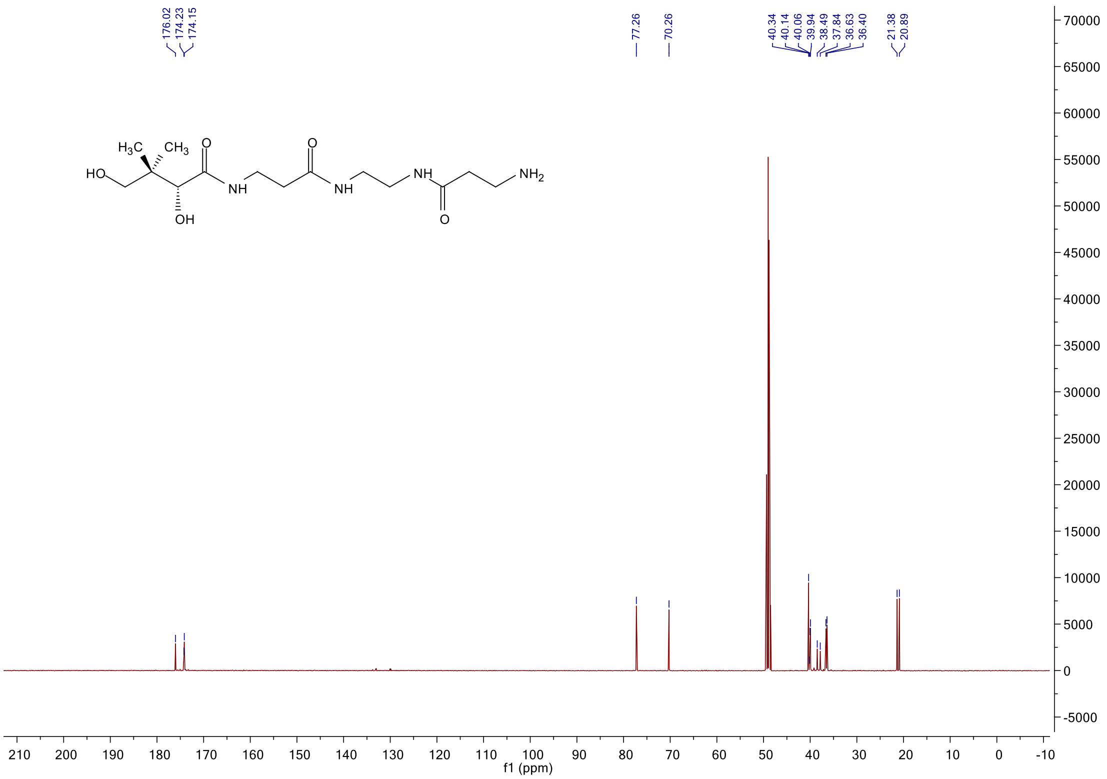

#### $^{1}$H NMR spectrum of **12**

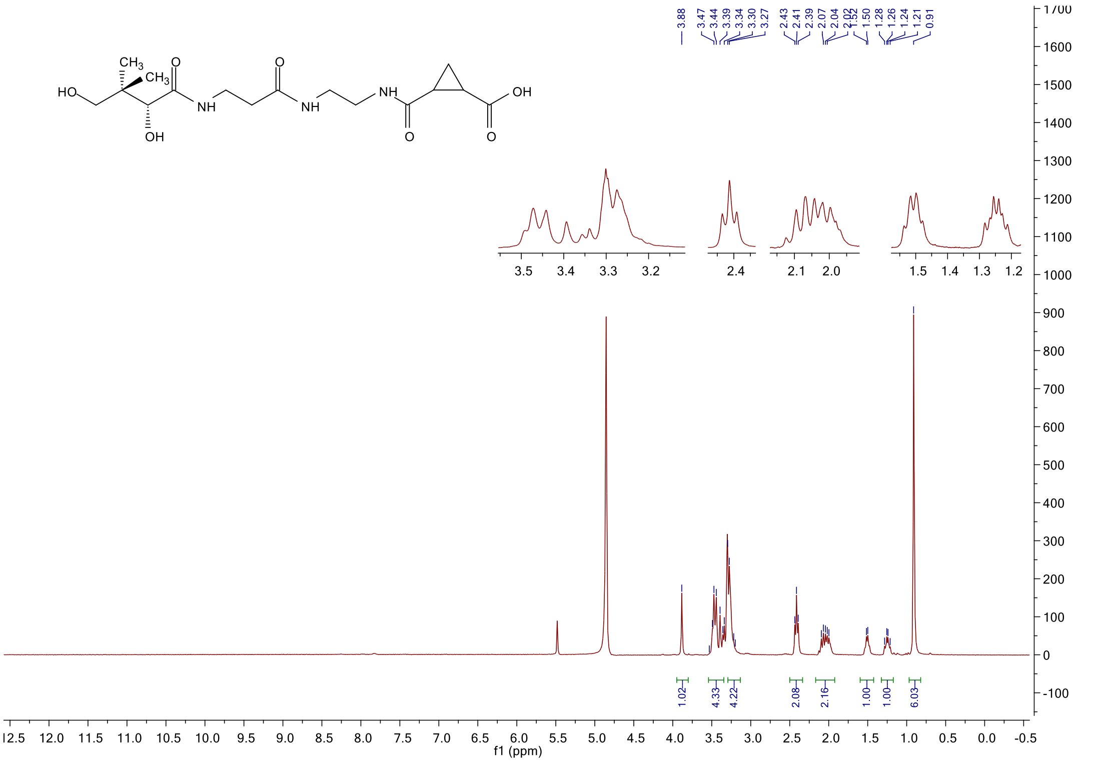

#### $^{13}$C NMR spectrum of **12**

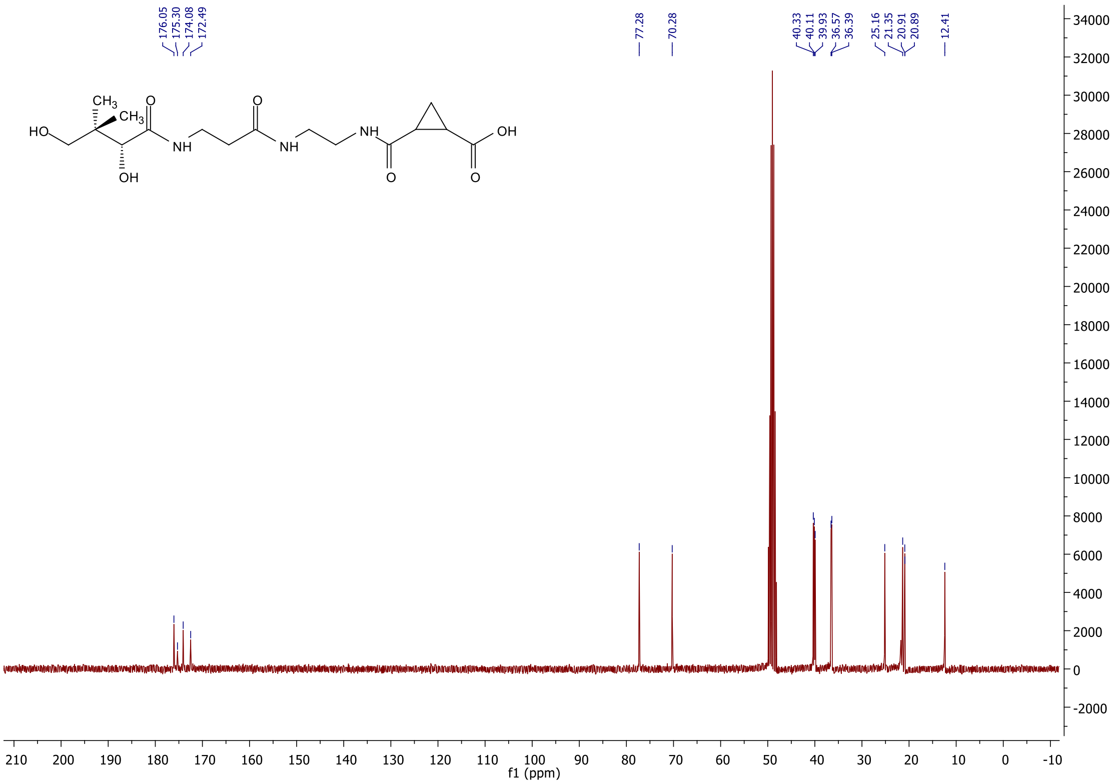

#### [1]H NMR spectrum of **15**

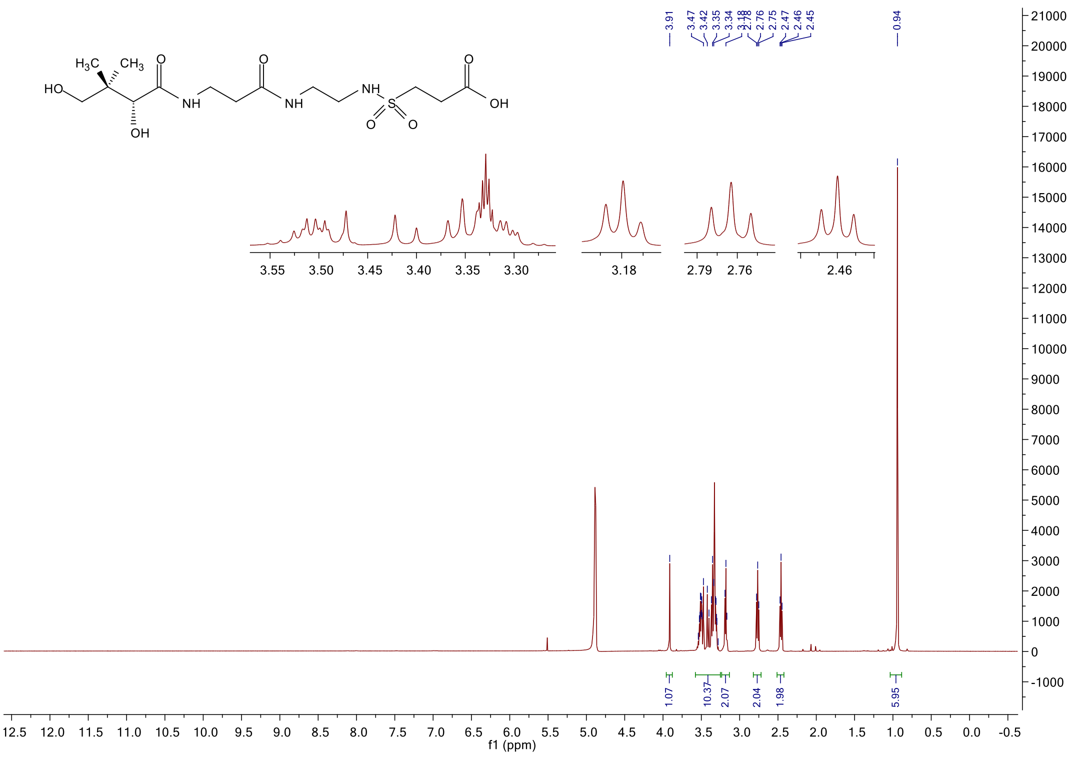

#### [13]C NMR spectrum of **15**

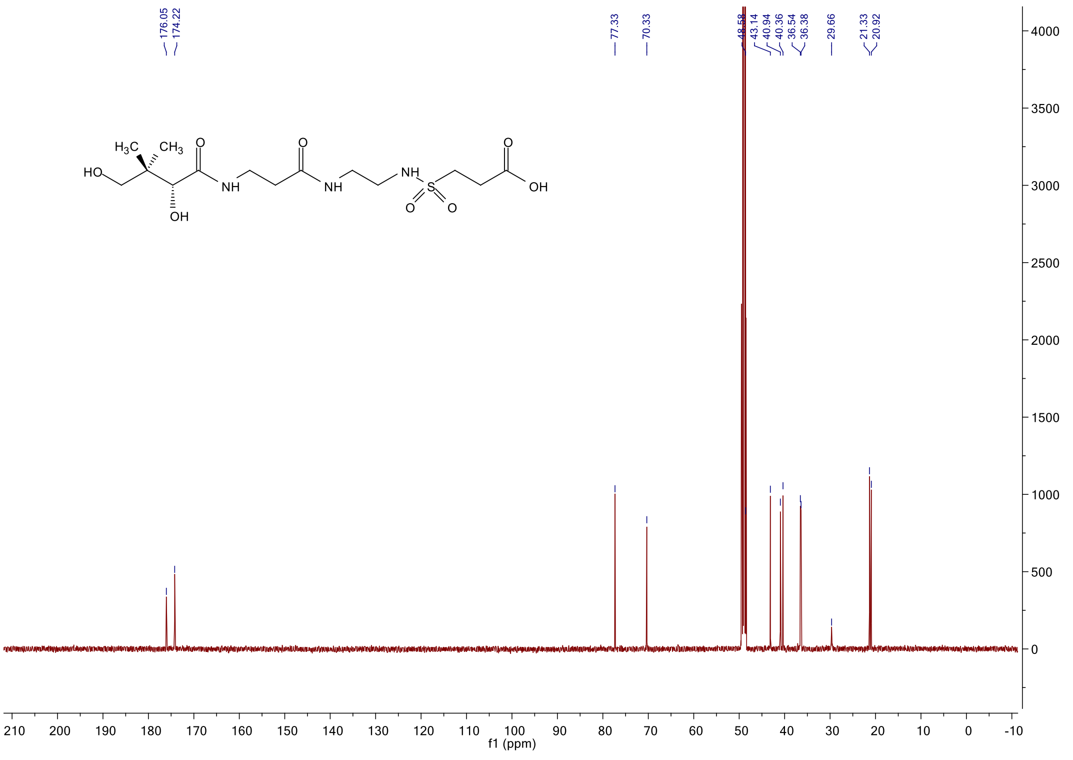

#### $^{1}$H NMR spectrum of **16**

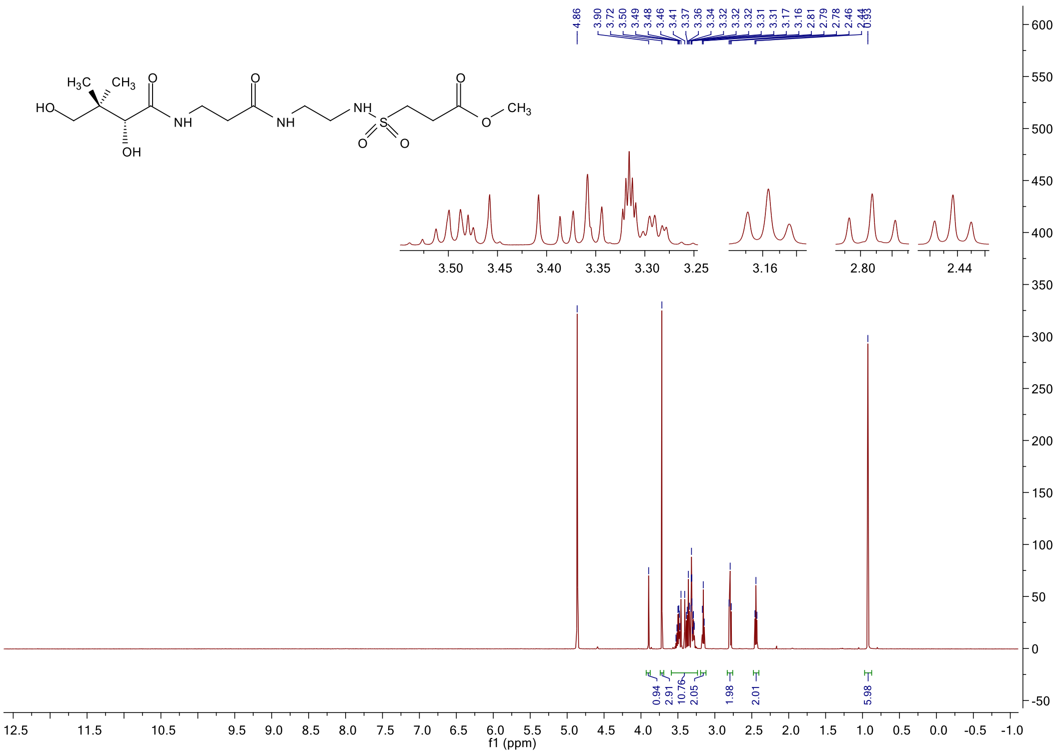

#### $^{13}$C NMR spectrum of **16**

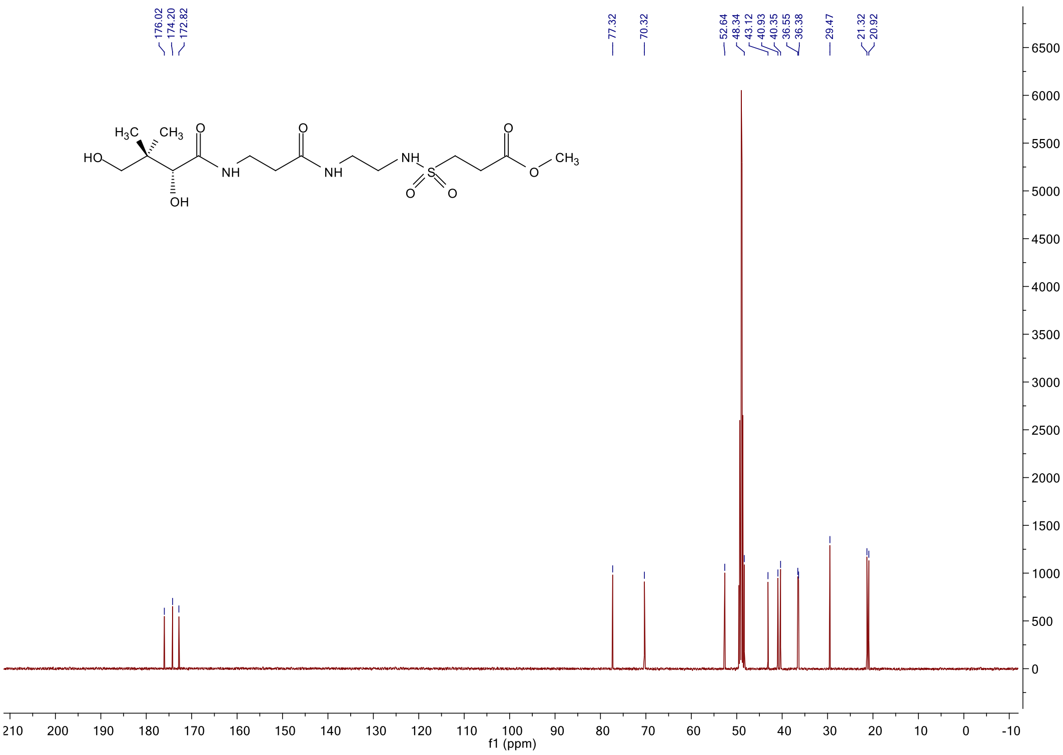

#### [1]H NMR spectrum of **22a**

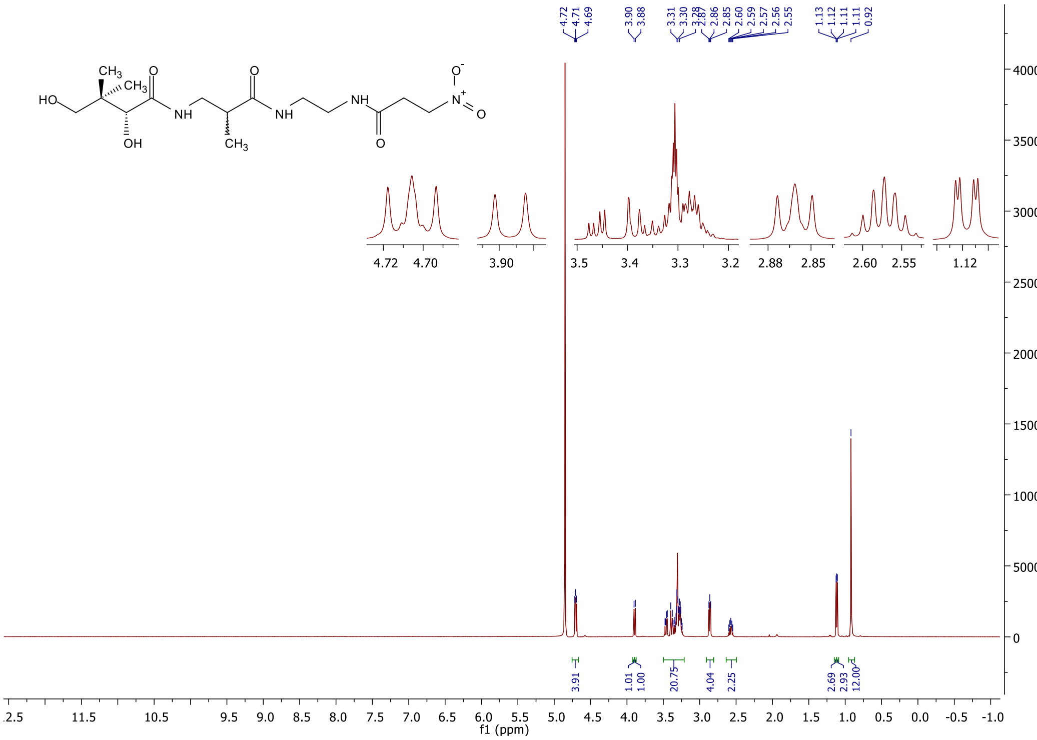

#### [13]C NMR spectrum of **22a**

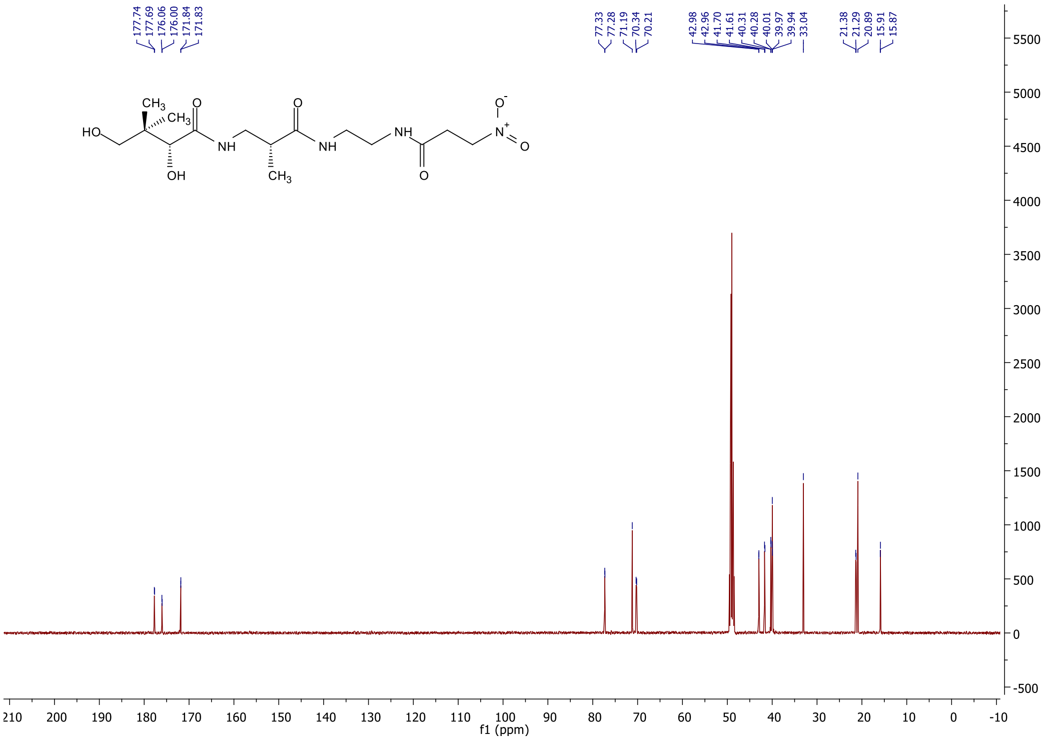

#### $^1$H NMR spectrum of **22b**

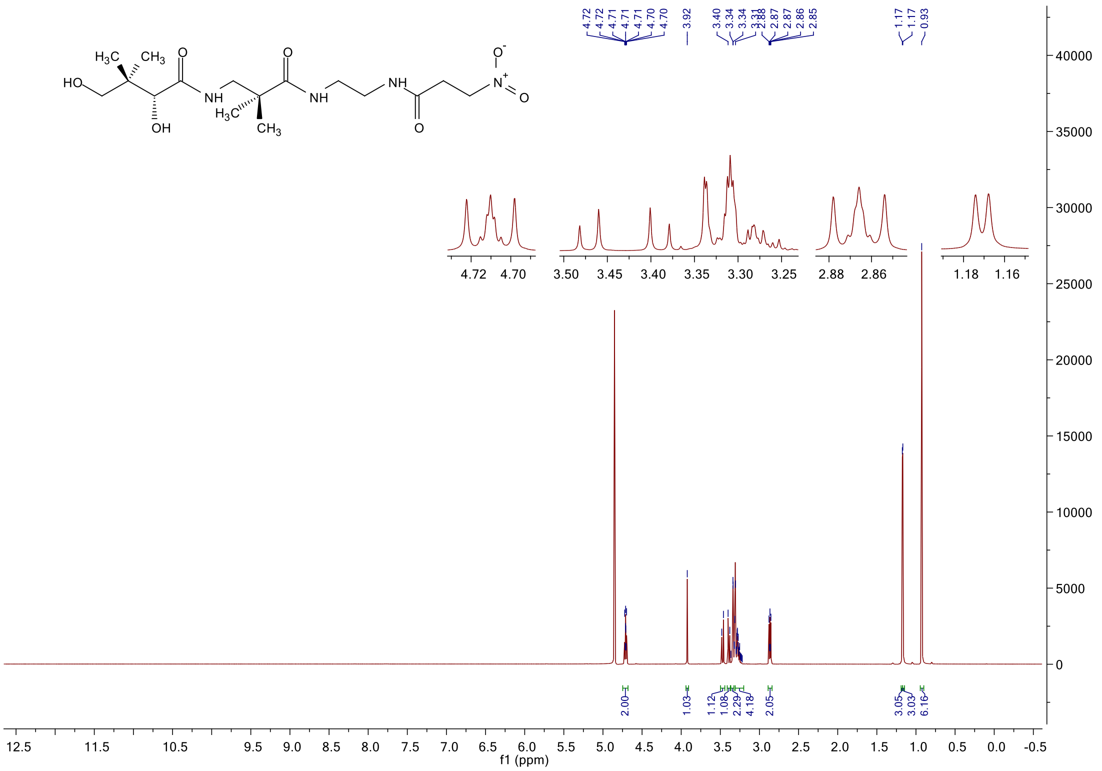

#### $^{13}$C NMR spectrum of **22b**

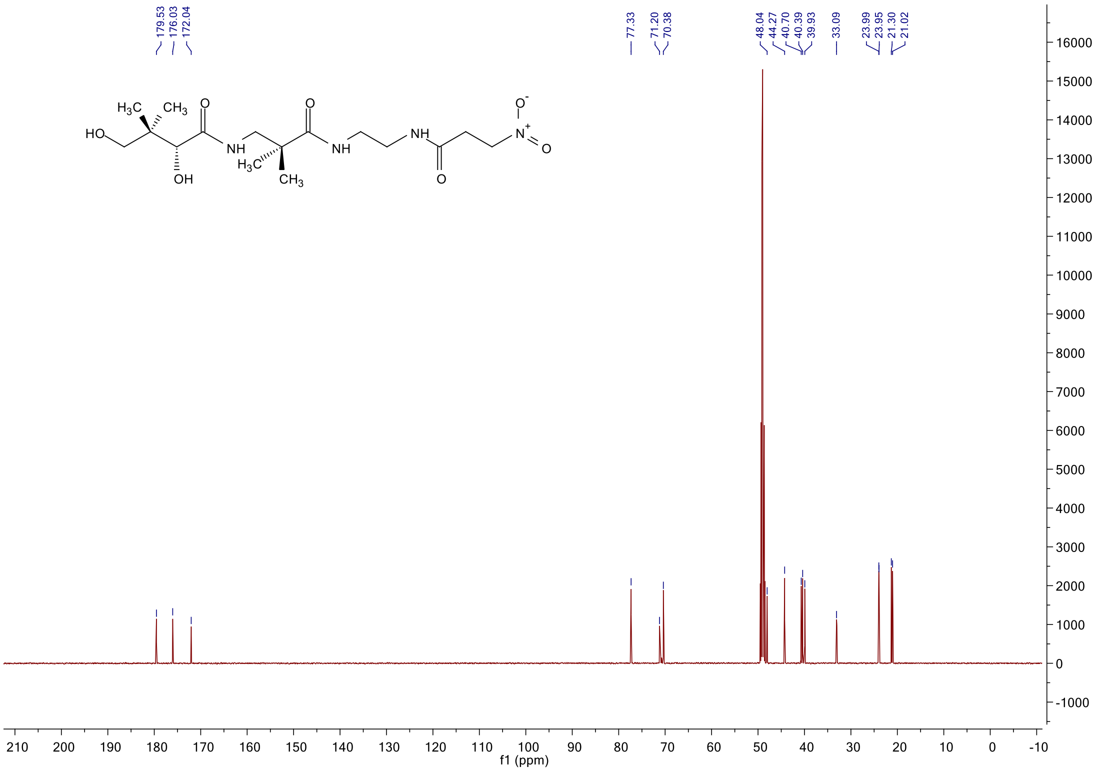

#### [1]H NMR spectrum of **22c**

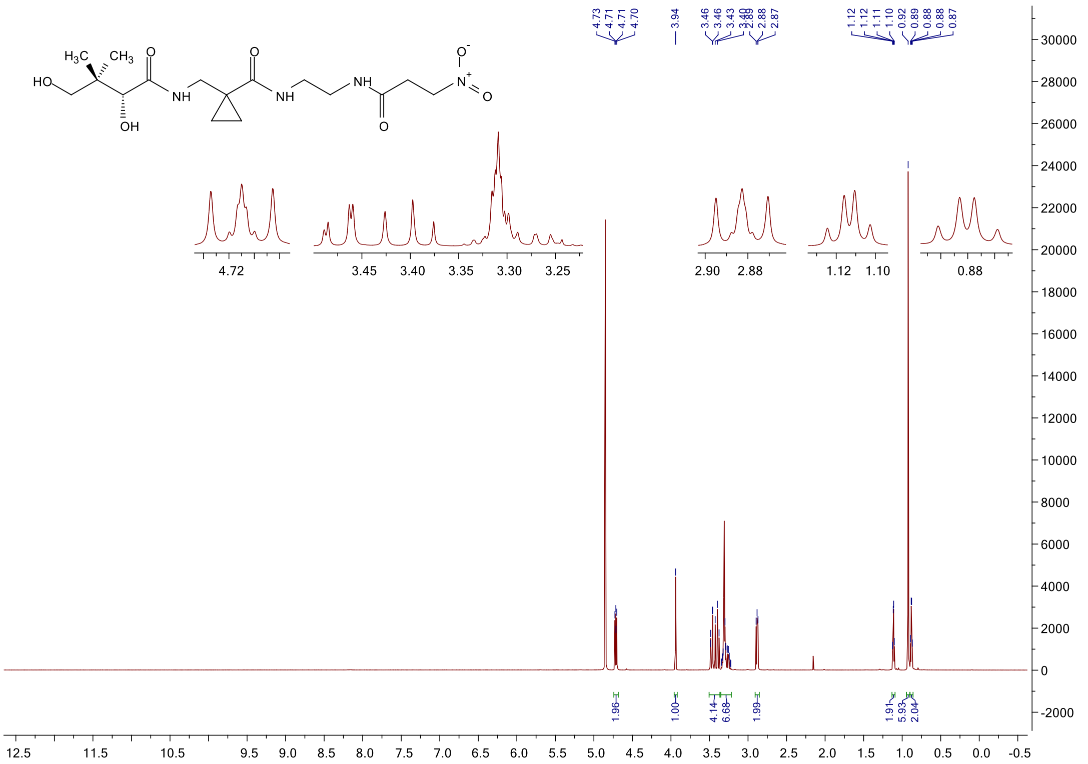

#### [13]C NMR spectrum of **22c**

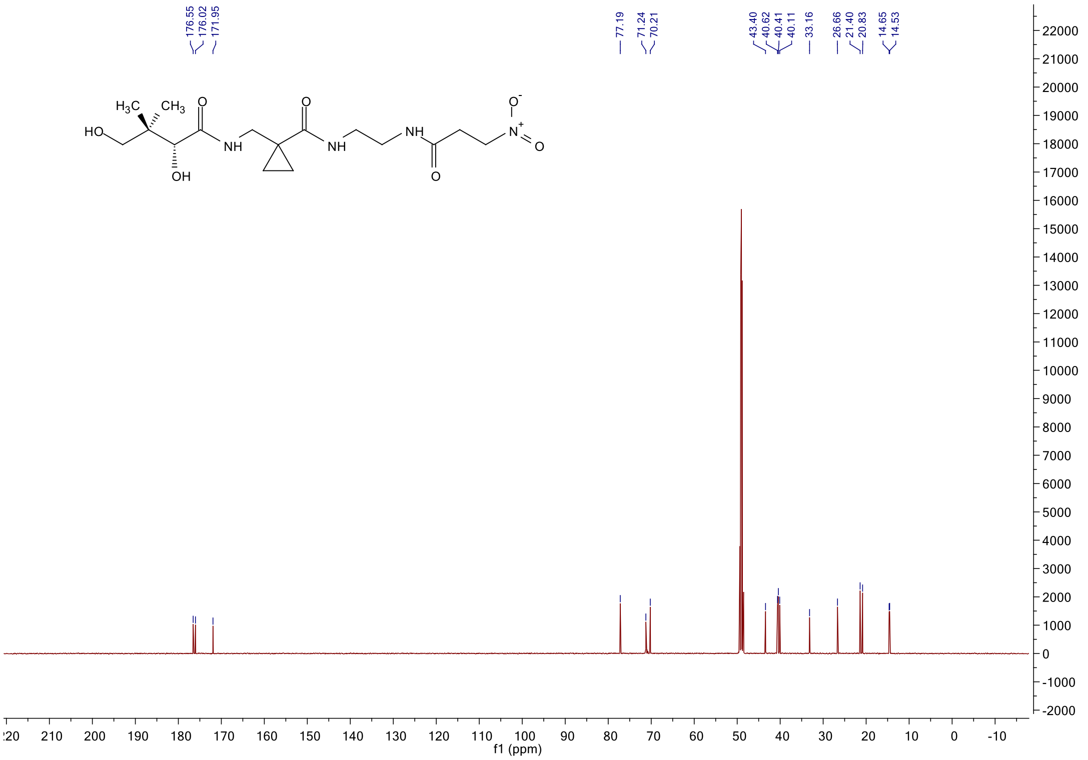

[1]H spectrum of **TFA-ethylenediamine hydrochloride**

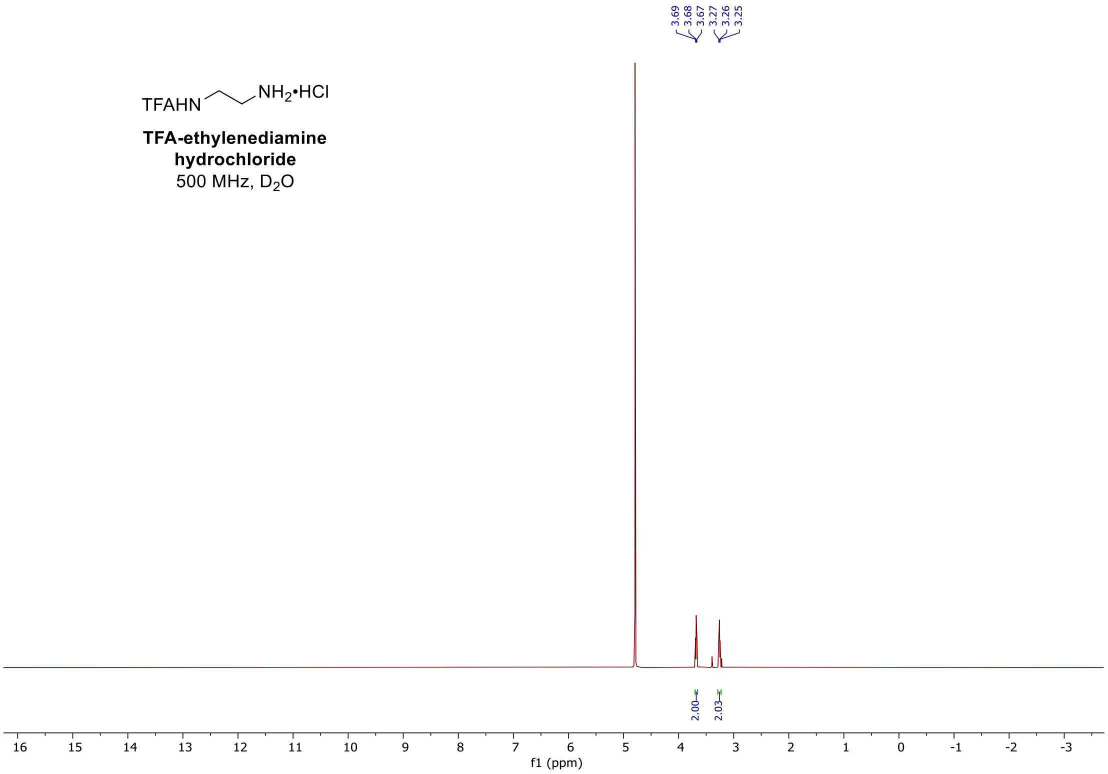

[1]H spectrum of **BCL-1-105**

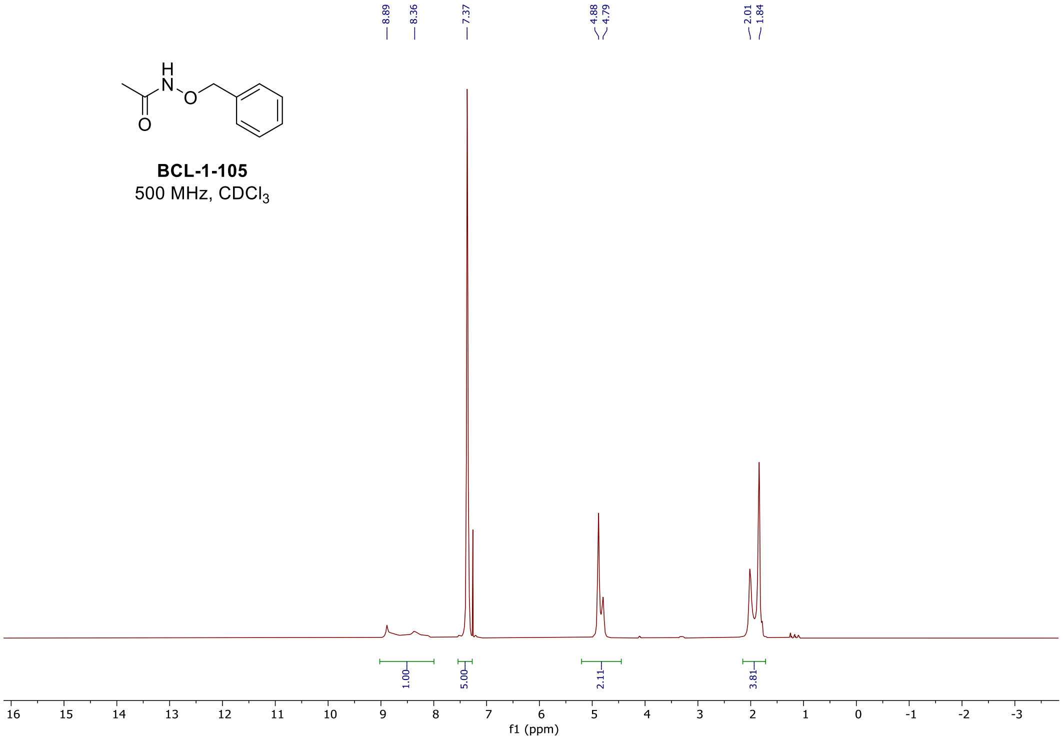

$^{1}$H spectrum of **BCL-1-110**

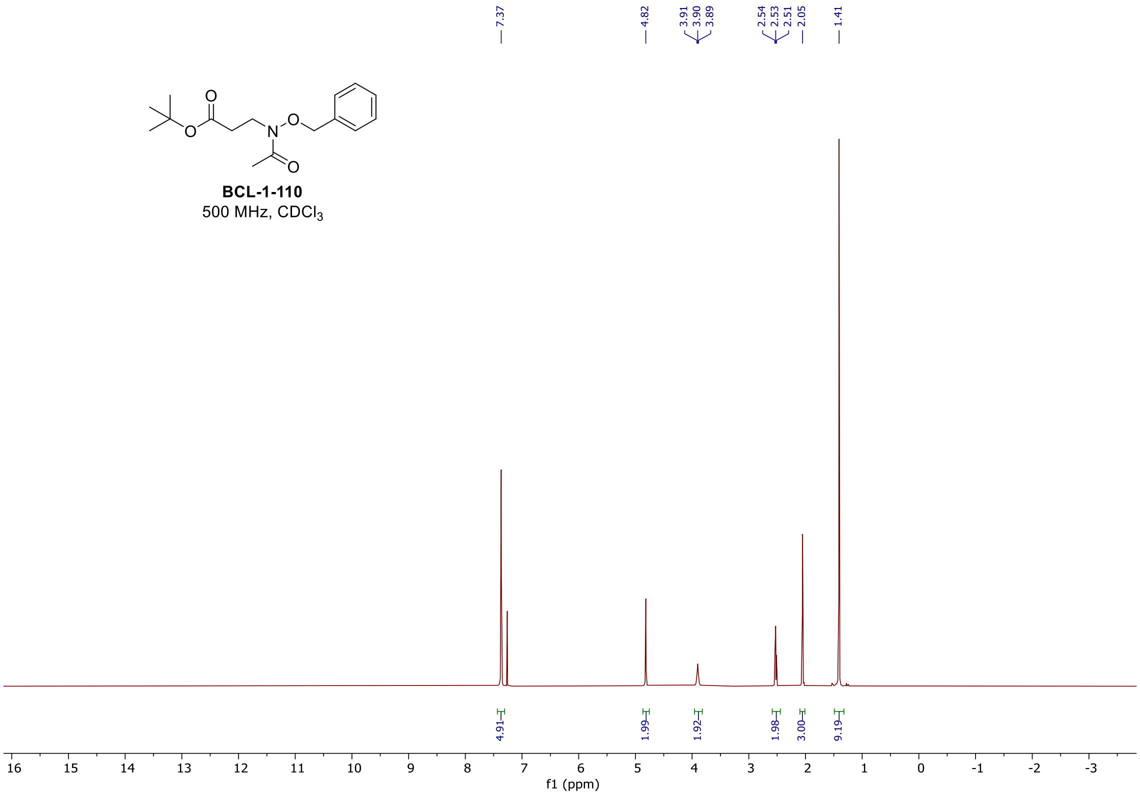

$^{1}$H spectrum of **BCL-1-115**

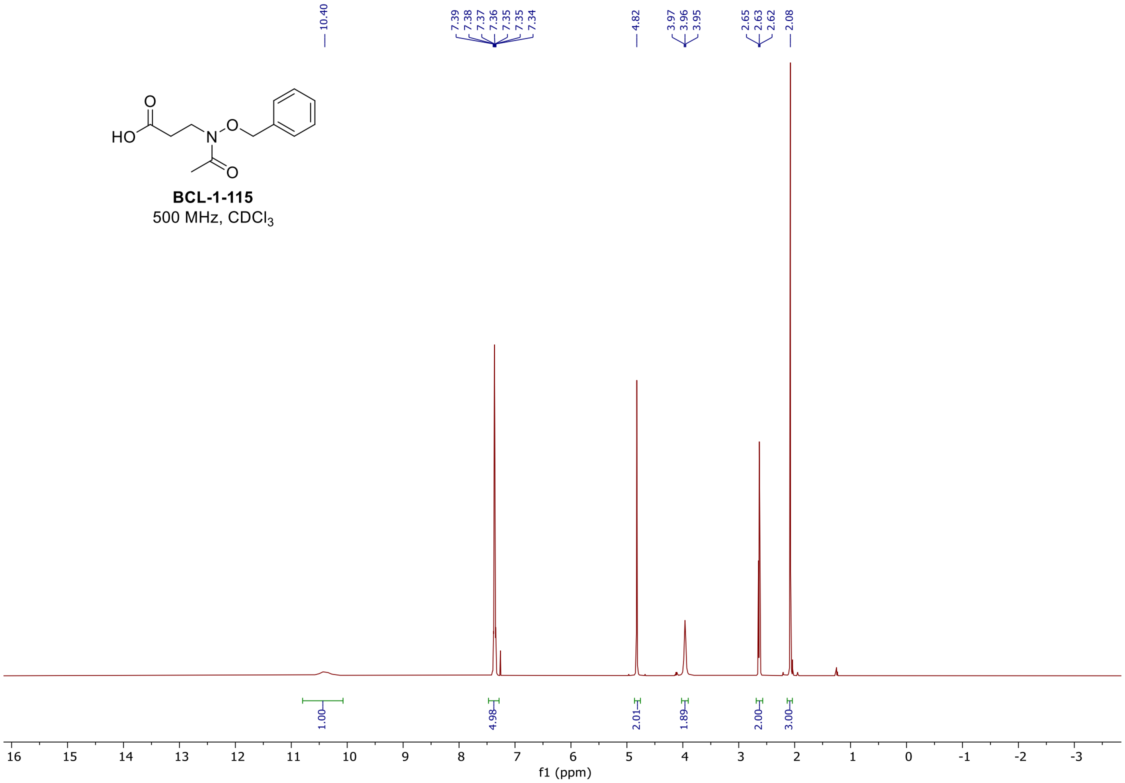

#### $^1$H spectrum of **BCL-2-83**

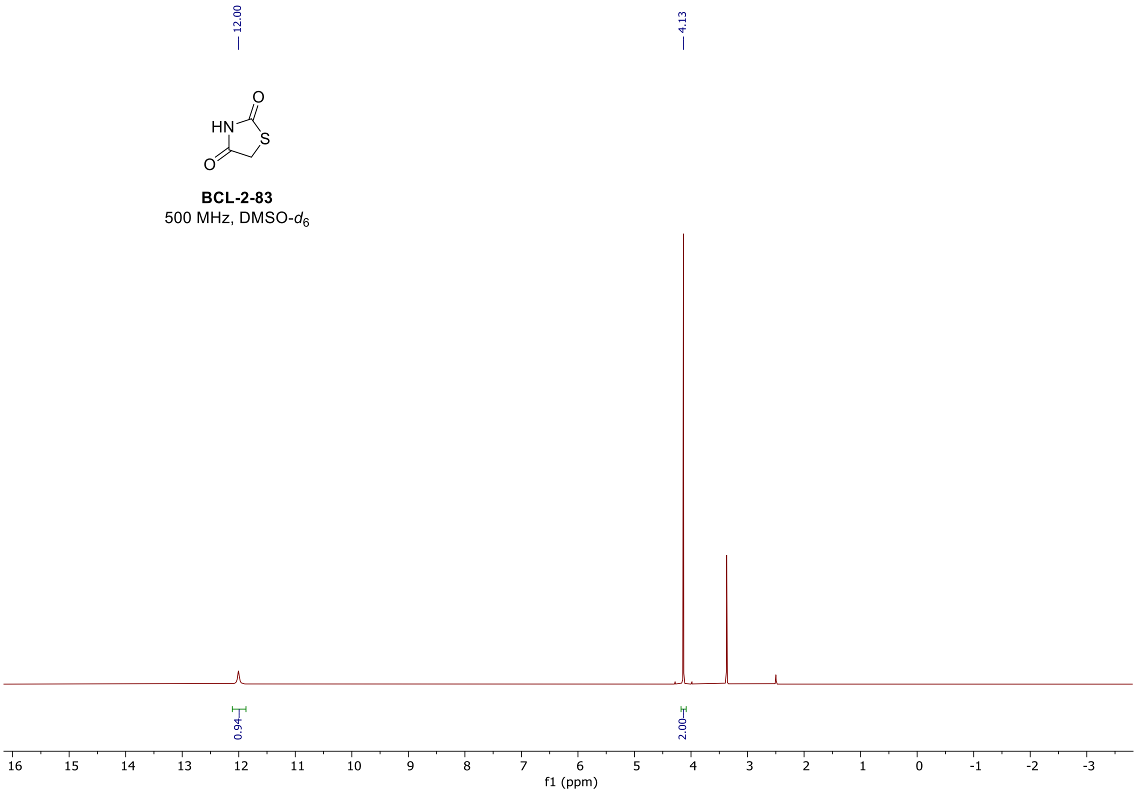

#### $^1$H spectrum of **BCL-2-93**

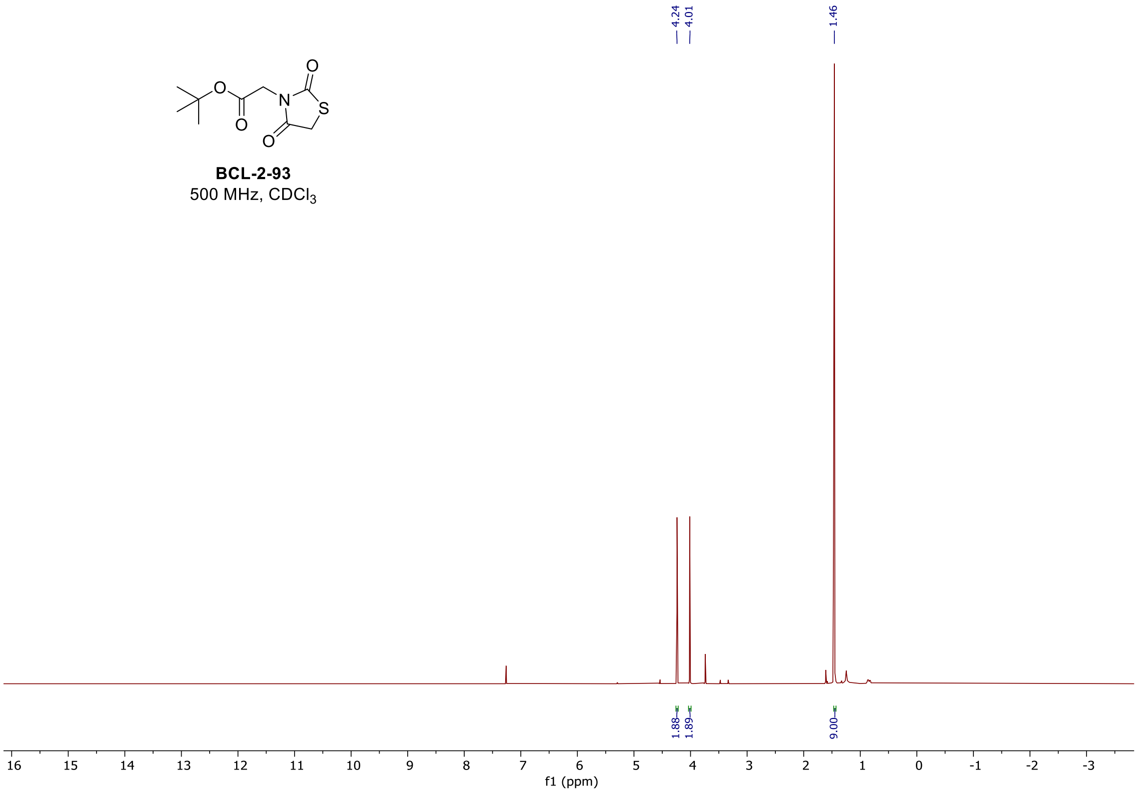

#### [1]H spectrum of **BCL-2-101**

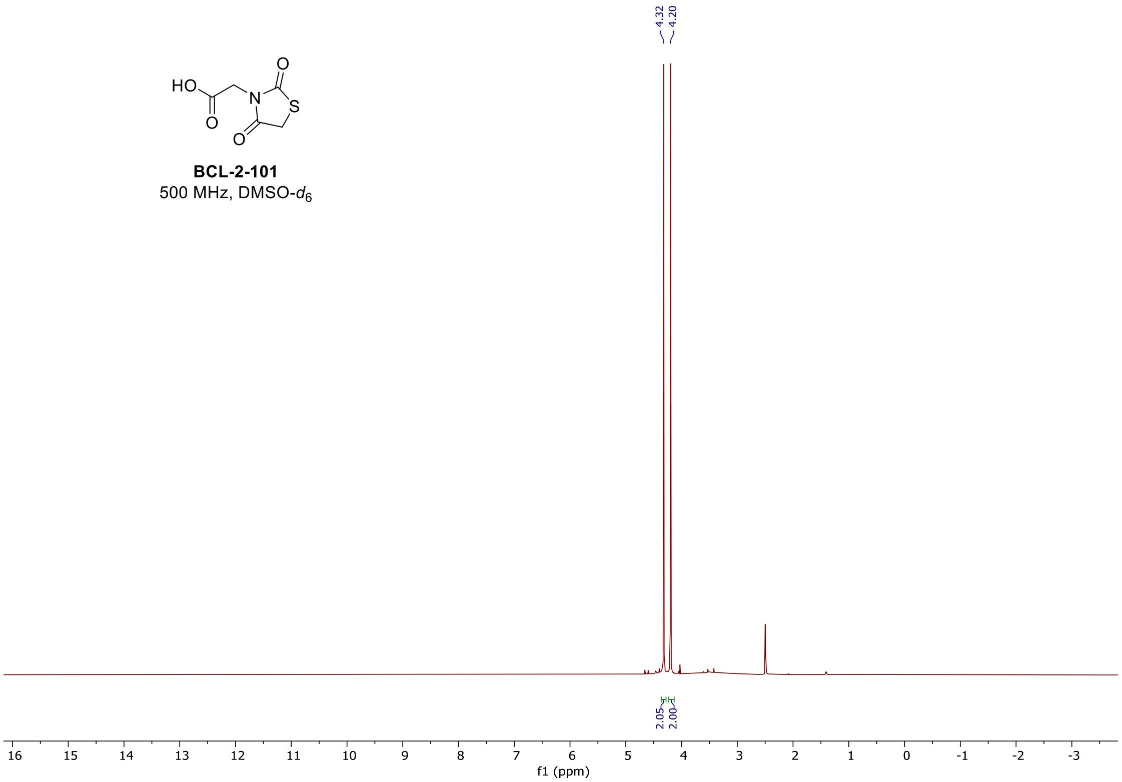

#### [1]H spectrum of **BCL-4-10**

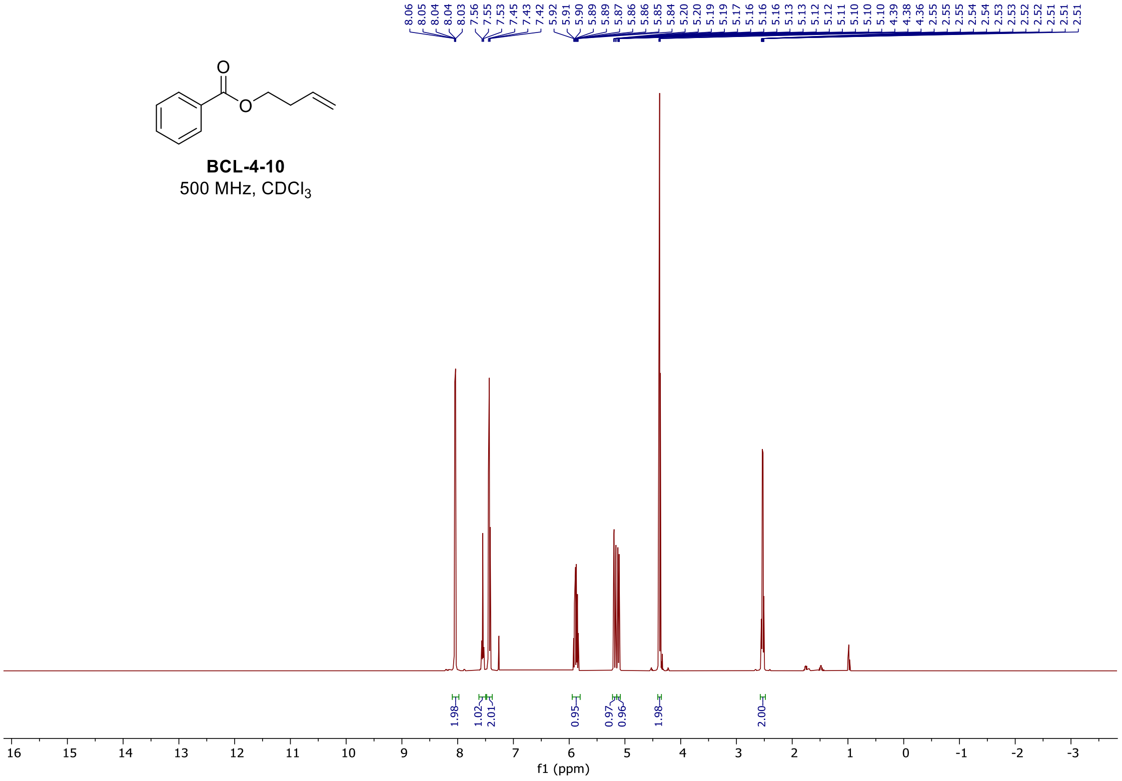

#### $^1H$ and $^{19}F$ spectra of **BCL-4-38**

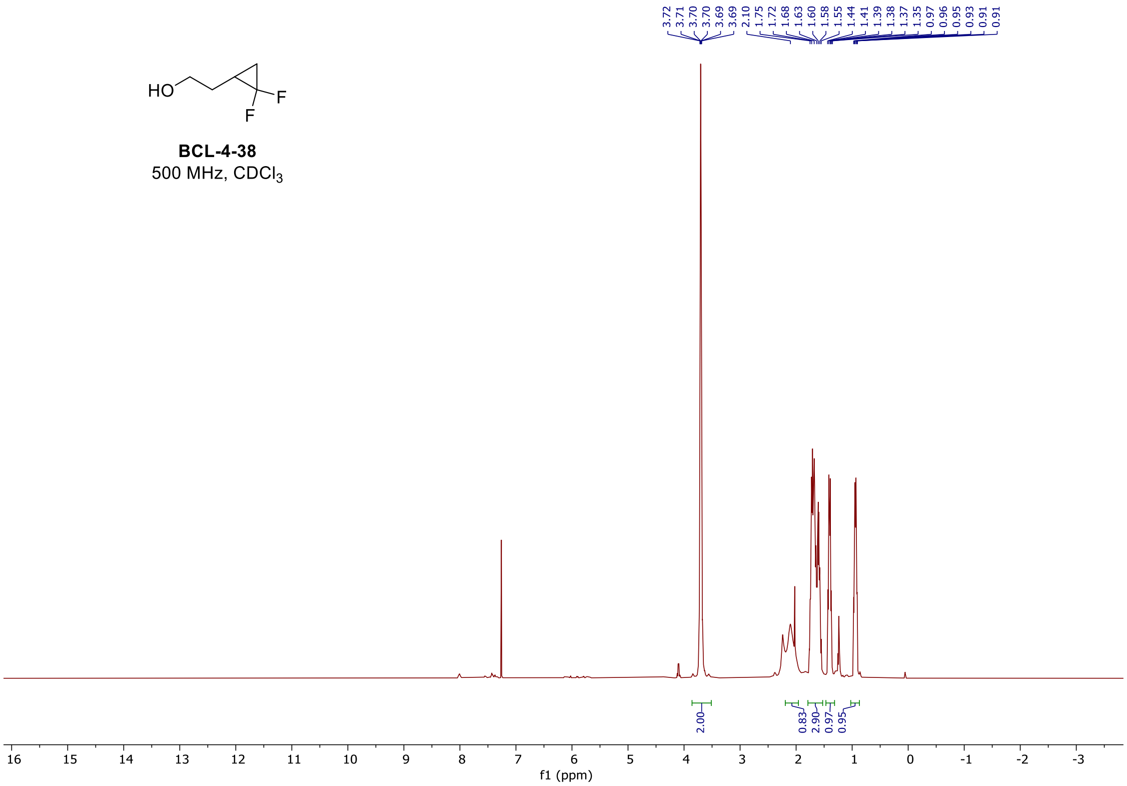

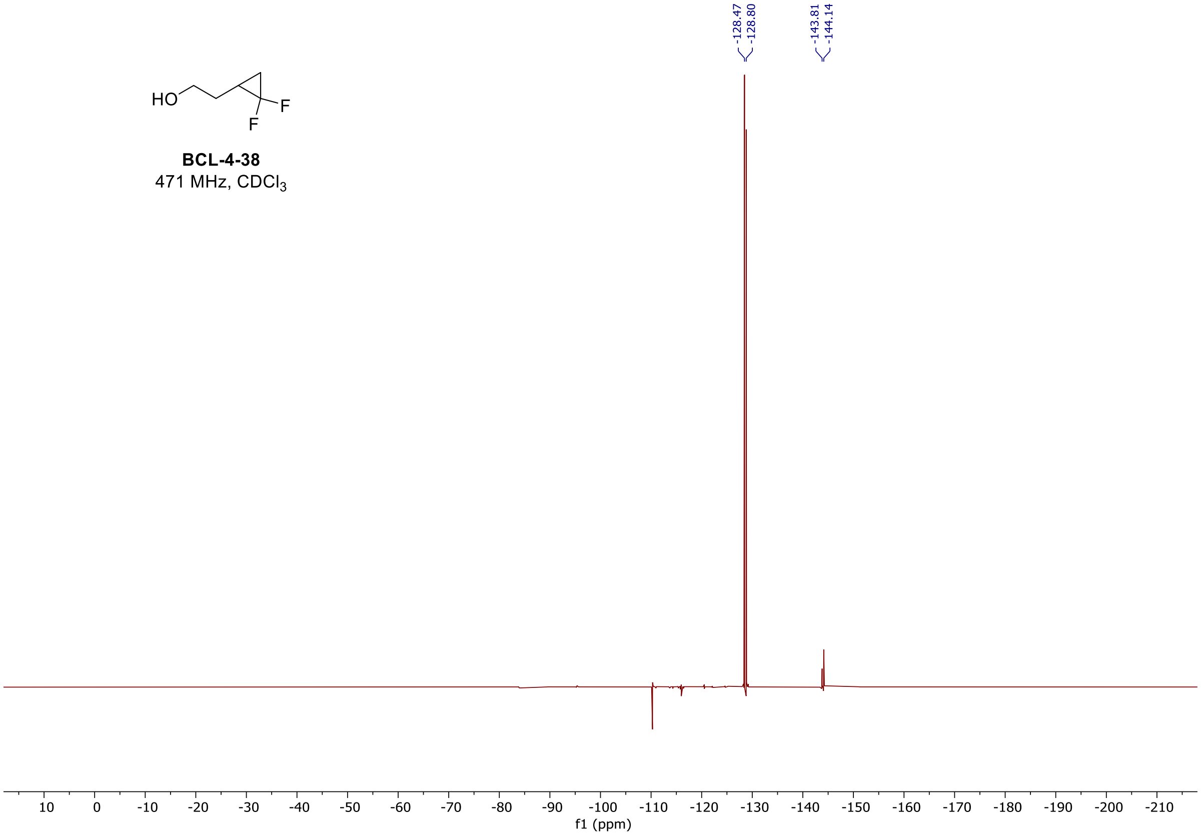

$^{1}H$ and $^{19}F$ spectra of **BCL-4-40**

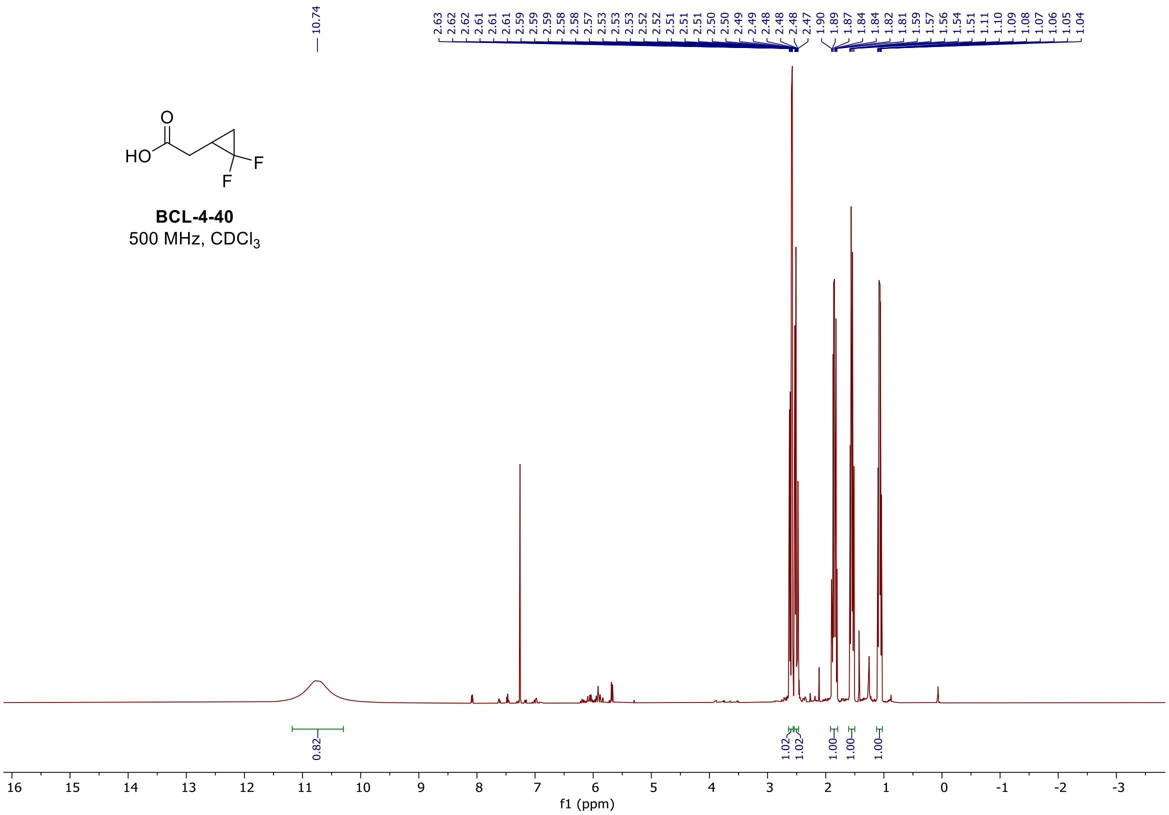

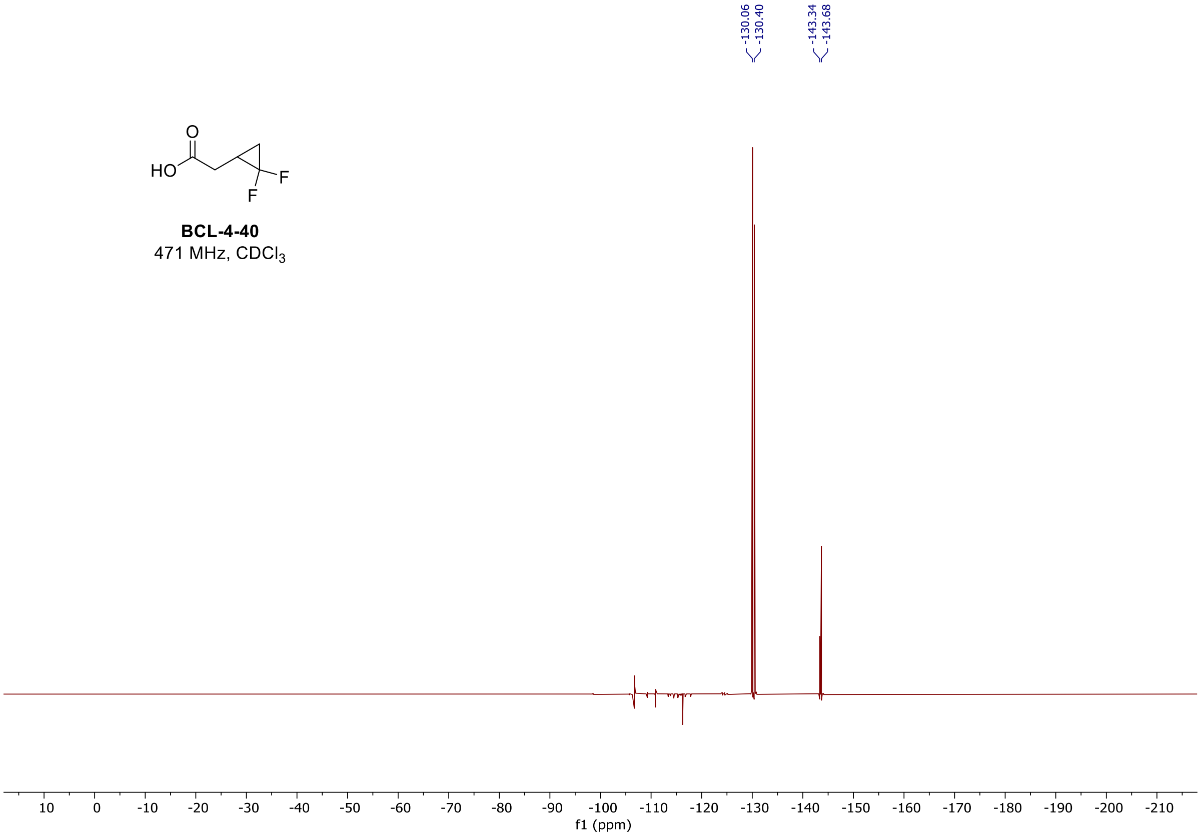

#### $^1$H spectrum of **BCL-4-50**

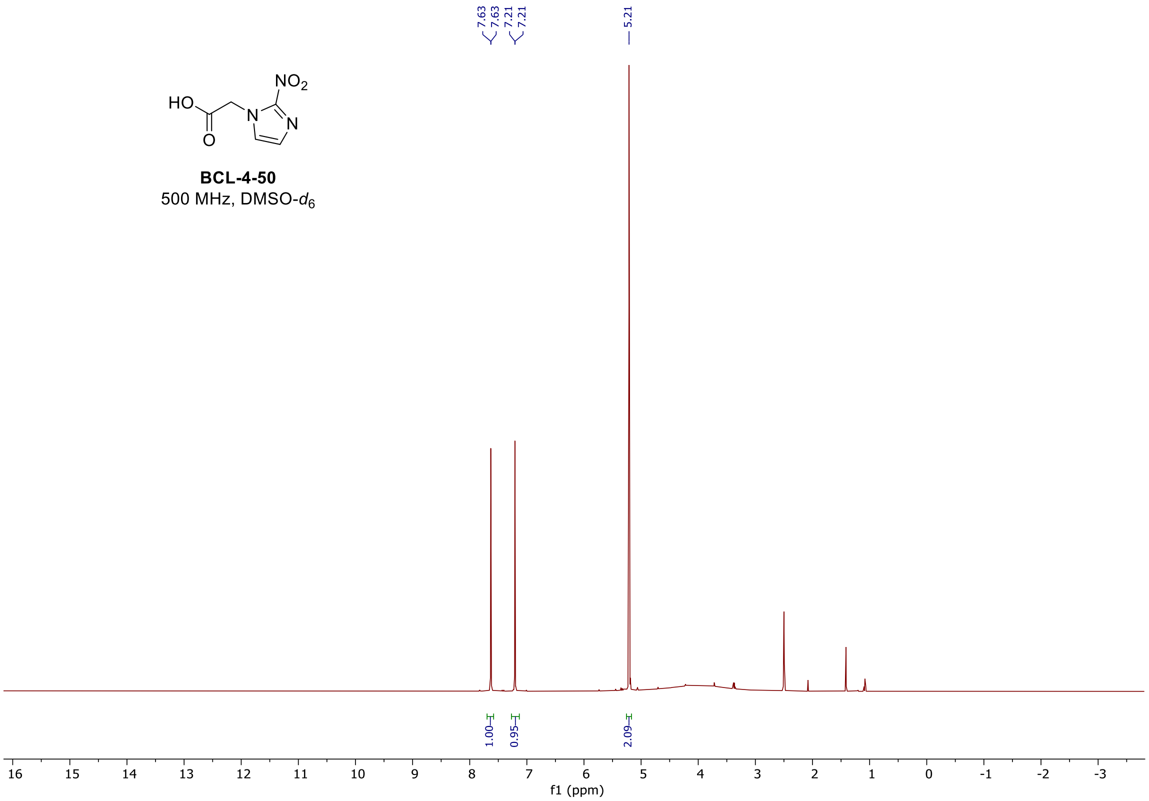

#### $^1$H spectrum of **BCL-4-53**

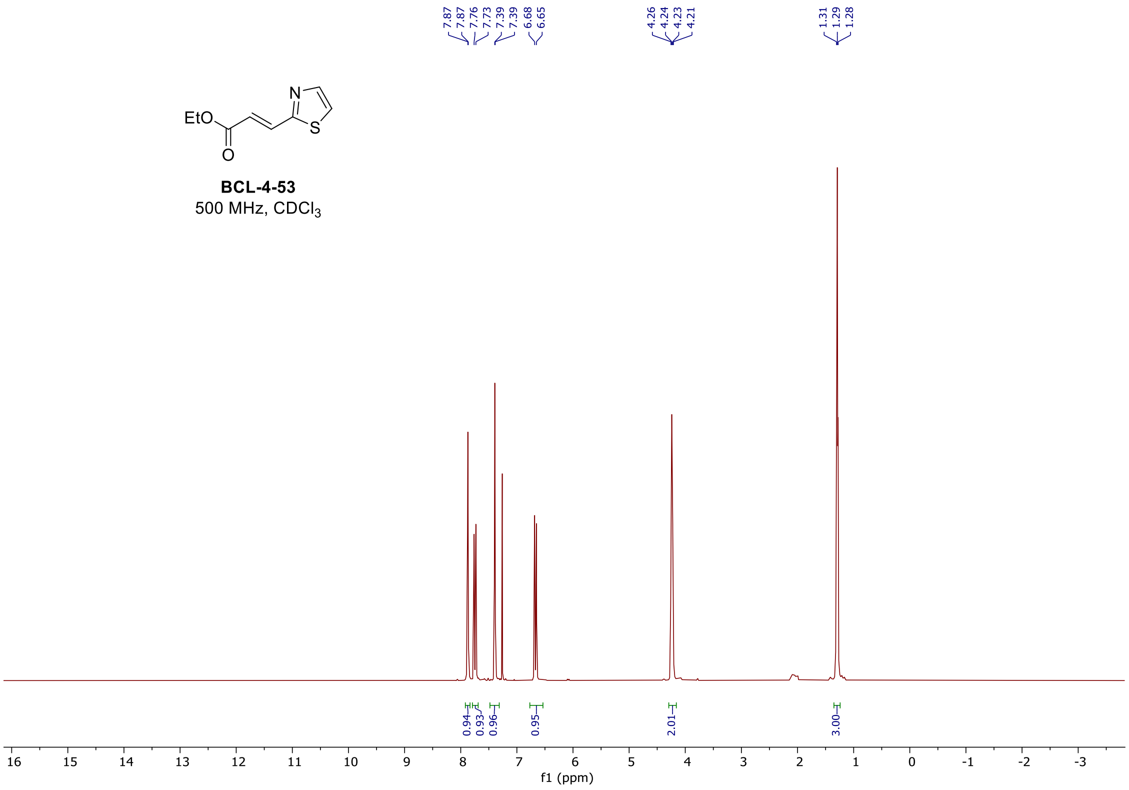

#### $^1$H spectrum of **BCL-4-65**

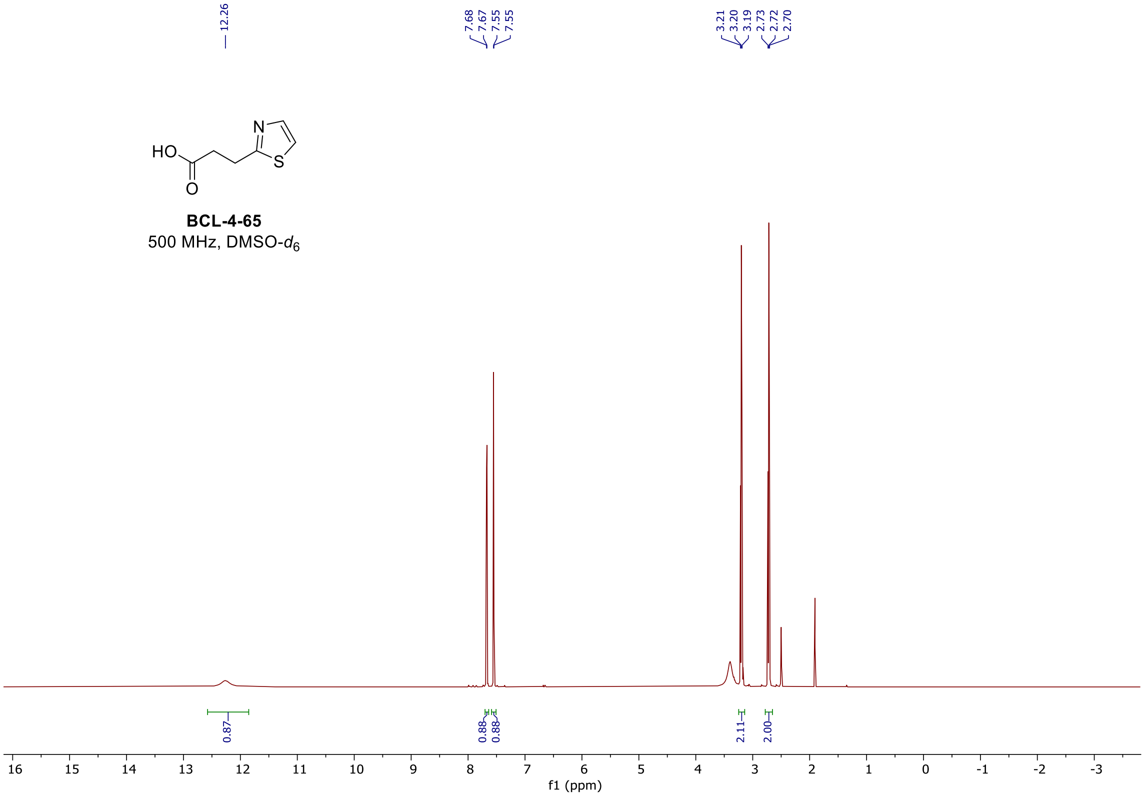

#### $^1$H spectrum of **BCL-7-40**

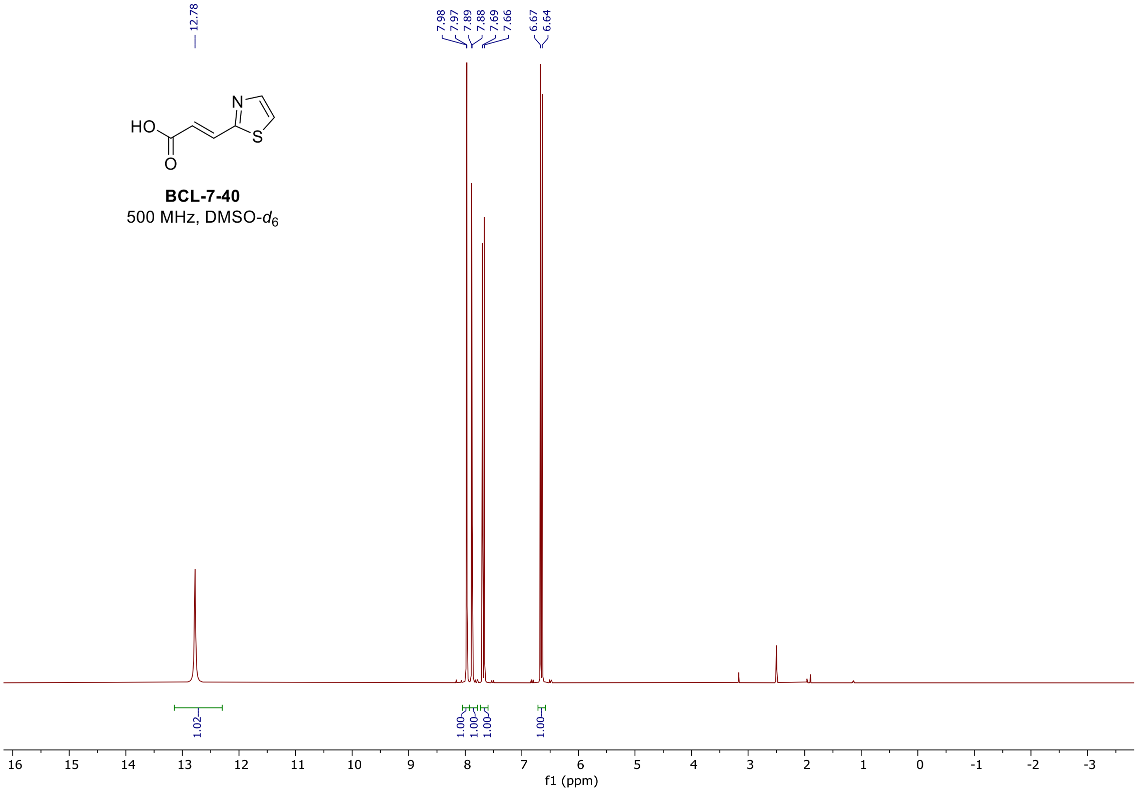

#### $^{1}H$ spectrum of **BCL-4-69**

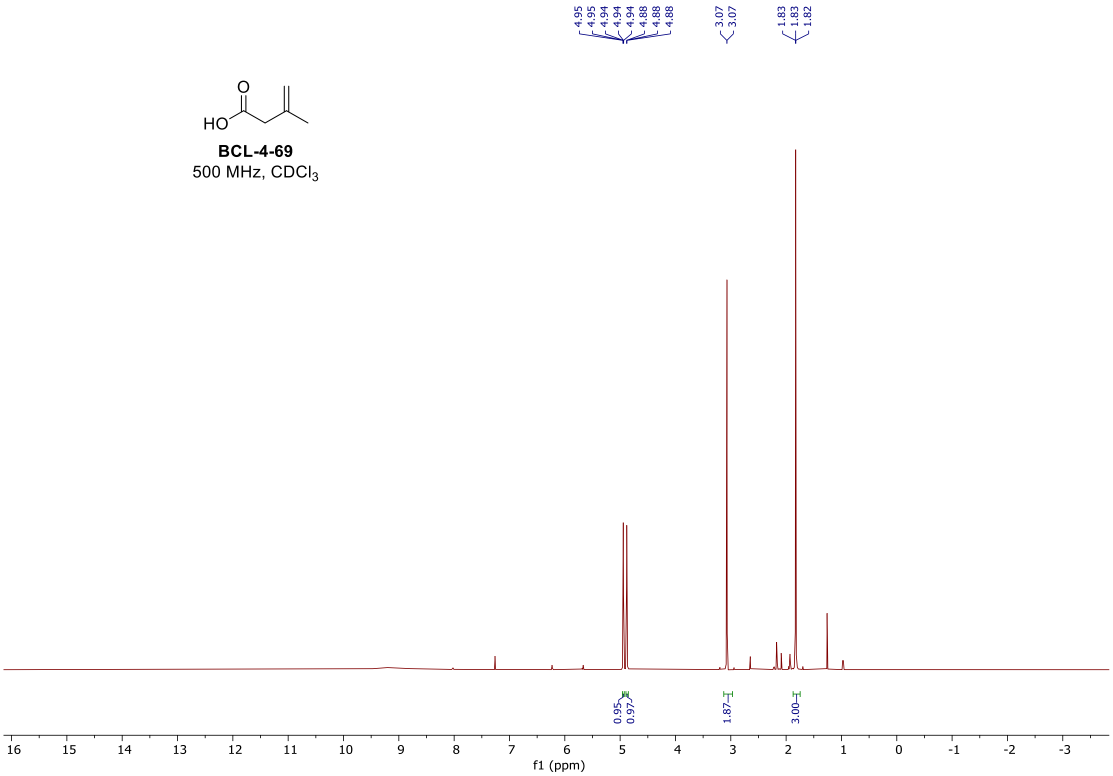

#### $^{1}H$ and $^{19}F$ spectra of **BCL-4-84**

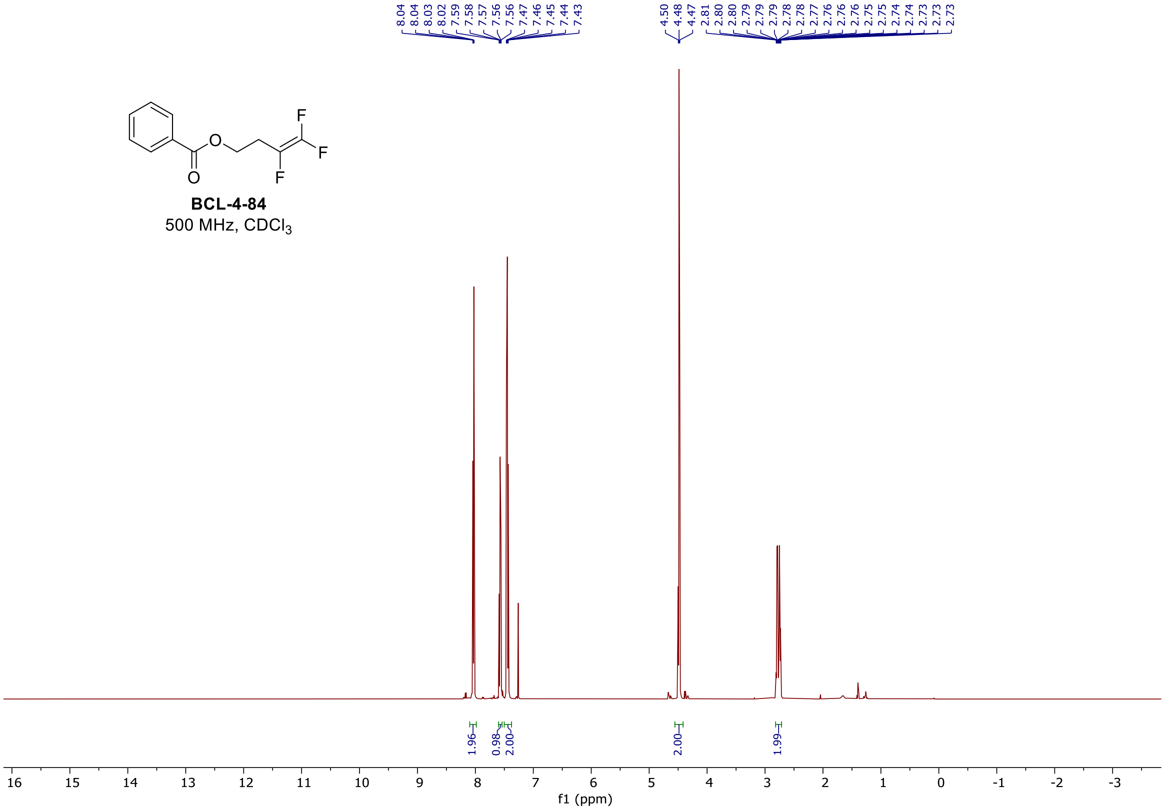

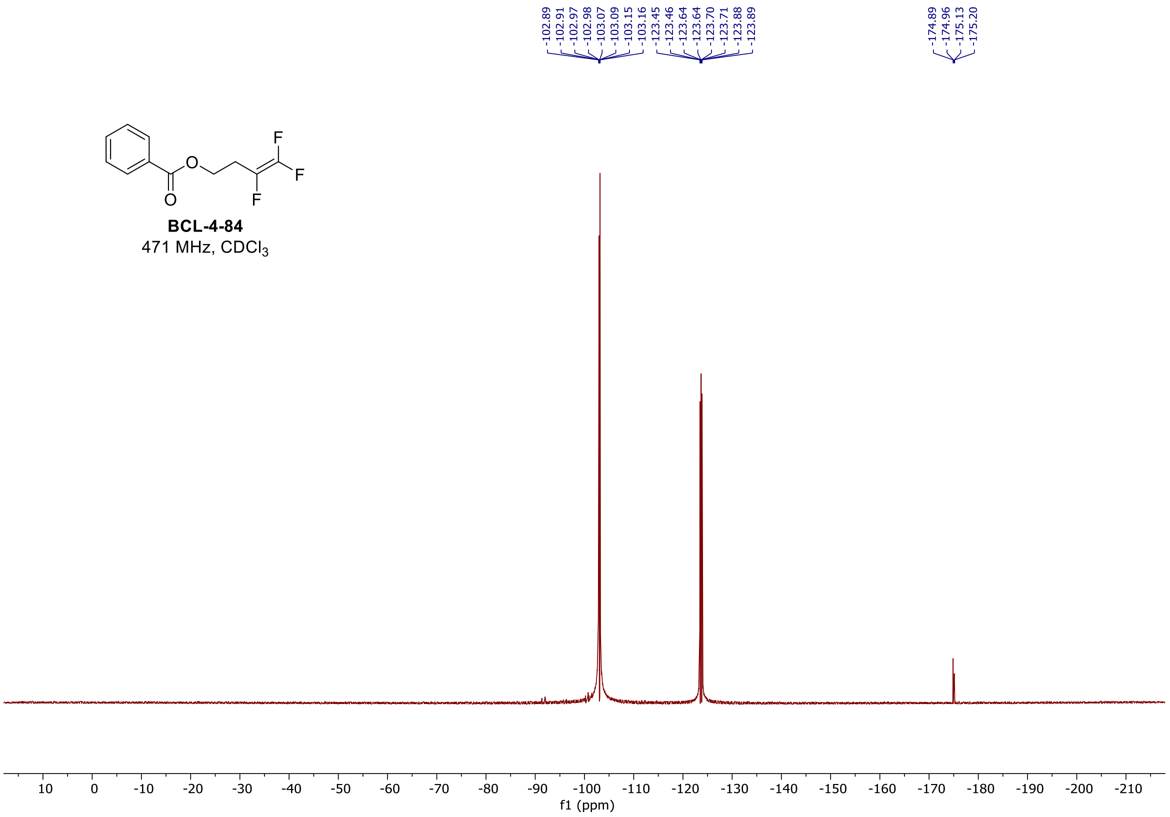

### [1]H spectrum of **BCL-4-101**

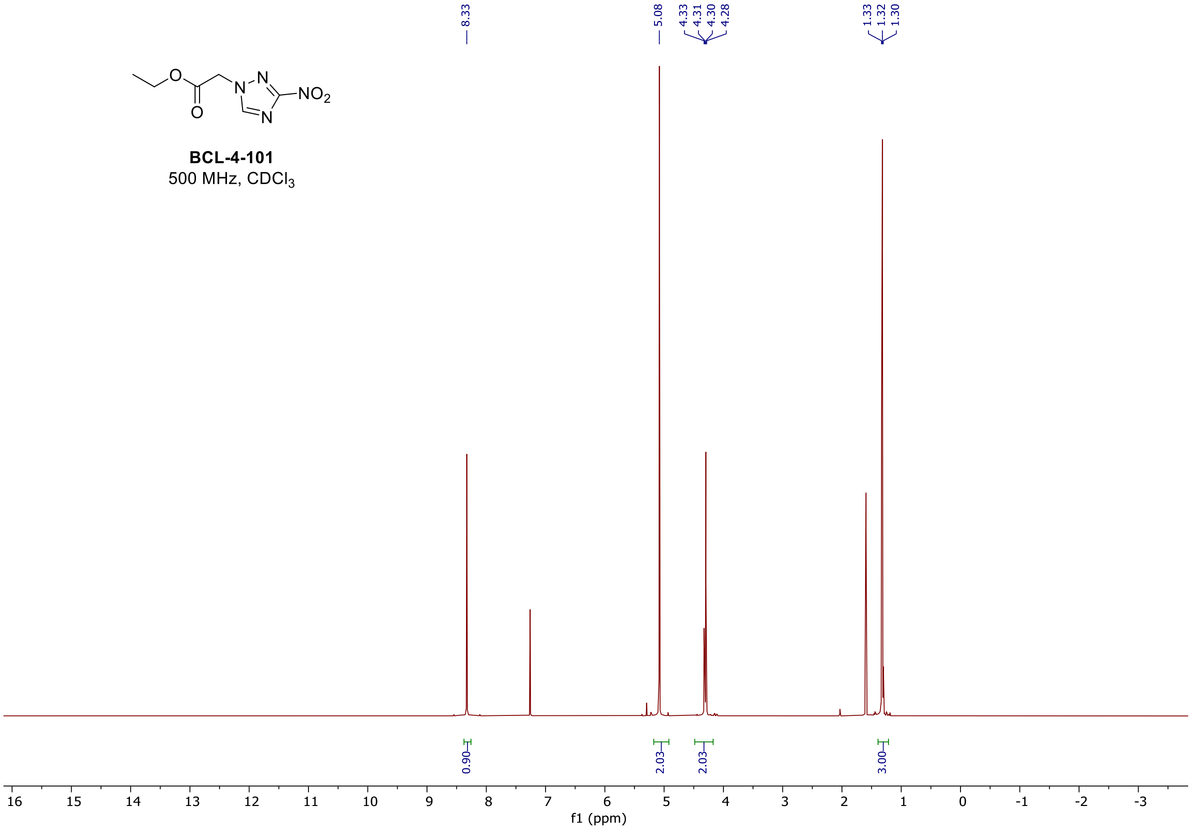

$^{1}$H spectrum of **BCL-4-107**

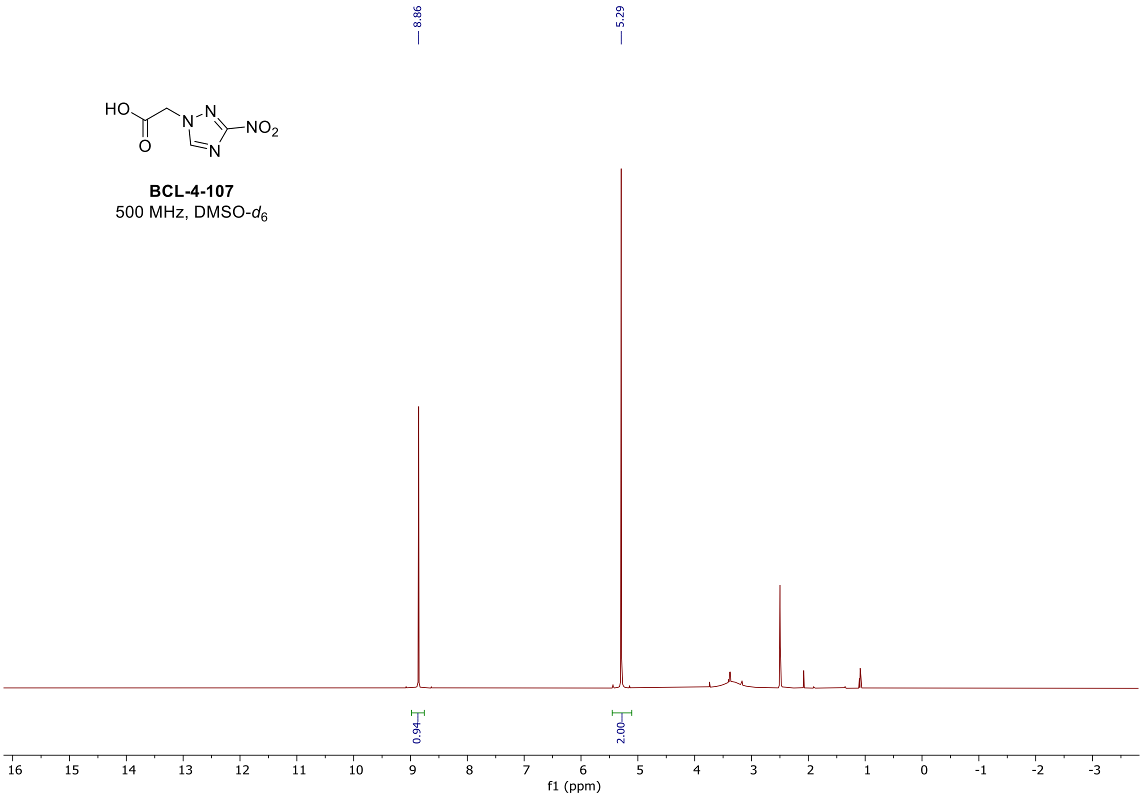

$^{1}$H spectrum of **BCL-4-82**

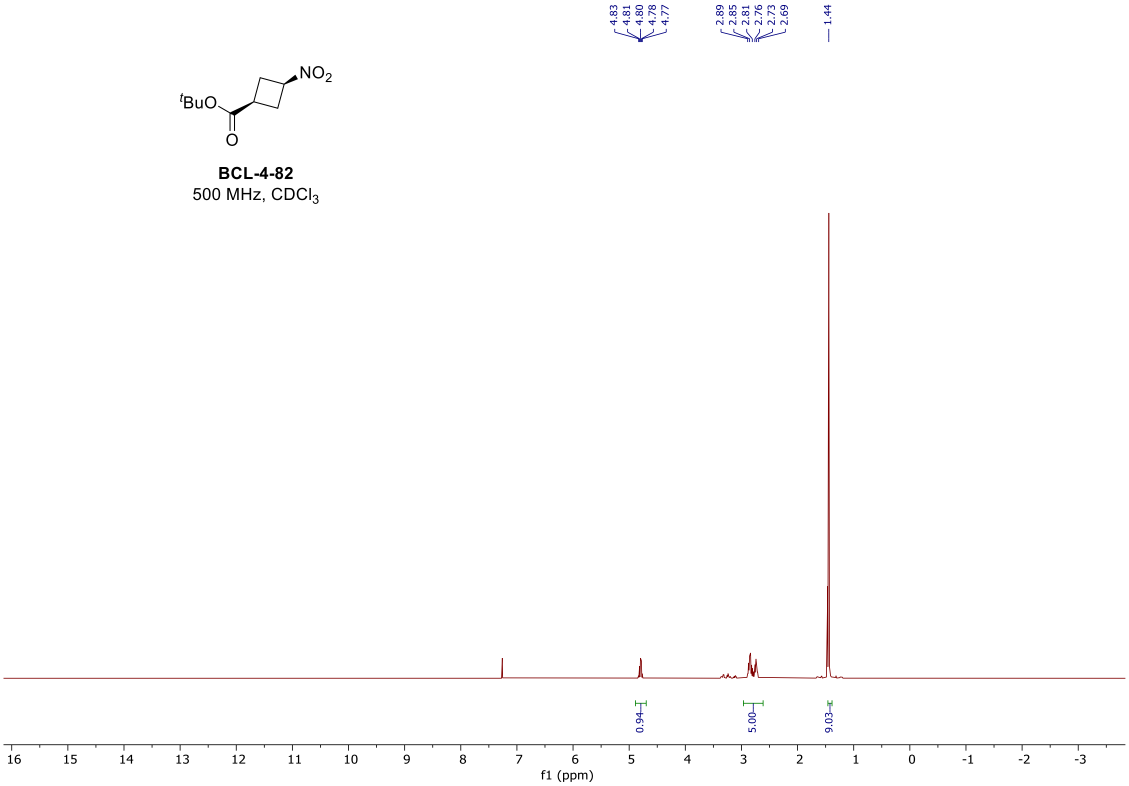

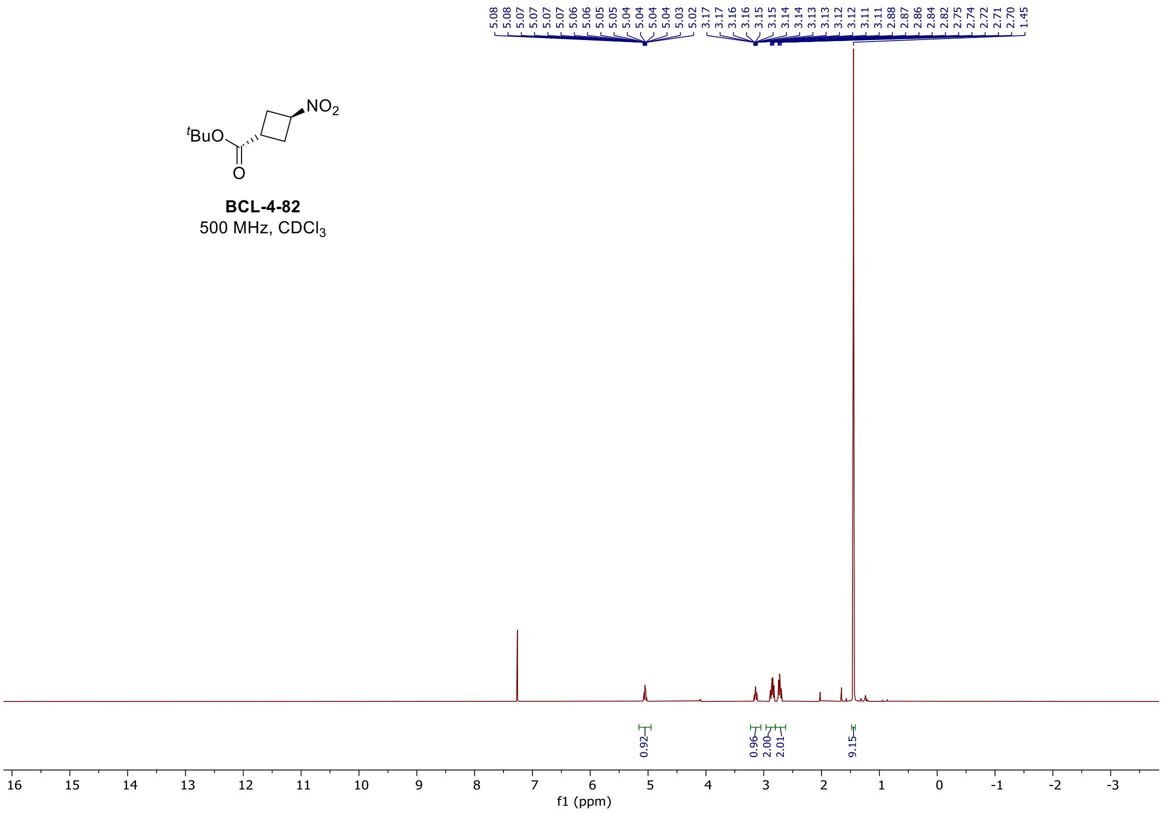

### $^{1}$H and $^{13}$C spectra of **BCL-5-52**

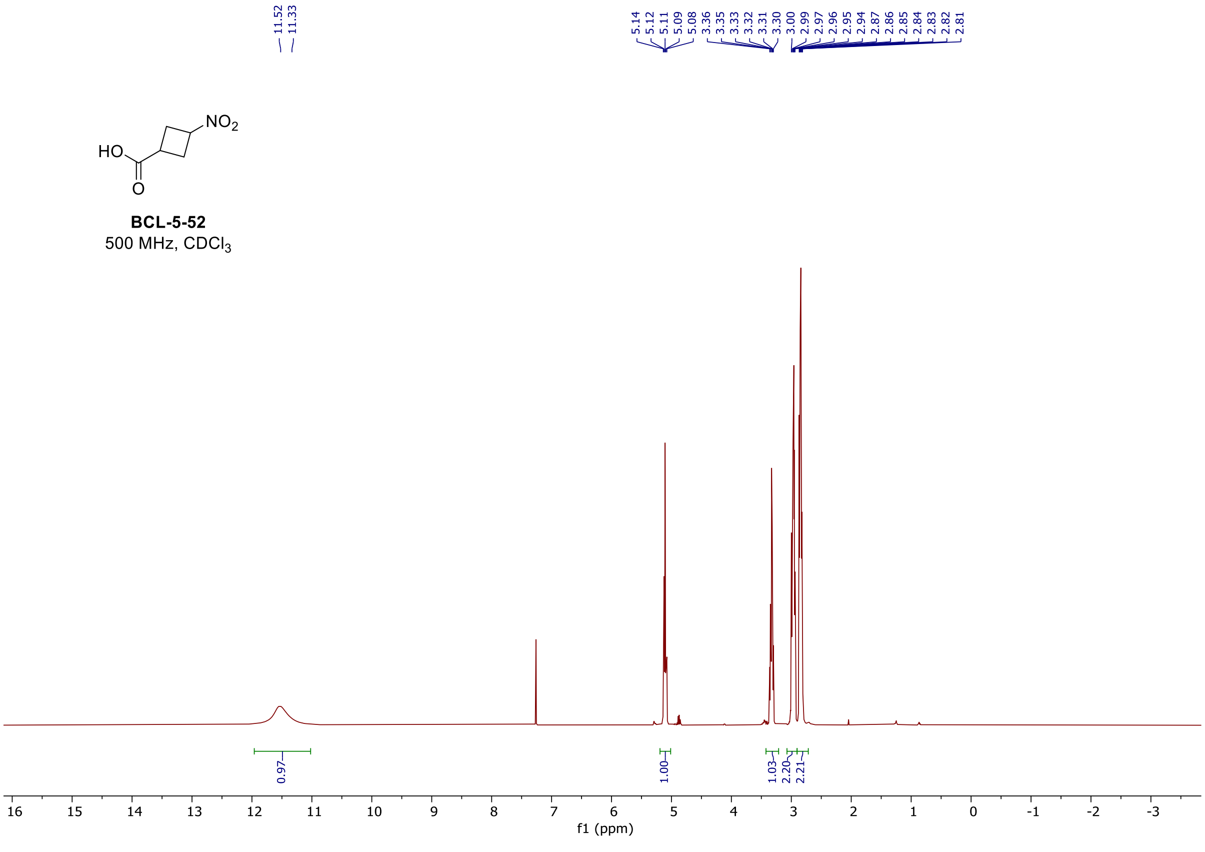

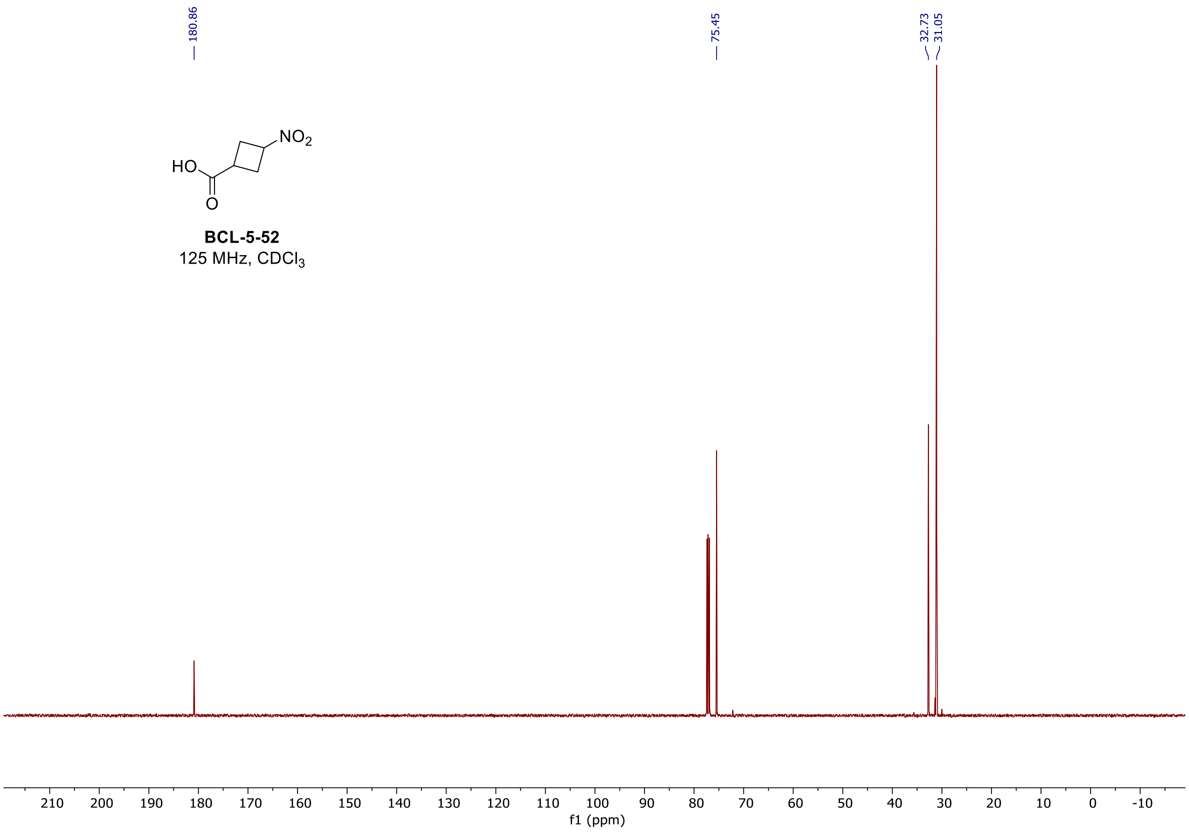

#### $^{1}$H spectrum of **BCL-7-14**

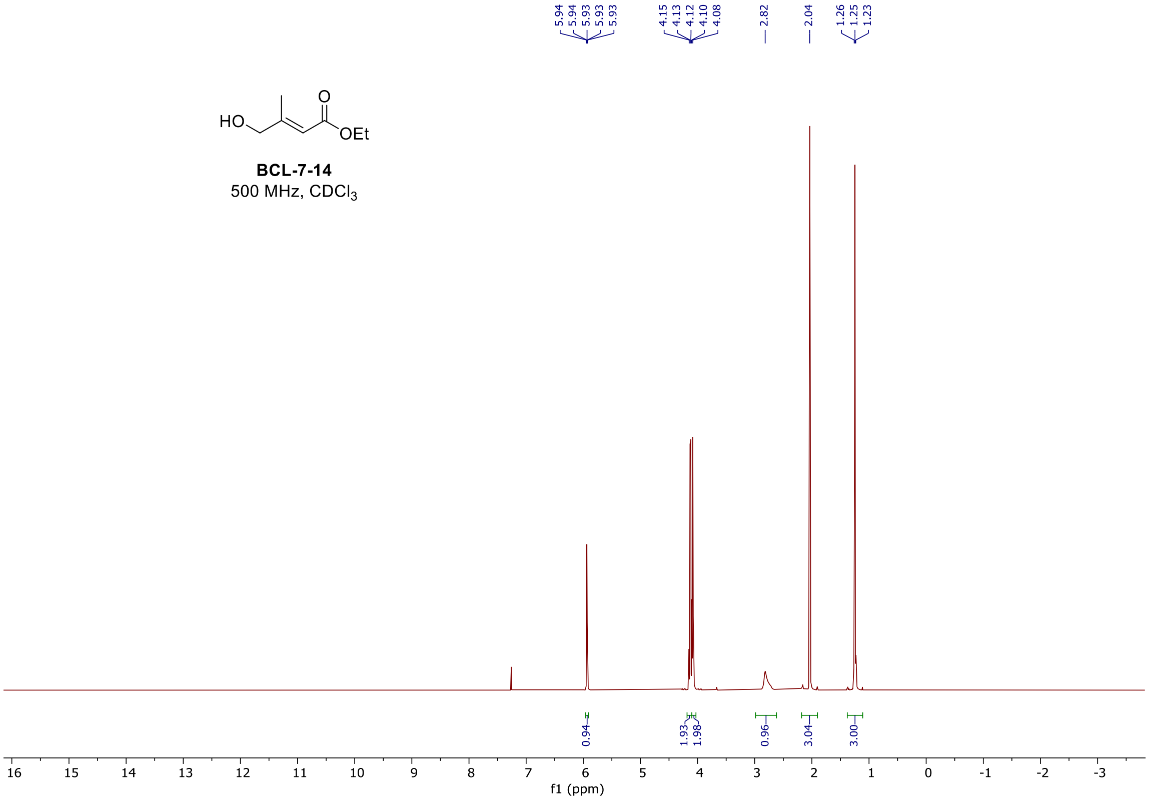

$^{1}$H spectrum of **BCL-7-19**

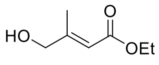

**BCL-7-19**
500 MHz,DMSO-$d_6$

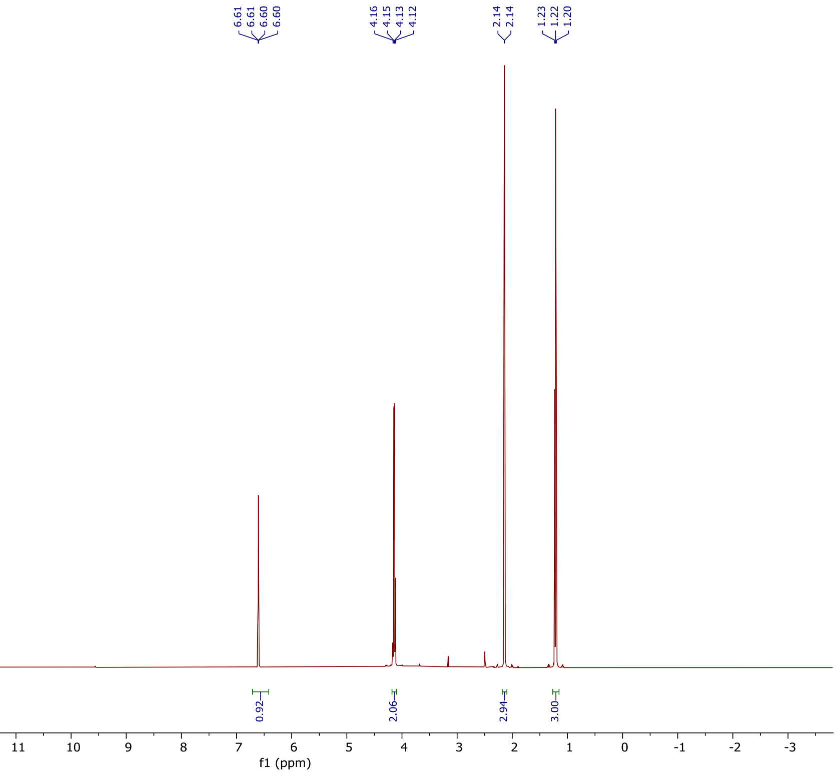

#### $^{1}H$ and $^{19}F$ spectra of **BCL-7-66**

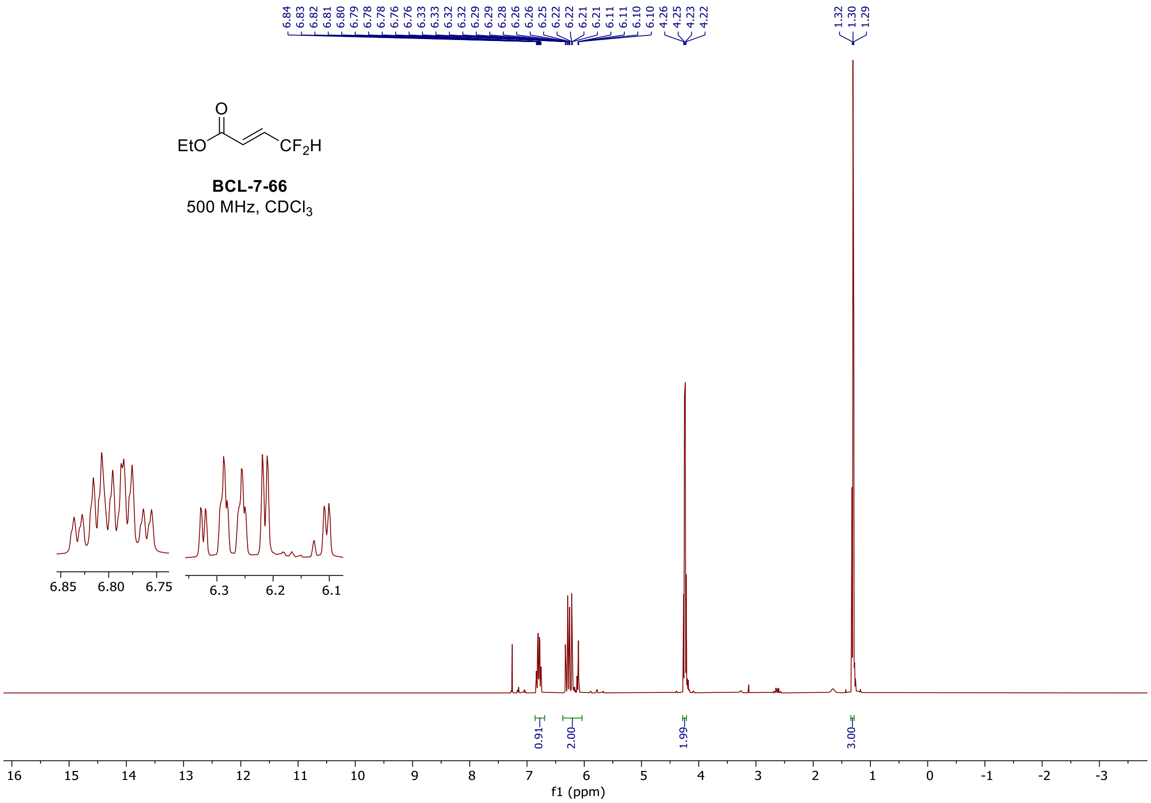

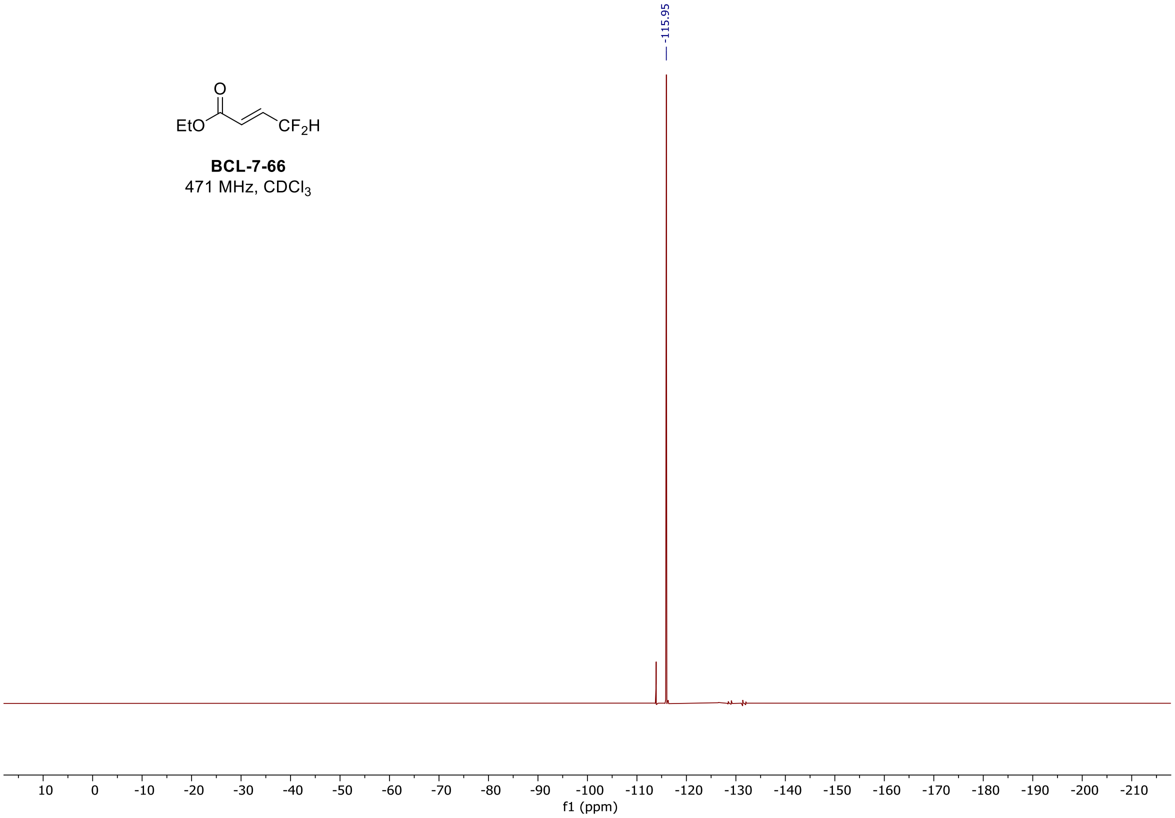

$^{1}H$ and $^{19}F$ spectra of **BCL-7-59**

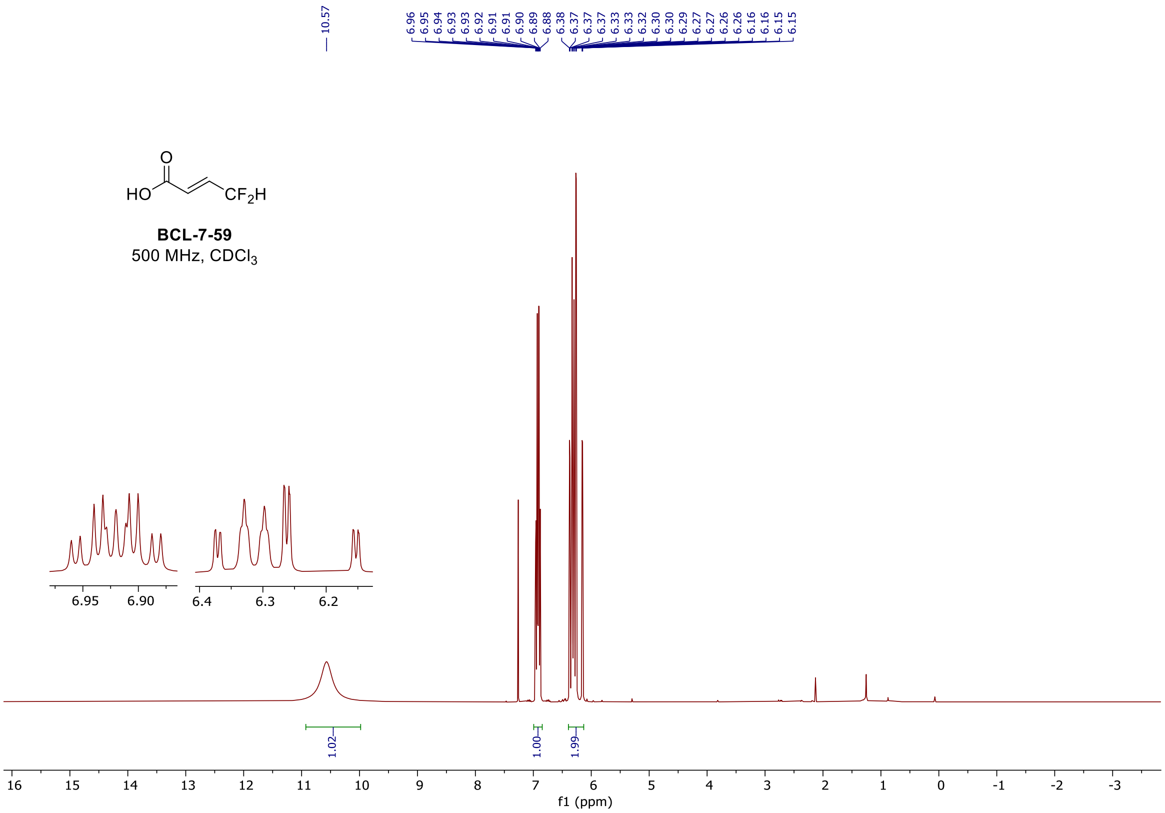

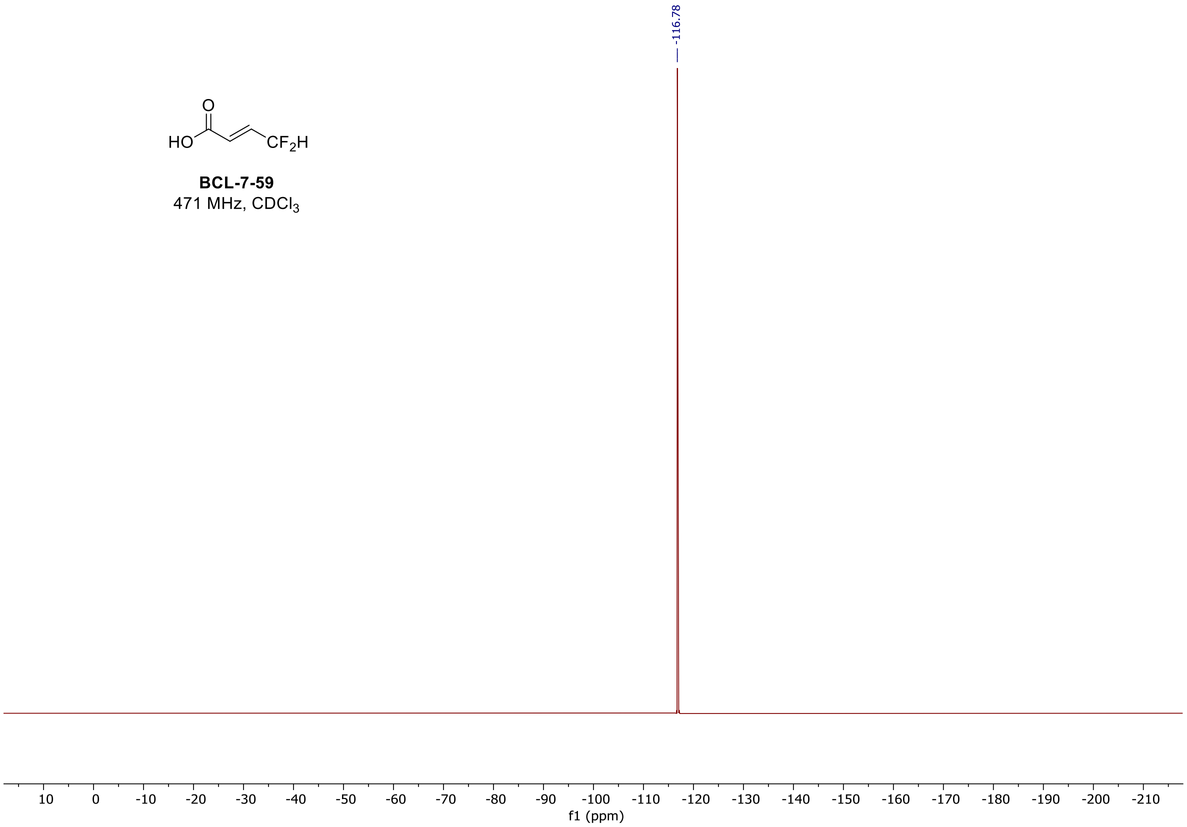

$^{1}H$ and $^{13}C$ spectra of **BCL-1-124**

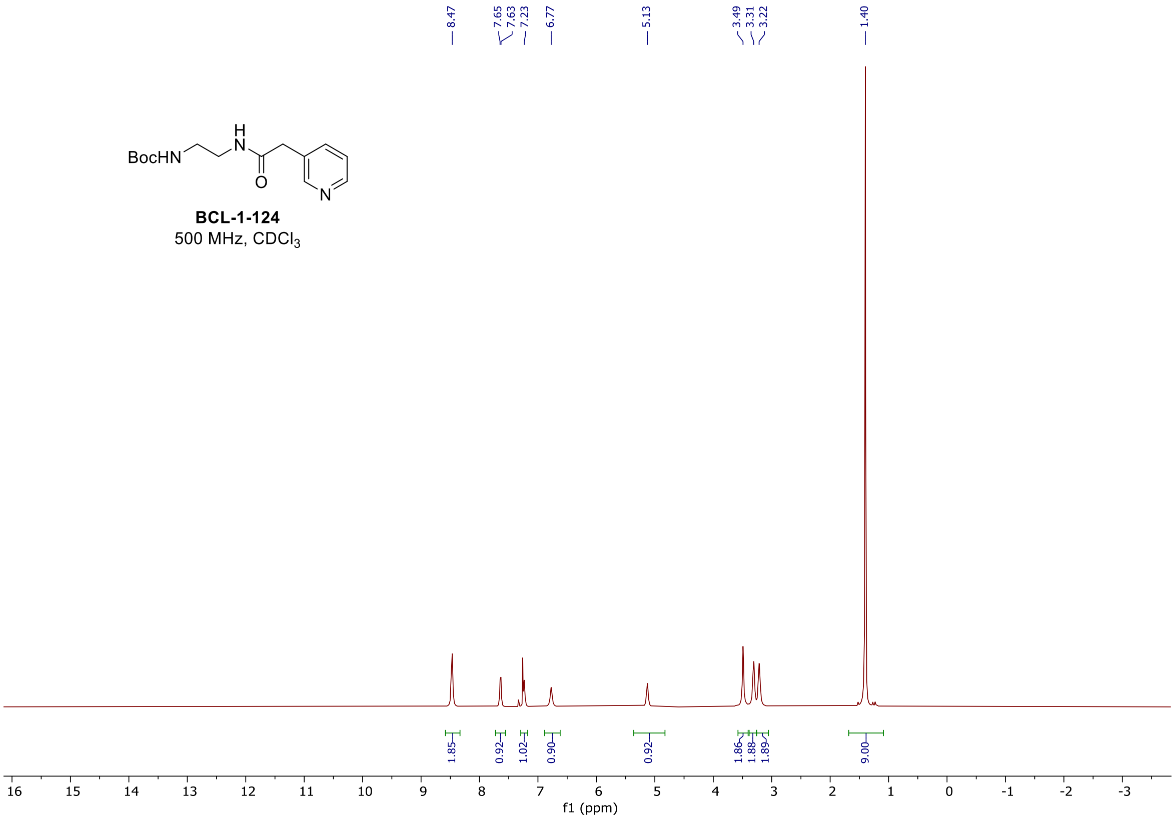

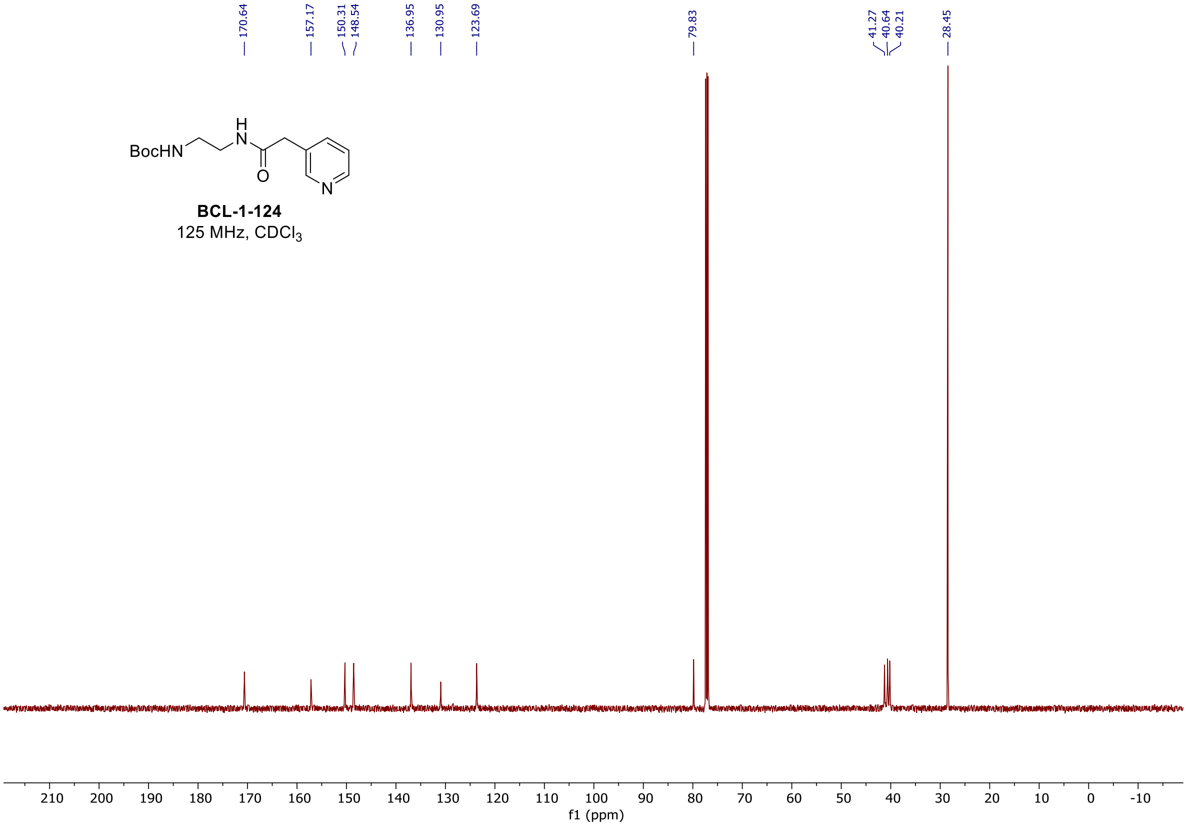

[1]H and [13]C spectra of **BCL-1-133**

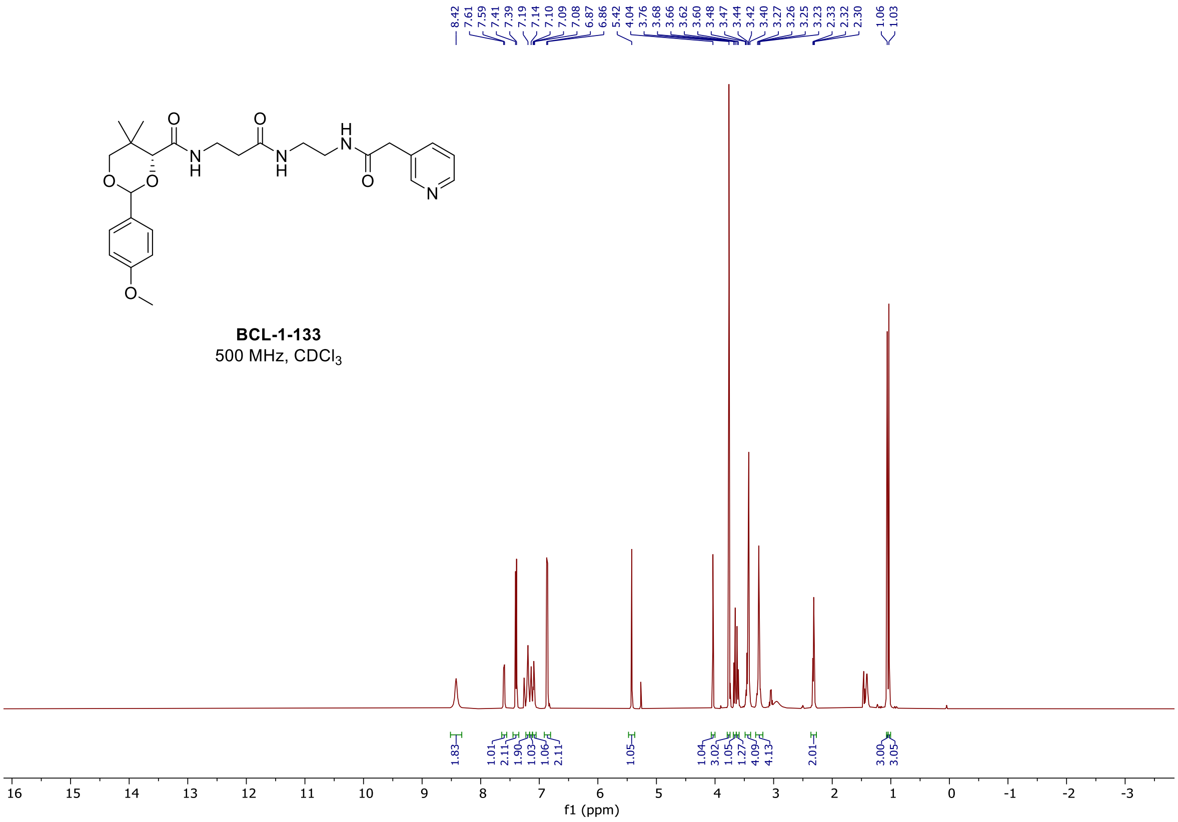

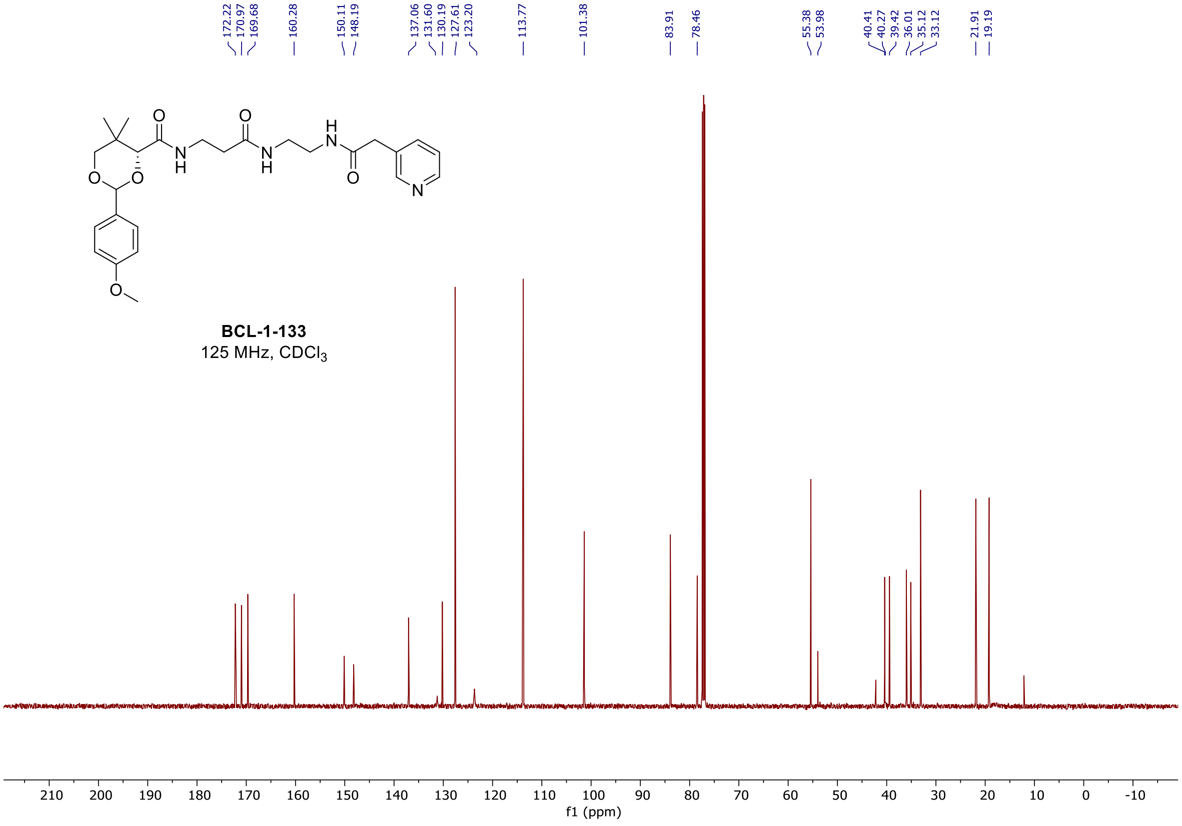

$^{1}H$ and $^{13}C$ spectra of **BCL-1-144**

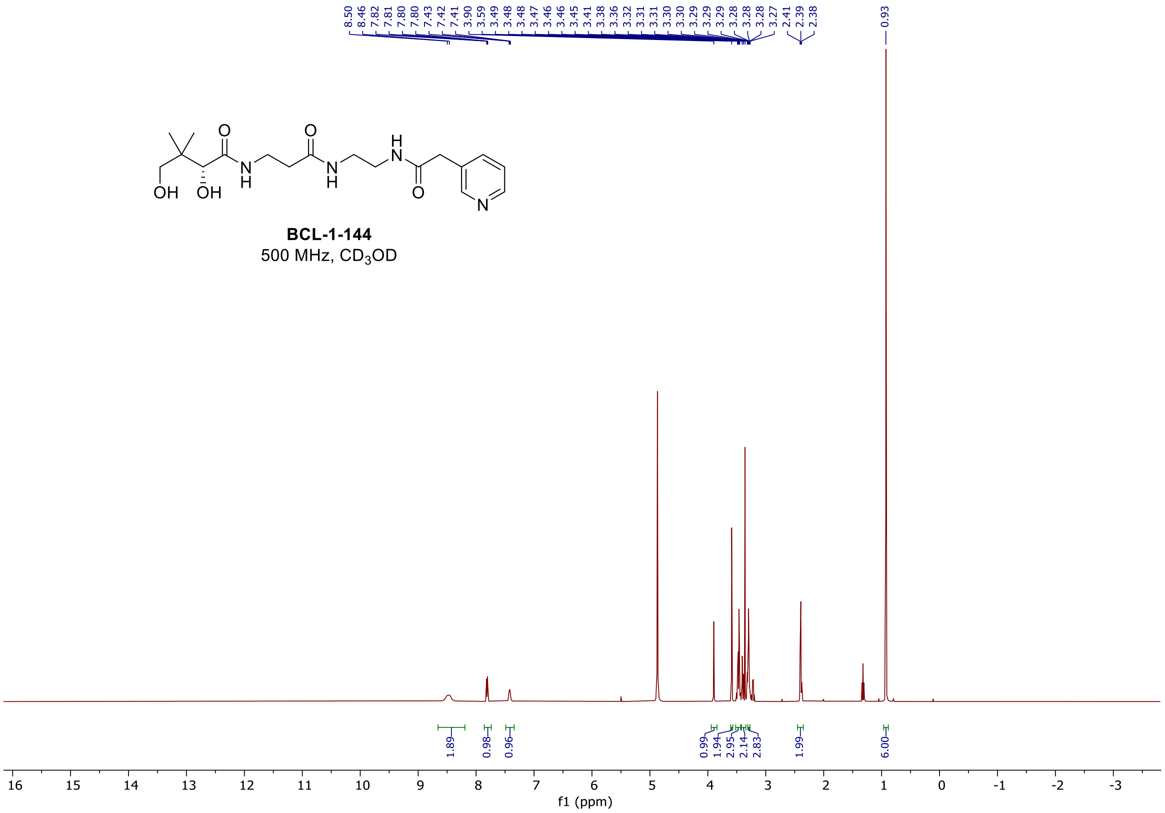

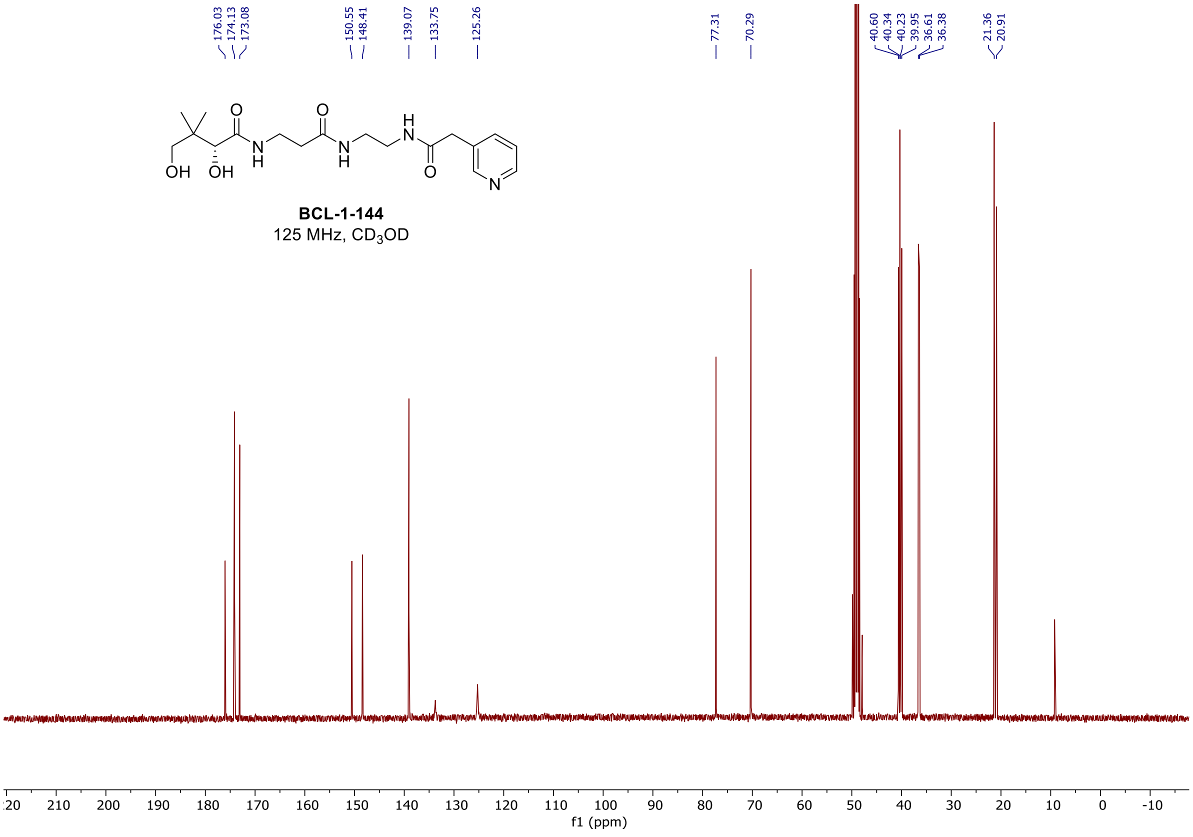

#### [1]H and [13]C spectra of **BCL-2-22**

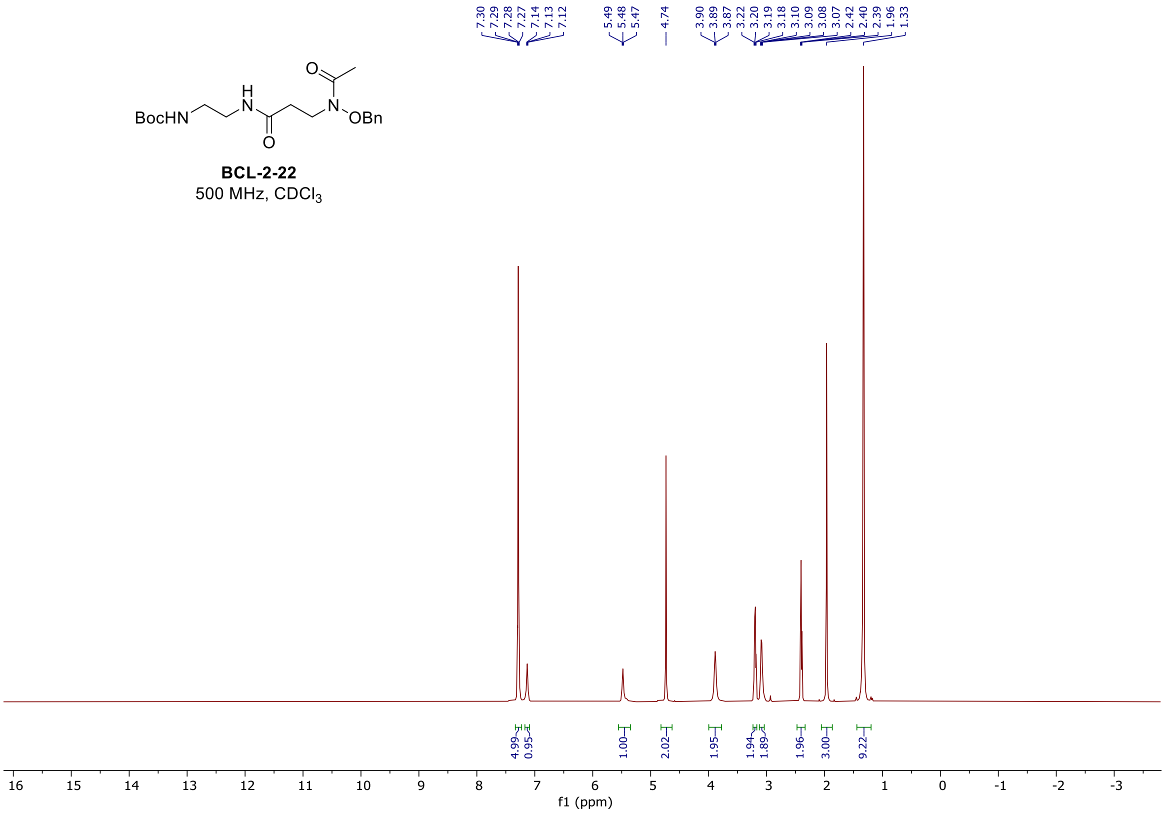

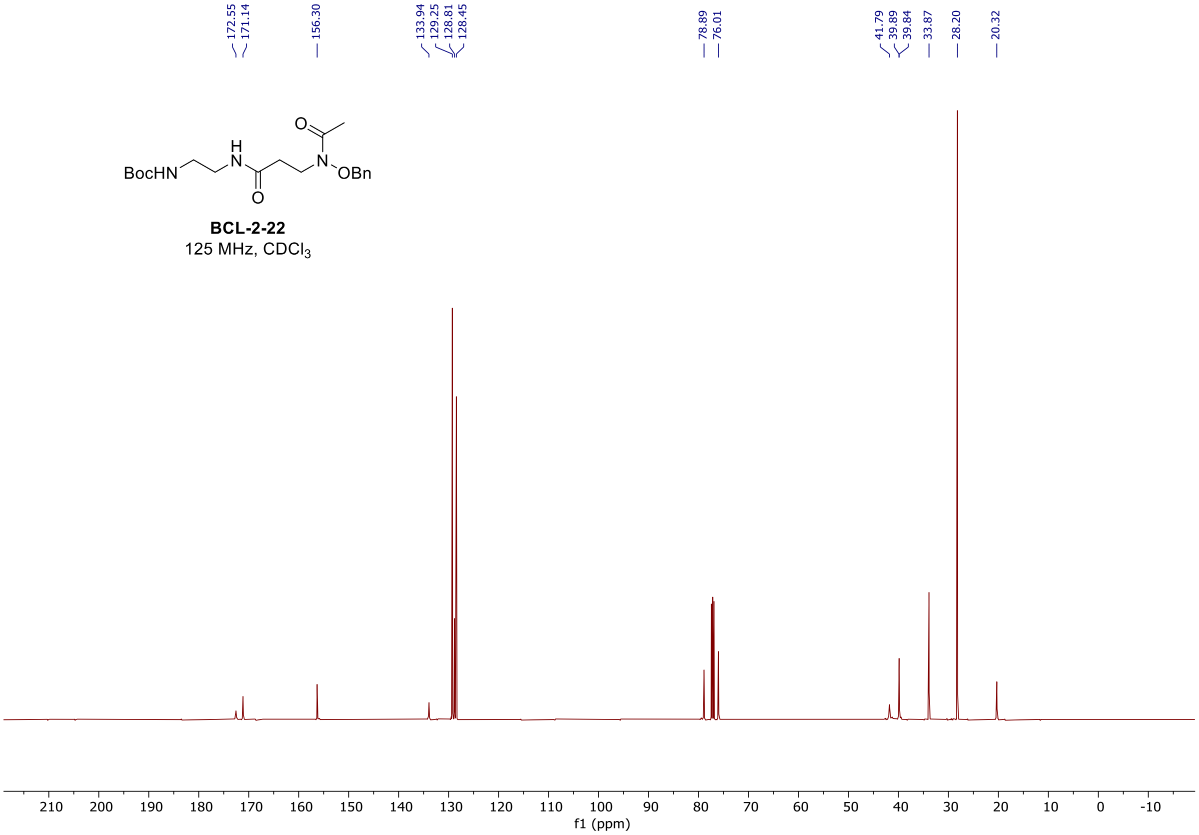

#### $^{1}$H and $^{13}$C spectra of **BCL-2-36**

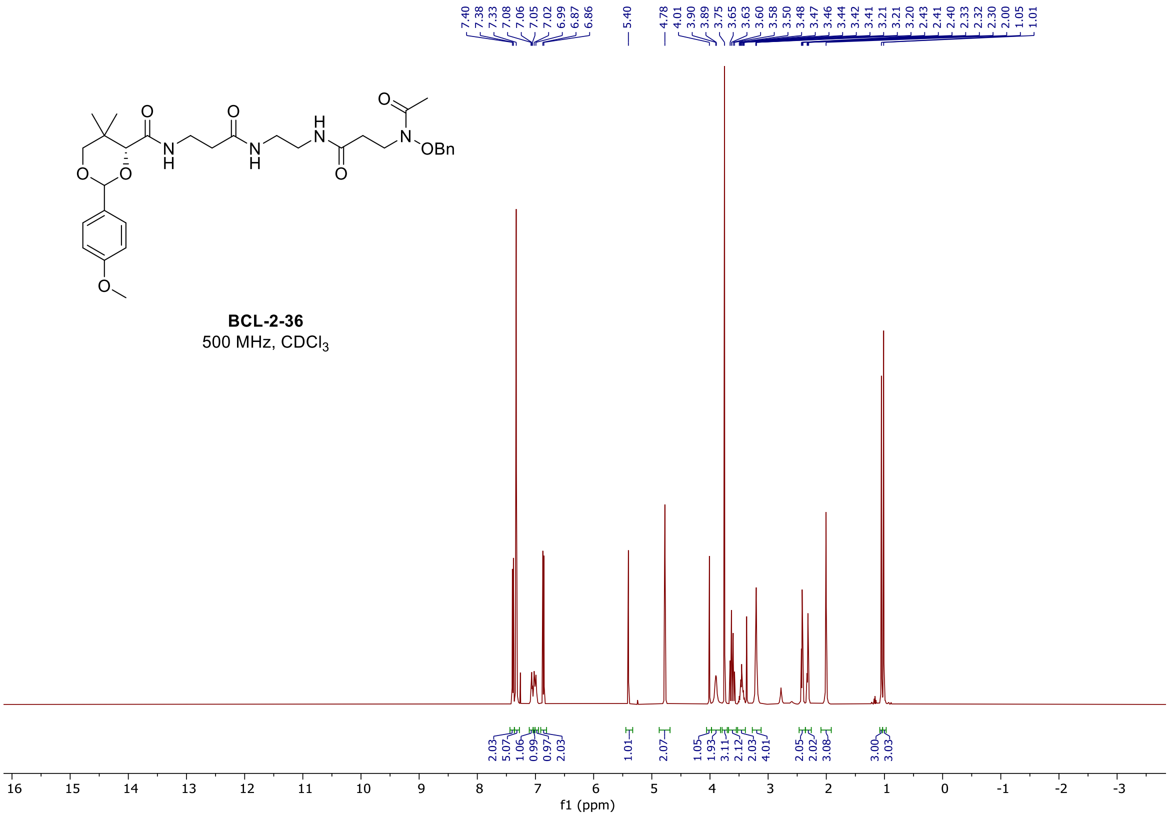

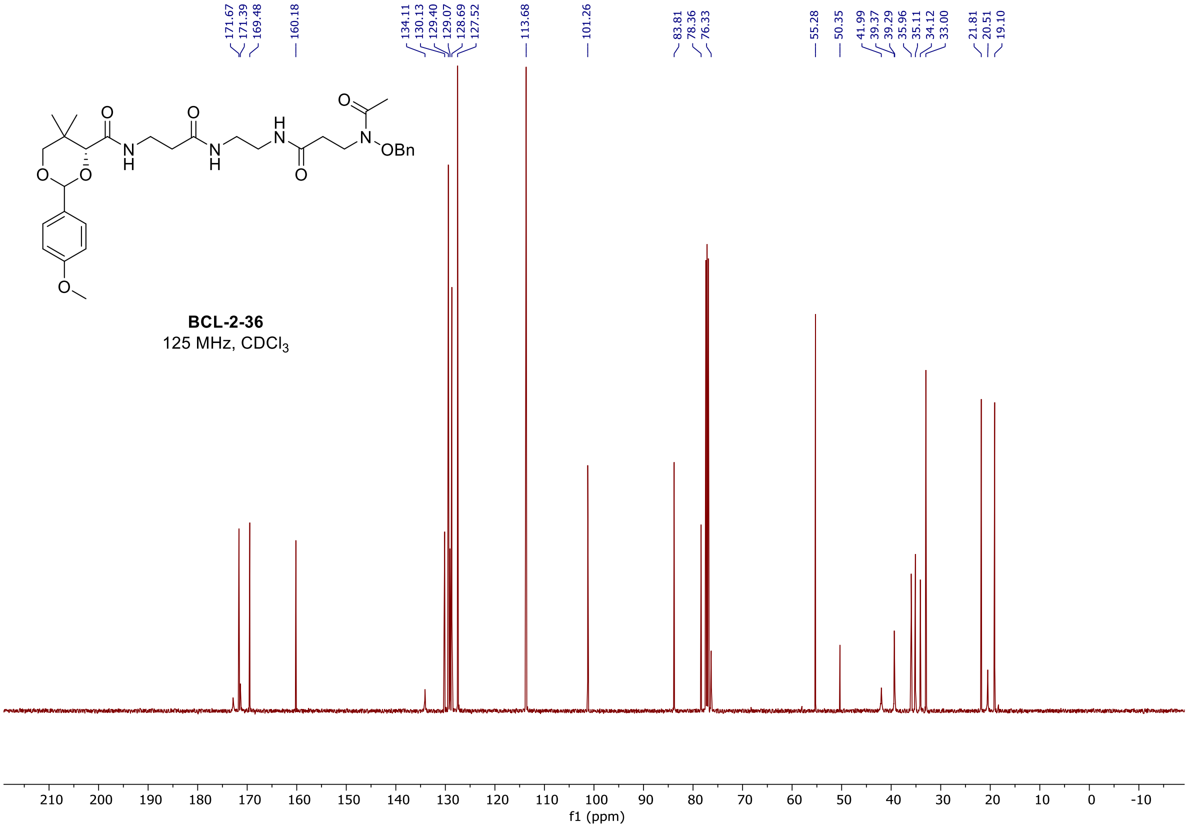

<sup></sup>

#### $^{1}$H and $^{13}$C spectra of **BCL-2-56**

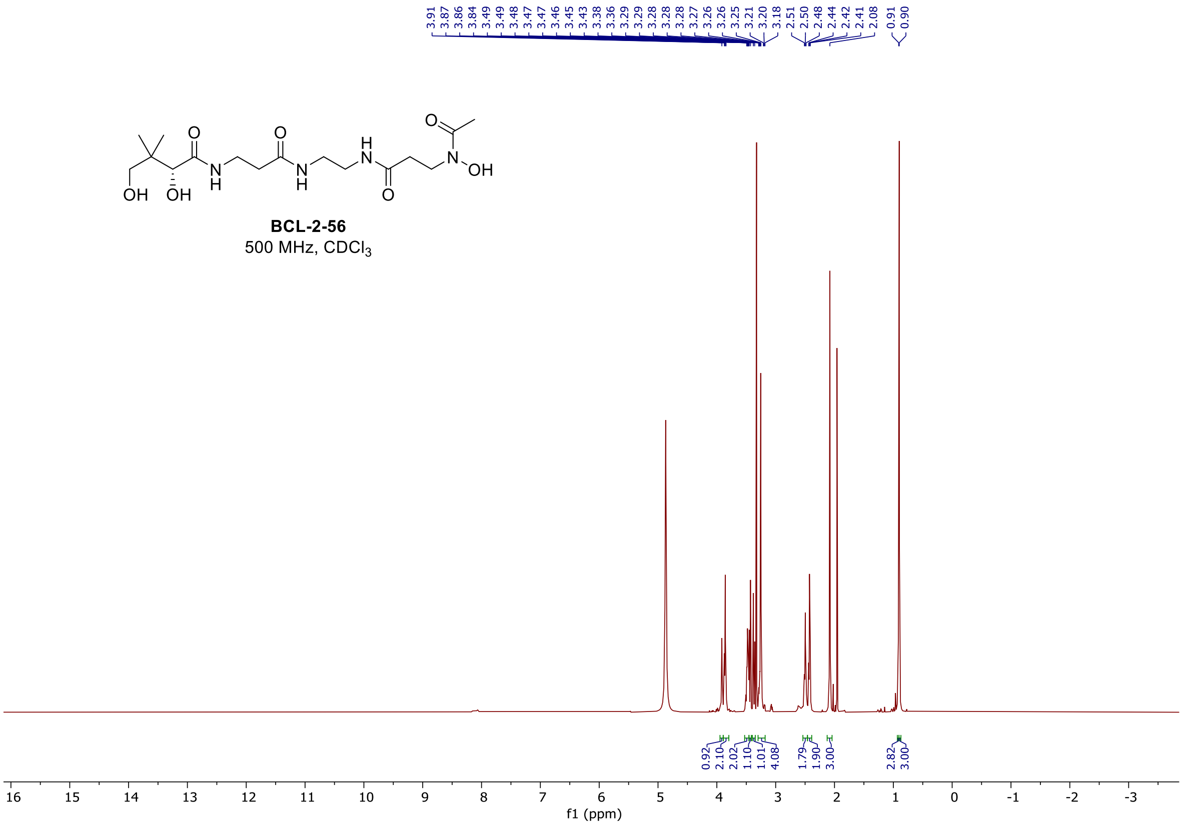

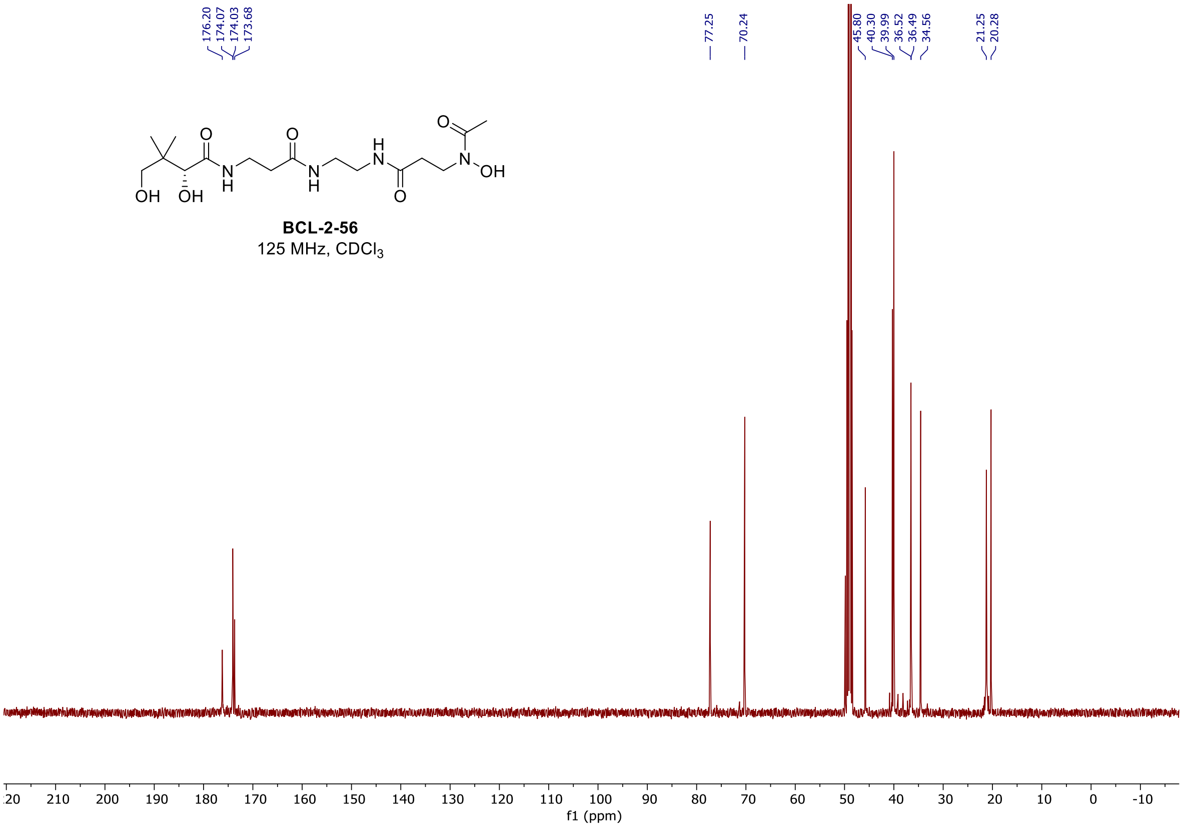

$^1$H and $^{13}$C spectra of **BCL-4-4**

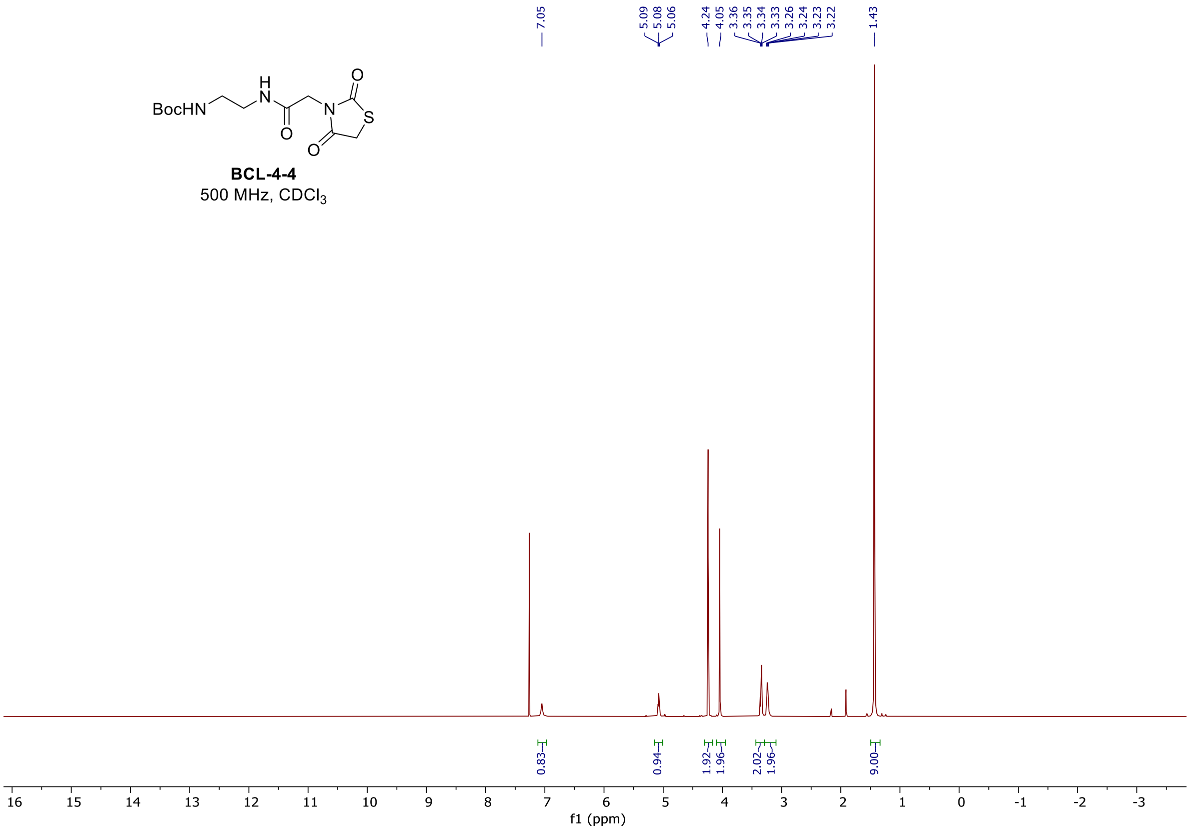

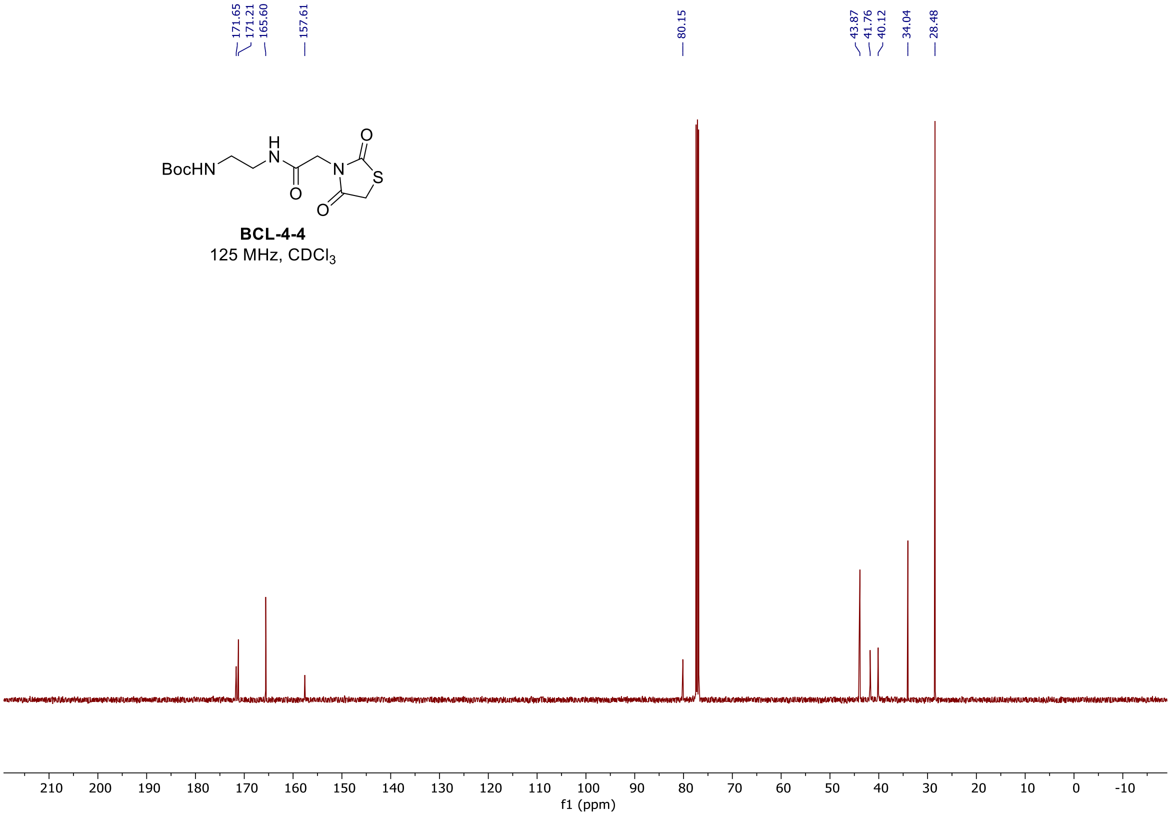

$^1H$ and $^{13}C$ spectra of **BCL-4-6**

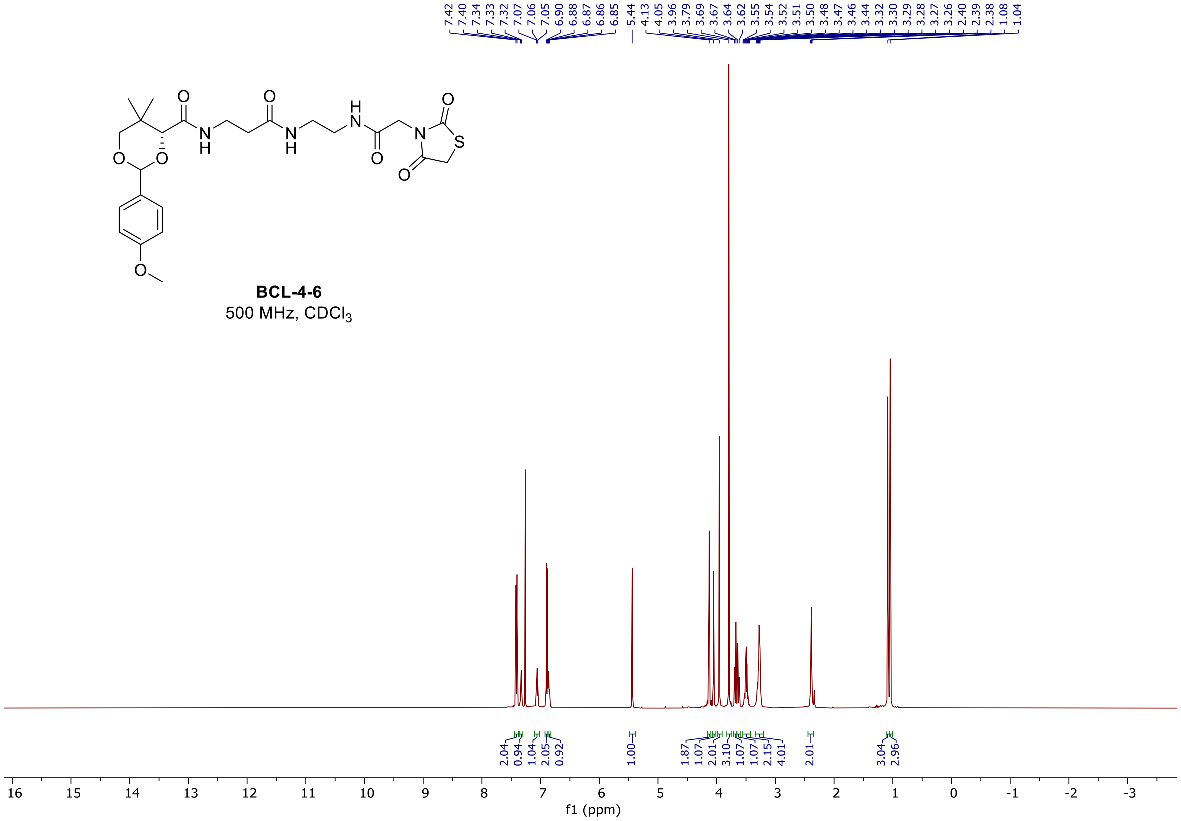

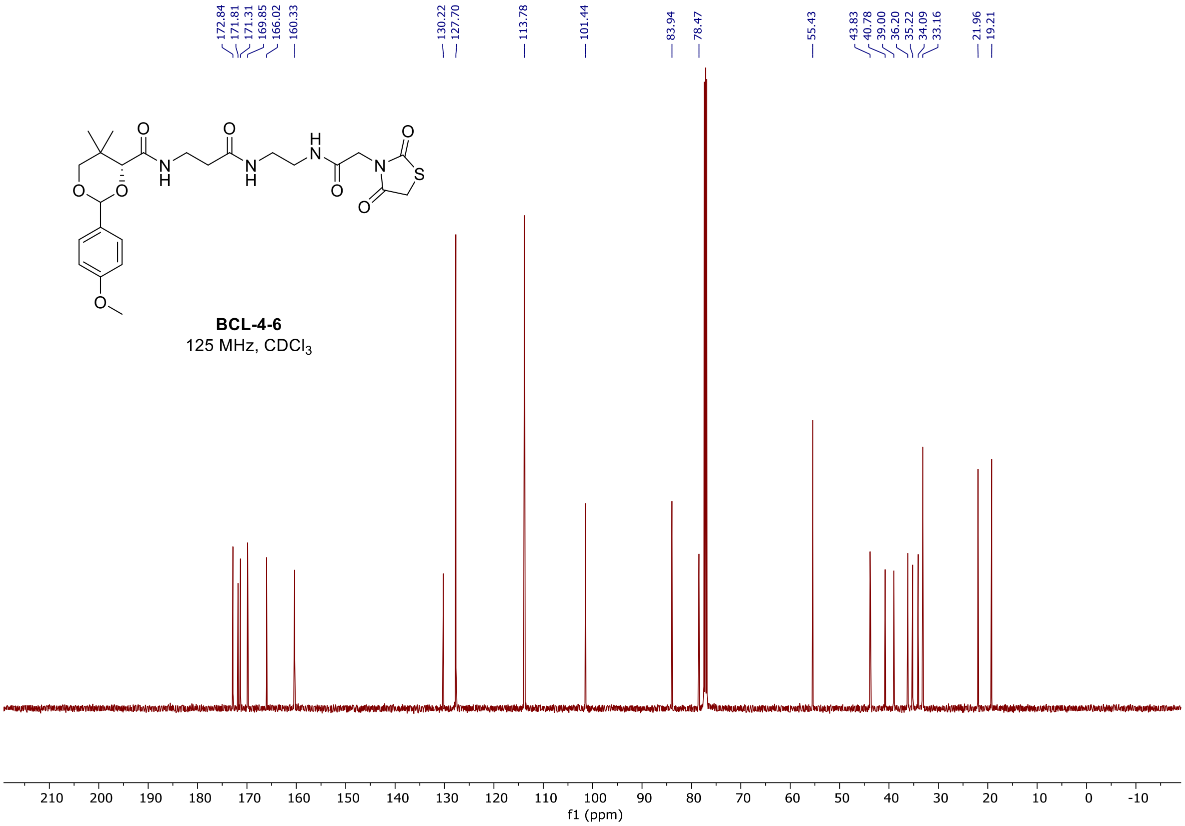

### $^{1}H$ and $^{13}C$ spectra of **BCL-4-11**

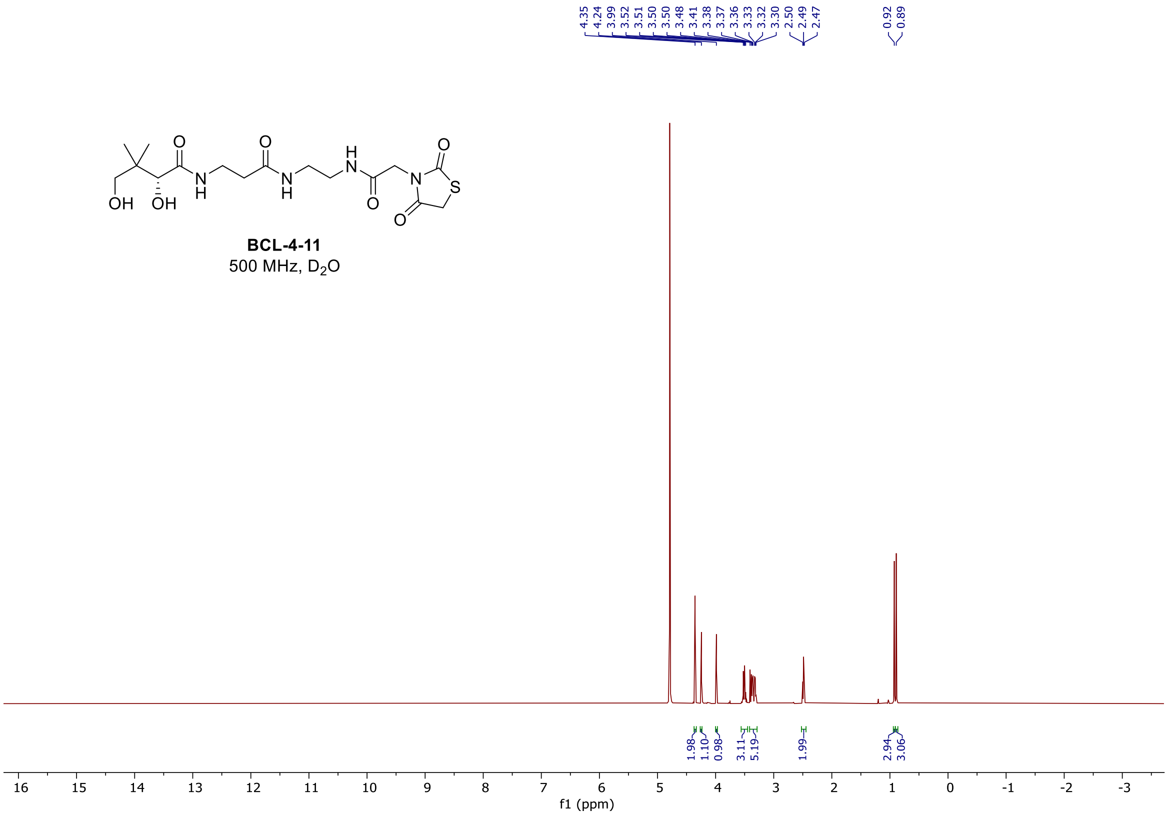

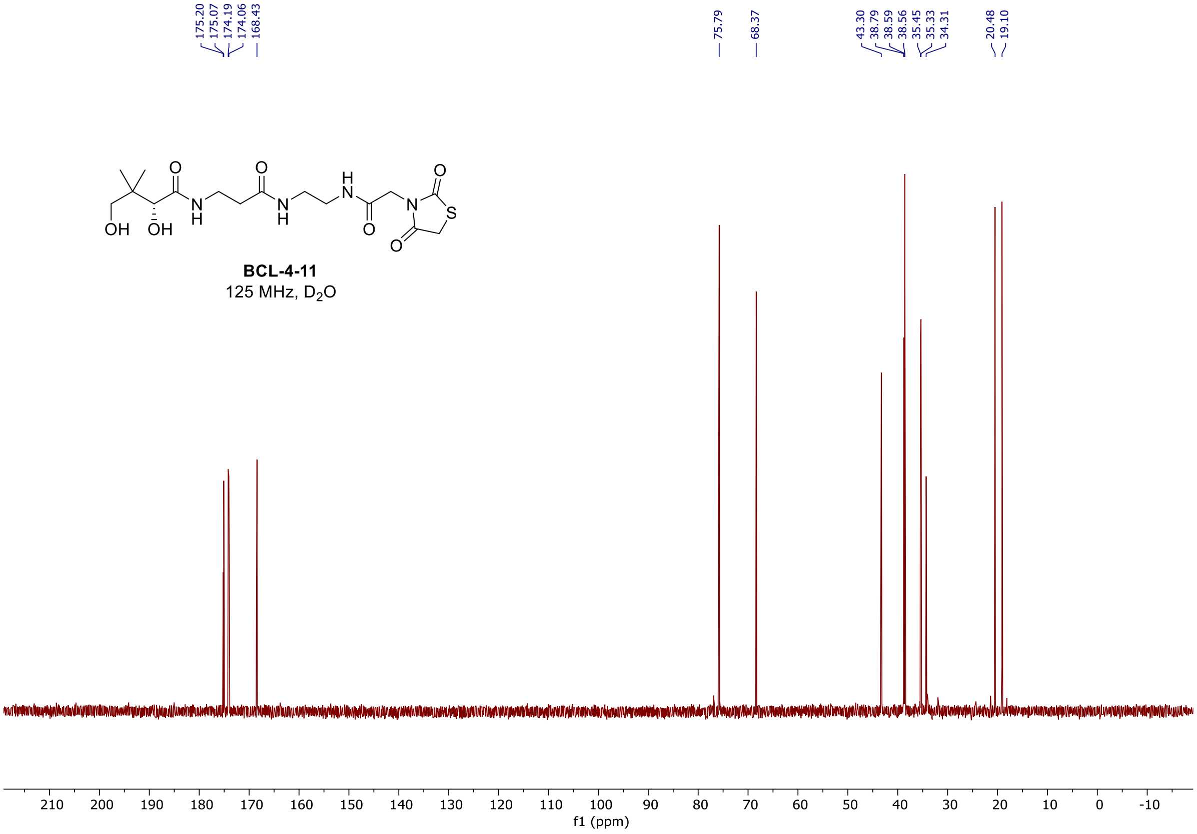

#### $^1$H, $^{19}$F, and $^{13}$C spectra of **BCL-4-43**

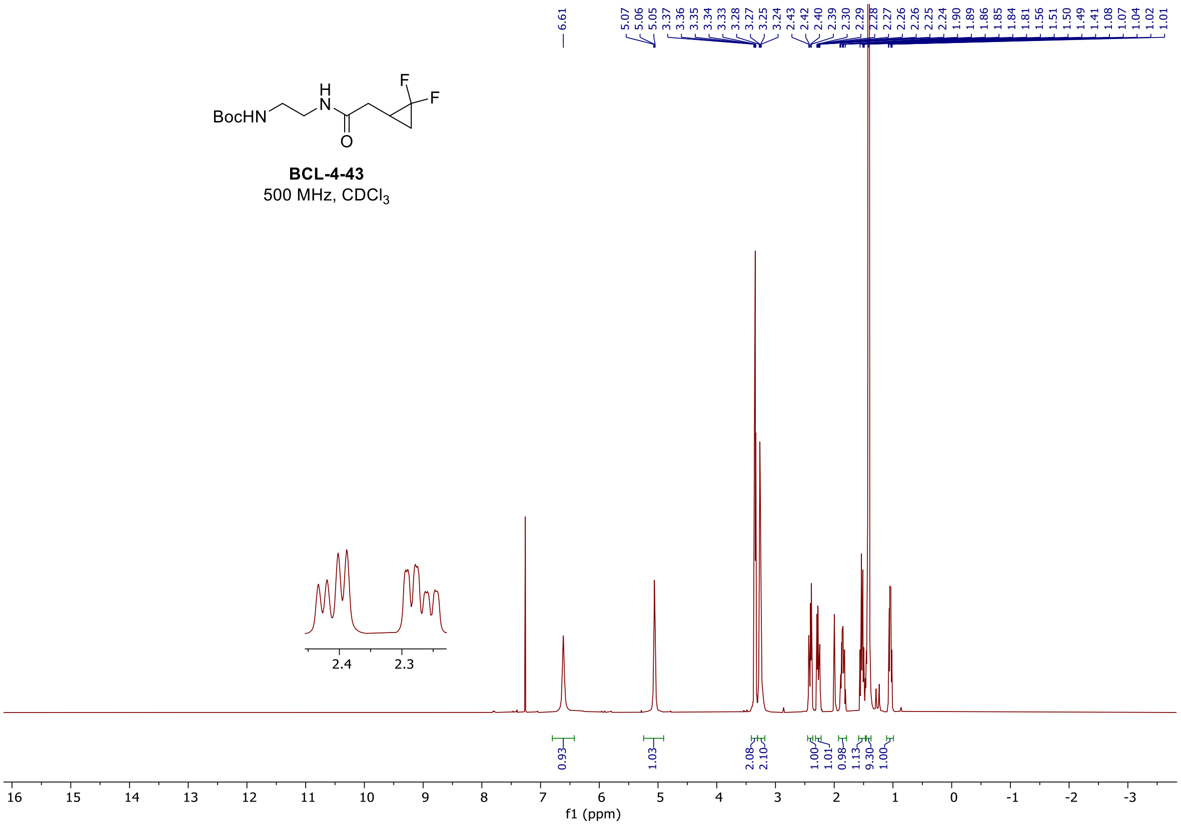

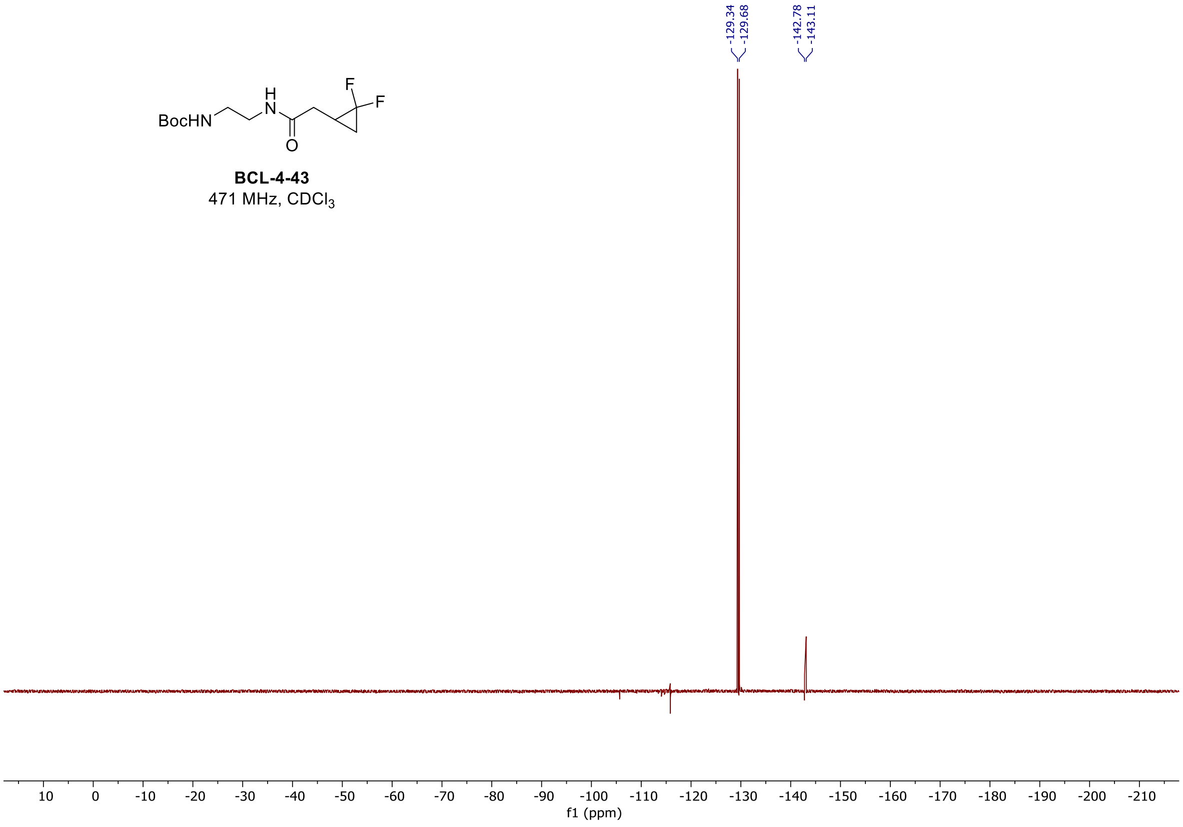

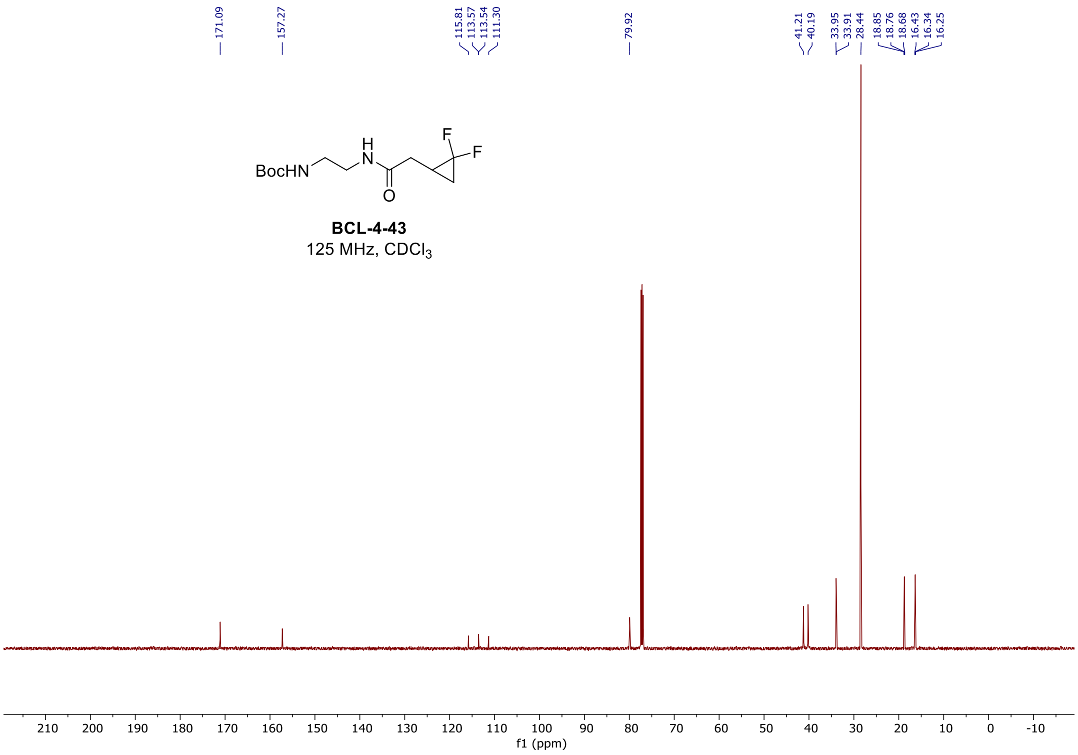

#### $^{1}H$, $^{19}F$, and $^{13}C$ spectra of **BCL-4-47**

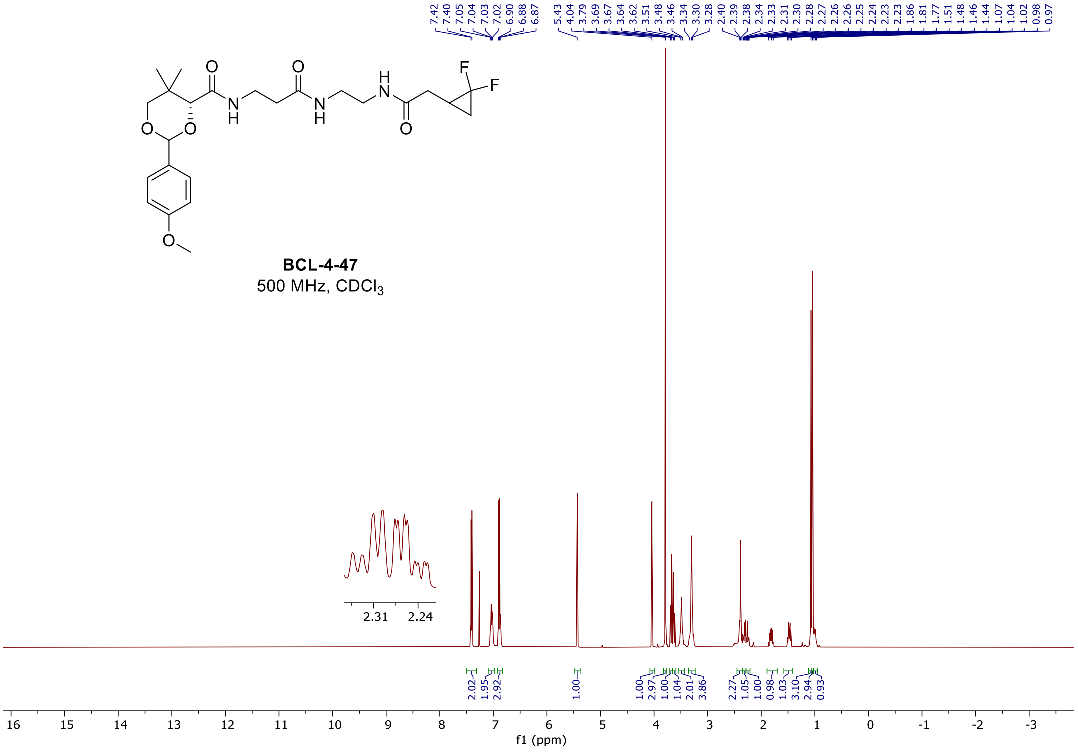

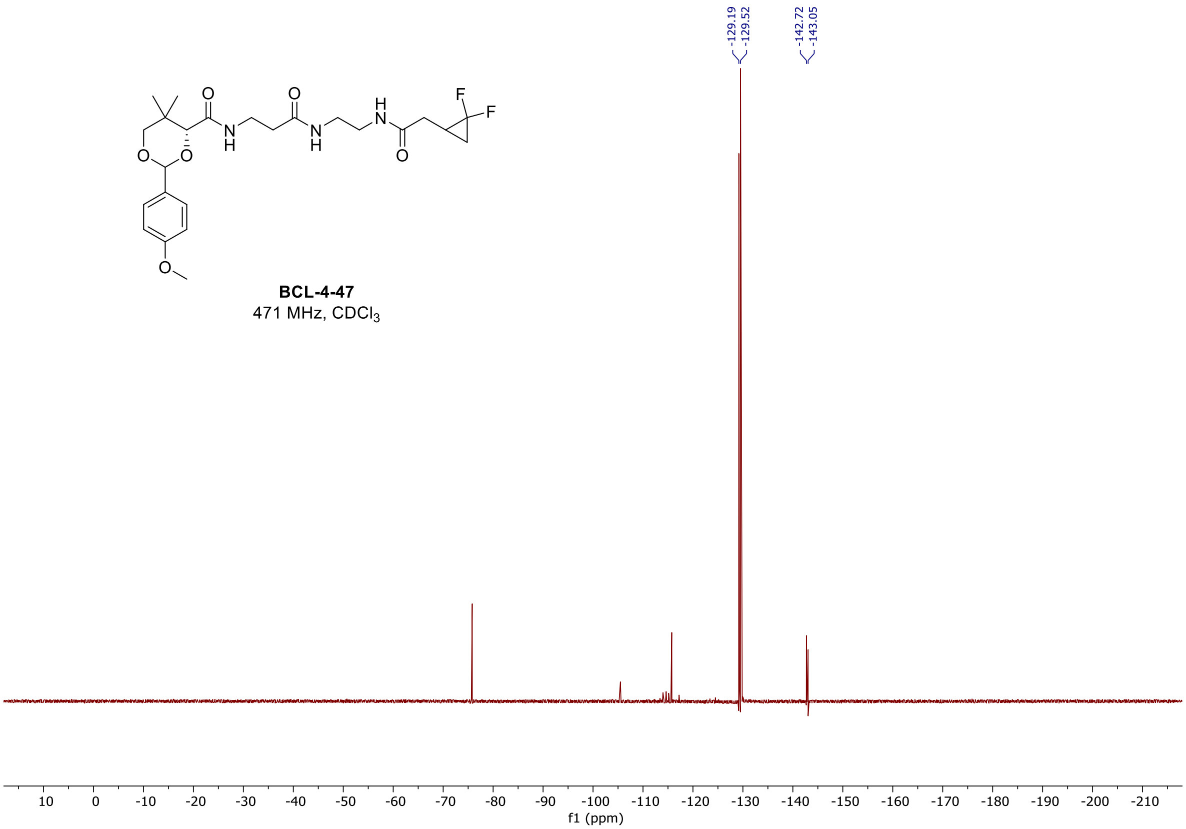

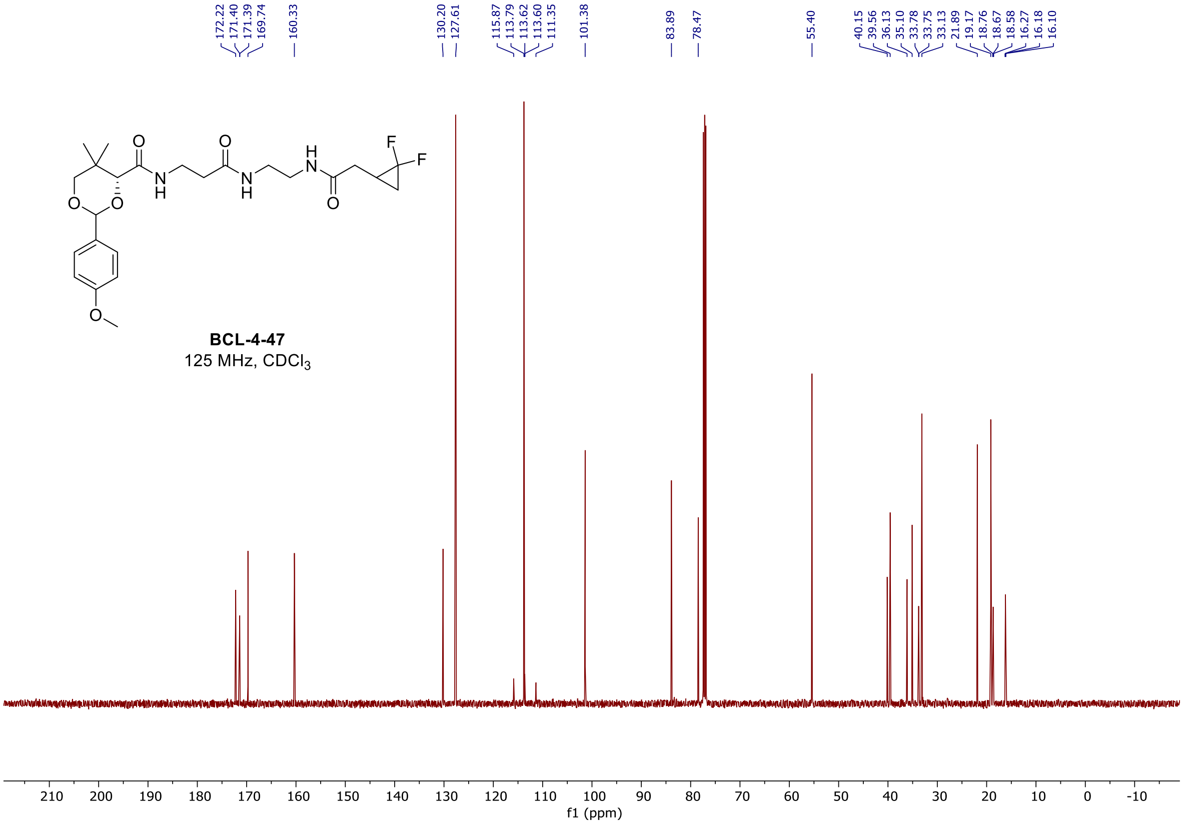

#### $^{1}$H, $^{19}$F, and $^{13}$C spectra of **BCL-4-52**

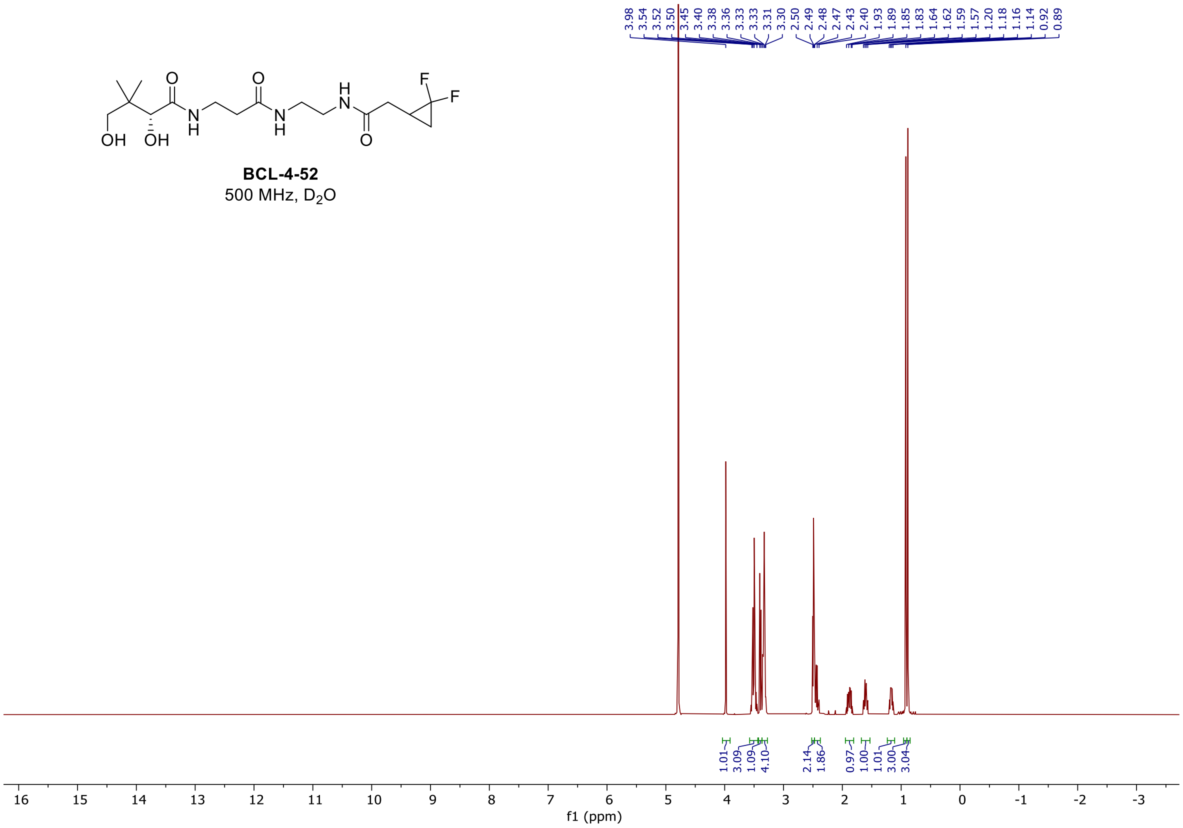
BCL-4-52
500 MHz, D2O
f1 (ppm)

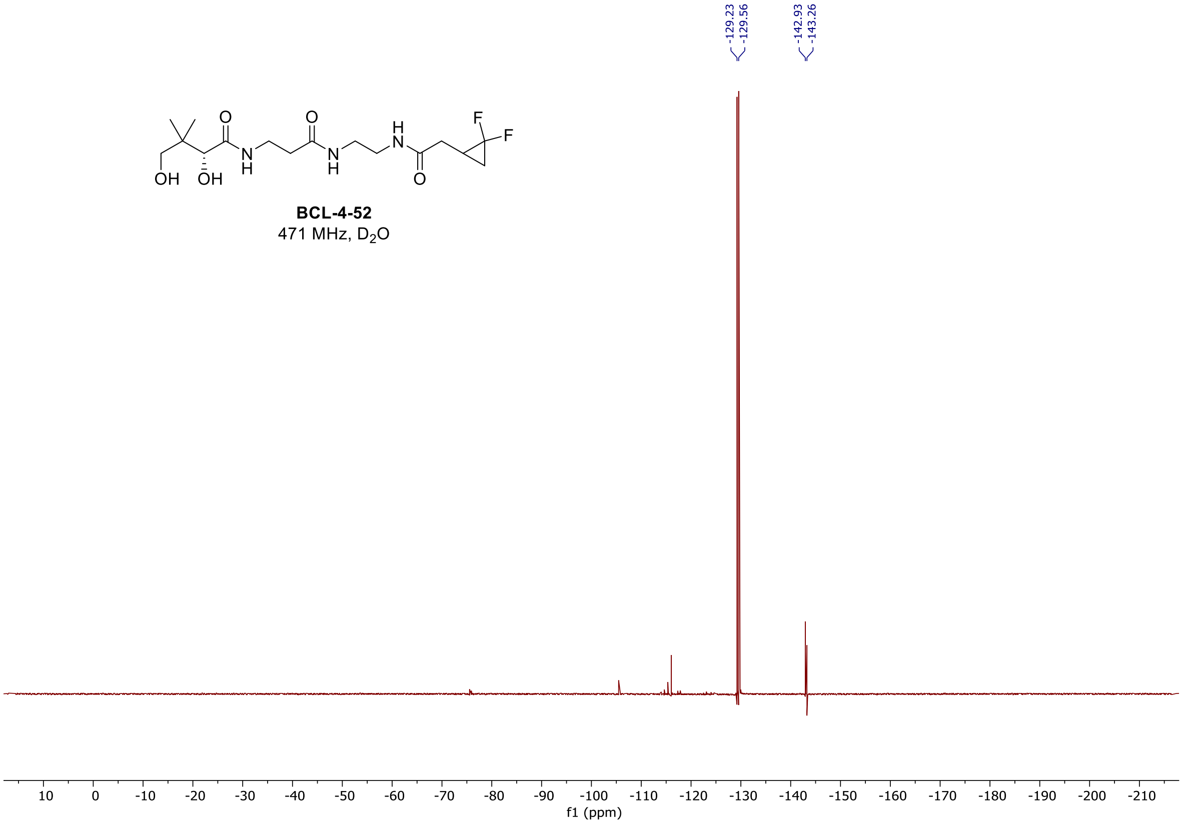
BCL-4-52
471 MHz, D2O
f1 (ppm)

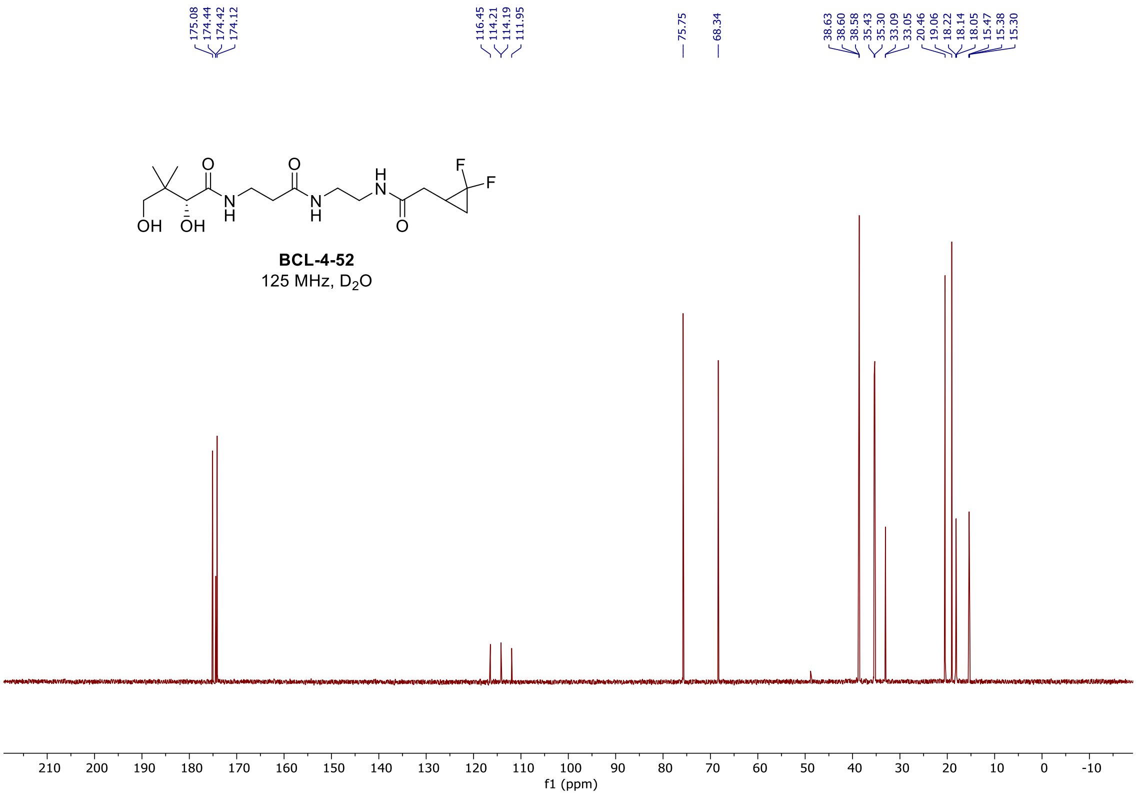

#### $^1$H, and $^{13}$C spectra of **BCL-4-51**

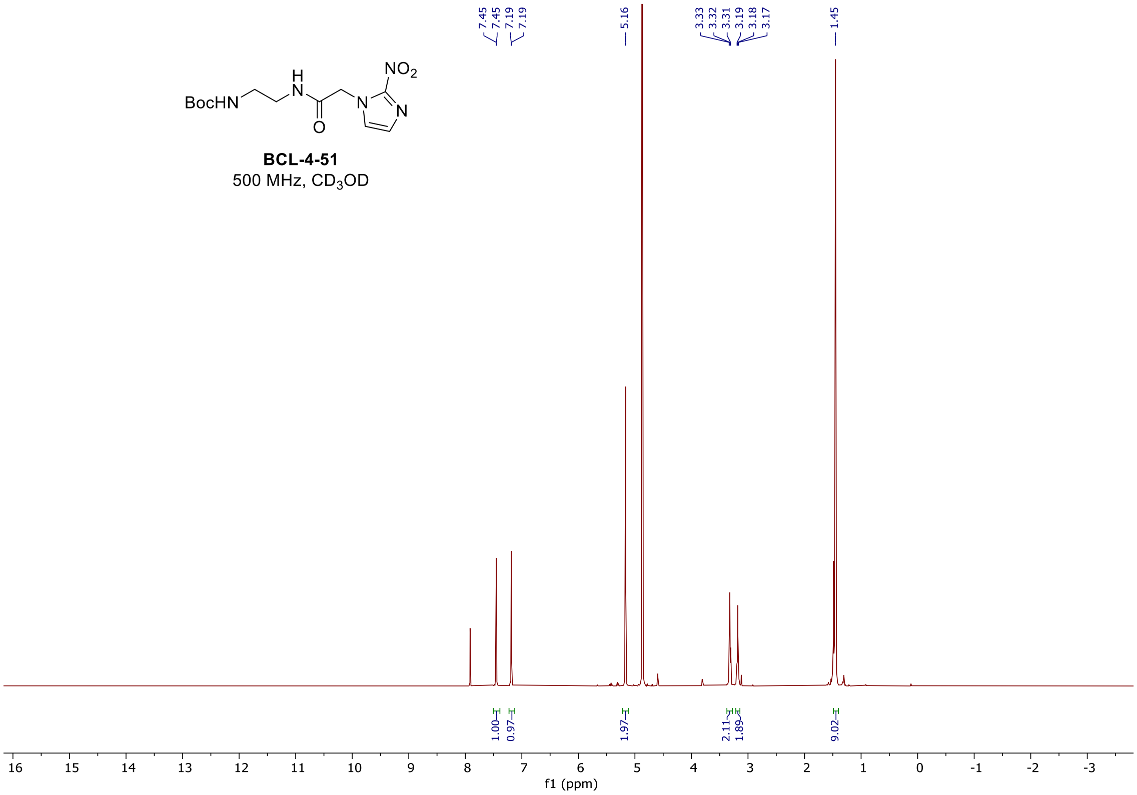

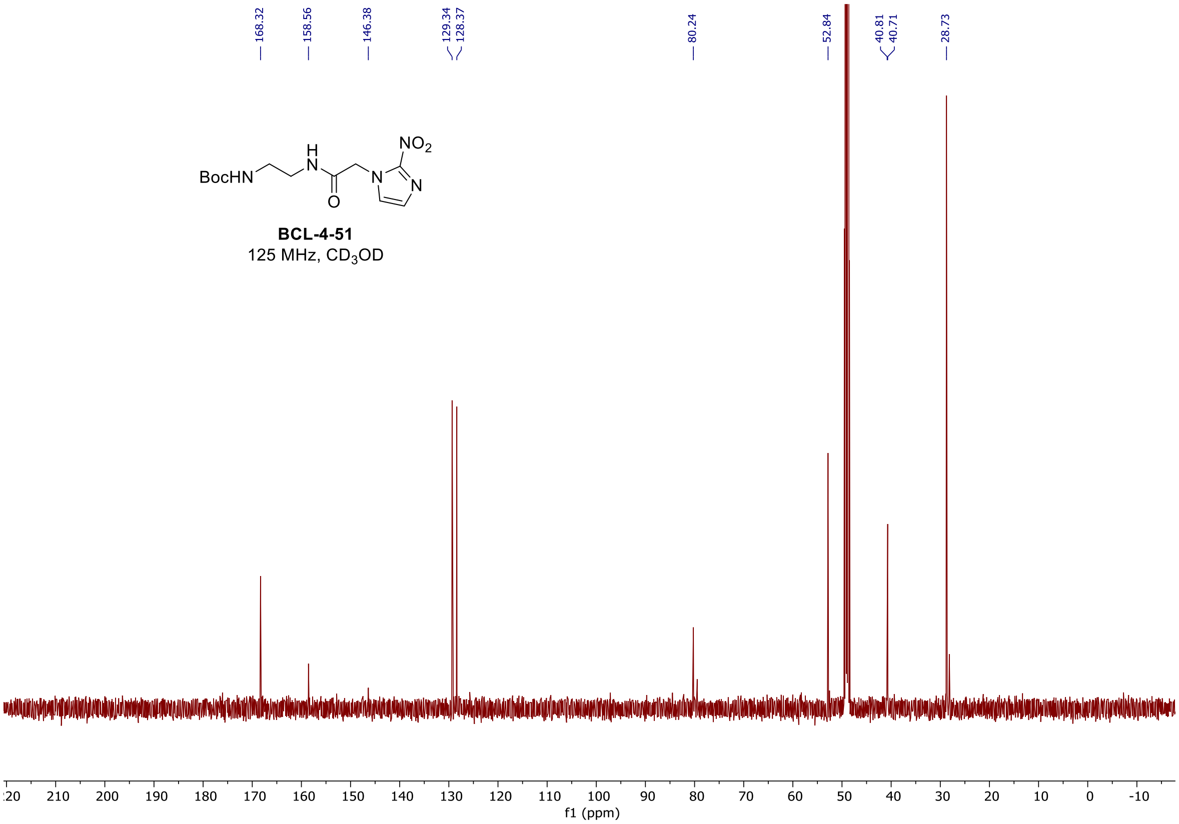

#### [1]H, and [13]C spectra of **BCL-4-61**

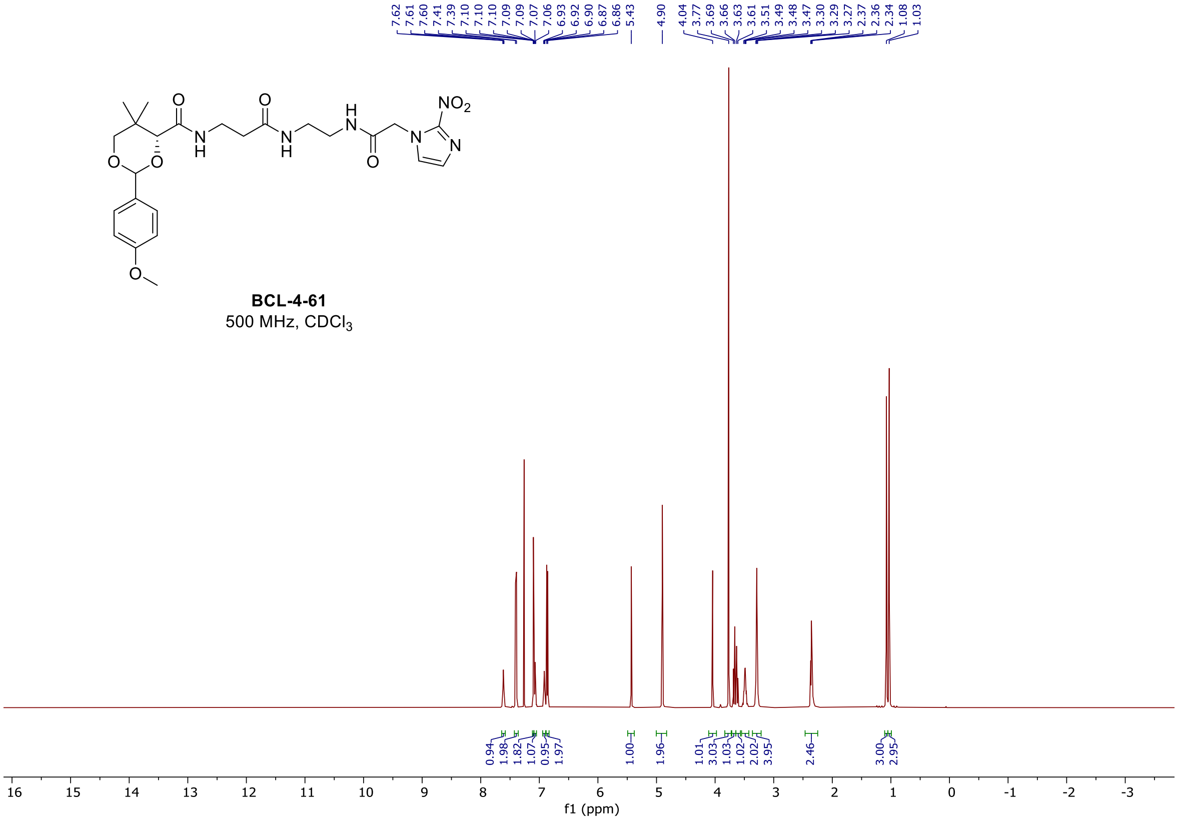

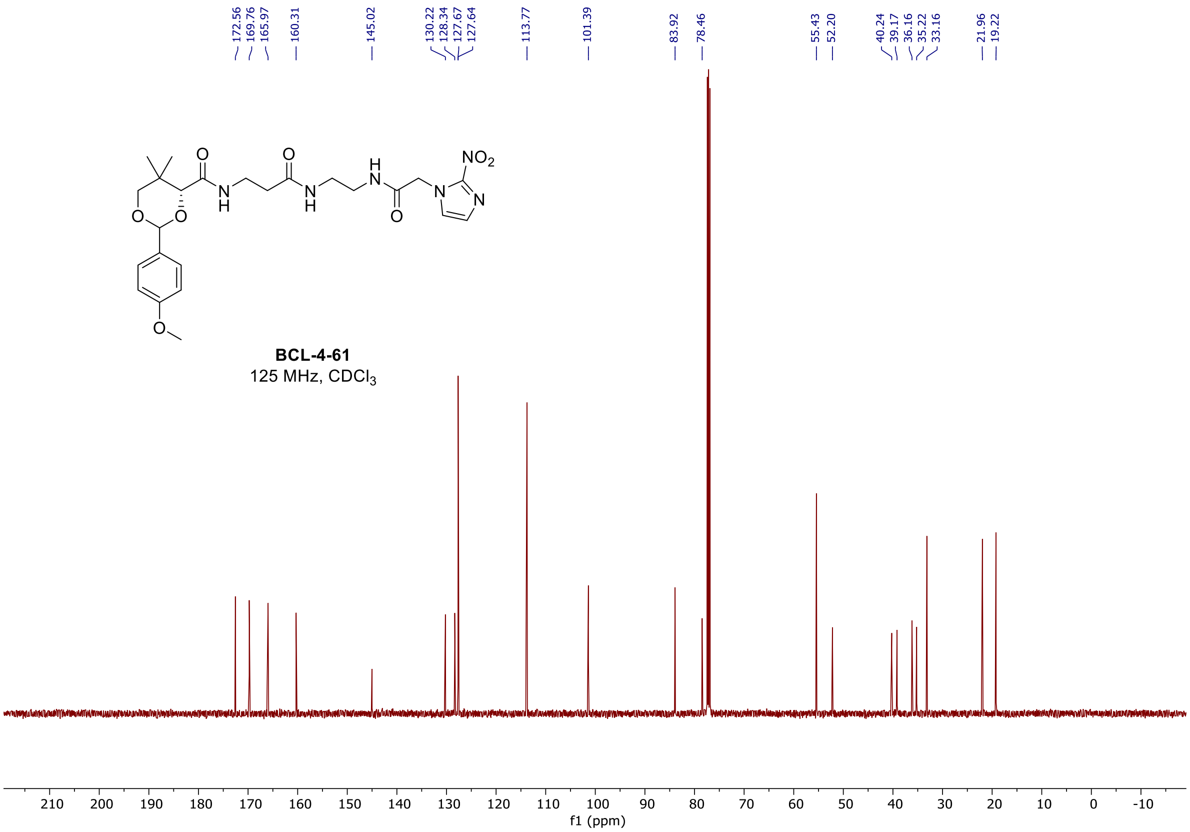
172.56
169.76
165.97
160.31
145.02
130.22
128.34
127.67
127.64
113.77
101.39
83.92
78.46
55.43
52.20
40.24
39.17
36.16
35.22
33.16
21.96
19.22
BCL-4-61
125 MHz, CDCl3
210 200 190 180 170 160 150 140 130 120 110 100 90 80 70 60 50 40 30 20 10 0 -10
f1 (ppm)

$^{1}H$, and $^{13}C$ spectra of **BCL-4-66**

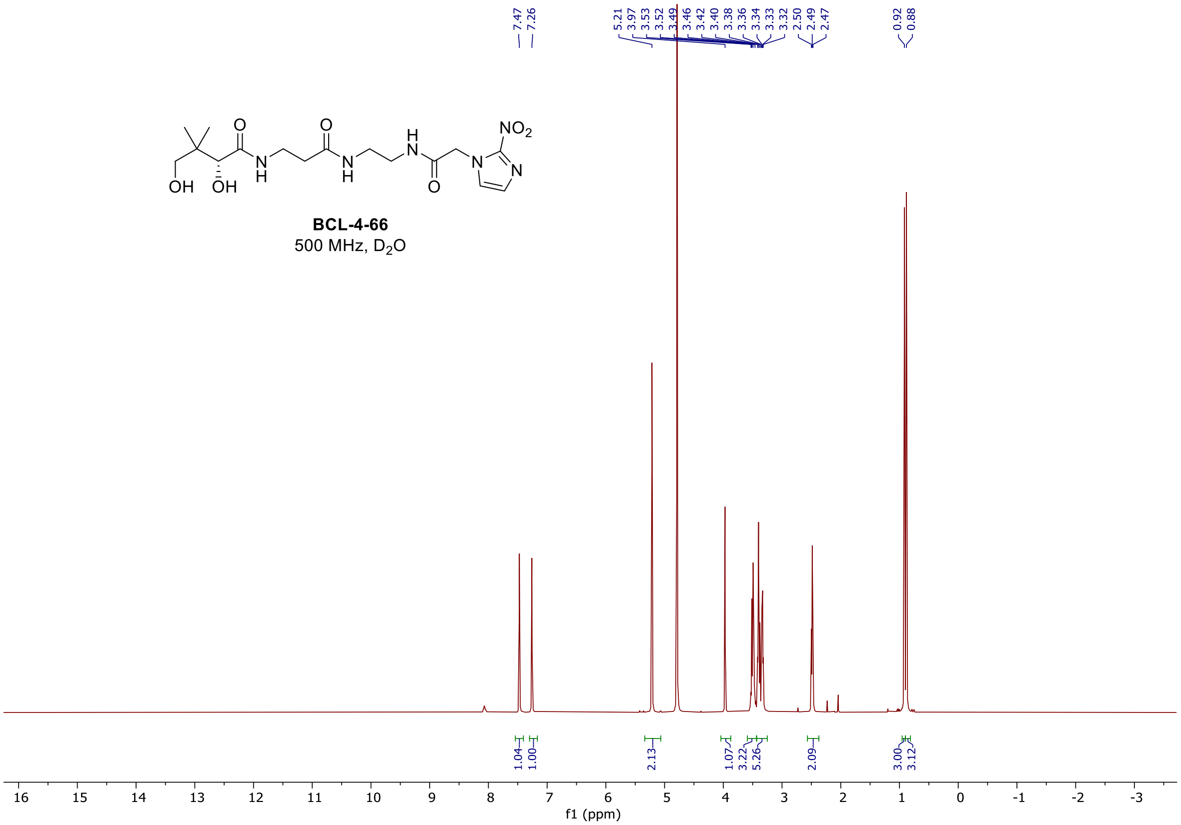

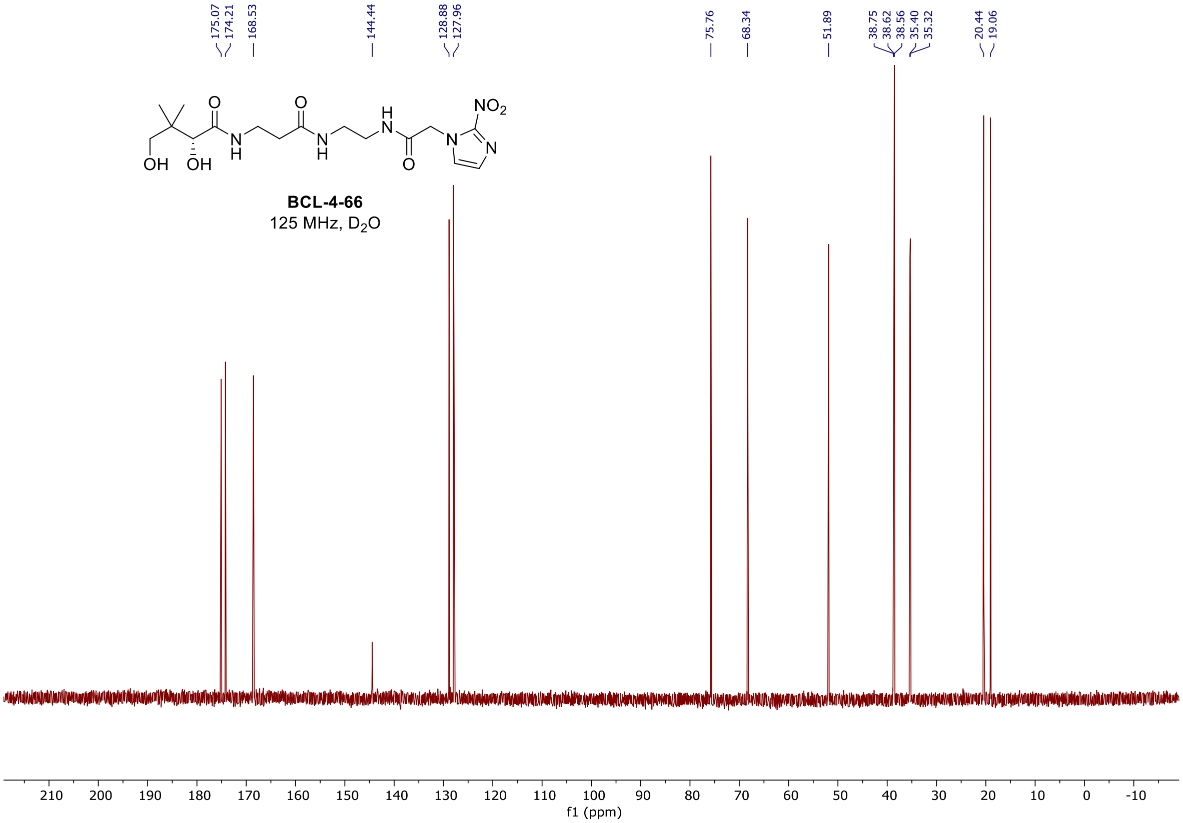

#### [1]H, and [13]C spectra of **BCL-4-72**

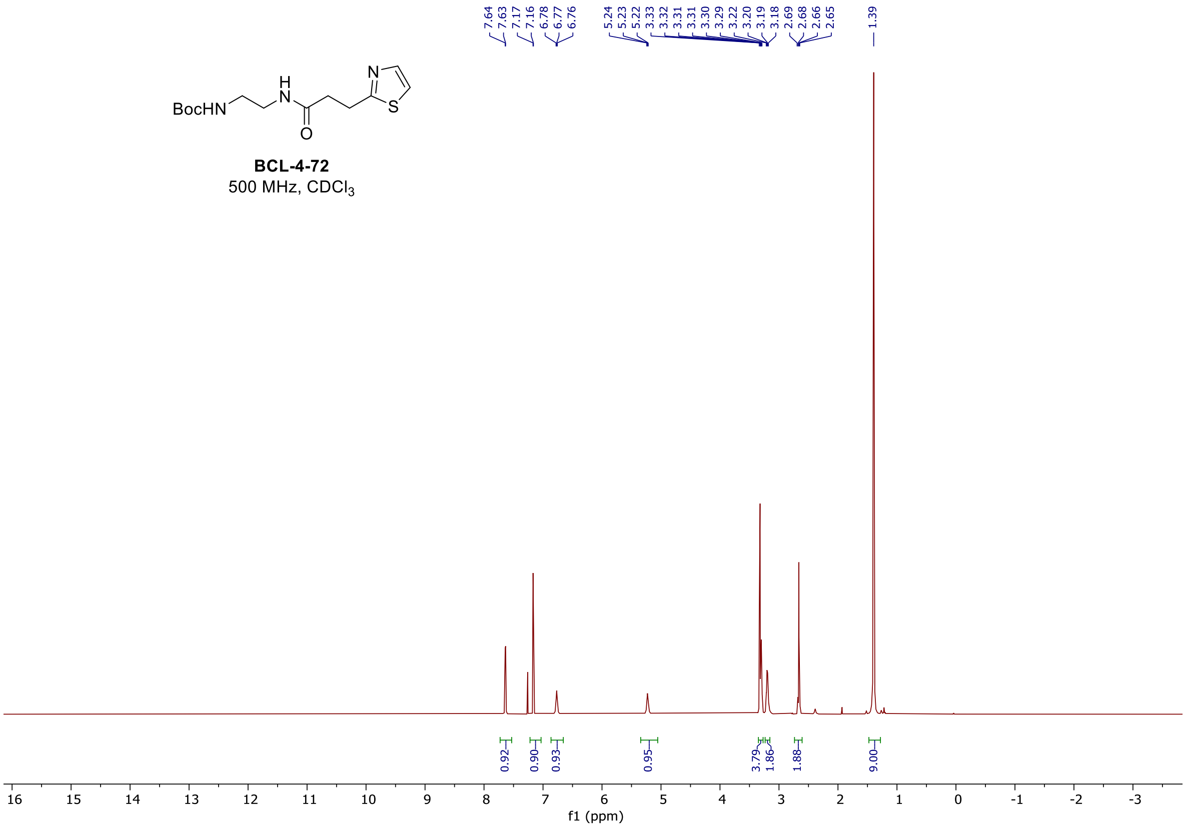

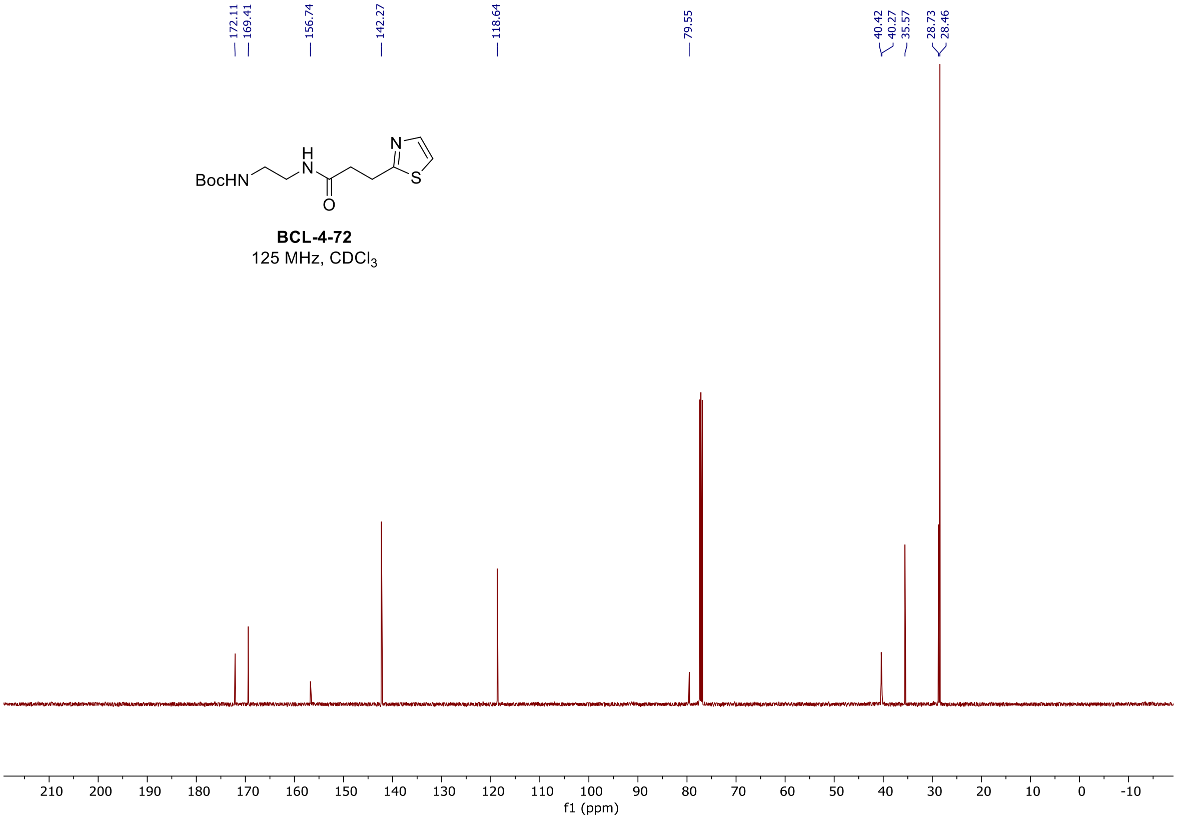

$^{1}H$, and $^{13}C$ spectra of **BCL-4-76**

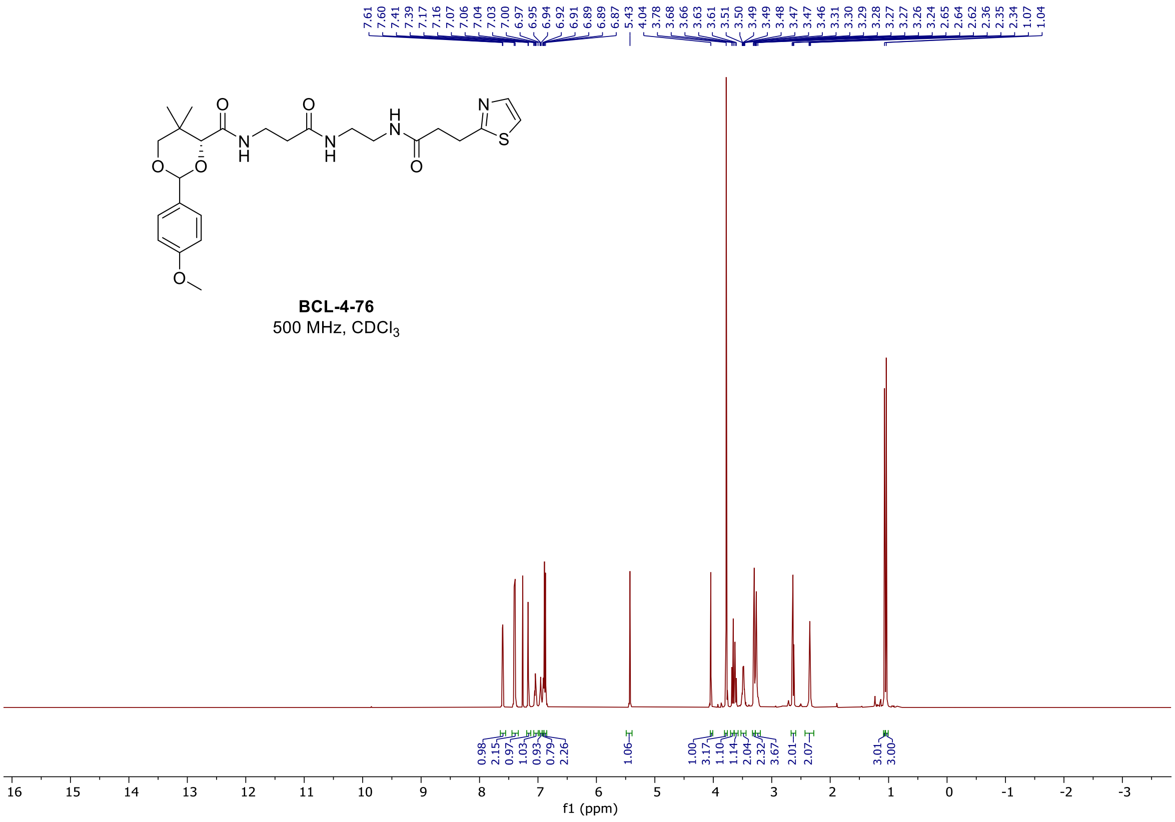

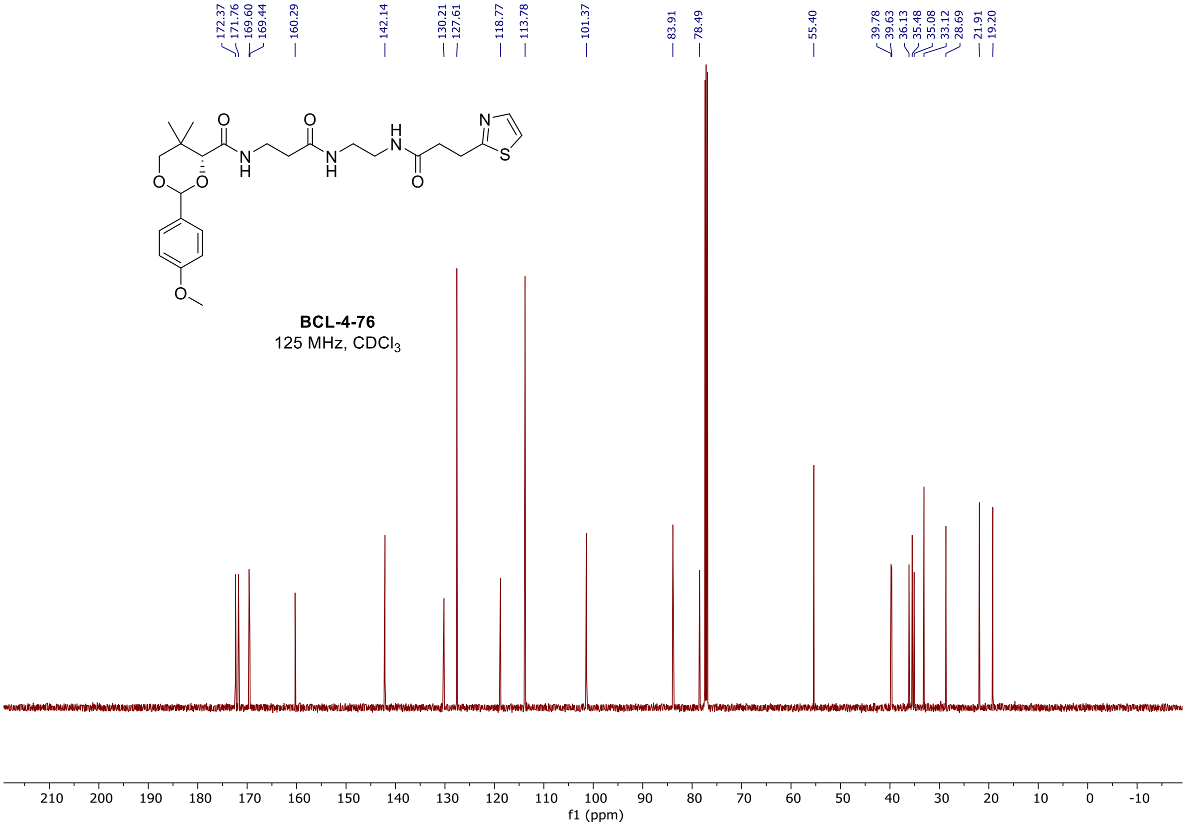

$^{1}H$, and $^{13}C$ spectra of **BCL-4-83**

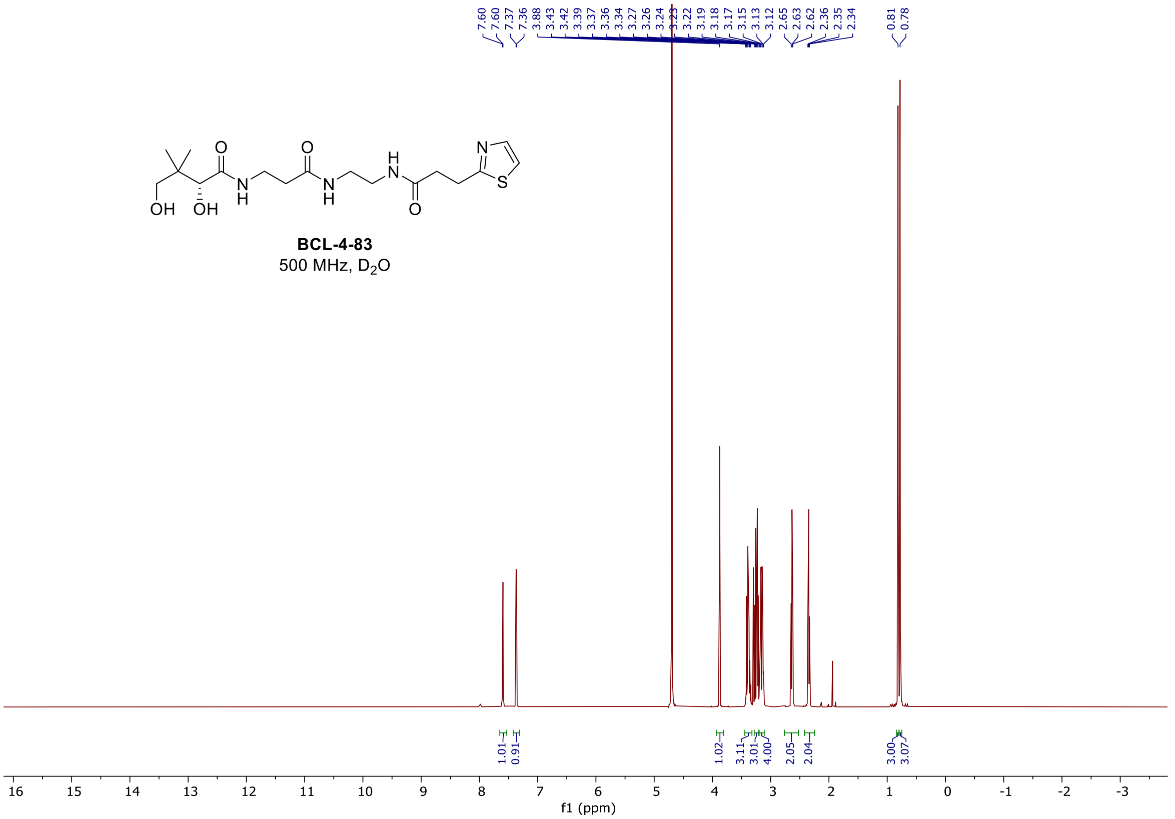

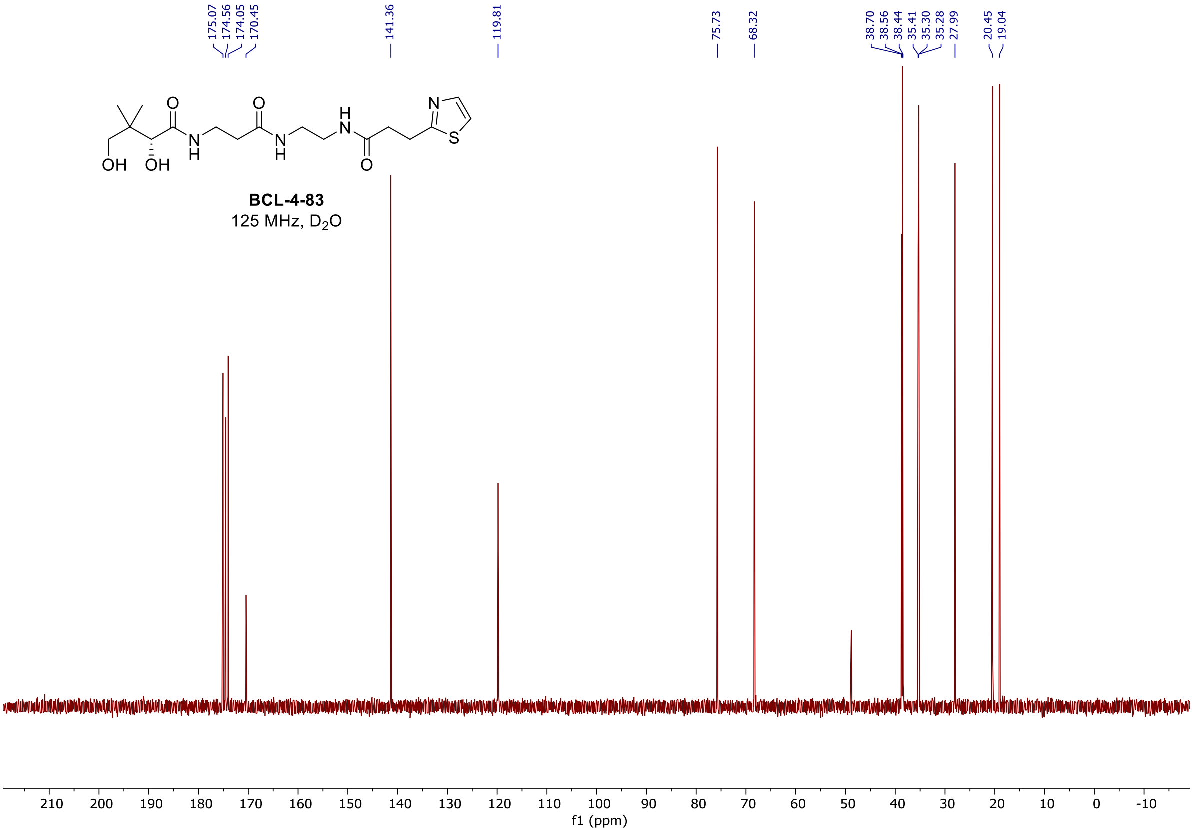

### $^{1}H$, and $^{13}C$ spectra of **BCL-4-86**

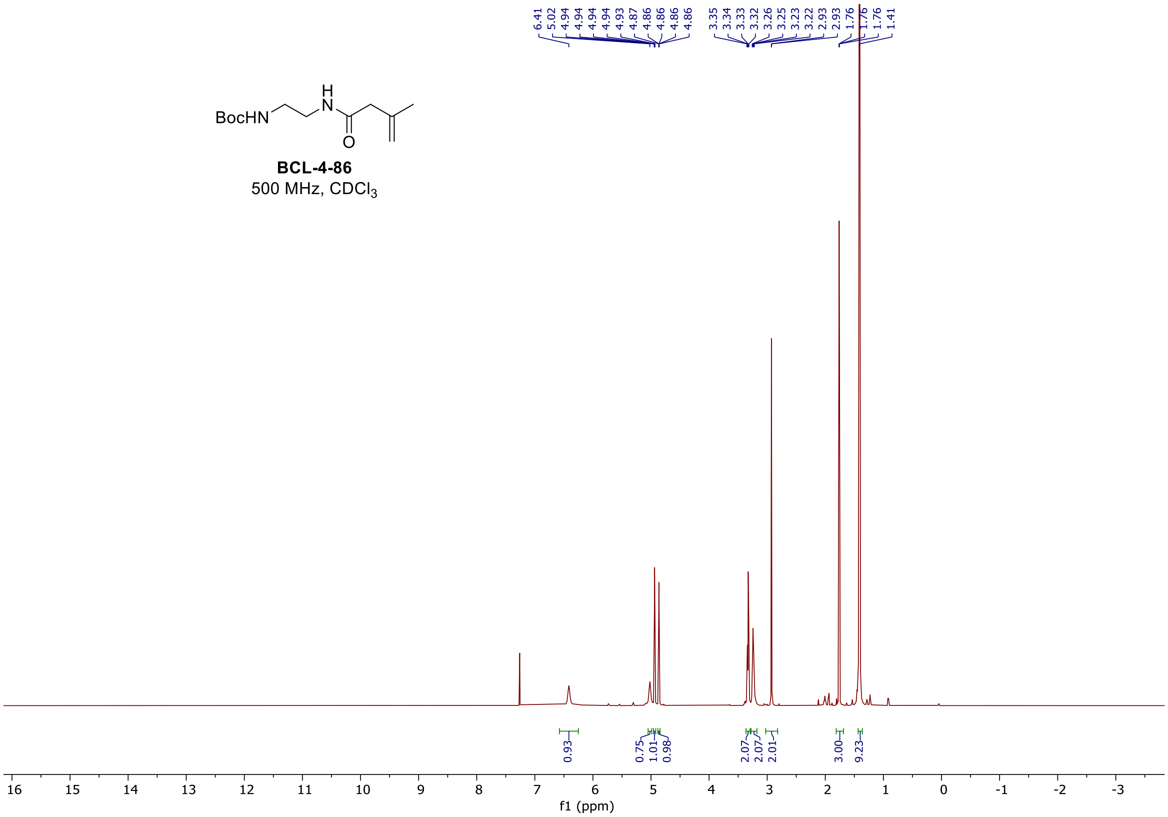

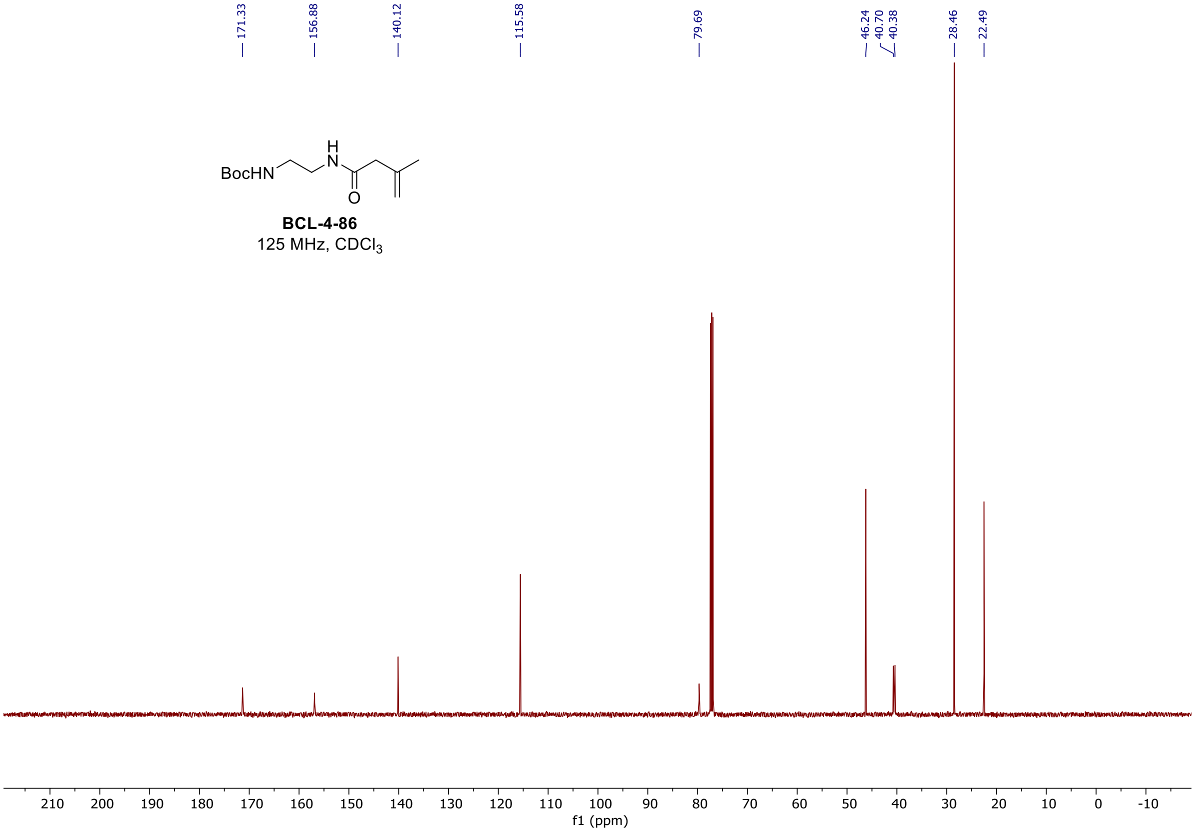

$^{1}H$, and $^{13}C$ spectra of **BCL-4-91**

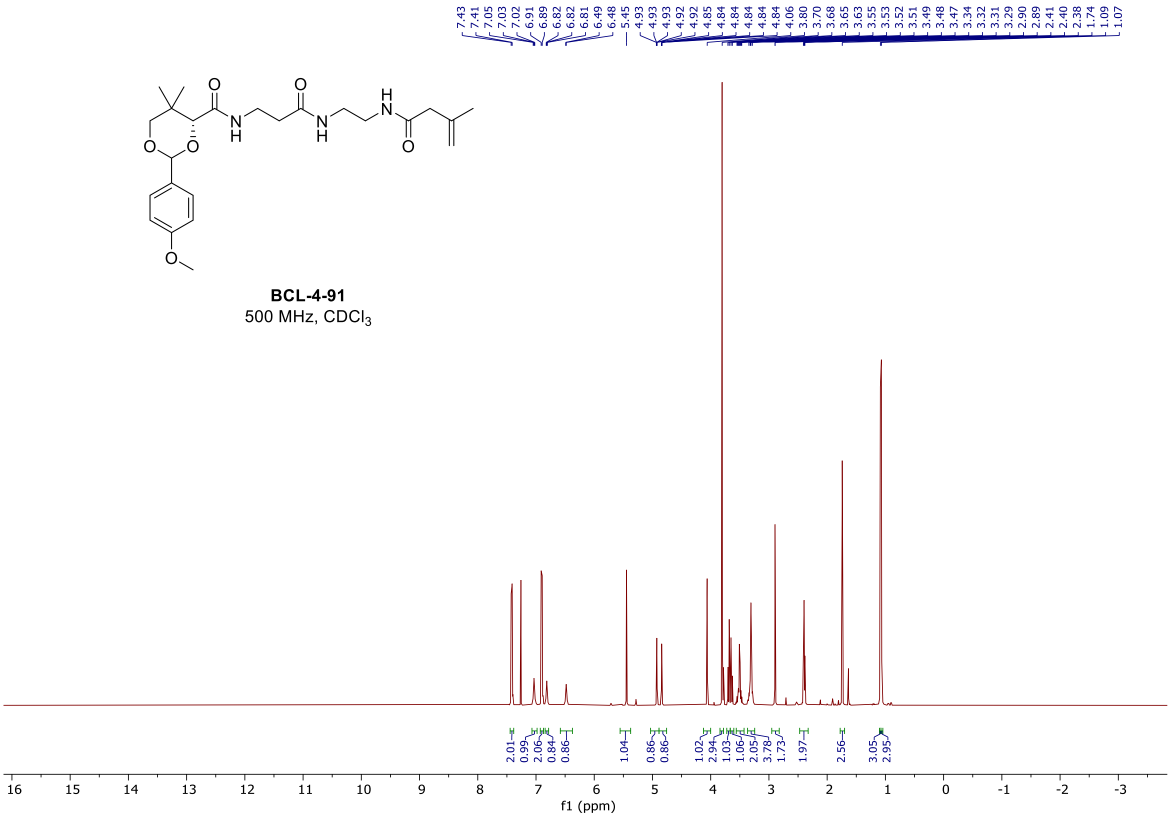

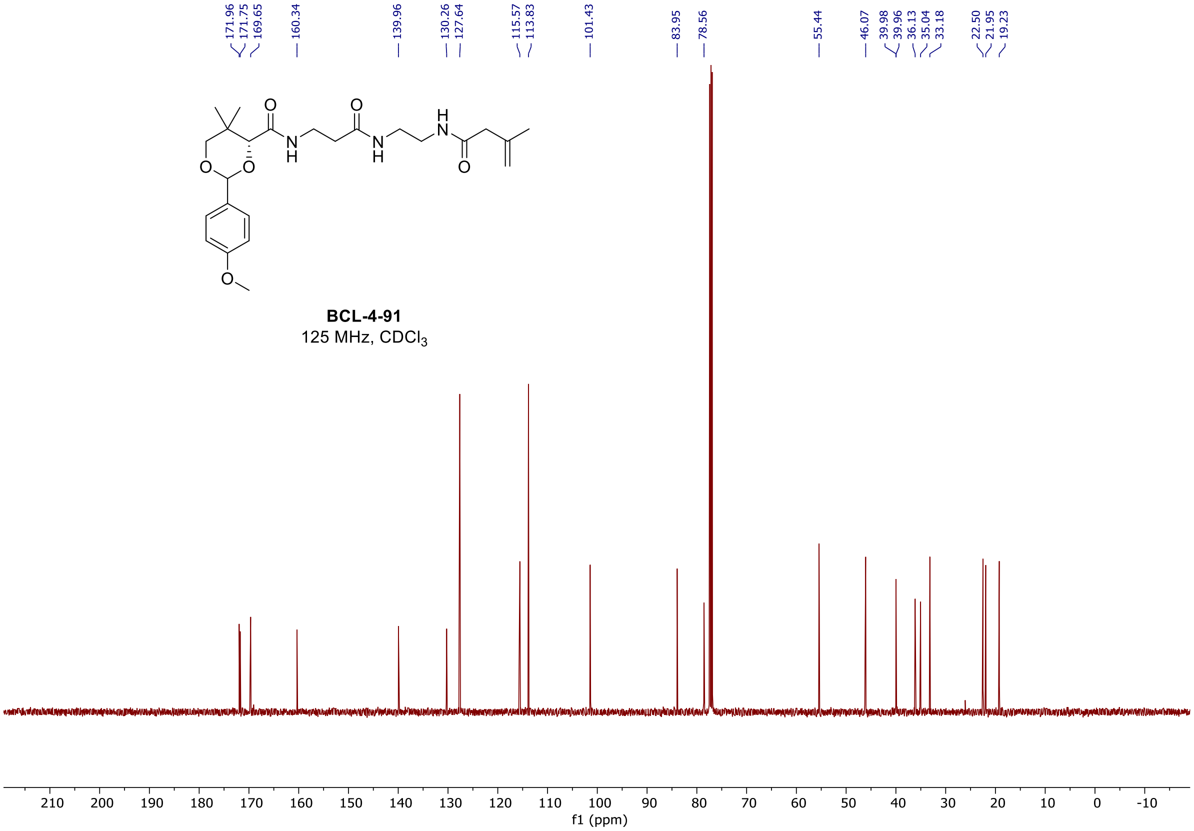

[1]H, and [13]C spectra of **BCL-4-95**

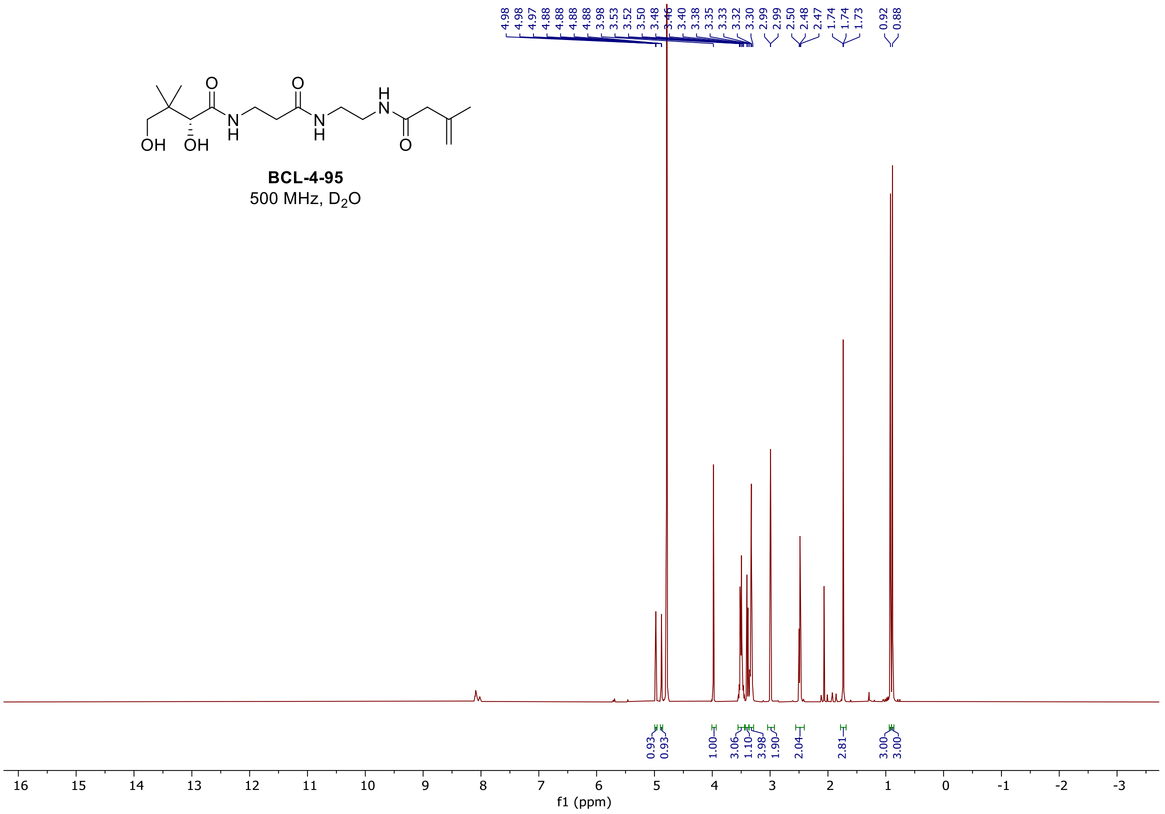

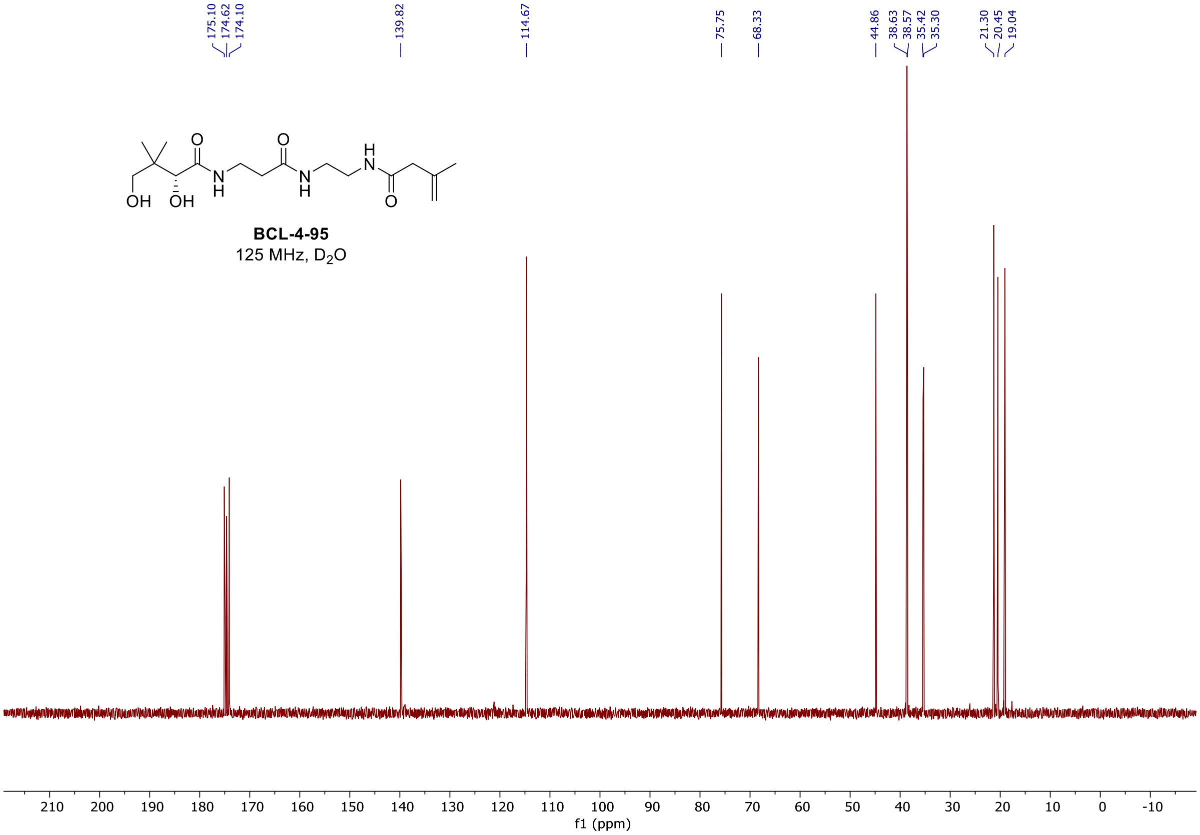

$^{1}$H, and $^{13}$C spectra of **BCL-4-99**

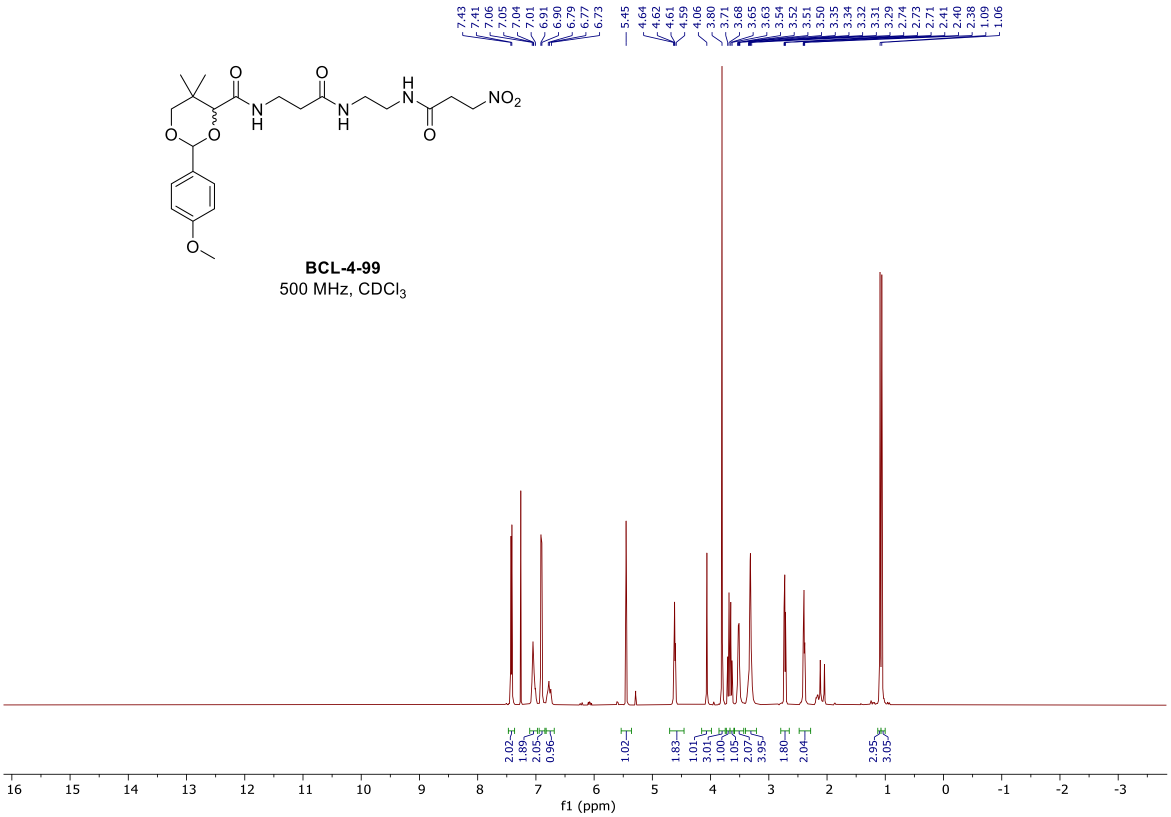

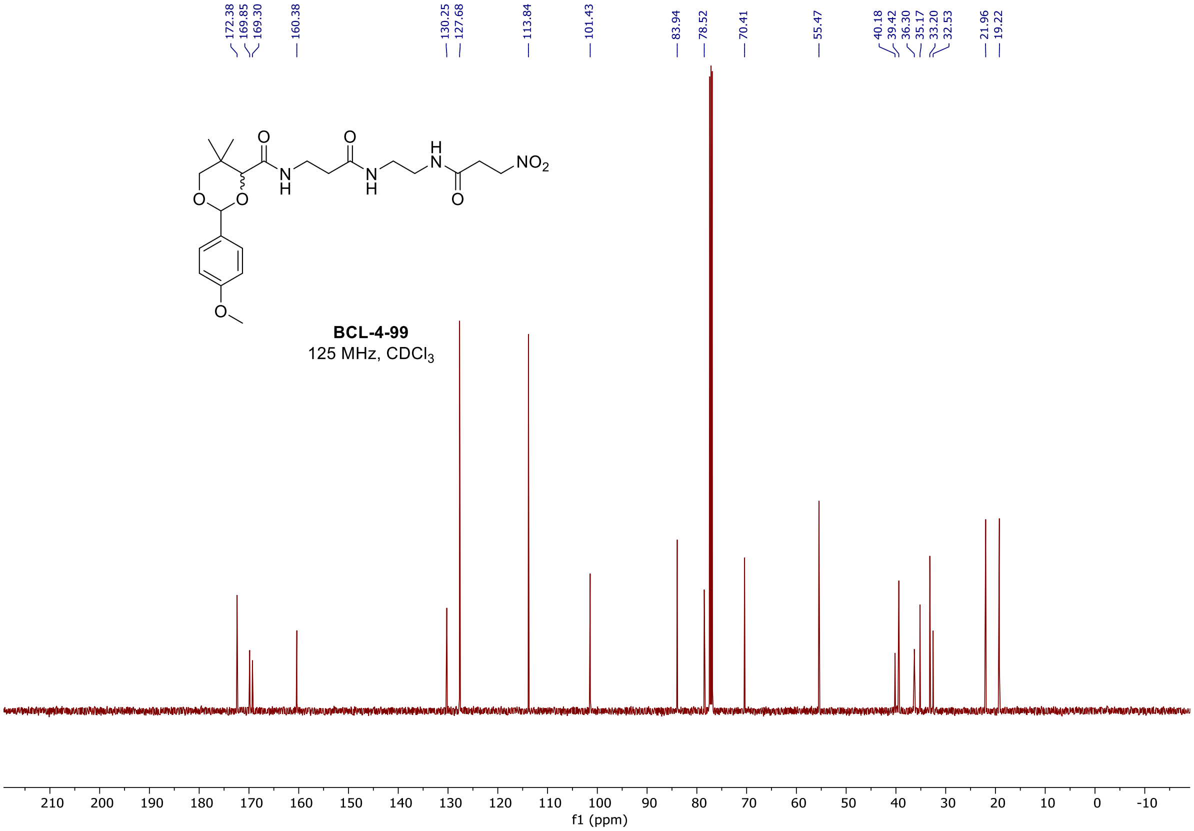

$^{1}H$, $^{19}F$, and $^{13}C$ spectra of **BCL-4-103**

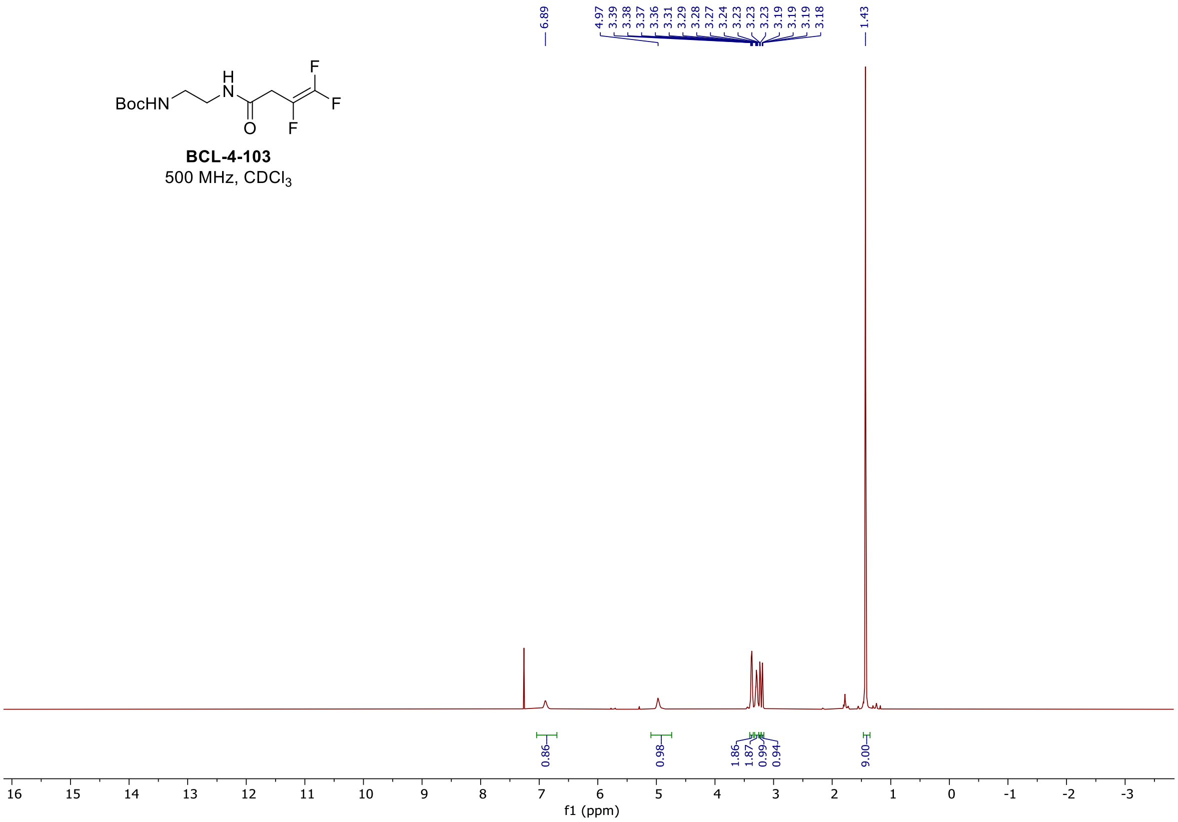

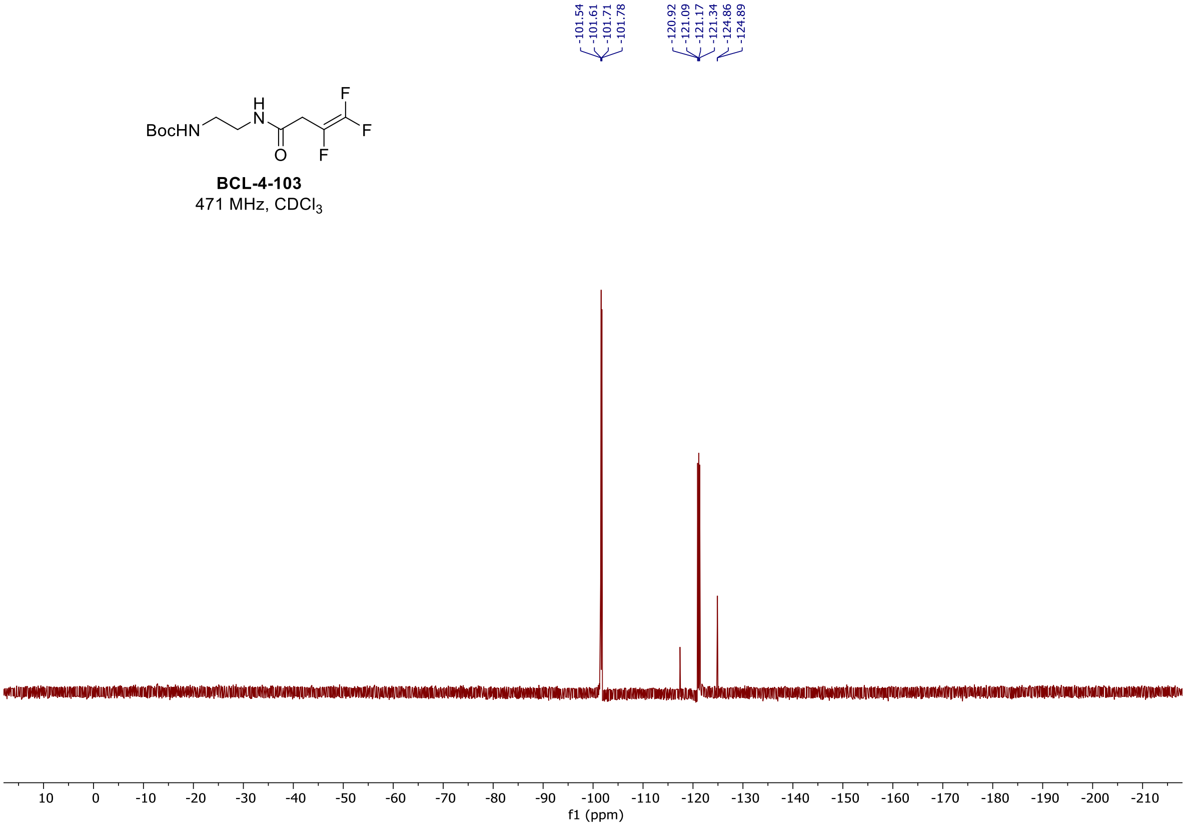

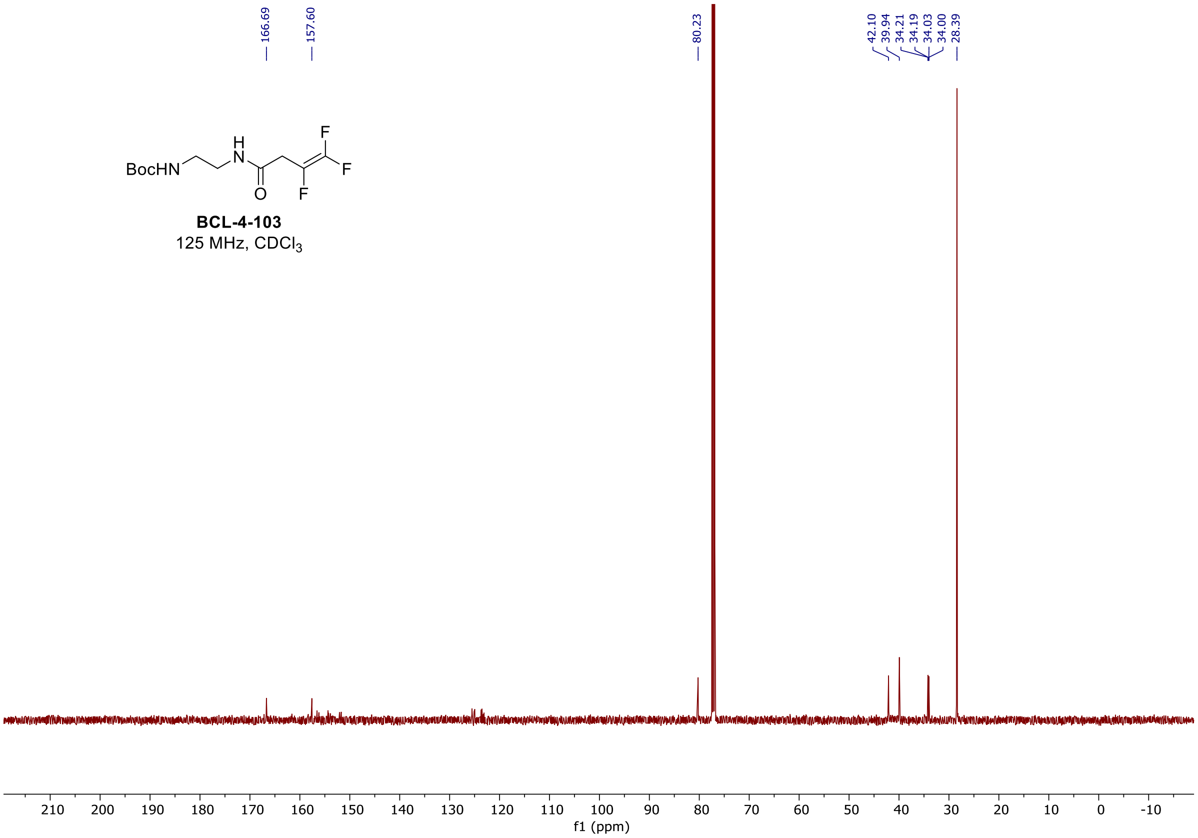

#### $^{1}H$, $^{19}F$, and $^{13}C$ spectra of **BCL-4-112**

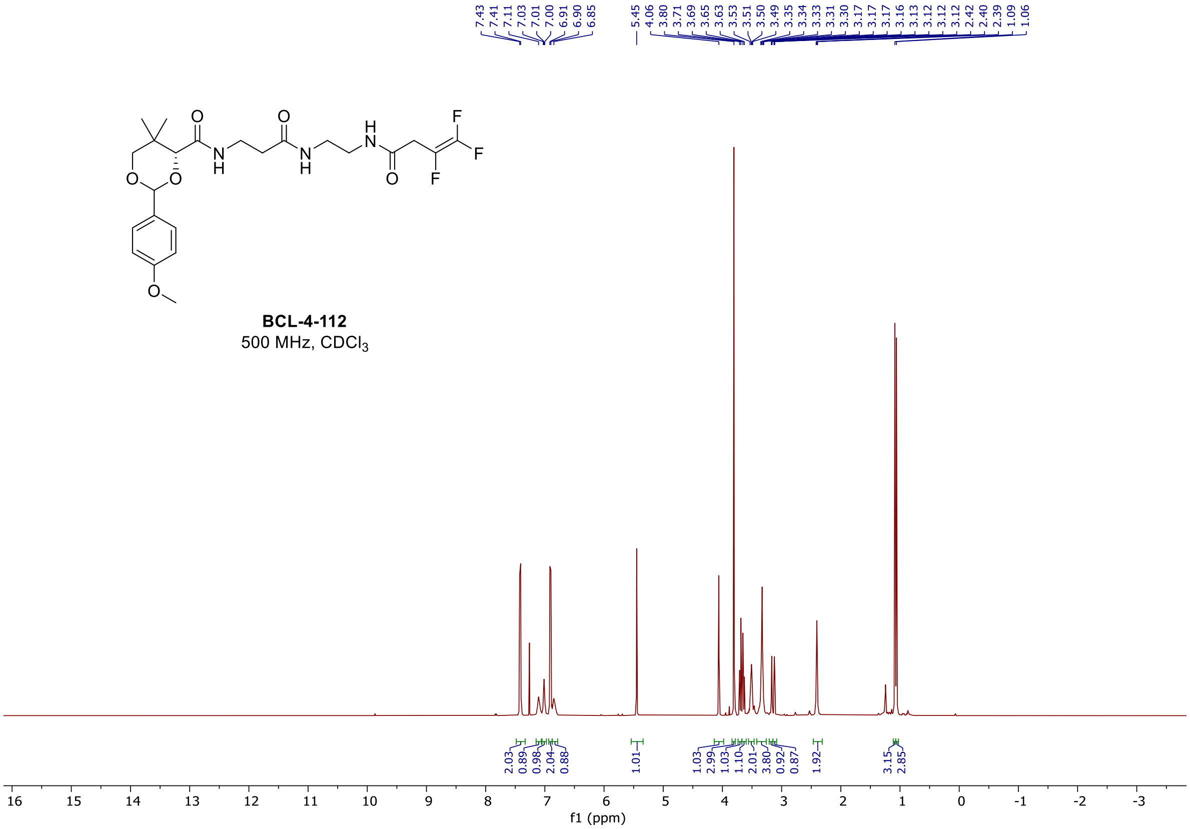

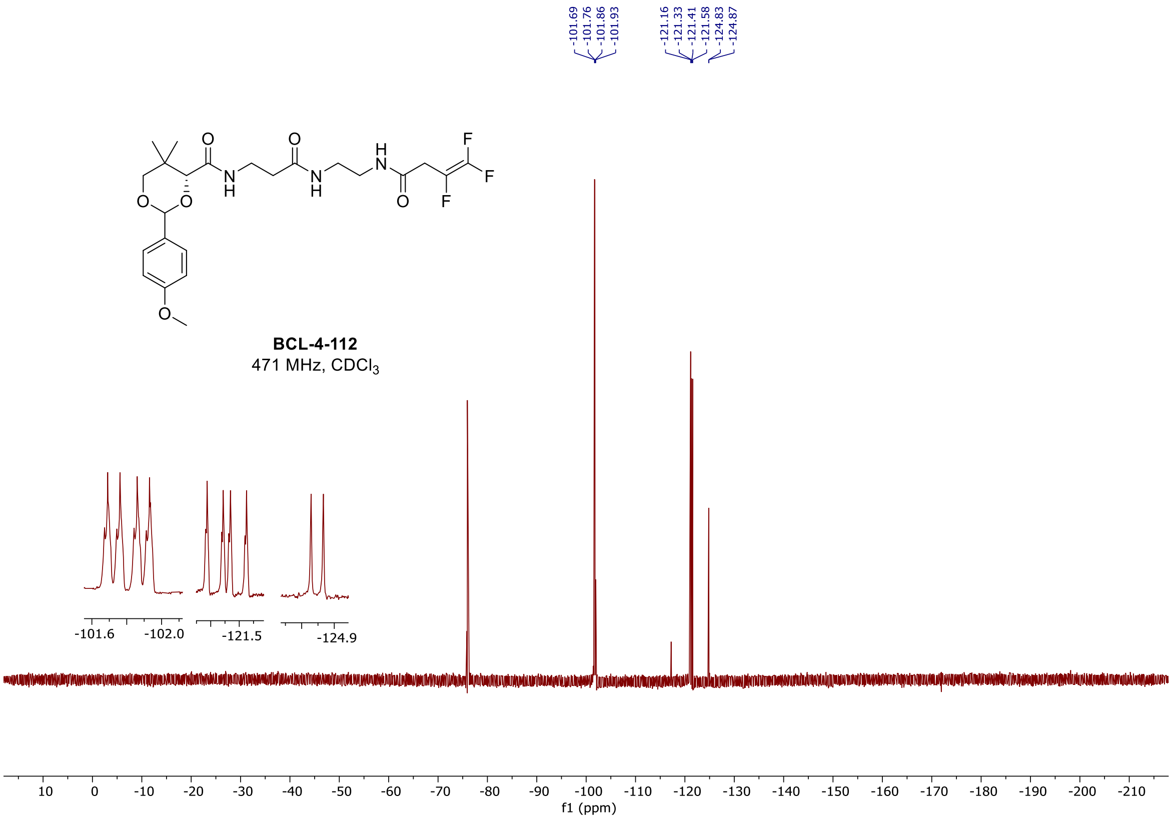

-101.69
-101.76
-101.86
-101.93
-121.16
-121.33
-121.41
-121.58
-124.83
-124.87
BCL-4-112
471 MHz, CDCl3
-101.6
-102.0
-121.5
-124.9
f1 (ppm)

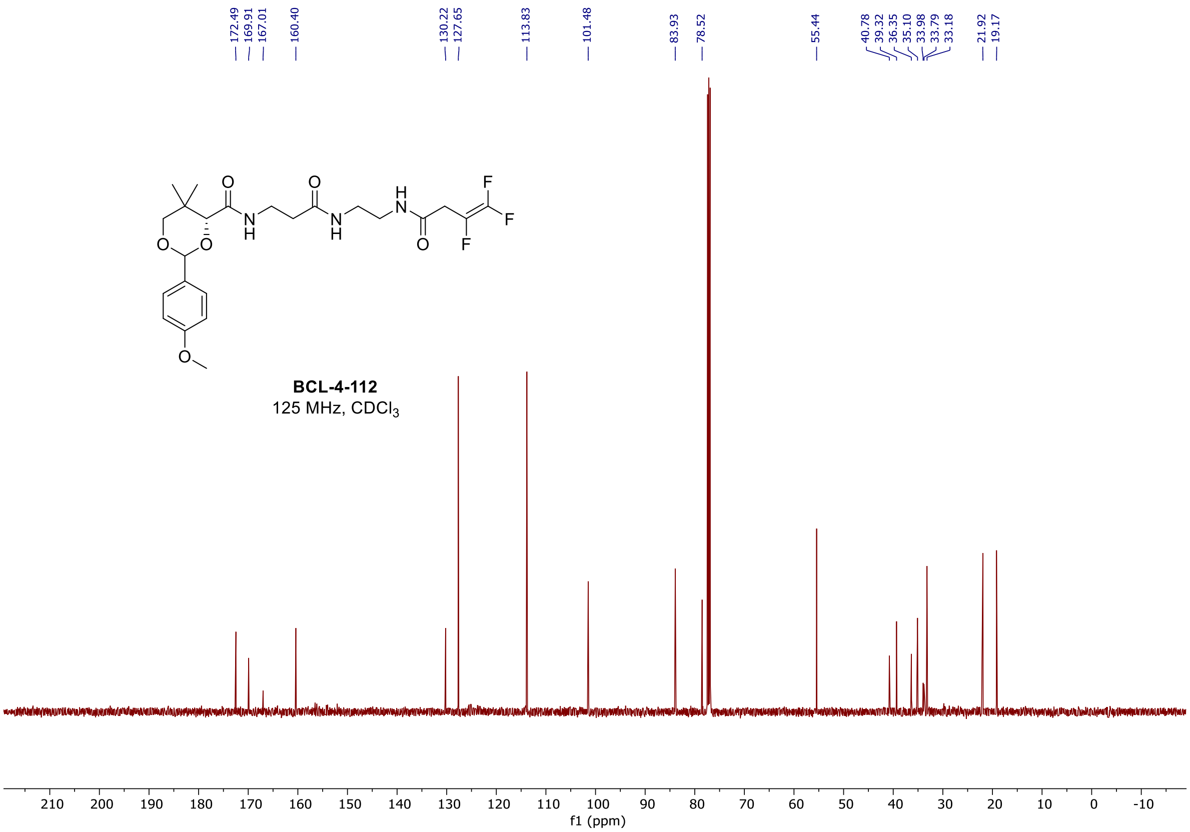

172.49
169.91
167.01
160.40
130.22
127.65
113.83
101.48
83.93
78.52
55.44
40.78
39.32
36.35
35.10
33.98
33.79
33.18
21.92
19.17
BCL-4-112
125 MHz, CDCl3
f1 (ppm)

<sup></sup>

#### $^{1}$H, $^{19}$F, and $^{13}$C spectra of **BCL-4-119**

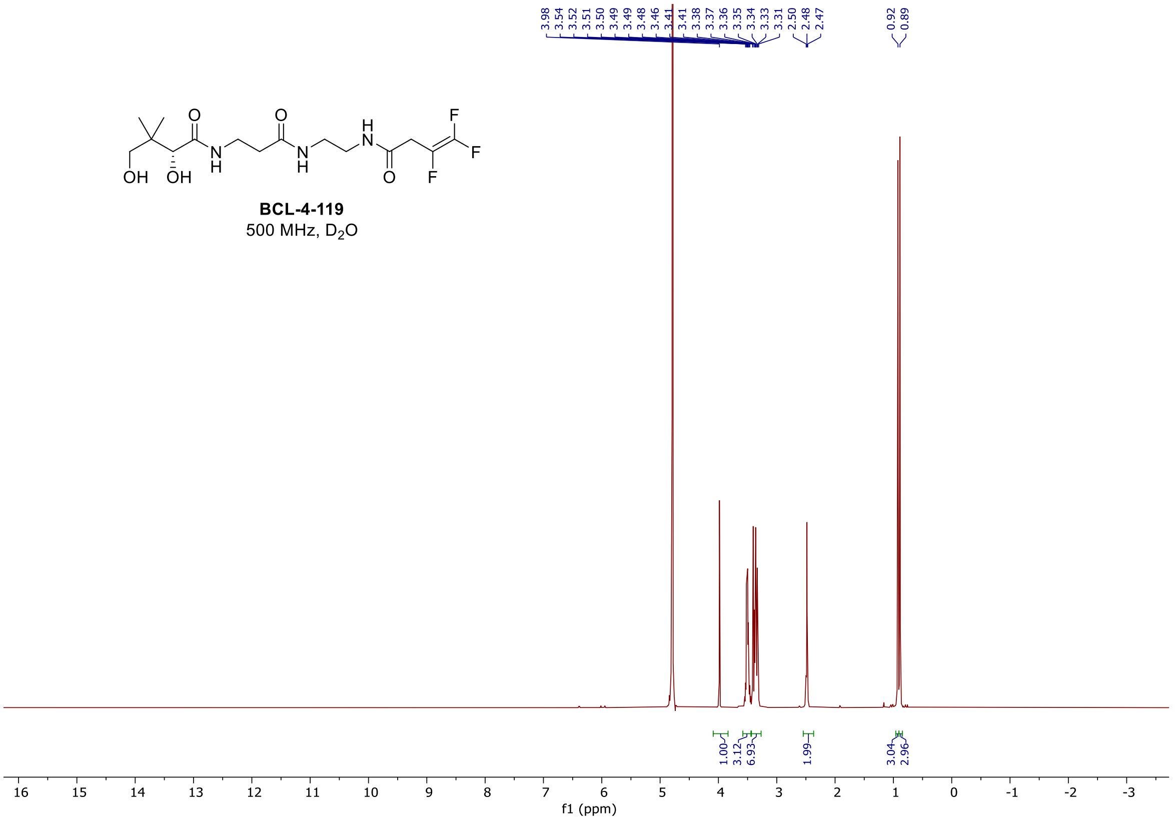

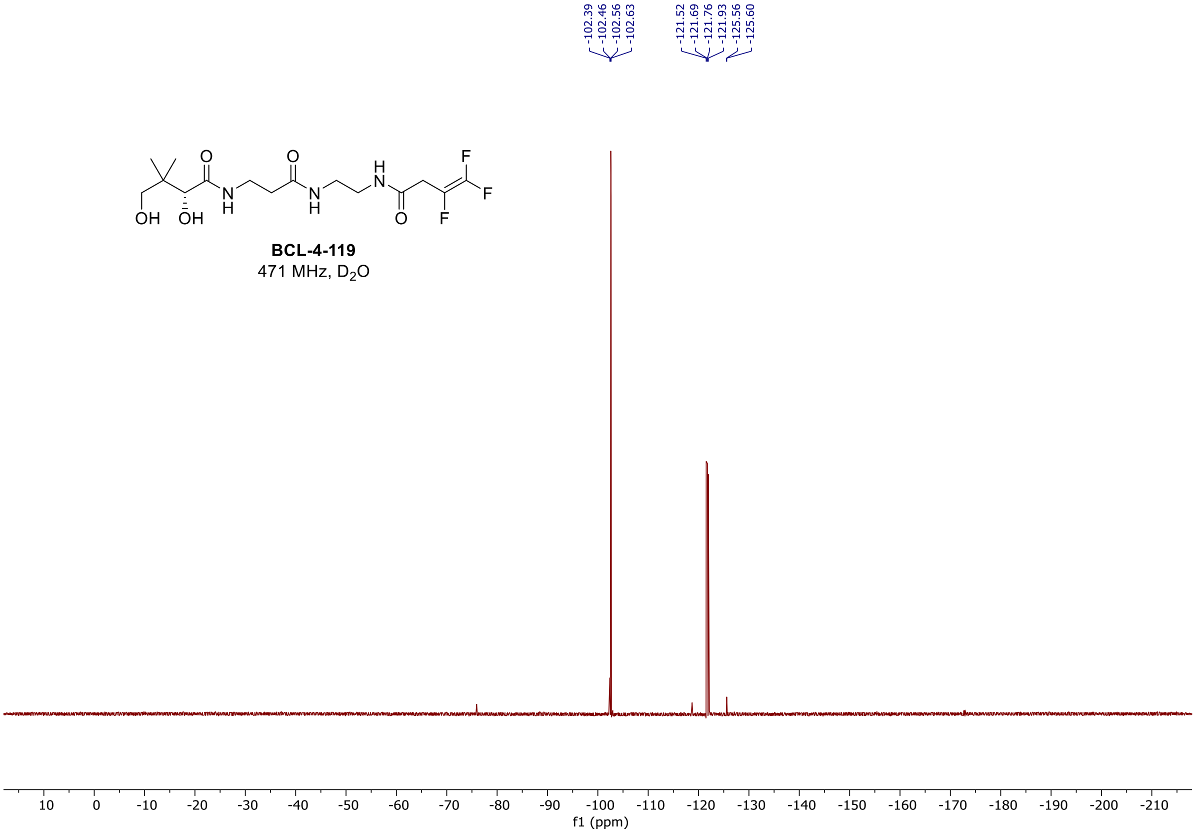

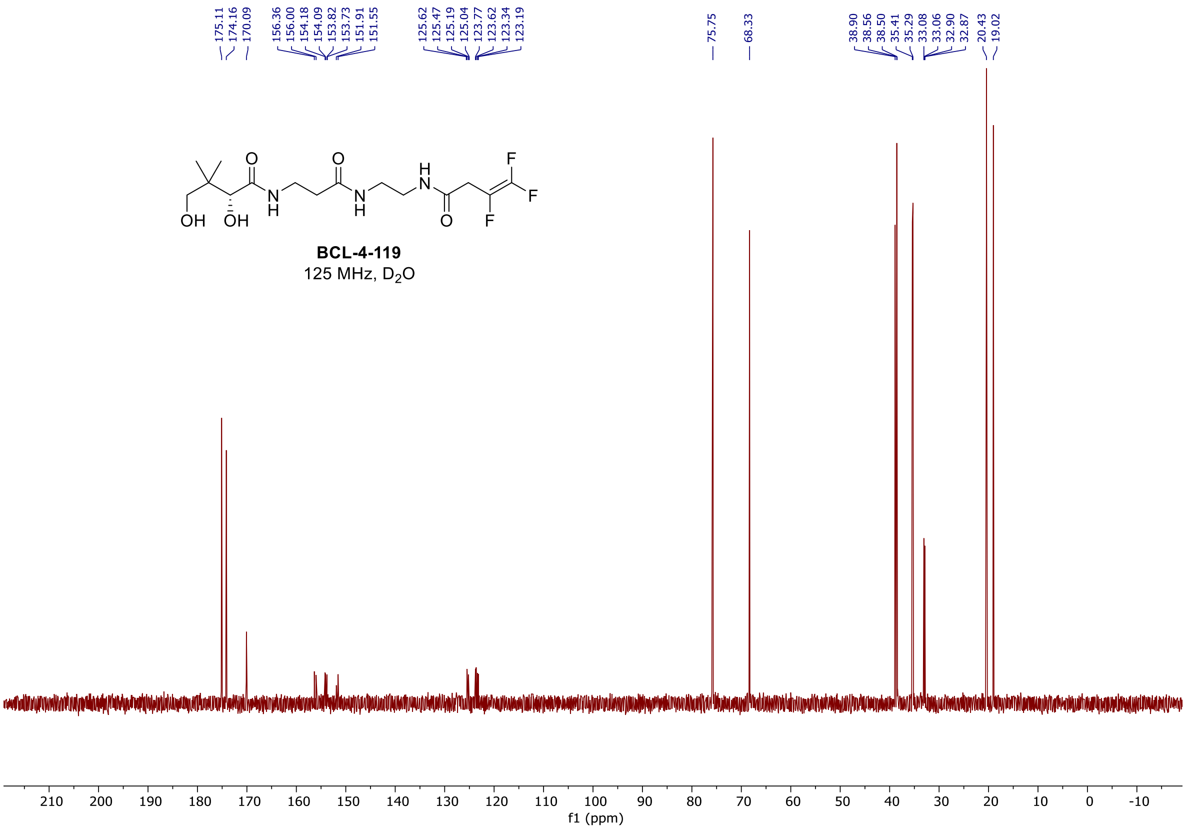

$^{1}H$ and $^{13}C$ spectra of **BCL-4-124**

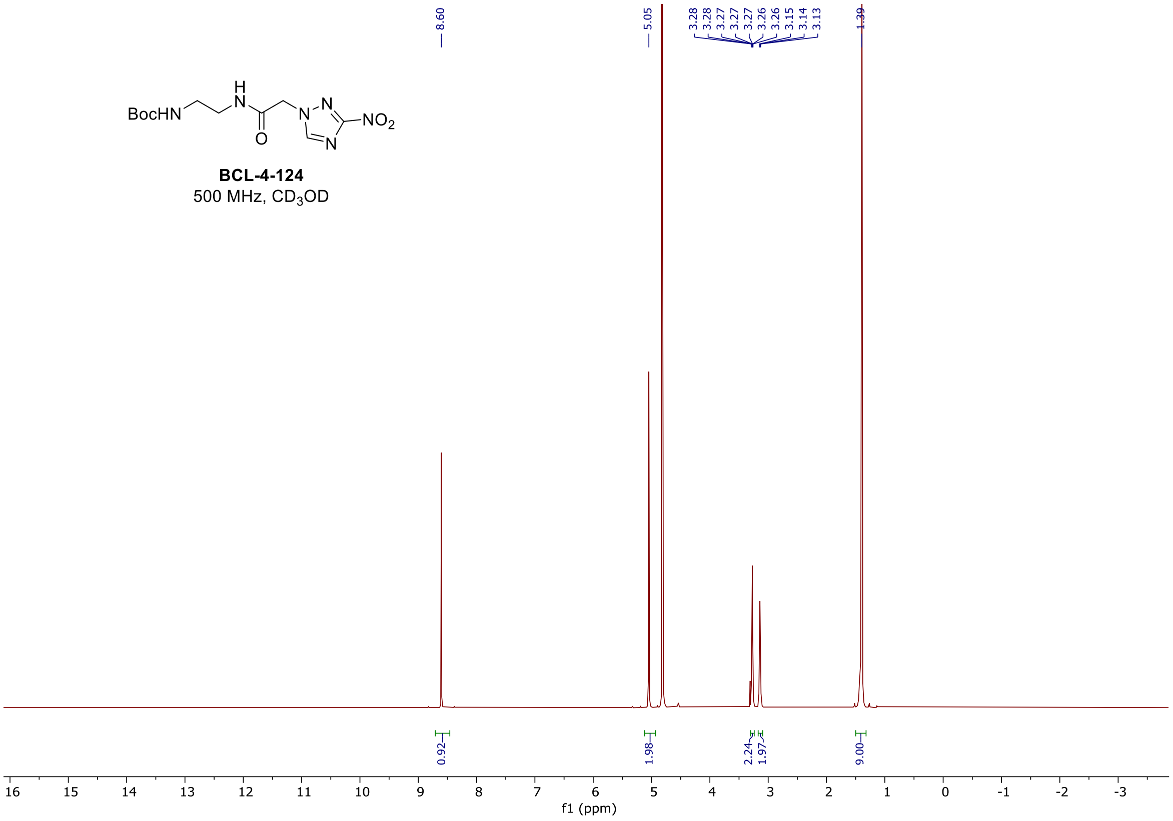

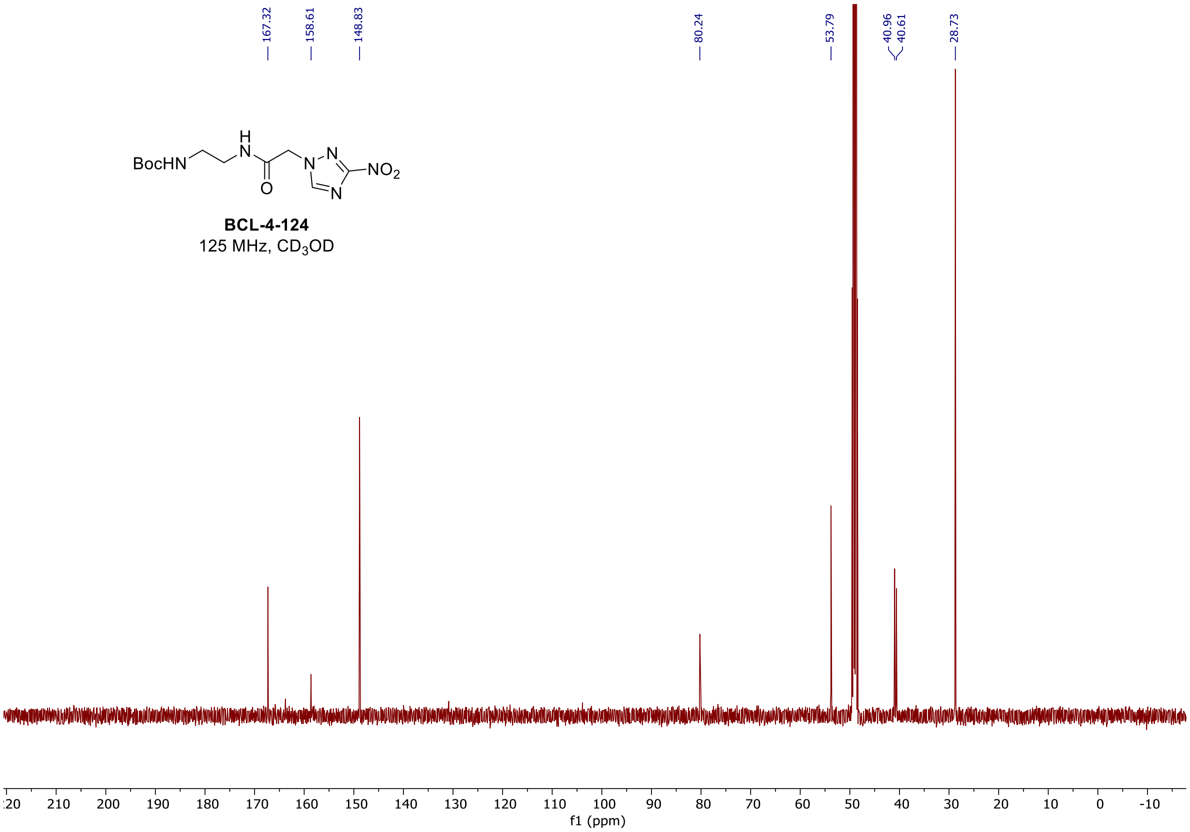

#### $^{1}H$ and $^{13}C$ spectra of **BCL-4-126**

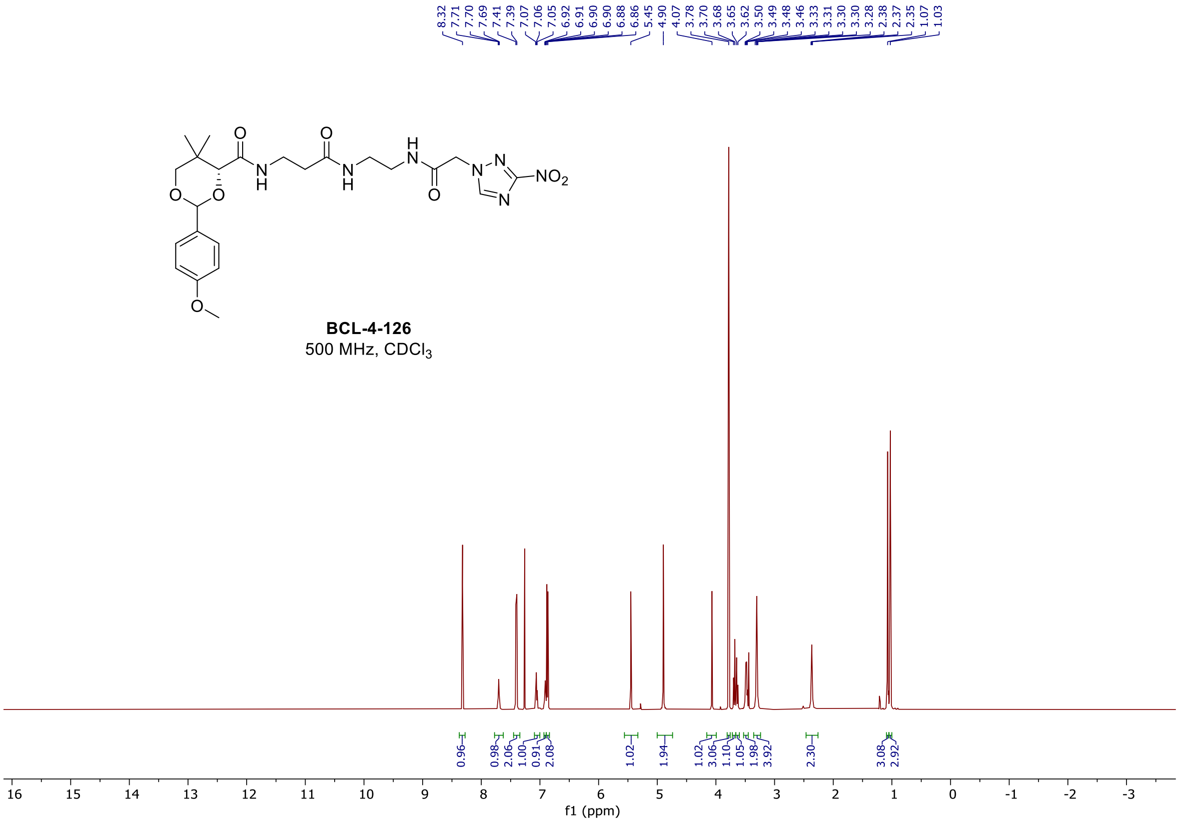

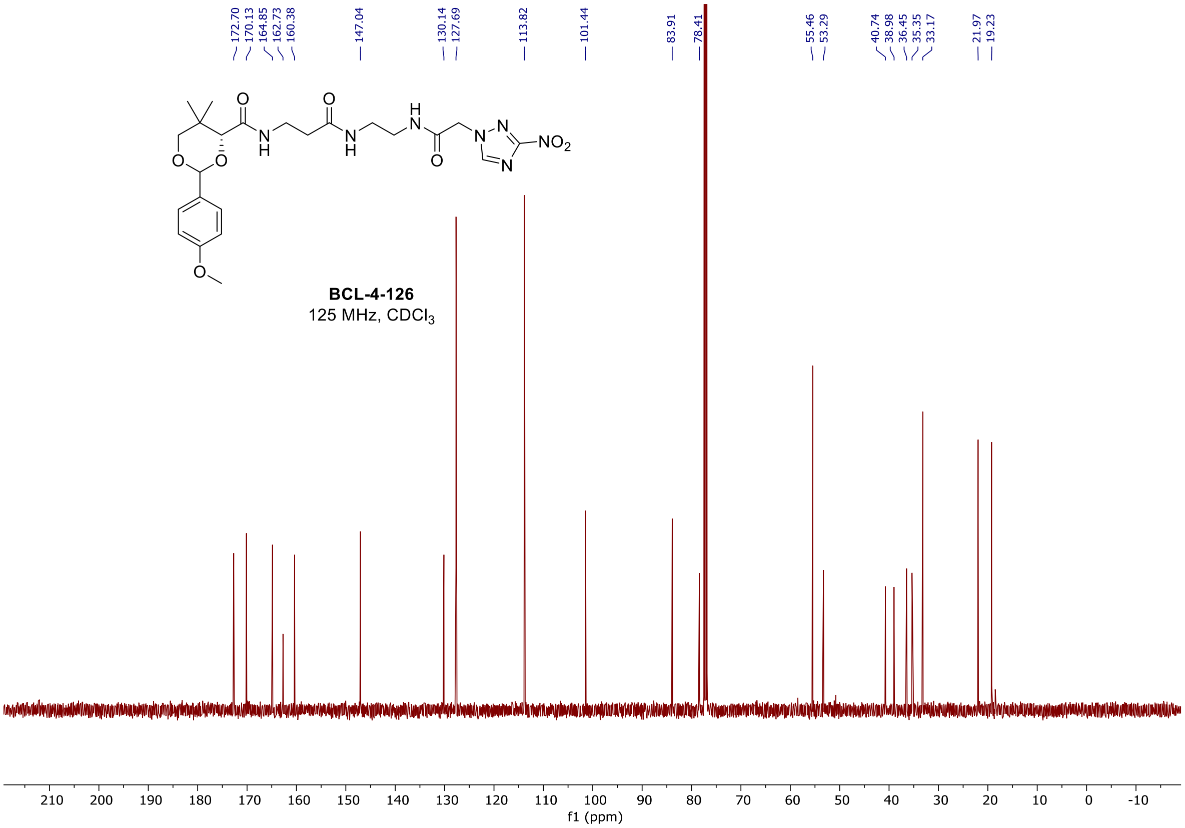

[1]H and [13]C spectra of **BCL-4-138**

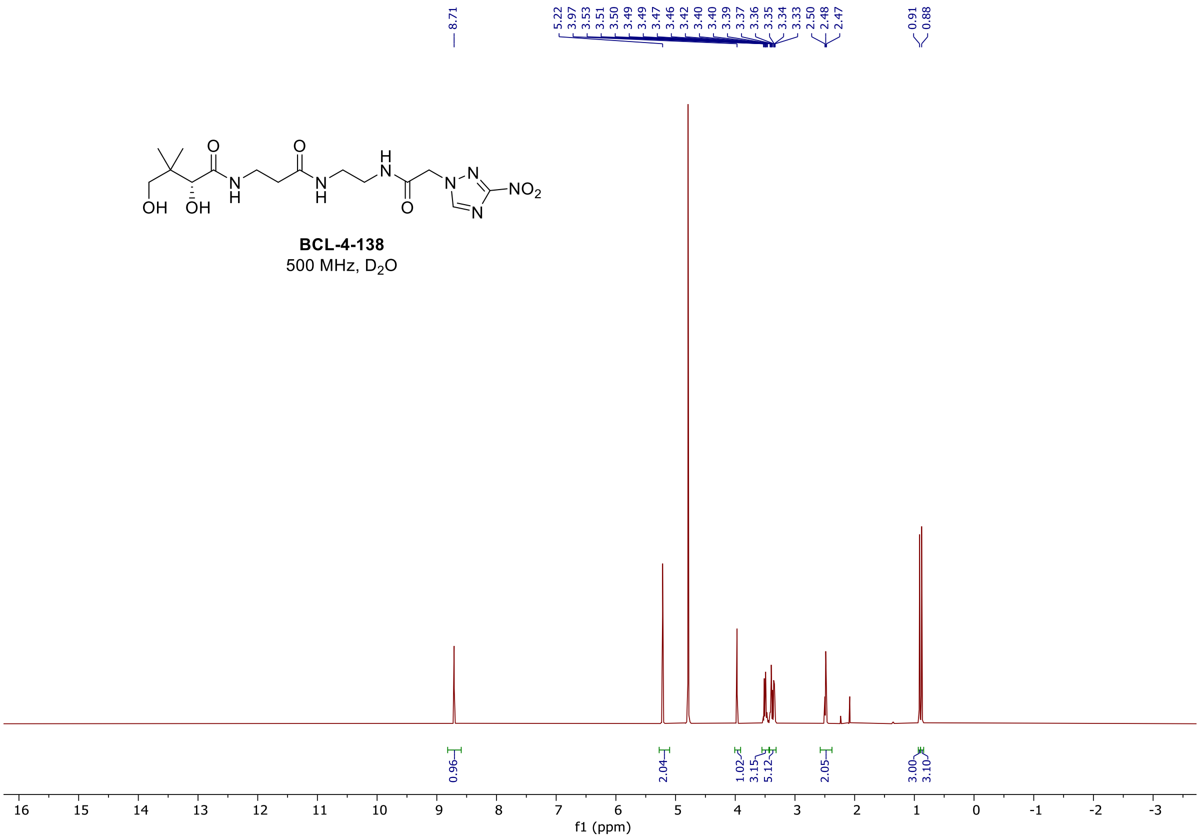

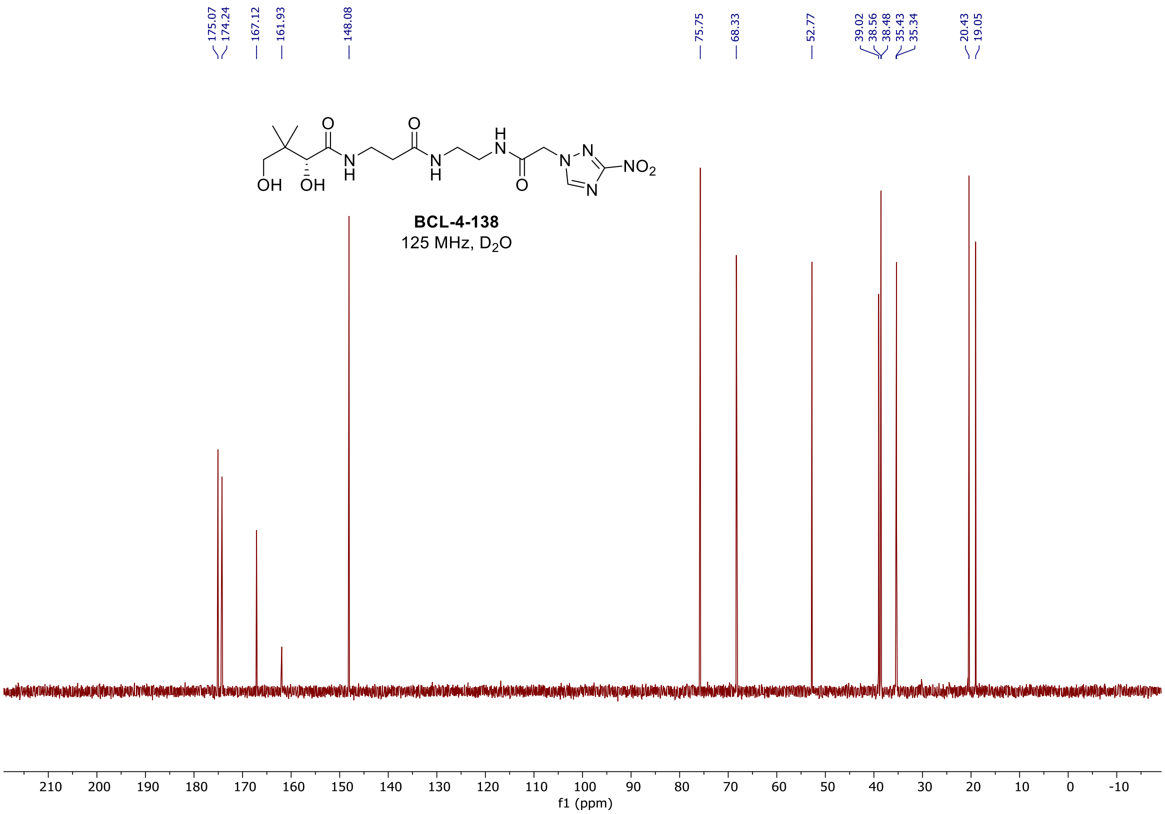

#### $^{1}H$, $^{19}F$, and $^{13}C$ spectra of **BCL-4-132**

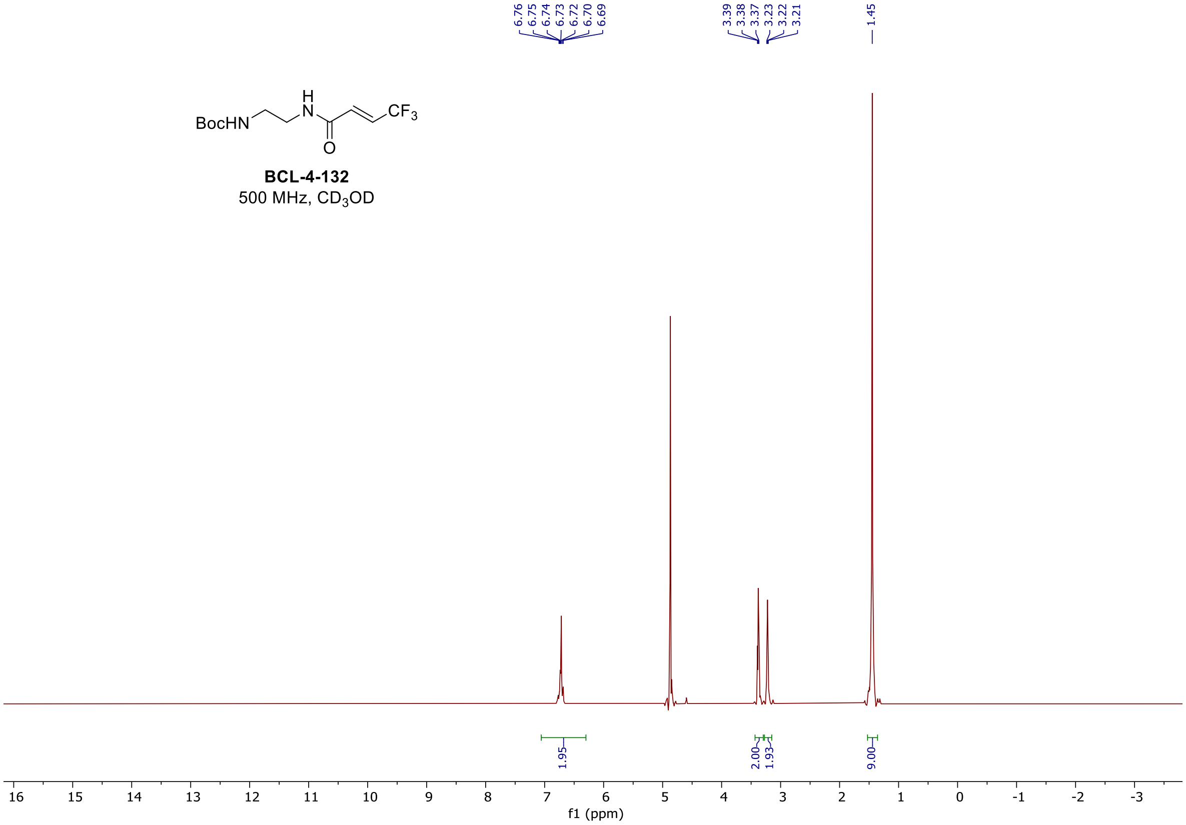

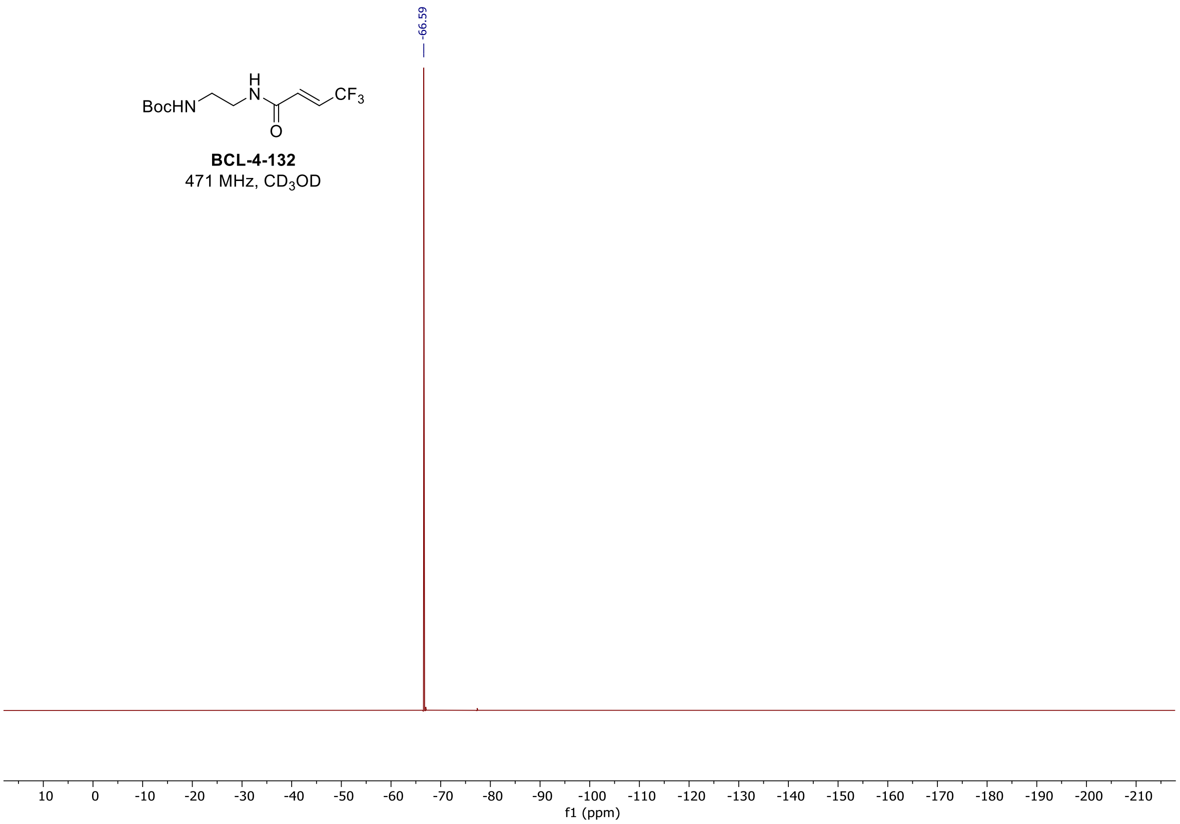

-66.59
BocHN
H
N
O
CF3
BCL-4-132
471 MHz, CD3OD
10 0 -10 -20 -30 -40 -50 -60 -70 -80 -90 -100 -110 -120 -130 -140 -150 -160 -170 -180 -190 -200 -210
f1 (ppm)

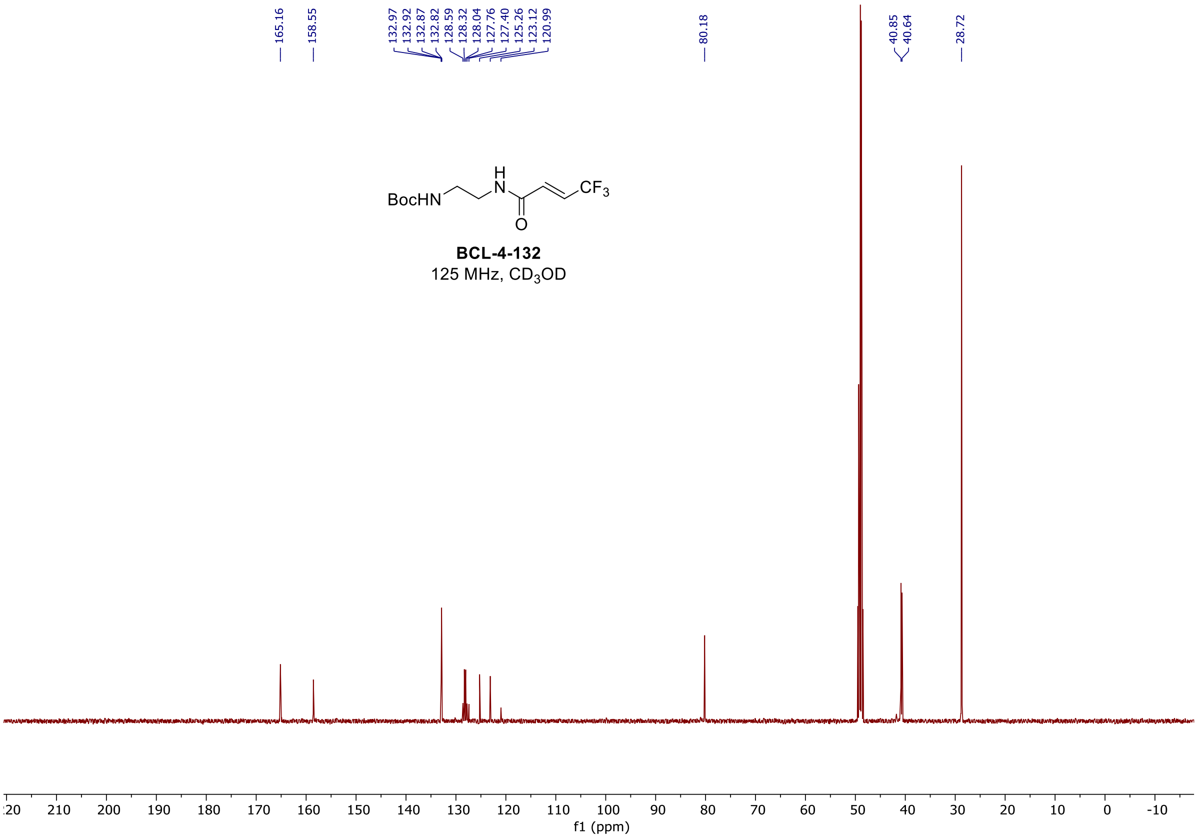

165.16
158.55
132.97
132.92
132.87
132.82
128.59
128.32
128.04
127.76
127.40
125.26
123.12
120.99
80.18
40.85
40.64
28.72
BocHN
H
N
O
CF3
BCL-4-132
125 MHz, CD3OD
20 210 200 190 180 170 160 150 140 130 120 110 100 90 80 70 60 50 40 30 20 10 0 -10
f1 (ppm)

<u>$^{1}$H, $^{19}$F, and $^{13}$C spectra of **BCL-4-144**</u>

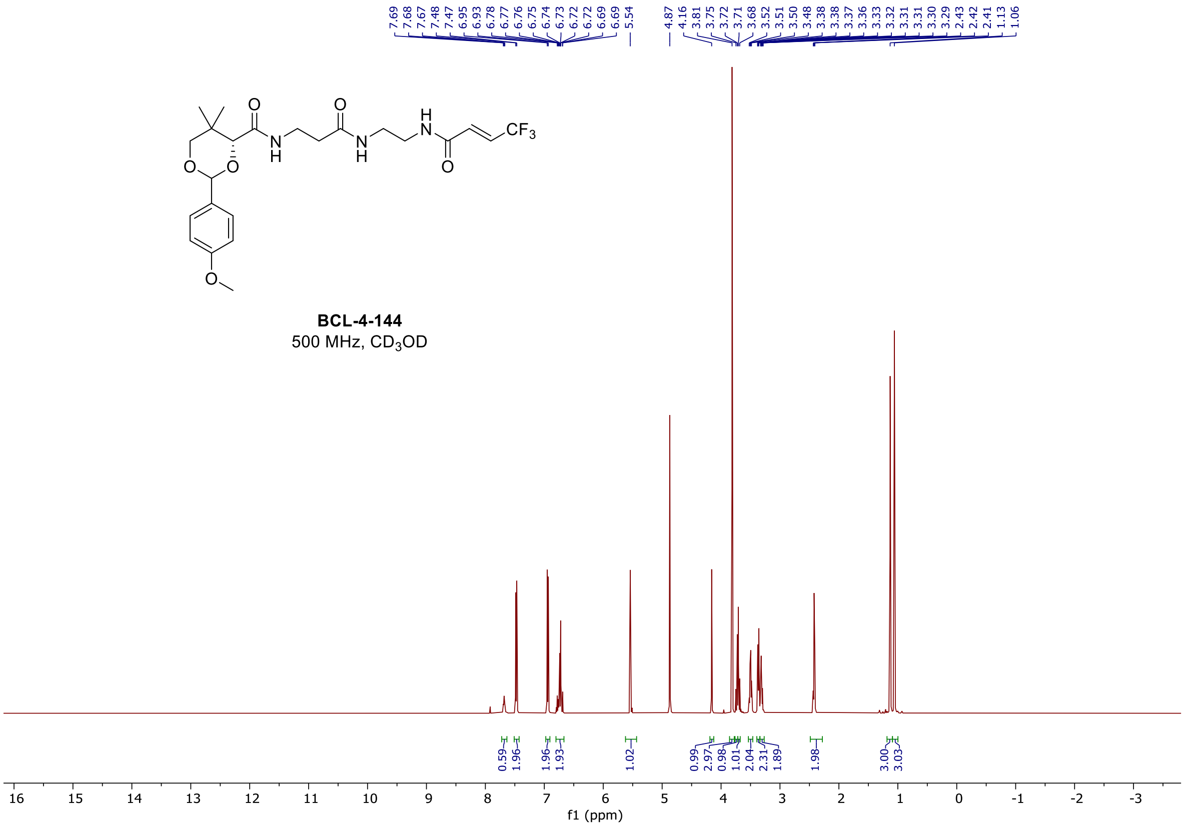

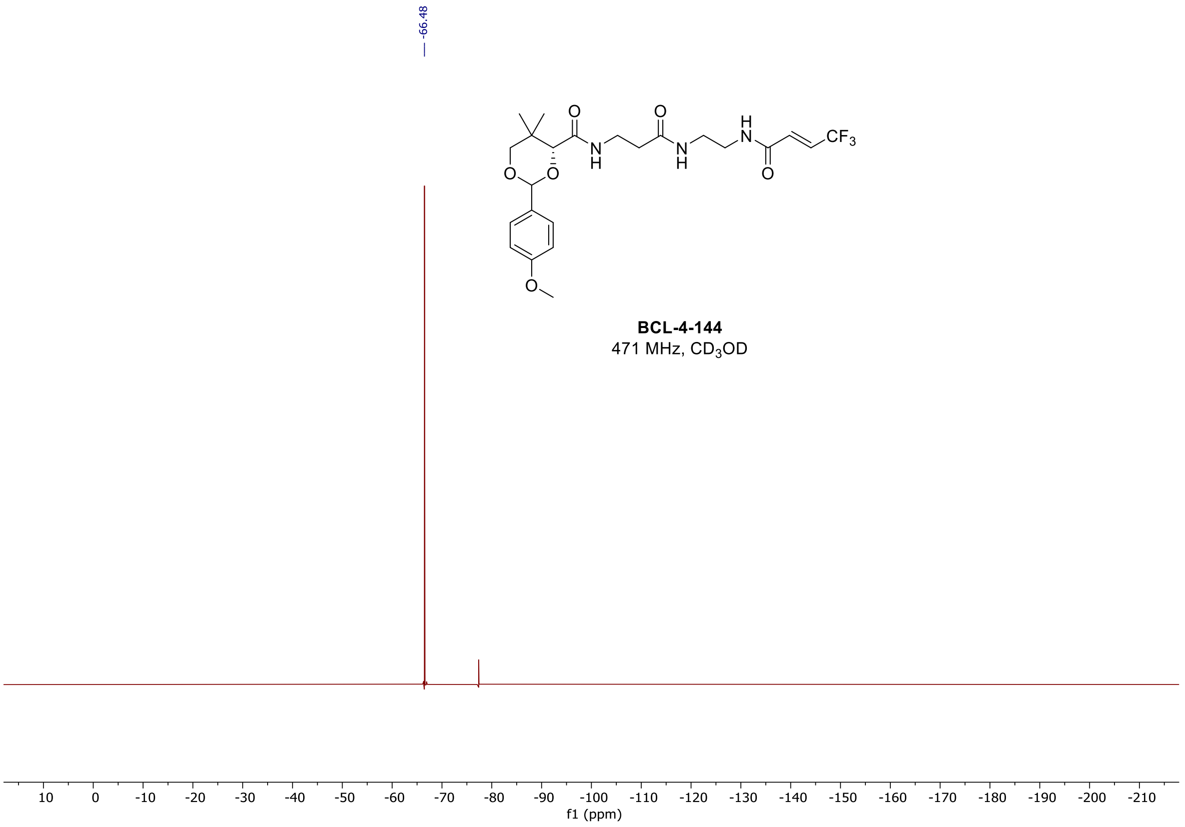

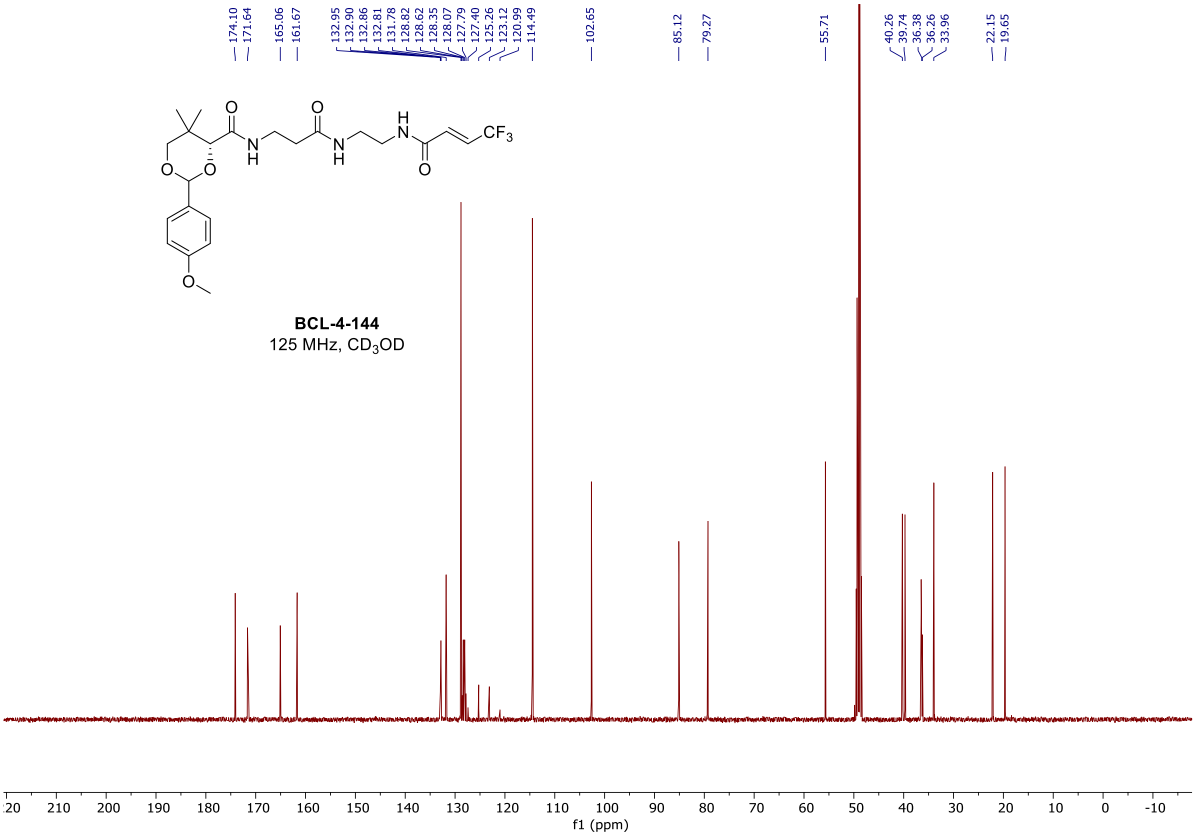

<u>$^{1}$H, $^{19}$F, and $^{13}$C spectra of **BCL-4-148**</u>

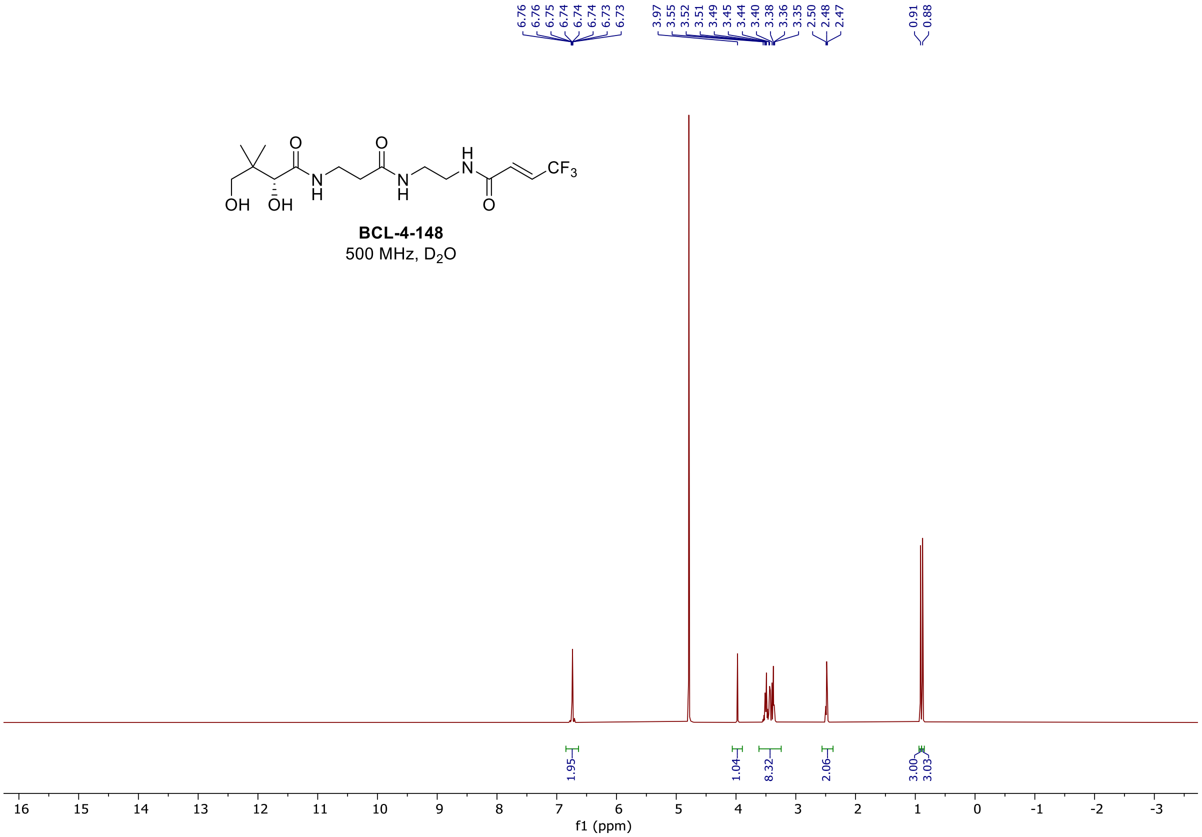

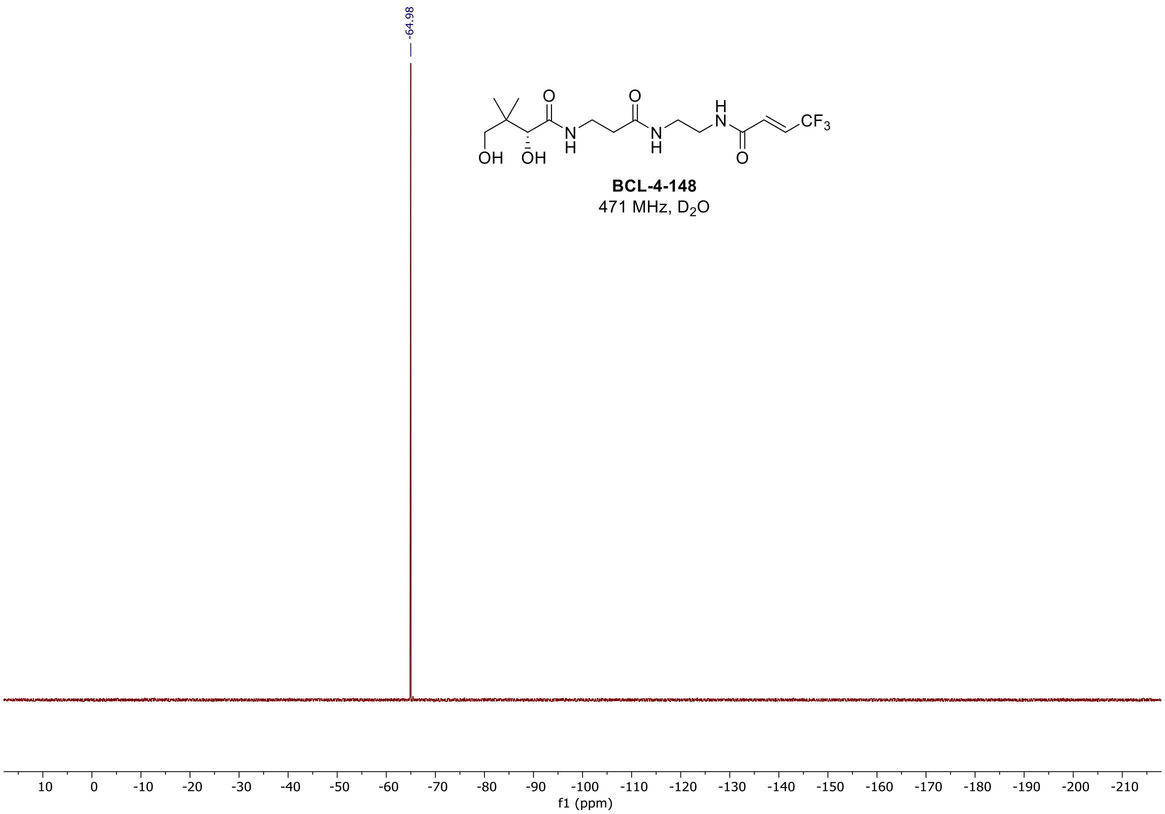
-64.98
BCL-4-148
471 MHz, D2O
f1 (ppm)

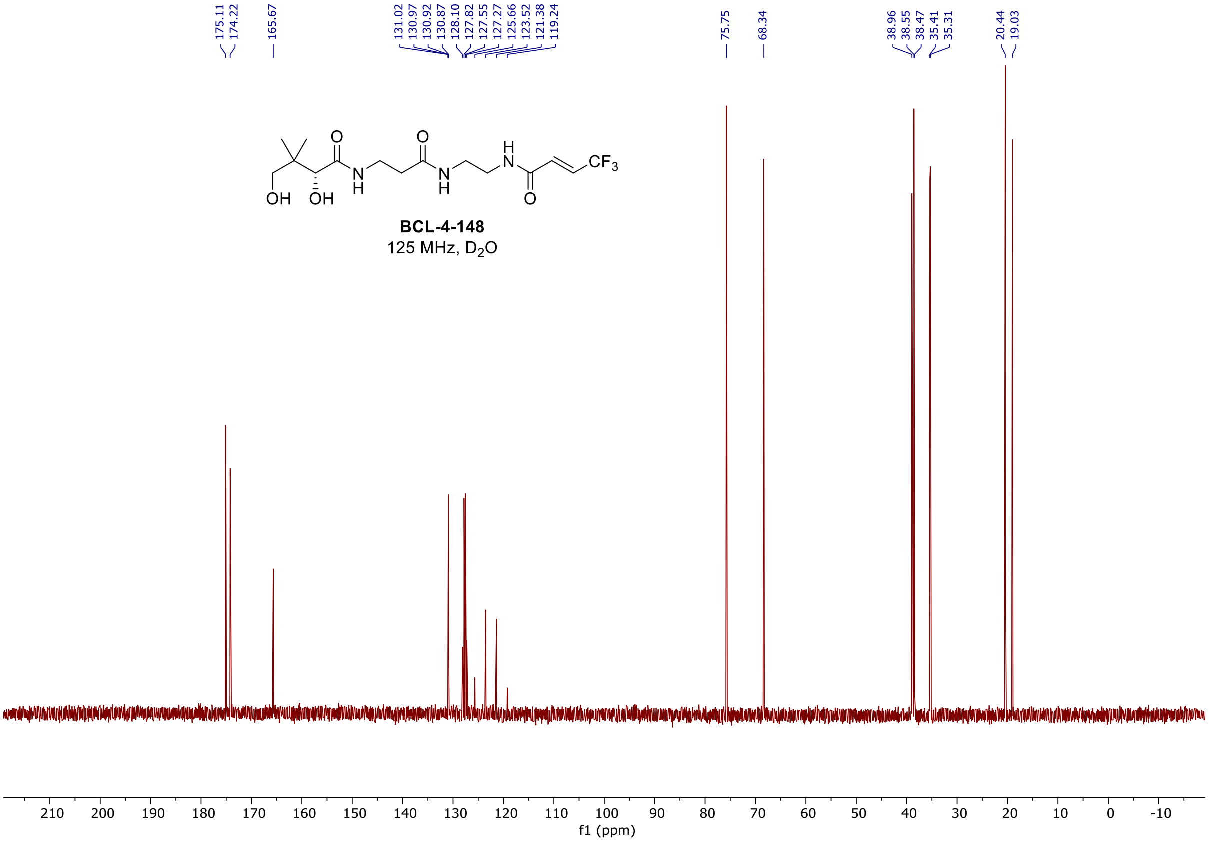
175.11
174.22
165.67
131.02
130.97
130.92
130.87
128.10
127.82
127.55
127.27
125.66
123.52
121.38
119.24
75.75
68.34
38.96
38.55
38.47
35.41
35.31
20.44
19.03
BCL-4-148
125 MHz, D2O
f1 (ppm)

#### $^1$H and $^{13}$C spectra of **BCL-4-150**

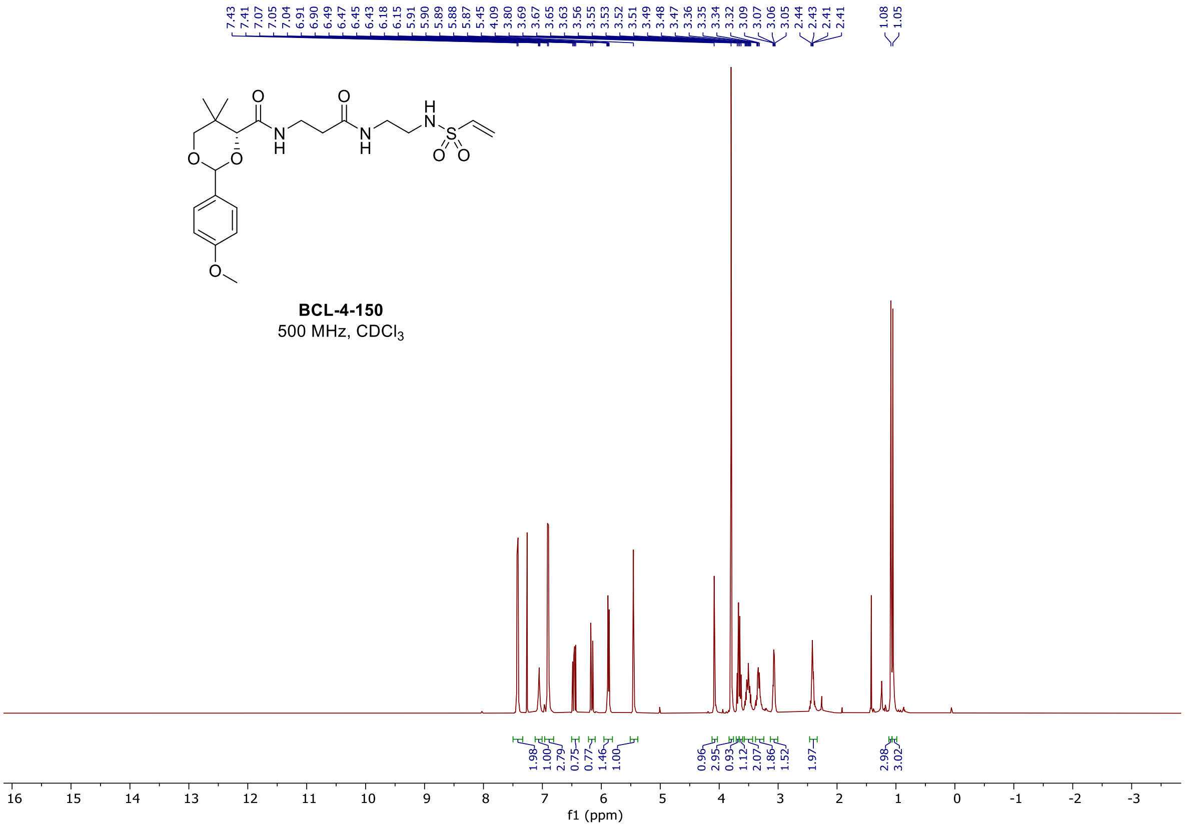

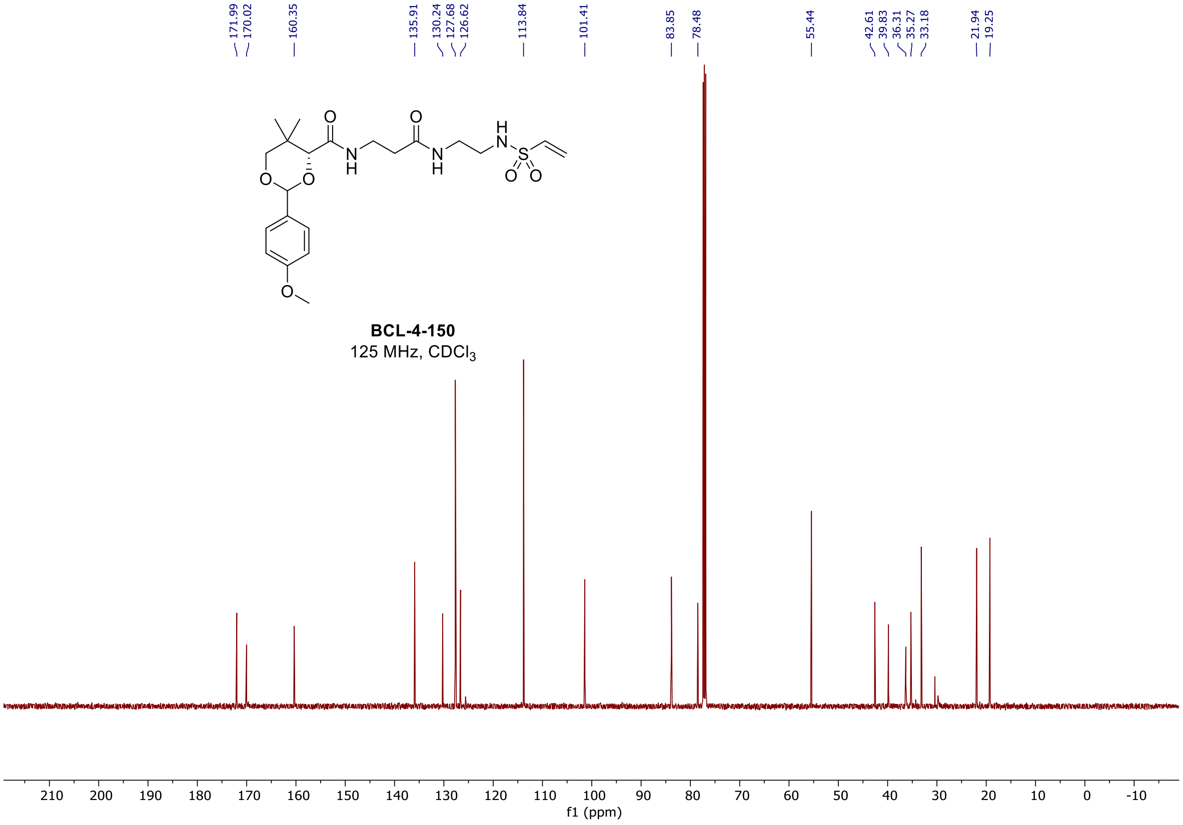

### $^{1}H$ and $^{13}C$ spectra of **BCL-4-154**

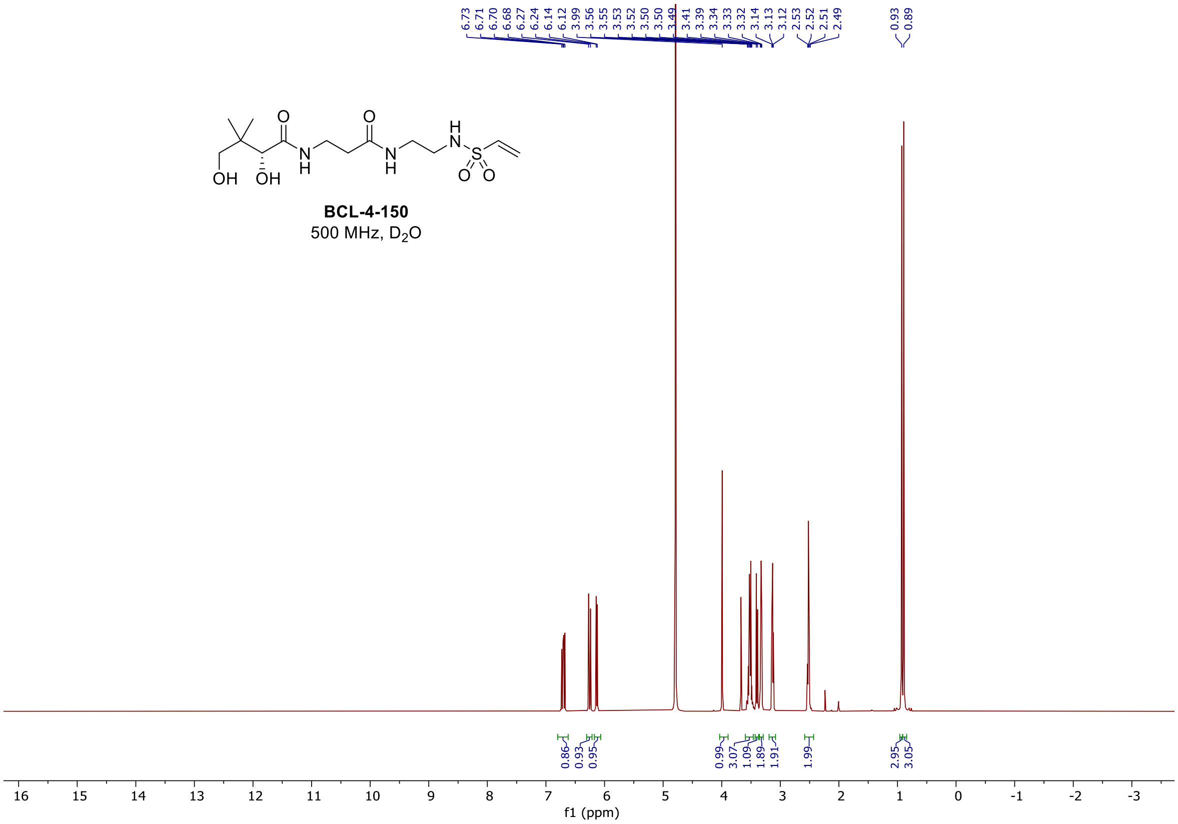

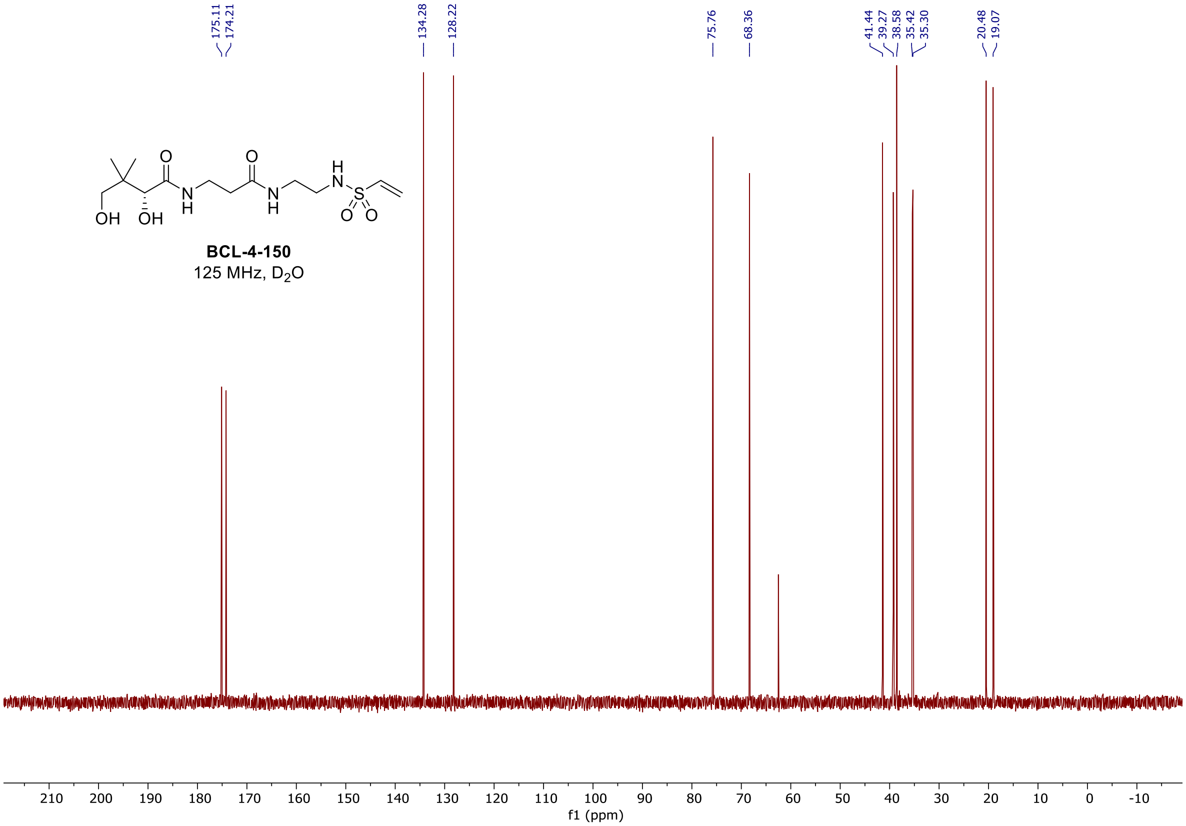

[1]H and [13]C spectra of **BCL-5-2**

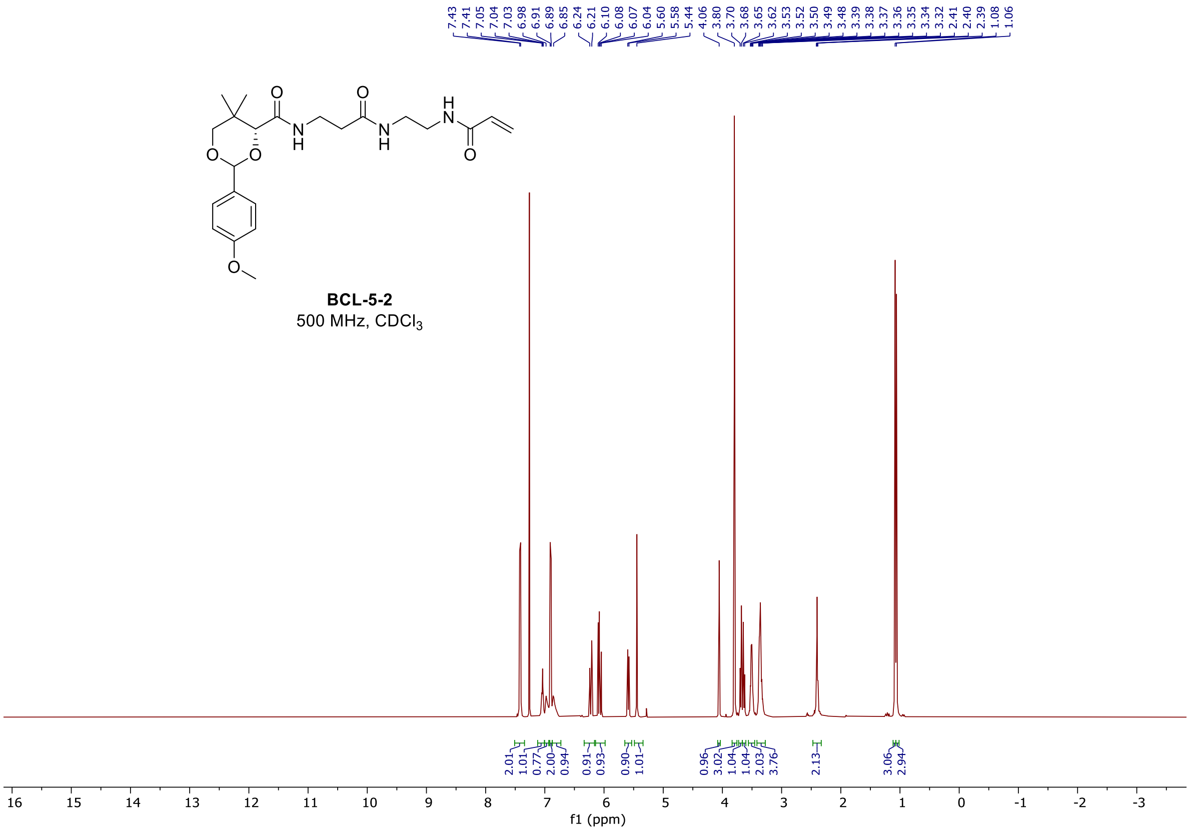

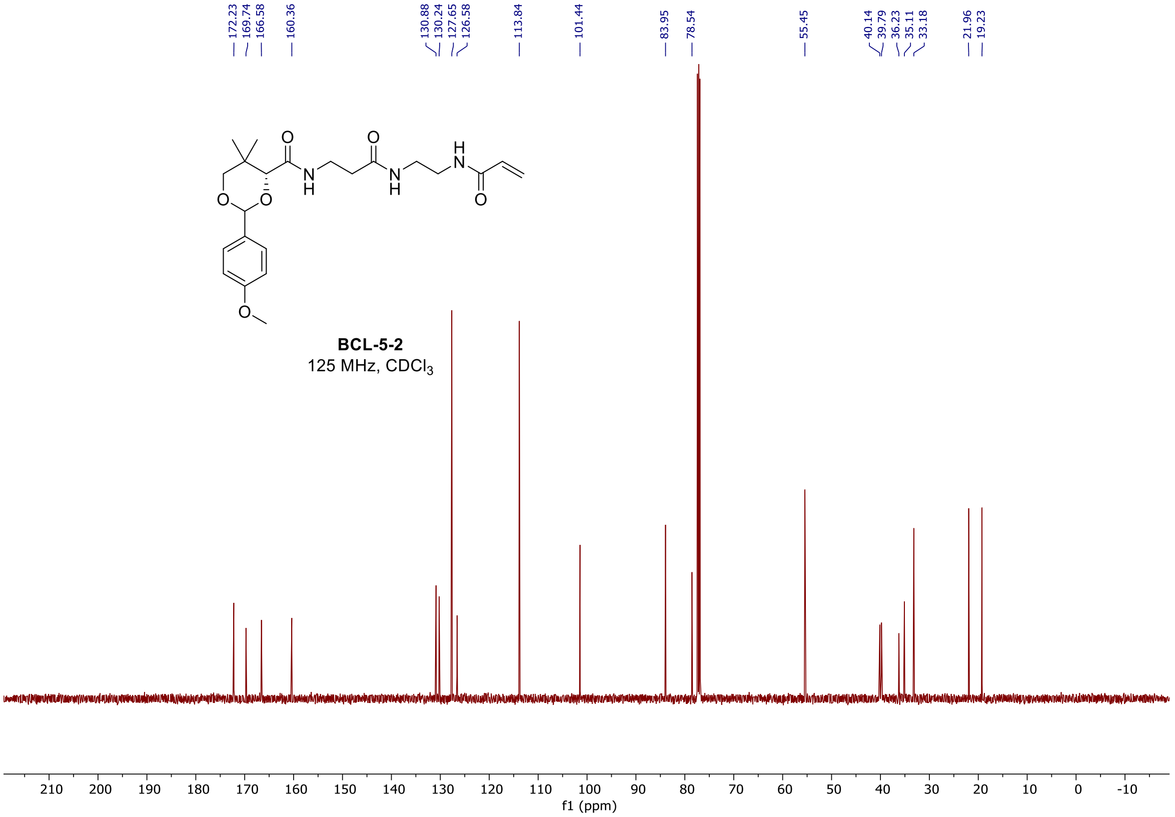

$^1H$ and $^{13}C$ spectra of **BCL-5-9**

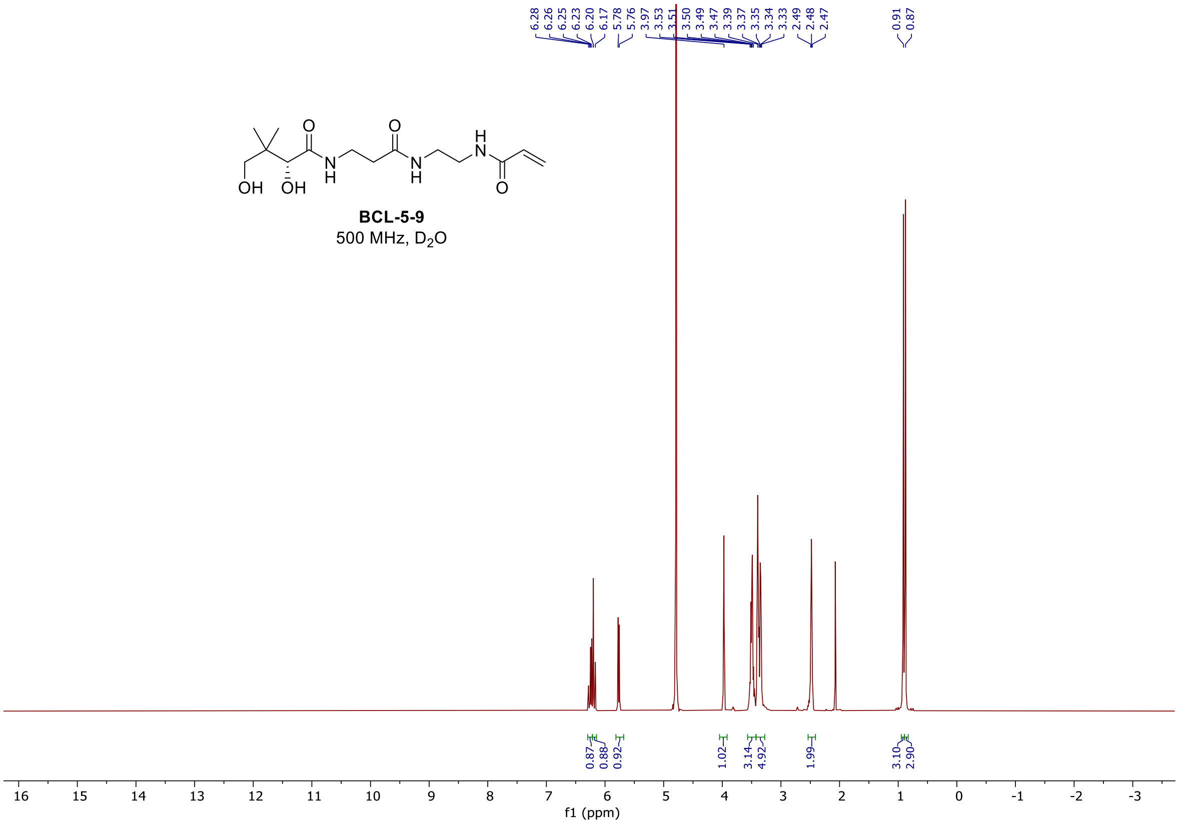

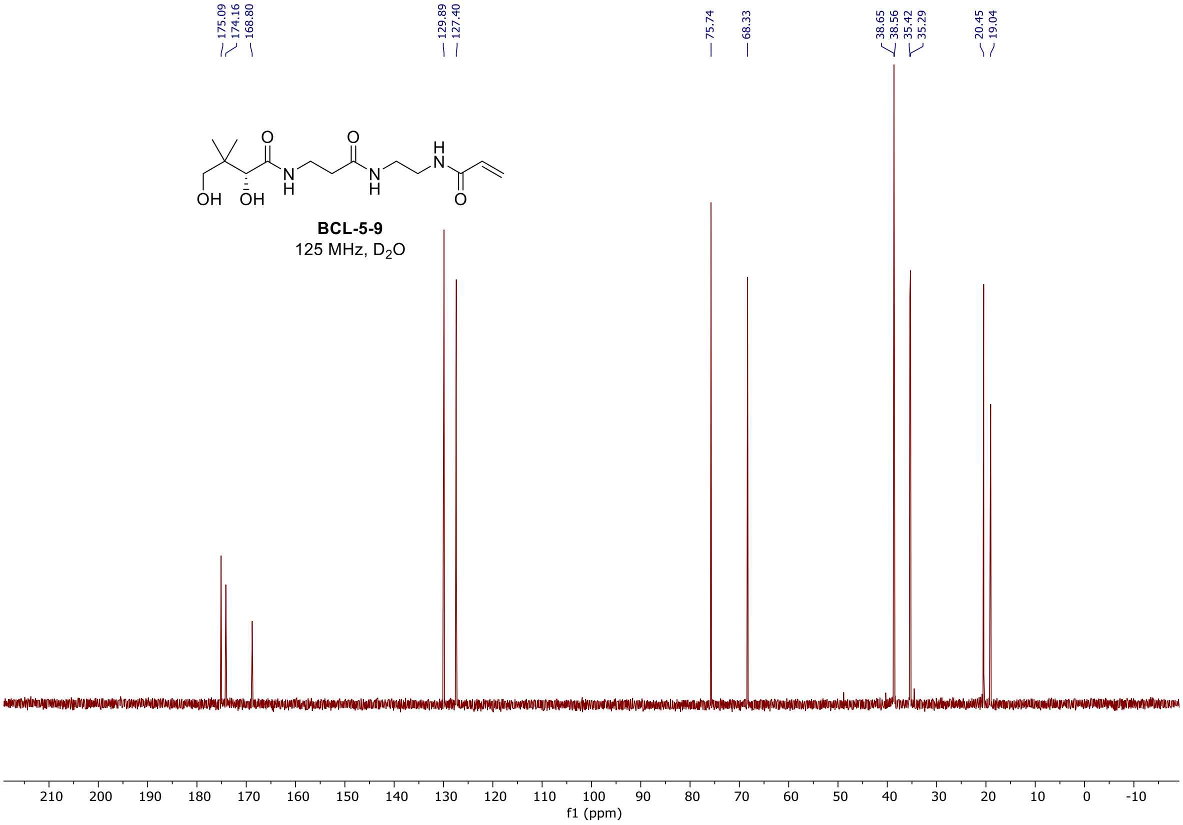

[1]H and [13]C spectra of **BCL-5-18**

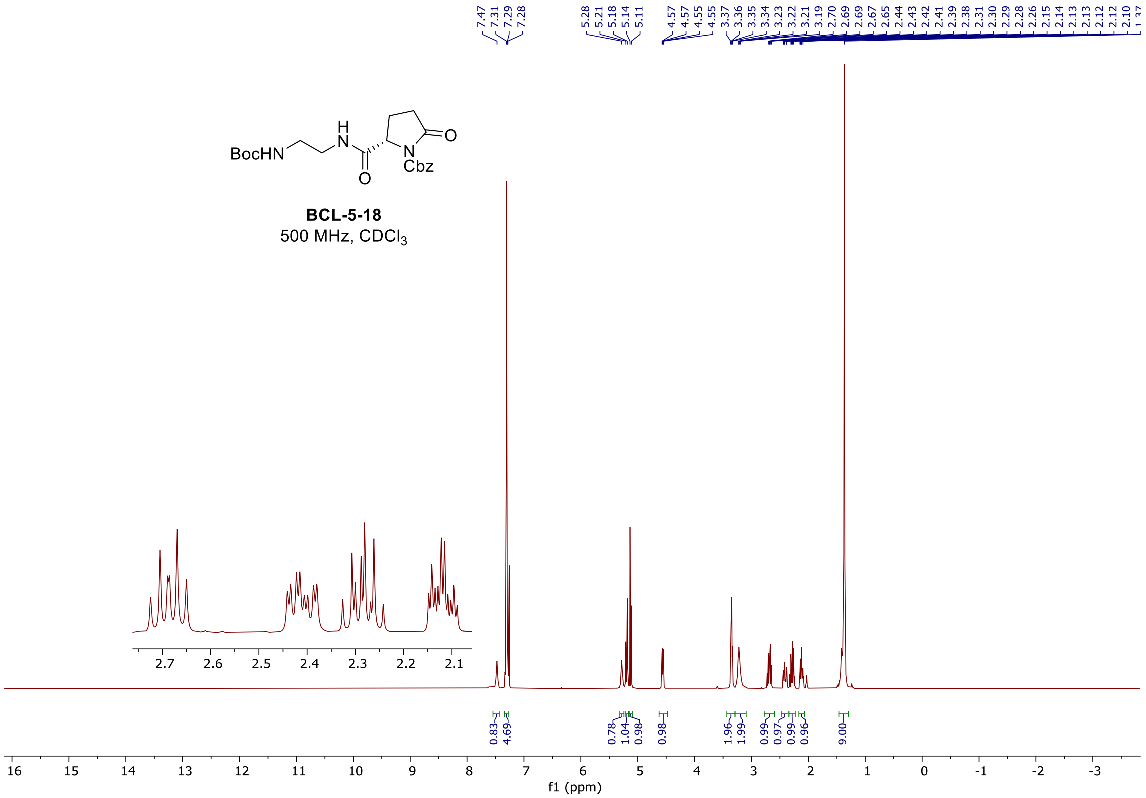

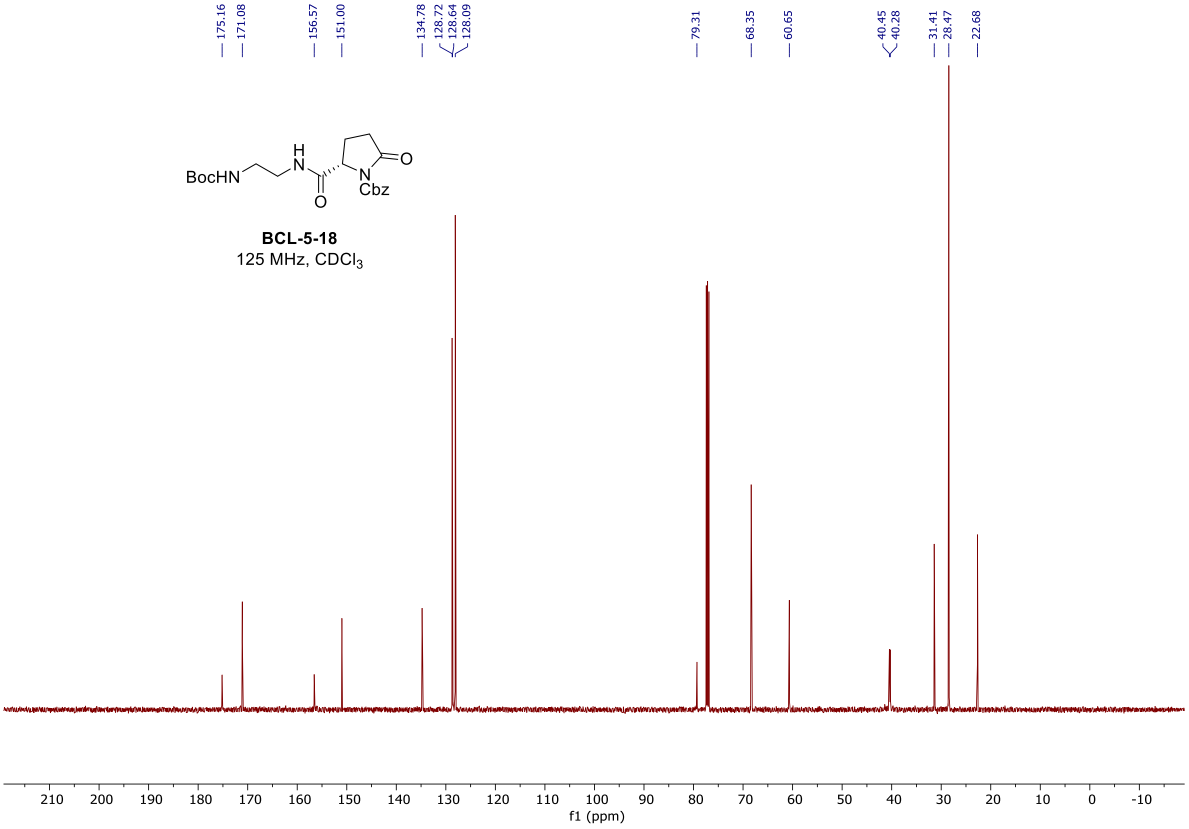

<sup></sup>

$^{1}$H and $^{13}$C spectra of **BCL-5-23**

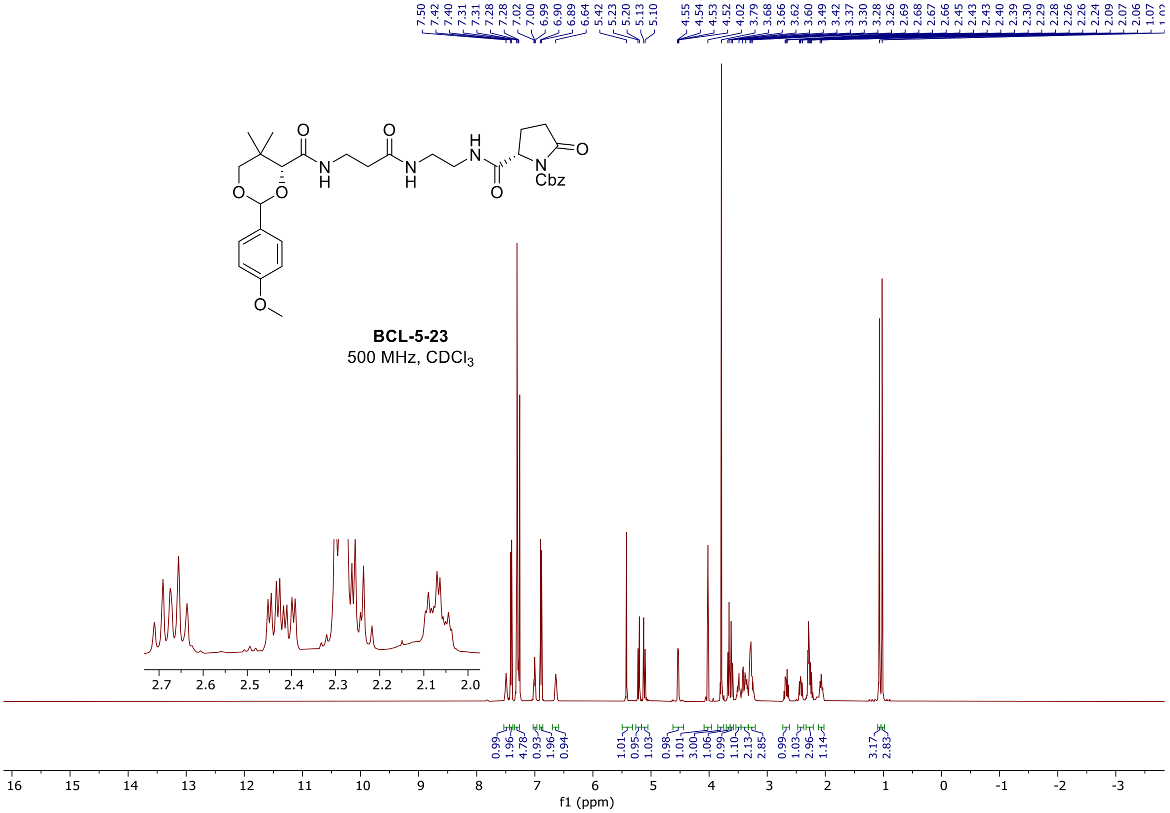

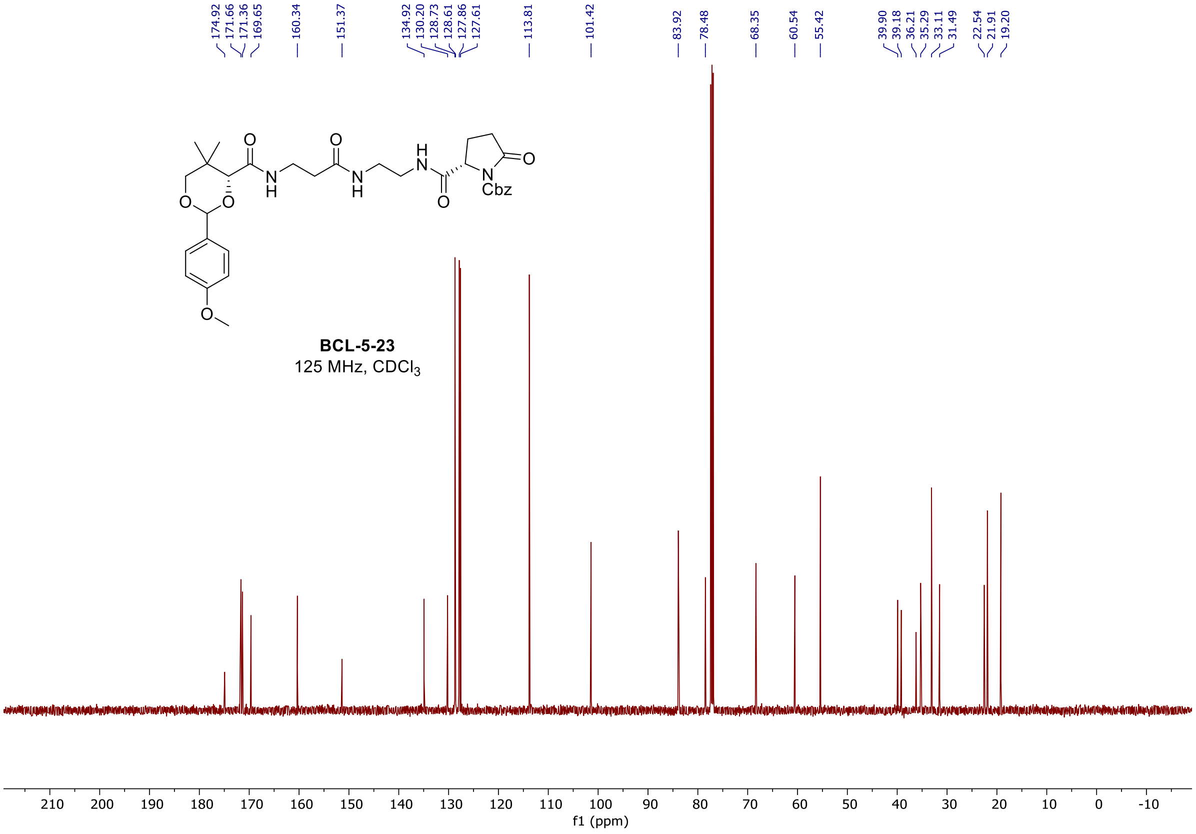

[1]H and [13]C spectra of **BCL-5-29**

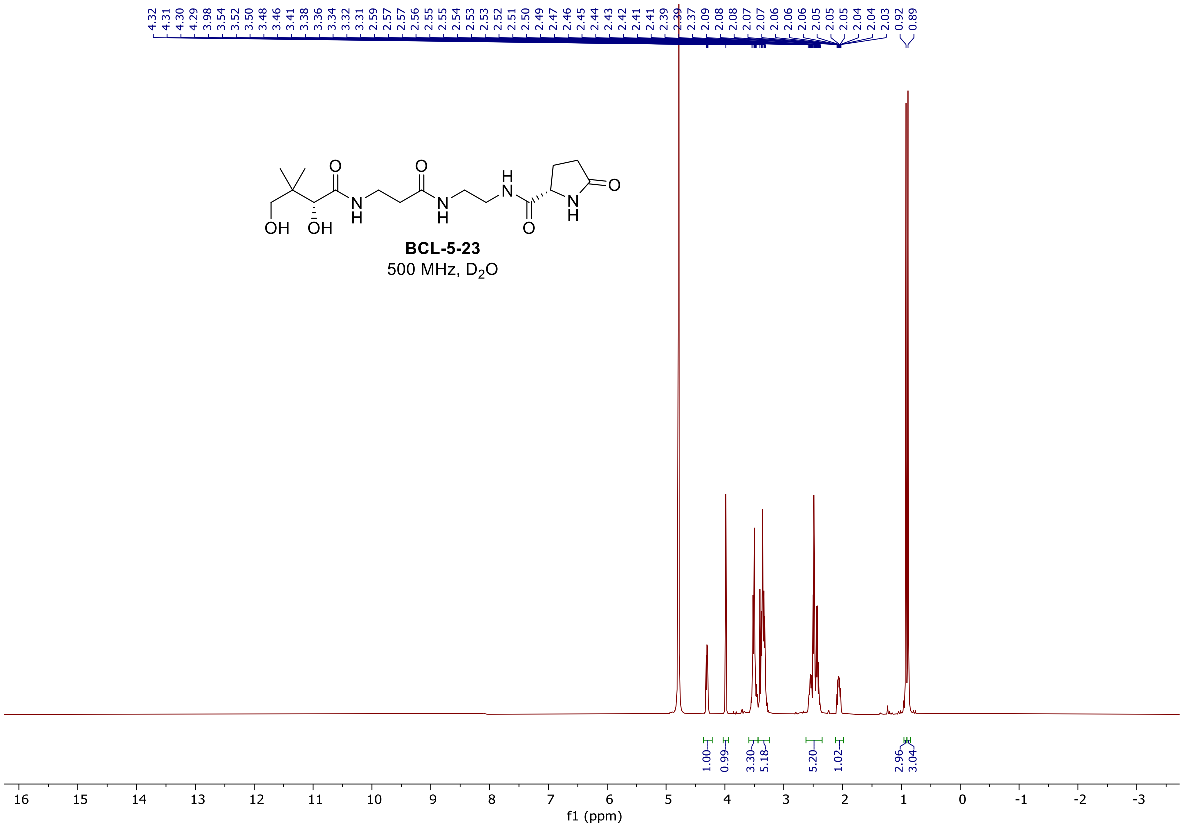

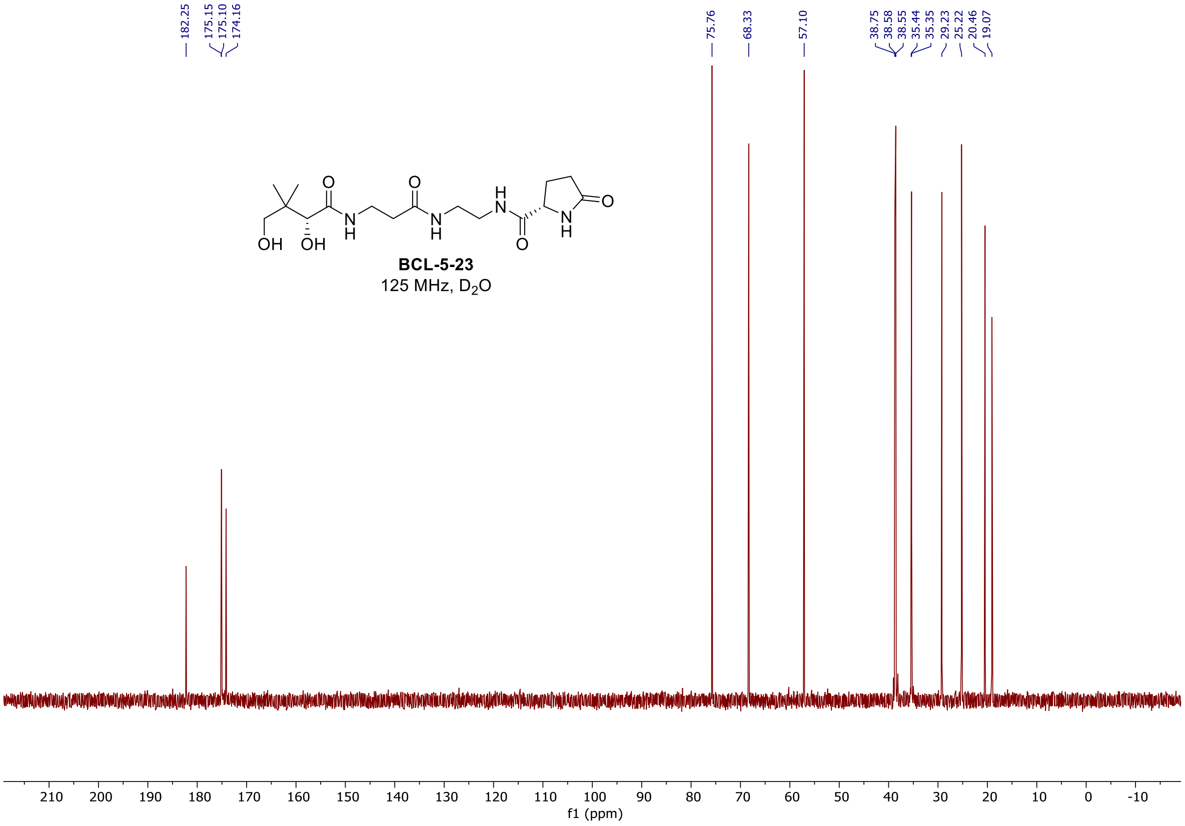

<sup></sup>

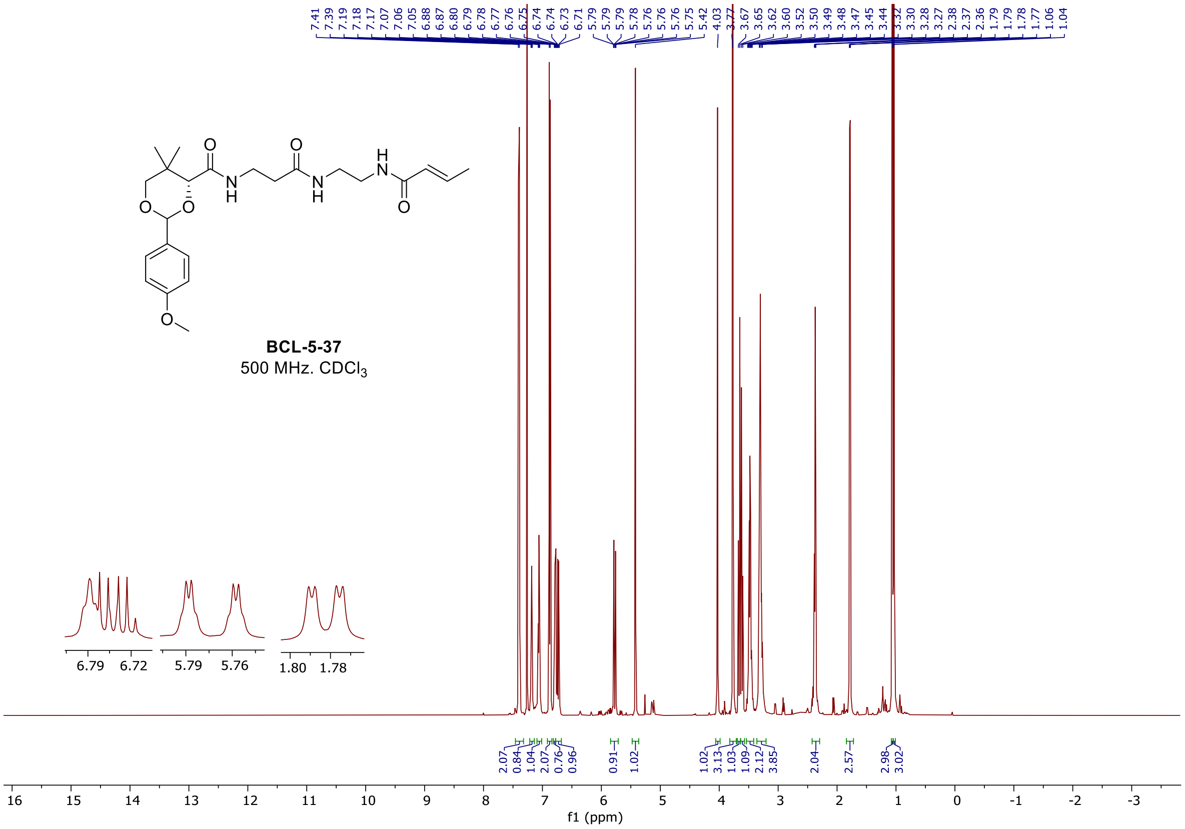

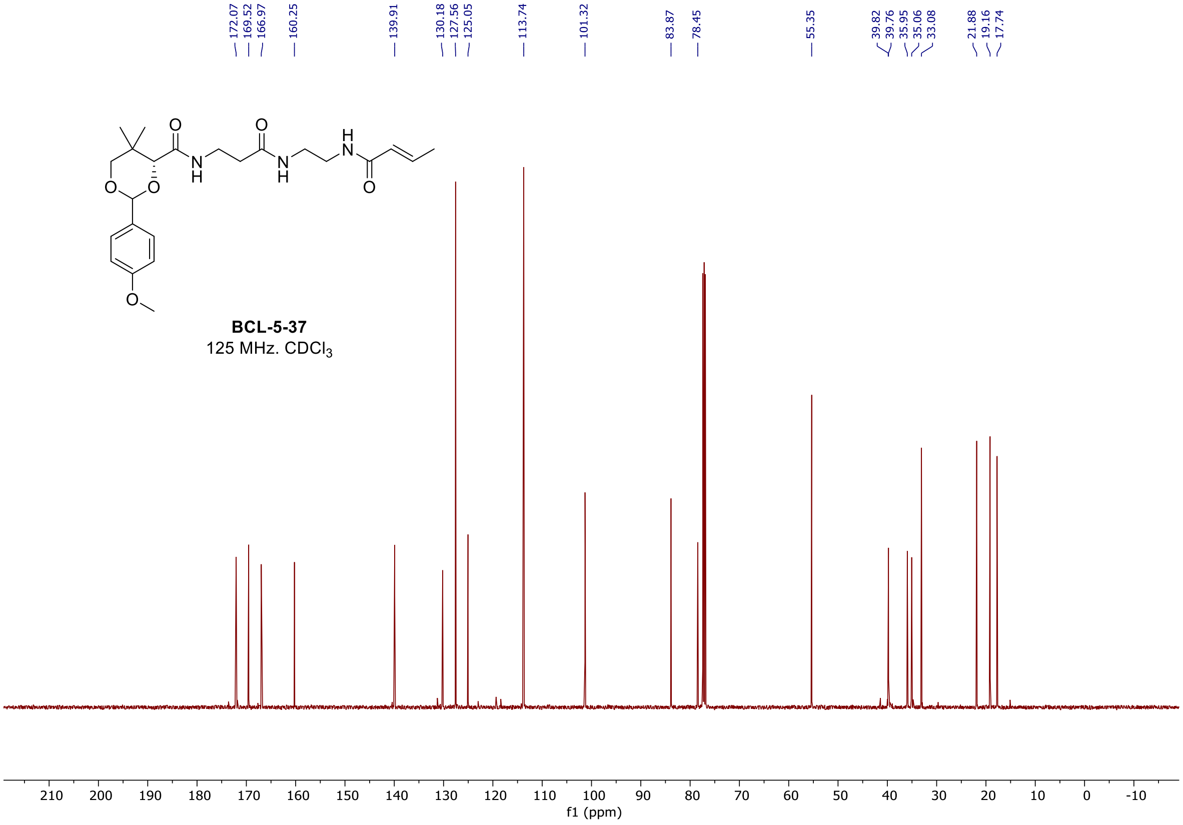

[1]H and [13]C spectra of **BCL-5-41**

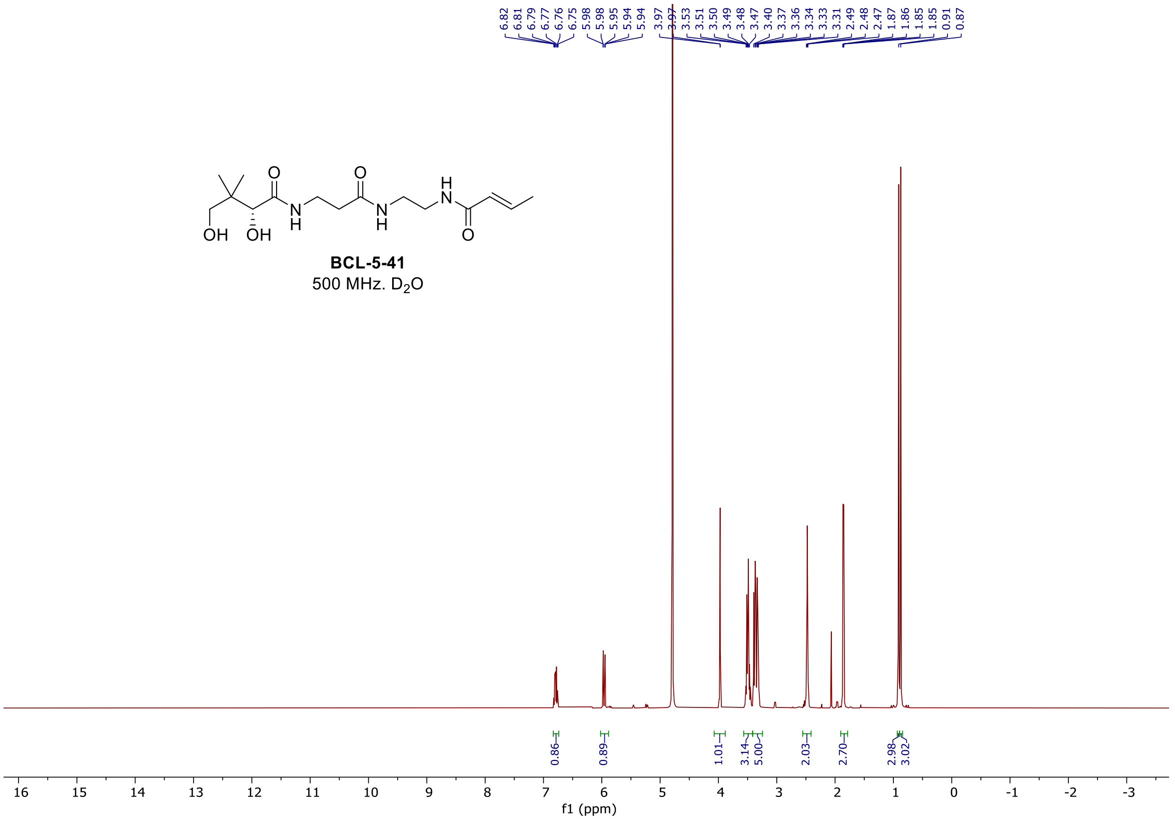

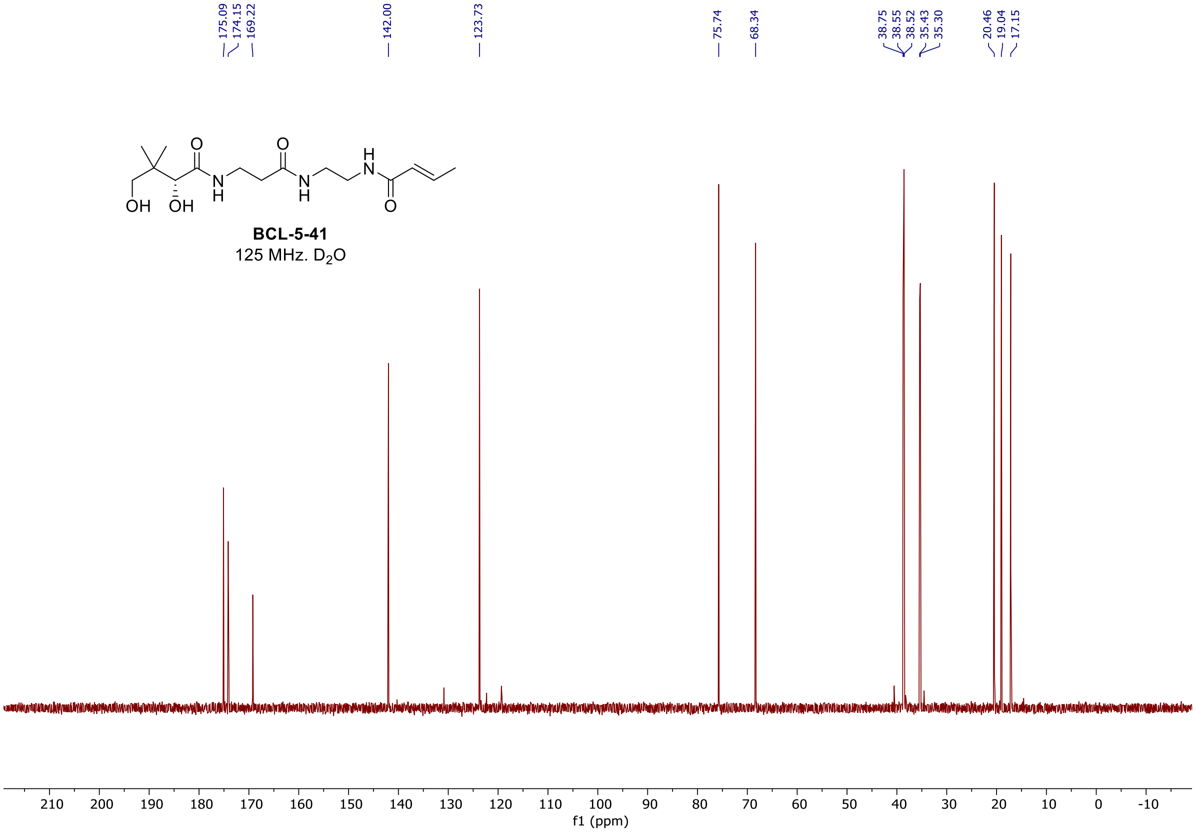

[1]H, [19]F, and [13]C spectra of **BCL-5-43**

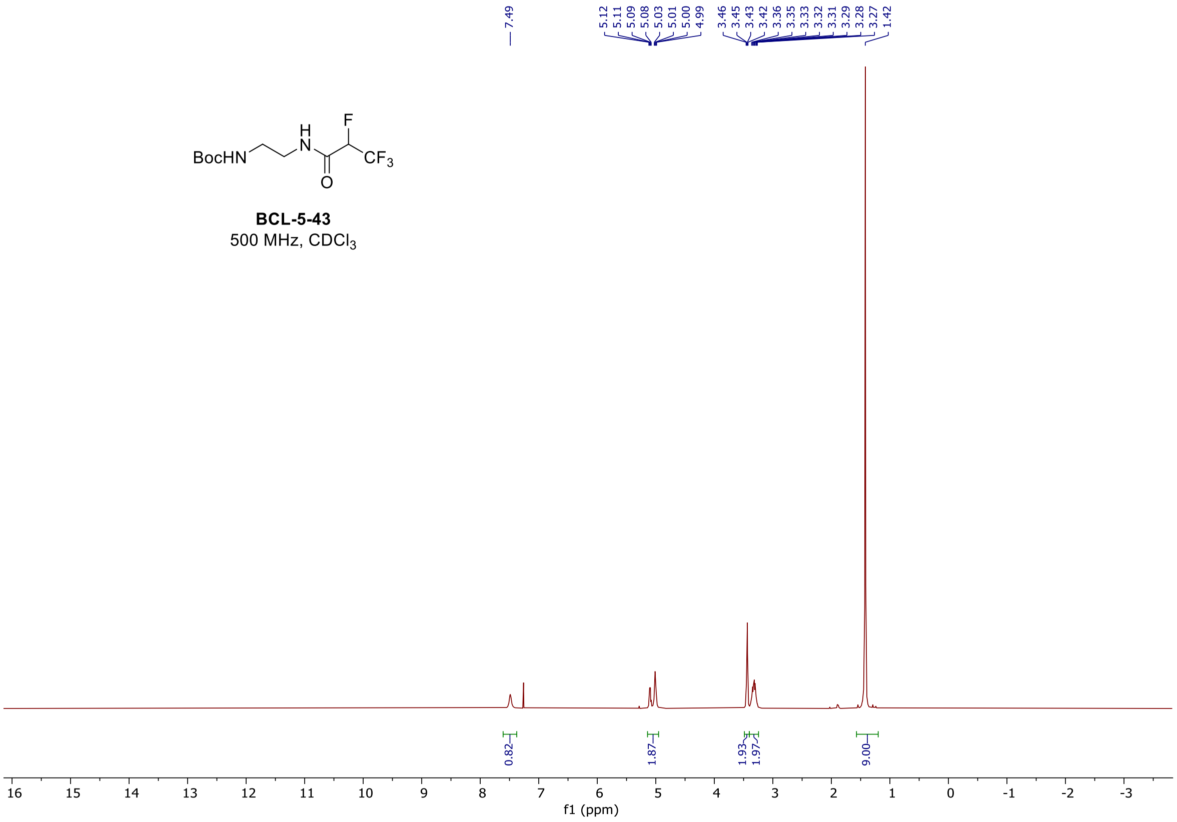

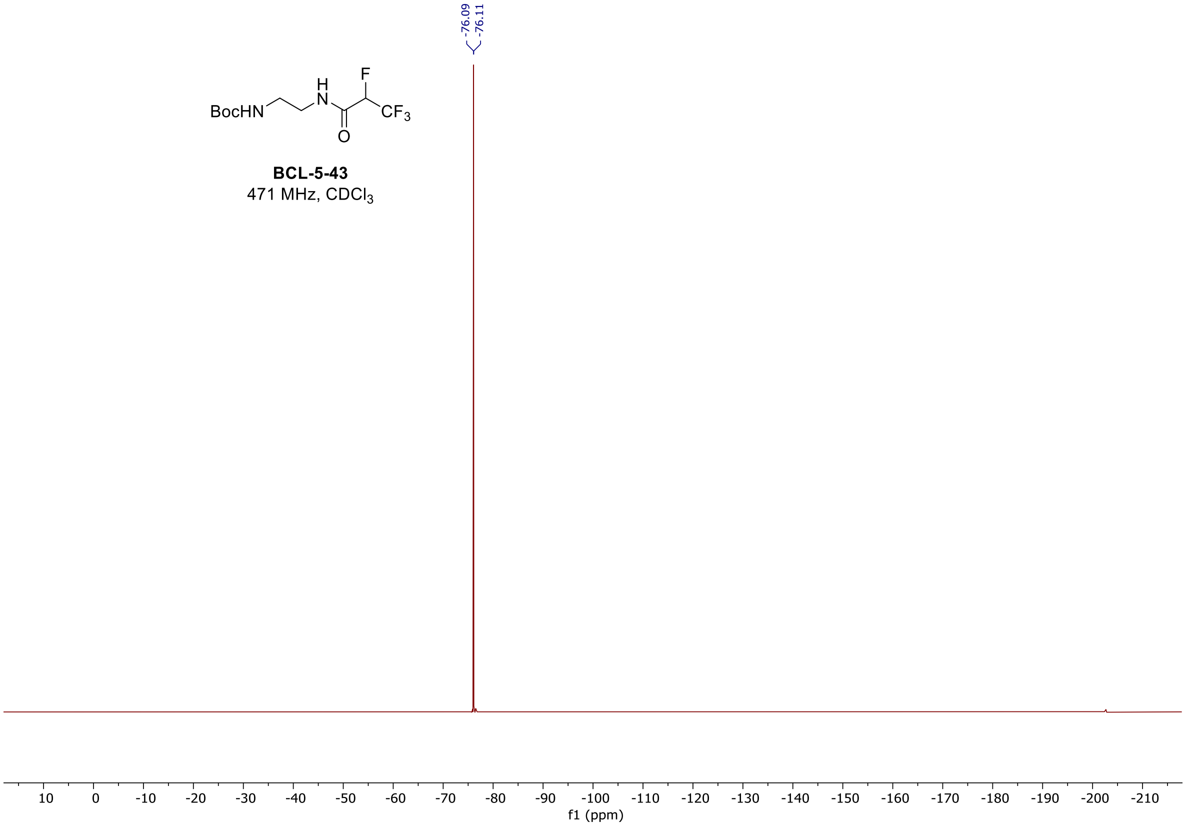

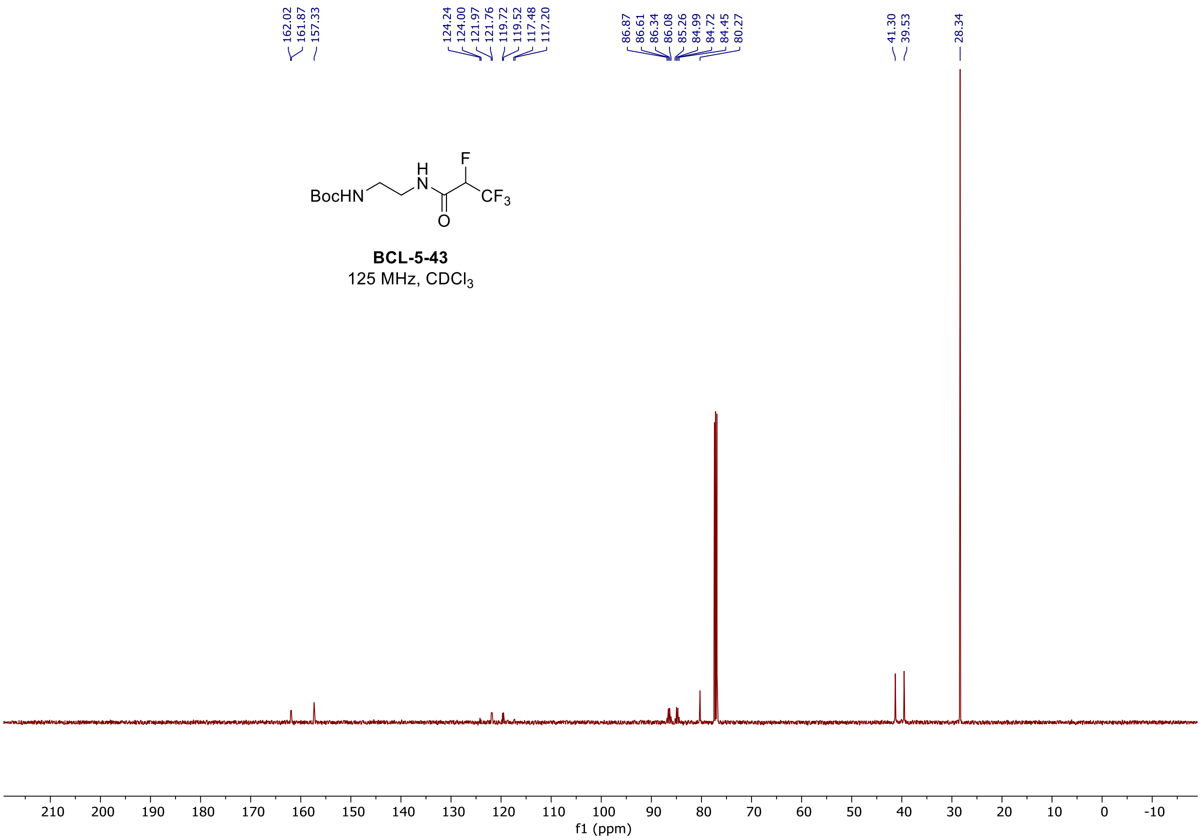

#### $^{1}H$, $^{19}F$, and $^{13}C$ spectra of **BCL-5-48**

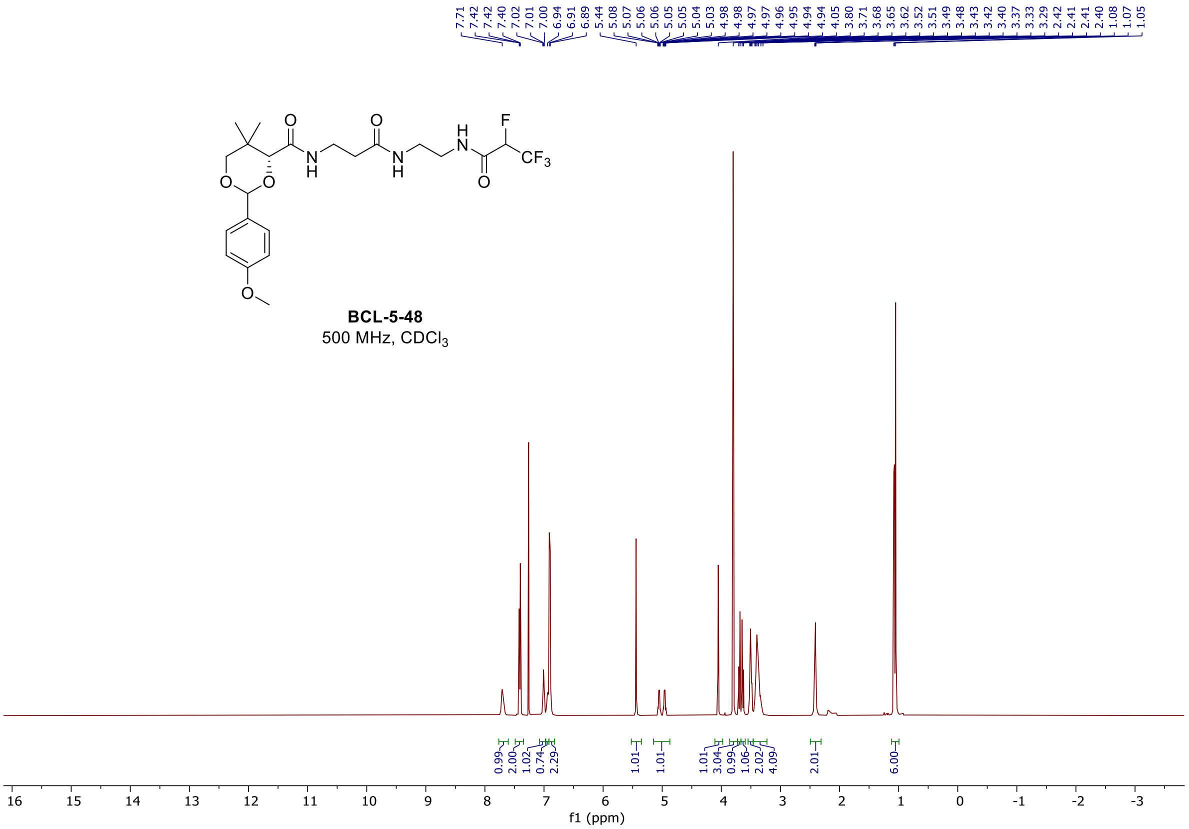

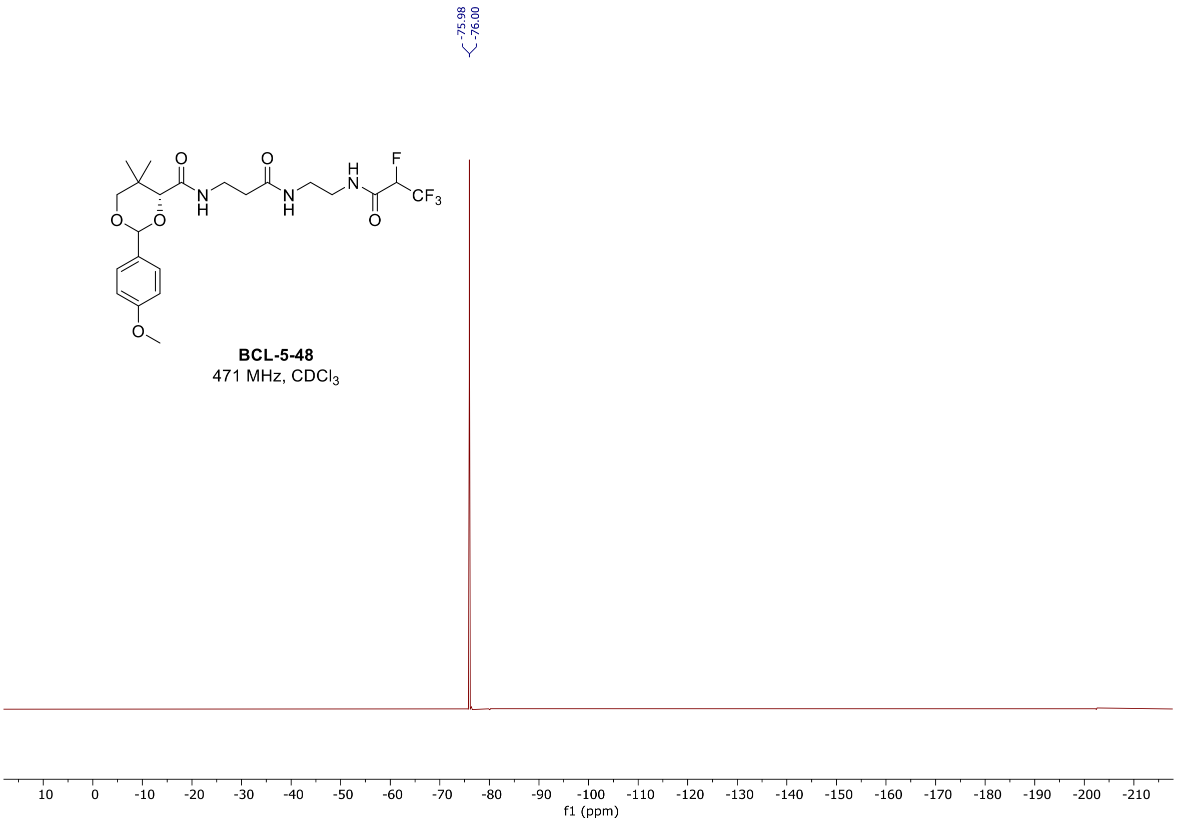
-75.98
-76.00
F
O
O
H
N
N
H
N
H
CF3
O
O
O
O
BCL-5-48
471 MHz, CDCl3
f1 (ppm)

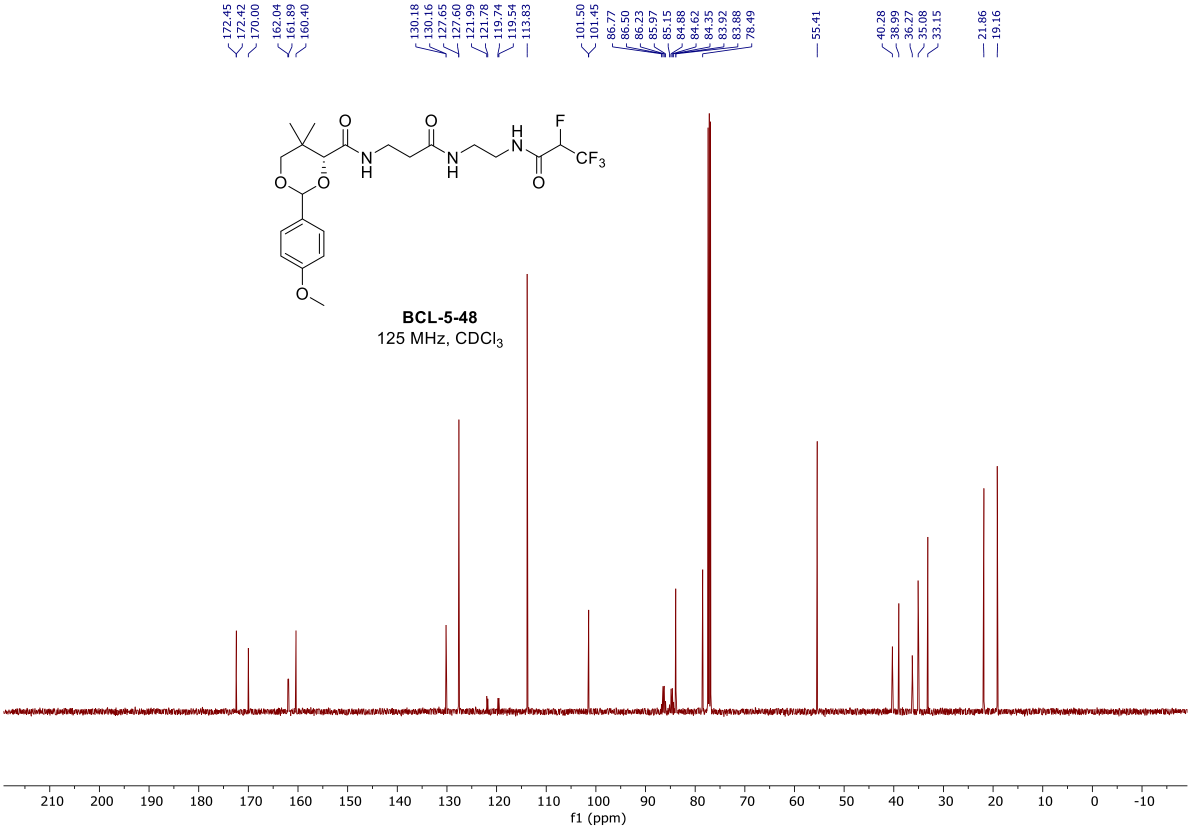
172.45
172.42
170.00
162.04
161.89
160.40
130.18
130.16
127.65
127.60
121.99
121.78
119.74
119.54
113.83
101.50
101.45
86.77
86.50
86.23
85.97
85.15
84.88
84.62
84.35
83.92
83.88
78.49
55.41
40.28
38.99
36.27
35.08
33.15
21.86
19.16
BCL-5-48
125 MHz, CDCl3
f1 (ppm)

$^{1}$H, $^{19}$F, and $^{13}$C spectra of **BCL-5-54**

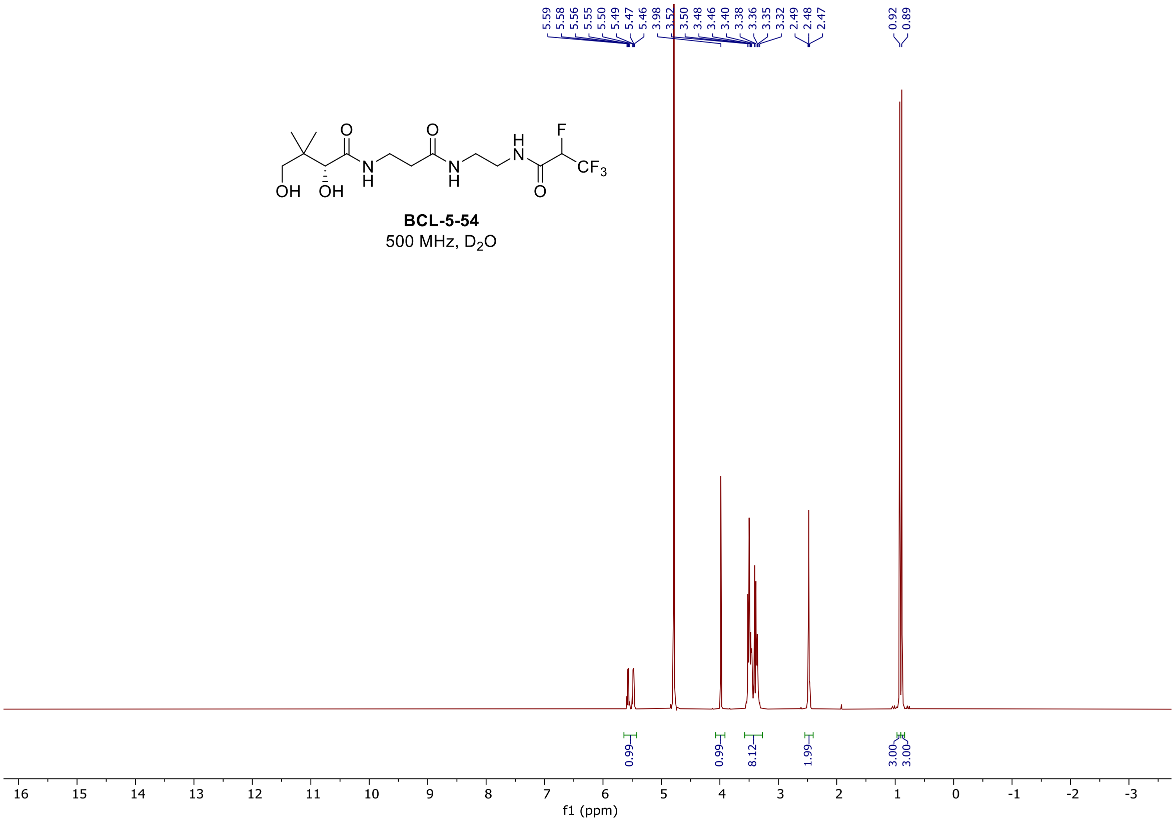

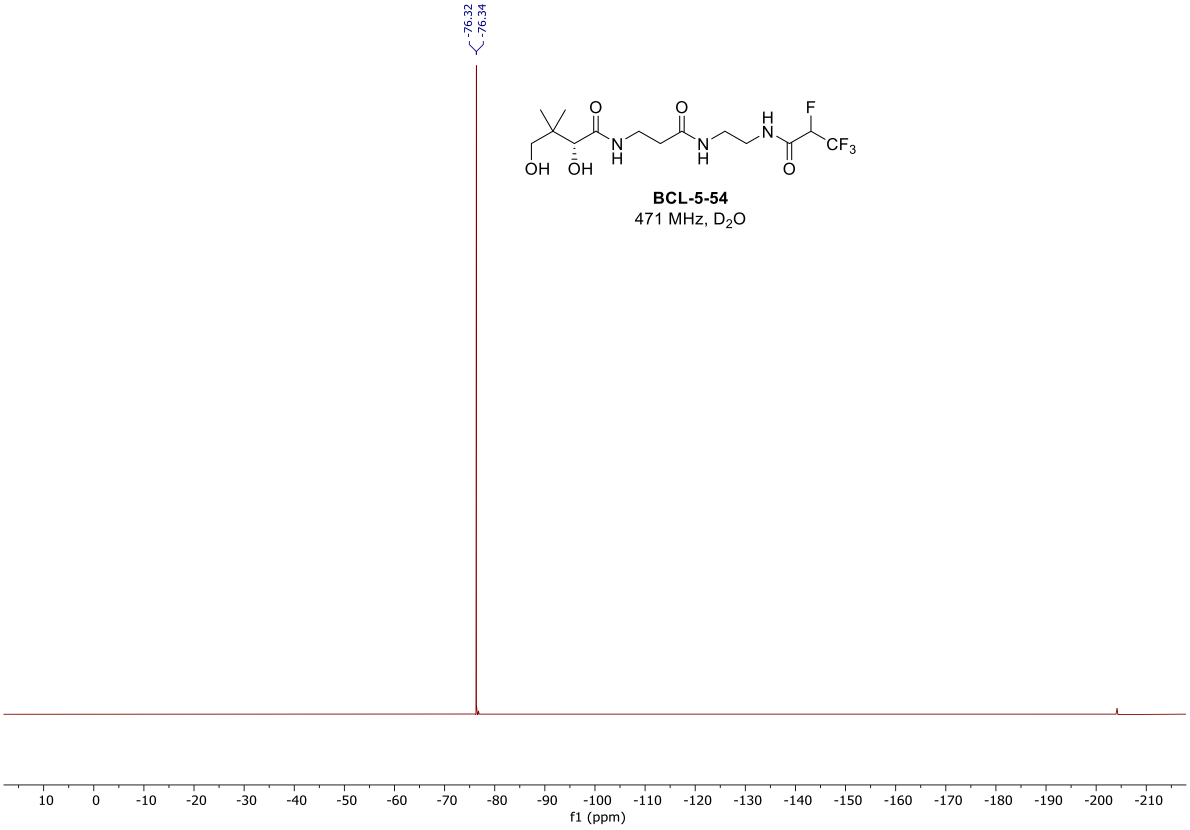

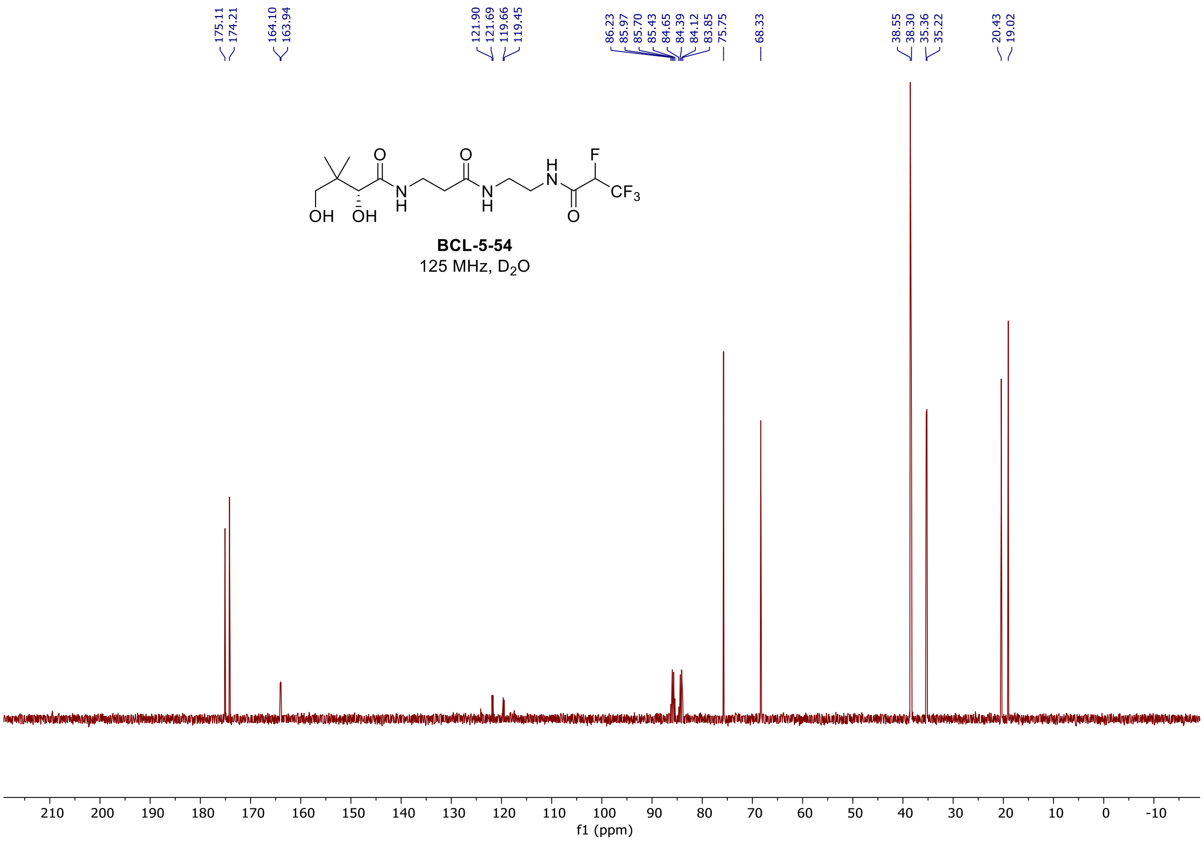

#### [1]H and [13]C spectra of **BCL-5-63**

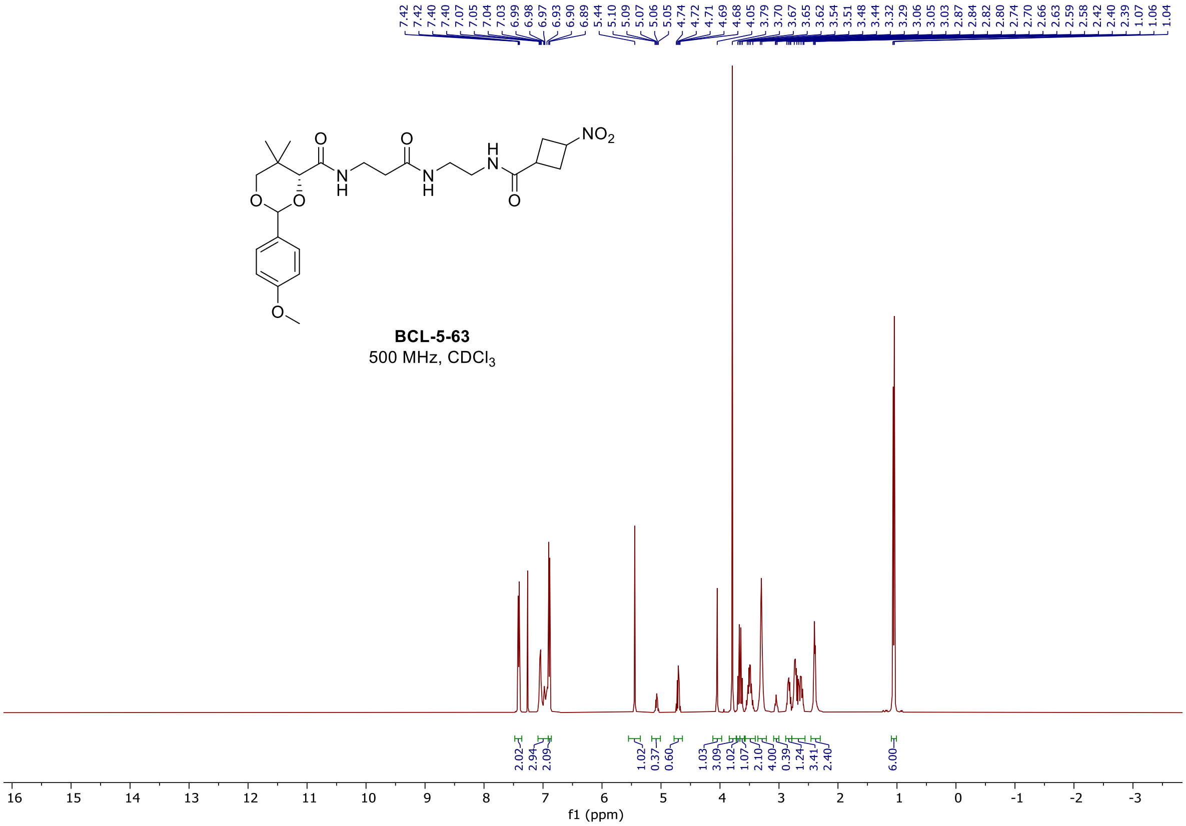

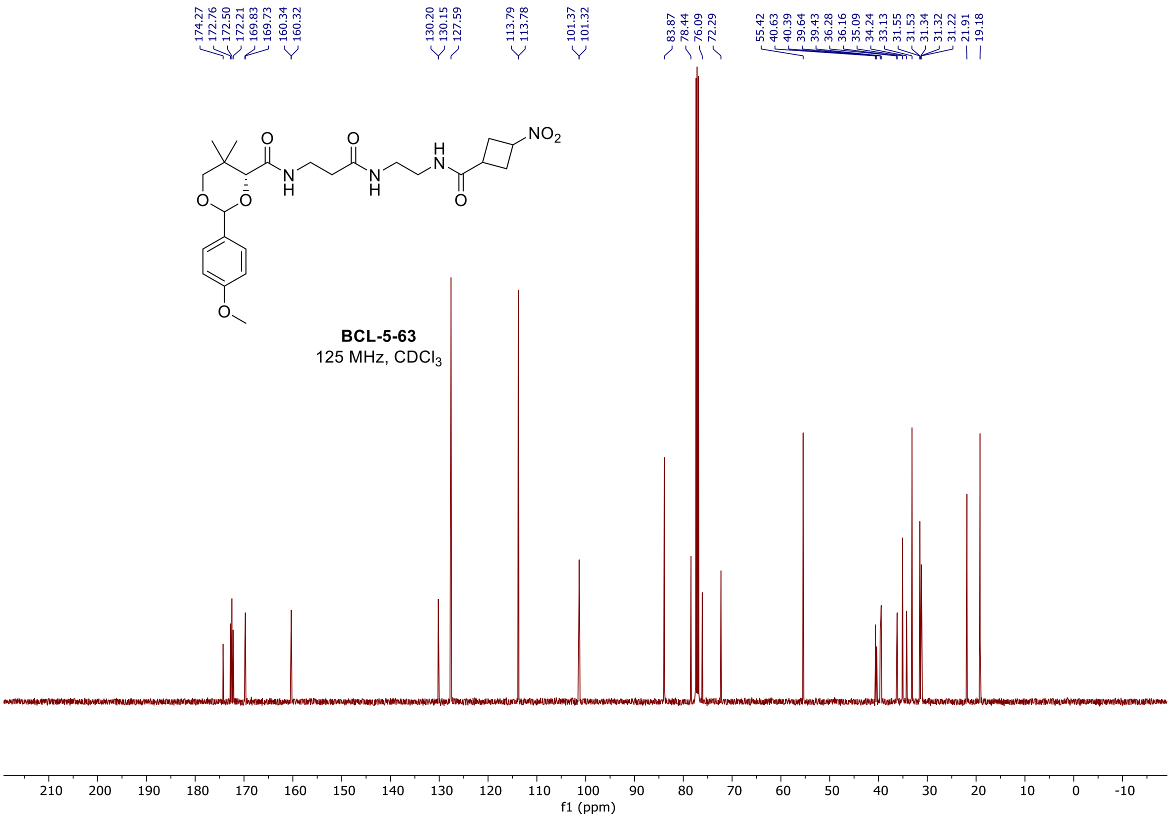

$^1H$ and $^{13}C$ spectra of **BCL-5-67**

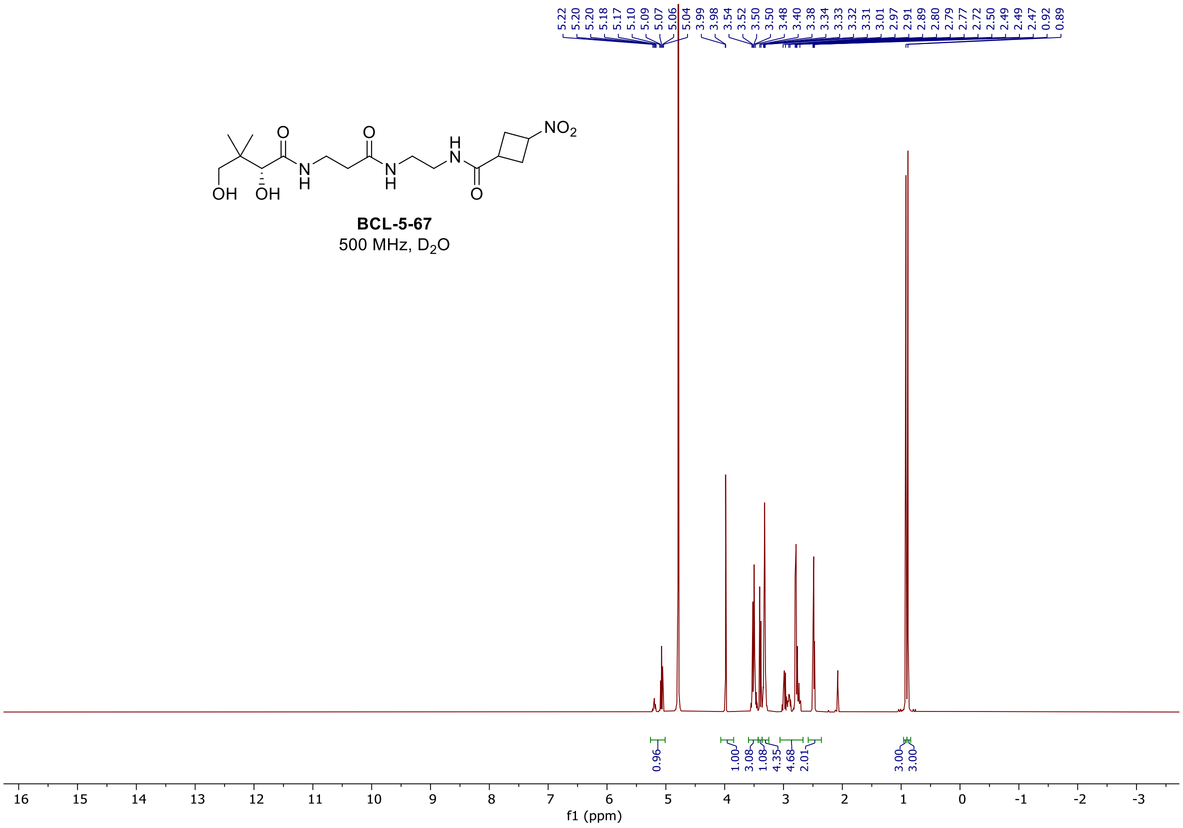

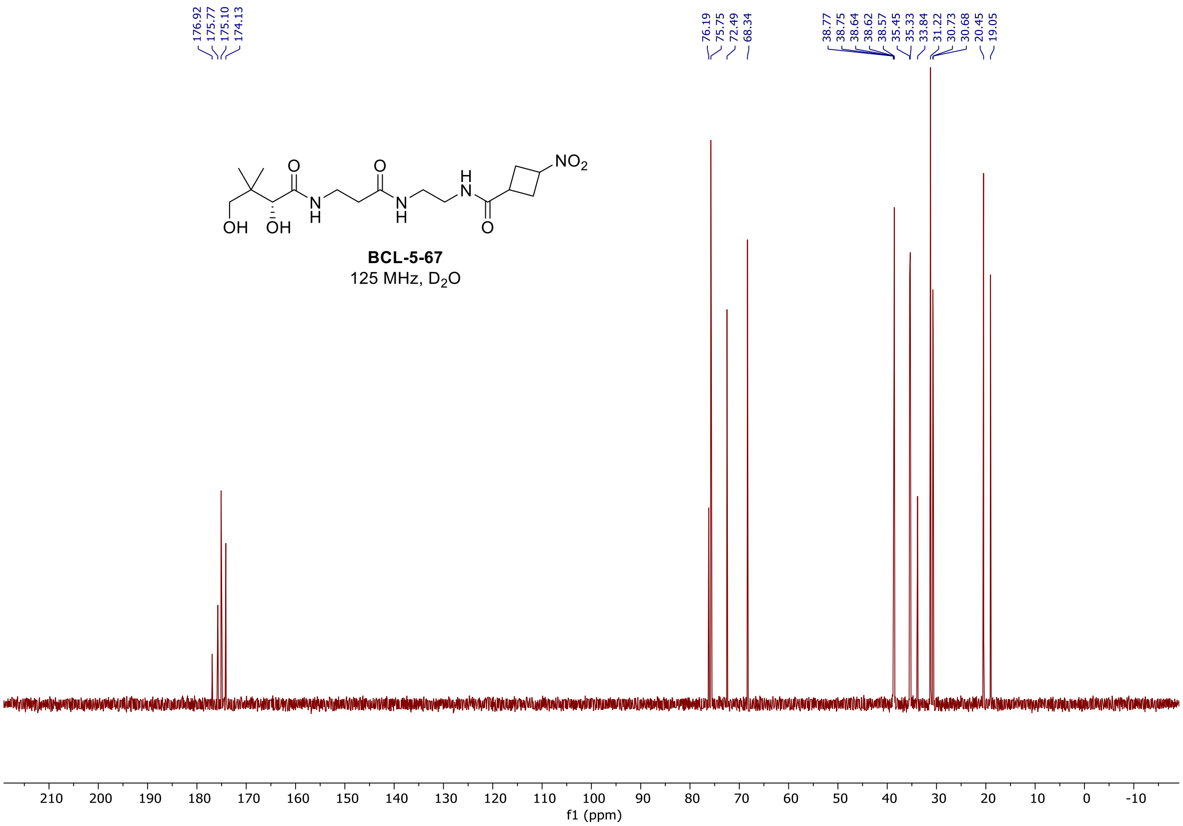

<u>$^{1}$H and $^{13}$C spectra of **BCL-7-1**</u>

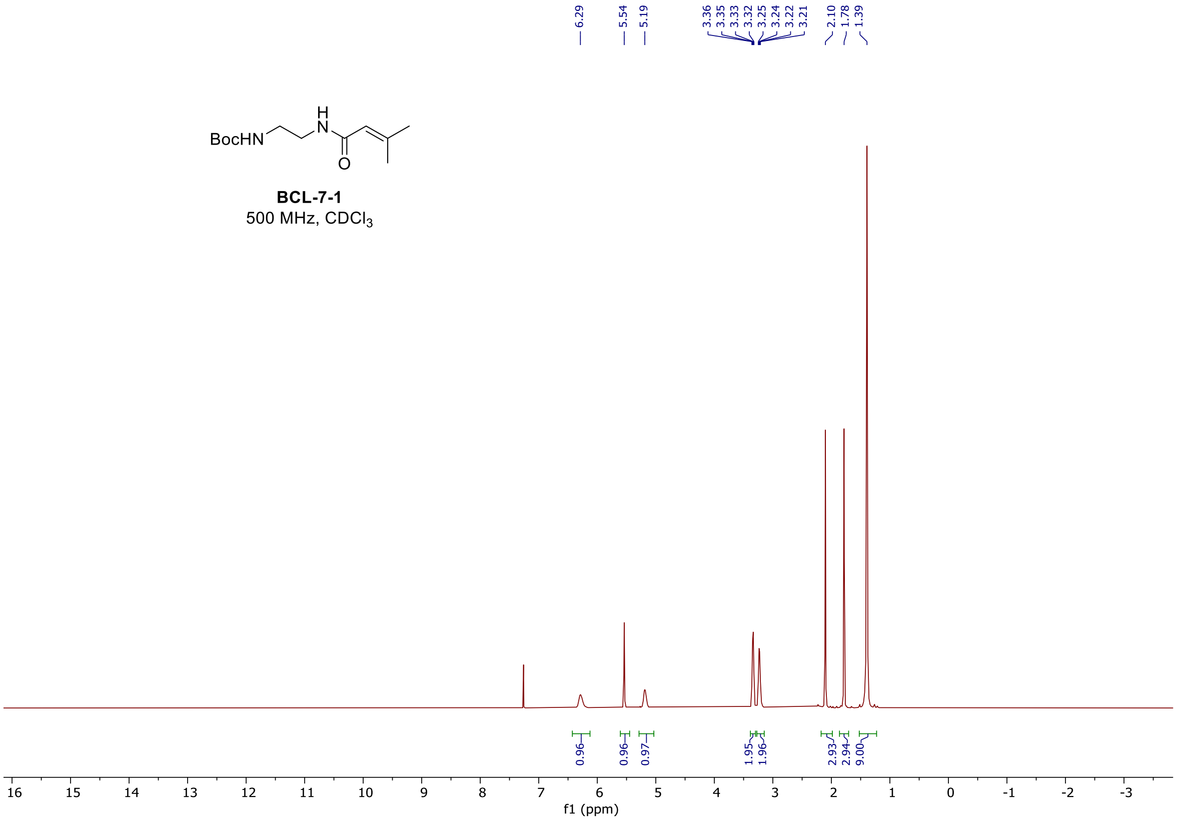

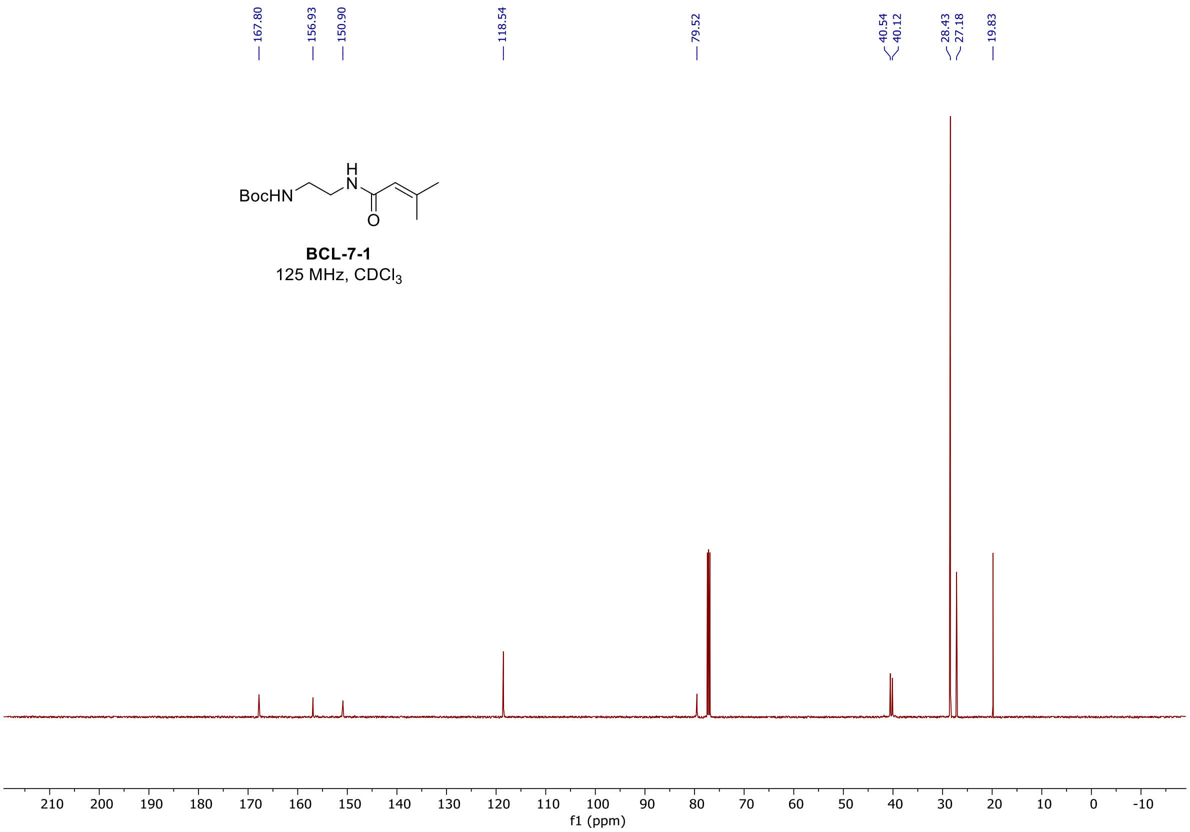

#### [1]H and [13]C spectra of **BCL-7-6**

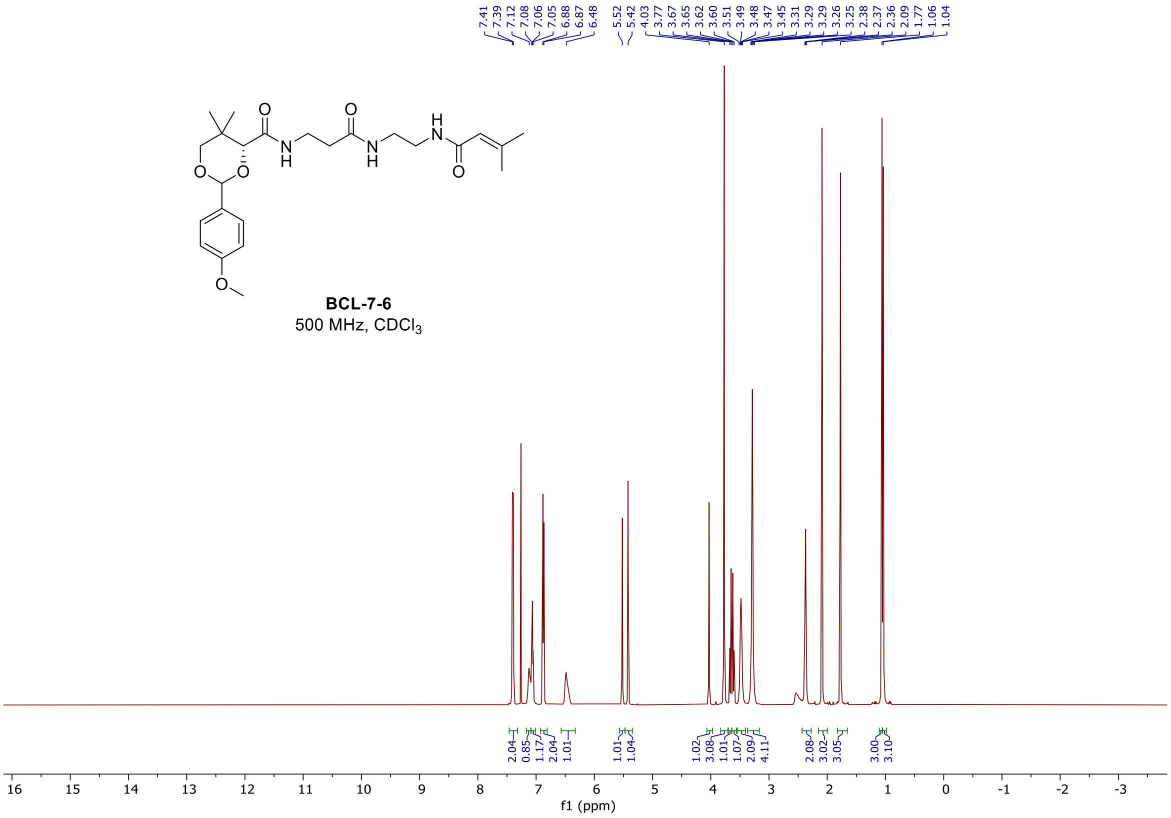

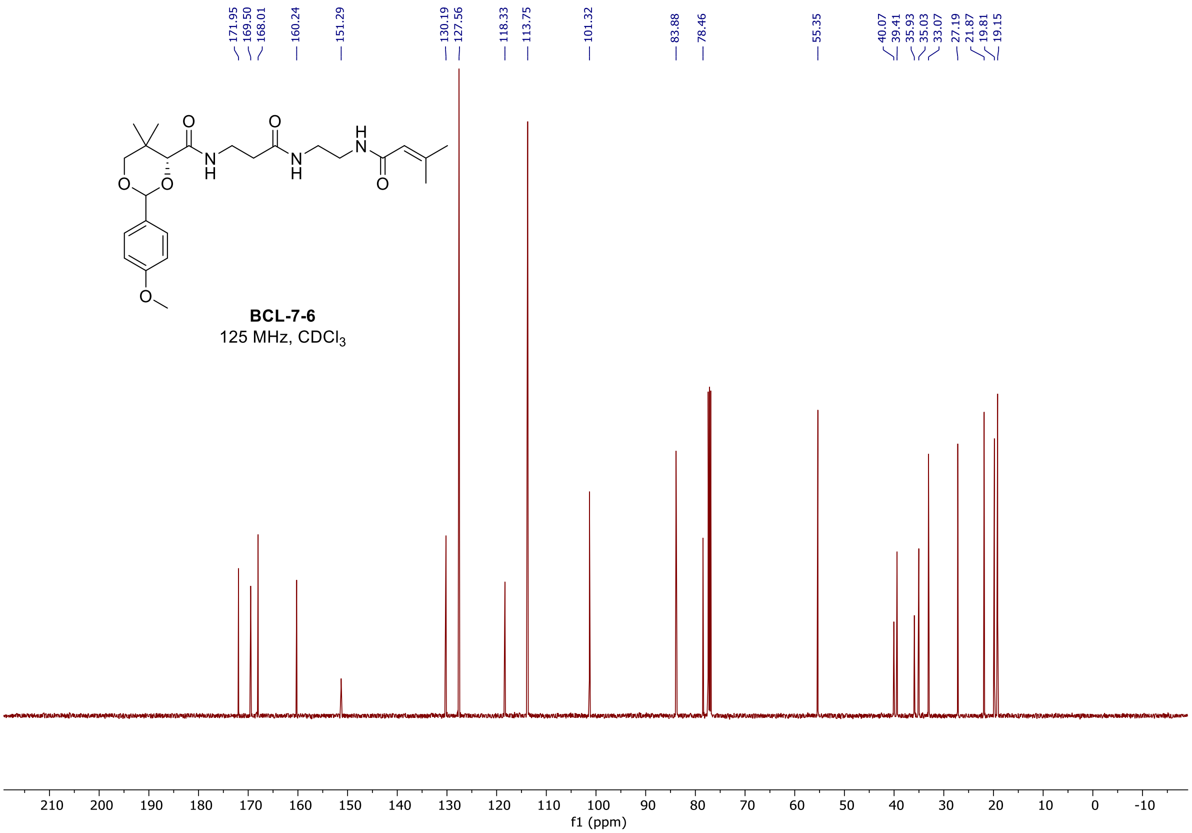

#### [1]H and [13]C spectra of **BCL-7-11**

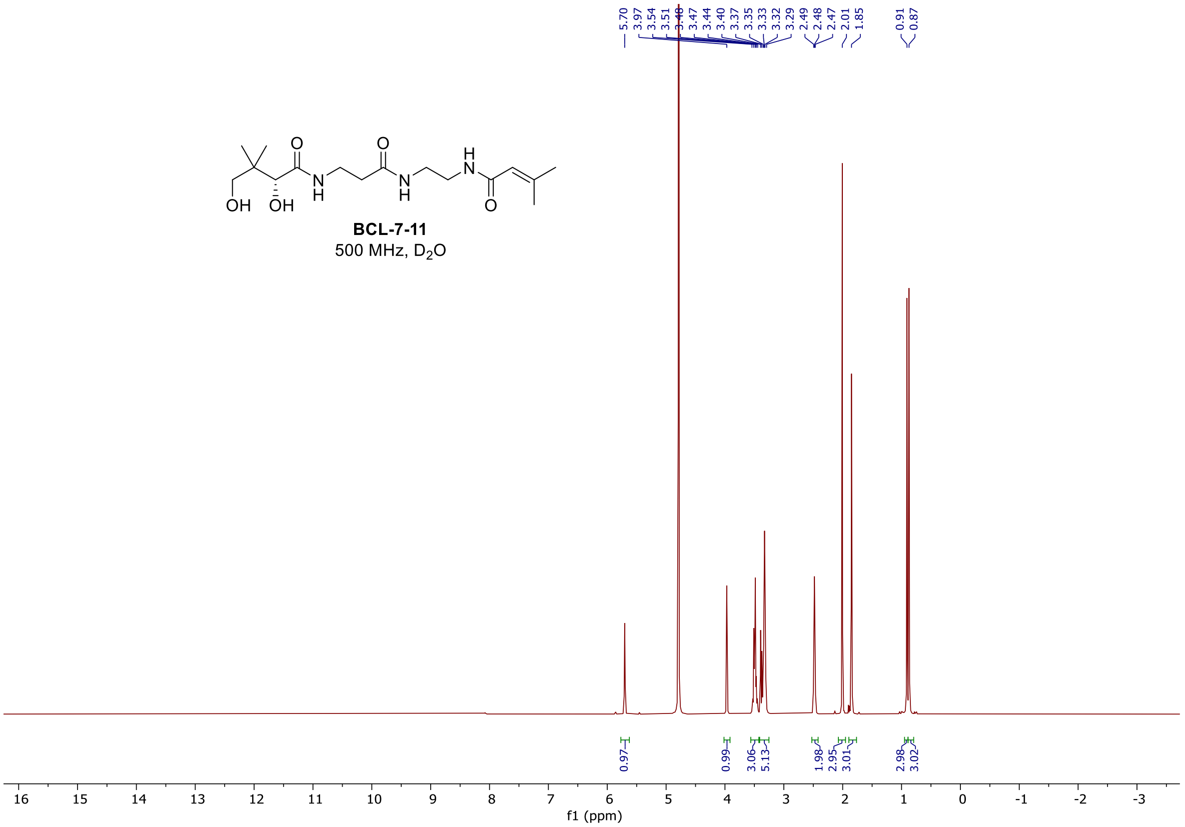

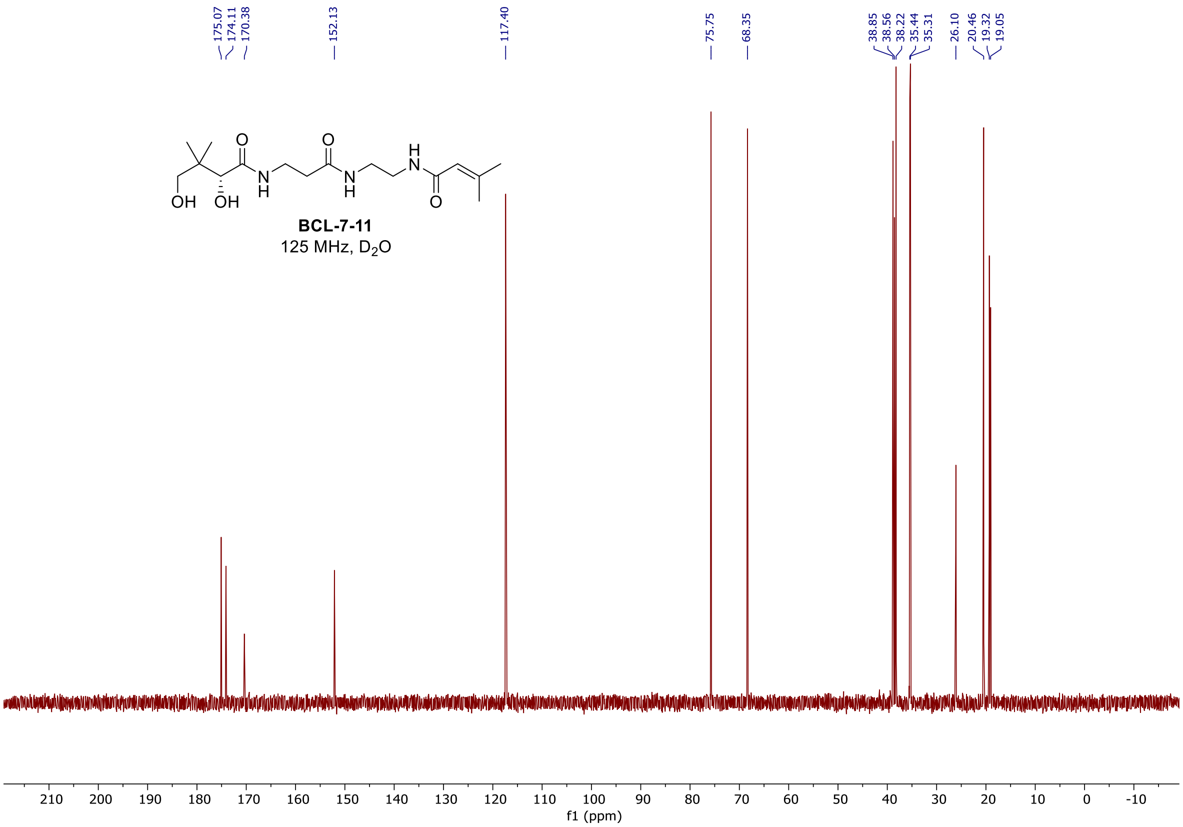

#### [1]H and [13]C spectra of **BCL-7-24**

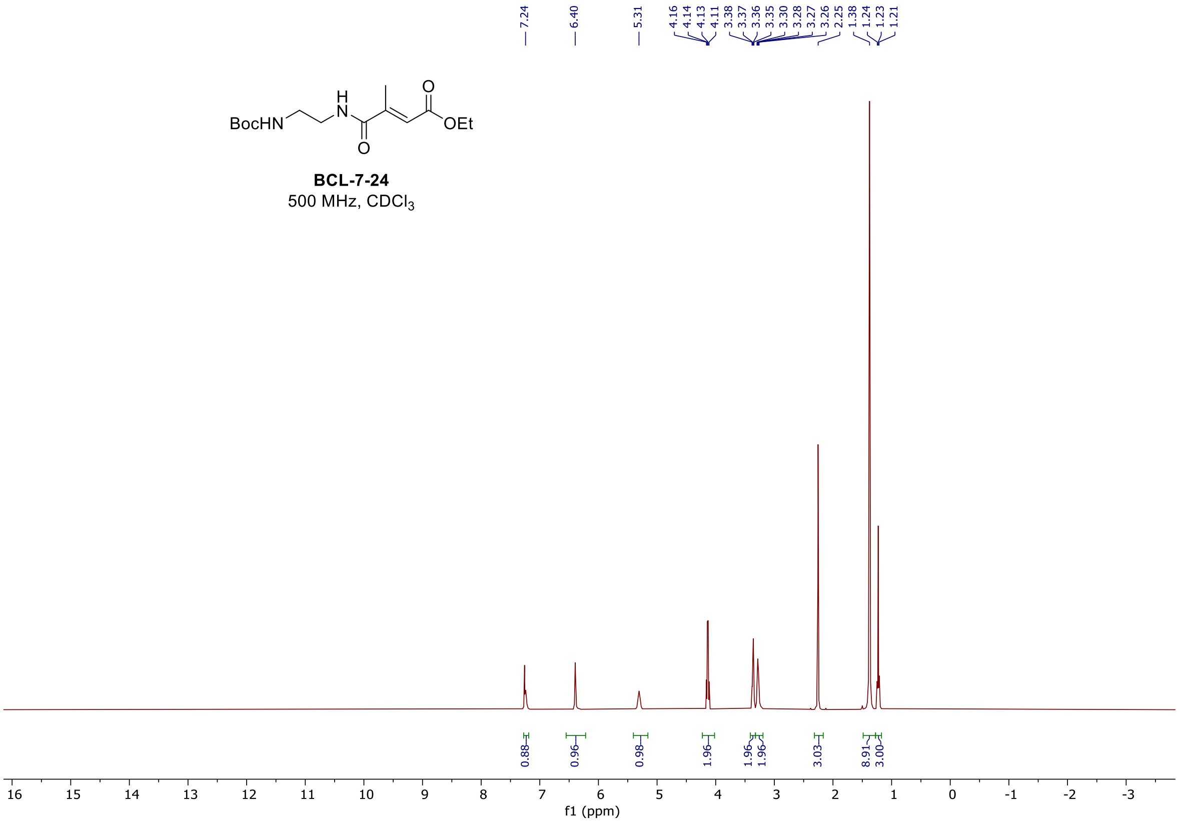

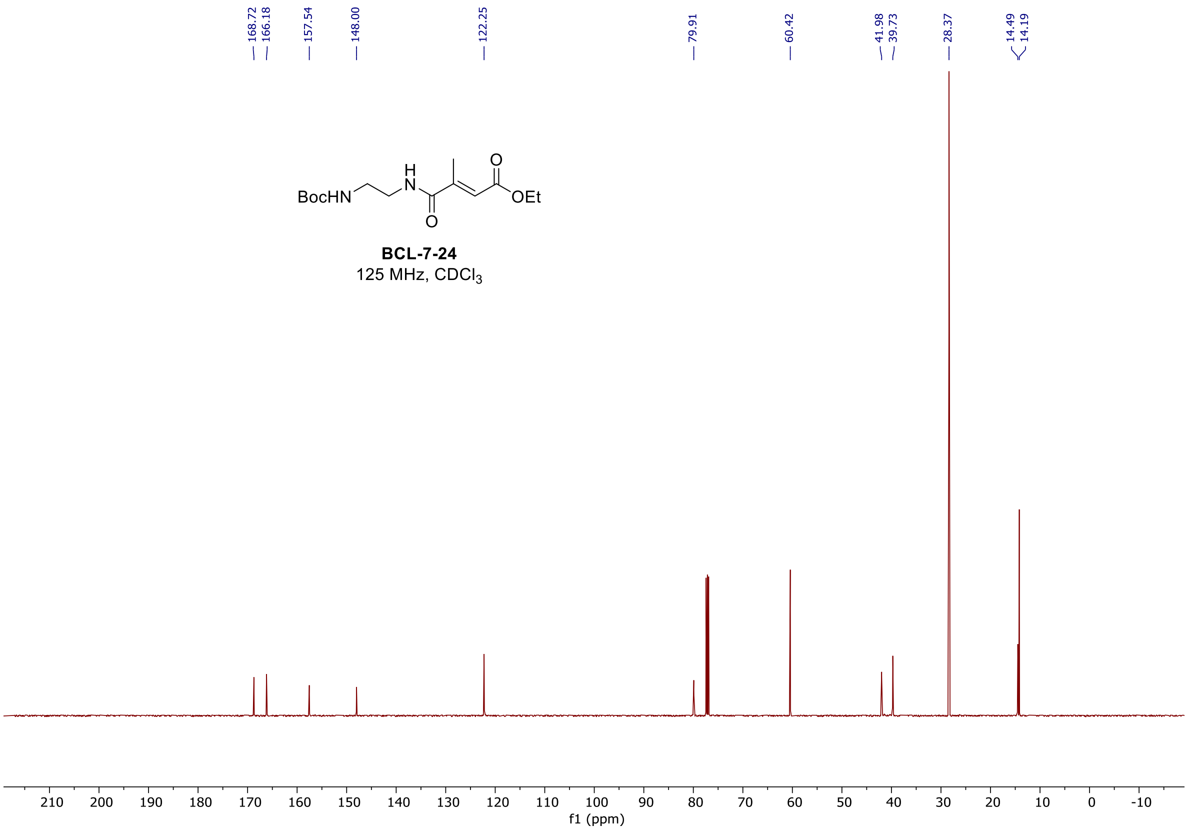

#### <u>$^{1}H$ and $^{13}C$ spectra of **BCL-7-28**</u>

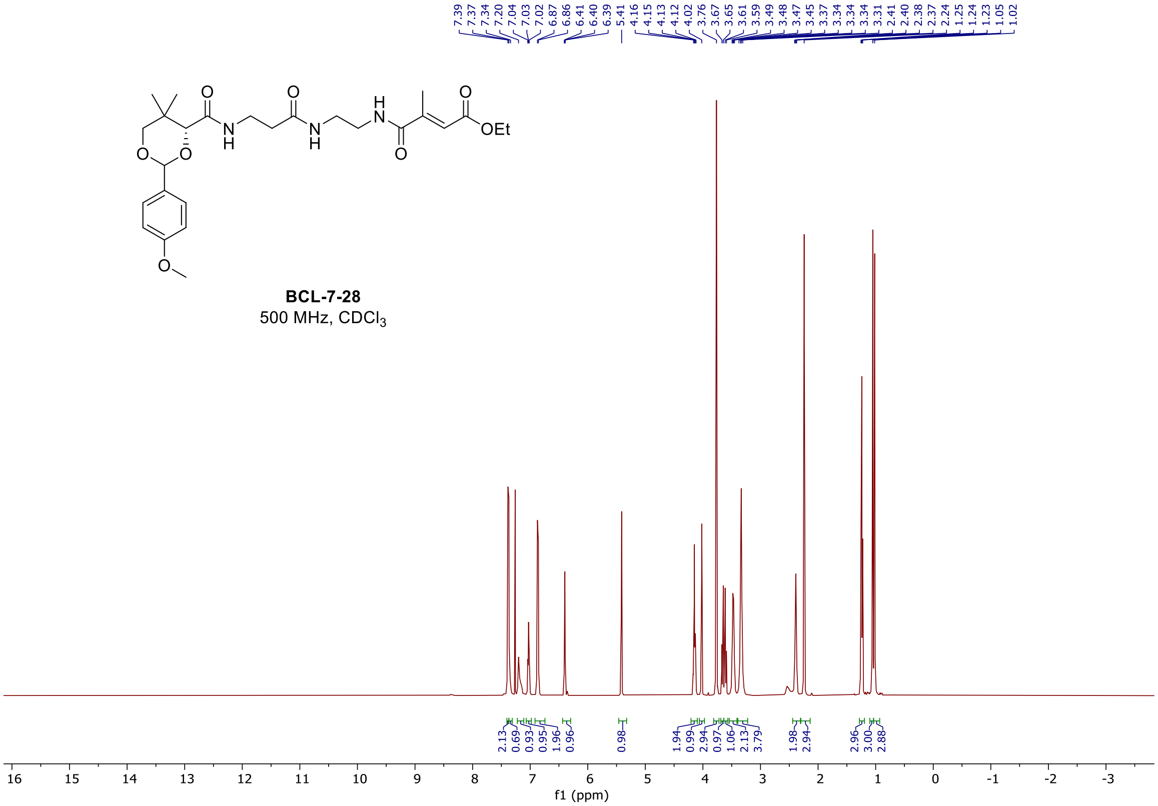

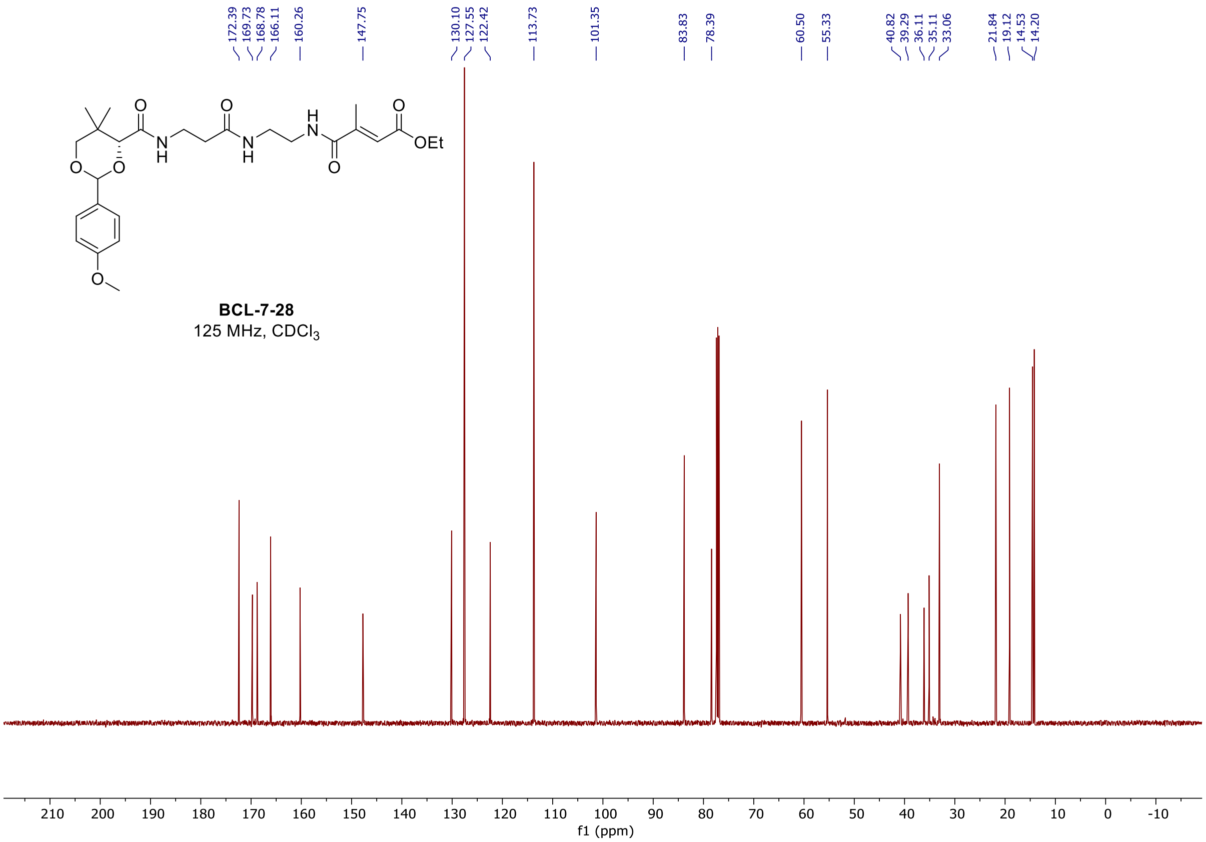

<u>$^1$H and $^{13}$C spectra of **BCL-7-31**</u>

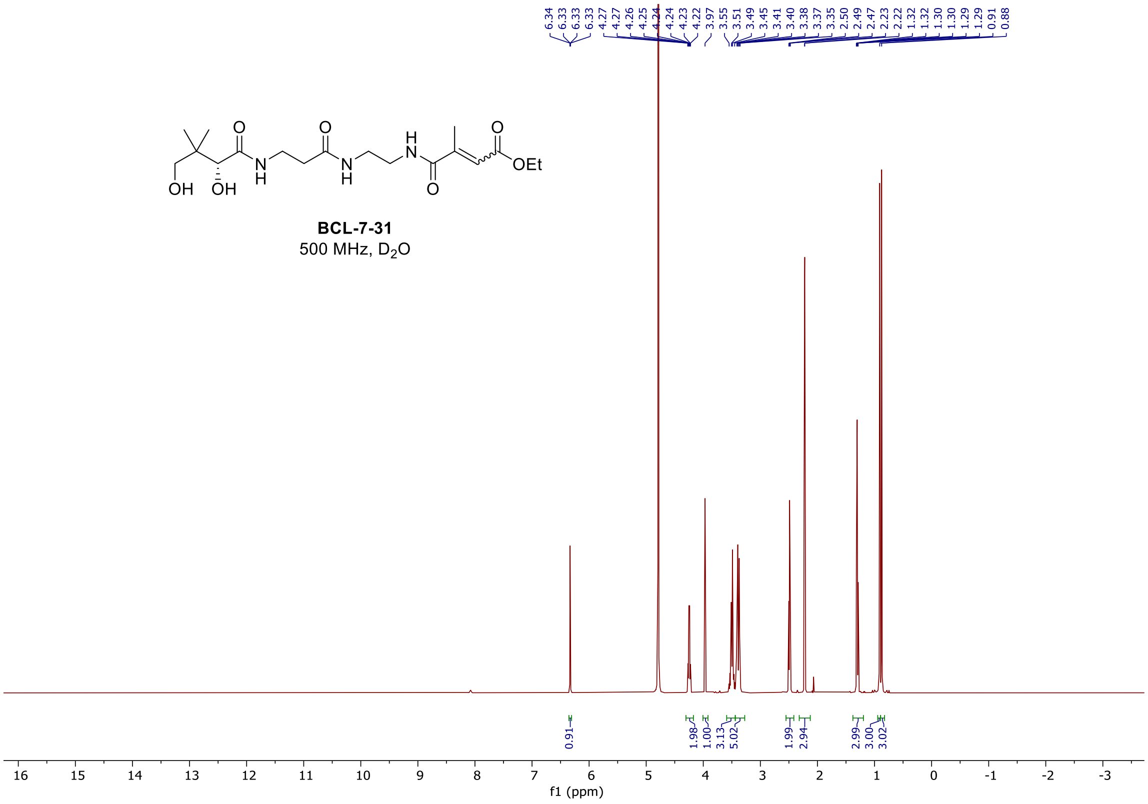

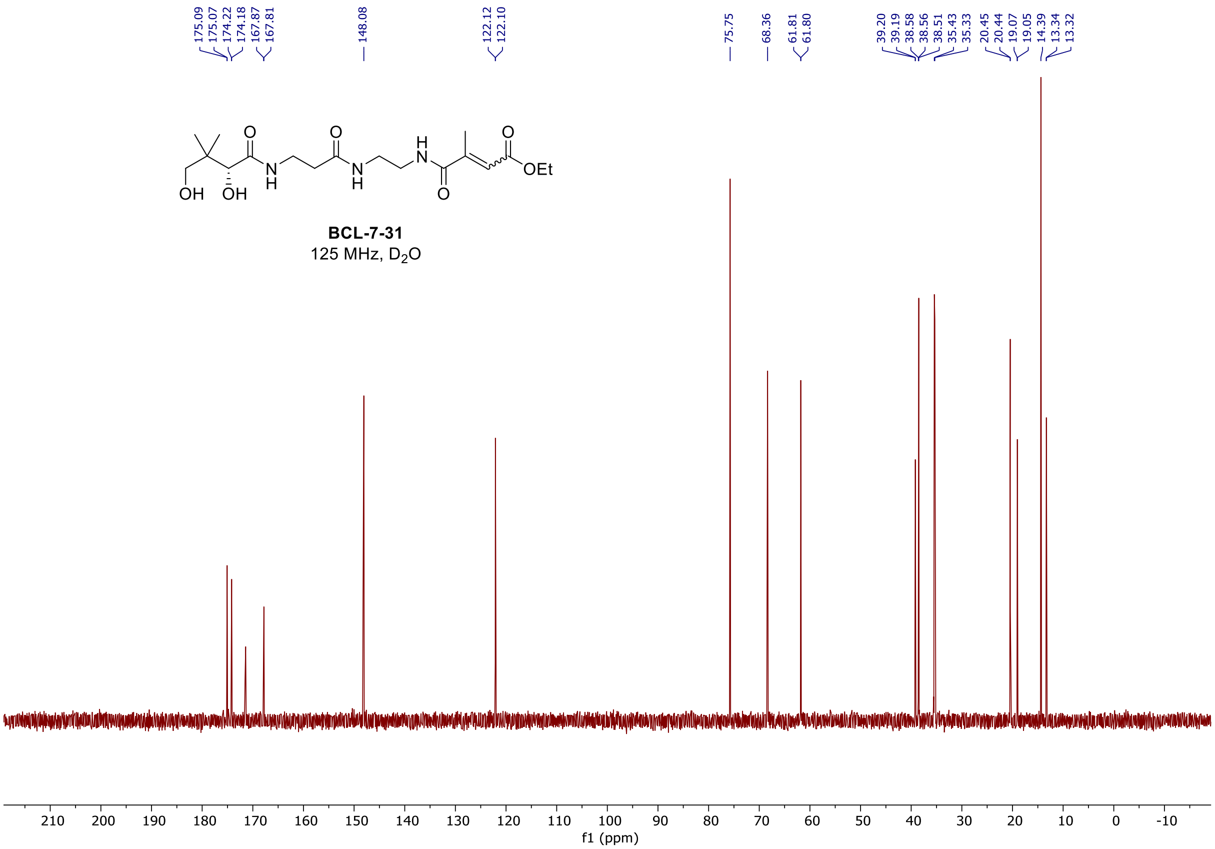

<u>$^{1}$H, $^{19}$F, and $^{13}$C spectra of **BCL-7-33**</u>

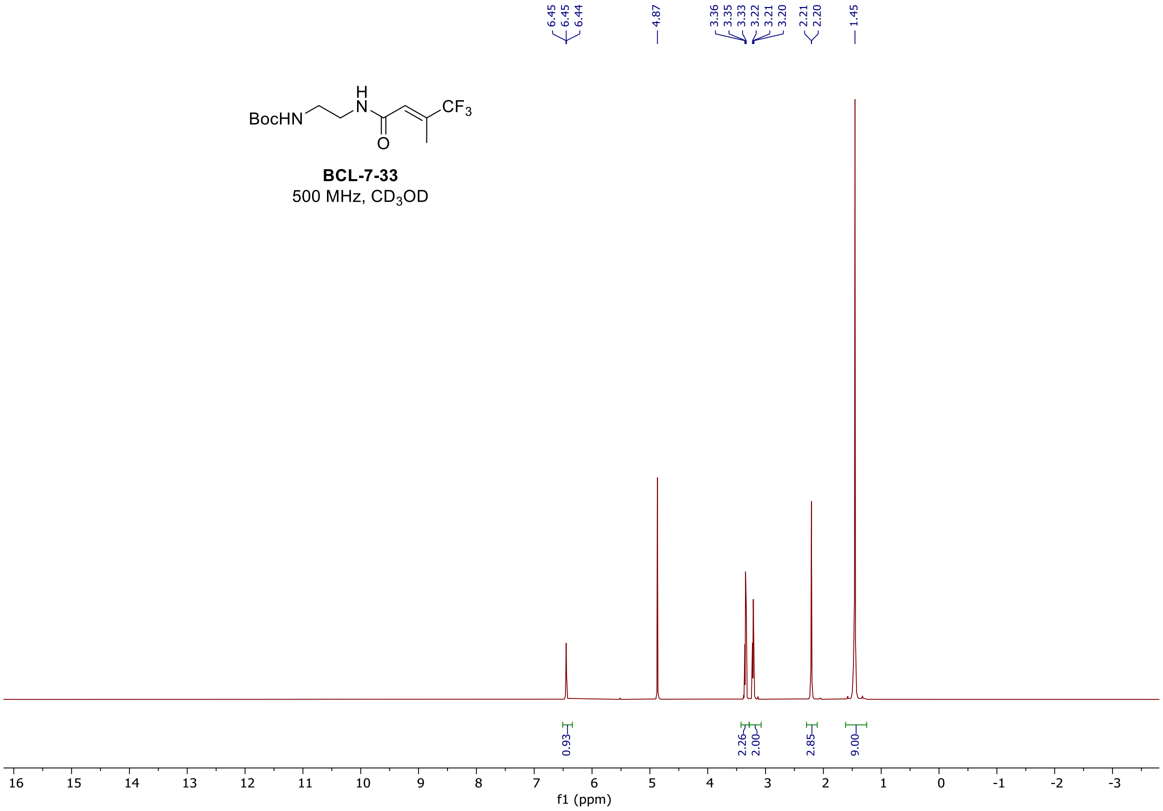

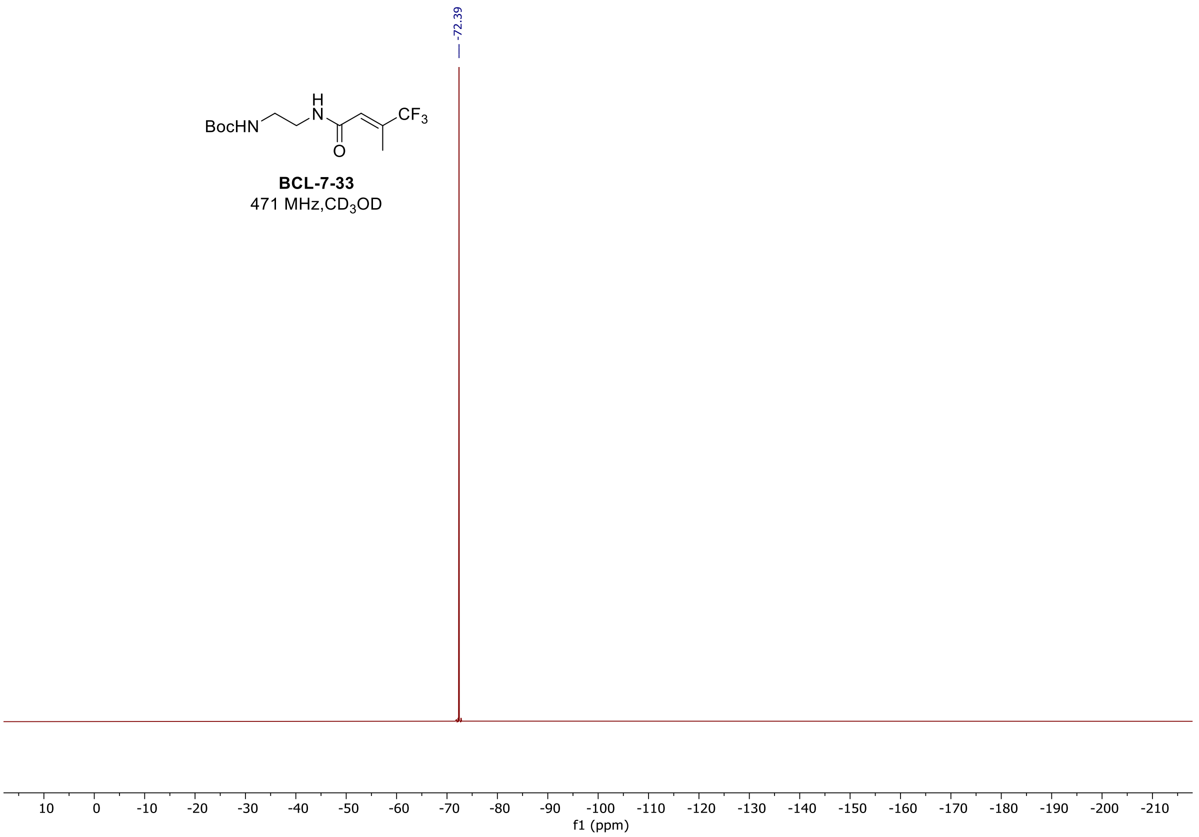
-72.39
BocHN
H
N
CF3
O
BCL-7-33
471 MHz,CD3OD
f1 (ppm)

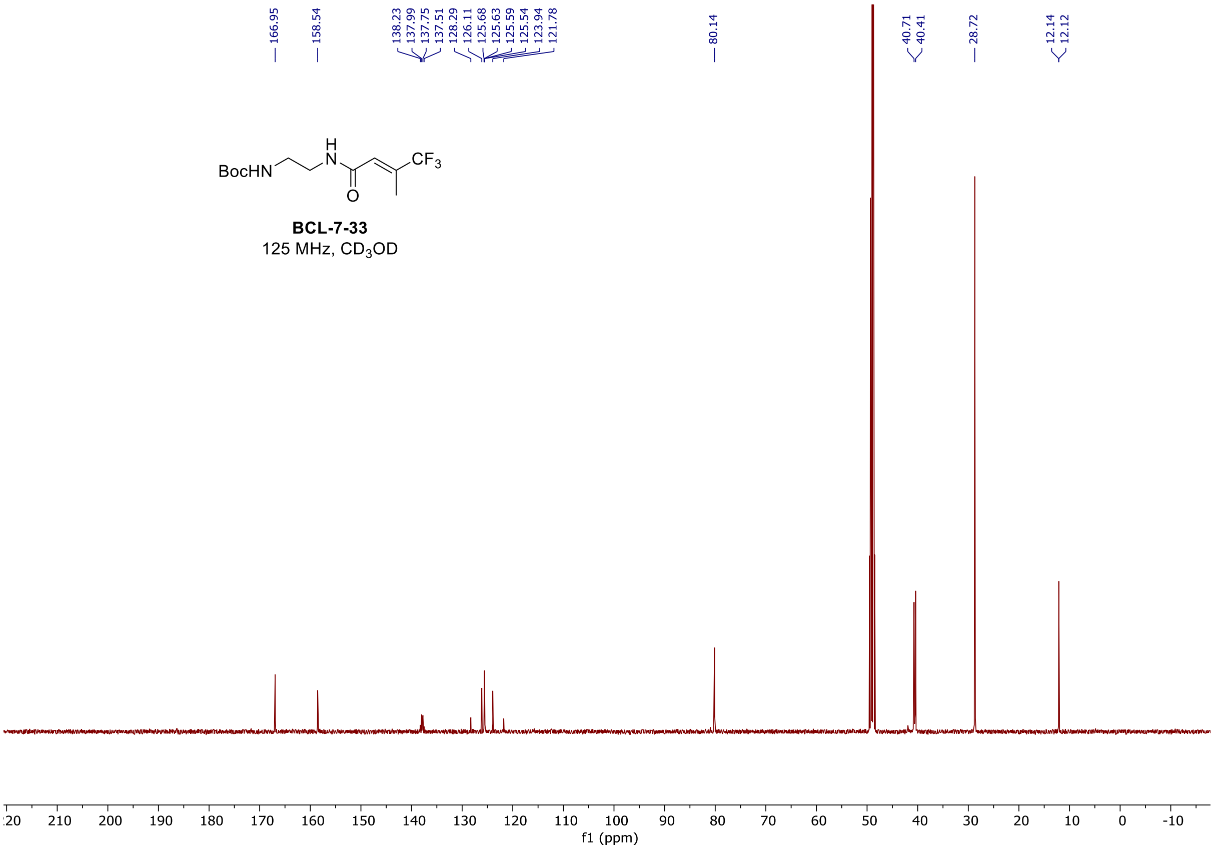
166.95
158.54
138.23
137.99
137.75
137.51
128.29
126.11
125.68
125.63
125.59
125.54
123.94
121.78
80.14
40.71
40.41
28.72
12.14
12.12
BocHN
H
N
CF3
O
BCL-7-33
125 MHz, CD3OD
f1 (ppm)

$^1H$, $^{19}F$, and $^{13}C$ spectra of **BCL-7-37**

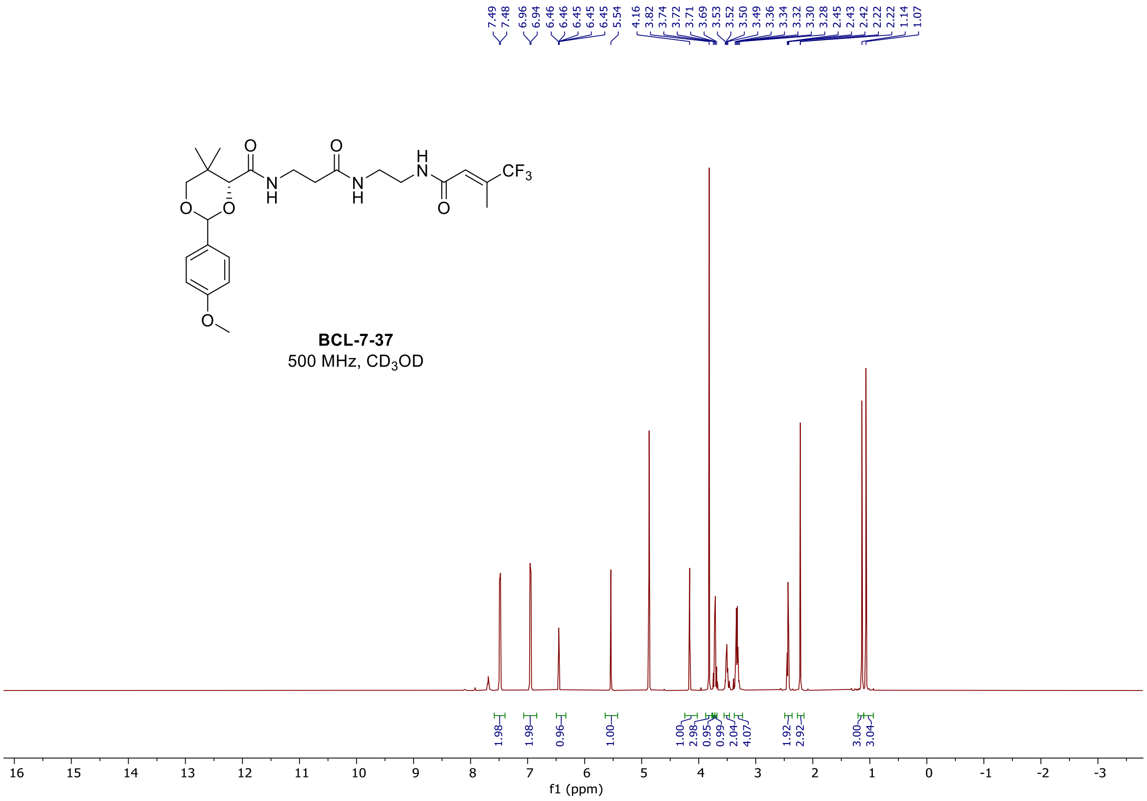

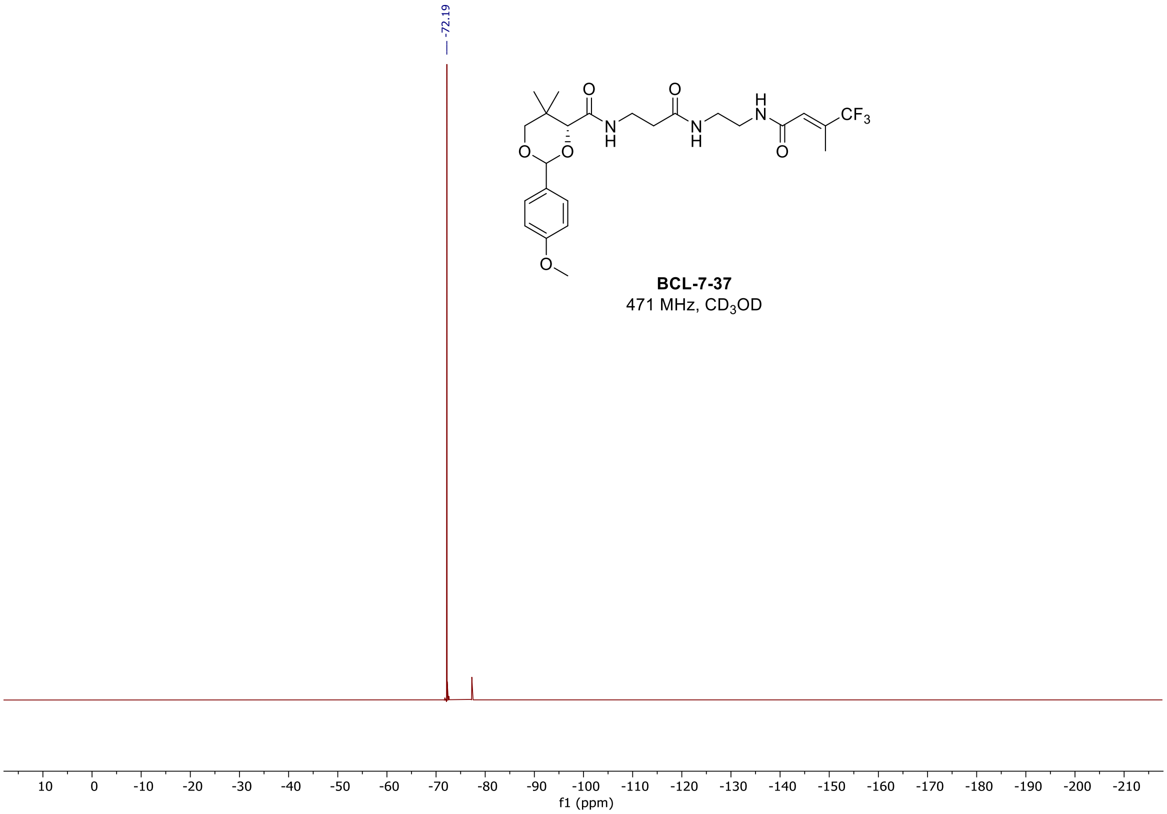

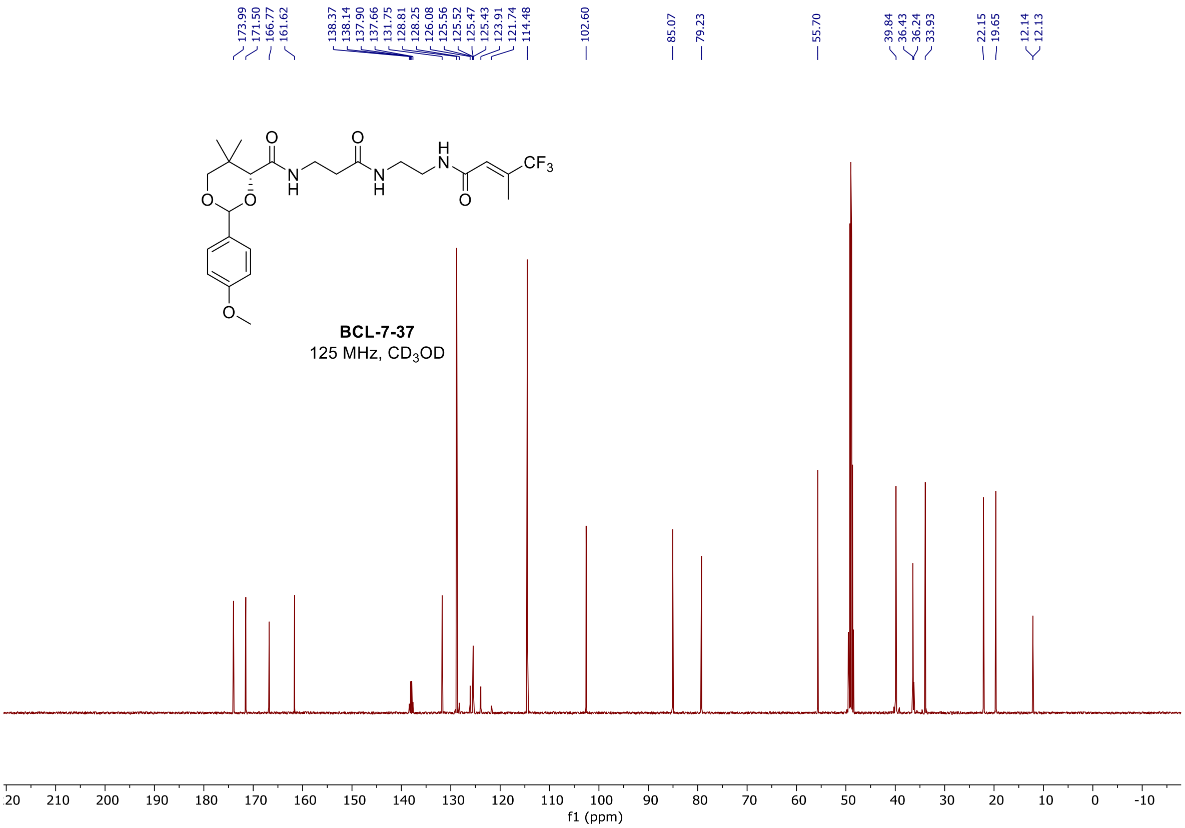

$^{1}$H, $^{19}$F, and $^{13}$C spectra of **BCL-7-45**

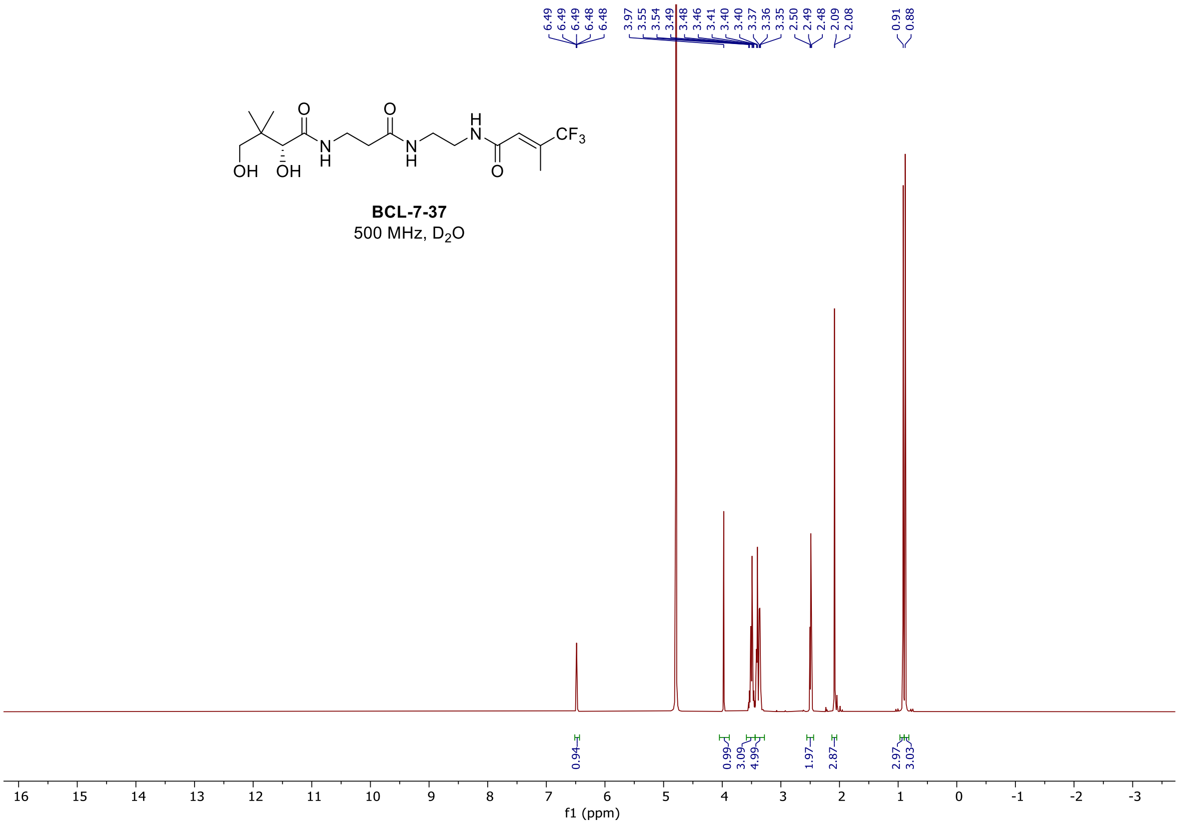

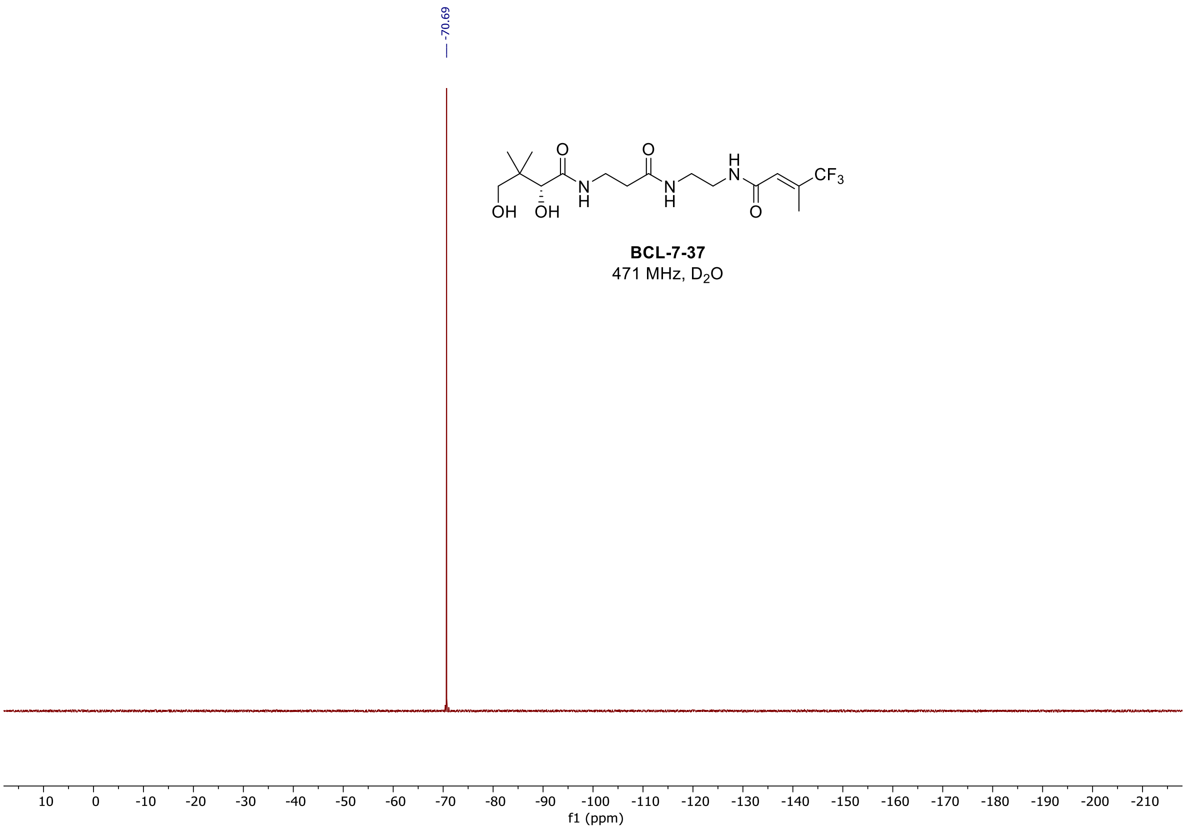
-70.69
BCL-7-37
471 MHz, D2O
f1 (ppm)

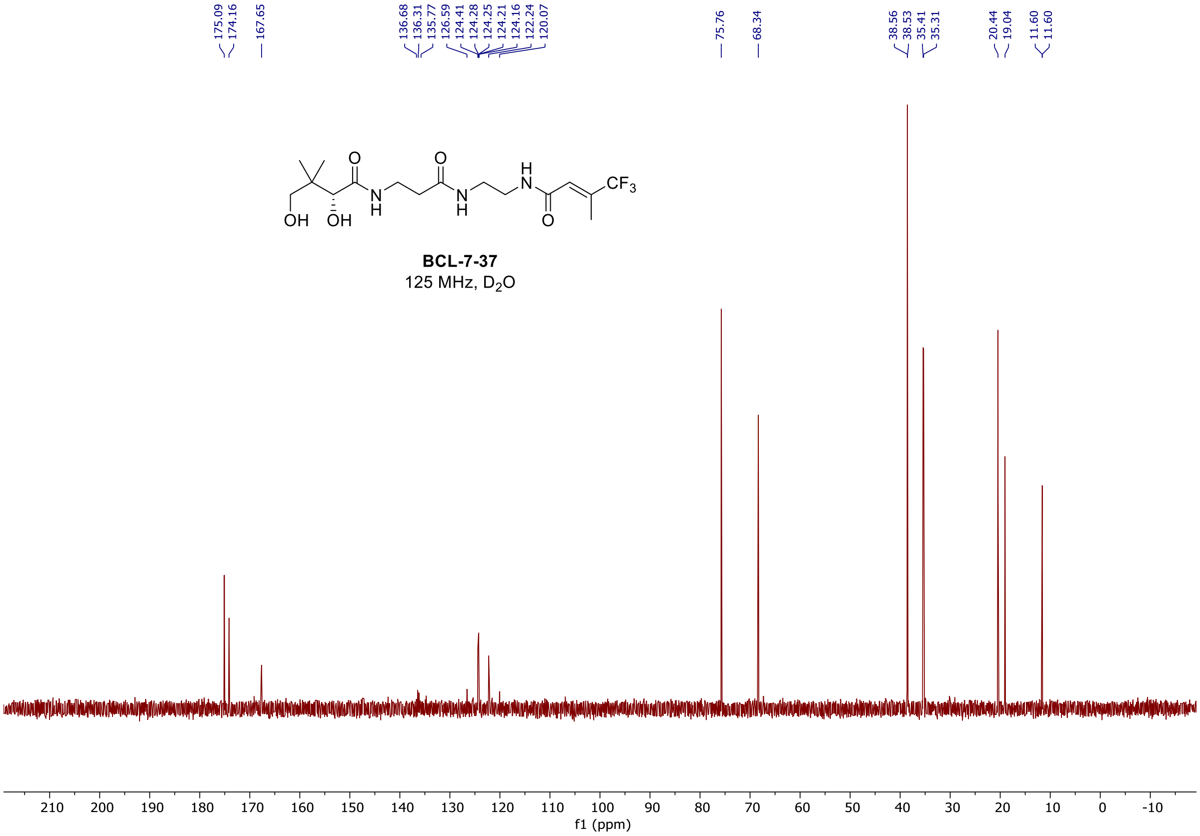
175.09
174.16
167.65
136.68
136.31
135.77
126.59
124.41
124.28
124.25
124.21
124.16
122.24
120.07
75.76
68.34
38.56
38.53
35.41
35.31
20.44
19.04
11.60
11.60
BCL-7-37
125 MHz, D2O
f1 (ppm)

#### $^{1}H$ and $^{13}C$ spectra of **BCL-7-46**

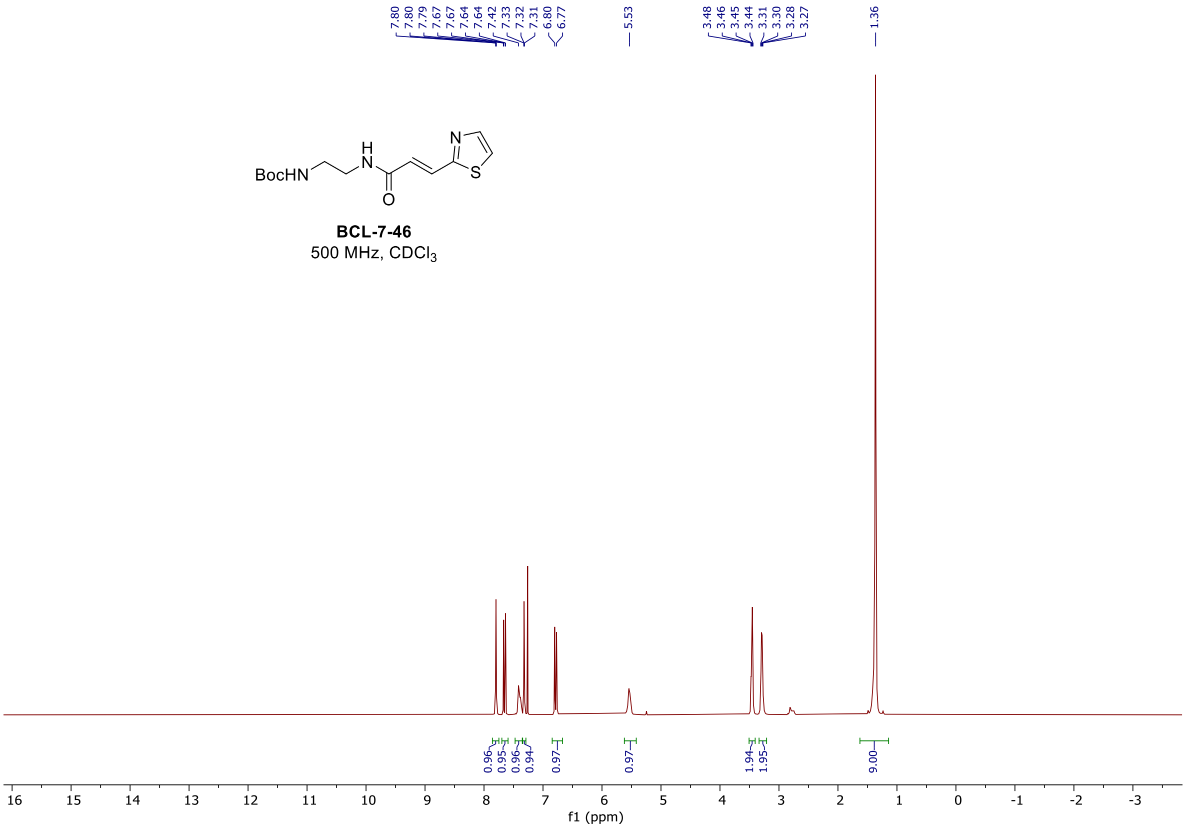

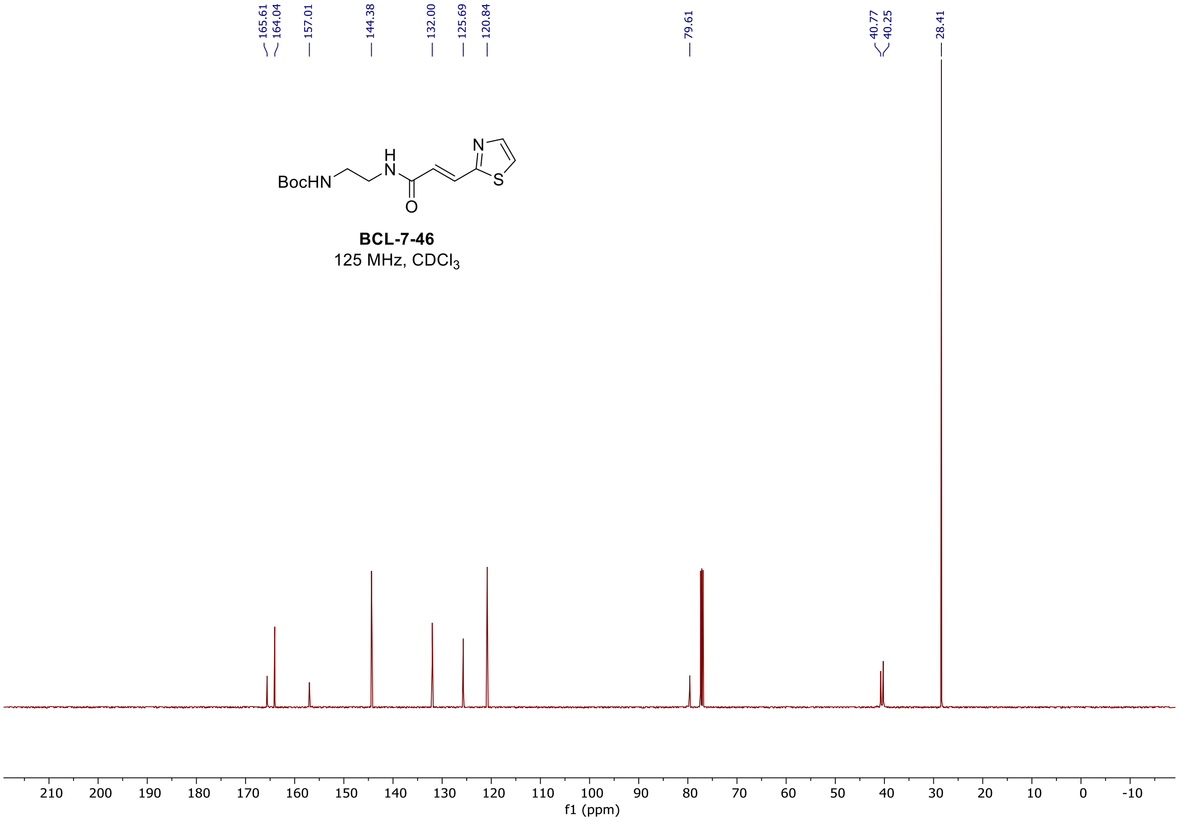

#### $^{1}H$ and $^{13}C$ spectra of **BCL-7-53**

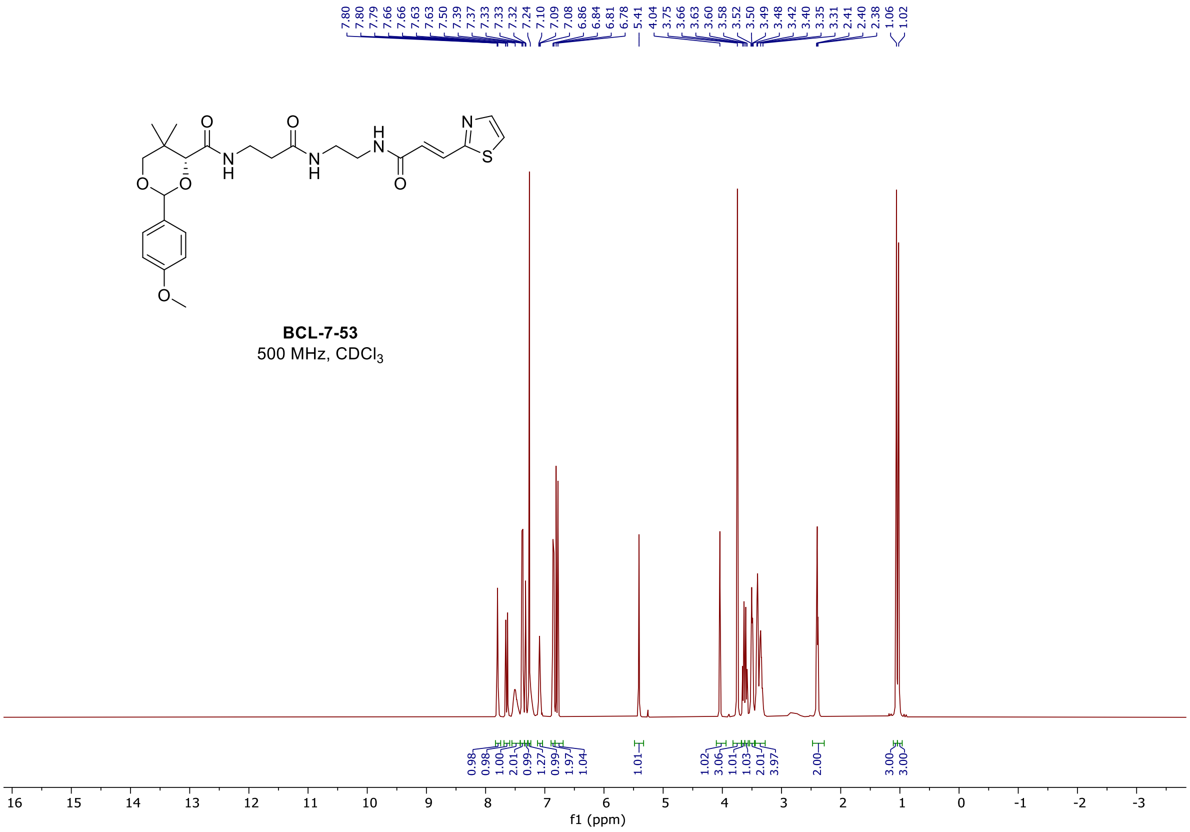

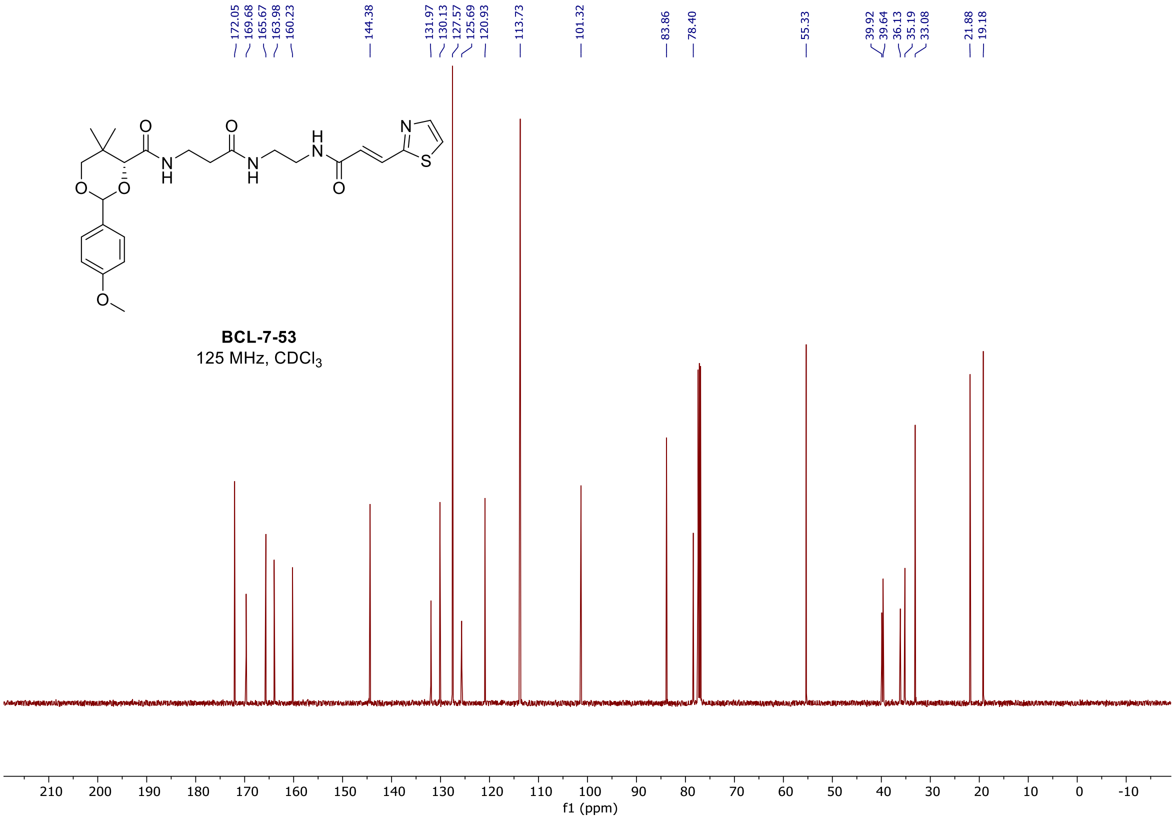

#### $^{1}H$ and $^{13}C$ spectra of **BCL-7-58**

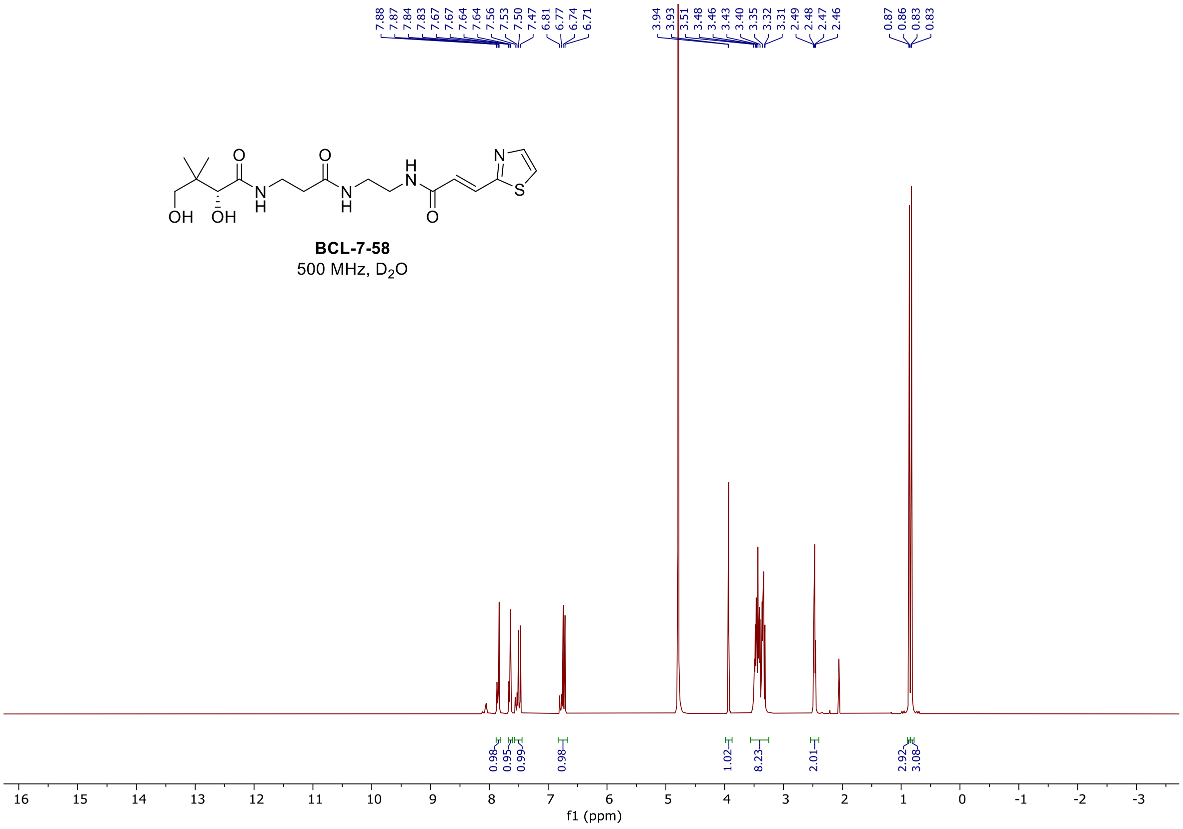

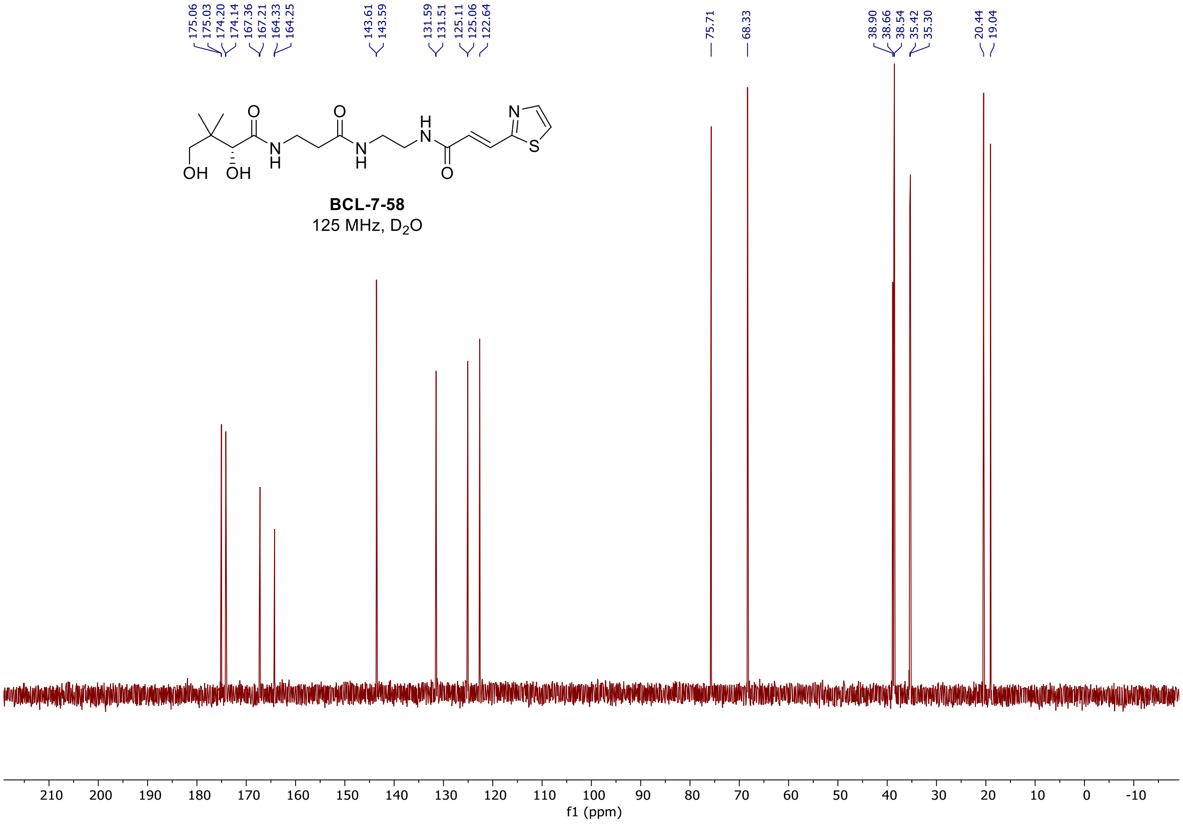

<u>$^{1}$H and $^{13}$C spectra of **BCL-7-17**</u>

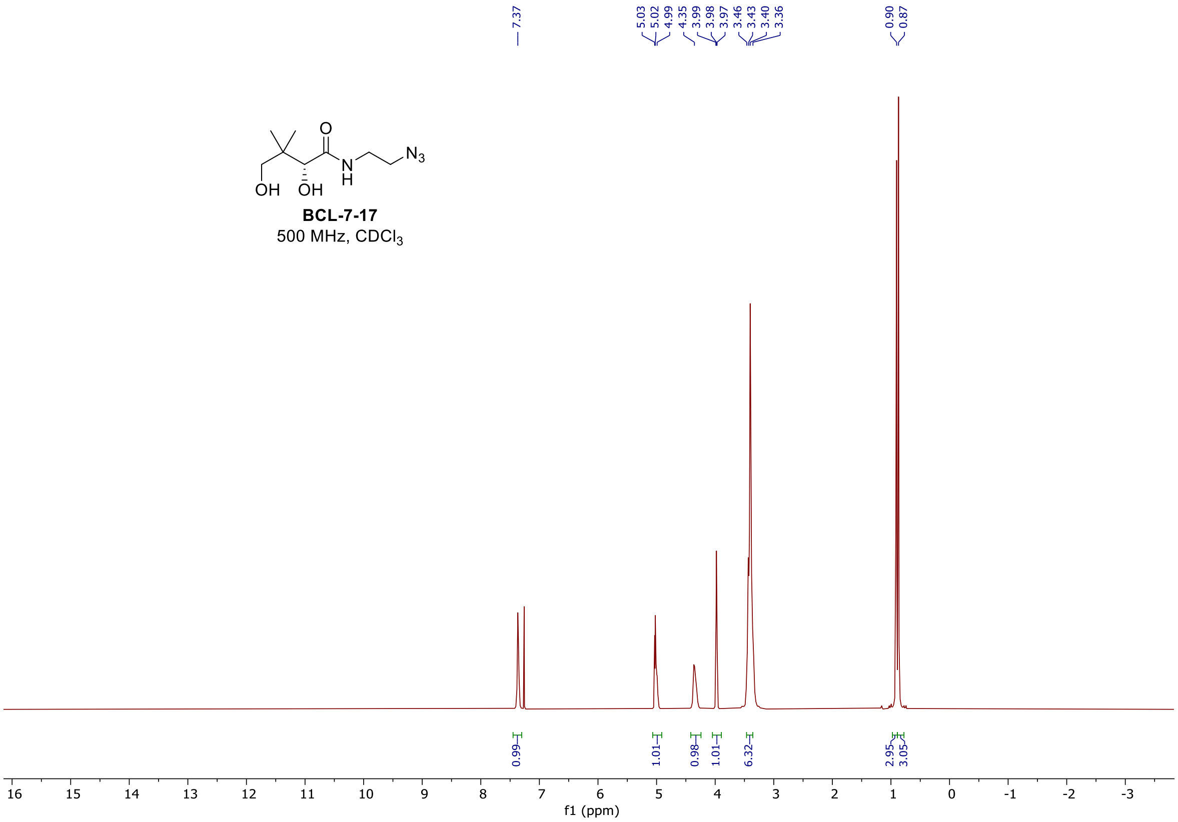

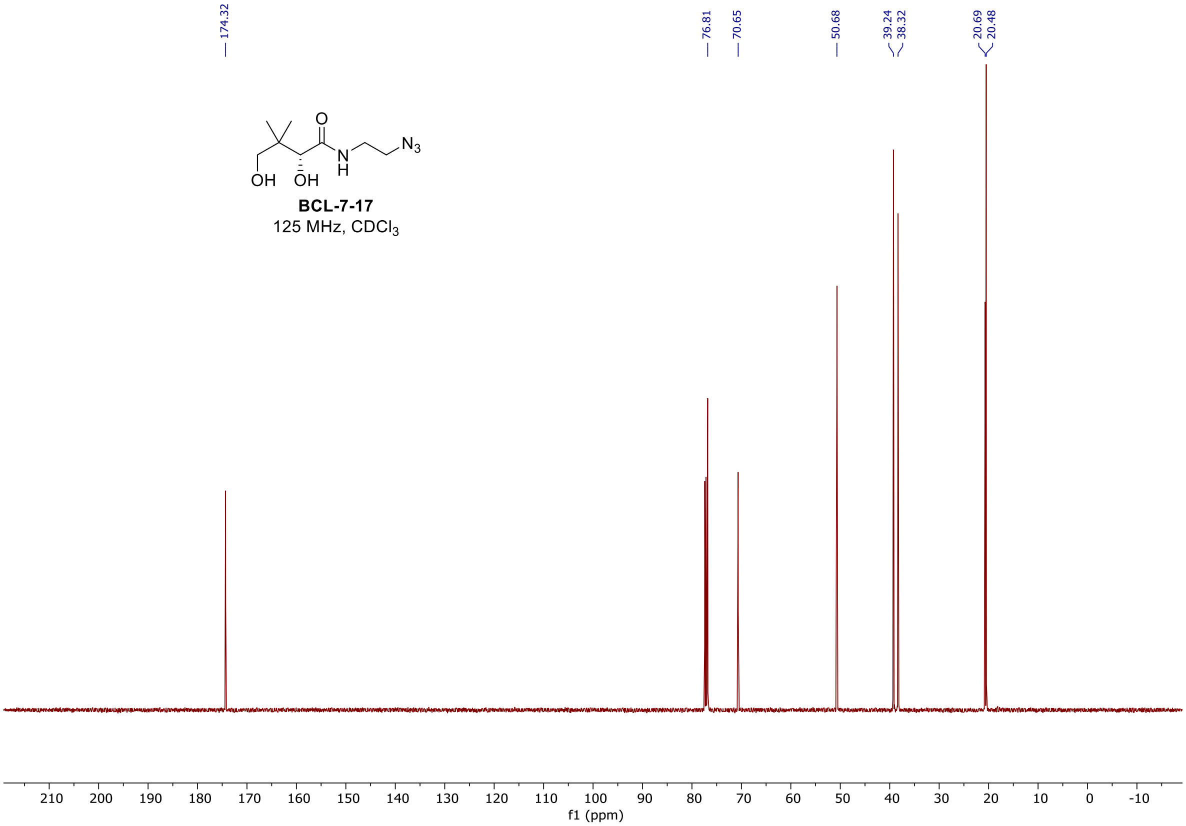

[1]H and [13]C spectra of **BCL-7-9**

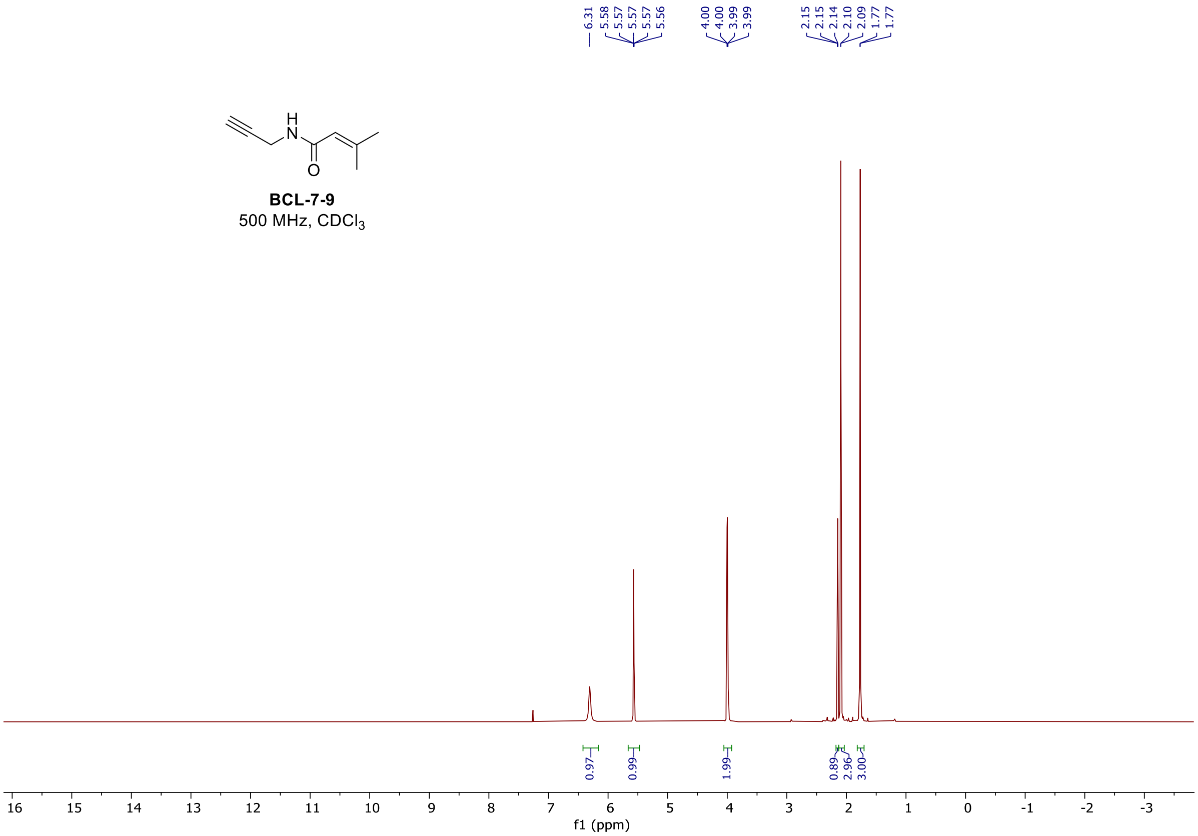

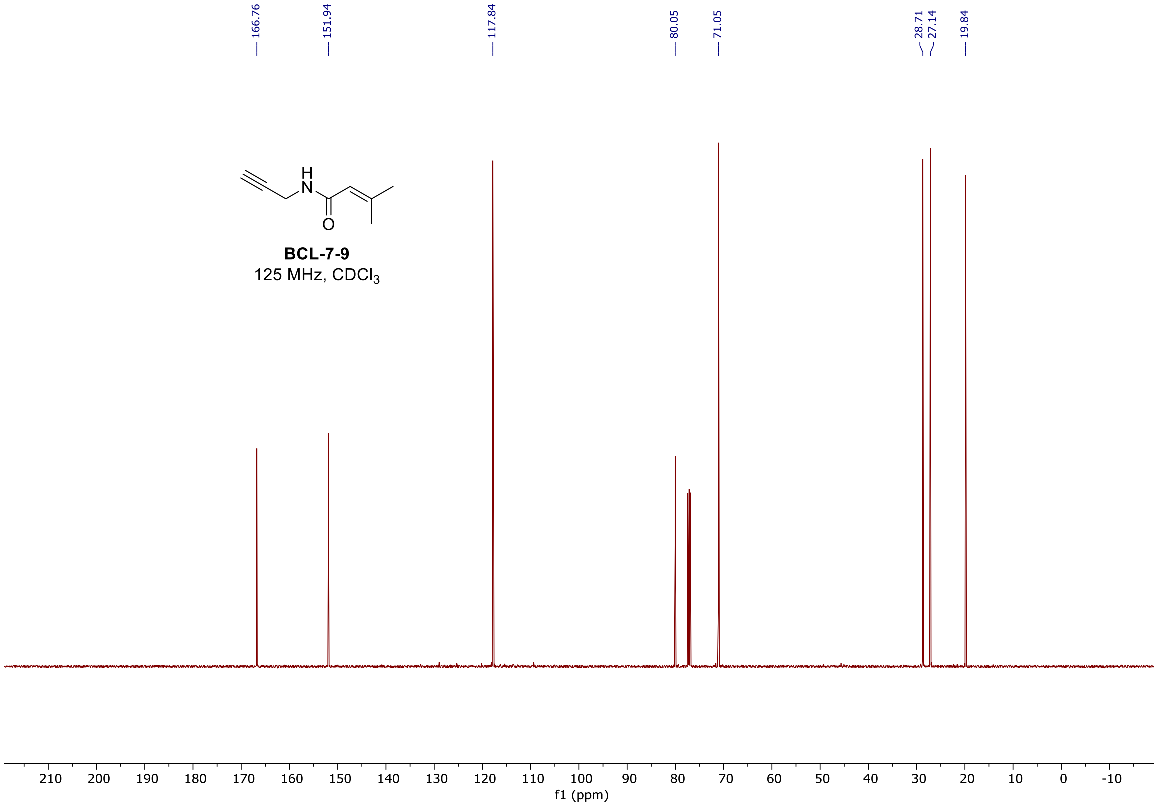

[1]H and [13]C spectra of **BCL-7-75**

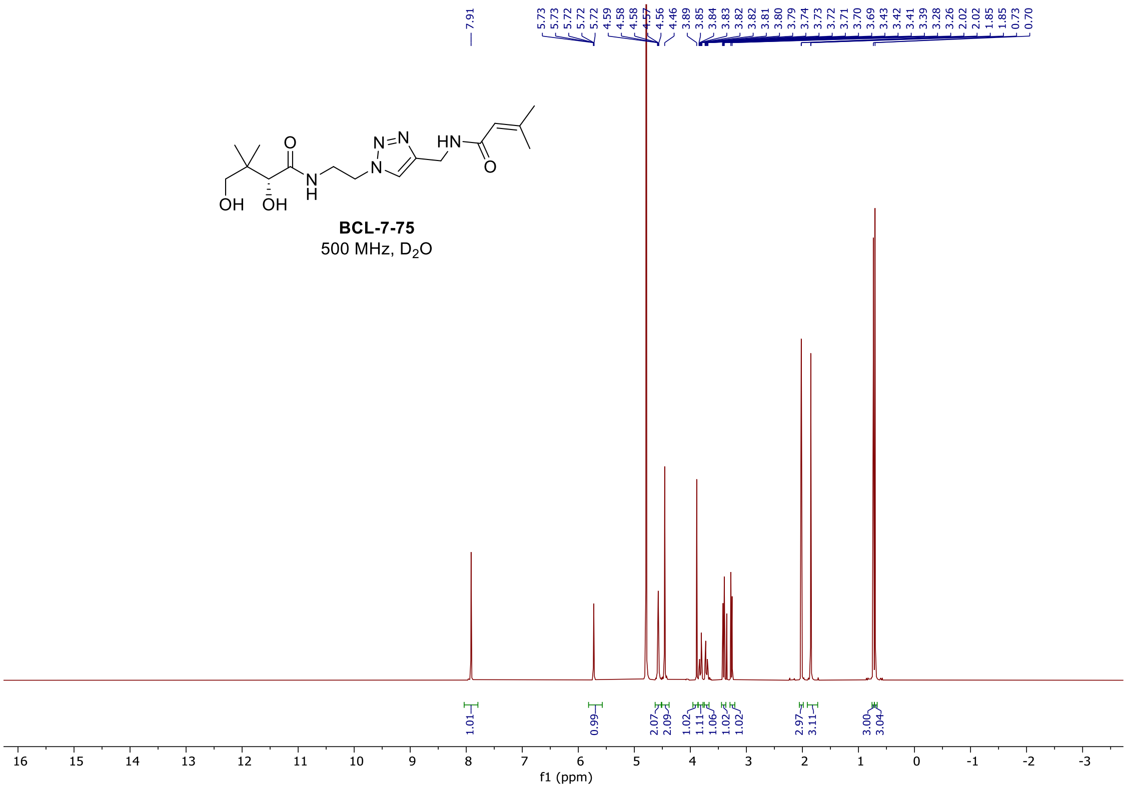

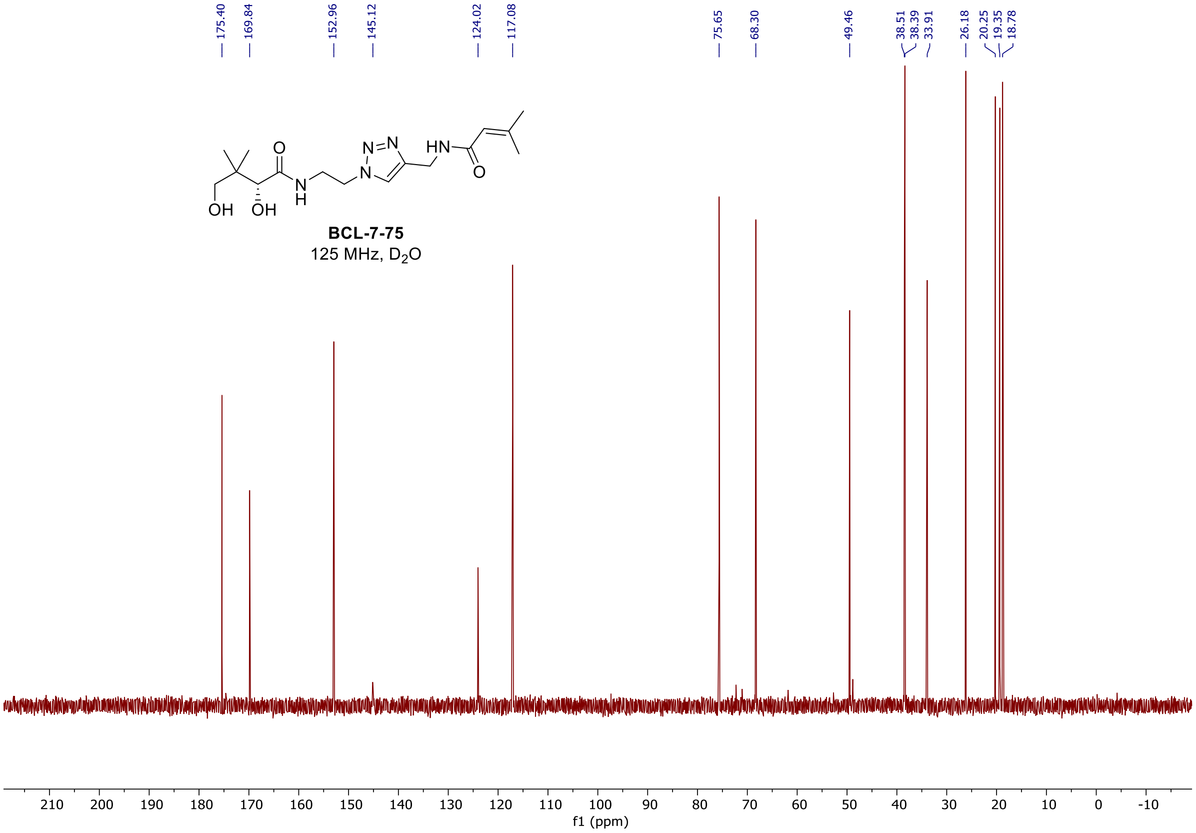

$^{1}H$, $^{19}F$, and $^{13}C$ spectra of **BCL-7-60**

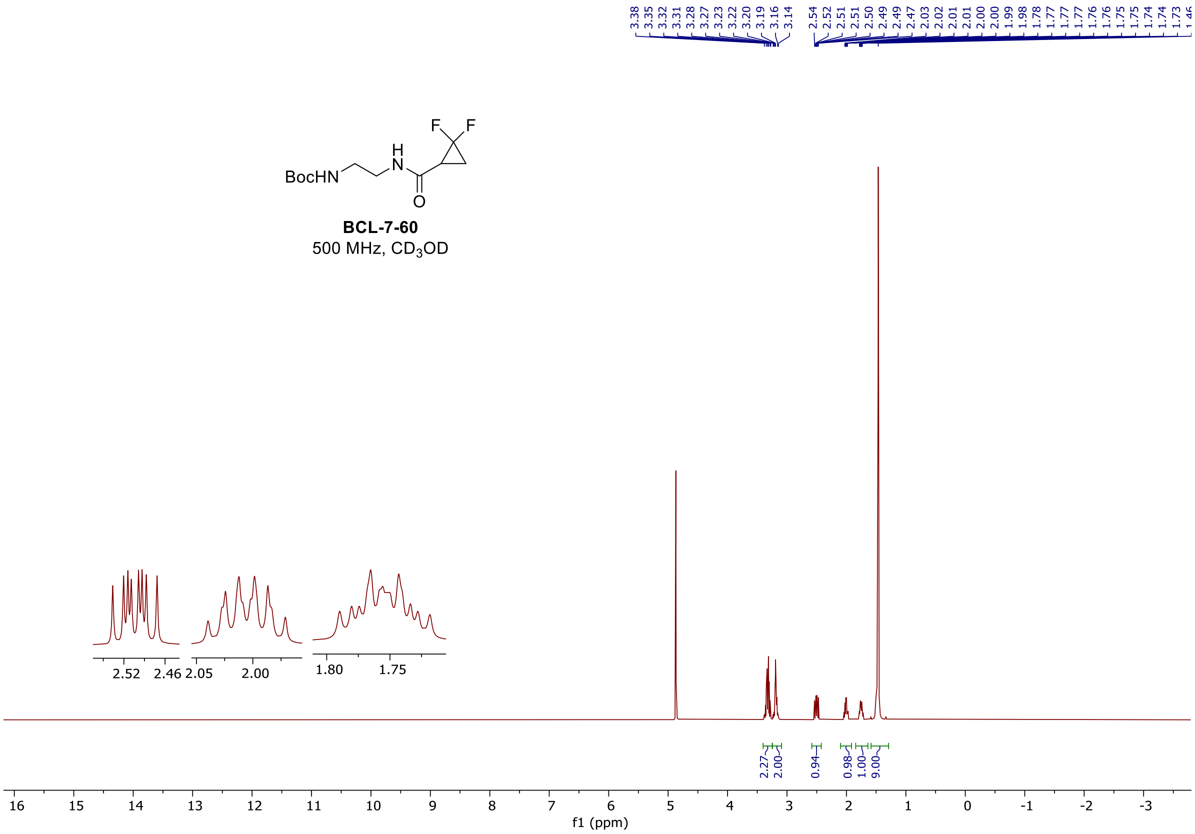

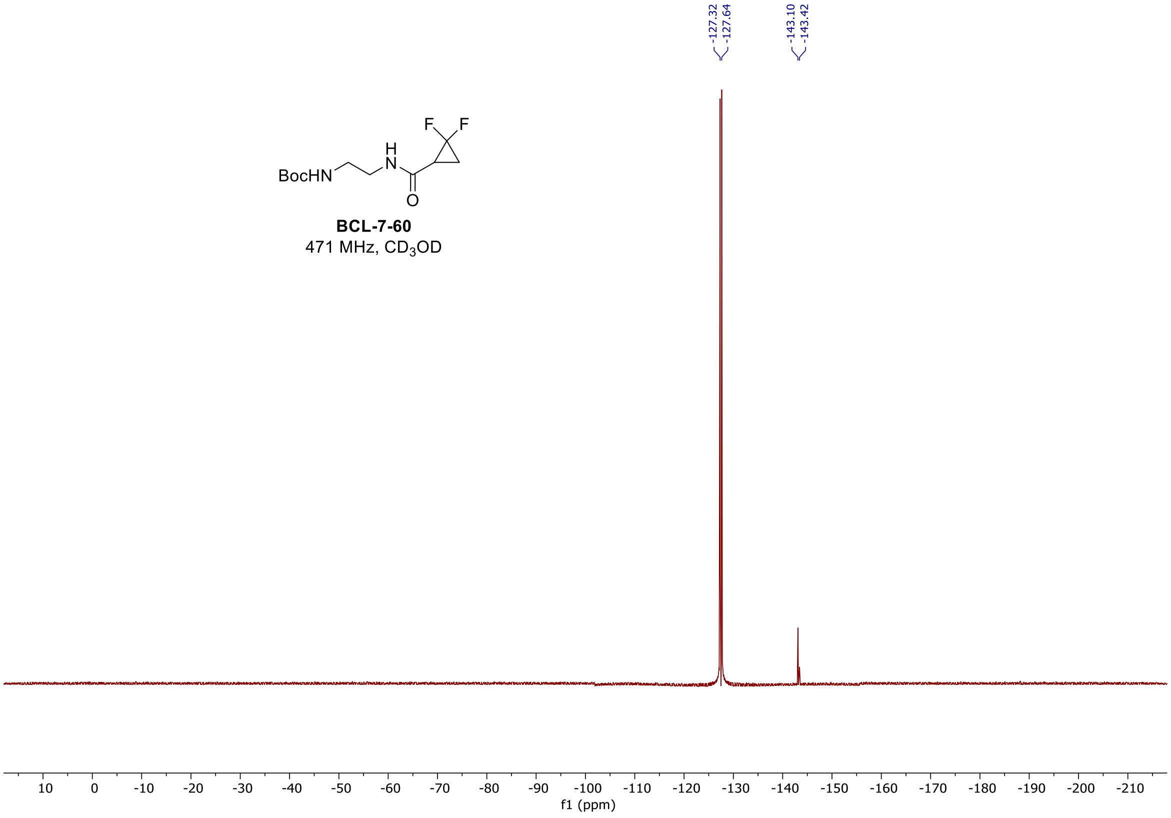

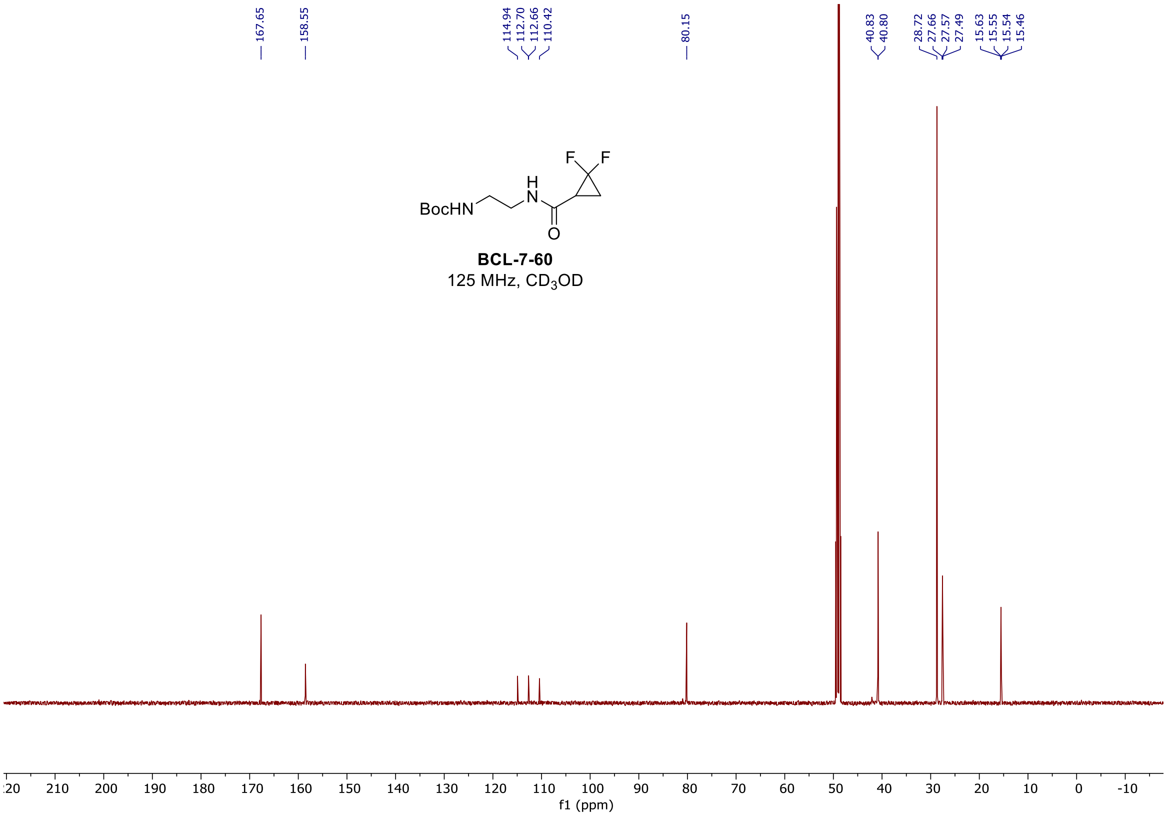

<u>[1]H, [19]F, and [13]C spectra of **BCL-7-67**</u>

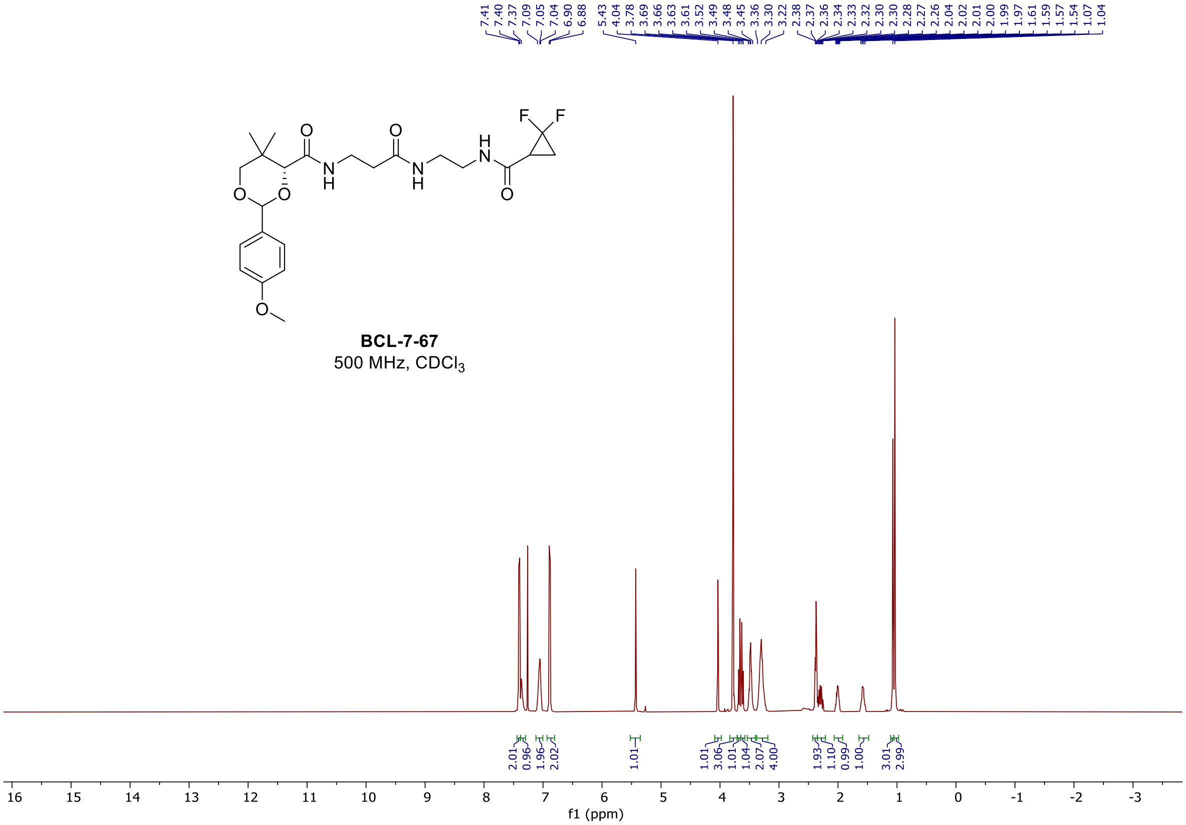

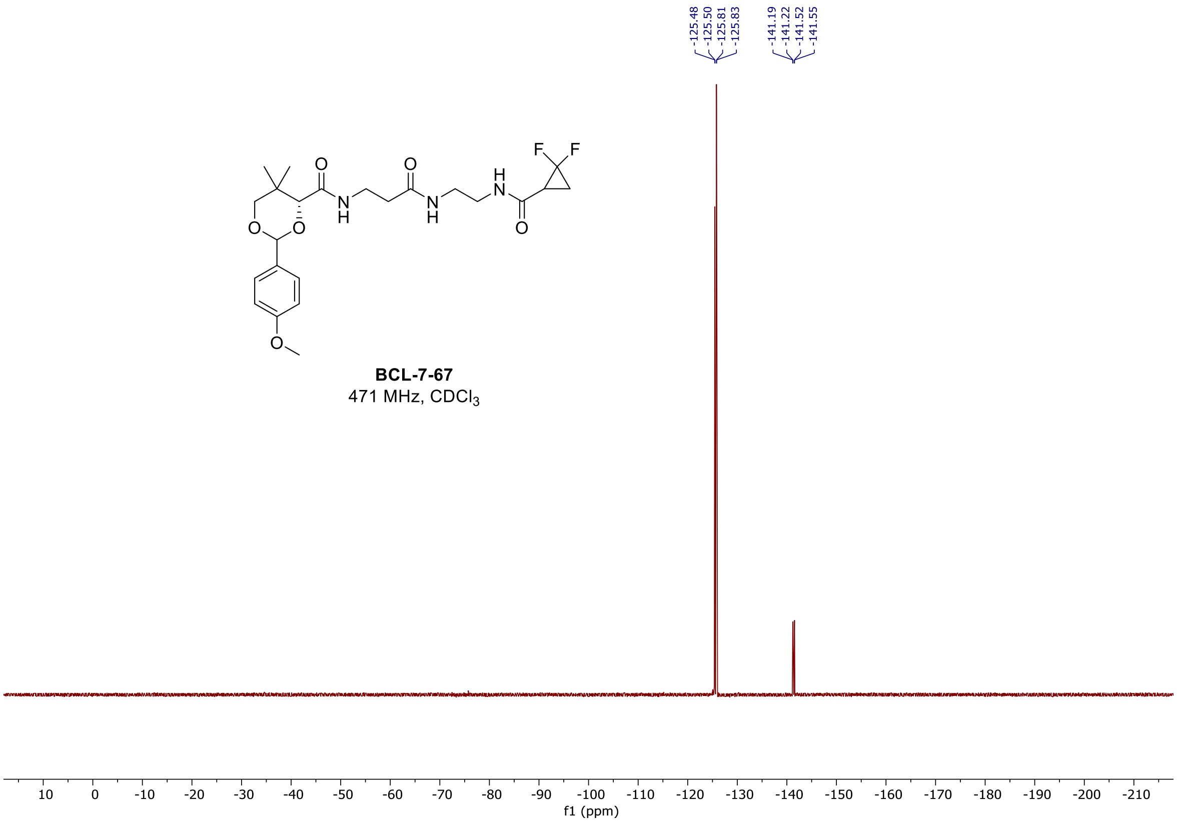
-125.48
-125.50
-125.81
-125.83
-141.19
-141.22
-141.52
-141.55
F
F
O
O
N
H
H
N
O
O
O
N
H
O
O
**BCL-7-67**
471 MHz, $CDCl_3$
10 0 -10 -20 -30 -40 -50 -60 -70 -80 -90 -100 -110 -120 -130 -140 -150 -160 -170 -180 -190 -200 -210
f1 (ppm)

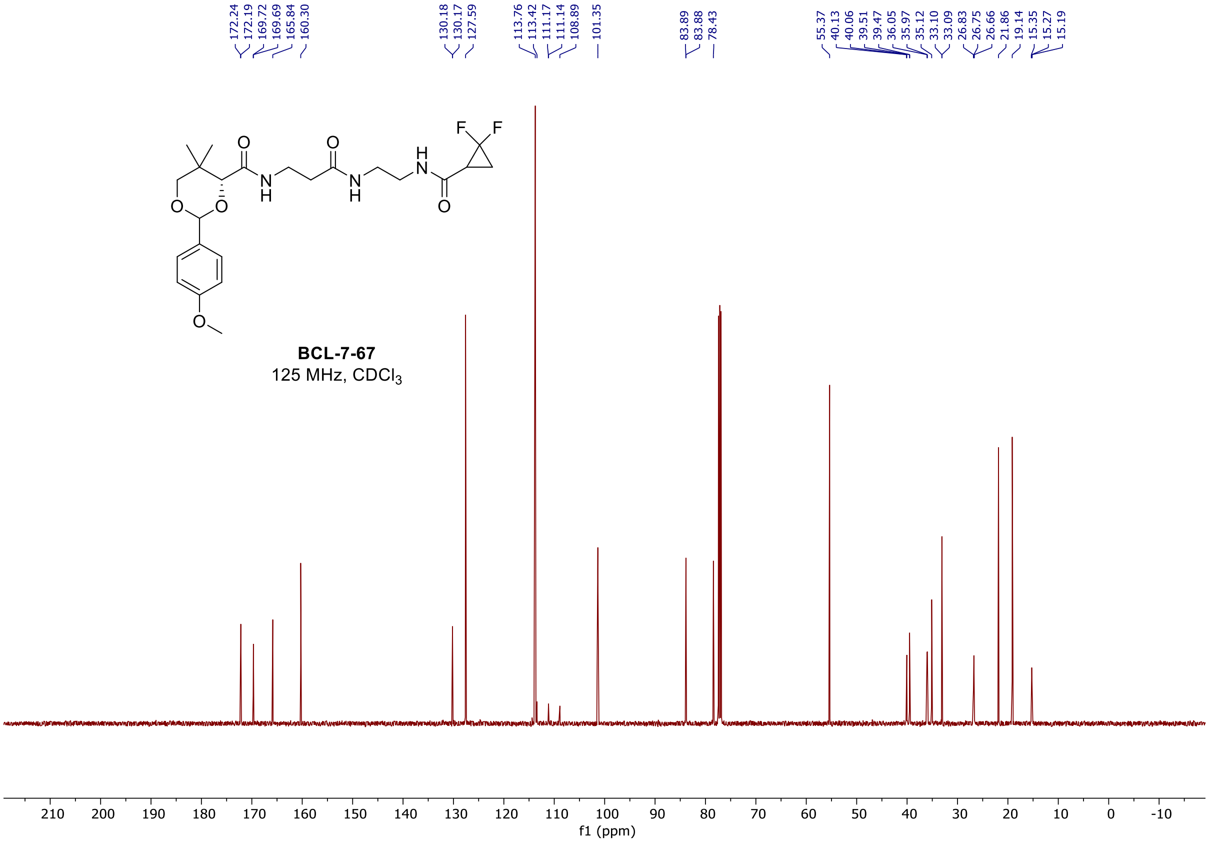
172.24
172.19
169.72
169.69
165.84
160.30
130.18
130.17
127.59
113.76
113.42
111.17
111.14
108.89
101.35
83.89
83.88
78.43
55.37
40.13
40.06
39.51
39.47
36.05
35.97
35.12
33.10
33.09
26.83
26.75
26.66
21.86
19.14
15.35
15.27
15.19
F
F
O
O
N
H
H
N
O
O
O
N
H
O
O
**BCL-7-67**
125 MHz, $CDCl_3$
210 200 190 180 170 160 150 140 130 120 110 100 90 80 70 60 50 40 30 20 10 0 -10
f1 (ppm)

[1]H, [19]F, and [13]C spectra of **BCL-7-76**

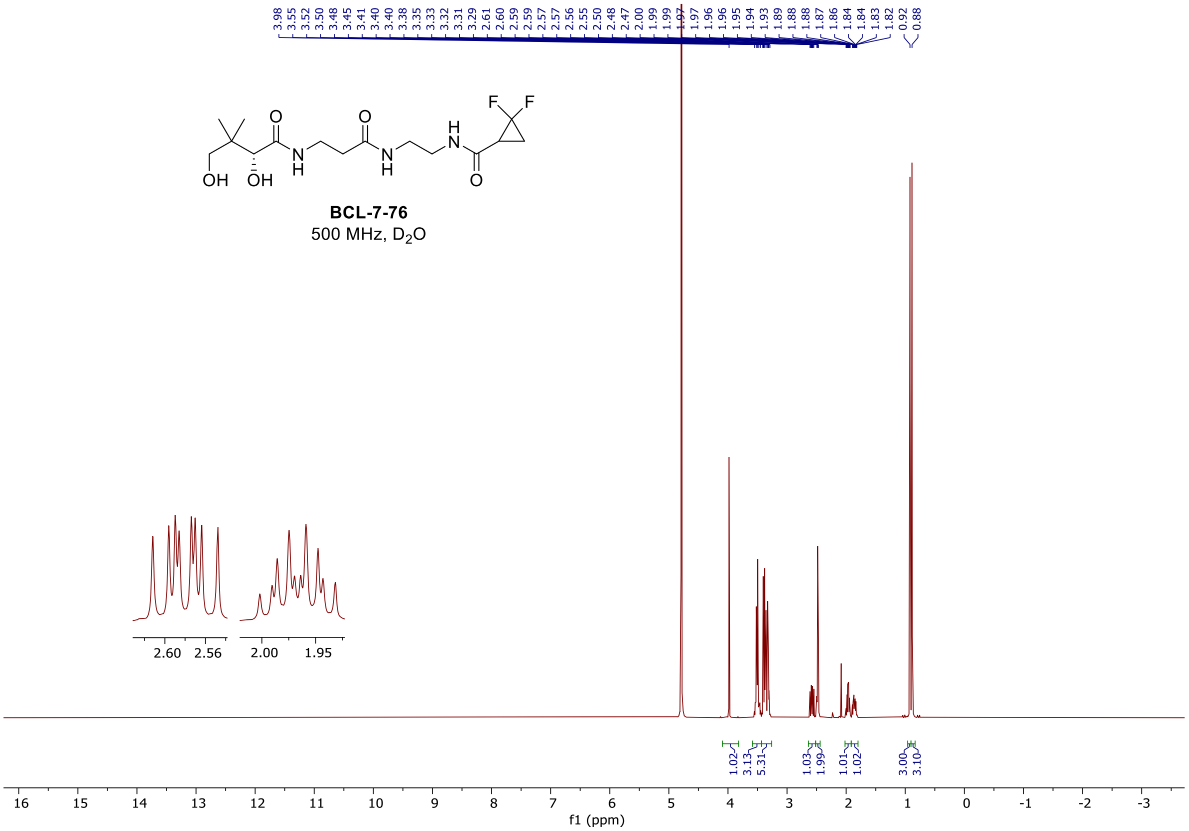

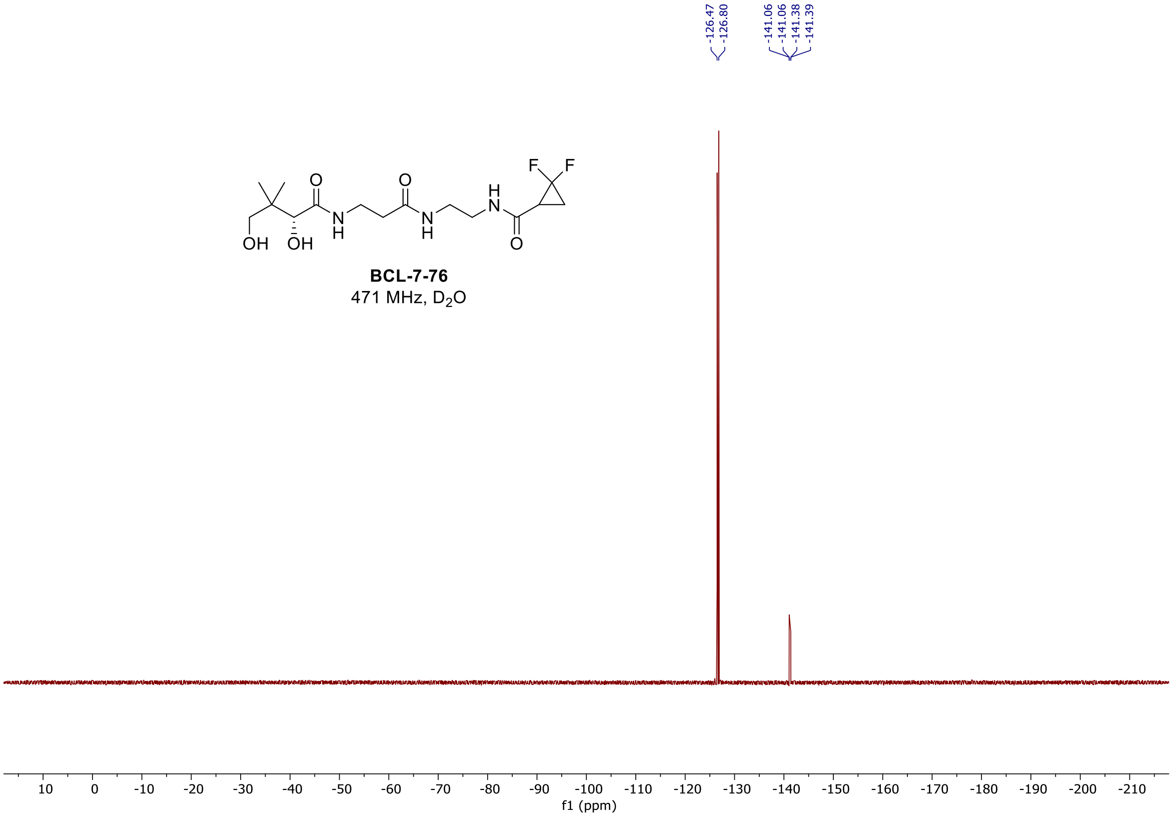

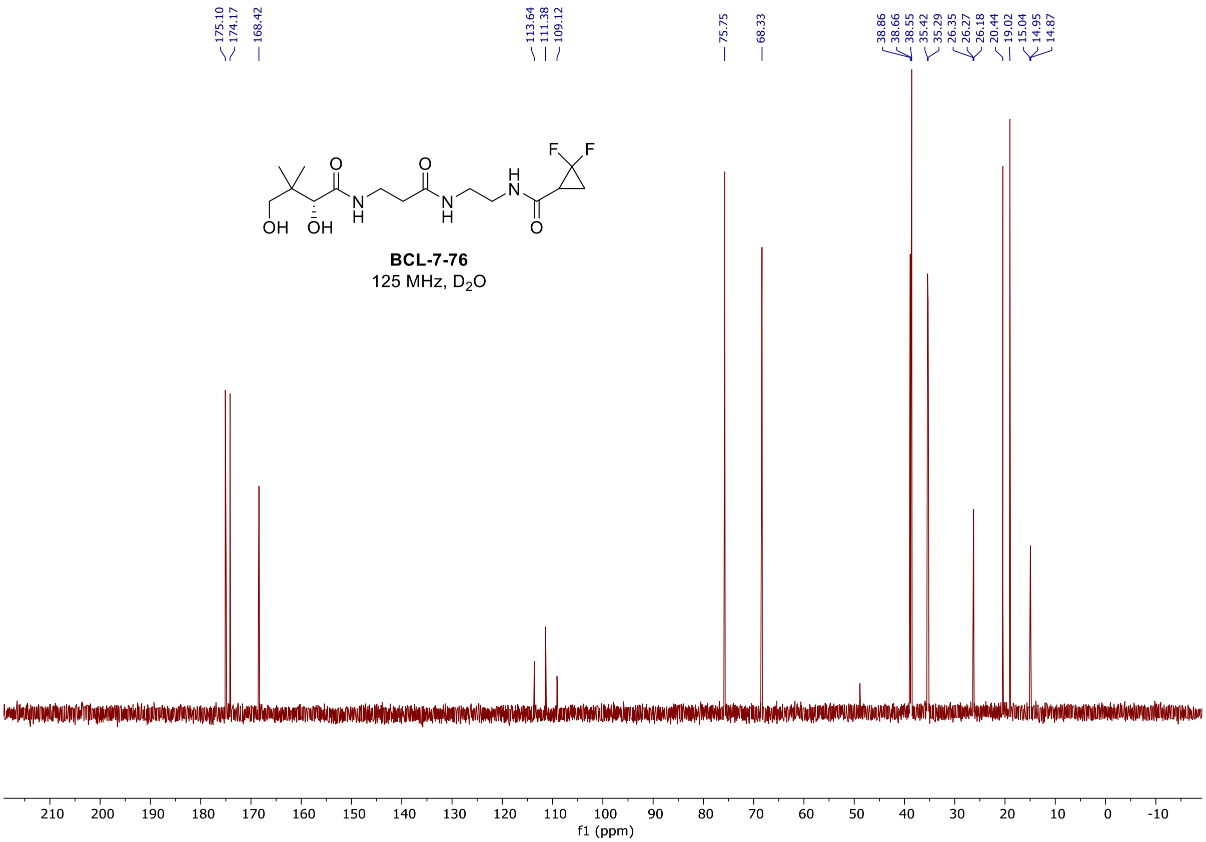

#### $^{1}H$, $^{19}F$, and $^{13}C$ spectra of **BCL-7-70**

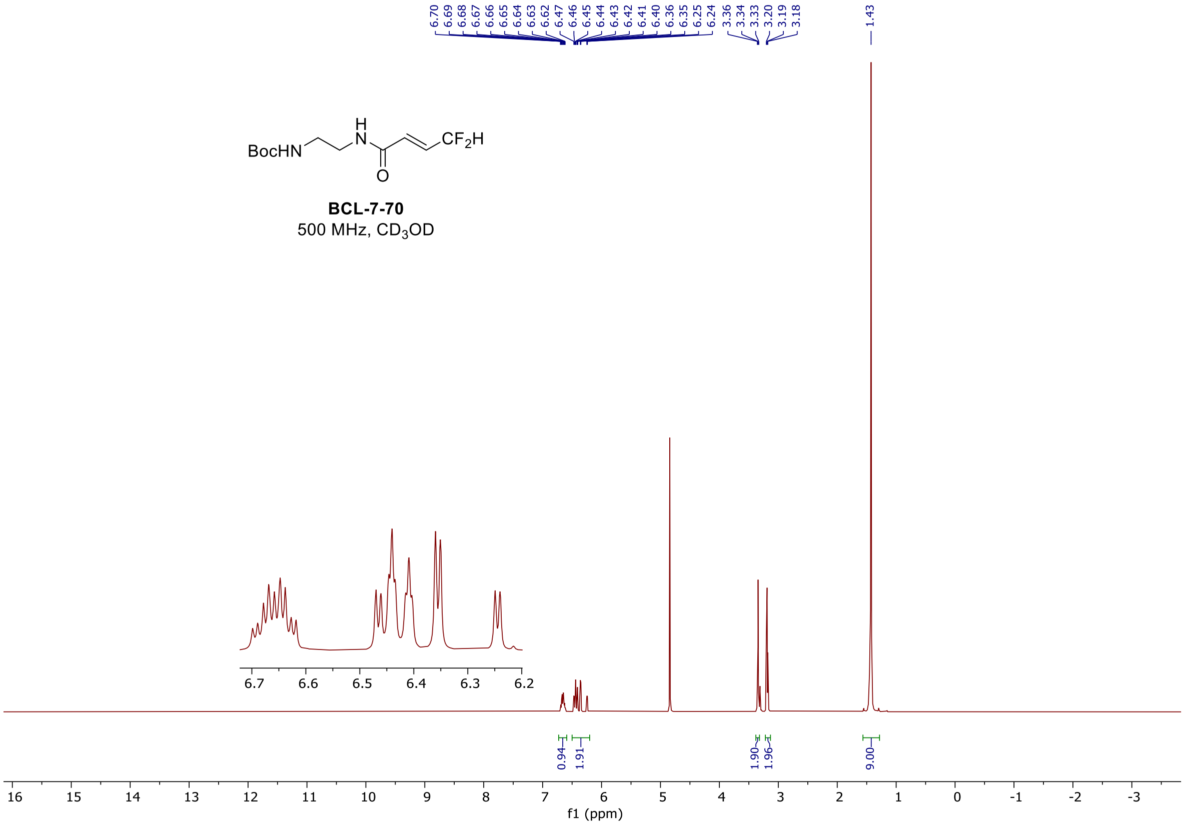

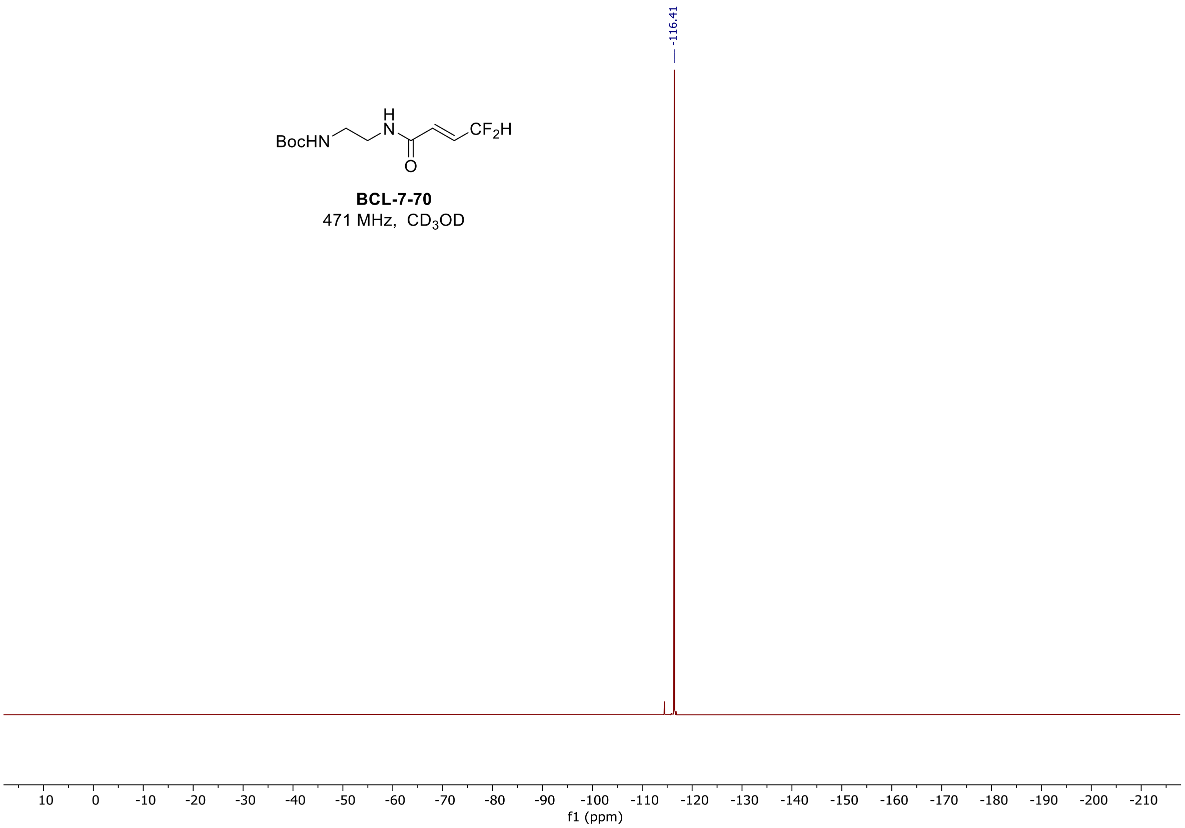
BocHN
H
N
O
CF2H
BCL-7-70
471 MHz, CD3OD
-116.41
10
0
-10
-20
-30
-40
-50
-60
-70
-80
-90
-100
-110
-120
-130
-140
-150
-160
-170
-180
-190
-200
-210
f1 (ppm)

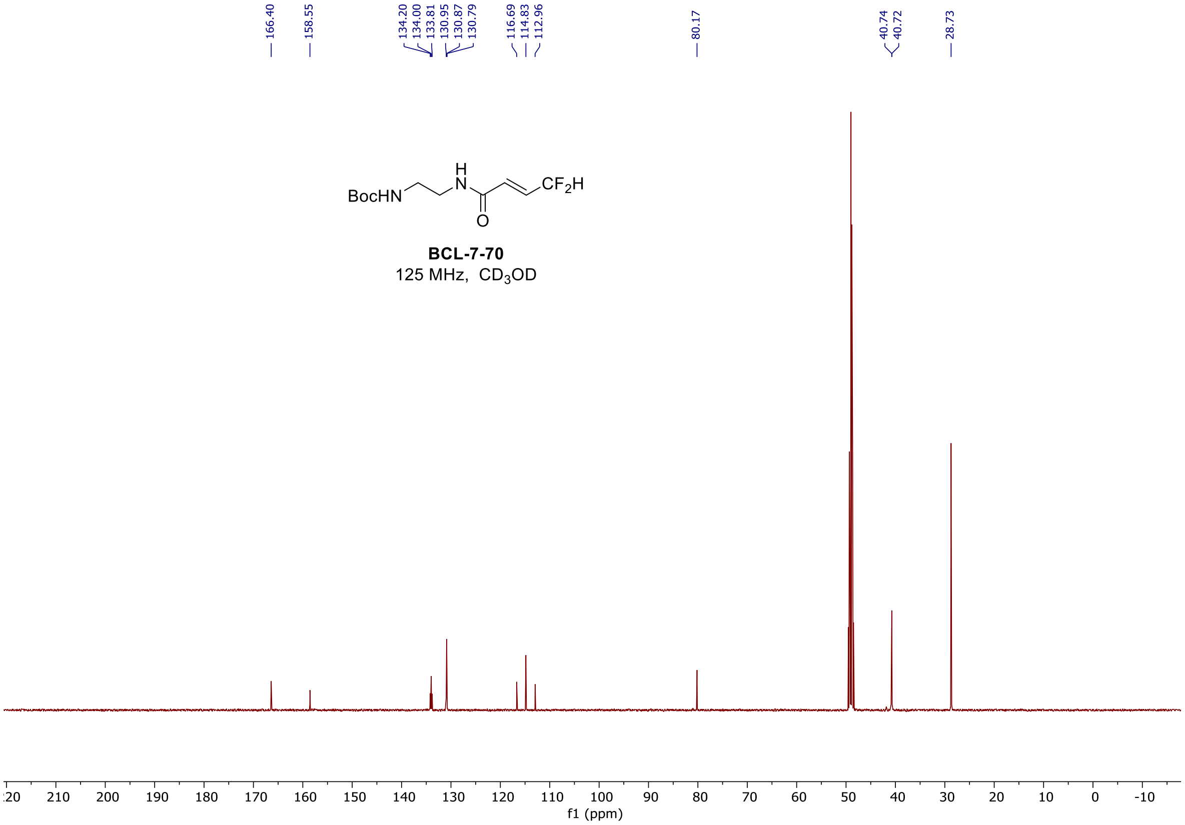
166.40
158.55
134.20
134.00
133.81
130.95
130.87
130.79
116.69
114.83
112.96
80.17
40.74
40.72
28.73
BocHN
H
N
O
CF2H
BCL-7-70
125 MHz, CD3OD
210
200
190
180
170
160
150
140
130
120
110
100
90
80
70
60
50
40
30
20
10
0
-10
f1 (ppm)

[1]H, [19]F, and [13]C spectra of **BCL-7-73**

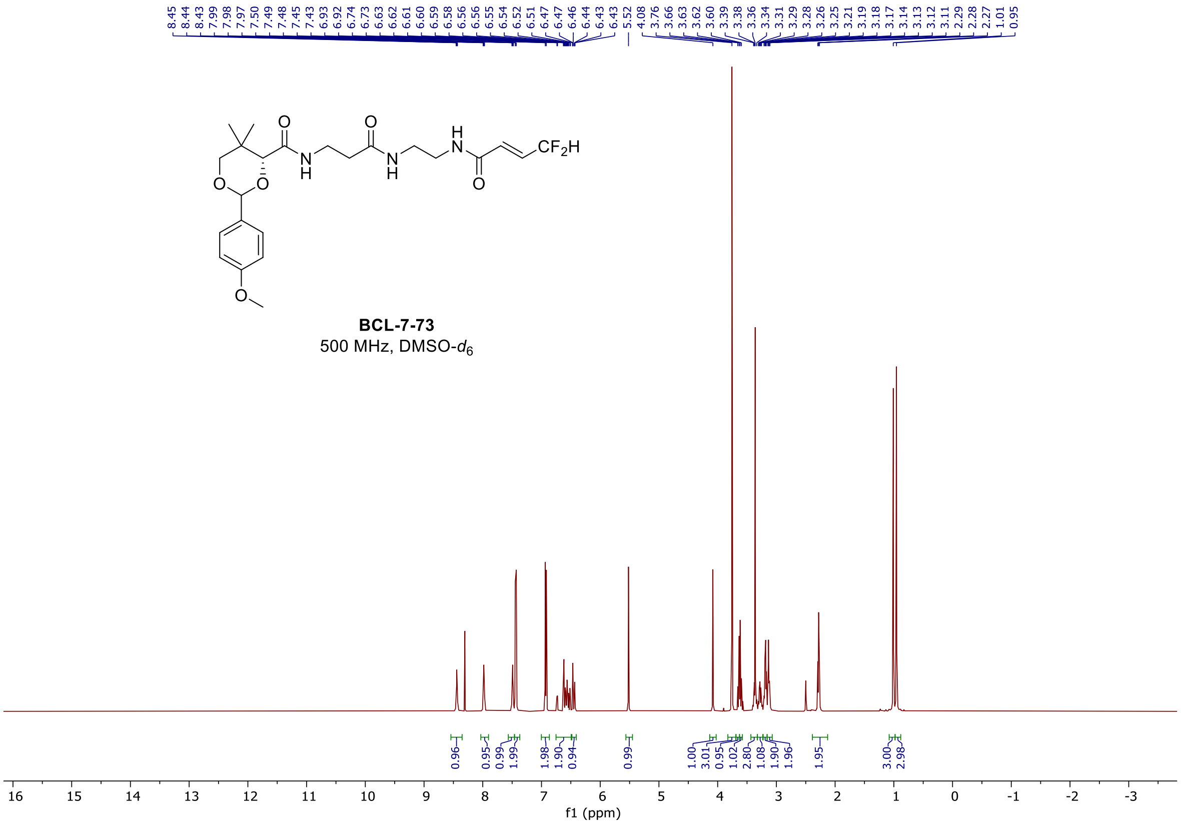

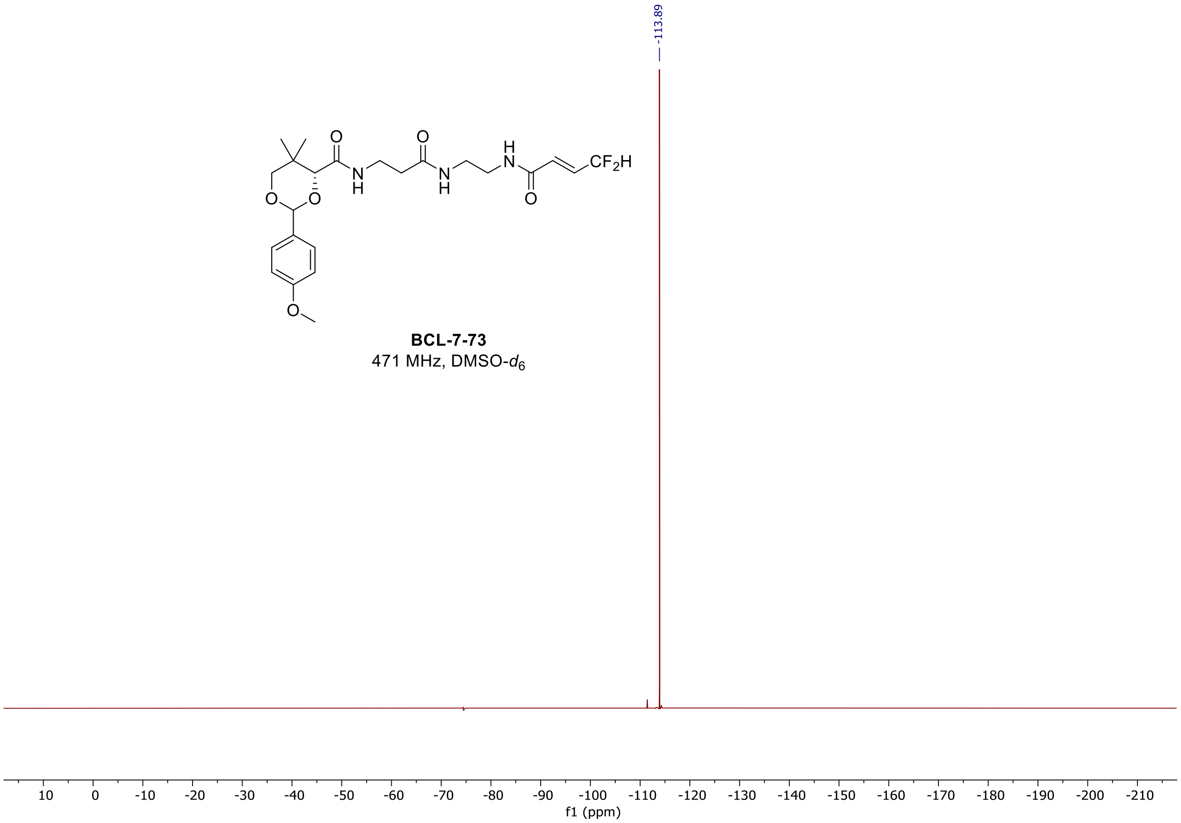

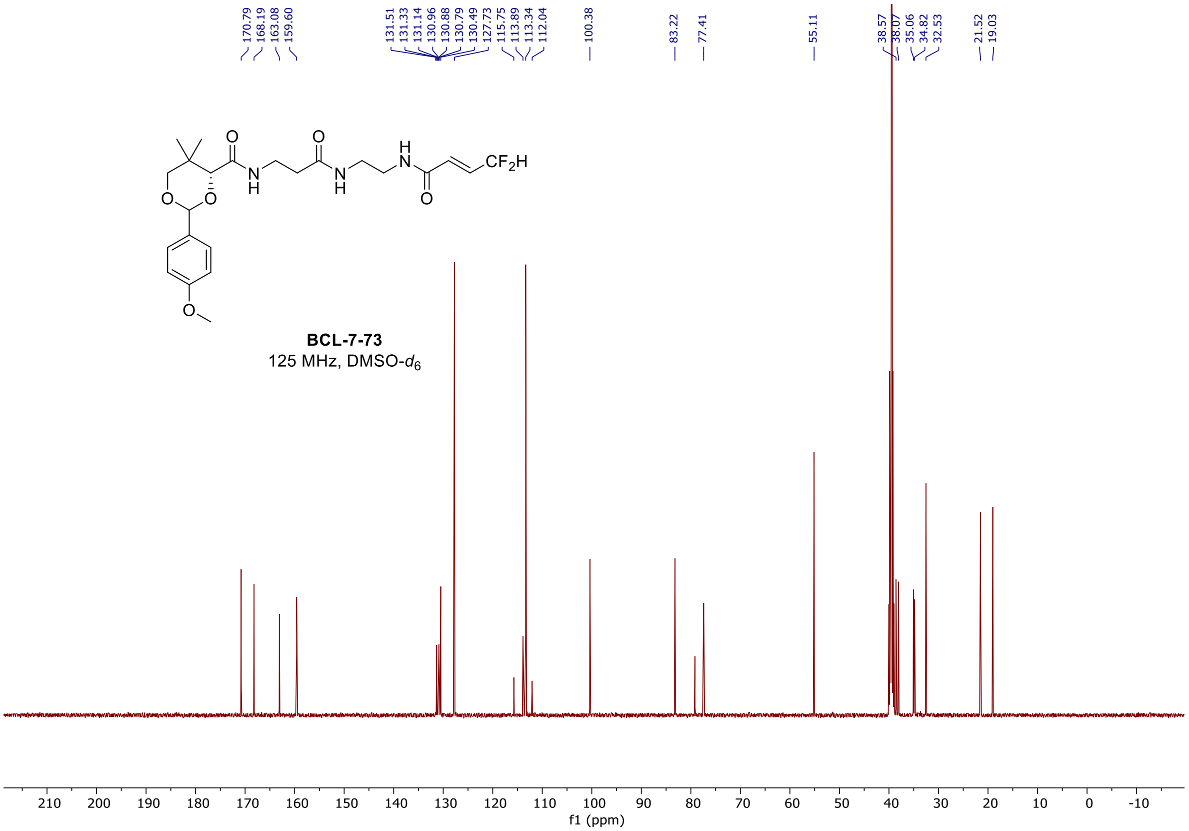

#### $^{1}$H, $^{19}$F, and $^{13}$C spectra of **BCL-7-77**

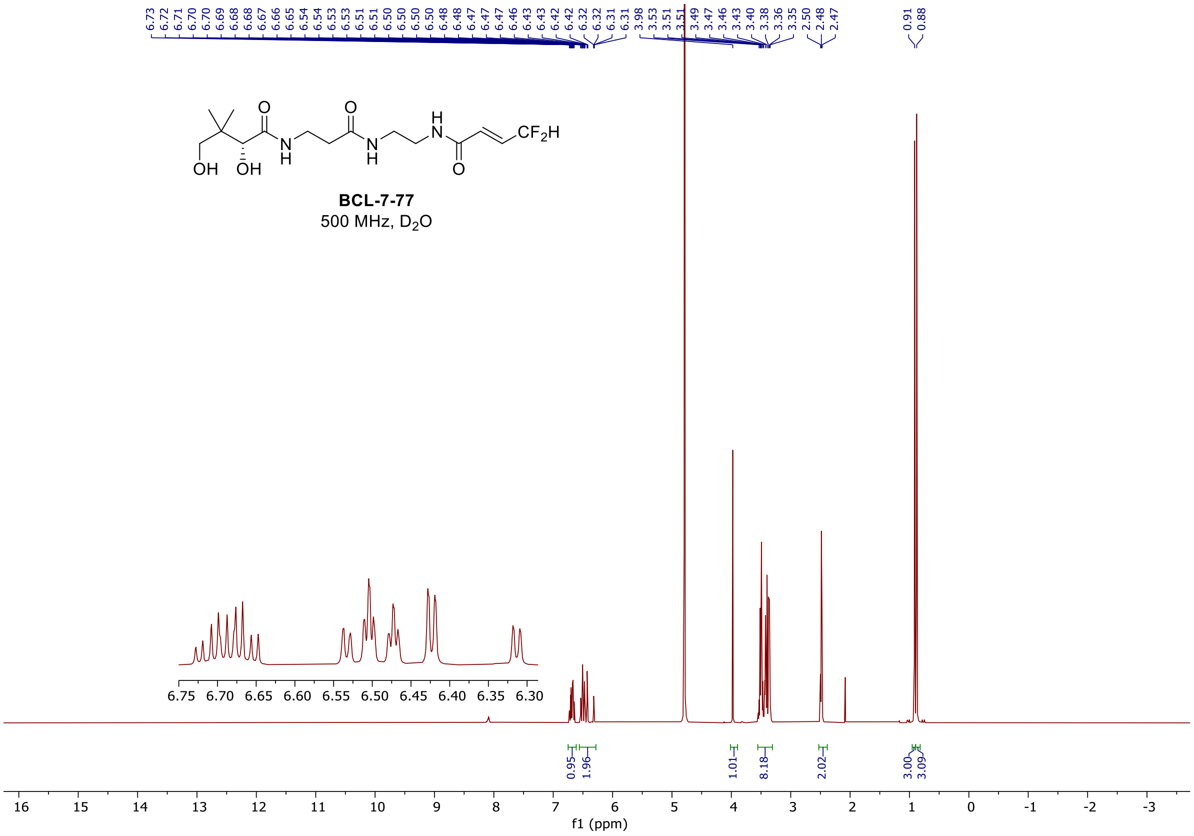

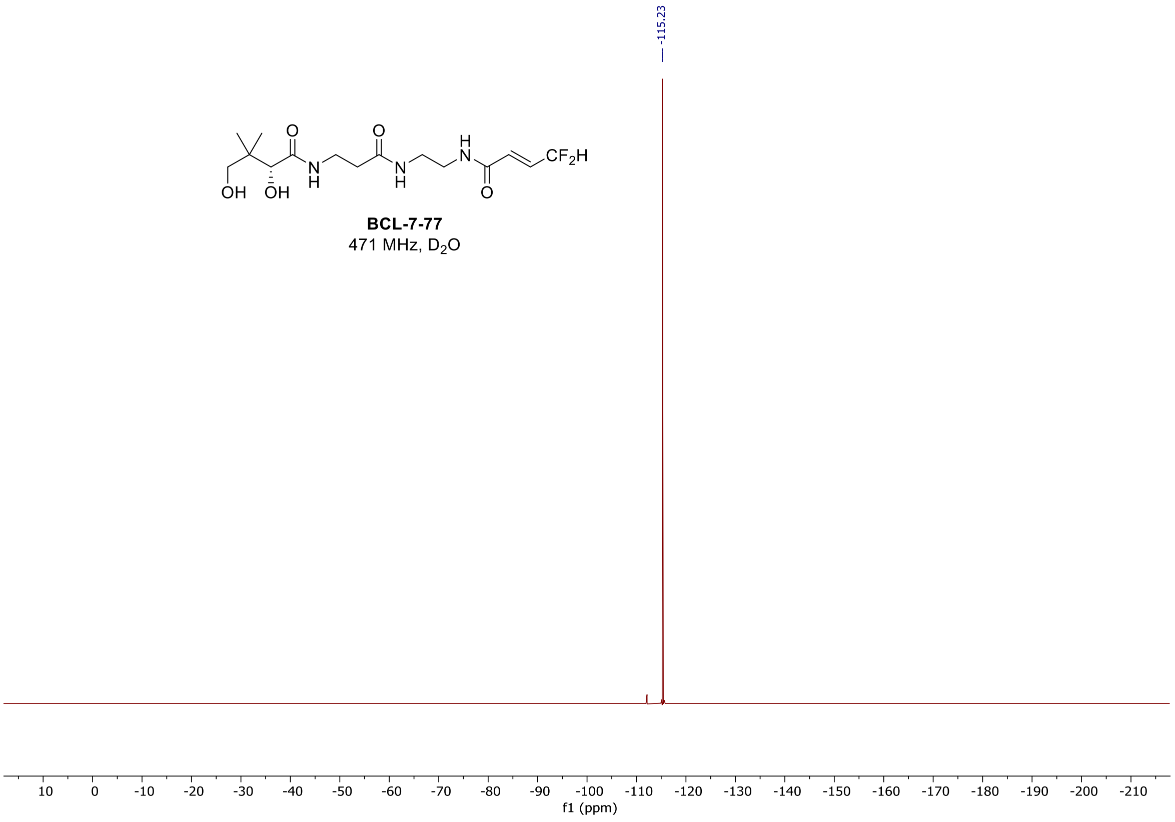
-115.23
BCL-7-77
471 MHz, $D_2O$
f1 (ppm)

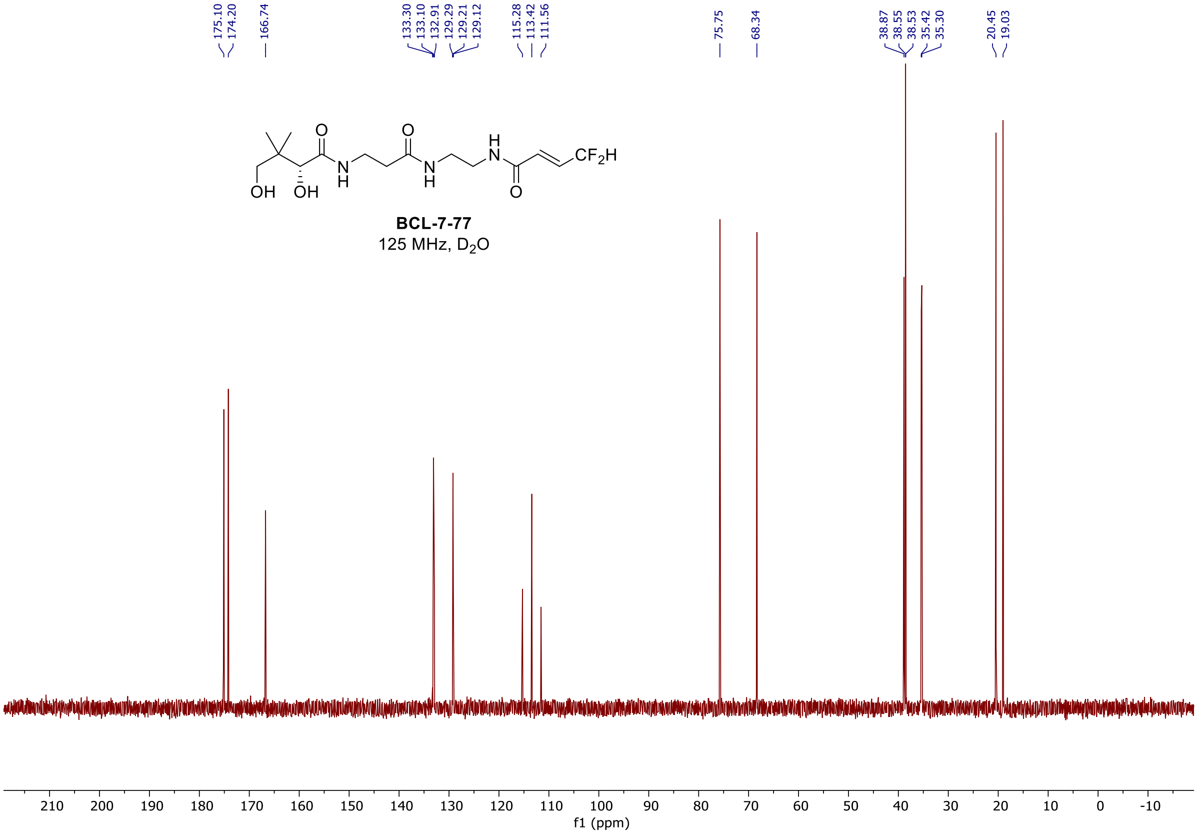
175.10
174.20
166.74
133.30
133.10
132.91
129.29
129.21
129.12
115.28
113.42
111.56
75.75
68.34
38.87
38.55
38.53
35.42
35.30
20.45
19.03
BCL-7-77
125 MHz, $D_2O$
f1 (ppm)

#### $^{1}H$ and $^{13}C\{^{1}H\}$ NMR spectra of **BCL-07-042**

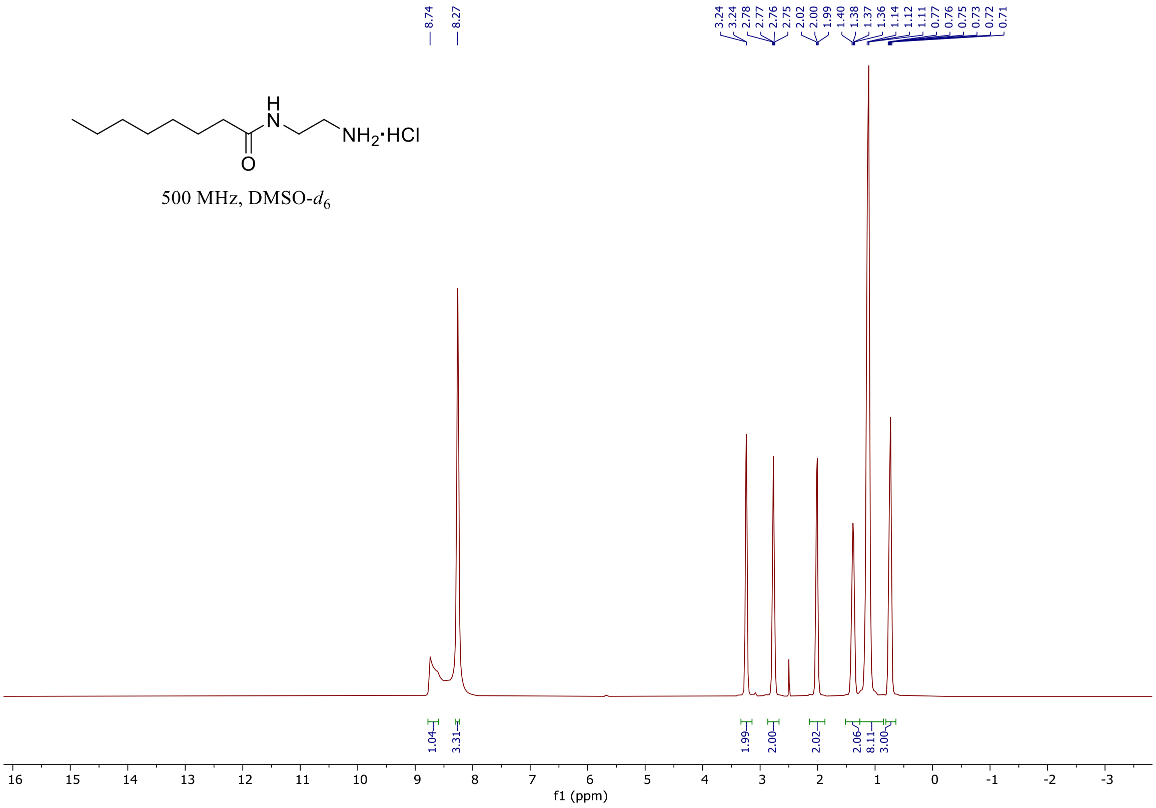

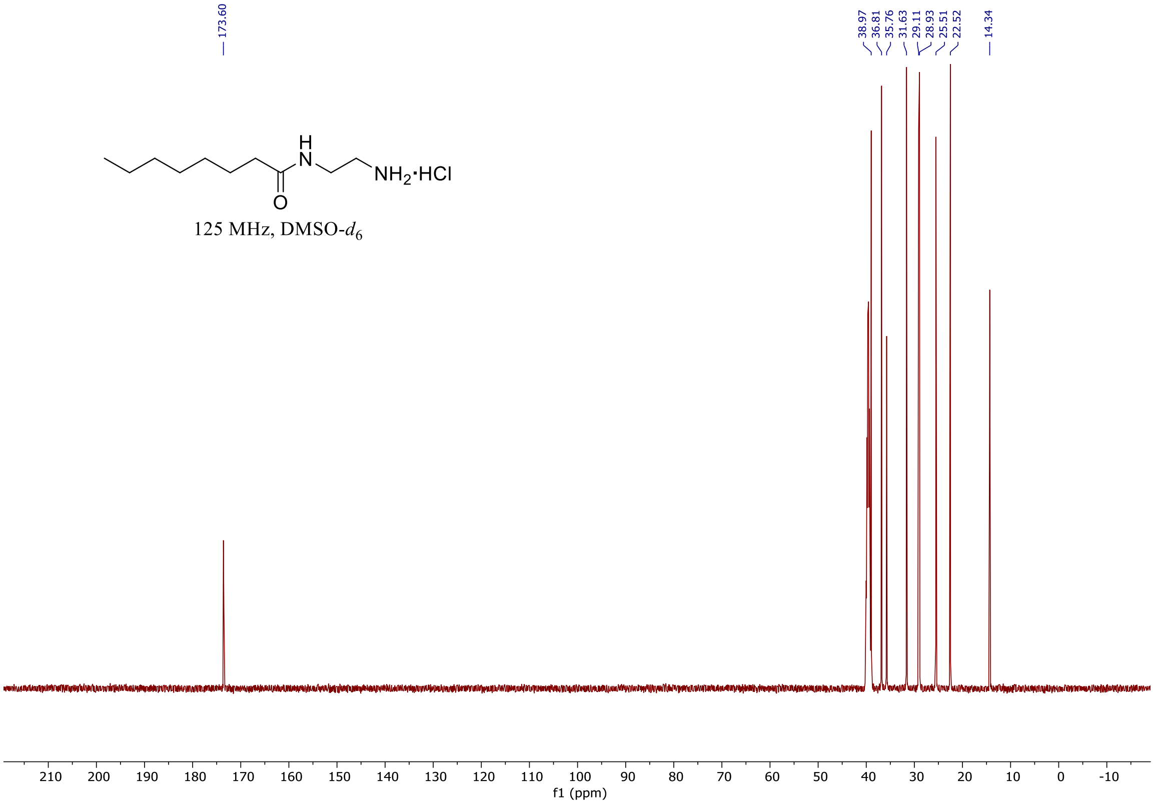

$^{1}H$ and $^{13}C\{^{1}H\}$ NMR spectra of **BCL-07-049**

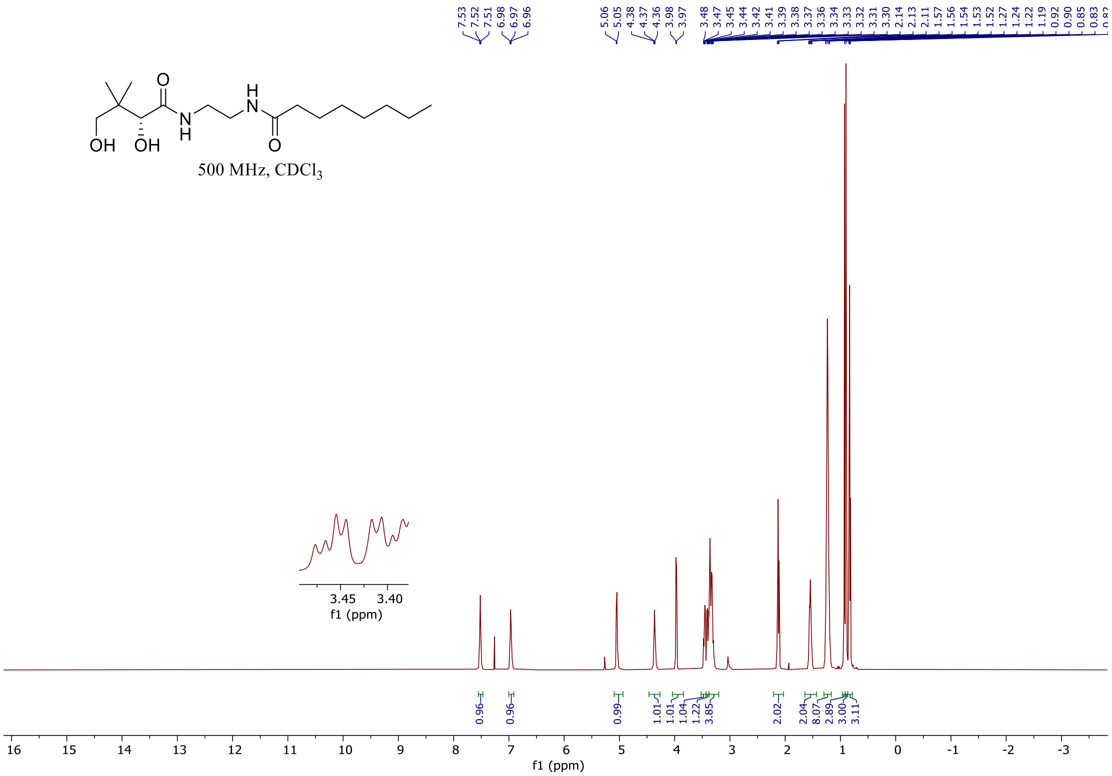

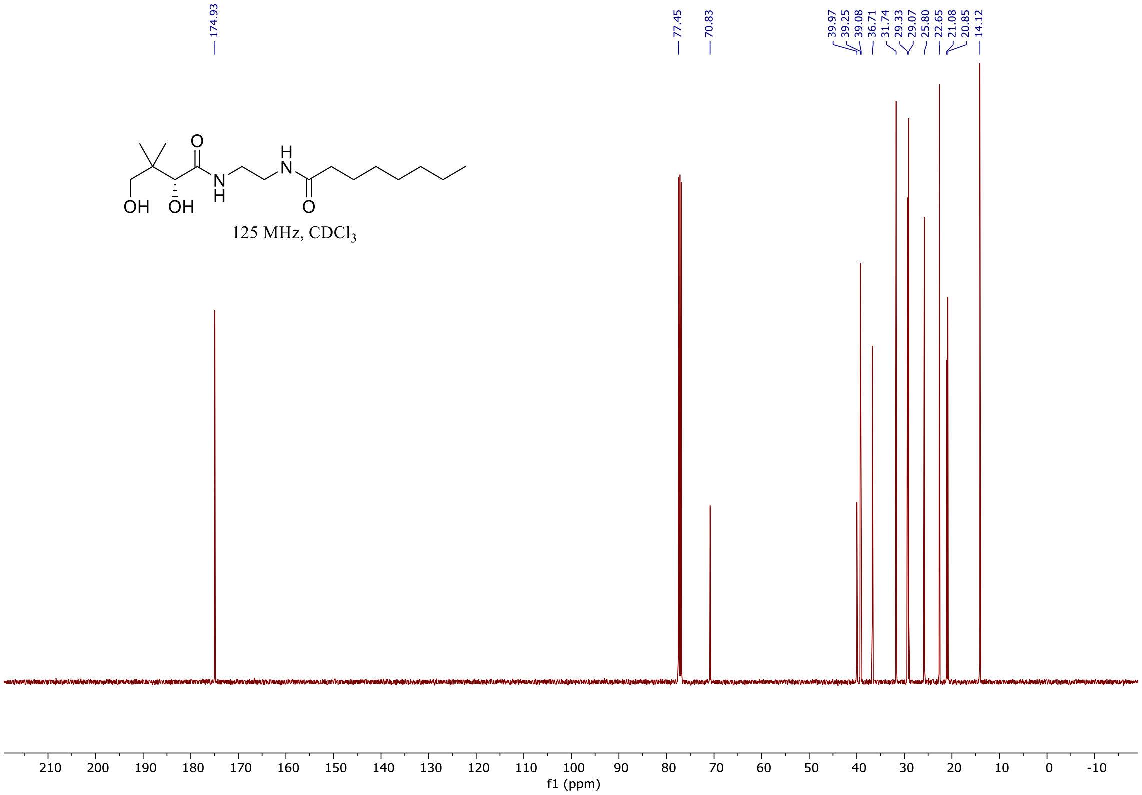

#### $^{1}$H and $^{13}$C spectra of **ABA-1-11**

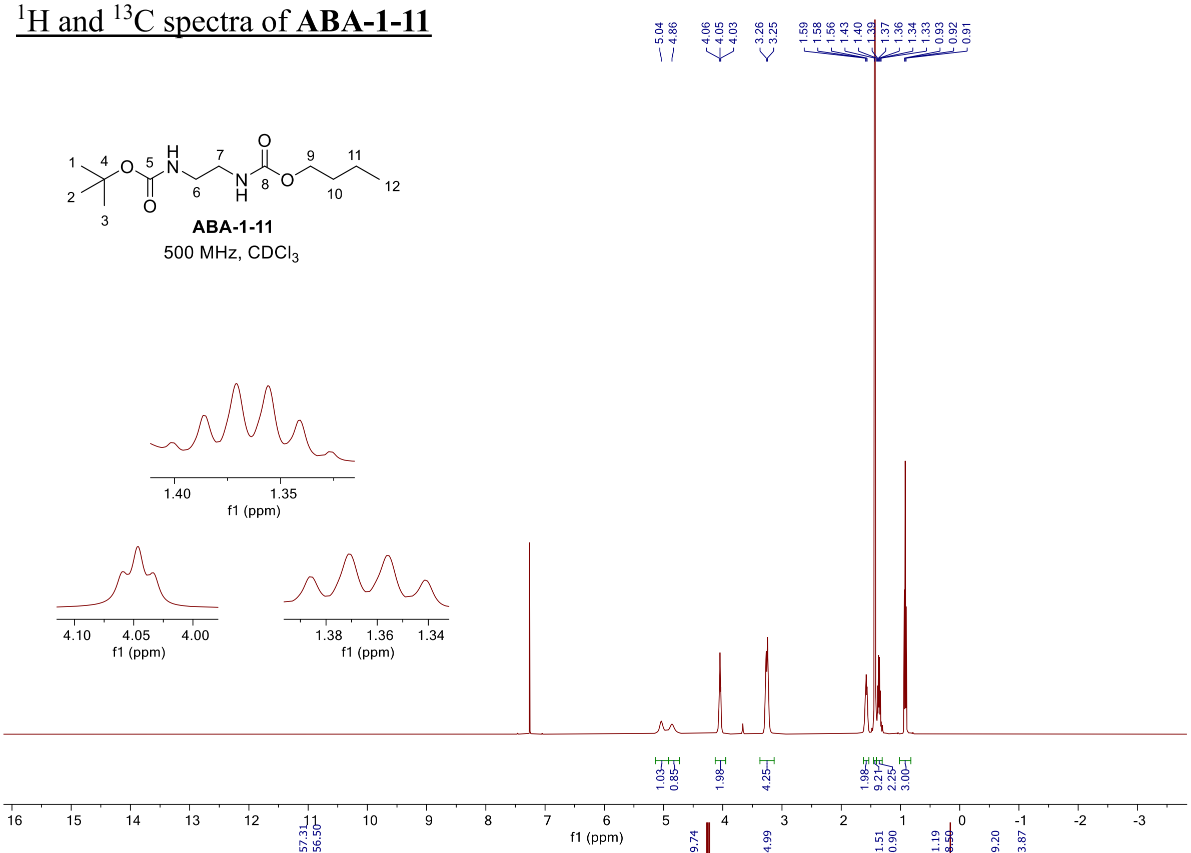

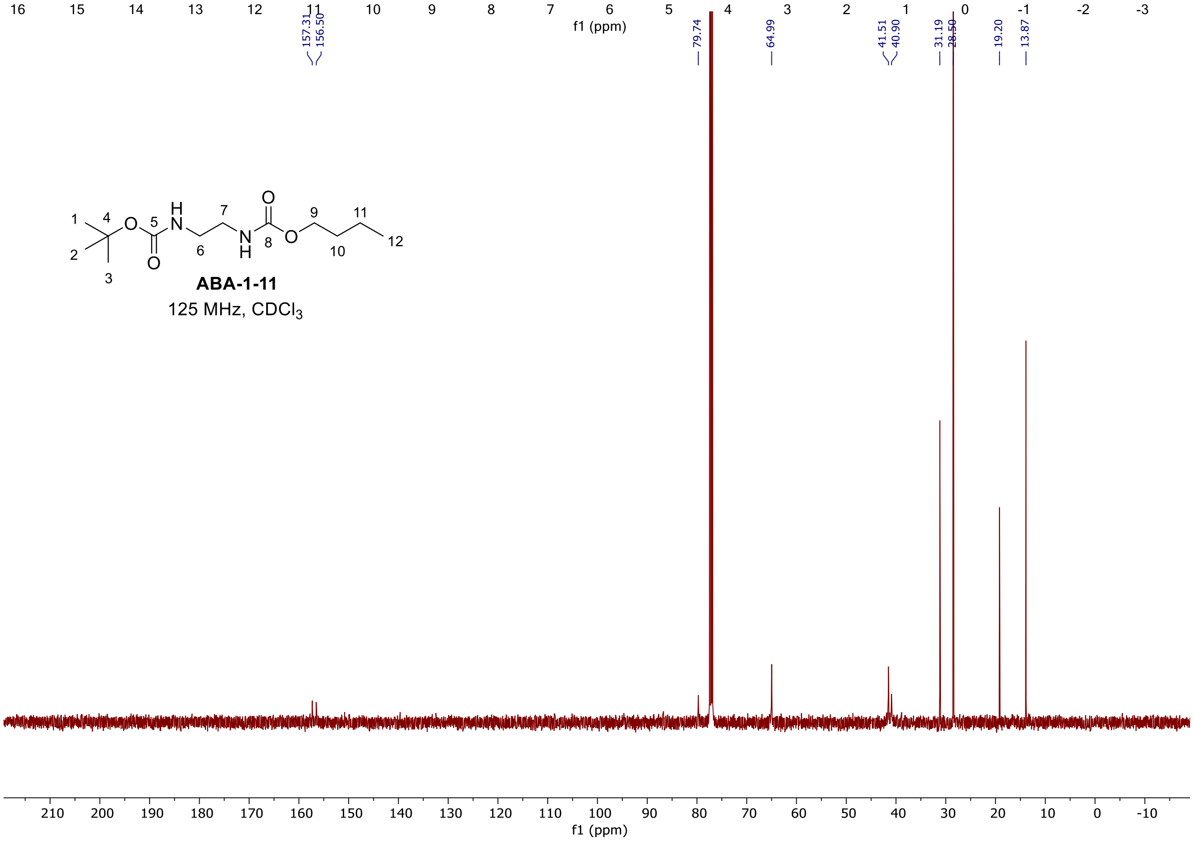

[1]H and [13]C spectra of **ABA-1-13**

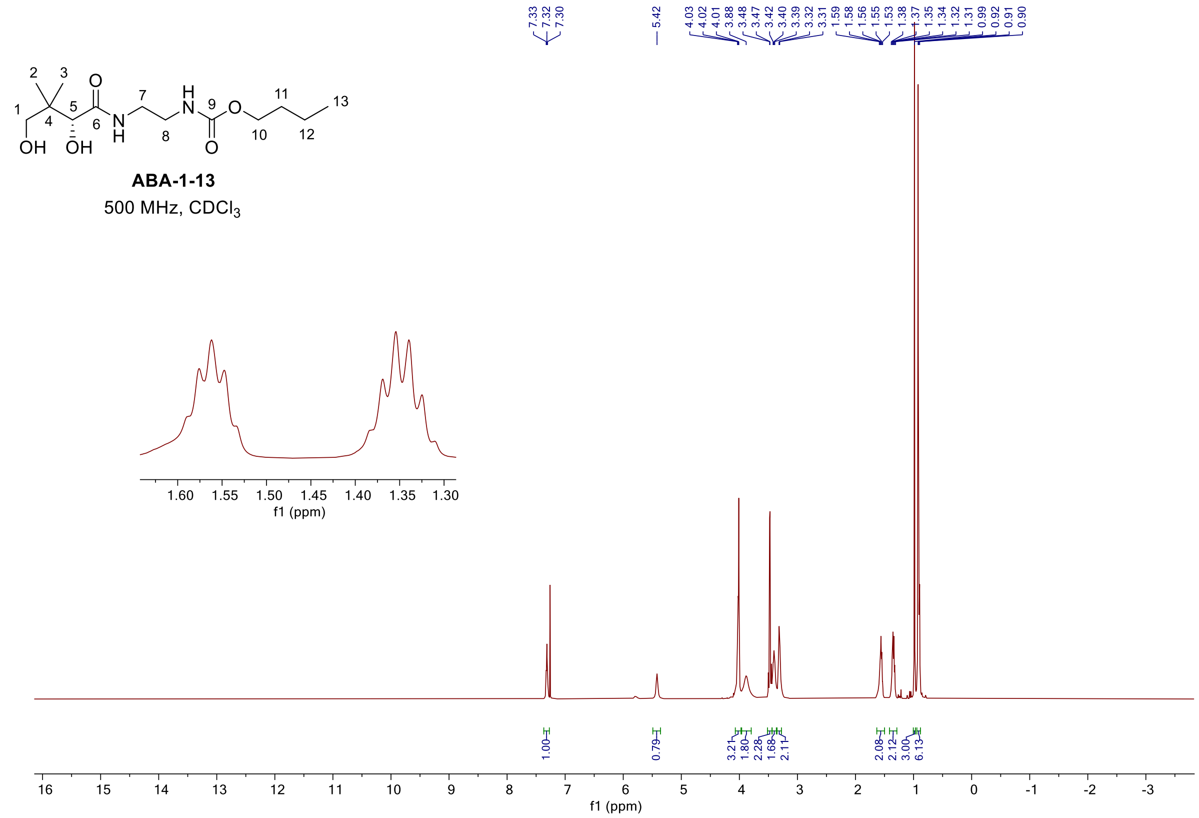

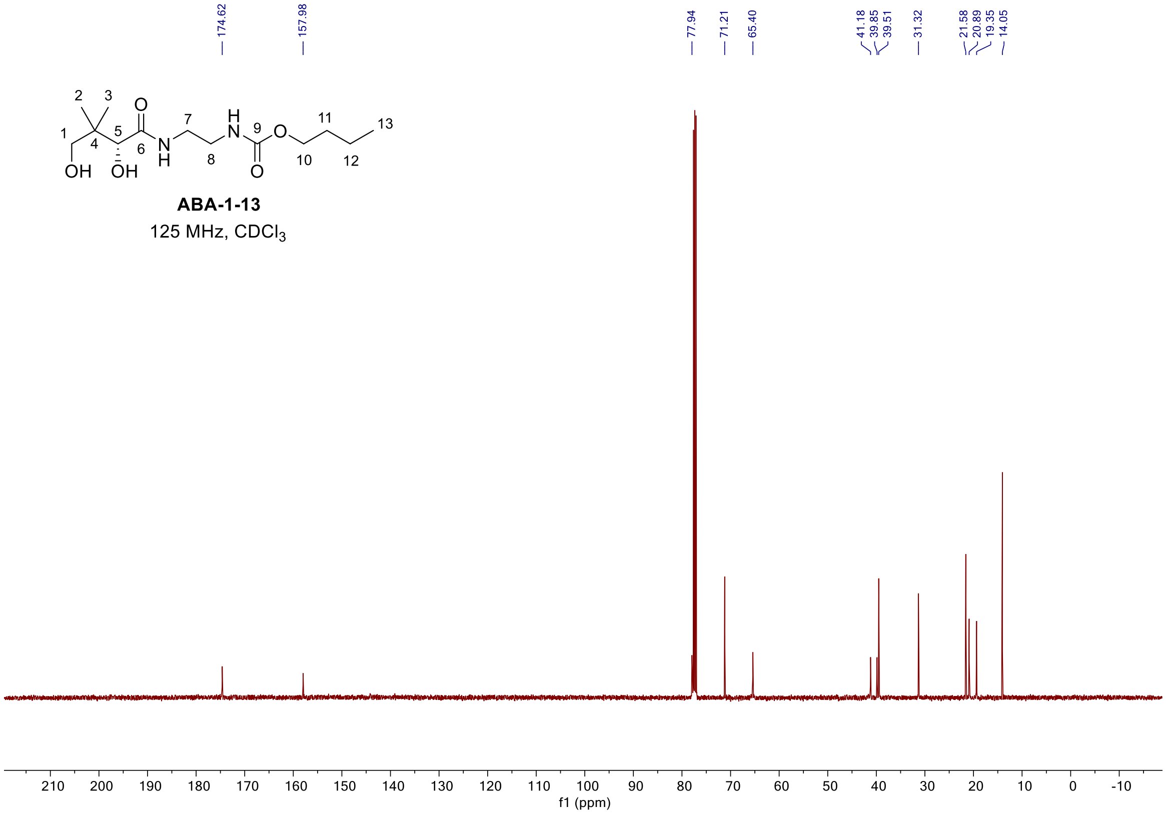

<sup></sup>

### [1]H, [19]F, and [13]C spectra of **ABA-1-1**

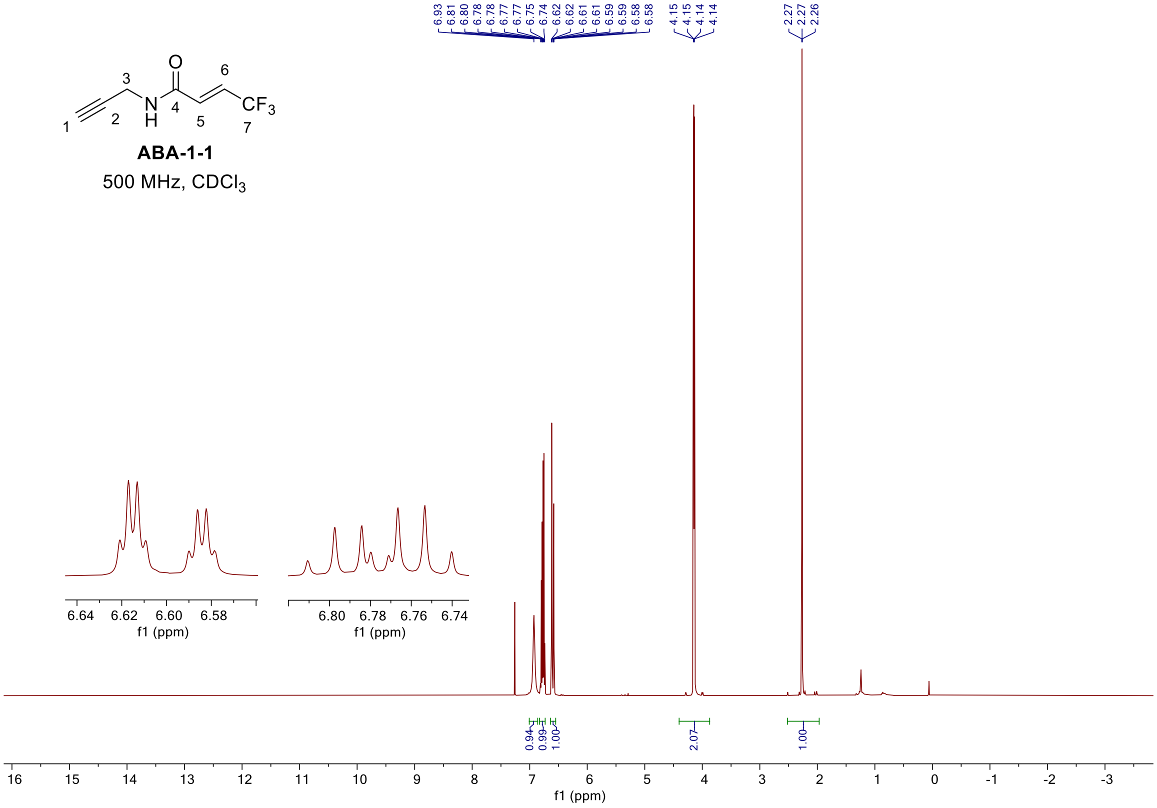

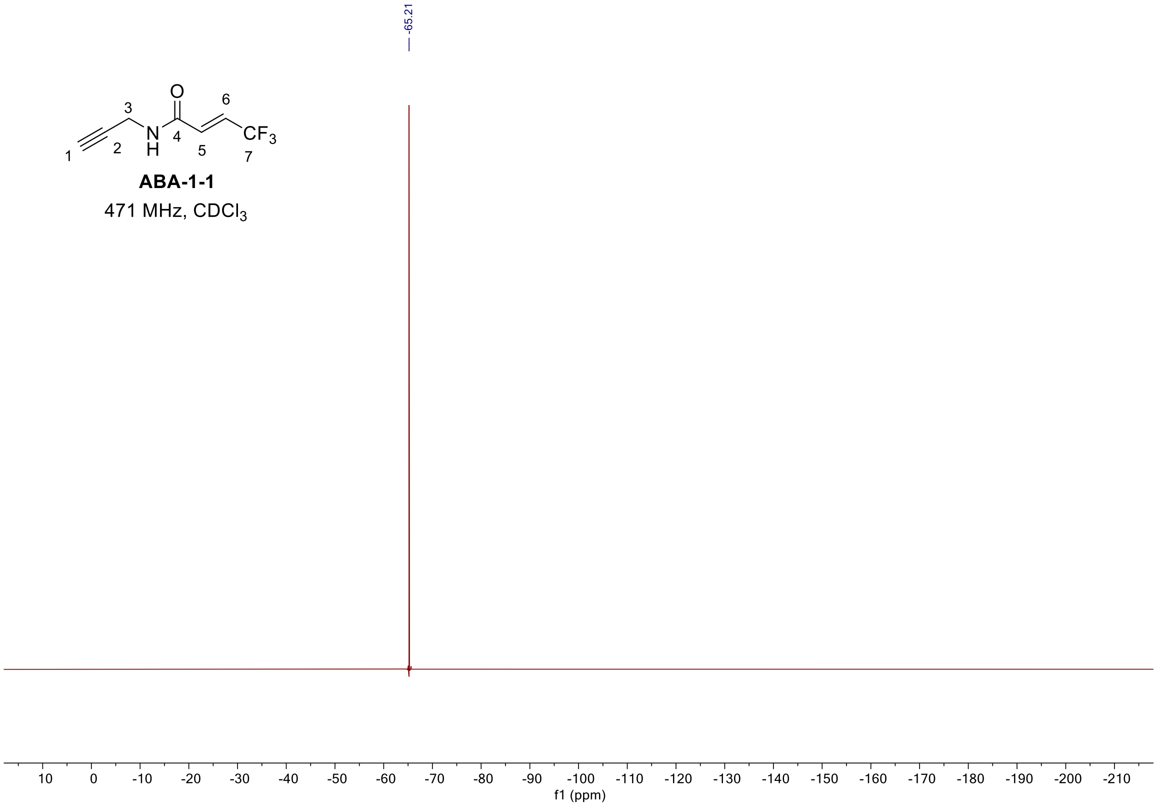

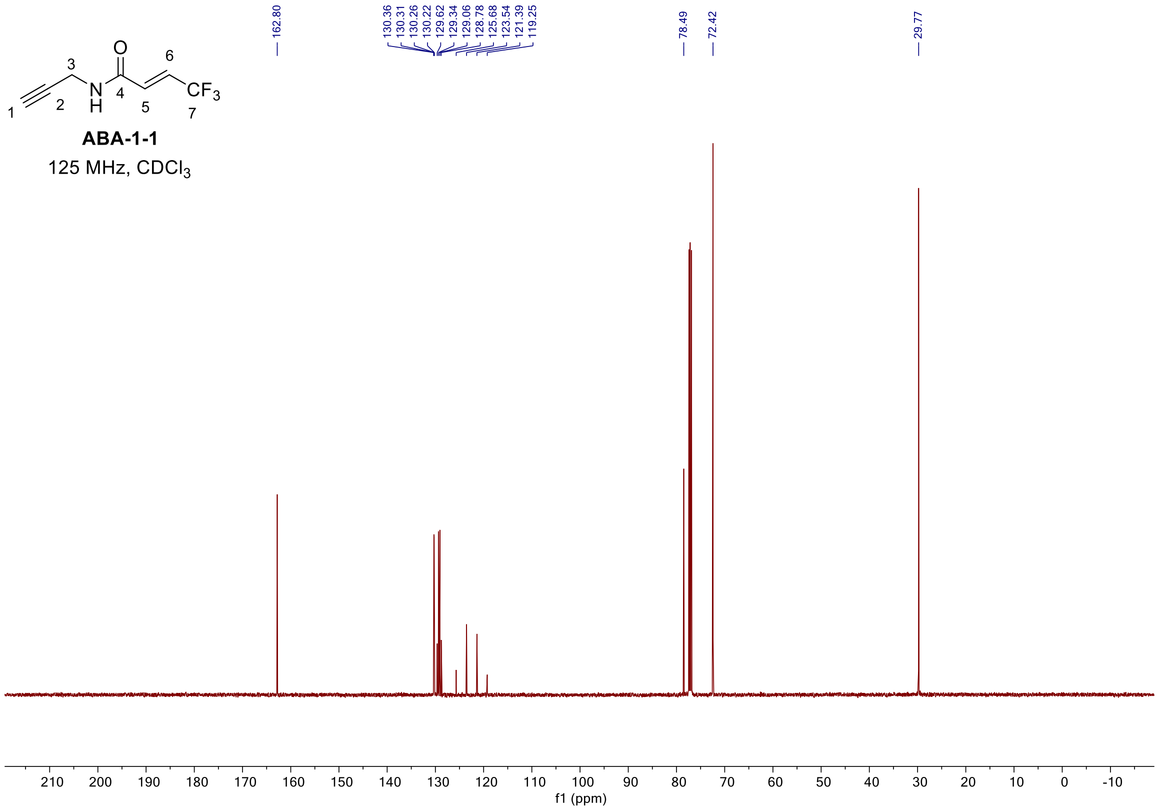

162.80
130.36
130.31
130.26
130.22
129.62
129.34
129.06
128.78
125.68
123.54
121.39
119.25
78.49
72.42
29.77
ABA-1-1
125 MHz, CDCl3
f1 (ppm)

[1]H and [13]C spectra of **ABA-1-3**

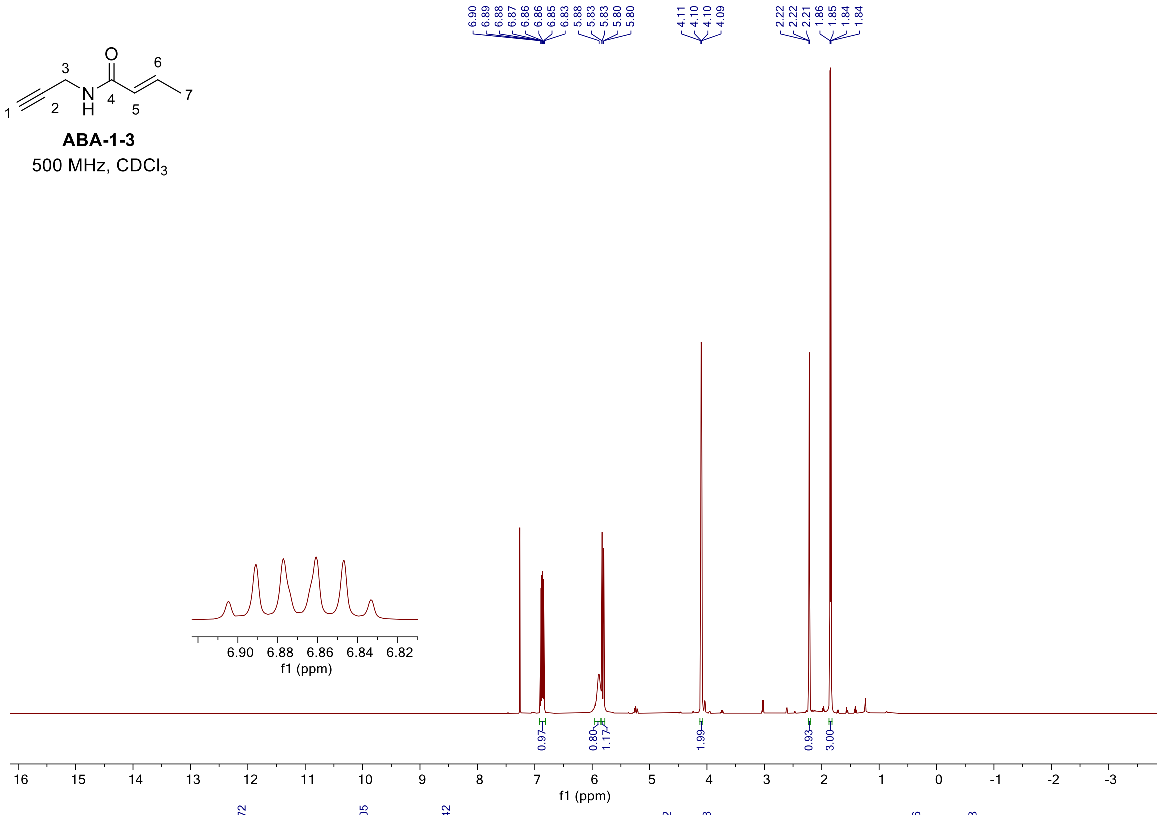

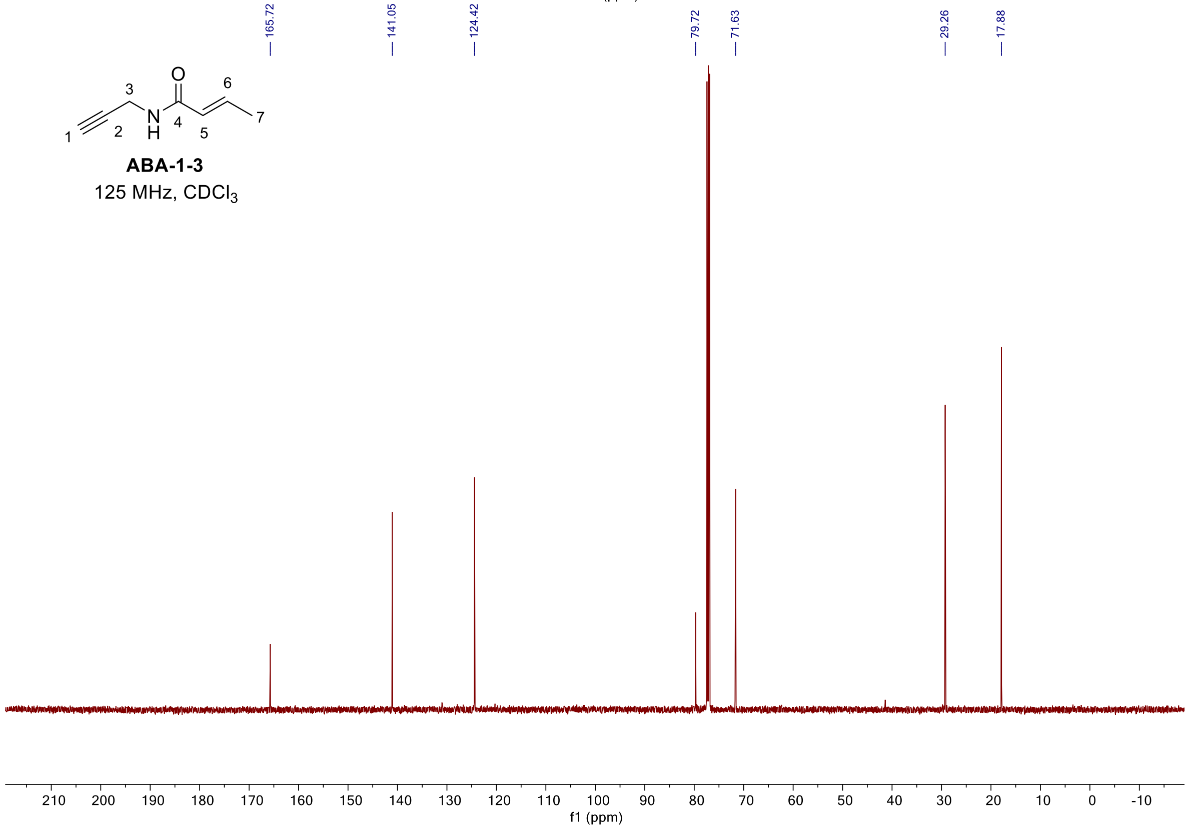

$^1H$, $^{19}F$, and $^{13}C$ spectra of **ABA-1-23**

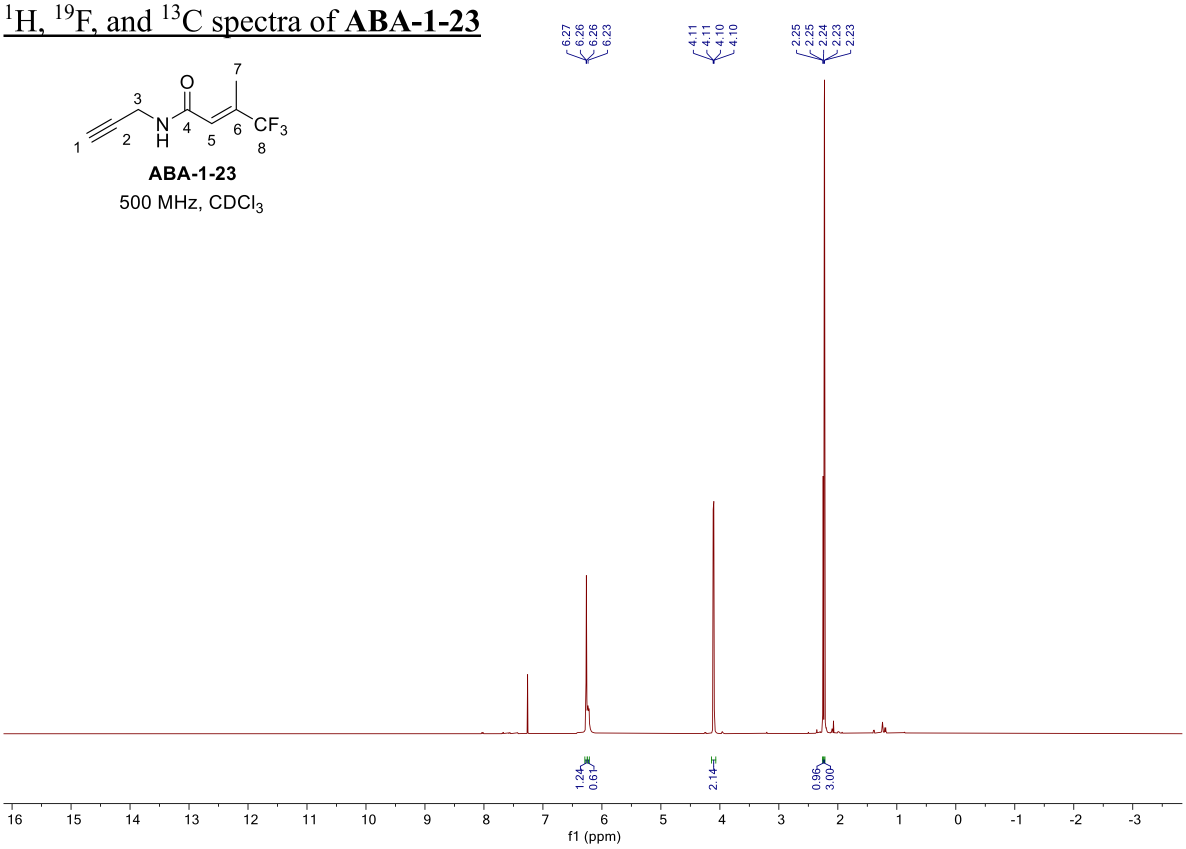

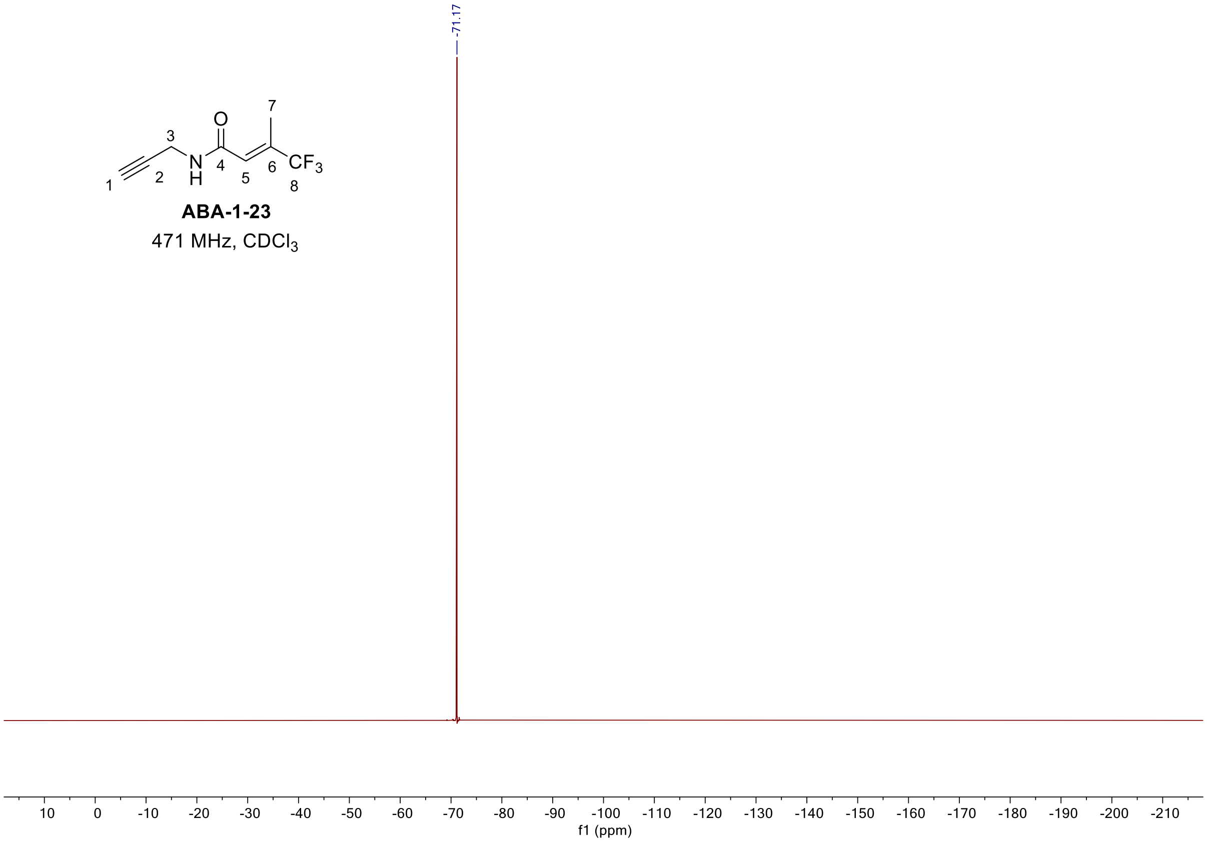

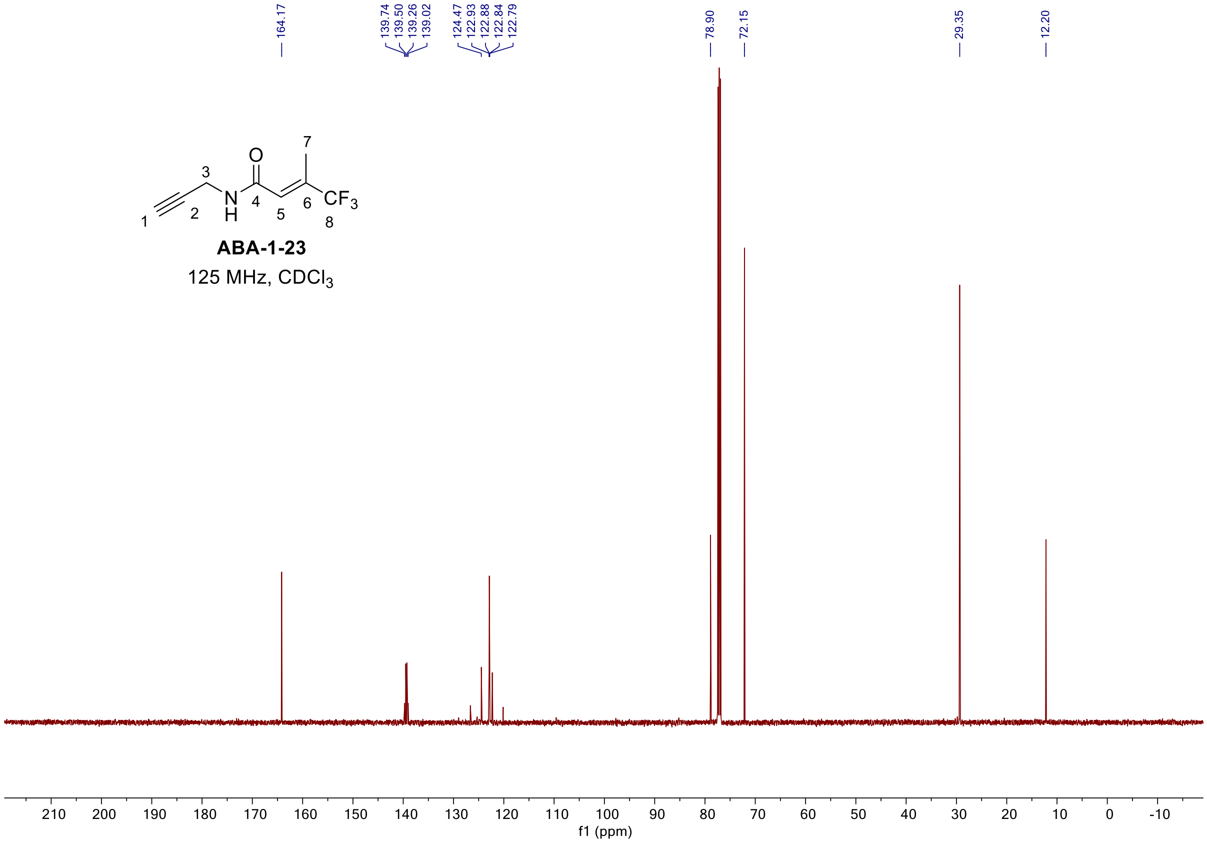
164.17
139.74
139.50
139.26
139.02
124.47
122.93
122.88
122.84
122.79
78.90
72.15
29.35
12.20
ABA-1-23
125 MHz, CDCl3
210
200
190
180
170
160
150
140
130
120
110
100
90
80
70
60
50
40
30
20
10
0
-10
f1 (ppm)

#### $^{1}H$ and $^{13}C$ spectra of **ABA-1-9**

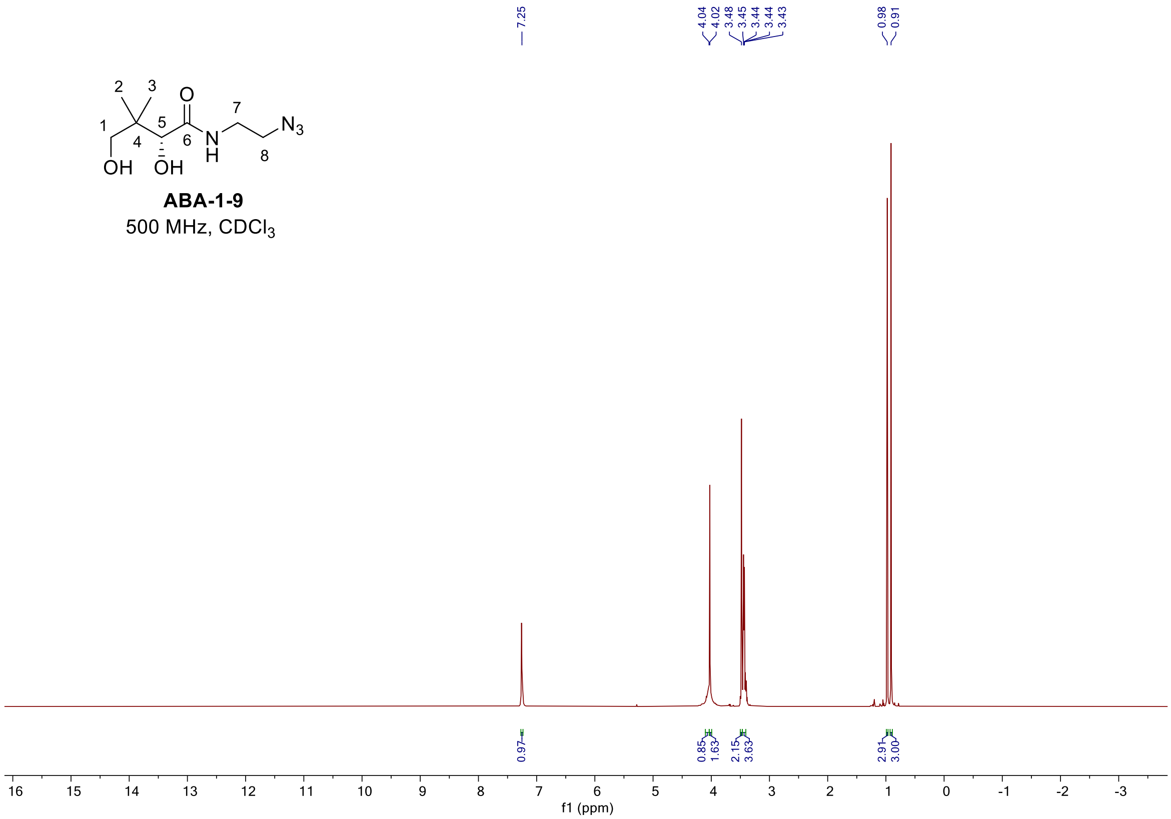

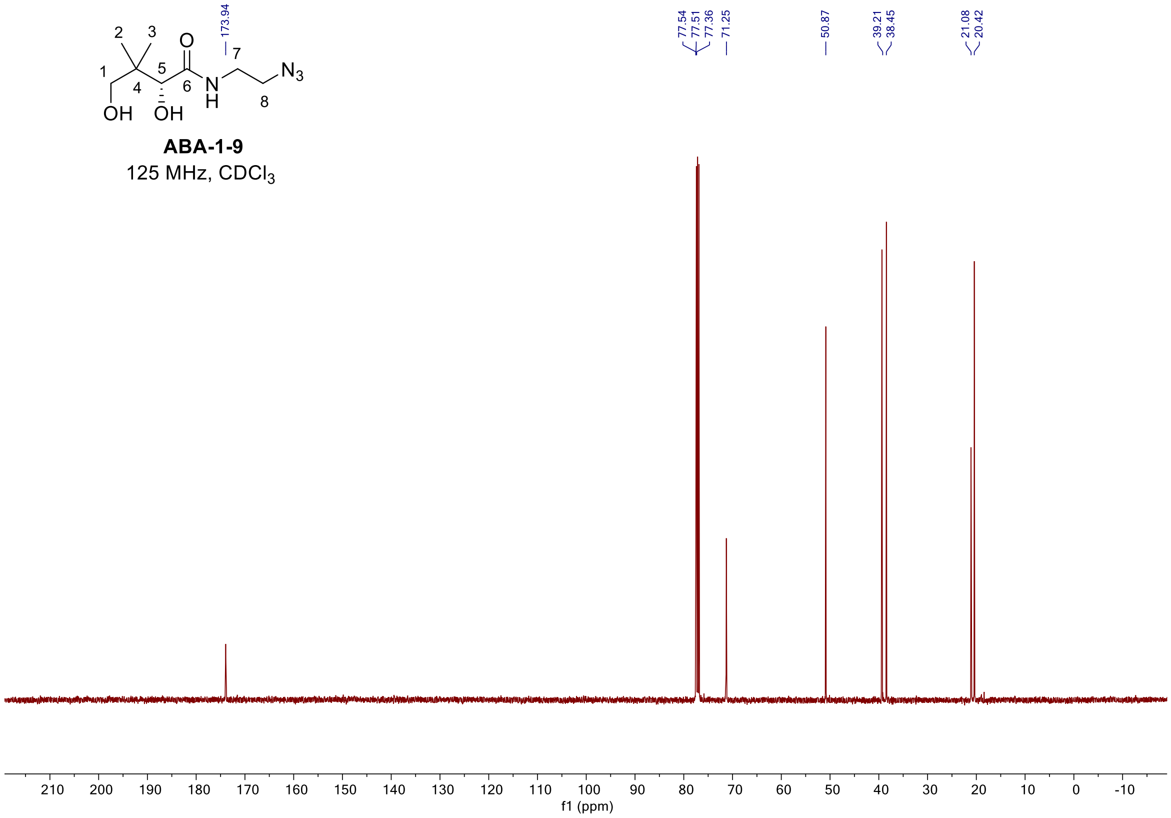

#### $^{1}H$ and $^{13}C$ spectra of **ABA-1-18**

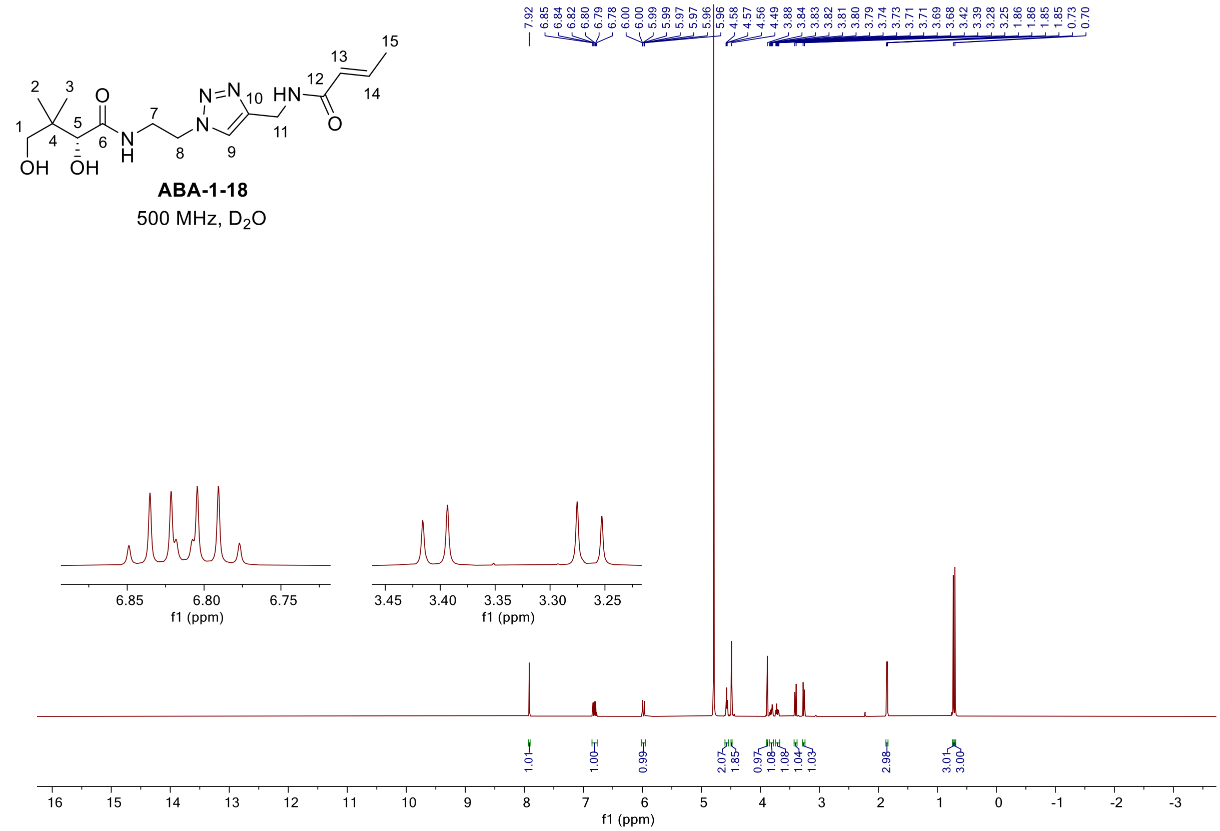

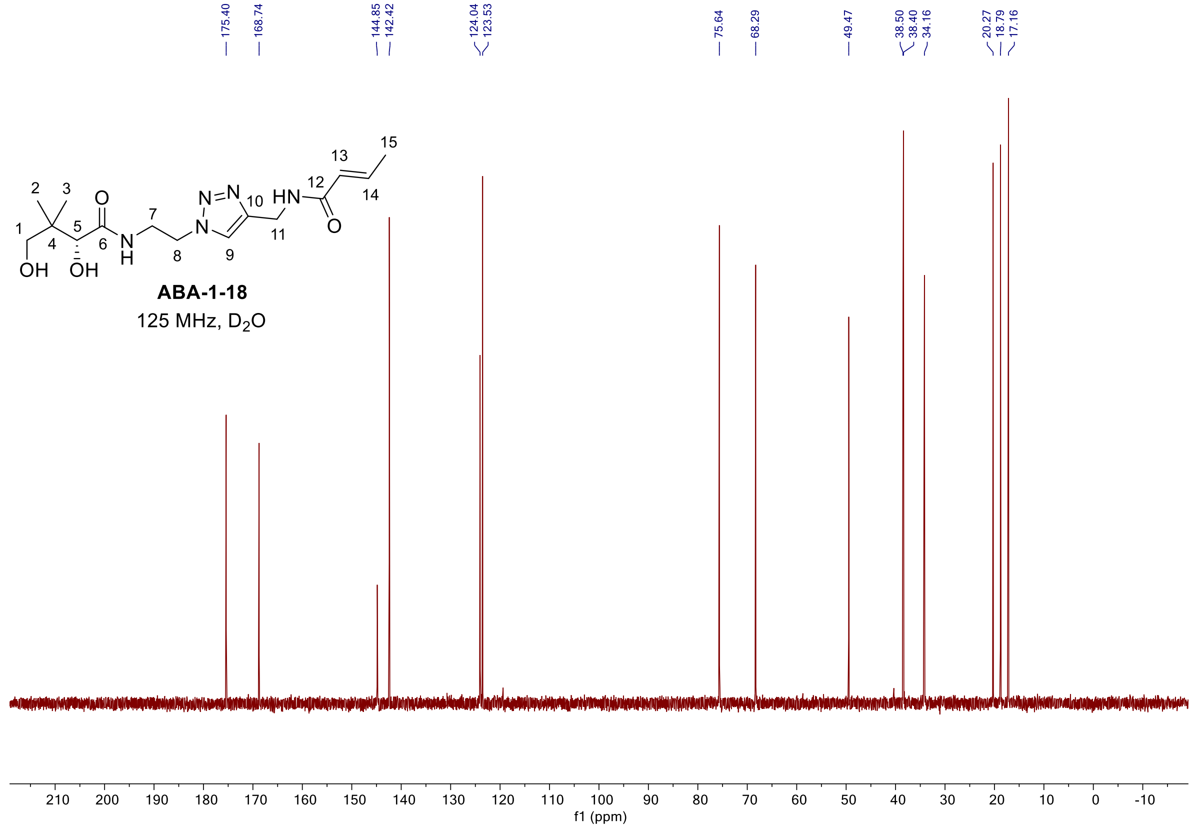

$^{1}H$, $^{19}F$, and $^{13}C$ spectra of **ABA-1-19 (4-12)**

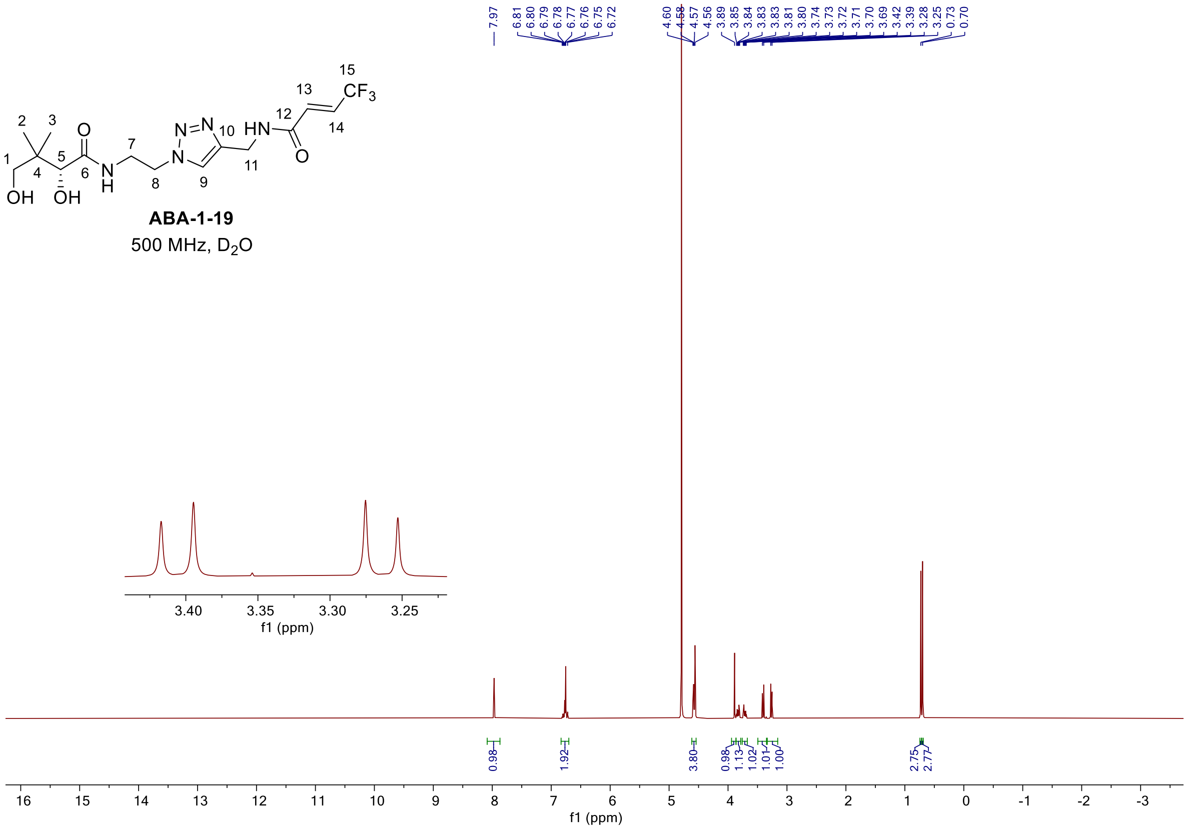

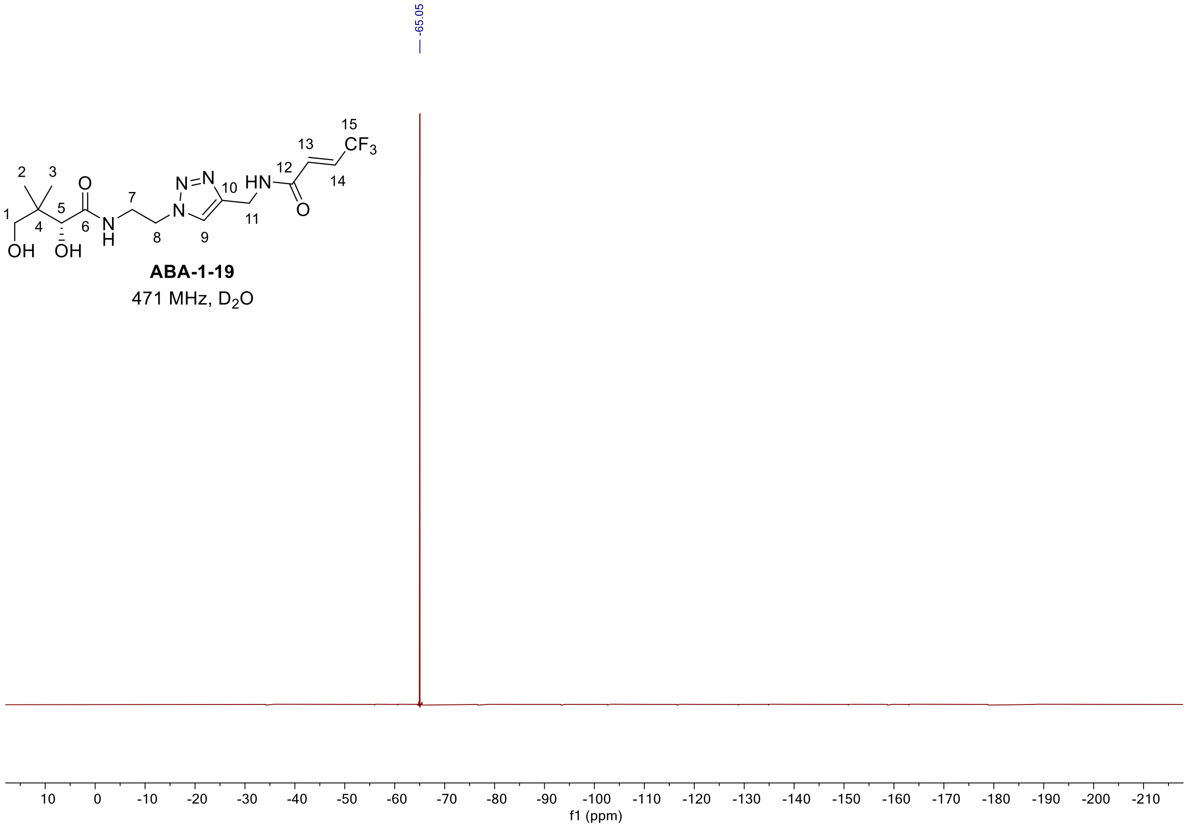

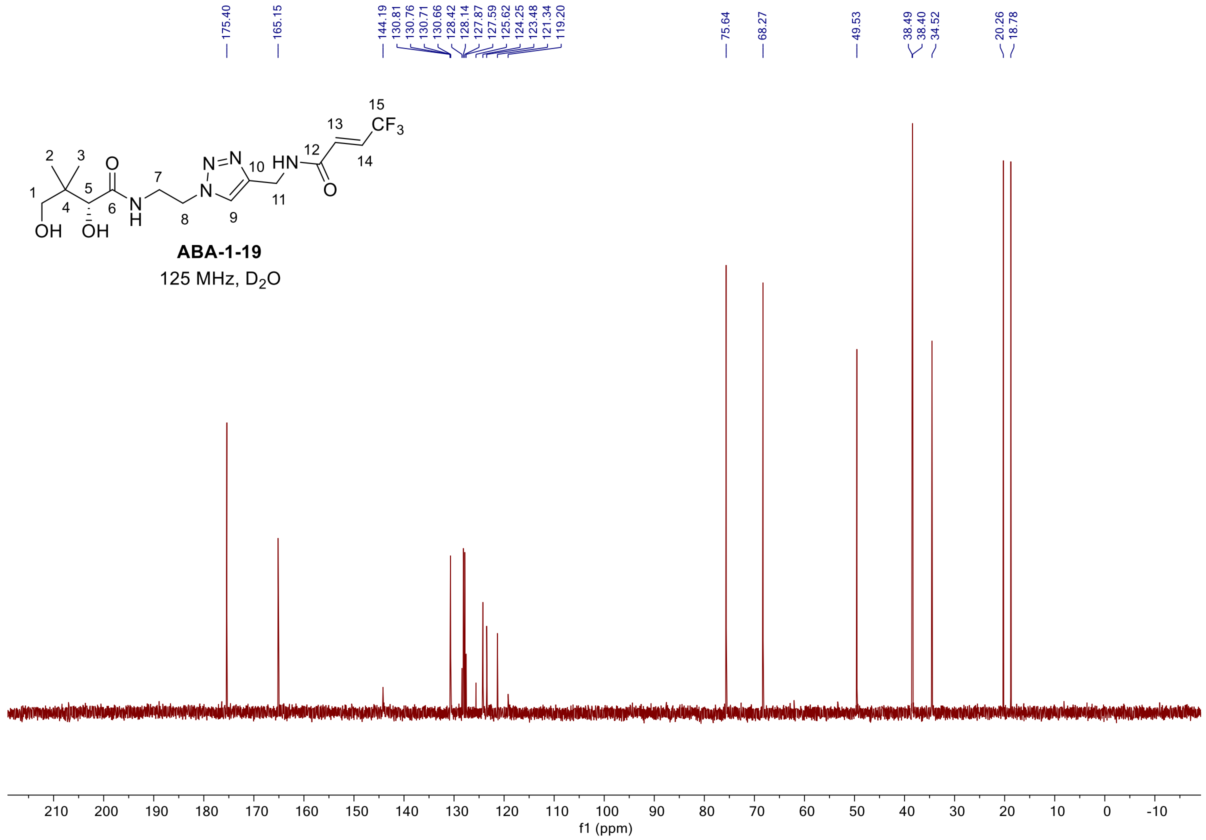

175.40
165.15
144.19
130.81
130.76
130.71
130.66
128.42
128.14
127.87
127.59
125.62
124.25
123.48
121.34
119.20
75.64
68.27
49.53
38.49
38.40
34.52
20.26
18.78
ABA-1-19
125 MHz, D2O
210 200 190 180 170 160 150 140 130 120 110 100 90 80 70 60 50 40 30 20 10 0 -10
f1 (ppm)

$^{1}H$, $^{19}F$, and $^{13}C$ spectra of **ABA-1-26 (4-13)**

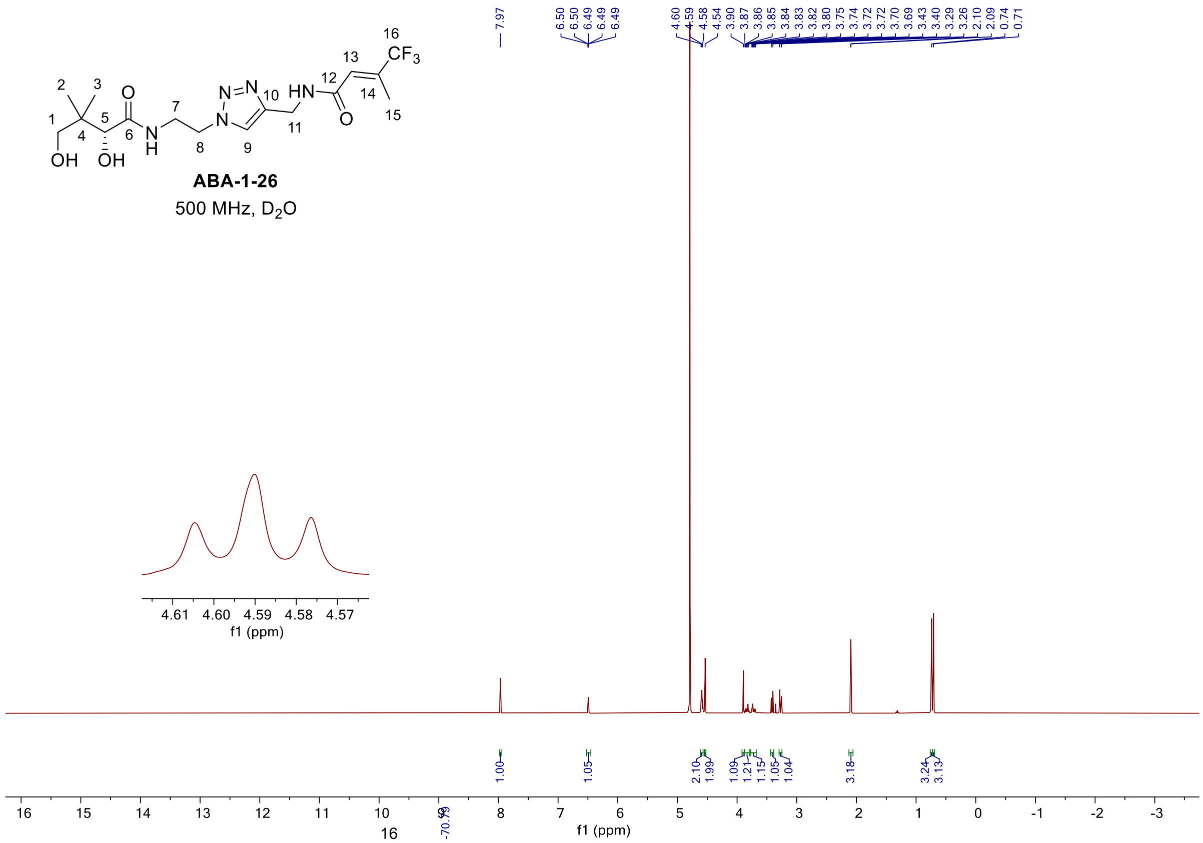

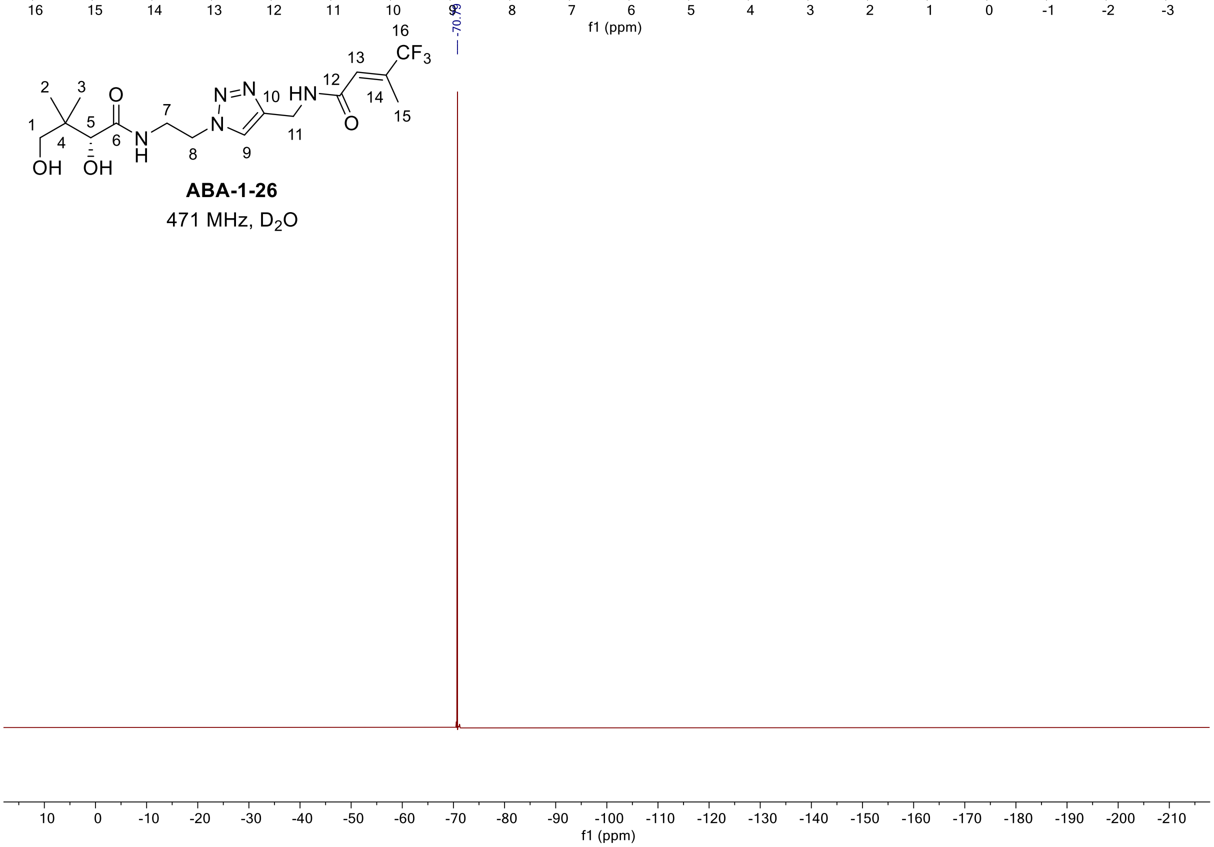

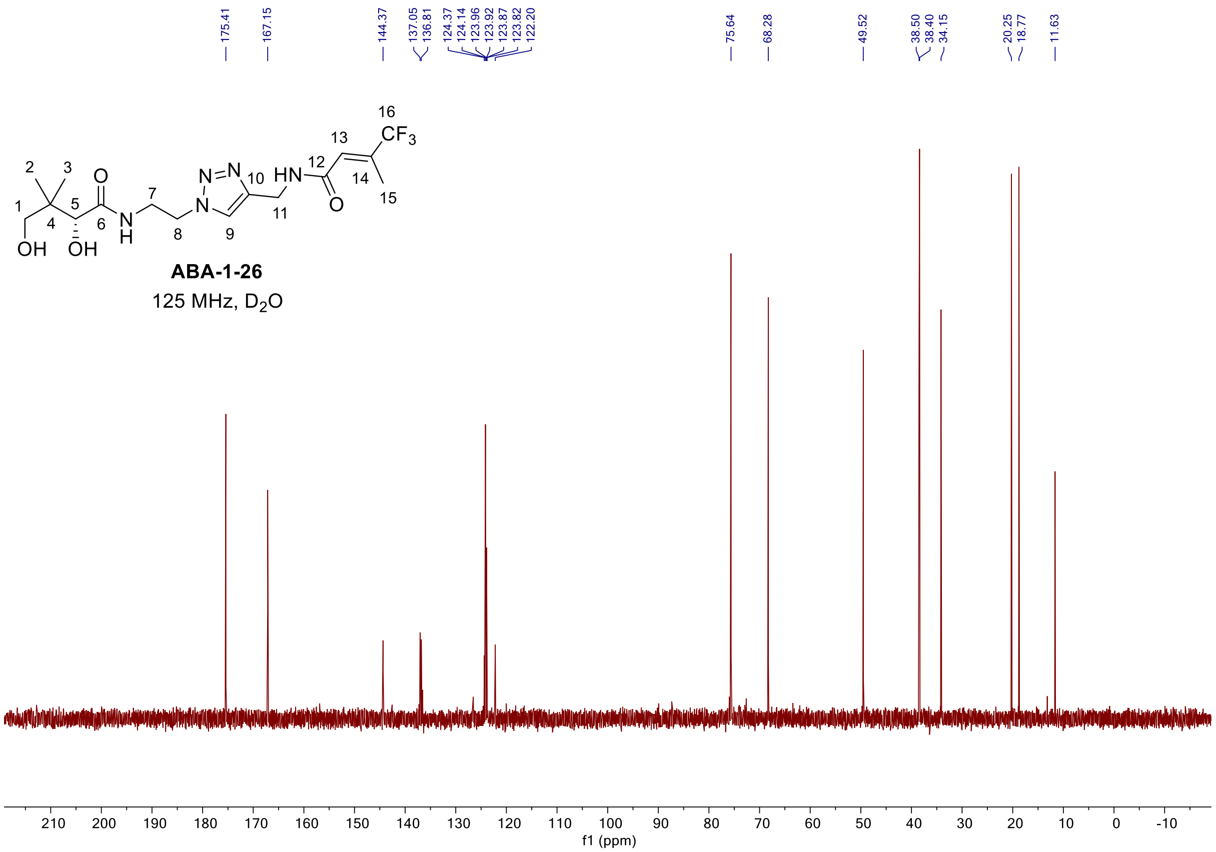

175.41
167.15
144.37
137.05
136.81
124.37
124.14
123.96
123.92
123.87
123.82
122.20
75.64
68.28
49.52
38.50
38.40
34.15
20.25
18.77
11.63
ABA-1-26
125 MHz, D2O
f1 (ppm)

[1]H and [13]C spectra of **ABA-1-32**

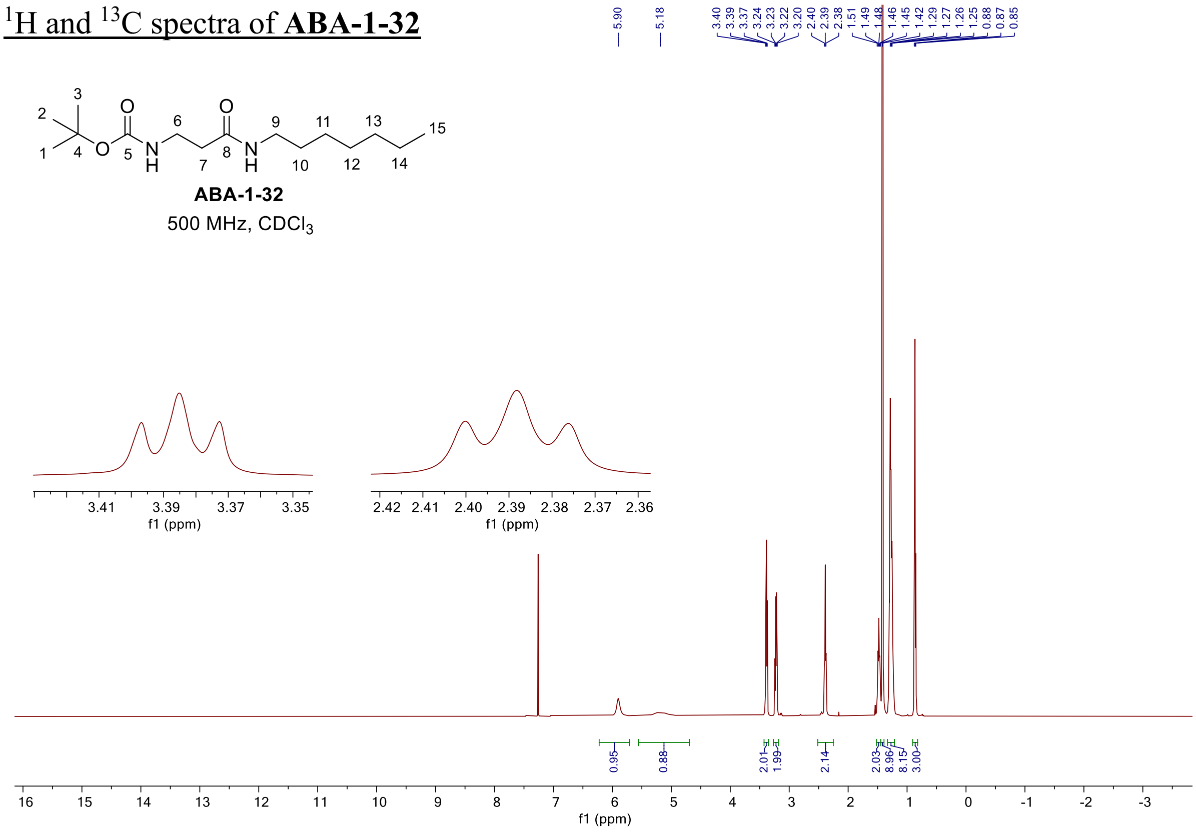

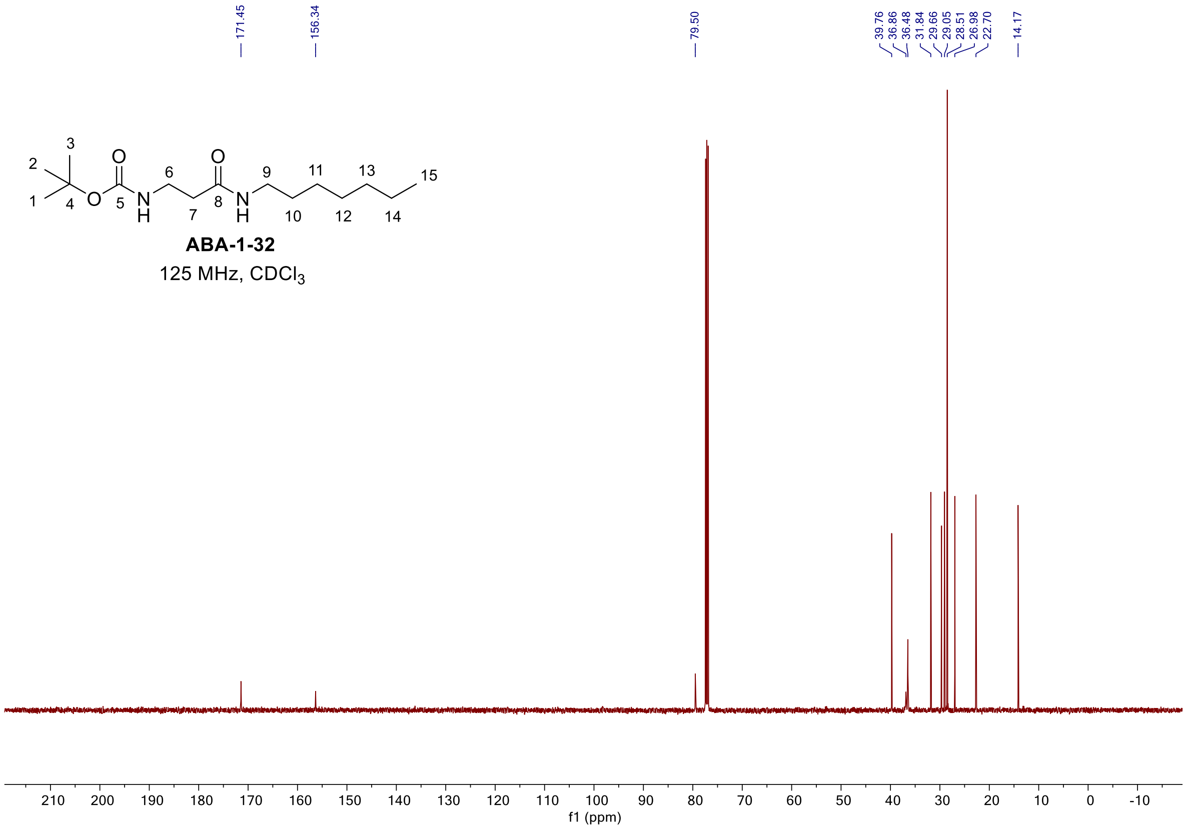

$^{1}H$ and $^{13}C$ spectra of **ABA-1-34 (6-13)**

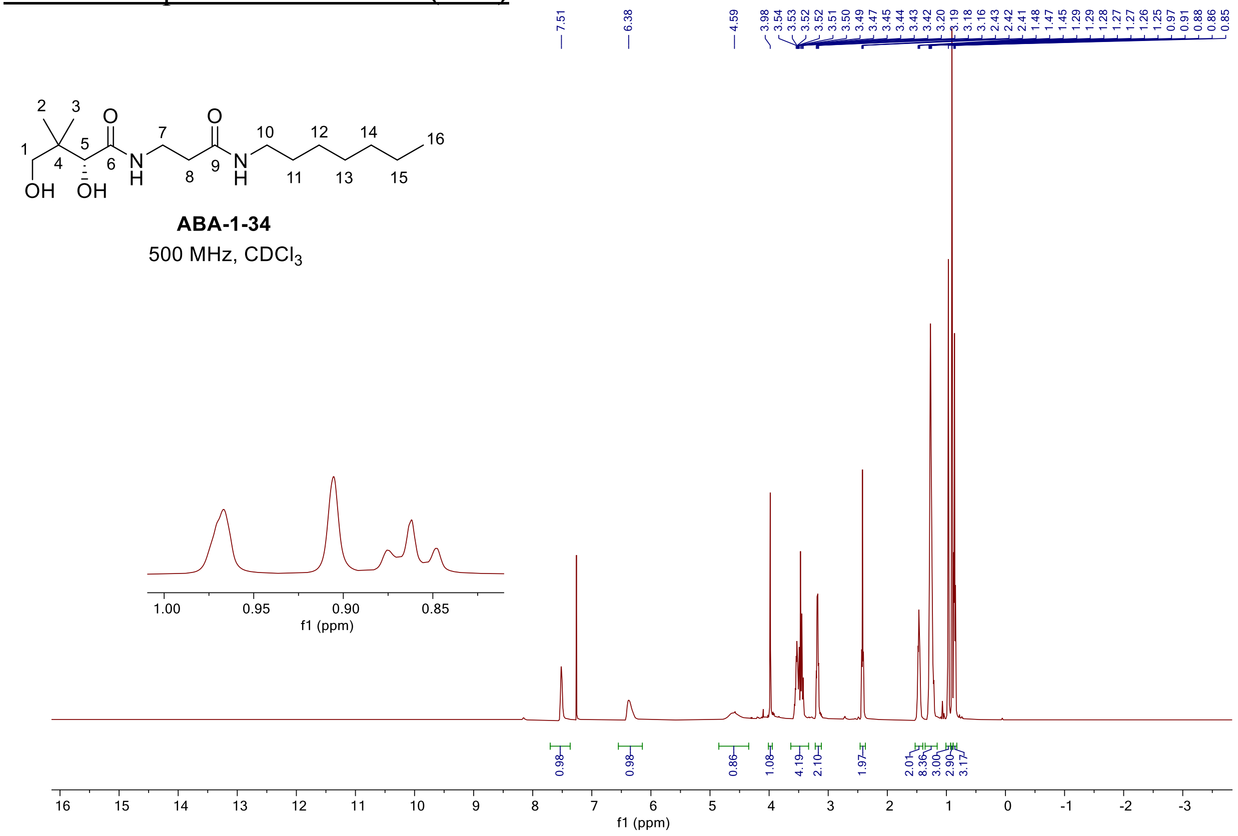

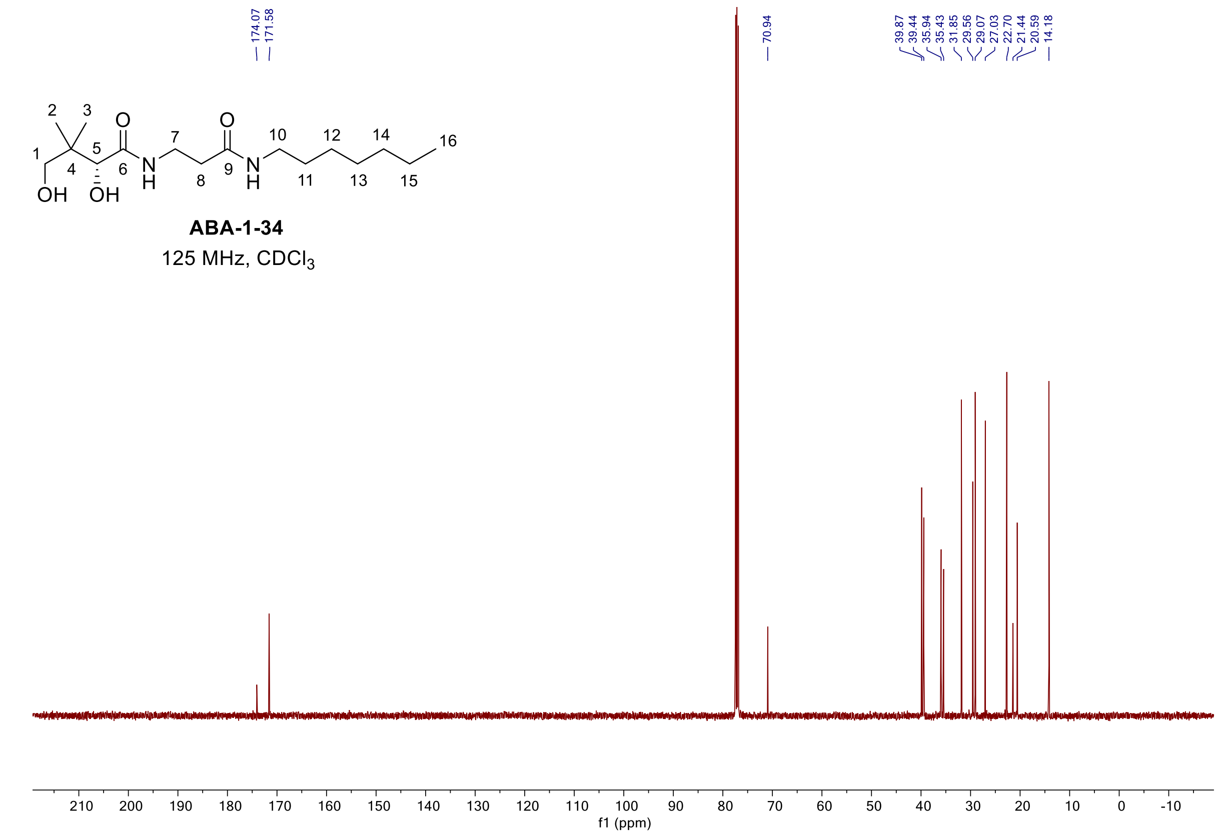

$^{1}H$ and $^{13}C$ spectra of **SCC-3-98**

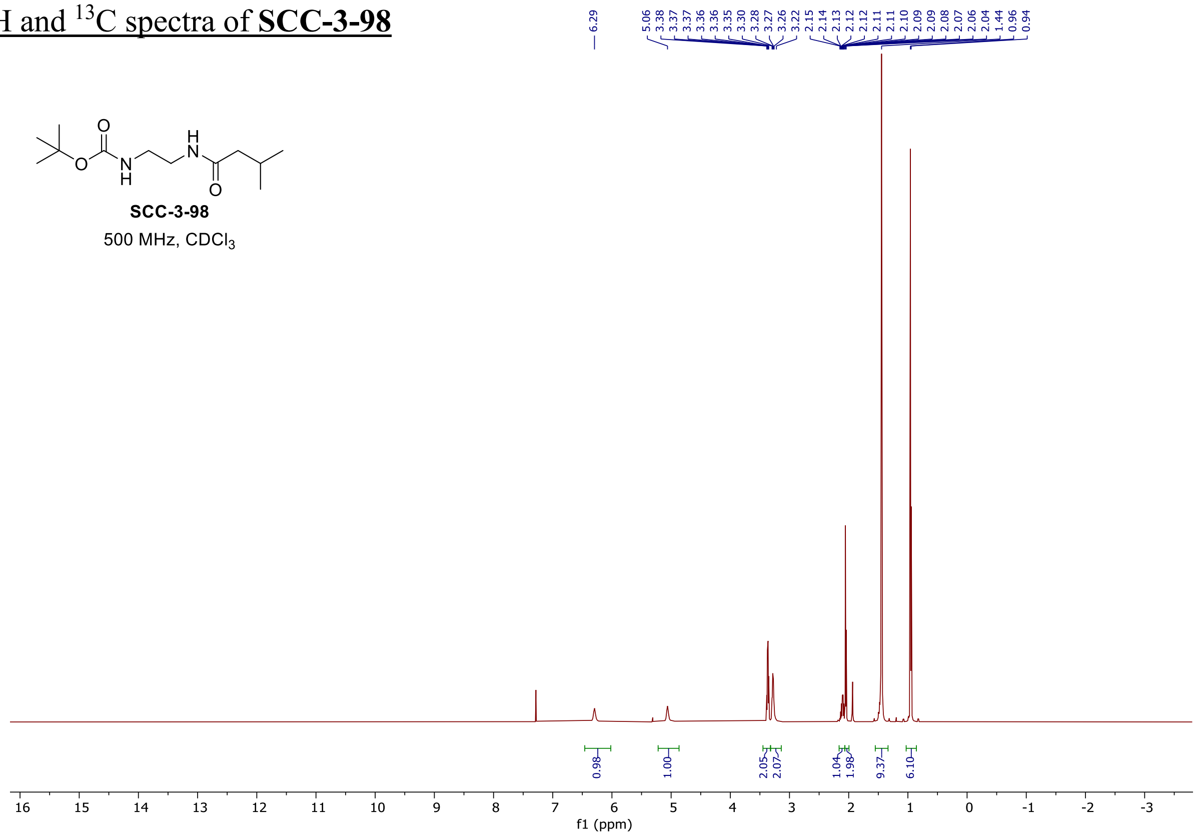

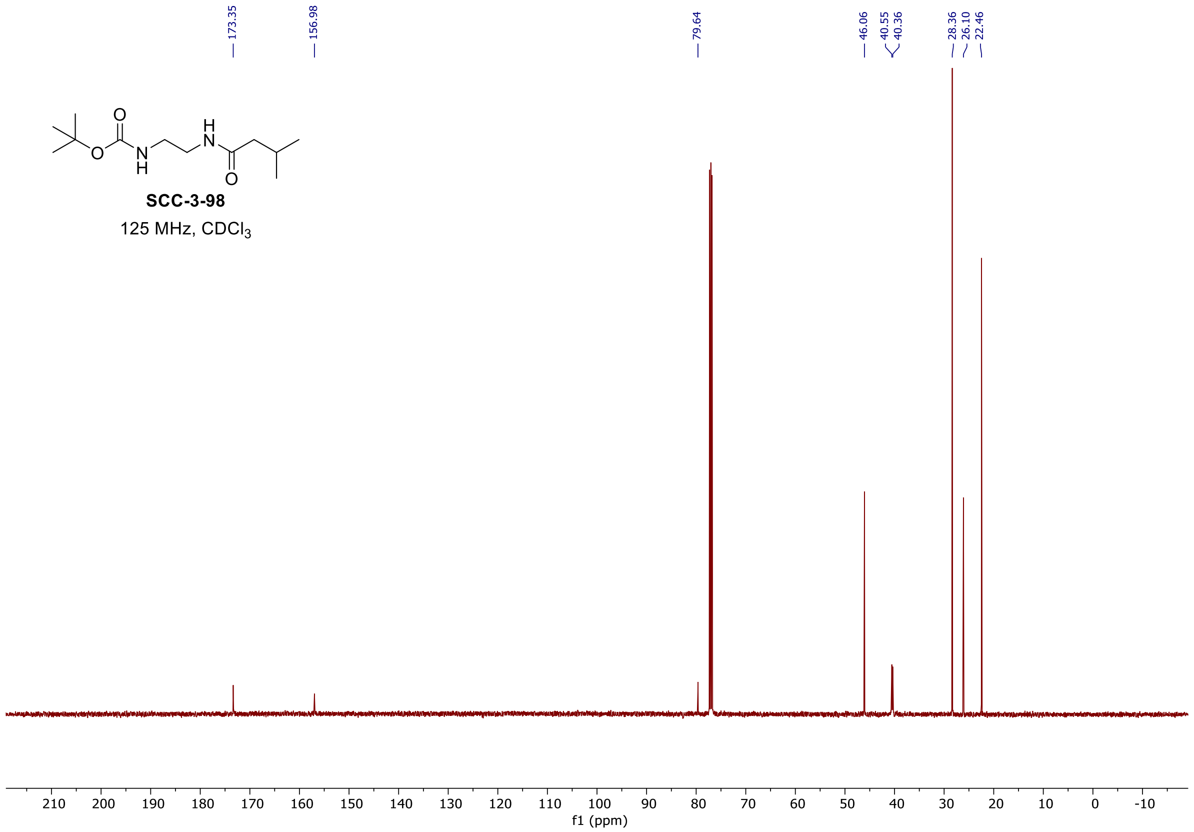

<u>$^{1}$H and $^{13}$C spectra of **SCC-3-93**</u>

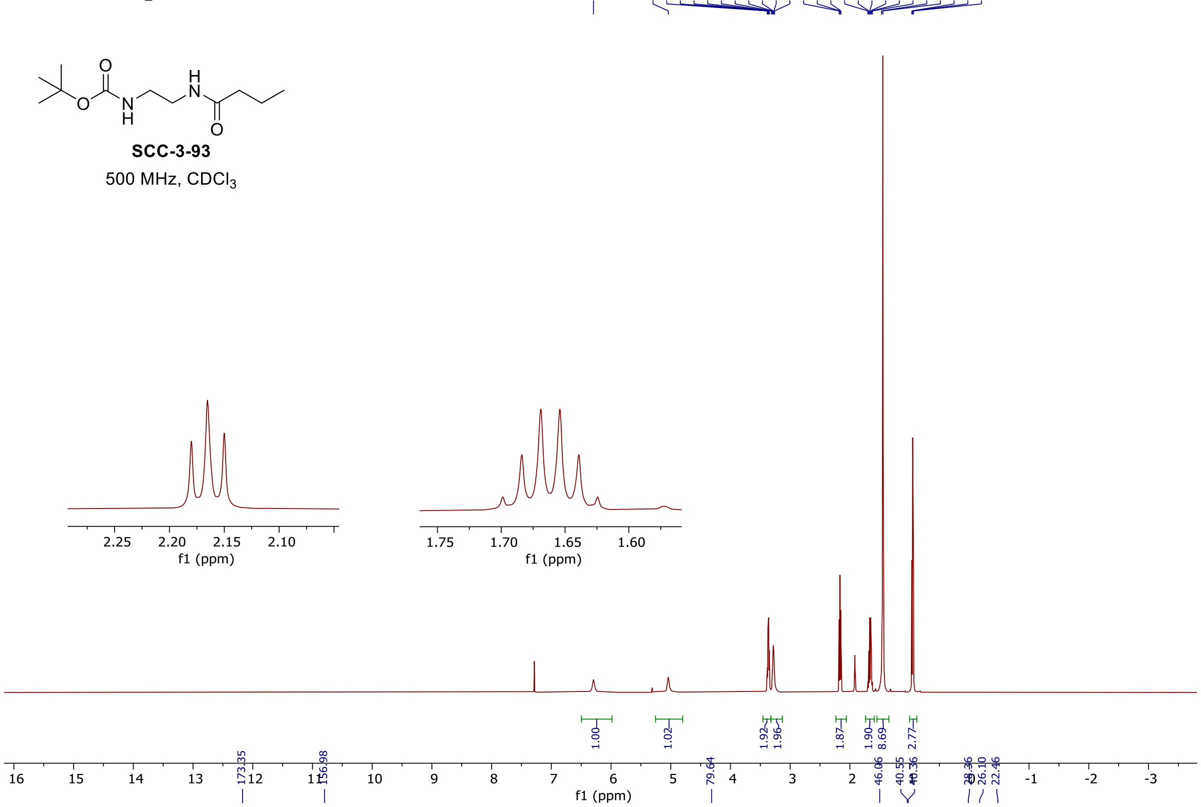

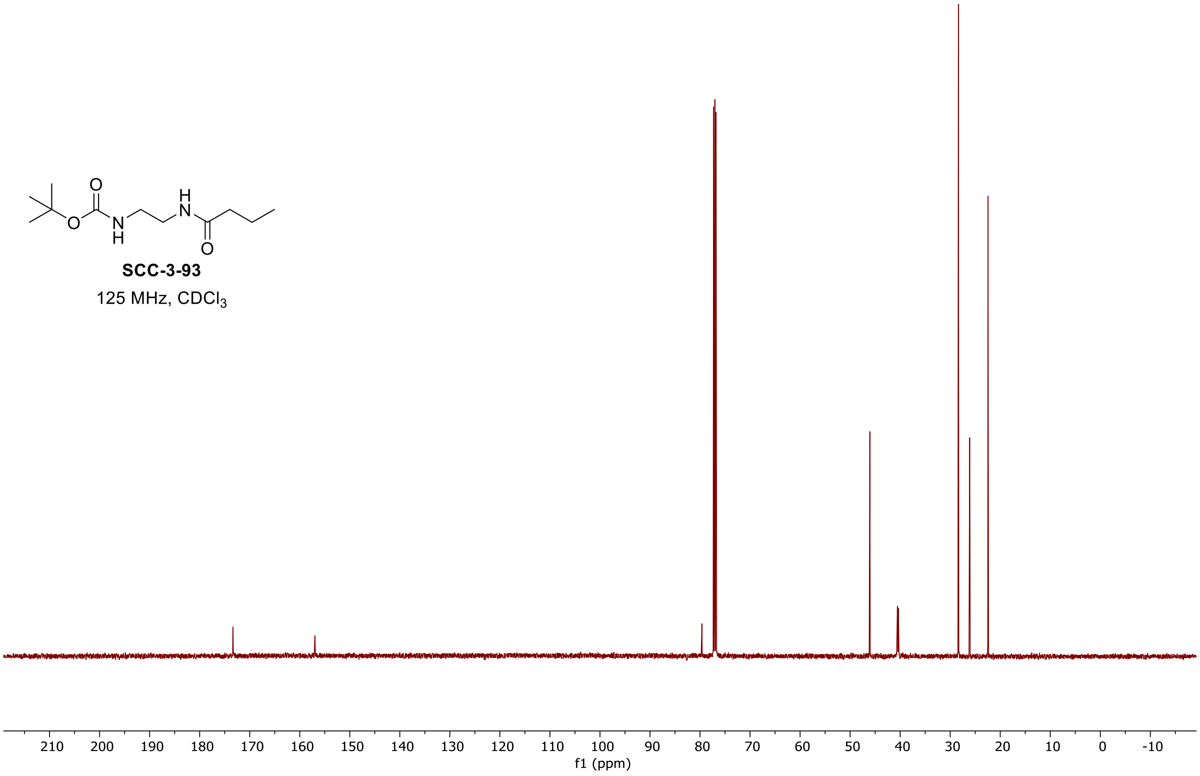

$^{1}$H and $^{13}$C spectra of **SCC-2-96**

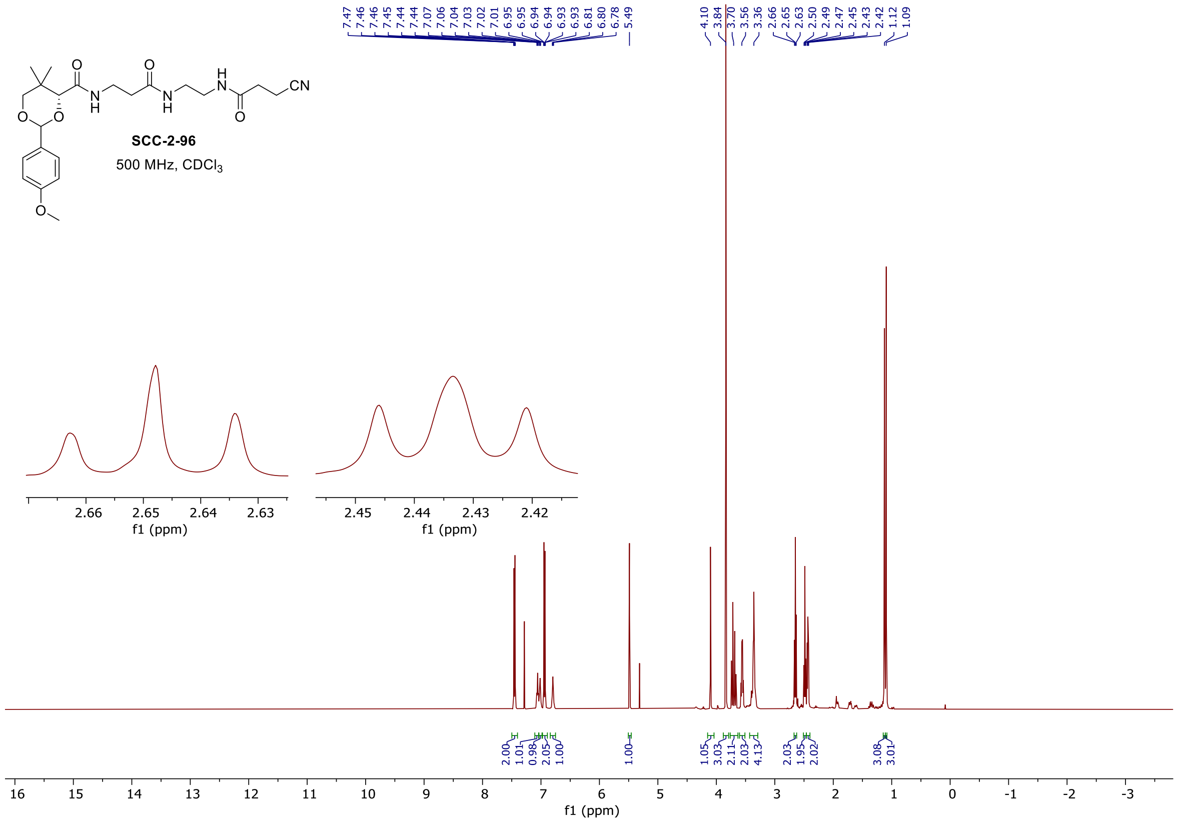

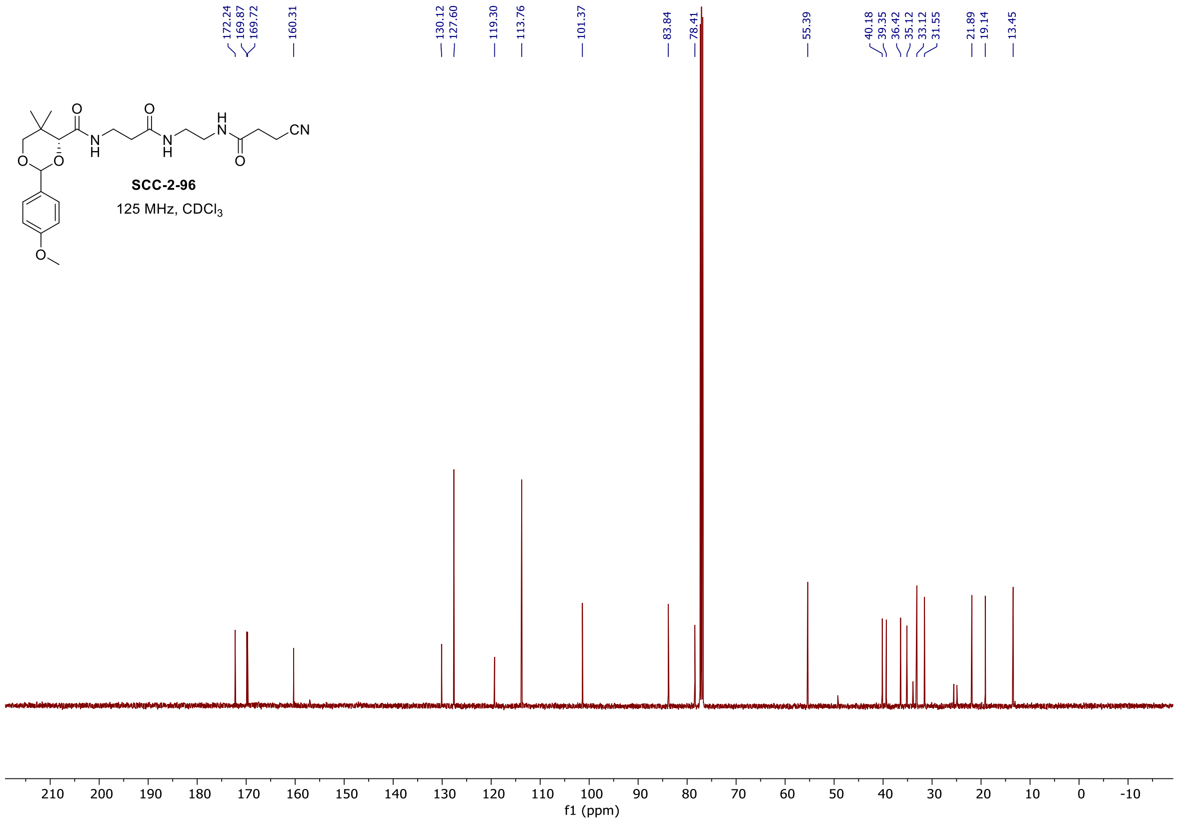

[1]H and [13]C spectra of **SCC-3-43**

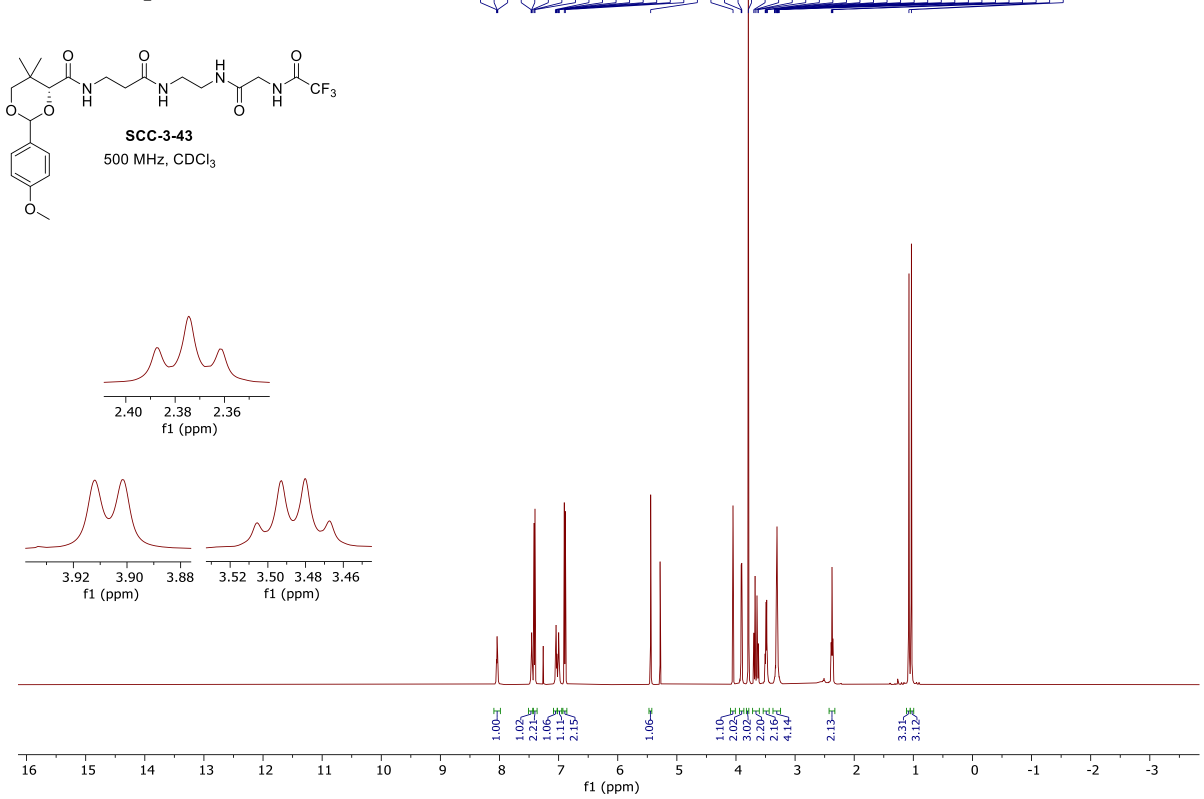

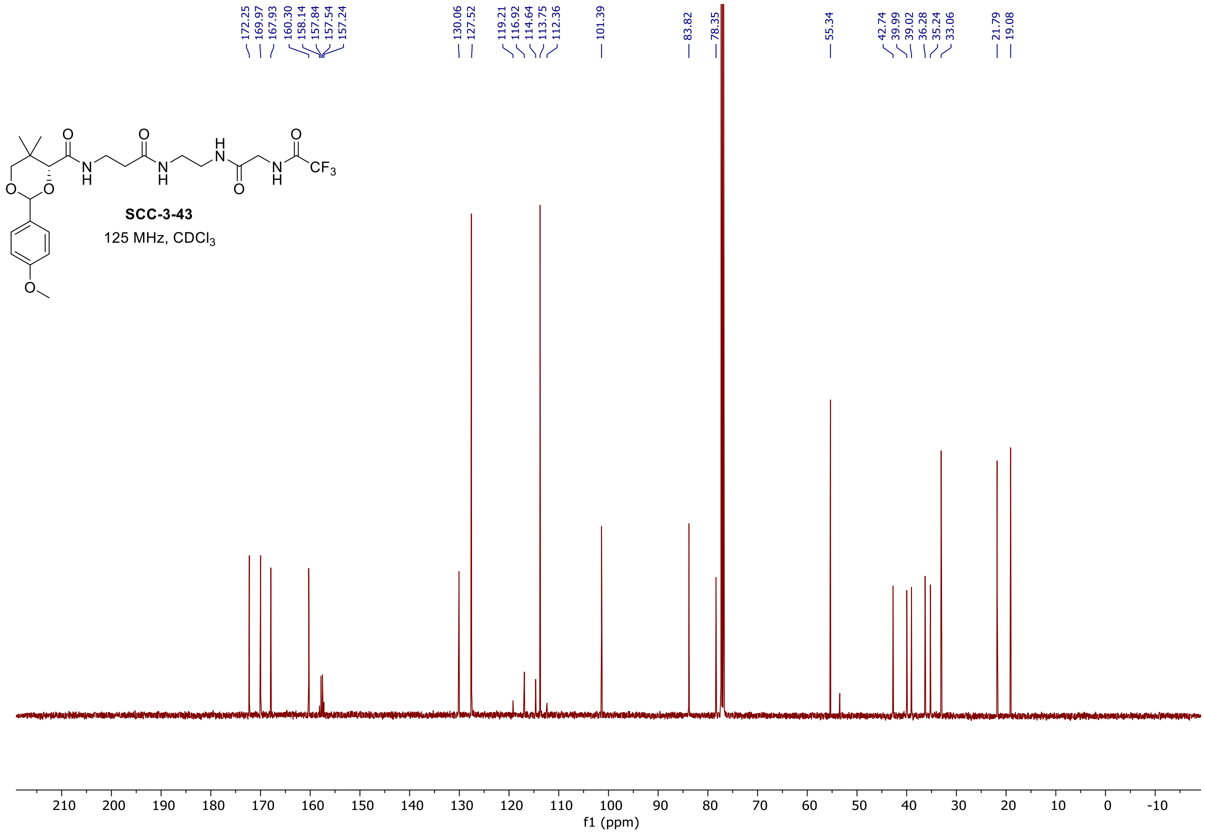

[1]H and [13]C spectra of **SCC-3-95**

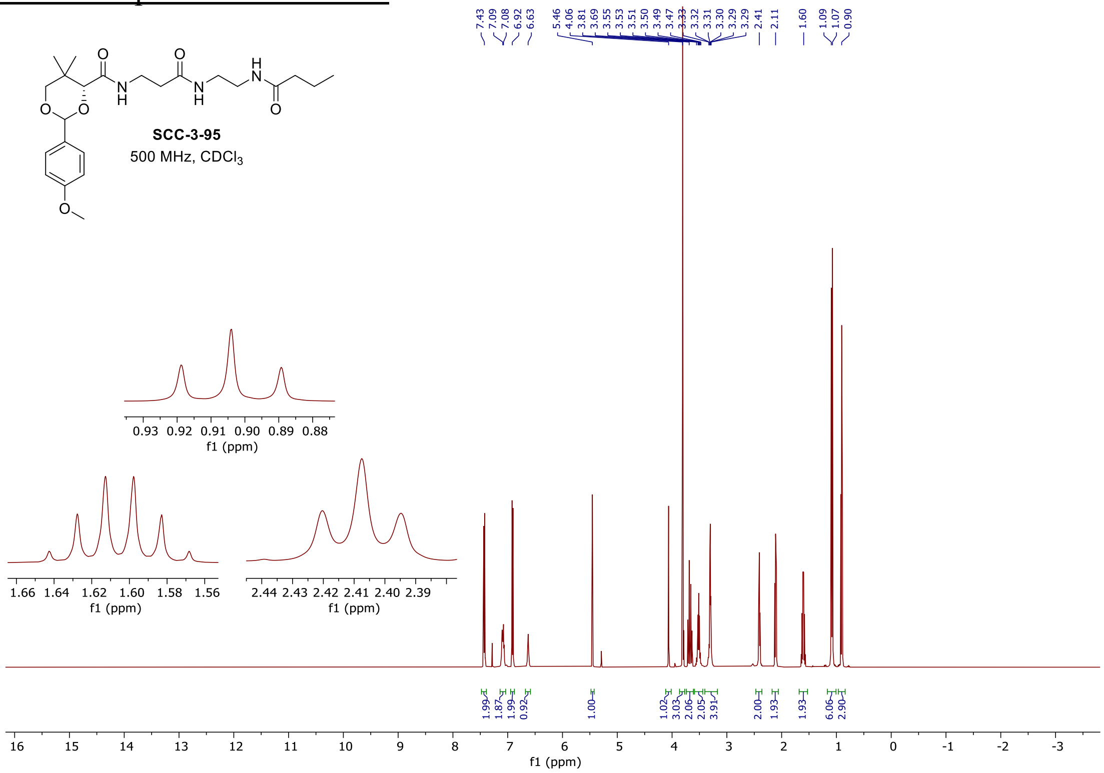

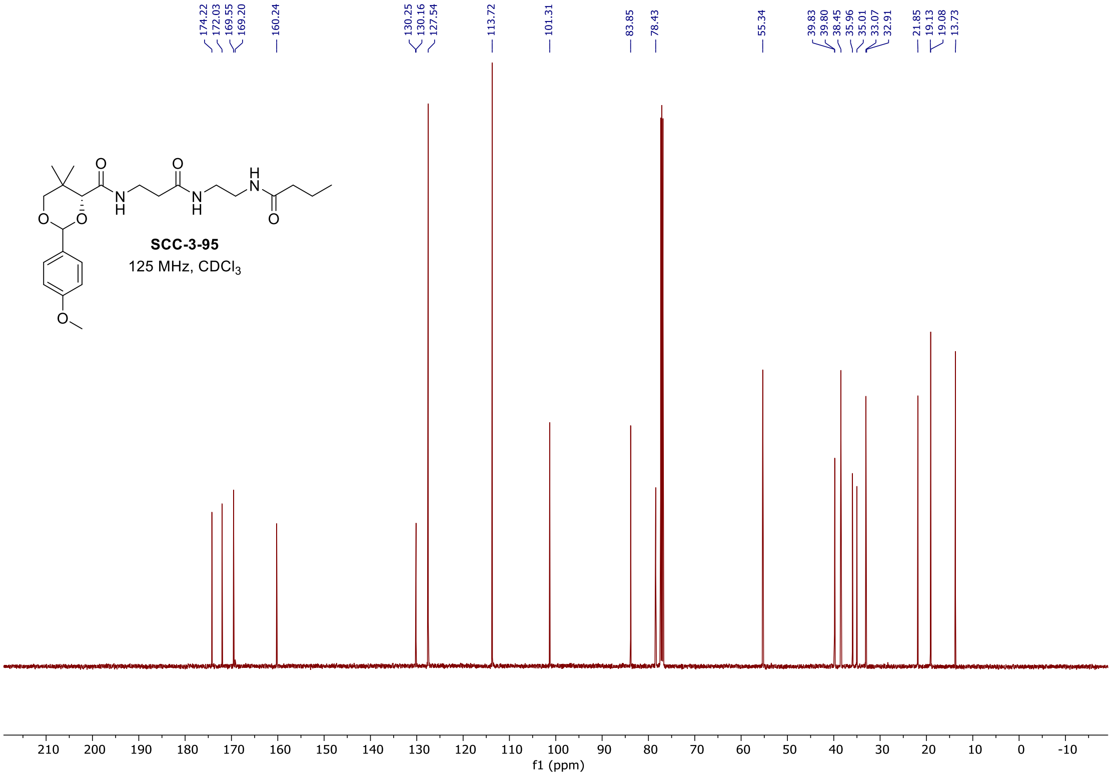

### $^{1}$H and $^{13}$C spectra of **SCC-3-99**

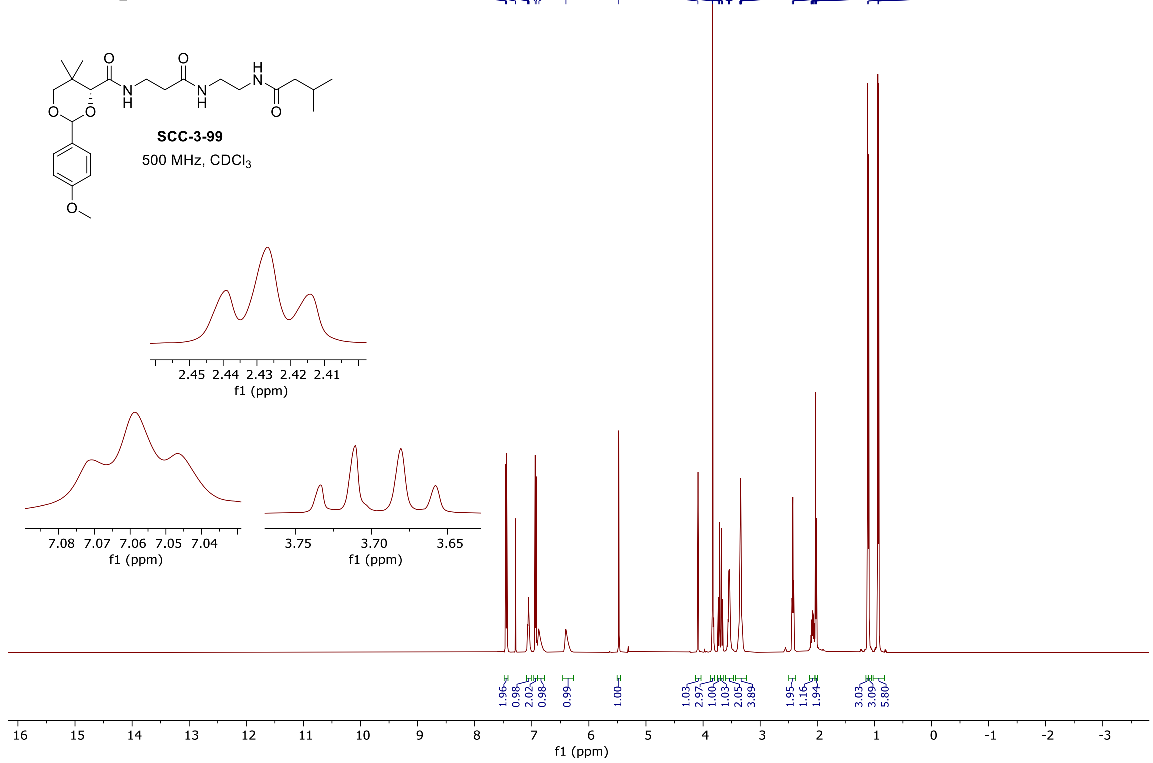

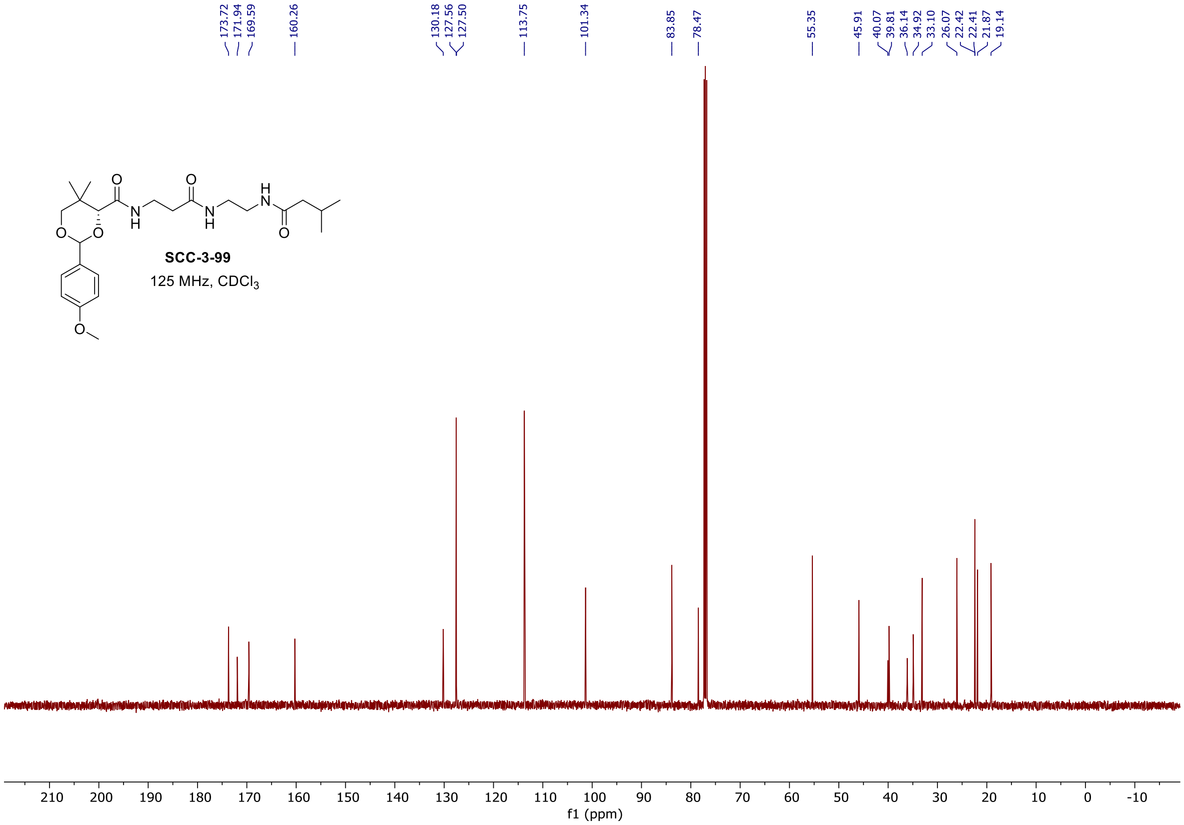

### $^{1}$H and $^{13}$C spectra of **SCC-2-99**

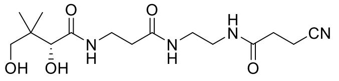

**SCC-2-99**

500 MHz, $CD_3OD$

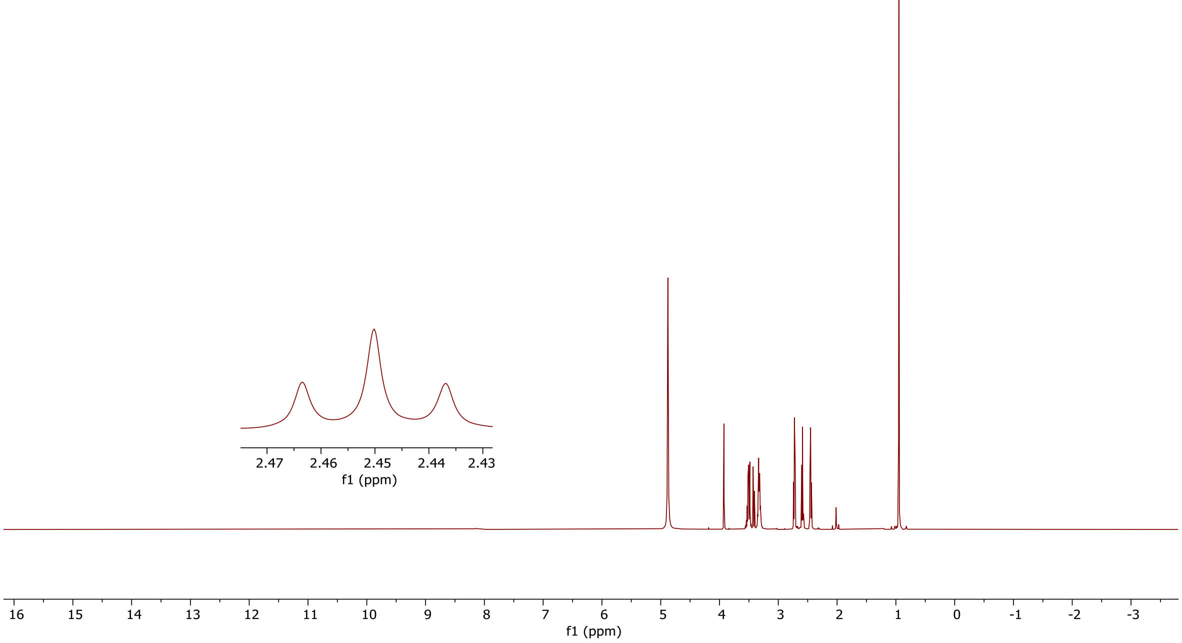

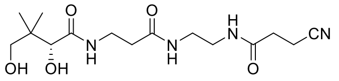

**SCC-2-99**

125 MHz, $CD_3OD$

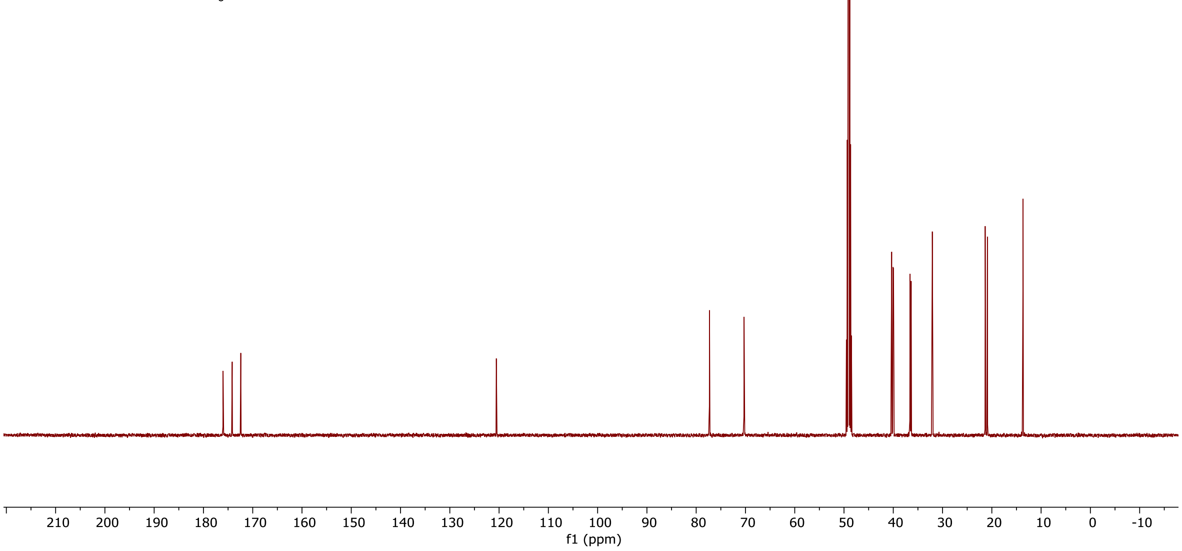

[1]H and [13]C spectra of **SCC-3-103**

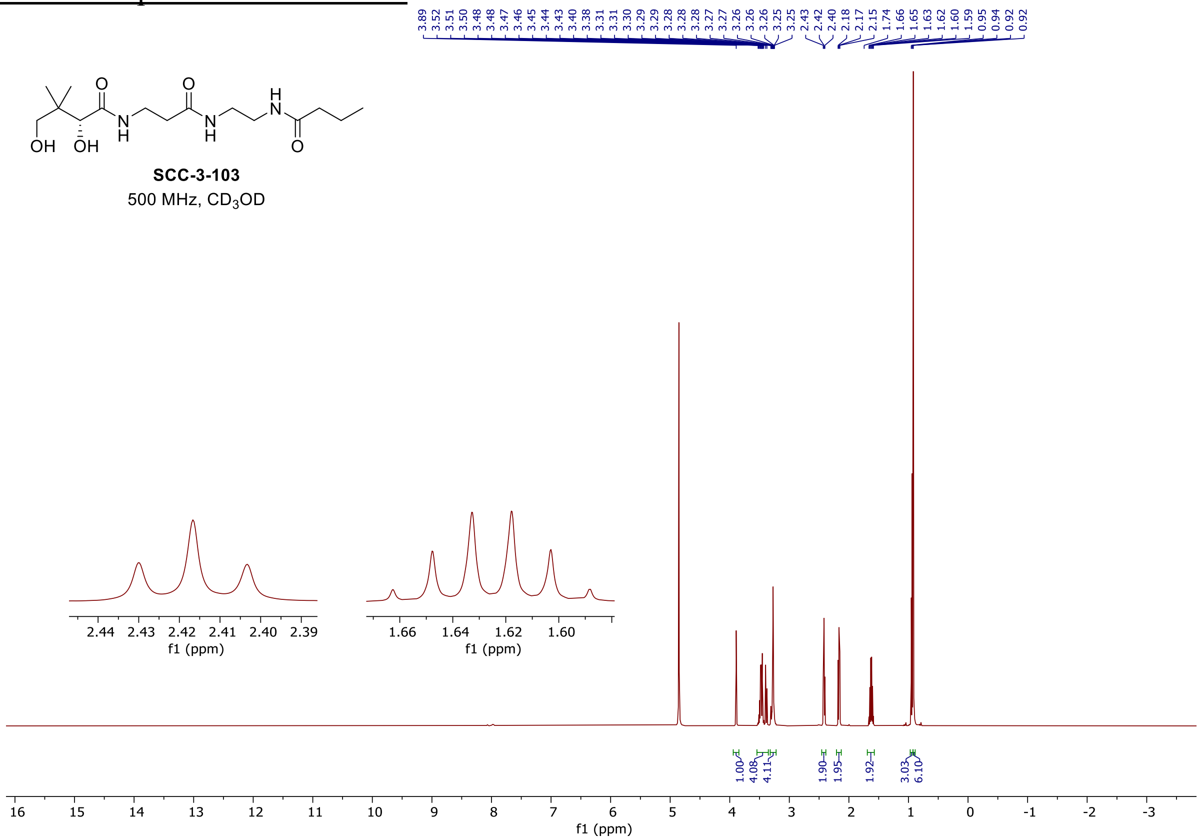

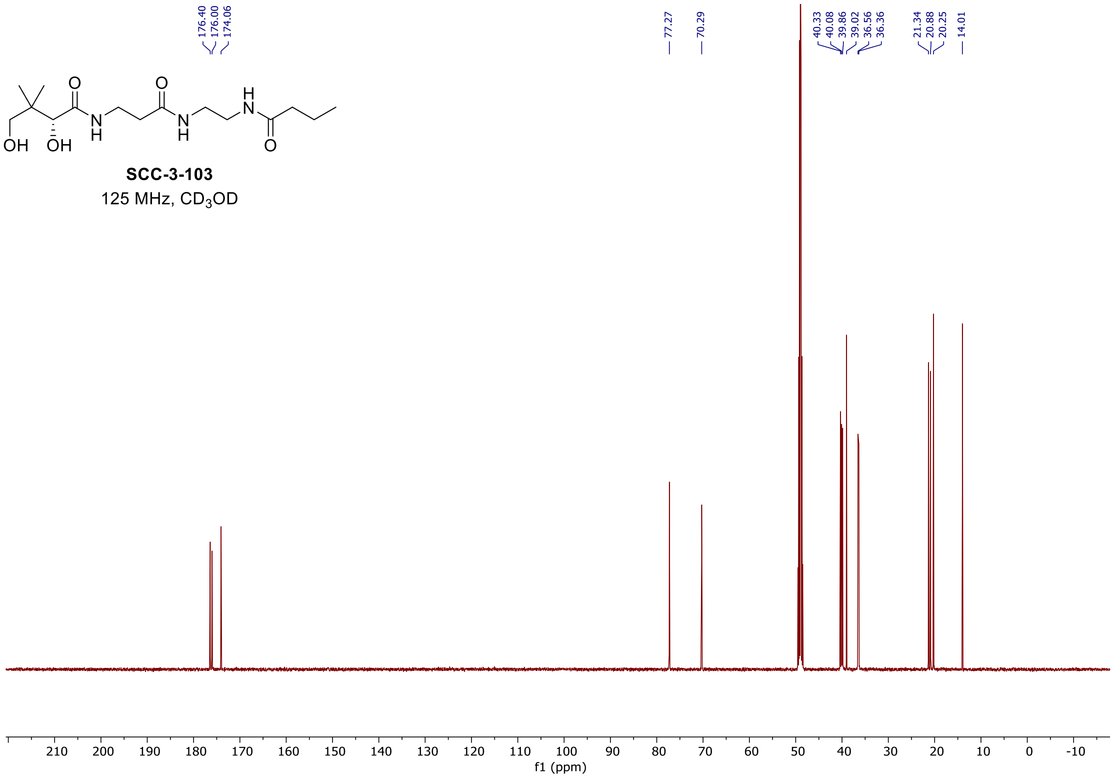

<u>[1]H and [13]C spectra of **SCC-3-102**</u>

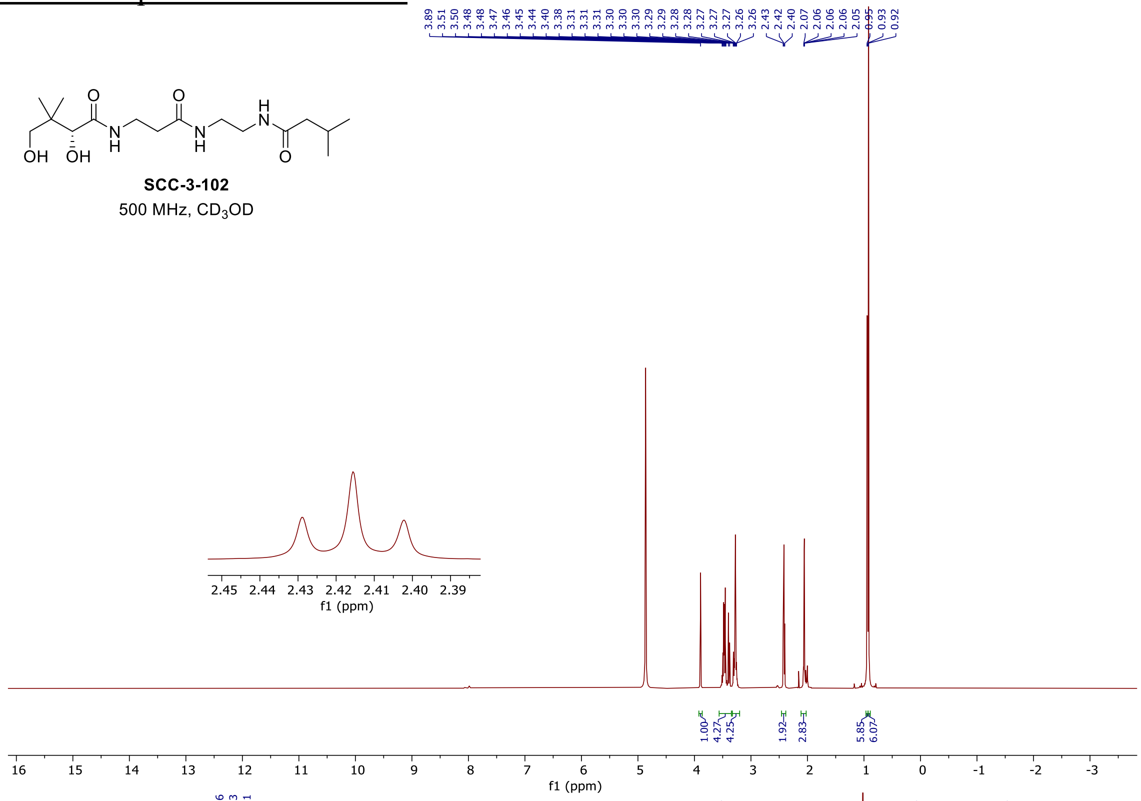

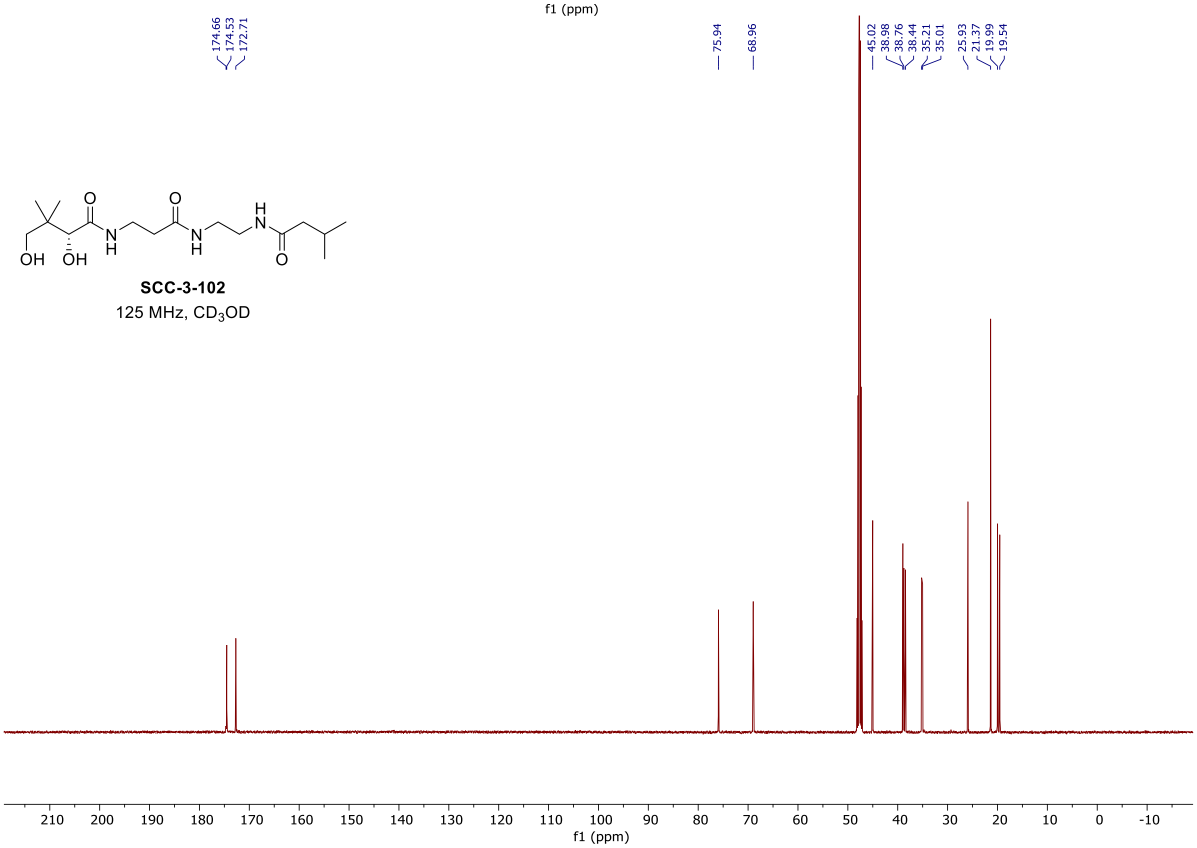

#### [1]H and [13]C spectra of **SCC-3-64**

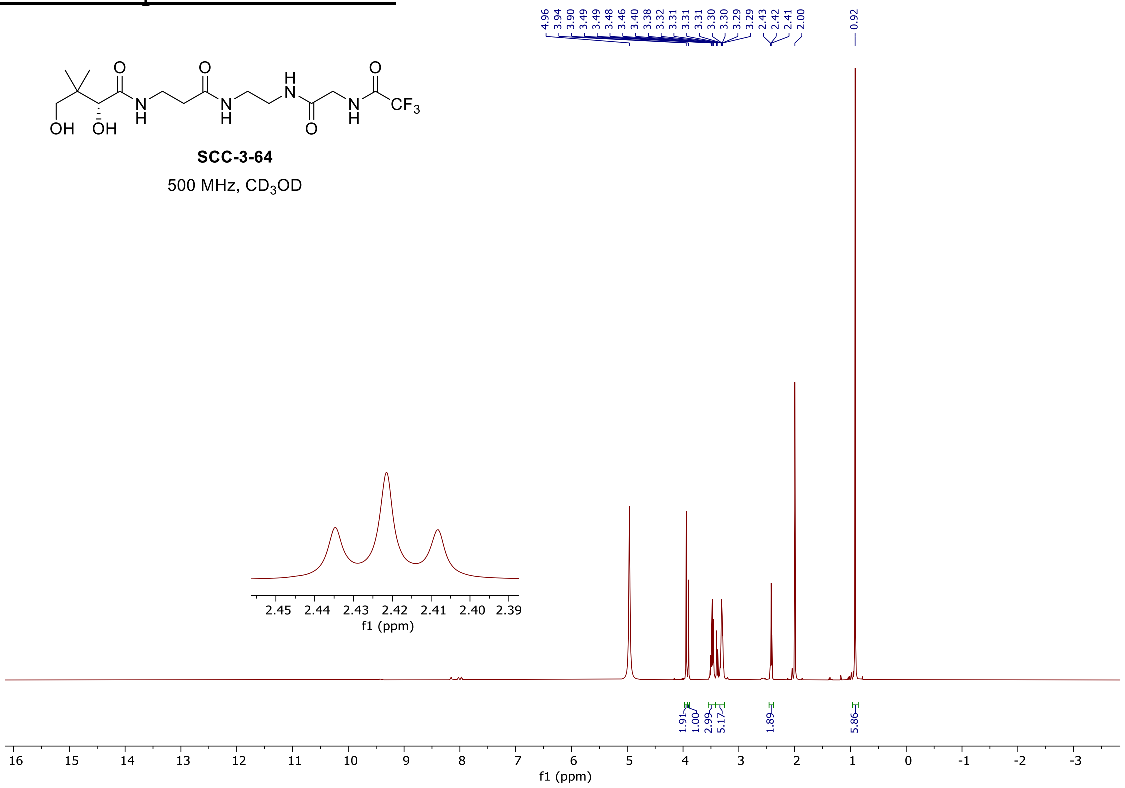

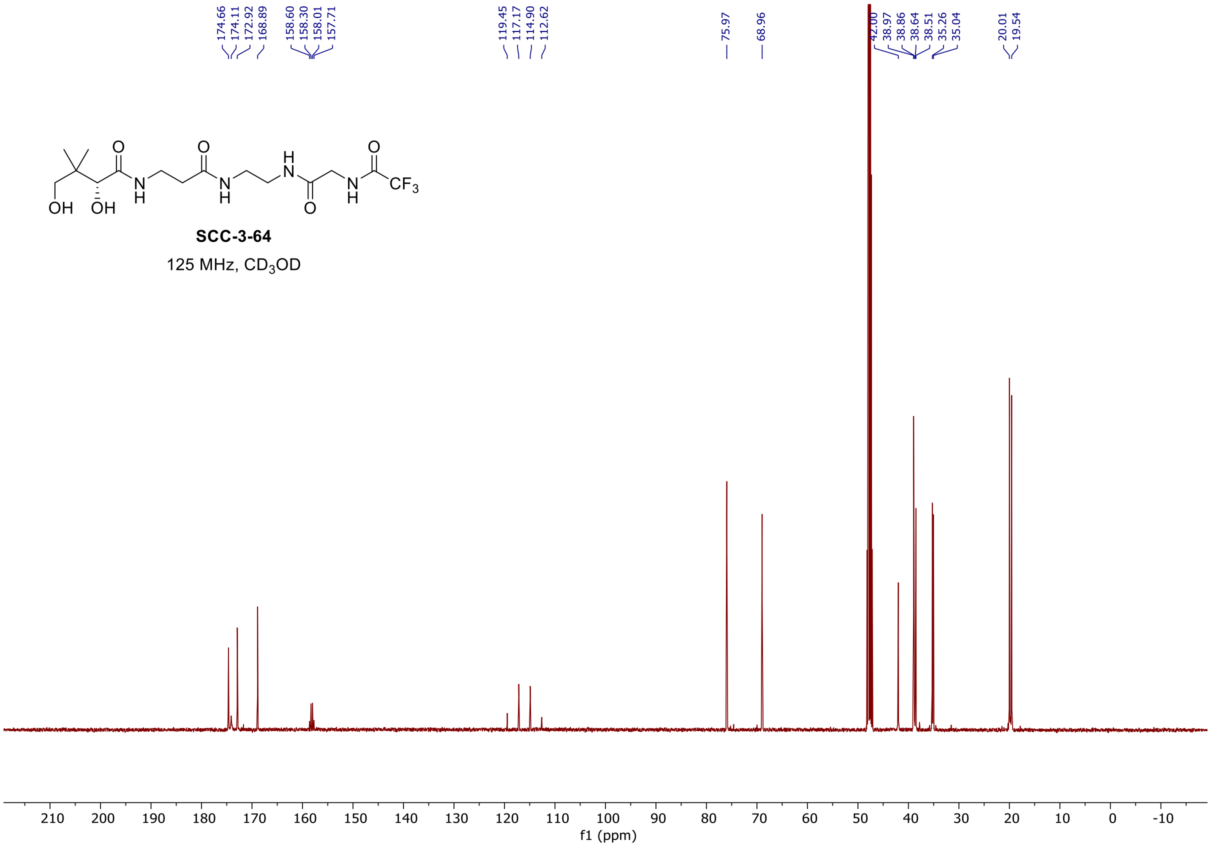
